# Wave succession in the pandemic clone of *Vibrio parahaemolyticus* driven by gene loss

**DOI:** 10.1101/2024.06.03.596356

**Authors:** Chao Yang, Hongling Qiu, Sarah L. Svensson, Chengpei Ni, Song Gao, Zhizhou Jia, Huiqi Wen, Li Xie, Wenxuan Xu, Yujiao Qin, Shuzhu Lin, Jiancheng Wang, Yiquan Zhang, Yinghui Li, Min Jiang, Xiaolu Shi, Qinghua Hu, Zhemin Zhou, Yanjie Chao, Ruifu Yang, Yujun Cui, Jaime Martinez-Urtaza, Hui Wang, Daniel Falush

## Abstract

While spontaneous mutation and gene acquisition are well-established drivers of pathogen adaptation, the role of gene loss remains underexplored. Here, we investigated the emergence and diversification of the pandemic clone (PC) of *Vibrio parahaemolyticus* through large-scale phylogenomic analysis of 8,684 global isolates. The PC rapidly acquired multiple marker genes and genomic islands, subsequently diverging into successive sublineages mediating independent waves of cross-country transmission, as also observed in *V. cholerae*. Wave succession in the last two decades was driven by loss of putrescine utilization (Puu) genes, conferring phenotypic advantages for environmental adaptation (enhanced biofilm formation) and human transmission (increased cell adhesion/intestinal colonization, reduced virulence), consistent with the virulence trade-off hypothesis. We identified Puu-gene loss in multiple bacterial genera, with effects on biofilm and adhesion replicated in *V. cholerae* and *Escherichia coli*, suggesting convergent evolution and universal phenotypic effects. Our results highlight the indispensable role of gene loss in bacterial pathogen adaptation.

## Introduction

Microbial pathogens have recurrently emerged from environments or animals, with a few leading to disease epidemics or pandemics that pose notable public health challenges^1,2^. Understanding the genetic basis of pathogen emergence and adaptive evolution is critical for disease control and prevention, especially for pandemic pathogens. *Vibrio*, Gram-negative, halophilic bacteria that are ubiquitous in marine and aquatic habitats, are ideal models for investigating pathogen emergence and evolution. Among >100 *Vibrio* species, approximately 12 cause human infections^3^. Two species, *V. cholerae* and *V. parahaemolyticus*, are associated with disease pandemics, with the ongoing seventh cholera pandemic attributed to El Tor (7PET) isolates. *V. cholerae* 7PET was inferred to originate from a pre-pandemic strain that subsequently underwent explosive diversification - gaining multiple genetic elements within a short timeframe (∼15 years), including the El Tor type cholera toxin prophage and *Vibrio* seventh pandemic islands (VSP-I and VSP-II)^4,5^. During the seventh pandemic, three successive sub-lineages of 7PET were identified, corresponding to the three independent but overlapping waves of global transmission^5,6^, which may be driven by antibiotic resistance^7^.

*V. parahaemolyticus* is the leading cause of seafood associated infections worldwide, with an increase in incidence over the last two decades. Its geographical range has also been expanding, possibly due to global warming^3,8,9^. The emergence of a pandemic clone (PC), characterized by serotype O3:K6, substantially changed *V. parahaemolyticus* epidemiology^10–12^. The O3:K6 PC strain was first isolated in 1995 from a traveler returning to Japan from Indonesia, and in 1996, it caused a disease outbreak in Calcutta, India^10^. Subsequently, it rapidly spread worldwide and came to dominate infections in many regions, along with rapid antigenic divergence (49 serotypes reported)^10–12^. Molecular and epidemiological surveillances has provided evidence that the PC may have originated from a serotype O3:K6 non-pathogenic strain after acquisition of multiple marker genes (e.g. virulence regulatory gene sequence *toxRS/new*^13^, phage f237 and its unique open reading frame *orf8*^14^) and marker genomic islands (VPaI-1 to VPaI-7), along with several virulence-associated loci such as those encoding Type 3 or 6 secretion systems (T3SS2 or T6SS1)^15,16^. However, how these events happened, the driving forces, and mechanisms underlying these events remain unclear. Currently, there is a lack of a comprehensive genomic view on the origin and evolution of the PC.

Leveraging a large-scale, global genome dataset of *V. parahaemolyticus*, which includes strains from six continents and 34 countries, we comprehensively reconstructed the evolutionary history of the PC. Intriguingly, we identified several similarities between PCs of *V. parahaemolyticus* and *V. cholerae*. However, distinct from *V. cholerae*, we demonstrated that gene-loss in the putrescine utilization pathway has driven the recent global wave succession event by conferring phenotypic advantages, and these phenotypic effects are likely universal among bacteria.

## Results

### From pre-pandemic (pre-PC) to PC

We assembled a total of 8,684 high-quality (completeness >90% and contamination <5%) genomes of global *V. parahaemolyticus* isolates, including 333 newly sequenced genomes (**Supplementary Table 1**). To characterize population structure and identify PC isolates, we constructed a maximum-likelihood (ML) phylogenetic tree based on core-genome single-nucleotide polymorphisms (SNPs). Based on the ML tree and pairwise SNP-distance, 2,815 were assigned as PC isolates (mean SNP-distance: 437), 39 as pre-PC (close relatives of PC, mean SNP-distance to PC: 15,463), with the remaining being non-PC isolates (mean SNP-distance to PC: 55,554, **Fig. 1a**).

**Figure 1.**
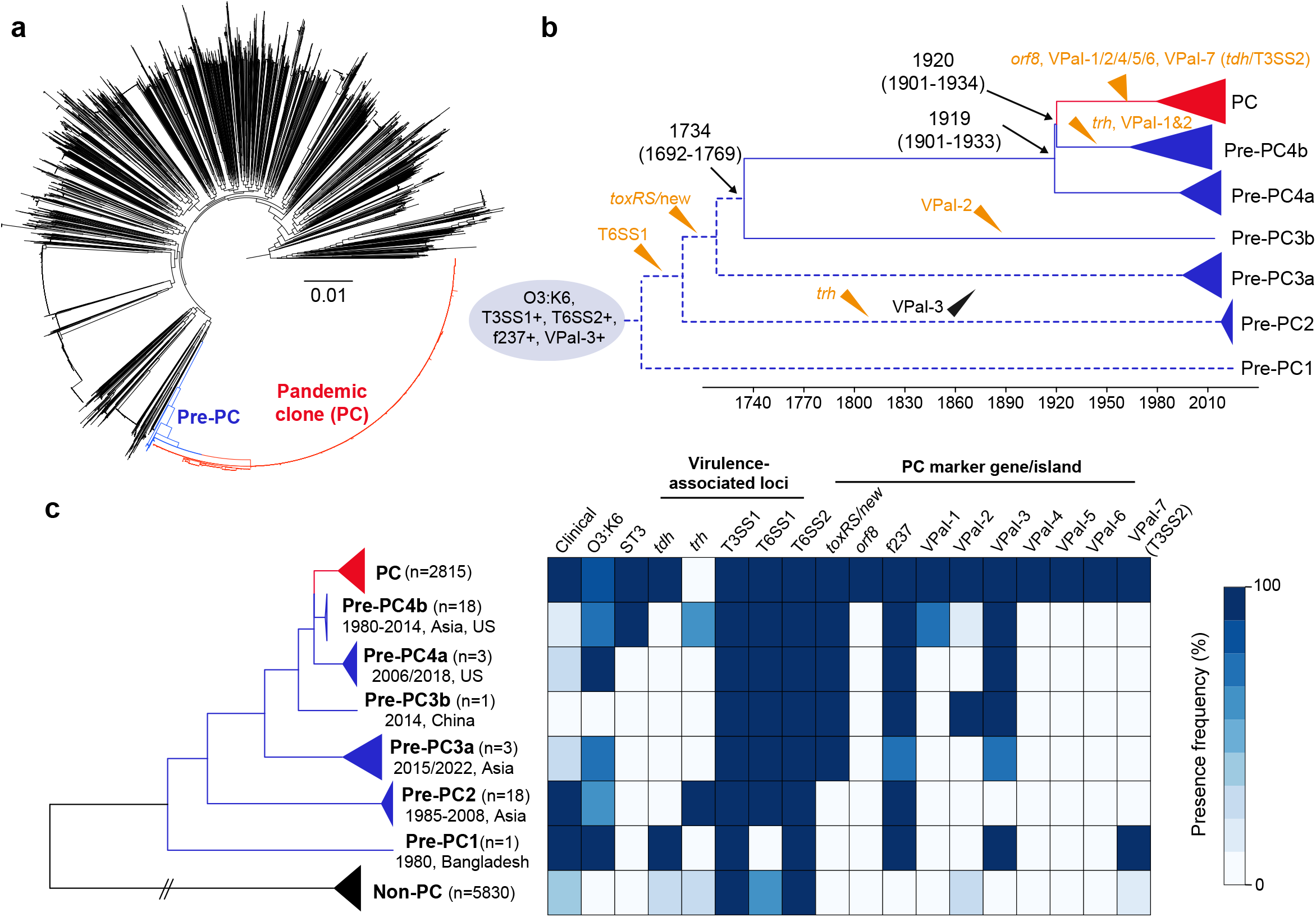
From pre-pandemic (pre-PC) to pandemic clone (PC). **a**, Maximum likelihood tree of 8,684 global isolates. Pre-PC and PC isolates are colored in blue and red, respectively. **b**, Inferred evolutionary history from pre-PC to PC. The status of known PC marker genes/islands in the ancestral strain is indicated by blue ellipses, while subsequent gains or losses are represented by orange and black triangles. The numbers next to tree nodes indicate inferred divergence times and 95% confidence intervals. Dashed lines represent lineages with unknown origination times due to a lack of significant temporal signal. **c**, Distribution of virulence-associated loci and known PC marker genes/islands among pre-PC, PC and non-PC strains. Double slashes indicate artificially shortened phylogenetic branches. Heatmap fill colors represent the frequency of presence for each locus across groups.

Pre-PC isolates were further split into four sub-groups, designated pre-PC1 to pre-PC4, respectively (**Fig. 1b**,**c**). Pre-PC1 to pre-PC3 were exclusively identified in Asia, with pre-PC1 and pre-PC2 being clinical isolates, while pre-PC3 were mainly environmental strains. The closest relative of the PC, pre-PC4, was represented mainly (13/16) by environmental isolates from Asia (Japan and China) and the US. The serotypes of most pre-PC isolates (33/39) were predicted to be O3:K6, and sequence types (STs) of pre-PC4b (a subset of pre-PC4) isolates were ST3 except one strain (**Fig. 1c** and **Supplementary Table 2**).

Among pre-PC and PC isolates, we identified a total of 88,754 core-genome SNPs, with 94% attributed to homologous recombination, suggesting that recombination was the primary driver of divergence from pre-PC to PC. We further assessed the temporal signal using the ML tree based on recombination-stripped SNPs and strain isolation dates. A significant signal (*p*<0.01) was detected among strains from pre-PC3b to PC, allowing inference of the dated phylogeny (**Fig. 1b**,**2b**). The most recent common ancestor (MRCA) of the PC was estimated at 1983 (95% confidence interval [CI]: 1979-1985), 12 years preceding its first discovery in 1995. The strains are expected to be highly homogeneous at the point of their origin, but early PC strains from 1996 already showed some diversity (an average of 38 non-recombined SNPs), consistent with the long time-interval between their origination and first discovery (**Fig. 2b**).

**Figure 2.**
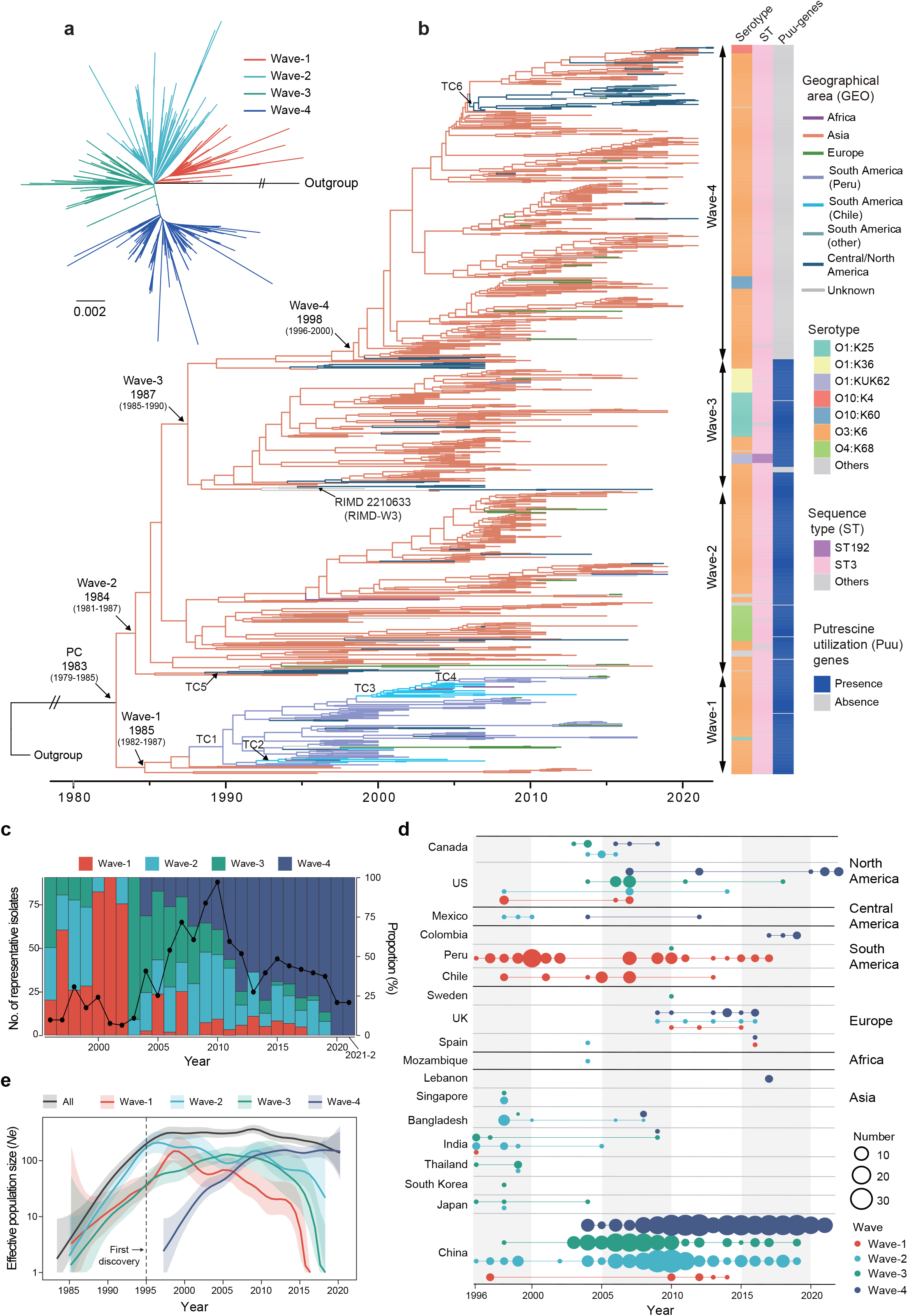
Fine-scale population structure and spatiotemporal/population dynamics of the PC. **a**, Maximum likelihood tree of 942 representative PC strains. The colors represent different waves. Double slashes indicate artificially truncated branches. **b**, Dated phylogeny of the PC (left) and the geographical, subtyping and distribution of genes in the putrescine utilization (Puu) pathway of strains (right). The colors of tree branches indicate geographic areas, and numbers next to tree nodes indicate inferred divergence times and 95% confidence intervals. TCs indicate transmissions resulting in local colonization. **c**, Temporal dynamics of four waves. Bar colors indicate the frequency of different waves. The line represents the number of representative strains. **d**, Temporal distribution of representative strains in different countries and regions. The size of circles scales with the number of representative strains. **e**, Effective population size dynamics of different waves of the PC. Background shading represents 95% confidence intervals.

We investigated the distribution of known PC marker genes/islands and inferred the evolutionary history from pre-PC to PC (**Fig. 1b**,**c** and **Extended Data Fig. 1)**. The PC likely originated from a serotype O3:K6 pathogenic strain encoding T3SS1, T6SS2, phage f237 and VPaI-3, and subsequently acquired T6SS1 and the *toxRS/new* sequence. Between 1920 (MRCA of pre-PC4b and PC) and 1983 (MRCA-PC), MRCA-PC gained multiple known marker genes/islands distributed across distinct genomic loci, including *orf8* and six out of the seven VPaIs, as well as the gene encoding the well-known virulence factor thermostable direct hemolysin, *tdh*. Additionally, 86 point mutations (34 synonymous; 38 non-synonymous) and 80 homologous recombination events (524 kb, 10% of the reference genome) were inferred along this branch. Moreover, *trh* (*tdh*-related hemolysin), VPaI-1 and VPaI-2 were independently gained or lost by pre-PC strains.

We further investigated the source of these marker genes/islands by searching against non-PC genomes. All of them were identified in non-PC isolates, although at low frequency **(Fig. 1c, Extended Data Fig. 2**), indicating that these elements are more likely the result of intra-species transfer from diverse non-PC *V. parahaemolyticus* strains than of cross-species transfer from other *Vibrio* or bacterial species. Moreover, the differing distribution patterns of these marker genes/islands among pre-PC and PC strains suggest that they were unlikely to have been acquired simultaneously in a single burst. Apart from marker genes/islands, six PC-specific genes were identified, three of which were within the superintegron (**Supplementary Table 3**).

### Fine-scale population structure of PC

To reduce the effect of repeat sampling (strains from disease outbreaks) and exclude outlier strains, we generated a representative subset of 942 PC isolates for further analysis (see methods), spanning 20 countries across five continents isolated between 1996 and 2022 (**Extended Data Fig. 3**). Among the representative isolates, we identified 26,377 core-genome SNPs, with 66% attributed to homologous recombination. Two recombination hotspots were identified, including the known O:K antigen region^17^ and a novel hotspot (VP1883-1891) encoding a type II toxin-antitoxin system and a cold shock protein. There were 281 homoplastic non-recombined SNPs, of which 26 were convergently identified 6-18 times in 16 genes associated with quorum sensing, environmental response and virulence (**Extended Data Fig. 4** and **Supplementary Table 4**).

We constructed an ML tree and dated phylogeny for PC representative strains (**Fig. 2a**,**b**). By integration with hierarchical clustering (**Extended Data Fig. 5**), we split the PC into four sub-lineages corresponding to independent waves of cross-country transmission (**Fig. 2a-d**), which was also observed in the *V. cholerae* 7PET^5^. Following the nomenclature of *V. cholerae*, we designated these sub-lineages as wave-1 to wave-4. The estimated MRCA times for wave-1 to wave-3 were close (1984-1987), while wave-4 emerged more recently in 1998.

We identified 11 serotypes and three STs in representative strains (**Fig. 2b**). While O3:K6 was dominant (79%), serotype conversions were identified, particularly in wave-2 and wave-3. ST3 (96%) remained the predominant ST. Several marker genes/islands (VPaI-1, 4, 5 and *orf8*) were lost in 2%-8% of representative isolates, with the others being conserved (**Extended Data Fig. 6a**). Notably, the genomic island VPaI-7, which encodes *tdh* and T3SS2, showed high differentiation between different waves (**Extended Data Fig. 6b**). Additionally, we observed a conserved antimicrobial resistance (AMR) gene profile (**Extended Data Fig. 6a**). Apart from intrinsic ampicillin resistance-related genes, no recently acquired gene was identified in >1% of representative isolates.

### Wave succession and global transmission

Remarkably, we identified a clear ‘wave succession’ pattern: wave-1 to wave-4 succeeded each other temporally as the major wave in China, where sampling is sufficient. Similar trends were also seen in other countries with ≥10 representative strains, including the US, UK, Canada and Bangladesh, suggesting a possible global pattern. In contrast, in two South American countries, Peru and Chile, wave-1 has remained dominant for over a decade (**Fig. 2c**,**d**).

Wave-4 frequency has been increasing since its emergence, ultimately becoming predominant from 2012 onward. Since its origin, the effective population size (*N*_*e*_) of the PC increased exponentially until 1998 (**Fig. 2e**). Subsequently, *N*_*e*_ fluctuated and showed a decreasing trend after 2010. Population dynamics of waves was consistent with temporal dynamics, with the *N*_*e*_ peaking alternatively for different waves. Post-2010, the *N*_*e*_ of wave-4 exceeded others and remained stable, while *N*_*e*_ of other waves declined rapidly.

We inferred the geographical transmission history of the PC by phylogeographical analysis, focusing on cross-continent events. A total of 67 transmission events were identified, mostly transient and not leading to local colonization (**Extended Data Fig. 7**). Only six ‘TC’ events (Transmissions resulting in local Colonization that persisted for ≥3 years) were identified (**Fig. 2b** and **Extended Data Fig. 7**). Wave-1 in South America likely sourced from a single TC event from Asia to Peru in 1990 (TC1), much earlier than its first discovery in South America (1997). Subsequently, wave-1 spread from Peru to Chile and backward transmissions were also identified (TC2-4). Additionally, two TC events from Asia to North America were identified (TC5-6) in other waves.

### Puu-gene loss is associated with wave-4 replacement

To investigate the genetic basis of the wave-4 replacement of other waves, we comprehensively identified wave-4-specific variations and examined their functions. A total of 40 variations were identified (**Fig. 3a** and **Extended Data Table 1**), including 29 SNPs (located in 26 genes and 3 intergenic regions), two small indels (located in 2 genes) and most notably, the absence of nine adjacent genes (VP1775-VP1786). Of the nine genes, five are annotated as part of the putrescine utilization (Puu, VP1775-1781) pathway, one encodes a Na^+^/H^+^ antiporter (VP1782), and the others encode hypothetical or DUF3291 domain-containing proteins (VP1783-1786).

The fixation of gene loss is expected to be adaptive and selection-driven in free-living species, which are generally believed to have large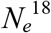. Moreover, *Vibrio* species, which are abundant in marine environments, have been broadly estimated to exhibit large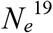. Therefore, we hypothesize that the fixation of Puu-pathway gene loss in *V. parahaemolyticus* confers phenotypic advantages.

The enzymes encoded by Puu-pathway genes degrade the polyamine putrescine into 4-aminobutanoate (GABA) or spermidine (**Fig. 3b**). Specifically, the *puuC* gene is involved in three sub-pathways in the conversion of putrescine to GABA, and its deletion is expected to completely block this conversion, mimicking the effect of a complete deletion of Puu-genes **(Fig. 3b)**. Therefore, we generated a *puuC* deletion mutant in the wildtype strain RIMD2210633 (wave-3, hereafter RIMD-W3) to explore the function and consequences of loss of Puu-genes in *V. parahaemolyticus*.

### Phenotypic effects of Puu-gene loss

#### Biofilm formation

We first tested the phenotypic effects of Puu-gene loss on biofilm formation, as the Puu-pathway has been reported to be involved biofilm formation in *Pseudomonas aeruginosa*^20^. As reported in *P. aeruginosa*, Δ*puuC* showed significantly (*p*<0.01) enhanced biofilm formation in M9 minimal media after 48 hours of culture (**Fig. 3c**), while complementation of the deletion strain *in trans* (C-Δ*puuC*) significantly decreased biofilm formation.

We next performed transcriptomic sequencing to identify differentially expressed genes (DEGs) that may underlie the increased biofilm of Δ*puuC*, as well as to explore other Puu-gene-loss related phenotypes. In line with increased biofilm formation of Δ*puuC*, we identified multiple DEGs between wildtype and Δ*puuC* associated with biofilm or biofilm-related pathways, including c-di-GMP and quorum sensing (**Fig. 3d, Supplementary Table 5**). We next quantified intracellular c-di-GMP levels in wildtype and Δ*puuC* during growth by introducing a dual-fluorescence reporter plasmid. Consistent with differential expression of several c-di-GMP-related enzymes and higher biofilm formation, c-di-GMP levels were significantly higher in Δ*puuC* compared to wildtype in stationary phase (**Fig. 3e**).

#### Cell adhesion and intestinal colonization

The differential expression analysis provided clues as to the effects of Puu-gene-loss on other phenotypes in addition to biofilm formation. Specifically, the most significant category of DEGs (Δ*puuC vs*. RIMD-W3) were genes encoding CupE-fimbriae and Flp-pili (**Fig. 3d)**, which are generally related to cell adhesion for pathogens^21^. Motivated by these observations, we performed Caco-2 cell adhesion assays and found significantly enhanced adhesion for Δ*puuC* compared to wildtype (**Fig. 3f**). We also tested whether increased cell adhesion might transform into enhanced intestinal colonization. We constructed bioluminescent wildtype and Δ*puuC* strains and traced their colonization of mice after oral gavage using *in vivo* imaging. At 5 hours post-infection, mice infected with Δ*puuC* exhibited stronger luminescence compared to wildtype (**Fig. 3g**). Bacteria were then rapidly cleared, with no detectable luminescence observed for both wildtype and mutant strains by 10 hours post-infection.

#### Virulence

An unexpected observation was the downregulation of the key virulence gene, *tdh*, in Δ*puuC* (2.6-fold, *p*-adj=0.02, **Fig. 3d**), suggesting that Puu-gene-loss might reduce virulence. Accordingly, we compared the lethality of wildtype and Δ*puuC* mutant strains in mice following intraperitoneal infection. Consistent with *tdh* downregulation, Δ*puuC* exhibited reduced lethality (**Fig. 3h**). These findings together revealed the potential impact of Puu-gene-loss on phenotypes related to human infection, including cell adhesion, intestinal colonization and virulence.

### Phenotypes of wave-4 strains

The phenotypic experiments above provide clues about the selective advantages of wave-4 strains. To link the phenotypes related to Puu-gene-loss observed in a Δ*puuC* wave-3 strain with the behavior of natural wave-4 strains, we next compared 12 natural strains, including six wave-1 to wave-3 (2 strains each) and six wave-4 strains, for the same phenotypes as described above. Consistent with the above phenotypic differences between wildtype and mutant strains established above, the wave-4 strains exhibited enhanced biofilms and cell adhesion, compared with wave-1 to wave-3 strains (**Fig. 4a**). Moreover, we complemented a wave-4 strain (VPCZ58) with the nine genes (VP1775-VP1786) that are absent in wave-4 strains and identified significantly reduced biofilm formation and adhesion compared to the wildtype strain (**Fig. 4b**), providing further evidence that the Puu-genes are responsible for the observed phenotypic differences.

**Figure 3.**
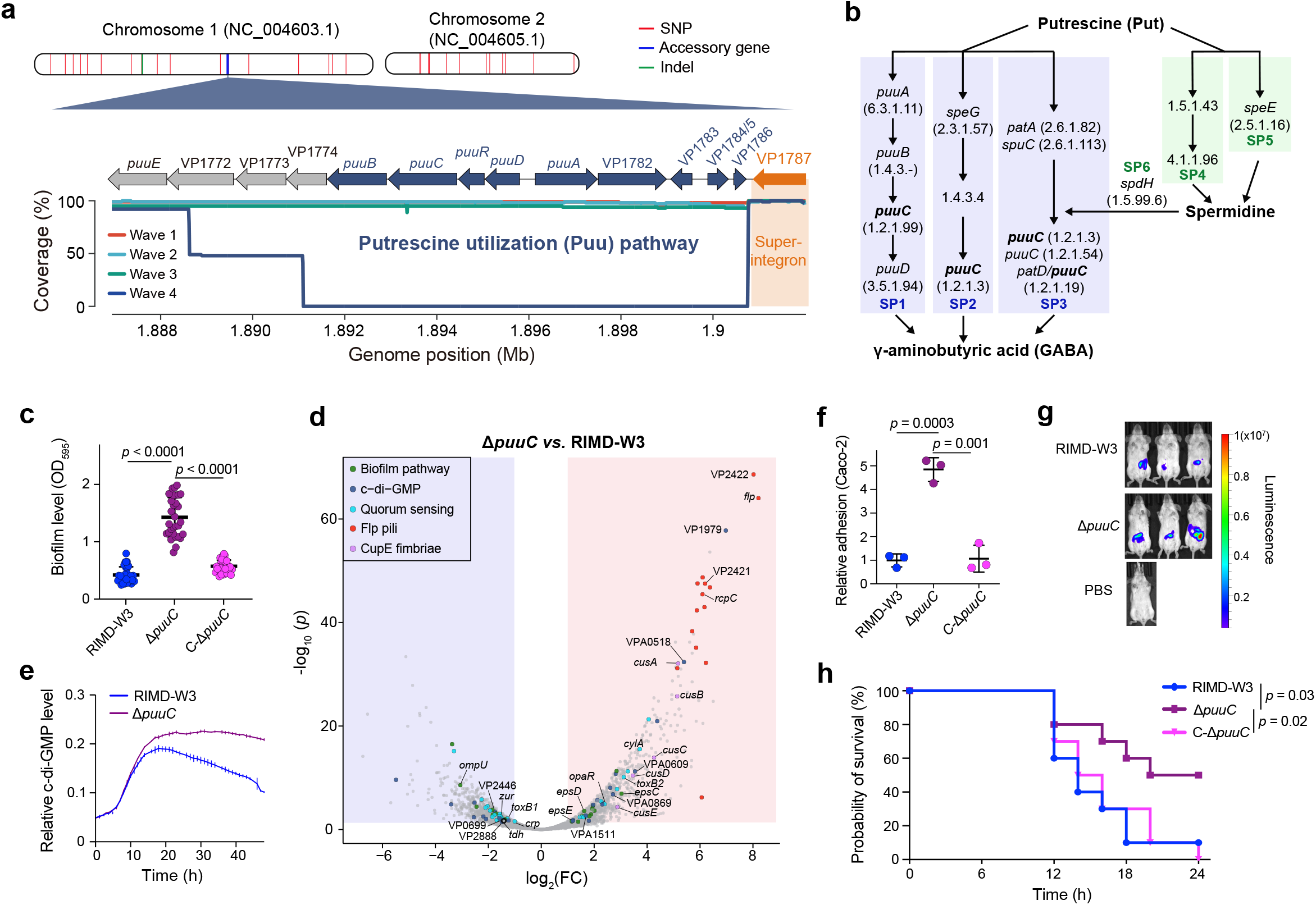
Phenotypic effects of gene-loss in the putrescine utilization (Puu) pathway. **a**, Positions of wave-4 specific variations on the reference genome (top) and gene structure and coverage of Puu-genes (colored in blue) in different waves (bottom). **b**, Summary of Puu-genes and metabolites based on KEGG annotations. The *puuC* gene, which appears in three sub-pathway, was highlighted in bold. **c**, Biofilm levels in M9 minimal media with 1% NaCl measured by optical density at 595 nm (OD_595_) of wildtype (RIMD-W3), mutant (Δ*puuC*) and complement strains (C-Δ*puuC*). **d**, Transcriptomic volcano plots showing differentially expressed genes (DEGs). The colors represent genes of specific category as in the legend. Red and blue backgrounds represent up- and down-regulated genes in Δ*puuC* compared to wildtype strain. **e**, c-di-GMP dynamics of wildtype (RIMD-W3) and mutant (Δ*puuC*) strains. **f**, The adhesion ability of strains to Caco-2 cells relative to wildtype (RIMD-W3). **g**, *In vivo* bioluminescent imaging of mice at 5 hours post-infection by gavage. **h**, Survival curves of mice by intraperitoneal infection. Mann–Whitney U tests (two-sided) were used for pairwise comparisons except for panel h, where log-rank (Mantel-Cox) tests were applied. *: *p*<0.05, **: *p*<0.01. Panels c, e, and f show mean ± SD as lines.

**Figure 4.**
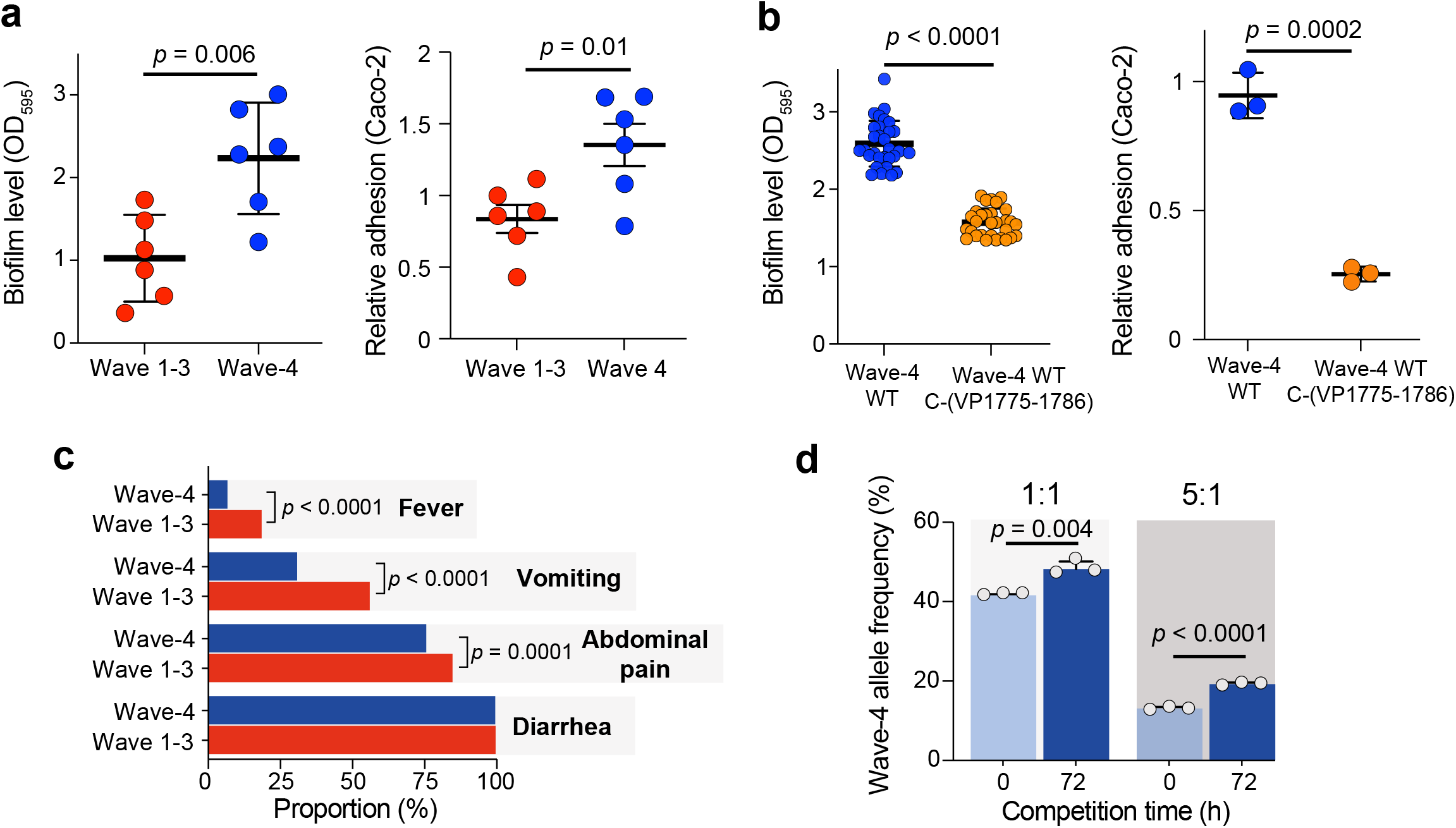
Phenotypes and clinical symptoms of wave1-3 and wave-4 strains. **a**, Biofilm level and adhesion ability between wave1-3 and wave-4 strains. **b**, Biofilm level and adhesion ability between the wave-4 wildtype strain (VPCZ58, wave-4 WT) and its complemented strain (wave-4 WT C-(VP1775-1786)) carrying the nine Puu-related genes (VP1775-1786). **c**, Clinical symptoms of wave-4 and other wave-associated patients. **d**, Wave-4 specific allele frequency after 72 hours of competition under seawater with chitin conditions. The light blue and dark blue bars represent the mixture of natural strains before and after competition, respectively. The Mann–Whitney U tests (two-sided) were used for pairwise comparisons, except for panel c, where Chi-squared tests were applied. *: *p*<0.05, **: *p*<0.01. Panels a, b, and d show mean ± SD as lines.

Additionally, we analyzed the clinical symptom data of 1,966 PC-related patients reported in our previous study^22^, and found that wave-4 was associated with milder symptoms, including significantly lower rates of fever, vomiting and abdominal pain (**Fig. 4c**), consistent with reduced virulence observed in our mouse infection experiment (**Fig. 3h**).

We hypothesized that these phenotypic differences mediated by Puu-gene-loss might also confer competitive advantages of wave-4 strains in the environment. Therefore, we performed competition experiments for natural strains in seawater with chitin as a nutrition source, mimicking a possible natural environment. We mixed wave-4 strains with those from other waves in equal (1:1) or different (1:5) ratios. By targeted PCR amplification of a gene with a wave-4-specific SNP allele (VPA0427 A919G), we quantified the relative abundance of wave-4 based on the frequency of the wave-4-specific allele. After 72 hours of competition, we measured a significant increase in the frequency of wave-4-specific alleles for both mixing ratios (**Fig. 4d**), demonstrating the competitive advantage of wave-4 under the tested condition.

### Puu-gene-loss in bacteria

The convergent loss of Puu-genes and/or its fixation in specific bacterial lineages would provide evidence supporting selection as the driving force. We analyzed the prevalence of Puu-genes among 108,526 genomes from 36 bacterial species, including *Vibrio*, common pathogens, and probiotics. Puu-genes were generally not conserved among bacteria, with part of the pathway lost in some species (**Fig. 5** and **Extended Data Table 2**). Notably, the absence/pseudogenization of Puu-genes in a subset of strains within species was convergently identified across multiple species (**Fig. 5b** and **Extended Data Fig. 8**). Furthermore, this absence/pseudogenization was fixed or nearly fixed in a specific lineage for species such as *Escherichia coli, Shigella boydii, S. sonnei* and *P. aeruginosa*, or convergently occurred in two or more lineages among species such as *Campylobacter coli, Yersinia pestis, Bifidobacterium longum* and *Lactobacillus helveticus*. Additionally, Puu-gene-loss was not only identified among *V. parahaemolyticus* PC strains, but also in the *V. cholerae* 7PET **(Fig. 5b)**.

To test if Puu-gene-loss has a similar phenotypic effect in other bacteria, we generated *puuC* deletion strains in *V. cholerae* (E7946 background) and *E. coli* (MG1655 background) and tested their biofilm formation and cell adhesion abilities. Consistent with the results in *V. parahaemolyticus*, the Δ*puuC* mutant strains in both species exhibited significantly enhanced biofilms and adhesion compared to their parental wildtype strains (**Fig. 5b**). Together, these results indicate that Puu-gene-loss is under selection in some species, and that the associated phenotypic consequences are potentially universal across bacterial species.

**Figure 5.**
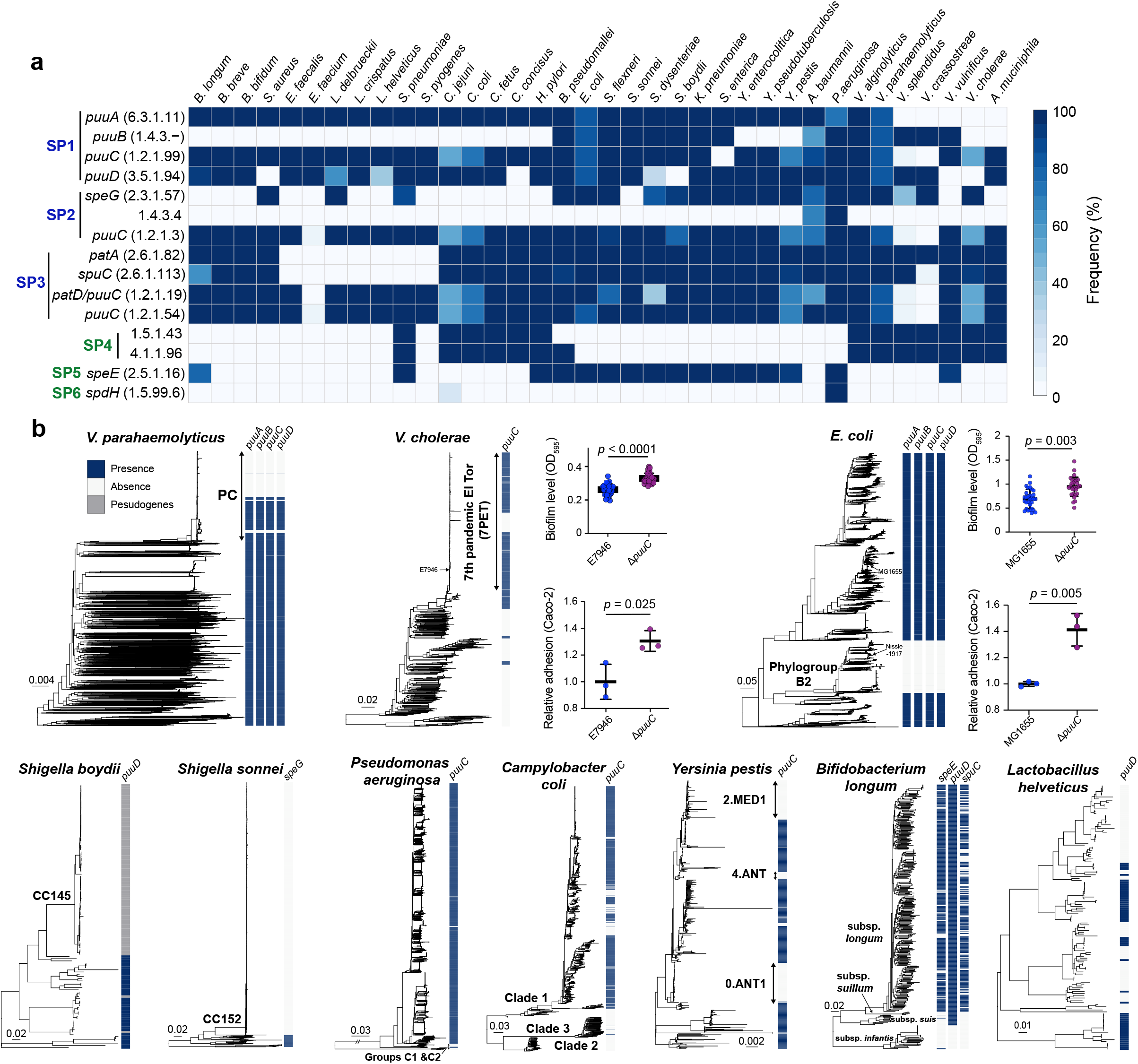
Puu-genes in 36 bacterial species. **a**, Prevalence of Puu-genes in 36 species. Genes and sub-pathways (SP) on the left are as in Fig. 3b. The colors (right) indicate the frequency of genes. **b**, Phylogenetic distribution of Puu-genes in ten species and the biofilm and adhesion of wildtype (E7946 and MG1655) and Δ*puuC* mutant strains in *V. cholerae* and *E. coli*. Mann–Whitney U tests (two-sided) were used for pairwise comparisons. *: *p*<0.05, **: *p*<0.01. Lines represent mean ± SD.

## Discussion

Leveraging a large-scale genome dataset of global isolates, we present the most comprehensive genomic view to date on the origin and evolution of the *V. parahaemolyticus* PC. Our findings suggest that the PC likely originated from a serotype O3:K6 pathogenic strain through gain of multiple genes/islands within a short timeframe (∼60 years) from *V. parahaemolyticus*, rather than from non-pathogenic strains or from other *Vibrio* species^11^. Notably, several characteristics previously considered to be specific to the PC^10–12^, including ST3, *toxRS*/new, and VPaI-1 to VPaI-3, were also present in pre-PC strains, while the *orf8* gene remained unique to the PC, thus remaining as a reliable marker. Moreover, we described the fine-scale population structure and spatiotemporal dynamics of the PC, and identified ‘wave succession’ with potential underlying molecular mechanisms. Together, our findings substantially enhance the current understanding of how epidemic lineages of *V. parahaemolyticus* emerge and evolve.

During the evolution of the *V. parahaemolyticus* PC, we observed intriguing parallels with the *V. cholerae* 7PET and other pathogens. First, we observed gain of genes/islands, including key virulence genes (*tdh* and *ctxAB*) and marker genomic islands (six VPaIs and two VSPs) for the *V. parahaemolyticus* PC, as has been reported for the *V. cholerae* 7PET^4,5^. This phenomenon has also been observed during human colonization by other pathogens^23^ and in the evolution of metabolic pathways across prokaryotic lineages^24^. Frequent human-mediated cross-regional transmission and adaptation to new environments are considered potential drivers for the *V. cholerae* 7PET^4^, which might also apply to the *V. parahaemolyticus* PC. Second, for both PCs from pandemic *Vibrio* species, the inferred times of origin (1983 and 1955^4^) and introduction to specific geographical regions (e.g., South America: 1990 and 1989^25^) were much earlier than their first epidemiological observations (1995 and 1961; South America: 1997 and 1991). Three possible explanations for this include inadequate surveillance, variability in the molecular clock rate^26^ with higher early mutation rates^27,28^, and the possibility of early cryptic disease epidemics, with specific events such as El Niño or warming in coastal areas triggering a sudden surge in cases and thereby drawing epidemiological attention. While the third explanation does not apply to *V. cholerae* due to the high risk and strict surveillance of cholera, all explanations are applicable to *V. parahaemolyticus*. Third, divergence into successive sublineages mediating waves of infections, identified in both pandemic *Vibrio* and other pathogens such as SARS-CoV-2^29^. While antimicrobial resistance likely drove the wave succession of *V. cholerae* 7PET^7^, *V. parahaemolyticus* exhibits low resistance rates to first-line antibiotics, indicating a different driving factor.

The importance of biofilm formation in bacterial environmental adaptation and survival under adverse conditions (e.g., antibiotic exposure) is well recognized^30^. Therefore, the enhanced biofilms mediated by Puu-gene-loss may improve the environmental adaptability of wave-4 strains, as suggested by our competition experiment results. Moreover, the ancestral strains of the PC, as well as most non-PC strains, carry the complete Puu-pathway genes, indicating that the loss of Puu-genes occurred recently, possibly in response to environmental changes. Putrescine serves multiple physiological functions in diverse organisms, including roles in cell growth and proliferation^31^, and has been particularly reported to promote acid stress tolerance^32,33^. The loss of Puu-genes might have been selected for by recent increases in ocean or coastal acidification.

Human activities are pivotal in pathogen transmission. Increased cell adhesion and intestinal colonization could directly facilitate human-mediated transmission, while reduced virulence, associated with milder clinical symptoms, may lower the likelihood of seeking medical attention, thereby promoting cryptic transmission. Notably, increased adhesion and colonization are not contradictory to reduced virulence, as different infection routes in the experiments (gavage *vs*. intraperitoneal injection) and their underlying mechanisms are known to differ^34^.

The effect of the Puu-pathway on biofilms is likely a broader phenomenon among bacteria than previously recognized. We have demonstrated that Puu-gene-loss enhances biofilms in three different species. Furthermore, Puu-gene-loss in *P. aeruginosa* also leads to enhanced biofilms^20^, and *E. coli* phylogroup B2, characterized by absence of *puuA*-*D* (**Fig. 5b**) exhibits a stronger biofilm-formation ability than other phylogroups^35,36^, providing further evidence for a more universal effect.

For the first time, we have established links between Puu-gene-loss, cell adhesion and intestinal colonization, which are also likely universal across bacteria. We demonstrated that Puu-gene-loss enhances cell adhesion in three different species and promotes intestinal colonization in *V. parahaemolyticus*. Furthermore, *E. coli* phylogroup B2 strains are often associated with urinary tract infections^37^ and show increased persistence in the intestines of human infants^38^, both of which is consistent with the hypothesis of increased adhesion and colonization mediated by Puu-gene-loss. Interestingly, the probiotic *E. coli* strain Nissle 1917, known for its strong capacity for human gut colonization^39^, belongs to phylogroup B2 and lacks Puu-genes (**Fig. 5b**).

The convergent loss or pseudogenization of Puu-gene within species (e.g., *C. coli*) and across different bacterial species, along with its lineage-specific fixation (e.g., in *E. coli*), indicate that these events are more likely selection-driven in these species and may confer fitness advantages. Puu-gene loss/pseudogenization was uniquely fixed in major pathogenic clonal complexes (CC) of *S. boydii* (CC145) and *S. sonnei* (CC152)^40^, and recently fixed in *V. parahaemolyticus* wave-4, suggesting potential fitness advantages for human infection or transmission. However, this appears to not always be the case, as Puu-gene loss/pseudogenization has not occurred in major pathogenic lineages of *C. coli* and *P. aeruginosa*. Additionally, Puu-gene loss/pseudogenization appears to be associated with lineage divergence in *E. coli* and *C. coli*, and Puu-gene gain is potentially linked to human adaptation in the probiotic *B. longum*, with Puu-genes predominantly identified in adult human strains and absent in infant or non-human strains. These observations together indicate that while Puu-gene loss/pseudogenization is likely selection-driven in some species, the underlying driving forces vary across different species.

While the significance of spontaneous mutation and gene acquisition in pathogen evolution is well recognized^41–43^, the role of gene loss remains underexplored. Gene loss is pervasive across all life kingdoms^44,45^. In bacteria, it is commonly observed in symbionts and host-dependent pathogens, where it is generally considered neutral and driven by genetic drift associated with transmission bottlenecks^46,47^, while loss of several specific genes has been experimentally validated to be associated with pathogen virulence^48–50^. Notably, gene loss affecting polyamine metabolism has also been reported in the pathogen *Shigella*, where the polyamine cadaverine was found to inhibit enterotoxin activity^48^. This aligns with our findings in *V. parahaemolyticus*, in which elevated levels of a different polyamine, putrescine, are associated with reduced virulence. Together, these findings suggest a potentially broader role for polyamines (e.g., cadaverine and putrescine) in modulating bacterial pathogenesis.

Free-living bacteria with a large *N*_*e*_ serve as better models for investigating adaptive evolution driven by gene loss, as such fixation is more likely to be adaptative and selection-driven. Some free-living bacteria, such as the abundant open-ocean bacterioplankton *Prochlorococcus*, exhibit extensive gene loss, which has been attributed to accelerated evolution and fitness advantages conferred by genomic and metabolic “streamlining”^51,52^. Moreover, previous studies of gene loss-mediated bacterial adaptation, whether for symbionts, host-dependent pathogens, or free-living species, generally come with the cost of transition from generalists capable of surviving in diverse environments to specialists adapted to specific ecological niches^49–52^. The exact niche of the *V. parahaemolyticus* PC remains unclear but it is more likely a generalist due to widespread distribution in marine and aquatic habitats. If we consider different geographical regions as macro-niches, the wave-4 strains characterized by Puu-gene-loss have spread globally, without clear evidence supporting its transition from a generalist to specialist strategy. Therefore, our study presents a novel pattern of gene loss-mediated adaptation in free-living species.

In summary, our study provides the most comprehensive genomic view on the origin and evolution of the *V. parahaemolyticus* PC. We identified several interesting common features between the evolution of pandemic clones in two *Vibrio* species, but the driving forces underlying wave succession events appear distinct. While spontaneous mutation and gene gain-mediated antibiotic resistance may drive the *V. cholerae* 7PET succession^7^, we demonstrated that Puu-gene loss, which confers phenotypic advantages related to environmental adaptation and human transmission, has likely driven the recent *V. parahaemolyticus* wave succession. An interesting finding is that *V. parahaemolyticus* wave succession has come along with reduced virulence, which aligns with the trade-off hypothesis of pathogen virulence evolution^53^. We also found that Puu-gene-loss is general and frequent among bacteria, selection-driven in multiple species, with its phenotypic impacts likely universal across species. Furthermore, Puu-gene-loss may play key roles in bacterial evolution, including lineage divergence (e.g., *E. coli, C. coli* and *B. longum*), human adaptation (e.g., *B. longum*) and the epidemic expansion of specific pathogenic lineages (e.g., *V. parahaemolyticus, S. boydii* and *S. sonnei*). Taken together, our study underscores the indispensable role of gene loss in the adaptative evolution of bacterial pathogens.

## Methods

### *V. parahaemolyticus* genome dataset

A total of 8,684 high-quality genomes were included in this study, including 333 newly sequenced and 8,351 publicly available genomes (**Supplementary Table 1**). Public genomes (up to November 2022) were downloaded from National Center for Biotechnology Information GenBank or Sequence Read Archive (SRA) database, with metadata extracted from corresponding BioSample information or the literature. New genomes were obtained from archived strains isolated from multiple countries, including China, Japan, Senegal, Peru and Chile.

Whole genome sequencing of new strains was performed using Illumina NovaSeq or Miseq platforms. New and public sequencing reads were trimmed using Trimmomatic v0.39^54^ and were *de novo* assembled using shovill v1.1.0 pipeline (https://github.com/tseemann/shovill). The quality assessment of genomes was performed using CheckM v1.1.3^55^. Whole genome average nucleotide identity (ANI) against the reference genome of *V. parahaemolyticus* (RIMD2210633, accession numbers: NC_004603.1 and NC_004605.1) was calculated using fastANI v1.1^56^. Only high-quality genomes (completeness >90% or contamination <5%) with ANI >95% were included in further analysis.

### Phylogenetic and phylogeographical analysis

Genome sequences were aligned against the reference genome using MUMmer v3.23^57^ to generate whole-genome alignments. Core-genome (regions present in >99% strains) single-nucleotide polymorphisms (SNPs) were identified using SNP-sites v2.5.1^58^. SNPs located in repetitive regions identified by TRF v4.07b^59^ or BLASTN self-search were excluded. Non-repetitive SNPs were used to construct maximum-likelihood (ML) phylogenetic trees using FastTree v2.1.10^60^ for all the isolates.

For both pre-PC and PC strains, recombination regions and associated length were detected using Gubbins v3.1.3^61^. Homoplastic SNPs and nutations (non-recombined SNPs) along the phylogenetic branch were identified using SNPPar v1.0^62^ based on ancestral state reconstruction of these variations. Non-recombined and non-homoplastic SNPs were used for the construction of ML tree using RAxML-NG v1.0.1^63^ with the GTR+Γ model. Phylogenetic trees were visualized using ggtree^64^. Hierarchical clustering analysis were performed using fastBAPS v1.0.6^65^ to assign strains into subgroups. The transitions in the isolation countries of strains in the phylogenetic tree are considered cross-country events. These were identified manually, and all events are shown in detail in Extended Data Fig. 7. TC was defined as Transmissions resulting in local Colonization (persisted for ≥3 years).

### Temporal analysis and population size dynamics

Temporal signals among pre-PC and PC isolates were assessed based on the recombination-stripped ML tree using TempEst v1.5.3^66^, by calculating the correlation between root-to-tip distances and strain isolation dates (years). A significant temporal signal was detected among strains from pre-PC3b to PC (*p*<0.01, *R*=0.62), whereas inclusion of pre-PC3a and earlier pre-PC strains eliminated the significant signal. A significant temporal signal was also detected within PC strains alone (*p*<0.01, *R*=0.57). For strains showing a significant temporal signal, we applied BactDating v1.1.0^67^ to infer the dated phylogenetic trees with the additive relaxed clock (ARC) model. The effective sample sizes (ESS) of the estimated parameters were all above 200. Effective population size dynamics was inferred using skygrowth v0.3.1^68^ based on dated trees with default parameters.

### Serotype and MLST typing

*In silico* O- and K-antigen genotyping was performed using the *V. parahaemolyticus* Kaptive database^69^. Multi-locus sequence typing (MLST) was performed using mlst (https://github.com/tseemann/mlst) to scan genome sequences against the PubMLST databases.

### Representative genome dataset of PC

We generated a representative subset of 942 PC isolates. First, to reduce the effect of repeat sampling such as strains from disease outbreaks, we selected the genome with maximum N50 from a cluster of strains with ≤6 SNPs from same country and within a year, representing putative outbreaks^22^. Second, we calculated root-to-tip distances of phylogenetic trees based on non-recombined SNPs, and removed putative hypermutators with extreme root-to-tip distances and branch lengths (**Extended Data Fig. 3**).

### Core-pangenome

All PC representative genomes were re-annotated using Prokka v1.13^70^, and the annotation outputs (GFF3 files) were used in Panaroo v1.2.8^71^ to identify the core-pangenome and gene presence/absence.

### Wave-4 specific variations

We analyzed three types of variations, including SNPs, gene presence/absence and unitigs. The SNPs and gene presence/absence data were extracted from the preceding analysis. The unitig matrix was determined using unitig-caller v1.2.1 (https://github.com/bacpop/unitig-caller). Unitig sequences were aligned against assembled genomes using BLASTN to identify corresponding genomic positions and related genes. Wave-4 specific variations were defined as alleles, genes or unitigs present/absent in >95% of wave-4 strains and <5% of non-wave-4 strains.

### *puuC* gene in the reference genome

The *puuC* gene (VP_RS08540/VP1776-7) in the reference genome RIMD2210633 was annotated as a pseudogene due to a frameshift caused by insertion of one nucleotide within a homopolymeric region (three consecutive adenines). However, we performed both Sanger sequencing and Illumina re-sequencing of this region in our RIMD2210633 strain. Both sequencing methods confirmed absence of this nucleotide and an intact *puuC* ORF in our strain, with no frameshift detected. Given the low sequencing depth of the original reference genome (average 8.9×) and the elevated error rate associated with homopolymeric regions, we hypothesize that the frameshift reported in the reference genome is likely caused by a sequencing error.

### Puu-genes in bacteria

We investigated the prevalence of Puu-genes in 36 bacterial species, including *Vibrio*, common pathogens, and probiotics. We included a total of 108,526 genomes (62-24,205 genomes per species, **Supplementary Table 6**) downloaded from Genome Taxonomy Database (GTDB, release 207)^72^. Puu-related protein sequences in *E. coli, P. aeruginosa* and *V. parahaemolyticus* were used as initial queries, which were searched against all genomes of the selected 36 species using TBLASTN to identify the reference genes of each species (**Extended Data Table 2**). Nucleotide and protein sequences of the reference genes for each species were then searched against all genomes of the corresponding species using BLASTN and TBLASTN. We combined BLASTN and TBLASTN results to identify the presence, absence or pseudogenization of genes, with identify >80% and coverage >70% as cutoffs. A gene was classified as present if a BLASTN or TBLASTN match was identified. If only a BLASTN match was identified, the gene was classified as a pseudogene; otherwise, it was considered absent.

### Bacterial strains and growth conditions

A total of 12 *V. parahaemolyticus* representative strains (pairwise SNP distances: >6) were used for laboratory experiments, including reference strain RIMD2210633 (wave-3) and 11 previously or newly sequenced PC strains. Additionally, *V. cholerae* strain E7946 (El Tor, Ogawa) and *E. coli* strain MG1655 (K12) were included. We chose *E. coli* and *V. cholerae* for two main reasons: our detection of natural strains with Puu-gene loss, and the availability of strains in our laboratory. In addition, *V. cholerae* is another *Vibrio* species associated with disease pandemics, and *E. coli* is a widely used model organism. *V. parahaemolyticus* strains were grown routinely in rich broth (LB with 3% NaCl) with shaking or on agar plates at 30**°**C aerobically. *E. coli* and *V. cholerae* strains were cultured in LB at 37°C.

### Mutant construction

Deletion mutants of *V. parahaemolyticus* were constructed by homologous recombination and counterselection with the suicide plasmid pDM4 as previously described^73^, with 1kb of homology upstream and downstream of the desired mutation. For *V. cholerae*, a similar approach with pDM4 was used as for *V. parahaemolyticus*, except *E. coli* MFD*pir* was used as donor strain for conjugation. *E. coli* deletion mutants were constructed using λ red recombination as previously described^74^. *V. parahaemolyticus* deletion strains were complemented with the corresponding gene *in trans* on plasmid pMMB207 using the Ptac promoter. All deletion and complement clones were verified by colony PCR. All constructed strains and plasmids are listed in **Supplementary Table 7**.

### Biofilm assays

Log-phase cultures were inoculated into M9 medium with 1% NaCl in 96-well plates (Kingmorn biotechnology) and incubated under static conditions at 37°C for 48 hours. Following incubation, culture medium was discarded, and 96-well plates were washed with 0.9% NaCl twice, dried at 60 °C for 30 min, and stained with 100 μl 0.1% (w/v) crystal violet solution for 10 min. After staining, the plates were washed with 0.9% NaCl three times. The bound crystal violet was resolubilized with 100 μl 95% ethanol for 10 min and quantified via OD_595_ with BioTek Synergy H1 (Agilent Technologies). Each strain was tested with 30 technical replicates.

### RNA extraction and transcriptome analysis

Strains were grown to stationary phase in M9 medium with 1% NaCl and harvested after treatment with stop mix (5% phenol, 95% ethanol). Total RNA was extracted from cells using the hot-phenol method as described previously^75^. RNA was DNase I-digested and rRNA depleted. cDNA libraries were constructed at Novogene Co. Ltd. (Beijing, China) and sequenced on an Illumina NovaSeq platform.

Sequencing reads were trimmed using Trimmomatic v0.39^54^ and then mapped to the reference genome using BWA v0.7.17^76^. The number of reads assigned to genes were counted using featureCounts v2.0.6^77^. Differentially expressed genes and pathway enrichment analysis were performed using R packages DESeq2 v1.34.0^78^ and ClusterProfiler v4.2.2^79^.

### c-di-GMP assay

To measure intracellular c-di-GMP, we used a previously published riboswitch-based reporter system^80^. Log-phase cultures of strains carrying the reporter were inoculated into M9 medium with 1% NaCl and 50 µg/ml spectinomycin in 96-well plates (six replicate wells per strain). Additionally, six wells of blank control with 200 µl uninoculated medium were included. The plate was incubated in a BioTek Synergy H1 plate reader (Agilent Technologies) at 37°C for 48 hours. OD_600_ and fluorescence (455 nm excitation/489 nm emission for AmCyan and 540 nm excitation/574 nm emission for TurboRFP) were measured every 15 min. The average values of the control wells were subtracted from each measurement (OD_600_, AmCyan, and TurboRFP) of the strains. The mean values were used to plot growth curves and the whisker length represents standard deviation.

### Cell adhesion

Log-phage strain cultures were washed with PBS and then resuspended in cell culture medium without antibiotics. Caco-2 cells grown in 24-well plates were infected with the bacterial suspension at an MOI of 10:1. After 45 min, cells were washed three times with PBS to remove the non-attached bacteria and then lysed with 0.1% saponin for 10 min. The number of cell-associated bacteria was determined by plating serial dilutions of the lysate on LB agar plates. The percentage of cell-associated bacteria was calculated as (recovered CFU/inoculated CFU) × 100%, and the results were presented as adhesion relative to wild-type strain (RIMD-W3).

### Mouse colonization

Female BALB/c mice aged 6-8 weeks were used to test strain colonization. Mice were randomly divided into three groups, including a control group and two groups inoculated with different strains (three mice each treated group). All mice were fasted for 8 hours prior to gavage.

We introduced a plasmid with a *lux* reporters (pWHQ205, encoding luciferase and substrate, constructed in this study) into two strains (RIMD-W3 and Δ*puuC*), respectively. Strains carrying *lux* reporters were cultured in LB medium with 3% NaCl supplemented with chloramphenicol (25 µg/ml) to logarithmic phase (OD_600_=0.6∼0.8) at 30°C. Strain concentration was then adjusted to 3×10^10^ CFU/ml with PBS, and each mouse was gavaged with 0.3 ml of the bacterial suspension. At 5 and 10 hours post-infection, the mice were anesthetized with 3% isoflurane for 5 minutes and exposed for 1 minute using the PerkinElmer IVIS Spectrum imaging system (E-Heng Import and Export Co. Ltd).

### Mouse lethality

Female BALB/c mice aged 4-5 weeks were used for lethality experiment, which were randomly divided into three groups treated by different strains, with each consisting of 10 mice. Log-phage strain cultures were washed three times with PBS and then adjusted to a concentration of 5×10^7^ CFU/ml with PBS. Mice were intraperitoneally injected with 100 μl of strain suspension, and the number of mice that died at a specific time point was record. The experimental protocols of colonization and lethality of mice were approved by the animal ethics committee of Shanghai Institute of Immunity and Infection (No. A2023011).

### Growth competition assay

Log-phage cultures of 12 natural strains, including six wave-1/2/3 and six wave-4 strains (**Supplementary Table 7**), were individually diluted to an OD_600_ of 0.08. Subsequently, the diluted cultures of wave-4 strains were mixed with those of wave-1 to wave-3 strains in equal (1:1) or different (1:5) ratios. Given the ecological relevance of chitin as a major nutrient source for *Vibrio* spp. in marine environments, and its role as a sole carbon source in aquatic habitats^81^, competitions were conducted in natural seawater (collected from Jinshan, Shanghai) supplemented with 0.5% chitin (Sigma). Mixed cultures were incubated at 30_°C under static conditions for 72_h.

Genomic DNA of harvested cultures was extracted using the TIANamp Bacteria DNA kit (Tiangen Biotech). PCR primers VPA0427SNPU (AGCGACGACGCCAGAGAAAG) and VPA0427SNPD (GGTGTCTAAAGTACGGTGCAG) were used to amplify the gene VPA0427, which contains a wave-4 specific SNP allele (A919G). The PCR amplification was performed for 30 cycles, with the parameters: 1) 3 min. at 95°C, 2) 30 sec. at 95°C, 3) 30 sec. at 58°C, 4) 45 sec. at 72°C, 5) repeat steps 2-4 for 30 cycles, 6) 5 min. at 72°C.

The PCR amplification products were used for library construction at Novogene Co. Ltd. (Beijing, China) and sequenced on an Illumina NovaSeq platform. Sequencing reads are trimmed using Trimmomatic v0.39^54^ and were then mapped to *V. parahaemolyticus* reference genome using BWA v0.7.17^76^. The frequency of the wave-4 specific allele was calculated to represent the relative abundance of wave-4 strains.

### Statistical analysis

Datasets were compared using the Mann-Whitney U test for continuous data and Chi-squared tests for categorical data. Log-rank (Mantel-Cox) tests were used for survival curve comparison. p-values less than 0.05 were considered statistically significant.

## Data availability

The sequencing data have been deposited in the NCBI SRA or GenBank database under accession number PRJNA1117214 and PRJNA1062747. Background information of sequenced isolate is listed in Supplementary Tables 1-2. Source data are provided with this paper.

## Acknowledgements

This work was supported by National Key Research and Development Program of China (No. 2022YFC2304700 to F.D., 2022YFD2101500 to H.W.), National Natural Science Foundation of China (No. 32270003 and No. 32000008 to C.Y., No. 32170640 and No.32211550014 to D.F., No. 32250610209 to S.S., 82030099 to H.W.), Youth Innovation Promotion Association, Chinese Academy of Sciences (No. 2022278 to C.Y.), Ministerio de Ciencia e Innovación (Spain) Grant PID2021-127107NB-I00 (J.M.U.), Generalitat de Catalunya Grant 2021 SGR 00526 (J.M.U.), Shanghai Rising-Star Program (No. 23QA1410500 to C.Y.), Shanghai Public Health System Construction Three-Year Action Plan (No. GWVI-11.1-43 to H.W.), Wuxi Science and Technology Development Fund’s ‘Light of Taihu Lake’ Science and Technology Research Program (Basic Research) (No. K20231033 to C.N., No. K20231045 to S.G.), and Innovative research team of high-level local universities in Shanghai. We are grateful to the labs of Prof. Qiyao Wang, Kim Orth, and Jin He for providing strains and plasmids. We thank Prof. Lei Pan for valuable comments.

## Author contributions

C.Y., J.M.-U. and D.F. designed, initiated and coordinated the study. J.M.-U., L.X, Y.L., M.J., X.S. and Q.H. contributed to data collection. C.Y., Z.J., J.W., and W.X. performed bioinformatics analysis. H.Q, S.S., C.N., S.G., Z.J., H.W., Y.Q. and S.L. performed experiments. All authors contributed to interpretation of the data. C.Y. wrote the first draft of the paper and H.Q., S.S., Y.Z., Z.Z., Y.C., J.M.-U., H.W., R.Y., Y.C. and D.F. reviewed and revised the paper. All authors read and approved the final manuscript.

