## Extended Data Figures for "Wave succession in the pandemic clone of *Vibrio parahaemolyticus* driven by gene loss"

**Extended Data Figure 1. Phylogenetic tree of pre-pandemic (pre-PC) strains and sequence coverage of virulence-associated loci and pandemic clone (PC) marker genes/genomic islands.**

Heatmap colors to the right indicate BLASTN-based sequence coverage of each loci, gene or island.

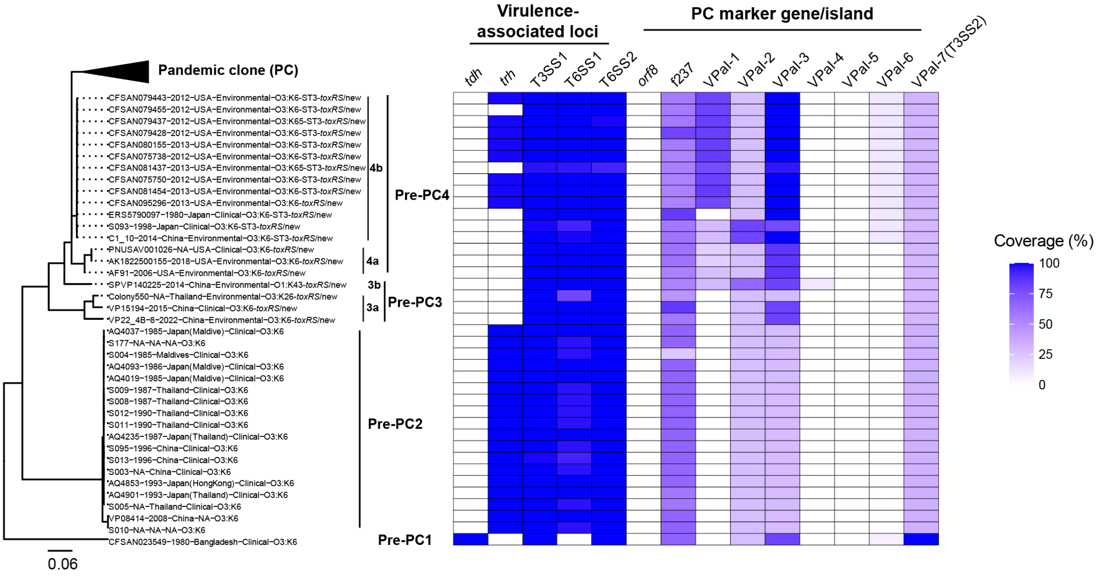

**Extended Data Figure 2. Phylogenetic distribution of virulence-associated loci and pandemic clone (PC) marker genes/genomic islands among PC, pre-PC, and non-PC strains.**

Virulence-associated loci: *tdh*, T3SS1, T6SS1, and T6SS2. PC marker genes/genomic islands: *toxRS*/new, *orf8*, f237, and VPaI-1 to VPaI-7. Bar colors on the right indicate strain classification (PC, pre-PC, or non-PC).

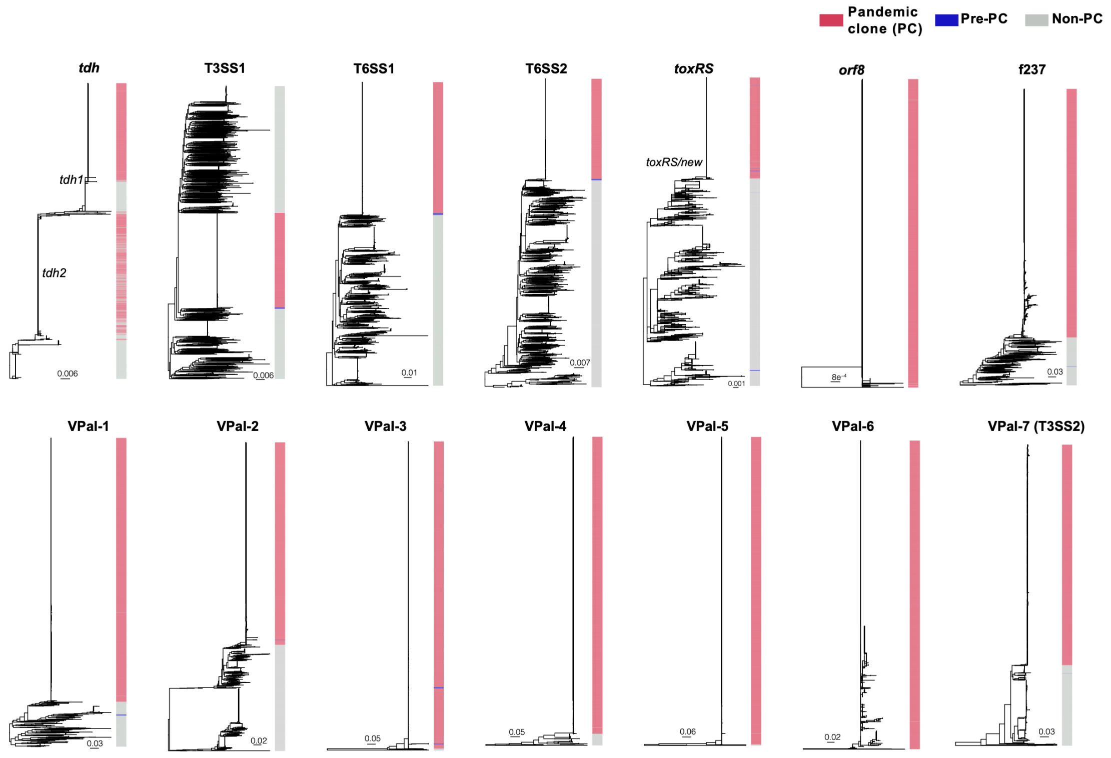

**Extended Data Figure 3. Tempo-geographical distribution of pandemic clone (PC) strains and putative hypermutators.**

a) Temporal distribution of PC strains. b) Geographical distribution of PC strains. Colors in panels a and b indicate strain classifications as shown in the legend at the top. c) Root-to-tip distances (left) and phylogenetic branch lengths (right) of putative hypermutator strains. Putative hypermutators are highlighted in red.

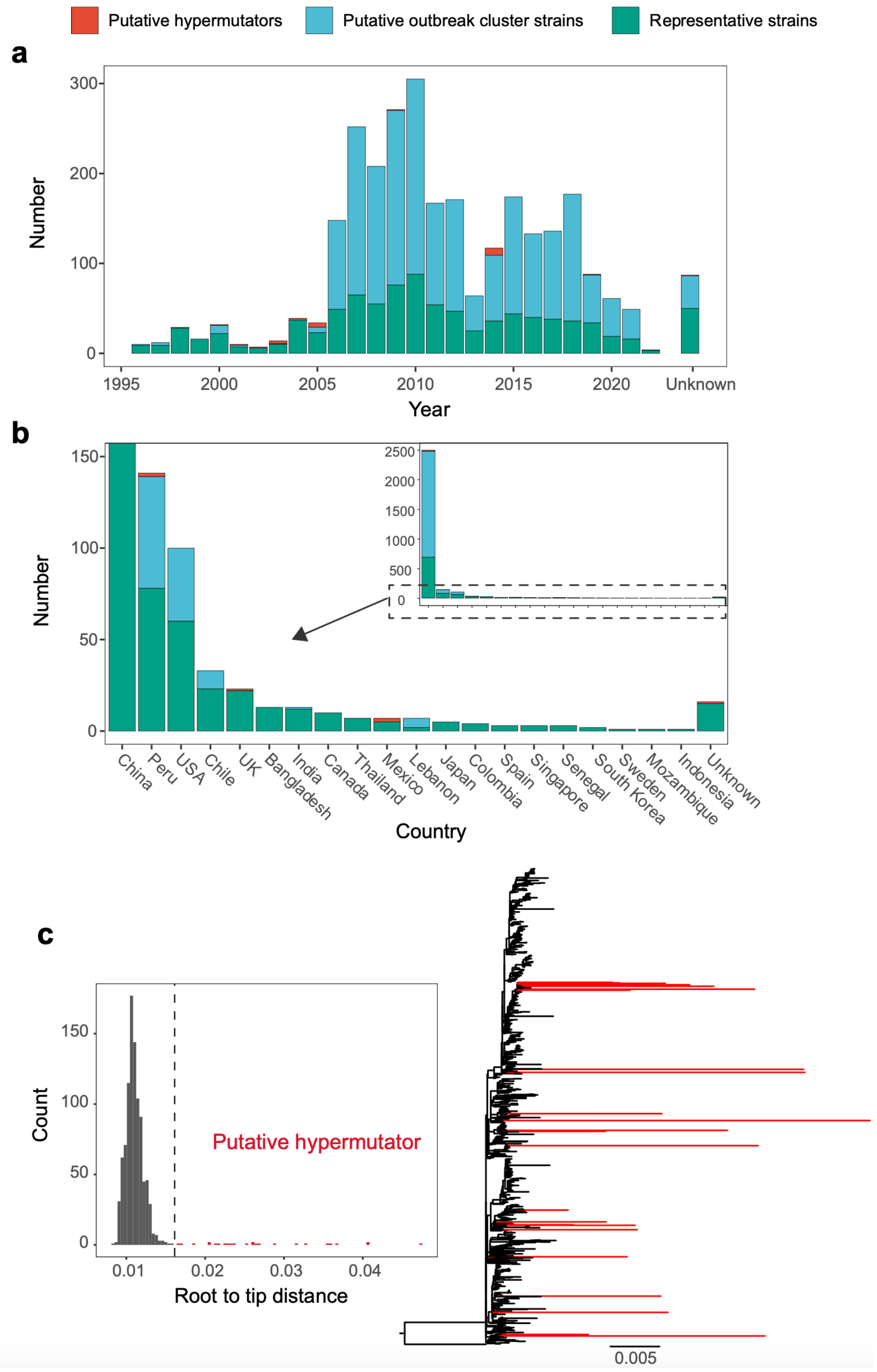

**Extended Data Figure 4. Distribution of homologous recombination events (a) and homoplastic SNP in pandemic clone (PC) strains (b).**

Phylogenetic trees based on non-recombined and non-homoplastic SNPs are shown on the left. In panel a, the top box summarizes the number and classification of recombination events (see legend at the top), while the bottom box displays their distribution across strains/nodes, represented by red bars. In panel b, bar colors indicate the status of SNPs or gaps, as defined in the legend at the bottom.

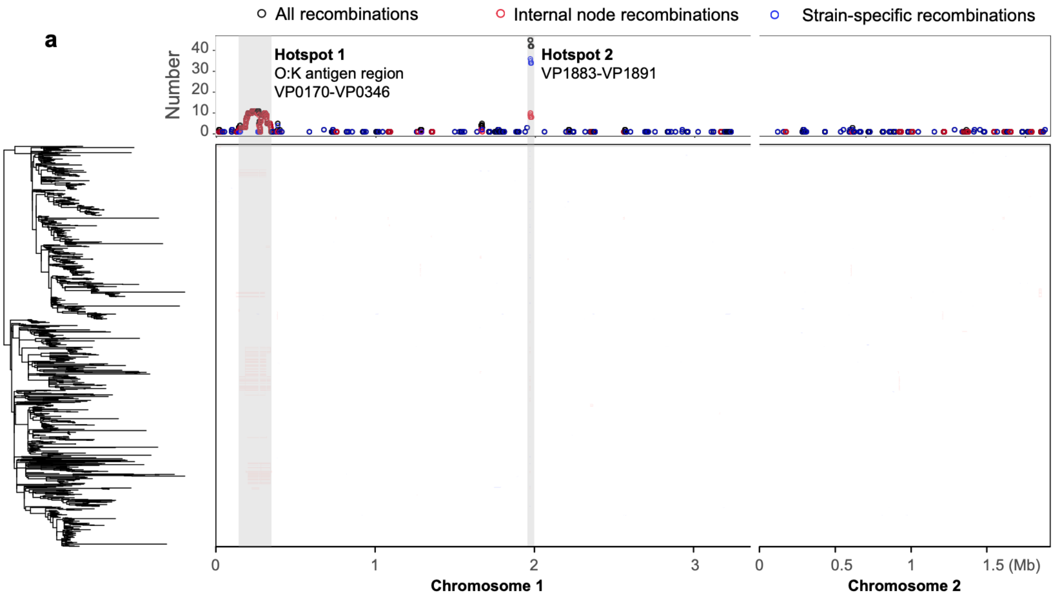


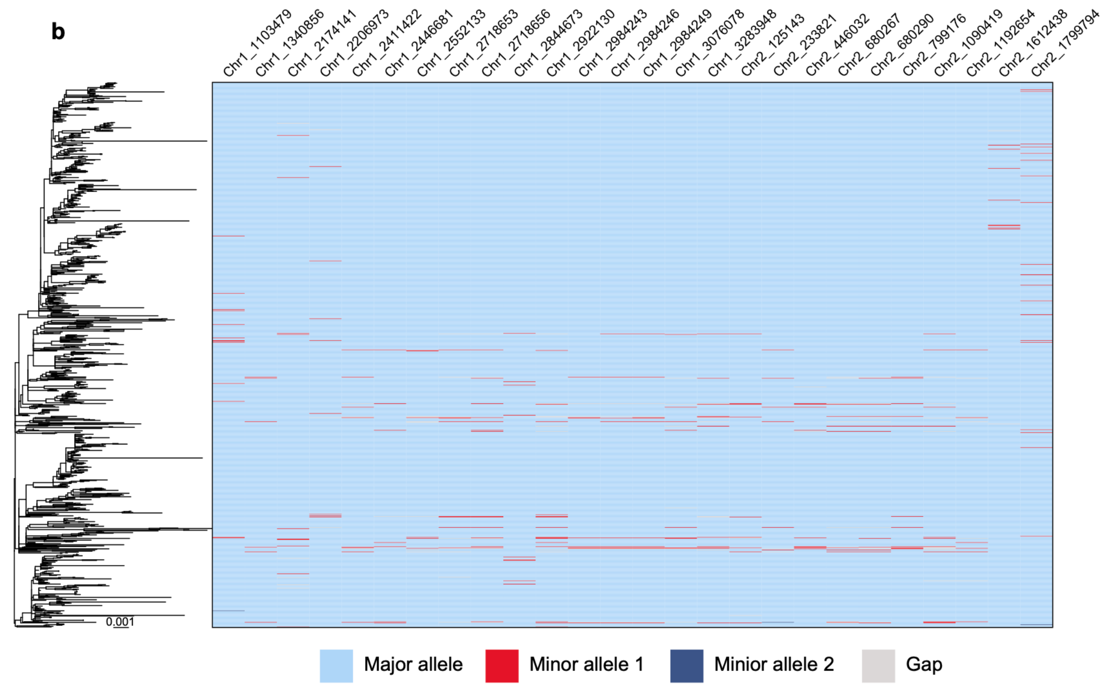

**Extended Data Figure 5. Maximum likelihood (ML) phylogeny and fastBAPS hierarchical clustering of representative pandemic clone (PC) strains.**
The branch colors of the ML tree on the left indicate the four waves defined in this study. The blue bars on the right display the fastBAPS hierarchical clustering results of PC strains.

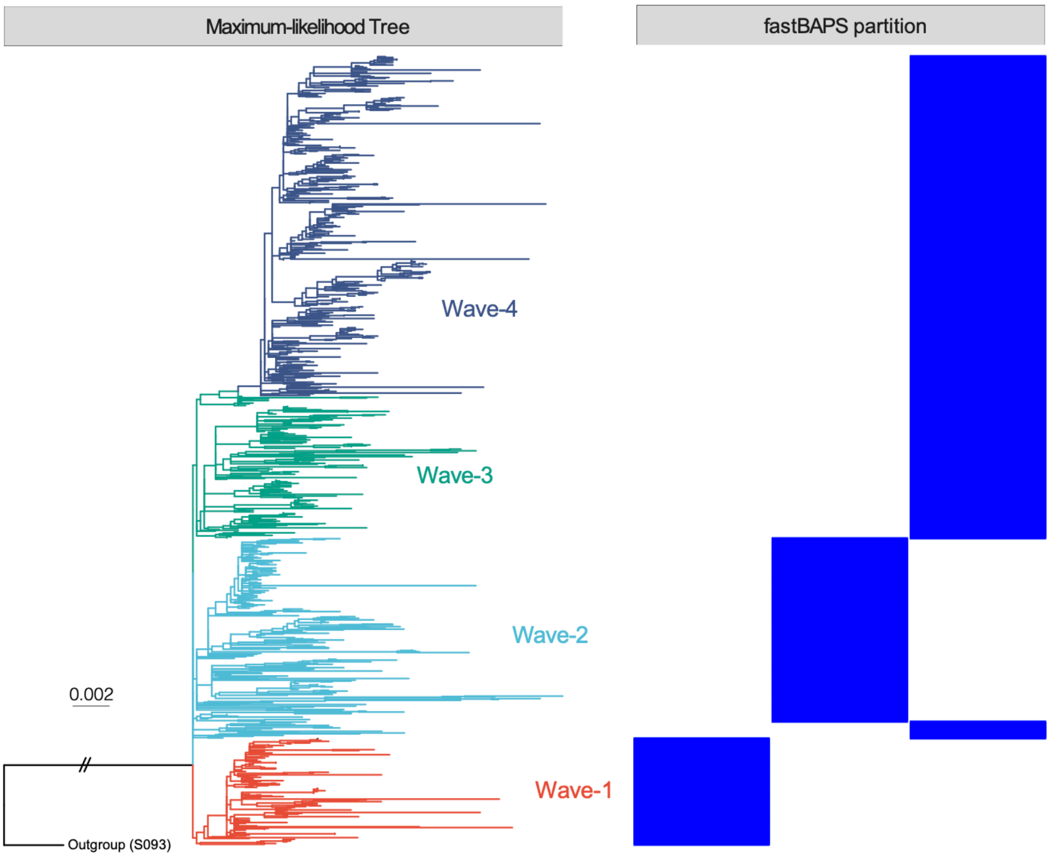

**Extended Data Figure 6. a) Phylogenetic distribution of pandemic clone (PC) marker genes, virulence-associated loci, antimicrobial resistance genes, and other accessory genes. b) Phylogenetic tree of virulence-associated loci and PC marker genes/islands.**

Branch colors in the ML tree (panel a) indicate the four waves defined in this study. Blue and white bars on the right denote the presence or absence of each locus/gene/island. In panel b, the bar colors on the right represent strain classification (four waves). Wave-specific variations are highlighted.

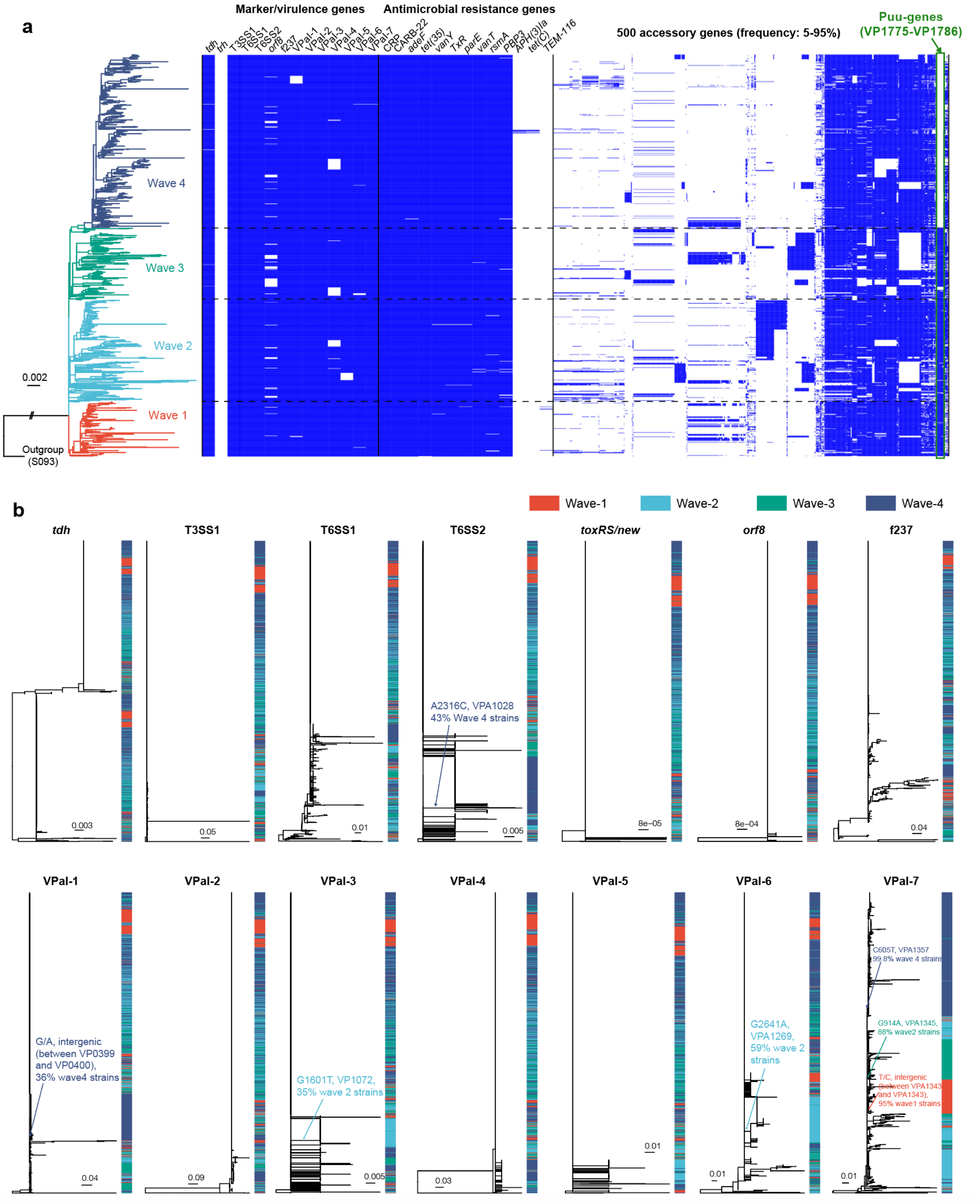

**Extended Data Figure 7. Global transmission events of the pandemic clone.**

a) Summary of global transmission events. Dashed lines represent transient transmission not leading to local colonization. Solid lines represent transmissions resulting in local colonization (TC). The colors indicate different waves. Pie charts indicate the composition of waves of countries. The size of circles scale with the number of representative strains. b) Inferred transmission events (T1-T67) of different waves. The branch colors indicate the geographical regions of strains. Specific transmission events are labeled on the dated phylogenetic tree.

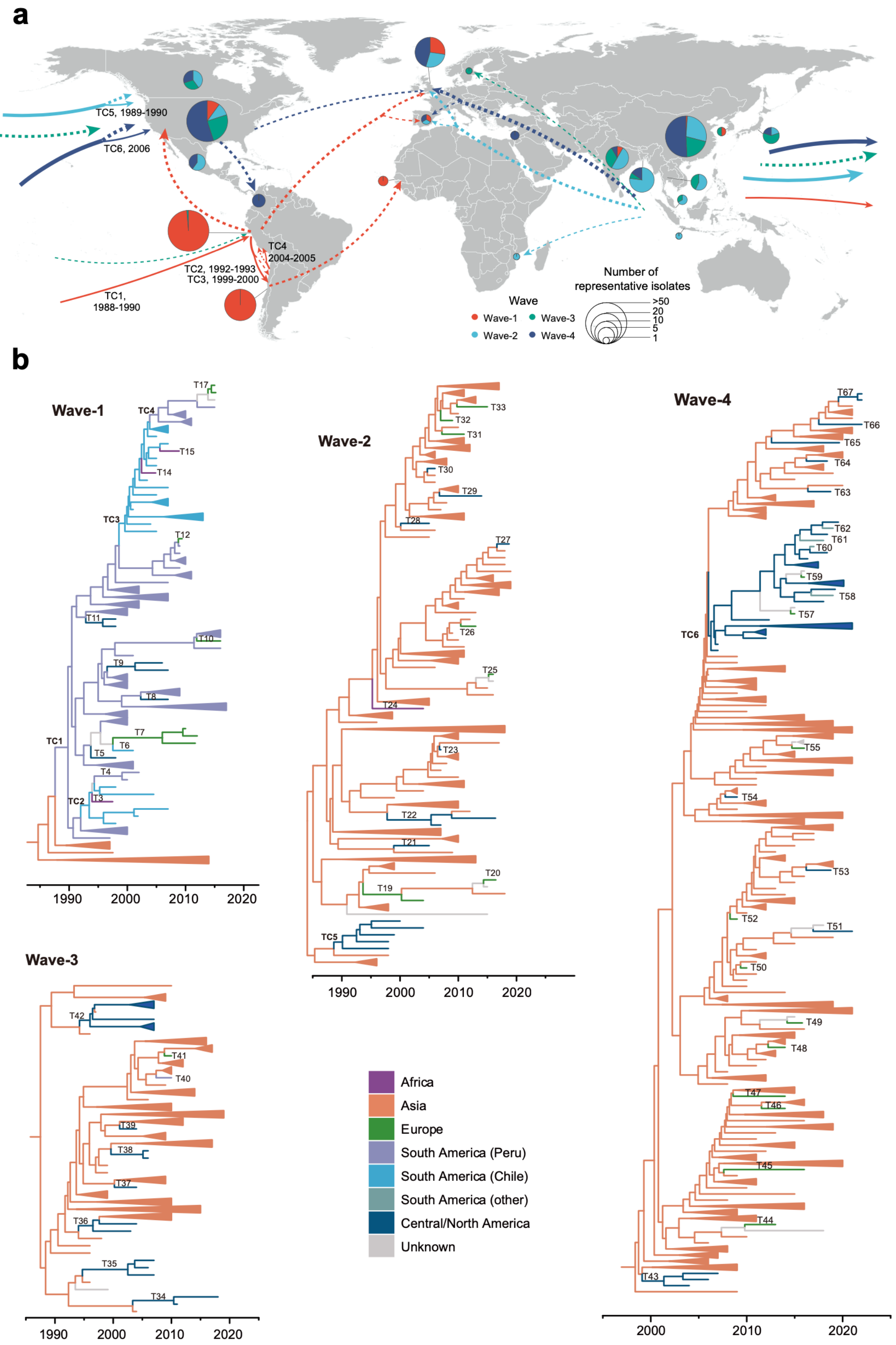

**Extended Data Fig. 8. Phylogenetic trees based on core-genome SNPs and distribution of Puu-genes in 10 bacterial species.**

The bar colors on the right side of the tree indicate the presence, absence, or pseudogenization of Puu-genes, as shown in the legend. Double slashes indicate artificially shortened branches.

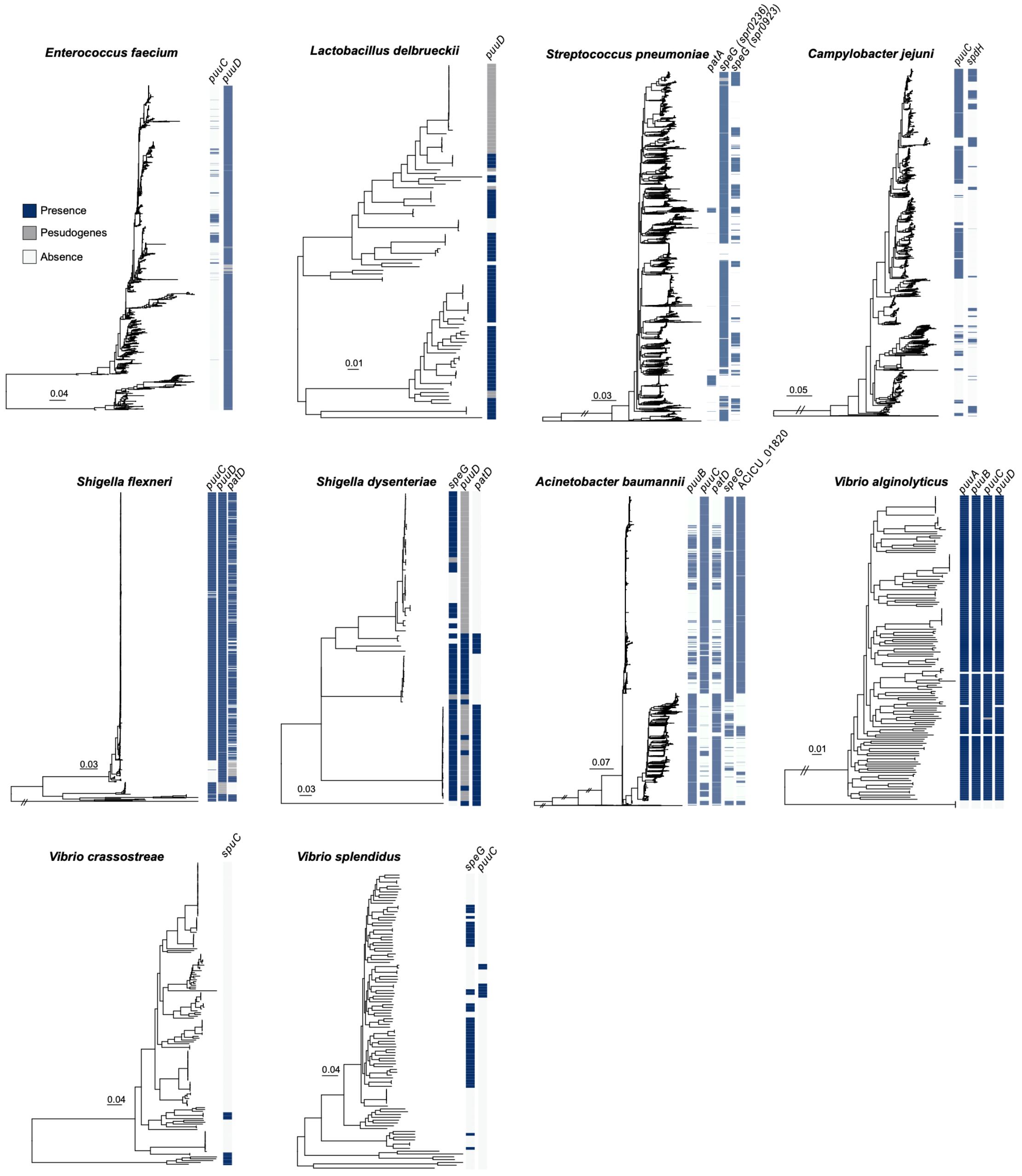
