## Supplementary Table 1 for "Wave succession in the pandemic clone of *Vibrio parahaemolyticus* driven by gene loss"

### Supplementary Table 1: Metadata and Genomic Characteristics of 8684 *Vibrio parahaemolyticus* Genomes.

| ID | Isolate_identifiers | BioSample | Accession | Classfication | Data source | Year | Country | Strain source | Reference coverage | Completeness(%) | Contamination(%) | Average nucleotide identity (ANI) | Genome length(bp) | GC% | No. of contigs | N50 |
|---|---|---|---|---|---|---|---|---|---|---|---|---|---|---|---|---|
| 004-02_NA_Peru_Clin_Clin | 004-02 | SAMN40891098 | PRJNA1062747 | PC | This study | 2002 | Peru | Clinical | 98.6 | 100 | 0.03 | 99.9904 | 5172805 | 45.12 | 89 | 198175 |
| 020-02_NA_Peru_Clin_Clin | 020-02 | SAMN40891099 | PRJNA1062747 | PC | This study | 2002 | Peru | Clinical | 98.6 | 100 | 0 | 99.9924 | 5183728 | 45.12 | 57 | 392233 |
| 056-01_NA_Peru_Clin_Clin | 056-01 | SAMN40891102 | PRJNA1062747 | PC | This study | 2001 | Peru | Clinical | 98.6 | 100 | 0 | 99.9907 | 5098297 | 45.29 | 59 | 334356 |
| 085-02_NA_Peru_Clin_Clin | 085-02 | SAMN40891103 | PRJNA1062747 | PC | This study | 2002 | Peru | Clinical | 98.7 | 100 | 0 | 99.9937 | 5102272 | 45.29 | 60 | 312811 |
| 182-02_NA_Peru_Clin_Clin | 182-02 | SAMN40891104 | PRJNA1062747 | PC | This study | 2002 | Peru | Clinical | 98.6 | 100 | 0 | 99.9906 | 5177250 | 45.12 | 61 | 336355 |
| 210_NA_Chile_NA_NA | ATC210 | SAMN40891111 | PRJNA1062747 | PC | This study | NA | Chile | NA | 98.7 | 100 | 0 | 99.992 | 5100712 | 45.29 | 53 | 401005 |
| 224_NA_Chile_NA_NA | ATC224 | SAMN40891107 | PRJNA1062747 | PC | This study | NA | Chile | NA | 97.3 | 100 | 0 | 99.9929 | 5032200 | 45.31 | 61 | 268333 |
| 240-02_NA_Peru_Clin_Clin | 240-02 | SAMN40891105 | PRJNA1062747 | PC | This study | 2002 | Peru | Clinical | 98.7 | 100 | 0 | 99.9948 | 5101837 | 45.29 | 58 | 334356 |
| 275-99_NA_Peru_Clin_Clin | 275-99 | SAMN40891642 | PRJNA1062747 | PC | This study | 1999 | Peru | Clinical | 98.6 | 100 | 0.14 | 99.9874 | 5210577 | 45.07 | 78 | 223831 |
| 276-99_NA_Peru_Clin_Clin | 276-99 | SAMN40891093 | PRJNA1062747 | PC | This study | 1999 | Peru | Clinical | 98.8 | 100 | 0 | 99.9942 | 5104628 | 45.29 | 57 | 347413 |
| 278-99_NA_Peru_Clin_Clin | 278-99 | SAMN40891094 | PRJNA1062747 | PC | This study | 1999 | Peru | Clinical | 98.6 | 100 | 0 | 99.9875 | 5180717 | 45.13 | 57 | 340345 |
| 357-99_NA_Peru_Clin_Clin | 357-99 | TBD | PRJNA1062747 | PC | This study | 1999 | Peru | Clinical | 98.7 | 100 | 0 | 99.9923 | 5189051 | 45.11 | 60 | 334356 |
| 39_5_NA_Chile_NA_NA | PMC39.5 | SAMN40891110 | PRJNA1062747 | PC | This study | 2005 | Chile | NA | 98.7 | 100 | 0 | 99.9944 | 5103511 | 45.29 | 55 | 461882 |
| 405-00_NA_Peru_Clin_Clin | 405-00 | SAMN40891095 | PRJNA1062747 | PC | This study | 2000 | Peru | Clinical | 98 | 99.73 | 0 | 99.9811 | 5071641 | 45.28 | 58 | 304117 |
| 429-00_NA_Peru_Clin_Clin | 429-00 | SAMN40891096 | PRJNA1062747 | PC | This study | 2000 | Peru | Clinical | 98.6 | 100 | 0.01 | 99.9874 | 5237354 | 45.32 | 58 | 427280 |
| 461-00_NA_Peru_Clin_Clin | 461-00 | SAMN40891090 | PRJNA1062747 | PC | This study | 2000 | Peru | Clinical | 98.8 | 100 | 0 | 99.994 | 5104271 | 45.29 | 61 | 289565 |
| 462-00_NA_Peru_Clin_Clin | 462-00 | SAMN40891091 | PRJNA1062747 | PC | This study | 2000 | Peru | Clinical | 98.6 | 100 | 0 | 99.9896 | 5098843 | 45.29 | 59 | 334356 |
| 463-01_NA_Peru_Clin_Clin | 463-01 | TBD | PRJNA1062747 | PC | This study | 2001 | Peru | Clinical | 98.8 | 100 | 0.01 | 99.9957 | 5193381 | 45.32 | 54 | 542852 |
| 48_1_NA_Chile_NA_NA | 48_1 | TBD | PRJNA1062747 | PC | This study | 2001 | Chile | NA | 98.6 | 100 | 0 | 99.991 | 5097840 | 45.29 | 54 | 312788 |
| 49_5-V2_NA_Chile_NA_NA | 49_5-V2 | TBD | PRJNA1062747 | PC | This study | 2005 | Chile | NA | 98.7 | 100 | 0 | 99.988 | 5103103 | 45.29 | 61 | 312826 |
| 511-00_NA_Peru_Clin_Clin | 511-00 | TBD | PRJNA1062747 | PC | This study | 2000 | Peru | Clinical | 98.6 | 100 | 0.01 | 99.9943 | 5188426 | 45.32 | 58 | 461842 |
| 518-01_NA_Peru_Clin_Clin | 518-01 | TBD | PRJNA1062747 | PC | This study | 2001 | Peru | Clinical | 98.4 | 100 | 0 | 99.9857 | 5085458 | 45.28 | 83 | 152539 |
| 568-00_NA_Peru_Clin_Clin | 568-00 | SAMN40891100 | PRJNA1062747 | PC | This study | 2000 | Peru | Clinical | 98.7 | 100 | 0 | 99.9927 | 5102914 | 45.29 | 53 | 312835 |
| 57_5_NA_Chile_NA_NA | PMC57.5 | SAMN40891639 | PRJNA1062747 | PC | This study | 2005 | Chile | NA | 98.7 | 100 | 0 | 99.989 | 5106457 | 45.29 | 56 | 427605 |
| 572-01_NA_Peru_Clin_Clin | 572-01 | SAMN40891097 | PRJNA1062747 | PC | This study | 2001 | Peru | Clinical | 98.6 | 100 | 0 | 99.9935 | 5175923 | 45.13 | 56 | 427036 |
| 61_NA_Chile_Amb_Amb | 61 | TBD | PRJNA1062747 | PC | This study | NA | Chile | NA | 98.6 | 100 | 0 | 99.9933 | 5097953 | 45.29 | 54 | 433028 |
| 61-V2_NA_Chile_Amb_NA | 61-V2 | TBD | PRJNA1062747 | PC | This study | NA | Chile | NA | 98.7 | 100 | 0 | 99.994 | 5101970 | 45.29 | 53 | 505531 |
| 698-99_NA_Peru_Clin_Clin | 698-99 | SAMN40891643 | PRJNA1062747 | PC | This study | 1999 | Peru | Clinical | 98.5 | 99.57 | 0.18 | 99.9855 | 5098461 | 45.29 | 154 | 71939 |
| 706-00_NA_Peru_Clin_Clin | 706-00 | SAMN40891101 | PRJNA1062747 | PC | This study | 2000 | Peru | Clinical | 98.7 | 100 | 0 | 99.987 | 5104047 | 45.29 | 56 | 347630 |
| 763-97_NA_Peru_Clin_Clin | 763-97 | SAMN40891084 | PRJNA1062747 | PC | This study | 1997 | Peru | Clinical | 98.7 | 100 | 0.01 | 99.9943 | 5193685 | 45.32 | 56 | 312677 |
| 776-00_NA_Peru_Clin_Clin | 776-00 | TBD | PRJNA1062747 | PC | This study | 2000 | Peru | Clinical | 98.6 | 100 | 0 | 99.9942 | 5139531 | 45.33 | 53 | 334356 |
| 780-98_NA_Peru_Clin_Clin | A10_780-98 | SAMN40891081 | PRJNA1062747 | PC | This study | 1998 | Peru | Clinical | 98.5 | 100 | 0 | 99.9918 | 5098463 | 45.29 | 53 | 461842 |
| 784-98_NA_Peru_Clin_Clin | 784-98 | SAMN40891082 | PRJNA1062747 | PC | This study | 1998 | Peru | Clinical | 98.7 | 100 | 0.14 | 99.9929 | 5217388 | 45.07 | 58 | 428132 |
| 790-97_NA_Peru_Clin_Clin | 790-97 | SAMN40891085 | PRJNA1062747 | PC | This study | 1997 | Peru | Clinical | 98.7 | 100 | 0 | 99.9942 | 5149414 | 45.24 | 60 | 334356 |
| 859-07_NA_Peru_Clin_Clin | 859-07 | SAMN40891108 | PRJNA1062747 | PC | This study | 2007 | Peru | Clinical | 98.7 | 100 | 0 | 99.9911 | 5102814 | 45.29 | 54 | 433108 |
| 875-97_NA_Peru_Clin_Clin | 875-97 | SAMN40891086 | PRJNA1062747 | PC | This study | 1997 | Peru | Clinical | 98.8 | 100 | 0 | 99.9948 | 5104770 | 45.29 | 64 | 289565 |
| 948-97_NA_Peru_Clin_Clin | 948-97 | SAMN40891088 | PRJNA1062747 | PC | This study | 1997 | Peru | Clinical | 98.6 | 99.91 | 0 | 99.9898 | 5184291 | 45.11 | 62 | 340201 |
| 971-98_NA_Peru_Clin_Clin | 971-98 | SAMN40891083 | PRJNA1062747 | PC | This study | 1998 | Peru | Clinical | 98.6 | 100 | 0 | 99.9954 | 5095988 | 45.29 | 57 | 334356 |
| AQ4019_NA_Japan_NA_NA | AQ4019 | TBD | PRJNA1062747/PRJNA1117214 | PrePC | This study | 1985 | Japan(Maldive) | Clinical | 93.5 | 100 | 0.07 | 99.2257 | 5054735 | 45.35 | 49 | 451451 |
| AQ4037_NA_Japan_NA_NA | AQ4037 | TBD | PRJNA1062747/PRJNA1117214 | PrePC | This study | 1985 | Japan(Maldive) | Clinical | 93.6 | 100 | 0.07 | 99.2317 | 5060240 | 45.35 | 52 | 458868 |
| AQ4093_NA_Japan_NA_NA | AQ4093 | TBD | PRJNA1062747/PRJNA1117214 | PrePC | This study | 1986 | Japan(Maldive) | Clinical | 93.6 | 100 | 0.07 | 99.2452 | 5055806 | 45.35 | 50 | 458740 |
| AQ4235_NA_Japan_NA_NA | AQ4235 | TBD | PRJNA1062747/PRJNA1117214 | PrePC | This study | 1987 | Japan(Thailand) | Clinical | 93.7 | 100 | 0 | 99.2288 | 5101465 | 45.33 | 51 | 388603 |
| AQ4853_NA_Japan_NA_NA | AQ4853 | TBD | PRJNA1062747/PRJNA1117214 | PrePC | This study | 1993 | Japan(HongKong) | Clinical | 93.7 | 100 | 0 | 99.2489 | 5035089 | 45.35 | 54 | 459125 |
| AQ4901_NA_Japan_NA_NA | AQ4901 | SAMN40891092 | PRJNA1062747/PRJNA1117214 | PrePC | This study | 1993 | Japan(Thailand) | Clinical | 93.6 | 100 | 0 | 99.2548 | 5035633 | 45.35 | 54 | 388418 |

| | | | | | | | | | | | | | | | | |
|---|---|---|---|---|---|---|---|---|---|---|---|---|---|---|---|---|
| C15_124-07__NA_Peru_Clin_Clin | C15_124-07 | TBD | PRJNA1062747 | PC | This study | 2007 | Peru | Clinical | 98.7 | 100 | 0 | 99.9936 | 5186851 | 45.13 | 52 | 432839 |
| C20_NA_Peru_Clin_Clin | C20 | TBD | PRJNA1062747 | PC | This study | NA | Peru | Clinical | 97.5 | 100 | 0 | 99.9306 | 5091241 | 45.3 | 44 | 472737 |
| C21_1262-07__NA_Peru_Clin_Clin | C21_1262-07 | TBD | PRJNA1062747 | PC | This study | 2007 | Peru | Clinical | 98.5 | 100 | 0 | 99.9908 | 5173425 | 45.12 | 59 | 347446 |
| LS326_NA_Senegal_Amb_Prawns | LS326 | TBD | PRJNA1062747/PRJNA1117214 | PC | This study | NA | Senegal | Env | 98.7 | 100 | 0 | 99.9931 | 5101784 | 45.29 | 67 | 348596 |
| LS328_NA_Senegal_Amb_Prawns | LS328 | TBD | PRJNA1062747/PRJNA1117214 | PC | This study | NA | Senegal | Env | 98.7 | 100 | 0 | 99.9876 | 5102566 | 45.29 | 53 | 506556 |
| LS329_NA_Senegal_Amb_Prawns | LS329 | TBD | PRJNA1062747/PRJNA1117214 | PC | This study | NA | Senegal | Env | 98.6 | 99.95 | 0 | 99.9938 | 5097825 | 45.29 | 53 | 366679 |
| MS104_NA_Senegal_Amb_Mussel | MS104 | TBD | PRJNA1062747/PRJNA1117214 | NonPC | This study | NA | Senegal | Env | 91.9 | 100 | 0.05 | 98.2868 | 5145944 | 45.23 | 39 | 522200 |
| PM57_7_NA_Chile_Amb_Amb | PM57 | TBD | PRJNA1062747 | PC | This study | 2007 | Chile | Env | 98.6 | 100 | 0 | 99.9937 | 5099140 | 45.29 | 55 | 461842 |
| PMC1_7_NA_Chile_NA_NA | PMC1 | TBD | PRJNA1062747 | PC | This study | 2007 | Chile | NA | 98.6 | 100 | 0 | 99.9924 | 5092496 | 45.29 | 55 | 461881 |
| PMC16_7_NA_Chile_NA_NA | PMC16.7 | SAMN40891106 | PRJNA1062747 | PC | This study | 2007 | Chile | NA | 98.5 | 100 | 0 | 99.993 | 5091206 | 45.28 | 73 | 267905 |
| PMC18_7_NA_Chile_NA_NA | PMC18 | TBD | PRJNA1062747 | PC | This study | 2007 | Chile | NA | 98.7 | 100 | 0 | 99.9934 | 5103372 | 45.29 | 59 | 334356 |
| PMC19_7_NA_Chile_Amb_Amb | PMC19 | TBD | PRJNA1062747 | PC | This study | 2007 | Chile | Env | 98.6 | 100 | 0 | 99.991 | 5100980 | 45.29 | 61 | 427031 |
| PMC28_7_NA_Chile_Amb_Amb | PMC28.7 | SAMN40891662 | PRJNA1062747 | PC | This study | 2007 | Chile | Env | 98.6 | 100 | 0 | 99.9895 | 5104595 | 45.29 | 55 | 432726 |
| PMC29_7_NA_Chile_NA_NA | PMC29 | TBD | PRJNA1062747 | PC | This study | 2007 | Chile | NA | 98.8 | 100 | 0 | 99.9937 | 5100848 | 45.29 | 54 | 461881 |
| PMC50_7_NA_Chile_NA_NA | PMC50.7 | SAMN40891660 | PRJNA1062747 | PC | This study | 2007 | Chile | NA | 98.6 | 100 | 0 | 99.9896 | 5098103 | 45.29 | 54 | 432726 |
| PMC51_7_NA_Chile_NA_NA | PMC51 | TBD | PRJNA1062747 | PC | This study | 2007 | Chile | NA | 98.8 | 100 | 0 | 99.9943 | 5102910 | 45.29 | 55 | 461881 |
| PMC60_7_NA_Chile_Amb_Amb | PMC60 | TBD | PRJNA1062747 | PC | This study | 2007 | Chile | Env | 98.6 | 100 | 0 | 99.9927 | 5097856 | 45.29 | 59 | 394767 |
| PMC70_7_NA_Chile_NA_NA | PMC70.7 | SAMN40891663 | PRJNA1062747 | PC | This study | 2007 | Chile | NA | 98.7 | 100 | 0 | 99.9886 | 5102025 | 45.28 | 59 | 505531 |
| PMC73_7_NA_Chile_Amb_Amb | PMC73.7 | SAMN40891109 | PRJNA1062747 | PC | This study | 2007 | Chile | Env | 98.6 | 100 | 0 | 99.9906 | 5096747 | 45.29 | 56 | 312654 |
| Vp_Peru_498-01_Peru_2001_C | 498-01 | SAMN40891645 | PRJNA1062747 | PC | This study | 2001 | Peru | Clinical | 98.1 | 99.73 | 0 | 99.9883 | 5078164 | 45.27 | 85 | 312502 |
| Vp_Peru_568-01_Peru_2001_C | 568-01 | SAMN40891646 | PRJNA1062747 | PC | This study | 2001 | Peru | Clinical | 98.4 | 100 | 0 | 99.9912 | 5170362 | 45.12 | 64 | 288851 |
| Vp_Peru_706_Peru_2000_C | 706 | TBD | PRJNA1062747 | PC | This study | 2000 | Peru | Clinical | 98.4 | 100 | 0.14 | 99.9788 | 5207845 | 45.07 | 63 | 288853 |
| Vp_Peru_CFSAN062237_2000_Peru_C | CFSAN062237 | TBD | PRJNA1062747 | PC | This study | 2000 | Peru | Clinical | 98.3 | 100 | 0 | 99.9648 | 5154819 | 45.27 | 71 | 250941 |
| Vp_Peru_CFSAN062238_2000_Peru_C | CFSAN062238 | TBD | PRJNA1062747 | PC | This study | 2000 | Peru | Clinical | 98.4 | 100 | 0.14 | 99.9885 | 5217527 | 45.07 | 59 | 288853 |
| Vp_Peru_CFSAN062240_2007_Peru_C | CFSAN062240 | TBD | PRJNA1062747 | PC | This study | 2007 | Peru | Clinical | 98.2 | 100 | 0 | 99.9811 | 5082137 | 45.3 | 42 | 312466 |
| Vp_Peru_CFSAN062241_2000_Peru_C | CFSAN062241 | TBD | PRJNA1062747 | PC | This study | 2000 | Peru | Clinical | 94 | 100 | 0.09 | 99.9941 | 5130185 | 45.37 | 57 | 312374 |
| Vp_Peru_CFSAN062243_2000_Peru_C | CFSAN062243 | TBD | PRJNA1062747 | PC | This study | 2000 | Peru | Clinical | 98.4 | 100 | 0 | 99.991 | 5171428 | 45.12 | 59 | 288853 |
| Vp_Peru_CFSAN062244_2000_Peru_C | CFSAN062244 | TBD | PRJNA1062747 | PC | This study | 2000 | Peru | Clinical | 98.4 | 100 | 0 | 99.9838 | 5091911 | 45.29 | 60 | 288853 |
| Vp_Peru_CFSAN062245_2000_Peru_C | CFSAN062245 | TBD | PRJNA1062747 | PC | This study | 2000 | Peru | Clinical | 98.4 | 100 | 0.07 | 99.9903 | 5097932 | 45.27 | 69 | 288853 |
| Vp_Peru_CFSAN062251_2007_Peru_C | CFSAN062251 | TBD | PRJNA1062747 | PC | This study | 2007 | Peru | Clinical | 97.3 | 100 | 0 | 99.9377 | 5094357 | 45.29 | 47 | 347272 |
| Vp_Peru_CFSAN062252_2007_Peru_C | CFSAN062252 | TBD | PRJNA1062747 | PC | This study | 2007 | Peru | Clinical | 98.2 | 100 | 0 | 99.9794 | 5159908 | 45.13 | 67 | 271828 |
| Vp_Peru_CFSAN062256_2000_Peru_C | CFSAN062256 | TBD | PRJNA1062747 | PC | This study | 2000 | Peru | Clinical | 95.4 | 100 | 0 | 99.9908 | 5094324 | 45.29 | 58 | 288853 |
| Vp_Peru_CFSAN062257_2000_Peru_C | CFSAN062257 | TBD | PRJNA1062747 | PC | This study | 2000 | Peru | Clinical | 98.4 | 100 | 0 | 99.9874 | 5171777 | 45.11 | 53 | 312582 |
| Vp_Peru_CFSAN062258_2007_Peru_C | CFSAN062258 | TBD | PRJNA1062747 | PC | This study | 2007 | Peru | Clinical | 98.5 | 100 | 0 | 99.9872 | 5177129 | 45.13 | 58 | 288856 |
| Vp_Peru_CFSAN062267_2009_Peru_C | CFSAN062267 | TBD | PRJNA1062747 | PC | This study | 2009 | Peru | Clinical | 98.5 | 100 | 0 | 99.9811 | 5175993 | 45.13 | 72 | 288855 |
| Vp_Peru_CFSAN062268_2007_Peru_C | CFSAN062268 | TBD | PRJNA1062747 | PC | This study | 2007 | Peru | Clinical | 98.2 | 100 | 0 | 99.9901 | 5175166 | 45.12 | 60 | 312381 |
| Vp_Peru_CFSAN062269_2000_Peru_C | CFSAN062269 | TBD | PRJNA1062747 | PC | This study | 2000 | Peru | Clinical | 98.4 | 100 | 0.09 | 99.9846 | 5093442 | 45.29 | 66 | 201346 |
| Vp_Peru_CFSAN062270_2000_Peru_C | CFSAN062270 | TBD | PRJNA1062747 | PC | This study | 2000 | Peru | Clinical | 98.4 | 100 | 0 | 99.9908 | 5182042 | 45.1 | 59 | 312388 |
| Vp_Peru_CFSAN062271_2015_Peru_C | CFSAN062271 | TBD | PRJNA1062747 | PC | This study | 2015 | Peru | Clinical | 98.4 | 100 | 0.09 | 99.9905 | 5244617 | 45.27 | 69 | 153020 |
| Vp_Peru_CFSAN062272_2010_Peru_C | CFSAN062272 | TBD | PRJNA1062747 | PC | This study | 2010 | Peru | Clinical | 98.4 | 100 | 0 | 99.9895 | 5162450 | 45.13 | 78 | 199729 |
| Vp_Peru_CFSAN062278_2010_Peru_C | CFSAN062278 | TBD | PRJNA1062747 | PC | This study | 2010 | Peru | Clinical | 98.5 | 100 | 0 | 99.994 | 5099091 | 45.29 | 59 | 312374 |
| Vp_Peru_CFSAN062279_2015_Peru_C | CFSAN062279 | TBD | PRJNA1062747 | PC | This study | 2015 | Peru | Clinical | 98.3 | 100 | 0.09 | 99.9897 | 5233847 | 45.28 | 82 | 161001 |
| Vp_Peru_CFSAN062297_2010_Peru_C | CFSAN062297 | TBD | PRJNA1062747 | PC | This study | 2010 | Peru | Clinical | 98.4 | 100 | 0 | 99.9848 | 5175162 | 45.11 | 60 | 312452 |
| Vp_Peru_CFSAN062298_2015_Peru_C | CFSAN062298 | TBD | PRJNA1062747 | PC | This study | 2015 | Peru | Clinical | 98.4 | 100 | 0.01 | 99.9813 | 5258136 | 45.15 | 71 | 267908 |
| Vp_Peru_CFSAN062299_2015_Peru_C | CFSAN062299 | TBD | PRJNA1062747 | PC | This study | 2015 | Peru | Clinical | 98.4 | 100 | 0.01 | 99.9897 | 5259564 | 45.15 | 64 | 312395 |
| Vp_Peru_CFSAN062301_2016_Peru_C | CFSAN062301 | TBD | PRJNA1062747 | PC | This study | 2016 | Peru | Clinical | 97.3 | 99.55 | 0.28 | 99.9888 | 5060053 | 45.28 | 372 | 32692 |
| Vp_Peru_CFSAN062308_2010_Peru_C | CFSAN062308 | TBD | PRJNA1062747 | PC | This study | 2010 | Peru | Clinical | 97.3 | 100 | 0.01 | 99.8893 | 5291884 | 45.3 | 176 | 246412 |
| Vp_Peru_CFSAN062318_2000_Peru_C | CFSAN062318 | TBD | PRJNA1062747 | PC | This study | 2000 | Peru | Clinical | 98.4 | 100 | 0.09 | 99.9935 | 5130078 | 45.38 | 64 | 288855 |

| | | | | | | | | | | | | | | | | |
|---|---|---|---|---|---|---|---|---|---|---|---|---|---|---|---|---|
| Vp_Peru_CFSAN062324_2010_Peru_C | CFSAN062324 | TBD | PRJNA1062747 | PC | This study | 2010 | Peru | Clinical | 98.4 | 100 | 0 | 99.9799 | 5165450 | 45.12 | 52 | 312374 |
| Vp_Peru_CFSAN062327_2015_Peru_C | CFSAN062327 | TBD | PRJNA1062747 | PC | This study | 2015 | Peru | Clinical | 98.4 | 100 | 0.01 | 99.9784 | 5260434 | 45.15 | 56 | 394629 |
| Vp_Peru_CFSAN062329_2010_Peru_C | CFSAN062329 | TBD | PRJNA1062747 | PC | This study | 2010 | Peru | Clinical | 98.5 | 100 | 0 | 99.9839 | 5099560 | 45.29 | 60 | 289055 |
| Vp_Peru_CFSAN062331_2010_Peru_C | CFSAN062331 | TBD | PRJNA1062747 | PC | This study | 2010 | Peru | Clinical | 98.4 | 100 | 0 | 99.992 | 5094730 | 45.29 | 60 | 314066 |
| Vp_Peru_CFSAN062333_2013_Peru_C | CFSAN062333 | TBD | PRJNA1062747 | PC | This study | 2013 | Peru | Clinical | 98.4 | 100 | 0.09 | 99.9893 | 5352219 | 45.19 | 195 | 280995 |
| Vp_Peru_CFSAN062337_2010_Peru_C | CFSAN062337 | TBD | PRJNA1062747 | PC | This study | 2010 | Peru | Clinical | 98.5 | 100 | 0 | 99.9856 | 5168389 | 45.13 | 65 | 288857 |
| Vp_Peru_CFSAN062348_2016_Peru_C | CFSAN062348 | TBD | PRJNA1062747 | PC | This study | 2016 | Peru | Clinical | 98.4 | 99.73 | 0.09 | 99.9805 | 5235149 | 45.27 | 70 | 288853 |
| Vp_Peru_CFSAN062349_2016_Peru_C | CFSAN062349 | TBD | PRJNA1062747 | PC | This study | 2016 | Peru | Clinical | 98.4 | 100 | 0.09 | 99.9879 | 5238650 | 45.26 | 69 | 294787 |
| Vp_Peru_CFSAN062351_2015_Peru_C | CFSAN062351 | TBD | PRJNA1062747 | PC | This study | 2015 | Peru | Clinical | 98.4 | 100 | 0.09 | 99.9819 | 5246267 | 45.27 | 60 | 288971 |
| Vp_Peru_CFSAN062354_2010_Peru_C | CFSAN062354 | TBD | PRJNA1062747 | PC | This study | 2010 | Peru | Clinical | 98.4 | 100 | 0.12 | 99.9939 | 5096580 | 45.28 | 61 | 267685 |
| Vp_Peru_CFSAN062355_2010_Peru_C | CFSAN062355 | TBD | PRJNA1062747 | PC | This study | 2010 | Peru | Clinical | 98.5 | 100 | 0 | 99.9869 | 5168836 | 45.13 | 56 | 294788 |
| Vp_Peru_CFSAN062364_2016_Peru_C | CFSAN062364 | TBD | PRJNA1062747 | PC | This study | 2016 | Peru | Clinical | 98.4 | 100 | 0 | 99.9857 | 5115284 | 45.27 | 64 | 312423 |
| Vp_Peru_CFSAN062365_2016_Peru_C | CFSAN062365 | TBD | PRJNA1062747 | PC | This study | 2016 | Peru | Clinical | 98.5 | 100 | 0 | 99.9929 | 5117304 | 45.28 | 57 | 347272 |
| Vp_Peru_CFSAN062367_2010_Peru_C | CFSAN062367 | TBD | PRJNA1062747 | PC | This study | 2010 | Peru | Clinical | 98.5 | 100 | 0 | 99.9898 | 5169782 | 45.13 | 57 | 394720 |
| Vp_Peru_CFSAN062368_2010_Peru_C | CFSAN062368 | TBD | PRJNA1062747 | PC | This study | 2010 | Peru | Clinical | 98.3 | 99.68 | 0.05 | 99.9884 | 5173591 | 45.11 | 121 | 340010 |
| Vp_Peru_G7_2015_Lima_C | G7 | TBD | PRJNA1062747 | PC | This study | 2015 | Peru | Clinical | 98.9 | 100 | 0.09 | 99.9851 | 5271206 | 45.29 | 69 | 402146 |
| Vp_Peru_H7_NA_NA_C | H7 | TBD | PRJNA1062747 | PC | This study | NA | Peru | Clinical | 98.8 | 100 | 0.83 | 99.9652 | 5579826 | 45.08 | 488 | 310034 |
| Vp_Peru_P682_2009_Peru_E | P682 | TBD | PRJNA1062747 | PC | This study | 2009 | Peru | Env | 98.3 | 100 | 0 | 99.9929 | 5090692 | 45.29 | 109 | 157162 |
| Vp_Peru_P729_2009_Peru_E | P729 | TBD | PRJNA1062747 | PC | This study | 2009 | Peru | Env | 98.4 | 100 | 0 | 99.9823 | 5094250 | 45.29 | 94 | 157889 |
| Vp_Peru_P860_2009_Peru_E | P860 | TBD | PRJNA1062747 | PC | This study | 2009 | Peru | Env | 97.8 | 100 | 0.27 | 99.9914 | 5084286 | 45.29 | 147 | 95820 |
| Vp_Peru_Peru-288_2001_Peru_C | Peru-288 | TBD | PRJNA1062747 | PC | This study | 2001 | Peru | Clinical | 98.2 | 100 | 0.01 | 99.9875 | 5173942 | 45.3 | 79 | 198215 |
| Vp_Peru_Peru-466_Peru_1996_NA | Peru-466-2 | TBD | PRJNA1062747 | PC | This study | 1996 | Peru | NA | 97.2 | 95.47 | 0.61 | 99.9886 | 5037488 | 45.36 | 149 | 81497 |
| Vp_Peru_vp1027-00_2000_NA_C | vp1027-00 | TBD | PRJNA1062747 | PC | This study | 2000 | Peru | Clinical | 98.6 | 100 | 0 | 99.9894 | 5195821 | 45.17 | 123 | 124878 |
| Vp_Peru_vp1202-11_2011_Peru_Lima_C | vp1202-11 | TBD | PRJNA1062747 | PC | This study | 2011 | Peru | Clinical | 98.7 | 100 | 0 | 99.9816 | 5192518 | 45.14 | 73 | 318936 |
| Vp_Peru_vp1218-11_2011_Peru_Lima_C | vp1218-11 | TBD | PRJNA1062747 | PC | This study | 2011 | Peru | Clinical | 98.7 | 100 | 0 | 99.9822 | 5211436 | 45.13 | 92 | 221913 |
| Vp_Peru_vp1247-09_2009_Peru_Lima_C | vp1247-09 | TBD | PRJNA1062747 | PC | This study | 2009 | Peru | Clinical | 98.8 | 100 | 0 | 99.9834 | 5201547 | 45.15 | 71 | 428501 |
| Vp_Peru_vp1254-07_2007_NA_C | vp1254-07 | TBD | PRJNA1062747 | PC | This study | 2007 | Peru | Clinical | 98.5 | 100 | 0 | 99.9794 | 5110095 | 45.32 | 70 | 187927 |
| Vp_Peru_vp1437-07_2007_Peru_Lima_C | vp1437-07 | TBD | PRJNA1062747 | PC | This study | 2007 | Peru | Clinical | 98.7 | 100 | 0 | 99.9812 | 5196023 | 45.16 | 108 | 126802 |
| Vp_Peru_vp202-11_2010_Peru_Lima_C | vp202-11 | TBD | PRJNA1062747 | PC | This study | 2010 | Peru | Clinical | 98.7 | 100 | 0 | 99.9849 | 5195673 | 45.14 | 84 | 340300 |
| Vp_Peru_vp21254-07_2007_NA_C | vp21254-07 | TBD | PRJNA1062747 | PC | This study | 2007 | Peru | Clinical | 98.4 | 100 | 0 | 99.9884 | 5110604 | 45.32 | 78 | 185492 |
| Vp_Peru_vp21260-07_2007_Peru_Lambayeque_C | vp21260-07 | TBD | PRJNA1062747 | PC | This study | 2007 | Peru | Clinical | 98.5 | 100 | 0 | 99.9816 | 5193602 | 45.15 | 74 | 269984 |
| Vp_Peru_vp2434-00_2000_NA_C | vp2434-00 | TBD | PRJNA1062747 | PC | This study | 2000 | Peru | Clinical | 98.8 | 100 | 0.14 | 99.9733 | 5244838 | 45.1 | 80 | 428063 |
| Vp_Peru_vp2568-00_2000_Peru_Loreto_C | vp2568-00 | TBD | PRJNA1062747 | PC | This study | 2000 | Peru | Clinical | 98.8 | 100 | 0 | 99.9849 | 5127271 | 45.31 | 74 | 347141 |
| Vp_Peru_vp276-15_2015_Peru_Lima_C | vp276-15 | TBD | PRJNA1062747 | PC | This study | 2015 | Peru | Clinical | 98.8 | 100 | 0.09 | 99.9792 | 5266797 | 45.29 | 76 | 216845 |
| Vp_Peru_vp437-07_2007_Peru_Lima_C | vp437-07 | TBD | PRJNA1062747 | PC | This study | 2007 | Peru | Clinical | 98.8 | 100 | 0 | 99.9855 | 5203012 | 45.14 | 79 | 310942 |
| VP08414 | VP08414 | SAMN41562421 | JBDZBP000000000 | PrePC | This study | 2008 | China | NA | 94.6 | 100 | 0.14 | 99.259 | 5140545 | 45.4 | 2 | 3337472 |
| VP16 | VP16 | SAMN41562387 | JBDZCX000000000 | PC | This study | NA | China | Env | 98.5 | 100 | 0 | 99.9785 | 5093390 | 45.29 | 57 | 401463 |
| VP19002 | VP19002 | SAMN41562422 | JBDZBO000000000 | PC | This study | 2019 | China | Clinical | 98.1 | 100 | 0 | 99.9927 | 5083123 | 45.28 | 53 | 439152 |
| VP19003 | VP19003 | SAMN41562423 | JBDZBN000000000 | PC | This study | 2019 | China | Clinical | 98.5 | 100 | 0 | 99.9795 | 5091711 | 45.29 | 53 | 461881 |
| VP19004 | VP19004 | SAMN41562424 | JBDZBM000000000 | PC | This study | 2019 | China | Clinical | 98.4 | 100 | 0 | 99.9816 | 5110070 | 45.28 | 51 | 505531 |
| VP19005 | VP19005 | SAMN41562425 | JBDZBL000000000 | PC | This study | 2019 | China | Clinical | 98.2 | 100 | 0 | 99.9877 | 5116798 | 45.32 | 44 | 505531 |
| VP19007 | VP19007 | SAMN41562426 | JBDZBK000000000 | PC | This study | 2019 | China | Clinical | 98.1 | 100 | 0 | 99.9923 | 5081716 | 45.28 | 53 | 334356 |
| VP19008 | VP19008 | SAMN41562427 | JBDZBJ000000000 | PC | This study | 2019 | China | Clinical | 98.5 | 100 | 0 | 99.9888 | 5139396 | 45.3 | 44 | 542844 |
| VP19009 | VP19009 | SAMN41562428 | JBDZBI000000000 | PC | This study | 2019 | China | Clinical | 98.4 | 100 | 0 | 99.9847 | 5110135 | 45.28 | 49 | 542850 |
| VP19010 | VP19010 | SAMN41562429 | JBDZBH000000000 | PC | This study | 2019 | China | Clinical | 98.5 | 100 | 0 | 99.991 | 5094946 | 45.29 | 51 | 461882 |
| VP19019 | VP19019 | SAMN41562430 | JBDZBG000000000 | PC | This study | 2019 | China | Clinical | 97.8 | 100 | 0 | 99.9278 | 5157754 | 45.15 | 41 | 461882 |
| VP19020 | VP19020 | SAMN41562431 | JBDZBF000000000 | PC | This study | 2019 | China | Clinical | 96.5 | 100 | 0 | 99.965 | 5161471 | 45.28 | 44 | 432948 |
| VP19021 | VP19021 | SAMN41562432 | JBDZBE000000000 | PC | This study | 2019 | China | Clinical | 97.8 | 100 | 0 | 99.9464 | 5155729 | 45.14 | 42 | 461882 |

| | | | | | | | | | | | | | | | |
|---|---|---|---|---|---|---|---|---|---|---|---|---|---|---|---|
| VP19024 | VP19024 | SAMN41562433 | JBDZBD000000000 | PC | This study | 2019 | China | Clinical | 98.5 | 100 | 0 | 99.9811 | 5113731 | 45.26 | 45 | 505531 |
| VP19025 | VP19025 | SAMN41562434 | JBDZBC000000000 | PC | This study | 2019 | China | Clinical | 98.5 | 100 | 0.27 | 99.9892 | 5120625 | 45.26 | 46 | 505531 |
| VP19026 | VP19026 | SAMN41562435 | JBDZBB000000000 | PC | This study | 2019 | China | Clinical | 98.4 | 100 | 0 | 99.9841 | 5110026 | 45.29 | 51 | 461882 |
| VP19027 | VP19027 | SAMN41562436 | JBDZBA000000000 | PC | This study | 2019 | China | Clinical | 98.4 | 100 | 0 | 99.9848 | 5110586 | 45.28 | 53 | 374185 |
| VP19029 | VP19029 | SAMN41562437 | JBDZAZ000000000 | PC | This study | 2019 | China | Clinical | 98.5 | 100 | 0 | 99.9822 | 5220820 | 45.08 | 52 | 542845 |
| VP19030 | VP19030 | SAMN41562438 | JBDZAY000000000 | PC | This study | 2019 | China | Clinical | 98.5 | 100 | 0 | 99.9858 | 5115359 | 45.26 | 47 | 473251 |
| VP19031 | VP19031 | SAMN41562439 | JBDZAX000000000 | PC | This study | 2019 | China | Clinical | 98.5 | 100 | 0 | 99.9845 | 5113403 | 45.26 | 49 | 473282 |
| VP19032 | VP19032 | SAMN41562440 | JBDZAW000000000 | PC | This study | 2019 | China | Clinical | 98.5 | 100 | 0 | 99.9821 | 5092977 | 45.29 | 52 | 461882 |
| VP19034 | VP19034 | SAMN41562441 | JBDZAV000000000 | PC | This study | 2019 | China | Clinical | 98.4 | 100 | 0 | 99.9864 | 5110367 | 45.28 | 50 | 461882 |
| VP19037 | VP19037 | SAMN41562442 | JBDZAU000000000 | PC | This study | 2019 | China | Clinical | 97.8 | 100 | 0 | 99.9278 | 5156901 | 45.15 | 39 | 473292 |
| VP19038 | VP19038 | SAMN41562443 | JBDZAT000000000 | PC | This study | 2019 | China | Clinical | 96.6 | 100 | 0 | 99.9856 | 5111373 | 45.28 | 52 | 505531 |
| VP19039 | VP19039 | SAMN41562444 | JBDZAS000000000 | PC | This study | 2019 | China | Clinical | 98.1 | 100 | 0 | 99.9926 | 5082785 | 45.28 | 54 | 432937 |
| VP19040 | VP19040 | SAMN41562445 | JBDZAR000000000 | PC | This study | 2019 | China | Clinical | 97.8 | 100 | 0 | 99.9365 | 5157428 | 45.15 | 44 | 362044 |
| VP19041 | VP19041 | SAMN41562446 | JBDZAQ000000000 | PC | This study | 2019 | China | Clinical | 97.8 | 100 | 0 | 99.9241 | 5156523 | 45.15 | 43 | 505521 |
| VP19042 | VP19042 | SAMN41562447 | JBDZAP000000000 | PC | This study | 2019 | China | Clinical | 97.8 | 100 | 0 | 99.9466 | 5157085 | 45.14 | 41 | 461882 |
| VP19043 | VP19043 | SAMN41562448 | JBDZAO000000000 | PC | This study | 2019 | China | Clinical | 98.5 | 100 | 0 | 99.9831 | 5092194 | 45.29 | 50 | 473044 |
| VP19044 | VP19044 | SAMN41562449 | JBDZAN000000000 | PC | This study | 2019 | China | Clinical | 98.4 | 100 | 0 | 99.9843 | 5110444 | 45.29 | 51 | 374185 |
| VP19045 | VP19045 | SAMN41562450 | JBDZAM000000000 | PC | This study | 2019 | China | Clinical | 98.5 | 100 | 0 | 99.989 | 5095514 | 45.29 | 56 | 505531 |
| VP19048 | VP19048 | SAMN41562451 | JBDZAL000000000 | PC | This study | 2019 | China | Clinical | 98.5 | 100 | 0.27 | 99.9889 | 5120961 | 45.27 | 58 | 432923 |
| VP19049 | VP19049 | SAMN41562452 | JBDZAK000000000 | PC | This study | 2019 | China | Clinical | 98.5 | 100 | 0 | 99.9645 | 5161632 | 45.22 | 52 | 505531 |
| VP19050 | VP19050 | SAMN41562453 | JBDZAJ000000000 | PC | This study | 2019 | China | Clinical | 98.5 | 100 | 0 | 99.9807 | 5114635 | 45.26 | 50 | 505531 |
| VP19071 | VP19071 | SAMN41562454 | JBDZAI000000000 | PC | This study | 2019 | China | Clinical | 97.8 | 100 | 0 | 99.9242 | 5157751 | 45.15 | 37 | 505521 |
| VP19074 | VP19074 | SAMN41562455 | JBDZAH000000000 | PC | This study | 2019 | China | Clinical | 98 | 100 | 0 | 99.9549 | 5145869 | 45.27 | 37 | 542846 |
| VP19075 | VP19075 | SAMN41562456 | JBDZAG000000000 | PC | This study | 2019 | China | Clinical | 98.4 | 100 | 0 | 99.9831 | 5110784 | 45.29 | 54 | 461882 |
| VP19076 | VP19076 | SAMN41562457 | JBDZAF000000000 | PC | This study | 2019 | China | Clinical | 98.5 | 100 | 0 | 99.9798 | 5196418 | 45.1 | 52 | 472957 |
| VP19077 | VP19077 | SAMN41562458 | JBDZAE000000000 | PC | This study | 2019 | China | Clinical | 98.5 | 100 | 0.2 | 99.9778 | 5131800 | 45.21 | 55 | 455248 |
| VP19079 | VP19079 | SAMN41562459 | JBDZAD000000000 | PC | This study | 2019 | China | Clinical | 98.5 | 100 | 0 | 99.9887 | 5113536 | 45.26 | 46 | 542845 |
| VP19090 | VP19090 | SAMN41562460 | JBDZAC000000000 | PC | This study | 2019 | China | Clinical | 98.5 | 100 | 0 | 99.9882 | 5139951 | 45.23 | 52 | 330588 |
| VP19091 | VP19091 | SAMN41562461 | JBDZAB000000000 | PC | This study | 2019 | China | Clinical | 98.1 | 100 | 0 | 99.9861 | 5083467 | 45.28 | 50 | 506568 |
| VP19092 | VP19092 | SAMN41562462 | JBDZAA000000000 | PC | This study | 2019 | China | Clinical | 98.5 | 100 | 0 | 99.986 | 5097733 | 45.29 | 50 | 334348 |
| VP19093 | VP19093 | SAMN41562463 | JBDYZZ000000000 | PC | This study | 2019 | China | Clinical | 98.5 | 100 | 0.27 | 99.9829 | 5158535 | 45.22 | 47 | 505531 |
| VP19095 | VP19095 | SAMN41562464 | JBDYZY000000000 | PC | This study | 2019 | China | Clinical | 98.4 | 100 | 0 | 99.985 | 5110301 | 45.28 | 51 | 505531 |
| VP19097 | VP19097 | SAMN41562465 | JBDYZX000000000 | PC | This study | 2019 | China | Clinical | 98.5 | 100 | 0 | 99.9883 | 5112099 | 45.28 | 49 | 542851 |
| VP19110 | VP19110 | SAMN41562466 | JBDYZW000000000 | PC | This study | 2019 | China | Clinical | 98.1 | 100 | 0 | 99.9863 | 5087943 | 45.28 | 58 | 374300 |
| VP19111 | VP19111 | SAMN41562467 | JBDYZV000000000 | PC | This study | 2019 | China | Clinical | 98.5 | 100 | 0 | 99.9818 | 5097099 | 45.29 | 49 | 461882 |
| VP19137 | VP19137 | SAMN41562468 | JBDYZU000000000 | PC | This study | 2019 | China | Clinical | 98.4 | 100 | 0 | 99.9805 | 5110262 | 45.29 | 48 | 505531 |
| VP19139 | VP19139 | SAMN41562469 | JBDYZT000000000 | PC | This study | 2019 | China | Clinical | 98.5 | 100 | 0.09 | 99.9809 | 5178462 | 45.21 | 52 | 374095 |
| VP19142 | VP19142 | SAMN41562470 | JBDYZS000000000 | PC | This study | 2019 | China | Clinical | 98.4 | 100 | 0 | 99.9849 | 5085152 | 45.29 | 46 | 506556 |
| VP19143 | VP19143 | SAMN41562471 | JBDYZR000000000 | PC | This study | 2019 | China | Clinical | 98.4 | 100 | 0 | 99.9785 | 5110451 | 45.28 | 50 | 542849 |
| VP19146 | VP19146 | SAMN41562472 | JBDYZQ000000000 | PC | This study | 2019 | China | Clinical | 98.5 | 100 | 0.07 | 99.97 | 5131323 | 45.21 | 50 | 401590 |
| VP19164 | VP19164 | SAMN41562473 | JBDYZP000000000 | PC | This study | 2019 | China | Clinical | 98.3 | 100 | 0 | 99.9802 | 5147202 | 45.28 | 50 | 505531 |
| VP19166 | VP19166 | SAMN41562474 | JBDYZO000000000 | PC | This study | 2019 | China | Clinical | 98.4 | 100 | 0 | 99.9765 | 5110428 | 45.29 | 54 | 461882 |
| VP19167 | VP19167 | SAMN41562475 | JBDYZN000000000 | PC | This study | 2019 | China | Clinical | 97.8 | 100 | 0 | 99.8978 | 5136257 | 45.27 | 56 | 461935 |
| VP19169 | VP19169 | SAMN41562476 | JBDYZM000000000 | PC | This study | 2019 | China | Clinical | 96.3 | 100 | 0 | 99.9734 | 5155789 | 45.27 | 41 | 432974 |
| VP19170 | VP19170 | SAMN41562477 | JBDYZL000000000 | PC | This study | 2019 | China | Clinical | 98 | 100 | 0 | 99.976 | 5158447 | 45.28 | 42 | 580628 |
| VP19171 | VP19171 | SAMN41562478 | JBDYZK000000000 | PC | This study | 2019 | China | Clinical | 98.5 | 100 | 0 | 99.9815 | 5091381 | 45.29 | 48 | 461882 |
| VP19205 | VP19205 | SAMN41562479 | JBDYZJ000000000 | PC | This study | 2019 | China | Clinical | 98.1 | 100 | 0 | 99.9928 | 5086572 | 45.28 | 52 | 439140 |
| VP19206 | VP19206 | SAMN41562480 | JBDYZI000000000 | PC | This study | 2019 | China | Clinical | 98.4 | 100 | 0 | 99.977 | 5110749 | 45.28 | 49 | 505531 |

| | | | | | | | | | | | | | | | | |
|---|---|---|---|---|---|---|---|---|---|---|---|---|---|---|---|---|
| VP19211 | VP19211 | SAMN41562481 | JBDYZH000000000 | PC | This study | 2019 | China | Clinical | 98.1 | 100 | 0.01 | 99.9805 | 5080470 | 45.29 | 42 | 505531 |
| VP19235 | VP19235 | SAMN41562482 | JBDYZG000000000 | PC | This study | 2019 | China | Clinical | 98.1 | 100 | 0.14 | 99.9743 | 5101149 | 45.31 | 57 | 444620 |
| VP19237 | VP19237 | SAMN41562483 | JBDYZF000000000 | PC | This study | 2019 | China | Clinical | 98.5 | 100 | 0 | 99.9875 | 5093858 | 45.29 | 55 | 385753 |
| VP19238 | VP19238 | SAMN41562484 | JBDYZE000000000 | PC | This study | 2019 | China | Clinical | 98.5 | 100 | 0 | 99.9869 | 5111898 | 45.26 | 55 | 504026 |
| VP19241 | VP19241 | SAMN41562485 | JBDYZD000000000 | PC | This study | 2019 | China | Clinical | 98.2 | 100 | 0.27 | 99.9783 | 5110793 | 45.32 | 44 | 399282 |
| VP19244 | VP19244 | SAMN41562486 | JBDYZC000000000 | PC | This study | 2019 | China | Clinical | 97.8 | 100 | 0 | 99.9292 | 5156910 | 45.15 | 45 | 461882 |
| VP19247 | VP19247 | SAMN41562487 | JBDYZB000000000 | PC | This study | 2019 | China | Clinical | 98.1 | 100 | 0 | 99.9799 | 5081552 | 45.3 | 49 | 334347 |
| VP19248 | VP19248 | SAMN41562488 | JBDYZA000000000 | PC | This study | 2019 | China | Clinical | 98.1 | 100 | 0 | 99.9841 | 5087842 | 45.28 | 57 | 312691 |
| VP19249 | VP19249 | SAMN41562489 | JBDYYZ000000000 | PC | This study | 2019 | China | Clinical | 98.4 | 100 | 0 | 99.9846 | 5112415 | 45.29 | 53 | 461882 |
| VP19250 | VP19250 | SAMN41562490 | JBDYYY000000000 | PC | This study | 2019 | China | Clinical | 98.4 | 100 | 0 | 99.9664 | 5145134 | 45.29 | 53 | 374262 |
| VP19251 | VP19251 | SAMN41562491 | JBDYYX000000000 | PC | This study | 2019 | China | Clinical | 98.1 | 100 | 0 | 99.986 | 5082971 | 45.28 | 52 | 505543 |
| VP19252 | VP19252 | SAMN41562492 | JBDYYW000000000 | PC | This study | 2019 | China | Clinical | 97.8 | 100 | 0.01 | 99.9429 | 5247370 | 45.17 | 46 | 360905 |
| VP19253 | VP19253 | SAMN41562493 | JBDYYV000000000 | PC | This study | 2019 | China | Clinical | 98.5 | 100 | 0 | 99.9858 | 5112578 | 45.29 | 58 | 374351 |
| VP19254 | VP19254 | SAMN41562494 | JBDYYU000000000 | PC | This study | 2019 | China | Clinical | 98.5 | 100 | 0 | 99.9852 | 5176426 | 45.11 | 52 | 374096 |
| VP19255 | VP19255 | SAMN41562495 | JBDYYT000000000 | PC | This study | 2019 | China | Clinical | 98.5 | 100 | 0 | 99.9824 | 5172703 | 45.11 | 47 | 506556 |
| VP19256 | VP19256 | SAMN41562496 | JBDYYS000000000 | PC | This study | 2019 | China | Clinical | 98.5 | 100 | 0 | 99.9793 | 5173318 | 45.1 | 44 | 505531 |
| VP19257 | VP19257 | SAMN41562497 | JBDYYR000000000 | PC | This study | 2019 | China | Clinical | 98 | 100 | 0 | 99.9673 | 5153190 | 45.27 | 40 | 580652 |
| VP19258 | VP19258 | SAMN41562498 | JBDYYQ000000000 | PC | This study | 2019 | China | Clinical | 98.2 | 100 | 0 | 99.9871 | 5087201 | 45.28 | 52 | 432977 |
| VP19260 | VP19260 | SAMN41562499 | JBDYYP000000000 | PC | This study | 2019 | China | Clinical | 98.1 | 100 | 0 | 99.9785 | 5078436 | 45.29 | 35 | 505531 |
| VP19263 | VP19263 | SAMN41562500 | JBDYYO000000000 | PC | This study | 2019 | China | Clinical | 98.4 | 100 | 0 | 99.9855 | 5212169 | 45.13 | 44 | 542678 |
| VP19267 | VP19267 | SAMN41562501 | JBDYYN000000000 | PC | This study | 2019 | China | Clinical | 98.4 | 100 | 0 | 99.9818 | 5088995 | 45.29 | 48 | 374281 |
| VP19273 | VP19273 | SAMN41562502 | JBDYYM000000000 | PC | This study | 2019 | China | Clinical | 98.4 | 100 | 0 | 99.9943 | 5084027 | 45.28 | 50 | 461842 |
| VP19275 | VP19275 | SAMN41562503 | JBDYYL000000000 | PC | This study | 2019 | China | Clinical | 98.2 | 100 | 0 | 99.9847 | 5136627 | 45.31 | 46 | 461842 |
| VP19276 | VP19276 | SAMN41562504 | JBDYYK000000000 | PC | This study | 2019 | China | Clinical | 98.3 | 100 | 0 | 99.9837 | 5105358 | 45.28 | 53 | 347594 |
| VP20006 | VP20006 | SAMN41562505 | JBDYYJ000000000 | PC | This study | 2020 | China | Clinical | 98.4 | 100 | 0 | 99.9774 | 5111636 | 45.28 | 49 | 506556 |
| VP20016 | VP20016 | SAMN41562506 | JBDYYI000000000 | PC | This study | 2020 | China | Clinical | 98.3 | 100 | 0 | 99.9826 | 5091305 | 45.29 | 47 | 542843 |
| VP20021 | VP20021 | SAMN41562507 | JBDYYH000000000 | PC | This study | 2020 | China | Clinical | 98.5 | 100 | 0 | 99.9854 | 5138137 | 45.24 | 58 | 461882 |
| VP20022 | VP20022 | SAMN41562508 | JBDYYG000000000 | PC | This study | 2020 | China | Clinical | 98.5 | 100 | 0 | 99.9889 | 5111198 | 45.28 | 48 | 542851 |
| VP20023 | VP20023 | SAMN41562509 | JBDYYF000000000 | PC | This study | 2020 | China | Clinical | 98.3 | 100 | 0 | 99.9841 | 5107437 | 45.28 | 54 | 432902 |
| VP20024 | VP20024 | SAMN41562510 | JBDYYE000000000 | PC | This study | 2020 | China | Clinical | 98.1 | 100 | 0 | 99.9851 | 5085025 | 45.28 | 50 | 505650 |
| VP20025 | VP20025 | SAMN41562511 | JBDYYD000000000 | PC | This study | 2020 | China | Clinical | 97.9 | 100 | 0 | 99.9767 | 5150635 | 45.27 | 42 | 334356 |
| VP20026 | VP20026 | SAMN41562512 | JBDYYC000000000 | PC | This study | 2020 | China | Clinical | 98.1 | 100 | 0 | 99.9921 | 5087712 | 45.28 | 55 | 334356 |
| VP20029 | VP20029 | SAMN41562513 | JBDYYB000000000 | PC | This study | 2020 | China | Clinical | 96.9 | 100 | 0 | 99.9832 | 5110501 | 45.28 | 55 | 374351 |
| VP20030 | VP20030 | SAMN41562514 | JBDYYA000000000 | PC | This study | 2020 | China | Clinical | 98.4 | 100 | 0 | 99.9748 | 5109846 | 45.29 | 54 | 505531 |
| VP20031 | VP20031 | SAMN41562515 | JBDYXZ000000000 | PC | This study | 2020 | China | Clinical | 98 | 100 | 0 | 99.989 | 5080924 | 45.28 | 59 | 300493 |
| VP20032 | VP20032 | SAMN41562516 | JBDYXY000000000 | PC | This study | 2020 | China | Clinical | 98 | 100 | 0 | 99.9926 | 5081151 | 45.28 | 56 | 334356 |
| VP20033 | VP20033 | SAMN41562517 | JBDYXX000000000 | PC | This study | 2020 | China | Clinical | 98 | 100 | 0 | 99.9707 | 5141283 | 45.28 | 41 | 456913 |
| VP20035 | VP20035 | SAMN41562518 | JBDYXW000000000 | PC | This study | 2020 | China | Clinical | 96.7 | 100 | 0 | 99.9852 | 5110426 | 45.28 | 58 | 374185 |
| VP20036 | VP20036 | SAMN41562519 | JBDYXV000000000 | PC | This study | 2020 | China | Clinical | 98.4 | 100 | 0 | 99.9801 | 5111688 | 45.28 | 57 | 374351 |
| VP20037 | VP20037 | SAMN41562520 | JBDYXU000000000 | PC | This study | 2020 | China | Clinical | 97.8 | 100 | 0.36 | 99.9069 | 5240147 | 45.31 | 53 | 394915 |
| VP20039 | VP20039 | SAMN41562521 | JBDYXT000000000 | PC | This study | 2020 | China | Clinical | 97.8 | 100 | 0.36 | 99.8984 | 5242458 | 45.31 | 57 | 312666 |
| VP20077 | VP20077 | SAMN41562522 | JBDYXS000000000 | PC | This study | 2020 | China | Clinical | 96.3 | 100 | 0 | 99.9145 | 5115802 | 45.28 | 50 | 374871 |
| VP20083 | VP20083 | SAMN41562523 | JBDYXR000000000 | PC | This study | 2020 | China | Clinical | 97.9 | 100 | 0.36 | 99.9113 | 5197045 | 45.34 | 57 | 461882 |
| VP20085 | VP20085 | SAMN41562524 | JBDYXQ000000000 | PC | This study | 2020 | China | Clinical | 97.9 | 100 | 0.36 | 99.9196 | 5196233 | 45.34 | 53 | 461882 |
| VP20090 | VP20090 | SAMN41562525 | JBDYXP000000000 | PC | This study | 2020 | China | Clinical | 98.1 | 100 | 0 | 99.9912 | 5086583 | 45.28 | 53 | 461881 |
| VP20092 | VP20092 | SAMN41562526 | JBDYXO000000000 | PC | This study | 2020 | China | Clinical | 96.7 | 100 | 0 | 99.9754 | 5111070 | 45.28 | 52 | 461881 |
| VP20093 | VP20093 | SAMN41562527 | JBDYXN000000000 | PC | This study | 2020 | China | Clinical | 96.9 | 100 | 0 | 99.9857 | 5110221 | 45.28 | 51 | 505531 |
| VP20094 | VP20094 | SAMN41562528 | JBDYXM000000000 | PC | This study | 2020 | China | Clinical | 98.5 | 100 | 0 | 99.9895 | 5109731 | 45.28 | 52 | 473156 |

| | | | | | | | | | | | | | | | | |
|---|---|---|---|---|---|---|---|---|---|---|---|---|---|---|---|---|
| VP20095 | VP20095 | SAMN41562529 | JBDYXL000000000 | PC | This study | 2020 | China | Clinical | 97.9 | 100 | 0 | 99.9112 | 5118266 | 45.28 | 49 | 506556 |
| VP20097 | VP20097 | SAMN41562530 | JBDYXK000000000 | PC | This study | 2020 | China | Clinical | 98.1 | 100 | 0 | 99.9854 | 5093965 | 45.28 | 41 | 542837 |
| VP20098 | VP20098 | SAMN41562531 | JBDYXJ000000000 | PC | This study | 2020 | China | Clinical | 96.8 | 100 | 0 | 99.9748 | 5109926 | 45.28 | 54 | 461881 |
| VP20105 | VP20105 | SAMN41562532 | JBDYXI000000000 | PC | This study | 2020 | China | Clinical | 98.4 | 100 | 0 | 99.9781 | 5153461 | 45.34 | 60 | 374189 |
| VP20106 | VP20106 | SAMN41562533 | JBDYXH000000000 | PC | This study | 2020 | China | Clinical | 98.5 | 100 | 0 | 99.985 | 5111892 | 45.28 | 46 | 461882 |
| VP20107 | VP20107 | SAMN41562534 | JBDYXG000000000 | PC | This study | 2020 | China | Clinical | 97.9 | 100 | 0 | 99.9129 | 5117738 | 45.28 | 44 | 505531 |
| VP20108 | VP20108 | SAMN41562535 | JBDYXF000000000 | PC | This study | 2020 | China | Clinical | 97.9 | 100 | 0 | 99.9215 | 5117910 | 45.28 | 50 | 473308 |
| VP20109 | VP20109 | SAMN41562536 | JBDYXE000000000 | PC | This study | 2020 | China | Clinical | 98.5 | 100 | 0 | 99.9717 | 5224543 | 45.08 | 53 | 542845 |
| VP20110 | VP20110 | SAMN41562537 | JBDYXD000000000 | PC | This study | 2020 | China | Clinical | 96.3 | 100 | 0 | 99.9203 | 5116645 | 45.28 | 53 | 461882 |
| VP20158 | VP20158 | SAMN41562538 | JBDYXC000000000 | PC | This study | 2020 | China | Clinical | 98.5 | 100 | 0 | 99.9892 | 5173320 | 45.11 | 55 | 334348 |
| VP20159 | VP20159 | SAMN41562539 | JBDYXB000000000 | PC | This study | 2020 | China | Clinical | 98.4 | 100 | 0 | 99.9855 | 5109392 | 45.28 | 51 | 505540 |
| VP20160 | VP20160 | SAMN41562540 | JBDYXA000000000 | PC | This study | 2020 | China | Clinical | 97.9 | 100 | 0 | 99.9194 | 5118090 | 45.28 | 46 | 505531 |
| VP20161 | VP20161 | SAMN41562541 | JBDYWZ000000000 | PC | This study | 2020 | China | Clinical | 98.4 | 100 | 0 | 99.982 | 5110932 | 45.28 | 55 | 461882 |
| VP20162 | VP20162 | SAMN41562542 | JBDYWY000000000 | PC | This study | 2020 | China | Clinical | 97.9 | 100 | 0 | 99.912 | 5117078 | 45.28 | 50 | 401605 |
| VP20164 | VP20164 | SAMN41562543 | JBDYWX000000000 | PC | This study | 2020 | China | Clinical | 98.2 | 100 | 0 | 99.9822 | 5074646 | 45.31 | 46 | 506542 |
| VP20165 | VP20165 | SAMN41562544 | JBDYWW000000000 | PC | This study | 2020 | China | Clinical | 98 | 100 | 0 | 99.9713 | 5154821 | 45.27 | 38 | 580640 |
| VP20171 | VP20171 | SAMN41562545 | JBDYWV000000000 | PC | This study | 2020 | China | Clinical | 97.9 | 100 | 0 | 99.9207 | 5118111 | 45.28 | 49 | 472693 |
| VP21015 | VP21015 | SAMN41562546 | JBDYWU000000000 | PC | This study | 2021 | China | Clinical | 98 | 100 | 0 | 99.9645 | 5160819 | 45.27 | 37 | 580647 |
| VP21019 | VP21019 | SAMN41562547 | JBDYWT000000000 | PC | This study | 2021 | China | Clinical | 97.9 | 100 | 0 | 99.9185 | 5119177 | 45.29 | 49 | 505531 |
| VP21021 | VP21021 | SAMN41562548 | JBDYWS000000000 | PC | This study | 2021 | China | Clinical | 98.6 | 100 | 0 | 99.9774 | 5248727 | 45.05 | 54 | 399329 |
| VP21023 | VP21023 | SAMN41562549 | JBDYWR000000000 | PC | This study | 2021 | China | Clinical | 97.9 | 100 | 0 | 99.9161 | 5121166 | 45.29 | 52 | 401502 |
| VP21024 | VP21024 | SAMN41562550 | JBDYWQ000000000 | PC | This study | 2021 | China | Clinical | 97.9 | 100 | 0 | 99.9127 | 5120124 | 45.28 | 52 | 426055 |
| VP21026 | VP21026 | SAMN41562551 | JBDYWP000000000 | PC | This study | 2021 | China | Clinical | 97.9 | 100 | 0 | 99.9112 | 5119231 | 45.28 | 52 | 426067 |
| VP21032 | VP21032 | SAMN41562552 | JBDYWO000000000 | PC | This study | 2021 | China | Clinical | 98.5 | 100 | 0 | 99.9785 | 5248192 | 45.05 | 51 | 507128 |
| VP21034 | VP21034 | SAMN41562553 | JBDYWN000000000 | PC | This study | 2021 | China | Clinical | 97.9 | 100 | 0 | 99.9215 | 5119985 | 45.28 | 56 | 396124 |
| VP21039 | VP21039 | SAMN41562554 | JBDYWM000000000 | PC | This study | 2021 | China | Clinical | 96.8 | 100 | 1.05 | 99.9928 | 5154658 | 45.28 | 150 | 81613 |
| VP21040 | VP21040 | SAMN41562555 | JBDYWL000000000 | PC | This study | 2021 | China | Clinical | 98.6 | 100 | 0 | 99.9838 | 5113173 | 45.28 | 53 | 506656 |
| VP21041 | VP21041 | SAMN41562556 | JBDYWK000000000 | PC | This study | 2021 | China | Clinical | 98.2 | 100 | 0 | 99.9904 | 5075641 | 45.31 | 58 | 289616 |
| VP21057 | VP21057 | SAMN41562557 | JBDYWJ000000000 | PC | This study | 2021 | China | Clinical | 97.8 | 100 | 0 | 99.9164 | 5231723 | 45.19 | 52 | 340447 |
| VP21059 | VP21059 | SAMN41562558 | JBDYWI000000000 | PC | This study | 2021 | China | Clinical | 97.9 | 100 | 0 | 99.9192 | 5117315 | 45.28 | 48 | 505531 |
| VP21061 | VP21061 | SAMN41562559 | JBDYWH000000000 | PC | This study | 2021 | China | Clinical | 97.9 | 100 | 0 | 99.9164 | 5131607 | 45.27 | 48 | 461882 |
| VP21065 | VP21065 | SAMN41562560 | JBDYWG000000000 | PC | This study | 2021 | China | Clinical | 97.9 | 100 | 0 | 99.9197 | 5119282 | 45.28 | 52 | 473617 |
| VP21067 | VP21067 | SAMN41562561 | JBDYWF000000000 | PC | This study | 2021 | China | Clinical | 97.9 | 100 | 0 | 99.9112 | 5132595 | 45.27 | 44 | 506556 |
| VP21068 | VP21068 | SAMN41562562 | JBDYWE000000000 | PC | This study | 2021 | China | Clinical | 96.2 | 100 | 0 | 99.9661 | 5154605 | 45.28 | 36 | 542852 |
| VP21070 | VP21070 | SAMN41562563 | JBDYWD000000000 | PC | This study | 2021 | China | Clinical | 98.5 | 100 | 0 | 99.9873 | 5091473 | 45.29 | 45 | 461882 |
| VP21071 | VP21071 | SAMN41562564 | JBDYWC000000000 | PC | This study | 2021 | China | Clinical | 97.9 | 100 | 0 | 99.913 | 5174799 | 45.28 | 50 | 506556 |
| VP21072 | VP21072 | SAMN41562565 | JBDYWB000000000 | PC | This study | 2021 | China | Clinical | 97.9 | 100 | 0 | 99.9133 | 5174198 | 45.28 | 50 | 506556 |
| VP21074 | VP21074 | SAMN41562566 | JBDYWA000000000 | PC | This study | 2021 | China | Clinical | 97.8 | 100 | 0 | 99.9212 | 5170550 | 45.28 | 48 | 505491 |
| VP21076 | VP21076 | SAMN41562567 | JBDYVZ000000000 | PC | This study | 2021 | China | Clinical | 97.9 | 100 | 0 | 99.9128 | 5125814 | 45.27 | 49 | 505531 |
| VP21077 | VP21077 | SAMN41562568 | JBDYVY000000000 | PC | This study | 2021 | China | Clinical | 97.9 | 100 | 0 | 99.9188 | 5117881 | 45.28 | 48 | 506556 |
| VP21078 | VP21078 | SAMN41562569 | JBDYVX000000000 | PC | This study | 2021 | China | Clinical | 97.7 | 100 | 0 | 99.9204 | 5127245 | 45.27 | 45 | 461842 |
| VP21079 | VP21079 | SAMN41562570 | JBDYVW000000000 | PC | This study | 2021 | China | Clinical | 97.8 | 100 | 0 | 99.9188 | 5131813 | 45.27 | 44 | 506556 |
| VP21080 | VP21080 | SAMN41562571 | JBDYVV000000000 | PC | This study | 2021 | China | Clinical | 97.9 | 100 | 0 | 99.9196 | 5173083 | 45.28 | 51 | 461882 |
| VP21081 | VP21081 | SAMN41562572 | JBDYVU000000000 | PC | This study | 2021 | China | Clinical | 97.9 | 100 | 0 | 99.9202 | 5131218 | 45.27 | 42 | 827757 |
| VP21082 | VP21082 | SAMN41562573 | JBDZCY000000000 | PC | This study | 2021 | China | Clinical | 97.9 | 100 | 0 | 99.9138 | 5132507 | 45.27 | 46 | 506556 |
| VP21083 | VP21083 | SAMN41562574 | JBDYVT000000000 | PC | This study | 2021 | China | Clinical | 97.9 | 100 | 0 | 99.9123 | 5131507 | 45.27 | 41 | 827643 |
| VP21084 | VP21084 | SAMN41562575 | JBDYVS000000000 | PC | This study | 2021 | China | Clinical | 97.8 | 100 | 0 | 99.9118 | 5130604 | 45.27 | 44 | 549545 |
| VP21086 | VP21086 | SAMN41562576 | JBDYVR000000000 | PC | This study | 2021 | China | Clinical | 97.8 | 100 | 0 | 99.9202 | 5129220 | 45.27 | 45 | 505491 |

| | | | | | | | | | | | | | | | | |
|---|---|---|---|---|---|---|---|---|---|---|---|---|---|---|---|---|
| VP21087 | VP21087 | SAMN41562577 | JBDYVQ000000000 | PC | This study | 2021 | China | Clinical | 97.8 | 100 | 0 | 99.921 | 5128646 | 45.27 | 45 | 472971 |
| VP21088 | VP21088 | SAMN41562578 | JBDYVP000000000 | PC | This study | 2021 | China | Clinical | 97.8 | 100 | 0 | 99.9114 | 5130602 | 45.27 | 43 | 506556 |
| VP21089 | VP21089 | SAMN41562579 | JBDYVO000000000 | PC | This study | 2021 | China | Clinical | 97.9 | 100 | 0 | 99.911 | 5131538 | 45.27 | 44 | 827643 |
| VP21090 | VP21090 | SAMN41562580 | JBDYVN000000000 | PC | This study | 2021 | China | Clinical | 97.8 | 100 | 0 | 99.9104 | 5130223 | 45.27 | 44 | 827644 |
| VP21091 | VP21091 | SAMN41562581 | JBDYVM000000000 | PC | This study | 2021 | China | Clinical | 97.9 | 100 | 0 | 99.9102 | 5131587 | 45.27 | 43 | 549545 |
| VP21092 | VP21092 | SAMN41562582 | JBDYVL000000000 | PC | This study | 2021 | China | Clinical | 97.9 | 100 | 0 | 99.9197 | 5132119 | 45.27 | 46 | 549545 |
| VP21095 | VP21095 | SAMN41562583 | JBDYVK000000000 | PC | This study | 2021 | China | Clinical | 97.9 | 100 | 0 | 99.91 | 5131153 | 45.27 | 41 | 506556 |
| VP21096 | VP21096 | SAMN41562584 | JBDYVJ000000000 | PC | This study | 2021 | China | Clinical | 97.9 | 100 | 0 | 99.9194 | 5132039 | 45.27 | 42 | 828005 |
| VP21097 | VP21097 | SAMN41562585 | JBDYVI000000000 | PC | This study | 2021 | China | Clinical | 97.9 | 100 | 0 | 99.9102 | 5132922 | 45.27 | 42 | 549545 |
| VP21098 | VP21098 | SAMN41562586 | JBDYVH000000000 | PC | This study | 2021 | China | Clinical | 97.8 | 100 | 0 | 99.912 | 5131112 | 45.27 | 45 | 549545 |
| VP21099 | VP21099 | SAMN41562587 | JBDYVG000000000 | PC | This study | 2021 | China | Clinical | 97.8 | 100 | 0 | 99.9043 | 5132076 | 45.27 | 49 | 506724 |
| VP21100 | VP21100 | SAMN41562588 | JBDYVF000000000 | PC | This study | 2021 | China | Clinical | 97.8 | 100 | 0 | 99.9206 | 5129965 | 45.27 | 47 | 461842 |
| VP21101 | VP21101 | SAMN41562589 | JBDYVE000000000 | PC | This study | 2021 | China | Clinical | 97.9 | 100 | 0 | 99.9193 | 5130623 | 45.27 | 45 | 401470 |
| VP21102 | VP21102 | SAMN41562590 | JBDYVD000000000 | PC | This study | 2021 | China | Clinical | 97.8 | 100 | 0 | 99.9222 | 5122176 | 45.28 | 51 | 399142 |
| VP21103 | VP21103 | SAMN41562591 | JBDYVC000000000 | PC | This study | 2021 | China | Clinical | 97.9 | 100 | 0 | 99.9103 | 5131271 | 45.27 | 41 | 827643 |
| VP22_4B-8 | VP22_4B-8 | SAMN41562592 | JBDYVB000000000 | PrePC | This study | 2022 | China | Env | 94.4 | 100 | 0 | 99.4893 | 5221526 | 45.28 | 54 | 826891 |
| VP4 | VP4 | SAMN41562388 | JBDZCW000000000 | PC | This study | NA | China | NA | 98.5 | 100 | 0 | 99.9856 | 5092040 | 45.29 | 56 | 401480 |
| VP5 | VP5 | SAMN41562389 | JBDZCV000000000 | PC | This study | NA | China | NA | 98.5 | 100 | 0 | 99.9866 | 5092693 | 45.29 | 56 | 425673 |
| VPCZ1 | VPCZ1 | SAMN41562390 | JBDZCU000000000 | PC | This study | 2013 | China | Clinical | 98 | 100 | 0 | 99.9914 | 5139546 | 45.25 | 58 | 401604 |
| VPCZ10 | VPCZ10 | SAMN41562391 | JBDZCT000000000 | PC | This study | 2014 | China | Clinical | 98.1 | 100 | 0 | 99.9905 | 5107011 | 45.32 | 51 | 401580 |
| VPCZ106 | VPCZ106 | SAMN41562392 | JBDZCS000000000 | PC | This study | 2014 | China | Clinical | 97.8 | 100 | 0 | 99.9065 | 5171198 | 45.3 | 58 | 426253 |
| VPCZ107 | VPCZ107 | SAMN41562393 | JBDZCR000000000 | PC | This study | 2014 | China | Clinical | 97.8 | 100 | 0 | 99.9152 | 5135001 | 45.27 | 56 | 426253 |
| VPCZ11 | VPCZ11 | SAMN41562394 | JBDZCQ000000000 | PC | This study | 2014 | China | Clinical | 96.5 | 100 | 0 | 99.99 | 5073268 | 45.3 | 45 | 404249 |
| VPCZ13 | VPCZ13 | SAMN41562395 | JBDZCP000000000 | PC | This study | 2014 | China | Clinical | 96.3 | 100 | 0 | 99.9588 | 5139283 | 45.28 | 48 | 401464 |
| VPCZ14 | VPCZ14 | SAMN41562396 | JBDZCO000000000 | PC | This study | 2014 | China | Clinical | 96.5 | 100 | 0 | 99.9902 | 5071567 | 45.3 | 45 | 401462 |
| VPCZ17 | VPCZ17 | SAMN41562397 | JBDZCN000000000 | PC | This study | 2014 | China | Clinical | 96.5 | 100 | 0 | 99.9895 | 5072180 | 45.3 | 49 | 401462 |
| VPCZ18 | VPCZ18 | SAMN41562398 | JBDZCM000000000 | PC | This study | 2014 | China | Clinical | 96.5 | 100 | 0 | 99.9907 | 5071847 | 45.3 | 46 | 394965 |
| VPCZ19 | VPCZ19 | SAMN41562399 | JBDZCL000000000 | PC | This study | 2014 | China | Clinical | 98.5 | 100 | 0 | 99.9838 | 5091762 | 45.29 | 56 | 425757 |
| VPCZ2 | VPCZ2 | SAMN41562400 | JBDZCK000000000 | PC | This study | 2013 | China | Clinical | 98 | 100 | 0 | 99.9913 | 5139272 | 45.25 | 55 | 440955 |
| VPCZ21 | VPCZ21 | SAMN41562401 | JBDZCJ000000000 | PC | This study | 2014 | China | Clinical | 96.4 | 100 | 0 | 99.9547 | 5139538 | 45.28 | 50 | 334356 |
| VPCZ23 | VPCZ23 | SAMN41562402 | JBDZCI000000000 | PC | This study | 2014 | China | Clinical | 96.4 | 100 | 0 | 99.9766 | 5138801 | 45.28 | 42 | 401463 |
| VPCZ25 | VPCZ25 | SAMN41562403 | JBDZCH000000000 | PC | This study | 2014 | China | Clinical | 96.4 | 100 | 0 | 99.9775 | 5138737 | 45.28 | 44 | 401464 |
| VPCZ27 | VPCZ27 | SAMN41562404 | JBDZCG000000000 | PC | This study | 2013 | China | Clinical | 98.7 | 100 | 0 | 99.9882 | 5112410 | 45.28 | 57 | 427752 |
| VPCZ28 | VPCZ28 | SAMN41562405 | JBDZCF000000000 | PC | This study | 2014 | China | Clinical | 98.7 | 100 | 0 | 99.9801 | 5103087 | 45.29 | 56 | 401477 |
| VPCZ29 | VPCZ29 | SAMN41562406 | JBDZCE000000000 | PC | This study | 2014 | China | Clinical | 97.9 | 100 | 0 | 99.9546 | 5139369 | 45.28 | 46 | 401463 |
| VPCZ3 | VPCZ3 | SAMN41562407 | JBDZCD000000000 | PC | This study | 2013 | China | Clinical | 98 | 100 | 0 | 99.9915 | 5139519 | 45.25 | 61 | 394978 |
| VPCZ31 | VPCZ31 | SAMN41562408 | JBDZCC000000000 | PC | This study | 2014 | China | Clinical | 98 | 100 | 0 | 99.9765 | 5139354 | 45.28 | 42 | 401463 |
| VPCZ32 | VPCZ32 | SAMN41562409 | JBDZCB000000000 | PC | This study | 2014 | China | Clinical | 97.9 | 100 | 0 | 99.9675 | 5139474 | 45.28 | 44 | 401597 |
| VPCZ33 | VPCZ33 | SAMN41562410 | JBDZCA000000000 | PC | This study | 2014 | China | Clinical | 95.1 | 100 | 0.02 | 99.9375 | 5138996 | 45.28 | 44 | 401463 |
| VPCZ35 | VPCZ35 | SAMN41562411 | JBDZBZ000000000 | PC | This study | 2014 | China | Clinical | 98.1 | 100 | 0 | 99.9904 | 5072223 | 45.3 | 48 | 394963 |
| VPCZ36 | VPCZ36 | SAMN41562412 | JBDZBY000000000 | PC | This study | 2014 | China | Clinical | 96.4 | 100 | 0 | 99.9685 | 5138618 | 45.28 | 44 | 401460 |
| VPCZ38 | VPCZ38 | SAMN41562413 | JBDZBX000000000 | PC | This study | 2013 | China | Clinical | 97.8 | 100 | 0 | 99.9213 | 5164102 | 45.25 | 54 | 476575 |
| VPCZ4 | VPCZ4 | SAMN41562414 | JBDZBW000000000 | PC | This study | 2013 | China | Clinical | 98 | 100 | 0 | 99.9903 | 5150175 | 45.26 | 60 | 401470 |
| VPCZ58 | VPCZ58 | SAMN41562415 | JBDZBV000000000 | PC | This study | 2014 | China | Clinical | 98.5 | 100 | 0 | 99.9856 | 5092142 | 45.29 | 56 | 401470 |
| VPCZ66 | VPCZ66 | SAMN41562416 | JBDZBU000000000 | PC | This study | 2013 | China | Clinical | 98.7 | 100 | 0 | 99.9798 | 5102197 | 45.29 | 58 | 401477 |
| VPCZ82 | VPCZ82 | SAMN41562417 | JBDZBT000000000 | PC | This study | 2014 | China | Clinical | 97.9 | 100 | 0 | 99.9633 | 5139431 | 45.28 | 47 | 395284 |
| VPCZ9 | VPCZ9 | SAMN41562418 | JBDZBS000000000 | PC | This study | 2013 | China | Clinical | 98.7 | 100 | 0 | 99.9916 | 5137353 | 45.28 | 54 | 431654 |
| VPCZ93 | VPCZ93 | SAMN41562419 | JBDZBR000000000 | PC | This study | 2014 | China | Clinical | 97.8 | 100 | 0.01 | 99.9073 | 5193706 | 45.32 | 56 | 401470 |

| | | | | | | | | | | | | | | | | |
|---|---|---|---|---|---|---|---|---|---|---|---|---|---|---|---|---|
| VPCZ97 | VPCZ97 | SAMN41562420 | JBDZBQ000000000 | PC | This study | 2014 | China | Clinical | 97.5 | 100 | 0 | 99.8699 | 5207235 | 45.27 | 50 | 478920 |
| RIMD_2210633 | RIMD_2210633 | SAMD00058707 | GCF_000196095.1 | PC | Public | 1996 | Japan | Clinical | 100 | 100 | 0 | 100 | 5165770 | 45.37 | 2 | 3288558 |
| SAMD00009556-GCF_000591515.1 | DRS013261 | SAMD00009556 | GCF_000591515.1 | NonPC | Public | 2014 | Thailand | NA | 88.9 | 100 | 0.79 | 98.1355 | 5211318 | 45.16 | 64 | 271160 |
| SAMD00009557-GCF_000591555.1 | DRS013263 | SAMD00009557 | GCF_000591555.1 | NonPC | Public | NA | Thailand | NA | 91.8 | 100 | 0.31 | 98.3622 | 5209504 | 45.18 | 50 | 329189 |
| SAMD00009558-GCA_000591475.1 | DRS013260 | SAMD00009558 | GCA_000591475.1 | NonPC | Public | NA | Thailand | Env | 91.6 | 100 | 0.27 | 98.2475 | 5618686 | 45.17 | 82 | 266480 |
| SAMD00009559-GCF_000591455.1 | DRS013259 | SAMD00009559 | GCF_000591455.1 | NonPC | Public | 2014 | Thailand | NA | 91.1 | 100 | 0.32 | 98.1739 | 5272528 | 45.28 | 104 | 314261 |
| SAMD00009560-GCF_000591495.1 | DRS013258 | SAMD00009560 | GCF_000591495.1 | NonPC | Public | 2014 | Thailand | NA | 91.8 | 100 | 0.41 | 98.3669 | 5229840 | 45.33 | 58 | 560774 |
| SAMD00018682-GCF_000813305.1 | NBRC_12711 | SAMD00018682 | GCF_000813305.1 | NonPC | Public | NA | Japan | NA | 90.1 | 100 | 0.05 | 98.3736 | 4991873 | 45.28 | 39 | 381903 |
| SAMEA3443777-GCF_900460415.1 | ERS743103 | SAMEA3443777 | GCF_900460415.1 | NonPC | Public | NA | UK | NA | 94 | 99.73 | 1.09 | 98.3669 | 5375494 | 45.27 | 3 | 2708224 |
| SAMEA3470405-ERR2532053 | NCTC10886 | SAMEA3470405 | ERR2532053 | NonPC | Public | NA | NA | NA | 90 | 96.23 | 1.49 | 98.2106 | 5227093 | 45.18 | 21 | 578458 |
| SAMEA3649022-GCF_900460535.1 | ERS956171 | SAMEA3649022 | GCF_900460535.1 | NonPC | Public | 1973 | NA | NA | 91.6 | 99.86 | 0.05 | 98.3135 | 5168732 | 45.34 | 5 | 3297116 |
| SAMEA5317561-GCF_900682685.1 | ERS3124799 | SAMEA5317561 | GCF_900682685.1 | NonPC | Public | 2009 | Brazil | NA | 91 | 98.62 | 2.74 | 98.3425 | 5028471 | 45.25 | 596 | 18070 |
| SAMEA5540085-ERR3255955 | ERS3342146 | SAMEA5540085 | ERR3255955 | NonPC | Public | NA | UK | NA | 90.5 | 100 | 0.11 | 98.3115 | 5006744 | 45.39 | 41 | 409438 |
| SAMEA5540086-ERR3255956 | ERS3342147 | SAMEA5540086 | ERR3255956 | PC | Public | NA | UK | NA | 98.6 | 100 | 0.14 | 99.9881 | 5116485 | 45.27 | 52 | 290606 |
| SAMEA5540087-ERR3255957 | ERS3342148 | SAMEA5540087 | ERR3255957 | NonPC | Public | NA | UK | NA | 91.8 | 100 | 0.24 | 98.3241 | 5261027 | 45.27 | 44 | 482347 |
| SAMEA5540088-ERR3255958 | ERS3342149 | SAMEA5540088 | ERR3255958 | NonPC | Public | NA | UK | NA | 88.4 | 100 | 0.47 | 98.2404 | 5069980 | 45.2 | 50 | 583955 |
| SAMEA5851835-GCF_902385965.1 | ERS3640361 | SAMEA5851835 | GCF_902385965.1 | NonPC | Public | NA | Bangladesh | Clinical | 90.9 | 100 | 0 | 98.3741 | 5262465 | 45.29 | 4 | 3378377 |
| SAMEA7061695-GCF_903986635.1 | BET6 | SAMEA7061695 | GCF_903986635.1 | NonPC | Public | 2018 | Brazil | NA | 91.1 | 100 | 0 | 98.4017 | 5140846 | 45.2 | 78 | 258264 |
| SAMEA7061697-GCF_903986745.1 | BET14 | SAMEA7061697 | GCF_903986745.1 | NonPC | Public | 2018 | Brazil | NA | 88.1 | 100 | 0.43 | 98.102 | 5110157 | 45.21 | 71 | 178539 |
| SAMEA7061699-GCF_903986865.1 | DRD10 | SAMEA7061699 | GCF_903986865.1 | NonPC | Public | 2018 | Brazil | NA | 87.5 | 100 | 0.4 | 97.7277 | 5154733 | 45.24 | 71 | 229171 |
| SAMEA8102990-GCA_905331545.1 | 20521 | SAMEA8102990 | GCA_905331545.1 | NonPC | Public | 1980 | Japan | Clinical | 89.3 | 100 | 0.37 | 98.4174 | 4914971 | 45.36 | 203 | 82256 |
| SAMEA8102992-GCA_905331555.1 | 20528 | SAMEA8102992 | GCA_905331555.1 | NonPC | Public | 1980 | Japan | Clinical | 89.7 | 100 | 0.12 | 98.4064 | 4931543 | 45.36 | 140 | 157677 |
| SAMEA8102993-GCA_905331575.1 | 20535 | SAMEA8102993 | GCA_905331575.1 | NonPC | Public | 1980 | Japan | Clinical | 89.7 | 100 | 0.12 | 98.3195 | 5125862 | 45.15 | 144 | 117963 |
| SAMEA8102994-GCA_905331585.1 | 20538 | SAMEA8102994 | GCA_905331585.1 | NonPC | Public | 1980 | Japan | Clinical | 92.6 | 99.73 | 0.16 | 98.3115 | 5148019 | 45.21 | 115 | 103602 |
| SAMEA8102995-GCA_905331615.1 | 20542 | SAMEA8102995 | GCA_905331615.1 | NonPC | Public | 1980 | Japan | Clinical | 90.2 | 100 | 0 | 98.3423 | 5017165 | 45.29 | 123 | 135034 |
| SAMEA8102996-GCA_905331595.1 | 20545 | SAMEA8102996 | GCA_905331595.1 | NonPC | Public | 1980 | Japan | Clinical | 90.2 | 100 | 0.05 | 98.3654 | 4962659 | 45.3 | 124 | 140431 |
| SAMEA8102997-GCA_905331605.1 | 20556 | SAMEA8102997 | GCA_905331605.1 | NonPC | Public | 1980 | Japan | Clinical | 91.3 | 100 | 0.15 | 98.4035 | 5127491 | 45.24 | 130 | 179113 |
| SAMEA8102998-GCA_905331625.1 | 20564 | SAMEA8102998 | GCA_905331625.1 | NonPC | Public | 1980 | Japan | Clinical | 92.5 | 100 | 0.65 | 98.4619 | 5040318 | 45.25 | 102 | 118539 |
| SAMEA8102999-GCA_905331645.1 | 20565 | SAMEA8102999 | GCA_905331645.1 | NonPC | Public | 1980 | Japan | Clinical | 91.4 | 99.95 | 0.25 | 98.3713 | 5171411 | 45.04 | 174 | 130140 |
| SAMEA8103000-GCA_905331635.1 | 20566 | SAMEA8103000 | GCA_905331635.1 | NonPC | Public | 1980 | Japan | Clinical | 91.6 | 99.9 | 0.09 | 98.4138 | 5084768 | 45.22 | 189 | 92817 |
| SAMEA8103001-GCA_905331655.1 | ERS5790079 | SAMEA8103001 | GCA_905331655.1 | NonPC | Public | 2010 | China | Clinical | 90.6 | 100 | 0.37 | 98.3142 | 5360015 | 45.16 | 184 | 73669 |
| SAMEA8103002-GCA_905331665.1 | ERS5790080 | SAMEA8103002 | GCA_905331665.1 | NonPC | Public | 2012 | China | Clinical | 91.1 | 100 | 0.09 | 98.4458 | 4998028 | 45.34 | 142 | 104152 |
| SAMEA8103003-GCA_905331695.1 | ERS5790081 | SAMEA8103003 | GCA_905331695.1 | NonPC | Public | 2012 | China | Clinical | 91.4 | 100 | 0.21 | 98.3674 | 4972417 | 45.38 | 154 | 69530 |
| SAMEA8103004-GCA_905331715.1 | ERS5790082 | SAMEA8103004 | GCA_905331715.1 | NonPC | Public | 2016 | China | Clinical | 90.8 | 100 | 0.11 | 98.3644 | 5021538 | 45.29 | 173 | 88827 |
| SAMEA8103005-GCA_905331845.1 | ERS5790083 | SAMEA8103005 | GCA_905331845.1 | NonPC | Public | 2016 | China | Clinical | 90.7 | 100 | 0.2 | 98.405 | 5056797 | 45.19 | 174 | 88609 |
| SAMEA8103006-GCA_905331875.1 | ERS5790084 | SAMEA8103006 | GCA_905331875.1 | NonPC | Public | 2016 | China | Clinical | 91.3 | 100 | 0.1 | 98.4733 | 4894534 | 45.4 | 145 | 128989 |
| SAMEA8103007-GCA_905331935.1 | ERS5790085 | SAMEA8103007 | GCA_905331935.1 | NonPC | Public | 2016 | China | Clinical | 88.9 | 100 | 0.35 | 98.392 | 5075576 | 45.17 | 159 | 90271 |
| SAMEA8103008-GCA_905331955.1 | ERS5790086 | SAMEA8103008 | GCA_905331955.1 | NonPC | Public | 2016 | China | Clinical | 90.9 | 99.97 | 0.16 | 98.4731 | 5034417 | 45.25 | 182 | 88744 |
| SAMEA8103009-GCA_905331915.1 | ERS5790087 | SAMEA8103009 | GCA_905331915.1 | NonPC | Public | 2016 | China | Clinical | 92.6 | 99.9 | 0.3 | 98.4771 | 5103069 | 45.16 | 185 | 79095 |
| SAMEA8103010-GCA_905331905.1 | ERS5790088 | SAMEA8103010 | GCA_905331905.1 | NonPC | Public | 2016 | China | Clinical | 89.5 | 100 | 0.08 | 98.3291 | 5182582 | 45.15 | 131 | 138125 |
| SAMEA8103011-GCA_905331945.1 | ERS5790089 | SAMEA8103011 | GCA_905331945.1 | NonPC | Public | 2016 | China | Clinical | 92 | 100 | 0.47 | 98.4184 | 5164947 | 45.27 | 135 | 102813 |
| SAMEA8103012-GCA_905331925.1 | ERS5790090 | SAMEA8103012 | GCA_905331925.1 | NonPC | Public | 2016 | China | Clinical | 88.1 | 100 | 0.33 | 98.3817 | 4983248 | 45.2 | 192 | 92828 |
| SAMEA8103013-GCA_905331895.1 | ERS5790091 | SAMEA8103013 | GCA_905331895.1 | NonPC | Public | 2016 | China | Clinical | 89.5 | 99.91 | 0.51 | 98.3963 | 5070243 | 45.17 | 112 | 107714 |
| SAMEA8103014-GCA_905331965.1 | ERS5790092 | SAMEA8103014 | GCA_905331965.1 | NonPC | Public | 2016 | China | Clinical | 90.9 | 100 | 0.1 | 98.4178 | 5074054 | 45.2 | 190 | 95151 |
| SAMEA8103015-GCA_905331995.1 | ERS5790093 | SAMEA8103015 | GCA_905331995.1 | NonPC | Public | 1980 | Japan | Clinical | 92.1 | 100 | 0.14 | 98.3256 | 5142793 | 45.18 | 35 | 401934 |
| SAMEA8103016-GCA_905331885.1 | ERS5790094 | SAMEA8103016 | GCA_905331885.1 | NonPC | Public | 1980 | Japan | Clinical | 91.9 | 100 | 0.46 | 98.2374 | 5079860 | 45.19 | 31 | 603376 |
| SAMEA8103017-GCA_905331815.1 | ERS5790095 | SAMEA8103017 | GCA_905331815.1 | NonPC | Public | 1980 | Japan | Clinical | 93 | 100 | 0.19 | 98.3631 | 5097725 | 45.27 | 34 | 427715 |
| SAMEA8103018-GCA_905331765.1 | ERS5790096 | SAMEA8103018 | GCA_905331765.1 | PC | Public | NA | Japan | Clinical | 97.7 | 100 | 0.01 | 99.9059 | 5192887 | 45.32 | 41 | 426363 |

| | | | | | | | | | | | | | | | | |
|---|---|---|---|---|---|---|---|---|---|---|---|---|---|---|---|---|
| SAMEA8103019-GCA_905332005.1 | ERS5790097 | SAMEA8103019 | GCA_905332005.1 | PrePC | Public | 1980 | Japan | Clinical | 94.8 | 100 | 0.08 | 99.8464 | 5021139 | 45.41 | 57 | 353994 |
| SAMEA8103020-GCA_905331975.1 | ERS5790098 | SAMEA8103020 | GCA_905331975.1 | NonPC | Public | 1980 | Japan | Clinical | 92.3 | 100 | 0 | 98.4066 | 5140215 | 45.15 | 38 | 499849 |
| SAMEA8103021-GCA_905331985.1 | ERS5790099 | SAMEA8103021 | GCA_905331985.1 | NonPC | Public | 1980 | Japan | Clinical | 92.3 | 100 | 0.05 | 98.336 | 4990978 | 45.3 | 37 | 547356 |
| SAMEA8103022-GCA_905331855.1 | ERS5790100 | SAMEA8103022 | GCA_905331855.1 | NonPC | Public | 1980 | Japan | Clinical | 91.4 | 100 | 0.31 | 98.3016 | 5136065 | 45.29 | 52 | 405122 |
| SAMEA8103023-GCA_905331835.1 | ERS5790101 | SAMEA8103023 | GCA_905331835.1 | NonPC | Public | 1980 | Japan | Clinical | 90.7 | 99.18 | 0.03 | 98.3064 | 5106629 | 45.24 | 44 | 312626 |
| SAMEA8103024-GCA_905331865.1 | ERS5790102 | SAMEA8103024 | GCA_905331865.1 | NonPC | Public | 1980 | Japan | Clinical | 90.9 | 100 | 0 | 98.3385 | 5054905 | 45.32 | 42 | 377072 |
| SAMEA8103025-GCA_905331825.1 | ERS5790103 | SAMEA8103025 | GCA_905331825.1 | NonPC | Public | 1980 | Japan | Clinical | 91.8 | 100 | 0.01 | 98.3842 | 5180062 | 45.27 | 33 | 417353 |
| SAMEA8103026-GCA_905331805.1 | ERS5790104 | SAMEA8103026 | GCA_905331805.1 | NonPC | Public | 2010 | China | Clinical | 91.3 | 100 | 0.04 | 98.3585 | 4975337 | 45.41 | 47 | 522212 |
| SAMEA8103027-GCA_905331785.1 | ERS5790105 | SAMEA8103027 | GCA_905331785.1 | PC | Public | 2012 | China | Clinical | 97.3 | 100 | 0 | 99.8541 | 5191259 | 45.25 | 47 | 373095 |
| SAMEA8103028-GCA_905331795.1 | ERS5790106 | SAMEA8103028 | GCA_905331795.1 | NonPC | Public | 2012 | China | Clinical | 87.6 | 100 | 0.44 | 98.3019 | 4977473 | 45.25 | 54 | 368025 |
| SAMEA8103029-GCA_905331775.1 | ERS5790107 | SAMEA8103029 | GCA_905331775.1 | PC | Public | 2016 | China | Clinical | 97.7 | 100 | 0 | 99.9222 | 5162116 | 45.24 | 59 | 373695 |
| SAMEA8103030-GCA_905331755.1 | ERS5790108 | SAMEA8103030 | GCA_905331755.1 | NonPC | Public | 2016 | China | Clinical | 92.2 | 100 | 0.05 | 98.1177 | 5118485 | 45.24 | 33 | 470297 |
| SAMEA8103031-GCA_905331745.1 | ERS5790109 | SAMEA8103031 | GCA_905331745.1 | NonPC | Public | 2016 | China | Clinical | 91.2 | 100 | 0.05 | 98.3434 | 5092778 | 45.25 | 52 | 315329 |
| SAMEA8103032-GCA_905331735.1 | ERS5790110 | SAMEA8103032 | GCA_905331735.1 | PC | Public | 2016 | China | Clinical | 97.4 | 100 | 0.07 | 99.9042 | 5180936 | 45.28 | 55 | 373744 |
| SAMEA8103033-GCA_905331725.1 | ERS5790111 | SAMEA8103033 | GCA_905331725.1 | NonPC | Public | 2016 | China | Clinical | 87.9 | 100 | 0.43 | 98.2147 | 5112043 | 45.15 | 76 | 557312 |
| SAMEA8103035-GCA_905331685.1 | ERS5790113 | SAMEA8103035 | GCA_905331685.1 | NonPC | Public | 2016 | China | Clinical | 92.9 | 100 | 0.38 | 98.4408 | 5104663 | 45.23 | 49 | 200323 |
| SAMEA8103036-GCA_905331705.1 | ERS5790114 | SAMEA8103036 | GCA_905331705.1 | NonPC | Public | 2016 | China | Clinical | 89.7 | 100 | 0.27 | 98.4239 | 5051817 | 45.34 | 37 | 411517 |
| SAMN01816333-GCF_000707585.2 | K1203 | SAMN01816333 | GCF_000707585.2 | NonPC | Public | 2004 | USA | Env | 91.5 | 99.73 | 0.09 | 98.283 | 5264419 | 45.13 | 74 | 315966 |
| SAMN01816334-GCF_002072925.1 | NY3547 | SAMN01816334 | GCF_002072925.1 | NonPC | Public | 1998 | USA | Env | 91.6 | 100 | 0.27 | 98.2961 | 5157847 | 45.17 | 71 | 221575 |
| SAMN01816336-GCF_000707605.2 | 98-513-F52 | SAMN01816336 | GCF_000707605.2 | NonPC | Public | 1998 | USA | Env | 90.5 | 100 | 0.2 | 98.2138 | 5085227 | 45.17 | 71 | 193593 |
| SAMN01923801-GCF_000522065.2 | PVCHO_Peru-288 | SAMN01923801 | GCF_000522065.2 | PC | Public | 2001 | Peru | Clinical | 98.2 | 100 | 0.01 | 99.9875 | 5173942 | 45.3 | 79 | 198215 |
| SAMN01923802-GCF_000571915.2 | EKP-021 | SAMN01923802 | GCF_000571915.2 | PC | Public | 2008 | Bangladesh | NA | 97.7 | 94.79 | 0.07 | 99.867 | 5213526 | 45.28 | 56 | 477789 |
| SAMN01923803-GCF_000558925.2 | PVCHO_VP2007-007 | SAMN01923803 | GCF_000558925.2 | NonPC | Public | 2007 | USA | NA | 90.3 | 96.75 | 0.08 | 98.1232 | 5111574 | 45.23 | 51 | 308313 |
| SAMN01923807-GCF_000522045.2 | PVCHO_SBR10290 | SAMN01923807 | GCF_000522045.2 | NonPC | Public | 1997 | USA | Clinical | 91.6 | 99.41 | 0 | 98.2922 | 5130638 | 45.27 | 45 | 464757 |
| SAMN01923894-GCF_000524535.2 | 861 | SAMN01923894 | GCF_000524535.2 | PC | Public | 2006 | USA | NA | 98.4 | 98.57 | 0 | 99.9707 | 5170911 | 45.28 | 51 | 473747 |
| SAMN01923895-GCF_000593325.2 | PVCHO_VPTS-2009 | SAMN01923895 | GCF_000593325.2 | NonPC | Public | 2009 | USA | NA | 89.7 | 100 | 0.26 | 98.299 | 5071843 | 45.32 | 59 | 311642 |
| SAMN01923897-GCF_000522025.2 | IDH02189 | SAMN01923897 | GCF_000522025.2 | PC | Public | 2009 | India | Clinical | 98 | 99.56 | 0 | 99.9875 | 5088693 | 45.28 | 59 | 395465 |
| SAMN01923903-GCF_000522005.2 | EKP-028 | SAMN01923903 | GCF_000522005.2 | PC | Public | 2008 | Bangladesh | Env | 98.2 | 94.1 | 0 | 99.9055 | 5190077 | 45.29 | 58 | 443052 |
| SAMN01924102-SRR748523 | 605 | SAMN01924102 | SRR748523 | PC | Public | 2006 | USA | NA | 98.3 | 100 | 0 | 99.9774 | 5144235 | 45.28 | 57 | 317807 |
| SAMN01924103-SRR748524 | 3256 | SAMN01924103 | SRR748524 | NonPC | Public | 2007 | USA | NA | 91.4 | 100 | 0 | 98.3603 | 5074895 | 45.28 | 44 | 718922 |
| SAMN01924111-GCF_000521825.2 | IDH02640 | SAMN01924111 | GCF_000521825.2 | PC | Public | 2009 | India | Clinical | 98 | 99.33 | 0 | 99.925 | 5211636 | 45.12 | 58 | 501700 |
| SAMN01924113-GCF_000558885.2 | PVCHO_V14/01 | SAMN01924113 | GCF_000558885.2 | PC | Public | 2001 | Chile | NA | 98.6 | 100 | 0.14 | 99.9738 | 5249071 | 45.1 | 64 | 267813 |
| SAMN01924114-GCF_000593345.2 | PVCHO_VPTS-2010 | SAMN01924114 | GCF_000593345.2 | NonPC | Public | 2010 | USA | NA | 92 | 92.43 | 0.35 | 98.2717 | 5284886 | 45.1 | 32 | 546552 |
| SAMN01924261-GCF_000500755.2 | 12310 | SAMN01924261 | GCF_000500755.2 | NonPC | Public | 2006 | USA | Clinical | 91.4 | 99.56 | 0 | 98.3204 | 5133199 | 45.26 | 40 | 719498 |
| SAMN01924275-GCF_000510585.2 | EKP-008 | SAMN01924275 | GCF_000510585.2 | NonPC | Public | 2007 | Bangladesh | NA | 92.8 | 98.23 | 0 | 98.3111 | 5304418 | 45.16 | 38 | 514908 |
| SAMN01924294-SRR748858 | VP-48 | SAMN01924294 | SRR748858 | NonPC | Public | 1996 | India | NA | 91.4 | 100 | 0 | 98.3747 | 5109260 | 45.3 | 46 | 285923 |
| SAMN01924295-SRR748859 | 10296 | SAMN01924295 | SRR748859 | NonPC | Public | 1997 | USA | NA | 91.5 | 100 | 0 | 98.263 | 5093788 | 45.27 | 38 | 548388 |
| SAMN01933054-GCF_000593285.2 | PVCHO_VP-48 | SAMN01933054 | GCF_000593285.2 | PC | Public | 1996 | India | NA | 98.7 | 98.33 | 0 | 99.9813 | 5116196 | 45.31 | 50 | 474978 |
| SAMN01940373-GCF_000593365.2 | PVCHO_VPTS-2010_2 | SAMN01940373 | GCF_000593365.2 | NonPC | Public | 2010 | USA | NA | 87.8 | 93.91 | 0.3 | 97.985 | 5062057 | 45.37 | 53 | 417603 |
| SAMN01940374-GCF_000593305.2 | PVCHO_VPCR-2009 | SAMN01940374 | GCF_000593305.2 | NonPC | Public | 2009 | USA | NA | 91.8 | 100 | 0.12 | 98.3356 | 5087696 | 45.31 | 58 | 391331 |
| SAMN01990931-SRR748210 | 50 | SAMN01990931 | SRR748210 | NonPC | Public | 2006 | USA | NA | 90.2 | 100 | 0.14 | 98.2196 | 5057152 | 45.16 | 48 | 425485 |
| SAMN02179882-GCF_000430405.1 | FDA_R31 | SAMN02179882 | GCF_000430405.1 | NonPC | Public | 2007 | USA | Env | 90.1 | 100 | 0 | 98.0533 | 5223562 | 45.3 | 2 | 3362228 |
| SAMN02179884-GCF_000430425.1 | CDC_K4557 | SAMN02179884 | GCF_000430425.1 | NonPC | Public | 2006 | USA | Clinical | 92.6 | 100 | 0 | 98.4157 | 5138575 | 45.36 | 2 | 3331580 |
| SAMN02183022-GCF_000454165.1 | PVCHO_VP2007-095 | SAMN02183022 | GCF_000454165.1 | NonPC | Public | 2007 | USA | NA | 90.3 | 99.73 | 0.04 | 98.2917 | 5175863 | 45.33 | 21 | 485232 |
| SAMN02189718-GCF_000454265.1 | NIHCB0603 | SAMN02189718 | GCF_000454265.1 | PC | Public | 2006 | Bangladesh | NA | 99.8 | 99.91 | 0.27 | 99.9845 | 5225215 | 45.38 | 13 | 1292484 |
| SAMN02189719-GCF_000454245.1 | 3259 | SAMN02189719 | GCF_000454245.1 | NonPC | Public | 2007 | USA | NA | 93.9 | 100 | 0 | 98.329 | 5360869 | 45.25 | 6 | 3370862 |
| SAMN02189720-GCF_000477475.1 | NIHCB0757 | SAMN02189720 | GCF_000477475.1 | NonPC | Public | 2006 | Bangladesh | NA | 90.9 | 100 | 0 | 98.3741 | 5262465 | 45.29 | 4 | 3378377 |
| SAMN02189721-GCF_000454225.1 | PVCHO_VP250 | SAMN02189721 | GCF_000454225.1 | PC | Public | 1998 | India | NA | 99.1 | 100 | 0.27 | 99.9431 | 5202656 | 45.37 | 5 | 3317379 |

| | | | | | | | | | | | | | | | | |
|---|---|---|---|---|---|---|---|---|---|---|---|---|---|---|---|---|
| SAMN02190054-GCF_000454455.1 | 949 | SAMN02190054 | GCF_000454455.1 | PC | Public | 2006 | USA | NA | 99.8 | 100 | 0.14 | 99.9922 | 5179240 | 45.39 | 7 | 972111 |
| SAMN02190055-GCF_000454145.1 | PVCHO_VP-NY4 | SAMN02190055 | GCF_000454145.1 | PC | Public | 1997 | India | NA | 99.7 | 100 | 0 | 99.9777 | 5247801 | 45.39 | 14 | 1120023 |
| SAMN02190056-GCF_000454185.1 | PVCHO_VP232 | SAMN02190056 | GCF_000454185.1 | PC | Public | 1998 | India | NA | 97.4 | 99.43 | 0.09 | 99.9365 | 5123959 | 45.4 | 52 | 171658 |
| SAMN02190057-GCF_000454205.1 | 10290 | SAMN02190057 | GCF_000454205.1 | NonPC | Public | 1997 | USA | NA | 92.3 | 100 | 0.23 | 98.3099 | 5183035 | 45.4 | 13 | 2604594 |
| SAMN02204303-GCA_000877625.1 | 49 | SAMN02204303 | GCA_000877625.1 | NonPC | Public | 2007 | USA | Env | 90.7 | 97.42 | 1.39 | 98.3516 | 5119579 | 45.29 | 325 | 29474 |
| SAMN02204304-GCF_000877415.2 | VP551 | SAMN02204304 | GCF_000877415.2 | PC | Public | 2007 | USA | Env | 97.5 | 99.18 | 0 | 99.9687 | 5098994 | 45.28 | 58 | 260061 |
| SAMN02204305-GCA_000877605.1 | VP766 | SAMN02204305 | GCA_000877605.1 | NonPC | Public | 2007 | USA | Env | 90.3 | 98.43 | 0.68 | 98.2577 | 5184533 | 45.22 | 290 | 32385 |
| SAMN02204306-GCF_000877405.1 | 846 | SAMN02204306 | GCF_000877405.1 | NonPC | Public | 2007 | USA | Env | 91.4 | 100 | 0 | 98.3111 | 5070127 | 45.27 | 69 | 264478 |
| SAMN02204307-GCF_000877485.2 | 863 | SAMN02204307 | GCF_000877485.2 | PC | Public | 2007 | USA | Env | 98.4 | 100 | 0 | 99.969 | 5156364 | 45.29 | 58 | 321400 |
| SAMN02204308-GCF_000877475.2 | 930 | SAMN02204308 | GCF_000877475.2 | PC | Public | 2007 | USA | Env | 98.4 | 100 | 0 | 99.9863 | 5103160 | 45.33 | 59 | 259639 |
| SAMN02204309-GCF_000877495.1 | 3324 | SAMN02204309 | GCF_000877495.1 | NonPC | Public | 2007 | USA | Clinical | 91.4 | 100 | 0 | 98.3268 | 5088910 | 45.25 | 59 | 348180 |
| SAMN02204311-GCF_000877595.2 | 3631 | SAMN02204311 | GCF_000877595.2 | NonPC | Public | 2007 | USA | Clinical | 89.7 | 99.46 | 0.87 | 98.2999 | 5064925 | 45.31 | 290 | 121810 |
| SAMN02204312-GCA_000877755.1 | 3644 | SAMN02204312 | GCA_000877755.1 | NonPC | Public | 2007 | USA | Clinical | 88.3 | 98.88 | 0.61 | 98.073 | 5003880 | 45.31 | 332 | 31249 |
| SAMN02204313-GCF_000877765.2 | 3646 | SAMN02204313 | GCF_000877765.2 | NonPC | Public | 2007 | USA | Clinical | 89.9 | 98.64 | 0.33 | 98.2819 | 5116657 | 45.18 | 70 | 307422 |
| SAMN02204314-GCF_000877535.1 | 12315 | SAMN02204314 | GCF_000877535.1 | NonPC | Public | 2006 | USA | Clinical | 91.4 | 100 | 0 | 98.3343 | 5078852 | 45.26 | 72 | 270580 |
| SAMN02204315-GCA_000877675.1 | 901128 | SAMN02204315 | GCA_000877675.1 | NonPC | Public | 1997 | USA | Clinical | 89.5 | 98.15 | 0.39 | 98.1945 | 5232629 | 45.17 | 332 | 31795 |
| SAMN02204316-GCF_000877425.1 | 97-10290 | SAMN02204316 | GCF_000877425.1 | NonPC | Public | 1997 | USA | Clinical | 91.5 | 100 | 0 | 98.3132 | 5100708 | 45.26 | 51 | 236714 |
| SAMN02204317-GCF_000877685.1 | EN2910 | SAMN02204317 | GCF_000877685.1 | NonPC | Public | 2000 | USA | Clinical | 91.5 | 100 | 0 | 98.3121 | 5088775 | 45.26 | 37 | 398882 |
| SAMN02204318-GCA_000877715.1 | EN9701072 | SAMN02204318 | GCA_000877715.1 | NonPC | Public | 1997 | USA | Clinical | 87.9 | 98.53 | 0.54 | 98.113 | 4943293 | 45.29 | 356 | 27006 |
| SAMN02204319-GCA_000877725.1 | EN9701121 | SAMN02204319 | GCA_000877725.1 | NonPC | Public | 1997 | USA | Clinical | 90.2 | 98.54 | 0.66 | 98.2613 | 5084119 | 45.25 | 309 | 32971 |
| SAMN02204320-GCF_000877555.1 | EN9701173 | SAMN02204320 | GCF_000877555.1 | NonPC | Public | 1997 | USA | Clinical | 91.5 | 100 | 0 | 98.3087 | 5089104 | 45.26 | 42 | 440310 |
| SAMN02204321-GCF_000877565.1 | EN9901310 | SAMN02204321 | GCF_000877565.1 | NonPC | Public | 1999 | USA | Clinical | 91.5 | 100 | 0 | 98.2903 | 5088251 | 45.26 | 49 | 439695 |
| SAMN02252525-GCF_000701045.2 | CFSAN001595 | SAMN02252525 | GCF_000701045.2 | NonPC | Public | 1996 | Peru | Clinical | 92.3 | 100 | 0.19 | 98.36 | 5097454 | 45.23 | 41 | 406461 |
| SAMN02338865-GCF_000492095.1 | S001 | SAMN02338865 | GCF_000492095.1 | NonPC | Public | NA | Thailand | NA | 89.7 | 99.89 | 1.6 | 98.2837 | 5007559 | 45.39 | 421 | 24000 |
| SAMN02338866-GCF_000492075.1 | S002 | SAMN02338866 | GCF_000492075.1 | NonPC | Public | 1999 | China | NA | 88.8 | 99.71 | 0.66 | 98.3463 | 5013889 | 45.09 | 502 | 20468 |
| SAMN02338867-GCF_000492055.1 | S003 | SAMN02338867 | GCF_000492055.1 | PrePC | Public | NA | China | Clinical | 93.1 | 99.73 | 1.36 | 99.2696 | 4986349 | 45.33 | 414 | 21622 |
| SAMN02338868-GCF_000492035.1 | S004 | SAMN02338868 | GCF_000492035.1 | PrePC | Public | 1985 | Maldives | Clinical | 92.8 | 99.82 | 0.82 | 99.3044 | 4987344 | 45.32 | 634 | 14023 |
| SAMN02338869-GCF_000492015.1 | S005 | SAMN02338869 | GCF_000492015.1 | PrePC | Public | NA | Thailand | Clinical | 93.1 | 99.77 | 0.62 | 99.2671 | 4996782 | 45.33 | 366 | 28304 |
| SAMN02338870-GCF_000491995.1 | S008 | SAMN02338870 | GCF_000491995.1 | PrePC | Public | 1987 | Thailand | Clinical | 93.1 | 99.64 | 0.48 | 99.2726 | 5056265 | 45.3 | 387 | 24941 |
| SAMN02338871-GCF_000491975.1 | S009 | SAMN02338871 | GCF_000491975.1 | PrePC | Public | 1987 | Thailand | Clinical | 93.1 | 99.98 | 0.6 | 99.2616 | 5058825 | 45.3 | 461 | 19460 |
| SAMN02338872-GCF_000491955.1 | S011 | SAMN02338872 | GCF_000491955.1 | PrePC | Public | 1990 | Thailand | Clinical | 93 | 99.91 | 0.46 | 99.2545 | 5055045 | 45.31 | 322 | 30146 |
| SAMN02338873-GCF_000491935.1 | S012 | SAMN02338873 | GCF_000491935.1 | PrePC | Public | 1990 | Thailand | Clinical | 93.1 | 99.94 | 0.95 | 99.2626 | 5059926 | 45.3 | 481 | 18090 |
| SAMN02338874-GCF_000491915.1 | S013 | SAMN02338874 | GCF_000491915.1 | PrePC | Public | 1996 | China | Clinical | 92.6 | 99.73 | 0.82 | 99.3092 | 4981412 | 45.44 | 347 | 31330 |
| SAMN02338875-GCF_000491895.1 | S014 | SAMN02338875 | GCF_000491895.1 | NonPC | Public | 1992 | China | NA | 92 | 99.98 | 1.01 | 98.4501 | 5139007 | 45.21 | 477 | 18650 |
| SAMN02338876-GCF_000491875.1 | S015 | SAMN02338876 | GCF_000491875.1 | NonPC | Public | 1992 | China | NA | 91.9 | 99.98 | 0.78 | 98.4348 | 5131263 | 45.2 | 490 | 17831 |
| SAMN02338877-GCF_000491855.1 | S016 | SAMN02338877 | GCF_000491855.1 | NonPC | Public | 1992 | China | NA | 92.3 | 99.98 | 1.05 | 98.4403 | 5083882 | 45.24 | 524 | 15462 |
| SAMN02338878-GCF_000491835.1 | S017 | SAMN02338878 | GCF_000491835.1 | NonPC | Public | 1992 | China | NA | 91.9 | 99.95 | 0.41 | 98.422 | 5158071 | 45.14 | 285 | 36112 |
| SAMN02338879-GCF_000491815.1 | S018 | SAMN02338879 | GCF_000491815.1 | NonPC | Public | 1993 | China | NA | 92.3 | 99.98 | 0.64 | 98.4353 | 5122600 | 45.16 | 614 | 15209 |
| SAMN02338880-GCF_000491795.1 | S019 | SAMN02338880 | GCF_000491795.1 | NonPC | Public | 1998 | USA | NA | 90.9 | 99.95 | 0.82 | 98.3286 | 5056371 | 45.2 | 392 | 23909 |
| SAMN02338881-GCF_000491775.1 | S020 | SAMN02338881 | GCF_000491775.1 | NonPC | Public | 1994 | China | NA | 90.1 | 99.98 | 1.34 | 98.3517 | 5157889 | 45.18 | 342 | 32890 |
| SAMN02338882-GCF_000491755.1 | S021 | SAMN02338882 | GCF_000491755.1 | NonPC | Public | 1999 | India | NA | 90.3 | 99.62 | 0.4 | 98.3255 | 5183664 | 45.26 | 460 | 21666 |
| SAMN02338883-GCF_000491735.1 | S022 | SAMN02338883 | GCF_000491735.1 | NonPC | Public | 1984 | Japan | NA | 92.5 | 99.94 | 0.22 | 98.4348 | 5192613 | 45.2 | 346 | 32171 |
| SAMN02338884-GCF_000491715.1 | S023 | SAMN02338884 | GCF_000491715.1 | NonPC | Public | 1994 | China | NA | 92.8 | 99.91 | 1.66 | 98.4818 | 5090798 | 45.21 | 583 | 15161 |
| SAMN02338885-GCF_000491695.1 | S024 | SAMN02338885 | GCF_000491695.1 | NonPC | Public | 1998 | China | NA | 92.2 | 99.98 | 0.02 | 98.42 | 5069074 | 45.18 | 376 | 22981 |
| SAMN02338886-GCF_000491675.1 | S025 | SAMN02338886 | GCF_000491675.1 | NonPC | Public | 1992 | China | NA | 91.9 | 99.98 | 0.7 | 98.4219 | 5158158 | 45.14 | 297 | 34958 |
| SAMN02338887-GCF_000491655.1 | S026 | SAMN02338887 | GCF_000491655.1 | NonPC | Public | NA | Thailand | NA | 91.7 | 99.74 | 0.44 | 98.4197 | 5043549 | 45.25 | 382 | 22512 |
| SAMN02338888-GCF_000491635.1 | S027 | SAMN02338888 | GCF_000491635.1 | NonPC | Public | NA | USA | NA | 91.9 | 99.62 | 0.85 | 98.4202 | 5094965 | 45.19 | 408 | 20530 |
| SAMN02338889-GCF_000491615.1 | S028 | SAMN02338889 | GCF_000491615.1 | NonPC | Public | NA | Thailand | NA | 92.6 | 99.95 | 0.19 | 98.4259 | 5042426 | 45.26 | 344 | 32572 |

| | | | | | | | | | | | | | | | | |
|---|---|---|---|---|---|---|---|---|---|---|---|---|---|---|---|---|
| SAMN02338890-GCF_000491595.1 | S029 | SAMN02338890 | GCF_000491595.1 | NonPC | Public | 1981 | Spain | NA | 88.8 | 99.73 | 0.6 | 98.4108 | 4918302 | 45.54 | 426 | 24807 |
| SAMN02338891-GCF_000491575.1 | S030 | SAMN02338891 | GCF_000491575.1 | NonPC | Public | 1999 | India | NA | 91.7 | 99.86 | 0.63 | 98.3161 | 5070237 | 45.25 | 413 | 24210 |
| SAMN02338892-GCF_000491555.1 | S031 | SAMN02338892 | GCF_000491555.1 | NonPC | Public | 1984 | Japan | NA | 91.7 | 99.85 | 0.35 | 98.4291 | 4972305 | 45.32 | 294 | 35058 |
| SAMN02338893-GCF_000491535.1 | S032 | SAMN02338893 | GCF_000491535.1 | NonPC | Public | 1998 | Philippines | NA | 92.3 | 99.81 | 1.53 | 98.4072 | 5087285 | 45.2 | 668 | 13893 |
| SAMN02338894-GCF_000491515.1 | S033 | SAMN02338894 | GCF_000491515.1 | NonPC | Public | 1994 | China | NA | 92.3 | 99.85 | 0.16 | 98.4356 | 5142761 | 45.19 | 382 | 29792 |
| SAMN02338895-GCF_000491495.1 | S034 | SAMN02338895 | GCF_000491495.1 | NonPC | Public | 1999 | India | NA | 92.3 | 99.81 | 0.07 | 98.4499 | 5123789 | 45.18 | 383 | 25656 |
| SAMN02338896-GCF_000491475.1 | S035 | SAMN02338896 | GCF_000491475.1 | NonPC | Public | 1984 | Japan | NA | 92.4 | 99.98 | 0.44 | 98.469 | 5092158 | 45.19 | 450 | 21594 |
| SAMN02338897-GCF_000491455.1 | S036 | SAMN02338897 | GCF_000491455.1 | NonPC | Public | NA | Thailand | NA | 91.3 | 99.98 | 0.18 | 98.4615 | 5008748 | 45.26 | 418 | 20685 |
| SAMN02338898-GCF_000491435.1 | S037 | SAMN02338898 | GCF_000491435.1 | NonPC | Public | 1994 | USA | NA | 90.9 | 99.65 | 2.19 | 98.324 | 5029130 | 45.26 | 692 | 12391 |
| SAMN02338899-GCF_000491415.1 | S038 | SAMN02338899 | GCF_000491415.1 | NonPC | Public | 1982 | USA | NA | 90.9 | 99.68 | 0.89 | 98.326 | 5075757 | 45.24 | 715 | 11986 |
| SAMN02338900-GCF_000491395.1 | S039 | SAMN02338900 | GCF_000491395.1 | NonPC | Public | NA | China | NA | 88.9 | 99.71 | 0.93 | 98.3679 | 4988754 | 45.23 | 684 | 13001 |
| SAMN02338901-GCF_000491375.1 | S040 | SAMN02338901 | GCF_000491375.1 | NonPC | Public | 1990 | Thailand | NA | 89.4 | 99.89 | 0.77 | 98.3453 | 4961865 | 45.26 | 433 | 19879 |
| SAMN02338902-GCF_000491355.1 | S041 | SAMN02338902 | GCF_000491355.1 | NonPC | Public | 1993 | China | NA | 92.5 | 99.54 | 0.48 | 98.4108 | 5060248 | 45.26 | 611 | 14941 |
| SAMN02338903-GCF_000491335.1 | S042 | SAMN02338903 | GCF_000491335.1 | NonPC | Public | 1984 | Japan | NA | 92.5 | 99.78 | 1.3 | 98.4303 | 5082663 | 45.25 | 637 | 14103 |
| SAMN02338904-GCF_000491315.1 | S043 | SAMN02338904 | GCF_000491315.1 | NonPC | Public | NA | Thailand | NA | 90.6 | 99.85 | 0.68 | 98.3055 | 5070140 | 45.17 | 634 | 14730 |
| SAMN02338905-GCF_000491295.1 | S044 | SAMN02338905 | GCF_000491295.1 | NonPC | Public | 1990 | Thailand | NA | 90.6 | 99.75 | 0.68 | 98.4259 | 5017231 | 45.31 | 660 | 13374 |
| SAMN02338906-GCF_000491275.1 | S045 | SAMN02338906 | GCF_000491275.1 | NonPC | Public | 1990 | Thailand | NA | 90.6 | 99.84 | 0.45 | 98.4111 | 5019993 | 45.32 | 439 | 21335 |
| SAMN02338907-GCF_000491255.1 | S046 | SAMN02338907 | GCF_000491255.1 | NonPC | Public | 1982 | Spain | NA | 89.8 | 99.92 | 1.47 | 98.2817 | 5180931 | 45.21 | 818 | 11177 |
| SAMN02338908-GCF_000491235.1 | S047 | SAMN02338908 | GCF_000491235.1 | NonPC | Public | 1981 | Spain | NA | 87.9 | 99.95 | 0.55 | 98.3579 | 5046699 | 45.23 | 495 | 24746 |
| SAMN02338909-GCF_000491215.1 | S048 | SAMN02338909 | GCF_000491215.1 | NonPC | Public | 1997 | USA | NA | 90.7 | 99.98 | 0.33 | 98.3343 | 5191346 | 45.17 | 457 | 20813 |
| SAMN02338910-GCF_000491195.1 | S049 | SAMN02338910 | GCF_000491195.1 | NonPC | Public | 1984 | Japan | NA | 92.3 | 99.71 | 1.34 | 98.407 | 5195371 | 45.03 | 604 | 16173 |
| SAMN02338911-GCF_000491175.1 | S052 | SAMN02338911 | GCF_000491175.1 | NonPC | Public | 1975 | Spain | NA | 89.8 | 99.2 | 1.18 | 98.3 | 4970536 | 45.34 | 511 | 18199 |
| SAMN02338912-GCF_000491155.1 | S053 | SAMN02338912 | GCF_000491155.1 | NonPC | Public | 1990 | Thailand | NA | 90.4 | 99.66 | 1.14 | 98.4518 | 4958528 | 45.41 | 649 | 13539 |
| SAMN02338913-GCF_000491135.1 | S054 | SAMN02338913 | GCF_000491135.1 | NonPC | Public | NA | Thailand | NA | 91.6 | 98.47 | 1.45 | 98.3354 | 5146238 | 45.22 | 576 | 14490 |
| SAMN02338914-GCF_000491115.1 | S055 | SAMN02338914 | GCF_000491115.1 | NonPC | Public | 1994 | China | NA | 91.2 | 99.73 | 0.67 | 98.434 | 5015346 | 45.31 | 343 | 35392 |
| SAMN02338915-GCF_000491095.1 | S056 | SAMN02338915 | GCF_000491095.1 | NonPC | Public | 1994 | China | NA | 91.6 | 99.24 | 2.67 | 98.3399 | 5216460 | 45.2 | 713 | 12396 |
| SAMN02338916-GCF_000491075.1 | S057 | SAMN02338916 | GCF_000491075.1 | NonPC | Public | 1994 | China | NA | 91.5 | 99.95 | 0.55 | 98.4341 | 5095226 | 45.17 | 482 | 20175 |
| SAMN02338917-GCF_000491055.1 | S058 | SAMN02338917 | GCF_000491055.1 | NonPC | Public | 1970 | Japan | NA | 91.1 | 99.85 | 0.61 | 98.4456 | 5007009 | 45.25 | 427 | 24420 |
| SAMN02338918-GCF_000491035.1 | S060 | SAMN02338918 | GCF_000491035.1 | NonPC | Public | 1992 | China | NA | 91.1 | 99 | 1.66 | 98.4444 | 4909808 | 45.39 | 646 | 13928 |
| SAMN02338919-GCF_000491015.1 | S062 | SAMN02338919 | GCF_000491015.1 | PC | Public | 1998 | Singapore | NA | 97.1 | 99.44 | 0.33 | 99.9454 | 5037173 | 45.26 | 374 | 22532 |
| SAMN02338920-GCF_000490995.1 | S063 | SAMN02338920 | GCF_000490995.1 | PC | Public | 1998 | China | NA | 97 | 99.85 | 0.48 | 99.9233 | 5098640 | 45.22 | 428 | 22691 |
| SAMN02338921-GCF_000490975.1 | S064 | SAMN02338921 | GCF_000490975.1 | PC | Public | 1998 | China | NA | 97.6 | 99.81 | 1.05 | 99.9956 | 5034158 | 45.27 | 574 | 14862 |
| SAMN02338922-GCF_000490955.1 | S065 | SAMN02338922 | GCF_000490955.1 | PC | Public | 1998 | China | NA | 96.9 | 99.98 | 0.11 | 99.9337 | 5032170 | 45.27 | 422 | 21258 |
| SAMN02338923-GCF_000490935.1 | S066 | SAMN02338923 | GCF_000490935.1 | PC | Public | 1997 | China | NA | 97.6 | 99.67 | 0.55 | 99.986 | 5031213 | 45.27 | 544 | 16046 |
| SAMN02338924-GCF_000490915.1 | S067 | SAMN02338924 | GCF_000490915.1 | PC | Public | 1997 | China | NA | 97.7 | 99.71 | 0.14 | 99.9885 | 5037978 | 45.26 | 428 | 21213 |
| SAMN02338925-GCF_000490895.1 | S068 | SAMN02338925 | GCF_000490895.1 | PC | Public | 1997 | China | NA | 97.7 | 99.69 | 0.23 | 99.9815 | 5105765 | 45.11 | 441 | 21152 |
| SAMN02338926-GCF_000490875.1 | S069 | SAMN02338926 | GCF_000490875.1 | PC | Public | NA | Thailand | Clinical | 97.5 | 99.64 | 0.09 | 99.986 | 5028624 | 45.27 | 335 | 28922 |
| SAMN02338927-GCF_000490855.1 | S070 | SAMN02338927 | GCF_000490855.1 | PC | Public | NA | Thailand | Clinical | 97.5 | 99.74 | 0.12 | 99.9953 | 5025807 | 45.29 | 547 | 15138 |
| SAMN02338928-GCF_000490835.1 | S071 | SAMN02338928 | GCF_000490835.1 | PC | Public | 1998 | Bangladesh | NA | 97 | 99.68 | 0.49 | 99.9371 | 5057184 | 45.24 | 336 | 28640 |
| SAMN02338929-GCF_000490815.1 | S072 | SAMN02338929 | GCF_000490815.1 | PC | Public | 1998 | Bangladesh | NA | 97.7 | 99.98 | 0.23 | 99.9962 | 5049210 | 45.25 | 349 | 27675 |
| SAMN02338930-GCF_000490795.1 | S073 | SAMN02338930 | GCF_000490795.1 | NonPC | Public | 1997 | Thailand | NA | 92.5 | 99.98 | 0.58 | 98.4405 | 5032891 | 45.28 | 443 | 23605 |
| SAMN02338931-GCF_000490775.1 | S074 | SAMN02338931 | GCF_000490775.1 | PC | Public | 1997 | China | NA | 97.6 | 99.98 | 0.31 | 99.9839 | 5111173 | 45.11 | 600 | 14150 |
| SAMN02338932-GCF_000490755.1 | S075 | SAMN02338932 | GCF_000490755.1 | PC | Public | 1999 | China | NA | 97.5 | 99.94 | 0.73 | 99.9714 | 5084568 | 45.25 | 627 | 14151 |
| SAMN02338933-GCF_000490735.1 | S076 | SAMN02338933 | GCF_000490735.1 | PC | Public | 1999 | China | NA | 97.6 | 99.98 | 0.31 | 99.9525 | 5085445 | 45.25 | 365 | 25100 |
| SAMN02338934-GCF_000490715.1 | S077 | SAMN02338934 | GCF_000490715.1 | PC | Public | 1999 | China | NA | 97.5 | 99.94 | 0.36 | 99.9697 | 5084409 | 45.25 | 462 | 18729 |
| SAMN02338935-GCF_000490695.1 | S078 | SAMN02338935 | GCF_000490695.1 | PC | Public | 1999 | China | NA | 97.6 | 99.95 | 0.25 | 99.9841 | 5085900 | 45.25 | 379 | 23501 |
| SAMN02338936-GCF_000490675.1 | S079 | SAMN02338936 | GCF_000490675.1 | PC | Public | NA | Indonesia | Clinical | 97.6 | 99.94 | 0.43 | 99.985 | 5079842 | 45.22 | 560 | 16129 |
| SAMN02338937-GCF_000490655.1 | S081 | SAMN02338937 | GCF_000490655.1 | PC | Public | NA | South_Korea | Clinical | 97.7 | 99.67 | 0.48 | 99.9965 | 5031080 | 45.27 | 407 | 22930 |

| | | | | | | | | | | | | | | | | |
|---|---|---|---|---|---|---|---|---|---|---|---|---|---|---|---|---|
| SAMN02338938-GCF_000490635.1 | S082 | SAMN02338938 | GCF_000490635.1 | PC | Public | NA | Thailand | Clinical | 96.5 | 99.13 | 0.64 | 99.9946 | 4970929 | 45.28 | 466 | 18688 |
| SAMN02338939-GCF_000490615.1 | S083 | SAMN02338939 | GCF_000490615.1 | PC | Public | 1998 | Japan | NA | 97.7 | 99.88 | 0.46 | 99.9766 | 5082517 | 45.21 | 350 | 28639 |
| SAMN02338940-GCF_000490595.1 | S086 | SAMN02338940 | GCF_000490595.1 | PC | Public | 1999 | Thailand | NA | 97.1 | 99.57 | 0.49 | 99.9295 | 5039274 | 45.26 | 381 | 25865 |
| SAMN02338941-GCF_000490575.1 | S087 | SAMN02338941 | GCF_000490575.1 | PC | Public | 1998 | Singapore | NA | 97 | 99.82 | 0.7 | 99.9178 | 5052727 | 45.24 | 361 | 27007 |
| SAMN02338942-GCF_000490555.1 | S088 | SAMN02338942 | GCF_000490555.1 | PC | Public | 1998 | Singapore | NA | 97 | 99.95 | 1.07 | 99.9419 | 5053483 | 45.24 | 326 | 30513 |
| SAMN02338943-GCF_000490535.1 | S090 | SAMN02338943 | GCF_000490535.1 | PC | Public | 1999 | China | NA | 96 | 99.25 | 1.43 | 99.9452 | 4990744 | 45.39 | 575 | 16448 |
| SAMN02338944-GCF_000490515.1 | S091 | SAMN02338944 | GCF_000490515.1 | PC | Public | 1999 | India | NA | 96.9 | 99.56 | 1.49 | 99.9174 | 5099591 | 45.25 | 634 | 14759 |
| SAMN02338945-GCF_000493715.1 | S092 | SAMN02338945 | GCF_000493715.1 | PC | Public | 1996 | China | NA | 97.7 | 99.78 | 0.57 | 99.9925 | 5135145 | 45.07 | 369 | 28070 |
| SAMN02338946-GCF_000490485.1 | S093 | SAMN02338946 | GCF_000490485.1 | PrePC | Public | 1998 | Japan | Clinical | 93.6 | 99.59 | 0.74 | 99.8762 | 4920314 | 45.43 | 382 | 29016 |
| SAMN02338947-GCF_000490465.1 | S094 | SAMN02338947 | GCF_000490465.1 | PC | Public | 1996 | Thailand | NA | 97.7 | 99.98 | 0.22 | 99.9956 | 5029906 | 45.27 | 416 | 22106 |
| SAMN02338948-GCF_000490445.1 | S095 | SAMN02338948 | GCF_000490445.1 | PrePC | Public | 1996 | China | Clinical | 93.1 | 99.91 | 0.27 | 99.2621 | 5060177 | 45.31 | 407 | 25401 |
| SAMN02338949-GCF_000493225.1 | S096 | SAMN02338949 | GCF_000493225.1 | NonPC | Public | 1999 | South_Korea | NA | 91.7 | 99.82 | 1.71 | 98.3373 | 5135645 | 45.12 | 541 | 17831 |
| SAMN02338950-GCF_000490415.1 | S097 | SAMN02338950 | GCF_000490415.1 | NonPC | Public | 1992 | China | NA | 90.2 | 99.96 | 1.01 | 98.3757 | 4996423 | 45.34 | 439 | 23502 |
| SAMN02338951-GCF_000490395.1 | S098 | SAMN02338951 | GCF_000490395.1 | NonPC | Public | 1997 | USA | NA | 90.8 | 99.85 | 0.39 | 98.2301 | 5270002 | 45.21 | 574 | 17405 |
| SAMN02338952-GCF_000490375.1 | S100 | SAMN02338952 | GCF_000490375.1 | NonPC | Public | 1990 | USA | NA | 89.8 | 99.98 | 0.36 | 98.3288 | 5004126 | 45.17 | 439 | 21352 |
| SAMN02338953-GCF_000490355.1 | S101 | SAMN02338953 | GCF_000490355.1 | NonPC | Public | 1990 | USA | NA | 89.1 | 99.98 | 0.11 | 98.1646 | 4976676 | 45.28 | 478 | 19913 |
| SAMN02338954-GCF_000490335.1 | S104 | SAMN02338954 | GCF_000490335.1 | NonPC | Public | 1997 | USA | NA | 90.5 | 99.98 | 0.69 | 98.2698 | 5145846 | 45.19 | 627 | 14072 |
| SAMN02338955-GCF_000490315.1 | S105 | SAMN02338955 | GCF_000490315.1 | NonPC | Public | 1984 | Japan | NA | 90.5 | 99.09 | 2.69 | 98.434 | 5079200 | 45.11 | 662 | 14631 |
| SAMN02338956-GCF_000490295.1 | S106 | SAMN02338956 | GCF_000490295.1 | NonPC | Public | 1990 | Thailand | NA | 89.6 | 99.98 | 0.33 | 98.2799 | 5053204 | 45.3 | 548 | 15464 |
| SAMN02338957-GCF_000490275.1 | S108 | SAMN02338957 | GCF_000490275.1 | NonPC | Public | 1990 | Thailand | NA | 89.6 | 99.89 | 0.11 | 98.3783 | 5050272 | 45.3 | 592 | 15384 |
| SAMN02338958-GCF_000490255.1 | S109 | SAMN02338958 | GCF_000490255.1 | NonPC | Public | 1993 | China | NA | 89.7 | 99.73 | 0 | 98.3249 | 5020392 | 45.33 | 300 | 38985 |
| SAMN02338959-GCF_000490235.1 | S110 | SAMN02338959 | GCF_000490235.1 | NonPC | Public | 1999 | India | NA | 90.2 | 99.98 | 0.72 | 98.4207 | 5095381 | 45.23 | 595 | 14255 |
| SAMN02338960-GCF_000490215.1 | S111 | SAMN02338960 | GCF_000490215.1 | NonPC | Public | 1990 | Thailand | NA | 90.1 | 99.91 | 0.38 | 98.3972 | 5115103 | 45.2 | 586 | 15938 |
| SAMN02338961-GCF_000490195.1 | S112 | SAMN02338961 | GCF_000490195.1 | NonPC | Public | 1951 | Japan | NA | 90.1 | 99.98 | 0.42 | 98.4064 | 5003321 | 45.27 | 586 | 16061 |
| SAMN02338962-GCF_000490175.1 | S113 | SAMN02338962 | GCF_000490175.1 | NonPC | Public | 1990 | India | NA | 90 | 99.68 | 1.27 | 98.4773 | 5079169 | 45.26 | 643 | 15260 |
| SAMN02338963-GCF_000490155.1 | S114 | SAMN02338963 | GCF_000490155.1 | NonPC | Public | 1998 | India | NA | 90.2 | 99.98 | 1.31 | 98.3786 | 5180804 | 45.06 | 611 | 15194 |
| SAMN02338964-GCF_000490135.1 | S115 | SAMN02338964 | GCF_000490135.1 | NonPC | Public | 1991 | Thailand | NA | 90.5 | 99.98 | 0.27 | 98.3389 | 5108717 | 45.18 | 740 | 11577 |
| SAMN02338965-GCF_000490115.1 | S116 | SAMN02338965 | GCF_000490115.1 | NonPC | Public | NA | Thailand | NA | 92.5 | 99.98 | 0.73 | 98.3588 | 5054893 | 45.22 | 692 | 11871 |
| SAMN02338966-GCF_000490095.1 | S117 | SAMN02338966 | GCF_000490095.1 | NonPC | Public | 1984 | Japan | NA | 90.6 | 99.85 | 0.07 | 98.4054 | 5035586 | 45.3 | 677 | 12496 |
| SAMN02338967-GCF_000490075.1 | S118 | SAMN02338967 | GCF_000490075.1 | NonPC | Public | 1984 | Japan | NA | 91.7 | 99.89 | 0.72 | 98.4373 | 5110002 | 45.12 | 643 | 12725 |
| SAMN02338968-GCF_000490055.1 | S119 | SAMN02338968 | GCF_000490055.1 | NonPC | Public | 1999 | China | NA | 91.8 | 99.95 | 0.82 | 98.3764 | 5175377 | 45.05 | 642 | 13018 |
| SAMN02338969-GCF_000490035.1 | S120 | SAMN02338969 | GCF_000490035.1 | NonPC | Public | 1990 | Thailand | NA | 89.9 | 99.82 | 0.7 | 98.3779 | 5117413 | 45.26 | 682 | 13137 |
| SAMN02338970-GCF_000490015.1 | S121 | SAMN02338970 | GCF_000490015.1 | NonPC | Public | NA | Thailand | NA | 92.5 | 99.82 | 0.49 | 98.3738 | 5031191 | 45.25 | 652 | 12282 |
| SAMN02338971-GCF_000489995.1 | S122 | SAMN02338971 | GCF_000489995.1 | NonPC | Public | 1999 | India | NA | 92 | 99.89 | 0.57 | 98.3811 | 5058414 | 45.25 | 643 | 12750 |
| SAMN02338972-GCF_000489975.1 | S123 | SAMN02338972 | GCF_000489975.1 | NonPC | Public | 1984 | Japan | NA | 91.9 | 99.85 | 0.58 | 98.4414 | 5042126 | 45.26 | 656 | 12246 |
| SAMN02338973-GCF_000489955.1 | S124 | SAMN02338973 | GCF_000489955.1 | NonPC | Public | 1992 | China | NA | 92.2 | 99.71 | 0.15 | 98.3363 | 5122465 | 45.22 | 637 | 12697 |
| SAMN02338974-GCF_000489935.1 | S125 | SAMN02338974 | GCF_000489935.1 | NonPC | Public | 1997 | USA | NA | 90.9 | 99.98 | 0.15 | 98.4247 | 5067929 | 45.28 | 333 | 27219 |
| SAMN02338975-GCF_000489915.1 | S126 | SAMN02338975 | GCF_000489915.1 | PC | Public | NA | China | Clinical | 97.7 | 99.94 | 0.18 | 99.9941 | 5036556 | 45.27 | 338 | 27924 |
| SAMN02338976-GCF_000489895.1 | S128 | SAMN02338976 | GCF_000489895.1 | NonPC | Public | 1999 | India | NA | 90.8 | 99.98 | 0.4 | 98.4347 | 4934757 | 45.39 | 334 | 30162 |
| SAMN02338977-GCF_000489875.1 | S129 | SAMN02338977 | GCF_000489875.1 | NonPC | Public | 1984 | Japan | NA | 90 | 99.71 | 0.9 | 98.4087 | 4981119 | 45.38 | 480 | 23715 |
| SAMN02338978-GCF_000489855.1 | S130 | SAMN02338978 | GCF_000489855.1 | NonPC | Public | 2003 | NA | NA | 91.6 | 99.94 | 0.85 | 98.3728 | 5167017 | 45.27 | 364 | 26145 |
| SAMN02338979-GCF_000489835.1 | S131 | SAMN02338979 | GCF_000489835.1 | PC | Public | 2003 | NA | Clinical | 97.7 | 99.98 | 0.5 | 99.9854 | 5107430 | 45.11 | 355 | 26782 |
| SAMN02338980-GCF_000489815.1 | S132 | SAMN02338980 | GCF_000489815.1 | NonPC | Public | 2005 | China | NA | 92.4 | 99.89 | 0.15 | 98.4154 | 5087488 | 45.23 | 364 | 28517 |
| SAMN02338981-GCF_000489795.1 | S133 | SAMN02338981 | GCF_000489795.1 | PC | Public | 2005 | China | NA | 96.9 | 99.94 | 0.32 | 99.9835 | 5006605 | 45.29 | 375 | 22687 |
| SAMN02338982-GCF_000489775.1 | S134 | SAMN02338982 | GCF_000489775.1 | NonPC | Public | 2005 | China | NA | 91.6 | 99.94 | 0.7 | 98.4308 | 5162307 | 45.27 | 368 | 27182 |
| SAMN02338983-GCF_000489755.1 | S135 | SAMN02338983 | GCF_000489755.1 | PC | Public | 2003 | China | NA | 97.3 | 99.98 | 0.39 | 99.9805 | 5033236 | 45.28 | 353 | 25931 |
| SAMN02338984-GCF_000489735.1 | S136 | SAMN02338984 | GCF_000489735.1 | PC | Public | 2004 | China | NA | 97.5 | 99.85 | 0.19 | 99.9828 | 5036939 | 45.26 | 400 | 21577 |
| SAMN02338985-GCF_000489715.1 | S137 | SAMN02338985 | GCF_000489715.1 | PC | Public | 2005 | China | NA | 96.5 | 98.98 | 1 | 99.9872 | 4977105 | 45.37 | 588 | 15592 |

| | | | | | | | | | | | | | | | | |
|---|---|---|---|---|---|---|---|---|---|---|---|---|---|---|---|---|
| SAMN02338986-GCF_000489695.1 | S138 | SAMN02338986 | GCF_000489695.1 | PC | Public | 2007 | China | NA | 97.7 | 99.94 | 0.17 | 99.9928 | 5107186 | 45.11 | 416 | 22106 |
| SAMN02338987-GCF_000489675.1 | S139 | SAMN02338987 | GCF_000489675.1 | NonPC | Public | 2006 | China | NA | 90.3 | 99.98 | 0.39 | 98.4238 | 4985005 | 45.28 | 482 | 22603 |
| SAMN02338988-GCF_000489655.1 | S140 | SAMN02338988 | GCF_000489655.1 | NonPC | Public | 2006 | China | NA | 90.7 | 99.98 | 0.23 | 98.3976 | 5095143 | 45.21 | 443 | 27809 |
| SAMN02338989-GCF_000489635.1 | S141 | SAMN02338989 | GCF_000489635.1 | NonPC | Public | 2006 | China | NA | 89 | 99.89 | 0.64 | 98.3027 | 5074610 | 45.29 | 406 | 25799 |
| SAMN02338990-GCF_000489615.1 | S142 | SAMN02338990 | GCF_000489615.1 | NonPC | Public | 2006 | China | NA | 87.9 | 99.94 | 0.8 | 98.2706 | 5140032 | 45.04 | 480 | 21560 |
| SAMN02338991-GCF_000489595.1 | S143 | SAMN02338991 | GCF_000489595.1 | NonPC | Public | 2006 | China | NA | 89.5 | 99.94 | 0.14 | 98.4319 | 5041558 | 45.32 | 401 | 25903 |
| SAMN02338992-GCF_000489575.1 | S144 | SAMN02338992 | GCF_000489575.1 | NonPC | Public | 2006 | China | NA | 89.5 | 99.94 | 0.3 | 98.4251 | 5043659 | 45.32 | 408 | 27181 |
| SAMN02338993-GCF_000489555.1 | S145 | SAMN02338993 | GCF_000489555.1 | NonPC | Public | 2006 | China | NA | 90.2 | 99.94 | 0.42 | 98.3579 | 5152255 | 45.36 | 396 | 23804 |
| SAMN02338994-GCF_000489535.1 | S146 | SAMN02338994 | GCF_000489535.1 | NonPC | Public | 2006 | China | NA | 89.3 | 99.98 | 0.23 | 98.3455 | 4994195 | 45.3 | 435 | 24916 |
| SAMN02338995-GCF_000489515.1 | S147 | SAMN02338995 | GCF_000489515.1 | NonPC | Public | 2006 | China | NA | 89 | 99.71 | 1.2 | 98.3631 | 5089815 | 45.36 | 556 | 17214 |
| SAMN02338996-GCF_000489495.1 | S148 | SAMN02338996 | GCF_000489495.1 | NonPC | Public | 2006 | China | NA | 88.7 | 99.76 | 0.7 | 98.3675 | 4998866 | 45.28 | 571 | 16810 |
| SAMN02338997-GCF_000489475.1 | S150 | SAMN02338997 | GCF_000489475.1 | NonPC | Public | 2006 | China | NA | 89.4 | 99.91 | 1.38 | 98.4331 | 5037906 | 45.33 | 521 | 18544 |
| SAMN02338998-GCF_000489455.1 | S151 | SAMN02338998 | GCF_000489455.1 | NonPC | Public | 2006 | China | NA | 89.4 | 99.94 | 0.84 | 98.4137 | 5039338 | 45.33 | 518 | 18813 |
| SAMN02338999-GCF_000489435.1 | S152 | SAMN02338999 | GCF_000489435.1 | NonPC | Public | 2006 | China | NA | 90.6 | 99.89 | 0.41 | 98.3281 | 4989879 | 45.39 | 522 | 15084 |
| SAMN02339000-GCF_000489415.1 | S153 | SAMN02339000 | GCF_000489415.1 | NonPC | Public | 2006 | China | NA | 89.5 | 99.6 | 0.73 | 98.4618 | 5040674 | 45.33 | 528 | 18172 |
| SAMN02339001-GCF_000489395.1 | S154 | SAMN02339001 | GCF_000489395.1 | NonPC | Public | 2006 | China | NA | 89.2 | 99.71 | 1.55 | 98.4917 | 4986339 | 45.4 | 580 | 15518 |
| SAMN02339002-GCF_000489375.1 | S155 | SAMN02339002 | GCF_000489375.1 | NonPC | Public | 2006 | China | NA | 89.4 | 99.88 | 0.34 | 98.4346 | 5039238 | 45.33 | 511 | 19961 |
| SAMN02339003-GCF_000489355.1 | S156 | SAMN02339003 | GCF_000489355.1 | NonPC | Public | 2006 | China | NA | 89.4 | 99.94 | 0.27 | 98.4569 | 5040063 | 45.33 | 517 | 17919 |
| SAMN02339004-GCF_000489335.1 | S157 | SAMN02339004 | GCF_000489335.1 | NonPC | Public | 2006 | China | NA | 91.8 | 99.94 | 0.38 | 98.3829 | 5016174 | 45.27 | 539 | 17263 |
| SAMN02339005-GCF_000489315.1 | S158 | SAMN02339005 | GCF_000489315.1 | NonPC | Public | 2006 | China | NA | 89.5 | 99.98 | 0.14 | 98.4178 | 5041116 | 45.32 | 427 | 23944 |
| SAMN02339006-GCF_000489295.1 | S159 | SAMN02339006 | GCF_000489295.1 | NonPC | Public | 2006 | China | NA | 91.8 | 99.94 | 0.52 | 98.3864 | 5020516 | 45.27 | 430 | 24030 |
| SAMN02339007-GCF_000489275.1 | S160 | SAMN02339007 | GCF_000489275.1 | NonPC | Public | 2006 | China | NA | 88 | 99.4 | 0.5 | 98.334 | 4756268 | 45.47 | 448 | 17257 |
| SAMN02339008-GCF_000489255.1 | S161 | SAMN02339008 | GCF_000489255.1 | NonPC | Public | 2006 | China | NA | 89.5 | 99.94 | 0.27 | 98.3881 | 5041812 | 45.32 | 415 | 24446 |
| SAMN02339009-GCF_000489235.1 | S162 | SAMN02339009 | GCF_000489235.1 | NonPC | Public | 2007 | Thailand | NA | 90 | 99.94 | 0.26 | 98.3915 | 5023906 | 45.26 | 459 | 20676 |
| SAMN02339010-GCF_000489215.1 | S163 | SAMN02339010 | GCF_000489215.1 | NonPC | Public | 2007 | Malaysia | NA | 90.1 | 99.89 | 0.62 | 98.4254 | 5150062 | 45.34 | 435 | 23895 |
| SAMN02339011-GCF_000489195.1 | S164 | SAMN02339011 | GCF_000489195.1 | NonPC | Public | 2007 | USA | NA | 91.3 | 99.95 | 0.44 | 98.4859 | 4962474 | 45.33 | 469 | 20188 |
| SAMN02339012-GCF_000489175.1 | S165 | SAMN02339012 | GCF_000489175.1 | NonPC | Public | 2007 | China | NA | 87.5 | 99.98 | 0.6 | 98.3389 | 4984370 | 45.35 | 474 | 22119 |
| SAMN02339013-GCF_000489155.1 | S166 | SAMN02339013 | GCF_000489155.1 | NonPC | Public | 2007 | China | NA | 90 | 99.98 | 0.33 | 98.352 | 4983170 | 45.36 | 450 | 23056 |
| SAMN02339014-GCF_000489135.1 | S167 | SAMN02339014 | GCF_000489135.1 | NonPC | Public | 2007 | China | NA | 90.4 | 99.98 | 0.12 | 98.3158 | 5143354 | 45.28 | 426 | 21981 |
| SAMN02339015-GCF_000489115.1 | S168 | SAMN02339015 | GCF_000489115.1 | NonPC | Public | 2007 | China | NA | 89 | 99.98 | 0.03 | 98.3558 | 5009287 | 45.29 | 291 | 39979 |
| SAMN02339016-GCF_000489095.1 | S170 | SAMN02339016 | GCF_000489095.1 | NonPC | Public | 2007 | China | NA | 89.5 | 99.98 | 0 | 98.4484 | 5041715 | 45.32 | 310 | 35762 |
| SAMN02339017-GCF_000489075.1 | S171 | SAMN02339017 | GCF_000489075.1 | NonPC | Public | 2007 | China | NA | 89.3 | 99.85 | 0.3 | 98.3638 | 5124338 | 45.18 | 265 | 35949 |
| SAMN02339018-GCF_000489055.1 | S172 | SAMN02339018 | GCF_000489055.1 | NonPC | Public | 2007 | China | NA | 89.4 | 99.98 | 0.2 | 98.3871 | 5027540 | 45.35 | 279 | 37669 |
| SAMN02339019-GCF_000489035.1 | S173 | SAMN02339019 | GCF_000489035.1 | NonPC | Public | 2007 | China | NA | 89.4 | 99.98 | 0.37 | 98.3109 | 4921086 | 45.35 | 309 | 37764 |
| SAMN02339020-GCF_000489015.1 | S174 | SAMN02339020 | GCF_000489015.1 | NonPC | Public | 2007 | China | NA | 91 | 99.94 | 0.18 | 98.4444 | 4980648 | 45.33 | 319 | 37577 |
| SAMN02368223-GCF_001726265.1 | BCW_3100 | SAMN02368223 | GCF_001726265.1 | NonPC | Public | 2007 | USA | Env | 89.7 | 100 | 0.14 | 98.2716 | 5076064 | 45.25 | 69 | 402719 |
| SAMN02368224-GCF_001726195.1 | BCW_3101 | SAMN02368224 | GCF_001726195.1 | NonPC | Public | 2007 | USA | Env | 90.4 | 100 | 0.07 | 98.2855 | 5036171 | 45.29 | 91 | 342982 |
| SAMN02368225-GCF_001726185.1 | BCW_3102 | SAMN02368225 | GCF_001726185.1 | NonPC | Public | 2007 | USA | Env | 90.3 | 100 | 0.07 | 98.2532 | 5190611 | 45.24 | 86 | 368783 |
| SAMN02368226-GCF_001726245.1 | BCW_3104 | SAMN02368226 | GCF_001726245.1 | NonPC | Public | 2007 | USA | Env | 90.7 | 100 | 0.14 | 98.1679 | 5040822 | 45.21 | 82 | 321940 |
| SAMN02368227-GCF_001726255.1 | BCW_3105 | SAMN02368227 | GCF_001726255.1 | NonPC | Public | 2007 | USA | Env | 90.7 | 100 | 0.07 | 98.3198 | 5104427 | 45.21 | 101 | 179238 |
| SAMN02368228-GCF_001726345.1 | BCW_3106 | SAMN02368228 | GCF_001726345.1 | NonPC | Public | 2007 | USA | Env | 90.3 | 100 | 0.41 | 98.2059 | 5039638 | 45.19 | 76 | 284110 |
| SAMN02368229-GCF_001726275.1 | BCW_3107 | SAMN02368229 | GCF_001726275.1 | NonPC | Public | 2007 | USA | Env | 91.3 | 100 | 0.65 | 98.3818 | 4999863 | 45.29 | 63 | 400267 |
| SAMN02368230-GCF_001726355.1 | BCW_3108 | SAMN02368230 | GCF_001726355.1 | NonPC | Public | 2007 | USA | Env | 89.8 | 100 | 0.22 | 98.3317 | 5116384 | 45.17 | 56 | 477172 |
| SAMN02368231-GCF_001726335.1 | BCW_3109 | SAMN02368231 | GCF_001726335.1 | NonPC | Public | 2007 | USA | Env | 90.7 | 100 | 0.14 | 98.2115 | 5051740 | 45.2 | 85 | 313874 |
| SAMN02368232-GCF_001726325.1 | BCW_3110 | SAMN02368232 | GCF_001726325.1 | NonPC | Public | 2007 | USA | Env | 88.6 | 100 | 0 | 98.1246 | 4938203 | 45.38 | 76 | 295296 |
| SAMN02368233-GCF_001726405.1 | BCW_3111 | SAMN02368233 | GCF_001726405.1 | NonPC | Public | 2007 | USA | Env | 89.3 | 100 | 0 | 98.1024 | 5120255 | 45.24 | 89 | 294632 |
| SAMN02368234-GCF_001726415.1 | BCW_3112 | SAMN02368234 | GCF_001726415.1 | NonPC | Public | 2007 | USA | Env | 89.2 | 100 | 0 | 98.0703 | 5115080 | 45.24 | 93 | 294642 |
| SAMN02368235-GCF_001726445.1 | BCW_3113 | SAMN02368235 | GCF_001726445.1 | NonPC | Public | 2007 | USA | Env | 88.9 | 100 | 0 | 98.0686 | 5054005 | 45.31 | 86 | 186217 |

| | | | | | | | | | | | | | | | | |
|---|---|---|---|---|---|---|---|---|---|---|---|---|---|---|---|---|
| SAMN02368236-GCF_001726435.1 | BCW_3114 | SAMN02368236 | GCF_001726435.1 | NonPC | Public | 2007 | USA | Env | 91.2 | 100 | 0.27 | 98.2841 | 5033940 | 45.27 | 62 | 350943 |
| SAMN02368237-GCF_001726495.1 | BCW_3116 | SAMN02368237 | GCF_001726495.1 | NonPC | Public | 2007 | USA | Env | 88.9 | 100 | 0.14 | 98.1621 | 4964409 | 45.28 | 69 | 253663 |
| SAMN02368238-GCF_001726515.1 | BCW_3117 | SAMN02368238 | GCF_001726515.1 | NonPC | Public | 2007 | USA | Env | 90.6 | 100 | 0.14 | 98.1594 | 5043964 | 45.18 | 89 | 313873 |
| SAMN02368239-GCF_001726565.1 | BCW_3119 | SAMN02368239 | GCF_001726565.1 | NonPC | Public | 2007 | USA | Env | 91.4 | 100 | 0.49 | 98.3637 | 5178707 | 45.06 | 51 | 323835 |
| SAMN02368240-GCF_001726575.1 | BCW_3120 | SAMN02368240 | GCF_001726575.1 | NonPC | Public | 2007 | USA | Env | 91.5 | 100 | 0.49 | 98.3762 | 5178432 | 45.06 | 57 | 324616 |
| SAMN02368241-GCF_001726595.1 | BCW_3121 | SAMN02368241 | GCF_001726595.1 | NonPC | Public | 2007 | USA | Env | 91.5 | 100 | 0.49 | 98.4057 | 5177472 | 45.06 | 56 | 333784 |
| SAMN02368242-GCF_001726615.1 | BCW_3122 | SAMN02368242 | GCF_001726615.1 | NonPC | Public | 2007 | USA | Env | 91.5 | 100 | 0.49 | 98.3763 | 5177808 | 45.07 | 57 | 323969 |
| SAMN02368243-GCF_001726655.1 | BCW_3123 | SAMN02368243 | GCF_001726655.1 | NonPC | Public | 2007 | USA | Env | 91.4 | 100 | 0.49 | 98.3772 | 5176850 | 45.07 | 55 | 324103 |
| SAMN02368244-GCF_001726645.1 | BCW_3124 | SAMN02368244 | GCF_001726645.1 | NonPC | Public | 2007 | USA | Env | 91.3 | 100 | 0.27 | 98.2779 | 5092334 | 45.28 | 75 | 314409 |
| SAMN02368245-GCF_001726695.1 | BCW_3126 | SAMN02368245 | GCF_001726695.1 | NonPC | Public | 2007 | USA | Env | 90.9 | 100 | 0.08 | 98.2517 | 5252579 | 45.07 | 85 | 256849 |
| SAMN02368247-GCF_001726725.1 | BCW_3128 | SAMN02368247 | GCF_001726725.1 | NonPC | Public | 2007 | USA | Env | 89.5 | 100 | 0 | 98.0822 | 5165289 | 45.16 | 91 | 192245 |
| SAMN02368248-GCF_001726735.1 | BCW_3129 | SAMN02368248 | GCF_001726735.1 | NonPC | Public | 2007 | USA | Env | 91 | 100 | 0.41 | 98.3947 | 5114878 | 45.21 | 68 | 347208 |
| SAMN02368250-GCF_001726755.1 | BCW_3131 | SAMN02368250 | GCF_001726755.1 | NonPC | Public | 2007 | USA | Env | 88.9 | 100 | 0 | 98.1544 | 5056540 | 45.36 | 79 | 207247 |
| SAMN02368251-GCF_001726775.1 | BCW_3132 | SAMN02368251 | GCF_001726775.1 | NonPC | Public | 2007 | USA | Env | 90.6 | 100 | 0.27 | 98.3694 | 5033520 | 45.31 | 73 | 521828 |
| SAMN02368252-GCF_001726805.1 | BCW_3133 | SAMN02368252 | GCF_001726805.1 | NonPC | Public | 2007 | USA | Env | 91.5 | 100 | 0 | 98.334 | 5004954 | 45.32 | 56 | 313333 |
| SAMN02368253-GCF_001726815.1 | BCW_3134 | SAMN02368253 | GCF_001726815.1 | NonPC | Public | 2007 | USA | Env | 91.5 | 100 | 0 | 98.3601 | 5009581 | 45.32 | 57 | 313262 |
| SAMN02368254-GCF_001726845.1 | BCW_3135 | SAMN02368254 | GCF_001726845.1 | NonPC | Public | 2007 | USA | Env | 91.5 | 100 | 0 | 98.3276 | 5006232 | 45.32 | 49 | 443710 |
| SAMN02368255-GCF_001726885.1 | BCW_3136 | SAMN02368255 | GCF_001726885.1 | NonPC | Public | 2007 | USA | Env | 89.1 | 100 | 0.13 | 98.1995 | 4885078 | 45.41 | 83 | 192233 |
| SAMN02368256-GCF_001726855.1 | BCW_3137 | SAMN02368256 | GCF_001726855.1 | NonPC | Public | 2007 | USA | Env | 90.9 | 100 | 0.15 | 98.2692 | 5169004 | 45.23 | 71 | 408986 |
| SAMN02368257-GCF_001726895.1 | BCW_3138 | SAMN02368257 | GCF_001726895.1 | NonPC | Public | 2007 | USA | Env | 90.9 | 100 | 0.15 | 98.2606 | 5174835 | 45.23 | 60 | 409230 |
| SAMN02368259-GCF_001726915.1 | BCW_3140 | SAMN02368259 | GCF_001726915.1 | NonPC | Public | 2007 | Canada | Env | 91.9 | 100 | 0.05 | 98.3611 | 5108886 | 45.21 | 56 | 431216 |
| SAMN02368261-GCF_001726985.1 | BCW_3144 | SAMN02368261 | GCF_001726985.1 | NonPC | Public | 2007 | Canada | Env | 91.9 | 100 | 0.05 | 98.3645 | 5107343 | 45.21 | 55 | 349147 |
| SAMN02368263-GCF_001727025.1 | BCW_3147 | SAMN02368263 | GCF_001727025.1 | NonPC | Public | 2007 | Canada | Env | 91.9 | 100 | 0.05 | 98.3671 | 5112712 | 45.22 | 55 | 349026 |
| SAMN02368264-GCF_001727065.1 | BCW_3149 | SAMN02368264 | GCF_001727065.1 | NonPC | Public | 2007 | Canada | Env | 88.5 | 100 | 0.11 | 98.1383 | 4918007 | 45.34 | 75 | 234627 |
| SAMN02368265-GCF_001727045.1 | BCW_3150 | SAMN02368265 | GCF_001727045.1 | NonPC | Public | 2007 | USA | Env | 89.9 | 100 | 0 | 98.0308 | 5064921 | 45.23 | 110 | 163207 |
| SAMN02368266-GCF_001727105.1 | BCW_3151 | SAMN02368266 | GCF_001727105.1 | NonPC | Public | 2007 | USA | Env | 91.1 | 99.73 | 0.08 | 98.3536 | 5123058 | 45.26 | 66 | 244992 |
| SAMN02368267-GCF_001727115.1 | BCW_3152 | SAMN02368267 | GCF_001727115.1 | NonPC | Public | 2007 | USA | Env | 88.8 | 100 | 0 | 98.1272 | 4885698 | 45.42 | 78 | 227454 |
| SAMN02368268-GCF_001975475.1 | BCW_3153 | SAMN02368268 | GCF_001975475.1 | NonPC | Public | 2007 | USA | Env | 91.9 | 100 | 0 | 98.3331 | 5043245 | 45.26 | 67 | 211278 |
| SAMN02368269-GCF_001727125.1 | BCW_3154 | SAMN02368269 | GCF_001727125.1 | NonPC | Public | 2007 | USA | Env | 91.3 | 100 | 0 | 98.246 | 5126807 | 45.18 | 89 | 313100 |
| SAMN02368270-GCF_001727155.1 | BCW_3155 | SAMN02368270 | GCF_001727155.1 | NonPC | Public | 2007 | USA | Env | 92.1 | 100 | 0 | 98.2903 | 5050663 | 45.36 | 94 | 324832 |
| SAMN02368271-GCF_001727185.1 | BCW_3156 | SAMN02368271 | GCF_001727185.1 | NonPC | Public | 2007 | USA | Env | 89.6 | 100 | 0 | 98.0973 | 5083259 | 45.25 | 89 | 168542 |
| SAMN02368272-GCF_001727195.1 | BCW_3157 | SAMN02368272 | GCF_001727195.1 | NonPC | Public | 2007 | USA | Env | 89.6 | 100 | 0 | 98.101 | 5095963 | 45.25 | 91 | 141391 |
| SAMN02368273-GCF_001727205.1 | BCW_3158 | SAMN02368273 | GCF_001727205.1 | NonPC | Public | 2007 | USA | Env | 89.6 | 100 | 0 | 98.0879 | 5092215 | 45.24 | 96 | 160436 |
| SAMN02368274-GCF_001727265.1 | BCW_3159 | SAMN02368274 | GCF_001727265.1 | NonPC | Public | 2007 | USA | Env | 88.4 | 100 | 0.3 | 98.1552 | 4929698 | 45.38 | 97 | 314716 |
| SAMN02368275-GCF_001727295.1 | BCW_3161 | SAMN02368275 | GCF_001727295.1 | NonPC | Public | 2007 | USA | Env | 88.6 | 100 | 0 | 98.0738 | 5062642 | 45.24 | 116 | 160364 |
| SAMN02368276-GCF_001727305.1 | BCW_3163 | SAMN02368276 | GCF_001727305.1 | NonPC | Public | 2007 | USA | Env | 89.7 | 100 | 0 | 98.0831 | 5053606 | 45.25 | 105 | 198940 |
| SAMN02368277-GCF_001727345.1 | BCW_3164 | SAMN02368277 | GCF_001727345.1 | NonPC | Public | 2007 | USA | Env | 89.7 | 100 | 0 | 98.1047 | 5050786 | 45.25 | 103 | 191682 |
| SAMN02368278-GCF_001727365.1 | BCW_3166 | SAMN02368278 | GCF_001727365.1 | NonPC | Public | 2006 | USA | Clinical | 91.5 | 100 | 0.14 | 98.3387 | 5138627 | 45.18 | 102 | 190507 |
| SAMN02368279-GCF_001727725.1 | BCW_3167 | SAMN02368279 | GCF_001727725.1 | NonPC | Public | 2006 | USA | Clinical | 91.5 | 100 | 0 | 98.405 | 5062918 | 45.29 | 60 | 361205 |
| SAMN02368280-GCF_001727735.1 | BCW_3168 | SAMN02368280 | GCF_001727735.1 | NonPC | Public | 2006 | USA | Clinical | 91 | 100 | 0.22 | 98.2739 | 5020238 | 45.27 | 83 | 314740 |
| SAMN02368281-GCF_001727745.1 | BCW_3169 | SAMN02368281 | GCF_001727745.1 | NonPC | Public | 2006 | USA | Clinical | 91.9 | 100 | 0.22 | 98.2642 | 5317980 | 45.05 | 101 | 202283 |
| SAMN02368282-GCF_001727775.1 | BCW_3170 | SAMN02368282 | GCF_001727775.1 | NonPC | Public | 2006 | USA | Clinical | 88.9 | 100 | 0 | 98.1227 | 5000115 | 45.38 | 76 | 207438 |
| SAMN02368283-GCF_001727845.1 | BCW_3174 | SAMN02368283 | GCF_001727845.1 | NonPC | Public | 2006 | USA | Clinical | 89.8 | 100 | 0.22 | 98.3291 | 5068984 | 45.27 | 70 | 248530 |
| SAMN02368284-GCF_001727885.1 | BCW_3176 | SAMN02368284 | GCF_001727885.1 | NonPC | Public | 2006 | USA | Clinical | 91.3 | 100 | 0 | 98.3433 | 5063895 | 45.26 | 68 | 548241 |
| SAMN02368286-GCF_001727925.1 | BCW_3178 | SAMN02368286 | GCF_001727925.1 | NonPC | Public | 2006 | USA | Clinical | 89.8 | 100 | 0.14 | 98.3485 | 5094315 | 45.32 | 80 | 392296 |
| SAMN02368287-GCF_001727915.1 | BCW_3179 | SAMN02368287 | GCF_001727915.1 | NonPC | Public | 2006 | USA | Clinical | 89.7 | 99.95 | 0.19 | 98.2483 | 5273779 | 45.12 | 89 | 355269 |
| SAMN02368288-GCF_001727975.1 | BCW_3180 | SAMN02368288 | GCF_001727975.1 | NonPC | Public | 2006 | USA | Clinical | 89.6 | 99.95 | 0.19 | 98.2612 | 5267172 | 45.13 | 85 | 355176 |
| SAMN02368289-GCF_001728005.1 | BCW_3182 | SAMN02368289 | GCF_001728005.1 | PC | Public | 2007 | USA | Clinical | 98 | 100 | 0 | 99.987 | 5145965 | 45.11 | 79 | 312581 |

| | | | | | | | | | | | | | | | | |
|---|---|---|---|---|---|---|---|---|---|---|---|---|---|---|---|---|
| SAMN02368290-GCF_001727995.1 | BCW_3183 | SAMN02368290 | GCF_001727995.1 | NonPC | Public | 2006 | USA | Clinical | 89.8 | 100 | 0 | 98.2503 | 5184421 | 45.18 | 84 | 322967 |
| SAMN02368291-GCF_001728045.1 | BCW_3184 | SAMN02368291 | GCF_001728045.1 | NonPC | Public | 2007 | USA | Clinical | 89.2 | 100 | 0.34 | 98.3247 | 5031868 | 45.3 | 82 | 293993 |
| SAMN02368292-GCF_001728065.1 | BCW_3185 | SAMN02368292 | GCF_001728065.1 | NonPC | Public | 2007 | USA | Clinical | 89.2 | 100 | 0.34 | 98.3303 | 5031530 | 45.3 | 83 | 302174 |
| SAMN02368293-GCF_001728085.1 | BCW_3186 | SAMN02368293 | GCF_001728085.1 | NonPC | Public | 2006 | USA | Clinical | 91 | 100 | 0 | 98.4481 | 4950924 | 45.38 | 59 | 322414 |
| SAMN02368294-GCF_001728095.1 | BCW_3187 | SAMN02368294 | GCF_001728095.1 | NonPC | Public | 2007 | USA | Clinical | 90.8 | 100 | 0.32 | 98.3419 | 5170313 | 45.25 | 75 | 238239 |
| SAMN02368295-GCF_001728125.1 | BCW_3188 | SAMN02368295 | GCF_001728125.1 | NonPC | Public | 2007 | USA | Clinical | 89 | 100 | 0 | 98.1024 | 4916386 | 45.38 | 96 | 177698 |
| SAMN02368297-GCF_001728135.1 | BCW_3190 | SAMN02368297 | GCF_001728135.1 | NonPC | Public | 2006 | USA | Clinical | 91.7 | 100 | 0 | 98.4112 | 5115199 | 45.28 | 97 | 438284 |
| SAMN02368298-GCF_001728155.1 | BCW_3191 | SAMN02368298 | GCF_001728155.1 | PC | Public | 2006 | USA | Clinical | 97.7 | 100 | 0.05 | 99.9246 | 5103276 | 45.23 | 83 | 242331 |
| SAMN02368299-GCF_001728175.1 | BCW_3192 | SAMN02368299 | GCF_001728175.1 | PC | Public | 2006 | USA | Clinical | 97.7 | 100 | 0.05 | 99.9175 | 5102218 | 45.23 | 78 | 259047 |
| SAMN02368300-GCF_001728205.1 | BCW_3193 | SAMN02368300 | GCF_001728205.1 | PC | Public | 2007 | USA | Clinical | 98.2 | 100 | 0 | 99.9874 | 5109310 | 45.26 | 66 | 301866 |
| SAMN02368301-GCF_001728215.1 | BCW_3194 | SAMN02368301 | GCF_001728215.1 | NonPC | Public | 2007 | USA | Clinical | 90.2 | 100 | 0.05 | 98.3723 | 4950427 | 45.38 | 54 | 467008 |
| SAMN02368302-GCF_001728235.1 | BCW_3195 | SAMN02368302 | GCF_001728235.1 | NonPC | Public | 2007 | USA | Clinical | 90.1 | 100 | 0.05 | 98.3716 | 4940595 | 45.39 | 82 | 466965 |
| SAMN02368303-GCF_001728255.1 | BCW_3196 | SAMN02368303 | GCF_001728255.1 | NonPC | Public | 2007 | USA | Clinical | 89.4 | 100 | 0 | 98.1144 | 5014009 | 45.31 | 96 | 150907 |
| SAMN02368304-GCF_001728275.1 | BCW_3197 | SAMN02368304 | GCF_001728275.1 | NonPC | Public | 2007 | USA | Clinical | 88.9 | 100 | 0.06 | 98.1299 | 5072545 | 45.15 | 88 | 192573 |
| SAMN02368305-GCF_001728295.1 | BCW_3198 | SAMN02368305 | GCF_001728295.1 | NonPC | Public | 2007 | USA | Clinical | 91.8 | 100 | 0 | 98.3463 | 5118772 | 45.28 | 87 | 316428 |
| SAMN02368306-GCF_001728325.1 | BCW_3199 | SAMN02368306 | GCF_001728325.1 | NonPC | Public | 2007 | USA | Clinical | 91 | 100 | 0.35 | 98.2349 | 5129854 | 45.16 | 105 | 261043 |
| SAMN02368307-GCF_001728335.1 | BCW_3200 | SAMN02368307 | GCF_001728335.1 | NonPC | Public | 2007 | USA | Clinical | 89.8 | 100 | 0.04 | 98.2638 | 5154560 | 45.19 | 93 | 339585 |
| SAMN02368308-GCF_001728345.1 | BCW_3201 | SAMN02368308 | GCF_001728345.1 | NonPC | Public | NA | USA | Clinical | 90.6 | 100 | 0 | 98.3836 | 5154314 | 45.2 | 74 | 215075 |
| SAMN02368309-GCF_001728405.1 | BCW_3202 | SAMN02368309 | GCF_001728405.1 | NonPC | Public | 2007 | USA | Clinical | 91.3 | 100 | 0 | 98.3301 | 5056257 | 45.28 | 76 | 548384 |
| SAMN02368310-GCF_001727655.1 | BCW_3203 | SAMN02368310 | GCF_001727655.1 | NonPC | Public | NA | USA | Clinical | 90.7 | 100 | 0 | 98.3335 | 5166391 | 45.21 | 74 | 245528 |
| SAMN02368311-GCF_001727625.1 | BCW_3204 | SAMN02368311 | GCF_001727625.1 | NonPC | Public | 2007 | USA | Clinical | 91.4 | 100 | 0 | 98.3364 | 5071813 | 45.28 | 75 | 536709 |
| SAMN02368312-GCF_001727575.1 | BCW_3205 | SAMN02368312 | GCF_001727575.1 | NonPC | Public | 2007 | USA | Clinical | 91.3 | 100 | 0 | 98.3598 | 5064405 | 45.28 | 75 | 548238 |
| SAMN02368313-GCF_001727545.1 | BCW_3206 | SAMN02368313 | GCF_001727545.1 | NonPC | Public | 2007 | USA | Clinical | 87.8 | 99.18 | 0.33 | 98.2211 | 4954280 | 45.3 | 50 | 385892 |
| SAMN02368314-GCF_001727505.1 | BCW_3207 | SAMN02368314 | GCF_001727505.1 | NonPC | Public | 2007 | USA | Clinical | 90.5 | 100 | 0.14 | 98.1805 | 5033501 | 45.2 | 78 | 314904 |
| SAMN02368315-GCF_001727485.1 | BCW_3208 | SAMN02368315 | GCF_001727485.1 | NonPC | Public | 2007 | USA | Clinical | 91.4 | 100 | 0 | 98.3485 | 5079373 | 45.27 | 92 | 315048 |
| SAMN02368316-GCF_001727475.1 | BCW_3209 | SAMN02368316 | GCF_001727475.1 | NonPC | Public | NA | USA | Clinical | 89.8 | 100 | 0 | 98.276 | 5214531 | 45.2 | 79 | 322095 |
| SAMN02368317-GCF_001727415.1 | BCW_3210 | SAMN02368317 | GCF_001727415.1 | NonPC | Public | NA | USA | Clinical | 89.8 | 100 | 0 | 98.3009 | 5200325 | 45.22 | 60 | 322199 |
| SAMN02368318-GCF_001727395.1 | BCW_3211 | SAMN02368318 | GCF_001727395.1 | NonPC | Public | 2007 | USA | Clinical | 89.8 | 100 | 0 | 98.0623 | 5257378 | 45.16 | 246 | 130569 |
| SAMN02368319-GCF_001727405.1 | BCW_3212 | SAMN02368319 | GCF_001727405.1 | NonPC | Public | 2007 | USA | Clinical | 89.7 | 100 | 0 | 98.1168 | 5043160 | 45.34 | 121 | 139910 |
| SAMN02368320-GCF_001727665.1 | BCW_3213 | SAMN02368320 | GCF_001727665.1 | NonPC | Public | NA | USA | Clinical | 91.3 | 100 | 0 | 98.3485 | 5075163 | 45.27 | 75 | 322231 |
| SAMN02368321-GCF_001728425.1 | BCW_3215 | SAMN02368321 | GCF_001728425.1 | NonPC | Public | 2007 | USA | Clinical | 90.9 | 100 | 0.46 | 98.3537 | 5059053 | 45.25 | 70 | 539086 |
| SAMN02368322-GCF_001728465.1 | BCW_3216 | SAMN02368322 | GCF_001728465.1 | NonPC | Public | 2007 | USA | Clinical | 91.5 | 100 | 0 | 98.2905 | 5134599 | 45.26 | 70 | 548356 |
| SAMN02368323-GCF_001728485.1 | BCW_3217 | SAMN02368323 | GCF_001728485.1 | NonPC | Public | 2007 | USA | Clinical | 91.3 | 100 | 0 | 98.3233 | 5114185 | 45.26 | 72 | 548245 |
| SAMN02368324-GCF_001727585.1 | BCW_3220 | SAMN02368324 | GCF_001727585.1 | NonPC | Public | 2007 | USA | Clinical | 91.3 | 100 | 0 | 98.3628 | 5064302 | 45.28 | 72 | 536614 |
| SAMN02368325-GCF_001727385.1 | BCW_3221 | SAMN02368325 | GCF_001727385.1 | NonPC | Public | 2007 | USA | Clinical | 91.3 | 100 | 0 | 98.3613 | 5062191 | 45.29 | 86 | 525700 |
| SAMN02368326-GCF_001727465.1 | BCW_3222 | SAMN02368326 | GCF_001727465.1 | NonPC | Public | 2007 | USA | Clinical | 90.7 | 100 | 0 | 98.3667 | 5157372 | 45.2 | 71 | 215075 |
| SAMN02368327-GCA_015819965.1 | BCW_3223 | SAMN02368327 | GCA_015819965.1 | NonPC | Public | NA | USA | Clinical | 91.5 | 100 | 0 | 98.3177 | 5086116 | 45.29 | 39 | 522047 |
| SAMN02368328-GCF_001727565.1 | BCW_3224 | SAMN02368328 | GCF_001727565.1 | NonPC | Public | 2007 | USA | Clinical | 91.3 | 100 | 0 | 98.368 | 5055780 | 45.28 | 87 | 548245 |
| SAMN02368329-GCF_001727705.1 | BCW_3225 | SAMN02368329 | GCF_001727705.1 | NonPC | Public | 2007 | USA | Clinical | 90.7 | 100 | 0 | 98.4126 | 5138941 | 45.21 | 76 | 313186 |
| SAMN02368330-GCF_001728495.1 | BCW_3226 | SAMN02368330 | GCF_001728495.1 | NonPC | Public | 2007 | USA | Clinical | 92.2 | 100 | 0.05 | 98.3286 | 5019487 | 45.25 | 69 | 539098 |
| SAMN02368331-GCF_001728545.1 | BCW_3227 | SAMN02368331 | GCF_001728545.1 | NonPC | Public | NA | USA | Clinical | 91.3 | 100 | 0 | 98.3573 | 5068150 | 45.27 | 75 | 548195 |
| SAMN02368332-GCF_001728505.1 | BCW_3228 | SAMN02368332 | GCF_001728505.1 | NonPC | Public | 2007 | USA | Clinical | 91.3 | 100 | 0 | 98.3346 | 5066595 | 45.29 | 69 | 536921 |
| SAMN02368333-GCF_001728585.1 | BCW_3230 | SAMN02368333 | GCF_001728585.1 | NonPC | Public | 2007 | USA | Clinical | 91.4 | 100 | 0 | 98.3399 | 5068146 | 45.27 | 76 | 322224 |
| SAMN02368334-GCF_001728625.1 | BCW_3231 | SAMN02368334 | GCF_001728625.1 | PC | Public | 2007 | USA | Clinical | 97.3 | 100 | 0 | 99.9247 | 5090244 | 45.27 | 68 | 275983 |
| SAMN02368335-GCF_001728575.1 | BCW_3232 | SAMN02368335 | GCF_001728575.1 | NonPC | Public | NA | USA | Clinical | 89.1 | 100 | 0 | 98.0472 | 5056227 | 45.25 | 114 | 191195 |
| SAMN02368336-GCF_001728645.1 | BCW_3233 | SAMN02368336 | GCF_001728645.1 | NonPC | Public | 2007 | USA | Clinical | 89.8 | 100 | 0.04 | 98.2664 | 5161896 | 45.19 | 66 | 339585 |
| SAMN02368338-GCA_015819865.1 | BCW_3237 | SAMN02368338 | GCA_015819865.1 | NonPC | Public | 2007 | USA | Clinical | 90.8 | 100 | 0 | 98.3709 | 5254646 | 45.29 | 7 | 3378122 |
| SAMN02368339-GCF_001728695.1 | BCW_3238 | SAMN02368339 | GCF_001728695.1 | NonPC | Public | 2007 | USA | Clinical | 91.3 | 100 | 0 | 98.3171 | 5064665 | 45.26 | 78 | 548240 |

| | | | | | | | | | | | | | | | | |
|---|---|---|---|---|---|---|---|---|---|---|---|---|---|---|---|---|
| SAMN02368340-GCF_001728725.1 | BCW_3239 | SAMN02368340 | GCF_001728725.1 | NonPC | Public | 2007 | USA | Clinical | 90.5 | 100 | 0.05 | 98.2416 | 5057055 | 45.34 | 69 | 517784 |
| SAMN02435812-GCF_000500105.2 | 10296 | SAMN02435812 | GCF_000500105.2 | NonPC | Public | 1997 | USA | Clinical | 91.6 | 98.54 | 0 | 98.2954 | 5116072 | 45.27 | 33 | 730466 |
| SAMN02436183-GCF_000182465.1 | ACKB | SAMN02436183 | GCF_000182465.1 | PC | Public | 2005 | India | NA | 96.4 | 94.17 | 0 | 99.9862 | 5028244 | 45.35 | 164 | 62978 |
| SAMN02436220-GCF_000182345.1 | ACFM | SAMN02436220 | GCF_000182345.1 | PC | Public | 1996 | Peru | NA | 97.2 | 95.47 | 0.61 | 99.9886 | 5037488 | 45.36 | 149 | 81497 |
| SAMN02436265-GCF_000182385.1 | ACFO | SAMN02436265 | GCF_000182385.1 | PC | Public | 1998 | Bangladesh | NA | 98.5 | 99.17 | 0 | 99.9316 | 5199902 | 45.32 | 54 | 346246 |
| SAMN02469946-GCF_000510605.2 | 970107 | SAMN02469946 | GCF_000510605.2 | NonPC | Public | 1997 | USA | Env | 91.2 | 100 | 0 | 98.3467 | 5102571 | 45.3 | 85 | 139797 |
| SAMN02469951-GCF_000347495.1 | AOCL | SAMN02469951 | GCF_000347495.1 | NonPC | Public | 2008 | Malaysia | Env | 89.8 | 99.98 | 0.82 | 98.294 | 5178715 | 45.31 | 60 | 262059 |
| SAMN02470919-GCF_000315135.1 | AMRZ | SAMN02470919 | GCF_000315135.1 | NonPC | Public | 2009 | South_Korea | Env | 90.7 | 100 | 0.11 | 98.2677 | 5241845 | 45.23 | 60 | 237357 |
| SAMN02471128-GCF_000500505.1 | AXNJ | SAMN02471128 | GCF_000500505.1 | PC | Public | 2007 | China | Clinical | 98.3 | 100 | 0 | 99.9929 | 5114414 | 45.26 | 66 | 232863 |
| SAMN02471129-GCA_000500385.1 | VIP4-0445 | SAMN02471129 | GCA_000500385.1 | PC | Public | 2008 | China | Clinical | 95 | 94.55 | 1.51 | 99.9087 | 4936248 | 45.28 | 827 | 9963 |
| SAMN02471130-GCF_000500425.1 | AXNN | SAMN02471130 | GCF_000500425.1 | NonPC | Public | 2008 | China | Clinical | 92.9 | 100 | 0.18 | 98.3659 | 5056047 | 45.29 | 58 | 375030 |
| SAMN02471131-GCF_000500525.1 | AXNQ | SAMN02471131 | GCF_000500525.1 | NonPC | Public | 2006 | China | Env | 90.9 | 100 | 0.34 | 98.3892 | 5174299 | 45.2 | 66 | 211270 |
| SAMN02471132-GCF_000500365.1 | AXNK | SAMN02471132 | GCF_000500365.1 | PC | Public | 2008 | China | Clinical | 98.3 | 100 | 0 | 99.9896 | 5131692 | 45.26 | 69 | 188126 |
| SAMN02471133-GCF_000500405.1 | AXNM | SAMN02471133 | GCF_000500405.1 | PC | Public | 2008 | China | Clinical | 98.2 | 100 | 0 | 99.9926 | 5088510 | 45.27 | 68 | 235555 |
| SAMN02471134-GCF_000500485.1 | AXNR | SAMN02471134 | GCF_000500485.1 | NonPC | Public | 2008 | China | Env | 90.9 | 100 | 0.08 | 98.2663 | 5264927 | 45.2 | 85 | 180240 |
| SAMN02471136-GCA_000500465.1 | VIP4-0443 | SAMN02471136 | GCA_000500465.1 | NonPC | Public | 2008 | China | Env | 84.6 | 95.33 | 1.8 | 98.1635 | 4842411 | 45.34 | 1076 | 7337 |
| SAMN02471137-GCF_000500545.1 | AXNS | SAMN02471137 | GCF_000500545.1 | NonPC | Public | 2008 | China | Env | 91.1 | 100 | 0.15 | 98.3568 | 5353554 | 45.17 | 74 | 305898 |
| SAMN02471143-GCA_000388025.1 | v110 | SAMN02471143 | GCA_000388025.1 | NonPC | Public | 2010 | China | Clinical | 89.7 | 96.86 | 0.6 | 98.1751 | 5481570 | 45.2 | 85 | 323637 |
| SAMN02584906-GCA_000523375.1 | M0605 | SAMN02584906 | GCA_000523375.1 | NonPC | Public | 2013 | Mexico | Env | 90.8 | 98.39 | 0.34 | 98.1185 | 5424137 | 45.24 | 96 | 121988 |
| SAMN02597366-GCF_000519405.2 | 3256 | SAMN02597366 | GCF_000519405.2 | NonPC | Public | 2007 | USA | Clinical | 91.4 | 99.61 | 0 | 98.286 | 5082699 | 45.29 | 40 | 719354 |
| SAMN02597367-GCF_000519385.2 | 50 | SAMN02597367 | GCF_000519385.2 | NonPC | Public | 2006 | USA | Clinical | 90.2 | 100 | 0.14 | 98.198 | 5052413 | 45.17 | 56 | 425429 |
| SAMN02597368-GCF_000519365.2 | 605 | SAMN02597368 | GCF_000519365.2 | PC | Public | 2006 | USA | Env | 98.3 | 100 | 0 | 99.9756 | 5141263 | 45.27 | 66 | 317752 |
| SAMN02597369-GCF_000516875.2 | AZKN | SAMN02597369 | GCF_000516875.2 | PC | Public | 2004 | Mozambique | Clinical | 98.3 | 100 | 0 | 99.9846 | 5180880 | 45.29 | 50 | 542793 |
| SAMN02604302-GCF_000328405.1 | BB22OP | SAMN02604302 | GCF_000328405.1 | NonPC | Public | 1980 | Bangladesh | NA | 93.3 | 100 | 0.19 | 98.3875 | 5103524 | 45.36 | 2 | 3297305 |
| SAMN02641511-GCF_000525005.2 | EKP-026 | SAMN02641511 | GCF_000525005.2 | PC | Public | 2008 | Bangladesh | Env | 98.4 | 99.48 | 0 | 99.9862 | 5101635 | 45.29 | 64 | 289829 |
| SAMN02666028-GCA_000662375.1 | VP49 | SAMN02666028 | GCA_000662375.1 | NonPC | Public | 2008 | India | Env | 90.6 | 95.28 | 0.58 | 98.2489 | 5047822 | 45.28 | 137 | 68277 |
| SAMN02712350-GCF_000958575.1 | K1461 | SAMN02712350 | GCF_000958575.1 | NonPC | Public | 2004 | USA | Clinical | 91.5 | 99.63 | 0.04 | 98.3102 | 5177972 | 45.33 | 65 | 303097 |
| SAMN02712351-GCF_000958585.1 | K1275 | SAMN02712351 | GCF_000958585.1 | NonPC | Public | 2004 | USA | Clinical | 92.3 | 100 | 0.07 | 98.3527 | 5114306 | 45.36 | 63 | 301883 |
| SAMN02712352-GCF_000958565.1 | SG176 | SAMN02712352 | GCF_000958565.1 | NonPC | Public | 2006 | USA | NA | 89.5 | 99.66 | 0.09 | 98.3318 | 4952407 | 45.38 | 48 | 394204 |
| SAMN02712353-GCF_000958655.1 | J-C2-34 | SAMN02712353 | GCF_000958655.1 | NonPC | Public | 1998 | USA | NA | 89.1 | 99.7 | 0.09 | 98.0437 | 5150449 | 45.49 | 91 | 268853 |
| SAMN02712354-GCF_000958595.1 | AF91 | SAMN02712354 | GCF_000958595.1 | PrePC | Public | 2006 | USA | Env | 94.5 | 99.46 | 0.42 | 99.7974 | 5176263 | 45.33 | 111 | 269871 |
| SAMN02712355-GCF_000958645.1 | 22702 | SAMN02712355 | GCF_000958645.1 | NonPC | Public | 1998 | USA | NA | 88.9 | 100 | 0 | 98.0663 | 4955222 | 45.45 | 43 | 454662 |
| SAMN02741348-GCF_000707045.2 | CFSAN001611 | SAMN02741348 | GCF_000707045.2 | NonPC | Public | 1997 | USA | Env | 91.6 | 100 | 0 | 98.2963 | 5194181 | 45.19 | 48 | 269754 |
| SAMN02741349-GCF_000706825.2 | CFSAN001612 | SAMN02741349 | GCF_000706825.2 | NonPC | Public | 1990 | USA | Clinical | 91.6 | 100 | 0 | 98.2883 | 5097276 | 45.27 | 33 | 482345 |
| SAMN02741350-GCF_000707625.2 | CFSAN001613 | SAMN02741350 | GCF_000707625.2 | NonPC | Public | 1997 | USA | Clinical | 91.5 | 100 | 0 | 98.2756 | 5103112 | 45.27 | 44 | 411598 |
| SAMN02741351-GCF_000707885.2 | CFSAN001614 | SAMN02741351 | GCF_000707885.2 | NonPC | Public | 2004 | USA | Env | 91.6 | 100 | 0 | 98.296 | 5235701 | 45.16 | 60 | 461837 |
| SAMN02741352-GCF_002072885.1 | CFSAN001615 | SAMN02741352 | GCF_002072885.1 | NonPC | Public | 1990 | USA | Env | 91.5 | 100 | 0 | 98.2993 | 5093169 | 45.27 | 44 | 482421 |
| SAMN02741353-GCF_002072845.1 | CFSAN001616 | SAMN02741353 | GCF_002072845.1 | NonPC | Public | 1982 | USA | Env | 91.5 | 100 | 0 | 98.272 | 5136539 | 45.24 | 101 | 132439 |
| SAMN02741354-GCF_000707245.2 | CFSAN001617 | SAMN02741354 | GCF_000707245.2 | NonPC | Public | 1997 | USA | Clinical | 91.1 | 100 | 0.05 | 98.2616 | 5200027 | 45.19 | 40 | 386259 |
| SAMN02741355-GCF_000707525.2 | CFSAN001618 | SAMN02741355 | GCF_000707525.2 | NonPC | Public | 1990 | USA | Clinical | 91.5 | 100 | 0 | 98.2981 | 5094093 | 45.27 | 37 | 727942 |
| SAMN02741356-GCF_000707545.2 | CFSAN001619 | SAMN02741356 | GCF_000707545.2 | NonPC | Public | 1988 | USA | Env | 91.5 | 100 | 0 | 98.2984 | 5093394 | 45.27 | 38 | 447225 |
| SAMN02741357-GCF_000707565.2 | CFSAN001620 | SAMN02741357 | GCF_000707565.2 | NonPC | Public | 1998 | USA | Env | 91.5 | 100 | 0 | 98.2761 | 5094276 | 45.27 | 50 | 315430 |
| SAMN02741358-GCF_000707225.2 | CFSAN006129 | SAMN02741358 | GCF_000707225.2 | NonPC | Public | 2012 | USA | Clinical | 91.4 | 100 | 0.07 | 98.3019 | 5088743 | 45.27 | 84 | 218641 |
| SAMN02741359-GCF_000706925.1 | CFSAN006130 | SAMN02741359 | GCF_000706925.1 | NonPC | Public | 2013 | USA | Clinical | 91.4 | 100 | 0.27 | 98.3523 | 5096750 | 45.27 | 56 | 418201 |
| SAMN02741360-GCF_000707325.2 | CFSAN006131 | SAMN02741360 | GCF_000707325.2 | NonPC | Public | 2013 | USA | Clinical | 91.4 | 100 | 0 | 98.3066 | 5089845 | 45.27 | 60 | 226574 |
| SAMN02741361-GCF_000707845.1 | CFSAN006132 | SAMN02741361 | GCF_000707845.1 | NonPC | Public | 2013 | USA | Clinical | 91.4 | 99.86 | 0 | 98.3046 | 5091393 | 45.27 | 63 | 277454 |
| SAMN02741362-GCF_000707865.2 | CFSAN006133 | SAMN02741362 | GCF_000707865.2 | NonPC | Public | 2013 | USA | Clinical | 91.4 | 100 | 0 | 98.3223 | 5090866 | 45.27 | 52 | 344873 |
| SAMN02741363-GCF_000706845.2 | CFSAN006134 | SAMN02741363 | GCF_000706845.2 | NonPC | Public | 2013 | USA | Clinical | 91.4 | 100 | 0 | 98.3102 | 5146392 | 45.25 | 60 | 475379 |

| | | | | | | | | | | | | | | | | |
|---|---|---|---|---|---|---|---|---|---|---|---|---|---|---|---|---|
| SAMN02741364-GCF_000706865.2 | CFSAN006135 | SAMN02741364 | GCF_000706865.2 | NonPC | Public | 2013 | USA | Clinical | 91.5 | 100 | 0 | 98.2965 | 5090134 | 45.27 | 51 | 403575 |
| SAMN02741365-GCF_000707405.1 | CFSAN007429 | SAMN02741365 | GCF_000707405.1 | NonPC | Public | 2012 | USA | Clinical | 89.8 | 100 | 0.04 | 98.253 | 5156130 | 45.21 | 44 | 384893 |
| SAMN02741366-GCF_000707425.2 | CFSAN007430 | SAMN02741366 | GCF_000707425.2 | NonPC | Public | 2012 | USA | Clinical | 89.7 | 100 | 0.04 | 98.2471 | 5151483 | 45.21 | 55 | 341757 |
| SAMN02741367-GCF_000707385.2 | CFSAN007431 | SAMN02741367 | GCF_000707385.2 | NonPC | Public | 2012 | USA | Clinical | 89.8 | 100 | 0.04 | 98.2448 | 5217639 | 45.19 | 65 | 342528 |
| SAMN02741368-GCF_000707445.2 | CFSAN007432 | SAMN02741368 | GCF_000707445.2 | NonPC | Public | 2013 | USA | Clinical | 89.8 | 100 | 0.04 | 98.2469 | 5177534 | 45.2 | 41 | 546942 |
| SAMN02741369-GCF_000707465.2 | CFSAN007433 | SAMN02741369 | GCF_000707465.2 | NonPC | Public | 2013 | USA | Clinical | 89.8 | 100 | 0.18 | 98.2429 | 5388113 | 44.86 | 47 | 387475 |
| SAMN02741370-GCF_000707485.2 | CFSAN007434 | SAMN02741370 | GCF_000707485.2 | NonPC | Public | 2013 | USA | Clinical | 89.8 | 100 | 0.04 | 98.2264 | 5217140 | 45.19 | 42 | 360870 |
| SAMN02741371-GCF_000707505.2 | CFSAN007435 | SAMN02741371 | GCF_000707505.2 | NonPC | Public | 2013 | USA | Clinical | 89.3 | 100 | 0.04 | 98.2722 | 5137415 | 45.2 | 65 | 341544 |
| SAMN02741372-GCF_000706885.2 | CFSAN007436 | SAMN02741372 | GCF_000706885.2 | NonPC | Public | 2013 | USA | Clinical | 89.7 | 100 | 0.04 | 98.2888 | 5150773 | 45.21 | 44 | 384892 |
| SAMN02741373-GCF_000707165.1 | CFSAN007437 | SAMN02741373 | GCF_000707165.1 | NonPC | Public | 2012 | USA | Clinical | 90.9 | 100 | 0.12 | 98.1952 | 5139188 | 45.24 | 103 | 315055 |
| SAMN02741374-GCF_000707145.1 | CFSAN007438 | SAMN02741374 | GCF_000707145.1 | NonPC | Public | 2012 | USA | Clinical | 89.6 | 100 | 0.05 | 98.2913 | 5066853 | 45.35 | 30 | 389883 |
| SAMN02741375-GCF_000707025.1 | CFSAN007439 | SAMN02741375 | GCF_000707025.1 | NonPC | Public | 2012 | USA | Clinical | 88.8 | 100 | 0.06 | 98.1341 | 5062660 | 45.15 | 59 | 218861 |
| SAMN02741376-GCF_000707005.2 | CFSAN007440 | SAMN02741376 | GCF_000707005.2 | NonPC | Public | 2013 | USA | Clinical | 88.6 | 100 | 0.06 | 98.1622 | 5052686 | 45.16 | 56 | 231302 |
| SAMN02741377-GCF_000706945.2 | CFSAN007441 | SAMN02741377 | GCF_000706945.2 | NonPC | Public | 2012 | USA | Clinical | 89.2 | 100 | 0.05 | 98.1335 | 4965004 | 45.35 | 87 | 216488 |
| SAMN02741378-GCF_000707305.2 | CFSAN007442 | SAMN02741378 | GCF_000707305.2 | NonPC | Public | 2012 | USA | Clinical | 90.4 | 100 | 0.14 | 98.1939 | 5065123 | 45.19 | 77 | 281184 |
| SAMN02741379-GCF_000707105.2 | CFSAN007443 | SAMN02741379 | GCF_000707105.2 | NonPC | Public | 2012 | USA | Clinical | 91.1 | 100 | 0.27 | 98.2735 | 5081378 | 45.25 | 48 | 514403 |
| SAMN02741380-GCF_000707065.2 | CFSAN007444 | SAMN02741380 | GCF_000707065.2 | NonPC | Public | 2012 | USA | Clinical | 91.5 | 100 | 0 | 98.4065 | 4978443 | 45.4 | 45 | 412611 |
| SAMN02741381-GCF_000707125.2 | CFSAN007445 | SAMN02741381 | GCF_000707125.2 | NonPC | Public | 2012 | USA | Clinical | 89.1 | 100 | 0 | 98.0779 | 5018203 | 45.27 | 81 | 182322 |
| SAMN02741382-GCF_000707685.2 | CFSAN007446 | SAMN02741382 | GCF_000707685.2 | NonPC | Public | 2012 | USA | Clinical | 91.1 | 100 | 0.14 | 98.3657 | 4994370 | 45.34 | 64 | 271255 |
| SAMN02741383-GCF_000707705.2 | CFSAN007447 | SAMN02741383 | GCF_000707705.2 | NonPC | Public | 2012 | USA | Clinical | 91.5 | 100 | 0 | 98.3917 | 4998468 | 45.32 | 50 | 409016 |
| SAMN02741384-GCF_000707725.2 | CFSAN007448 | SAMN02741384 | GCF_000707725.2 | NonPC | Public | 2012 | USA | Clinical | 88.3 | 100 | 0.72 | 98.2514 | 5371853 | 45.05 | 95 | 172418 |
| SAMN02741385-GCF_000707745.2 | CFSAN007449 | SAMN02741385 | GCF_000707745.2 | PC | Public | 2012 | USA | Clinical | 98.3 | 100 | 0 | 99.9716 | 5119321 | 45.26 | 55 | 346912 |
| SAMN02741386-GCF_000707765.2 | CFSAN007450 | SAMN02741386 | GCF_000707765.2 | PC | Public | 2012 | USA | Clinical | 98.2 | 100 | 0 | 99.9843 | 5114004 | 45.25 | 65 | 288413 |
| SAMN02741387-GCF_000707805.2 | CFSAN007451 | SAMN02741387 | GCF_000707805.2 | PC | Public | 2012 | USA | Clinical | 98.3 | 100 | 0 | 99.9878 | 5121158 | 45.27 | 49 | 401590 |
| SAMN02741388-GCF_000707785.2 | CFSAN007452 | SAMN02741388 | GCF_000707785.2 | NonPC | Public | 2010 | USA | Clinical | 91.5 | 100 | 0 | 98.3982 | 5127281 | 45.15 | 37 | 527441 |
| SAMN02741389-GCF_000707825.2 | CFSAN007453 | SAMN02741389 | GCF_000707825.2 | NonPC | Public | 2010 | USA | Clinical | 91.4 | 100 | 0 | 98.3615 | 5124758 | 45.16 | 46 | 367090 |
| SAMN02741390-GCF_000707905.2 | CFSAN007454 | SAMN02741390 | GCF_000707905.2 | NonPC | Public | 2010 | USA | Env | 91.6 | 100 | 0 | 98.3399 | 5017786 | 45.33 | 36 | 445027 |
| SAMN02741391-GCF_000706985.2 | CFSAN007455 | SAMN02741391 | GCF_000706985.2 | NonPC | Public | 2013 | USA | Clinical | 90.4 | 100 | 0.35 | 98.2915 | 5152595 | 45.19 | 42 | 735315 |
| SAMN02741392-GCF_000707285.2 | CFSAN007456 | SAMN02741392 | GCF_000707285.2 | NonPC | Public | 2010 | USA | Env | 89.3 | 100 | 0.26 | 98.3118 | 5206921 | 45.24 | 56 | 480822 |
| SAMN02741393-GCF_000707085.2 | CFSAN007457 | SAMN02741393 | GCF_000707085.2 | NonPC | Public | 2010 | USA | Env | 91.4 | 99.86 | 0.08 | 98.3196 | 5183716 | 45.17 | 47 | 417837 |
| SAMN02741394-GCF_000707365.2 | CFSAN007458 | SAMN02741394 | GCF_000707365.2 | NonPC | Public | 2010 | USA | Env | 90.4 | 100 | 0.14 | 98.2017 | 5061948 | 45.2 | 52 | 321455 |
| SAMN02741395-GCF_000707185.2 | CFSAN007459 | SAMN02741395 | GCF_000707185.2 | NonPC | Public | 2010 | USA | Env | 91.4 | 99.93 | 0.08 | 98.2406 | 5205568 | 45.15 | 45 | 425650 |
| SAMN02741396-GCF_000706965.2 | CFSAN007460 | SAMN02741396 | GCF_000706965.2 | NonPC | Public | 2013 | USA | Clinical | 91.4 | 100 | 0 | 98.3261 | 5089463 | 45.27 | 58 | 418110 |
| SAMN02741397-GCF_000707205.2 | CFSAN007461 | SAMN02741397 | GCF_000707205.2 | NonPC | Public | 2013 | USA | Clinical | 91.5 | 100 | 0 | 98.2866 | 5091663 | 45.27 | 55 | 548102 |
| SAMN02741398-GCF_000706905.1 | CFSAN007462 | SAMN02741398 | GCF_000706905.1 | NonPC | Public | 2013 | USA | Clinical | 91.5 | 100 | 0 | 98.2816 | 5090919 | 45.27 | 47 | 447367 |
| SAMN02741399-GCF_000707645.2 | CFSAN012491 | SAMN02741399 | GCF_000707645.2 | NonPC | Public | 2010 | USA | Env | 91.4 | 100 | 0 | 98.3744 | 5126325 | 45.15 | 35 | 526001 |
| SAMN02741400-GCF_000707665.2 | CFSAN012492 | SAMN02741400 | GCF_000707665.2 | NonPC | Public | 2010 | USA | Env | 91.5 | 100 | 0 | 98.3543 | 5128956 | 45.16 | 34 | 485136 |
| SAMN02741401-GCF_000707265.2 | CFSAN012493 | SAMN02741401 | GCF_000707265.2 | NonPC | Public | 2010 | USA | Env | 91.4 | 99.93 | 0 | 98.3811 | 5123258 | 45.16 | 47 | 329201 |
| SAMN02741402-GCF_000707345.1 | CFSAN012494 | SAMN02741402 | GCF_000707345.1 | NonPC | Public | 2010 | USA | Env | 91.5 | 100 | 0 | 98.384 | 5127159 | 45.16 | 34 | 527658 |
| SAMN02781333-GCF_001270885.1 | ATC210 | SAMN02781333 | GCF_001270885.1 | PC | Public | 1998 | Chile | Clinical | 96.5 | 99.87 | 0 | 99.9824 | 5136577 | 45.33 | 22 | 838297 |
| SAMN02781334-GCF_001270975.1 | ATC220 | SAMN02781334 | GCF_001270975.1 | PC | Public | 1998 | Chile | Clinical | 99.3 | 100 | 0 | 99.9862 | 5145131 | 45.35 | 19 | 547874 |
| SAMN02781335-GCA_001270805.1 | PMA109.5 | SAMN02781335 | GCA_001270805.1 | PC | Public | 2005 | Chile | Env | 98.9 | 99.55 | 0.36 | 99.9536 | 5338346 | 45.29 | 20 | 582970 |
| SAMN02781336-GCF_001270895.1 | PMC14.7 | SAMN02781336 | GCF_001270895.1 | PC | Public | 2007 | Chile | Clinical | 98.9 | 99.73 | 0 | 99.9839 | 5133772 | 45.31 | 23 | 853734 |
| SAMN02781337-GCF_001270905.1 | PMC48 | SAMN02781337 | GCF_001270905.1 | PC | Public | 2004 | Chile | Clinical | 98.9 | 100 | 0 | 99.9836 | 5138290 | 45.34 | 13 | 992458 |
| SAMN02781338-GCF_001270815.1 | PMC58.5 | SAMN02781338 | GCF_001270815.1 | PC | Public | 2005 | Chile | Clinical | 99.3 | 100 | 0 | 99.9858 | 5154023 | 45.34 | 14 | 1090964 |
| SAMN02781339-GCF_001270825.1 | PMC58.7 | SAMN02781339 | GCF_001270825.1 | PC | Public | 2007 | Chile | Clinical | 99.3 | 99.95 | 0 | 99.9828 | 5162709 | 45.34 | 22 | 547261 |
| SAMN02781341-GCA_001270835.1 | PMA37.5 | SAMN02781341 | GCA_001270835.1 | PC | Public | 2005 | Chile | Env | 98.1 | 99.29 | 0.14 | 99.9807 | 5095862 | 45.25 | 92 | 175000 |
| SAMN02866418-GCA_000737635.1 | 13-028/A3 | SAMN02866418 | GCA_000737635.1 | NonPC | Public | 2013 | Viet_Nam | Env | 91.4 | 99.33 | 0.59 | 98.3456 | 5362496 | 45.3 | 201 | 103448 |

| | | | | | | | | | | | | | | | | |
|---|---|---|---|---|---|---|---|---|---|---|---|---|---|---|---|---|
| SAMN02867584-GCA_000732245.1 | 13-028/A2 | SAMN02867584 | GCA_000732245.1 | NonPC | Public | 2013 | Viet_Nam | Env | 89.3 | 99.51 | 0.64 | 98.2685 | 4917813 | 45.38 | 123 | 143936 |
| SAMN02910583-GCF_000736335.1 | NCKU_TV_3HP | SAMN02910583 | GCF_000736335.1 | NonPC | Public | 1999 | Thailand | Env | 91.8 | 100 | 0 | 98.2553 | 5537284 | 45.21 | 85 | 266574 |
| SAMN02910584-GCF_000736315.1 | NCKU_TV_5HP | SAMN02910584 | GCF_000736315.1 | NonPC | Public | 1999 | Thailand | Env | 91.8 | 100 | 0 | 98.2687 | 5532049 | 45.22 | 76 | 266695 |
| SAMN02910585-GCF_000736325.1 | NCKU_CV_CHN | SAMN02910585 | GCF_000736325.1 | NonPC | Public | 2010 | China | Env | 91.2 | 100 | 0.32 | 98.2299 | 5311226 | 45.26 | 126 | 551331 |
| SAMN02910586-GCF_000736345.1 | NCKU_TN_S02 | SAMN02910586 | GCF_000736345.1 | NonPC | Public | 2008 | Thailand | Env | 90.9 | 100 | 0.41 | 98.3306 | 5405810 | 45.21 | 87 | 520118 |
| SAMN02912085-GCF_000732995.1 | FIM-S1392- | SAMN02912085 | GCF_000732995.1 | NonPC | Public | 2014 | Mexico | Env | 91.3 | 99.57 | 0.27 | 98.3058 | 5174919 | 45.32 | 14 | 1931617 |
| SAMN02912086-GCF_000732985.1 | FIM-S1708+ | SAMN02912086 | GCF_000732985.1 | NonPC | Public | 2014 | Mexico | Env | 89.9 | 100 | 0.39 | 98.2016 | 5246988 | 45.34 | 79 | 174266 |
| SAMN02952920-GCF_009649015.1 | 10329 | SAMN02952920 | GCF_009649015.1 | NonPC | Public | 1998 | USA | Clinical | 92.3 | 100 | 0 | 98.3175 | 5144090 | 45.32 | 2 | 3311004 |
| SAMN03081521-GCF_000568495.1 | UCM-V493 | SAMN03081521 | GCF_000568495.1 | NonPC | Public | 2002 | Spain | NA | 89 | 100 | 0.61 | 98.2774 | 5232572 | 45.33 | 3 | 3445790 |
| SAMN03107383-GCF_001023055.1 | MAVP-26 | SAMN03107383 | GCF_001023055.1 | NonPC | Public | 2013 | NA | Clinical | 91.6 | 100 | 0 | 98.3056 | 5112012 | 45.26 | 28 | 809118 |
| SAMN03107385-GCF_001023105.1 | MAVP-36 | SAMN03107385 | GCF_001023105.1 | NonPC | Public | 2013 | NA | Clinical | 91.7 | 100 | 0.09 | 98.3117 | 5193467 | 45.27 | 31 | 809400 |
| SAMN03107386-GCF_001007525.1 | MAVP-E | SAMN03107386 | GCF_001007525.1 | NonPC | Public | NA | NA | Clinical | 89.7 | 100 | 0.52 | 98.2392 | 5250566 | 45.23 | 148 | 339816 |
| SAMN03140317-GCF_001433415.1 | FORC_004 | SAMN03140317 | GCF_001433415.1 | NonPC | Public | 2014 | South_Korea | Env | 92.2 | 100 | 0.11 | 98.2659 | 5171661 | 45.48 | 3 | 3348850 |
| SAMN03140318-GCF_001304775.1 | FORC_006 | SAMN03140318 | GCF_001304775.1 | NonPC | Public | 2014 | South_Korea | Env | 88.4 | 100 | 0.36 | 98.2115 | 5094100 | 45.37 | 2 | 3376731 |
| SAMN03159469-GCF_001244315.1 | FORC_008 | SAMN03159469 | GCF_001244315.1 | NonPC | Public | 2014 | South_Korea | Env | 91.8 | 100 | 0.08 | 98.2817 | 5038168 | 45.42 | 2 | 3266132 |
| SAMN03177808-GCF_001023155.1 | MAVP-M | SAMN03177808 | GCF_001023155.1 | NonPC | Public | 2011 | NA | Clinical | 89.7 | 100 | 0.08 | 98.1993 | 5204499 | 45.09 | 146 | 402360 |
| SAMN03177809-GCF_001007545.1 | MAVP-V | SAMN03177809 | GCF_001007545.1 | NonPC | Public | 2011 | NA | Clinical | 91.5 | 100 | 0.01 | 98.2967 | 5200026 | 45.27 | 29 | 764045 |
| SAMN03177810-GCF_001007535.1 | MAVP-45 | SAMN03177810 | GCF_001007535.1 | NonPC | Public | 2013 | NA | Clinical | 91.6 | 100 | 0 | 98.3451 | 5111481 | 45.27 | 20 | 812214 |
| SAMN03177811-GCF_001023125.1 | CT4287 | SAMN03177811 | GCF_001023125.1 | NonPC | Public | 2013 | NA | Env | 90.6 | 100 | 0 | 98.3138 | 5231329 | 45.19 | 38 | 558312 |
| SAMN03195665-GCF_000786835.1 | T12739 | SAMN03195665 | GCF_000786835.1 | NonPC | Public | 2007 | Canada | Clinical | 89.9 | 100 | 0.51 | 98.3565 | 5005773 | 45.3 | 62 | 179503 |
| SAMN03195667-GCF_000786845.1 | T9109 | SAMN03195667 | GCF_000786845.1 | NonPC | Public | 2007 | Canada | Clinical | 91.4 | 100 | 1.01 | 98.3555 | 5065711 | 45.28 | 45 | 484812 |
| SAMN03195675-GCF_000786855.1 | 4.2548 | SAMN03195675 | GCF_000786855.1 | NonPC | Public | 2004 | Canada | Clinical | 91.4 | 100 | 0.11 | 98.271 | 5471842 | 45.06 | 54 | 244392 |
| SAMN03195677-GCF_000786865.1 | 9.5357 | SAMN03195677 | GCF_000786865.1 | NonPC | Public | 2009 | Canada | Clinical | 90.2 | 100 | 3.19 | 98.3199 | 5180823 | 45.21 | 67 | 387375 |
| SAMN03255431-GCF_001188185.2 | FDA:FDAARGOS_51 | SAMN03255431 | GCF_001188185.2 | NonPC | Public | 1998 | USA | Clinical | 92.3 | 100 | 0 | 98.3298 | 5149217 | 45.33 | 2 | 3316147 |
| SAMN03255432-GCF_001188135.2 | FDA:FDAARGOS_52 | SAMN03255432 | GCF_001188135.2 | NonPC | Public | 1998 | USA | Env | 91.6 | 100 | 0.14 | 98.1682 | 5173745 | 45.26 | 4 | 3055024 |
| SAMN03255433-GCF_001188035.2 | FDA:FDAARGOS_53 | SAMN03255433 | GCF_001188035.2 | NonPC | Public | 2004 | USA | Env | 92.5 | 100 | 0.2 | 98.3134 | 5318119 | 45.23 | 8 | 972395 |
| SAMN03287713-GCF_000878815.1 | 04-1290 | SAMN03287713 | GCF_000878815.1 | NonPC | Public | 2004 | Canada | Clinical | 91.5 | 100 | 0.09 | 98.3166 | 5132984 | 45.21 | 72 | 548500 |
| SAMN03287714-GCF_000878785.1 | 09-3216 | SAMN03287714 | GCF_000878785.1 | NonPC | Public | 2009 | Canada | Clinical | 91.5 | 100 | 0 | 98.3408 | 5090133 | 45.29 | 53 | 548674 |
| SAMN03287715-GCF_000878805.1 | 10-4241 | SAMN03287715 | GCF_000878805.1 | NonPC | Public | 2006 | Canada | Clinical | 91.5 | 100 | 0 | 98.2795 | 5097015 | 45.28 | 38 | 548500 |
| SAMN03287757-GCF_000878755.1 | 10-4242 | SAMN03287757 | GCF_000878755.1 | NonPC | Public | 2006 | Canada | Clinical | 91.2 | 100 | 0.24 | 98.3411 | 5117538 | 45.26 | 51 | 403279 |
| SAMN03287758-GCF_000878725.1 | 10-4245 | SAMN03287758 | GCF_000878725.1 | NonPC | Public | 2006 | Canada | Clinical | 91.5 | 100 | 0.05 | 98.3188 | 5087132 | 45.29 | 47 | 423844 |
| SAMN03287759-GCF_000878705.1 | 10-4246 | SAMN03287759 | GCF_000878705.1 | NonPC | Public | 2006 | Canada | Clinical | 91.5 | 100 | 0 | 98.3534 | 5088681 | 45.29 | 50 | 423840 |
| SAMN03287760-GCF_000878665.1 | 10-4247 | SAMN03287760 | GCF_000878665.1 | NonPC | Public | 2006 | Canada | Clinical | 91.5 | 100 | 0 | 98.3275 | 5115078 | 45.27 | 61 | 462063 |
| SAMN03287761-GCF_000878675.1 | 10-4248 | SAMN03287761 | GCF_000878675.1 | NonPC | Public | 2006 | Canada | Clinical | 91.5 | 100 | 0.02 | 98.3282 | 5088257 | 45.29 | 52 | 548500 |
| SAMN03287762-GCF_000878735.1 | 10-4274 | SAMN03287762 | GCF_000878735.1 | NonPC | Public | 2005 | Canada | Clinical | 91.5 | 100 | 0.52 | 98.2826 | 5094836 | 45.28 | 44 | 548500 |
| SAMN03287763-GCF_000878645.1 | 10-4288 | SAMN03287763 | GCF_000878645.1 | NonPC | Public | 2003 | Canada | Clinical | 91.5 | 100 | 0 | 98.3169 | 5103957 | 45.27 | 46 | 548500 |
| SAMN03287764-GCF_000878595.1 | 10-4293 | SAMN03287764 | GCF_000878595.1 | NonPC | Public | 2002 | Canada | Clinical | 91.7 | 100 | 0 | 98.3334 | 5196520 | 45.19 | 44 | 444499 |
| SAMN03287765-GCF_000878565.1 | 10-4298 | SAMN03287765 | GCF_000878565.1 | NonPC | Public | 2001 | Canada | Clinical | 91.9 | 100 | 0.14 | 98.2645 | 5225557 | 45.18 | 56 | 426178 |
| SAMN03287766-GCF_000878575.1 | 10-4303 | SAMN03287766 | GCF_000878575.1 | NonPC | Public | 2000 | Canada | Clinical | 91.7 | 100 | 0 | 98.3147 | 5099537 | 45.27 | 34 | 437054 |
| SAMN03287767-GCF_000878585.1 | 10-7197 | SAMN03287767 | GCF_000878585.1 | NonPC | Public | 2008 | Canada | Clinical | 91.5 | 100 | 0 | 98.3143 | 5086376 | 45.29 | 44 | 475405 |
| SAMN03325855-GCF_001700835.1 | CHN25 | SAMN03325855 | GCF_001700835.1 | NonPC | Public | 2011 | China | Env | 90.6 | 100 | 0.07 | 98.288 | 5443401 | 45.23 | 5 | 3416467 |
| SAMN03349598-GCF_000951795.1 | 04-2549 | SAMN03349598 | GCF_000951795.1 | PC | Public | 2004 | Canada | Clinical | 98.4 | 100 | 0.2 | 99.9828 | 5105512 | 45.3 | 75 | 411035 |
| SAMN03349602-GCF_000960565.1 | 07-2965 | SAMN03349602 | GCF_000960565.1 | NonPC | Public | 2007 | Canada | Clinical | 90.9 | 100 | 0.04 | 98.2792 | 5206189 | 45.22 | 57 | 557313 |
| SAMN03349603-GCF_000960645.1 | 08-0278 | SAMN03349603 | GCF_000960645.1 | NonPC | Public | 2008 | Canada | Clinical | 89.7 | 100 | 1.06 | 98.3142 | 5154387 | 45.03 | 56 | 402713 |
| SAMN03349604-GCF_000960665.1 | 08-7626 | SAMN03349604 | GCF_000960665.1 | NonPC | Public | 2008 | Canada | Clinical | 91 | 100 | 0.6 | 98.2719 | 5188330 | 45.19 | 70 | 315699 |
| SAMN03349605-GCF_000960655.1 | 09-3217 | SAMN03349605 | GCF_000960655.1 | NonPC | Public | 2009 | Canada | Clinical | 89.2 | 100 | 0.27 | 98.061 | 5036314 | 45.27 | 93 | 319036 |
| SAMN03349606-GCF_000960685.1 | 09-4435 | SAMN03349606 | GCF_000960685.1 | PC | Public | 2009 | Canada | Clinical | 98.4 | 100 | 0 | 99.9903 | 5102456 | 45.28 | 67 | 288608 |
| SAMN03349607-GCF_001006115.1 | 09-4663 | SAMN03349607 | GCF_001006115.1 | NonPC | Public | 2009 | Canada | Clinical | 90.9 | 100 | 0.33 | 98.2983 | 5188104 | 45.18 | 74 | 403392 |

| | | | | | | | | | | | | | | | | |
|---|---|---|---|---|---|---|---|---|---|---|---|---|---|---|---|---|
| SAMN03349608-GCF_001006125.1 | 10-4244 | SAMN03349608 | GCF_001006125.1 | NonPC | Public | 2006 | Canada | Clinical | 90.4 | 100 | 0.61 | 98.274 | 5240907 | 45.17 | 56 | 383602 |
| SAMN03349609-GCF_001006105.1 | 10-4251 | SAMN03349609 | GCF_001006105.1 | PC | Public | 2006 | Canada | Clinical | 98.3 | 100 | 0.35 | 99.9565 | 5200339 | 45.11 | 107 | 157866 |
| SAMN03349610-GCF_001006195.1 | 10-4255 | SAMN03349610 | GCF_001006195.1 | NonPC | Public | 2006 | Canada | Clinical | 89.4 | 100 | 0.36 | 98.0958 | 5043842 | 45.28 | 77 | 239950 |
| SAMN03349611-GCF_001006185.1 | 10-4287 | SAMN03349611 | GCF_001006185.1 | NonPC | Public | 2003 | Canada | Clinical | 91.2 | 100 | 2.5 | 98.2715 | 5160841 | 45.2 | 62 | 509297 |
| SAMN03349612-GCF_001006205.1 | 10-7205 | SAMN03349612 | GCF_001006205.1 | NonPC | Public | 2008 | Canada | Clinical | 91 | 100 | 0.63 | 98.297 | 5189627 | 45.19 | 73 | 315571 |
| SAMN03358815-GCA_015820245.1 | BCW_3098 | SAMN03358815 | GCA_015820245.1 | NonPC | Public | NA | NA | Env | 91.6 | 100 | 0.08 | 98.3591 | 5153950 | 45.27 | 91 | 448033 |
| SAMN03358816-GCF_001726165.1 | BCW_3099 | SAMN03358816 | GCF_001726165.1 | NonPC | Public | 2007 | USA | Env | 89.7 | 100 | 0.15 | 98.3038 | 5174516 | 45.28 | 59 | 460669 |
| SAMN03358817-GCF_001726175.1 | BCW_3103 | SAMN03358817 | GCF_001726175.1 | NonPC | Public | 2007 | USA | Env | 89.7 | 100 | 0 | 98.1063 | 5054587 | 45.25 | 105 | 191541 |
| SAMN03358818-GCF_001726485.1 | BCW_3115 | SAMN03358818 | GCF_001726485.1 | NonPC | Public | 2007 | USA | Env | 88.6 | 100 | 0.35 | 98.1553 | 4970708 | 45.33 | 72 | 314803 |
| SAMN03358819-GCF_001726535.1 | BCW_3118 | SAMN03358819 | GCF_001726535.1 | NonPC | Public | 2007 | USA | Env | 91.6 | 100 | 0 | 98.344 | 5035158 | 45.28 | 60 | 401329 |
| SAMN03358820-GCF_001726685.1 | BCW_3125 | SAMN03358820 | GCF_001726685.1 | NonPC | Public | 2007 | USA | Env | 88.7 | 100 | 0.05 | 98.1072 | 4940635 | 45.34 | 87 | 207497 |
| SAMN03358821-GCF_001726935.1 | BCW_3141 | SAMN03358821 | GCF_001726935.1 | NonPC | Public | 2007 | Canada | Env | 88.6 | 100 | 0.11 | 98.1282 | 4922746 | 45.33 | 82 | 246453 |
| SAMN03358822-GCF_001726965.1 | BCW_3143 | SAMN03358822 | GCF_001726965.1 | NonPC | Public | 2007 | Canada | Env | 91.9 | 100 | 0.05 | 98.3631 | 5110782 | 45.21 | 55 | 413198 |
| SAMN03358823-GCF_001726975.1 | BCW_3146 | SAMN03358823 | GCF_001726975.1 | NonPC | Public | 2007 | Canada | Env | 91.9 | 100 | 0.14 | 98.3879 | 5144621 | 45.19 | 56 | 348887 |
| SAMN03358824-GCF_001727055.1 | BCW_3148 | SAMN03358824 | GCF_001727055.1 | NonPC | Public | 2007 | Canada | Env | 91.9 | 100 | 0.05 | 98.3628 | 5109709 | 45.21 | 51 | 413181 |
| SAMN03358825-GCF_001727275.1 | BCW_3160 | SAMN03358825 | GCF_001727275.1 | NonPC | Public | 2007 | USA | Env | 88.6 | 100 | 0 | 98.1067 | 5072118 | 45.24 | 136 | 177799 |
| SAMN03358826-GCF_001727245.1 | BCW_3162 | SAMN03358826 | GCF_001727245.1 | NonPC | Public | 2007 | USA | Env | 88.6 | 100 | 0 | 98.1028 | 5071913 | 45.24 | 131 | 183086 |
| SAMN03358827-GCF_001727805.1 | BCW_3171 | SAMN03358827 | GCF_001727805.1 | NonPC | Public | 2006 | USA | Clinical | 92 | 100 | 0.22 | 98.2453 | 5328804 | 45.06 | 99 | 208879 |
| SAMN03358828-GCF_001727815.1 | BCW_3172 | SAMN03358828 | GCF_001727815.1 | PC | Public | 2006 | USA | Clinical | 98.3 | 100 | 0 | 99.9928 | 5144946 | 45.3 | 89 | 259160 |
| SAMN03358829-GCF_001727825.1 | BCW_3173 | SAMN03358829 | GCF_001727825.1 | PC | Public | 2006 | USA | Clinical | 98.3 | 100 | 0 | 99.9933 | 5142252 | 45.3 | 85 | 268238 |
| SAMN03358830-GCF_001727895.1 | BCW_3175 | SAMN03358830 | GCF_001727895.1 | NonPC | Public | 2006 | USA | Clinical | 91.3 | 100 | 0 | 98.3089 | 5078582 | 45.26 | 81 | 548356 |
| SAMN03358831-GCF_001727965.1 | BCW_3181 | SAMN03358831 | GCF_001727965.1 | NonPC | Public | 2006 | USA | Clinical | 91.1 | 100 | 0 | 98.3213 | 5021660 | 45.35 | 74 | 519873 |
| SAMN03358832-GCF_001728375.1 | BCW_3214 | SAMN03358832 | GCF_001728375.1 | NonPC | Public | 2007 | USA | Clinical | 90 | 99.73 | 0.22 | 98.3599 | 5154351 | 45.2 | 101 | 192823 |
| SAMN03358833-GCF_001728415.1 | BCW_3218 | SAMN03358833 | GCF_001728415.1 | NonPC | Public | 2007 | USA | Clinical | 91.3 | 100 | 0 | 98.3123 | 5073998 | 45.27 | 86 | 548384 |
| SAMN03358834-GCF_001727645.1 | BCW_3219 | SAMN03358834 | GCF_001727645.1 | NonPC | Public | 2007 | USA | Clinical | 90.8 | 100 | 0.14 | 98.2656 | 5075905 | 45.23 | 88 | 312872 |
| SAMN03358835-GCF_001728565.1 | BCW_3229 | SAMN03358835 | GCF_001728565.1 | NonPC | Public | 2007 | USA | Clinical | 90.9 | 100 | 0.05 | 98.2912 | 5137243 | 45.21 | 82 | 321792 |
| SAMN03358836-GCF_001728655.1 | BCW_3235 | SAMN03358836 | GCF_001728655.1 | NonPC | Public | 2007 | USA | Clinical | 91.7 | 100 | 0 | 98.2536 | 5091239 | 45.26 | 96 | 208616 |
| SAMN03358837-GCF_001728665.1 | BCW_3236 | SAMN03358837 | GCF_001728665.1 | NonPC | Public | 2007 | USA | Clinical | 91.7 | 100 | 0 | 98.2429 | 5084458 | 45.26 | 84 | 208616 |
| SAMN03358838-GCF_001728745.1 | BCW_3240 | SAMN03358838 | GCF_001728745.1 | NonPC | Public | NA | USA | Clinical | 91.3 | 100 | 0 | 98.3222 | 5131444 | 45.28 | 86 | 548356 |
| SAMN03358839-GCF_001728755.1 | BCW_3241 | SAMN03358839 | GCF_001728755.1 | NonPC | Public | 2007 | USA | Clinical | 90.7 | 100 | 0 | 98.4014 | 5145112 | 45.21 | 55 | 315632 |
| SAMN03358840-GCA_015820125.1 | BCW_3242 | SAMN03358840 | GCA_015820125.1 | NonPC | Public | NA | NA | Env | 89.5 | 100 | 0.05 | 98.413 | 4842526 | 45.45 | 68 | 446853 |
| SAMN03372139-GCA_004137845.1 | CAIM_320 | SAMN03372139 | GCA_004137845.1 | NonPC | Public | 1950 | Japan | Env | 90.6 | 98.49 | 0.46 | 98.3157 | 5063368 | 45.23 | 70 | 237891 |
| SAMN03452289-GCF_000975195.1 | 04-2551 | SAMN03452289 | GCF_000975195.1 | PC | Public | 2004 | Canada | Clinical | 98.7 | 100 | 0.5 | 99.9934 | 5180034 | 45.36 | 63 | 339951 |
| SAMN03452290-GCF_000972045.1 | 07-1339 | SAMN03452290 | GCF_000972045.1 | PC | Public | 2007 | Canada | Clinical | 98.6 | 100 | 0 | 99.9716 | 5244148 | 45.11 | 68 | 373794 |
| SAMN03452291-GCF_000974905.1 | 09-3218 | SAMN03452291 | GCF_000974905.1 | NonPC | Public | 2009 | Canada | Clinical | 91 | 100 | 0.33 | 98.2939 | 5196469 | 45.19 | 67 | 418558 |
| SAMN03452292-GCF_000972125.1 | 09-4434 | SAMN03452292 | GCF_000972125.1 | NonPC | Public | 2009 | Canada | Clinical | 91 | 100 | 0.33 | 98.2892 | 5186534 | 45.18 | 56 | 425238 |
| SAMN03452293-GCF_000972055.1 | 09-4660 | SAMN03452293 | GCF_000972055.1 | NonPC | Public | 2009 | Canada | Clinical | 90.9 | 100 | 0.33 | 98.2799 | 5187132 | 45.18 | 59 | 315795 |
| SAMN03452294-GCF_000972035.1 | 09-4664 | SAMN03452294 | GCF_000972035.1 | NonPC | Public | 2009 | Canada | Clinical | 91 | 100 | 0.33 | 98.3294 | 5187023 | 45.18 | 61 | 425466 |
| SAMN03452299-GCF_000972025.1 | 09-4681 | SAMN03452299 | GCF_000972025.1 | NonPC | Public | 2009 | Canada | Clinical | 91.7 | 100 | 0.29 | 98.2605 | 5171306 | 45.25 | 31 | 536325 |
| SAMN03452300-GCF_000972105.1 | 10-4243 | SAMN03452300 | GCF_000972105.1 | NonPC | Public | 2006 | Canada | Clinical | 90.4 | 100 | 0.41 | 98.2469 | 5243884 | 45.16 | 49 | 525588 |
| SAMN03457164-GCF_001636035.1 | FORC_014 | SAMN03457164 | GCF_001636035.1 | NonPC | Public | 2015 | South_Korea | Env | 91.8 | 100 | 0.05 | 98.2403 | 5289960 | 45.32 | 3 | 3241330 |
| SAMN03458146-GCF_001011015.1 | ATCC_17802 | SAMN03458146 | GCF_001011015.1 | NonPC | Public | 1951 | Japan | Clinical | 90.7 | 100 | 0.33 | 98.3468 | 5067729 | 45.25 | 51 | 366570 |
| SAMN03465090-GCF_002072975.1 | VN-0028 | SAMN03465090 | GCF_002072975.1 | NonPC | Public | 1995 | Germany | Clinical | 91 | 100 | 0.54 | 98.2449 | 5153221 | 45.21 | 80 | 166252 |
| SAMN03470168-GCF_002072985.1 | VN-0029 | SAMN03470168 | GCF_002072985.1 | NonPC | Public | 1995 | Germany | Clinical | 89.9 | 100 | 0.48 | 98.337 | 5195632 | 45.12 | 55 | 237978 |
| SAMN03470171-GCF_002073025.1 | VN-5003 | SAMN03470171 | GCF_002073025.1 | NonPC | Public | 2009 | Germany | Env | 91.5 | 100 | 0.15 | 98.2395 | 4982635 | 45.38 | 54 | 261848 |
| SAMN03470181-GCF_002073005.1 | VN-0293 | SAMN03470181 | GCF_002073005.1 | NonPC | Public | 2011 | Germany | Env | 89.5 | 100 | 0.9 | 98.2749 | 5224548 | 45.09 | 86 | 217637 |
| SAMN03646983-GCF_001013435.1 | VH3 | SAMN03646983 | GCF_001013435.1 | NonPC | Public | 2007 | Greece | Env | 89.5 | 100 | 0 | 98.2973 | 4955051 | 45.36 | 67 | 370587 |
| SAMN03701448-GCF_001758605.1 | 378 | SAMN03701448 | GCF_001758605.1 | NonPC | Public | 2014 | South_Korea | Clinical | 90.5 | 100 | 0.05 | 98.3044 | 5015214 | 45.44 | 2 | 3253462 |

| | | | | | | | | | | | | | | | | |
|---|---|---|---|---|---|---|---|---|---|---|---|---|---|---|---|---|
| SAMN03735013-GCF_001497485.1 | K23 | SAMN03735013 | GCF_001497485.1 | NonPC | Public | 2013 | India | Env | 90.1 | 99.93 | 0.27 | 98.3097 | 5084087 | 45.34 | 58 | 410730 |
| SAMN03766034-GCF_001682175.1 | MAVP-Q | SAMN03766034 | GCF_001682175.1 | NonPC | Public | 2011 | NA | Clinical | 90.8 | 100 | 0.04 | 98.2406 | 5257288 | 45.29 | 2 | 3369172 |
| SAMN03795177-GCF_001267555.1 | ISF-01-07 | SAMN03795177 | GCF_001267555.1 | NonPC | Public | 2010 | Canada | Env | 92.4 | 100 | 0.05 | 98.3777 | 5043382 | 45.29 | 69 | 450650 |
| SAMN03795178-GCF_001267595.1 | ISF-25-6 | SAMN03795178 | GCF_001267595.1 | NonPC | Public | 2010 | Canada | Env | 89.3 | 100 | 0.34 | 98.2241 | 5069601 | 45.24 | 51 | 408976 |
| SAMN03795180-GCF_001267965.1 | RM-13-3 | SAMN03795180 | GCF_001267965.1 | NonPC | Public | 2015 | Canada | Env | 92.3 | 100 | 0.66 | 98.3639 | 5229823 | 45.27 | 48 | 427514 |
| SAMN03795181-GCF_001267655.1 | RM-17-6 | SAMN03795181 | GCF_001267655.1 | NonPC | Public | 2015 | Canada | Env | 90.7 | 100 | 0.63 | 98.2253 | 5237488 | 45.22 | 47 | 458622 |
| SAMN03795184-GCF_001268005.1 | S195-7 | SAMN03795184 | GCF_001268005.1 | NonPC | Public | 2007 | Canada | Env | 91.5 | 100 | 0.35 | 98.2339 | 5215691 | 45.21 | 60 | 324181 |
| SAMN03795185-GCF_001268015.1 | S349-10 | SAMN03795185 | GCF_001268015.1 | NonPC | Public | 2010 | Canada | Env | 91.2 | 100 | 0.23 | 98.3063 | 5218518 | 45.2 | 53 | 273670 |
| SAMN03795186-GCF_001267625.1 | S383-6 | SAMN03795186 | GCF_001267625.1 | NonPC | Public | 2011 | Canada | Env | 90.9 | 100 | 0.07 | 98.272 | 5095043 | 45.31 | 48 | 461802 |
| SAMN03795187-GCF_001267635.1 | S448-16 | SAMN03795187 | GCF_001267635.1 | NonPC | Public | 2012 | Canada | Env | 91 | 100 | 0.2 | 98.2537 | 5106840 | 45.32 | 50 | 523860 |
| SAMN03799829-GCF_001273555.1 | RM-14-5 | SAMN03799829 | GCF_001273555.1 | NonPC | Public | 2014 | Canada | Env | 90.5 | 100 | 0 | 98.2504 | 5054091 | 45.23 | 78 | 279336 |
| SAMN03799830-GCF_001273575.1 | ISF-29-3 | SAMN03799830 | GCF_001273575.1 | NonPC | Public | 2011 | Canada | Env | 90.9 | 100 | 0.49 | 98.3072 | 5361039 | 45.09 | 75 | 255897 |
| SAMN03799832-GCF_001270125.1 | HS-13-1 | SAMN03799832 | GCF_001270125.1 | NonPC | Public | 2014 | Canada | Env | 91.7 | 100 | 0 | 98.3736 | 5088377 | 45.27 | 78 | 401486 |
| SAMN03799834-GCF_001280635.1 | ISF-54-12 | SAMN03799834 | GCF_001280635.1 | NonPC | Public | 2011 | Canada | Env | 89.1 | 100 | 0 | 98.1284 | 5036473 | 45.44 | 62 | 335149 |
| SAMN03799835-GCF_001270285.1 | ISF-77-01 | SAMN03799835 | GCF_001270285.1 | NonPC | Public | 2011 | Canada | Env | 91 | 100 | 0.12 | 98.3552 | 5030433 | 45.31 | 62 | 518506 |
| SAMN03799836-GCF_001280645.1 | ISF-94-1 | SAMN03799836 | GCF_001280645.1 | NonPC | Public | 2011 | Canada | Env | 89.4 | 100 | 0.14 | 98.3072 | 5048972 | 45.29 | 40 | 407861 |
| SAMN03799839-GCF_001273635.1 | S357-21 | SAMN03799839 | GCF_001273635.1 | NonPC | Public | 2010 | Canada | Env | 89.6 | 100 | 0 | 98.11 | 5055485 | 45.27 | 99 | 173413 |
| SAMN03799840-GCF_001280655.1 | S372-5 | SAMN03799840 | GCF_001280655.1 | NonPC | Public | 2011 | Canada | Env | 90.5 | 100 | 0.19 | 98.244 | 5141951 | 45.12 | 66 | 480510 |
| SAMN03799842-GCF_001270235.1 | S440-7 | SAMN03799842 | GCF_001270235.1 | NonPC | Public | 2012 | Canada | Env | 90.5 | 100 | 0.14 | 98.1741 | 5121772 | 45.21 | 66 | 470844 |
| SAMN03799853-GCF_001270215.1 | S487-4 | SAMN03799853 | GCF_001270215.1 | NonPC | Public | 2013 | Canada | Env | 89.9 | 100 | 0.04 | 98.2763 | 5169202 | 45.22 | 70 | 431578 |
| SAMN03800013-GCF_001270145.1 | S499-7 | SAMN03800013 | GCF_001270145.1 | NonPC | Public | 2013 | Canada | Env | 90.9 | 100 | 0.07 | 98.2736 | 5102682 | 45.31 | 62 | 523860 |
| SAMN03941060-GCF_001695955.1 | 428/00 | SAMN03941060 | GCF_001695955.1 | NonPC | Public | 1998 | Spain | Clinical | 92.2 | 100 | 0.65 | 98.3911 | 5138507 | 45.19 | 66 | 145778 |
| SAMN03941061-GCF_001695995.1 | 30824 | SAMN03941061 | GCF_001695995.1 | NonPC | Public | 1999 | Spain | Clinical | 92.2 | 100 | 0.65 | 98.4166 | 5075961 | 45.24 | 61 | 185164 |
| SAMN03941062-GCF_001695945.1 | 44569 | SAMN03941062 | GCF_001695945.1 | PC | Public | 2004 | Spain | Clinical | 98 | 100 | 0 | 99.9935 | 5073596 | 45.29 | 60 | 267687 |
| SAMN03941063-GCF_001695935.1 | CFSAN018755 | SAMN03941063 | GCF_001695935.1 | NonPC | Public | 2002 | Spain | Env | 91.7 | 100 | 0.14 | 98.3351 | 5112517 | 45.34 | 76 | 379292 |
| SAMN03941064-GCF_001696005.1 | CFSAN018756 | SAMN03941064 | GCF_001696005.1 | NonPC | Public | 2003 | Spain | Env | 88.2 | 100 | 0.39 | 98.2583 | 4969236 | 45.33 | 64 | 246923 |
| SAMN03941065-GCF_001696015.1 | 906-97 | SAMN03941065 | GCF_001696015.1 | PC | Public | 1997 | Peru | Clinical | 98.5 | 100 | 0 | 99.987 | 5099401 | 45.29 | 72 | 288855 |
| SAMN03941067-GCF_001696115.1 | CFSAN001174 | SAMN03941067 | GCF_001696115.1 | NonPC | Public | 2004 | USA | Env | 91.1 | 100 | 0.05 | 98.2756 | 5148437 | 45.19 | 88 | 122650 |
| SAMN03941068-GCF_001696085.1 | CFSAN018760 | SAMN03941068 | GCF_001696085.1 | NonPC | Public | 2004 | USA | Env | 88.9 | 100 | 0.14 | 98.1136 | 4986443 | 45.28 | 61 | 252511 |
| SAMN03941069-GCF_001696065.1 | CFSAN018761 | SAMN03941069 | GCF_001696065.1 | NonPC | Public | 2004 | USA | Env | 89.8 | 99.73 | 0.22 | 98.2767 | 5284763 | 45.11 | 37 | 564826 |
| SAMN03941070-GCF_001696155.1 | CFSAN018762 | SAMN03941070 | GCF_001696155.1 | NonPC | Public | 2004 | USA | Env | 89.2 | 99.73 | 0.67 | 98.3497 | 5309218 | 45.2 | 504 | 21464 |
| SAMN03941071-GCF_001696135.1 | CFSAN018763 | SAMN03941071 | GCF_001696135.1 | NonPC | Public | 2004 | USA | Env | 89.5 | 100 | 0 | 98.0638 | 5045122 | 45.28 | 81 | 216660 |
| SAMN03941072-GCF_001696145.1 | CFSAN018764 | SAMN03941072 | GCF_001696145.1 | NonPC | Public | 2004 | USA | Env | 91.1 | 100 | 0 | 98.2591 | 5139686 | 45.23 | 46 | 642262 |
| SAMN03941073-GCF_001696165.1 | CFSAN018765 | SAMN03941073 | GCF_001696165.1 | NonPC | Public | 2004 | USA | Env | 91.1 | 100 | 0.01 | 98.2875 | 5245157 | 45.23 | 60 | 256190 |
| SAMN03941074-GCF_001696215.1 | CFSAN018766 | SAMN03941074 | GCF_001696215.1 | NonPC | Public | 2004 | USA | Env | 89.9 | 99.93 | 0.05 | 98.2823 | 5043748 | 45.3 | 60 | 348215 |
| SAMN03941075-GCF_001696225.1 | CFSAN018767 | SAMN03941075 | GCF_001696225.1 | NonPC | Public | 2004 | USA | Env | 89.5 | 100 | 0 | 98.1107 | 5085389 | 45.28 | 85 | 185639 |
| SAMN03945128-GCF_002198215.1 | CDC__K5280 | SAMN03945128 | GCF_002198215.1 | NonPC | Public | 2007 | USA | Clinical | 91.3 | 100 | 0.27 | 98.3255 | 5080467 | 45.28 | 73 | 322295 |
| SAMN03945129-GCF_002198195.1 | CDC__K5629 | SAMN03945129 | GCF_002198195.1 | NonPC | Public | 2007 | USA | Clinical | 91.5 | 99.86 | 0.14 | 98.3115 | 5092943 | 45.27 | 74 | 217798 |
| SAMN03945130-GCF_002198275.1 | 2012V-1076 | SAMN03945130 | GCF_002198275.1 | NonPC | Public | 2012 | USA | Clinical | 91.4 | 99.86 | 0 | 98.2948 | 5090222 | 45.27 | 56 | 418031 |
| SAMN03945131-GCF_002198055.1 | 2012V-1132 | SAMN03945131 | GCF_002198055.1 | NonPC | Public | 2012 | USA | Clinical | 91.4 | 100 | 0.41 | 98.3109 | 5091707 | 45.27 | 70 | 331637 |
| SAMN03945132-GCF_002198075.1 | 2012V-1131 | SAMN03945132 | GCF_002198075.1 | NonPC | Public | 2012 | USA | Clinical | 91.5 | 100 | 0 | 98.3159 | 5093005 | 45.27 | 56 | 354544 |
| SAMN03945133-GCF_002198295.1 | 2012V-1109 | SAMN03945133 | GCF_002198295.1 | NonPC | Public | 2012 | USA | Clinical | 91.4 | 100 | 0.54 | 98.2865 | 5089969 | 45.27 | 87 | 258524 |
| SAMN03945134-GCF_002198335.1 | 2012V-1108 | SAMN03945134 | GCF_002198335.1 | NonPC | Public | 2012 | USA | Clinical | 91.4 | 100 | 0 | 98.3509 | 5090757 | 45.27 | 51 | 286856 |
| SAMN03945135-GCF_002198135.1 | 2012V-1121 | SAMN03945135 | GCF_002198135.1 | NonPC | Public | 2012 | USA | Clinical | 91.5 | 100 | 0 | 98.3067 | 5090206 | 45.27 | 65 | 267989 |
| SAMN03945136-GCF_002198175.1 | 2012V-1134 | SAMN03945136 | GCF_002198175.1 | NonPC | Public | 2012 | USA | Clinical | 91.4 | 100 | 0 | 98.3058 | 5088765 | 45.27 | 90 | 179123 |
| SAMN03945137-GCF_002198185.1 | 2012V-1103 | SAMN03945137 | GCF_002198185.1 | NonPC | Public | 2012 | USA | Clinical | 91.5 | 100 | 0 | 98.3362 | 5079992 | 45.28 | 37 | 561815 |
| SAMN03945138-GCF_002198305.1 | CDC__K4639-1 | SAMN03945138 | GCF_002198305.1 | NonPC | Public | 2006 | USA | Clinical | 91.4 | 99.73 | 0.27 | 98.2913 | 5088752 | 45.27 | 56 | 419059 |
| SAMN03945139-GCF_002198255.1 | CDC__AM47612 | SAMN03945139 | GCF_002198255.1 | NonPC | Public | 2011 | USA | Clinical | 91.4 | 100 | 0 | 98.3197 | 5078491 | 45.29 | 44 | 465477 |

| | | | | | | | | | | | | | | | | |
|---|---|---|---|---|---|---|---|---|---|---|---|---|---|---|---|---|
| SAMN03945140-GCF_002198375.1 | CDC__M12-106 | SAMN03945140 | GCF_002198375.1 | NonPC | Public | 2012 | USA | Clinical | 91.4 | 100 | 0 | 98.292 | 5090052 | 45.27 | 55 | 418153 |
| SAMN03945141-GCF_002198415.1 | AM51556 | SAMN03945141 | GCF_002198415.1 | NonPC | Public | 2012 | USA | Clinical | 91.4 | 100 | 0 | 98.3134 | 5089807 | 45.27 | 55 | 322763 |
| SAMN03945142-GCF_002198485.1 | AM51816 | SAMN03945142 | GCF_002198485.1 | NonPC | Public | 2012 | USA | Clinical | 91.4 | 99.86 | 0 | 98.3448 | 5129824 | 45.27 | 90 | 420016 |
| SAMN03945143-GCF_002198355.1 | AM51866 | SAMN03945143 | GCF_002198355.1 | NonPC | Public | 2012 | USA | Clinical | 91.5 | 99.86 | 0.27 | 98.3392 | 5077472 | 45.29 | 51 | 403545 |
| SAMN03945144-GCF_002198435.1 | AM51867 | SAMN03945144 | GCF_002198435.1 | NonPC | Public | 2012 | USA | Clinical | 91.4 | 100 | 0 | 98.3267 | 5079908 | 45.29 | 86 | 315302 |
| SAMN03945145-GCF_002198515.1 | CDC__A8962 | SAMN03945145 | GCF_002198515.1 | NonPC | Public | 2008 | USA | Clinical | 91.6 | 100 | 0 | 98.3213 | 5109512 | 45.25 | 86 | 157810 |
| SAMN03945146-GCF_002198395.1 | CDC__121898 | SAMN03945146 | GCF_002198395.1 | NonPC | Public | 2012 | USA | Clinical | 91.1 | 99.73 | 0.01 | 98.2929 | 5165041 | 45.29 | 180 | 100778 |
| SAMN03945147-GCF_002198525.1 | CDC__101325304 | SAMN03945147 | GCF_002198525.1 | NonPC | Public | 2012 | USA | Clinical | 91.4 | 99.86 | 0 | 98.3018 | 5086482 | 45.27 | 53 | 461832 |
| SAMN03945148-GCF_002198475.1 | CFSAN022330 | SAMN03945148 | GCF_002198475.1 | NonPC | Public | 2012 | Spain | Clinical | 91.7 | 100 | 0.41 | 98.2545 | 5185327 | 45.22 | 61 | 297386 |
| SAMN03945149-GCF_002198555.1 | CFSAN022331 | SAMN03945149 | GCF_002198555.1 | NonPC | Public | 2012 | Spain | Clinical | 91.8 | 100 | 0 | 98.2775 | 5185431 | 45.22 | 58 | 391044 |
| SAMN03945150-GCF_002198565.1 | CFSAN022332 | SAMN03945150 | GCF_002198565.1 | NonPC | Public | 2012 | Spain | Clinical | 91.8 | 100 | 0 | 98.2907 | 5188756 | 45.22 | 57 | 406461 |
| SAMN03945151-GCF_002198445.1 | CFSAN022334 | SAMN03945151 | GCF_002198445.1 | NonPC | Public | 2012 | Spain | Clinical | 91.7 | 99.86 | 0 | 98.2892 | 5180235 | 45.21 | 70 | 314813 |
| SAMN03945152-GCF_002198575.1 | CFSAN022335 | SAMN03945152 | GCF_002198575.1 | NonPC | Public | 2012 | Spain | Clinical | 91.8 | 100 | 0 | 98.2472 | 5186069 | 45.22 | 56 | 461627 |
| SAMN03945153-GCF_002198615.1 | CFSAN022336 | SAMN03945153 | GCF_002198615.1 | NonPC | Public | 2012 | Spain | Clinical | 91.7 | 100 | 0 | 98.2802 | 5181843 | 45.22 | 67 | 297386 |
| SAMN03993998-SRR4016792 | PNUSAV000009 | SAMN03993998 | SRR4016792 | NonPC | Public | NA | USA | Clinical | 91.2 | 100 | 0.31 | 98.3597 | 5021748 | 45.36 | 44 | 363327 |
| SAMN03996250-GCF_001558495.2 | ATCC_17802 | SAMN03996250 | GCF_001558495.2 | NonPC | Public | 1951 | Japan | Env | 91.7 | 100 | 0.05 | 98.393 | 5152461 | 45.33 | 2 | 3286802 |
| SAMN03996335-GCF_001471485.2 | FDA:FDAARGOS_120 | SAMN03996335 | GCF_001471485.2 | NonPC | Public | NA | USA | Clinical | 92.1 | 100 | 0.59 | 98.233 | 5332466 | 45.25 | 3 | 1907420 |
| SAMN04018860-SRR4016793 | PNUSAV000010 | SAMN04018860 | SRR4016793 | NonPC | Public | NA | USA | Clinical | 90.6 | 100 | 0.05 | 98.3112 | 4980219 | 45.35 | 47 | 525797 |
| SAMN04145915-GCF_001634135.1 | 281-09 | SAMN04145915 | GCF_001634135.1 | NonPC | Public | 2009 | Peru | Env | 92.9 | 100 | 0 | 98.4183 | 5129398 | 45.22 | 42 | 280966 |
| SAMN04145916-GCF_001634165.1 | 283-09 | SAMN04145916 | GCF_001634165.1 | NonPC | Public | 2009 | Peru | Clinical | 92.9 | 100 | 0 | 98.4157 | 5126292 | 45.22 | 49 | 313618 |
| SAMN04145917-GCF_001634105.1 | C220-09 | SAMN04145917 | GCF_001634105.1 | NonPC | Public | 2009 | Peru | Clinical | 92.9 | 100 | 0 | 98.4206 | 5179121 | 45.23 | 34 | 453287 |
| SAMN04145918-GCF_001634125.1 | C224-09 | SAMN04145918 | GCF_001634125.1 | NonPC | Public | 2009 | Peru | Clinical | 92.8 | 100 | 0 | 98.4016 | 5123197 | 45.22 | 47 | 404432 |
| SAMN04145919-GCF_001634185.1 | C226-09 | SAMN04145919 | GCF_001634185.1 | NonPC | Public | 2009 | Peru | Clinical | 93 | 100 | 0 | 98.4105 | 5130775 | 45.22 | 35 | 391859 |
| SAMN04145920-GCF_001634205.1 | C244-09 | SAMN04145920 | GCF_001634205.1 | NonPC | Public | 2009 | Peru | Clinical | 92.9 | 99.93 | 0 | 98.4034 | 5136853 | 45.22 | 35 | 453287 |
| SAMN04145921-GCF_001634215.1 | C235 | SAMN04145921 | GCF_001634215.1 | NonPC | Public | 2009 | Peru | Clinical | 92.9 | 100 | 0 | 98.4201 | 5126956 | 45.22 | 35 | 467870 |
| SAMN04145922-GCF_001633985.1 | CFSAN025059 | SAMN04145922 | GCF_001633985.1 | NonPC | Public | 2009 | Peru | Clinical | 92.9 | 100 | 0 | 98.3957 | 5126321 | 45.22 | 40 | 376159 |
| SAMN04145923-GCF_001634015.1 | C237 | SAMN04145923 | GCF_001634015.1 | NonPC | Public | 2009 | Peru | Clinical | 92.8 | 100 | 0 | 98.4055 | 5126855 | 45.22 | 52 | 265563 |
| SAMN04145924-GCF_001634245.1 | 239-09 | SAMN04145924 | GCF_001634245.1 | NonPC | Public | 2009 | Peru | Clinical | 92.9 | 100 | 0 | 98.4185 | 5129762 | 45.22 | 34 | 467872 |
| SAMN04145925-GCF_001633825.1 | 241-09 | SAMN04145925 | GCF_001633825.1 | NonPC | Public | 2009 | Peru | Clinical | 92.9 | 99.86 | 0 | 98.4151 | 5128953 | 45.22 | 35 | 467870 |
| SAMN04145926-GCF_001633845.1 | 245-09 | SAMN04145926 | GCF_001633845.1 | NonPC | Public | 2009 | Peru | Clinical | 92.9 | 100 | 0.01 | 98.3961 | 5220707 | 45.25 | 35 | 455995 |
| SAMN04145927-GCF_001633995.1 | CFSAN025064 | SAMN04145927 | GCF_001633995.1 | NonPC | Public | 2009 | Peru | Clinical | 92.9 | 100 | 0 | 98.3864 | 5130119 | 45.22 | 40 | 337873 |
| SAMN04145928-GCF_001633835.1 | CFSAN025065 | SAMN04145928 | GCF_001633835.1 | NonPC | Public | 2009 | Peru | Clinical | 92.9 | 100 | 0 | 98.396 | 5126976 | 45.22 | 44 | 284637 |
| SAMN04145929-GCF_001633855.1 | 285-09 | SAMN04145929 | GCF_001633855.1 | NonPC | Public | 2009 | Peru | Clinical | 93 | 100 | 0 | 98.4062 | 5132315 | 45.22 | 39 | 330290 |
| SAMN04145930-GCF_001633905.1 | 287-09 | SAMN04145930 | GCF_001633905.1 | NonPC | Public | 2009 | Peru | Clinical | 92.9 | 99.93 | 0 | 98.3991 | 5130983 | 45.22 | 45 | 366735 |
| SAMN04145931-GCF_001633915.1 | 379-09 | SAMN04145931 | GCF_001633915.1 | NonPC | Public | 2009 | Peru | Clinical | 92.8 | 100 | 0 | 98.3988 | 5125547 | 45.22 | 69 | 221959 |
| SAMN04145932-GCF_001633935.1 | CFSAN029653 | SAMN04145932 | GCF_001633935.1 | NonPC | Public | 2009 | Peru | Env | 92.5 | 100 | 0 | 98.3822 | 5122821 | 45.23 | 112 | 101266 |
| SAMN04145933-GCF_001633955.1 | CFSAN029654 | SAMN04145933 | GCF_001633955.1 | NonPC | Public | 2009 | Peru | Clinical | 92.8 | 100 | 0.27 | 98.4189 | 5126587 | 45.22 | 104 | 152415 |
| SAMN04145934-GCF_001634035.1 | CFSAN029656 | SAMN04145934 | GCF_001634035.1 | NonPC | Public | 2009 | Peru | Env | 92.8 | 100 | 0 | 98.3766 | 5127031 | 45.22 | 80 | 145779 |
| SAMN04229570-SRR4016795 | PNUSAV000011 | SAMN04229570 | SRR4016795 | NonPC | Public | NA | USA | Clinical | 91.5 | 100 | 0 | 98.3041 | 5096595 | 45.27 | 43 | 719200 |
| SAMN04229571-SRR4016797 | PNUSAV000012 | SAMN04229571 | SRR4016797 | NonPC | Public | NA | USA | Clinical | 89.9 | 100 | 0.2 | 98.2308 | 5494468 | 44.93 | 42 | 564236 |
| SAMN04229572-SRR4016798 | PNUSAV000013 | SAMN04229572 | SRR4016798 | NonPC | Public | NA | USA | Clinical | 91.5 | 100 | 0 | 98.2972 | 5096842 | 45.27 | 49 | 462160 |
| SAMN04241617-GCF_001879585.1 | FORC_022 | SAMN04241617 | GCF_001879585.1 | NonPC | Public | NA | South_Korea | Env | 92.7 | 98.77 | 0.05 | 98.351 | 5339691 | 45.25 | 3 | 3358191 |
| SAMN04252309-SRR4016799 | PNUSAV000014 | SAMN04252309 | SRR4016799 | NonPC | Public | NA | USA | Clinical | 91.5 | 100 | 0 | 98.2489 | 5107986 | 45.28 | 56 | 717452 |
| SAMN04252310-SRR4018049 | PNUSAV000015 | SAMN04252310 | SRR4018049 | NonPC | Public | NA | USA | Clinical | 89.8 | 100 | 0.04 | 98.2712 | 5162458 | 45.22 | 39 | 547107 |
| SAMN04285689-GCF_001887055.1 | FORC_018 | SAMN04285689 | GCF_001887055.1 | NonPC | Public | NA | South_Korea | Env | 91.9 | 100 | 0.08 | 98.2737 | 5045560 | 45.43 | 2 | 3273522 |
| SAMN04293498-GCF_001559885.1 | 09-4661 | SAMN04293498 | GCF_001559885.1 | NonPC | Public | 2009 | Canada | Clinical | 91 | 100 | 0.38 | 98.3148 | 5185731 | 45.18 | 66 | 403281 |
| SAMN04296855-GCF_001559895.1 | A1EZ919 | SAMN04296855 | GCF_001559895.1 | NonPC | Public | 2001 | Canada | Clinical | 91.6 | 100 | 0 | 98.2968 | 5094419 | 45.27 | 31 | 462093 |
| SAMN04304554-SRR4016801 | PNUSAV000016 | SAMN04304554 | SRR4016801 | NonPC | Public | NA | USA | Clinical | 89 | 100 | 0.4 | 98.2919 | 5222032 | 45.24 | 48 | 538026 |

| | | | | | | | | | | | | | | | | |
|---|---|---|---|---|---|---|---|---|---|---|---|---|---|---|---|---|
| SAMN04308162-GCF_001559805.1 | A4EZ700 | SAMN04308162 | GCF_001559805.1 | NonPC | Public | 2004 | Canada | Clinical | 89.3 | 100 | 0 | 98.0531 | 5127080 | 45.21 | 76 | 319036 |
| SAMN04320985-GCF_001584415.1 | A4EZ703 | SAMN04320985 | GCF_001584415.1 | NonPC | Public | 2004 | Canada | Clinical | 90.5 | 100 | 0.14 | 98.2693 | 5146304 | 45.23 | 34 | 525437 |
| SAMN04327442-GCF_001595575.1 | A4EZ724 | SAMN04327442 | GCF_001595575.1 | NonPC | Public | 2004 | Canada | Clinical | 89.4 | 100 | 0.27 | 98.1201 | 5058285 | 45.25 | 72 | 292301 |
| SAMN04327443-GCF_001595565.1 | A4EZ927 | SAMN04327443 | GCF_001595565.1 | PC | Public | 2004 | Canada | Clinical | 97.7 | 100 | 0 | 99.9475 | 5104589 | 45.27 | 48 | 373935 |
| SAMN04338417-SRR3002506 | NA | SAMN04338417 | SRR3002506 | PC | Public | 2013 | Chile | Clinical | 98.8 | 99.97 | 0 | 99.9921 | 5114426 | 45.3 | 29 | 543019 |
| SAMN04349747-GCF_001541615.1 | Gxfcg07_003_7 | SAMN04349747 | GCF_001541615.1 | PC | Public | 2007 | China | Clinical | 97.7 | 100 | 0 | 99.9081 | 5153303 | 45.29 | 49 | 323009 |
| SAMN04349756-GCF_001541625.1 | Gxnn09_048 | SAMN04349756 | GCF_001541625.1 | NonPC | Public | 2009 | China | Clinical | 92.3 | 100 | 0.19 | 98.3974 | 5134259 | 45.09 | 41 | 603826 |
| SAMN04358131-GCF_001609435.1 | 05-4792 | SAMN04358131 | GCF_001609435.1 | NonPC | Public | 2005 | Canada | Clinical | 90.8 | 100 | 0 | 98.3075 | 5140425 | 45.27 | 49 | 465886 |
| SAMN04358132-GCF_001609375.1 | C142 | SAMN04358132 | GCF_001609375.1 | NonPC | Public | 2008 | Canada | Clinical | 90.9 | 100 | 0.6 | 98.2919 | 5210898 | 45.22 | 52 | 334240 |
| SAMN04358133-GCF_001609395.1 | C143 | SAMN04358133 | GCF_001609395.1 | NonPC | Public | 2008 | Canada | Clinical | 91.5 | 100 | 0 | 98.3256 | 5078280 | 45.29 | 30 | 548226 |
| SAMN04358134-GCF_001609415.1 | C144 | SAMN04358134 | GCF_001609415.1 | NonPC | Public | 2008 | Canada | Clinical | 91.5 | 100 | 0 | 98.31 | 5078762 | 45.29 | 34 | 548500 |
| SAMN04358135-GCF_001609345.1 | C145 | SAMN04358135 | GCF_001609345.1 | NonPC | Public | 2008 | Canada | Clinical | 90.9 | 100 | 0.6 | 98.2794 | 5176962 | 45.19 | 47 | 403418 |
| SAMN04358136-GCF_001609445.1 | C146 | SAMN04358136 | GCF_001609445.1 | NonPC | Public | 2008 | Canada | Clinical | 90.2 | 100 | 0.05 | 98.2973 | 5212184 | 45.18 | 51 | 404244 |
| SAMN04358199-GCF_001609495.1 | C147 | SAMN04358199 | GCF_001609495.1 | NonPC | Public | 2008 | Canada | Clinical | 91.5 | 100 | 0 | 98.3032 | 5078771 | 45.29 | 36 | 521092 |
| SAMN04358200-GCF_001609475.1 | C148 | SAMN04358200 | GCF_001609475.1 | NonPC | Public | 2008 | Canada | Clinical | 89.4 | 100 | 0 | 98.0855 | 5065366 | 45.25 | 80 | 146049 |
| SAMN04358201-GCF_001601695.1 | C150 | SAMN04358201 | GCF_001601695.1 | NonPC | Public | 2008 | Canada | Clinical | 90.9 | 100 | 0.6 | 98.2863 | 5209864 | 45.22 | 53 | 334486 |
| SAMN04377087-GCF_001609555.1 | 04-2192 | SAMN04377087 | GCF_001609555.1 | NonPC | Public | 2004 | Canada | Clinical | 90 | 100 | 0 | 98.2156 | 5245730 | 45.19 | 35 | 464267 |
| SAMN04377152-GCF_001609575.1 | 09-4666 | SAMN04377152 | GCF_001609575.1 | NonPC | Public | 2009 | Canada | Clinical | 90.9 | 100 | 0.33 | 98.2863 | 5172328 | 45.18 | 44 | 425238 |
| SAMN04377384-GCF_001609595.1 | A5Z273 | SAMN04377384 | GCF_001609595.1 | PC | Public | 2005 | Canada | Clinical | 98.2 | 99.73 | 0 | 99.9854 | 5098054 | 45.28 | 44 | 373921 |
| SAMN04377387-GCF_001609515.1 | A5Z652 | SAMN04377387 | GCF_001609515.1 | NonPC | Public | 2005 | Canada | Clinical | 91.4 | 100 | 0 | 98.3167 | 5082632 | 45.28 | 32 | 556943 |
| SAMN04377388-GCF_001609525.1 | A5Z853 | SAMN04377388 | GCF_001609525.1 | PC | Public | 2005 | Canada | Clinical | 98.4 | 100 | 0.27 | 99.9897 | 5142950 | 45.27 | 41 | 427483 |
| SAMN04377550-GCF_001609615.1 | A5Z860 | SAMN04377550 | GCF_001609615.1 | NonPC | Public | 2005 | Canada | Clinical | 89.2 | 100 | 0 | 98.0939 | 5094605 | 45.25 | 68 | 319006 |
| SAMN04377551-GCF_001609695.1 | A5Z878 | SAMN04377551 | GCF_001609695.1 | NonPC | Public | 2005 | Canada | Clinical | 91.5 | 100 | 0 | 98.3298 | 5079776 | 45.29 | 36 | 548591 |
| SAMN04377552-GCF_001609635.1 | A5Z905 | SAMN04377552 | GCF_001609635.1 | NonPC | Public | 2005 | Canada | Clinical | 91.5 | 100 | 0 | 98.2513 | 5226793 | 45.2 | 31 | 570484 |
| SAMN04377553-GCF_001609675.1 | A5Z924 | SAMN04377553 | GCF_001609675.1 | NonPC | Public | 2005 | Canada | Clinical | 91.4 | 100 | 0 | 98.3152 | 5076261 | 45.29 | 38 | 376542 |
| SAMN04378285-GCF_001609655.1 | C140 | SAMN04378285 | GCF_001609655.1 | NonPC | Public | 2008 | Canada | Clinical | 93 | 100 | 0.05 | 98.3411 | 5046146 | 45.32 | 29 | 636695 |
| SAMN04382997-GCF_001609715.1 | A4EZ964 | SAMN04382997 | GCF_001609715.1 | NonPC | Public | 2004 | Canada | Clinical | 91.9 | 100 | 0 | 98.25 | 5101569 | 45.28 | 49 | 407937 |
| SAMN04407163-SRR4016804 | PNUSAV000017 | SAMN04407163 | SRR4016804 | NonPC | Public | NA | USA | Clinical | 89.7 | 100 | 0 | 98.24 | 5186911 | 45.22 | 61 | 266237 |
| SAMN04417569-GCF_001610595.1 | 04-2550 | SAMN04417569 | GCF_001610595.1 | NonPC | Public | 2004 | Canada | Clinical | 91.5 | 100 | 0 | 98.2799 | 5099458 | 45.29 | 40 | 537513 |
| SAMN04417570-GCF_001608505.1 | 05-3133 | SAMN04417570 | GCF_001608505.1 | NonPC | Public | 2005 | Canada | Clinical | 89.3 | 100 | 0 | 98.0693 | 5022875 | 45.28 | 64 | 292212 |
| SAMN04417571-GCF_001608525.1 | 09-4436 | SAMN04417571 | GCF_001608525.1 | NonPC | Public | 2009 | Canada | Clinical | 89.7 | 99.95 | 0.04 | 98.2916 | 5151693 | 45.21 | 41 | 431727 |
| SAMN04422010-GCF_001608575.1 | 09-4665 | SAMN04422010 | GCF_001608575.1 | NonPC | Public | 2009 | Canada | Clinical | 90.9 | 100 | 0.33 | 98.2948 | 5169946 | 45.2 | 55 | 307466 |
| SAMN04422011-GCF_001608655.1 | A0EZ608 | SAMN04422011 | GCF_001608655.1 | NonPC | Public | 2000 | Canada | Clinical | 91.6 | 100 | 0 | 98.2923 | 5096832 | 45.27 | 28 | 548505 |
| SAMN04422012-GCF_001608605.1 | A0EZ664 | SAMN04422012 | GCF_001608605.1 | NonPC | Public | 2000 | Canada | Clinical | 91.1 | 100 | 0.05 | 98.2556 | 5151804 | 45.21 | 37 | 561155 |
| SAMN04422014-GCF_001608705.1 | A0EZ713 | SAMN04422014 | GCF_001608705.1 | NonPC | Public | 2000 | Canada | Clinical | 91.1 | 100 | 0.05 | 98.2452 | 5151966 | 45.21 | 42 | 461183 |
| SAMN04422016-GCF_001608735.1 | A2EZ614 | SAMN04422016 | GCF_001608735.1 | NonPC | Public | 2002 | Canada | Clinical | 89.3 | 100 | 0 | 98.0651 | 5065489 | 45.25 | 67 | 254300 |
| SAMN04422017-GCF_001608675.1 | A2EZ715 | SAMN04422017 | GCF_001608675.1 | NonPC | Public | 2002 | Canada | Clinical | 91.6 | 100 | 0 | 98.3188 | 5103078 | 45.27 | 31 | 548679 |
| SAMN04422023-GCF_001608765.1 | A2EZ743 | SAMN04422023 | GCF_001608765.1 | NonPC | Public | 2002 | Canada | Clinical | 90.4 | 100 | 0.14 | 98.2416 | 5063270 | 45.18 | 34 | 526416 |
| SAMN04422047-GCF_001608785.1 | A3EZ136 | SAMN04422047 | GCF_001608785.1 | PC | Public | 2003 | Canada | Clinical | 98.2 | 100 | 0 | 99.9839 | 5089443 | 45.3 | 45 | 373928 |
| SAMN04422049-GCF_001608755.1 | A5Z1022 | SAMN04422049 | GCF_001608755.1 | NonPC | Public | 2005 | Canada | Clinical | 91.3 | 100 | 0 | 98.3259 | 5128457 | 45.25 | 35 | 443357 |
| SAMN04422051-GCF_001608685.1 | F4395 | SAMN04422051 | GCF_001608685.1 | NonPC | Public | 2006 | Canada | Clinical | 91.5 | 100 | 0 | 98.3289 | 5134947 | 45.27 | 34 | 548500 |
| SAMN04422052-GCF_001608815.1 | F30368 | SAMN04422052 | GCF_001608815.1 | NonPC | Public | 2006 | Canada | Clinical | 91.9 | 100 | 0 | 98.4377 | 5090449 | 45.25 | 30 | 524972 |
| SAMN04422054-GCF_001608595.1 | H11523 | SAMN04422054 | GCF_001608595.1 | NonPC | Public | 2006 | Canada | Clinical | 91.5 | 100 | 0 | 98.2726 | 5077113 | 45.29 | 34 | 548500 |
| SAMN04422055-GCF_001608835.1 | F63267 | SAMN04422055 | GCF_001608835.1 | PC | Public | 2006 | Canada | Clinical | 98.4 | 100 | 0 | 99.9869 | 5121492 | 45.29 | 47 | 427352 |
| SAMN04422063-GCF_001608855.1 | W501 | SAMN04422063 | GCF_001608855.1 | NonPC | Public | 2006 | Canada | Clinical | 88.6 | 100 | 0.05 | 98.283 | 4963903 | 45.31 | 40 | 509565 |
| SAMN04422093-GCF_001608865.1 | T8994 | SAMN04422093 | GCF_001608865.1 | NonPC | Public | 2006 | Canada | Clinical | 91.4 | 100 | 0 | 98.28 | 5076710 | 45.29 | 34 | 548500 |
| SAMN04422094-GCF_001608615.1 | H64024 | SAMN04422094 | GCF_001608615.1 | NonPC | Public | 2006 | Canada | Clinical | 91.5 | 100 | 0 | 98.3194 | 5080527 | 45.29 | 33 | 548500 |
| SAMN04422095-GCF_001608545.1 | M59787 | SAMN04422095 | GCF_001608545.1 | NonPC | Public | 2006 | Canada | Clinical | 91.4 | 100 | 0 | 98.3338 | 5076618 | 45.29 | 34 | 462077 |

| | | | | | | | | | | | | | | | | |
|---|---|---|---|---|---|---|---|---|---|---|---|---|---|---|---|---|
| SAMN04441010-GCF_001608945.1 | 07-2964 | SAMN04441010 | GCF_001608945.1 | NonPC | Public | 2007 | Canada | Clinical | 92.8 | 100 | 0 | 98.4123 | 5120702 | 45.2 | 32 | 415597 |
| SAMN04441011-GCF_001609025.1 | 09-1772 | SAMN04441011 | GCF_001609025.1 | NonPC | Public | 2009 | Canada | Clinical | 90.9 | 99.93 | 0.33 | 98.3125 | 5173304 | 45.18 | 53 | 315980 |
| SAMN04441012-GCF_001608935.1 | 09-3219 | SAMN04441012 | GCF_001608935.1 | NonPC | Public | 2009 | Canada | Clinical | 91.5 | 100 | 0 | 98.3241 | 5074950 | 45.28 | 38 | 462063 |
| SAMN04441013-GCF_001609015.1 | 09-4662 | SAMN04441013 | GCF_001609015.1 | NonPC | Public | 2009 | Canada | Clinical | 90.9 | 100 | 0.6 | 98.2729 | 5176120 | 45.18 | 47 | 315437 |
| SAMN04441014-GCF_001609055.1 | A0EZ383 | SAMN04441014 | GCF_001609055.1 | NonPC | Public | 2000 | Canada | Clinical | 91.5 | 100 | 0.17 | 98.3186 | 5158965 | 45.27 | 37 | 475179 |
| SAMN04441015-GCF_001609075.1 | A1EZ679 | SAMN04441015 | GCF_001609075.1 | NonPC | Public | 2001 | Canada | Clinical | 91.6 | 100 | 0 | 98.2881 | 5110204 | 45.27 | 31 | 548505 |
| SAMN04441016-GCF_001609095.1 | A2EZ523 | SAMN04441016 | GCF_001609095.1 | NonPC | Public | 2002 | Canada | Clinical | 91.5 | 100 | 0 | 98.2871 | 5093534 | 45.27 | 33 | 462105 |
| SAMN04441017-GCF_001609105.1 | A3EZ634 | SAMN04441017 | GCF_001609105.1 | NonPC | Public | 2003 | Canada | Clinical | 91.1 | 100 | 0.05 | 98.2162 | 5147837 | 45.2 | 41 | 461307 |
| SAMN04441018-GCF_001608995.1 | A3EZ710 | SAMN04441018 | GCF_001608995.1 | NonPC | Public | 2003 | Canada | Clinical | 89.2 | 100 | 0 | 98.0443 | 5122387 | 45.21 | 77 | 319006 |
| SAMN04441020-GCF_001608975.1 | A3EZ711 | SAMN04441020 | GCF_001608975.1 | NonPC | Public | 2003 | Canada | Clinical | 89.3 | 100 | 0 | 98.054 | 5127994 | 45.21 | 72 | 319035 |
| SAMN04441021-GCF_001609135.1 | A3EZ770 | SAMN04441021 | GCF_001609135.1 | NonPC | Public | 2003 | Canada | Clinical | 91.1 | 100 | 0.05 | 98.2787 | 5177526 | 45.18 | 47 | 542663 |
| SAMN04441022-GCF_001609155.1 | A3EZ799 | SAMN04441022 | GCF_001609155.1 | NonPC | Public | 2003 | Canada | Clinical | 89.2 | 100 | 0 | 98.0888 | 5044575 | 45.26 | 73 | 292206 |
| SAMN04441023-GCF_001609175.1 | A3EZ936 | SAMN04441023 | GCF_001609175.1 | NonPC | Public | 2003 | Canada | Clinical | 90.2 | 100 | 0.19 | 98.281 | 5245252 | 45.18 | 51 | 387604 |
| SAMN04441024-GCF_001609185.1 | A1EZ952 | SAMN04441024 | GCF_001609185.1 | NonPC | Public | 2001 | Canada | Clinical | 89.4 | 100 | 0 | 98.0651 | 5066903 | 45.25 | 65 | 254552 |
| SAMN04441025-GCF_001608915.1 | F1419 | SAMN04441025 | GCF_001608915.1 | NonPC | Public | 2006 | Canada | Clinical | 89.1 | 99.95 | 0 | 98.1104 | 5031127 | 45.29 | 64 | 244430 |
| SAMN04441026-GCF_001608895.1 | H18983 | SAMN04441026 | GCF_001608895.1 | NonPC | Public | 2006 | Canada | Clinical | 91.5 | 100 | 0 | 98.282 | 5085869 | 45.29 | 48 | 548500 |
| SAMN04875528-GCF_002073775.2 | FDA:FDAARGOS_191 | SAMN04875528 | GCF_002073775.2 | PC | Public | 1996 | India | Clinical | 100 | 100 | 0 | 99.9955 | 5173292 | 45.39 | 2 | 3295711 |
| SAMN05161900-GCF_013420895.1 | v70 | SAMN05161900 | GCF_013420895.1 | NonPC | Public | 2009 | Malaysia | NA | 91.2 | 100 | 0.26 | 98.3955 | 5277960 | 45.04 | 34 | 660363 |
| SAMN05163299-SRR4018050 | PNUSAV000018 | SAMN05163299 | SRR4018050 | NonPC | Public | NA | USA | Clinical | 92.3 | 100 | 0.05 | 98.3237 | 5139903 | 45.23 | 49 | 450060 |
| SAMN05194681-GCF_001856035.1 | 06-043 | SAMN05194681 | GCF_001856035.1 | NonPC | Public | 2006 | Mozambique | Env | 91 | 100 | 0 | 98.3717 | 5320621 | 45.07 | 54 | 330334 |
| SAMN05194682-SRR3624770 | 920 | SAMN05194682 | SRR3624770 | PC | Public | 2007 | USA | Env | 98.4 | 100 | 0 | 99.9953 | 5098408 | 45.35 | 47 | 505937 |
| SAMN05194683-SRR3624774 | 752 | SAMN05194683 | SRR3624774 | PC | Public | 2007 | USA | Env | 98.7 | 100 | 0 | 99.9776 | 5163109 | 45.29 | 48 | 505937 |
| SAMN05195061-SRR3624775 | 783 | SAMN05195061 | SRR3624775 | PC | Public | 2007 | USA | Env | 98.7 | 100 | 0 | 99.977 | 5163244 | 45.29 | 50 | 505937 |
| SAMN05195070-SRR3628261 | 658 | SAMN05195070 | SRR3628261 | PC | Public | 2007 | USA | Env | 98.7 | 100 | 0 | 99.9807 | 5162104 | 45.29 | 48 | 542987 |
| SAMN05195071-SRR3624776 | 805 | SAMN05195071 | SRR3624776 | NonPC | Public | 2007 | USA | Env | 91.2 | 100 | 0.08 | 98.2642 | 5333617 | 45.19 | 43 | 381728 |
| SAMN05195118-SRR3628260 | 3328 | SAMN05195118 | SRR3628260 | NonPC | Public | 2007 | USA | Clinical | 90.1 | 100 | 0 | 98.3709 | 5186846 | 45.21 | 25 | 547955 |
| SAMN05195119-SRR3628270 | 27 | SAMN05195119 | SRR3628270 | NonPC | Public | 2007 | USA | Env | 90.4 | 100 | 0.14 | 98.2048 | 5069628 | 45.18 | 31 | 874371 |
| SAMN05195120-SRR3628271 | 765 | SAMN05195120 | SRR3628271 | PC | Public | 2007 | USA | Env | 98.7 | 100 | 0.13 | 99.9782 | 5161096 | 45.29 | 46 | 401783 |
| SAMN05195240-GCF_001707995.1 | KVp10 | SAMN05195240 | GCF_001707995.1 | NonPC | Public | 2007 | Sweden | Env | 88.1 | 98.64 | 0.01 | 98.0991 | 5008534 | 45.36 | 57 | 237565 |
| SAMN05216829-SRR3649460 | In711 | SAMN05216829 | SRR3649460 | NonPC | Public | 2007 | India | Env | 91.7 | 100 | 0.09 | 98.3599 | 5119135 | 45.28 | 20 | 1387433 |
| SAMN05220835-SRR3655232 | 4635 | SAMN05220835 | SRR3655232 | PC | Public | 2007 | USA | Clinical | 98.6 | 100 | 0 | 99.9935 | 5099666 | 45.29 | 43 | 505937 |
| SAMN05220836-SRR3655233 | 4610 | SAMN05220836 | SRR3655233 | PC | Public | 2007 | USA | Clinical | 98.1 | 100 | 0 | 99.9305 | 5202658 | 45.11 | 50 | 433150 |
| SAMN05220837-SRR3655243 | 4703 | SAMN05220837 | SRR3655243 | PC | Public | 2007 | USA | Clinical | 98.5 | 100 | 0 | 99.9913 | 5114996 | 45.29 | 44 | 543011 |
| SAMN05271621-GCA_015806495.1 | PNUSAV000019 | SAMN05271621 | GCA_015806495.1 | NonPC | Public | NA | USA | Clinical | 91.2 | 100 | 0.82 | 98.2978 | 5144143 | 45.24 | 103 | 107423 |
| SAMN05301360-GCF_002144515.1 | 04-2548_1 | SAMN05301360 | GCF_002144515.1 | NonPC | Public | NA | Canada | Clinical | 91.4 | 100 | 0 | 98.3041 | 5457389 | 45.07 | 66 | 233416 |
| SAMN05301458-GCF_002144535.1 | 04-2548_100 | SAMN05301458 | GCF_002144535.1 | NonPC | Public | NA | Canada | Clinical | 91.5 | 100 | 0 | 98.2571 | 5464342 | 45.08 | 72 | 231483 |
| SAMN05301460-GCF_002144575.1 | 09-3216_1 | SAMN05301460 | GCF_002144575.1 | NonPC | Public | NA | Canada | Clinical | 91.4 | 100 | 0 | 98.3066 | 5078959 | 45.28 | 35 | 419836 |
| SAMN05301461-GCF_002144555.1 | 09-3216_100 | SAMN05301461 | GCF_002144555.1 | NonPC | Public | NA | Canada | Clinical | 91.4 | 100 | 0 | 98.3222 | 5191233 | 45.19 | 30 | 562116 |
| SAMN05301462-GCF_002144585.1 | Day_1_HS-13-1 | SAMN05301462 | GCF_002144585.1 | NonPC | Public | NA | NA | Env | 91.7 | 100 | 0 | 98.4199 | 5070397 | 45.26 | 40 | 401451 |
| SAMN05301509-GCF_002144625.1 | Day_1_HS-06-05 | SAMN05301509 | GCF_002144625.1 | NonPC | Public | NA | Canada | Env | 91.8 | 100 | 0.08 | 98.3411 | 5052979 | 45.37 | 41 | 331056 |
| SAMN05386387-GCF_001856105.1 | CAIM_1772 | SAMN05386387 | GCF_001856105.1 | NonPC | Public | 2005 | Mexico | Env | 89.4 | 100 | 0.71 | 98.307 | 5124623 | 45.22 | 112 | 136711 |
| SAMN05442492-GCF_001704915.1 | R10B2_71 | SAMN05442492 | GCF_001704915.1 | NonPC | Public | 1997 | USA | Env | 88.9 | 100 | 3.25 | 98.0592 | 5147063 | 45.18 | 145 | 131833 |
| SAMN05557942-SRR4018052 | PNUSAV000020 | SAMN05557942 | SRR4018052 | NonPC | Public | NA | USA | Clinical | 90.3 | 100 | 0 | 98.343 | 5006561 | 45.33 | 48 | 365656 |
| SAMN05557943-SRR4018053 | PNUSAV000021 | SAMN05557943 | SRR4018053 | NonPC | Public | NA | USA | Clinical | 89.9 | 100 | 0.04 | 98.2888 | 5175469 | 45.21 | 37 | 547113 |
| SAMN05570712-GCA_015818155.1 | PRJNA245882 | SAMN05570712 | GCA_015818155.1 | NonPC | Public | 2013 | USA | Clinical | 89.9 | 100 | 0.04 | 98.244 | 5192244 | 45.2 | 54 | 404314 |
| SAMN05570713-GCA_015819155.1 | CFSAN026734 | SAMN05570713 | GCA_015819155.1 | NonPC | Public | 2014 | USA | Clinical | 89.8 | 100 | 0.05 | 98.2652 | 5258918 | 45.24 | 50 | 433461 |
| SAMN05578009-GCF_002209595.1 | MAVP-A | SAMN05578009 | GCF_002209595.1 | NonPC | Public | 2010 | NA | Clinical | 89.8 | 100 | 0.04 | 98.2855 | 5258924 | 45.15 | 97 | 125700 |
| SAMN05578010-GCF_002209665.1 | MAVP-L | SAMN05578010 | GCF_002209665.1 | NonPC | Public | 2010 | NA | Clinical | 89.9 | 100 | 0.04 | 98.2514 | 5165475 | 45.21 | 38 | 513108 |

| | | | | | | | | | | | | | | | | |
|---|---|---|---|---|---|---|---|---|---|---|---|---|---|---|---|---|
| SAMN05578011-GCF_002209605.1 | MAVP-P | SAMN05578011 | GCF_002209605.1 | NonPC | Public | 2010 | NA | Clinical | 89.9 | 100 | 0.04 | 98.3079 | 5175068 | 45.22 | 41 | 388871 |
| SAMN05578012-GCF_002209625.1 | MAVP-T | SAMN05578012 | GCF_002209625.1 | NonPC | Public | 2010 | NA | Clinical | 89.8 | 100 | 0.04 | 98.2713 | 5161617 | 45.22 | 34 | 472965 |
| SAMN05578013-GCF_002209675.1 | MAVP4 | SAMN05578013 | GCF_002209675.1 | NonPC | Public | 2013 | NA | Clinical | 89.8 | 99.97 | 0.1 | 98.2879 | 5211934 | 45.17 | 28 | 773710 |
| SAMN05578014-GCF_002209765.1 | MAVP30 | SAMN05578014 | GCF_002209765.1 | NonPC | Public | 2013 | NA | Clinical | 89.9 | 100 | 0.04 | 98.2562 | 5171230 | 45.21 | 65 | 384922 |
| SAMN05578015-GCF_002209685.1 | MAVP39 | SAMN05578015 | GCF_002209685.1 | NonPC | Public | 2013 | NA | Clinical | 89.9 | 100 | 0.04 | 98.2303 | 5175449 | 45.21 | 73 | 315088 |
| SAMN05578016-GCF_002209735.1 | MAVP56 | SAMN05578016 | GCF_002209735.1 | NonPC | Public | 2013 | NA | Clinical | 90 | 100 | 0.26 | 98.2305 | 5468369 | 44.99 | 84 | 245909 |
| SAMN05578017-GCF_002209775.1 | MAVP74 | SAMN05578017 | GCF_002209775.1 | NonPC | Public | 2014 | NA | Clinical | 89.8 | 100 | 0.04 | 98.238 | 5174347 | 45.21 | 35 | 579011 |
| SAMN05578018-GCF_002209785.1 | MAVP75 | SAMN05578018 | GCF_002209785.1 | NonPC | Public | 2014 | NA | Clinical | 89.8 | 99.86 | 0.04 | 98.2587 | 5170491 | 45.21 | 44 | 341999 |
| SAMN05578019-GCF_002210265.1 | MAVP78 | SAMN05578019 | GCF_002210265.1 | NonPC | Public | 2014 | NA | Clinical | 89.9 | 100 | 0.04 | 98.2428 | 5172720 | 45.21 | 37 | 438828 |
| SAMN05578020-GCA_015780155.1 | MAVP90 | SAMN05578020 | GCA_015780155.1 | NonPC | Public | 2015 | NA | Clinical | 89.4 | 100 | 0.58 | 98.2467 | 5152277 | 45.25 | 158 | 71964 |
| SAMN05578021-GCA_015779665.1 | MAVP94 | SAMN05578021 | GCA_015779665.1 | NonPC | Public | 2015 | NA | Clinical | 89.5 | 100 | 0.38 | 98.2518 | 5158801 | 45.23 | 144 | 77052 |
| SAMN05578022-GCA_015779895.1 | MAVP109 | SAMN05578022 | GCA_015779895.1 | NonPC | Public | 2015 | NA | Clinical | 89.5 | 100 | 0.61 | 98.2485 | 5208324 | 45.18 | 131 | 86677 |
| SAMN05578023-GCA_015779705.1 | MAVP112 | SAMN05578023 | GCA_015779705.1 | NonPC | Public | 2015 | NA | Clinical | 90.9 | 99.73 | 0.27 | 98.3326 | 5065937 | 45.33 | 252 | 44251 |
| SAMN05579852-GCF_002209725.2 | MAVP-Q | SAMN05579852 | GCF_002209725.2 | NonPC | Public | 2011 | NA | Clinical | 90.8 | 100 | 0.04 | 98.2408 | 5257294 | 45.29 | 2 | 3369175 |
| SAMN05591502-GCF_001720365.1 | Klin | SAMN05591502 | GCF_001720365.1 | PC | Public | 2010 | Sweden | Clinical | 95.7 | 100 | 0 | 99.8885 | 5104502 | 45.29 | 56 | 260007 |
| SAMN05713723-GCA_015818515.1 | PNUSAV000022 | SAMN05713723 | GCA_015818515.1 | NonPC | Public | NA | USA | Clinical | 90 | 100 | 0.31 | 98.312 | 5155652 | 45.27 | 56 | 466593 |
| SAMN05713797-GCF_002221395.1 | CTVP27C | SAMN05713797 | GCF_002221395.1 | NonPC | Public | 2013 | NA | Clinical | 88.1 | 100 | 0.04 | 98.2564 | 5256712 | 45.21 | 34 | 478671 |
| SAMN05713798-GCF_002221385.1 | CTVP31C | SAMN05713798 | GCF_002221385.1 | NonPC | Public | 2013 | NA | Clinical | 89.9 | 100 | 0.04 | 98.2633 | 5273069 | 45.19 | 42 | 388605 |
| SAMN05713799-GCF_002221365.1 | CTVP34C | SAMN05713799 | GCF_002221365.1 | NonPC | Public | 2013 | NA | Clinical | 89.8 | 100 | 0.04 | 98.2717 | 5228251 | 45.19 | 44 | 328253 |
| SAMN05713800-GCF_002221345.1 | MEVP12 | SAMN05713800 | GCF_002221345.1 | NonPC | Public | 2015 | NA | Clinical | 88.2 | 100 | 0.04 | 98.2628 | 5166141 | 45.22 | 50 | 277090 |
| SAMN05713801-GCF_002221325.1 | MEVP14 | SAMN05713801 | GCF_002221325.1 | NonPC | Public | 2015 | NA | Clinical | 90.4 | 100 | 0.15 | 98.1675 | 5124877 | 45.22 | 143 | 344839 |
| SAMN05727869-GCF_001901565.1 | CICESE-170 | SAMN05727869 | GCF_001901565.1 | PC | Public | 1998 | Mexico | Clinical | 96.1 | 98.35 | 1.5 | 99.9687 | 5016302 | 45.31 | 663 | 12547 |
| SAMN05791359-SRR4290004 | PNUSAV000027 | SAMN05791359 | SRR4290004 | NonPC | Public | NA | USA | Clinical | 91.5 | 100 | 0 | 98.303 | 5132126 | 45.24 | 26 | 845755 |
| SAMN05791360-SRR4290003 | PNUSAV000026 | SAMN05791360 | SRR4290003 | NonPC | Public | NA | USA | Clinical | 89.9 | 100 | 0.04 | 98.2176 | 5243567 | 45.18 | 36 | 782140 |
| SAMN05791361-SRR4290002 | PNUSAV000025 | SAMN05791361 | SRR4290002 | NonPC | Public | NA | USA | Clinical | 91.2 | 100 | 0 | 98.3859 | 5100270 | 45.23 | 36 | 387382 |
| SAMN05791362-SRR4290001 | PNUSAV000024 | SAMN05791362 | SRR4290001 | NonPC | Public | NA | USA | Clinical | 90 | 100 | 0.04 | 98.2281 | 5243906 | 45.18 | 35 | 782140 |
| SAMN05791363-SRR4290006 | PNUSAV000028 | SAMN05791363 | SRR4290006 | NonPC | Public | NA | USA | Clinical | 90 | 100 | 0.04 | 98.2343 | 5242761 | 45.18 | 38 | 438141 |
| SAMN05804984-GCA_015818495.1 | PNUSAV000023 | SAMN05804984 | GCA_015818495.1 | NonPC | Public | NA | USA | Clinical | 90.3 | 100 | 0 | 98.3194 | 4964065 | 45.3 | 71 | 259107 |
| SAMN05845065-GCA_015818535.1 | PNUSAV000029 | SAMN05845065 | GCA_015818535.1 | NonPC | Public | NA | USA | Clinical | 90.9 | 100 | 0.33 | 98.3132 | 5189824 | 45.21 | 112 | 101012 |
| SAMN05858272-GCF_002018565.1 | PMC53.7 | SAMN05858272 | GCF_002018565.1 | NonPC | Public | 2007 | Chile | NA | 91.6 | 100 | 0.35 | 98.251 | 5097280 | 45.34 | 27 | 543576 |
| SAMN05858273-GCF_002018625.1 | PMA37.5 | SAMN05858273 | GCF_002018625.1 | PC | Public | 2005 | Chile | Clinical | 98.8 | 99.86 | 0 | 99.9709 | 5130113 | 45.29 | 48 | 327973 |
| SAMN05858274-GCF_002018645.1 | PMA54.13 | SAMN05858274 | GCF_002018645.1 | NonPC | Public | 2013 | Chile | Env | 91.6 | 100 | 0.03 | 98.2764 | 5132501 | 45.27 | 28 | 537826 |
| SAMN05858275-GCF_002018635.1 | PMA11.14 | SAMN05858275 | GCF_002018635.1 | NonPC | Public | 2014 | Chile | Env | 90 | 100 | 0 | 98.259 | 5131642 | 45.29 | 25 | 474599 |
| SAMN05858276-GCF_002018735.1 | PMA12.14 | SAMN05858276 | GCF_002018735.1 | NonPC | Public | 2014 | Chile | Env | 89.4 | 100 | 1.15 | 98.2933 | 5246644 | 45.16 | 24 | 1208324 |
| SAMN05858277-GCF_002018725.1 | PMA14.14 | SAMN05858277 | GCF_002018725.1 | NonPC | Public | 2014 | Chile | Env | 88.8 | 97 | 0.17 | 98.2643 | 5176280 | 45.08 | 30 | 521587 |
| SAMN05858278-GCF_002018705.1 | PMA21.14 | SAMN05858278 | GCF_002018705.1 | NonPC | Public | 2014 | Chile | Env | 90.2 | 100 | 0 | 98.3237 | 5140414 | 45.32 | 25 | 441483 |
| SAMN05858279-GCF_002018775.1 | PMA31.14 | SAMN05858279 | GCF_002018775.1 | NonPC | Public | 2014 | Chile | Env | 90 | 100 | 0 | 98.2868 | 5132441 | 45.31 | 22 | 474182 |
| SAMN05858280-GCF_002018695.1 | PMA32.14 | SAMN05858280 | GCF_002018695.1 | NonPC | Public | 2014 | Chile | Env | 90.2 | 100 | 0 | 98.2711 | 5146683 | 45.32 | 24 | 707727 |
| SAMN05858281-GCA_002018615.1 | PMA1.15 | SAMN05858281 | GCA_002018615.1 | NonPC | Public | 2015 | Chile | Env | 89.3 | 99.68 | 0.44 | 98.2359 | 5231763 | 45.17 | 130 | 77968 |
| SAMN05858282-GCF_002018535.1 | PMA2.15 | SAMN05858282 | GCF_002018535.1 | NonPC | Public | 2015 | Chile | Env | 91.7 | 98.18 | 1 | 98.4052 | 4967963 | 45.35 | 209 | 42215 |
| SAMN05858283-GCF_002018555.1 | PMA3.15 | SAMN05858283 | GCF_002018555.1 | NonPC | Public | 2015 | Chile | Env | 91 | 99.39 | 0 | 98.3057 | 5035150 | 45.32 | 80 | 137938 |
| SAMN05890675-GCF_003408895.1 | L70 | SAMN05890675 | GCF_003408895.1 | PC | Public | 2016 | China | NA | 97 | 100 | 0.01 | 99.9879 | 5107316 | 45.27 | 92 | 152401 |
| SAMN05895041-GCA_015818335.1 | PNUSAV000030 | SAMN05895041 | GCA_015818335.1 | NonPC | Public | NA | USA | Clinical | 91.5 | 100 | 0.08 | 98.3163 | 5058289 | 45.28 | 59 | 383211 |
| SAMN05895042-GCA_015818595.1 | PNUSAV000031 | SAMN05895042 | GCA_015818595.1 | NonPC | Public | NA | USA | Clinical | 87.4 | 100 | 0.38 | 98.1362 | 5042963 | 45.22 | 109 | 144775 |
| SAMN05895043-GCA_015818415.1 | PNUSAV000032 | SAMN05895043 | GCA_015818415.1 | NonPC | Public | NA | USA | Clinical | 88.7 | 100 | 0.11 | 98.2061 | 5188251 | 45.36 | 94 | 315668 |
| SAMN05895044-GCA_015818375.1 | PNUSAV000033 | SAMN05895044 | GCA_015818375.1 | NonPC | Public | NA | USA | Clinical | 91.7 | 100 | 0 | 98.3521 | 5102719 | 45.24 | 43 | 545410 |
| SAMN05895045-GCA_015818395.1 | PNUSAV000034 | SAMN05895045 | GCA_015818395.1 | NonPC | Public | NA | USA | Clinical | 89.6 | 99.98 | 0.08 | 98.3225 | 5189921 | 45.21 | 50 | 688953 |
| SAMN05895046-GCA_015818315.1 | PNUSAV000035 | SAMN05895046 | GCA_015818315.1 | NonPC | Public | NA | USA | Clinical | 91.8 | 100 | 0 | 98.3635 | 5104667 | 45.33 | 61 | 285297 |

| | | | | | | | | | | | | | | | | |
|---|---|---|---|---|---|---|---|---|---|---|---|---|---|---|---|---|
| SAMN05895047-SRR5057373 | PNUSAV000036 | SAMN05895047 | SRR5057373 | NonPC | Public | NA | USA | Clinical | 89.8 | 100 | 0.04 | 98.2333 | 5260082 | 45.21 | 45 | 341756 |
| SAMN05935522-GCF_001895505.1 | 100135 | SAMN05935522 | GCF_001895505.1 | NonPC | Public | 2010 | China | Clinical | 92.4 | 100 | 0.05 | 98.3354 | 5033313 | 45.26 | 31 | 478263 |
| SAMN05935523-GCF_001895615.1 | 100136 | SAMN05935523 | GCF_001895615.1 | NonPC | Public | 2008 | China | Clinical | 92.4 | 100 | 0.19 | 98.3465 | 5057222 | 45.29 | 25 | 602694 |
| SAMN05935524-GCF_001895665.1 | 100138 | SAMN05935524 | GCF_001895665.1 | PC | Public | 2008 | China | Clinical | 97.6 | 100 | 0 | 99.9421 | 5090660 | 45.29 | 43 | 288996 |
| SAMN05935525-GCF_001895585.1 | 100139 | SAMN05935525 | GCF_001895585.1 | NonPC | Public | 2014 | China | Clinical | 92.3 | 100 | 0.19 | 98.3796 | 5028841 | 45.25 | 39 | 604186 |
| SAMN05935526-GCF_001895705.1 | 100142 | SAMN05935526 | GCF_001895705.1 | NonPC | Public | 2010 | China | Clinical | 92.3 | 100 | 0.19 | 98.3699 | 5044382 | 45.26 | 31 | 547540 |
| SAMN05935527-GCF_001895515.1 | 100143 | SAMN05935527 | GCF_001895515.1 | NonPC | Public | 2013 | China | Clinical | 92.4 | 100 | 0.19 | 98.3414 | 5066319 | 45.29 | 36 | 652725 |
| SAMN05935528-GCF_001895685.1 | 100144 | SAMN05935528 | GCF_001895685.1 | NonPC | Public | 2014 | China | Clinical | 92.3 | 100 | 0.19 | 98.3651 | 5050670 | 45.22 | 36 | 495576 |
| SAMN05935529-GCF_001895695.1 | 100145 | SAMN05935529 | GCF_001895695.1 | PC | Public | 2013 | China | Clinical | 98 | 100 | 0 | 99.9792 | 5077475 | 45.3 | 41 | 490820 |
| SAMN05935530-GCF_001895745.1 | 100146 | SAMN05935530 | GCF_001895745.1 | NonPC | Public | 2009 | China | Clinical | 92.3 | 100 | 0.19 | 98.3438 | 5047819 | 45.26 | 32 | 630435 |
| SAMN05935531-GCF_001895535.1 | 100147 | SAMN05935531 | GCF_001895535.1 | NonPC | Public | 2009 | China | Clinical | 92.5 | 100 | 0.05 | 98.3554 | 5073646 | 45.27 | 37 | 569591 |
| SAMN05935532-GCF_001895525.1 | 100148 | SAMN05935532 | GCF_001895525.1 | NonPC | Public | 2009 | China | Clinical | 92.3 | 100 | 0.19 | 98.3398 | 5049550 | 45.26 | 28 | 571338 |
| SAMN05935533-GCF_001895345.1 | 100149 | SAMN05935533 | GCF_001895345.1 | NonPC | Public | 2015 | China | Clinical | 92.3 | 100 | 0.19 | 98.359 | 5037388 | 45.24 | 43 | 803378 |
| SAMN05935534-GCF_001895425.1 | 100150 | SAMN05935534 | GCF_001895425.1 | PC | Public | 2008 | China | Clinical | 97.8 | 100 | 0 | 99.9447 | 5091431 | 45.28 | 37 | 449421 |
| SAMN05935535-GCF_001895595.1 | 100151 | SAMN05935535 | GCF_001895595.1 | PC | Public | 2010 | China | Clinical | 97.5 | 100 | 0 | 99.9108 | 5160566 | 45.29 | 39 | 401400 |
| SAMN05935536-GCF_001895435.1 | 100152 | SAMN05935536 | GCF_001895435.1 | PC | Public | 2010 | China | Clinical | 98.7 | 100 | 0 | 99.9879 | 5159755 | 45.28 | 49 | 503358 |
| SAMN05935537-GCF_001895445.1 | 100153 | SAMN05935537 | GCF_001895445.1 | PC | Public | 2009 | China | Clinical | 98.3 | 100 | 0 | 99.9879 | 5089150 | 45.29 | 45 | 267664 |
| SAMN05935538-GCF_001895365.1 | 100154 | SAMN05935538 | GCF_001895365.1 | NonPC | Public | 2012 | China | Clinical | 92.6 | 100 | 0.05 | 98.2839 | 5128688 | 45.25 | 27 | 603320 |
| SAMN05935539-GCF_001895465.1 | 100156 | SAMN05935539 | GCF_001895465.1 | NonPC | Public | 2013 | China | Clinical | 92.5 | 100 | 0.19 | 98.3628 | 5073718 | 45.29 | 39 | 605610 |
| SAMN05935540-GCF_001895635.1 | 100157 | SAMN05935540 | GCF_001895635.1 | NonPC | Public | 2009 | China | Clinical | 92.3 | 100 | 0.19 | 98.3359 | 5050108 | 45.26 | 32 | 842035 |
| SAMN05935541-GCF_001895355.1 | 100158 | SAMN05935541 | GCF_001895355.1 | NonPC | Public | 2015 | China | Clinical | 92.3 | 100 | 0.19 | 98.3633 | 5032300 | 45.24 | 32 | 1084024 |
| SAMN05967526-GCA_015818465.1 | PNUSAV000037 | SAMN05967526 | GCA_015818465.1 | NonPC | Public | NA | USA | Clinical | 91.4 | 99.93 | 0 | 98.3249 | 5129931 | 45.22 | 25 | 899192 |
| SAMN05977307-GCF_001951115.1 | 1682 | SAMN05977307 | GCF_001951115.1 | NonPC | Public | 2004 | Norway | Clinical | 90.9 | 99.73 | 0 | 98.5015 | 4708005 | 45.74 | 2 | 3013583 |
| SAMN06009032-GCA_015818435.1 | PNUSAV000038 | SAMN06009032 | GCA_015818435.1 | NonPC | Public | NA | USA | Clinical | 92 | 100 | 0.63 | 98.3721 | 5146326 | 45.12 | 49 | 398982 |
| SAMN06009033-GCA_015818555.1 | PNUSAV000039 | SAMN06009033 | GCA_015818555.1 | NonPC | Public | NA | USA | Clinical | 88.8 | 100 | 0.06 | 98.1245 | 5052944 | 45.18 | 50 | 464076 |
| SAMN06009034-GCA_016306225.1 | PNUSAV000040 | SAMN06009034 | GCA_016306225.1 | NonPC | Public | NA | USA | Clinical | 89.4 | 100 | 0.31 | 98.3725 | 5093744 | 45.29 | 69 | 433085 |
| SAMN06009035-GCA_015818455.1 | PNUSAV000041 | SAMN06009035 | GCA_015818455.1 | PC | Public | NA | USA | Clinical | 97.5 | 100 | 0 | 99.9352 | 5184337 | 45.12 | 61 | 336290 |
| SAMN06042541-GCF_002220925.1 | G149 | SAMN06042541 | GCF_002220925.1 | NonPC | Public | 2007 | NA | Env | 89.6 | 100 | 0.12 | 98.3179 | 5101037 | 45.27 | 25 | 798860 |
| SAMN06042542-GCF_002220935.1 | G6928 | SAMN06042542 | GCF_002220935.1 | NonPC | Public | 2015 | NA | Env | 89.6 | 100 | 0.25 | 98.2787 | 5214042 | 45.13 | 38 | 385344 |
| SAMN06042543-GCF_002220915.1 | MA561 | SAMN06042543 | GCF_002220915.1 | NonPC | Public | 2015 | NA | Env | 89.6 | 100 | 0.25 | 98.2351 | 5234540 | 45.13 | 34 | 530527 |
| SAMN06042545-GCF_002220985.3 | MAVP-R | SAMN06042545 | GCF_002220985.3 | NonPC | Public | 2011 | NA | Clinical | 90.2 | 100 | 0.39 | 98.2675 | 5249405 | 45.19 | 2 | 3468665 |
| SAMN06042546-GCF_002220905.1 | MAVP-90 | SAMN06042546 | GCF_002220905.1 | NonPC | Public | 2015 | NA | Clinical | 89.2 | 100 | 0.04 | 98.2584 | 5127903 | 45.24 | 104 | 97905 |
| SAMN06042547-GCF_002220995.1 | MAVP-94 | SAMN06042547 | GCF_002220995.1 | NonPC | Public | 2015 | NA | Clinical | 89.3 | 100 | 0.04 | 98.2863 | 5134782 | 45.21 | 79 | 118688 |
| SAMN06042548-GCF_002221015.1 | MAVP-109 | SAMN06042548 | GCF_002221015.1 | NonPC | Public | 2015 | NA | Clinical | 89.9 | 100 | 0.04 | 98.2915 | 5231020 | 45.19 | 61 | 200281 |
| SAMN06042549-GCF_002221005.1 | MAVP-112 | SAMN06042549 | GCF_002221005.1 | NonPC | Public | 2015 | NA | Clinical | 90 | 100 | 0.04 | 98.2672 | 5285634 | 45.2 | 42 | 328132 |
| SAMN06061934-GCF_001996365.2 | MVP1 | SAMN06061934 | GCF_001996365.2 | NonPC | Public | 2016 | Malaysia | Env | 92.3 | 100 | 0 | 98.2648 | 5389211 | 45.3 | 3 | 3446010 |
| SAMN06066407-GCF_002013995.1 | MVP2 | SAMN06066407 | GCF_002013995.1 | NonPC | Public | 2016 | Malaysia | Env | 91.1 | 100 | 0 | 98.3058 | 5273104 | 45.36 | 84 | 129821 |
| SAMN06066420-GCF_002014005.1 | MVP4 | SAMN06066420 | GCF_002014005.1 | NonPC | Public | 2016 | Malaysia | Env | 89.9 | 100 | 0.27 | 98.2906 | 5268039 | 45.37 | 115 | 94227 |
| SAMN06066428-GCF_002014055.1 | MVP6 | SAMN06066428 | GCF_002014055.1 | NonPC | Public | 2016 | Malaysia | Env | 90.8 | 100 | 0 | 98.2938 | 5195990 | 45.5 | 190 | 47800 |
| SAMN06066439-GCF_002014015.1 | MVP9 | SAMN06066439 | GCF_002014015.1 | NonPC | Public | 2016 | Malaysia | Env | 89.3 | 100 | 0.05 | 98.3224 | 4967664 | 45.51 | 115 | 87871 |
| SAMN06076928-GCF_001998785.1 | ATCC_17802 | SAMN06076928 | GCF_001998785.1 | NonPC | Public | 1951 | Japan | Clinical | 90.7 | 100 | 0.05 | 98.354 | 5071046 | 45.25 | 64 | 309969 |
| SAMN06076958-GCA_015818275.1 | CFSAN022891 | SAMN06076958 | GCA_015818275.1 | NonPC | Public | 2004 | USA | Env | 89.9 | 100 | 0.07 | 98.1035 | 5156837 | 45.33 | 14 | 1771206 |
| SAMN06076959-GCA_015818355.1 | CFSAN022892 | SAMN06076959 | GCA_015818355.1 | NonPC | Public | 2004 | USA | Env | 91.1 | 99.95 | 0.05 | 98.2365 | 5144129 | 45.21 | 28 | 566363 |
| SAMN06076960-GCA_015818295.1 | 48262 | SAMN06076960 | GCA_015818295.1 | NonPC | Public | 1990 | USA | Clinical | 89.3 | 100 | 0 | 98.073 | 5024075 | 45.28 | 58 | 292211 |
| SAMN06076961-GCA_015818235.1 | CFSAN022894 | SAMN06076961 | GCA_015818235.1 | NonPC | Public | 2001 | USA | Clinical | 89.3 | 99.95 | 0 | 98.0736 | 5059503 | 45.28 | 77 | 319036 |
| SAMN06076962-GCA_015819565.1 | 049-2A3 | SAMN06076962 | GCA_015819565.1 | NonPC | Public | 1997 | USA | Env | 91.6 | 100 | 0.38 | 98.3835 | 5234220 | 45.23 | 61 | 521863 |
| SAMN06076963-GCA_015818175.1 | CFSAN022896 | SAMN06076963 | GCA_015818175.1 | NonPC | Public | 2001 | USA | Env | 90.7 | 100 | 0 | 98.2687 | 5057637 | 45.27 | 45 | 556640 |
| SAMN06076964-GCA_015818215.1 | CFSAN022897 | SAMN06076964 | GCA_015818215.1 | NonPC | Public | 1993 | USA | Env | 89.2 | 100 | 0.11 | 98.146 | 5019491 | 45.32 | 72 | 371397 |

| | | | | | | | | | | | | | | | | |
|---|---|---|---|---|---|---|---|---|---|---|---|---|---|---|---|---|
| SAMN06076965-GCA_015818255.1 | CFSAN022898 | SAMN06076965 | GCA_015818255.1 | NonPC | Public | 2001 | USA | Env | 90.7 | 100 | 0 | 98.2789 | 5060622 | 45.26 | 43 | 511446 |
| SAMN06076966-GCA_015819525.1 | 9546257 | SAMN06076966 | GCA_015819525.1 | NonPC | Public | 1995 | USA | Clinical | 90.6 | 100 | 0.14 | 98.2332 | 5055534 | 45.18 | 45 | 548562 |
| SAMN06076967-GCA_015818105.1 | 98-506-B102 | SAMN06076967 | GCA_015818105.1 | NonPC | Public | 1998 | USA | Env | 88.9 | 100 | 0 | 98.1499 | 5123987 | 45.22 | 59 | 452507 |
| SAMN06076968-GCA_015819505.1 | 98-506-B103 | SAMN06076968 | GCA_015819505.1 | NonPC | Public | 1998 | USA | Env | 88.9 | 100 | 0 | 98.1919 | 5111579 | 45.23 | 66 | 437830 |
| SAMN06076969-GCA_015819465.1 | 98-513-F51 | SAMN06076969 | GCA_015819465.1 | NonPC | Public | 1998 | USA | Env | 91.4 | 100 | 0.14 | 98.1606 | 5166771 | 45.24 | 8 | 1817145 |
| SAMN06076970-GCA_015819435.1 | 98-548-D11 | SAMN06076970 | GCA_015819435.1 | NonPC | Public | 1998 | USA | Env | 90.6 | 100 | 0.14 | 98.2115 | 5063164 | 45.17 | 36 | 490092 |
| SAMN06076971-GCA_015817995.1 | 98-605-A9 | SAMN06076971 | GCA_015817995.1 | NonPC | Public | 1998 | USA | Env | 89 | 100 | 0 | 98.1807 | 5078113 | 45.26 | 86 | 368122 |
| SAMN06076972-GCA_015819395.1 | 98-605-A10 | SAMN06076972 | GCA_015819395.1 | NonPC | Public | 1998 | USA | Env | 89.1 | 100 | 0 | 98.1439 | 5087445 | 45.25 | 78 | 405700 |
| SAMN06076973-GCA_015818075.1 | 99-524-A9 | SAMN06076973 | GCA_015818075.1 | NonPC | Public | 1999 | USA | Env | 91.4 | 100 | 0.04 | 98.2837 | 5073555 | 45.25 | 68 | 373496 |
| SAMN06076974-GCA_015819315.1 | 99-780-C12 | SAMN06076974 | GCA_015819315.1 | NonPC | Public | 1999 | USA | Env | 89.1 | 100 | 0.03 | 98.0993 | 4962556 | 45.33 | 67 | 235187 |
| SAMN06076975-GCA_015819355.1 | CFSAN023522 | SAMN06076975 | GCA_015819355.1 | NonPC | Public | 1999 | USA | Env | 92.4 | 100 | 0.29 | 98.3209 | 5070780 | 45.3 | 37 | 511597 |
| SAMN06076976-GCA_015817855.1 | CFSAN023523 | SAMN06076976 | GCA_015817855.1 | NonPC | Public | 2000 | USA | Env | 89.1 | 100 | 0.08 | 98.1589 | 5001439 | 45.36 | 77 | 180005 |
| SAMN06076977-GCA_015818035.1 | CFSAN023524 | SAMN06076977 | GCA_015818035.1 | NonPC | Public | 2000 | USA | Env | 89.2 | 100 | 0.05 | 98.1624 | 5096223 | 45.28 | 66 | 256974 |
| SAMN06076978-GCA_015818095.1 | CFSAN023525 | SAMN06076978 | GCA_015818095.1 | NonPC | Public | 2000 | USA | Env | 90 | 100 | 0.19 | 98.037 | 5185821 | 45.37 | 29 | 1138702 |
| SAMN06076979-GCA_015818015.1 | CFSAN023526 | SAMN06076979 | GCA_015818015.1 | NonPC | Public | 2000 | USA | Env | 90 | 100 | 0 | 98.0344 | 5174187 | 45.34 | 8 | 1138428 |
| SAMN06076980-GCA_015818135.1 | CFSAN023527 | SAMN06076980 | GCA_015818135.1 | NonPC | Public | 2000 | USA | Env | 89.3 | 100 | 0.14 | 98.1175 | 5207073 | 45.21 | 71 | 238399 |
| SAMN06076981-GCA_015817815.1 | CFSAN023528 | SAMN06076981 | GCA_015817815.1 | NonPC | Public | 2000 | USA | Env | 89.3 | 100 | 0.14 | 98.1233 | 5055170 | 45.27 | 65 | 238399 |
| SAMN06076982-GCA_015819285.1 | CFSAN023529 | SAMN06076982 | GCA_015819285.1 | NonPC | Public | 1999 | USA | Env | 89.5 | 100 | 0 | 98.1116 | 5040691 | 45.3 | 95 | 200252 |
| SAMN06076984-GCA_015819215.1 | CFSAN023531 | SAMN06076984 | GCA_015819215.1 | NonPC | Public | 1999 | USA | Env | 90.6 | 100 | 0.14 | 98.1616 | 5086542 | 45.17 | 34 | 525379 |
| SAMN06076985-GCA_015819095.1 | CFSAN023532 | SAMN06076985 | GCA_015819095.1 | NonPC | Public | 1999 | USA | Env | 90.6 | 100 | 0.14 | 98.1752 | 5088684 | 45.17 | 34 | 526265 |
| SAMN06076986-GCA_015819055.1 | CFSAN023533 | SAMN06076986 | GCA_015819055.1 | PC | Public | 1996 | India | Clinical | 99.9 | 100 | 0 | 99.9944 | 5162993 | 45.37 | 3 | 3288181 |
| SAMN06076987-GCA_015818995.1 | CFSAN023535 | SAMN06076987 | GCA_015818995.1 | PC | Public | 1996 | India | Clinical | 99.9 | 100 | 0 | 99.9952 | 5167742 | 45.36 | 4 | 3287277 |
| SAMN06076988-GCA_015818975.1 | CFSAN023536 | SAMN06076988 | GCA_015818975.1 | PC | Public | 1996 | India | Clinical | 99.9 | 100 | 0 | 99.9947 | 5162977 | 45.37 | 3 | 3287984 |
| SAMN06076989-GCA_015818875.1 | CFSAN023537 | SAMN06076989 | GCA_015818875.1 | PC | Public | 1997 | India | Clinical | 98.2 | 100 | 0 | 99.9722 | 5146017 | 45.28 | 41 | 507703 |
| SAMN06076990-GCA_015818955.1 | AN-8373 | SAMN06076990 | GCA_015818955.1 | PC | Public | 1998 | Bangladesh | Clinical | 99.5 | 99.73 | 0 | 99.9959 | 5154691 | 45.35 | 6 | 3288001 |
| SAMN06076991-GCA_015818795.1 | CFSAN023540 | SAMN06076991 | GCA_015818795.1 | PC | Public | 1998 | South_Korea | Clinical | 98.2 | 99.97 | 4.09 | 99.9923 | 5210225 | 45.29 | 51 | 274170 |
| SAMN06076992-GCA_015818835.1 | CFSAN023541 | SAMN06076992 | GCA_015818835.1 | PC | Public | 1998 | USA | Clinical | 98.2 | 100 | 0 | 99.9745 | 5150702 | 45.28 | 41 | 444402 |
| SAMN06076993-GCA_015819075.1 | BAC-98-3372 | SAMN06076993 | GCA_015819075.1 | PC | Public | 1998 | USA | Clinical | 99.9 | 100 | 0 | 99.9956 | 5168526 | 45.36 | 4 | 3287948 |
| SAMN06076994-GCA_015819025.1 | BAC-98-3374 | SAMN06076994 | GCA_015819025.1 | PC | Public | 1998 | USA | Clinical | 100 | 100 | 0 | 99.9942 | 5251443 | 45.19 | 9 | 3287894 |
| SAMN06076995-GCA_015818935.1 | BAC-98-4092 | SAMN06076995 | GCA_015818935.1 | PC | Public | 1998 | USA | Clinical | 99.9 | 100 | 0 | 99.9937 | 5249429 | 45.19 | 9 | 3287830 |
| SAMN06076996-GCA_015818855.1 | AN-5034 | SAMN06076996 | GCA_015818855.1 | PC | Public | 1998 | Bangladesh | Clinical | 98.8 | 100 | 0 | 99.933 | 5217959 | 45.31 | 16 | 1006395 |
| SAMN06076997-GCA_015818815.1 | AO-24491 | SAMN06076997 | GCA_015818815.1 | PC | Public | 1999 | Bangladesh | Clinical | 97.7 | 100 | 0 | 99.9411 | 5132826 | 45.29 | 40 | 545189 |
| SAMN06076998-GCA_015818775.1 | CFSAN023547 | SAMN06076998 | GCA_015818775.1 | PC | Public | 1999 | Thailand | Clinical | 97.4 | 99.73 | 0 | 99.9257 | 5129218 | 45.26 | 43 | 461427 |
| SAMN06076999-GCA_015819015.1 | AN-16000 | SAMN06076999 | GCA_015819015.1 | PC | Public | 1998 | Bangladesh | Clinical | 99.2 | 100 | 0.27 | 99.9352 | 5204005 | 45.35 | 8 | 1391732 |
| SAMN06077000-GCA_015818905.1 | CFSAN023549 | SAMN06077000 | GCA_015818905.1 | PrePC | Public | 1980 | Bangladesh | Clinical | 94.4 | 99.73 | 0.14 | 99.149 | 5029688 | 45.34 | 49 | 461912 |
| SAMN06077001-GCA_015818895.1 | CFSAN023550 | SAMN06077001 | GCA_015818895.1 | NonPC | Public | 2005 | Chile | Env | 91.2 | 100 | 0.35 | 98.3377 | 5018974 | 45.37 | 49 | 399296 |
| SAMN06077002-GCA_015818735.1 | CFSAN023551 | SAMN06077002 | GCA_015818735.1 | NonPC | Public | 2005 | Chile | Env | 91.3 | 100 | 0.22 | 98.3462 | 5229550 | 45.09 | 39 | 558772 |
| SAMN06077003-GCA_015818755.1 | CFSAN023552 | SAMN06077003 | GCA_015818755.1 | NonPC | Public | 2005 | Chile | Env | 90.4 | 100 | 0.27 | 98.3771 | 4988830 | 45.39 | 49 | 563943 |
| SAMN06077004-GCA_015818695.1 | CFSAN023553 | SAMN06077004 | GCA_015818695.1 | NonPC | Public | 2005 | Chile | Env | 91 | 100 | 0.08 | 98.3108 | 5199350 | 45.33 | 51 | 531409 |
| SAMN06077005-GCA_015818715.1 | CFSAN023554 | SAMN06077005 | GCA_015818715.1 | PC | Public | 2005 | Chile | Env | 99.9 | 100 | 0 | 99.9956 | 5173876 | 45.35 | 5 | 1458437 |
| SAMN06077006-GCA_015817895.1 | CFSAN023555 | SAMN06077006 | GCA_015817895.1 | NonPC | Public | 2005 | Chile | Env | 90.9 | 100 | 0.4 | 98.3386 | 4975221 | 45.41 | 48 | 408879 |
| SAMN06077007-GCA_015817935.1 | CFSAN023556 | SAMN06077007 | GCA_015817935.1 | PC | Public | 2005 | Chile | Env | 99 | 100 | 0.07 | 99.9869 | 5256208 | 45.28 | 22 | 494557 |
| SAMN06077008-GCA_015817955.1 | CFSAN023557 | SAMN06077008 | GCA_015817955.1 | NonPC | Public | 2004 | Chile | Env | 88 | 100 | 0.33 | 98.3538 | 5037064 | 45.28 | 55 | 475954 |
| SAMN06077010-GCA_015817975.1 | CFSAN023559 | SAMN06077010 | GCA_015817975.1 | NonPC | Public | 2004 | Chile | Env | 91.5 | 100 | 0.35 | 98.3635 | 5127257 | 45.29 | 38 | 580631 |
| SAMN06077011-GCA_015818675.1 | CFSAN023560 | SAMN06077011 | GCA_015818675.1 | NonPC | Public | 2004 | Chile | Env | 90.1 | 100 | 0 | 98.3016 | 4975984 | 45.41 | 60 | 710324 |
| SAMN06077012-GCA_015818055.1 | CFSAN023561 | SAMN06077012 | GCA_015818055.1 | NonPC | Public | 2004 | Chile | Env | 91.5 | 100 | 0.48 | 98.378 | 5111350 | 45.31 | 67 | 298580 |
| SAMN06077013-GCA_015817755.1 | CFSAN023562 | SAMN06077013 | GCA_015817755.1 | NonPC | Public | 2004 | Chile | Env | 91.1 | 100 | 0.1 | 98.2913 | 5007484 | 45.36 | 37 | 620888 |
| SAMN06077014-GCA_015817735.1 | CFSAN023563 | SAMN06077014 | GCA_015817735.1 | PC | Public | 1999 | Thailand | Clinical | 97.9 | 100 | 0 | 99.9421 | 5159946 | 45.24 | 56 | 403044 |

| | | | | | | | | | | | | | | | | |
|---|---|---|---|---|---|---|---|---|---|---|---|---|---|---|---|---|
| SAMN06077015-GCA_015817835.1 | CFSAN023564 | SAMN06077015 | GCA_015817835.1 | PC | Public | 1998 | Japan | Clinical | 97.5 | 100 | 0 | 99.9417 | 5076836 | 45.27 | 35 | 507855 |
| SAMN06077016-GCA_015817775.1 | AN-2189 | SAMN06077016 | GCA_015817775.1 | PC | Public | 1998 | Bangladesh | Clinical | 97.9 | 100 | 0 | 99.9421 | 5121288 | 45.28 | 55 | 427828 |
| SAMN06077017-GCA_015817595.1 | AP-11243 | SAMN06077017 | GCA_015817595.1 | PC | Public | 2000 | Bangladesh | Clinical | 99.2 | 100 | 0.27 | 99.9295 | 5204070 | 45.35 | 8 | 3316320 |
| SAMN06107071-GCF_003408935.1 | 271 | SAMN06107071 | GCF_003408935.1 | NonPC | Public | 2015 | China | Env | 88.4 | 99.66 | 0.71 | 98.4072 | 4883222 | 45.34 | 250 | 37510 |
| SAMN06110481-GCF_002150185.1 | xtf19 | SAMN06110481 | GCF_002150185.1 | NonPC | Public | 2013 | China | Env | 91.4 | 100 | 0.27 | 98.3143 | 5128492 | 45.23 | 75 | 421386 |
| SAMN06110496-GCF_002150175.1 | xtf22 | SAMN06110496 | GCF_002150175.1 | NonPC | Public | 2013 | China | Env | 90 | 100 | 0.27 | 98.3386 | 5014511 | 45.33 | 54 | 857198 |
| SAMN06112607-GCF_002150155.1 | xtf27 | SAMN06112607 | GCF_002150155.1 | NonPC | Public | 2013 | China | Env | 92.2 | 100 | 0.14 | 98.2965 | 5223584 | 45.12 | 40 | 645623 |
| SAMN06115936-GCF_002072835.1 | CFSAN026730 | SAMN06115936 | GCF_002072835.1 | NonPC | Public | 2013 | USA | Clinical | 91.5 | 100 | 0 | 98.3103 | 5092810 | 45.27 | 56 | 548157 |
| SAMN06127921-GCF_002153945.1 | xtf37 | SAMN06127921 | GCF_002153945.1 | NonPC | Public | 2013 | China | Env | 91.5 | 100 | 0.68 | 98.2937 | 5287423 | 45.12 | 93 | 170860 |
| SAMN06130233-GCF_002150335.1 | GIMxtf40-2013.07 | SAMN06130233 | GCF_002150335.1 | NonPC | Public | 2013 | China | Env | 91.3 | 100 | 0.08 | 98.326 | 5211236 | 45.27 | 56 | 549560 |
| SAMN06131432-GCF_002150315.1 | GIMxtf41-2013.07 | SAMN06131432 | GCF_002150315.1 | NonPC | Public | 2013 | China | Env | 91.3 | 100 | 0.08 | 98.3036 | 5209742 | 45.27 | 59 | 484037 |
| SAMN06133346-GCF_002153925.1 | GIMxtf47-2013.12 | SAMN06133346 | GCF_002153925.1 | NonPC | Public | 2013 | China | Env | 92.5 | 100 | 3.51 | 98.3247 | 5394055 | 45.34 | 144 | 96662 |
| SAMN06134384-GCF_002153875.1 | GIMxtf133-2013.12 | SAMN06134384 | GCF_002153875.1 | NonPC | Public | 2013 | China | Env | 91.1 | 100 | 0.27 | 98.2242 | 5440974 | 45.32 | 157 | 280864 |
| SAMN06134688-GCF_002153895.1 | GIMxtf230-2012.11 | SAMN06134688 | GCF_002153895.1 | NonPC | Public | 2012 | China | Env | 92.1 | 100 | 0.27 | 98.3612 | 5033236 | 45.31 | 41 | 508047 |
| SAMN06140976-GCF_002154015.1 | GIMxtf294-2013.07 | SAMN06140976 | GCF_002154015.1 | NonPC | Public | 2013 | China | Env | 91.7 | 100 | 0.09 | 98.3699 | 5172403 | 45.31 | 77 | 411289 |
| SAMN06143023-GCF_001913695.1 | GIMxtf283-2012.12 | SAMN06143023 | GCF_001913695.1 | NonPC | Public | 2012 | China | Env | 90.9 | 100 | 0.19 | 98.3227 | 5219743 | 45.2 | 48 | 439580 |
| SAMN06158907-GCF_002153975.1 | GIMxtf296-2013.05 | SAMN06158907 | GCF_002153975.1 | NonPC | Public | 2013 | China | Env | 91.6 | 100 | 0.2 | 98.3488 | 5226646 | 45.34 | 111 | 525237 |
| SAMN06160634-GCA_015817695.1 | PNUSAV000045 | SAMN06160634 | GCA_015817695.1 | NonPC | Public | NA | USA | Clinical | 91.1 | 100 | 0.14 | 98.3872 | 5064031 | 45.39 | 80 | 189025 |
| SAMN06160635-GCA_015817715.1 | PNUSAV000046 | SAMN06160635 | GCA_015817715.1 | NonPC | Public | NA | USA | Clinical | 91.6 | 100 | 0 | 98.3706 | 5017248 | 45.32 | 28 | 451116 |
| SAMN06160636-GCA_015817635.1 | PNUSAV000047 | SAMN06160636 | GCA_015817635.1 | NonPC | Public | NA | USA | Clinical | 90.4 | 100 | 0.33 | 98.2752 | 5155002 | 45.28 | 75 | 385813 |
| SAMN06160637-GCA_015817615.1 | PNUSAV000048 | SAMN06160637 | GCA_015817615.1 | NonPC | Public | NA | USA | Clinical | 90.2 | 100 | 0.31 | 98.3019 | 5263405 | 45 | 79 | 407054 |
| SAMN06160639-GCA_015817665.1 | PNUSAV000050 | SAMN06160639 | GCA_015817665.1 | NonPC | Public | NA | USA | Clinical | 89.9 | 100 | 0 | 98.3418 | 4931752 | 45.43 | 77 | 332220 |
| SAMN06161405-GCF_002154055.1 | GIMxtf336-2013.07 | SAMN06161405 | GCF_002154055.1 | NonPC | Public | 2013 | China | Env | 91.6 | 100 | 0.27 | 98.3388 | 5110250 | 45.31 | 57 | 861660 |
| SAMN06161545-GCF_001913785.1 | GIMxtf381-2013.06 | SAMN06161545 | GCF_001913785.1 | NonPC | Public | 2013 | China | Env | 91 | 100 | 2.82 | 98.3533 | 5273588 | 45.27 | 99 | 405631 |
| SAMN06162268-GCF_001913735.1 | GIMxtfL61-2011.05 | SAMN06162268 | GCF_001913735.1 | PC | Public | 2011 | China | Clinical | 98.5 | 100 | 0.01 | 99.9835 | 5193336 | 45.32 | 66 | 426047 |
| SAMN06163187-GCF_001913745.1 | GIMxtfL65-2011.05 | SAMN06163187 | GCF_001913745.1 | PC | Public | 2011 | China | Clinical | 98.5 | 100 | 0.01 | 99.9884 | 5193136 | 45.32 | 64 | 426017 |
| SAMN06166873-GCF_001913775.1 | GIMxtfL71-2011.05 | SAMN06166873 | GCF_001913775.1 | NonPC | Public | 2011 | China | Clinical | 92 | 100 | 0 | 98.4133 | 5103058 | 45.25 | 35 | 636384 |
| SAMN06169655-GCF_002150465.1 | GIMxtfL83-2011.05 | SAMN06169655 | GCF_002150465.1 | NonPC | Public | 2011 | China | Clinical | 88.2 | 100 | 0.49 | 98.2533 | 5247295 | 45.28 | 60 | 414853 |
| SAMN06169820-GCF_001921805.1 | GIMxtfL88-2011.05 | SAMN06169820 | GCF_001921805.1 | NonPC | Public | 2011 | China | Clinical | 88.5 | 100 | 0.39 | 98.2267 | 5076715 | 45.23 | 48 | 1367986 |
| SAMN06214593-GCA_015817375.1 | CFSAN025071 | SAMN06214593 | GCA_015817375.1 | NonPC | Public | 2006 | Spain | Env | 89.2 | 100 | 0.05 | 98.3284 | 5011873 | 45.3 | 50 | 334451 |
| SAMN06214594-GCA_015817565.1 | CFSAN025072 | SAMN06214594 | GCA_015817565.1 | NonPC | Public | 2007 | Spain | Env | 90.8 | 100 | 0 | 98.2886 | 5049827 | 45.27 | 45 | 692586 |
| SAMN06214595-GCA_015817795.1 | CFSAN025073 | SAMN06214595 | GCA_015817795.1 | NonPC | Public | 2006 | Spain | Env | 89.2 | 100 | 0.05 | 98.3295 | 5007150 | 45.3 | 54 | 334451 |
| SAMN06214596-GCA_015817465.1 | CFSAN025074 | SAMN06214596 | GCA_015817465.1 | NonPC | Public | 2006 | Spain | Env | 92.7 | 99.73 | 0 | 98.4173 | 5214002 | 45.17 | 37 | 464571 |
| SAMN06214597-GCA_015817235.1 | CFSAN025075 | SAMN06214597 | GCA_015817235.1 | NonPC | Public | 2007 | Spain | Env | 89.2 | 100 | 0.05 | 98.3423 | 5007071 | 45.3 | 58 | 329435 |
| SAMN06214598-GCA_015816535.1 | CFSAN025076 | SAMN06214598 | GCA_015816535.1 | NonPC | Public | 2007 | Spain | Env | 91.4 | 100 | 0.27 | 98.353 | 5165215 | 45.26 | 118 | 261266 |
| SAMN06214599-GCA_015817295.1 | CFSAN025077 | SAMN06214599 | GCA_015817295.1 | NonPC | Public | 2006 | Spain | Env | 91.6 | 100 | 0.49 | 98.2797 | 5259103 | 45.12 | 63 | 401792 |
| SAMN06214600-GCA_015817215.1 | CFSAN025078 | SAMN06214600 | GCA_015817215.1 | NonPC | Public | 2007 | Spain | Env | 91.4 | 100 | 0.27 | 98.3044 | 5168135 | 45.25 | 118 | 286762 |
| SAMN06214601-GCA_015817115.1 | CFSAN025079 | SAMN06214601 | GCA_015817115.1 | NonPC | Public | 2007 | Spain | Env | 90.5 | 100 | 0 | 98.2698 | 5182545 | 45.14 | 40 | 541627 |
| SAMN06214602-GCA_015817315.1 | CFSAN022333 | SAMN06214602 | GCA_015817315.1 | NonPC | Public | 2012 | Spain | Clinical | 91.3 | 100 | 0.53 | 98.2711 | 5416300 | 44.88 | 50 | 360405 |
| SAMN06214603-GCA_015816595.1 | CFSAN022337 | SAMN06214603 | GCA_015816595.1 | NonPC | Public | 2012 | Spain | Clinical | 89.1 | 100 | 0.05 | 98.3486 | 4997729 | 45.3 | 65 | 275128 |
| SAMN06214604-GCA_015817445.1 | CFSAN029659 | SAMN06214604 | GCA_015817445.1 | NonPC | Public | 2007 | Spain | Env | 92.8 | 99.73 | 0 | 98.4149 | 5217807 | 45.17 | 40 | 458434 |
| SAMN06214605-GCA_015817415.1 | CFSAN029660 | SAMN06214605 | GCA_015817415.1 | NonPC | Public | 2006 | Spain | Env | 89.6 | 100 | 0.83 | 98.3031 | 5179208 | 45.18 | 34 | 544471 |
| SAMN06214606-GCA_015816555.1 | 118 | SAMN06214606 | GCA_015816555.1 | NonPC | Public | 2012 | Spain | Clinical | 91.2 | 100 | 0.27 | 98.3187 | 5144440 | 45.18 | 57 | 360405 |
| SAMN06214607-GCA_015817335.1 | 119 | SAMN06214607 | GCA_015817335.1 | NonPC | Public | 2012 | Spain | Clinical | 91.2 | 100 | 0.27 | 98.3201 | 5134310 | 45.19 | 59 | 360405 |
| SAMN06214608-SRR5163834 | 113477 | SAMN06214608 | SRR5163834 | NonPC | Public | 2015 | Spain | Clinical | 90.6 | 100 | 0 | 98.232 | 5185965 | 45.3 | 38 | 345819 |
| SAMN06214609-GCA_015817655.1 | CFSAN053627 | SAMN06214609 | GCA_015817655.1 | NonPC | Public | 2016 | Spain | Clinical | 90.4 | 100 | 0 | 98.2611 | 5161429 | 45.29 | 39 | 346562 |
| SAMN06214610-GCA_015816435.1 | CFSAN053626 | SAMN06214610 | GCA_015816435.1 | PC | Public | 2016 | Spain | Clinical | 98.1 | 100 | 0 | 99.993 | 5100864 | 45.28 | 68 | 426433 |
| SAMN06214611-GCA_015817515.1 | AMC_317 | SAMN06214611 | GCA_015817515.1 | PC | Public | 2016 | Spain | Clinical | 99.9 | 100 | 0 | 99.9925 | 5187194 | 45.34 | 5 | 3286689 |

| | | | | | | | | | | | | | | | | |
|---|---|---|---|---|---|---|---|---|---|---|---|---|---|---|---|---|
| SAMN06214612-GCA_015816575.1 | AMC_325 | SAMN06214612 | GCA_015816575.1 | NonPC | Public | 2016 | Spain | Clinical | 91.2 | 100 | 0.27 | 98.2962 | 5137553 | 45.19 | 59 | 360405 |
| SAMN06310878-GCF_003085735.1 | 20130629002S01 | SAMN06310878 | GCF_003085735.1 | NonPC | Public | 2013 | China | Env | 92.3 | 100 | 0.27 | 98.3442 | 5404824 | 45.33 | 4 | 3394092 |
| SAMN06490120-GCF_002144645.1 | 13-028/A2 | SAMN06490120 | GCF_002144645.1 | NonPC | Public | 2013 | Viet_Nam | Env | 89 | 99.86 | 0.62 | 98.3149 | 4904435 | 45.38 | 330 | 29144 |
| SAMN06546662-GCF_002114595.1 | 12-009A/1335 | SAMN06546662 | GCF_002114595.1 | NonPC | Public | 2012 | Viet_Nam | Env | 91 | 98.94 | 0.52 | 98.4293 | 5310627 | 45.2 | 410 | 30833 |
| SAMN06546663-GCF_002114585.1 | 12-297/B | SAMN06546663 | GCF_002114585.1 | NonPC | Public | 2012 | Viet_Nam | Env | 91.1 | 100 | 0.87 | 98.3759 | 5334719 | 45.29 | 385 | 37471 |
| SAMN06546664-GCF_002114605.1 | 13-306D/4 | SAMN06546664 | GCF_002114605.1 | NonPC | Public | 2013 | Mexico | Env | 90.8 | 99.64 | 0.94 | 98.2294 | 5236816 | 45.27 | 388 | 27902 |
| SAMN06607466-GCA_015817385.1 | PNUSAV000044 | SAMN06607466 | GCA_015817385.1 | NonPC | Public | NA | USA | Clinical | 91.5 | 100 | 0.08 | 98.2861 | 5200378 | 45.17 | 19 | 1131871 |
| SAMN06607467-GCA_015817435.1 | PNUSAV000051 | SAMN06607467 | GCA_015817435.1 | NonPC | Public | NA | USA | Clinical | 91.2 | 100 | 0.14 | 98.4059 | 5059972 | 45.28 | 40 | 681958 |
| SAMN06917270-GCF_002221225.1 | G4186 | SAMN06917270 | GCF_002221225.1 | NonPC | Public | 2013 | NA | Env | 90.3 | 100 | 0.14 | 98.174 | 5045181 | 45.2 | 40 | 423176 |
| SAMN06917271-GCF_002221165.1 | MAVP-14 | SAMN06917271 | GCF_002221165.1 | NonPC | Public | 2013 | NA | Clinical | 90.4 | 100 | 0.15 | 98.1675 | 5125559 | 45.22 | 144 | 344839 |
| SAMN06917272-GCF_002221305.1 | MAVP-71 | SAMN06917272 | GCF_002221305.1 | NonPC | Public | 2014 | NA | Clinical | 89.5 | 100 | 0 | 98.0711 | 5036752 | 45.27 | 50 | 367453 |
| SAMN06917273-GCF_002221175.1 | MAVP-46 | SAMN06917273 | GCF_002221175.1 | NonPC | Public | 2013 | NA | Clinical | 88.8 | 100 | 0.06 | 98.1144 | 5104502 | 45.16 | 55 | 434737 |
| SAMN06917274-GCF_002221185.1 | MAVP-50 | SAMN06917274 | GCF_002221185.1 | NonPC | Public | 2013 | NA | Clinical | 91.9 | 100 | 0 | 98.2483 | 5117363 | 45.28 | 44 | 500691 |
| SAMN06917275-GCF_002221285.1 | MAVP-67 | SAMN06917275 | GCF_002221285.1 | NonPC | Public | 2014 | NA | Clinical | 89.8 | 100 | 0 | 98.2526 | 5143603 | 45.24 | 35 | 557757 |
| SAMN06917276-GCF_002221425.1 | MAVP-99 | SAMN06917276 | GCF_002221425.1 | NonPC | Public | 2015 | NA | Clinical | 89.9 | 100 | 0.22 | 98.2803 | 5156447 | 45.19 | 42 | 231995 |
| SAMN06917277-GCF_002221245.1 | G3578 | SAMN06917277 | GCF_002221245.1 | NonPC | Public | 2013 | NA | Env | 90.4 | 100 | 0 | 98.2654 | 5222070 | 45.16 | 25 | 1287787 |
| SAMN06917278-GCF_002221265.1 | MAVP-21 | SAMN06917278 | GCF_002221265.1 | NonPC | Public | 2013 | NA | Clinical | 89.9 | 100 | 0 | 98.3284 | 5220871 | 45.24 | 48 | 432240 |
| SAMN06917279-GCF_002221145.1 | CTVP43C | SAMN06917279 | GCF_002221145.1 | NonPC | Public | 2013 | NA | Clinical | 91.3 | 100 | 0 | 98.3874 | 5067622 | 45.29 | 37 | 314858 |
| SAMN06917281-GCF_002238235.1 | MAVP-25 | SAMN06917281 | GCF_002238235.1 | NonPC | Public | 2013 | NA | Clinical | 89.7 | 100 | 0.08 | 98.2443 | 5181753 | 45.17 | 63 | 166456 |
| SAMN06917282-GCF_002221085.1 | CTVP35C | SAMN06917282 | GCF_002221085.1 | NonPC | Public | 2013 | NA | Clinical | 89.3 | 100 | 0.12 | 98.3499 | 5024761 | 45.39 | 45 | 460036 |
| SAMN06917283-GCF_002221075.1 | MAVP-G | SAMN06917283 | GCF_002221075.1 | NonPC | Public | 2011 | NA | Clinical | 90.2 | 100 | 0.22 | 98.3113 | 5157417 | 45.17 | 70 | 206191 |
| SAMN06917284-GCF_002221065.1 | MAVP-108 | SAMN06917284 | GCF_002221065.1 | NonPC | Public | 2015 | NA | Clinical | 90.2 | 100 | 0.2 | 98.2589 | 5103891 | 45.23 | 29 | 483451 |
| SAMN07185541-GCA_015817175.1 | 14/692_A_1_VP | SAMN07185541 | GCA_015817175.1 | NonPC | Public | 2009 | UK | Env | 91.7 | 100 | 0.22 | 98.41 | 5148005 | 45.26 | 66 | 325252 |
| SAMN07185542-GCA_015817265.1 | 14/1073_H_VP | SAMN07185542 | GCA_015817265.1 | NonPC | Public | 2010 | UK | Env | 91.7 | 100 | 0.22 | 98.4032 | 5143550 | 45.27 | 54 | 714419 |
| SAMN07185543-GCA_015817195.1 | 14/559_B_VP | SAMN07185543 | GCA_015817195.1 | NonPC | Public | 2010 | UK | Env | 91.7 | 100 | 0.22 | 98.3885 | 5148806 | 45.26 | 61 | 327691 |
| SAMN07185544-GCA_015817255.1 | 14/1499_VP | SAMN07185544 | GCA_015817255.1 | NonPC | Public | 2010 | UK | Env | 88.9 | 100 | 0 | 98.1087 | 5080959 | 45.31 | 80 | 202309 |
| SAMN07185545-GCA_015817135.1 | 14/1498_F_VP | SAMN07185545 | GCA_015817135.1 | NonPC | Public | 2010 | UK | Env | 89.8 | 100 | 0.03 | 98.1335 | 5185440 | 45.36 | 43 | 724548 |
| SAMN07185546-GCA_015817155.1 | 14/1072_D_VP | SAMN07185546 | GCA_015817155.1 | NonPC | Public | 2014 | UK | Env | 91.7 | 100 | 0.22 | 98.364 | 5148970 | 45.26 | 56 | 454097 |
| SAMN07185547-SRR5639917 | CFSAN029648 | SAMN07185547 | SRR5639917 | NonPC | Public | 2005 | UK | Clinical | 91.2 | 99.8 | 0 | 98.3426 | 5112336 | 45.29 | 53 | 239146 |
| SAMN07185548-GCA_015817095.1 | CFSAN029649 | SAMN07185548 | GCA_015817095.1 | NonPC | Public | 1980 | UK | Clinical | 92.6 | 100 | 0.65 | 98.4756 | 5043913 | 45.27 | 48 | 171546 |
| SAMN07185549-GCA_015817555.1 | CFSAN029650 | SAMN07185549 | GCA_015817555.1 | NonPC | Public | 1972 | UK | Clinical | 92.7 | 99.46 | 0 | 98.4228 | 5214412 | 45.17 | 50 | 208382 |
| SAMN07185550-GCA_015817425.1 | CFSAN029651 | SAMN07185550 | GCA_015817425.1 | PC | Public | 2010 | UK | Env | 99.9 | 100 | 0.14 | 99.996 | 5180933 | 45.35 | 13 | 1309403 |
| SAMN07267284-SRR5933709 | JN-7 | SAMN07267284 | SRR5933709 | NonPC | Public | 2014 | China | Env | 91.6 | 100 | 0.36 | 98.3312 | 5270349 | 45.33 | 37 | 815458 |
| SAMN07267286-GCF_003351885.1 | PB1937 | SAMN07267286 | GCF_003351885.1 | NonPC | Public | 2012 | China | Env | 92.1 | 97.68 | 0 | 98.2691 | 5580188 | 45.34 | 4 | 3567562 |
| SAMN07270562-SRR5933708 | LNM-3 | SAMN07270562 | SRR5933708 | NonPC | Public | 2014 | China | Env | 91.5 | 100 | 0.14 | 98.3445 | 5401479 | 45.12 | 66 | 459164 |
| SAMN07270668-SRR5933712 | DX-1 | SAMN07270668 | SRR5933712 | NonPC | Public | 2015 | China | Env | 91.1 | 100 | 0.3 | 98.2912 | 5404459 | 45.04 | 80 | 404265 |
| SAMN07279725-SRR5765325 | MAVP-55 | SAMN07279725 | SRR5765325 | NonPC | Public | 2013 | USA | Clinical | 91.7 | 100 | 0.29 | 98.2895 | 5252347 | 45.17 | 54 | 243658 |
| SAMN07279726-SRR5765297 | CTVP1C | SAMN07279726 | SRR5765297 | NonPC | Public | 2012 | USA | Clinical | 91.7 | 100 | 0.27 | 98.3057 | 5161941 | 45.17 | 58 | 254243 |
| SAMN07279727-SRR5765299 | CTVP28C | SAMN07279727 | SRR5765299 | NonPC | Public | 2013 | USA | Clinical | 89.9 | 100 | 0.27 | 98.332 | 5039237 | 45.37 | 41 | 280106 |
| SAMN07279728-SRR5765331 | MAVP-93 | SAMN07279728 | SRR5765331 | NonPC | Public | 2015 | USA | Clinical | 91.5 | 100 | 0 | 98.3719 | 5026563 | 45.35 | 42 | 277761 |
| SAMN07279729-SRR5765313 | CTVP11C | SAMN07279729 | SRR5765313 | NonPC | Public | 2012 | USA | Clinical | 91.6 | 100 | 0.03 | 98.2869 | 5179677 | 45.21 | 79 | 239711 |
| SAMN07279730-SRR5765323 | MAVP-98 | SAMN07279730 | SRR5765323 | NonPC | Public | 2015 | USA | Clinical | 90.8 | 100 | 0.22 | 98.2614 | 5130817 | 45.22 | 66 | 402060 |
| SAMN07279731-SRR5765306 | MAVP-F | SAMN07279731 | SRR5765306 | NonPC | Public | 2011 | USA | Clinical | 89.7 | 100 | 0 | 98.2727 | 4973927 | 45.33 | 70 | 375618 |
| SAMN07279732-SRR5765291 | MEVP13 | SAMN07279732 | SRR5765291 | NonPC | Public | 2015 | USA | Clinical | 89.9 | 100 | 0.22 | 98.305 | 5146123 | 45.18 | 43 | 284160 |
| SAMN07279733-SRR5765295 | MAVP-1 | SAMN07279733 | SRR5765295 | NonPC | Public | 2013 | USA | Clinical | 91.5 | 100 | 0 | 98.3064 | 5094069 | 45.27 | 59 | 269344 |
| SAMN07279734-SRR5765304 | MAVP-13 | SAMN07279734 | SRR5765304 | NonPC | Public | 2013 | USA | Clinical | 89.5 | 100 | 0.11 | 98.2943 | 4996097 | 45.36 | 29 | 621835 |
| SAMN07279735-SRR5765293 | CTVP37C | SAMN07279735 | SRR5765293 | NonPC | Public | 2013 | USA | Clinical | 90.1 | 100 | 0.12 | 98.3411 | 5020997 | 45.39 | 43 | 460037 |
| SAMN07279736-SRR5765298 | CTVP32C | SAMN07279736 | SRR5765298 | NonPC | Public | 2013 | USA | Clinical | 90.7 | 100 | 0 | 98.3671 | 5236918 | 45.27 | 43 | 243178 |

| | | | | | | | | | | | | | | | | |
|---|---|---|---|---|---|---|---|---|---|---|---|---|---|---|---|---|
| SAMN07279737-SRR5765326 | MAVP-66 | SAMN07279737 | SRR5765326 | NonPC | Public | 2014 | USA | Clinical | 90.8 | 100 | 0 | 98.3803 | 5127744 | 45.23 | 50 | 379827 |
| SAMN07279738-SRR5765309 | CTVP17C | SAMN07279738 | SRR5765309 | NonPC | Public | 2012 | USA | Clinical | 92 | 100 | 0.05 | 98.2618 | 5327298 | 45.28 | 51 | 276716 |
| SAMN07279739-SRR5765328 | MAVP-76 | SAMN07279739 | SRR5765328 | NonPC | Public | 2014 | USA | Clinical | 91.9 | 100 | 0.03 | 98.3781 | 5050087 | 45.38 | 56 | 393739 |
| SAMN07279740-SRR5765301 | MAVP-U | SAMN07279740 | SRR5765301 | NonPC | Public | 2011 | USA | Clinical | 92 | 100 | 0 | 98.3624 | 5136679 | 45.29 | 70 | 316978 |
| SAMN07279741-SRR5765315 | CT4264 | SAMN07279741 | SRR5765315 | NonPC | Public | 2013 | USA | Clinical | 89.9 | 100 | 0.05 | 98.3561 | 4966025 | 45.36 | 45 | 394585 |
| SAMN07279742-SRR5765311 | CTVP14C | SAMN07279742 | SRR5765311 | NonPC | Public | 2012 | USA | Clinical | 88.3 | 100 | 0.76 | 98.1267 | 5258215 | 45.21 | 168 | 57024 |
| SAMN07279743-SRR5765290 | MEVP15 | SAMN07279743 | SRR5765290 | NonPC | Public | 2015 | USA | Clinical | 91 | 100 | 0.2 | 98.2729 | 5090635 | 45.23 | 58 | 194816 |
| SAMN07279744-SRR5765302 | MAVP-S | SAMN07279744 | SRR5765302 | NonPC | Public | 2011 | USA | Clinical | 90.5 | 100 | 0.08 | 98.3179 | 5066459 | 45.32 | 409 | 22032 |
| SAMN07279745-SRR5765307 | MAVP-D | SAMN07279745 | SRR5765307 | NonPC | Public | 2011 | USA | Clinical | 91.3 | 100 | 0.11 | 98.2866 | 5156547 | 45.32 | 380 | 26020 |
| SAMN07279746-SRR5765324 | MAVP-53 | SAMN07279746 | SRR5765324 | NonPC | Public | 2013 | USA | Clinical | 91.2 | 100 | 0.08 | 98.3407 | 5043474 | 45.33 | 37 | 411087 |
| SAMN07279747-SRR5765312 | CTVP12C | SAMN07279747 | SRR5765312 | NonPC | Public | 2012 | USA | Clinical | 93 | 100 | 0.01 | 98.3727 | 5219568 | 45.25 | 54 | 264920 |
| SAMN07279748-SRR5765319 | MAVP-K | SAMN07279748 | SRR5765319 | NonPC | Public | 2011 | USA | Clinical | 91.4 | 100 | 0 | 98.3774 | 5126413 | 45.15 | 52 | 349568 |
| SAMN07279749-SRR5765322 | MAVP-95 | SAMN07279749 | SRR5765322 | NonPC | Public | 2015 | USA | Clinical | 90.4 | 100 | 0.05 | 98.2649 | 5084146 | 45.21 | 37 | 402417 |
| SAMN07279750-SRR5765330 | MAVP-91 | SAMN07279750 | SRR5765330 | NonPC | Public | 2015 | USA | Clinical | 91.3 | 100 | 0 | 98.409 | 5057906 | 45.31 | 68 | 168909 |
| SAMN07279751-SRR5765300 | MAVP-X | SAMN07279751 | SRR5765300 | NonPC | Public | 2011 | USA | Clinical | 91.6 | 100 | 0.08 | 98.3326 | 5123995 | 45.23 | 52 | 245246 |
| SAMN07279752-SRR5765294 | MAVP-10 | SAMN07279752 | SRR5765294 | NonPC | Public | 2013 | USA | Clinical | 90.6 | 100 | 0.4 | 98.2038 | 5099929 | 45.32 | 45 | 1139883 |
| SAMN07279753-SRR5765318 | MAVP-J | SAMN07279753 | SRR5765318 | NonPC | Public | 2011 | USA | Clinical | 91 | 100 | 0.22 | 98.333 | 5088064 | 45.26 | 47 | 453482 |
| SAMN07279754-SRR5765321 | MEVP7 | SAMN07279754 | SRR5765321 | NonPC | Public | 2014 | USA | Clinical | 91.5 | 100 | 0.08 | 98.2055 | 5073113 | 45.28 | 38 | 437196 |
| SAMN07279756-SRR5765329 | MAVP-87 | SAMN07279756 | SRR5765329 | NonPC | Public | 2015 | USA | Clinical | 89 | 100 | 0.09 | 98.1159 | 5088826 | 45.26 | 59 | 287273 |
| SAMN07279757-SRR5765292 | CTVP3C | SAMN07279757 | SRR5765292 | NonPC | Public | 2012 | USA | Clinical | 89.4 | 100 | 0.05 | 98.0981 | 5093219 | 45.25 | 90 | 136462 |
| SAMN07279759-SRR5765305 | MAVP110 | SAMN07279759 | SRR5765305 | NonPC | Public | 2015 | USA | Clinical | 89 | 100 | 0 | 98.0842 | 4946724 | 45.47 | 49 | 315099 |
| SAMN07279760-SRR5765316 | CT24E | SAMN07279760 | SRR5765316 | NonPC | Public | 2013 | USA | Clinical | 89.6 | 100 | 0 | 98.1113 | 4941073 | 45.37 | 86 | 240899 |
| SAMN07279761-SRR5765308 | CTVP19C | SAMN07279761 | SRR5765308 | NonPC | Public | 2013 | USA | Clinical | 90.3 | 100 | 0.14 | 98.2115 | 5096645 | 45.17 | 43 | 547108 |
| SAMN07279762-SRR5765314 | CTVP10C | SAMN07279762 | SRR5765314 | NonPC | Public | 2012 | USA | Clinical | 88.7 | 100 | 0.03 | 98.0433 | 5162186 | 45.18 | 52 | 262954 |
| SAMN07279763-SRR5765320 | MEVP6 | SAMN07279763 | SRR5765320 | NonPC | Public | 2013 | USA | Clinical | 87.5 | 100 | 0.93 | 98.0817 | 5075940 | 45.24 | 60 | 209098 |
| SAMN07279764-SRR5765296 | CTVP21C | SAMN07279764 | SRR5765296 | NonPC | Public | 2013 | USA | Clinical | 92 | 100 | 0 | 98.2894 | 5031788 | 45.33 | 61 | 209727 |
| SAMN07279765-SRR5765310 | CTVP15C | SAMN07279765 | SRR5765310 | NonPC | Public | 2012 | USA | Clinical | 91.4 | 100 | 0 | 98.3261 | 5096849 | 45.22 | 66 | 223319 |
| SAMN07279766-SRR5765327 | MAVP-73 | SAMN07279766 | SRR5765327 | NonPC | Public | 2014 | USA | Clinical | 91.1 | 100 | 0 | 98.4099 | 5094870 | 45.22 | 34 | 452362 |
| SAMN07281453-GCA_015817055.1 | PNUSAV000054 | SAMN07281453 | GCA_015817055.1 | NonPC | Public | NA | USA | Clinical | 90.9 | 100 | 0.33 | 98.2925 | 5254763 | 45.25 | 115 | 104723 |
| SAMN07291586-GCF_002807115.1 | MAVP398 | SAMN07291586 | GCF_002807115.1 | NonPC | Public | 2015 | USA | Env | 91.2 | 100 | 0.27 | 98.2903 | 5087559 | 45.38 | 72 | 148777 |
| SAMN07327345-GCA_015816975.1 | PNUSAV000060 | SAMN07327345 | GCA_015816975.1 | PC | Public | NA | USA | Clinical | 97.9 | 100 | 0 | 99.9715 | 5161481 | 45.23 | 39 | 401520 |
| SAMN07327346-GCA_015816995.1 | PNUSAV000061 | SAMN07327346 | GCA_015816995.1 | PC | Public | NA | USA | Clinical | 97.9 | 100 | 0 | 99.97 | 5171460 | 45.21 | 45 | 314443 |
| SAMN07337843-GCF_006378735.1 | A1_1 | SAMN07337843 | GCF_006378735.1 | NonPC | Public | 2014 | China | Env | 91.5 | 100 | 0.28 | 98.337 | 5203868 | 45.26 | 77 | 306101 |
| SAMN07337845-GCF_006378875.1 | A1_3 | SAMN07337845 | GCF_006378875.1 | NonPC | Public | 2014 | China | Env | 90.5 | 100 | 0.27 | 98.3408 | 5068122 | 45.33 | 165 | 89247 |
| SAMN07337846-GCF_006378705.1 | A1_4 | SAMN07337846 | GCF_006378705.1 | NonPC | Public | 2014 | China | Env | 91.9 | 100 | 2.94 | 98.3215 | 5311389 | 45.26 | 88 | 172628 |
| SAMN07337847-GCF_006378865.1 | A1_6 | SAMN07337847 | GCF_006378865.1 | NonPC | Public | 2014 | China | Env | 89.6 | 100 | 0.03 | 98.3518 | 5008413 | 45.44 | 63 | 239655 |
| SAMN07337848-GCF_006379005.1 | A1_7 | SAMN07337848 | GCF_006379005.1 | NonPC | Public | 2014 | China | Env | 90.6 | 100 | 0.32 | 98.2942 | 5088332 | 45.27 | 66 | 292306 |
| SAMN07337849-GCF_006378995.1 | A1_8 | SAMN07337849 | GCF_006378995.1 | NonPC | Public | 2014 | China | Env | 90.6 | 100 | 0.61 | 98.2857 | 5222852 | 45.36 | 243 | 79348 |
| SAMN07337850-GCF_006378975.1 | A1_9 | SAMN07337850 | GCF_006378975.1 | NonPC | Public | 2014 | China | Env | 91.1 | 100 | 0.28 | 98.3958 | 5122274 | 45.43 | 216 | 66604 |
| SAMN07337851-GCF_006378675.1 | A2_1 | SAMN07337851 | GCF_006378675.1 | NonPC | Public | 2014 | China | Env | 90.7 | 100 | 0.05 | 98.338 | 5002823 | 45.32 | 80 | 210773 |
| SAMN07337852-GCF_006378855.1 | A2_2 | SAMN07337852 | GCF_006378855.1 | NonPC | Public | 2014 | China | Env | 89.1 | 99.84 | 0.14 | 98.2443 | 4960574 | 45.34 | 67 | 232916 |
| SAMN07337853-GCF_006378945.1 | A2_3 | SAMN07337853 | GCF_006378945.1 | NonPC | Public | 2014 | China | Env | 90.2 | 100 | 0.27 | 98.4114 | 4853953 | 45.54 | 250 | 49362 |
| SAMN07337854-GCF_006378955.1 | A2_4 | SAMN07337854 | GCF_006378955.1 | NonPC | Public | 2014 | China | Env | 89.9 | 100 | 0.27 | 98.2831 | 5004134 | 45.46 | 117 | 155622 |
| SAMN07337855-GCF_006378615.1 | A2_5 | SAMN07337855 | GCF_006378615.1 | NonPC | Public | 2014 | China | Env | 90.9 | 100 | 0 | 98.3688 | 5109041 | 45.26 | 133 | 134408 |
| SAMN07337856-GCF_006378935.1 | A2_6 | SAMN07337856 | GCF_006378935.1 | NonPC | Public | 2014 | China | Env | 89.8 | 100 | 0.05 | 98.3478 | 5034518 | 45.33 | 66 | 373545 |
| SAMN07337857-GCF_006378635.1 | A2_7 | SAMN07337857 | GCF_006378635.1 | NonPC | Public | 2014 | China | Env | 89.7 | 100 | 0.12 | 98.2937 | 4975875 | 45.46 | 297 | 58872 |
| SAMN07337858-GCF_006378835.1 | A2_8 | SAMN07337858 | GCF_006378835.1 | NonPC | Public | 2014 | China | Env | 88.7 | 99.73 | 0.11 | 98.4089 | 4877832 | 45.62 | 427 | 24928 |
| SAMN07337859-GCF_006377205.1 | A2_9 | SAMN07337859 | GCF_006377205.1 | NonPC | Public | 2014 | China | Env | 90.8 | 100 | 2.27 | 98.2282 | 5277957 | 45.35 | 188 | 74182 |

| | | | | | | | | | | | | | | | | |
|---|---|---|---|---|---|---|---|---|---|---|---|---|---|---|---|---|
| SAMN07337860-GCF_006378795.1 | A3_2 | SAMN07337860 | GCF_006378795.1 | NonPC | Public | 2014 | China | Env | 87.4 | 99.18 | 0.76 | 98.3841 | 4854663 | 45.81 | 791 | 12290 |
| SAMN07337861-GCF_006378785.1 | A3_3 | SAMN07337861 | GCF_006378785.1 | NonPC | Public | 2014 | China | Env | 88.5 | 100 | 0.36 | 98.2739 | 5121653 | 45.19 | 96 | 221668 |
| SAMN07337862-GCF_006378805.1 | A3_4 | SAMN07337862 | GCF_006378805.1 | NonPC | Public | 2014 | China | Env | 91.9 | 100 | 0.24 | 98.3285 | 5238807 | 45.24 | 92 | 190516 |
| SAMN07337863-GCF_006381695.1 | A3_5 | SAMN07337863 | GCF_006381695.1 | NonPC | Public | 2014 | China | Env | 90.8 | 100 | 0.41 | 98.197 | 5228982 | 45.33 | 190 | 76032 |
| SAMN07337864-GCF_006377575.1 | A3_6 | SAMN07337864 | GCF_006377575.1 | NonPC | Public | 2014 | China | Env | 90.9 | 100 | 0.35 | 98.343 | 5234000 | 45.17 | 145 | 85592 |
| SAMN07337865-GCF_006378655.1 | A3_7 | SAMN07337865 | GCF_006378655.1 | NonPC | Public | 2014 | China | Env | 87 | 99.93 | 0.37 | 98.2937 | 4912151 | 45.6 | 564 | 20109 |
| SAMN07337866-GCF_006378765.1 | A3_8 | SAMN07337866 | GCF_006378765.1 | NonPC | Public | 2014 | China | Env | 89.2 | 100 | 0.13 | 98.3884 | 5101119 | 45.44 | 287 | 52918 |
| SAMN07337867-GCF_006377345.1 | A3_9 | SAMN07337867 | GCF_006377345.1 | NonPC | Public | 2014 | China | Env | 88.9 | 100 | 0.54 | 98.3774 | 4896659 | 45.47 | 252 | 43539 |
| SAMN07337868-GCF_006377565.1 | A4_1 | SAMN07337868 | GCF_006377565.1 | NonPC | Public | 2014 | China | Env | 91 | 100 | 0 | 98.425 | 5047138 | 45.49 | 442 | 29062 |
| SAMN07337869-GCF_006377545.1 | A4_2 | SAMN07337869 | GCF_006377545.1 | NonPC | Public | 2014 | China | Env | 89.4 | 100 | 0.14 | 98.3752 | 4975060 | 45.41 | 428 | 30894 |
| SAMN07337870-GCF_006377285.1 | A4_3 | SAMN07337870 | GCF_006377285.1 | NonPC | Public | 2014 | China | Env | 89.2 | 99.73 | 0 | 98.3146 | 4942225 | 45.59 | 247 | 44582 |
| SAMN07337871-GCF_006377305.1 | A4_4 | SAMN07337871 | GCF_006377305.1 | NonPC | Public | 2014 | China | Env | 87.9 | 99.64 | 0.41 | 98.4247 | 4814419 | 45.68 | 547 | 18657 |
| SAMN07337872-GCF_006377195.1 | A4_5 | SAMN07337872 | GCF_006377195.1 | NonPC | Public | 2014 | China | Env | 89.8 | 100 | 0.11 | 98.2954 | 5073945 | 45.33 | 57 | 283243 |
| SAMN07337873-GCF_006377295.1 | A4_6 | SAMN07337873 | GCF_006377295.1 | NonPC | Public | 2014 | China | Env | 90.6 | 100 | 0.11 | 98.3516 | 4997170 | 45.35 | 71 | 347376 |
| SAMN07337874-GCF_006377535.1 | A4_9 | SAMN07337874 | GCF_006377535.1 | NonPC | Public | 2014 | China | Env | 88.2 | 99.93 | 0.33 | 98.2615 | 5150692 | 45.3 | 55 | 285202 |
| SAMN07337875-GCF_006377495.1 | A5_1 | SAMN07337875 | GCF_006377495.1 | NonPC | Public | 2014 | China | Env | 89.5 | 99.95 | 0.1 | 98.3376 | 5151481 | 45.29 | 77 | 399240 |
| SAMN07337876-GCF_006377265.1 | A5_3 | SAMN07337876 | GCF_006377265.1 | NonPC | Public | 2014 | China | Env | 89.4 | 100 | 0.03 | 98.2981 | 4984474 | 45.33 | 111 | 192988 |
| SAMN07337877-GCF_006377145.1 | A5_4 | SAMN07337877 | GCF_006377145.1 | NonPC | Public | 2014 | China | Env | 91.1 | 100 | 0 | 98.361 | 5046602 | 45.43 | 162 | 88293 |
| SAMN07337878-GCF_006377475.1 | A5_7 | SAMN07337878 | GCF_006377475.1 | NonPC | Public | 2014 | China | Env | 90.4 | 100 | 0.69 | 98.323 | 5278218 | 45.39 | 222 | 67729 |
| SAMN07337879-GCF_006377105.1 | A5_8 | SAMN07337879 | GCF_006377105.1 | NonPC | Public | 2014 | China | Env | 89.2 | 100 | 2.78 | 98.2714 | 5074167 | 45.44 | 168 | 75935 |
| SAMN07337880-GCF_006377455.1 | B1_1 | SAMN07337880 | GCF_006377455.1 | NonPC | Public | 2014 | China | Env | 87.1 | 100 | 0.5 | 98.3233 | 4984628 | 45.41 | 340 | 34436 |
| SAMN07337881-GCF_006377085.1 | B1_10 | SAMN07337881 | GCF_006377085.1 | NonPC | Public | 2014 | China | Env | 89.4 | 100 | 0.02 | 98.3577 | 5008345 | 45.33 | 57 | 385458 |
| SAMN07337882-GCF_006377395.1 | B1_3 | SAMN07337882 | GCF_006377395.1 | NonPC | Public | 2014 | China | Env | 90.5 | 100 | 0.03 | 98.3399 | 4950089 | 45.4 | 109 | 91420 |
| SAMN07337883-GCF_006377385.1 | B1_4 | SAMN07337883 | GCF_006377385.1 | NonPC | Public | 2014 | China | Env | 89.1 | 99.73 | 0.03 | 98.3871 | 4935609 | 45.65 | 640 | 17133 |
| SAMN07337884-GCF_006377115.1 | B1_5 | SAMN07337884 | GCF_006377115.1 | NonPC | Public | 2014 | China | Env | 90.2 | 100 | 0.17 | 98.3297 | 5092771 | 45.39 | 356 | 37807 |
| SAMN07337885-GCF_006377405.1 | B1_6 | SAMN07337885 | GCF_006377405.1 | NonPC | Public | 2014 | China | Env | 90.7 | 100 | 0.03 | 98.3256 | 5099984 | 45.23 | 128 | 151448 |
| SAMN07337886-GCF_006377445.1 | B1_7 | SAMN07337886 | GCF_006377445.1 | NonPC | Public | 2014 | China | Env | 89.7 | 100 | 0.5 | 98.2799 | 4973507 | 45.31 | 78 | 370957 |
| SAMN07337887-GCF_006377225.1 | B1_8 | SAMN07337887 | GCF_006377225.1 | NonPC | Public | 2014 | China | Env | 89.2 | 100 | 0.68 | 98.3316 | 5004110 | 45.64 | 335 | 34917 |
| SAMN07337888-GCF_006377095.1 | B1_9 | SAMN07337888 | GCF_006377095.1 | NonPC | Public | 2014 | China | Env | 89.8 | 100 | 0.37 | 98.3965 | 5038868 | 45.48 | 488 | 24009 |
| SAMN07337889-GCF_006377075.1 | B10_1 | SAMN07337889 | GCF_006377075.1 | NonPC | Public | 2014 | China | Env | 87.5 | 100 | 0.33 | 98.3629 | 4990493 | 45.49 | 496 | 24772 |
| SAMN07337890-GCF_006377375.1 | B10_2 | SAMN07337890 | GCF_006377375.1 | NonPC | Public | 2014 | China | Env | 87.8 | 100 | 0.06 | 98.3803 | 4803854 | 45.71 | 584 | 17105 |
| SAMN07337891-GCF_006377025.1 | B10_3 | SAMN07337891 | GCF_006377025.1 | NonPC | Public | 2014 | China | Env | 88.7 | 99.73 | 0.05 | 98.3746 | 4890739 | 45.62 | 394 | 26939 |
| SAMN07337892-GCF_006376835.1 | B10_4 | SAMN07337892 | GCF_006376835.1 | NonPC | Public | 2014 | China | Env | 89.4 | 100 | 0.4 | 98.3545 | 4996407 | 45.34 | 287 | 45220 |
| SAMN07337893-GCA_006377235.1 | B10_6 | SAMN07337893 | GCA_006377235.1 | NonPC | Public | 2014 | China | Env | 87.7 | 97.91 | 1.63 | 98.497 | 4922276 | 45.62 | 1324 | 6725 |
| SAMN07337894-GCF_006376815.1 | B10_7 | SAMN07337894 | GCF_006376815.1 | NonPC | Public | 2014 | China | Env | 87.9 | 99.23 | 2.82 | 98.4581 | 4918638 | 45.76 | 1135 | 8563 |
| SAMN07337895-GCF_006376995.1 | B10_8 | SAMN07337895 | GCF_006376995.1 | NonPC | Public | 2014 | China | Env | 91.1 | 100 | 0.11 | 98.3235 | 5122896 | 45.35 | 151 | 90934 |
| SAMN07337896-GCF_006376975.1 | B10_9 | SAMN07337896 | GCF_006376975.1 | NonPC | Public | 2014 | China | Env | 89.3 | 99.45 | 1.92 | 98.34 | 5222078 | 45.24 | 610 | 16787 |
| SAMN07337897-GCF_006376765.1 | B2_1 | SAMN07337897 | GCF_006376765.1 | NonPC | Public | 2014 | China | Env | 90.1 | 99.93 | 0.03 | 98.361 | 5077025 | 45.42 | 387 | 39154 |
| SAMN07337898-GCF_006376775.1 | B2_10 | SAMN07337898 | GCF_006376775.1 | NonPC | Public | 2014 | China | Env | 91.1 | 99.96 | 0.03 | 98.4642 | 4951412 | 45.34 | 89 | 125674 |
| SAMN07337899-GCF_006377005.1 | B2_2 | SAMN07337899 | GCF_006377005.1 | NonPC | Public | 2014 | China | Env | 88.2 | 100 | 0.3 | 98.1687 | 5132468 | 45.18 | 127 | 124621 |
| SAMN07337900-GCF_006376915.1 | B2_3 | SAMN07337900 | GCF_006376915.1 | NonPC | Public | 2014 | China | Env | 88.2 | 100 | 0.49 | 98.1116 | 5240841 | 44.98 | 97 | 332368 |
| SAMN07337901-GCF_006376895.1 | B2_4 | SAMN07337901 | GCF_006376895.1 | NonPC | Public | 2014 | China | Env | 87.8 | 100 | 0.49 | 98.2547 | 5089823 | 45.28 | 456 | 37262 |
| SAMN07337902-GCF_006376755.1 | B2_5 | SAMN07337902 | GCF_006376755.1 | NonPC | Public | 2014 | China | Env | 90.3 | 99.86 | 0.03 | 98.3803 | 5054517 | 45.4 | 348 | 39312 |
| SAMN07337903-GCF_006376985.1 | B2_6 | SAMN07337903 | GCF_006376985.1 | NonPC | Public | 2014 | China | Env | 89 | 100 | 0.16 | 98.3865 | 4891197 | 45.56 | 219 | 53802 |
| SAMN07337904-GCF_006375565.1 | B2_7 | SAMN07337904 | GCF_006375565.1 | NonPC | Public | 2014 | China | Env | 89.5 | 100 | 0.21 | 98.3149 | 4958838 | 45.39 | 55 | 314869 |
| SAMN07337905-GCF_006376905.1 | B2_8 | SAMN07337905 | GCF_006376905.1 | NonPC | Public | 2014 | China | Env | 87.8 | 99.6 | 0.27 | 98.454 | 4725461 | 45.79 | 513 | 20407 |
| SAMN07337906-GCF_006376745.1 | B2_9 | SAMN07337906 | GCF_006376745.1 | NonPC | Public | 2014 | China | Env | 88.7 | 99.49 | 1.84 | 98.3934 | 4915368 | 45.64 | 864 | 12111 |
| SAMN07337907-GCF_006375555.1 | B3_1 | SAMN07337907 | GCF_006375555.1 | NonPC | Public | 2014 | China | Env | 86.5 | 99.39 | 1.2 | 98.3887 | 4790641 | 45.66 | 610 | 18715 |

| | | | | | | | | | | | | | | | | |
|---|---|---|---|---|---|---|---|---|---|---|---|---|---|---|---|---|
| SAMN07337908-GCF_006376705.1 | B3_10 | SAMN07337908 | GCF_006376705.1 | NonPC | Public | 2014 | China | Env | 89.8 | 99.76 | 0.2 | 98.493 | 4747699 | 45.71 | 647 | 14165 |
| SAMN07337909-GCF_006375285.1 | B3_2 | SAMN07337909 | GCF_006375285.1 | NonPC | Public | 2014 | China | Env | 90.5 | 100 | 0.12 | 98.3213 | 5145197 | 45.29 | 227 | 67629 |
| SAMN07337910-GCF_006376875.1 | B3_3 | SAMN07337910 | GCF_006376875.1 | NonPC | Public | 2014 | China | Env | 90.8 | 100 | 0.03 | 98.3225 | 5204109 | 45.18 | 106 | 207745 |
| SAMN07337911-GCF_006375525.1 | B3_4 | SAMN07337911 | GCF_006375525.1 | NonPC | Public | 2014 | China | Env | 86.5 | 99.39 | 0.58 | 98.3321 | 4881401 | 45.65 | 771 | 13931 |
| SAMN07337912-GCF_006375515.1 | B3_5 | SAMN07337912 | GCF_006375515.1 | NonPC | Public | 2014 | China | Env | 88.9 | 99.63 | 0.41 | 98.4876 | 4809173 | 45.78 | 714 | 14629 |
| SAMN07337913-GCF_006376855.1 | B3_6 | SAMN07337913 | GCF_006376855.1 | NonPC | Public | 2014 | China | Env | 87.3 | 98.63 | 0.67 | 98.4523 | 4739626 | 45.88 | 939 | 9740 |
| SAMN07337914-GCF_006375495.1 | B3_7 | SAMN07337914 | GCF_006375495.1 | NonPC | Public | 2014 | China | Env | 89.7 | 99.68 | 0.33 | 98.4681 | 4778575 | 45.75 | 595 | 16864 |
| SAMN07337916-GCF_006375275.1 | B3_9 | SAMN07337916 | GCF_006375275.1 | NonPC | Public | 2014 | China | Env | 90.4 | 100 | 0 | 98.3954 | 5044160 | 45.42 | 242 | 42957 |
| SAMN07337917-GCA_006375435.1 | B4_1 | SAMN07337917 | GCA_006375435.1 | NonPC | Public | 2014 | China | Env | 88.3 | 99.37 | 3.51 | 98.484 | 4920093 | 45.77 | 1033 | 9148 |
| SAMN07337918-GCF_006375455.1 | B4_10 | SAMN07337918 | GCF_006375455.1 | NonPC | Public | 2014 | China | Env | 89.1 | 99.88 | 0.94 | 98.4913 | 4797573 | 45.87 | 891 | 9622 |
| SAMN07337919-GCF_006375445.1 | B4_2 | SAMN07337919 | GCF_006375445.1 | NonPC | Public | 2014 | China | Env | 89.7 | 99.73 | 0.16 | 98.4024 | 5045729 | 45.48 | 588 | 19617 |
| SAMN07337920-GCF_006375385.1 | B4_3 | SAMN07337920 | GCF_006375385.1 | NonPC | Public | 2014 | China | Env | 87 | 99.59 | 0.65 | 98.4255 | 4801971 | 45.62 | 754 | 14640 |
| SAMN07337921-GCF_006375405.1 | B4_5 | SAMN07337921 | GCF_006375405.1 | NonPC | Public | 2014 | China | Env | 91.8 | 100 | 0.05 | 98.2932 | 5176546 | 45.19 | 105 | 205247 |
| SAMN07337922-GCF_006381675.1 | B4_6 | SAMN07337922 | GCF_006381675.1 | NonPC | Public | 2014 | China | Env | 88.9 | 99.33 | 1.3 | 98.4187 | 4912809 | 45.63 | 870 | 11480 |
| SAMN07337923-GCF_006375395.1 | B4_7 | SAMN07337923 | GCF_006375395.1 | NonPC | Public | 2014 | China | Env | 87.3 | 98.16 | 1.34 | 98.4401 | 4770595 | 45.84 | 1059 | 7926 |
| SAMN07337924-GCF_006375215.1 | B4_8 | SAMN07337924 | GCF_006375215.1 | NonPC | Public | 2014 | China | Env | 88.3 | 99.44 | 1.54 | 98.4818 | 4629416 | 45.93 | 933 | 9067 |
| SAMN07337925-GCF_006375195.1 | B4_9 | SAMN07337925 | GCF_006375195.1 | NonPC | Public | 2014 | China | Env | 88.7 | 99.52 | 0.07 | 98.4289 | 4919261 | 45.71 | 713 | 14218 |
| SAMN07337926-GCF_006375345.1 | B5_1 | SAMN07337926 | GCF_006375345.1 | NonPC | Public | 2014 | China | Env | 90.6 | 100 | 0.03 | 98.3024 | 5144426 | 45.31 | 277 | 50460 |
| SAMN07337927-GCF_006375075.1 | B5_10 | SAMN07337927 | GCF_006375075.1 | NonPC | Public | 2014 | China | Env | 91.4 | 100 | 1.49 | 98.3285 | 5283253 | 45.26 | 62 | 205652 |
| SAMN07337928-GCF_006374755.1 | B5_2 | SAMN07337928 | GCF_006374755.1 | NonPC | Public | 2014 | China | Env | 87.7 | 100 | 0.46 | 98.2935 | 4930158 | 45.41 | 505 | 21492 |
| SAMN07337929-GCF_006375205.1 | B5_3 | SAMN07337929 | GCF_006375205.1 | NonPC | Public | 2014 | China | Env | 86.4 | 98.24 | 2.42 | 98.3434 | 4858447 | 45.62 | 1160 | 8423 |
| SAMN07337930-GCF_006375335.1 | B5_4 | SAMN07337930 | GCF_006375335.1 | NonPC | Public | 2014 | China | Env | 89.8 | 99.67 | 1.01 | 98.4053 | 5082046 | 45.41 | 166 | 119223 |
| SAMN07337931-GCF_006375325.1 | B5_5 | SAMN07337931 | GCF_006375325.1 | NonPC | Public | 2014 | China | Env | 88.7 | 99.25 | 1 | 98.4793 | 4897295 | 45.65 | 871 | 11729 |
| SAMN07337932-GCF_006375035.1 | B5_6 | SAMN07337932 | GCF_006375035.1 | NonPC | Public | 2014 | China | Env | 90.7 | 100 | 0.08 | 98.4125 | 5142554 | 45.18 | 240 | 51305 |
| SAMN07337933-GCF_006375125.1 | B5_8 | SAMN07337933 | GCF_006375125.1 | NonPC | Public | 2014 | China | Env | 90.8 | 100 | 0.03 | 98.2947 | 5197684 | 45.17 | 107 | 239868 |
| SAMN07337934-GCF_006375045.1 | B5_9 | SAMN07337934 | GCF_006375045.1 | NonPC | Public | 2014 | China | Env | 90.4 | 99.73 | 4.96 | 98.3109 | 5318074 | 45.57 | 713 | 19000 |
| SAMN07337935-GCF_006374465.1 | B6_1 | SAMN07337935 | GCF_006374465.1 | NonPC | Public | 2014 | China | Env | 89.8 | 99.66 | 0.16 | 98.4657 | 4928461 | 45.68 | 666 | 18524 |
| SAMN07337936-GCF_006375175.1 | B6_10 | SAMN07337936 | GCF_006375175.1 | NonPC | Public | 2014 | China | Env | 86.9 | 99.34 | 0.05 | 98.4443 | 4697519 | 45.95 | 808 | 10737 |
| SAMN07337937-GCF_006375115.1 | B6_2 | SAMN07337937 | GCF_006375115.1 | NonPC | Public | 2014 | China | Env | 90.5 | 99.73 | 0.54 | 98.41 | 4974810 | 45.47 | 307 | 49233 |
| SAMN07337938-GCF_006375005.1 | B6_3 | SAMN07337938 | GCF_006375005.1 | NonPC | Public | 2014 | China | Env | 91 | 100 | 0 | 98.4727 | 4911180 | 45.54 | 252 | 45160 |
| SAMN07337939-GCF_006375165.1 | B6_4 | SAMN07337939 | GCF_006375165.1 | NonPC | Public | 2014 | China | Env | 89.6 | 100 | 0.14 | 98.3309 | 5278012 | 45.43 | 179 | 71142 |
| SAMN07337940-GCF_006375315.1 | B6_5 | SAMN07337940 | GCF_006375315.1 | NonPC | Public | 2014 | China | Env | 89.8 | 100 | 2.93 | 98.3484 | 5338893 | 45.3 | 89 | 296900 |
| SAMN07337941-GCF_006374415.1 | B6_6 | SAMN07337941 | GCF_006374415.1 | NonPC | Public | 2014 | China | Env | 90 | 100 | 0.47 | 98.4552 | 5009435 | 45.62 | 441 | 24183 |
| SAMN07337942-GCF_006375025.1 | B6_7 | SAMN07337942 | GCF_006375025.1 | NonPC | Public | 2014 | China | Env | 87.9 | 98.79 | 2.36 | 98.4289 | 5007923 | 45.57 | 1049 | 10526 |
| SAMN07337943-GCF_006374735.1 | B6_8 | SAMN07337943 | GCF_006374735.1 | NonPC | Public | 2014 | China | Env | 90.5 | 100 | 0.14 | 98.3454 | 5108026 | 45.5 | 385 | 29413 |
| SAMN07337944-GCF_006374995.1 | B6_9 | SAMN07337944 | GCF_006374995.1 | NonPC | Public | 2014 | China | Env | 89.7 | 100 | 0.09 | 98.2999 | 4900626 | 45.47 | 207 | 67408 |
| SAMN07337945-GCF_006374955.1 | B7_2 | SAMN07337945 | GCF_006374955.1 | NonPC | Public | 2014 | China | Env | 89.3 | 100 | 0.28 | 98.316 | 5146393 | 45.39 | 171 | 67446 |
| SAMN07337946-GCF_006374405.1 | B7_3 | SAMN07337946 | GCF_006374405.1 | NonPC | Public | 2014 | China | Env | 87.9 | 99.3 | 0.42 | 98.4893 | 4809180 | 45.91 | 892 | 10009 |
| SAMN07337947-GCF_006374925.1 | B7_4 | SAMN07337947 | GCF_006374925.1 | NonPC | Public | 2014 | China | Env | 89.7 | 99.86 | 0.91 | 98.3588 | 4901133 | 45.33 | 355 | 33382 |
| SAMN07337948-GCF_006374915.1 | B7_5 | SAMN07337948 | GCF_006374915.1 | NonPC | Public | 2014 | China | Env | 89.4 | 100 | 0.6 | 98.2666 | 5126436 | 45.22 | 141 | 102525 |
| SAMN07337949-GCF_006374895.1 | B7_6 | SAMN07337949 | GCF_006374895.1 | NonPC | Public | 2014 | China | Env | 88.3 | 99.73 | 0.27 | 98.3691 | 4804734 | 45.75 | 464 | 21982 |
| SAMN07337950-GCF_006374395.1 | B7_7 | SAMN07337950 | GCF_006374395.1 | NonPC | Public | 2014 | China | Env | 88.9 | 99.73 | 0 | 98.3938 | 4793079 | 45.65 | 392 | 30119 |
| SAMN07337951-GCF_006374375.1 | B8_1 | SAMN07337951 | GCF_006374375.1 | NonPC | Public | 2014 | China | Env | 88.1 | 100 | 0.27 | 98.4396 | 4805810 | 45.72 | 527 | 21955 |
| SAMN07337952-GCF_006374745.1 | B8_10 | SAMN07337952 | GCF_006374745.1 | NonPC | Public | 2014 | China | Env | 89.9 | 100 | 0.03 | 98.3603 | 4969576 | 45.39 | 87 | 179359 |
| SAMN07337953-GCF_006374675.1 | B8_2 | SAMN07337953 | GCF_006374675.1 | NonPC | Public | 2014 | China | Env | 89.6 | 100 | 0.25 | 98.3158 | 5037585 | 45.33 | 90 | 131067 |
| SAMN07337954-GCF_006374695.1 | B8_3 | SAMN07337954 | GCF_006374695.1 | NonPC | Public | 2014 | China | Env | 89.2 | 100 | 0.27 | 98.3885 | 4939101 | 45.53 | 219 | 56076 |
| SAMN07337955-GCF_006374665.1 | B8_4 | SAMN07337955 | GCF_006374665.1 | NonPC | Public | 2014 | China | Env | 90.7 | 100 | 0.07 | 98.3424 | 5006518 | 45.34 | 89 | 295600 |
| SAMN07337956-GCF_006374355.1 | B8_5 | SAMN07337956 | GCF_006374355.1 | NonPC | Public | 2014 | China | Env | 90.6 | 100 | 0.09 | 98.4153 | 4933921 | 45.43 | 386 | 37463 |

| | | | | | | | | | | | | | | | | |
|---|---|---|---|---|---|---|---|---|---|---|---|---|---|---|---|---|
| SAMN07337957-GCF_006374875.1 | B8_6 | SAMN07337957 | GCF_006374875.1 | NonPC | Public | 2014 | China | Env | 91.8 | 100 | 0.58 | 98.3137 | 5266595 | 45.1 | 78 | 256928 |
| SAMN07337958-GCF_006374635.1 | B8_7 | SAMN07337958 | GCF_006374635.1 | NonPC | Public | 2014 | China | Env | 90.8 | 99.73 | 0 | 98.4363 | 4906026 | 45.55 | 330 | 41649 |
| SAMN07337959-GCF_006374605.1 | B8_8 | SAMN07337959 | GCF_006374605.1 | NonPC | Public | 2014 | China | Env | 88.2 | 100 | 0.3 | 98.1935 | 5176160 | 45.13 | 101 | 203599 |
| SAMN07337960-GCF_006374595.1 | B8_9 | SAMN07337960 | GCF_006374595.1 | NonPC | Public | 2014 | China | Env | 91.2 | 99.73 | 0 | 98.4413 | 4891724 | 45.57 | 243 | 49866 |
| SAMN07337961-GCF_006374585.1 | B9_1 | SAMN07337961 | GCF_006374585.1 | NonPC | Public | 2014 | China | Env | 89 | 100 | 0.27 | 98.3321 | 5233469 | 45.48 | 285 | 49811 |
| SAMN07337962-GCF_006374335.1 | B9_10 | SAMN07337962 | GCF_006374335.1 | NonPC | Public | 2014 | China | Env | 91.1 | 100 | 0 | 98.3811 | 5006523 | 45.32 | 170 | 72226 |
| SAMN07337963-GCF_006374555.1 | B9_2 | SAMN07337963 | GCF_006374555.1 | NonPC | Public | 2014 | China | Env | 86.2 | 99.41 | 0.67 | 98.4044 | 4839358 | 45.71 | 703 | 15834 |
| SAMN07337964-GCF_006374865.1 | B9_3 | SAMN07337964 | GCF_006374865.1 | NonPC | Public | 2014 | China | Env | 89.2 | 99.59 | 2.3 | 98.3439 | 4955129 | 45.32 | 78 | 419729 |
| SAMN07337965-GCF_006374845.1 | B9_4 | SAMN07337965 | GCF_006374845.1 | NonPC | Public | 2014 | China | Env | 89.8 | 99.73 | 0.27 | 98.46 | 4894798 | 45.65 | 528 | 21278 |
| SAMN07337966-GCF_006374815.1 | B9_5 | SAMN07337966 | GCF_006374815.1 | NonPC | Public | 2014 | China | Env | 87.8 | 100 | 0.3 | 98.2063 | 5064354 | 45.34 | 223 | 61434 |
| SAMN07337967-GCF_006374345.1 | B9_6 | SAMN07337967 | GCF_006374345.1 | NonPC | Public | 2014 | China | Env | 89.1 | 100 | 0.32 | 98.3085 | 4946747 | 45.43 | 143 | 81111 |
| SAMN07337968-GCF_006374545.1 | B9_7 | SAMN07337968 | GCF_006374545.1 | NonPC | Public | 2014 | China | Env | 90.1 | 100 | 0.05 | 98.2851 | 5169839 | 45.18 | 80 | 258756 |
| SAMN07337969-GCF_006374805.1 | B9_8 | SAMN07337969 | GCF_006374805.1 | NonPC | Public | 2014 | China | Env | 88.2 | 100 | 2.93 | 98.2911 | 5170531 | 45.41 | 95 | 120421 |
| SAMN07337970-GCF_006374505.1 | B9_9 | SAMN07337970 | GCF_006374505.1 | NonPC | Public | 2014 | China | Env | 91 | 99.93 | 0 | 98.3571 | 5015269 | 45.29 | 156 | 82409 |
| SAMN07337971-GCF_006374305.1 | C1_10 | SAMN07337971 | GCF_006374305.1 | PrePC | Public | 2014 | China | Env | 94.6 | 100 | 0.14 | 99.8158 | 5194622 | 45.23 | 66 | 340500 |
| SAMN07337972-GCF_006374285.1 | C1_2 | SAMN07337972 | GCF_006374285.1 | NonPC | Public | 2014 | China | Env | 89.6 | 100 | 0.05 | 98.3673 | 5002189 | 45.36 | 103 | 373804 |
| SAMN07337973-GCF_006374255.1 | C1_3 | SAMN07337973 | GCF_006374255.1 | NonPC | Public | 2014 | China | Env | 88 | 99.84 | 1.08 | 98.1961 | 5078479 | 45.21 | 310 | 42326 |
| SAMN07337974-GCF_006374495.1 | C1_4 | SAMN07337974 | GCF_006374495.1 | NonPC | Public | 2014 | China | Env | 86 | 99.62 | 0.67 | 98.4074 | 4727376 | 45.83 | 671 | 14024 |
| SAMN07337975-GCF_006374245.1 | C1_5 | SAMN07337975 | GCF_006374245.1 | NonPC | Public | 2014 | China | Env | 91.3 | 100 | 0.1 | 98.432 | 4945657 | 45.39 | 77 | 309705 |
| SAMN07337977-GCF_006374775.1 | C1_7 | SAMN07337977 | GCF_006374775.1 | NonPC | Public | 2014 | China | Env | 91.5 | 100 | 0.94 | 98.3116 | 5244361 | 45.17 | 89 | 248339 |
| SAMN07337978-GCF_006373735.1 | C1_8 | SAMN07337978 | GCF_006373735.1 | NonPC | Public | 2014 | China | Env | 91.6 | 100 | 1.15 | 98.3378 | 5283762 | 45.17 | 101 | 248348 |
| SAMN07337979-GCF_006374225.1 | C1_9 | SAMN07337979 | GCF_006374225.1 | NonPC | Public | 2014 | China | Env | 90.2 | 100 | 0.36 | 98.3256 | 5027285 | 45.37 | 60 | 298006 |
| SAMN07337980-GCF_006374215.1 | C10_4 | SAMN07337980 | GCF_006374215.1 | NonPC | Public | 2014 | China | Env | 89.7 | 100 | 0 | 98.3337 | 4896430 | 45.52 | 344 | 29377 |
| SAMN07337981-GCF_006373715.1 | C10_5 | SAMN07337981 | GCF_006373715.1 | NonPC | Public | 2014 | China | Env | 91.4 | 100 | 0.08 | 98.2528 | 5047259 | 45.26 | 99 | 200798 |
| SAMN07337982-GCF_006374185.1 | C2_1 | SAMN07337982 | GCF_006374185.1 | NonPC | Public | 2014 | China | Env | 90.9 | 100 | 0.26 | 98.2058 | 5290960 | 45.13 | 71 | 291487 |
| SAMN07337983-GCF_006374155.1 | C2_2 | SAMN07337983 | GCF_006374155.1 | NonPC | Public | 2014 | China | Env | 88 | 100 | 1.2 | 98.2678 | 5064414 | 45.47 | 71 | 317098 |
| SAMN07337984-GCF_006374135.1 | C2_4 | SAMN07337984 | GCF_006374135.1 | NonPC | Public | 2014 | China | Env | 91.3 | 100 | 3.38 | 98.3149 | 5305939 | 45.32 | 220 | 63623 |
| SAMN07337985-GCF_006373705.1 | C2_5 | SAMN07337985 | GCF_006373705.1 | NonPC | Public | 2014 | China | Env | 91.4 | 100 | 0.03 | 98.3707 | 5030118 | 45.32 | 70 | 219134 |
| SAMN07337986-GCF_006373675.1 | C2_6 | SAMN07337986 | GCF_006373675.1 | NonPC | Public | 2014 | China | Env | 87.8 | 100 | 0.36 | 98.2922 | 4988539 | 45.54 | 149 | 80333 |
| SAMN07337987-GCF_006373975.1 | C2_8 | SAMN07337987 | GCF_006373975.1 | NonPC | Public | 2014 | China | Env | 91.8 | 100 | 0.14 | 98.3679 | 5330400 | 45.07 | 108 | 177426 |
| SAMN07337988-GCF_006374125.1 | C3_10 | SAMN07337988 | GCF_006374125.1 | NonPC | Public | 2014 | China | Env | 87.8 | 100 | 0.45 | 98.313 | 5125002 | 45.18 | 351 | 51298 |
| SAMN07337989-GCF_006373945.1 | C3_2 | SAMN07337989 | GCF_006373945.1 | NonPC | Public | 2014 | China | Env | 91.8 | 100 | 0.23 | 98.4167 | 5162555 | 45.29 | 54 | 295449 |
| SAMN07337990-GCF_006373935.1 | C3_3 | SAMN07337990 | GCF_006373935.1 | NonPC | Public | 2014 | China | Env | 91 | 100 | 0.29 | 98.2618 | 5282172 | 45.23 | 161 | 190004 |
| SAMN07337992-GCF_006373905.1 | C3_5 | SAMN07337992 | GCF_006373905.1 | NonPC | Public | 2014 | China | Env | 89.4 | 100 | 0.19 | 98.2335 | 5339612 | 45.28 | 113 | 176252 |
| SAMN07337993-GCF_006373875.1 | C3_7 | SAMN07337993 | GCF_006373875.1 | NonPC | Public | 2014 | China | Env | 89.2 | 100 | 0.31 | 98.2759 | 5062812 | 45.22 | 60 | 452635 |
| SAMN07337994-GCF_006373845.1 | C3_9 | SAMN07337994 | GCF_006373845.1 | NonPC | Public | 2014 | China | Env | 88.7 | 100 | 0.09 | 98.2553 | 5236181 | 45.31 | 60 | 304611 |
| SAMN07337995-GCF_006374055.1 | C4_10 | SAMN07337995 | GCF_006374055.1 | NonPC | Public | 2014 | China | Env | 90 | 100 | 2.48 | 98.3195 | 4978917 | 45.37 | 193 | 51436 |
| SAMN07337996-GCF_006374095.1 | C4_2 | SAMN07337996 | GCF_006374095.1 | NonPC | Public | 2014 | China | Env | 91.2 | 100 | 0 | 98.3658 | 5033577 | 45.34 | 82 | 257980 |
| SAMN07337997-GCF_006374045.1 | C4_3 | SAMN07337997 | GCF_006374045.1 | NonPC | Public | 2014 | China | Env | 88.6 | 100 | 0.09 | 98.3436 | 5004885 | 45.32 | 51 | 382059 |
| SAMN07337998-GCF_006373645.1 | C4_4 | SAMN07337998 | GCF_006373645.1 | NonPC | Public | 2014 | China | Env | 91 | 100 | 0 | 98.3518 | 5014746 | 45.38 | 134 | 101569 |
| SAMN07337999-GCF_006373825.1 | C4_5 | SAMN07337999 | GCF_006373825.1 | NonPC | Public | 2014 | China | Env | 90.2 | 100 | 0.89 | 98.4618 | 5092174 | 45.68 | 346 | 36000 |
| SAMN07338000-GCF_006373635.1 | C4_6 | SAMN07338000 | GCF_006373635.1 | NonPC | Public | 2014 | China | Env | 89.5 | 100 | 0.15 | 98.218 | 5149541 | 45.28 | 52 | 314408 |
| SAMN07338001-GCF_006374075.1 | C4_7 | SAMN07338001 | GCF_006374075.1 | NonPC | Public | 2014 | China | Env | 89.5 | 100 | 0.35 | 98.25 | 5271357 | 45.25 | 109 | 158213 |
| SAMN07338002-GCF_006373815.1 | C4_8 | SAMN07338002 | GCF_006373815.1 | NonPC | Public | 2014 | China | Env | 90 | 100 | 0.05 | 98.3243 | 4947366 | 45.4 | 57 | 316962 |
| SAMN07338003-GCF_006374035.1 | C4_9 | SAMN07338003 | GCF_006374035.1 | NonPC | Public | 2014 | China | Env | 89.1 | 99.73 | 0.01 | 98.3458 | 4925014 | 45.51 | 436 | 31417 |
| SAMN07338004-GCF_006373995.1 | C5_1 | SAMN07338004 | GCF_006373995.1 | NonPC | Public | 2014 | China | Env | 90.9 | 100 | 0.27 | 98.3299 | 5020572 | 45.33 | 95 | 157083 |
| SAMN07338005-GCF_006373785.1 | C5_10 | SAMN07338005 | GCF_006373785.1 | NonPC | Public | 2014 | China | Env | 87.2 | 100 | 0.42 | 98.2168 | 4973052 | 45.32 | 84 | 218399 |
| SAMN07338006-GCF_006373955.1 | C5_2 | SAMN07338006 | GCF_006373955.1 | NonPC | Public | 2014 | China | Env | 89.7 | 100 | 0 | 98.2982 | 5068235 | 45.26 | 85 | 368907 |

| | | | | | | | | | | | | | | | | |
|---|---|---|---|---|---|---|---|---|---|---|---|---|---|---|---|---|
| SAMN07338007-GCF_006373775.1 | C5_3 | SAMN07338007 | GCF_006373775.1 | NonPC | Public | 2014 | China | Env | 89.7 | 100 | 0 | 98.303 | 5066624 | 45.26 | 89 | 368929 |
| SAMN07338008-GCF_006373625.1 | C5_4 | SAMN07338008 | GCF_006373625.1 | NonPC | Public | 2014 | China | Env | 91 | 100 | 0 | 98.3634 | 5014837 | 45.39 | 152 | 82118 |
| SAMN07338009-GCF_006373615.1 | C5_5 | SAMN07338009 | GCF_006373615.1 | NonPC | Public | 2014 | China | Env | 89 | 100 | 1.98 | 98.3824 | 5229901 | 45.27 | 732 | 22681 |
| SAMN07338010-GCF_006373585.1 | C5_6 | SAMN07338010 | GCF_006373585.1 | NonPC | Public | 2014 | China | Env | 89.4 | 100 | 0.09 | 98.2123 | 5135994 | 45.32 | 65 | 382343 |
| SAMN07338011-GCF_006373545.1 | C5_7 | SAMN07338011 | GCF_006373545.1 | NonPC | Public | 2014 | China | Env | 91.1 | 100 | 0.27 | 98.3107 | 5079309 | 45.28 | 89 | 157091 |
| SAMN07338012-GCF_006373555.1 | C5_8 | SAMN07338012 | GCF_006373555.1 | NonPC | Public | 2014 | China | Env | 88.3 | 100 | 1.12 | 98.2692 | 5134776 | 45.21 | 105 | 241788 |
| SAMN07338013-GCF_006373505.1 | C5_9 | SAMN07338013 | GCF_006373505.1 | NonPC | Public | 2014 | China | Env | 91.3 | 100 | 0 | 98.3945 | 4975272 | 45.4 | 54 | 251188 |
| SAMN07338014-GCF_006373515.1 | C6_1 | SAMN07338014 | GCF_006373515.1 | NonPC | Public | 2014 | China | Env | 90.7 | 100 | 2.26 | 98.3679 | 5126574 | 45.49 | 422 | 28321 |
| SAMN07338015-GCF_006373495.1 | C6_10 | SAMN07338015 | GCF_006373495.1 | NonPC | Public | 2014 | China | Env | 89.3 | 100 | 0 | 98.3234 | 5084230 | 45.17 | 126 | 151664 |
| SAMN07338016-GCF_006373465.1 | C6_2 | SAMN07338016 | GCF_006373465.1 | NonPC | Public | 2014 | China | Env | 89.3 | 100 | 0.05 | 98.363 | 4925202 | 45.38 | 78 | 256003 |
| SAMN07338017-GCF_006373455.1 | C6_4 | SAMN07338017 | GCF_006373455.1 | NonPC | Public | 2014 | China | Env | 88.2 | 100 | 0.36 | 98.2607 | 5022500 | 45.31 | 128 | 76735 |
| SAMN07338018-GCF_006373385.1 | C6_5 | SAMN07338018 | GCF_006373385.1 | NonPC | Public | 2014 | China | Env | 89.7 | 100 | 0 | 98.3108 | 5026709 | 45.32 | 104 | 381563 |
| SAMN07338019-GCF_006373395.1 | C6_6 | SAMN07338019 | GCF_006373395.1 | NonPC | Public | 2014 | China | Env | 89 | 100 | 0.05 | 98.3563 | 4894185 | 45.47 | 156 | 74262 |
| SAMN07338020-GCF_006373375.1 | C6_7 | SAMN07338020 | GCF_006373375.1 | NonPC | Public | 2014 | China | Env | 88.9 | 100 | 0.05 | 98.3339 | 4878084 | 45.51 | 252 | 44159 |
| SAMN07338021-GCF_006373365.1 | C6_8 | SAMN07338021 | GCF_006373365.1 | NonPC | Public | 2014 | China | Env | 89 | 99.86 | 1.31 | 98.3419 | 5145446 | 45.24 | 597 | 22092 |
| SAMN07338022-GCF_006373345.1 | C6_9 | SAMN07338022 | GCF_006373345.1 | NonPC | Public | 2014 | China | Env | 89.8 | 100 | 0 | 98.2593 | 5298072 | 45.13 | 75 | 285111 |
| SAMN07338023-GCF_006373315.1 | C7_1 | SAMN07338023 | GCF_006373315.1 | NonPC | Public | 2014 | China | Env | 90.1 | 100 | 0 | 98.2919 | 5113264 | 45.23 | 127 | 94082 |
| SAMN07338024-GCF_006373325.1 | C7_10 | SAMN07338024 | GCF_006373325.1 | NonPC | Public | 2014 | China | Env | 88.4 | 99.59 | 0.14 | 98.3799 | 4787233 | 45.74 | 463 | 21157 |
| SAMN07338025-GCF_006373285.1 | C7_2 | SAMN07338025 | GCF_006373285.1 | NonPC | Public | 2014 | China | Env | 89.6 | 100 | 0 | 98.3632 | 5199508 | 45.18 | 59 | 371244 |
| SAMN07338026-GCF_006373265.1 | C7_3 | SAMN07338026 | GCF_006373265.1 | NonPC | Public | 2014 | China | Env | 90.8 | 100 | 0.11 | 98.3815 | 5072745 | 45.38 | 89 | 253873 |
| SAMN07338027-GCF_006373235.1 | C7_4 | SAMN07338027 | GCF_006373235.1 | NonPC | Public | 2014 | China | Env | 89.7 | 100 | 0 | 98.3308 | 5035688 | 45.42 | 168 | 70735 |
| SAMN07338028-GCF_006373225.1 | C7_5 | SAMN07338028 | GCF_006373225.1 | NonPC | Public | 2014 | China | Env | 88.1 | 100 | 0.4 | 98.2453 | 5055662 | 45.19 | 100 | 365325 |
| SAMN07338029-GCF_006373185.1 | C7_6 | SAMN07338029 | GCF_006373185.1 | NonPC | Public | 2014 | China | Env | 90.7 | 100 | 0 | 98.3947 | 5052317 | 45.29 | 76 | 314616 |
| SAMN07338030-GCF_006373195.1 | C7_7 | SAMN07338030 | GCF_006373195.1 | NonPC | Public | 2014 | China | Env | 88.3 | 100 | 0.38 | 98.4799 | 4844204 | 45.66 | 402 | 24583 |
| SAMN07338031-GCF_006373175.1 | C7_9 | SAMN07338031 | GCF_006373175.1 | NonPC | Public | 2014 | China | Env | 91.3 | 100 | 0.13 | 98.3305 | 5236068 | 45.26 | 55 | 341283 |
| SAMN07338032-GCF_006373155.1 | C8_2 | SAMN07338032 | GCF_006373155.1 | NonPC | Public | 2014 | China | Env | 89.3 | 100 | 0.05 | 98.3347 | 4932075 | 45.36 | 58 | 264572 |
| SAMN07338033-GCF_006373115.1 | C8_3 | SAMN07338033 | GCF_006373115.1 | NonPC | Public | 2014 | China | Env | 89.5 | 99.91 | 0.34 | 98.3074 | 5006939 | 45.43 | 62 | 371057 |
| SAMN07338034-GCF_006373095.1 | C8_4 | SAMN07338034 | GCF_006373095.1 | NonPC | Public | 2014 | China | Env | 89.7 | 100 | 0.41 | 98.2606 | 5316681 | 45.27 | 133 | 202256 |
| SAMN07338035-GCF_006373055.1 | C8_5 | SAMN07338035 | GCF_006373055.1 | NonPC | Public | 2014 | China | Env | 90.6 | 99.73 | 4.96 | 98.2177 | 5036252 | 45.31 | 235 | 42434 |
| SAMN07338036-GCF_006373015.1 | C8_6 | SAMN07338036 | GCF_006373015.1 | NonPC | Public | 2014 | China | Env | 88.6 | 100 | 0 | 98.3889 | 4866358 | 45.63 | 505 | 23870 |
| SAMN07338037-GCF_006373025.1 | C8_7 | SAMN07338037 | GCF_006373025.1 | NonPC | Public | 2014 | China | Env | 89.1 | 99.64 | 0.51 | 98.4402 | 4830478 | 45.6 | 550 | 21069 |
| SAMN07338038-GCF_006373045.1 | C8_8 | SAMN07338038 | GCF_006373045.1 | NonPC | Public | 2014 | China | Env | 89.2 | 100 | 0.05 | 98.2792 | 5026409 | 45.38 | 157 | 84356 |
| SAMN07338039-GCF_006373035.1 | D1_1 | SAMN07338039 | GCF_006373035.1 | NonPC | Public | 2014 | China | Env | 88.5 | 100 | 0 | 98.3584 | 4851876 | 45.56 | 356 | 35468 |
| SAMN07338040-GCF_006372965.1 | D1_3 | SAMN07338040 | GCF_006372965.1 | NonPC | Public | 2014 | China | Env | 89.1 | 100 | 0.31 | 98.2764 | 4994054 | 45.27 | 58 | 259506 |
| SAMN07338041-GCF_006372945.1 | D1_4 | SAMN07338041 | GCF_006372945.1 | NonPC | Public | 2014 | China | Env | 91.6 | 100 | 0.05 | 98.3533 | 5050319 | 45.33 | 123 | 126144 |
| SAMN07338042-GCF_006372915.1 | D1_7 | SAMN07338042 | GCF_006372915.1 | NonPC | Public | 2014 | China | Env | 90.8 | 100 | 0 | 98.3604 | 5045373 | 45.47 | 207 | 66427 |
| SAMN07338043-GCF_006372925.1 | D1_8 | SAMN07338043 | GCF_006372925.1 | NonPC | Public | 2014 | China | Env | 89.4 | 100 | 0 | 98.3348 | 5049601 | 45.48 | 235 | 59680 |
| SAMN07338044-GCF_006372905.1 | D2_2 | SAMN07338044 | GCF_006372905.1 | NonPC | Public | 2014 | China | Env | 89.3 | 100 | 0.33 | 98.3038 | 4946008 | 45.36 | 67 | 445089 |
| SAMN07338045-GCF_006372885.1 | D2_4 | SAMN07338045 | GCF_006372885.1 | NonPC | Public | 2014 | China | Env | 89.6 | 100 | 0.14 | 98.3335 | 4964653 | 45.39 | 48 | 314876 |
| SAMN07338046-GCF_006372865.1 | D2_7 | SAMN07338046 | GCF_006372865.1 | NonPC | Public | 2014 | China | Env | 86.9 | 100 | 0.64 | 98.2933 | 4854211 | 45.68 | 635 | 16366 |
| SAMN07338047-GCF_006372855.1 | D2_8 | SAMN07338047 | GCF_006372855.1 | NonPC | Public | 2014 | China | Env | 90.1 | 100 | 0.05 | 98.2931 | 5216418 | 45.31 | 113 | 98398 |
| SAMN07338048-GCF_006372795.1 | D2_9 | SAMN07338048 | GCF_006372795.1 | NonPC | Public | 2014 | China | Env | 91.3 | 100 | 0.03 | 98.2986 | 5097558 | 45.25 | 98 | 262129 |
| SAMN07338049-GCF_006372785.1 | D3_2 | SAMN07338049 | GCF_006372785.1 | NonPC | Public | 2014 | China | Env | 88.2 | 100 | 0.36 | 98.2522 | 5019115 | 45.34 | 86 | 130662 |
| SAMN07338050-GCF_006372775.1 | D3_5 | SAMN07338050 | GCF_006372775.1 | NonPC | Public | 2014 | China | Env | 88.1 | 100 | 0.37 | 98.251 | 5042232 | 45.25 | 112 | 233085 |
| SAMN07338051-GCF_006372765.1 | D3_6 | SAMN07338051 | GCF_006372765.1 | NonPC | Public | 2014 | China | Env | 86.9 | 99.73 | 0.66 | 98.3289 | 4855167 | 45.56 | 591 | 19844 |
| SAMN07338052-GCF_006372735.1 | D4_1 | SAMN07338052 | GCF_006372735.1 | NonPC | Public | 2014 | China | Env | 88.2 | 99.42 | 0.35 | 98.4297 | 4713241 | 45.85 | 842 | 10672 |
| SAMN07338053-GCF_006372705.1 | D4_3 | SAMN07338053 | GCF_006372705.1 | NonPC | Public | 2014 | China | Env | 90.9 | 100 | 0.1 | 98.3848 | 5054607 | 45.31 | 143 | 90426 |
| SAMN07338054-GCF_006372715.1 | D4_4 | SAMN07338054 | GCF_006372715.1 | NonPC | Public | 2014 | China | Env | 91.3 | 100 | 0.54 | 98.3956 | 5473703 | 45.21 | 127 | 90506 |

| | | | | | | | | | | | | | | | | |
|---|---|---|---|---|---|---|---|---|---|---|---|---|---|---|---|---|
| SAMN07338055-GCF_006372695.1 | D4_5 | SAMN07338055 | GCF_006372695.1 | NonPC | Public | 2014 | China | Env | 87.8 | 100 | 0.3 | 98.2731 | 5095682 | 45.25 | 72 | 239452 |
| SAMN07338056-GCF_006372635.1 | D4_6 | SAMN07338056 | GCF_006372635.1 | NonPC | Public | 2014 | China | Env | 88.1 | 100 | 0.4 | 98.3226 | 4916770 | 45.34 | 55 | 379785 |
| SAMN07338057-GCF_006372625.1 | D4_7 | SAMN07338057 | GCF_006372625.1 | NonPC | Public | 2014 | China | Env | 87.2 | 99.73 | 0.61 | 98.2826 | 5273254 | 45.39 | 363 | 41474 |
| SAMN07338058-GCF_006372615.1 | D5_1 | SAMN07338058 | GCF_006372615.1 | NonPC | Public | 2014 | China | Env | 89 | 100 | 2.51 | 98.3368 | 4956579 | 45.6 | 217 | 61211 |
| SAMN07338059-GCF_006372595.1 | D5_3 | SAMN07338059 | GCF_006372595.1 | NonPC | Public | 2014 | China | Env | 90.9 | 100 | 0 | 98.4298 | 5210936 | 45.57 | 305 | 47168 |
| SAMN07338060-GCF_006372575.1 | D5_4 | SAMN07338060 | GCF_006372575.1 | NonPC | Public | 2014 | China | Env | 90.8 | 100 | 0.36 | 98.377 | 4933733 | 45.43 | 126 | 127551 |
| SAMN07338061-GCF_006372535.1 | D5_6 | SAMN07338061 | GCF_006372535.1 | NonPC | Public | 2014 | China | Env | 89.3 | 100 | 0 | 98.2623 | 4997191 | 45.29 | 65 | 372907 |
| SAMN07338062-GCF_006372565.1 | D5_7 | SAMN07338062 | GCF_006372565.1 | NonPC | Public | 2014 | China | Env | 90.5 | 99.66 | 0.43 | 98.4064 | 4933479 | 45.6 | 468 | 23859 |
| SAMN07338063-GCF_006372485.1 | D5_9 | SAMN07338063 | GCF_006372485.1 | NonPC | Public | 2014 | China | Env | 90.6 | 100 | 0.27 | 98.3626 | 5066189 | 45.33 | 72 | 179156 |
| SAMN07338064-GCF_006372465.1 | D6_1 | SAMN07338064 | GCF_006372465.1 | NonPC | Public | 2014 | China | Env | 88.9 | 100 | 0.27 | 98.3876 | 4975806 | 45.35 | 137 | 93392 |
| SAMN07338065-GCF_006372455.1 | D6_2 | SAMN07338065 | GCF_006372455.1 | NonPC | Public | 2014 | China | Env | 87.7 | 99.73 | 0.27 | 98.4534 | 4780541 | 45.7 | 488 | 21520 |
| SAMN07338066-GCF_006372445.1 | D6_3 | SAMN07338066 | GCF_006372445.1 | NonPC | Public | 2014 | China | Env | 90.8 | 99.98 | 0 | 98.429 | 5009781 | 45.53 | 380 | 28830 |
| SAMN07338067-GCF_006372405.1 | D6_4 | SAMN07338067 | GCF_006372405.1 | NonPC | Public | 2014 | China | Env | 88.3 | 100 | 0.47 | 98.2577 | 5004990 | 45.27 | 85 | 423635 |
| SAMN07338068-GCF_006372385.1 | D6_5 | SAMN07338068 | GCF_006372385.1 | NonPC | Public | 2014 | China | Env | 90.6 | 100 | 0.48 | 98.4676 | 4869790 | 45.62 | 442 | 23866 |
| SAMN07338069-GCF_006372395.1 | D6_6 | SAMN07338069 | GCF_006372395.1 | NonPC | Public | 2014 | China | Env | 91.6 | 100 | 0.14 | 98.3447 | 5451901 | 45.17 | 147 | 183172 |
| SAMN07338070-GCF_006372545.1 | D6_7 | SAMN07338070 | GCF_006372545.1 | NonPC | Public | 2014 | China | Env | 89.7 | 100 | 0.28 | 98.3569 | 5069505 | 45.25 | 127 | 101492 |
| SAMN07338071-GCF_006372185.1 | D6_8 | SAMN07338071 | GCF_006372185.1 | NonPC | Public | 2014 | China | Env | 88.7 | 100 | 0.05 | 98.358 | 4980304 | 45.38 | 66 | 392052 |
| SAMN07338072-GCF_006372365.1 | D7_1 | SAMN07338072 | GCF_006372365.1 | NonPC | Public | 2014 | China | Env | 88 | 99.55 | 0.59 | 98.4271 | 4723962 | 45.67 | 596 | 17302 |
| SAMN07338073-GCF_006372165.1 | D7_2 | SAMN07338073 | GCF_006372165.1 | NonPC | Public | 2014 | China | Env | 91.7 | 100 | 0.27 | 98.3328 | 5095568 | 45.28 | 175 | 120644 |
| SAMN07338074-GCF_006372145.1 | D7_3 | SAMN07338074 | GCF_006372145.1 | NonPC | Public | 2014 | China | Env | 90.6 | 100 | 0.08 | 98.3466 | 5009072 | 45.48 | 312 | 37299 |
| SAMN07338075-GCF_006372095.1 | D7_4 | SAMN07338075 | GCF_006372095.1 | NonPC | Public | 2014 | China | Env | 89.7 | 100 | 0.11 | 98.2821 | 5098965 | 45.18 | 84 | 337407 |
| SAMN07338076-GCF_006372325.1 | D7_6 | SAMN07338076 | GCF_006372325.1 | NonPC | Public | 2014 | China | Env | 89.4 | 100 | 0.33 | 98.3223 | 4931804 | 45.34 | 50 | 379622 |
| SAMN07338077-GCF_006372085.1 | D7_7 | SAMN07338077 | GCF_006372085.1 | NonPC | Public | 2014 | China | Env | 88.1 | 100 | 0.8 | 98.2678 | 4916242 | 45.34 | 120 | 128116 |
| SAMN07338078-GCF_006372075.1 | D7_8 | SAMN07338078 | GCF_006372075.1 | NonPC | Public | 2014 | China | Env | 89.9 | 99.66 | 0.33 | 98.4953 | 4790536 | 45.77 | 537 | 19811 |
| SAMN07338079-GCF_006372055.1 | D8_1 | SAMN07338079 | GCF_006372055.1 | NonPC | Public | 2014 | China | Env | 87.7 | 100 | 0.15 | 98.3744 | 4773685 | 45.63 | 250 | 41808 |
| SAMN07338080-GCF_006372305.1 | D8_2 | SAMN07338080 | GCF_006372305.1 | NonPC | Public | 2014 | China | Env | 87.8 | 100 | 0.1 | 98.3505 | 4793296 | 45.59 | 222 | 52145 |
| SAMN07338081-GCF_006372035.1 | D8_3 | SAMN07338081 | GCF_006372035.1 | NonPC | Public | 2014 | China | Env | 88.2 | 100 | 0.15 | 98.3447 | 4873577 | 45.37 | 55 | 245329 |
| SAMN07338083-GCF_006372275.1 | D8_5 | SAMN07338083 | GCF_006372275.1 | NonPC | Public | 2014 | China | Env | 90.2 | 100 | 0.41 | 98.387 | 4997426 | 45.54 | 415 | 30747 |
| SAMN07338084-GCF_006372265.1 | D8_6 | SAMN07338084 | GCF_006372265.1 | NonPC | Public | 2014 | China | Env | 90.7 | 100 | 0 | 98.387 | 5089973 | 45.28 | 90 | 168771 |
| SAMN07338085-GCF_006372255.1 | D8_7 | SAMN07338085 | GCF_006372255.1 | NonPC | Public | 2014 | China | Env | 89.9 | 100 | 0 | 98.4607 | 4927104 | 45.56 | 426 | 30223 |
| SAMN07338086-GCF_006372045.1 | D8_8 | SAMN07338086 | GCF_006372045.1 | NonPC | Public | 2014 | China | Env | 89.4 | 100 | 0.36 | 98.385 | 4944584 | 45.53 | 248 | 51960 |
| SAMN07338087-GCF_006372015.1 | D8_9 | SAMN07338087 | GCF_006372015.1 | NonPC | Public | 2014 | China | Env | 91.5 | 100 | 0.41 | 98.2644 | 5232978 | 45.32 | 92 | 209078 |
| SAMN07338088-GCF_006371975.1 | D9_2 | SAMN07338088 | GCF_006371975.1 | NonPC | Public | 2014 | China | Env | 88.3 | 100 | 0.71 | 98.279 | 5065447 | 45.17 | 95 | 254734 |
| SAMN07338089-GCF_006371945.1 | D9_3 | SAMN07338089 | GCF_006371945.1 | NonPC | Public | 2014 | China | Env | 87.6 | 100 | 0.36 | 98.2752 | 4971901 | 45.46 | 290 | 45545 |
| SAMN07338090-GCF_006371955.1 | D9_4 | SAMN07338090 | GCF_006371955.1 | NonPC | Public | 2014 | China | Env | 88.4 | 100 | 0.71 | 98.2693 | 5079856 | 45.18 | 88 | 254636 |
| SAMN07338091-GCF_006371915.1 | D9_6 | SAMN07338091 | GCF_006371915.1 | NonPC | Public | 2014 | China | Env | 90.6 | 99.91 | 0.27 | 98.3859 | 4979052 | 45.42 | 177 | 78453 |
| SAMN07338092-GCF_006371905.1 | E1_10 | SAMN07338092 | GCF_006371905.1 | NonPC | Public | 2014 | China | Env | 88.3 | 100 | 0.66 | 98.2626 | 5074467 | 45.23 | 92 | 197185 |
| SAMN07338093-GCF_006372235.1 | E1_5 | SAMN07338093 | GCF_006372235.1 | NonPC | Public | 2014 | China | Env | 89.4 | 99.93 | 0.07 | 98.2708 | 5099671 | 45.31 | 79 | 209120 |
| SAMN07338094-GCF_006372155.1 | E1_9 | SAMN07338094 | GCF_006372155.1 | NonPC | Public | 2014 | China | Env | 91.3 | 100 | 0.54 | 98.3374 | 5400416 | 45.36 | 141 | 198998 |
| SAMN07338095-GCF_006371865.1 | E10_8 | SAMN07338095 | GCF_006371865.1 | NonPC | Public | 2014 | China | Env | 86.7 | 99.69 | 1.26 | 98.2778 | 5062298 | 45.57 | 688 | 18973 |
| SAMN07338096-GCF_006371685.1 | E10_9 | SAMN07338096 | GCF_006371685.1 | NonPC | Public | 2014 | China | Env | 90.7 | 100 | 0.26 | 98.3227 | 5184711 | 45.15 | 80 | 314247 |
| SAMN07338097-GCF_006371825.1 | E2_10 | SAMN07338097 | GCF_006371825.1 | NonPC | Public | 2014 | China | Env | 89.2 | 100 | 0 | 98.3506 | 5013688 | 45.46 | 221 | 67838 |
| SAMN07338098-GCF_006371845.1 | E2_2 | SAMN07338098 | GCF_006371845.1 | NonPC | Public | 2014 | China | Env | 88.8 | 99.41 | 0.05 | 98.3813 | 4875015 | 45.61 | 391 | 27583 |
| SAMN07338099-GCF_006371835.1 | E2_4 | SAMN07338099 | GCF_006371835.1 | NonPC | Public | 2014 | China | Env | 89.7 | 100 | 0.19 | 98.2578 | 5101118 | 45.23 | 161 | 72997 |
| SAMN07338100-GCF_006371655.1 | E3_1 | SAMN07338100 | GCF_006371655.1 | NonPC | Public | 2014 | China | Env | 90.5 | 100 | 0.11 | 98.3932 | 5005223 | 45.53 | 359 | 34569 |
| SAMN07338101-GCF_006371635.1 | E3_10 | SAMN07338101 | GCF_006371635.1 | NonPC | Public | 2014 | China | Env | 90 | 100 | 0.08 | 98.2689 | 5117462 | 45.3 | 99 | 299328 |
| SAMN07338102-GCF_006371805.1 | E3_6 | SAMN07338102 | GCF_006371805.1 | NonPC | Public | 2014 | China | Env | 91.4 | 99.66 | 3.1 | 98.3473 | 5166013 | 45.27 | 400 | 26298 |
| SAMN07338103-GCF_006371625.1 | E4_10 | SAMN07338103 | GCF_006371625.1 | NonPC | Public | 2014 | China | Env | 87.9 | 100 | 2.9 | 98.2094 | 5104846 | 45.34 | 98 | 216437 |

| | | | | | | | | | | | | | | | | |
|---|---|---|---|---|---|---|---|---|---|---|---|---|---|---|---|---|
| SAMN07338104-GCF_006371575.1 | E4_2 | SAMN07338104 | GCF_006371575.1 | NonPC | Public | 2014 | China | Env | 89.8 | 100 | 0.04 | 98.3021 | 5085867 | 45.44 | 242 | 52162 |
| SAMN07338105-GCF_006371585.1 | E4_3 | SAMN07338105 | GCF_006371585.1 | NonPC | Public | 2014 | China | Env | 87.7 | 100 | 0.22 | 98.3778 | 4854400 | 45.75 | 570 | 16906 |
| SAMN07338106-GCF_006371565.1 | E4_4 | SAMN07338106 | GCF_006371565.1 | NonPC | Public | 2014 | China | Env | 87.9 | 100 | 0.36 | 98.2455 | 5019417 | 45.33 | 121 | 115162 |
| SAMN07338107-GCF_006371545.1 | E4_9 | SAMN07338107 | GCF_006371545.1 | NonPC | Public | 2014 | China | Env | 91.2 | 100 | 0.05 | 98.33 | 5036756 | 45.31 | 91 | 161897 |
| SAMN07338108-GCF_006371495.1 | E5_8 | SAMN07338108 | GCF_006371495.1 | NonPC | Public | 2014 | China | Env | 88 | 100 | 0.39 | 98.1721 | 5056150 | 45.21 | 94 | 233354 |
| SAMN07338109-GCF_006371485.1 | E6_4 | SAMN07338109 | GCF_006371485.1 | NonPC | Public | 2014 | China | Env | 90.9 | 100 | 0.05 | 98.3594 | 5070489 | 45.2 | 184 | 76214 |
| SAMN07338110-GCF_006371465.1 | E7_10 | SAMN07338110 | GCF_006371465.1 | NonPC | Public | 2014 | China | Env | 90.8 | 100 | 0.54 | 98.3289 | 5137702 | 45.43 | 294 | 45316 |
| SAMN07338111-GCF_006371435.1 | E7_9 | SAMN07338111 | GCF_006371435.1 | NonPC | Public | 2014 | China | Env | 86.3 | 99.19 | 1.55 | 98.4066 | 4783444 | 45.68 | 812 | 12485 |
| SAMN07338112-GCF_006371445.1 | E9_7 | SAMN07338112 | GCF_006371445.1 | NonPC | Public | 2014 | China | Env | 90.1 | 99.73 | 0 | 98.4602 | 5014432 | 45.6 | 475 | 23008 |
| SAMN07338113-GCF_006371455.1 | E9_8 | SAMN07338113 | GCF_006371455.1 | NonPC | Public | 2014 | China | Env | 91.2 | 100 | 0.05 | 98.3162 | 5124211 | 45.24 | 75 | 296613 |
| SAMN07338114-GCF_006371795.1 | F1_1 | SAMN07338114 | GCF_006371795.1 | NonPC | Public | 2014 | China | Env | 88.9 | 100 | 0.05 | 98.2783 | 5091319 | 45.31 | 88 | 212430 |
| SAMN07338115-GCF_006371405.1 | F1_10 | SAMN07338115 | GCF_006371405.1 | NonPC | Public | 2014 | China | Env | 90.1 | 100 | 0.05 | 98.3485 | 5070500 | 45.33 | 150 | 80003 |
| SAMN07338116-GCF_006371755.1 | F1_2 | SAMN07338116 | GCF_006371755.1 | NonPC | Public | 2014 | China | Env | 91 | 100 | 0.05 | 98.3219 | 5208910 | 45.23 | 103 | 178028 |
| SAMN07338117-GCF_006371395.1 | F1_3 | SAMN07338117 | GCF_006371395.1 | NonPC | Public | 2014 | China | Env | 89.5 | 100 | 0 | 98.3104 | 4965809 | 45.33 | 65 | 367488 |
| SAMN07338118-GCF_006371735.1 | F1_4 | SAMN07338118 | GCF_006371735.1 | NonPC | Public | 2014 | China | Env | 91.1 | 100 | 0.05 | 98.3248 | 5222742 | 45.21 | 79 | 297598 |
| SAMN07338119-GCF_006371355.1 | F1_5 | SAMN07338119 | GCF_006371355.1 | NonPC | Public | 2014 | China | Env | 89.8 | 100 | 0.05 | 98.3528 | 5229417 | 45.21 | 78 | 320507 |
| SAMN07338120-GCF_006371345.1 | F1_6 | SAMN07338120 | GCF_006371345.1 | NonPC | Public | 2014 | China | Env | 88.9 | 100 | 2.49 | 98.3712 | 5073714 | 45.36 | 90 | 326689 |
| SAMN07338121-GCF_006371715.1 | F1_7 | SAMN07338121 | GCF_006371715.1 | NonPC | Public | 2014 | China | Env | 90.2 | 100 | 0.05 | 98.3321 | 5073406 | 45.3 | 151 | 78214 |
| SAMN07338122-GCF_006371705.1 | F1_8 | SAMN07338122 | GCF_006371705.1 | NonPC | Public | 2014 | China | Env | 90.4 | 100 | 0.05 | 98.2983 | 5126775 | 45.19 | 77 | 177520 |
| SAMN07338123-GCF_006371325.1 | F1_9 | SAMN07338123 | GCF_006371325.1 | NonPC | Public | 2014 | China | Env | 89.7 | 100 | 0.05 | 98.3609 | 4981097 | 45.5 | 357 | 32790 |
| SAMN07338124-GCF_006371305.1 | F10_1 | SAMN07338124 | GCF_006371305.1 | NonPC | Public | 2014 | China | Env | 90.5 | 100 | 0 | 98.3267 | 5188250 | 45.16 | 81 | 375766 |
| SAMN07338125-GCF_006371095.1 | F10_10 | SAMN07338125 | GCF_006371095.1 | NonPC | Public | 2014 | China | Env | 89.3 | 100 | 0.01 | 98.3035 | 4990426 | 45.42 | 195 | 65021 |
| SAMN07338126-GCF_006371115.1 | F10_2 | SAMN07338126 | GCF_006371115.1 | NonPC | Public | 2014 | China | Env | 89.1 | 100 | 2.51 | 98.349 | 4985614 | 45.45 | 62 | 390613 |
| SAMN07338127-GCF_006371055.1 | F10_3 | SAMN07338127 | GCF_006371055.1 | NonPC | Public | 2014 | China | Env | 91.3 | 100 | 0 | 98.3644 | 5179427 | 45.1 | 122 | 190822 |
| SAMN07338128-GCF_006371065.1 | F10_4 | SAMN07338128 | GCF_006371065.1 | NonPC | Public | 2014 | China | Env | 89.1 | 100 | 0.27 | 98.2274 | 5002737 | 45.39 | 184 | 72750 |
| SAMN07338129-GCF_006371005.1 | F10_5 | SAMN07338129 | GCF_006371005.1 | NonPC | Public | 2014 | China | Env | 90.7 | 100 | 0.1 | 98.3521 | 4982271 | 45.34 | 120 | 105412 |
| SAMN07338130-GCF_006371275.1 | F10_6 | SAMN07338130 | GCF_006371275.1 | NonPC | Public | 2014 | China | Env | 89.6 | 99.73 | 0.01 | 98.3558 | 5022992 | 45.35 | 97 | 221284 |
| SAMN07338131-GCF_006371015.1 | F10_7 | SAMN07338131 | GCF_006371015.1 | NonPC | Public | 2014 | China | Env | 90.9 | 100 | 0 | 98.3531 | 5142467 | 45.28 | 72 | 178264 |
| SAMN07338132-GCF_006371265.1 | F10_8 | SAMN07338132 | GCF_006371265.1 | NonPC | Public | 2014 | China | Env | 91.2 | 100 | 2.21 | 98.3969 | 5191029 | 45.26 | 77 | 206938 |
| SAMN07338134-GCF_006370725.1 | F2_1 | SAMN07338134 | GCF_006370725.1 | NonPC | Public | 2014 | China | Env | 90.3 | 100 | 0.05 | 98.347 | 5116569 | 45.18 | 86 | 166456 |
| SAMN07338135-GCF_006371215.1 | F2_10 | SAMN07338135 | GCF_006371215.1 | PC | Public | 2014 | China | Env | 98 | 100 | 0 | 99.993 | 5067129 | 45.27 | 80 | 262079 |
| SAMN07338136-GCF_006370745.1 | F2_2 | SAMN07338136 | GCF_006370745.1 | NonPC | Public | 2014 | China | Env | 89.6 | 100 | 0.33 | 98.363 | 4978813 | 45.51 | 343 | 35177 |
| SAMN07338137-GCF_006370945.1 | F2_3 | SAMN07338137 | GCF_006370945.1 | NonPC | Public | 2014 | China | Env | 89.3 | 100 | 0.05 | 98.3215 | 5080772 | 45.29 | 164 | 81828 |
| SAMN07338138-GCF_006370715.1 | F2_4 | SAMN07338138 | GCF_006370715.1 | NonPC | Public | 2014 | China | Env | 90.4 | 100 | 0.05 | 98.2903 | 5126466 | 45.19 | 84 | 177520 |
| SAMN07338139-GCF_006371235.1 | F2_5 | SAMN07338139 | GCF_006371235.1 | NonPC | Public | 2014 | China | Env | 89.6 | 100 | 0.12 | 98.3749 | 4981143 | 45.47 | 465 | 23839 |
| SAMN07338140-GCF_006370975.1 | F2_6 | SAMN07338140 | GCF_006370975.1 | NonPC | Public | 2014 | China | Env | 90.1 | 100 | 1.8 | 98.3541 | 5101267 | 45.33 | 275 | 46919 |
| SAMN07338142-GCF_006371225.1 | F2_8 | SAMN07338142 | GCF_006371225.1 | PC | Public | 2014 | China | Env | 98.1 | 100 | 1.09 | 99.991 | 5155416 | 45.29 | 82 | 259512 |
| SAMN07338143-GCF_006370655.1 | F2_9 | SAMN07338143 | GCF_006370655.1 | PC | Public | 2014 | China | Env | 97.6 | 99.46 | 0 | 99.9941 | 5040360 | 45.28 | 131 | 136127 |
| SAMN07338145-GCF_006370685.1 | F3_10 | SAMN07338145 | GCF_006370685.1 | PC | Public | 2014 | China | Env | 97.9 | 100 | 0 | 99.9933 | 5058616 | 45.27 | 83 | 259512 |
| SAMN07338146-GCF_006370915.1 | F3_2 | SAMN07338146 | GCF_006370915.1 | PC | Public | 2014 | China | Env | 98.1 | 100 | 0 | 99.9875 | 5071547 | 45.28 | 89 | 259516 |
| SAMN07338147-GCF_006370895.1 | F3_3 | SAMN07338147 | GCF_006370895.1 | PC | Public | 2014 | China | Env | 98.1 | 99.95 | 0 | 99.984 | 5071440 | 45.28 | 84 | 199079 |
| SAMN07338148-GCF_006370855.1 | F3_4 | SAMN07338148 | GCF_006370855.1 | PC | Public | 2014 | China | Env | 98.1 | 100 | 0 | 99.9889 | 5069597 | 45.28 | 85 | 261244 |
| SAMN07338149-GCF_006370835.1 | F3_5 | SAMN07338149 | GCF_006370835.1 | PC | Public | 2014 | China | Env | 98.1 | 100 | 0 | 99.988 | 5069799 | 45.28 | 93 | 186910 |
| SAMN07338151-GCF_006370635.1 | F3_7 | SAMN07338151 | GCF_006370635.1 | PC | Public | 2014 | China | Env | 97.7 | 100 | 0 | 99.9895 | 5044635 | 45.34 | 151 | 94818 |
| SAMN07338152-GCF_006371125.1 | F3_8 | SAMN07338152 | GCF_006371125.1 | PC | Public | 2014 | China | Env | 98.2 | 100 | 0 | 99.9937 | 5073175 | 45.28 | 88 | 243534 |
| SAMN07338153-GCF_006370625.1 | F3_9 | SAMN07338153 | GCF_006370625.1 | PC | Public | 2014 | China | Env | 98.1 | 100 | 0 | 99.9939 | 5072417 | 45.28 | 81 | 259520 |
| SAMN07338154-GCF_006370805.1 | F4_1 | SAMN07338154 | GCF_006370805.1 | NonPC | Public | 2014 | China | Env | 90.4 | 100 | 4.14 | 98.3056 | 5279289 | 45.27 | 85 | 166473 |
| SAMN07338155-GCF_006370785.1 | F4_10 | SAMN07338155 | GCF_006370785.1 | NonPC | Public | 2014 | China | Env | 88.2 | 98.86 | 0.47 | 98.462 | 4937873 | 45.68 | 655 | 18644 |

| | | | | | | | | | | | | | | | | |
|---|---|---|---|---|---|---|---|---|---|---|---|---|---|---|---|---|
| SAMN07338156-GCF_006370775.1 | F4_2 | SAMN07338156 | GCF_006370775.1 | NonPC | Public | 2014 | China | Env | 87 | 99.82 | 0.61 | 98.3626 | 4842485 | 45.64 | 580 | 19777 |
| SAMN07338157-GCF_006370595.1 | F4_3 | SAMN07338157 | GCF_006370595.1 | NonPC | Public | 2014 | China | Env | 90.7 | 100 | 0.04 | 98.454 | 4895309 | 45.6 | 430 | 29022 |
| SAMN07338158-GCF_006371105.1 | F4_4 | SAMN07338158 | GCF_006371105.1 | NonPC | Public | 2014 | China | Env | 89.5 | 100 | 0.03 | 98.238 | 5064342 | 45.23 | 84 | 231036 |
| SAMN07338160-GCF_006369665.1 | F4_6 | SAMN07338160 | GCF_006369665.1 | NonPC | Public | 2014 | China | Env | 89.2 | 100 | 0.27 | 98.3505 | 5035947 | 45.33 | 137 | 94941 |
| SAMN07338161-GCF_006370755.1 | F4_7 | SAMN07338161 | GCF_006370755.1 | NonPC | Public | 2014 | China | Env | 88.6 | 100 | 0.36 | 98.3854 | 5126934 | 45.48 | 576 | 21603 |
| SAMN07338162-GCF_006369375.1 | F4_8 | SAMN07338162 | GCF_006369375.1 | NonPC | Public | 2014 | China | Env | 89.2 | 100 | 0.05 | 98.3741 | 5018824 | 45.37 | 89 | 140572 |
| SAMN07338163-GCF_006369075.1 | F4_9 | SAMN07338163 | GCF_006369075.1 | NonPC | Public | 2014 | China | Env | 89.5 | 100 | 0 | 98.3511 | 4928694 | 45.48 | 357 | 38481 |
| SAMN07338164-GCF_006369095.1 | F5_1 | SAMN07338164 | GCF_006369095.1 | NonPC | Public | 2014 | China | Env | 89.5 | 100 | 0.87 | 98.3866 | 5032451 | 45.51 | 359 | 29415 |
| SAMN07338165-GCF_006369055.1 | F5_10 | SAMN07338165 | GCF_006369055.1 | NonPC | Public | 2014 | China | Env | 89.8 | 100 | 0.03 | 98.31 | 5042930 | 45.31 | 69 | 189168 |
| SAMN07338166-GCF_006368995.1 | F5_3 | SAMN07338166 | GCF_006368995.1 | NonPC | Public | 2014 | China | Env | 90.4 | 100 | 0.05 | 98.3266 | 4931061 | 45.46 | 207 | 59877 |
| SAMN07338167-GCF_006369005.1 | F5_4 | SAMN07338167 | GCF_006369005.1 | NonPC | Public | 2014 | China | Env | 89.5 | 100 | 0.23 | 98.2576 | 5069221 | 45.3 | 98 | 111688 |
| SAMN07338168-GCF_006368955.1 | F5_5 | SAMN07338168 | GCF_006368955.1 | NonPC | Public | 2014 | China | Env | 89.5 | 100 | 0 | 98.3763 | 4962044 | 45.41 | 92 | 139651 |
| SAMN07338169-GCF_006369645.1 | F5_6 | SAMN07338169 | GCF_006369645.1 | NonPC | Public | 2014 | China | Env | 89.4 | 99.73 | 0.15 | 98.3531 | 5015916 | 45.34 | 187 | 81740 |
| SAMN07338170-GCF_006369655.1 | F5_7 | SAMN07338170 | GCF_006369655.1 | NonPC | Public | 2014 | China | Env | 88.9 | 100 | 0.54 | 98.4366 | 4957452 | 45.52 | 522 | 28089 |
| SAMN07338171-GCF_006369355.1 | F5_8 | SAMN07338171 | GCF_006369355.1 | NonPC | Public | 2014 | China | Env | 89.1 | 99.73 | 0.11 | 98.4459 | 4874878 | 45.6 | 336 | 37720 |
| SAMN07338172-GCF_006369625.1 | F5_9 | SAMN07338172 | GCF_006369625.1 | NonPC | Public | 2014 | China | Env | 88.7 | 99.69 | 0.61 | 98.4819 | 4923797 | 45.52 | 508 | 26696 |
| SAMN07338173-GCF_006368915.1 | F6_1 | SAMN07338173 | GCF_006368915.1 | NonPC | Public | 2014 | China | Env | 89.5 | 100 | 0.14 | 98.2865 | 5280327 | 45.24 | 185 | 68349 |
| SAMN07338174-GCF_006369365.1 | F6_2 | SAMN07338174 | GCF_006369365.1 | NonPC | Public | 2014 | China | Env | 88.6 | 100 | 0.05 | 98.354 | 4874159 | 45.64 | 306 | 43971 |
| SAMN07338175-GCF_006368945.1 | F6_3 | SAMN07338175 | GCF_006368945.1 | NonPC | Public | 2014 | China | Env | 89.8 | 100 | 0.33 | 98.469 | 4808680 | 45.62 | 355 | 31693 |
| SAMN07338176-GCF_006369615.1 | F6_4 | SAMN07338176 | GCF_006369615.1 | NonPC | Public | 2014 | China | Env | 89.5 | 100 | 0.05 | 98.3793 | 4941249 | 45.44 | 54 | 285859 |
| SAMN07338177-GCF_006368935.1 | F6_5 | SAMN07338177 | GCF_006368935.1 | NonPC | Public | 2014 | China | Env | 91.3 | 100 | 0.36 | 98.3314 | 5073915 | 45.26 | 85 | 221148 |
| SAMN07338178-GCF_006368925.1 | F6_6 | SAMN07338178 | GCF_006368925.1 | NonPC | Public | 2014 | China | Env | 88.4 | 99.46 | 0.18 | 98.3798 | 4856632 | 45.59 | 568 | 21472 |
| SAMN07338179-GCF_006369335.1 | F6_7 | SAMN07338179 | GCF_006369335.1 | NonPC | Public | 2014 | China | Env | 89.1 | 99.73 | 0.27 | 98.4095 | 4875675 | 45.64 | 430 | 30527 |
| SAMN07338180-GCF_006368875.1 | F6_8 | SAMN07338180 | GCF_006368875.1 | NonPC | Public | 2014 | China | Env | 90.8 | 100 | 0 | 98.4403 | 4949070 | 45.47 | 205 | 67808 |
| SAMN07338181-GCF_006369565.1 | F6_9 | SAMN07338181 | GCF_006369565.1 | NonPC | Public | 2014 | China | Env | 90.1 | 100 | 0 | 98.3522 | 5265014 | 45.21 | 158 | 87224 |
| SAMN07338182-GCF_006369535.1 | F7_10 | SAMN07338182 | GCF_006369535.1 | NonPC | Public | 2014 | China | Env | 91.8 | 100 | 0.03 | 98.4037 | 5050438 | 45.29 | 62 | 466576 |
| SAMN07338183-GCF_006369325.1 | F7_2 | SAMN07338183 | GCF_006369325.1 | NonPC | Public | 2014 | China | Env | 90.8 | 100 | 0.41 | 98.4878 | 4975835 | 45.51 | 374 | 31291 |
| SAMN07338184-GCF_006369545.1 | F7_3 | SAMN07338184 | GCF_006369545.1 | NonPC | Public | 2014 | China | Env | 88 | 100 | 0.77 | 98.2966 | 4969552 | 45.31 | 197 | 66223 |
| SAMN07338185-GCF_006369525.1 | F7_4 | SAMN07338185 | GCF_006369525.1 | NonPC | Public | 2014 | China | Env | 91.2 | 100 | 0.36 | 98.4421 | 5033767 | 45.42 | 295 | 41869 |
| SAMN07338186-GCF_006368845.1 | F7_6 | SAMN07338186 | GCF_006368845.1 | NonPC | Public | 2014 | China | Env | 88.3 | 100 | 0.66 | 98.2116 | 5068848 | 45.27 | 140 | 98690 |
| SAMN07338187-GCF_006369515.1 | F7_7 | SAMN07338187 | GCF_006369515.1 | NonPC | Public | 2014 | China | Env | 87.1 | 100 | 0.51 | 98.2838 | 5053548 | 45.54 | 489 | 27004 |
| SAMN07338188-GCF_006369315.1 | F7_8 | SAMN07338188 | GCF_006369315.1 | NonPC | Public | 2014 | China | Env | 91.5 | 100 | 0.54 | 98.3683 | 5224664 | 45.28 | 64 | 315040 |
| SAMN07338189-GCF_006369475.1 | F7_9 | SAMN07338189 | GCF_006369475.1 | NonPC | Public | 2014 | China | Env | 91.2 | 100 | 1.7 | 98.3884 | 5097123 | 45.35 | 143 | 100217 |
| SAMN07338190-GCF_006369285.1 | F8_1 | SAMN07338190 | GCF_006369285.1 | NonPC | Public | 2014 | China | Env | 89 | 100 | 0.27 | 98.3247 | 4927262 | 45.36 | 96 | 140213 |
| SAMN07338191-GCF_006369265.1 | F8_10 | SAMN07338191 | GCF_006369265.1 | NonPC | Public | 2014 | China | Env | 89.7 | 100 | 0.05 | 98.3116 | 5047945 | 45.36 | 51 | 408535 |
| SAMN07338192-GCF_006369235.1 | F8_2 | SAMN07338192 | GCF_006369235.1 | NonPC | Public | 2014 | China | Env | 91.6 | 100 | 0.03 | 98.4105 | 5068344 | 45.26 | 52 | 316396 |
| SAMN07338193-GCF_006369205.1 | F8_3 | SAMN07338193 | GCF_006369205.1 | NonPC | Public | 2014 | China | Env | 91.1 | 100 | 0 | 98.2283 | 5133828 | 45.27 | 81 | 191589 |
| SAMN07338194-GCF_006369185.1 | F8_4 | SAMN07338194 | GCF_006369185.1 | NonPC | Public | 2014 | China | Env | 90.7 | 99.46 | 0 | 98.4558 | 4918252 | 45.57 | 331 | 30048 |
| SAMN07338195-GCF_006369455.1 | F8_5 | SAMN07338195 | GCF_006369455.1 | NonPC | Public | 2014 | China | Env | 92.3 | 100 | 0.05 | 98.3445 | 5236970 | 45.29 | 152 | 97362 |
| SAMN07338196-GCF_006368835.1 | F8_6 | SAMN07338196 | GCF_006368835.1 | NonPC | Public | 2014 | China | Env | 89.3 | 100 | 0.14 | 98.3246 | 4923455 | 45.47 | 148 | 72093 |
| SAMN07338197-GCF_006369195.1 | F8_7 | SAMN07338197 | GCF_006369195.1 | NonPC | Public | 2014 | China | Env | 88.9 | 99.73 | 0.05 | 98.3763 | 4960033 | 45.54 | 343 | 37409 |
| SAMN07338198-GCF_006369175.1 | F8_8 | SAMN07338198 | GCF_006369175.1 | NonPC | Public | 2014 | China | Env | 89.6 | 100 | 0 | 98.3173 | 4999359 | 45.35 | 71 | 412050 |
| SAMN07338199-GCF_006368815.1 | F8_9 | SAMN07338199 | GCF_006368815.1 | NonPC | Public | 2014 | China | Env | 88.7 | 100 | 0.54 | 98.3835 | 4957759 | 45.48 | 302 | 40695 |
| SAMN07338200-GCF_006369105.1 | F9_1 | SAMN07338200 | GCF_006369105.1 | NonPC | Public | 2014 | China | Env | 89.2 | 100 | 0 | 98.3109 | 5048708 | 45.31 | 75 | 238517 |
| SAMN07338201-GCF_006368785.1 | F9_10 | SAMN07338201 | GCF_006368785.1 | NonPC | Public | 2014 | China | Env | 90.3 | 100 | 0.41 | 98.361 | 5329999 | 45.19 | 94 | 118389 |
| SAMN07338202-GCF_006369115.1 | F9_2 | SAMN07338202 | GCF_006369115.1 | NonPC | Public | 2014 | China | Env | 91.6 | 100 | 0 | 98.3458 | 5226233 | 45.22 | 110 | 110123 |
| SAMN07338203-GCF_006369435.1 | F9_3 | SAMN07338203 | GCF_006369435.1 | NonPC | Public | 2014 | China | Env | 89.8 | 100 | 0.05 | 98.283 | 4955771 | 45.48 | 130 | 81690 |
| SAMN07338204-GCF_006369425.1 | F9_4 | SAMN07338204 | GCF_006369425.1 | NonPC | Public | 2014 | China | Env | 90.2 | 100 | 4.77 | 98.3927 | 5393618 | 45.22 | 89 | 118389 |

| | | | | | | | | | | | | | | | | |
|---|---|---|---|---|---|---|---|---|---|---|---|---|---|---|---|---|
| SAMN07338205-GCF_006368755.1 | F9_5 | SAMN07338205 | GCF_006368755.1 | NonPC | Public | 2014 | China | Env | 89.4 | 100 | 0.11 | 98.2869 | 4998646 | 45.3 | 77 | 283428 |
| SAMN07338206-GCF_006368575.1 | F9_6 | SAMN07338206 | GCF_006368575.1 | NonPC | Public | 2014 | China | Env | 91.4 | 100 | 0.03 | 98.3071 | 5132639 | 45.25 | 96 | 324925 |
| SAMN07338207-GCF_006368795.1 | F9_7 | SAMN07338207 | GCF_006368795.1 | NonPC | Public | 2014 | China | Env | 89.4 | 100 | 0.34 | 98.2589 | 5239244 | 45.42 | 238 | 80044 |
| SAMN07338208-GCF_006368565.1 | F9_8 | SAMN07338208 | GCF_006368565.1 | NonPC | Public | 2014 | China | Env | 89.6 | 100 | 3.95 | 98.3464 | 5139428 | 45.37 | 73 | 202994 |
| SAMN07338209-GCF_006368365.1 | F9_9 | SAMN07338209 | GCF_006368365.1 | NonPC | Public | 2014 | China | Env | 89.6 | 100 | 1.77 | 98.3528 | 5042615 | 45.42 | 181 | 71186 |
| SAMN07338210-GCF_006368545.1 | G1_1 | SAMN07338210 | GCF_006368545.1 | NonPC | Public | 2014 | China | Env | 89.3 | 100 | 0.05 | 98.3605 | 5029412 | 45.34 | 250 | 60148 |
| SAMN07338211-GCF_006368725.1 | G1_10 | SAMN07338211 | GCF_006368725.1 | PC | Public | 2014 | China | Env | 98 | 99.95 | 0 | 99.9845 | 5130880 | 45.28 | 98 | 152446 |
| SAMN07338212-GCF_006368705.1 | G1_2 | SAMN07338212 | GCF_006368705.1 | NonPC | Public | 2014 | China | Env | 89.1 | 100 | 0.27 | 98.3196 | 4952358 | 45.47 | 194 | 56346 |
| SAMN07338213-GCF_006368505.1 | G1_3 | SAMN07338213 | GCF_006368505.1 | PC | Public | 2014 | China | Env | 98 | 99.97 | 0 | 99.9835 | 5062460 | 45.3 | 109 | 149982 |
| SAMN07338214-GCF_006368495.1 | G1_4 | SAMN07338214 | GCF_006368495.1 | PC | Public | 2014 | China | Env | 98.1 | 100 | 0.27 | 99.9871 | 5073097 | 45.28 | 91 | 198328 |
| SAMN07338215-GCF_006368335.1 | G1_5 | SAMN07338215 | GCF_006368335.1 | PC | Public | 2014 | China | Env | 98 | 99.73 | 0 | 99.9869 | 5062986 | 45.28 | 106 | 151206 |
| SAMN07338216-GCF_006368345.1 | G1_6 | SAMN07338216 | GCF_006368345.1 | PC | Public | 2014 | China | Env | 97.9 | 100 | 0 | 99.9915 | 5059234 | 45.28 | 97 | 168136 |
| SAMN07338217-GCF_006368485.1 | G1_7 | SAMN07338217 | GCF_006368485.1 | PC | Public | 2014 | China | Env | 95.3 | 100 | 0.27 | 99.9909 | 4873327 | 45.66 | 503 | 21909 |
| SAMN07338218-GCF_006368305.1 | G1_8 | SAMN07338218 | GCF_006368305.1 | PC | Public | 2014 | China | Env | 98.1 | 100 | 0 | 99.9898 | 5068363 | 45.28 | 90 | 198152 |
| SAMN07338219-GCF_006368435.1 | G1_9 | SAMN07338219 | GCF_006368435.1 | PC | Public | 2014 | China | Env | 96.7 | 100 | 0.27 | 99.9925 | 4968894 | 45.5 | 298 | 41781 |
| SAMN07338220-GCF_006368645.1 | G2_1 | SAMN07338220 | GCF_006368645.1 | PC | Public | 2014 | China | Env | 96 | 100 | 0.54 | 99.9863 | 4933902 | 45.6 | 382 | 30627 |
| SAMN07338221-GCF_006368425.1 | G2_2 | SAMN07338221 | GCF_006368425.1 | PC | Public | 2014 | China | Env | 98.2 | 100 | 0 | 99.9938 | 5072448 | 45.28 | 86 | 198168 |
| SAMN07338222-GCF_006368265.1 | G2_3 | SAMN07338222 | GCF_006368265.1 | NonPC | Public | 2014 | China | Env | 91.1 | 100 | 0 | 98.2853 | 4983683 | 45.36 | 83 | 197305 |
| SAMN07338223-GCF_006368665.1 | G2_5 | SAMN07338223 | GCF_006368665.1 | NonPC | Public | 2014 | China | Env | 90.9 | 100 | 0.6 | 98.3923 | 5389948 | 45.25 | 90 | 188342 |
| SAMN07338224-GCF_006368625.1 | G2_6 | SAMN07338224 | GCF_006368625.1 | NonPC | Public | 2014 | China | Env | 89.4 | 100 | 2.66 | 98.3468 | 5035798 | 45.36 | 72 | 255295 |
| SAMN07338225-GCF_006368635.1 | G2_7 | SAMN07338225 | GCF_006368635.1 | NonPC | Public | 2014 | China | Env | 88.2 | 99.73 | 0.02 | 98.4408 | 4930452 | 45.75 | 491 | 22075 |
| SAMN07338226-GCF_006368255.1 | S010 | SAMN07338226 | GCF_006368255.1 | PrePC | Public | NA | NA | NA | 93.1 | 99.98 | 0.7 | 99.2512 | 5014236 | 45.32 | 353 | 28944 |
| SAMN07338227-GCF_006368375.1 | S061 | SAMN07338227 | GCF_006368375.1 | PC | Public | NA | NA | Clinical | 97.5 | 99.57 | 0.35 | 99.9682 | 5030563 | 45.27 | 416 | 20681 |
| SAMN07338228-GCF_006368415.1 | S089 | SAMN07338228 | GCF_006368415.1 | PC | Public | NA | NA | Clinical | 97 | 99.65 | 0.52 | 99.9403 | 5051883 | 45.24 | 438 | 20481 |
| SAMN07338229-GCF_006368235.1 | S107 | SAMN07338229 | GCF_006368235.1 | NonPC | Public | NA | NA | Clinical | 89.6 | 99.44 | 0.71 | 98.2814 | 4977621 | 45.31 | 619 | 13562 |
| SAMN07338230-GCF_006368225.1 | S149 | SAMN07338230 | GCF_006368225.1 | NonPC | Public | NA | NA | Clinical | 87.7 | 99.89 | 1.05 | 98.2918 | 5084661 | 45.21 | 612 | 16780 |
| SAMN07338231-GCF_006368595.1 | S175 | SAMN07338231 | GCF_006368595.1 | PC | Public | NA | NA | Clinical | 97.2 | 99.71 | 0.28 | 99.9966 | 5005911 | 45.27 | 374 | 23888 |
| SAMN07338232-GCF_006368215.1 | S176 | SAMN07338232 | GCF_006368215.1 | NonPC | Public | NA | NA | Clinical | 92.5 | 99.98 | 0.11 | 98.4419 | 5055268 | 45.27 | 267 | 37751 |
| SAMN07338233-GCF_006368175.1 | S177 | SAMN07338233 | GCF_006368175.1 | PrePC | Public | NA | NA | NA | 93.1 | 99.98 | 0.35 | 99.2879 | 5019669 | 45.32 | 265 | 37612 |
| SAMN07338234-GCF_006368385.1 | S178 | SAMN07338234 | GCF_006368385.1 | NonPC | Public | NA | NA | Clinical | 91.5 | 99.98 | 0.23 | 98.402 | 5088258 | 45.25 | 330 | 34070 |
| SAMN07341911-GCA_015816935.1 | PNUSAV000057 | SAMN07341911 | GCA_015816935.1 | NonPC | Public | NA | USA | Clinical | 89.5 | 100 | 0 | 98.0487 | 5039789 | 45.26 | 58 | 291897 |
| SAMN07341912-GCA_015817035.1 | PNUSAV000058 | SAMN07341912 | GCA_015817035.1 | NonPC | Public | NA | USA | Clinical | 90.8 | 100 | 0 | 98.2745 | 5078036 | 45.25 | 48 | 557287 |
| SAMN07341913-GCA_016306285.1 | PNUSAV000059 | SAMN07341913 | GCA_016306285.1 | NonPC | Public | NA | USA | Clinical | 89.9 | 100 | 0.04 | 98.2455 | 5200880 | 45.21 | 32 | 784491 |
| SAMN07375601-GCA_015816695.1 | PNUSAV000062 | SAMN07375601 | GCA_015816695.1 | NonPC | Public | NA | USA | Clinical | 88.8 | 100 | 0 | 98.0809 | 4978040 | 45.31 | 78 | 222031 |
| SAMN07428725-GCA_015816875.1 | PNUSAV000064 | SAMN07428725 | GCA_015816875.1 | NonPC | Public | NA | USA | Clinical | 91.5 | 99.98 | 0 | 98.3727 | 4979347 | 45.34 | 54 | 530131 |
| SAMN07428726-GCA_015816915.1 | PNUSAV000065 | SAMN07428726 | GCA_015816915.1 | NonPC | Public | NA | USA | Clinical | 88.5 | 100 | 0.33 | 98.2633 | 5174094 | 45.19 | 88 | 654003 |
| SAMN07428728-GCA_015816895.1 | PNUSAV000063 | SAMN07428728 | GCA_015816895.1 | NonPC | Public | NA | USA | Clinical | 91.1 | 100 | 0.19 | 98.3272 | 5234278 | 45.34 | 39 | 1860971 |
| SAMN07440990-GCA_015816795.1 | PNUSAV000068 | SAMN07440990 | GCA_015816795.1 | NonPC | Public | NA | USA | Clinical | 90.7 | 100 | 0.15 | 98.2652 | 5072350 | 45.25 | 56 | 451230 |
| SAMN07457115-GCA_015816615.1 | PNUSAV000070 | SAMN07457115 | GCA_015816615.1 | NonPC | Public | NA | USA | Clinical | 90.7 | 100 | 0.22 | 98.2724 | 5109480 | 45.24 | 56 | 518451 |
| SAMN07457116-GCA_015816475.1 | PNUSAV000071 | SAMN07457116 | GCA_015816475.1 | NonPC | Public | NA | USA | Clinical | 91.2 | 100 | 0 | 98.4211 | 5095654 | 45.23 | 33 | 541720 |
| SAMN07457117-GCA_015816515.1 | PNUSAV000072 | SAMN07457117 | GCA_015816515.1 | NonPC | Public | NA | USA | Clinical | 90.3 | 100 | 0.14 | 98.21 | 5103264 | 45.14 | 30 | 685827 |
| SAMN07500766-GCA_015816335.1 | PNUSAV000074 | SAMN07500766 | GCA_015816335.1 | NonPC | Public | NA | USA | Clinical | 91.5 | 100 | 0 | 98.2915 | 5146847 | 45.24 | 46 | 549973 |
| SAMN07500768-GCA_015816225.1 | PNUSAV000076 | SAMN07500768 | GCA_015816225.1 | NonPC | Public | NA | USA | Clinical | 91.6 | 100 | 0 | 98.3105 | 5140720 | 45.25 | 41 | 549973 |
| SAMN07500775-GCA_015816775.1 | PNUSAV000073 | SAMN07500775 | GCA_015816775.1 | NonPC | Public | NA | USA | Clinical | 91.6 | 100 | 0 | 98.2709 | 5199123 | 45.25 | 30 | 561124 |
| SAMN07500777-GCA_015816495.1 | PNUSAV000077 | SAMN07500777 | GCA_015816495.1 | NonPC | Public | NA | USA | Clinical | 90.8 | 100 | 0.04 | 98.2786 | 5263934 | 45.28 | 5 | 3368846 |
| SAMN07500778-GCA_015816445.1 | PNUSAV000075 | SAMN07500778 | GCA_015816445.1 | NonPC | Public | NA | USA | Clinical | 91.5 | 100 | 0 | 98.2925 | 5146249 | 45.24 | 46 | 549973 |
| SAMN07526020-GCA_015816755.1 | PNUSAV000080 | SAMN07526020 | GCA_015816755.1 | NonPC | Public | NA | USA | Clinical | 91.5 | 100 | 0 | 98.2901 | 5139181 | 45.24 | 44 | 549973 |
| SAMN07535984-GCA_015816735.1 | PNUSAV000078 | SAMN07535984 | GCA_015816735.1 | NonPC | Public | NA | USA | Clinical | 89.9 | 100 | 0.04 | 98.2573 | 5229635 | 45.19 | 40 | 392622 |

| | | | | | | | | | | | | | | | | |
|---|---|---|---|---|---|---|---|---|---|---|---|---|---|---|---|---|
| SAMN07535985-GCA_015816715.1 | PNUSAV000079 | SAMN07535985 | GCA_015816715.1 | NonPC | Public | NA | USA | Clinical | 91.1 | 100 | 0.08 | 98.2665 | 5052756 | 45.27 | 77 | 323057 |
| SAMN07559594-GCA_015816825.1 | PNUSAV000055 | SAMN07559594 | GCA_015816825.1 | PC | Public | NA | USA | Clinical | 98.3 | 100 | 0 | 99.9855 | 5101365 | 45.28 | 43 | 288887 |
| SAMN07604357-GCA_015806335.1 | PNUSAV000081 | SAMN07604357 | GCA_015806335.1 | NonPC | Public | NA | USA | Clinical | 91.3 | 100 | 0.96 | 98.2745 | 5168746 | 45.26 | 74 | 180521 |
| SAMN07607826-GCA_015816675.1 | PNUSAV000083 | SAMN07607826 | GCA_015816675.1 | NonPC | Public | NA | USA | Clinical | 91.5 | 100 | 0 | 98.2596 | 5147118 | 45.24 | 43 | 549973 |
| SAMN07610980-GCA_015816635.1 | PNUSAV000084 | SAMN07610980 | GCA_015816635.1 | NonPC | Public | NA | USA | Clinical | 91.6 | 100 | 0 | 98.2922 | 5139733 | 45.24 | 44 | 549973 |
| SAMN07629009-GCF_003612715.1 | FORC_071 | SAMN07629009 | GCF_003612715.1 | NonPC | Public | 2017 | South_Korea | Env | 91.4 | 100 | 0.14 | 98.334 | 5173301 | 45.29 | 2 | 3318228 |
| SAMN07629020-GCF_003612695.1 | FORC_072 | SAMN07629020 | GCF_003612695.1 | NonPC | Public | 2017 | South_Korea | Env | 89.5 | 100 | 0.39 | 98.236 | 5188238 | 45.3 | 2 | 3402675 |
| SAMN07637872-GCF_002803235.1 | M1-1 | SAMN07637872 | GCF_002803235.1 | NonPC | Public | 2014 | Viet_Nam | Env | 91.4 | 100 | 0 | 98.3332 | 5474991 | 45.03 | 187 | 170129 |
| SAMN07661832-GCA_015816415.1 | PNUSAV000094 | SAMN07661832 | GCA_015816415.1 | NonPC | Public | NA | USA | Clinical | 89.6 | 100 | 0 | 98.0382 | 5041251 | 45.28 | 73 | 207360 |
| SAMN07666370-GCA_015816375.1 | PNUSAV000086 | SAMN07666370 | GCA_015816375.1 | NonPC | Public | NA | USA | Clinical | 90.6 | 100 | 0 | 98.2623 | 5282710 | 45.17 | 48 | 450130 |
| SAMN07666371-GCA_015816305.1 | PNUSAV000087 | SAMN07666371 | GCA_015816305.1 | NonPC | Public | NA | USA | Clinical | 90.8 | 100 | 0 | 98.2579 | 5073639 | 45.25 | 47 | 511462 |
| SAMN07672045-GCA_015816355.1 | PNUSAV000088 | SAMN07672045 | GCA_015816355.1 | NonPC | Public | NA | USA | Clinical | 91.5 | 100 | 0 | 98.2837 | 5150055 | 45.24 | 44 | 549973 |
| SAMN07680340-GCF_002504185.1 | HA2 | SAMN07680340 | GCF_002504185.1 | NonPC | Public | 2012 | China | Env | 90.4 | 100 | 0 | 98.2525 | 5278641 | 44.96 | 2 | 3423876 |
| SAMN07682410-GCA_015816265.1 | PNUSAV000097 | SAMN07682410 | GCA_015816265.1 | PC | Public | NA | USA | Clinical | 97.9 | 100 | 0 | 99.9566 | 5152373 | 45.23 | 31 | 449415 |
| SAMN07730948-GCA_015814645.1 | PNUSAV000100 | SAMN07730948 | GCA_015814645.1 | NonPC | Public | NA | USA | Clinical | 91.5 | 100 | 0 | 98.2705 | 5139081 | 45.24 | 53 | 329876 |
| SAMN07734741-GCA_015815945.1 | PNUSAV000099 | SAMN07734741 | GCA_015815945.1 | NonPC | Public | NA | USA | Clinical | 90.7 | 100 | 0.15 | 98.3071 | 5071696 | 45.25 | 42 | 530396 |
| SAMN07757022-GCA_015814565.1 | PNUSAV000103 | SAMN07757022 | GCA_015814565.1 | NonPC | Public | NA | USA | Clinical | 91.5 | 100 | 0 | 98.2903 | 5139024 | 45.24 | 44 | 549973 |
| SAMN07780145-GCA_015814585.1 | PNUSAV000105 | SAMN07780145 | GCA_015814585.1 | NonPC | Public | 2017 | USA | Env | 90.3 | 100 | 0 | 98.3238 | 4964199 | 45.3 | 55 | 325678 |
| SAMN07782186-SRR6176852 | PNUSAV000104 | SAMN07782186 | SRR6176852 | NonPC | Public | NA | USA | Clinical | 91.5 | 100 | 0 | 98.294 | 5130735 | 45.25 | 73 | 216448 |
| SAMN07788146-GCA_015815825.1 | PNUSAV000106 | SAMN07788146 | GCA_015815825.1 | NonPC | Public | NA | USA | Clinical | 91.5 | 100 | 0 | 98.3082 | 5136829 | 45.26 | 72 | 217139 |
| SAMN07788147-GCA_015815925.1 | PNUSAV000107 | SAMN07788147 | GCA_015815925.1 | NonPC | Public | NA | USA | Clinical | 91.5 | 100 | 0 | 98.2504 | 5136803 | 45.25 | 78 | 148974 |
| SAMN07806869-GCA_015816005.1 | PNUSAV000108 | SAMN07806869 | GCA_015816005.1 | NonPC | Public | NA | USA | Clinical | 91.6 | 100 | 0 | 98.3229 | 5147974 | 45.25 | 57 | 549960 |
| SAMN07818924-GCF_002887545.1 | PSU5579 | SAMN07818924 | GCF_002887545.1 | NonPC | Public | 2015 | Thailand | Env | 91 | 100 | 0 | 98.3525 | 5229426 | 45.33 | 44 | 268222 |
| SAMN07818995-GCA_015815985.1 | PNUSAV000110 | SAMN07818995 | GCA_015815985.1 | NonPC | Public | NA | USA | Clinical | 91.6 | 100 | 0 | 98.3075 | 5138093 | 45.24 | 50 | 548505 |
| SAMN07830308-GCA_016577205.1 | PNUSAV000111 | SAMN07830308 | GCA_016577205.1 | NonPC | Public | NA | USA | Clinical | 91.3 | 100 | 1.03 | 98.2997 | 5208698 | 45.33 | 97 | 200002 |
| SAMN07830309-GCA_015815675.1 | PNUSAV000112 | SAMN07830309 | GCA_015815675.1 | NonPC | Public | NA | USA | Clinical | 91.5 | 100 | 0 | 98.2987 | 5139075 | 45.25 | 47 | 403495 |
| SAMN07830310-GCA_015815965.1 | PNUSAV000113 | SAMN07830310 | GCA_015815965.1 | NonPC | Public | NA | USA | Clinical | 89.2 | 100 | 0.27 | 98.0624 | 5074038 | 45.24 | 63 | 233622 |
| SAMN07830311-GCA_015815885.1 | PNUSAV000114 | SAMN07830311 | GCA_015815885.1 | NonPC | Public | NA | USA | Clinical | 90.8 | 100 | 0.26 | 98.3638 | 5127347 | 45.36 | 76 | 276049 |
| SAMN07830312-GCA_015815835.1 | PNUSAV000115 | SAMN07830312 | GCA_015815835.1 | NonPC | Public | NA | USA | Clinical | 90 | 100 | 0.04 | 98.2335 | 5228562 | 45.19 | 44 | 330174 |
| SAMN07830313-GCA_015815865.1 | PNUSAV000116 | SAMN07830313 | GCA_015815865.1 | NonPC | Public | NA | USA | Clinical | 90.1 | 100 | 0.46 | 98.3131 | 5232407 | 45.25 | 31 | 432353 |
| SAMN07830314-GCA_015815895.1 | PNUSAV000118 | SAMN07830314 | GCA_015815895.1 | NonPC | Public | NA | USA | Clinical | 90 | 100 | 0.04 | 98.2623 | 5233138 | 45.2 | 39 | 547251 |
| SAMN07831547-GCA_015815465.1 | PNUSAV000126 | SAMN07831547 | GCA_015815465.1 | NonPC | Public | NA | USA | Clinical | 91.5 | 100 | 0 | 98.2921 | 5139388 | 45.24 | 45 | 553160 |
| SAMN07831548-GCA_015815565.1 | PNUSAV000127 | SAMN07831548 | GCA_015815565.1 | NonPC | Public | NA | USA | Clinical | 91.5 | 100 | 0 | 98.2753 | 5147368 | 45.24 | 48 | 480669 |
| SAMN07831549-GCA_015815585.1 | PNUSAV000129 | SAMN07831549 | GCA_015815585.1 | NonPC | Public | NA | USA | Clinical | 91.5 | 100 | 0 | 98.2864 | 5099321 | 45.27 | 39 | 548498 |
| SAMN07831550-GCA_015815705.1 | PNUSAV000130 | SAMN07831550 | GCA_015815705.1 | NonPC | Public | NA | USA | Clinical | 91.5 | 100 | 0 | 98.2816 | 5139729 | 45.24 | 48 | 422857 |
| SAMN07831551-GCA_015815785.1 | PNUSAV000131 | SAMN07831551 | GCA_015815785.1 | NonPC | Public | NA | USA | Clinical | 91.5 | 100 | 0 | 98.2831 | 5139420 | 45.24 | 45 | 549973 |
| SAMN07831552-GCA_015815665.1 | PNUSAV000132 | SAMN07831552 | GCA_015815665.1 | NonPC | Public | NA | USA | Clinical | 91.5 | 100 | 0 | 98.2879 | 5139863 | 45.24 | 44 | 549973 |
| SAMN07831553-GCA_015815805.1 | PNUSAV000133 | SAMN07831553 | GCA_015815805.1 | NonPC | Public | NA | USA | Clinical | 90.4 | 100 | 0 | 98.3074 | 5020507 | 45.27 | 53 | 526037 |
| SAMN07831554-GCA_015815625.1 | PNUSAV000134 | SAMN07831554 | GCA_015815625.1 | NonPC | Public | NA | USA | Clinical | 90.9 | 100 | 0.6 | 98.2819 | 5180166 | 45.18 | 63 | 421304 |
| SAMN07831555-GCA_015815725.1 | PNUSAV000128 | SAMN07831555 | GCA_015815725.1 | NonPC | Public | NA | USA | Clinical | 91.5 | 100 | 0 | 98.2998 | 5139407 | 45.24 | 44 | 553160 |
| SAMN07838934-GCA_015815645.1 | PNUSAV000092 | SAMN07838934 | GCA_015815645.1 | NonPC | Public | NA | USA | Clinical | 90.4 | 100 | 0 | 98.3604 | 4965211 | 45.29 | 54 | 533547 |
| SAMN07838935-GCA_015815735.1 | PNUSAV000093 | SAMN07838935 | GCA_015815735.1 | NonPC | Public | NA | USA | Clinical | 91.5 | 100 | 0 | 98.3208 | 5173458 | 45.22 | 29 | 556449 |
| SAMN07838949-GCA_015815765.1 | PNUSAV000101 | SAMN07838949 | GCA_015815765.1 | PC | Public | NA | USA | Clinical | 97.9 | 100 | 0 | 99.9825 | 5174756 | 45.21 | 49 | 302629 |
| SAMN07838950-GCA_015815605.1 | PNUSAV000102 | SAMN07838950 | GCA_015815605.1 | NonPC | Public | NA | USA | Clinical | 91.5 | 100 | 0 | 98.3277 | 5092202 | 45.27 | 29 | 549859 |
| SAMN07956717-GCF_003174825.1 | 16-B3PA-006 | SAMN07956717 | GCF_003174825.1 | NonPC | Public | 2016 | Viet_Nam | Env | 91.6 | 100 | 0.6 | 98.5823 | 5184396 | 45.65 | 190 | 83566 |
| SAMN07964850-GCA_015815425.1 | PNUSAV000119 | SAMN07964850 | GCA_015815425.1 | NonPC | Public | NA | USA | Clinical | 91.6 | 100 | 0 | 98.3075 | 5138397 | 45.25 | 54 | 234360 |
| SAMN07964851-GCA_015815345.1 | PNUSAV000120 | SAMN07964851 | GCA_015815345.1 | NonPC | Public | NA | USA | Clinical | 91.5 | 100 | 0 | 98.2595 | 5135356 | 45.26 | 73 | 221306 |
| SAMN07964852-GCA_015815485.1 | PNUSAV000121 | SAMN07964852 | GCA_015815485.1 | NonPC | Public | NA | USA | Clinical | 91.5 | 100 | 0 | 98.2763 | 5138230 | 45.25 | 54 | 267089 |

| | | | | | | | | | | | | | | | | |
|---|---|---|---|---|---|---|---|---|---|---|---|---|---|---|---|---|
| SAMN07964853-GCA_015815305.1 | PNUSAV000122 | SAMN07964853 | GCA_015815305.1 | NonPC | Public | NA | USA | Clinical | 91.5 | 100 | 0 | 98.2583 | 5135603 | 45.26 | 66 | 289756 |
| SAMN07964854-GCA_015815515.1 | PNUSAV000123 | SAMN07964854 | GCA_015815515.1 | NonPC | Public | NA | USA | Clinical | 91.6 | 100 | 0 | 98.2806 | 5139084 | 45.24 | 41 | 548505 |
| SAMN07964855-GCA_015815405.1 | PNUSAV000124 | SAMN07964855 | GCA_015815405.1 | NonPC | Public | NA | USA | Clinical | 91.5 | 100 | 0 | 98.2816 | 5138753 | 45.24 | 42 | 553160 |
| SAMN07964856-GCA_015815505.1 | PNUSAV000125 | SAMN07964856 | GCA_015815505.1 | NonPC | Public | NA | USA | Clinical | 91.5 | 100 | 0 | 98.2832 | 5139028 | 45.24 | 49 | 404359 |
| SAMN08001963-GCA_015815445.1 | PNUSAV000145 | SAMN08001963 | GCA_015815445.1 | NonPC | Public | NA | USA | Clinical | 90.7 | 100 | 0.14 | 98.1631 | 5055412 | 45.21 | 57 | 197144 |
| SAMN08007025-GCF_002775145.1 | 1911C | SAMN08007025 | GCF_002775145.1 | NonPC | Public | 2008 | USA | Env | 87.8 | 100 | 0.3 | 98.0523 | 4988624 | 45.33 | 36 | 326797 |
| SAMN08007337-GCA_015815225.1 | PNUSAV000146 | SAMN08007337 | GCA_015815225.1 | NonPC | Public | NA | USA | Clinical | 91.1 | 100 | 0.21 | 98.2874 | 5199346 | 45.25 | 31 | 674169 |
| SAMN08007338-SRR6293335 | PNUSAV000147 | SAMN08007338 | SRR6293335 | NonPC | Public | NA | USA | Clinical | 90.8 | 100 | 0.11 | 98.2156 | 5060305 | 45.3 | 44 | 756034 |
| SAMN08007339-GCA_015815355.1 | PNUSAV000148 | SAMN08007339 | GCA_015815355.1 | NonPC | Public | NA | USA | Clinical | 91.5 | 100 | 0.14 | 98.3264 | 5044897 | 45.28 | 54 | 516355 |
| SAMN08016203-GCA_015815145.1 | PNUSAV000137 | SAMN08016203 | GCA_015815145.1 | NonPC | Public | NA | USA | Clinical | 89.9 | 100 | 0.05 | 98.3155 | 5009923 | 45.36 | 64 | 479524 |
| SAMN08016204-GCA_015815365.1 | PNUSAV000142 | SAMN08016204 | GCA_015815365.1 | NonPC | Public | NA | USA | Clinical | 90.5 | 100 | 0.05 | 98.3502 | 4938561 | 45.49 | 45 | 715349 |
| SAMN08016205-GCA_015815315.1 | PNUSAV000143 | SAMN08016205 | GCA_015815315.1 | NonPC | Public | NA | USA | Clinical | 89.7 | 100 | 0.49 | 98.2384 | 5238580 | 45.18 | 47 | 315032 |
| SAMN08016206-GCA_015815285.1 | PNUSAV000144 | SAMN08016206 | GCA_015815285.1 | NonPC | Public | NA | USA | Clinical | 91.5 | 100 | 0.15 | 98.3135 | 5116358 | 45.26 | 44 | 548242 |
| SAMN08024706-GCA_015815245.1 | PNUSAV000117 | SAMN08024706 | GCA_015815245.1 | NonPC | Public | NA | USA | Clinical | 91.2 | 100 | 0 | 98.284 | 5166499 | 45.19 | 56 | 340448 |
| SAMN08024707-GCA_015815265.1 | PNUSAV000149 | SAMN08024707 | GCA_015815265.1 | NonPC | Public | NA | USA | Clinical | 89.9 | 100 | 0.04 | 98.2459 | 5216020 | 45.2 | 43 | 241631 |
| SAMN08027481-GCA_015814705.1 | PNUSAV000150 | SAMN08027481 | GCA_015814705.1 | NonPC | Public | NA | USA | Clinical | 92 | 100 | 0.35 | 98.2938 | 5126217 | 45.23 | 61 | 215883 |
| SAMN08104782-GCA_020410965.1 | PNUSAV000151 | SAMN08104782 | GCA_020410965.1 | NonPC | Public | NA | USA | Clinical | 90.6 | 100 | 3.85 | 98.3297 | 5362543 | 45.32 | 66 | 233348 |
| SAMN08104783-GCA_020410925.1 | PNUSAV000152 | SAMN08104783 | GCA_020410925.1 | NonPC | Public | NA | USA | Clinical | 89.9 | 100 | 1.27 | 98.3638 | 5081629 | 45.27 | 68 | 311099 |
| SAMN08113866-GCA_015816185.1 | PNUSAV000138 | SAMN08113866 | GCA_015816185.1 | NonPC | Public | NA | USA | Clinical | 91.3 | 100 | 0.05 | 98.3938 | 5012951 | 45.41 | 53 | 452967 |
| SAMN08113867-GCA_015815165.1 | PNUSAV000139 | SAMN08113867 | GCA_015815165.1 | NonPC | Public | NA | USA | Clinical | 91.5 | 100 | 0 | 98.2989 | 5101433 | 45.27 | 37 | 557043 |
| SAMN08113868-GCA_015814725.1 | PNUSAV000140 | SAMN08113868 | GCA_015814725.1 | PC | Public | NA | USA | Clinical | 97.8 | 100 | 0 | 99.9316 | 5148887 | 45.24 | 32 | 512926 |
| SAMN08113869-GCA_015814665.1 | PNUSAV000141 | SAMN08113869 | GCA_015814665.1 | NonPC | Public | NA | USA | Clinical | 91.1 | 100 | 0.17 | 98.3978 | 4995916 | 45.45 | 34 | 1231410 |
| SAMN08130118-GCA_015815205.1 | PNUSAV000153 | SAMN08130118 | GCA_015815205.1 | NonPC | Public | 2014 | USA | Clinical | 91.5 | 100 | 0 | 98.2931 | 5100378 | 45.27 | 40 | 548498 |
| SAMN08130119-GCA_015815085.1 | PNUSAV000154 | SAMN08130119 | GCA_015815085.1 | NonPC | Public | 2016 | USA | Clinical | 90.4 | 100 | 0 | 98.3058 | 4967263 | 45.29 | 59 | 524598 |
| SAMN08162841-GCF_003186355.1 | KS17.S5-1 | SAMN08162841 | GCF_003186355.1 | NonPC | Public | 2017 | Malaysia | Env | 91.2 | 100 | 0.05 | 98.307 | 5296440 | 45.21 | 57 | 405134 |
| SAMN08166448-GCF_003186805.1 | ST17.P5-S1 | SAMN08166448 | GCF_003186805.1 | NonPC | Public | 2017 | Malaysia | Env | 91.3 | 100 | 0.86 | 98.235 | 5446640 | 45.25 | 103 | 199471 |
| SAMN08180259-GCA_015814985.1 | PNUSAV000159 | SAMN08180259 | GCA_015814985.1 | NonPC | Public | NA | USA | Clinical | 89.3 | 100 | 0 | 98.0591 | 5064753 | 45.27 | 78 | 319036 |
| SAMN08180260-GCA_015815105.1 | PNUSAV000160 | SAMN08180260 | GCA_015815105.1 | NonPC | Public | NA | USA | Clinical | 91.5 | 100 | 0 | 98.3148 | 5130585 | 45.26 | 39 | 548500 |
| SAMN08180261-GCA_015814905.1 | PNUSAV000161 | SAMN08180261 | GCA_015814905.1 | NonPC | Public | NA | USA | Clinical | 89.4 | 100 | 0 | 98.0602 | 5123668 | 45.25 | 80 | 380592 |
| SAMN08180262-GCA_015815045.1 | PNUSAV000162 | SAMN08180262 | GCA_015815045.1 | NonPC | Public | NA | USA | Clinical | 91.5 | 100 | 0 | 98.2776 | 5148217 | 45.25 | 25 | 1263981 |
| SAMN08180263-GCA_015815005.1 | PNUSAV000163 | SAMN08180263 | GCA_015815005.1 | NonPC | Public | NA | USA | Clinical | 90.9 | 100 | 0.6 | 98.2759 | 5182513 | 45.19 | 58 | 403418 |
| SAMN08180264-GCA_015814925.1 | PNUSAV000164 | SAMN08180264 | GCA_015814925.1 | NonPC | Public | NA | USA | Clinical | 89.4 | 100 | 0.27 | 98.0668 | 5071961 | 45.24 | 59 | 379650 |
| SAMN08180265-GCA_015815125.1 | PNUSAV000165 | SAMN08180265 | GCA_015815125.1 | NonPC | Public | NA | USA | Clinical | 89.4 | 100 | 0.27 | 98.0793 | 5071595 | 45.24 | 64 | 306811 |
| SAMN08182975-GCA_015814955.1 | PNUSAV000135 | SAMN08182975 | GCA_015814955.1 | NonPC | Public | NA | USA | Clinical | 89.4 | 100 | 0 | 98.0953 | 5073126 | 45.28 | 111 | 126662 |
| SAMN08182993-GCA_015815025.1 | PNUSAV000167 | SAMN08182993 | GCA_015815025.1 | NonPC | Public | NA | USA | Clinical | 91.4 | 100 | 0 | 98.3514 | 5017438 | 45.39 | 106 | 128404 |
| SAMN08204774-GCF_003798505.1 | Lovell_lab_04 | SAMN08204774 | GCF_003798505.1 | NonPC | Public | 2013 | USA | Env | 89.2 | 100 | 0.27 | 98.0808 | 4989423 | 45.38 | 87 | 190904 |
| SAMN08225440-GCA_015814885.1 | C21_(1262-07) | SAMN08225440 | GCA_015814885.1 | NonPC | Public | 2007 | Peru | Clinical | 92.2 | 100 | 0.46 | 98.3573 | 5022946 | 45.25 | 37 | 547499 |
| SAMN08225451-GCA_015814785.1 | C4-324-95 | SAMN08225451 | GCA_015814785.1 | NonPC | Public | 1995 | Peru | Clinical | 92.6 | 99.73 | 0.11 | 98.3365 | 5141146 | 45.23 | 50 | 450591 |
| SAMN08225452-GCA_015814855.1 | C5-326-95 | SAMN08225452 | GCA_015814855.1 | NonPC | Public | 1995 | Peru | Clinical | 92.6 | 100 | 0.11 | 98.3534 | 5141689 | 45.23 | 52 | 662339 |
| SAMN08225453-GCA_015814825.1 | C6-267-96 | SAMN08225453 | GCA_015814825.1 | NonPC | Public | 1996 | Peru | Clinical | 92.5 | 100 | 0.11 | 98.347 | 5198123 | 45.22 | 54 | 526822 |
| SAMN08225454-GCA_015814845.1 | C7-212-96 | SAMN08225454 | GCA_015814845.1 | NonPC | Public | 1996 | Peru | Clinical | 92.3 | 100 | 0.19 | 98.3456 | 5098035 | 45.23 | 27 | 698164 |
| SAMN08225455-GCA_015815065.1 | C9-1257-07 | SAMN08225455 | GCA_015815065.1 | NonPC | Public | 2007 | Peru | Clinical | 92.2 | 100 | 0.46 | 98.3386 | 5023903 | 45.25 | 33 | 545947 |
| SAMN08225457-GCA_015814805.1 | CFSAN029655 | SAMN08225457 | GCA_015814805.1 | PC | Public | 2009 | Peru | Env | 99.9 | 100 | 0 | 99.9958 | 5168009 | 45.36 | 14 | 841949 |
| SAMN08225458-GCA_015814745.1 | CFSAN029652 | SAMN08225458 | GCA_015814745.1 | PC | Public | 2009 | Peru | Env | 99.9 | 100 | 0 | 99.9944 | 5167901 | 45.36 | 15 | 1127066 |
| SAMN08225459-GCA_015814685.1 | CFSAN029657 | SAMN08225459 | GCA_015814685.1 | PC | Public | 2009 | Peru | Env | 99.9 | 100 | 0 | 99.9956 | 5168713 | 45.36 | 3 | 3287991 |
| SAMN08235473-GCA_015816105.1 | PNUSAV000157 | SAMN08235473 | GCA_015816105.1 | NonPC | Public | NA | USA | Clinical | 91.7 | 100 | 0.29 | 98.2995 | 5115185 | 45.27 | 33 | 417976 |
| SAMN08275786-GCA_015814765.1 | PNUSAV000168 | SAMN08275786 | GCA_015814765.1 | NonPC | Public | NA | USA | Clinical | 90 | 100 | 0.04 | 98.2744 | 5227549 | 45.2 | 52 | 247982 |
| SAMN08275787-GCA_015816145.1 | PNUSAV000169 | SAMN08275787 | GCA_015816145.1 | NonPC | Public | NA | USA | Clinical | 89.9 | 100 | 0.04 | 98.2603 | 5227637 | 45.21 | 72 | 164979 |

| | | | | | | | | | | | | | | | | |
|---|---|---|---|---|---|---|---|---|---|---|---|---|---|---|---|---|
| SAMN08329233-GCF_002880595.1 | MVP3 | SAMN08329233 | GCF_002880595.1 | NonPC | Public | 2016 | Malaysia | Env | 90.1 | 100 | 0.27 | 98.3434 | 5337197 | 45.26 | 64 | 384183 |
| SAMN08329234-GCF_002880555.1 | MVP5 | SAMN08329234 | GCF_002880555.1 | NonPC | Public | 2016 | Malaysia | Env | 89.8 | 100 | 0.27 | 98.2774 | 5300835 | 45.26 | 100 | 229987 |
| SAMN08329235-GCF_002880505.1 | MVP10 | SAMN08329235 | GCF_002880505.1 | NonPC | Public | 2016 | Malaysia | Env | 89.5 | 100 | 0.26 | 98.3145 | 5004679 | 45.39 | 40 | 485472 |
| SAMN08329236-GCF_002880575.1 | NA1 | SAMN08329236 | GCF_002880575.1 | NonPC | Public | 2017 | Malaysia | Env | 91.8 | 100 | 0.14 | 98.2526 | 5505729 | 45.29 | 82 | 202086 |
| SAMN08329237-GCF_002880475.1 | NA4 | SAMN08329237 | GCF_002880475.1 | NonPC | Public | 2017 | Malaysia | Env | 88.3 | 100 | 0.3 | 98.2494 | 5066938 | 45.26 | 42 | 887465 |
| SAMN08329238-GCF_002880495.1 | NA6 | SAMN08329238 | GCF_002880495.1 | NonPC | Public | 2017 | Malaysia | Env | 90.9 | 100 | 0.54 | 98.353 | 5470115 | 45.22 | 99 | 148165 |
| SAMN08329239-GCF_002880415.1 | NA7 | SAMN08329239 | GCF_002880415.1 | NonPC | Public | 2017 | Malaysia | Env | 88.1 | 100 | 0.3 | 98.2203 | 5054162 | 45.26 | 60 | 387482 |
| SAMN08329240-GCF_002880535.1 | NA8 | SAMN08329240 | GCF_002880535.1 | NonPC | Public | 2017 | Malaysia | Env | 90.9 | 100 | 0.54 | 98.3425 | 5473387 | 45.21 | 69 | 311576 |
| SAMN08329241-GCF_002880435.1 | NA9 | SAMN08329241 | GCF_002880435.1 | NonPC | Public | 2017 | Malaysia | Env | 88.3 | 100 | 0.3 | 98.2622 | 5067933 | 45.26 | 41 | 1156426 |
| SAMN08329242-GCF_002880455.1 | NB07 | SAMN08329242 | GCF_002880455.1 | NonPC | Public | 2017 | Malaysia | Env | 90.9 | 100 | 0.54 | 98.3288 | 5477693 | 45.21 | 79 | 308724 |
| SAMN08329243-GCF_002880375.1 | NB08 | SAMN08329243 | GCF_002880375.1 | NonPC | Public | 2017 | Malaysia | Env | 90.9 | 100 | 0.54 | 98.3652 | 5475102 | 45.21 | 95 | 259462 |
| SAMN08329244-GCF_002880355.1 | NB09 | SAMN08329244 | GCF_002880355.1 | NonPC | Public | 2017 | Malaysia | Env | 90.9 | 100 | 0.54 | 98.3317 | 5466869 | 45.22 | 88 | 308483 |
| SAMN08329245-GCF_002880385.1 | NB10 | SAMN08329245 | GCF_002880385.1 | NonPC | Public | 2017 | Malaysia | Env | 90.9 | 100 | 0.54 | 98.3226 | 5474429 | 45.21 | 68 | 310637 |
| SAMN08329246-GCF_002880315.1 | NB11 | SAMN08329246 | GCF_002880315.1 | NonPC | Public | 2017 | Malaysia | Env | 90.9 | 100 | 0.54 | 98.3305 | 5472865 | 45.21 | 104 | 247474 |
| SAMN08329247-GCF_002880335.1 | NB12 | SAMN08329247 | GCF_002880335.1 | NonPC | Public | 2017 | Malaysia | Env | 90.9 | 100 | 0.54 | 98.2817 | 5464241 | 45.21 | 44 | 451962 |
| SAMN08329248-GCF_002880295.1 | NB13 | SAMN08329248 | GCF_002880295.1 | NonPC | Public | 2017 | Malaysia | Env | 90.9 | 100 | 0.54 | 98.332 | 5476168 | 45.21 | 75 | 398758 |
| SAMN08329249-GCF_002880275.1 | ND11 | SAMN08329249 | GCF_002880275.1 | NonPC | Public | 2017 | Malaysia | Env | 91.6 | 100 | 0.15 | 98.2565 | 5580136 | 45.25 | 66 | 202086 |
| SAMN08329250-GCF_002880255.1 | ND12 | SAMN08329250 | GCF_002880255.1 | NonPC | Public | 2017 | Malaysia | Env | 91.8 | 100 | 0.14 | 98.2669 | 5499508 | 45.29 | 78 | 189547 |
| SAMN08329251-GCF_002880225.1 | ND13 | SAMN08329251 | GCF_002880225.1 | NonPC | Public | 2017 | Malaysia | Env | 91.8 | 100 | 0.14 | 98.2676 | 5507382 | 45.28 | 88 | 189547 |
| SAMN08329252-GCF_002880215.1 | ND14 | SAMN08329252 | GCF_002880215.1 | NonPC | Public | 2017 | Malaysia | Env | 90.9 | 100 | 0.54 | 98.3346 | 5476115 | 45.21 | 64 | 399294 |
| SAMN08329253-GCF_002880195.1 | ND15 | SAMN08329253 | GCF_002880195.1 | NonPC | Public | 2017 | Malaysia | Env | 90.9 | 100 | 0.54 | 98.3361 | 5475418 | 45.21 | 87 | 295691 |
| SAMN08329254-GCF_002880175.1 | ND16 | SAMN08329254 | GCF_002880175.1 | NonPC | Public | 2017 | Malaysia | Env | 91.8 | 100 | 0.14 | 98.2683 | 5503110 | 45.28 | 69 | 205901 |
| SAMN08329255-GCF_002880155.1 | ND17 | SAMN08329255 | GCF_002880155.1 | NonPC | Public | 2017 | Malaysia | Env | 90.9 | 100 | 0.54 | 98.3192 | 5475415 | 45.21 | 85 | 295691 |
| SAMN08329256-GCF_002880095.1 | ND18 | SAMN08329256 | GCF_002880095.1 | NonPC | Public | 2017 | Malaysia | Env | 90.9 | 100 | 0.54 | 98.3098 | 5467141 | 45.21 | 57 | 313562 |
| SAMN08329257-GCF_002880135.1 | ND19 | SAMN08329257 | GCF_002880135.1 | NonPC | Public | 2017 | Malaysia | Env | 91.8 | 100 | 0.14 | 98.259 | 5501562 | 45.28 | 95 | 202086 |
| SAMN08329258-GCF_002880085.1 | ND20 | SAMN08329258 | GCF_002880085.1 | NonPC | Public | 2017 | Malaysia | Env | 90.9 | 100 | 0.54 | 98.3368 | 5473683 | 45.21 | 85 | 201716 |
| SAMN08329259-GCF_002880015.1 | ND21 | SAMN08329259 | GCF_002880015.1 | NonPC | Public | 2017 | Malaysia | Env | 90.9 | 99.91 | 0.54 | 98.3291 | 5473291 | 45.2 | 90 | 295525 |
| SAMN08329260-GCF_002880075.1 | ND22 | SAMN08329260 | GCF_002880075.1 | NonPC | Public | 2017 | Malaysia | Env | 91.9 | 100 | 0.14 | 98.249 | 5498980 | 45.29 | 94 | 189547 |
| SAMN08329261-GCF_002880035.1 | ND23 | SAMN08329261 | GCF_002880035.1 | NonPC | Public | 2017 | Malaysia | Env | 90.9 | 100 | 0.54 | 98.3327 | 5467535 | 45.21 | 69 | 247474 |
| SAMN08329262-GCF_002880025.1 | ND24 | SAMN08329262 | GCF_002880025.1 | NonPC | Public | 2017 | Malaysia | Env | 91.8 | 100 | 0.14 | 98.2486 | 5507062 | 45.29 | 101 | 180175 |
| SAMN08361201-GCA_015816165.1 | CFSAN074483 | SAMN08361201 | GCA_015816165.1 | NonPC | Public | 2012 | USA | Env | 88.7 | 100 | 0.35 | 98.1978 | 4953270 | 45.36 | 46 | 262126 |
| SAMN08361212-GCA_015814525.1 | CFSAN074484 | SAMN08361212 | GCA_015814525.1 | NonPC | Public | 2012 | USA | Env | 88.6 | 100 | 0.35 | 98.1988 | 4945129 | 45.35 | 36 | 358438 |
| SAMN08370010-GCA_015816045.1 | CFSAN026727 | SAMN08370010 | GCA_015816045.1 | NonPC | Public | 2013 | USA | Clinical | 89.7 | 100 | 0 | 98.1203 | 5085832 | 45.34 | 91 | 208593 |
| SAMN08370011-GCA_015816085.1 | CFSAN026728 | SAMN08370011 | GCA_015816085.1 | NonPC | Public | 2013 | USA | Clinical | 87.8 | 100 | 0.25 | 98.1086 | 5057456 | 45.27 | 102 | 262267 |
| SAMN08370012-GCA_015816025.1 | CFSAN026729 | SAMN08370012 | GCA_015816025.1 | NonPC | Public | 2013 | USA | Clinical | 91.5 | 100 | 0 | 98.3351 | 5096605 | 45.27 | 33 | 461989 |
| SAMN08370013-GCA_015814455.1 | CFSAN026730 | SAMN08370013 | GCA_015814455.1 | NonPC | Public | 2013 | USA | Clinical | 91.5 | 100 | 0 | 98.315 | 5096763 | 45.27 | 31 | 462006 |
| SAMN08370014-GCA_015814605.1 | CFSAN026731 | SAMN08370014 | GCA_015814605.1 | NonPC | Public | 2013 | USA | Clinical | 90.9 | 100 | 0.05 | 98.3638 | 5178085 | 45.31 | 31 | 1849186 |
| SAMN08370015-GCA_015814625.1 | CFSAN026732 | SAMN08370015 | GCA_015814625.1 | PC | Public | 2014 | USA | Clinical | 99.7 | 100 | 0.27 | 99.9873 | 5225471 | 45.38 | 13 | 2928409 |
| SAMN08370016-GCA_015816065.1 | CFSAN026733 | SAMN08370016 | GCA_015816065.1 | NonPC | Public | 2014 | USA | Clinical | 88.3 | 100 | 0.52 | 98.1297 | 5074733 | 45.41 | 66 | 271322 |
| SAMN08469470-GCA_015811865.1 | CFSAN075755 | SAMN08469470 | GCA_015811865.1 | NonPC | Public | 2012 | USA | Env | 91.1 | 100 | 0.41 | 98.4072 | 5075864 | 45.23 | 45 | 533428 |
| SAMN08469471-GCA_015811965.1 | CFSAN075756 | SAMN08469471 | GCA_015811965.1 | NonPC | Public | 2012 | USA | Env | 89.2 | 100 | 0 | 98.0698 | 5083825 | 45.28 | 65 | 287386 |
| SAMN08469472-GCA_015811985.1 | CFSAN075738 | SAMN08469472 | GCA_015811985.1 | PrePC | Public | 2012 | USA | Env | 94.9 | 100 | 0.08 | 99.8453 | 5088550 | 45.28 | 41 | 547275 |
| SAMN08469473-GCA_015812125.1 | CFSAN075737 | SAMN08469473 | GCA_015812125.1 | NonPC | Public | 2012 | USA | Env | 90.5 | 100 | 0.14 | 98.2506 | 5155164 | 45.23 | 34 | 487880 |
| SAMN08469474-GCA_015811805.1 | CFSAN075740 | SAMN08469474 | GCA_015811805.1 | NonPC | Public | 2012 | USA | Env | 91.2 | 100 | 2.86 | 98.3121 | 5356493 | 45.22 | 76 | 285810 |
| SAMN08469475-GCA_015811845.1 | CFSAN075739 | SAMN08469475 | GCA_015811845.1 | NonPC | Public | 2012 | USA | Env | 90.5 | 100 | 0 | 98.287 | 5250464 | 45.22 | 79 | 265467 |
| SAMN08469476-GCA_015808375.1 | CFSAN075733 | SAMN08469476 | GCA_015808375.1 | NonPC | Public | 2012 | USA | Env | 88.6 | 100 | 0.36 | 98.2093 | 4871666 | 45.48 | 93 | 170920 |
| SAMN08469477-GCA_015811825.1 | CFSAN075734 | SAMN08469477 | GCA_015811825.1 | NonPC | Public | 2012 | USA | Env | 88.7 | 100 | 0.38 | 98.2151 | 4910838 | 45.41 | 80 | 210613 |
| SAMN08469478-GCA_015812045.1 | CFSAN075736 | SAMN08469478 | GCA_015812045.1 | NonPC | Public | 2012 | USA | Env | 88.9 | 99.73 | 0.14 | 98.1344 | 4952179 | 45.32 | 53 | 237573 |

| | | | | | | | | | | | | | | | | |
|---|---|---|---|---|---|---|---|---|---|---|---|---|---|---|---|---|
| SAMN08469544-GCA_015811895.1 | CFSAN075729 | SAMN08469544 | GCA_015811895.1 | NonPC | Public | 2012 | USA | Env | 90.1 | 100 | 0.22 | 98.2153 | 5048047 | 45.33 | 64 | 175619 |
| SAMN08469545-GCA_015810425.1 | CFSAN075745 | SAMN08469545 | GCA_015810425.1 | NonPC | Public | 2012 | USA | Env | 88.9 | 99.73 | 0.14 | 98.1308 | 4942271 | 45.33 | 65 | 227800 |
| SAMN08469546-GCA_015812165.1 | CFSAN075744 | SAMN08469546 | GCA_015812165.1 | NonPC | Public | 2012 | USA | Env | 89.1 | 100 | 0 | 98.096 | 5093709 | 45.28 | 64 | 313531 |
| SAMN08469547-GCA_015810465.1 | CFSAN075747 | SAMN08469547 | GCA_015810465.1 | NonPC | Public | 2012 | USA | Env | 91.9 | 100 | 0.14 | 98.4353 | 5343371 | 45.08 | 72 | 301296 |
| SAMN08469548-GCA_015811885.1 | CFSAN075753 | SAMN08469548 | GCA_015811885.1 | NonPC | Public | 2012 | USA | Env | 89.2 | 100 | 0 | 98.1154 | 5084313 | 45.28 | 64 | 313531 |
| SAMN08469549-GCA_015812145.1 | CFSAN075741 | SAMN08469549 | GCA_015812145.1 | NonPC | Public | 2012 | USA | Env | 90.5 | 100 | 0 | 98.3158 | 5247842 | 45.23 | 82 | 286711 |
| SAMN08469550-GCA_015809395.1 | CFSAN075752 | SAMN08469550 | GCA_015809395.1 | NonPC | Public | 2012 | USA | Env | 88.6 | 100 | 0.35 | 98.1445 | 4959018 | 45.35 | 62 | 241953 |
| SAMN08469552-GCA_015811705.1 | CFSAN075732 | SAMN08469552 | GCA_015811705.1 | NonPC | Public | 2012 | USA | Env | 89.3 | 100 | 0 | 98.1338 | 5022760 | 45.33 | 57 | 281926 |
| SAMN08469555-GCA_015812225.1 | CFSAN075742 | SAMN08469555 | GCA_015812225.1 | NonPC | Public | 2012 | USA | Env | 89.2 | 100 | 0 | 98.0978 | 5247144 | 45.21 | 63 | 287387 |
| SAMN08469556-GCA_015811925.1 | CFSAN075743 | SAMN08469556 | GCA_015811925.1 | NonPC | Public | 2012 | USA | Env | 91 | 100 | 2.86 | 98.3148 | 5376212 | 45.19 | 52 | 336406 |
| SAMN08469557-GCA_015811755.1 | CFSAN075731 | SAMN08469557 | GCA_015811755.1 | NonPC | Public | 2012 | USA | Env | 89.3 | 100 | 0 | 98.1643 | 5014926 | 45.34 | 59 | 281926 |
| SAMN08469558-GCA_015811785.1 | CFSAN075749 | SAMN08469558 | GCA_015811785.1 | NonPC | Public | 2012 | USA | Env | 91.3 | 100 | 0.07 | 98.3082 | 5101697 | 45.29 | 46 | 376855 |
| SAMN08469559-GCA_015811725.1 | CFSAN075730 | SAMN08469559 | GCA_015811725.1 | NonPC | Public | 2012 | USA | Env | 90.2 | 100 | 0.22 | 98.1956 | 5054840 | 45.32 | 58 | 234817 |
| SAMN08469560-GCA_015811945.1 | CFSAN075746 | SAMN08469560 | GCA_015811945.1 | NonPC | Public | 2012 | USA | Env | 88.9 | 99.73 | 0.14 | 98.1259 | 4952513 | 45.3 | 39 | 330796 |
| SAMN08469561-GCA_015812185.1 | CFSAN075751 | SAMN08469561 | GCA_015812185.1 | NonPC | Public | 2012 | USA | Env | 91.1 | 100 | 0.08 | 98.3119 | 5226946 | 45.28 | 79 | 312686 |
| SAMN08469562-GCA_015812005.1 | CFSAN075750 | SAMN08469562 | GCA_015812005.1 | PrePC | Public | 2012 | USA | Env | 94.9 | 100 | 0.08 | 99.8425 | 5090997 | 45.29 | 42 | 430792 |
| SAMN08473219-GCA_015813125.1 | CFSAN075860 | SAMN08473219 | GCA_015813125.1 | NonPC | Public | 2011 | USA | Env | 90.8 | 100 | 0 | 98.2484 | 5110660 | 45.24 | 74 | 289081 |
| SAMN08473220-GCA_015813845.1 | CFSAN075869 | SAMN08473220 | GCA_015813845.1 | NonPC | Public | 2012 | USA | Env | 91.2 | 100 | 0.12 | 98.2954 | 5121512 | 45.3 | 45 | 749996 |
| SAMN08473221-GCA_015814405.1 | CFSAN075870 | SAMN08473221 | GCA_015814405.1 | NonPC | Public | 2012 | USA | Env | 90.8 | 100 | 0.08 | 98.2979 | 5087255 | 45.4 | 30 | 1221832 |
| SAMN08473223-GCA_015813705.1 | CFSAN075866 | SAMN08473223 | GCA_015813705.1 | NonPC | Public | 2012 | USA | Env | 90.4 | 100 | 0 | 98.2532 | 5021487 | 45.44 | 48 | 955675 |
| SAMN08473224-GCA_015813885.1 | CFSAN075867 | SAMN08473224 | GCA_015813885.1 | NonPC | Public | 2012 | USA | Env | 90.4 | 100 | 0 | 98.2877 | 5021538 | 45.45 | 51 | 700372 |
| SAMN08473225-GCA_015813945.1 | CFSAN075872 | SAMN08473225 | GCA_015813945.1 | NonPC | Public | 2012 | USA | Env | 89.1 | 100 | 0.39 | 98.2819 | 5016268 | 45.43 | 41 | 1529695 |
| SAMN08473226-GCA_015814325.1 | CFSAN075808 | SAMN08473226 | GCA_015814325.1 | NonPC | Public | 2010 | USA | Env | 88.4 | 100 | 0.36 | 98.1869 | 5325044 | 45 | 50 | 383278 |
| SAMN08473227-GCA_015814345.1 | CFSAN075822 | SAMN08473227 | GCA_015814345.1 | NonPC | Public | 2011 | USA | Env | 91.3 | 100 | 0 | 98.2357 | 5156375 | 45.24 | 52 | 440691 |
| SAMN08473228-GCA_015814505.1 | CFSAN075823 | SAMN08473228 | GCA_015814505.1 | NonPC | Public | 2011 | USA | Env | 90.9 | 100 | 0.22 | 98.3113 | 5120290 | 45.23 | 51 | 545869 |
| SAMN08473230-GCA_015813805.1 | CFSAN075813 | SAMN08473230 | GCA_015813805.1 | NonPC | Public | 2010 | USA | Env | 88.4 | 100 | 0.47 | 98.1927 | 5335875 | 45.02 | 59 | 366182 |
| SAMN08473231-GCA_015813245.1 | CFSAN075801 | SAMN08473231 | GCA_015813245.1 | NonPC | Public | 2009 | USA | Env | 92.1 | 100 | 0.49 | 98.2451 | 5277728 | 45.15 | 52 | 605694 |
| SAMN08473235-GCA_015813745.1 | CFSAN075815 | SAMN08473235 | GCA_015813745.1 | NonPC | Public | 2010 | USA | Env | 92 | 100 | 0.49 | 98.2573 | 5267912 | 45.15 | 55 | 575153 |
| SAMN08473236-GCA_015813785.1 | CFSAN075814 | SAMN08473236 | GCA_015813785.1 | NonPC | Public | 2010 | USA | Env | 88.5 | 100 | 0.41 | 98.1909 | 5341195 | 45.02 | 50 | 366192 |
| SAMN08473237-GCA_015813045.1 | CFSAN075817 | SAMN08473237 | GCA_015813045.1 | NonPC | Public | 2010 | USA | Env | 92.6 | 100 | 0 | 98.3343 | 5155849 | 45.33 | 61 | 481482 |
| SAMN08473238-GCA_015813865.1 | CFSAN075816 | SAMN08473238 | GCA_015813865.1 | NonPC | Public | 2010 | USA | Env | 92 | 100 | 0.49 | 98.2682 | 5268483 | 45.16 | 53 | 538263 |
| SAMN08473240-GCA_015814485.1 | CFSAN075841 | SAMN08473240 | GCA_015814485.1 | NonPC | Public | 2010 | USA | Env | 90.9 | 100 | 0.5 | 98.2761 | 5256215 | 45.19 | 67 | 489063 |
| SAMN08473241-GCA_015813685.1 | CFSAN075819 | SAMN08473241 | GCA_015813685.1 | NonPC | Public | 2011 | USA | Env | 90 | 100 | 0.08 | 98.1655 | 4994849 | 45.32 | 95 | 167194 |
| SAMN08473242-GCA_015813795.1 | CFSAN075818 | SAMN08473242 | GCA_015813795.1 | NonPC | Public | 2010 | USA | Env | 92.6 | 100 | 0 | 98.3783 | 5157775 | 45.34 | 61 | 481507 |
| SAMN08473243-GCA_015813325.1 | CFSAN075840 | SAMN08473243 | GCA_015813325.1 | NonPC | Public | 2010 | USA | Env | 92.2 | 100 | 0.58 | 98.259 | 5271768 | 45.21 | 51 | 575609 |
| SAMN08473244-GCA_015813065.1 | CFSAN075843 | SAMN08473244 | GCA_015813065.1 | NonPC | Public | 2010 | USA | Env | 92.2 | 100 | 0.49 | 98.2866 | 5224435 | 45.24 | 49 | 618662 |
| SAMN08473246-GCA_015814545.1 | CFSAN075821 | SAMN08473246 | GCA_015814545.1 | NonPC | Public | 2011 | USA | Env | 90.8 | 100 | 0.61 | 98.2462 | 5259724 | 45.07 | 44 | 360062 |
| SAMN08473247-GCA_016304305.1 | CFSAN075820 | SAMN08473247 | GCA_016304305.1 | NonPC | Public | 2011 | USA | Env | 90 | 100 | 0.08 | 98.1505 | 5000911 | 45.32 | 101 | 246571 |
| SAMN08473248-GCA_015814265.1 | CFSAN075833 | SAMN08473248 | GCA_015814265.1 | NonPC | Public | 2009 | USA | Env | 91 | 100 | 0.26 | 98.2456 | 5246071 | 45.24 | 44 | 285961 |
| SAMN08473249-GCA_015813545.1 | CFSAN075849 | SAMN08473249 | GCA_015813545.1 | NonPC | Public | 2011 | USA | Env | 90.8 | 100 | 0 | 98.2609 | 5101081 | 45.26 | 68 | 325645 |
| SAMN08473250-GCA_015814055.1 | CFSAN075850 | SAMN08473250 | GCA_015814055.1 | NonPC | Public | 2011 | USA | Env | 91.4 | 100 | 0 | 98.3894 | 5227489 | 45 | 93 | 313718 |
| SAMN08473251-GCA_015813665.1 | CFSAN075811 | SAMN08473251 | GCA_015813665.1 | NonPC | Public | 2010 | USA | Env | 88.4 | 100 | 0.72 | 98.2135 | 5336323 | 45.16 | 47 | 407588 |
| SAMN08473252-GCA_015813525.1 | CFSAN075848 | SAMN08473252 | GCA_015813525.1 | NonPC | Public | 2011 | USA | Env | 91.9 | 100 | 0.07 | 98.4576 | 5026147 | 45.45 | 47 | 770449 |
| SAMN08473253-GCA_015813305.1 | CFSAN075862 | SAMN08473253 | GCA_015813305.1 | NonPC | Public | 2011 | USA | Env | 90.7 | 100 | 0 | 98.3158 | 5237202 | 45.38 | 37 | 927767 |
| SAMN08473254-GCA_015813425.1 | CFSAN075847 | SAMN08473254 | GCA_015813425.1 | NonPC | Public | 2011 | USA | Env | 91.5 | 100 | 0.54 | 98.3769 | 5313742 | 45.37 | 69 | 249250 |
| SAMN08473255-GCA_015814105.1 | CFSAN075875 | SAMN08473255 | GCA_015814105.1 | NonPC | Public | 2012 | USA | Env | 90.4 | 100 | 0.08 | 98.3678 | 5042837 | 45.44 | 46 | 658473 |
| SAMN08473256-GCA_015814385.1 | CFSAN075803 | SAMN08473256 | GCA_015814385.1 | NonPC | Public | 2010 | USA | Env | 92.1 | 100 | 0.49 | 98.2882 | 5266624 | 45.16 | 56 | 605720 |
| SAMN08473257-GCA_015813465.1 | CFSAN075804 | SAMN08473257 | GCA_015813465.1 | NonPC | Public | 2010 | USA | Env | 92.2 | 100 | 0.49 | 98.2725 | 5273232 | 45.16 | 51 | 605710 |

| | | | | | | | | | | | | | | | | |
|---|---|---|---|---|---|---|---|---|---|---|---|---|---|---|---|---|
| SAMN08473258-GCA_015813185.1 | CFSAN075853 | SAMN08473258 | GCA_015813185.1 | NonPC | Public | 2010 | USA | Env | 89.1 | 100 | 0 | 98.2758 | 5137413 | 45.25 | 38 | 545025 |
| SAMN08473259-GCA_015814165.1 | CFSAN075825 | SAMN08473259 | GCA_015814165.1 | NonPC | Public | 2011 | USA | Env | 88.4 | 100 | 0.36 | 98.2185 | 5260725 | 45.11 | 38 | 445146 |
| SAMN08473260-GCA_015813645.1 | CFSAN075826 | SAMN08473260 | GCA_015813645.1 | NonPC | Public | 2011 | USA | Env | 87.8 | 100 | 0.41 | 98.2102 | 5343825 | 45.02 | 47 | 366192 |
| SAMN08473261-GCA_015813165.1 | CFSAN075827 | SAMN08473261 | GCA_015813165.1 | NonPC | Public | 2011 | USA | Env | 92.6 | 100 | 0.07 | 98.3966 | 5152301 | 45.36 | 61 | 481367 |
| SAMN08473262-GCA_015814285.1 | CFSAN075828 | SAMN08473262 | GCA_015814285.1 | NonPC | Public | 2012 | USA | Env | 92.1 | 100 | 0.54 | 98.2512 | 5257252 | 45.18 | 51 | 605564 |
| SAMN08473263-GCA_015813605.1 | CFSAN075829 | SAMN08473263 | GCA_015813605.1 | NonPC | Public | 2012 | USA | Env | 90.8 | 100 | 0.08 | 98.3106 | 5026885 | 45.29 | 59 | 517193 |
| SAMN08473264-GCA_015813505.1 | CFSAN075830 | SAMN08473264 | GCA_015813505.1 | NonPC | Public | 2012 | USA | Env | 92.1 | 100 | 0.49 | 98.2868 | 5223137 | 45.19 | 49 | 605686 |
| SAMN08473265-GCA_015813585.1 | CFSAN075852 | SAMN08473265 | GCA_015813585.1 | NonPC | Public | 2009 | USA | Env | 89.8 | 100 | 0.23 | 98.292 | 4995638 | 45.43 | 43 | 660422 |
| SAMN08473266-GCA_015813265.1 | CFSAN075800 | SAMN08473266 | GCA_015813265.1 | NonPC | Public | 2009 | USA | Env | 88.4 | 100 | 0.41 | 98.208 | 5314816 | 45.08 | 51 | 405404 |
| SAMN08473267-GCA_015814445.1 | CFSAN075799 | SAMN08473267 | GCA_015814445.1 | NonPC | Public | 2009 | USA | Env | 91.3 | 100 | 0.27 | 98.3339 | 5293990 | 45.01 | 109 | 290366 |
| SAMN08473268-GCA_015813085.1 | CFSAN075824 | SAMN08473268 | GCA_015813085.1 | NonPC | Public | 2011 | USA | Env | 90.7 | 100 | 0.28 | 98.3733 | 5121473 | 45.32 | 80 | 433721 |
| SAMN08473269-GCA_015813145.1 | CFSAN075844 | SAMN08473269 | GCA_015813145.1 | NonPC | Public | 2010 | USA | Env | 91.1 | 100 | 0 | 98.3971 | 4993056 | 45.42 | 50 | 641668 |
| SAMN08473270-GCA_015813485.1 | CFSAN075807 | SAMN08473270 | GCA_015813485.1 | NonPC | Public | 2010 | USA | Env | 88.4 | 100 | 0.36 | 98.237 | 5319948 | 45 | 50 | 366187 |
| SAMN08473271-GCA_015813005.1 | CFSAN075873 | SAMN08473271 | GCA_015813005.1 | NonPC | Public | 2012 | USA | Env | 90.7 | 100 | 0 | 98.4294 | 4986312 | 45.39 | 61 | 514405 |
| SAMN08473272-GCA_015813405.1 | CFSAN075845 | SAMN08473272 | GCA_015813405.1 | NonPC | Public | 2010 | USA | Env | 92.2 | 100 | 0.49 | 98.2289 | 5235360 | 45.2 | 43 | 605696 |
| SAMN08473273-GCA_015814425.1 | CFSAN075810 | SAMN08473273 | GCA_015814425.1 | NonPC | Public | 2010 | USA | Env | 90.7 | 100 | 0.39 | 98.323 | 5242188 | 45.19 | 54 | 443877 |
| SAMN08473274-GCA_015812325.1 | CFSAN075874 | SAMN08473274 | GCA_015812325.1 | NonPC | Public | 2012 | USA | Env | 88.6 | 100 | 0.03 | 98.1056 | 4915221 | 45.38 | 70 | 521764 |
| SAMN08473275-GCA_015813365.1 | CFSAN075846 | SAMN08473275 | GCA_015813365.1 | NonPC | Public | 2011 | USA | Env | 91.3 | 100 | 0 | 98.2414 | 5114616 | 45.25 | 73 | 560564 |
| SAMN08473276-GCA_015814245.1 | CFSAN075809 | SAMN08473276 | GCA_015814245.1 | NonPC | Public | 2010 | USA | Env | 88.4 | 100 | 0.36 | 98.2083 | 5274707 | 45.13 | 44 | 445146 |
| SAMN08473279-GCA_015814205.1 | CFSAN075878 | SAMN08473279 | GCA_015814205.1 | NonPC | Public | 2012 | USA | Env | 91.2 | 100 | 0.41 | 98.2575 | 5112217 | 45.24 | 74 | 252235 |
| SAMN08473280-GCA_015814125.1 | CFSAN075879 | SAMN08473280 | GCA_015814125.1 | NonPC | Public | 2012 | USA | Env | 90.7 | 100 | 0.05 | 98.3298 | 5016046 | 45.43 | 38 | 806511 |
| SAMN08473282-GCA_015814225.1 | CFSAN075854 | SAMN08473282 | GCA_015814225.1 | NonPC | Public | 2010 | USA | Env | 89.2 | 100 | 0 | 98.271 | 5137974 | 45.25 | 37 | 545024 |
| SAMN08473283-GCA_015813905.1 | CFSAN075861 | SAMN08473283 | GCA_015813905.1 | NonPC | Public | 2011 | USA | Env | 90.9 | 100 | 0 | 98.3741 | 4952652 | 45.47 | 32 | 1226893 |
| SAMN08473286-GCA_015814365.1 | CFSAN075871 | SAMN08473286 | GCA_015814365.1 | NonPC | Public | 2012 | USA | Env | 91.1 | 100 | 0.08 | 98.382 | 5054236 | 45.44 | 42 | 694067 |
| SAMN08473287-GCA_015812245.1 | CFSAN075835 | SAMN08473287 | GCA_015812245.1 | NonPC | Public | 2010 | USA | Env | 91.5 | 100 | 0.35 | 98.3361 | 5216294 | 45.1 | 28 | 545566 |
| SAMN08473289-GCA_015814065.1 | CFSAN075832 | SAMN08473289 | GCA_015814065.1 | NonPC | Public | 2012 | USA | Env | 88.4 | 100 | 0.36 | 98.2108 | 5256423 | 45.13 | 41 | 383275 |
| SAMN08473293-GCA_015813375.1 | CFSAN075837 | SAMN08473293 | GCA_015813375.1 | NonPC | Public | 2011 | USA | Env | 88.7 | 100 | 0 | 98.1749 | 4929717 | 45.4 | 59 | 522463 |
| SAMN08473295-GCA_015813315.1 | CFSAN075838 | SAMN08473295 | GCA_015813315.1 | NonPC | Public | 2012 | USA | Env | 91.9 | 100 | 0 | 98.3011 | 5093802 | 45.27 | 59 | 316626 |
| SAMN08473296-GCA_015814045.1 | CFSAN075836 | SAMN08473296 | GCA_015814045.1 | NonPC | Public | 2010 | USA | Env | 92.2 | 100 | 0.35 | 98.3301 | 5216195 | 45.1 | 25 | 568824 |
| SAMN08473298-GCA_015814305.1 | CFSAN075868 | SAMN08473298 | GCA_015814305.1 | NonPC | Public | 2012 | USA | Env | 90.9 | 100 | 0.03 | 98.2757 | 5158928 | 45.18 | 26 | 1184365 |
| SAMN08473299-GCA_015813765.1 | CFSAN075812 | SAMN08473299 | GCA_015813765.1 | NonPC | Public | 2010 | USA | Env | 88.4 | 100 | 0.41 | 98.1732 | 5305839 | 45.08 | 59 | 279303 |
| SAMN08473300-SRR6728015 | CFSAN075863 | SAMN08473300 | SRR6728015 | NonPC | Public | 2011 | USA | Env | 89 | 100 | 0 | 98.3166 | 5027642 | 45.39 | 28 | 561589 |
| SAMN08473301-GCA_015814025.1 | CFSAN075857 | SAMN08473301 | GCA_015814025.1 | NonPC | Public | 2010 | USA | Env | 90.6 | 100 | 0.63 | 98.2778 | 5172717 | 45.17 | 54 | 401479 |
| SAMN08473302-GCA_015814145.1 | CFSAN075856 | SAMN08473302 | GCA_015814145.1 | NonPC | Public | 2010 | USA | Env | 89.2 | 100 | 0 | 98.2369 | 5140602 | 45.24 | 35 | 545026 |
| SAMN08473303-GCA_015813285.1 | CFSAN075858 | SAMN08473303 | GCA_015813285.1 | NonPC | Public | 2010 | USA | Env | 92.3 | 100 | 0.35 | 98.3057 | 5235167 | 45.13 | 27 | 568794 |
| SAMN08473304-GCA_015813225.1 | CFSAN075859 | SAMN08473304 | GCA_015813225.1 | NonPC | Public | 2010 | USA | Env | 89.8 | 100 | 0.14 | 98.3438 | 5155515 | 45.31 | 62 | 482247 |
| SAMN08473305-GCA_015813565.1 | CFSAN075831 | SAMN08473305 | GCA_015813565.1 | NonPC | Public | 2012 | USA | Env | 88.4 | 100 | 0.36 | 98.2206 | 5309858 | 45.08 | 44 | 405404 |
| SAMN08473306-GCA_015814185.1 | CFSAN075851 | SAMN08473306 | GCA_015814185.1 | NonPC | Public | 2012 | USA | Env | 92.5 | 100 | 0.26 | 98.3214 | 5135015 | 45.39 | 34 | 783635 |
| SAMN08473307-GCA_015812985.1 | CFSAN075802 | SAMN08473307 | GCA_015812985.1 | NonPC | Public | 2010 | USA | Env | 90 | 100 | 0.05 | 98.2375 | 4980821 | 45.38 | 68 | 402139 |
| SAMN08473308-GCA_015813625.1 | CFSAN075806 | SAMN08473308 | GCA_015813625.1 | NonPC | Public | 2010 | USA | Env | 88.4 | 100 | 0.41 | 98.1763 | 5338643 | 45.02 | 57 | 407582 |
| SAMN08473309-GCA_015813445.1 | CFSAN075805 | SAMN08473309 | GCA_015813445.1 | NonPC | Public | 2010 | USA | Env | 88.4 | 100 | 0.41 | 98.1708 | 5340927 | 45.01 | 48 | 445146 |
| SAMN08473310-GCA_015813205.1 | CFSAN075842 | SAMN08473310 | GCA_015813205.1 | NonPC | Public | 2010 | USA | Env | 90.9 | 100 | 0.5 | 98.2395 | 5249949 | 45.2 | 70 | 489129 |
| SAMN08473311-GCA_015813955.1 | CFSAN075877 | SAMN08473311 | GCA_015813955.1 | NonPC | Public | 2012 | USA | Env | 91 | 100 | 0.37 | 98.2962 | 4977008 | 45.37 | 37 | 549454 |
| SAMN08473312-GCA_015813985.1 | CFSAN075876 | SAMN08473312 | GCA_015813985.1 | NonPC | Public | 2012 | USA | Env | 91.1 | 100 | 0.05 | 98.4018 | 5048183 | 45.22 | 50 | 578469 |
| SAMN08568263-GCA_015810605.1 | CFSAN075748 | SAMN08568263 | GCA_015810605.1 | NonPC | Public | 2012 | USA | Env | 88.6 | 100 | 0.39 | 98.0949 | 5116267 | 45.23 | 56 | 250588 |
| SAMN08627207-GCA_015812285.1 | PNUSAV000174 | SAMN08627207 | GCA_015812285.1 | PC | Public | NA | USA | Clinical | 98.4 | 100 | 0 | 99.988 | 5092352 | 45.29 | 60 | 229941 |
| SAMN08627210-GCA_015812265.1 | PNUSAV000173 | SAMN08627210 | GCA_015812265.1 | PC | Public | NA | USA | Clinical | 98.1 | 100 | 0 | 99.9865 | 5079477 | 45.32 | 113 | 94122 |
| SAMN08667565-GCF_003056675.1 | VPF-1 | SAMN08667565 | GCF_003056675.1 | PC | Public | 2017 | Lebanon | Clinical | 98.5 | 100 | 0 | 99.9909 | 5095004 | 45.29 | 42 | 427776 |

| | | | | | | | | | | | | | | | | |
|---|---|---|---|---|---|---|---|---|---|---|---|---|---|---|---|---|
| SAMN08667566-GCF_003057295.1 | VPF-2 | SAMN08667566 | GCF_003057295.1 | PC | Public | 2017 | Lebanon | Clinical | 98.5 | 100 | 0.34 | 99.9886 | 5098903 | 45.29 | 51 | 433180 |
| SAMN08667567-GCF_003056625.1 | VPF-3 | SAMN08667567 | GCF_003056625.1 | PC | Public | 2017 | Lebanon | Clinical | 98.6 | 100 | 0.12 | 99.9871 | 5138349 | 45.28 | 44 | 401761 |
| SAMN08667585-GCF_003056645.1 | VPF-4 | SAMN08667585 | GCF_003056645.1 | PC | Public | 2017 | Lebanon | Clinical | 98.6 | 100 | 0.13 | 99.9869 | 5099120 | 45.29 | 53 | 402137 |
| SAMN08667654-GCF_003057315.1 | VPF-5 | SAMN08667654 | GCF_003057315.1 | PC | Public | 2017 | Lebanon | Clinical | 98.6 | 100 | 0 | 99.9866 | 5099283 | 45.3 | 44 | 433180 |
| SAMN08667664-GCF_003056665.1 | VPF-6 | SAMN08667664 | GCF_003056665.1 | PC | Public | 2017 | Lebanon | Clinical | 98.6 | 100 | 0.16 | 99.9861 | 5138805 | 45.28 | 42 | 433354 |
| SAMN08667669-GCF_003056745.1 | VPF-7 | SAMN08667669 | GCF_003056745.1 | PC | Public | 2017 | Lebanon | Clinical | 98.6 | 100 | 0 | 99.9857 | 5099741 | 45.29 | 33 | 433258 |
| SAMN08744595-GCF_003076895.1 | APL-R14 | SAMN08744595 | GCF_003076895.1 | NonPC | Public | 2016 | NA | Env | 92.3 | 100 | 0.18 | 98.2429 | 5444136 | 45.27 | 5 | 3477001 |
| SAMN08774111-GCF_003119375.1 | APL-R13 | SAMN08774111 | GCF_003119375.1 | NonPC | Public | 2016 | NA | Env | 92.4 | 100 | 0.18 | 98.2494 | 5441285 | 45.27 | 5 | 3477006 |
| SAMN08851398-GCF_003047085.1 | S107-1 | SAMN08851398 | GCF_003047085.1 | NonPC | Public | NA | Canada | Env | 92.5 | 99.54 | 0.09 | 98.2993 | 5203326 | 45.39 | 2 | 3445421 |
| SAMN08902697-GCA_015809895.1 | CFSAN079457 | SAMN08902697 | GCA_015809895.1 | NonPC | Public | 2012 | USA | Env | 90.3 | 100 | 0.2 | 98.267 | 5156584 | 45.21 | 36 | 371852 |
| SAMN08902704-GCA_015809375.1 | CFSAN079454 | SAMN08902704 | GCA_015809375.1 | NonPC | Public | 2012 | USA | Env | 91 | 100 | 0.2 | 98.2633 | 5168837 | 45.2 | 42 | 371852 |
| SAMN08902705-GCA_015808395.1 | CFSAN079453 | SAMN08902705 | GCA_015808395.1 | NonPC | Public | 2012 | USA | Env | 91 | 100 | 0.2 | 98.2678 | 5169804 | 45.2 | 40 | 312111 |
| SAMN08902706-GCA_015808475.1 | CFSAN079456 | SAMN08902706 | GCA_015808475.1 | NonPC | Public | 2012 | USA | Env | 91 | 100 | 0 | 98.2516 | 5195496 | 45.2 | 45 | 443057 |
| SAMN08902707-GCA_015810665.1 | CFSAN079455 | SAMN08902707 | GCA_015810665.1 | PrePC | Public | 2012 | USA | Env | 94.8 | 100 | 0.35 | 99.8156 | 5095882 | 45.3 | 62 | 214727 |
| SAMN08902716-GCA_015810365.1 | CFSAN079448 | SAMN08902716 | GCA_015810365.1 | NonPC | Public | 2012 | USA | Env | 92.2 | 100 | 0.01 | 98.3518 | 5104355 | 45.53 | 58 | 331373 |
| SAMN08902719-GCA_015811385.1 | CFSAN079467 | SAMN08902719 | GCA_015811385.1 | NonPC | Public | 2012 | USA | Env | 90.6 | 100 | 0.82 | 98.2619 | 5277069 | 45.39 | 44 | 1087261 |
| SAMN08902720-GCA_015811605.1 | CFSAN079458 | SAMN08902720 | GCA_015811605.1 | NonPC | Public | 2012 | USA | Env | 90.3 | 100 | 0.2 | 98.304 | 5156015 | 45.21 | 41 | 324923 |
| SAMN08902723-GCA_015811685.1 | CFSAN079461 | SAMN08902723 | GCA_015811685.1 | NonPC | Public | 2012 | USA | Env | 91.5 | 100 | 0 | 98.2414 | 5099891 | 45.29 | 45 | 239382 |
| SAMN08902724-GCA_015811585.1 | CFSAN079462 | SAMN08902724 | GCA_015811585.1 | NonPC | Public | 2012 | USA | Env | 91.5 | 100 | 0 | 98.2406 | 5103878 | 45.29 | 41 | 381450 |
| SAMN08902725-GCA_015812725.1 | CFSAN079463 | SAMN08902725 | GCA_015812725.1 | NonPC | Public | 2012 | USA | Env | 91.1 | 100 | 0.41 | 98.3374 | 5252313 | 45.25 | 48 | 368434 |
| SAMN08902726-GCA_015811665.1 | CFSAN079464 | SAMN08902726 | GCA_015811665.1 | NonPC | Public | 2012 | USA | Env | 89.8 | 100 | 0.08 | 98.1643 | 5066996 | 45.27 | 100 | 162653 |
| SAMN08902729-GCA_015811345.1 | CFSAN079466 | SAMN08902729 | GCA_015811345.1 | NonPC | Public | 2012 | USA | Env | 91.1 | 100 | 0.41 | 98.3027 | 5257664 | 45.24 | 53 | 346956 |
| SAMN08902733-GCA_015810515.1 | CFSAN079433 | SAMN08902733 | GCA_015810515.1 | NonPC | Public | 2012 | USA | Env | 92.2 | 100 | 0.01 | 98.3813 | 5190099 | 45.47 | 65 | 364994 |
| SAMN08902738-GCA_015810765.1 | CFSAN079437 | SAMN08902738 | GCA_015810765.1 | PrePC | Public | 2012 | USA | Env | 94.8 | 100 | 0.35 | 99.794 | 5106245 | 45.3 | 59 | 231554 |
| SAMN08902754-GCA_015810805.1 | CFSAN079444 | SAMN08902754 | GCA_015810805.1 | NonPC | Public | 2012 | USA | Env | 91.7 | 100 | 0.08 | 98.3708 | 5150072 | 45.26 | 56 | 302348 |
| SAMN08902764-GCA_015809995.1 | CFSAN079440 | SAMN08902764 | GCA_015809995.1 | NonPC | Public | 2012 | USA | Env | 91.5 | 100 | 0 | 98.3392 | 5085028 | 45.29 | 37 | 418676 |
| SAMN08902766-GCA_015809485.1 | CFSAN079447 | SAMN08902766 | GCA_015809485.1 | NonPC | Public | 2012 | USA | Env | 91 | 100 | 2.86 | 98.2813 | 5327107 | 45.26 | 63 | 279609 |
| SAMN08902768-GCA_015808415.1 | CFSAN079446 | SAMN08902768 | GCA_015808415.1 | NonPC | Public | 2012 | USA | Env | 91.5 | 100 | 0.01 | 98.2986 | 5189795 | 45.28 | 47 | 536446 |
| SAMN08902770-GCA_015808355.1 | CFSAN079445 | SAMN08902770 | GCA_015808355.1 | NonPC | Public | 2012 | USA | Env | 91.5 | 100 | 0.01 | 98.2895 | 5192927 | 45.28 | 57 | 321951 |
| SAMN08902775-GCA_015810545.1 | CFSAN079443 | SAMN08902775 | GCA_015810545.1 | PrePC | Public | 2012 | USA | Env | 94.9 | 100 | 0.35 | 99.8105 | 5100080 | 45.29 | 55 | 277048 |
| SAMN08902776-GCA_015810235.1 | CFSAN079438 | SAMN08902776 | GCA_015810235.1 | NonPC | Public | 2012 | USA | Env | 91 | 100 | 2.86 | 98.3302 | 5321108 | 45.27 | 72 | 208608 |
| SAMN08929979-GCA_015811735.1 | PNUSAV000177 | SAMN08929979 | GCA_015811735.1 | NonPC | Public | NA | USA | Clinical | 91.5 | 100 | 0 | 98.3074 | 5100666 | 45.27 | 61 | 234484 |
| SAMN08965100-GCA_015808955.1 | CFSAN080170 | SAMN08965100 | GCA_015808955.1 | NonPC | Public | 2013 | USA | Env | 91 | 100 | 0 | 98.2187 | 5319869 | 45.18 | 98 | 180522 |
| SAMN08965101-GCA_015808545.1 | CFSAN080135 | SAMN08965101 | GCA_015808545.1 | NonPC | Public | 2013 | USA | Env | 90.1 | 100 | 0.19 | 98.3162 | 5207868 | 45.31 | 93 | 196737 |
| SAMN08965102-GCA_015808935.1 | CFSAN080169 | SAMN08965102 | GCA_015808935.1 | NonPC | Public | 2013 | USA | Env | 91.5 | 100 | 0.22 | 98.3961 | 5151627 | 45.27 | 80 | 134753 |
| SAMN08965103-GCA_015808975.1 | CFSAN080168 | SAMN08965103 | GCA_015808975.1 | NonPC | Public | 2013 | USA | Env | 91.5 | 100 | 1.17 | 98.3222 | 5178111 | 45.28 | 86 | 126898 |
| SAMN08965104-GCA_015807855.1 | CFSAN080129 | SAMN08965104 | GCA_015807855.1 | NonPC | Public | 2013 | USA | Env | 90.1 | 100 | 0.05 | 98.3473 | 5123691 | 45.33 | 117 | 156725 |
| SAMN08965105-GCA_015808815.1 | CFSAN080130 | SAMN08965105 | GCA_015808815.1 | NonPC | Public | 2013 | USA | Env | 88.6 | 100 | 0.35 | 98.1408 | 5018063 | 45.32 | 64 | 233545 |
| SAMN08965107-GCA_015805365.1 | CFSAN080177 | SAMN08965107 | GCA_015805365.1 | NonPC | Public | 2013 | USA | Env | 88.4 | 100 | 1.17 | 98.1791 | 5057283 | 45.38 | 124 | 81153 |
| SAMN08965108-GCA_015807745.1 | CFSAN080128 | SAMN08965108 | GCA_015807745.1 | NonPC | Public | 2013 | USA | Env | 90.9 | 100 | 0.2 | 98.2876 | 5139502 | 45.26 | 115 | 92031 |
| SAMN08965109-GCA_015808835.1 | CFSAN080174 | SAMN08965109 | GCA_015808835.1 | NonPC | Public | 2013 | USA | Env | 92.1 | 100 | 0.61 | 98.3738 | 5541277 | 44.9 | 162 | 91468 |
| SAMN08965110-GCA_015808535.1 | CFSAN080172 | SAMN08965110 | GCA_015808535.1 | NonPC | Public | 2013 | USA | Env | 89.5 | 100 | 0.33 | 98.2777 | 5189800 | 45.29 | 117 | 94463 |
| SAMN08965111-GCA_015809175.1 | CFSAN080132 | SAMN08965111 | GCA_015809175.1 | NonPC | Public | 2013 | USA | Env | 92 | 100 | 0.53 | 98.4597 | 5035717 | 45.31 | 99 | 123179 |
| SAMN08965112-GCA_015808715.1 | CFSAN080175 | SAMN08965112 | GCA_015808715.1 | NonPC | Public | 2013 | USA | Env | 88.5 | 100 | 1.09 | 98.1334 | 5076725 | 45.24 | 96 | 128872 |
| SAMN08965113-GCA_015807615.1 | CFSAN080134 | SAMN08965113 | GCA_015807615.1 | NonPC | Public | 2013 | USA | Env | 91.7 | 100 | 0.08 | 98.3588 | 5141256 | 45.29 | 70 | 198893 |
| SAMN08965114-GCA_015807375.1 | CFSAN080131 | SAMN08965114 | GCA_015807375.1 | NonPC | Public | 2013 | USA | Env | 88.6 | 100 | 0.35 | 98.1573 | 5103417 | 45.25 | 85 | 114777 |
| SAMN08965115-GCA_015808455.1 | CFSAN080176 | SAMN08965115 | GCA_015808455.1 | NonPC | Public | 2013 | USA | Env | 92.2 | 100 | 0.61 | 98.3216 | 5415881 | 45.02 | 116 | 145640 |
| SAMN08965121-GCA_015805325.1 | CFSAN080187 | SAMN08965121 | GCA_015805325.1 | NonPC | Public | 2013 | USA | Env | 89.6 | 100 | 4.31 | 98.2552 | 5392777 | 45.28 | 76 | 149598 |

| | | | | | | | | | | | | | | | | |
|---|---|---|---|---|---|---|---|---|---|---|---|---|---|---|---|---|
| SAMN08965123-GCA_015809155.1 | CFSAN080137 | SAMN08965123 | GCA_015809155.1 | NonPC | Public | 2013 | USA | Env | 88.9 | 99.73 | 0.14 | 98.1908 | 4948716 | 45.31 | 47 | 262951 |
| SAMN08965124-GCA_015808995.1 | CFSAN080133 | SAMN08965124 | GCA_015808995.1 | NonPC | Public | 2013 | USA | Env | 89.7 | 100 | 3.32 | 98.3364 | 5286208 | 45.24 | 73 | 163052 |
| SAMN08965125-GCA_015808695.1 | CFSAN080171 | SAMN08965125 | GCA_015808695.1 | NonPC | Public | 2013 | USA | Env | 90.8 | 100 | 0.89 | 98.2317 | 5161967 | 45.22 | 77 | 169782 |
| SAMN08965126-GCA_015809215.1 | CFSAN080111 | SAMN08965126 | GCA_015809215.1 | NonPC | Public | 2013 | USA | Env | 91.7 | 100 | 0.08 | 98.4003 | 5144469 | 45.27 | 62 | 232243 |
| SAMN08965127-GCA_015809135.1 | CFSAN080115 | SAMN08965127 | GCA_015809135.1 | NonPC | Public | 2013 | USA | Env | 91.2 | 100 | 0 | 98.3092 | 5190240 | 45.25 | 107 | 104898 |
| SAMN08965128-GCA_015809235.1 | CFSAN080152 | SAMN08965128 | GCA_015809235.1 | NonPC | Public | 2013 | USA | Env | 91.1 | 100 | 0.41 | 98.3005 | 5254895 | 45.24 | 60 | 293581 |
| SAMN08965129-GCA_015808875.1 | CFSAN080153 | SAMN08965129 | GCA_015808875.1 | NonPC | Public | 2013 | USA | Env | 89.2 | 100 | 0 | 98.0852 | 5129372 | 45.29 | 80 | 230411 |
| SAMN08965130-GCA_015807055.1 | CFSAN080190 | SAMN08965130 | GCA_015807055.1 | NonPC | Public | 2013 | USA | Env | 90.8 | 100 | 0.01 | 98.2671 | 5307572 | 45.28 | 152 | 77554 |
| SAMN08965131-GCA_015808575.1 | CFSAN080151 | SAMN08965131 | GCA_015808575.1 | NonPC | Public | 2013 | USA | Env | 91.4 | 100 | 0.08 | 98.2325 | 5065751 | 45.3 | 75 | 184807 |
| SAMN08965132-GCA_015807915.1 | CFSAN080113 | SAMN08965132 | GCA_015807915.1 | NonPC | Public | 2013 | USA | Env | 91.7 | 100 | 0.08 | 98.3577 | 5149945 | 45.27 | 50 | 402762 |
| SAMN08965133-GCA_015807035.1 | CFSAN080189 | SAMN08965133 | GCA_015807035.1 | NonPC | Public | 2013 | USA | Env | 89.9 | 100 | 1.02 | 98.2405 | 5281190 | 45.2 | 99 | 88648 |
| SAMN08965134-GCA_015807815.1 | CFSAN080116 | SAMN08965134 | GCA_015807815.1 | NonPC | Public | 2013 | USA | Env | 91.3 | 99.93 | 0 | 98.2978 | 5130533 | 45.33 | 47 | 320888 |
| SAMN08965136-GCA_015809095.1 | CFSAN080117 | SAMN08965136 | GCA_015809095.1 | NonPC | Public | 2013 | USA | Env | 90.4 | 100 | 0.2 | 98.2949 | 5158485 | 45.21 | 45 | 250100 |
| SAMN08965138-GCA_015808515.1 | CFSAN080156 | SAMN08965138 | GCA_015808515.1 | NonPC | Public | 2013 | USA | Env | 91.8 | 100 | 0.54 | 98.2617 | 5042907 | 45.39 | 49 | 542574 |
| SAMN08965139-GCA_015806735.1 | CFSAN080193 | SAMN08965139 | GCA_015806735.1 | NonPC | Public | 2013 | USA | Env | 92.1 | 100 | 1.55 | 98.3965 | 5180320 | 45.3 | 83 | 140312 |
| SAMN08965140-GCA_015807555.1 | CFSAN080155 | SAMN08965140 | GCA_015807555.1 | PrePC | Public | 2013 | USA | Env | 94.8 | 100 | 0.35 | 99.8321 | 5099200 | 45.28 | 54 | 189108 |
| SAMN08965141-GCA_015805255.1 | CFSAN080197 | SAMN08965141 | GCA_015805255.1 | NonPC | Public | 2013 | USA | Env | 89.6 | 100 | 0.83 | 98.2788 | 5178357 | 45.28 | 104 | 100839 |
| SAMN08965144-GCA_015808895.1 | CFSAN080157 | SAMN08965144 | GCA_015808895.1 | NonPC | Public | 2013 | USA | Env | 90.3 | 100 | 0.2 | 98.2726 | 5164550 | 45.21 | 70 | 180438 |
| SAMN08965146-GCA_015805215.1 | CFSAN080188 | SAMN08965146 | GCA_015805215.1 | NonPC | Public | 2013 | USA | Env | 89.2 | 100 | 1.2 | 98.2036 | 5098299 | 45.5 | 110 | 106387 |
| SAMN08965147-GCA_015807635.1 | CFSAN080154 | SAMN08965147 | GCA_015807635.1 | NonPC | Public | 2013 | USA | Env | 91.7 | 100 | 0.08 | 98.3748 | 5150310 | 45.26 | 50 | 236220 |
| SAMN08965149-GCA_015808675.1 | CFSAN080105 | SAMN08965149 | GCA_015808675.1 | NonPC | Public | 2013 | USA | Env | 92.7 | 100 | 0 | 98.3613 | 5284420 | 45.22 | 135 | 142716 |
| SAMN08965151-GCA_015809015.1 | CFSAN080106 | SAMN08965151 | GCA_015809015.1 | NonPC | Public | 2013 | USA | Env | 92.7 | 100 | 0 | 98.3388 | 5273279 | 45.23 | 128 | 130966 |
| SAMN08965152-GCA_015808615.1 | CFSAN080142 | SAMN08965152 | GCA_015808615.1 | NonPC | Public | 2013 | USA | Env | 89.9 | 100 | 0.04 | 98.2637 | 5224779 | 45.19 | 46 | 397629 |
| SAMN08965153-GCA_015807895.1 | CFSAN080107 | SAMN08965153 | GCA_015807895.1 | NonPC | Public | 2013 | USA | Env | 91.6 | 100 | 0.08 | 98.3706 | 5133207 | 45.31 | 82 | 145057 |
| SAMN08965154-GCA_015809035.1 | CFSAN080145 | SAMN08965154 | GCA_015809035.1 | NonPC | Public | 2013 | USA | Env | 89 | 100 | 0.14 | 98.1151 | 5022884 | 45.27 | 43 | 313599 |
| SAMN08965156-GCA_015806955.1 | CFSAN080178 | SAMN08965156 | GCA_015806955.1 | NonPC | Public | 2013 | USA | Env | 88.3 | 100 | 0.43 | 98.1913 | 5040927 | 45.3 | 146 | 66369 |
| SAMN08965157-GCA_015809075.1 | CFSAN080149 | SAMN08965157 | GCA_015809075.1 | NonPC | Public | 2013 | USA | Env | 90.1 | 100 | 0.22 | 98.2054 | 5044501 | 45.35 | 86 | 170559 |
| SAMN08965158-GCA_015808595.1 | CFSAN080150 | SAMN08965158 | GCA_015808595.1 | NonPC | Public | 2013 | USA | Env | 91.6 | 100 | 0.08 | 98.364 | 5137080 | 45.29 | 74 | 178120 |
| SAMN08965159-GCA_015806755.1 | CFSAN080180 | SAMN08965159 | GCA_015806755.1 | NonPC | Public | 2013 | USA | Env | 90 | 100 | 3.87 | 98.2178 | 5389922 | 45.1 | 135 | 95303 |
| SAMN08965160-GCA_015808755.1 | CFSAN080146 | SAMN08965160 | GCA_015808755.1 | NonPC | Public | 2013 | USA | Env | 89 | 100 | 0.14 | 98.1287 | 5023503 | 45.27 | 41 | 269251 |
| SAMN08965161-GCA_015806855.1 | CFSAN080181 | SAMN08965161 | GCA_015806855.1 | NonPC | Public | 2013 | USA | Env | 90.9 | 100 | 1.89 | 98.3333 | 5352088 | 45.21 | 106 | 105033 |
| SAMN08965162-GCA_015809275.1 | CFSAN080147 | SAMN08965162 | GCA_015809275.1 | NonPC | Public | 2013 | USA | Env | 92.2 | 100 | 0.27 | 98.3769 | 5322695 | 45.01 | 86 | 275952 |
| SAMN08965164-GCA_015805415.1 | CFSAN080179 | SAMN08965164 | GCA_015805415.1 | NonPC | Public | 2013 | USA | Env | 91.3 | 100 | 0.9 | 98.3417 | 5175788 | 45.36 | 109 | 88480 |
| SAMN08965165-GCA_015807695.1 | CFSAN080141 | SAMN08965165 | GCA_015807695.1 | NonPC | Public | 2013 | USA | Env | 89.9 | 100 | 0.04 | 98.2608 | 5225027 | 45.19 | 57 | 241812 |
| SAMN08965166-GCA_015807315.1 | CFSAN080196 | SAMN08965166 | GCA_015807315.1 | NonPC | Public | 2013 | USA | Env | 91.3 | 100 | 2.09 | 98.3331 | 5337486 | 45.34 | 164 | 68097 |
| SAMN08965167-GCA_015808795.1 | CFSAN080148 | SAMN08965167 | GCA_015808795.1 | NonPC | Public | 2013 | USA | Env | 88.6 | 100 | 0.35 | 98.1584 | 5028643 | 45.2 | 67 | 180115 |
| SAMN08965169-GCA_015807355.1 | CFSAN080162 | SAMN08965169 | GCA_015807355.1 | NonPC | Public | 2013 | USA | Env | 91 | 100 | 0.2 | 98.2813 | 5140095 | 45.22 | 62 | 193990 |
| SAMN08965173-GCA_015808635.1 | CFSAN080166 | SAMN08965173 | GCA_015808635.1 | NonPC | Public | 2013 | USA | Env | 89.7 | 100 | 1.4 | 98.2396 | 5200606 | 45.25 | 75 | 122587 |
| SAMN08965175-GCA_015808655.1 | CFSAN080163 | SAMN08965175 | GCA_015808655.1 | NonPC | Public | 2013 | USA | Env | 89 | 100 | 0.24 | 98.1205 | 5420876 | 44.92 | 121 | 185050 |
| SAMN08965176-GCA_015807765.1 | CFSAN080109 | SAMN08965176 | GCA_015807765.1 | NonPC | Public | 2013 | USA | Env | 91.1 | 100 | 0.08 | 98.3425 | 5341735 | 45.09 | 110 | 171182 |
| SAMN08965177-GCA_015807475.1 | CFSAN080110 | SAMN08965177 | GCA_015807475.1 | NonPC | Public | 2013 | USA | Env | 91.1 | 100 | 0.08 | 98.3195 | 5330597 | 45.11 | 126 | 149916 |
| SAMN08965178-GCA_015809115.1 | CFSAN080108 | SAMN08965178 | GCA_015809115.1 | NonPC | Public | 2013 | USA | Env | 91.6 | 100 | 0.08 | 98.3506 | 5140117 | 45.29 | 95 | 122132 |
| SAMN08965179-GCA_015807455.1 | CFSAN080158 | SAMN08965179 | GCA_015807455.1 | NonPC | Public | 2013 | USA | Env | 91 | 100 | 0.2 | 98.2528 | 5172619 | 45.21 | 55 | 199290 |
| SAMN08965180-GCA_015809055.1 | CFSAN080159 | SAMN08965180 | GCA_015809055.1 | NonPC | Public | 2013 | USA | Env | 91.5 | 100 | 0.35 | 98.2704 | 5123259 | 45.31 | 77 | 194515 |
| SAMN08965181-GCA_015807435.1 | CFSAN080160 | SAMN08965181 | GCA_015807435.1 | NonPC | Public | 2013 | USA | Env | 90.3 | 100 | 0.08 | 98.2773 | 5042721 | 45.44 | 72 | 179289 |
| SAMN08965182-GCA_015807535.1 | CFSAN080161 | SAMN08965182 | GCA_015807535.1 | NonPC | Public | 2013 | USA | Env | 91 | 100 | 0.2 | 98.3004 | 5144265 | 45.23 | 74 | 162970 |
| SAMN08965183-GCA_015808735.1 | CFSAN080167 | SAMN08965183 | GCA_015808735.1 | NonPC | Public | 2013 | USA | Env | 89.7 | 100 | 0.36 | 98.2454 | 5163696 | 45.23 | 78 | 123123 |
| SAMN08965184-GCA_015807755.1 | CFSAN080121 | SAMN08965184 | GCA_015807755.1 | NonPC | Public | 2013 | USA | Env | 89.8 | 100 | 0.08 | 98.1406 | 5138869 | 45.2 | 118 | 141359 |

| | | | | | | | | | | | | | | | | |
|---|---|---|---|---|---|---|---|---|---|---|---|---|---|---|---|---|
| SAMN08965185-GCA_015807995.1 | CFSAN080119 | SAMN08965185 | GCA_015807995.1 | NonPC | Public | 2013 | USA | Env | 89.9 | 100 | 0.04 | 98.2718 | 5224673 | 45.19 | 45 | 356183 |
| SAMN08965186-GCA_015807335.1 | CFSAN080120 | SAMN08965186 | GCA_015807335.1 | NonPC | Public | 2013 | USA | Env | 89.9 | 100 | 0.04 | 98.222 | 5216208 | 45.21 | 53 | 184575 |
| SAMN08965187-GCA_015809195.1 | CFSAN080125 | SAMN08965187 | GCA_015809195.1 | NonPC | Public | 2013 | USA | Env | 89.8 | 100 | 0.22 | 98.2348 | 5153071 | 45.23 | 43 | 206520 |
| SAMN08965188-GCA_015809335.1 | CFSAN080118 | SAMN08965188 | GCA_015809335.1 | NonPC | Public | 2013 | USA | Env | 91 | 100 | 0.2 | 98.2862 | 5158707 | 45.2 | 38 | 371852 |
| SAMN08965189-GCA_015809355.1 | CFSAN080112 | SAMN08965189 | GCA_015809355.1 | NonPC | Public | 2013 | USA | Env | 91.7 | 100 | 0.08 | 98.3508 | 5139502 | 45.29 | 64 | 197171 |
| SAMN08965190-GCA_015807835.1 | CFSAN080124 | SAMN08965190 | GCA_015807835.1 | NonPC | Public | 2013 | USA | Env | 89.8 | 100 | 0.22 | 98.2795 | 5145867 | 45.25 | 80 | 122393 |
| SAMN08965191-GCA_015807935.1 | CFSAN080123 | SAMN08965191 | GCA_015807935.1 | NonPC | Public | 2013 | USA | Env | 88.9 | 100 | 0.14 | 98.1321 | 4966764 | 45.31 | 72 | 180927 |
| SAMN08965192-GCA_015807875.1 | CFSAN080122 | SAMN08965192 | GCA_015807875.1 | NonPC | Public | 2013 | USA | Env | 91.7 | 100 | 0.05 | 98.3487 | 5329457 | 45.14 | 122 | 170813 |
| SAMN08965194-GCA_015808915.1 | CFSAN080165 | SAMN08965194 | GCA_015808915.1 | NonPC | Public | 2013 | USA | Env | 91.5 | 100 | 0.69 | 98.3446 | 5162878 | 45.29 | 90 | 108848 |
| SAMN09074393-GCA_015808215.1 | CFSAN079474 | SAMN09074393 | GCA_015808215.1 | NonPC | Public | 2013 | USA | Env | 91.7 | 100 | 0.08 | 98.364 | 5240067 | 45.18 | 59 | 211955 |
| SAMN09074605-GCA_015809315.1 | CFSAN079429 | SAMN09074605 | GCA_015809315.1 | NonPC | Public | 2012 | USA | Env | 88.6 | 100 | 0.35 | 98.1971 | 4938757 | 45.37 | 51 | 303798 |
| SAMN09074606-GCA_015808235.1 | CFSAN079431 | SAMN09074606 | GCA_015808235.1 | NonPC | Public | 2012 | USA | Env | 88 | 100 | 0.06 | 98.1129 | 5104773 | 45.34 | 105 | 176382 |
| SAMN09074607-GCA_015808175.1 | CFSAN079430 | SAMN09074607 | GCA_015808175.1 | NonPC | Public | 2012 | USA | Env | 89.2 | 100 | 0 | 98.1013 | 5075283 | 45.32 | 85 | 194901 |
| SAMN09074609-GCA_015808275.1 | CFSAN079432 | SAMN09074609 | GCA_015808275.1 | NonPC | Public | 2012 | USA | Env | 87.9 | 100 | 0.06 | 98.1081 | 5109934 | 45.32 | 77 | 224191 |
| SAMN09074611-GCA_015808055.1 | CFSAN079428 | SAMN09074611 | GCA_015808055.1 | PrePC | Public | 2012 | USA | Env | 94.8 | 100 | 0.35 | 99.8211 | 5104053 | 45.32 | 70 | 228767 |
| SAMN09074615-GCA_015808315.1 | CFSAN079468 | SAMN09074615 | GCA_015808315.1 | NonPC | Public | 2012 | USA | Env | 90.6 | 100 | 0.82 | 98.2597 | 5273878 | 45.39 | 46 | 887291 |
| SAMN09074618-GCA_015808015.1 | CFSAN079470 | SAMN09074618 | GCA_015808015.1 | NonPC | Public | 2012 | USA | Env | 90.6 | 100 | 0.54 | 98.2171 | 5223201 | 45.38 | 56 | 648555 |
| SAMN09074632-GCA_015808115.1 | CFSAN079471 | SAMN09074632 | GCA_015808115.1 | NonPC | Public | 2013 | USA | Env | 91.6 | 100 | 0.13 | 98.2361 | 5175907 | 45.34 | 43 | 1079133 |
| SAMN09074633-GCA_015808035.1 | CFSAN079436 | SAMN09074633 | GCA_015808035.1 | NonPC | Public | 2012 | USA | Env | 91.1 | 100 | 0.2 | 98.2625 | 5152968 | 45.22 | 52 | 249620 |
| SAMN09074634-GCA_015808135.1 | CFSAN079465 | SAMN09074634 | GCA_015808135.1 | NonPC | Public | 2012 | USA | Env | 91.1 | 100 | 2.86 | 98.3135 | 5335003 | 45.24 | 51 | 293581 |
| SAMN09074637-GCA_015808075.1 | CFSAN079472 | SAMN09074637 | GCA_015808075.1 | NonPC | Public | 2013 | USA | Env | 91.4 | 99.93 | 0 | 98.3019 | 5132944 | 45.33 | 36 | 535104 |
| SAMN09074639-GCA_015808295.1 | CFSAN079435 | SAMN09074639 | GCA_015808295.1 | NonPC | Public | 2012 | USA | Env | 91.5 | 100 | 0 | 98.2937 | 5082143 | 45.3 | 42 | 270153 |
| SAMN09074642-GCA_015807955.1 | CFSAN079434 | SAMN09074642 | GCA_015807955.1 | NonPC | Public | 2012 | USA | Env | 92.3 | 100 | 0.01 | 98.38 | 5193845 | 45.46 | 58 | 410874 |
| SAMN09074986-GCA_015808255.1 | CFSAN079427 | SAMN09074986 | GCA_015808255.1 | NonPC | Public | 2012 | USA | Env | 91 | 100 | 0.41 | 98.3303 | 5221817 | 45.31 | 90 | 147499 |
| SAMN09074990-GCA_015808195.1 | CFSAN079473 | SAMN09074990 | GCA_015808195.1 | NonPC | Public | 2013 | USA | Env | 91.7 | 100 | 0.08 | 98.3498 | 5241456 | 45.19 | 66 | 231348 |
| SAMN09074991-GCA_015808095.1 | CFSAN079475 | SAMN09074991 | GCA_015808095.1 | NonPC | Public | 2013 | USA | Env | 91.1 | 100 | 2.86 | 98.2958 | 5339892 | 45.24 | 46 | 402301 |
| SAMN09074992-GCA_015807975.1 | CFSAN079476 | SAMN09074992 | GCA_015807975.1 | NonPC | Public | 2013 | USA | Env | 91 | 100 | 0.41 | 98.2867 | 5234310 | 45.28 | 102 | 130937 |
| SAMN09075001-GCA_015808155.1 | CFSAN079469 | SAMN09075001 | GCA_015808155.1 | NonPC | Public | 2012 | USA | Env | 90.7 | 100 | 0.82 | 98.2262 | 5281511 | 45.38 | 34 | 3144583 |
| SAMN09206535-GCA_015807655.1 | PNUSAV000185 | SAMN09206535 | GCA_015807655.1 | NonPC | Public | NA | USA | Clinical | 91.6 | 100 | 0 | 98.2915 | 5147394 | 45.24 | 58 | 376929 |
| SAMN09242481-GCA_015805095.1 | CFSAN081466 | SAMN09242481 | GCA_015805095.1 | NonPC | Public | 2013 | USA | Env | 88.8 | 100 | 0 | 98.0968 | 4983277 | 45.3 | 71 | 187616 |
| SAMN09242482-GCA_015806215.1 | CFSAN081469 | SAMN09242482 | GCA_015806215.1 | NonPC | Public | 2014 | USA | Env | 90.5 | 100 | 0.24 | 98.2032 | 5247383 | 45.33 | 192 | 62736 |
| SAMN09242486-GCA_015806255.1 | CFSAN081465 | SAMN09242486 | GCA_015806255.1 | NonPC | Public | 2013 | USA | Env | 88.4 | 100 | 0.07 | 98.1181 | 4986987 | 45.36 | 113 | 101004 |
| SAMN09242487-GCA_015805315.1 | CFSAN081440 | SAMN09242487 | GCA_015805315.1 | NonPC | Public | 2013 | USA | Env | 91.4 | 100 | 0.35 | 98.3697 | 5189560 | 45.32 | 92 | 115535 |
| SAMN09242490-GCA_015806275.1 | CFSAN081448 | SAMN09242490 | GCA_015806275.1 | NonPC | Public | 2013 | USA | Env | 88.6 | 99.73 | 0.42 | 98.2169 | 4921253 | 45.41 | 107 | 144209 |
| SAMN09242492-GCA_015806175.1 | CFSAN081449 | SAMN09242492 | GCA_015806175.1 | NonPC | Public | 2013 | USA | Env | 88.6 | 100 | 0.52 | 98.1036 | 4924411 | 45.42 | 114 | 107015 |
| SAMN09242494-GCA_015805135.1 | CFSAN081447 | SAMN09242494 | GCA_015805135.1 | NonPC | Public | 2013 | USA | Env | 89.6 | 100 | 0.11 | 98.262 | 5222603 | 45.2 | 77 | 222726 |
| SAMN09242496-GCA_015806635.1 | CFSAN081446 | SAMN09242496 | GCA_015806635.1 | NonPC | Public | 2013 | USA | Env | 91.1 | 100 | 0.34 | 98.2595 | 5306929 | 45.24 | 101 | 129322 |
| SAMN09242499-GCA_015806195.1 | CFSAN081445 | SAMN09242499 | GCA_015806195.1 | NonPC | Public | 2013 | USA | Env | 88.9 | 100 | 0.64 | 98.1424 | 5074412 | 45.33 | 113 | 125755 |
| SAMN09242501-GCA_015805275.1 | CFSAN081451 | SAMN09242501 | GCA_015805275.1 | NonPC | Public | 2013 | USA | Env | 89.6 | 100 | 0.12 | 98.323 | 5273339 | 45.17 | 71 | 255955 |
| SAMN09242502-GCA_015806155.1 | CFSAN081450 | SAMN09242502 | GCA_015806155.1 | NonPC | Public | 2013 | USA | Env | 91 | 100 | 0.48 | 98.2738 | 5301820 | 45.2 | 69 | 174653 |
| SAMN09242504-GCA_015805225.1 | CFSAN081452 | SAMN09242504 | GCA_015805225.1 | NonPC | Public | 2013 | USA | Env | 89.6 | 100 | 0.12 | 98.3046 | 5171404 | 45.28 | 96 | 101352 |
| SAMN09242507-GCA_015806115.1 | CFSAN081459 | SAMN09242507 | GCA_015806115.1 | NonPC | Public | 2013 | USA | Env | 91 | 100 | 0.78 | 98.2726 | 5207408 | 45.33 | 101 | 150262 |
| SAMN09242509-GCA_015805175.1 | CFSAN081458 | SAMN09242509 | GCA_015805175.1 | NonPC | Public | 2013 | USA | Env | 88.5 | 100 | 0.35 | 98.18 | 5026984 | 45.24 | 102 | 153671 |
| SAMN09242510-GCA_015806595.1 | CFSAN081457 | SAMN09242510 | GCA_015806595.1 | NonPC | Public | 2013 | USA | Env | 91.1 | 100 | 0.07 | 98.2775 | 5254597 | 45.2 | 72 | 270728 |
| SAMN09242511-GCA_015806975.1 | CFSAN081454 | SAMN09242511 | GCA_015806975.1 | PrePC | Public | 2013 | USA | Env | 95 | 100 | 0.69 | 99.8344 | 5160765 | 45.29 | 41 | 450384 |
| SAMN09242513-GCA_015807195.1 | CFSAN081453 | SAMN09242513 | GCA_015807195.1 | NonPC | Public | 2013 | USA | Env | 89 | 100 | 0.54 | 98.2811 | 5139848 | 45.27 | 92 | 198330 |
| SAMN09242514-GCA_015806775.1 | CFSAN081473 | SAMN09242514 | GCA_015806775.1 | NonPC | Public | 2014 | USA | Env | 91 | 100 | 0.89 | 98.4171 | 5091048 | 45.25 | 85 | 149567 |
| SAMN09242515-GCA_015804195.1 | CFSAN081456 | SAMN09242515 | GCA_015804195.1 | NonPC | Public | 2013 | USA | Env | 88.2 | 100 | 0.78 | 98.1101 | 5055263 | 45.24 | 49 | 277612 |

| | | | | | | | | | | | | | | | | |
|---|---|---|---|---|---|---|---|---|---|---|---|---|---|---|---|---|
| SAMN09242516-GCA_015807235.1 | CFSAN081455 | SAMN09242516 | GCA_015807235.1 | NonPC | Public | 2013 | USA | Env | 91.1 | 100 | 0.41 | 98.4543 | 5088329 | 45.22 | 49 | 356390 |
| SAMN09242518-GCA_015805355.1 | CFSAN081438 | SAMN09242518 | GCA_015805355.1 | NonPC | Public | 2013 | USA | Env | 92.1 | 100 | 1.84 | 98.3861 | 5176941 | 45.34 | 48 | 255205 |
| SAMN09242519-GCA_015803975.1 | CFSAN081463 | SAMN09242519 | GCA_015803975.1 | NonPC | Public | 2013 | USA | Env | 91 | 100 | 0.23 | 98.2923 | 5361644 | 45.12 | 98 | 198623 |
| SAMN09242520-GCA_015807275.1 | CFSAN081471 | SAMN09242520 | GCA_015807275.1 | NonPC | Public | 2014 | USA | Env | 91.2 | 100 | 0.15 | 98.2331 | 5346156 | 45.25 | 93 | 172523 |
| SAMN09242522-GCA_015805455.1 | CFSAN081439 | SAMN09242522 | GCA_015805455.1 | NonPC | Public | 2013 | USA | Env | 91.5 | 100 | 0.35 | 98.3525 | 5175368 | 45.27 | 66 | 186632 |
| SAMN09242527-GCA_015805395.1 | CFSAN081437 | SAMN09242527 | GCA_015805395.1 | PrePC | Public | 2013 | USA | Env | 94.2 | 100 | 0.15 | 99.805 | 5125903 | 45.49 | 191 | 52390 |
| SAMN09242528-GCA_015806695.1 | CFSAN081470 | SAMN09242528 | GCA_015806695.1 | NonPC | Public | 2014 | USA | Env | 91.6 | 100 | 2.03 | 98.2362 | 5186653 | 45.26 | 48 | 359277 |
| SAMN09242530-GCA_015807395.1 | CFSAN081462 | SAMN09242530 | GCA_015807395.1 | NonPC | Public | 2013 | USA | Env | 91.3 | 100 | 0 | 98.3392 | 5109328 | 45.27 | 58 | 418536 |
| SAMN09242533-GCA_015806135.1 | CFSAN081460 | SAMN09242533 | GCA_015806135.1 | NonPC | Public | 2013 | USA | Env | 88.5 | 100 | 0.42 | 98.2142 | 5029366 | 45.26 | 123 | 172077 |
| SAMN09242537-GCA_015806895.1 | CFSAN081461 | SAMN09242537 | GCA_015806895.1 | NonPC | Public | 2013 | USA | Env | 91.1 | 100 | 0.36 | 98.3207 | 5097339 | 45.34 | 132 | 130369 |
| SAMN09242542-GCA_015806095.1 | CFSAN081444 | SAMN09242542 | GCA_015806095.1 | NonPC | Public | 2013 | USA | Env | 89.5 | 100 | 2.14 | 98.2667 | 5290806 | 45.49 | 409 | 48597 |
| SAMN09242547-GCA_015806985.1 | CFSAN081441 | SAMN09242547 | GCA_015806985.1 | NonPC | Public | 2013 | USA | Env | 91 | 100 | 0.61 | 98.2897 | 5355070 | 45.27 | 72 | 192866 |
| SAMN09242548-GCA_016385205.1 | CFSAN081443 | SAMN09242548 | GCA_016385205.1 | NonPC | Public | 2013 | USA | Env | 91.4 | 100 | 0.22 | 98.2793 | 5187132 | 45.22 | 102 | 150654 |
| SAMN09242549-GCA_015806875.1 | CFSAN081442 | SAMN09242549 | GCA_015806875.1 | NonPC | Public | 2013 | USA | Env | 91.5 | 100 | 2.87 | 98.2969 | 5309288 | 45.19 | 43 | 469149 |
| SAMN09280044-GCA_015805735.1 | 218340 | SAMN09280044 | GCA_015805735.1 | NonPC | Public | 2015 | NA | Clinical | 89.6 | 99.91 | 1.27 | 98.2077 | 5386861 | 45.12 | 116 | 125323 |
| SAMN09280045-GCA_015805655.1 | 218342 | SAMN09280045 | GCA_015805655.1 | PC | Public | 2015 | NA | Clinical | 98 | 100 | 3.2 | 99.9937 | 5176132 | 45.25 | 87 | 184919 |
| SAMN09280046-GCA_015805805.1 | 218346 | SAMN09280046 | GCA_015805805.1 | PC | Public | 2015 | NA | Clinical | 97.7 | 100 | 4.9 | 99.9712 | 5284672 | 45.27 | 110 | 103970 |
| SAMN09280047-GCA_015805855.1 | 218355 | SAMN09280047 | GCA_015805855.1 | PC | Public | 2015 | NA | Clinical | 97.6 | 100 | 2.82 | 99.9914 | 5202545 | 45.29 | 78 | 152529 |
| SAMN09280048-GCA_015805675.1 | 218360 | SAMN09280048 | GCA_015805675.1 | NonPC | Public | 2015 | NA | Clinical | 89.6 | 100 | 1.56 | 98.2666 | 5191416 | 45.15 | 87 | 179016 |
| SAMN09280049-GCA_015805575.1 | 218362 | SAMN09280049 | GCA_015805575.1 | NonPC | Public | 2015 | NA | Clinical | 88.2 | 100 | 2.12 | 98.2169 | 5201074 | 45.13 | 130 | 171579 |
| SAMN09280050-GCA_015805715.1 | 235118 | SAMN09280050 | GCA_015805715.1 | NonPC | Public | 2016 | NA | Clinical | 90.9 | 100 | 0.54 | 98.2927 | 5100696 | 45.25 | 143 | 84243 |
| SAMN09280051-GCA_015805695.1 | 242385 | SAMN09280051 | GCA_015805695.1 | PC | Public | 2016 | NA | Clinical | 97.7 | 100 | 3.66 | 99.9938 | 5162999 | 45.27 | 91 | 145869 |
| SAMN09280052-GCA_015805795.1 | 245518 | SAMN09280052 | GCA_015805795.1 | PC | Public | 2016 | NA | Clinical | 97.1 | 100 | 0.41 | 99.9155 | 5157160 | 45.28 | 168 | 71400 |
| SAMN09280054-GCA_015805755.1 | 399421 | SAMN09280054 | GCA_015805755.1 | PC | Public | 2015 | NA | Clinical | 97.2 | 100 | 3.32 | 99.9632 | 5239867 | 45.24 | 103 | 141759 |
| SAMN09280055-GCA_015805555.1 | 399423 | SAMN09280055 | GCA_015805555.1 | PC | Public | 2015 | NA | Clinical | 97.9 | 100 | 4.37 | 99.9933 | 5374779 | 45.3 | 81 | 179493 |
| SAMN09280056-GCA_015805615.1 | 399425 | SAMN09280056 | GCA_015805615.1 | NonPC | Public | 2015 | NA | Clinical | 90.5 | 100 | 4.02 | 98.3297 | 5250944 | 45.17 | 114 | 188568 |
| SAMN09280057-GCA_015805595.1 | 404285 | SAMN09280057 | GCA_015805595.1 | PC | Public | 2016 | NA | Clinical | 97.1 | 100 | 4.63 | 99.9675 | 5270911 | 45.22 | 85 | 176741 |
| SAMN09280059-GCA_015805775.1 | 404287 | SAMN09280059 | GCA_015805775.1 | NonPC | Public | 2016 | NA | Clinical | 91.2 | 100 | 3.24 | 98.4094 | 5151540 | 45.37 | 68 | 192804 |
| SAMN09280061-GCA_015805515.1 | 476479 | SAMN09280061 | GCA_015805515.1 | NonPC | Public | 2018 | NA | Clinical | 89.4 | 100 | 2.82 | 98.3093 | 5115745 | 45.31 | 138 | 82021 |
| SAMN09280065-GCA_015805975.1 | 513298 | SAMN09280065 | GCA_015805975.1 | NonPC | Public | 2018 | NA | Clinical | 88.1 | 100 | 1.96 | 98.2947 | 5015457 | 45.41 | 249 | 42221 |
| SAMN09280068-GCA_015807405.1 | 524168 | SAMN09280068 | GCA_015807405.1 | NonPC | Public | 2017 | NA | Clinical | 92.6 | 100 | 3.54 | 98.358 | 5218937 | 45.29 | 68 | 188262 |
| SAMN09280071-GCA_015805895.1 | 539844 | SAMN09280071 | GCA_015805895.1 | NonPC | Public | 2017 | NA | Clinical | 89.6 | 100 | 4.68 | 98.2699 | 5282119 | 45.24 | 80 | 181831 |
| SAMN09280073-GCA_015805875.1 | 539846 | SAMN09280073 | GCA_015805875.1 | PC | Public | 2017 | NA | Clinical | 98 | 100 | 2.72 | 99.9889 | 5355261 | 45.26 | 77 | 179174 |
| SAMN09295476-GCA_015805075.1 | CFSAN081912 | SAMN09295476 | GCA_015805075.1 | NonPC | Public | 2014 | USA | Env | 91.1 | 100 | 0.48 | 98.4204 | 5085974 | 45.23 | 64 | 197662 |
| SAMN09295478-GCA_015805955.1 | CFSAN081933 | SAMN09295478 | GCA_015805955.1 | NonPC | Public | 2014 | USA | Env | 90.9 | 100 | 3.32 | 98.2879 | 5353033 | 45.29 | 91 | 108066 |
| SAMN09295479-GCA_015806015.1 | CFSAN081932 | SAMN09295479 | GCA_015806015.1 | NonPC | Public | 2014 | USA | Env | 92.1 | 100 | 4.35 | 98.405 | 5300908 | 45.3 | 80 | 166816 |
| SAMN09295480-GCA_015803695.1 | CFSAN081956 | SAMN09295480 | GCA_015803695.1 | NonPC | Public | 2014 | USA | Env | 91 | 100 | 4.22 | 98.2739 | 5406357 | 45.27 | 42 | 349796 |
| SAMN09295481-GCA_015804395.1 | CFSAN081955 | SAMN09295481 | GCA_015804395.1 | NonPC | Public | 2014 | USA | Env | 90.4 | 100 | 3 | 98.2601 | 5270229 | 45.24 | 47 | 244325 |
| SAMN09295482-GCA_015805035.1 | CFSAN081954 | SAMN09295482 | GCA_015805035.1 | NonPC | Public | 2014 | USA | Env | 91.1 | 100 | 4.36 | 98.2436 | 5213165 | 45.33 | 42 | 263039 |
| SAMN09295483-GCA_015805115.1 | CFSAN081953 | SAMN09295483 | GCA_015805115.1 | NonPC | Public | 2014 | USA | Env | 88.8 | 99.73 | 0.36 | 98.1971 | 4961229 | 45.31 | 77 | 180631 |
| SAMN09295484-GCA_015806515.1 | CFSAN081919 | SAMN09295484 | GCA_015806515.1 | NonPC | Public | 2014 | USA | Env | 91 | 100 | 1.04 | 98.2217 | 5293712 | 45.29 | 80 | 130981 |
| SAMN09295485-GCA_015806415.1 | CFSAN081952 | SAMN09295485 | GCA_015806415.1 | NonPC | Public | 2014 | USA | Env | 88.7 | 99.73 | 0.2 | 98.174 | 4967224 | 45.33 | 80 | 150872 |
| SAMN09295486-GCA_015806575.1 | CFSAN081916 | SAMN09295486 | GCA_015806575.1 | NonPC | Public | 2014 | USA | Env | 90.9 | 100 | 1.68 | 98.4533 | 5103076 | 45.34 | 121 | 100219 |
| SAMN09295487-GCA_015807135.1 | CFSAN081938 | SAMN09295487 | GCA_015807135.1 | NonPC | Public | 2014 | USA | Env | 90.6 | 100 | 0.27 | 98.2882 | 5152747 | 45.27 | 136 | 89045 |
| SAMN09295488-GCA_015807245.1 | CFSAN081915 | SAMN09295488 | GCA_015807245.1 | NonPC | Public | 2014 | USA | Env | 90.9 | 100 | 0.48 | 98.3943 | 5079102 | 45.33 | 116 | 95039 |
| SAMN09295533-GCA_015804045.1 | CFSAN081950 | SAMN09295533 | GCA_015804045.1 | NonPC | Public | 2014 | USA | Env | 91.1 | 100 | 0.15 | 98.2175 | 5275120 | 45.22 | 63 | 270192 |
| SAMN09295739-GCA_015806395.1 | CFSAN081917 | SAMN09295739 | GCA_015806395.1 | NonPC | Public | 2014 | USA | Env | 91.4 | 100 | 0.99 | 98.2315 | 5158551 | 45.35 | 108 | 102597 |
| SAMN09295740-GCA_015806475.1 | CFSAN081939 | SAMN09295740 | GCA_015806475.1 | NonPC | Public | 2014 | USA | Env | 91.4 | 100 | 0.36 | 98.2071 | 5096209 | 45.34 | 109 | 91890 |

| | | | | | | | | | | | | | | | | |
|---|---|---|---|---|---|---|---|---|---|---|---|---|---|---|---|---|
| SAMN09296051-GCA_015807115.1 | CFSAN081930 | SAMN09296051 | GCA_015807115.1 | NonPC | Public | 2014 | USA | Env | 89.5 | 100 | 0.27 | 98.1838 | 5225784 | 45.34 | 60 | 170768 |
| SAMN09296172-GCA_015805195.1 | CFSAN081923 | SAMN09296172 | GCA_015805195.1 | NonPC | Public | 2014 | USA | Env | 90.9 | 100 | 0.55 | 98.2571 | 5267853 | 45.28 | 94 | 133011 |
| SAMN09296173-GCA_015806795.1 | CFSAN081945 | SAMN09296173 | GCA_015806795.1 | NonPC | Public | 2014 | USA | Env | 90.8 | 100 | 0.21 | 98.2675 | 5162753 | 45.37 | 188 | 49945 |
| SAMN09296174-GCA_015804135.1 | CFSAN081922 | SAMN09296174 | GCA_015804135.1 | NonPC | Public | 2014 | USA | Env | 91.3 | 100 | 0.15 | 98.2196 | 5086612 | 45.34 | 116 | 84478 |
| SAMN09296175-GCA_015806355.1 | CFSAN081944 | SAMN09296175 | GCA_015806355.1 | NonPC | Public | 2014 | USA | Env | 90.7 | 100 | 1.32 | 98.2832 | 5367837 | 45.27 | 142 | 86003 |
| SAMN09296176-GCA_015806615.1 | CFSAN081921 | SAMN09296176 | GCA_015806615.1 | NonPC | Public | 2014 | USA | Env | 91 | 100 | 1.36 | 98.2385 | 5084337 | 45.37 | 86 | 172378 |
| SAMN09296177-GCA_015806035.1 | CFSAN081943 | SAMN09296177 | GCA_015806035.1 | NonPC | Public | 2014 | USA | Env | 90.8 | 100 | 0.75 | 98.271 | 5304549 | 45.29 | 120 | 90226 |
| SAMN09296178-GCA_015807095.1 | CFSAN081929 | SAMN09296178 | GCA_015807095.1 | NonPC | Public | 2014 | USA | Env | 91.1 | 100 | 0 | 98.257 | 5271439 | 45.23 | 91 | 143594 |
| SAMN09296179-GCA_015807155.1 | CFSAN081931 | SAMN09296179 | GCA_015807155.1 | NonPC | Public | 2014 | USA | Env | 91 | 100 | 3.95 | 98.2774 | 5386276 | 45.3 | 83 | 125853 |
| SAMN09296181-GCA_015807015.1 | CFSAN081924 | SAMN09296181 | GCA_015807015.1 | NonPC | Public | 2014 | USA | Env | 91.1 | 100 | 0.07 | 98.2615 | 5250560 | 45.22 | 98 | 148161 |
| SAMN09296187-GCA_015807295.1 | CFSAN081927 | SAMN09296187 | GCA_015807295.1 | NonPC | Public | 2014 | USA | Env | 91.4 | 100 | 0.08 | 98.3588 | 5148818 | 45.29 | 89 | 118635 |
| SAMN09296189-GCA_015805055.1 | CFSAN081949 | SAMN09296189 | GCA_015805055.1 | NonPC | Public | 2014 | USA | Env | 91.1 | 100 | 0.44 | 98.2226 | 5315367 | 45.22 | 72 | 189982 |
| SAMN09296195-GCA_015806235.1 | CFSAN081926 | SAMN09296195 | GCA_015806235.1 | NonPC | Public | 2014 | USA | Env | 91 | 100 | 2.18 | 98.2523 | 5146692 | 45.37 | 89 | 117260 |
| SAMN09296196-GCA_015804155.1 | CFSAN081948 | SAMN09296196 | GCA_015804155.1 | NonPC | Public | 2014 | USA | Env | 90.4 | 100 | 1.08 | 98.2631 | 5371880 | 45.22 | 71 | 194714 |
| SAMN09296199-GCA_015806055.1 | CFSAN081925 | SAMN09296199 | GCA_015806055.1 | NonPC | Public | 2014 | USA | Env | 90.2 | 100 | 0.41 | 98.2654 | 5233624 | 45.4 | 277 | 34046 |
| SAMN09296202-GCA_015803955.1 | CFSAN081947 | SAMN09296202 | GCA_015803955.1 | NonPC | Public | 2014 | USA | Env | 90.9 | 100 | 2.81 | 98.3208 | 5513976 | 45.24 | 99 | 187907 |
| SAMN09296206-GCA_015806295.1 | CFSAN081942 | SAMN09296206 | GCA_015806295.1 | NonPC | Public | 2014 | USA | Env | 90.8 | 100 | 2 | 98.2781 | 5273833 | 45.34 | 131 | 80234 |
| SAMN09296209-GCA_015806435.1 | CFSAN081941 | SAMN09296209 | GCA_015806435.1 | NonPC | Public | 2014 | USA | Env | 88.3 | 100 | 2.26 | 98.1962 | 5067920 | 45.32 | 147 | 81626 |
| SAMN09296210-GCA_015806075.1 | CFSAN081951 | SAMN09296210 | GCA_015806075.1 | NonPC | Public | 2014 | USA | Env | 90.9 | 100 | 0.15 | 98.2376 | 5345465 | 45.28 | 106 | 115205 |
| SAMN09296217-GCA_015806555.1 | CFSAN081920 | SAMN09296217 | GCA_015806555.1 | NonPC | Public | 2014 | USA | Env | 91.1 | 100 | 0.07 | 98.2457 | 5066523 | 45.33 | 60 | 319502 |
| SAMN09296219-GCA_015806815.1 | CFSAN081918 | SAMN09296219 | GCA_015806815.1 | NonPC | Public | 2014 | USA | Env | 88.4 | 100 | 0.46 | 98.1341 | 4917606 | 45.41 | 107 | 103354 |
| SAMN09375013-GCA_015806675.1 | PNUSAV000179 | SAMN09375013 | GCA_015806675.1 | NonPC | Public | NA | USA | Clinical | 91.2 | 100 | 4.08 | 98.3514 | 5279846 | 45.33 | 61 | 190397 |
| SAMN09375021-GCA_015806715.1 | PNUSAV000180 | SAMN09375021 | GCA_015806715.1 | NonPC | Public | NA | USA | Clinical | 90.6 | 100 | 1.94 | 98.3552 | 5085547 | 45.31 | 84 | 196157 |
| SAMN09456007-GCA_015806835.1 | PNUSAV000187 | SAMN09456007 | GCA_015806835.1 | NonPC | Public | NA | USA | Clinical | 91.4 | 100 | 0.61 | 98.2907 | 5164103 | 45.25 | 41 | 718998 |
| SAMN09464982-GCA_015806915.1 | PNUSAV000188 | SAMN09464982 | GCA_015806915.1 | NonPC | Public | NA | USA | Clinical | 91.4 | 100 | 0.82 | 98.3132 | 5153811 | 45.24 | 41 | 719457 |
| SAMN09488304-GCA_015806535.1 | PNUSAV000192 | SAMN09488304 | GCA_015806535.1 | PC | Public | NA | USA | Clinical | 97.5 | 100 | 0.41 | 99.9653 | 5113367 | 45.28 | 87 | 192530 |
| SAMN09488305-GCA_015806375.1 | PNUSAV000190 | SAMN09488305 | GCA_015806375.1 | PC | Public | NA | USA | Clinical | 97.5 | 100 | 0.34 | 99.9685 | 5108455 | 45.27 | 86 | 179912 |
| SAMN09488306-GCA_015806455.1 | PNUSAV000189 | SAMN09488306 | GCA_015806455.1 | PC | Public | NA | USA | Clinical | 96.7 | 100 | 0.61 | 99.9723 | 5077529 | 45.39 | 192 | 73501 |
| SAMN09635686-GCA_015807175.1 | PNUSAV000191 | SAMN09635686 | GCA_015807175.1 | NonPC | Public | NA | USA | Clinical | 90.5 | 100 | 0.56 | 98.1657 | 5320618 | 45.13 | 44 | 262367 |
| SAMN09642115-GCA_015807075.1 | PNUSAV000195 | SAMN09642115 | GCA_015807075.1 | NonPC | Public | NA | USA | Clinical | 91.4 | 100 | 2.71 | 98.2942 | 5209684 | 45.25 | 50 | 269897 |
| SAMN09655865-GCA_015805015.1 | CFSAN083198 | SAMN09655865 | GCA_015805015.1 | NonPC | Public | 2018 | Venezuela | Env | 89.9 | 100 | 2.57 | 98.2522 | 5130374 | 45.38 | 41 | 461391 |
| SAMN09655866-GCA_015804215.1 | CFSAN083199 | SAMN09655866 | GCA_015804215.1 | NonPC | Public | 2018 | Venezuela | Env | 90 | 100 | 0.64 | 98.2732 | 5305347 | 45.32 | 32 | 548456 |
| SAMN09655880-GCA_015804975.1 | CFSAN083200 | SAMN09655880 | GCA_015804975.1 | NonPC | Public | 2018 | Venezuela | Env | 92.1 | 100 | 3.07 | 98.4082 | 5158412 | 45.33 | 51 | 197459 |
| SAMN09655881-GCA_015804955.1 | CFSAN083201 | SAMN09655881 | GCA_015804955.1 | NonPC | Public | 2018 | Venezuela | Env | 89.1 | 100 | 1.35 | 98.3001 | 5012539 | 45.44 | 37 | 291635 |
| SAMN09655882-GCA_015804235.1 | CFSAN083202 | SAMN09655882 | GCA_015804235.1 | NonPC | Public | 2018 | Venezuela | Env | 89.1 | 100 | 0.34 | 98.322 | 4990482 | 45.47 | 52 | 255669 |
| SAMN09655883-GCA_015804935.1 | CFSAN083197 | SAMN09655883 | GCA_015804935.1 | NonPC | Public | 2018 | Venezuela | Env | 89.8 | 100 | 4.75 | 98.2933 | 5186275 | 45.34 | 58 | 160904 |
| SAMN09664793-GCA_015806315.1 | PNUSAV000194 | SAMN09664793 | GCA_015806315.1 | PC | Public | NA | USA | Clinical | 96.7 | 100 | 0.07 | 99.9774 | 5086489 | 45.36 | 117 | 204020 |
| SAMN09671573-GCA_015804035.1 | PNUSAV000206 | SAMN09671573 | GCA_015804035.1 | PC | Public | NA | USA | Clinical | 97.8 | 100 | 0 | 99.97 | 5136822 | 45.28 | 83 | 186320 |
| SAMN09671574-GCA_015803895.1 | PNUSAV000205 | SAMN09671574 | GCA_015803895.1 | NonPC | Public | NA | USA | Clinical | 91.3 | 100 | 0.36 | 98.3393 | 5065736 | 45.37 | 93 | 179401 |
| SAMN09671576-GCA_015805145.1 | PNUSAV000203 | SAMN09671576 | GCA_015805145.1 | PC | Public | NA | USA | Clinical | 97.8 | 100 | 3.89 | 99.9681 | 5323645 | 45.3 | 67 | 204642 |
| SAMN09671577-GCA_015804095.1 | PNUSAV000202 | SAMN09671577 | GCA_015804095.1 | PC | Public | NA | USA | Clinical | 97.7 | 100 | 3.35 | 99.9682 | 5227877 | 45.31 | 80 | 175513 |
| SAMN09671578-GCA_015803915.1 | PNUSAV000201 | SAMN09671578 | GCA_015803915.1 | NonPC | Public | NA | USA | Clinical | 91.3 | 100 | 1.24 | 98.3293 | 5159694 | 45.25 | 69 | 194129 |
| SAMN09671587-GCA_015804115.1 | PNUSAV000207 | SAMN09671587 | GCA_015804115.1 | PC | Public | NA | USA | Clinical | 97.8 | 100 | 2.17 | 99.9691 | 5194689 | 45.3 | 84 | 179296 |
| SAMN09689475-GCA_015803755.1 | PNUSAV000186 | SAMN09689475 | GCA_015803755.1 | PC | Public | NA | USA | Clinical | 97.8 | 100 | 1.89 | 99.9606 | 5247919 | 45.28 | 62 | 293968 |
| SAMN09694269-GCF_009859405.1 | SC4 | SAMN09694269 | GCF_009859405.1 | NonPC | Public | 2016 | South_Korea | Env | 91.7 | 100 | 0.19 | 98.2962 | 5296544 | 45.35 | 3 | 3365393 |
| SAMN09694270-GCF_009859385.1 | SK | SAMN09694270 | GCF_009859385.1 | NonPC | Public | 2016 | South_Korea | Env | 91.8 | 100 | 0.19 | 98.2958 | 5302943 | 45.35 | 3 | 3355895 |
| SAMN09694271-GCF_009859365.1 | SM3 | SAMN09694271 | GCF_009859365.1 | NonPC | Public | 2016 | South_Korea | Env | 92.7 | 100 | 0.41 | 98.2747 | 5440354 | 45.24 | 5 | 3466068 |
| SAMN09694272-GCF_009859345.1 | SM4 | SAMN09694272 | GCF_009859345.1 | NonPC | Public | 2016 | South_Korea | Env | 92.7 | 100 | 0.41 | 98.2886 | 5444704 | 45.23 | 5 | 3466067 |

| | | | | | | | | | | | | | | | | |
|---|---|---|---|---|---|---|---|---|---|---|---|---|---|---|---|---|
| SAMN09694273-GCF_009859325.1 | WC15 | SAMN09694273 | GCF_009859325.1 | NonPC | Public | 2016 | South_Korea | Env | 91.8 | 100 | 0.19 | 98.2929 | 5298825 | 45.35 | 3 | 3365382 |
| SAMN09694274-GCF_009859305.1 | WS | SAMN09694274 | GCF_009859305.1 | NonPC | Public | 2016 | South_Korea | Env | 91.4 | 100 | 0 | 98.2993 | 5206684 | 45.43 | 4 | 3383458 |
| SAMN09694275-GCF_009859285.1 | WY3 | SAMN09694275 | GCF_009859285.1 | NonPC | Public | 2016 | South_Korea | Env | 91.8 | 100 | 0.19 | 98.3395 | 5307166 | 45.35 | 4 | 3366116 |
| SAMN09697267-GCA_015803875.1 | PNUSAV000216 | SAMN09697267 | GCA_015803875.1 | NonPC | Public | NA | USA | Clinical | 91.3 | 100 | 1 | 98.3009 | 5178283 | 45.26 | 73 | 131715 |
| SAMN09697268-GCA_015803935.1 | PNUSAV000215 | SAMN09697268 | GCA_015803935.1 | NonPC | Public | NA | USA | Clinical | 91.2 | 100 | 4.63 | 98.3051 | 5239967 | 45.32 | 86 | 177610 |
| SAMN09703853-GCA_016435625.1 | PNUSAV000228 | SAMN09703853 | GCA_016435625.1 | NonPC | Public | NA | USA | Clinical | 87.5 | 100 | 1.28 | 98.0315 | 5072443 | 45.35 | 72 | 182857 |
| SAMN09703856-GCA_015804165.1 | PNUSAV000224 | SAMN09703856 | GCA_015804165.1 | NonPC | Public | NA | USA | Clinical | 89.7 | 100 | 0.38 | 98.2786 | 5193425 | 45.2 | 52 | 283246 |
| SAMN09703857-GCA_015803815.1 | PNUSAV000223 | SAMN09703857 | GCA_015803815.1 | PC | Public | NA | USA | Clinical | 97.8 | 100 | 3.76 | 99.9676 | 5243804 | 45.28 | 64 | 291308 |
| SAMN09703859-GCA_015803795.1 | PNUSAV000222 | SAMN09703859 | GCA_015803795.1 | PC | Public | NA | USA | Clinical | 97.7 | 100 | 0.48 | 99.9688 | 5158479 | 45.3 | 89 | 200067 |
| SAMN09703860-GCA_015804015.1 | PNUSAV000221 | SAMN09703860 | GCA_015804015.1 | PC | Public | NA | USA | Clinical | 97.7 | 100 | 1.36 | 99.9657 | 5188488 | 45.28 | 55 | 290482 |
| SAMN09703861-GCA_015803775.1 | PNUSAV000197 | SAMN09703861 | GCA_015803775.1 | NonPC | Public | NA | USA | Clinical | 91.4 | 100 | 4.63 | 98.2746 | 5299179 | 45.26 | 52 | 269886 |
| SAMN09709537-GCA_015804995.1 | PNUSAV000220 | SAMN09709537 | GCA_015804995.1 | NonPC | Public | NA | USA | Clinical | 91.3 | 100 | 1.63 | 98.269 | 5187710 | 45.27 | 77 | 131540 |
| SAMN09724629-GCA_015804355.1 | PNUSAV000225 | SAMN09724629 | GCA_015804355.1 | NonPC | Public | NA | USA | Clinical | 91.3 | 100 | 0.59 | 98.3181 | 5142729 | 45.24 | 43 | 556709 |
| SAMN09724631-GCA_015804335.1 | PNUSAV000198 | SAMN09724631 | GCA_015804335.1 | NonPC | Public | NA | USA | Clinical | 91.4 | 100 | 1.7 | 98.3057 | 5204815 | 45.25 | 47 | 556720 |
| SAMN09726929-GCA_015804665.1 | PNUSAV000241 | SAMN09726929 | GCA_015804665.1 | NonPC | Public | NA | USA | Clinical | 89.1 | 100 | 0.83 | 98.0709 | 5110574 | 45.29 | 60 | 233228 |
| SAMN09726930-GCA_015804815.1 | PNUSAV000240 | SAMN09726930 | GCA_015804815.1 | NonPC | Public | NA | USA | Clinical | 90.8 | 100 | 1.35 | 98.3036 | 5201883 | 45.17 | 42 | 418297 |
| SAMN09726931-GCA_015804755.1 | PNUSAV000238 | SAMN09726931 | GCA_015804755.1 | NonPC | Public | NA | USA | Clinical | 91.4 | 100 | 0.48 | 98.2842 | 5153151 | 45.25 | 42 | 475308 |
| SAMN09726932-GCA_016377565.1 | PNUSAV000237 | SAMN09726932 | GCA_016377565.1 | NonPC | Public | NA | USA | Clinical | 91.4 | 100 | 0.27 | 98.2782 | 5141026 | 45.24 | 38 | 556708 |
| SAMN09726933-GCA_015804875.1 | PNUSAV000236 | SAMN09726933 | GCA_015804875.1 | NonPC | Public | NA | USA | Clinical | 91.4 | 100 | 4.77 | 98.2948 | 5321938 | 45.26 | 41 | 556708 |
| SAMN09726934-GCA_015804635.1 | PNUSAV000233 | SAMN09726934 | GCA_015804635.1 | NonPC | Public | NA | USA | Clinical | 91.4 | 100 | 1.91 | 98.2874 | 5228045 | 45.25 | 40 | 718934 |
| SAMN09726935-GCA_015804795.1 | PNUSAV000232 | SAMN09726935 | GCA_015804795.1 | NonPC | Public | NA | USA | Clinical | 91.4 | 100 | 0.5 | 98.2929 | 5153928 | 45.24 | 42 | 475251 |
| SAMN09726936-GCA_015804855.1 | PNUSAV000231 | SAMN09726936 | GCA_015804855.1 | NonPC | Public | NA | USA | Clinical | 91.3 | 100 | 2.72 | 98.2743 | 5206870 | 45.25 | 51 | 280401 |
| SAMN09726937-GCA_015804735.1 | PNUSAV000230 | SAMN09726937 | GCA_015804735.1 | NonPC | Public | NA | USA | Clinical | 91.3 | 100 | 0.95 | 98.3023 | 5155971 | 45.25 | 39 | 556708 |
| SAMN09726938-GCA_015804915.1 | PNUSAV000212 | SAMN09726938 | GCA_015804915.1 | NonPC | Public | NA | USA | Clinical | 89.5 | 100 | 1.13 | 98.2828 | 5194787 | 45.23 | 72 | 128567 |
| SAMN09726939-GCA_015804895.1 | PNUSAV000211 | SAMN09726939 | GCA_015804895.1 | PC | Public | NA | USA | Clinical | 97.4 | 100 | 3.19 | 99.9662 | 5196823 | 45.32 | 102 | 123189 |
| SAMN09737240-GCF_006383335.1 | VP830010 | SAMN09737240 | GCF_006383335.1 | NonPC | Public | 1983 | China | Clinical | 92.3 | 100 | 0.05 | 98.336 | 4990978 | 45.3 | 37 | 547356 |
| SAMN09737242-GCF_006383445.1 | VP150615 | SAMN09737242 | GCF_006383445.1 | NonPC | Public | 2015 | China | Clinical | 92.3 | 100 | 0.19 | 98.385 | 5032301 | 45.24 | 32 | 568816 |
| SAMN09737245-GCF_006383455.1 | VP161902 | SAMN09737245 | GCF_006383455.1 | NonPC | Public | 2016 | China | Clinical | 92.2 | 100 | 0.19 | 98.361 | 5021595 | 45.24 | 29 | 603271 |
| SAMN09737247-GCF_006383425.1 | VP140355 | SAMN09737247 | GCF_006383425.1 | NonPC | Public | 2014 | China | Clinical | 91.9 | 100 | 0.19 | 98.2769 | 5061807 | 45.27 | 37 | 547249 |
| SAMN09737248-GCF_006383325.1 | VP160483 | SAMN09737248 | GCF_006383325.1 | NonPC | Public | 2016 | China | Clinical | 92 | 100 | 0.19 | 98.3154 | 5066301 | 45.23 | 35 | 471444 |
| SAMN09742574-GCF_006383435.1 | MH16438 | SAMN09742574 | GCF_006383435.1 | NonPC | Public | 2016 | China | Clinical | 91.9 | 100 | 0.46 | 98.2746 | 5066271 | 45.2 | 29 | 603334 |
| SAMN09742575-GCF_006383315.1 | MH16778 | SAMN09742575 | GCF_006383315.1 | NonPC | Public | 2016 | China | Clinical | 91.9 | 100 | 0.46 | 98.2357 | 5079860 | 45.2 | 31 | 603376 |
| SAMN09742576-GCF_006383285.1 | MH16537 | SAMN09742576 | GCF_006383285.1 | NonPC | Public | 2016 | China | Clinical | 92.2 | 100 | 0.19 | 98.3795 | 5011029 | 45.26 | 30 | 603255 |
| SAMN09742577-GCF_006383245.1 | MH16256 | SAMN09742577 | GCF_006383245.1 | NonPC | Public | 2016 | China | Clinical | 92.2 | 100 | 0.05 | 98.1177 | 5118485 | 45.24 | 33 | 470297 |
| SAMN09742578-GCF_006383265.1 | FX16039 | SAMN09742578 | GCF_006383265.1 | NonPC | Public | 2016 | China | Clinical | 91.1 | 100 | 0.27 | 98.3714 | 4931985 | 45.4 | 19 | 868950 |
| SAMN09742579-GCF_006383355.1 | BS16021 | SAMN09742579 | GCF_006383355.1 | NonPC | Public | 2016 | China | Clinical | 89.7 | 100 | 0.27 | 98.412 | 5051817 | 45.34 | 37 | 411517 |
| SAMN09742580-GCF_006383225.1 | MH16126 | SAMN09742580 | GCF_006383225.1 | NonPC | Public | 2016 | China | Clinical | 93.1 | 100 | 0.05 | 98.36 | 5095087 | 45.34 | 33 | 425245 |
| SAMN09742581-GCF_006382975.1 | SJ16045 | SAMN09742581 | GCF_006382975.1 | NonPC | Public | 2016 | China | Clinical | 93.1 | 100 | 0.05 | 98.3956 | 5066014 | 45.31 | 33 | 516117 |
| SAMN09742582-GCF_006383385.1 | QP16184 | SAMN09742582 | GCF_006383385.1 | PC | Public | 2016 | China | Clinical | 97.4 | 100 | 0.07 | 99.9042 | 5180936 | 45.28 | 55 | 373744 |
| SAMN09742583-GCF_006382895.1 | QP16233 | SAMN09742583 | GCF_006382895.1 | PC | Public | 2016 | China | Clinical | 97.7 | 100 | 0 | 99.9222 | 5162116 | 45.24 | 59 | 373695 |
| SAMN09742584-GCF_006382955.1 | MH16341 | SAMN09742584 | GCF_006382955.1 | PC | Public | 2016 | China | Clinical | 97.3 | 100 | 0 | 99.8993 | 5100242 | 45.27 | 55 | 339399 |
| SAMN09742585-GCF_006383215.1 | HK16029 | SAMN09742585 | GCF_006383215.1 | NonPC | Public | 2016 | China | Clinical | 89.7 | 100 | 0 | 98.264 | 5141243 | 45.25 | 48 | 529796 |
| SAMN09742586-GCF_006382865.1 | BS16117 | SAMN09742586 | GCF_006382865.1 | NonPC | Public | 2016 | China | Clinical | 91.4 | 100 | 0.31 | 98.3003 | 5136065 | 45.29 | 52 | 405122 |
| SAMN09742587-GCF_006382855.1 | VP840119 | SAMN09742587 | GCF_006382855.1 | NonPC | Public | 1968 | Japan | Clinical | 92.1 | 100 | 0.14 | 98.3279 | 5142793 | 45.18 | 35 | 401934 |
| SAMN09742588-GCF_006383165.1 | QP17014 | SAMN09742588 | GCF_006383165.1 | NonPC | Public | 2017 | China | Clinical | 92.5 | 100 | 0.05 | 98.3426 | 5021509 | 45.27 | 25 | 539371 |
| SAMN09742589-GCF_006383155.1 | BS17035 | SAMN09742589 | GCF_006383155.1 | NonPC | Public | 2017 | China | Clinical | 92.5 | 100 | 0.05 | 98.3389 | 5133564 | 45.07 | 22 | 604414 |
| SAMN09742590-GCF_006382805.1 | PD17005 | SAMN09742590 | GCF_006382805.1 | NonPC | Public | 2017 | China | Clinical | 91.9 | 100 | 0.46 | 98.2802 | 5069087 | 45.2 | 25 | 603562 |
| SAMN09742591-GCF_006382785.1 | NH001 | SAMN09742591 | GCF_006382785.1 | NonPC | Public | 2009 | China | Clinical | 92.4 | 100 | 0.05 | 98.3416 | 5065309 | 45.27 | 23 | 603364 |

| | | | | | | | | | | | | | | | | |
|---|---|---|---|---|---|---|---|---|---|---|---|---|---|---|---|---|
| SAMN09742592-GCF_006382825.1 | SH100044 | SAMN09742592 | GCF_006382825.1 | NonPC | Public | 2010 | China | Clinical | 92.4 | 100 | 0.33 | 98.3082 | 5050830 | 45.26 | 23 | 603364 |
| SAMN09742593-GCF_006382765.1 | SH110082 | SAMN09742593 | GCF_006382765.1 | NonPC | Public | 2011 | China | Clinical | 92.4 | 100 | 0.05 | 98.3496 | 5085835 | 45.26 | 24 | 603400 |
| SAMN09742594-GCF_006382965.1 | 13-0386 | SAMN09742594 | GCF_006382965.1 | NonPC | Public | 2013 | China | Clinical | 92.4 | 99.86 | 0.05 | 98.3466 | 5020810 | 45.27 | 24 | 604070 |
| SAMN09742595-GCF_006382735.1 | SJ15027 | SAMN09742595 | GCF_006382735.1 | NonPC | Public | 2015 | China | Clinical | 92 | 100 | 0.19 | 98.2766 | 5059211 | 45.23 | 25 | 604057 |
| SAMN09742596-GCF_006382885.1 | QP14003 | SAMN09742596 | GCF_006382885.1 | NonPC | Public | 2014 | China | Clinical | 92.2 | 100 | 0.19 | 98.3645 | 5005638 | 45.26 | 28 | 450240 |
| SAMN09742597-GCF_006382755.1 | MH140255 | SAMN09742597 | GCF_006382755.1 | NonPC | Public | 2014 | China | Clinical | 91 | 100 | 0.27 | 98.3018 | 4998220 | 45.32 | 24 | 806519 |
| SAMN09742598-GCF_006382705.1 | F101 | SAMN09742598 | GCF_006382705.1 | NonPC | Public | 2008 | China | Clinical | 92.2 | 100 | 0.19 | 98.3548 | 5040100 | 45.29 | 27 | 603934 |
| SAMN09742599-GCF_006382695.1 | VP8-90 | SAMN09742599 | GCF_006382695.1 | NonPC | Public | 1990 | China | Clinical | 92.2 | 100 | 0.19 | 98.3677 | 5038170 | 45.29 | 25 | 603926 |
| SAMN09742602-GCF_006382925.1 | BSVP16064 | SAMN09742602 | GCF_006382925.1 | NonPC | Public | 2016 | China | Env | 91.4 | 100 | 0.03 | 98.2272 | 5142524 | 45.2 | 60 | 347079 |
| SAMN09742603-GCF_006382625.1 | BSVP16113 | SAMN09742603 | GCF_006382625.1 | NonPC | Public | 2016 | China | Env | 90.7 | 99.18 | 0.03 | 98.3216 | 5106629 | 45.24 | 44 | 312626 |
| SAMN09742604-GCF_006382645.1 | CMVP16045 | SAMN09742604 | GCF_006382645.1 | NonPC | Public | 2016 | China | Env | 90.9 | 100 | 0.14 | 98.3822 | 5090414 | 45.27 | 49 | 316034 |
| SAMN09742605-GCF_006382635.1 | VP120036 | SAMN09742605 | GCF_006382635.1 | NonPC | Public | 2012 | China | Env | 92.3 | 100 | 0.19 | 98.3661 | 5071319 | 45.24 | 27 | 749114 |
| SAMN09742665-GCF_006383415.1 | MHVP16536 | SAMN09742665 | GCF_006383415.1 | NonPC | Public | 2016 | China | Clinical | 91.9 | 100 | 0.19 | 98.2596 | 5054347 | 45.23 | 29 | 603335 |
| SAMN09745128-GCA_015804715.1 | PNUSAV000217 | SAMN09745128 | GCA_015804715.1 | NonPC | Public | NA | USA | Clinical | 91.4 | 100 | 0.71 | 98.2617 | 5144664 | 45.25 | 54 | 232261 |
| SAMN09745129-GCA_015804535.1 | PNUSAV000262 | SAMN09745129 | GCA_015804535.1 | NonPC | Public | NA | USA | Clinical | 91.4 | 100 | 4 | 98.2971 | 5244555 | 45.24 | 41 | 556708 |
| SAMN09745131-GCA_015803715.1 | PNUSAV000259 | SAMN09745131 | GCA_015803715.1 | NonPC | Public | NA | USA | Clinical | 92.1 | 100 | 4.14 | 98.3418 | 5158340 | 45.24 | 51 | 348797 |
| SAMN09745133-GCA_015804585.1 | PNUSAV000255 | SAMN09745133 | GCA_015804585.1 | PC | Public | NA | USA | Clinical | 98.2 | 100 | 2.1 | 99.9898 | 5224397 | 45.29 | 50 | 289345 |
| SAMN09745136-GCA_015804495.1 | PNUSAV000274 | SAMN09745136 | GCA_015804495.1 | NonPC | Public | NA | USA | Clinical | 91.3 | 100 | 0.07 | 98.268 | 5131479 | 45.24 | 49 | 475341 |
| SAMN09760593-GCA_015803735.1 | PNUSAV000249 | SAMN09760593 | GCA_015803735.1 | PC | Public | NA | USA | Clinical | 97.4 | 100 | 1.91 | 99.964 | 5153134 | 45.34 | 136 | 88542 |
| SAMN09760594-GCA_015804455.1 | PNUSAV000200 | SAMN09760594 | GCA_015804455.1 | NonPC | Public | NA | USA | Clinical | 90.5 | 100 | 2.18 | 98.3197 | 5122124 | 45.28 | 109 | 115529 |
| SAMN09760595-GCA_015803635.1 | PNUSAV000196 | SAMN09760595 | GCA_015803635.1 | NonPC | Public | NA | USA | Clinical | 91.4 | 100 | 4.44 | 98.2935 | 5357600 | 45.26 | 40 | 564957 |
| SAMN09768251-GCA_015802635.1 | PNUSAV000291 | SAMN09768251 | GCA_015802635.1 | NonPC | Public | NA | USA | Clinical | 89.2 | 100 | 4.9 | 98.2207 | 5256871 | 45.19 | 127 | 93982 |
| SAMN09768252-GCA_015802715.1 | PNUSAV000286 | SAMN09768252 | GCA_015802715.1 | NonPC | Public | NA | USA | Clinical | 90 | 100 | 0.07 | 98.2643 | 4972203 | 45.4 | 191 | 53027 |
| SAMN09768253-GCA_015803355.1 | PNUSAV000284 | SAMN09768253 | GCA_015803355.1 | NonPC | Public | NA | USA | Clinical | 89.9 | 100 | 0.07 | 98.3125 | 5037952 | 45.42 | 221 | 52967 |
| SAMN09768254-GCA_015803475.1 | PNUSAV000283 | SAMN09768254 | GCA_015803475.1 | PC | Public | NA | USA | Clinical | 97.4 | 100 | 0.27 | 99.9678 | 5117644 | 45.35 | 125 | 158970 |
| SAMN09783468-GCA_015803295.1 | PNUSAV000235 | SAMN09783468 | GCA_015803295.1 | NonPC | Public | NA | USA | Clinical | 90.8 | 100 | 3.69 | 98.2547 | 5282455 | 45.19 | 37 | 438811 |
| SAMN09783470-GCA_015802835.1 | PNUSAV000210 | SAMN09783470 | GCA_015802835.1 | NonPC | Public | NA | USA | Clinical | 91.1 | 100 | 0.54 | 98.2852 | 5112445 | 45.33 | 123 | 100771 |
| SAMN09783471-GCA_015803255.1 | PNUSAV000298 | SAMN09783471 | GCA_015803255.1 | NonPC | Public | NA | USA | Clinical | 90.6 | 100 | 4.51 | 98.3138 | 5248680 | 45.25 | 70 | 233704 |
| SAMN09787689-GCA_015803215.1 | PNUSAV000279 | SAMN09787689 | GCA_015803215.1 | NonPC | Public | NA | USA | Clinical | 91.1 | 100 | 2.45 | 98.3294 | 5168293 | 45.34 | 138 | 87417 |
| SAMN09787690-GCA_015803175.1 | PNUSAV000280 | SAMN09787690 | GCA_015803175.1 | NonPC | Public | NA | USA | Clinical | 91.2 | 100 | 0.42 | 98.2616 | 5135677 | 45.3 | 133 | 100771 |
| SAMN09787691-GCA_015803115.1 | PNUSAV000282 | SAMN09787691 | GCA_015803115.1 | NonPC | Public | NA | USA | Clinical | 91 | 100 | 0.14 | 98.2841 | 5112658 | 45.36 | 189 | 52927 |
| SAMN09818936-GCA_015802995.1 | PNUSAV000267 | SAMN09818936 | GCA_015802995.1 | NonPC | Public | 2018 | USA | Env | 91.3 | 100 | 1.24 | 98.291 | 5182239 | 45.23 | 46 | 244425 |
| SAMN09818964-GCA_015802915.1 | PNUSAV000265 | SAMN09818964 | GCA_015802915.1 | NonPC | Public | 2018 | USA | Env | 91.6 | 100 | 1.98 | 98.2724 | 5176858 | 45.28 | 59 | 203848 |
| SAMN09827874-GCA_015802975.1 | PNUSAV000269 | SAMN09827874 | GCA_015802975.1 | NonPC | Public | 2018 | USA | Env | 88.1 | 100 | 1.48 | 98.2477 | 5004774 | 45.35 | 75 | 137568 |
| SAMN09827888-GCA_015803015.1 | PNUSAV000268 | SAMN09827888 | GCA_015803015.1 | NonPC | Public | 2018 | USA | Env | 91.6 | 100 | 1.38 | 98.2452 | 5132133 | 45.33 | 63 | 190388 |
| SAMN09828185-GCA_015803135.1 | PNUSAV000272 | SAMN09828185 | GCA_015803135.1 | NonPC | Public | 2018 | USA | Env | 91.2 | 100 | 1.08 | 98.2853 | 5170445 | 45.19 | 87 | 230303 |
| SAMN09828187-GCA_015803155.1 | PNUSAV000270 | SAMN09828187 | GCA_015803155.1 | NonPC | Public | 2018 | USA | Env | 88.6 | 100 | 4.47 | 98.2442 | 5277402 | 45.26 | 86 | 220996 |
| SAMN09831787-GCA_015802935.1 | PNUSAV000305 | SAMN09831787 | GCA_015802935.1 | NonPC | Public | NA | USA | Clinical | 91.4 | 100 | 3.18 | 98.2663 | 5270045 | 45.27 | 73 | 198025 |
| SAMN09831788-GCA_015802895.1 | PNUSAV000304 | SAMN09831788 | GCA_015802895.1 | NonPC | Public | NA | USA | Clinical | 91.2 | 100 | 2.36 | 98.2824 | 5210725 | 45.3 | 126 | 88829 |
| SAMN09831789-GCA_015801935.1 | PNUSAV000303 | SAMN09831789 | GCA_015801935.1 | NonPC | Public | NA | USA | Clinical | 91.3 | 100 | 1.43 | 98.2898 | 5174978 | 45.26 | 65 | 195175 |
| SAMN09831790-GCA_015803095.1 | PNUSAV000288 | SAMN09831790 | GCA_015803095.1 | NonPC | Public | NA | USA | Clinical | 91.3 | 100 | 0.91 | 98.2847 | 5163944 | 45.25 | 53 | 239884 |
| SAMN09831791-GCA_015802955.1 | PNUSAV000275 | SAMN09831791 | GCA_015802955.1 | NonPC | Public | NA | USA | Clinical | 91.4 | 100 | 1.91 | 98.2835 | 5189937 | 45.24 | 39 | 556724 |
| SAMN09838635-GCA_015801835.1 | PNUSAV000332 | SAMN09838635 | GCA_015801835.1 | NonPC | Public | NA | USA | Clinical | 89 | 100 | 1.36 | 98.0855 | 5119231 | 45.36 | 133 | 82433 |
| SAMN09838636-GCA_016316105.1 | PNUSAV000302 | SAMN09838636 | GCA_016316105.1 | NonPC | Public | NA | USA | Clinical | 91.2 | 100 | 3.04 | 98.3002 | 5213666 | 45.27 | 88 | 126129 |
| SAMN09838637-GCA_015801715.1 | PNUSAV000281 | SAMN09838637 | GCA_015801715.1 | NonPC | Public | NA | USA | Clinical | 91.3 | 100 | 2.72 | 98.2791 | 5212552 | 45.26 | 54 | 204704 |
| SAMN09839615-GCA_015801235.1 | AK1822500155 | SAMN09839615 | GCA_015801235.1 | PrePC | Public | 2018 | USA | Env | 94.2 | 100 | 3 | 99.7364 | 5072778 | 45.42 | 41 | 444332 |
| SAMN09839616-GCA_015799555.1 | AK0924400197 | SAMN09839616 | GCA_015799555.1 | NonPC | Public | 2009 | USA | Env | 88.1 | 100 | 4.12 | 98.2715 | 5459017 | 44.99 | 75 | 156066 |
| SAMN09839618-GCA_015799615.1 | AK1008100229 | SAMN09839618 | GCA_015799615.1 | NonPC | Public | 2010 | USA | Env | 89.2 | 100 | 2.36 | 98.2301 | 5008538 | 45.35 | 69 | 184924 |

| | | | | | | | | | | | | | | | | |
|---|---|---|---|---|---|---|---|---|---|---|---|---|---|---|---|---|
| SAMN09839620-GCA_015800055.1 | AK1008100228 | SAMN09839620 | GCA_015800055.1 | NonPC | Public | 2010 | USA | Env | 89.2 | 100 | 3 | 98.248 | 5073597 | 45.35 | 63 | 240192 |
| SAMN09839634-GCA_015799595.1 | AK1008100226 | SAMN09839634 | GCA_015799595.1 | NonPC | Public | 2010 | USA | Env | 89.5 | 100 | 4.54 | 98.1384 | 5341267 | 45.2 | 69 | 197218 |
| SAMN09839642-GCA_015799735.1 | AK1822500158 | SAMN09839642 | GCA_015799735.1 | NonPC | Public | 2018 | USA | Env | 91.1 | 100 | 3.11 | 98.2624 | 5122235 | 45.38 | 41 | 677518 |
| SAMN09843540-GCA_018030705.1 | PNUSAV000333 | SAMN09843540 | GCA_018030705.1 | NonPC | Public | NA | USA | Clinical | 91.3 | 100 | 4.09 | 98.2999 | 5363587 | 45.25 | 97 | 117744 |
| SAMN09843542-GCA_015802675.1 | PNUSAV000318 | SAMN09843542 | GCA_015802675.1 | NonPC | Public | NA | USA | Clinical | 91.4 | 100 | 3.81 | 98.3185 | 5231820 | 45.26 | 39 | 722184 |
| SAMN09843543-GCA_015801735.1 | PNUSAV000301 | SAMN09843543 | GCA_015801735.1 | NonPC | Public | NA | USA | Clinical | 90.6 | 100 | 1.04 | 98.3324 | 5128228 | 45.2 | 61 | 410387 |
| SAMN09843544-GCA_015801875.1 | PNUSAV000300 | SAMN09843544 | GCA_015801875.1 | NonPC | Public | NA | USA | Clinical | 91.3 | 100 | 2.33 | 98.2893 | 5220884 | 45.26 | 61 | 270143 |
| SAMN09864325-GCA_015802655.1 | PNUSAV000306 | SAMN09864325 | GCA_015802655.1 | NonPC | Public | NA | USA | Clinical | 91.3 | 100 | 0.69 | 98.2951 | 5152626 | 45.24 | 72 | 183665 |
| SAMN09864335-GCA_015802475.1 | PNUSAV000313 | SAMN09864335 | GCA_015802475.1 | NonPC | Public | NA | USA | Clinical | 91.4 | 100 | 1.09 | 98.2985 | 5158991 | 45.24 | 48 | 475293 |
| SAMN09864337-GCA_016315445.1 | PNUSAV000312 | SAMN09864337 | GCA_016315445.1 | NonPC | Public | NA | USA | Clinical | 90.3 | 100 | 0.86 | 98.3201 | 5026602 | 45.27 | 38 | 521198 |
| SAMN09864368-GCA_015801635.1 | PNUSAV000340 | SAMN09864368 | GCA_015801635.1 | NonPC | Public | NA | USA | Clinical | 91.3 | 100 | 3.27 | 98.2675 | 5228258 | 45.26 | 70 | 200001 |
| SAMN09864371-GCA_015801555.1 | PNUSAV000336 | SAMN09864371 | GCA_015801555.1 | PC | Public | NA | USA | Clinical | 97.6 | 100 | 0.54 | 99.9672 | 5133780 | 45.3 | 100 | 129160 |
| SAMN09864373-GCA_015801695.1 | PNUSAV000339 | SAMN09864373 | GCA_015801695.1 | NonPC | Public | NA | USA | Clinical | 88.7 | 100 | 4.96 | 98.2824 | 5179525 | 45.25 | 56 | 245272 |
| SAMN09869694-GCA_015801335.1 | PNUSAV000349 | SAMN09869694 | GCA_015801335.1 | NonPC | Public | NA | USA | Clinical | 88.8 | 100 | 2.53 | 98.171 | 5116380 | 45.39 | 163 | 61030 |
| SAMN09869697-GCA_015801395.1 | PNUSAV000338 | SAMN09869697 | GCA_015801395.1 | NonPC | Public | NA | USA | Clinical | 90.2 | 100 | 2.45 | 98.2345 | 5136722 | 45.14 | 46 | 470066 |
| SAMN09869750-GCA_015801615.1 | PNUSAV000273 | SAMN09869750 | GCA_015801615.1 | NonPC | Public | 2018 | USA | Env | 91.3 | 100 | 0.56 | 98.3742 | 5180837 | 45.27 | 96 | 107134 |
| SAMN09874427-GCF_004006515.1 | VPD14 | SAMN09874427 | GCF_004006515.1 | PC | Public | 2012 | China | NA | 99.7 | 100 | 0 | 99.9824 | 5168640 | 45.38 | 2 | 3291121 |
| SAMN09878925-GCA_015801455.1 | 2018AW-0103 | SAMN09878925 | GCA_015801455.1 | PC | Public | NA | USA | Clinical | 97.6 | 100 | 0.61 | 99.9682 | 5152621 | 45.32 | 124 | 93893 |
| SAMN09878927-GCA_015801375.1 | 2018V-1064 | SAMN09878927 | GCA_015801375.1 | PC | Public | NA | USA | Clinical | 97.7 | 100 | 1.36 | 99.9665 | 5178379 | 45.31 | 101 | 101564 |
| SAMN09878947-GCF_003691525.1 | 160807 | SAMN09878947 | GCF_003691525.1 | NonPC | Public | 2016 | China | Env | 90.8 | 100 | 0.19 | 98.225 | 5396107 | 45.24 | 3 | 3508313 |
| SAMN09917440-GCA_015801155.1 | PNUSAV000347 | SAMN09917440 | GCA_015801155.1 | NonPC | Public | NA | USA | Clinical | 91.4 | 100 | 0.87 | 98.298 | 5215543 | 45.26 | 46 | 475225 |
| SAMN09917443-GCA_015800035.1 | PNUSAV000356 | SAMN09917443 | GCA_015800035.1 | PC | Public | NA | USA | Clinical | 97.4 | 100 | 1.02 | 99.9667 | 5159965 | 45.31 | 88 | 154479 |
| SAMN09917444-GCA_015801215.1 | PNUSAV000355 | SAMN09917444 | GCA_015801215.1 | NonPC | Public | NA | USA | Clinical | 90.6 | 100 | 0.89 | 98.3038 | 5126080 | 45.28 | 83 | 191687 |
| SAMN09917445-GCA_015801035.1 | PNUSAV000350 | SAMN09917445 | GCA_015801035.1 | NonPC | Public | NA | USA | Clinical | 91.1 | 100 | 1.36 | 98.2789 | 5194236 | 45.31 | 121 | 85188 |
| SAMN09917448-GCA_015803455.1 | PNUSAV000334 | SAMN09917448 | GCA_015803455.1 | NonPC | Public | NA | USA | Clinical | 91.3 | 100 | 0.73 | 98.2867 | 5192298 | 45.29 | 91 | 113953 |
| SAMN09927918-GCA_015801095.1 | PNUSAV000374 | SAMN09927918 | GCA_015801095.1 | NonPC | Public | NA | USA | Clinical | 91.4 | 100 | 0.34 | 98.2878 | 5134387 | 45.25 | 61 | 218257 |
| SAMN09927919-GCA_015801295.1 | PNUSAV000360 | SAMN09927919 | GCA_015801295.1 | NonPC | Public | NA | USA | Clinical | 91.4 | 100 | 4.16 | 98.2962 | 5312649 | 45.25 | 41 | 564960 |
| SAMN09937954-GCA_015801115.1 | PNUSAV000379 | SAMN09937954 | GCA_015801115.1 | NonPC | Public | NA | USA | Clinical | 89.4 | 100 | 2.86 | 98.2553 | 5238714 | 45.18 | 58 | 227439 |
| SAMN09937955-GCA_015799975.1 | PNUSAV000378 | SAMN09937955 | GCA_015799975.1 | NonPC | Public | NA | USA | Clinical | 90.6 | 100 | 3.27 | 98.266 | 5208947 | 45.23 | 38 | 436245 |
| SAMN09937956-GCA_015799935.1 | PNUSAV000377 | SAMN09937956 | GCA_015799935.1 | NonPC | Public | NA | USA | Clinical | 90.7 | 100 | 4.09 | 98.3135 | 5280339 | 45.25 | 40 | 342337 |
| SAMN09937957-GCA_015800715.1 | PNUSAV000368 | SAMN09937957 | GCA_015800715.1 | NonPC | Public | NA | USA | Clinical | 91.4 | 100 | 0.82 | 98.2838 | 5162150 | 45.24 | 52 | 343094 |
| SAMN09937960-GCA_015801175.1 | PNUSAV000367 | SAMN09937960 | GCA_015801175.1 | NonPC | Public | NA | USA | Clinical | 90.8 | 100 | 0.64 | 98.3433 | 5205760 | 45.27 | 24 | 447421 |
| SAMN09951148-GCA_015801315.1 | PNUSAV000371 | SAMN09951148 | GCA_015801315.1 | NonPC | Public | NA | USA | Clinical | 91.3 | 100 | 1.91 | 98.294 | 5196230 | 45.25 | 49 | 344611 |
| SAMN09951150-GCA_015800015.1 | PNUSAV000370 | SAMN09951150 | GCA_015800015.1 | NonPC | Public | NA | USA | Clinical | 91.3 | 100 | 4.09 | 98.3303 | 5267844 | 45.25 | 45 | 404526 |
| SAMN09951153-GCA_015800975.1 | PNUSAV000358 | SAMN09951153 | GCA_015800975.1 | NonPC | Public | NA | USA | Clinical | 91.4 | 100 | 1.84 | 98.3702 | 5229305 | 45.17 | 37 | 378165 |
| SAMN09984749-GCA_015799915.1 | PNUSAV000351 | SAMN09984749 | GCA_015799915.1 | NonPC | Public | NA | USA | Clinical | 91.4 | 100 | 3.68 | 98.2838 | 5232405 | 45.24 | 37 | 556708 |
| SAMN09984756-GCA_015801255.1 | PNUSAV000383 | SAMN09984756 | GCA_015801255.1 | NonPC | Public | NA | USA | Clinical | 91.4 | 100 | 1.5 | 98.3233 | 5156681 | 45.26 | 38 | 548329 |
| SAMN09984757-GCA_015799815.1 | PNUSAV000365 | SAMN09984757 | GCA_015799815.1 | NonPC | Public | NA | USA | Clinical | 91.4 | 100 | 4.56 | 98.2926 | 5270967 | 45.25 | 41 | 565044 |
| SAMN09984758-GCA_015799895.1 | PNUSAV000364 | SAMN09984758 | GCA_015799895.1 | NonPC | Public | NA | USA | Clinical | 89.2 | 100 | 2.53 | 98.072 | 5140362 | 45.27 | 65 | 233226 |
| SAMN09984857-GCA_015799755.1 | 2018V-1124 | SAMN09984857 | GCA_015799755.1 | NonPC | Public | NA | USA | Clinical | 91.3 | 100 | 0.2 | 98.2977 | 5128465 | 45.25 | 77 | 179082 |
| SAMN10033017-GCA_015800275.1 | PNUSAV000392 | SAMN10033017 | GCA_015800275.1 | NonPC | Public | NA | USA | Clinical | 91.4 | 100 | 0.36 | 98.296 | 5159357 | 45.26 | 41 | 556708 |
| SAMN10033018-GCA_015800355.1 | PNUSAV000391 | SAMN10033018 | GCA_015800355.1 | NonPC | Public | NA | USA | Clinical | 91.4 | 100 | 3.22 | 98.2717 | 5263787 | 45.25 | 45 | 475325 |
| SAMN10033019-GCA_016305405.1 | PNUSAV000389 | SAMN10033019 | GCA_016305405.1 | NonPC | Public | NA | USA | Clinical | 91.4 | 100 | 3 | 98.2902 | 5231233 | 45.25 | 39 | 556698 |
| SAMN10033020-GCA_015799695.1 | PNUSAV000388 | SAMN10033020 | GCA_015799695.1 | NonPC | Public | NA | USA | Clinical | 91.4 | 100 | 4.42 | 98.3051 | 5281496 | 45.26 | 41 | 556714 |
| SAMN10033021-GCA_015799715.1 | PNUSAV000346 | SAMN10033021 | GCA_015799715.1 | NonPC | Public | NA | USA | Clinical | 91.4 | 100 | 2.38 | 98.276 | 5227377 | 45.25 | 51 | 221918 |
| SAMN10033023-GCA_015800215.1 | PNUSAV000403 | SAMN10033023 | GCA_015800215.1 | NonPC | Public | NA | USA | Clinical | 91.4 | 100 | 0.89 | 98.3009 | 5177333 | 45.24 | 43 | 556587 |
| SAMN10033024-GCA_015800725.1 | PNUSAV000402 | SAMN10033024 | GCA_015800725.1 | NonPC | Public | NA | USA | Clinical | 90.6 | 100 | 1.47 | 98.2682 | 5232353 | 45.19 | 52 | 536977 |
| SAMN10033025-GCA_015800655.1 | PNUSAV000399 | SAMN10033025 | GCA_015800655.1 | NonPC | Public | NA | USA | Clinical | 91.3 | 100 | 0.35 | 98.3298 | 5141145 | 45.24 | 44 | 556700 |

| | | | | | | | | | | | | | | | | |
|---|---|---|---|---|---|---|---|---|---|---|---|---|---|---|---|---|
| SAMN10033028-GCA_015800235.1 | PNUSAV000397 | SAMN10033028 | GCA_015800235.1 | NonPC | Public | NA | USA | Clinical | 91.4 | 100 | 4.09 | 98.2902 | 5258071 | 45.25 | 42 | 475310 |
| SAMN10033031-GCA_015800295.1 | PNUSAV000394 | SAMN10033031 | GCA_015800295.1 | NonPC | Public | NA | USA | Clinical | 91.4 | 100 | 3.13 | 98.282 | 5235933 | 45.25 | 40 | 556708 |
| SAMN10033032-GCA_015800615.1 | PNUSAV000393 | SAMN10033032 | GCA_015800615.1 | NonPC | Public | NA | USA | Clinical | 91.4 | 100 | 3.95 | 98.3083 | 5286585 | 45.25 | 39 | 718992 |
| SAMN10037990-GCA_015800195.1 | PNUSAV000278 | SAMN10037990 | GCA_015800195.1 | NonPC | Public | NA | USA | Clinical | 91.1 | 100 | 2.53 | 98.2818 | 5201495 | 45.31 | 123 | 100878 |
| SAMN10038050-GCA_016303985.1 | PNUSAV000277 | SAMN10038050 | GCA_016303985.1 | NonPC | Public | NA | USA | Clinical | 91.3 | 100 | 2.82 | 98.3184 | 5212380 | 45.25 | 56 | 197329 |
| SAMN10038115-GCA_015800895.1 | PNUSAV000417 | SAMN10038115 | GCA_015800895.1 | NonPC | Public | NA | USA | Clinical | 91.5 | 100 | 1.78 | 98.3379 | 5253809 | 45.26 | 107 | 108422 |
| SAMN10038121-GCA_015800175.1 | PNUSAV000411 | SAMN10038121 | GCA_015800175.1 | NonPC | Public | NA | USA | Clinical | 91 | 100 | 3.53 | 98.2848 | 5233460 | 45.26 | 60 | 198526 |
| SAMN10038139-GCA_015800315.1 | PNUSAV000276 | SAMN10038139 | GCA_015800315.1 | NonPC | Public | NA | USA | Clinical | 91.4 | 100 | 3.54 | 98.2866 | 5269095 | 45.26 | 42 | 475296 |
| SAMN10038163-GCA_015800155.1 | PNUSAV000419 | SAMN10038163 | GCA_015800155.1 | NonPC | Public | NA | USA | Clinical | 91.4 | 100 | 2.52 | 98.4088 | 5272786 | 45.14 | 48 | 293226 |
| SAMN10038166-GCA_015800075.1 | PNUSAV000418 | SAMN10038166 | GCA_015800075.1 | NonPC | Public | NA | USA | Clinical | 91.9 | 100 | 0.74 | 98.3715 | 5192245 | 45.18 | 144 | 100699 |
| SAMN10038179-GCA_015800095.1 | PNUSAV000416 | SAMN10038179 | GCA_015800095.1 | NonPC | Public | NA | USA | Clinical | 89.8 | 100 | 3.41 | 98.2946 | 5248095 | 45.21 | 44 | 196761 |
| SAMN10056417-GCA_015800995.1 | PNUSAV000412 | SAMN10056417 | GCA_015800995.1 | NonPC | Public | NA | USA | Clinical | 91.2 | 100 | 1.36 | 98.2863 | 5180996 | 45.3 | 112 | 107966 |
| SAMN10061638-GCA_015800115.1 | PNUSAV000405 | SAMN10061638 | GCA_015800115.1 | NonPC | Public | NA | USA | Clinical | 91.3 | 100 | 0.27 | 98.3064 | 5194609 | 45.24 | 77 | 140362 |
| SAMN10061641-GCA_015800805.1 | PNUSAV000404 | SAMN10061641 | GCA_015800805.1 | NonPC | Public | NA | USA | Clinical | 91.3 | 100 | 0.07 | 98.3188 | 5127698 | 45.25 | 71 | 201804 |
| SAMN10076462-GCA_004324645.1 | V146 | SAMN10076462 | GCA_004324645.1 | NonPC | Public | 2016 | China | Env | 89.8 | 99.51 | 0.27 | 98.3398 | 5042453 | 45.29 | 58 | 385816 |
| SAMN10076535-GCA_004324805.1 | V147 | SAMN10076535 | GCA_004324805.1 | NonPC | Public | 2016 | China | Env | 91.4 | 98.18 | 0.42 | 98.3129 | 5220515 | 45.19 | 48 | 741792 |
| SAMN10076539-GCA_004324745.1 | V148 | SAMN10076539 | GCA_004324745.1 | NonPC | Public | 2016 | China | Env | 89.5 | 98.91 | 0.41 | 98.2021 | 5047540 | 45.31 | 46 | 767598 |
| SAMN10076540-GCA_004324735.1 | V187 | SAMN10076540 | GCA_004324735.1 | NonPC | Public | 2016 | China | Env | 87.6 | 98.65 | 1.24 | 98.0965 | 5081195 | 45.23 | 73 | 546050 |
| SAMN10076541-GCA_004324625.1 | V209 | SAMN10076541 | GCA_004324625.1 | NonPC | Public | 2015 | China | Env | 87.6 | 99.9 | 0.67 | 98.1067 | 5071841 | 45.23 | 64 | 535378 |
| SAMN10076542-GCA_004324525.1 | Vp10 | SAMN10076542 | GCA_004324525.1 | NonPC | Public | 2012 | China | Env | 91.4 | 99.15 | 0.73 | 98.3675 | 5004554 | 45.33 | 40 | 862363 |
| SAMN10076543-GCA_004324515.1 | Vp12 | SAMN10076543 | GCA_004324515.1 | NonPC | Public | 2012 | China | Env | 89.8 | 99.73 | 0.33 | 98.3007 | 4975492 | 45.31 | 55 | 455720 |
| SAMN10076544-GCA_004324505.1 | Vp2 | SAMN10076544 | GCA_004324505.1 | NonPC | Public | 2012 | China | Env | 88.3 | 99.32 | 0.36 | 98.2126 | 5133301 | 45.16 | 52 | 550410 |
| SAMN10076545-GCF_004324495.1 | Vp23 | SAMN10076545 | GCF_004324495.1 | NonPC | Public | 2013 | China | Env | 91.2 | 99.93 | 0.3 | 98.2094 | 5368262 | 45.23 | 140 | 250001 |
| SAMN10076547-GCA_004324435.1 | Vp28 | SAMN10076547 | GCA_004324435.1 | NonPC | Public | 2013 | China | Env | 91.6 | 98.85 | 0.37 | 98.252 | 5212325 | 45.23 | 58 | 325378 |
| SAMN10076548-GCA_004324675.1 | Vp3 | SAMN10076548 | GCA_004324675.1 | NonPC | Public | 2012 | China | Env | 91.7 | 99.3 | 0.33 | 98.2618 | 5112726 | 45.3 | 62 | 282892 |
| SAMN10076549-GCA_004324595.1 | Vp30 | SAMN10076549 | GCA_004324595.1 | NonPC | Public | 2013 | China | Env | 89.9 | 98.57 | 0.1 | 98.2439 | 5076869 | 45.34 | 55 | 499220 |
| SAMN10076550-GCA_004324425.1 | Vp34 | SAMN10076550 | GCA_004324425.1 | NonPC | Public | 2013 | China | Env | 89.7 | 99.1 | 0.05 | 98.3494 | 4936940 | 45.38 | 45 | 376913 |
| SAMN10076551-GCA_004324415.1 | Vp38 | SAMN10076551 | GCA_004324415.1 | NonPC | Public | 2013 | China | Env | 91.1 | 99.32 | 0.66 | 98.2926 | 5003788 | 45.31 | 56 | 416191 |
| SAMN10076552-GCA_004324575.1 | Vp43 | SAMN10076552 | GCA_004324575.1 | NonPC | Public | 2013 | China | Env | 88.2 | 99.08 | 0.76 | 98.2568 | 4973694 | 45.32 | 61 | 465069 |
| SAMN10076553-GCA_004324405.1 | Vp45 | SAMN10076553 | GCA_004324405.1 | NonPC | Public | 2013 | China | Env | 88.5 | 99.32 | 0.86 | 98.1505 | 5470146 | 45.03 | 98 | 327169 |
| SAMN10076554-GCA_004324395.1 | Vp46 | SAMN10076554 | GCA_004324395.1 | NonPC | Public | 2013 | China | Env | 92 | 98.92 | 0.78 | 98.316 | 5181815 | 45.26 | 87 | 399220 |
| SAMN10076555-GCF_004324655.1 | 743 | SAMN10076555 | GCF_004324655.1 | NonPC | Public | 2012 | China | Env | 89.7 | 99.3 | 0.54 | 98.1584 | 5325136 | 45.09 | 57 | 458770 |
| SAMN10076556-GCA_004324555.1 | Vp9 | SAMN10076556 | GCA_004324555.1 | NonPC | Public | 2012 | China | Env | 91.4 | 98.62 | 0.48 | 98.3371 | 5007318 | 45.34 | 46 | 386651 |
| SAMN10077887-GCA_015800855.1 | PNUSAV000434 | SAMN10077887 | GCA_015800855.1 | NonPC | Public | NA | USA | Clinical | 91 | 100 | 0.36 | 98.2858 | 5128078 | 45.34 | 161 | 75612 |
| SAMN10077891-GCA_015800955.1 | PNUSAV000433 | SAMN10077891 | GCA_015800955.1 | NonPC | Public | NA | USA | Clinical | 86.6 | 100 | 0.37 | 98.0956 | 4920172 | 45.58 | 314 | 25841 |
| SAMN10079516-GCA_015801015.1 | PNUSAV000431 | SAMN10079516 | GCA_015801015.1 | NonPC | Public | NA | USA | Clinical | 91.3 | 100 | 3.54 | 98.3142 | 5254311 | 45.26 | 95 | 117698 |
| SAMN10090358-GCA_015800395.1 | PNUSAV000429 | SAMN10090358 | GCA_015800395.1 | NonPC | Public | NA | USA | Clinical | 89.6 | 100 | 0.24 | 98.2204 | 5173669 | 45.23 | 138 | 115692 |
| SAMN10090359-GCA_015800455.1 | PNUSAV000421 | SAMN10090359 | GCA_015800455.1 | NonPC | Public | NA | USA | Clinical | 91.4 | 100 | 4.57 | 98.294 | 5308828 | 45.25 | 42 | 556713 |
| SAMN10090360-GCA_015800515.1 | PNUSAV000414 | SAMN10090360 | GCA_015800515.1 | NonPC | Public | NA | USA | Clinical | 91.1 | 100 | 1.16 | 98.2753 | 5207231 | 45.3 | 146 | 82130 |
| SAMN10090361-GCA_015800595.1 | PNUSAV000410 | SAMN10090361 | GCA_015800595.1 | NonPC | Public | NA | USA | Clinical | 90.3 | 100 | 2.6 | 98.3139 | 5178774 | 45.32 | 198 | 62589 |
| SAMN10090362-GCA_015800335.1 | PNUSAV000409 | SAMN10090362 | GCA_015800335.1 | NonPC | Public | NA | USA | Clinical | 91.1 | 100 | 1.63 | 98.2985 | 5165914 | 45.36 | 141 | 80341 |
| SAMN10090363-GCA_015800575.1 | PNUSAV000408 | SAMN10090363 | GCA_015800575.1 | NonPC | Public | NA | USA | Clinical | 91.1 | 100 | 2.45 | 98.2597 | 5193786 | 45.32 | 114 | 100771 |
| SAMN10090364-GCA_015800675.1 | PNUSAV000407 | SAMN10090364 | GCA_015800675.1 | NonPC | Public | NA | USA | Clinical | 91.2 | 100 | 2.72 | 98.2744 | 5206626 | 45.3 | 105 | 101013 |
| SAMN10090365-GCA_015800875.1 | PNUSAV000385 | SAMN10090365 | GCA_015800875.1 | NonPC | Public | NA | USA | Clinical | 91.5 | 100 | 2.45 | 98.356 | 5068719 | 45.41 | 83 | 139678 |
| SAMN10090366-GCA_015800415.1 | PNUSAV000375 | SAMN10090366 | GCA_015800415.1 | NonPC | Public | NA | USA | Clinical | 90.4 | 100 | 0.42 | 98.2111 | 5241408 | 45.22 | 134 | 95588 |
| SAMN10095495-GCA_015800555.1 | PNUSAV000440 | SAMN10095495 | GCA_015800555.1 | PC | Public | NA | USA | Clinical | 98.2 | 100 | 4.63 | 99.9898 | 5246127 | 45.31 | 82 | 140997 |
| SAMN10095496-GCA_016303805.1 | PNUSAV000439 | SAMN10095496 | GCA_016303805.1 | PC | Public | NA | USA | Clinical | 97.7 | 100 | 2.52 | 99.9872 | 5227464 | 45.39 | 156 | 75679 |
| SAMN10095498-GCA_015800795.1 | PNUSAV000459 | SAMN10095498 | GCA_015800795.1 | NonPC | Public | NA | USA | Clinical | 91.4 | 100 | 2.09 | 98.2917 | 5266982 | 45.24 | 41 | 475354 |

| | | | | | | | | | | | | | | | | |
|---|---|---|---|---|---|---|---|---|---|---|---|---|---|---|---|---|
| SAMN10095500-GCA_015800435.1 | PNUSAV000446 | SAMN10095500 | GCA_015800435.1 | PC | Public | NA | USA | Clinical | 98.1 | 100 | 2.09 | 99.9853 | 5186584 | 45.27 | 102 | 114341 |
| SAMN10095503-GCA_015800775.1 | PNUSAV000443 | SAMN10095503 | GCA_015800775.1 | PC | Public | NA | USA | Clinical | 98.3 | 100 | 4.36 | 99.9911 | 5298862 | 45.26 | 71 | 177461 |
| SAMN10095504-GCA_015800755.1 | PNUSAV000442 | SAMN10095504 | GCA_015800755.1 | PC | Public | NA | USA | Clinical | 98.2 | 100 | 4.36 | 99.9949 | 5342921 | 45.28 | 73 | 177502 |
| SAMN10095505-GCA_015800495.1 | PNUSAV000441 | SAMN10095505 | GCA_015800495.1 | PC | Public | NA | USA | Clinical | 98 | 100 | 4.09 | 99.9837 | 5227571 | 45.34 | 107 | 116258 |
| SAMN10096183-GCA_015799535.1 | PNUSAV000452 | SAMN10096183 | GCA_015799535.1 | NonPC | Public | NA | USA | Clinical | 90.8 | 100 | 1.04 | 98.261 | 5260262 | 45.26 | 81 | 130219 |
| SAMN10096184-GCA_015799575.1 | PNUSAV000449 | SAMN10096184 | GCA_015799575.1 | NonPC | Public | NA | USA | Clinical | 89.6 | 100 | 0.58 | 98.2554 | 5201659 | 45.2 | 115 | 111121 |
| SAMN10096185-GCA_015799635.1 | PNUSAV000448 | SAMN10096185 | GCA_015799635.1 | NonPC | Public | NA | USA | Clinical | 90.7 | 100 | 0.22 | 98.3081 | 5096188 | 45.23 | 106 | 100825 |
| SAMN10111973-GCA_015798735.1 | PNUSAV000425 | SAMN10111973 | GCA_015798735.1 | NonPC | Public | NA | USA | Clinical | 91.4 | 100 | 3.81 | 98.2892 | 5266308 | 45.26 | 44 | 354363 |
| SAMN10111974-GCA_015798655.1 | PNUSAV000424 | SAMN10111974 | GCA_015798655.1 | NonPC | Public | NA | USA | Clinical | 89.1 | 100 | 3.27 | 98.0902 | 5190307 | 45.33 | 65 | 233229 |
| SAMN10111975-GCA_015799455.1 | PNUSAV000423 | SAMN10111975 | GCA_015799455.1 | NonPC | Public | NA | USA | Clinical | 91.4 | 100 | 4.36 | 98.3014 | 5365498 | 45.26 | 42 | 286904 |
| SAMN10111977-GCA_015798795.1 | PNUSAV000422 | SAMN10111977 | GCA_015798795.1 | NonPC | Public | NA | USA | Clinical | 91.4 | 100 | 3 | 98.2897 | 5226853 | 45.25 | 42 | 475325 |
| SAMN10111979-GCA_015799495.1 | PNUSAV000415 | SAMN10111979 | GCA_015799495.1 | NonPC | Public | NA | USA | Clinical | 90 | 100 | 0.4 | 98.3001 | 5054082 | 45.32 | 56 | 436767 |
| SAMN10124231-GCA_015799355.1 | PNUSAV000335 | SAMN10124231 | GCA_015799355.1 | NonPC | Public | NA | USA | Clinical | 91.1 | 100 | 0.91 | 98.2898 | 5137583 | 45.3 | 123 | 88212 |
| SAMN10127616-GCA_015798775.1 | PNUSAV000466 | SAMN10127616 | GCA_015798775.1 | NonPC | Public | NA | USA | Clinical | 91.2 | 100 | 4.63 | 98.3186 | 5266855 | 45.3 | 99 | 114552 |
| SAMN10127617-GCA_015798675.1 | PNUSAV000437 | SAMN10127617 | GCA_015798675.1 | NonPC | Public | NA | USA | Clinical | 90.2 | 100 | 0.94 | 98.2557 | 5056323 | 45.34 | 161 | 76356 |
| SAMN10127618-GCA_015799475.1 | PNUSAV000436 | SAMN10127618 | GCA_015799475.1 | NonPC | Public | NA | USA | Clinical | 91 | 100 | 0.28 | 98.32 | 5076678 | 45.37 | 173 | 68582 |
| SAMN10127619-GCA_015798835.1 | PNUSAV000413 | SAMN10127619 | GCA_015798835.1 | NonPC | Public | NA | USA | Clinical | 91.3 | 100 | 3.41 | 98.252 | 5251963 | 45.29 | 96 | 113825 |
| SAMN10139624-GCA_015799435.1 | PNUSAV000432 | SAMN10139624 | GCA_015799435.1 | NonPC | Public | NA | USA | Clinical | 91.3 | 100 | 1.27 | 98.2856 | 5167151 | 45.25 | 62 | 218348 |
| SAMN10142585-GCA_015799395.1 | PNUSAV000465 | SAMN10142585 | GCA_015799395.1 | NonPC | Public | NA | USA | Clinical | 90.6 | 100 | 3.27 | 98.3137 | 5201646 | 45.26 | 84 | 191227 |
| SAMN10142586-GCA_015799375.1 | PNUSAV000463 | SAMN10142586 | GCA_015799375.1 | NonPC | Public | NA | USA | Clinical | 90.6 | 100 | 2.23 | 98.3678 | 5060079 | 45.47 | 126 | 128120 |
| SAMN10148613-GCA_015799255.1 | PNUSAV000455 | SAMN10148613 | GCA_015799255.1 | PC | Public | NA | USA | Clinical | 98.2 | 100 | 3.27 | 99.9911 | 5219152 | 45.29 | 75 | 144236 |
| SAMN10148616-GCA_015799335.1 | PNUSAV000430 | SAMN10148616 | GCA_015799335.1 | NonPC | Public | NA | USA | Clinical | 91.3 | 100 | 2.45 | 98.2764 | 5206633 | 45.26 | 54 | 238718 |
| SAMN10163150-GCA_015799235.1 | PNUSAV000480 | SAMN10163150 | GCA_015799235.1 | PC | Public | NA | USA | Clinical | 97.8 | 100 | 2.18 | 99.9704 | 5217485 | 45.29 | 75 | 199451 |
| SAMN10163151-GCA_015799215.1 | PNUSAV000479 | SAMN10163151 | GCA_015799215.1 | PC | Public | NA | USA | Clinical | 97.7 | 100 | 2.72 | 99.9656 | 5217809 | 45.29 | 83 | 160161 |
| SAMN10163154-GCA_015799155.1 | PNUSAV000474 | SAMN10163154 | GCA_015799155.1 | NonPC | Public | NA | USA | Clinical | 90.4 | 100 | 2.63 | 98.2701 | 5189349 | 45.26 | 32 | 534682 |
| SAMN10180436-GCA_015799135.1 | PNUSAV000382 | SAMN10180436 | GCA_015799135.1 | NonPC | Public | NA | USA | Clinical | 91.3 | 100 | 2.79 | 98.2811 | 5235773 | 45.27 | 64 | 204818 |
| SAMN10181216-GCA_015799015.1 | PNUSAV000401 | SAMN10181216 | GCA_015799015.1 | NonPC | Public | NA | USA | Clinical | 89.2 | 100 | 4.13 | 98.0481 | 5383425 | 45.1 | 78 | 207092 |
| SAMN10181217-GCA_015799075.1 | PNUSAV000400 | SAMN10181217 | GCA_015799075.1 | NonPC | Public | NA | USA | Clinical | 91.3 | 100 | 3.27 | 98.3095 | 5241952 | 45.28 | 80 | 123316 |
| SAMN10221803-GCA_015798915.1 | PNUSAV000482 | SAMN10221803 | GCA_015798915.1 | NonPC | Public | NA | USA | Clinical | 91.1 | 100 | 2.18 | 98.3059 | 5161848 | 45.33 | 140 | 85995 |
| SAMN10221805-GCA_015798995.1 | PNUSAV000481 | SAMN10221805 | GCA_015798995.1 | PC | Public | NA | USA | Clinical | 96.9 | 100 | 0.61 | 99.9848 | 5050653 | 45.45 | 266 | 38857 |
| SAMN10221808-GCA_015798935.1 | PNUSAV000478 | SAMN10221808 | GCA_015798935.1 | NonPC | Public | NA | USA | Clinical | 91.4 | 100 | 3.72 | 98.2866 | 5247442 | 45.25 | 42 | 351374 |
| SAMN10221812-GCA_015798955.1 | PNUSAV000477 | SAMN10221812 | GCA_015798955.1 | NonPC | Public | NA | USA | Clinical | 91.4 | 100 | 4.09 | 98.3185 | 5279304 | 45.28 | 41 | 461768 |
| SAMN10221815-GCA_015798875.1 | PNUSAV000476 | SAMN10221815 | GCA_015798875.1 | NonPC | Public | NA | USA | Clinical | 88.8 | 100 | 3.79 | 98.1475 | 5204283 | 45.18 | 51 | 195114 |
| SAMN10221819-GCA_015798895.1 | PNUSAV000475 | SAMN10221819 | GCA_015798895.1 | NonPC | Public | NA | USA | Clinical | 91.4 | 100 | 3 | 98.2885 | 5224982 | 45.25 | 39 | 467082 |
| SAMN10221821-GCA_015798855.1 | PNUSAV000470 | SAMN10221821 | GCA_015798855.1 | NonPC | Public | NA | USA | Clinical | 90.7 | 100 | 4.7 | 98.3548 | 5466398 | 45.29 | 29 | 375489 |
| SAMN10221826-GCA_015799035.1 | PNUSAV000469 | SAMN10221826 | GCA_015799035.1 | NonPC | Public | NA | USA | Clinical | 91.4 | 100 | 4.09 | 98.3004 | 5256885 | 45.26 | 38 | 807987 |
| SAMN10221842-GCA_015798975.1 | PNUSAV000492 | SAMN10221842 | GCA_015798975.1 | NonPC | Public | NA | USA | Clinical | 91.2 | 100 | 3.35 | 98.2766 | 5324500 | 45.14 | 85 | 176965 |
| SAMN10221845-GCA_015798635.1 | PNUSAV000491 | SAMN10221845 | GCA_015798635.1 | NonPC | Public | NA | USA | Clinical | 91.4 | 100 | 3.54 | 98.3526 | 5263493 | 45.17 | 43 | 264245 |
| SAMN10221847-GCA_015799095.1 | PNUSAV000483 | SAMN10221847 | GCA_015799095.1 | NonPC | Public | NA | USA | Clinical | 89.6 | 100 | 3.76 | 98.2741 | 5268457 | 45.28 | 130 | 74191 |
| SAMN10221860-GCA_015798515.1 | PNUSAV000488 | SAMN10221860 | GCA_015798515.1 | NonPC | Public | NA | USA | Clinical | 88.9 | 100 | 0.36 | 98.2924 | 5214466 | 45.35 | 357 | 30740 |
| SAMN10227695-GCA_015798555.1 | 2018V-1114 | SAMN10227695 | GCA_015798555.1 | NonPC | Public | NA | USA | Clinical | 91.3 | 100 | 4.5 | 98.2859 | 5262126 | 45.26 | 54 | 204873 |
| SAMN10227696-GCA_015796675.1 | PNUSAV000460 | SAMN10227696 | GCA_015796675.1 | NonPC | Public | NA | USA | Clinical | 91.3 | 100 | 4.09 | 98.2788 | 5275136 | 45.27 | 51 | 414513 |
| SAMN10227697-GCA_016301505.1 | PNUSAV000458 | SAMN10227697 | GCA_016301505.1 | NonPC | Public | NA | USA | Clinical | 90.8 | 100 | 2.78 | 98.2941 | 5273993 | 45.2 | 53 | 331689 |
| SAMN10227698-GCA_015796475.1 | PNUSAV000457 | SAMN10227698 | GCA_015796475.1 | NonPC | Public | NA | USA | Clinical | 90.8 | 100 | 3.94 | 98.3073 | 5363449 | 45.19 | 52 | 365265 |
| SAMN10227700-GCA_015796975.1 | PNUSAV000359 | SAMN10227700 | GCA_015796975.1 | NonPC | Public | NA | USA | Clinical | 91.2 | 100 | 1.99 | 98.3217 | 5183337 | 45.28 | 89 | 122508 |
| SAMN10231873-GCA_015796955.1 | PNUSAV000493 | SAMN10231873 | GCA_015796955.1 | NonPC | Public | NA | USA | Clinical | 90.6 | 100 | 0.82 | 98.2964 | 5111300 | 45.3 | 98 | 159071 |
| SAMN10233644-GCA_015778585.1 | YK38 | SAMN10233644 | GCA_015778585.1 | NonPC | Public | 2017 | China | Env | 91.7 | 100 | 3.72 | 98.3123 | 5678488 | 45.13 | 65 | 401250 |
| SAMN10239531-SRR8892836 | YK13 | SAMN10239531 | SRR8892836 | NonPC | Public | 2017 | China | Env | 88.5 | 100 | 0.33 | 98.2224 | 5133596 | 45.21 | 52 | 401987 |

| | | | | | | | | | | | | | | | | |
|---|---|---|---|---|---|---|---|---|---|---|---|---|---|---|---|---|
| SAMN10240051-SRR8892853 | YK33 | SAMN10240051 | SRR8892853 | NonPC | Public | 2017 | China | Env | 89.8 | 100 | 0.05 | 98.2923 | 5117467 | 45.28 | 54 | 450391 |
| SAMN10240085-SRR8892843 | PJ44 | SAMN10240085 | SRR8892843 | NonPC | Public | 2017 | China | Env | 91.8 | 100 | 0.27 | 98.3032 | 5386856 | 45.13 | 78 | 421761 |
| SAMN10240147-GCA_015778625.1 | PJ35 | SAMN10240147 | GCA_015778625.1 | PC | Public | 2017 | China | Env | 98.1 | 100 | 3.03 | 99.9839 | 5360837 | 45.27 | 52 | 394944 |
| SAMN10250995-GCA_015796795.1 | PNUSAV000499 | SAMN10250995 | GCA_015796795.1 | NonPC | Public | NA | USA | Clinical | 91.1 | 100 | 3.85 | 98.3386 | 5193824 | 45.34 | 90 | 114167 |
| SAMN10255399-SRR8079360 | YK-17 | SAMN10255399 | SRR8079360 | NonPC | Public | 2017 | China | Env | 89.9 | 100 | 0.15 | 98.2573 | 5156218 | 45.34 | 46 | 460053 |
| SAMN10255400-SRR8079362 | YK-33 | SAMN10255400 | SRR8079362 | NonPC | Public | 2017 | China | Env | 89.8 | 100 | 0.05 | 98.2816 | 5117467 | 45.28 | 54 | 450391 |
| SAMN10256549-GCA_015796495.1 | PNUSAV000512 | SAMN10256549 | GCA_015796495.1 | NonPC | Public | NA | USA | Clinical | 90.7 | 100 | 4.2 | 98.2461 | 5401122 | 45.18 | 80 | 153939 |
| SAMN10256550-GCA_015796775.1 | PNUSAV000505 | SAMN10256550 | GCA_015796775.1 | NonPC | Public | NA | USA | Clinical | 90.8 | 100 | 3.51 | 98.3444 | 5287049 | 45.21 | 74 | 175046 |
| SAMN10256551-GCA_015796875.1 | PNUSAV000503 | SAMN10256551 | GCA_015796875.1 | NonPC | Public | NA | USA | Clinical | 91.4 | 100 | 4.13 | 98.3259 | 5206931 | 45.3 | 50 | 319804 |
| SAMN10266904-GCA_015798455.1 | PNUSAV000516 | SAMN10266904 | GCA_015798455.1 | NonPC | Public | NA | USA | Clinical | 91 | 100 | 0.36 | 98.3127 | 5108860 | 45.36 | 199 | 59220 |
| SAMN10266905-GCA_015798245.1 | PNUSAV000515 | SAMN10266905 | GCA_015798245.1 | NonPC | Public | NA | USA | Clinical | 91.4 | 100 | 4.63 | 98.3187 | 5298873 | 45.26 | 36 | 722184 |
| SAMN10266906-GCA_015798475.1 | PNUSAV000501 | SAMN10266906 | GCA_015798475.1 | NonPC | Public | NA | USA | Clinical | 91.4 | 100 | 3.41 | 98.2961 | 5271281 | 45.26 | 38 | 564960 |
| SAMN10266907-GCA_015798175.1 | PNUSAV000500 | SAMN10266907 | GCA_015798175.1 | NonPC | Public | NA | USA | Clinical | 91.3 | 100 | 0.07 | 98.3162 | 5134321 | 45.24 | 45 | 475296 |
| SAMN10285715-GCA_015798235.1 | PNUSAV000517 | SAMN10285715 | GCA_015798235.1 | NonPC | Public | NA | USA | Clinical | 88.3 | 100 | 0.1 | 98.0886 | 5146336 | 45.31 | 169 | 56159 |
| SAMN10285717-GCA_015798195.1 | PNUSAV000384 | SAMN10285717 | GCA_015798195.1 | NonPC | Public | NA | USA | Clinical | 91.3 | 100 | 0.34 | 98.2926 | 5143071 | 45.25 | 89 | 167918 |
| SAMN10287211-GCA_015798135.1 | 1.210-16 | SAMN10287211 | GCA_015798135.1 | NonPC | Public | 2016 | Peru | Clinical | 91.8 | 100 | 0.27 | 98.2975 | 5239380 | 45.17 | 65 | 249360 |
| SAMN10287212-GCA_015798035.1 | 3.252-16 | SAMN10287212 | GCA_015798035.1 | NonPC | Public | 2015 | Peru | Clinical | 91.2 | 100 | 0.16 | 98.333 | 5190734 | 45.25 | 139 | 101949 |
| SAMN10287213-GCA_015798415.1 | 1.166-15 | SAMN10287213 | GCA_015798415.1 | NonPC | Public | 2016 | Peru | Clinical | 91.4 | 100 | 0.55 | 98.301 | 5209951 | 45.21 | 38 | 861249 |
| SAMN10287215-GCA_015798315.1 | 3.369-15 | SAMN10287215 | GCA_015798315.1 | NonPC | Public | 2011 | Peru | Env | 91.7 | 100 | 2.86 | 98.2827 | 5313055 | 45.17 | 41 | 769321 |
| SAMN10287216-GCA_015798275.1 | 1.146-15 | SAMN10287216 | GCA_015798275.1 | NonPC | Public | 2015 | Peru | Clinical | 91.4 | 100 | 2.49 | 98.3039 | 5284217 | 45.22 | 42 | 492690 |
| SAMN10287217-GCA_015798215.1 | 3.254-16 | SAMN10287217 | GCA_015798215.1 | NonPC | Public | 2015 | Peru | Clinical | 91.4 | 100 | 0.7 | 98.2603 | 5156385 | 45.41 | 49 | 452521 |
| SAMN10320211-GCA_015798295.1 | PNUSAV000518 | SAMN10320211 | GCA_015798295.1 | NonPC | Public | NA | USA | Clinical | 89.5 | 100 | 1.7 | 98.0805 | 5101089 | 45.3 | 78 | 152836 |
| SAMN10320212-GCA_015798055.1 | PNUSAV000386 | SAMN10320212 | GCA_015798055.1 | NonPC | Public | NA | USA | Clinical | 91.3 | 100 | 0 | 98.2789 | 5142238 | 45.25 | 76 | 159426 |
| SAMN10338902-GCA_015798015.1 | 617910 | SAMN10338902 | GCA_015798015.1 | PC | Public | 2018 | NA | Clinical | 97.3 | 100 | 1.06 | 99.9922 | 5180767 | 45.14 | 101 | 148362 |
| SAMN10341922-GCA_015798075.1 | PNUSAV000520 | SAMN10341922 | GCA_015798075.1 | NonPC | Public | NA | USA | Clinical | 91.2 | 100 | 0 | 98.3094 | 5121926 | 45.29 | 123 | 86547 |
| SAMN10390740-GCA_015797875.1 | 634852 | SAMN10390740 | GCA_015797875.1 | NonPC | Public | 2018 | NA | Clinical | 92.4 | 100 | 4.12 | 98.354 | 5315649 | 45.19 | 73 | 199748 |
| SAMN10391026-GCA_015797575.1 | PNUSAV000523 | SAMN10391026 | GCA_015797575.1 | PC | Public | NA | USA | Clinical | 98.1 | 100 | 4.36 | 99.9921 | 5219191 | 45.32 | 50 | 196275 |
| SAMN10391027-GCA_015798495.1 | PNUSAV000511 | SAMN10391027 | GCA_015798495.1 | NonPC | Public | NA | USA | Clinical | 89.1 | 100 | 3.81 | 98.0804 | 5238248 | 45.28 | 67 | 215497 |
| SAMN10391029-GCA_015797555.1 | PNUSAV000506 | SAMN10391029 | GCA_015797555.1 | NonPC | Public | NA | USA | Clinical | 89.1 | 100 | 2.45 | 98.0598 | 5163504 | 45.28 | 92 | 148828 |
| SAMN10395984-GCA_015797515.1 | PNUSAV000522 | SAMN10395984 | GCA_015797515.1 | NonPC | Public | NA | USA | Clinical | 91.1 | 100 | 0.82 | 98.3283 | 5148832 | 45.36 | 156 | 76154 |
| SAMN10397788-GCF_004121195.1 | NA2 | SAMN10397788 | GCF_004121195.1 | NonPC | Public | 2017 | Malaysia | Env | 91.8 | 100 | 0.14 | 98.2648 | 5502878 | 45.29 | 75 | 202086 |
| SAMN10397789-GCF_004121285.1 | ML1 | SAMN10397789 | GCF_004121285.1 | NonPC | Public | 2017 | Malaysia | Env | 91.8 | 100 | 0.33 | 98.3655 | 5379877 | 45.21 | 31 | 545476 |
| SAMN10397790-GCF_004121175.1 | ML2 | SAMN10397790 | GCF_004121175.1 | NonPC | Public | 2017 | Malaysia | Env | 91.9 | 100 | 0.05 | 98.4042 | 5341133 | 45.17 | 28 | 405399 |
| SAMN10397791-GCF_004121225.1 | ML3 | SAMN10397791 | GCF_004121225.1 | NonPC | Public | 2017 | Malaysia | Env | 91.8 | 100 | 0.33 | 98.4044 | 5380185 | 45.2 | 41 | 308335 |
| SAMN10397792-GCF_004121185.1 | ML4 | SAMN10397792 | GCF_004121185.1 | NonPC | Public | 2017 | Malaysia | Env | 91.9 | 100 | 0.33 | 98.4281 | 5384157 | 45.2 | 31 | 411668 |
| SAMN10397793-GCF_004121235.1 | ML5 | SAMN10397793 | GCF_004121235.1 | NonPC | Public | 2017 | Malaysia | Env | 90.1 | 100 | 0.54 | 98.2769 | 5161074 | 45.23 | 22 | 680062 |
| SAMN10397794-GCF_004121105.1 | ML6 | SAMN10397794 | GCF_004121105.1 | NonPC | Public | 2017 | Malaysia | Env | 91.9 | 100 | 0.05 | 98.3928 | 5343831 | 45.17 | 34 | 402519 |
| SAMN10397795-GCF_004121085.1 | ML7 | SAMN10397795 | GCF_004121085.1 | NonPC | Public | 2017 | Malaysia | Env | 91.7 | 99.82 | 0.33 | 98.4054 | 5375249 | 45.2 | 38 | 308335 |
| SAMN10397796-GCF_004121095.1 | ML8 | SAMN10397796 | GCF_004121095.1 | NonPC | Public | 2017 | Malaysia | Env | 91.7 | 99.82 | 0.33 | 98.4262 | 5374262 | 45.2 | 42 | 309464 |
| SAMN10397797-GCF_004121115.1 | ML9 | SAMN10397797 | GCF_004121115.1 | NonPC | Public | 2017 | Malaysia | Env | 91.8 | 100 | 0.05 | 98.4092 | 5343603 | 45.17 | 41 | 308335 |
| SAMN10417092-SRR8186505 | 17802 | SAMN10417092 | SRR8186505 | NonPC | Public | NA | China | NA | 90.8 | 100 | 0.05 | 98.3513 | 5068708 | 45.25 | 44 | 509646 |
| SAMN10419342-GCA_015797455.1 | PNUSAV000528 | SAMN10419342 | GCA_015797455.1 | NonPC | Public | NA | USA | Clinical | 90.6 | 100 | 1.44 | 98.3248 | 5132538 | 45.26 | 75 | 194010 |
| SAMN10419343-GCA_015797535.1 | PNUSAV000526 | SAMN10419343 | GCA_015797535.1 | NonPC | Public | NA | USA | Clinical | 90.6 | 100 | 2.18 | 98.2717 | 5140139 | 45.24 | 69 | 191922 |
| SAMN10419344-GCA_015797495.1 | PNUSAV000525 | SAMN10419344 | GCA_015797495.1 | PC | Public | NA | USA | Clinical | 98.3 | 100 | 3.72 | 99.9895 | 5324709 | 45.34 | 74 | 125288 |
| SAMN10426750-GCF_009738005.1 | MSR16 | SAMN10426750 | GCF_009738005.1 | NonPC | Public | 2016 | Bangladesh | Env | 91.7 | 100 | 0.27 | 98.2872 | 5377931 | 45.09 | 53 | 422921 |
| SAMN10426773-GCF_009738025.1 | MSR17 | SAMN10426773 | GCF_009738025.1 | NonPC | Public | 2017 | Bangladesh | Env | 91.3 | 100 | 0.27 | 98.2857 | 5232129 | 45.19 | 35 | 886735 |
| SAMN10435367-GCA_015798435.1 | PNUSAV000529 | SAMN10435367 | GCA_015798435.1 | NonPC | Public | NA | USA | Clinical | 91.4 | 100 | 0.36 | 98.2819 | 5143081 | 45.24 | 46 | 343335 |
| SAMN10436686-GCF_009883835.1 | 20140722001-1 | SAMN10436686 | GCF_009883835.1 | NonPC | Public | 2014 | China | Env | 93 | 100 | 0.49 | 98.3008 | 5563056 | 45.27 | 5 | 3421899 |

| | | | | | | | | | | | | | | | | |
|---|---|---|---|---|---|---|---|---|---|---|---|---|---|---|---|---|
| SAMN10436796-GCF_009883855.1 | 20140829008-1 | SAMN10436796 | GCF_009883855.1 | NonPC | Public | 2014 | China | Env | 92.3 | 99.97 | 0.49 | 98.2962 | 5420599 | 45.34 | 4 | 3358530 |
| SAMN10436807-GCF_009883895.1 | 20151116002-3 | SAMN10436807 | GCF_009883895.1 | NonPC | Public | 2015 | China | Env | 92.7 | 99.46 | 0.54 | 98.3379 | 5456849 | 45.2 | 4 | 3491101 |
| SAMN10436897-GCF_009883815.1 | 20140624012-1 | SAMN10436897 | GCF_009883815.1 | NonPC | Public | 2014 | China | Env | 92.9 | 99.66 | 0.22 | 98.3157 | 5347867 | 45.35 | 4 | 3338799 |
| SAMN10449341-GCA_015797395.1 | PNUSAV000538 | SAMN10449341 | GCA_015797395.1 | NonPC | Public | NA | USA | Clinical | 92.7 | 100 | 2.02 | 98.3594 | 5195572 | 45.31 | 122 | 101942 |
| SAMN10477346-GCA_015797195.1 | PNUSAV000543 | SAMN10477346 | GCA_015797195.1 | NonPC | Public | NA | USA | Clinical | 91.4 | 100 | 3.41 | 98.2655 | 5233230 | 45.24 | 41 | 556688 |
| SAMN10477347-GCA_015797415.1 | PNUSAV000541 | SAMN10477347 | GCA_015797415.1 | NonPC | Public | NA | USA | Clinical | 91.3 | 100 | 0.15 | 98.2951 | 5144641 | 45.27 | 93 | 132319 |
| SAMN10477350-GCA_015797375.1 | PNUSAV000510 | SAMN10477350 | GCA_015797375.1 | NonPC | Public | NA | USA | Clinical | 91.3 | 100 | 0.82 | 98.2887 | 5122418 | 45.29 | 78 | 195066 |
| SAMN10477351-GCA_015797315.1 | PNUSAV000509 | SAMN10477351 | GCA_015797315.1 | NonPC | Public | NA | USA | Clinical | 89.7 | 100 | 3.58 | 98.2658 | 5320740 | 45.2 | 45 | 229753 |
| SAMN10477352-GCA_015797355.1 | PNUSAV000508 | SAMN10477352 | GCA_015797355.1 | NonPC | Public | NA | USA | Clinical | 90.6 | 100 | 0.07 | 98.3401 | 5242021 | 45.29 | 75 | 121512 |
| SAMN10484929-GCA_015797255.1 | PNUSAV000533 | SAMN10484929 | GCA_015797255.1 | NonPC | Public | NA | USA | Clinical | 91.4 | 100 | 4.09 | 98.2753 | 5279680 | 45.26 | 42 | 564960 |
| SAMN10484933-GCA_015797175.1 | PNUSAV000544 | SAMN10484933 | GCA_015797175.1 | NonPC | Public | NA | USA | Clinical | 87.5 | 100 | 1.84 | 98.0959 | 5012369 | 45.32 | 58 | 326219 |
| SAMN10484934-GCA_015796595.1 | PNUSAV000540 | SAMN10484934 | GCA_015796595.1 | NonPC | Public | NA | USA | Clinical | 90.9 | 100 | 0.5 | 98.3447 | 5058645 | 45.3 | 56 | 359433 |
| SAMN10519626-GCA_015797275.1 | PNUSAV000547 | SAMN10519626 | GCA_015797275.1 | NonPC | Public | NA | USA | Clinical | 89.7 | 100 | 0.39 | 98.25 | 5404194 | 45.27 | 101 | 100671 |
| SAMN10585383-GCA_015795155.1 | PNUSAV000546 | SAMN10585383 | GCA_015795155.1 | NonPC | Public | NA | USA | Clinical | 89.4 | 100 | 0.07 | 98.114 | 5053121 | 45.34 | 77 | 179288 |
| SAMN10585384-GCA_015795115.1 | PNUSAV000545 | SAMN10585384 | GCA_015795115.1 | NonPC | Public | NA | USA | Clinical | 89 | 100 | 0.36 | 98.0978 | 5024766 | 45.29 | 85 | 181438 |
| SAMN10591529-GCF_004194515.1 | D3112 | SAMN10591529 | GCF_004194515.1 | NonPC | Public | 2015 | China | Env | 90.7 | 100 | 0 | 98.3496 | 5286292 | 45.35 | 4 | 3239077 |
| SAMN10689491-GCA_015794895.1 | PNUSAV000562 | SAMN10689491 | GCA_015794895.1 | NonPC | Public | NA | USA | Clinical | 89 | 100 | 0.82 | 98.0986 | 5175839 | 45.21 | 175 | 76893 |
| SAMN10702672-GCF_009665495.1 | 2012AW-0154 | SAMN10702672 | GCF_009665495.1 | NonPC | Public | NA | USA | Clinical | 92.2 | 100 | 0.22 | 98.2897 | 5397131 | 45.17 | 2 | 3639425 |
| SAMN10723998-GCA_015790495.1 | CFSAN090610 | SAMN10723998 | GCA_015790495.1 | NonPC | Public | 2018 | Venezuela | Env | 92 | 100 | 4.46 | 98.3342 | 5365865 | 45.25 | 41 | 336407 |
| SAMN10723999-GCA_015790515.1 | CFSAN090611 | SAMN10723999 | GCA_015790515.1 | NonPC | Public | 2018 | Venezuela | Env | 92.1 | 100 | 0.74 | 98.3547 | 5202715 | 45.22 | 41 | 437526 |
| SAMN10743323-SRR8455928 | vp100057 | SAMN10743323 | SRR8455928 | NonPC | Public | 2010 | China | Clinical | 92.3 | 100 | 0 | 98.3727 | 5070545 | 45.22 | 61 | 452188 |
| SAMN10761833-GCF_004028465.1 | ICDC-VP1329 | SAMN10761833 | GCF_004028465.1 | NonPC | Public | 2013 | China | Clinical | 91.1 | 100 | 0 | 98.3185 | 5196118 | 45.26 | 124 | 157489 |
| SAMN10781732-SRR10019360 | 20141124002 | SAMN10781732 | SRR10019360 | NonPC | Public | 2014 | China | Env | 91.8 | 100 | 0.27 | 98.278 | 5377966 | 45.11 | 73 | 460478 |
| SAMN10781733-SRR10019359 | 20140526002 | SAMN10781733 | SRR10019359 | NonPC | Public | 2014 | China | Env | 91.9 | 100 | 0.22 | 98.2908 | 5312756 | 45.27 | 68 | 266006 |
| SAMN10782547-SRR10019358 | 20140115002 | SAMN10782547 | SRR10019358 | NonPC | Public | 2014 | China | Env | 91.3 | 100 | 0.56 | 98.3246 | 5344231 | 45.28 | 75 | 212524 |
| SAMN10794260-GCA_015790955.1 | PNUSAV000564 | SAMN10794260 | GCA_015790955.1 | NonPC | Public | NA | USA | Clinical | 88.3 | 100 | 4.09 | 98.1171 | 5024205 | 45.5 | 143 | 65941 |
| SAMN10794262-GCA_015791115.1 | PNUSAV000566 | SAMN10794262 | GCA_015791115.1 | PC | Public | NA | USA | Clinical | 98.2 | 100 | 2.72 | 99.9927 | 5243541 | 45.28 | 54 | 289349 |
| SAMN10794263-GCA_015790895.1 | PNUSAV000565 | SAMN10794263 | GCA_015790895.1 | NonPC | Public | NA | USA | Clinical | 91.2 | 100 | 4.88 | 98.2242 | 5218263 | 45.4 | 206 | 63361 |
| SAMN10818954-GCF_014769095.1 | DFP1847 | SAMN10818954 | GCF_014769095.1 | NonPC | Public | 2016 | India | Env | 92.1 | 100 | 0.82 | 98.3124 | 5490669 | 45.14 | 46 | 324980 |
| SAMN10854974-GCA_015790415.1 | PNUSAV000567 | SAMN10854974 | GCA_015790415.1 | PC | Public | NA | USA | Clinical | 98.1 | 100 | 2.04 | 99.9753 | 5276319 | 45.3 | 71 | 158353 |
| SAMN10954690-GCA_015789425.1 | PNUSAV000572 | SAMN10954690 | GCA_015789425.1 | NonPC | Public | NA | USA | Clinical | 87 | 99.73 | 0.43 | 98.3011 | 4998371 | 45.38 | 128 | 100421 |
| SAMN11056377-GCF_008693745.1 | FDA:FDAARGOS_662 | SAMN11056377 | GCF_008693745.1 | NonPC | Public | NA | USA | Clinical | 92.5 | 100 | 0 | 98.3248 | 5154108 | 45.35 | 2 | 3320553 |
| SAMN11056382-GCF_008693625.1 | FDA:FDAARGOS_667 | SAMN11056382 | GCF_008693625.1 | NonPC | Public | NA | USA | Clinical | 89.4 | 100 | 0 | 98.0464 | 5003148 | 45.47 | 2 | 3260755 |
| SAMN11119395-GCA_015789295.1 | PNUSAV000575 | SAMN11119395 | GCA_015789295.1 | NonPC | Public | NA | USA | Clinical | 90.6 | 100 | 0.68 | 98.3402 | 5181478 | 45.24 | 103 | 188848 |
| SAMN11119398-GCA_015789325.1 | PNUSAV000552 | SAMN11119398 | GCA_015789325.1 | NonPC | Public | NA | USA | Clinical | 90.6 | 100 | 1.91 | 98.3224 | 5126856 | 45.24 | 66 | 217547 |
| SAMN11119401-GCA_015789375.1 | PNUSAV000551 | SAMN11119401 | GCA_015789375.1 | NonPC | Public | NA | USA | Clinical | 91 | 100 | 0.44 | 98.2993 | 5117423 | 45.34 | 163 | 68249 |
| SAMN11159227-GCA_015789105.1 | PNUSAV000580 | SAMN11159227 | GCA_015789105.1 | PC | Public | NA | USA | Clinical | 98 | 100 | 2.45 | 99.9942 | 5160616 | 45.34 | 82 | 130428 |
| SAMN11191138-GCA_015789135.1 | PNUSAV000584 | SAMN11191138 | GCA_015789135.1 | NonPC | Public | NA | USA | Clinical | 89.1 | 100 | 2.72 | 98.058 | 5233406 | 45.16 | 148 | 89361 |
| SAMN11250066-GCA_015789255.1 | PNUSAV000583 | SAMN11250066 | GCA_015789255.1 | NonPC | Public | NA | USA | Clinical | 89.2 | 100 | 1.09 | 98.0645 | 5197041 | 45.09 | 87 | 155568 |
| SAMN11265710-GCA_015789195.1 | PNUSAV000587 | SAMN11265710 | GCA_015789195.1 | NonPC | Public | NA | USA | Clinical | 89.7 | 100 | 0.46 | 98.3076 | 5415214 | 44.98 | 70 | 195308 |
| SAMN11267087-GCF_004923355.1 | 16763 | SAMN11267087 | GCF_004923355.1 | NonPC | Public | 1978 | Russia | Clinical | 89.5 | 100 | 0.42 | 98.2582 | 5314145 | 45.05 | 63 | 241423 |
| SAMN11299007-GCA_015789175.1 | PNUSAV000588 | SAMN11299007 | GCA_015789175.1 | NonPC | Public | NA | USA | Clinical | 88.9 | 100 | 2.72 | 98.0796 | 5091410 | 45.37 | 164 | 66060 |
| SAMN11321601-GCF_012276655.1 | VP_2HP | SAMN11321601 | GCF_012276655.1 | NonPC | Public | 2014 | Thailand | Env | 91.7 | 99.95 | 0.09 | 98.2438 | 5484088 | 45.19 | 138 | 183055 |
| SAMN11360081-GCA_015782515.1 | G9456 | SAMN11360081 | GCA_015782515.1 | NonPC | Public | 2017 | USA | NA | 89 | 100 | 0.3 | 98.0568 | 5035102 | 45.27 | 92 | 166933 |
| SAMN11388518-GCA_015782095.1 | G1449 | SAMN11388518 | GCA_015782095.1 | NonPC | Public | 2009 | USA | Env | 89.8 | 99.9 | 0.2 | 98.2862 | 5178493 | 45.29 | 111 | 339157 |
| SAMN11388521-SRR9734085 | G8 | SAMN11388521 | SRR9734085 | NonPC | Public | 2007 | USA | Env | 88.8 | 100 | 0.04 | 98.2229 | 5068068 | 45.34 | 85 | 185892 |
| SAMN11388522-SRR9734088 | G735 | SAMN11388522 | SRR9734088 | NonPC | Public | 2008 | USA | Env | 89.8 | 100 | 0 | 98.301 | 4974495 | 45.35 | 51 | 369311 |
| SAMN11388525-GCA_015784255.1 | MA303 | SAMN11388525 | GCA_015784255.1 | NonPC | Public | 2015 | USA | NA | 89.3 | 100 | 2.94 | 98.3074 | 5483882 | 44.97 | 74 | 168181 |

| | | | | | | | | | | | | | | | | |
|---|---|---|---|---|---|---|---|---|---|---|---|---|---|---|---|---|
| SAMN11388529-GCA_015783955.1 | MA739 | SAMN11388529 | GCA_015783955.1 | NonPC | Public | 2016 | USA | NA | 88.8 | 100 | 0.1 | 98.1587 | 5050023 | 45.18 | 96 | 112288 |
| SAMN11388533-GCA_015782395.1 | G79 | SAMN11388533 | GCA_015782395.1 | NonPC | Public | 2007 | USA | NA | 90.8 | 100 | 3.81 | 98.3258 | 5208880 | 45.3 | 77 | 319472 |
| SAMN11400008-GCA_015789035.1 | PNUSAV000597 | SAMN11400008 | GCA_015789035.1 | NonPC | Public | NA | USA | Clinical | 88.3 | 100 | 3.21 | 98.1264 | 5105459 | 45.33 | 88 | 122171 |
| SAMN11407924-GCA_015788955.1 | PNUSAV000598 | SAMN11407924 | GCA_015788955.1 | NonPC | Public | NA | USA | Clinical | 89 | 100 | 1.91 | 98.0449 | 5051261 | 45.38 | 195 | 51732 |
| SAMN11537008-GCF_013369315.1 | JS-8-11-2 | SAMN11537008 | GCF_013369315.1 | NonPC | Public | 2013 | USA | Env | 88.7 | 100 | 0 | 98.0489 | 4933605 | 45.41 | 60 | 215206 |
| SAMN11547019-GCF_009391175.1 | HZ11-012 | SAMN11547019 | GCF_009391175.1 | NonPC | Public | 2011 | China | Clinical | 91.9 | 100 | 0.19 | 98.3332 | 5018862 | 45.33 | 133 | 82438 |
| SAMN11547020-GCF_009391125.1 | HZ12-014 | SAMN11547020 | GCF_009391125.1 | NonPC | Public | 2012 | China | Clinical | 91.9 | 100 | 0.19 | 98.3428 | 5018045 | 45.34 | 155 | 73801 |
| SAMN11547021-GCF_009391145.1 | HZ13-025 | SAMN11547021 | GCF_009391145.1 | NonPC | Public | 2013 | China | Clinical | 92.3 | 100 | 0.19 | 98.3473 | 5061888 | 45.29 | 55 | 291734 |
| SAMN11547022-GCF_009391115.1 | HZ13-037 | SAMN11547022 | GCF_009391115.1 | NonPC | Public | 2013 | China | Clinical | 92.9 | 100 | 0.06 | 98.3432 | 5127496 | 45.4 | 141 | 69789 |
| SAMN11547023-GCF_009391075.1 | HZ13-054 | SAMN11547023 | GCF_009391075.1 | NonPC | Public | 2013 | China | Clinical | 93.1 | 100 | 0.05 | 98.3588 | 5067513 | 45.3 | 38 | 513701 |
| SAMN11547024-GCF_009391085.1 | HZ13-082 | SAMN11547024 | GCF_009391085.1 | NonPC | Public | 2013 | China | Clinical | 93 | 100 | 0.05 | 98.3313 | 5051040 | 45.33 | 70 | 190299 |
| SAMN11547025-GCF_009391045.1 | HZ13-113 | SAMN11547025 | GCF_009391045.1 | NonPC | Public | 2013 | China | Clinical | 93 | 100 | 0.05 | 98.3366 | 5073495 | 45.31 | 85 | 137036 |
| SAMN11547026-GCF_009391035.1 | HZ14-003 | SAMN11547026 | GCF_009391035.1 | NonPC | Public | 2014 | China | Clinical | 92.9 | 100 | 0.05 | 98.348 | 5037975 | 45.37 | 118 | 103808 |
| SAMN11547027-GCF_009391015.1 | HZ14-038 | SAMN11547027 | GCF_009391015.1 | NonPC | Public | 2014 | China | Clinical | 93 | 100 | 0.06 | 98.366 | 5137504 | 45.37 | 91 | 111987 |
| SAMN11547028-GCF_009390965.1 | HZ14-064 | SAMN11547028 | GCF_009390965.1 | NonPC | Public | 2014 | China | Clinical | 93 | 100 | 0.05 | 98.3837 | 5033452 | 45.38 | 119 | 93072 |
| SAMN11547029-GCF_009390985.1 | HZ15-003 | SAMN11547029 | GCF_009390985.1 | NonPC | Public | 2015 | China | Clinical | 92.9 | 100 | 0.05 | 98.3483 | 5064057 | 45.33 | 146 | 72765 |
| SAMN11547030-GCF_009390925.1 | HZ15-072 | SAMN11547030 | GCF_009390925.1 | NonPC | Public | 2015 | China | Clinical | 92.9 | 100 | 0.05 | 98.3476 | 5032828 | 45.38 | 123 | 88416 |
| SAMN11547031-GCF_009390945.1 | HZ15-131 | SAMN11547031 | GCF_009390945.1 | NonPC | Public | 2015 | China | Clinical | 92.9 | 100 | 0.05 | 98.299 | 5064334 | 45.38 | 122 | 94032 |
| SAMN11547032-GCF_009390915.1 | HZ16-008 | SAMN11547032 | GCF_009390915.1 | NonPC | Public | 2016 | China | Clinical | 93.1 | 100 | 0.05 | 98.3702 | 5066486 | 45.31 | 51 | 345864 |
| SAMN11547033-GCF_009390895.1 | HZ16-018 | SAMN11547033 | GCF_009390895.1 | NonPC | Public | 2016 | China | Clinical | 92 | 100 | 0.19 | 98.2572 | 5065890 | 45.23 | 45 | 603561 |
| SAMN11547034-GCF_009390865.1 | HZ16-044 | SAMN11547034 | GCF_009390865.1 | NonPC | Public | 2016 | China | Clinical | 92 | 100 | 1.25 | 98.2779 | 5109138 | 45.29 | 98 | 545128 |
| SAMN11547035-GCF_009390855.1 | HZ16-101 | SAMN11547035 | GCF_009390855.1 | NonPC | Public | 2016 | China | Clinical | 92 | 100 | 0.19 | 98.2426 | 5067505 | 45.22 | 52 | 439746 |
| SAMN11547036-GCF_009390825.1 | HZ16-115 | SAMN11547036 | GCF_009390825.1 | NonPC | Public | 2016 | China | Clinical | 92 | 100 | 0.46 | 98.2488 | 5078916 | 45.2 | 46 | 545128 |
| SAMN11547037-GCF_009390795.1 | HZ16-148 | SAMN11547037 | GCF_009390795.1 | NonPC | Public | 2016 | China | Clinical | 92 | 100 | 0.46 | 98.2473 | 5081585 | 45.21 | 54 | 287952 |
| SAMN11547038-GCF_009390805.1 | HZ16-159 | SAMN11547038 | GCF_009390805.1 | NonPC | Public | 2016 | China | Clinical | 93.2 | 100 | 0.05 | 98.3839 | 5083148 | 45.3 | 49 | 384340 |
| SAMN11547039-GCF_009390765.1 | HZ16-165 | SAMN11547039 | GCF_009390765.1 | NonPC | Public | 2016 | China | Clinical | 92 | 100 | 0.46 | 98.2734 | 5080133 | 45.2 | 51 | 603820 |
| SAMN11547040-GCF_009390755.1 | HZ16-167 | SAMN11547040 | GCF_009390755.1 | NonPC | Public | 2016 | China | Clinical | 93.2 | 100 | 0.05 | 98.3855 | 5058921 | 45.31 | 44 | 513872 |
| SAMN11547041-GCF_009390695.1 | HZ16-178 | SAMN11547041 | GCF_009390695.1 | NonPC | Public | 2016 | China | Clinical | 92 | 100 | 0.46 | 98.2687 | 5086689 | 45.18 | 56 | 603790 |
| SAMN11547042-GCF_009390725.1 | HZ16-187 | SAMN11547042 | GCF_009390725.1 | NonPC | Public | 2016 | China | Clinical | 92 | 100 | 0.46 | 98.2769 | 5080679 | 45.2 | 49 | 603592 |
| SAMN11547043-GCF_009390705.1 | HZ16-189 | SAMN11547043 | GCF_009390705.1 | NonPC | Public | 2016 | China | Clinical | 92 | 100 | 0.19 | 98.2558 | 5068041 | 45.23 | 46 | 603548 |
| SAMN11547044-GCF_009390675.1 | HZ16-194 | SAMN11547044 | GCF_009390675.1 | NonPC | Public | 2016 | China | Clinical | 92 | 100 | 0.46 | 98.2521 | 5080198 | 45.2 | 51 | 545128 |
| SAMN11547045-GCF_009390655.1 | HZ16-215 | SAMN11547045 | GCF_009390655.1 | NonPC | Public | 2016 | China | Clinical | 92 | 100 | 0.19 | 98.2853 | 5067988 | 45.23 | 48 | 495342 |
| SAMN11547046-GCF_009390605.1 | HZ16-237 | SAMN11547046 | GCF_009390605.1 | NonPC | Public | 2016 | China | Clinical | 92 | 100 | 0.5 | 98.2312 | 5078503 | 45.21 | 51 | 440123 |
| SAMN11547047-GCF_009390615.1 | HZ16-242 | SAMN11547047 | GCF_009390615.1 | NonPC | Public | 2016 | China | Clinical | 92 | 100 | 0.46 | 98.2646 | 5083338 | 45.21 | 48 | 603790 |
| SAMN11547048-GCF_009390595.1 | HZ16-248 | SAMN11547048 | GCF_009390595.1 | NonPC | Public | 2016 | China | Clinical | 92 | 100 | 0.46 | 98.292 | 5085389 | 45.2 | 43 | 545128 |
| SAMN11547049-GCF_009390555.1 | HZ16-259 | SAMN11547049 | GCF_009390555.1 | NonPC | Public | 2016 | China | Clinical | 92 | 100 | 0.46 | 98.2773 | 5079437 | 45.21 | 42 | 662127 |
| SAMN11547050-GCF_009390565.1 | HZ16-260 | SAMN11547050 | GCF_009390565.1 | NonPC | Public | 2016 | China | Clinical | 92 | 100 | 0.46 | 98.2398 | 5080408 | 45.21 | 47 | 546215 |
| SAMN11547051-GCF_009390515.1 | HZ16-270 | SAMN11547051 | GCF_009390515.1 | NonPC | Public | 2016 | China | Clinical | 92 | 100 | 0.46 | 98.2709 | 5084257 | 45.2 | 50 | 604274 |
| SAMN11547052-GCF_009390495.1 | HZ16-290 | SAMN11547052 | GCF_009390495.1 | NonPC | Public | 2016 | China | Clinical | 92 | 100 | 0.46 | 98.2462 | 5081644 | 45.2 | 50 | 545128 |
| SAMN11547053-GCF_009390465.1 | HZ16-307 | SAMN11547053 | GCF_009390465.1 | NonPC | Public | 2016 | China | Clinical | 92 | 100 | 0.74 | 98.2738 | 5078630 | 45.2 | 47 | 544915 |
| SAMN11547054-GCF_009390505.1 | HZ16-323 | SAMN11547054 | GCF_009390505.1 | NonPC | Public | 2016 | China | Clinical | 92 | 100 | 0.46 | 98.2632 | 5077337 | 45.2 | 51 | 603929 |
| SAMN11547055-GCF_009390455.1 | HZ16-329 | SAMN11547055 | GCF_009390455.1 | NonPC | Public | 2016 | China | Clinical | 92 | 100 | 0.46 | 98.2494 | 5090782 | 45.2 | 50 | 440111 |
| SAMN11547056-GCF_009390415.1 | HZ17-011 | SAMN11547056 | GCF_009390415.1 | NonPC | Public | 2017 | China | Clinical | 91.6 | 100 | 0.19 | 98.2085 | 5026779 | 45.31 | 121 | 85986 |
| SAMN11547057-GCF_009390385.1 | HZ17-017 | SAMN11547057 | GCF_009390385.1 | NonPC | Public | 2017 | China | Clinical | 91.7 | 100 | 0.46 | 98.2462 | 5052923 | 45.26 | 110 | 114586 |
| SAMN11547058-GCF_009390395.1 | HZ17-025 | SAMN11547058 | GCF_009390395.1 | NonPC | Public | 2017 | China | Clinical | 91.7 | 100 | 0.46 | 98.165 | 5051659 | 45.29 | 115 | 88869 |
| SAMN11547059-GCF_009390375.1 | HZ17-098 | SAMN11547059 | GCF_009390375.1 | NonPC | Public | 2017 | China | Clinical | 92.6 | 100 | 0.33 | 98.3303 | 5056956 | 45.4 | 172 | 68983 |
| SAMN11547060-GCF_009390355.1 | HZ17-110 | SAMN11547060 | GCF_009390355.1 | NonPC | Public | 2017 | China | Clinical | 92.8 | 100 | 0.19 | 98.3412 | 5061723 | 45.36 | 98 | 113014 |
| SAMN11547061-GCF_009390325.1 | HZ17-118 | SAMN11547061 | GCF_009390325.1 | NonPC | Public | 2017 | China | Clinical | 91.9 | 100 | 0.46 | 98.2626 | 5068915 | 45.22 | 83 | 142292 |

| | | | | | | | | | | | | | | | | |
|---|---|---|---|---|---|---|---|---|---|---|---|---|---|---|---|---|
| SAMN11547062-GCF_009390305.1 | HZ17-125 | SAMN11547062 | GCF_009390305.1 | NonPC | Public | 2017 | China | Clinical | 92.9 | 100 | 0.19 | 98.3345 | 5063594 | 45.34 | 100 | 98765 |
| SAMN11547063-GCF_009390295.1 | HZ17-129 | SAMN11547063 | GCF_009390295.1 | NonPC | Public | 2017 | China | Clinical | 92.9 | 100 | 0.19 | 98.3481 | 5055257 | 45.35 | 88 | 137878 |
| SAMN11547064-GCF_009390255.1 | HZ17-143 | SAMN11547064 | GCF_009390255.1 | NonPC | Public | 2017 | China | Clinical | 91.7 | 100 | 0.46 | 98.2662 | 5040826 | 45.29 | 156 | 62103 |
| SAMN11547065-GCF_009390265.1 | HZ17-150 | SAMN11547065 | GCF_009390265.1 | NonPC | Public | 2017 | China | Clinical | 91.6 | 100 | 0.46 | 98.2329 | 5041921 | 45.3 | 118 | 84534 |
| SAMN11547066-GCF_009390195.1 | HZ17-157 | SAMN11547066 | GCF_009390195.1 | NonPC | Public | 2017 | China | Clinical | 91.6 | 100 | 0.19 | 98.2122 | 5041191 | 45.28 | 103 | 114991 |
| SAMN11547067-GCF_009390215.1 | HZ17-162 | SAMN11547067 | GCF_009390215.1 | NonPC | Public | 2017 | China | Clinical | 91.9 | 100 | 0.19 | 98.265 | 5049555 | 45.27 | 98 | 122976 |
| SAMN11547068-GCF_009390205.1 | HZ17-167 | SAMN11547068 | GCF_009390205.1 | NonPC | Public | 2017 | China | Clinical | 91.7 | 100 | 0.2 | 98.2721 | 5126192 | 45.33 | 129 | 82583 |
| SAMN11547069-GCF_009390165.1 | HZ17-171 | SAMN11547069 | GCF_009390165.1 | NonPC | Public | 2017 | China | Clinical | 91.8 | 100 | 0.46 | 98.2389 | 5023973 | 45.26 | 109 | 118217 |
| SAMN11547070-GCF_009390155.1 | HZ17-180 | SAMN11547070 | GCF_009390155.1 | NonPC | Public | 2017 | China | Clinical | 90.6 | 100 | 0.33 | 98.2018 | 4926642 | 45.48 | 343 | 24643 |
| SAMN11547071-GCF_009390105.1 | HZ17-187 | SAMN11547071 | GCF_009390105.1 | NonPC | Public | 2017 | China | Clinical | 91.4 | 100 | 0.46 | 98.2176 | 5021964 | 45.35 | 178 | 57915 |
| SAMN11547072-GCF_009390115.1 | HZ17-191 | SAMN11547072 | GCF_009390115.1 | NonPC | Public | 2017 | China | Clinical | 91.3 | 100 | 0.46 | 98.2365 | 5024634 | 45.35 | 179 | 60736 |
| SAMN11547073-GCF_009390095.1 | HZ17-194 | SAMN11547073 | GCF_009390095.1 | NonPC | Public | 2017 | China | Clinical | 91.5 | 100 | 0.6 | 98.185 | 5036643 | 45.31 | 139 | 71879 |
| SAMN11547074-GCF_009390065.1 | HZ17-210 | SAMN11547074 | GCF_009390065.1 | NonPC | Public | 2017 | China | Clinical | 91.8 | 100 | 0.46 | 98.2395 | 5054141 | 45.26 | 116 | 104492 |
| SAMN11547075-GCF_009390055.1 | HZ17-212 | SAMN11547075 | GCF_009390055.1 | NonPC | Public | 2017 | China | Clinical | 91.6 | 100 | 0.46 | 98.2457 | 5043182 | 45.3 | 148 | 68753 |
| SAMN11547076-GCF_009390025.1 | HZ17-221 | SAMN11547076 | GCF_009390025.1 | NonPC | Public | 2017 | China | Clinical | 92.6 | 100 | 0.19 | 98.365 | 5056543 | 45.4 | 179 | 63033 |
| SAMN11547077-GCF_009390005.1 | HZ17-224 | SAMN11547077 | GCF_009390005.1 | NonPC | Public | 2017 | China | Clinical | 91.6 | 100 | 0.46 | 98.2068 | 5042708 | 45.29 | 131 | 78210 |
| SAMN11547078-GCF_009389965.1 | HZ17-253 | SAMN11547078 | GCF_009389965.1 | NonPC | Public | 2017 | China | Clinical | 91.7 | 100 | 0.46 | 98.2201 | 5051030 | 45.28 | 125 | 89096 |
| SAMN11547079-GCF_009389955.1 | HZ17-255 | SAMN11547079 | GCF_009389955.1 | NonPC | Public | 2017 | China | Clinical | 91.6 | 100 | 0.46 | 98.2231 | 5037863 | 45.31 | 166 | 57970 |
| SAMN11547080-GCF_009389975.1 | HZ17-265 | SAMN11547080 | GCF_009389975.1 | NonPC | Public | 2017 | China | Clinical | 91.8 | 100 | 0.46 | 98.2087 | 5060440 | 45.25 | 92 | 122976 |
| SAMN11547081-GCF_009389855.1 | HZ17-272 | SAMN11547081 | GCF_009389855.1 | NonPC | Public | 2017 | China | Clinical | 91.6 | 100 | 0.46 | 98.2286 | 5044087 | 45.3 | 148 | 84714 |
| SAMN11547082-GCF_009389865.1 | HZ17-290 | SAMN11547082 | GCF_009389865.1 | NonPC | Public | 2017 | China | Clinical | 91.9 | 100 | 0.46 | 98.2319 | 5063345 | 45.25 | 101 | 117620 |
| SAMN11547083-GCF_009389915.1 | HZ17-386 | SAMN11547083 | GCF_009389915.1 | NonPC | Public | 2017 | China | Env | 91.4 | 100 | 0 | 98.3259 | 5084719 | 45.3 | 186 | 65067 |
| SAMN11547084-GCF_009389895.1 | HZ17-395 | SAMN11547084 | GCF_009389895.1 | NonPC | Public | 2017 | China | Clinical | 91.7 | 100 | 0.19 | 98.2189 | 5035925 | 45.31 | 162 | 70323 |
| SAMN11547085-GCF_009389905.1 | HZ18-031 | SAMN11547085 | GCF_009389905.1 | NonPC | Public | 2018 | China | Clinical | 92.9 | 100 | 0.05 | 98.3331 | 5042529 | 45.36 | 100 | 128171 |
| SAMN11547086-GCF_009389775.1 | HZ18-037 | SAMN11547086 | GCF_009389775.1 | NonPC | Public | 2018 | China | Clinical | 92.9 | 100 | 0.05 | 98.3414 | 5032522 | 45.38 | 131 | 88225 |
| SAMN11547087-GCF_009389805.1 | HZ18-056 | SAMN11547087 | GCF_009389805.1 | NonPC | Public | 2018 | China | Clinical | 91.8 | 100 | 0.19 | 98.2476 | 5009793 | 45.28 | 115 | 108812 |
| SAMN11547088-GCF_009389785.1 | HZ18-094 | SAMN11547088 | GCF_009389785.1 | NonPC | Public | 2018 | China | Clinical | 92.7 | 100 | 0.05 | 98.315 | 5026105 | 45.39 | 130 | 82546 |
| SAMN11547089-GCF_009389755.1 | HZ18-096 | SAMN11547089 | GCF_009389755.1 | NonPC | Public | 2018 | China | Clinical | 92.2 | 100 | 0.19 | 98.3265 | 5024673 | 45.27 | 103 | 101495 |
| SAMN11547090-GCF_009389765.1 | HZ18-116 | SAMN11547090 | GCF_009389765.1 | PC | Public | 2018 | China | Clinical | 97.9 | 100 | 0 | 99.9719 | 5087076 | 45.35 | 147 | 78583 |
| SAMN11579495-GCF_013393845.1 | LVP1 | SAMN11579495 | GCF_013393845.1 | NonPC | Public | 2013 | China | Env | 89.5 | 100 | 0.39 | 98.149 | 5310635 | 45.17 | 2 | 3478693 |
| SAMN11579496-GCF_013393865.1 | LVP2 | SAMN11579496 | GCF_013393865.1 | NonPC | Public | 2013 | China | Env | 92.5 | 100 | 0.27 | 98.2694 | 5521375 | 45.03 | 2 | 3589443 |
| SAMN11579497-GCF_013393885.1 | LVP66 | SAMN11579497 | GCF_013393885.1 | NonPC | Public | 2013 | China | Env | 91.9 | 100 | 0.05 | 98.3606 | 5107724 | 45.41 | 2 | 3325904 |
| SAMN11618431-GCA_015788735.1 | PNUSAV000610 | SAMN11618431 | GCA_015788735.1 | NonPC | Public | NA | USA | Clinical | 89.2 | 100 | 3.54 | 98.0668 | 5297144 | 45.09 | 62 | 234013 |
| SAMN11620798-GCA_015783775.1 | G1334 | SAMN11620798 | GCA_015783775.1 | NonPC | Public | 2009 | USA | Env | 89.4 | 100 | 0.98 | 98.2752 | 5086455 | 45.28 | 63 | 184246 |
| SAMN11620799-GCA_015784275.1 | G4973 | SAMN11620799 | GCA_015784275.1 | NonPC | Public | 2014 | USA | Env | 91.6 | 100 | 0.2 | 98.3785 | 5094228 | 45.32 | 150 | 87027 |
| SAMN11890813-GCA_015790115.1 | 636292 | SAMN11890813 | GCA_015790115.1 | NonPC | Public | 2018 | NA | Clinical | 89.7 | 99.59 | 2.78 | 98.3566 | 5093985 | 45.25 | 135 | 179503 |
| SAMN11890816-GCA_015790165.1 | 652104 | SAMN11890816 | GCA_015790165.1 | PC | Public | 2018 | NA | Clinical | 97.6 | 100 | 4.63 | 99.9723 | 5366587 | 45.3 | 83 | 159139 |
| SAMN11959618-GCF_006370575.1 | GSJ2016VP009 | SAMN11959618 | GCF_006370575.1 | NonPC | Public | 2016 | China | Env | 88.3 | 100 | 0.4 | 98.3087 | 5042876 | 45.31 | 29 | 1012849 |
| SAMN12015781-GCA_015788015.1 | PNUSAV000633 | SAMN12015781 | GCA_015788015.1 | NonPC | Public | NA | USA | Clinical | 89.5 | 100 | 2.76 | 98.2686 | 5231515 | 45.3 | 112 | 86106 |
| SAMN12015782-GCA_015787935.1 | PNUSAV000631 | SAMN12015782 | GCA_015787935.1 | NonPC | Public | NA | USA | Clinical | 91.4 | 100 | 4.63 | 98.2848 | 5287354 | 45.26 | 40 | 556718 |
| SAMN12084703-GCA_015787915.1 | PNUSAV000640 | SAMN12084703 | GCA_015787915.1 | PC | Public | NA | USA | Clinical | 97.6 | 100 | 0 | 99.9783 | 5146924 | 45.38 | 108 | 105826 |
| SAMN12084709-GCA_015789555.1 | PNUSAV000638 | SAMN12084709 | GCA_015789555.1 | PC | Public | NA | USA | Clinical | 97.8 | 100 | 3 | 99.9837 | 5282886 | 45.34 | 69 | 154256 |
| SAMN12084717-GCA_015789615.1 | PNUSAV000639 | SAMN12084717 | GCA_015789615.1 | PC | Public | NA | USA | Clinical | 97.7 | 100 | 3.27 | 99.9819 | 5238031 | 45.34 | 73 | 140198 |
| SAMN12084723-GCA_015787895.1 | PNUSAV000637 | SAMN12084723 | GCA_015787895.1 | PC | Public | NA | USA | Clinical | 97.4 | 100 | 1.1 | 99.9746 | 5271521 | 45.42 | 135 | 82937 |
| SAMN12084766-GCA_015787955.1 | PNUSAV000641 | SAMN12084766 | GCA_015787955.1 | PC | Public | NA | USA | Clinical | 97.8 | 100 | 4.36 | 99.9887 | 5304508 | 45.35 | 72 | 156751 |
| SAMN12123413-GCF_006517795.1 | Vb0624 | SAMN12123413 | GCF_006517795.1 | NonPC | Public | 2015 | China | Env | 90.5 | 100 | 0.37 | 98.2566 | 5383500 | 45.32 | 2 | 3556649 |
| SAMN12136230-GCA_015789575.1 | PNUSAV000649 | SAMN12136230 | GCA_015789575.1 | NonPC | Public | NA | USA | Clinical | 91.4 | 100 | 4.36 | 98.3184 | 5289723 | 45.25 | 39 | 718998 |
| SAMN12136231-GCA_015789495.1 | PNUSAV000642 | SAMN12136231 | GCA_015789495.1 | NonPC | Public | NA | USA | Clinical | 91.4 | 100 | 4.63 | 98.2846 | 5300657 | 45.26 | 44 | 346430 |

| | | | | | | | | | | | | | | | | |
|---|---|---|---|---|---|---|---|---|---|---|---|---|---|---|---|---|
| SAMN12136232-GCA_015789415.1 | PNUSAV000643 | SAMN12136232 | GCA_015789415.1 | NonPC | Public | NA | USA | Clinical | 91.4 | 100 | 1.98 | 98.323 | 5218112 | 45.26 | 45 | 344577 |
| SAMN12141645-GCA_015789515.1 | PNUSAV000653 | SAMN12141645 | GCA_015789515.1 | NonPC | Public | NA | USA | Clinical | 91 | 100 | 0.82 | 98.3107 | 5129461 | 45.34 | 182 | 66337 |
| SAMN12180012-GCA_015787855.1 | PNUSAV000651 | SAMN12180012 | GCA_015787855.1 | NonPC | Public | NA | USA | Clinical | 88.9 | 100 | 3.66 | 98.0844 | 5045981 | 45.41 | 88 | 129604 |
| SAMN12211963-GCA_015787715.1 | PNUSAV000646 | SAMN12211963 | GCA_015787715.1 | NonPC | Public | NA | USA | Clinical | 89.7 | 100 | 0.49 | 98.3513 | 5029433 | 45.32 | 45 | 387589 |
| SAMN12211964-GCA_015789655.1 | PNUSAV000618 | SAMN12211964 | GCA_015789655.1 | PC | Public | NA | USA | Clinical | 97.5 | 100 | 1.91 | 99.9668 | 5173902 | 45.34 | 91 | 179672 |
| SAMN12211965-GCA_015789635.1 | PNUSAV000252 | SAMN12211965 | GCA_015789635.1 | NonPC | Public | NA | USA | Clinical | 89.8 | 100 | 1.4 | 98.248 | 5219644 | 45.2 | 53 | 230849 |
| SAMN12211966-GCA_015789875.1 | PNUSAV000251 | SAMN12211966 | GCA_015789875.1 | NonPC | Public | NA | USA | Clinical | 88.4 | 100 | 0.05 | 98.1314 | 4905625 | 45.44 | 66 | 136367 |
| SAMN12211985-GCA_015789815.1 | PNUSAV000654 | SAMN12211985 | GCA_015789815.1 | PC | Public | NA | USA | Clinical | 97.8 | 100 | 4.54 | 99.985 | 5348099 | 45.38 | 53 | 311511 |
| SAMN12212336-GCA_015789775.1 | PNUSAV000253 | SAMN12212336 | GCA_015789775.1 | PC | Public | NA | USA | Clinical | 97.8 | 100 | 3.27 | 99.9675 | 5234656 | 45.31 | 87 | 155476 |
| SAMN12242087-GCA_015789915.1 | PNUSAV000662 | SAMN12242087 | GCA_015789915.1 | NonPC | Public | NA | USA | Clinical | 90.1 | 100 | 4.32 | 98.21 | 5217795 | 45.16 | 42 | 469920 |
| SAMN12275090-GCA_015789755.1 | PNUSAV000665 | SAMN12275090 | GCA_015789755.1 | NonPC | Public | NA | USA | Clinical | 90.6 | 100 | 4.9 | 98.3025 | 5291296 | 45.28 | 80 | 183535 |
| SAMN12288857-GCA_015789835.1 | PNUSAV000596 | SAMN12288857 | GCA_015789835.1 | NonPC | Public | NA | USA | Clinical | 88.8 | 100 | 4.42 | 98.1472 | 5215349 | 45.19 | 47 | 268527 |
| SAMN12288859-GCA_016258225.1 | PNUSAV000615 | SAMN12288859 | GCA_016258225.1 | NonPC | Public | NA | USA | Clinical | 89.3 | 100 | 2.68 | 98.0644 | 5094155 | 45.35 | 78 | 150913 |
| SAMN12324285-GCA_015787755.1 | PNUSAV000668 | SAMN12324285 | GCA_015787755.1 | NonPC | Public | NA | USA | Clinical | 91.3 | 100 | 0.68 | 98.2997 | 5142952 | 45.25 | 70 | 201872 |
| SAMN12333865-GCA_015787455.1 | PNUSAV000674 | SAMN12333865 | GCA_015787455.1 | NonPC | Public | NA | USA | Clinical | 91.2 | 100 | 1.36 | 98.2918 | 5180177 | 45.31 | 112 | 108325 |
| SAMN12333872-GCA_015787545.1 | PNUSAV000685 | SAMN12333872 | GCA_015787545.1 | NonPC | Public | NA | USA | Clinical | 89.1 | 100 | 0.61 | 98.0913 | 5108008 | 45.25 | 81 | 173513 |
| SAMN12333874-GCA_016257805.1 | PNUSAV000683 | SAMN12333874 | GCA_016257805.1 | NonPC | Public | NA | USA | Clinical | 90.8 | 100 | 4.11 | 98.2777 | 5299766 | 45.19 | 47 | 399084 |
| SAMN12333875-GCA_015787475.1 | PNUSAV000675 | SAMN12333875 | GCA_015787475.1 | NonPC | Public | NA | USA | Clinical | 91.2 | 100 | 2.45 | 98.3009 | 5211876 | 45.29 | 110 | 96079 |
| SAMN12337027-GCA_015787655.1 | PNUSAV000678 | SAMN12337027 | GCA_015787655.1 | NonPC | Public | NA | USA | Clinical | 90.7 | 100 | 2.72 | 98.2792 | 5172515 | 45.25 | 82 | 184648 |
| SAMN12337177-GCA_015786375.1 | PNUSAV000664 | SAMN12337177 | GCA_015786375.1 | NonPC | Public | NA | USA | Clinical | 89.2 | 100 | 0.48 | 98.0618 | 5078259 | 45.24 | 84 | 192198 |
| SAMN12341753-GCA_015787065.1 | PNUSAV000681 | SAMN12341753 | GCA_015787065.1 | NonPC | Public | NA | USA | Clinical | 91 | 100 | 1.23 | 98.3078 | 5144159 | 45.32 | 178 | 56677 |
| SAMN12359103-GCF_009193065.1 | 731 | SAMN12359103 | GCF_009193065.1 | NonPC | Public | 2016 | China | Clinical | 91.8 | 100 | 0.46 | 98.2926 | 5068848 | 45.19 | 44 | 450036 |
| SAMN12364834-GCF_009991885.1 | Jan-64 | SAMN12364834 | GCF_009991885.1 | NonPC | Public | 2001 | Peru | Clinical | 92.9 | 100 | 0 | 98.3955 | 5129335 | 45.22 | 44 | 409540 |
| SAMN12364835-GCF_009991855.1 | 514-08 | SAMN12364835 | GCF_009991855.1 | NonPC | Public | 2008 | Peru | Clinical | 91.3 | 100 | 0 | 98.3864 | 5128651 | 45.22 | 58 | 284597 |
| SAMN12364836-GCF_009991845.1 | 1202-11 | SAMN12364836 | GCF_009991845.1 | PC | Public | 2011 | Peru | Clinical | 98.7 | 100 | 0 | 99.9816 | 5192518 | 45.14 | 73 | 318936 |
| SAMN12364837-GCF_009991825.1 | 613-13 | SAMN12364837 | GCF_009991825.1 | NonPC | Public | 2013 | Peru | Clinical | 91 | 100 | 0.14 | 98.2015 | 5136681 | 45.24 | 104 | 166701 |
| SAMN12364838-GCF_009991435.1 | 1168-13 | SAMN12364838 | GCF_009991435.1 | NonPC | Public | 2013 | Peru | Clinical | 89.4 | 100 | 0 | 98.17 | 5094901 | 45.2 | 76 | 247998 |
| SAMN12364839-GCF_009991445.1 | G1 | SAMN12364839 | GCF_009991445.1 | PC | Public | 2014 | Peru | Clinical | 98.8 | 99.73 | 0.14 | 99.9866 | 5234819 | 45.19 | 214 | 340270 |
| SAMN12364840-GCF_009991765.1 | 1833-14 | SAMN12364840 | GCF_009991765.1 | NonPC | Public | 2014 | Peru | Clinical | 91.2 | 100 | 0.07 | 98.3249 | 5171793 | 45.22 | 54 | 261308 |
| SAMN12364841-GCF_009991745.1 | 249-15 | SAMN12364841 | GCF_009991745.1 | PC | Public | 2015 | Peru | Clinical | 98.4 | 100 | 0.09 | 99.9905 | 5244617 | 45.27 | 69 | 153020 |
| SAMN12364842-GCF_009991805.1 | 276-15 | SAMN12364842 | GCF_009991805.1 | PC | Public | 2015 | Peru | Clinical | 98.4 | 100 | 0.09 | 99.9819 | 5246267 | 45.27 | 60 | 288971 |
| SAMN12364843-GCF_009991755.1 | 147-15 | SAMN12364843 | GCF_009991755.1 | NonPC | Public | 2015 | Peru | Clinical | 91.5 | 100 | 0.41 | 98.3164 | 5218963 | 45.22 | 70 | 278180 |
| SAMN12364844-GCF_009991725.1 | 146-15 | SAMN12364844 | GCF_009991725.1 | NonPC | Public | 2015 | Peru | Clinical | 91.6 | 100 | 0 | 98.2784 | 5239871 | 45.23 | 109 | 124705 |
| SAMN12364845-GCF_009991645.1 | 001-15 | SAMN12364845 | GCF_009991645.1 | NonPC | Public | 2015 | Peru | Clinical | 90.8 | 100 | 0.61 | 98.2811 | 5302058 | 45.09 | 57 | 315343 |
| SAMN12364846-GCF_009991685.1 | 164-16 | SAMN12364846 | GCF_009991685.1 | PC | Public | 2016 | Peru | Clinical | 98.5 | 100 | 0 | 99.9929 | 5117304 | 45.28 | 57 | 347272 |
| SAMN12364847-GCF_009991625.1 | 686-17 | SAMN12364847 | GCF_009991625.1 | PC | Public | 2017 | Peru | Clinical | 98 | 100 | 0 | 99.9021 | 5167836 | 45.3 | 80 | 245630 |
| SAMN12364848-GCF_009991675.1 | 2214-17 | SAMN12364848 | GCF_009991675.1 | PC | Public | 2017 | Peru | Clinical | 98.2 | 100 | 0 | 99.9434 | 5171831 | 45.31 | 73 | 402111 |
| SAMN12364849-GCF_009991665.1 | 710-17 | SAMN12364849 | GCF_009991665.1 | NonPC | Public | 2017 | Peru | Clinical | 91.1 | 100 | 0 | 98.2463 | 5057112 | 45.32 | 90 | 192334 |
| SAMN12386531-SRR9850089 | VP4974 | SAMN12386531 | SRR9850089 | NonPC | Public | 2014 | USA | NA | 89.6 | 100 | 0.63 | 98.2883 | 5123789 | 45.22 | 229 | 40137 |
| SAMN12404565-GCF_008271865.1 | PH698 | SAMN12404565 | GCF_008271865.1 | NonPC | Public | 2014 | Philippines | Env | 90.9 | 100 | 0.07 | 98.3107 | 5342783 | 45.46 | 62 | 193506 |
| SAMN12404860-GCA_015786875.1 | PNUSAV000754 | SAMN12404860 | GCA_015786875.1 | NonPC | Public | NA | USA | Clinical | 91.3 | 100 | 2.25 | 98.328 | 5248678 | 45.27 | 73 | 145376 |
| SAMN12404861-GCA_015787355.1 | PNUSAV000751 | SAMN12404861 | GCA_015787355.1 | NonPC | Public | NA | USA | Clinical | 91.3 | 100 | 4.9 | 98.3313 | 5339224 | 45.28 | 79 | 141461 |
| SAMN12404862-GCA_015787035.1 | PNUSAV000687 | SAMN12404862 | GCA_015787035.1 | NonPC | Public | NA | USA | Clinical | 91.3 | 100 | 1.64 | 98.2976 | 5169378 | 45.24 | 53 | 456955 |
| SAMN12404863-GCA_015787675.1 | PNUSAV000214 | SAMN12404863 | GCA_015787675.1 | NonPC | Public | NA | USA | Clinical | 91.3 | 100 | 1.36 | 98.2977 | 5197871 | 45.28 | 63 | 189730 |
| SAMN12419323-GCA_015787695.1 | PNUSAV000781 | SAMN12419323 | GCA_015787695.1 | NonPC | Public | NA | USA | Clinical | 91.4 | 100 | 0.62 | 98.2888 | 5164653 | 45.25 | 42 | 680372 |
| SAMN12422654-GCA_015787295.1 | PNUSAV000766 | SAMN12422654 | GCA_015787295.1 | NonPC | Public | NA | USA | Clinical | 91.3 | 100 | 1.24 | 98.3671 | 5077147 | 45.4 | 131 | 85573 |
| SAMN12495036-GCF_008039385.1 | HZ02-J7 | SAMN12495036 | GCF_008039385.1 | NonPC | Public | 2002 | China | Clinical | 92.3 | 100 | 0.05 | 98.3577 | 5017258 | 45.28 | 48 | 539211 |
| SAMN12495037-GCF_008039855.1 | HZ13-043 | SAMN12495037 | GCF_008039855.1 | NonPC | Public | 2013 | China | Clinical | 92.3 | 100 | 0.19 | 98.3441 | 5041074 | 45.25 | 52 | 407592 |

| | | | | | | | | | | | | | | | | |
|---|---|---|---|---|---|---|---|---|---|---|---|---|---|---|---|---|
| SAMN12495038-GCF_008039365.1 | HZ13-088 | SAMN12495038 | GCF_008039365.1 | NonPC | Public | 2013 | China | Clinical | 92.5 | 100 | 0.05 | 98.3116 | 5020959 | 45.26 | 38 | 602702 |
| SAMN12495039-GCF_008039345.1 | HZ13-102 | SAMN12495039 | GCF_008039345.1 | NonPC | Public | 2013 | China | Clinical | 93.1 | 100 | 0.05 | 98.3526 | 5062729 | 45.3 | 41 | 463546 |
| SAMN12495040-GCF_008039325.1 | HZ14-002 | SAMN12495040 | GCF_008039325.1 | NonPC | Public | 2014 | China | Clinical | 93.1 | 100 | 0.05 | 98.3561 | 5063217 | 45.3 | 45 | 513454 |
| SAMN12495041-GCF_008039285.1 | HZ15-046 | SAMN12495041 | GCF_008039285.1 | NonPC | Public | 2015 | China | Clinical | 93 | 100 | 0 | 98.4253 | 5126382 | 45.2 | 37 | 455052 |
| SAMN12495042-GCF_008039295.1 | HZ16-338 | SAMN12495042 | GCF_008039295.1 | NonPC | Public | 2016 | China | Clinical | 92.3 | 100 | 0.19 | 98.3462 | 5028859 | 45.24 | 53 | 471372 |
| SAMN12495043-GCF_008039275.1 | HZ16-515 | SAMN12495043 | GCF_008039275.1 | NonPC | Public | 2016 | China | Clinical | 92.3 | 100 | 0.19 | 98.375 | 5029673 | 45.23 | 49 | 471687 |
| SAMN12495044-GCF_008039245.1 | HZ17-018 | SAMN12495044 | GCF_008039245.1 | NonPC | Public | 2017 | China | Clinical | 92.1 | 100 | 0.05 | 98.2802 | 5123366 | 45.24 | 61 | 548712 |
| SAMN12495045-GCF_008039215.1 | HZ17-083 | SAMN12495045 | GCF_008039215.1 | NonPC | Public | 2017 | China | Clinical | 92.2 | 100 | 0.19 | 98.3572 | 5018692 | 45.25 | 47 | 399559 |
| SAMN12495046-GCF_008039225.1 | HZ17-381 | SAMN12495046 | GCF_008039225.1 | NonPC | Public | 2017 | China | Env | 91.5 | 100 | 0.33 | 98.3968 | 5019325 | 45.31 | 45 | 449975 |
| SAMN12495047-GCF_008039905.1 | HZ17-383 | SAMN12495047 | GCF_008039905.1 | NonPC | Public | 2017 | China | Env | 90.5 | 100 | 0.03 | 98.2813 | 5051563 | 45.31 | 37 | 1300409 |
| SAMN12495048-GCF_008039175.1 | HZ18-062 | SAMN12495048 | GCF_008039175.1 | NonPC | Public | 2018 | China | Clinical | 92.3 | 100 | 0.19 | 98.3649 | 5027629 | 45.24 | 48 | 602548 |
| SAMN12495049-GCF_008039155.1 | HZ18-100 | SAMN12495049 | GCF_008039155.1 | NonPC | Public | 2018 | China | Clinical | 93.1 | 100 | 0.05 | 98.3727 | 5065698 | 45.3 | 41 | 463552 |
| SAMN12504970-GCA_015786935.1 | PNUSAV000804 | SAMN12504970 | GCA_015786935.1 | NonPC | Public | NA | USA | Clinical | 91.3 | 100 | 2.86 | 98.279 | 5209432 | 45.26 | 69 | 205197 |
| SAMN12504973-GCA_015786265.1 | PNUSAV000753 | SAMN12504973 | GCA_015786265.1 | NonPC | Public | NA | USA | Clinical | 90.4 | 100 | 1.32 | 98.3 | 5151481 | 45.26 | 126 | 101704 |
| SAMN12504979-GCA_015786395.1 | PNUSAV000760 | SAMN12504979 | GCA_015786395.1 | NonPC | Public | NA | USA | Clinical | 91.3 | 100 | 4.63 | 98.2676 | 5329154 | 45.27 | 56 | 269760 |
| SAMN12504987-GCA_015786255.1 | PNUSAV000627 | SAMN12504987 | GCA_015786255.1 | NonPC | Public | NA | USA | Clinical | 91.4 | 100 | 0.32 | 98.3087 | 5145409 | 45.24 | 38 | 807987 |
| SAMN12534784-GCA_015787225.1 | PNUSAV000782 | SAMN12534784 | GCA_015787225.1 | NonPC | Public | NA | USA | Clinical | 90.3 | 100 | 4.68 | 98.192 | 5267187 | 45.17 | 41 | 425491 |
| SAMN12542225-GCF_008271735.1 | PH1339 | SAMN12542225 | GCF_008271735.1 | NonPC | Public | 2015 | Philippines | Env | 91.4 | 100 | 0.41 | 98.3277 | 5332603 | 45.21 | 57 | 294559 |
| SAMN12583379-GCA_015787255.1 | PNUSAV000824 | SAMN12583379 | GCA_015787255.1 | NonPC | Public | NA | USA | Clinical | 91.3 | 100 | 4.57 | 98.2659 | 5328691 | 45.26 | 49 | 237356 |
| SAMN12583380-GCA_015786075.1 | PNUSAV000811 | SAMN12583380 | GCA_015786075.1 | NonPC | Public | NA | USA | Clinical | 88.7 | 100 | 3.14 | 98.3011 | 5149170 | 45.32 | 47 | 201451 |
| SAMN12583381-GCA_015786095.1 | PNUSAV000809 | SAMN12583381 | GCA_015786095.1 | NonPC | Public | NA | USA | Clinical | 89.3 | 100 | 3.88 | 98.2105 | 5317852 | 45.31 | 53 | 267449 |
| SAMN12583385-GCA_015787275.1 | PNUSAV000808 | SAMN12583385 | GCA_015787275.1 | NonPC | Public | NA | USA | Clinical | 91.1 | 100 | 0.42 | 98.3384 | 5121279 | 45.28 | 45 | 431710 |
| SAMN12587483-GCA_015785995.1 | PNUSAV000645 | SAMN12587483 | GCA_015785995.1 | NonPC | Public | NA | USA | Clinical | 88.8 | 100 | 1.99 | 98.125 | 5170903 | 45.25 | 116 | 123618 |
| SAMN12587867-GCA_015785835.1 | PNUSAV000833 | SAMN12587867 | GCA_015785835.1 | NonPC | Public | NA | USA | Clinical | 90.4 | 100 | 0.86 | 98.3204 | 5129690 | 45.28 | 152 | 86777 |
| SAMN12587869-GCA_015785875.1 | PNUSAV000805 | SAMN12587869 | GCA_015785875.1 | NonPC | Public | NA | USA | Clinical | 89.6 | 100 | 2.01 | 98.2673 | 5222345 | 45.24 | 52 | 215988 |
| SAMN12587878-GCA_015785915.1 | PNUSAV000836 | SAMN12587878 | GCA_015785915.1 | NonPC | Public | NA | USA | Clinical | 90.5 | 100 | 0.7 | 98.281 | 5434833 | 45.03 | 198 | 51919 |
| SAMN12587880-GCA_015785895.1 | PNUSAV000841 | SAMN12587880 | GCA_015785895.1 | NonPC | Public | NA | USA | Clinical | 91.9 | 100 | 0.69 | 98.2853 | 5208025 | 45.22 | 111 | 101452 |
| SAMN12587882-GCA_015785855.1 | PNUSAV000837 | SAMN12587882 | GCA_015785855.1 | NonPC | Public | NA | USA | Clinical | 90.3 | 100 | 2.38 | 98.2264 | 5270835 | 45.28 | 187 | 59857 |
| SAMN12587883-GCA_015785795.1 | PNUSAV000840 | SAMN12587883 | GCA_015785795.1 | NonPC | Public | NA | USA | Clinical | 91.8 | 100 | 2.44 | 98.2696 | 5189357 | 45.3 | 135 | 88459 |
| SAMN12587887-GCA_015785755.1 | PNUSAV000867 | SAMN12587887 | GCA_015785755.1 | NonPC | Public | NA | USA | Clinical | 91.2 | 100 | 3.68 | 98.309 | 5237391 | 45.32 | 130 | 87843 |
| SAMN12587889-GCA_015785935.1 | PNUSAV000868 | SAMN12587889 | GCA_015785935.1 | NonPC | Public | NA | USA | Clinical | 91.2 | 100 | 0.54 | 98.3241 | 5146811 | 45.31 | 131 | 95011 |
| SAMN12587891-GCA_015785815.1 | PNUSAV000870 | SAMN12587891 | GCA_015785815.1 | NonPC | Public | NA | USA | Clinical | 91.2 | 100 | 1.23 | 98.2943 | 5180820 | 45.3 | 114 | 100493 |
| SAMN12587894-GCA_015784695.1 | PNUSAV000820 | SAMN12587894 | GCA_015784695.1 | NonPC | Public | NA | USA | Clinical | 91.1 | 100 | 1.09 | 98.2788 | 5163448 | 45.32 | 128 | 87402 |
| SAMN12587898-GCA_015784655.1 | PNUSAV000873 | SAMN12587898 | GCA_015784655.1 | NonPC | Public | NA | USA | Clinical | 89 | 100 | 0.34 | 98.1007 | 5073759 | 45.35 | 157 | 73415 |
| SAMN12587899-GCA_016253795.1 | PNUSAV000822 | SAMN12587899 | GCA_016253795.1 | NonPC | Public | NA | USA | Clinical | 89.6 | 100 | 4.13 | 98.2506 | 5285117 | 45.26 | 124 | 88705 |
| SAMN12587900-GCA_015784715.1 | PNUSAV000829 | SAMN12587900 | GCA_015784715.1 | NonPC | Public | NA | USA | Clinical | 91.4 | 100 | 3 | 98.3244 | 5225685 | 45.32 | 118 | 89938 |
| SAMN12587901-GCA_015784675.1 | PNUSAV000866 | SAMN12587901 | GCA_015784675.1 | NonPC | Public | NA | USA | Clinical | 91 | 100 | 0.36 | 98.2885 | 5138942 | 45.35 | 162 | 81037 |
| SAMN12588159-GCA_015784635.1 | PNUSAV000869 | SAMN12588159 | GCA_015784635.1 | NonPC | Public | NA | USA | Clinical | 91.2 | 100 | 2.45 | 98.2722 | 5246792 | 45.32 | 95 | 107825 |
| SAMN12588274-GCA_015786835.1 | PNUSAV000832 | SAMN12588274 | GCA_015786835.1 | NonPC | Public | NA | USA | Clinical | 90.6 | 100 | 2.91 | 98.408 | 5186023 | 45.26 | 128 | 80001 |
| SAMN12588275-GCA_015785355.1 | PNUSAV000834 | SAMN12588275 | GCA_015785355.1 | NonPC | Public | NA | USA | Clinical | 90.4 | 100 | 4.35 | 98.1925 | 5288448 | 45.21 | 142 | 93656 |
| SAMN12588289-GCA_015785595.1 | PNUSAV000872 | SAMN12588289 | GCA_015785595.1 | NonPC | Public | NA | USA | Clinical | 91 | 100 | 1.36 | 98.3173 | 5178907 | 45.36 | 146 | 83150 |
| SAMN12588290-GCA_015784575.1 | PNUSAV000821 | SAMN12588290 | GCA_015784575.1 | NonPC | Public | NA | USA | Clinical | 90.5 | 100 | 1.63 | 98.3397 | 5129473 | 45.28 | 101 | 116898 |
| SAMN12588291-GCA_016253695.1 | PNUSAV000838 | SAMN12588291 | GCA_016253695.1 | NonPC | Public | NA | USA | Clinical | 90.3 | 100 | 4.48 | 98.2579 | 5333462 | 45.11 | 141 | 98255 |
| SAMN12588292-GCA_015785575.1 | PNUSAV000835 | SAMN12588292 | GCA_015785575.1 | NonPC | Public | NA | USA | Clinical | 90.4 | 100 | 0.07 | 98.2851 | 5154106 | 45.31 | 139 | 85536 |
| SAMN12588308-GCA_015784795.1 | PNUSAV000871 | SAMN12588308 | GCA_015784795.1 | NonPC | Public | NA | USA | Clinical | 91.1 | 100 | 3.54 | 98.3336 | 5222028 | 45.34 | 127 | 100722 |
| SAMN12602752-GCA_016253655.1 | PNUSAV000910 | SAMN12602752 | GCA_016253655.1 | NonPC | Public | NA | USA | Clinical | 90.8 | 100 | 0.34 | 98.3284 | 5054917 | 45.46 | 251 | 41252 |
| SAMN12602754-GCA_015786775.1 | PNUSAV000880 | SAMN12602754 | GCA_015786775.1 | NonPC | Public | NA | USA | Clinical | 91.3 | 100 | 1.7 | 98.317 | 5181189 | 45.23 | 47 | 560450 |
| SAMN12609659-GCA_015786815.1 | PNUSAV000842 | SAMN12609659 | GCA_015786815.1 | NonPC | Public | NA | USA | Clinical | 91.4 | 100 | 1.26 | 98.306 | 5168571 | 45.24 | 41 | 762300 |

| | | | | | | | | | | | | | | | | |
|---|---|---|---|---|---|---|---|---|---|---|---|---|---|---|---|---|
| SAMN12614692-GCA_015786675.1 | PNUSAV000941 | SAMN12614692 | GCA_015786675.1 | NonPC | Public | NA | USA | Clinical | 90 | 100 | 1.21 | 98.2785 | 5032013 | 45.37 | 189 | 53701 |
| SAMN12614694-GCA_015786855.1 | PNUSAV000931 | SAMN12614694 | GCA_015786855.1 | NonPC | Public | NA | USA | Clinical | 90.4 | 100 | 3.34 | 98.2774 | 5226929 | 45.24 | 33 | 534718 |
| SAMN12620793-GCA_015786515.1 | PNUSAV000885 | SAMN12620793 | GCA_015786515.1 | NonPC | Public | NA | USA | Clinical | 90.6 | 100 | 3.72 | 98.2729 | 5216188 | 45.27 | 67 | 233803 |
| SAMN12620794-GCA_015786575.1 | PNUSAV000817 | SAMN12620794 | GCA_015786575.1 | NonPC | Public | NA | USA | Clinical | 91.3 | 100 | 4.77 | 98.3085 | 5245418 | 45.24 | 45 | 520829 |
| SAMN12627221-GCA_015786455.1 | PNUSAV000932 | SAMN12627221 | GCA_015786455.1 | NonPC | Public | NA | USA | Clinical | 91.2 | 100 | 1.67 | 98.299 | 5378024 | 45.21 | 133 | 88396 |
| SAMN12627222-GCA_015786475.1 | PNUSAV000812 | SAMN12627222 | GCA_015786475.1 | NonPC | Public | NA | USA | Clinical | 91.7 | 100 | 2.18 | 98.3964 | 5126364 | 45.25 | 69 | 153186 |
| SAMN12641087-GCA_015786415.1 | PNUSAV000942 | SAMN12641087 | GCA_015786415.1 | NonPC | Public | NA | USA | Clinical | 91.2 | 100 | 1.63 | 98.3104 | 5167105 | 45.31 | 131 | 85210 |
| SAMN12642305-GCA_015786435.1 | PNUSAV000814 | SAMN12642305 | GCA_015786435.1 | NonPC | Public | NA | USA | Clinical | 91.3 | 100 | 2.09 | 98.2822 | 5183210 | 45.26 | 66 | 194089 |
| SAMN12642391-GCA_015786335.1 | PNUSAV000865 | SAMN12642391 | GCA_015786335.1 | NonPC | Public | NA | USA | Clinical | 89.2 | 100 | 4.09 | 98.0691 | 5214902 | 45.25 | 68 | 233237 |
| SAMN12644902-GCA_015786495.1 | PNUSAV000862 | SAMN12644902 | GCA_015786495.1 | NonPC | Public | NA | USA | Clinical | 91.4 | 100 | 4.09 | 98.2927 | 5299952 | 45.25 | 42 | 516571 |
| SAMN12645948-GCF_009496475.1 | SEB2 | SAMN12645948 | GCF_009496475.1 | NonPC | Public | 2014 | Malaysia | Env | 91.6 | 100 | 0.27 | 98.2704 | 5580190 | 45.16 | 98 | 205165 |
| SAMN12647415-GCA_015785315.1 | PNUSAV000943 | SAMN12647415 | GCA_015785315.1 | NonPC | Public | NA | USA | Clinical | 89.9 | 100 | 4.4 | 98.2498 | 5418717 | 45.19 | 35 | 559056 |
| SAMN12648277-GCF_009764075.1 | 2010V-1106 | SAMN12648277 | GCF_009764075.1 | NonPC | Public | 2010 | USA | Clinical | 92.4 | 100 | 0 | 98.3282 | 5144797 | 45.35 | 2 | 3311269 |
| SAMN12648278-GCF_009764055.1 | 2012AW-0224 | SAMN12648278 | GCF_009764055.1 | NonPC | Public | 2012 | USA | Clinical | 89.4 | 100 | 0.72 | 98.2052 | 5425729 | 45.19 | 3 | 3535345 |
| SAMN12648279-GCF_012275005.1 | 2012V-1116 | SAMN12648279 | GCF_012275005.1 | NonPC | Public | NA | USA | Clinical | 91.2 | 100 | 0.14 | 98.1926 | 5177231 | 45.25 | 2 | 3383078 |
| SAMN12648280-GCF_012274985.1 | 2012V-1165 | SAMN12648280 | GCF_012274985.1 | NonPC | Public | 2012 | USA | Clinical | 89.6 | 100 | 0.34 | 98.1256 | 5161492 | 45.26 | 2 | 3411422 |
| SAMN12648281-GCF_009763625.1 | 2013V-1174 | SAMN12648281 | GCF_009763625.1 | NonPC | Public | 2013 | USA | Clinical | 91.3 | 100 | 0.14 | 98.1915 | 5163702 | 45.25 | 2 | 3369685 |
| SAMN12648282-GCF_009763645.1 | 2013V-1146 | SAMN12648282 | GCF_009763645.1 | NonPC | Public | 2013 | USA | Clinical | 92.4 | 100 | 0 | 98.2781 | 5197133 | 45.31 | 2 | 3343366 |
| SAMN12648283-GCF_009763605.1 | 2013V-1136 | SAMN12648283 | GCF_009763605.1 | NonPC | Public | NA | NA | Clinical | 92 | 99.76 | 0.2 | 98.2663 | 5170591 | 45.3 | 2 | 3397332 |
| SAMN12648284-GCF_014525355.1 | 2013V-1176 | SAMN12648284 | GCF_014525355.1 | NonPC | Public | 2013 | USA | Clinical | 90.7 | 100 | 0.04 | 98.2527 | 5328844 | 45.28 | 3 | 3310415 |
| SAMN12648285-GCF_009763565.1 | 2013V-1181 | SAMN12648285 | GCF_009763565.1 | NonPC | Public | 2013 | USA | Clinical | 91 | 100 | 0.04 | 98.2473 | 5292073 | 45.26 | 2 | 3418724 |
| SAMN12648286-GCF_009763545.1 | 2013V-1244 | SAMN12648286 | GCF_009763545.1 | NonPC | Public | 2013 | USA | Clinical | 92.3 | 99.93 | 0 | 98.365 | 5129940 | 45.36 | 2 | 3276195 |
| SAMN12648287-GCF_009763525.1 | 2014V-1066 | SAMN12648287 | GCF_009763525.1 | NonPC | Public | 2014 | USA | Clinical | 92.4 | 100 | 0 | 98.3515 | 5158934 | 45.34 | 2 | 3323763 |
| SAMN12648288-GCF_009763505.1 | 2014V-1125 | SAMN12648288 | GCF_009763505.1 | NonPC | Public | 2014 | USA | Clinical | 92.4 | 100 | 0 | 98.317 | 5158348 | 45.34 | 2 | 3323757 |
| SAMN12648289-GCF_009763025.1 | AM43962 | SAMN12648289 | GCF_009763025.1 | NonPC | Public | NA | USA | Clinical | 90.9 | 100 | 0 | 98.3353 | 5128087 | 45.42 | 2 | 3297364 |
| SAMN12648300-GCF_009763445.1 | 2012AW-0353 | SAMN12648300 | GCF_009763445.1 | NonPC | Public | 2012 | USA | Clinical | 92.2 | 100 | 0 | 98.3643 | 5087213 | 45.39 | 2 | 3332037 |
| SAMN12648301-GCF_009763425.1 | AM46865 | SAMN12648301 | GCF_009763425.1 | NonPC | Public | NA | USA | Clinical | 91.8 | 100 | 0.05 | 98.3358 | 5088684 | 45.46 | 2 | 3284504 |
| SAMN12648302-GCF_009763405.1 | AM51552 | SAMN12648302 | GCF_009763405.1 | NonPC | Public | NA | USA | Clinical | 92.3 | 100 | 0.03 | 98.4662 | 5136309 | 45.37 | 2 | 3293962 |
| SAMN12648303-GCF_012274865.1 | AM51557 | SAMN12648303 | GCF_012274865.1 | NonPC | Public | NA | USA | Clinical | 90.3 | 100 | 0 | 98.3171 | 5088562 | 45.39 | 2 | 3330514 |
| SAMN12648306-GCF_009763165.1 | 2015AW-0174 | SAMN12648306 | GCF_009763165.1 | NonPC | Public | 2015 | USA | Clinical | 92.4 | 100 | 0 | 98.3124 | 5160547 | 45.33 | 2 | 3326911 |
| SAMN12657534-GCA_015785435.1 | PNUSAV000953 | SAMN12657534 | GCA_015785435.1 | NonPC | Public | NA | USA | Clinical | 91.3 | 100 | 1.43 | 98.2985 | 5196659 | 45.28 | 88 | 125389 |
| SAMN12657537-GCA_015785475.1 | PNUSAV000954 | SAMN12657537 | GCA_015785475.1 | NonPC | Public | NA | USA | Clinical | 91.3 | 100 | 0.5 | 98.2853 | 5151032 | 45.28 | 105 | 102261 |
| SAMN12661200-GCA_015785375.1 | PNUSAV000939 | SAMN12661200 | GCA_015785375.1 | NonPC | Public | NA | USA | Clinical | 91.4 | 100 | 2.52 | 98.3165 | 5309363 | 45.25 | 50 | 444307 |
| SAMN12661210-GCA_015784965.1 | PNUSAV000940 | SAMN12661210 | GCA_015784965.1 | NonPC | Public | NA | USA | Clinical | 89.2 | 100 | 4.9 | 98.0563 | 5229640 | 45.24 | 69 | 318605 |
| SAMN12661287-GCA_015784395.1 | PNUSAV000952 | SAMN12661287 | GCA_015784395.1 | NonPC | Public | NA | USA | Clinical | 91.5 | 100 | 4.9 | 98.299 | 5297957 | 45.28 | 54 | 231963 |
| SAMN12661288-GCA_015785035.1 | PNUSAV000951 | SAMN12661288 | GCA_015785035.1 | NonPC | Public | NA | USA | Clinical | 91.5 | 100 | 3.27 | 98.2985 | 5236329 | 45.28 | 46 | 718943 |
| SAMN12661289-GCA_015784365.1 | PNUSAV000950 | SAMN12661289 | GCA_015784365.1 | PC | Public | NA | USA | Clinical | 97.7 | 100 | 4.09 | 99.9648 | 5337013 | 45.29 | 83 | 154463 |
| SAMN12668540-GCA_015784345.1 | PNUSAV000676 | SAMN12668540 | GCA_015784345.1 | NonPC | Public | NA | USA | Clinical | 89.3 | 100 | 1.43 | 98.077 | 5144471 | 45.24 | 66 | 244772 |
| SAMN12671079-GCF_009647345.1 | 3HP | SAMN12671079 | GCF_009647345.1 | NonPC | Public | 2014 | Thailand | Env | 91.8 | 100 | 0 | 98.2739 | 5531288 | 45.22 | 94 | 244558 |
| SAMN12671080-GCF_009645315.1 | SEA5 | SAMN12671080 | GCF_009645315.1 | NonPC | Public | 2014 | Malaysia | Env | 91.7 | 100 | 0.27 | 98.2481 | 5584932 | 45.17 | 99 | 271397 |
| SAMN12671081-GCF_009645245.1 | SAB6 | SAMN12671081 | GCF_009645245.1 | NonPC | Public | 2014 | Malaysia | Env | 91.7 | 100 | 0.27 | 98.297 | 5571725 | 45.17 | 99 | 213427 |
| SAMN12671083-GCF_009647695.1 | SDA4 | SAMN12671083 | GCF_009647695.1 | NonPC | Public | 2014 | Viet_Nam | Env | 91 | 100 | 0.46 | 98.3548 | 5344121 | 45.21 | 113 | 196190 |
| SAMN12671085-GCF_009647705.1 | SDA1 | SAMN12671085 | GCF_009647705.1 | NonPC | Public | 2014 | Viet_Nam | Env | 90.7 | 100 | 0.07 | 98.3228 | 5340816 | 45.27 | 53 | 344251 |
| SAMN12671088-GCF_009647755.1 | SA2D2 | SAMN12671088 | GCF_009647755.1 | NonPC | Public | 2014 | Malaysia | Env | 89 | 100 | 1.02 | 98.2386 | 5337335 | 45.1 | 70 | 334543 |
| SAMN12699037-GCA_015784455.1 | PNUSAV000955 | SAMN12699037 | GCA_015784455.1 | NonPC | Public | NA | USA | Clinical | 90.8 | 100 | 3.51 | 98.3327 | 5180112 | 45.3 | 90 | 151965 |
| SAMN12699056-GCA_015784415.1 | PNUSAV000965 | SAMN12699056 | GCA_015784415.1 | NonPC | Public | NA | USA | Clinical | 89.8 | 100 | 1.7 | 98.1862 | 5207812 | 45.17 | 62 | 213624 |
| SAMN12699060-GCA_015785195.1 | PNUSAV000970 | SAMN12699060 | GCA_015785195.1 | PC | Public | NA | USA | Clinical | 97.9 | 100 | 0.07 | 99.9793 | 5167100 | 45.2 | 176 | 93832 |
| SAMN12699061-GCA_015784915.1 | PNUSAV000968 | SAMN12699061 | GCA_015784915.1 | NonPC | Public | NA | USA | Clinical | 91.4 | 100 | 3.86 | 98.2896 | 5294530 | 45.24 | 43 | 461806 |

| | | | | | | | | | | | | | | | | |
|---|---|---|---|---|---|---|---|---|---|---|---|---|---|---|---|---|
| SAMN12715986-GCA_015783415.1 | PNUSAV000947 | SAMN12715986 | GCA_015783415.1 | NonPC | Public | NA | USA | Clinical | 91.3 | 100 | 2.73 | 98.2818 | 5241131 | 45.26 | 63 | 200108 |
| SAMN12715990-GCA_015785735.1 | PNUSAV000946 | SAMN12715990 | GCA_015785735.1 | NonPC | Public | NA | USA | Clinical | 89.1 | 100 | 2.72 | 98.1156 | 5151931 | 45.29 | 95 | 114778 |
| SAMN12715998-GCA_015784955.1 | PNUSAV000881 | SAMN12715998 | GCA_015784955.1 | NonPC | Public | NA | USA | Clinical | 91.5 | 100 | 0.32 | 98.3846 | 5063592 | 45.26 | 35 | 455607 |
| SAMN12716007-GCA_015784875.1 | PNUSAV000958 | SAMN12716007 | GCA_015784875.1 | NonPC | Public | NA | USA | Clinical | 89.9 | 100 | 0.32 | 98.3362 | 5170114 | 45.18 | 36 | 446537 |
| SAMN12716008-GCA_015784815.1 | PNUSAV000945 | SAMN12716008 | GCA_015784815.1 | NonPC | Public | NA | USA | Clinical | 91.3 | 100 | 2.25 | 98.3138 | 5207663 | 45.26 | 63 | 199612 |
| SAMN12716019-GCA_016248395.1 | PNUSAV000957 | SAMN12716019 | GCA_016248395.1 | NonPC | Public | NA | USA | Clinical | 91.1 | 100 | 0.15 | 98.2955 | 5170493 | 45.18 | 40 | 323127 |
| SAMN12716020-GCA_015783455.1 | PNUSAV000995 | SAMN12716020 | GCA_015783455.1 | NonPC | Public | NA | USA | Clinical | 91.2 | 100 | 0.77 | 98.2851 | 5130446 | 45.29 | 116 | 89037 |
| SAMN12716021-GCA_015783475.1 | PNUSAV000996 | SAMN12716021 | GCA_015783475.1 | NonPC | Public | NA | USA | Clinical | 91.3 | 100 | 1.16 | 98.2839 | 5150619 | 45.26 | 81 | 131534 |
| SAMN12716022-GCA_015784765.1 | PNUSAV000997 | SAMN12716022 | GCA_015784765.1 | PC | Public | NA | USA | Clinical | 97.8 | 100 | 1.28 | 99.9849 | 5250794 | 45.37 | 67 | 144869 |
| SAMN12721745-GCA_015783435.1 | PNUSAV000975 | SAMN12721745 | GCA_015783435.1 | NonPC | Public | NA | USA | Clinical | 89.9 | 100 | 0.89 | 98.259 | 5158242 | 45.3 | 115 | 100815 |
| SAMN12721751-GCA_015783595.1 | PNUSAV000974 | SAMN12721751 | GCA_015783595.1 | NonPC | Public | NA | USA | Clinical | 90.3 | 100 | 2.82 | 98.3122 | 5076546 | 45.27 | 37 | 294149 |
| SAMN12721753-GCA_015783655.1 | PNUSAV000964 | SAMN12721753 | GCA_015783655.1 | NonPC | Public | NA | USA | Clinical | 89.6 | 100 | 0.04 | 98.2802 | 5203343 | 45.2 | 38 | 438995 |
| SAMN12766994-GCA_015785775.1 | PNUSAV001023 | SAMN12766994 | GCA_015785775.1 | NonPC | Public | NA | USA | Clinical | 91.4 | 100 | 2.72 | 98.3241 | 5196136 | 45.32 | 90 | 117793 |
| SAMN12767034-GCA_015783635.1 | PNUSAV000963 | SAMN12767034 | GCA_015783635.1 | NonPC | Public | NA | USA | Clinical | 91.3 | 100 | 3 | 98.3173 | 5198600 | 45.34 | 125 | 88825 |
| SAMN12767263-GCA_015785455.1 | PNUSAV001006 | SAMN12767263 | GCA_015785455.1 | NonPC | Public | NA | USA | Clinical | 91.3 | 100 | 1.63 | 98.3013 | 5180363 | 45.25 | 61 | 239462 |
| SAMN12767278-GCA_015784315.1 | PNUSAV000908 | SAMN12767278 | GCA_015784315.1 | NonPC | Public | NA | USA | Clinical | 91.3 | 100 | 2.45 | 98.3322 | 5187466 | 45.31 | 79 | 167990 |
| SAMN12767281-GCA_015785155.1 | PNUSAV000905 | SAMN12767281 | GCA_015785155.1 | NonPC | Public | NA | USA | Clinical | 87.9 | 100 | 2.41 | 98.1357 | 4966744 | 45.38 | 110 | 93154 |
| SAMN12767291-GCA_015784615.1 | PNUSAV001010 | SAMN12767291 | GCA_015784615.1 | NonPC | Public | NA | USA | Clinical | 91.2 | 100 | 1.27 | 98.3379 | 5136268 | 45.32 | 107 | 114121 |
| SAMN12767298-GCA_015785515.1 | PNUSAV000906 | SAMN12767298 | GCA_015785515.1 | NonPC | Public | NA | USA | Clinical | 88.9 | 100 | 1.91 | 98.0551 | 5109456 | 45.31 | 144 | 77517 |
| SAMN12767337-GCA_015785555.1 | PNUSAV000909 | SAMN12767337 | GCA_015785555.1 | NonPC | Public | NA | USA | Clinical | 91.2 | 100 | 0.07 | 98.3306 | 5120535 | 45.33 | 109 | 96021 |
| SAMN12771411-GCA_015784495.1 | PNUSAV001022 | SAMN12771411 | GCA_015784495.1 | NonPC | Public | NA | USA | Clinical | 90.5 | 100 | 0.79 | 98.3072 | 5206028 | 45.23 | 112 | 91549 |
| SAMN12771765-GCA_015785175.1 | PNUSAV001018 | SAMN12771765 | GCA_015785175.1 | NonPC | Public | NA | USA | Clinical | 91.8 | 100 | 4.81 | 98.2738 | 5248330 | 45.28 | 58 | 267750 |
| SAMN12781801-GCA_015785255.1 | PNUSAV001013 | SAMN12781801 | GCA_015785255.1 | NonPC | Public | NA | USA | Clinical | 90.6 | 100 | 2.97 | 98.3193 | 5186740 | 45.2 | 53 | 434910 |
| SAMN12783979-GCA_015784535.1 | CFSAN095172 | SAMN12783979 | GCA_015784535.1 | NonPC | Public | 2013 | USA | Env | 91.5 | 100 | 0 | 98.2895 | 5227924 | 45.3 | 274 | 59568 |
| SAMN12783981-GCA_015784475.1 | CFSAN095176 | SAMN12783981 | GCA_015784475.1 | NonPC | Public | 2013 | USA | Env | 90.7 | 100 | 0.15 | 98.3454 | 5065816 | 45.46 | 255 | 39771 |
| SAMN12783995-GCA_015785095.1 | CFSAN095177 | SAMN12783995 | GCA_015785095.1 | NonPC | Public | 2013 | USA | Env | 91.1 | 100 | 0.42 | 98.3408 | 5121427 | 45.39 | 166 | 76764 |
| SAMN12784254-GCA_015785135.1 | CFSAN095171 | SAMN12784254 | GCA_015785135.1 | NonPC | Public | 2013 | USA | Env | 90.2 | 100 | 0.67 | 98.3209 | 5037927 | 45.43 | 277 | 39662 |
| SAMN12784264-GCA_015785115.1 | CFSAN095178 | SAMN12784264 | GCA_015785115.1 | NonPC | Public | 2013 | USA | Env | 90.9 | 100 | 0.15 | 98.3867 | 5090503 | 45.43 | 237 | 48128 |
| SAMN12784269-GCA_015786155.1 | CFSAN095191 | SAMN12784269 | GCA_015786155.1 | NonPC | Public | 2013 | USA | Env | 90.2 | 100 | 0.25 | 98.36 | 5079725 | 45.38 | 226 | 60047 |
| SAMN12784271-GCA_015784335.1 | CFSAN095189 | SAMN12784271 | GCA_015784335.1 | NonPC | Public | 2013 | USA | Env | 90.1 | 99.86 | 0.15 | 98.3795 | 5033086 | 45.44 | 265 | 54544 |
| SAMN12784420-GCA_015782455.1 | CFSAN095193 | SAMN12784420 | GCA_015782455.1 | NonPC | Public | 2013 | USA | Env | 89.8 | 100 | 0.34 | 98.3257 | 5087236 | 45.39 | 256 | 53053 |
| SAMN12797616-GCA_015784515.1 | PNUSAV001031 | SAMN12797616 | GCA_015784515.1 | NonPC | Public | NA | USA | Clinical | 91.4 | 100 | 4.63 | 98.2994 | 5317144 | 45.26 | 40 | 556725 |
| SAMN12797630-GCA_015785635.1 | PNUSAV001030 | SAMN12797630 | GCA_015785635.1 | NonPC | Public | NA | USA | Clinical | 91.3 | 100 | 1.63 | 98.3083 | 5201243 | 45.24 | 43 | 564948 |
| SAMN12797632-GCA_015784435.1 | PNUSAV001032 | SAMN12797632 | GCA_015784435.1 | NonPC | Public | NA | USA | Clinical | 91.4 | 100 | 4.5 | 98.3067 | 5279492 | 45.26 | 43 | 564960 |
| SAMN12822629-GCA_015784035.1 | CFSAN095174 | SAMN12822629 | GCA_015784035.1 | NonPC | Public | 2013 | USA | Env | 91.1 | 99.86 | 0 | 98.3184 | 5186952 | 45.37 | 328 | 50136 |
| SAMN12822663-GCA_015784075.1 | CFSAN095179 | SAMN12822663 | GCA_015784075.1 | NonPC | Public | 2013 | USA | Env | 90.6 | 100 | 0.15 | 98.4068 | 5061030 | 45.49 | 305 | 35925 |
| SAMN12827004-GCA_015783335.1 | PNUSAV001036 | SAMN12827004 | GCA_015783335.1 | NonPC | Public | NA | USA | Clinical | 90.6 | 100 | 1.43 | 98.332 | 5109636 | 45.25 | 66 | 402750 |
| SAMN12842754-GCA_015783055.1 | PNUSAV001025 | SAMN12842754 | GCA_015783055.1 | NonPC | Public | NA | USA | Clinical | 91.4 | 100 | 0.54 | 98.3021 | 5183048 | 45.25 | 52 | 225965 |
| SAMN12842757-GCA_015783205.1 | PNUSAV001026 | SAMN12842757 | GCA_015783205.1 | PrePC | Public | NA | USA | Clinical | 94.2 | 100 | 4.63 | 99.7306 | 5137880 | 45.44 | 41 | 536363 |
| SAMN12856669-GCA_015783935.1 | 2019V-1160 | SAMN12856669 | GCA_015783935.1 | NonPC | Public | NA | USA | Clinical | 91.2 | 100 | 1.63 | 98.2725 | 5182398 | 45.3 | 109 | 89463 |
| SAMN12857386-GCA_015783155.1 | PNUSAV001081 | SAMN12857386 | GCA_015783155.1 | NonPC | Public | NA | USA | Clinical | 91.8 | 100 | 1.16 | 98.4004 | 5287951 | 45.3 | 78 | 232978 |
| SAMN12857387-GCA_015783235.1 | PNUSAV001083 | SAMN12857387 | GCA_015783235.1 | NonPC | Public | NA | USA | Clinical | 89.7 | 100 | 0.29 | 98.3462 | 5236641 | 45.2 | 99 | 186895 |
| SAMN12857419-GCA_015783835.1 | PNUSAV001085 | SAMN12857419 | GCA_015783835.1 | NonPC | Public | NA | USA | Clinical | 89.6 | 100 | 3.9 | 98.3129 | 5137127 | 45.37 | 67 | 170856 |
| SAMN12857420-GCA_015782435.1 | PNUSAV001088 | SAMN12857420 | GCA_015782435.1 | NonPC | Public | NA | USA | Clinical | 92 | 100 | 1.78 | 98.2692 | 5250313 | 45.23 | 63 | 316206 |
| SAMN12857422-GCA_015783975.1 | PNUSAV001064 | SAMN12857422 | GCA_015783975.1 | NonPC | Public | NA | USA | Clinical | 91.3 | 100 | 4.36 | 98.291 | 5289150 | 45.28 | 84 | 124892 |
| SAMN12873332-GCA_015783815.1 | PNUSAV001062 | SAMN12873332 | GCA_015783815.1 | NonPC | Public | NA | USA | Clinical | 91.4 | 100 | 3.81 | 98.3317 | 5234922 | 45.25 | 39 | 762179 |
| SAMN12873335-GCA_016240055.1 | PNUSAV001029 | SAMN12873335 | GCA_016240055.1 | NonPC | Public | NA | USA | Clinical | 91.4 | 100 | 2.32 | 98.2803 | 5225003 | 45.24 | 39 | 556714 |
| SAMN12879735-GCA_015783715.1 | PNUSAV001076 | SAMN12879735 | GCA_015783715.1 | NonPC | Public | NA | USA | Clinical | 91.2 | 100 | 0.36 | 98.3145 | 5106609 | 45.35 | 130 | 88555 |

| | | | | | | | | | | | | | | | | |
|---|---|---|---|---|---|---|---|---|---|---|---|---|---|---|---|---|
| SAMN12882180-GCA_015783115.1 | PNUSAV001074 | SAMN12882180 | GCA_015783115.1 | NonPC | Public | NA | USA | Clinical | 88.8 | 100 | 2.31 | 98.1911 | 5053218 | 45.45 | 124 | 93610 |
| SAMN12905839-GCA_015783075.1 | PNUSAV001095 | SAMN12905839 | GCA_015783075.1 | NonPC | Public | NA | USA | Clinical | 89.3 | 100 | 0.37 | 98.34 | 4990088 | 45.48 | 233 | 35889 |
| SAMN12909086-GCA_015783015.1 | PNUSAV001098 | SAMN12909086 | GCA_015783015.1 | NonPC | Public | NA | USA | Clinical | 89.5 | 100 | 1.47 | 98.2495 | 5255780 | 45.28 | 169 | 70194 |
| SAMN12911616-GCA_015782975.1 | PNUSAV001037 | SAMN12911616 | GCA_015782975.1 | NonPC | Public | NA | USA | Clinical | 91.3 | 100 | 3.54 | 98.3237 | 5296706 | 45.25 | 41 | 556734 |
| SAMN12915777-GCA_015782915.1 | PNUSAV001091 | SAMN12915777 | GCA_015782915.1 | NonPC | Public | NA | USA | Clinical | 90.1 | 100 | 3.81 | 98.3086 | 5154731 | 45.39 | 146 | 73988 |
| SAMN12930360-GCA_015782755.1 | PNUSAV001102 | SAMN12930360 | GCA_015782755.1 | NonPC | Public | NA | USA | Clinical | 90.8 | 100 | 0.67 | 98.3339 | 5185185 | 45.17 | 55 | 307659 |
| SAMN12930362-GCA_015782955.1 | PNUSAV001101 | SAMN12930362 | GCA_015782955.1 | NonPC | Public | NA | USA | Clinical | 91.4 | 100 | 0.07 | 98.2921 | 5154376 | 45.23 | 47 | 423502 |
| SAMN13000254-GCA_015782875.1 | PNUSAV001104 | SAMN13000254 | GCA_015782875.1 | NonPC | Public | NA | USA | Clinical | 91.3 | 100 | 0.31 | 98.2905 | 5151348 | 45.24 | 40 | 556727 |
| SAMN13000255-GCA_015782895.1 | PNUSAV001105 | SAMN13000255 | GCA_015782895.1 | NonPC | Public | NA | USA | Clinical | 91.3 | 100 | 0 | 98.3293 | 5137692 | 45.24 | 40 | 556737 |
| SAMN13008947-GCA_015782935.1 | PNUSAV001100 | SAMN13008947 | GCA_015782935.1 | NonPC | Public | NA | USA | Clinical | 89.8 | 100 | 0.41 | 98.2438 | 5145059 | 45.25 | 47 | 463026 |
| SAMN13008948-GCA_015782835.1 | PNUSAV001099 | SAMN13008948 | GCA_015782835.1 | NonPC | Public | NA | USA | Clinical | 91.5 | 100 | 1.43 | 98.3143 | 5135108 | 45.31 | 63 | 259693 |
| SAMN13008949-GCA_015782775.1 | PNUSAV001060 | SAMN13008949 | GCA_015782775.1 | NonPC | Public | NA | USA | Clinical | 91.2 | 100 | 0.58 | 98.255 | 5225653 | 45.2 | 39 | 529649 |
| SAMN13008951-GCA_015782855.1 | PNUSAV001112 | SAMN13008951 | GCA_015782855.1 | NonPC | Public | 2019 | USA | Env | 88.1 | 100 | 0.34 | 98.0583 | 5568237 | 45.27 | 318 | 37441 |
| SAMN13011472-GCA_016229185.1 | PNUSAV001110 | SAMN13011472 | GCA_016229185.1 | NonPC | Public | NA | USA | Clinical | 91.1 | 100 | 4.63 | 98.3065 | 5199228 | 45.29 | 65 | 367469 |
| SAMN13039307-GCA_015782675.1 | PNUSAV001118 | SAMN13039307 | GCA_015782675.1 | NonPC | Public | NA | USA | Clinical | 91.3 | 100 | 3.81 | 98.2556 | 5295932 | 45.27 | 56 | 236780 |
| SAMN13039308-GCA_015782715.1 | PNUSAV001117 | SAMN13039308 | GCA_015782715.1 | NonPC | Public | NA | USA | Clinical | 91.1 | 100 | 0.54 | 98.297 | 5145743 | 45.32 | 162 | 67595 |
| SAMN13039666-GCA_015782595.1 | PNUSAV001122 | SAMN13039666 | GCA_015782595.1 | NonPC | Public | 2019 | USA | Env | 90.6 | 100 | 2.27 | 98.352 | 5261219 | 45.28 | 206 | 56801 |
| SAMN13039718-GCA_015782695.1 | PNUSAV001121 | SAMN13039718 | GCA_015782695.1 | NonPC | Public | 2019 | USA | Env | 88.6 | 100 | 1.77 | 98.0537 | 5398663 | 45.26 | 140 | 89376 |
| SAMN13075360-GCA_015782575.1 | PNUSAV001126 | SAMN13075360 | GCA_015782575.1 | NonPC | Public | NA | USA | Clinical | 90.5 | 100 | 3.88 | 98.2875 | 5236090 | 45.27 | 44 | 377228 |
| SAMN13092740-GCA_015782735.1 | PNUSAV001128 | SAMN13092740 | GCA_015782735.1 | NonPC | Public | NA | USA | Clinical | 90.7 | 100 | 1.42 | 98.3195 | 5206547 | 45.23 | 69 | 157412 |
| SAMN13112824-GCA_015782555.1 | PNUSAV001131 | SAMN13112824 | GCA_015782555.1 | NonPC | Public | NA | USA | Clinical | 89.6 | 100 | 2.45 | 98.2648 | 5177642 | 45.34 | 137 | 83322 |
| SAMN13156552-GCA_015782635.1 | PNUSAV001125 | SAMN13156552 | GCA_015782635.1 | NonPC | Public | NA | USA | Clinical | 89.4 | 100 | 2.26 | 98.1213 | 5190657 | 45.18 | 87 | 169390 |
| SAMN13181012-SRR10413229 | NA | SAMN13181012 | SRR10413229 | PC | Public | 2004 | Japan | NA | 98.4 | 100 | 0 | 99.9957 | 5073896 | 45.29 | 77 | 288633 |
| SAMN13181013-SRR10413228 | NA | SAMN13181013 | SRR10413228 | PC | Public | 2011 | USA | NA | 98.4 | 100 | 0 | 99.9902 | 5072148 | 45.29 | 77 | 288633 |
| SAMN13181014-SRR10413227 | NA | SAMN13181014 | SRR10413227 | PC | Public | 2018 | USA | NA | 98.4 | 100 | 0 | 99.9951 | 5074587 | 45.28 | 77 | 288633 |
| SAMN13207724-GCA_016226745.1 | CFSAN095290 | SAMN13207724 | GCA_016226745.1 | NonPC | Public | 2013 | USA | Env | 90.8 | 100 | 4.75 | 98.2614 | 5281735 | 45.23 | 78 | 126326 |
| SAMN13207727-GCA_015783795.1 | CFSAN095277 | SAMN13207727 | GCA_015783795.1 | NonPC | Public | 2013 | USA | Env | 89 | 100 | 0.64 | 98.0769 | 5128619 | 45.34 | 138 | 88408 |
| SAMN13207728-GCA_016226825.1 | CFSAN095239 | SAMN13207728 | GCA_016226825.1 | NonPC | Public | 2013 | USA | Env | 89.6 | 100 | 1.62 | 98.3208 | 5196814 | 45.27 | 117 | 93049 |
| SAMN13245767-GCA_015782275.1 | CFSAN095241 | SAMN13245767 | GCA_015782275.1 | NonPC | Public | 2013 | USA | Env | 89.7 | 100 | 1.31 | 98.2864 | 5213451 | 45.21 | 30 | 462516 |
| SAMN13245769-GCA_015782315.1 | CFSAN095232 | SAMN13245769 | GCA_015782315.1 | NonPC | Public | 2013 | USA | Env | 88.9 | 100 | 0.2 | 98.1542 | 4988156 | 45.27 | 59 | 362239 |
| SAMN13245770-GCA_015782295.1 | CFSAN095251 | SAMN13245770 | GCA_015782295.1 | NonPC | Public | 2013 | USA | Env | 91.6 | 100 | 0.08 | 98.2027 | 5106456 | 45.27 | 41 | 543021 |
| SAMN13245774-GCA_015782175.1 | CFSAN095238 | SAMN13245774 | GCA_015782175.1 | NonPC | Public | 2013 | USA | Env | 89.7 | 100 | 0.35 | 98.319 | 5165314 | 45.21 | 37 | 462516 |
| SAMN13245778-GCA_015782335.1 | CFSAN095246 | SAMN13245778 | GCA_015782335.1 | NonPC | Public | 2013 | USA | Env | 90.2 | 100 | 1.25 | 98.2777 | 5210305 | 45.21 | 26 | 507969 |
| SAMN13245779-GCA_015781535.1 | CFSAN095254 | SAMN13245779 | GCA_015781535.1 | NonPC | Public | 2013 | USA | Env | 92.3 | 100 | 0.64 | 98.3381 | 5410696 | 44.97 | 41 | 541887 |
| SAMN13245780-GCA_015782115.1 | CFSAN095248 | SAMN13245780 | GCA_015782115.1 | NonPC | Public | 2013 | USA | Env | 91.6 | 100 | 0.77 | 98.2262 | 5177761 | 45.26 | 41 | 571895 |
| SAMN13245782-GCA_016225145.1 | CFSAN095200 | SAMN13245782 | GCA_016225145.1 | NonPC | Public | 2013 | USA | Env | 91.3 | 100 | 0.61 | 98.2332 | 5035776 | 45.36 | 42 | 438046 |
| SAMN13245784-GCA_015782055.1 | CFSAN095247 | SAMN13245784 | GCA_015782055.1 | NonPC | Public | 2013 | USA | Env | 90.2 | 100 | 0.61 | 98.2856 | 5196008 | 45.2 | 32 | 507969 |
| SAMN13245786-GCA_015782135.1 | CFSAN095235 | SAMN13245786 | GCA_015782135.1 | NonPC | Public | 2013 | USA | Env | 89.7 | 100 | 0.29 | 98.2759 | 5163185 | 45.21 | 37 | 411328 |
| SAMN13245792-GCA_015782195.1 | CFSAN095234 | SAMN13245792 | GCA_015782195.1 | NonPC | Public | 2013 | USA | Env | 89.7 | 100 | 0.3 | 98.2519 | 5168587 | 45.21 | 35 | 439705 |
| SAMN13245794-GCA_015781515.1 | CFSAN095201 | SAMN13245794 | GCA_015781515.1 | NonPC | Public | 2013 | USA | Env | 89.7 | 100 | 0.42 | 98.3108 | 5195643 | 45.21 | 44 | 336499 |
| SAMN13245796-GCA_015781595.1 | CFSAN095244 | SAMN13245796 | GCA_015781595.1 | NonPC | Public | 2013 | USA | Env | 87.4 | 100 | 2.6 | 98.0898 | 5343500 | 45.06 | 106 | 337522 |
| SAMN13245799-GCA_015782255.1 | CFSAN095237 | SAMN13245799 | GCA_015782255.1 | NonPC | Public | 2013 | USA | Env | 89.7 | 100 | 0.69 | 98.3022 | 5171446 | 45.2 | 36 | 452781 |
| SAMN13245800-GCA_015782235.1 | CFSAN095252 | SAMN13245800 | GCA_015782235.1 | NonPC | Public | 2013 | USA | Env | 92.3 | 100 | 0.73 | 98.3725 | 5382104 | 44.95 | 36 | 548387 |
| SAMN13245804-GCA_015779745.1 | CFSAN095249 | SAMN13245804 | GCA_015779745.1 | NonPC | Public | 2013 | USA | Env | 88.6 | 100 | 1.17 | 98.1581 | 5083663 | 45.3 | 51 | 458026 |
| SAMN13245809-GCA_015781655.1 | CFSAN095240 | SAMN13245809 | GCA_015781655.1 | NonPC | Public | 2013 | USA | Env | 89.7 | 100 | 2.25 | 98.3019 | 5233791 | 45.2 | 31 | 549472 |
| SAMN13245812-GCA_015779775.1 | CFSAN095253 | SAMN13245812 | GCA_015779775.1 | NonPC | Public | 2013 | USA | Env | 92.3 | 100 | 0.27 | 98.3343 | 5380319 | 44.95 | 43 | 460282 |
| SAMN13245813-GCA_015782075.1 | CFSAN095233 | SAMN13245813 | GCA_015782075.1 | NonPC | Public | 2013 | USA | Env | 88.9 | 100 | 0.82 | 98.1461 | 4993040 | 45.28 | 56 | 335820 |
| SAMN13245814-GCA_015781615.1 | CFSAN095250 | SAMN13245814 | GCA_015781615.1 | NonPC | Public | 2013 | USA | Env | 91.6 | 100 | 1.35 | 98.2359 | 5192314 | 45.28 | 36 | 543021 |

| | | | | | | | | | | | | | | | | |
|---|---|---|---|---|---|---|---|---|---|---|---|---|---|---|---|---|
| SAMN13262823-GCA_015781565.1 | PNUSAV001135 | SAMN13262823 | GCA_015781565.1 | NonPC | Public | NA | USA | Clinical | 91.4 | 100 | 3.8 | 98.4057 | 5086560 | 45.4 | 47 | 409202 |
| SAMN13282664-SRR10441589 | 769807 | SAMN13282664 | SRR10441589 | NonPC | Public | 2019 | NA | Clinical | 91.4 | 100 | 0 | 98.2992 | 5117317 | 45.24 | 61 | 351325 |
| SAMN13282666-SRR10441577 | 786602 | SAMN13282666 | SRR10441577 | PC | Public | 2019 | NA | Clinical | 97.7 | 100 | 0.27 | 99.9777 | 5085246 | 45.3 | 59 | 290456 |
| SAMN13283446-GCA_015779685.1 | CFSAN095157 | SAMN13283446 | GCA_015779685.1 | NonPC | Public | 2013 | USA | Env | 91.3 | 100 | 0.14 | 98.2869 | 5141431 | 45.32 | 35 | 548414 |
| SAMN13283453-GCA_015779545.1 | CFSAN095261 | SAMN13283453 | GCA_015779545.1 | NonPC | Public | 2013 | USA | Env | 89.8 | 100 | 4.4 | 98.2726 | 5390394 | 45.2 | 40 | 349623 |
| SAMN13283455-GCA_015781855.1 | CFSAN095267 | SAMN13283455 | GCA_015781855.1 | NonPC | Public | 2013 | USA | Env | 89.7 | 100 | 0.73 | 98.2626 | 5304636 | 45.2 | 39 | 371805 |
| SAMN13283457-GCA_015781635.1 | CFSAN095153 | SAMN13283457 | GCA_015781635.1 | NonPC | Public | 2013 | USA | Env | 91.7 | 100 | 2.31 | 98.3592 | 5354084 | 45.25 | 39 | 385511 |
| SAMN13283460-GCA_015781875.1 | CFSAN095152 | SAMN13283460 | GCA_015781875.1 | NonPC | Public | 2013 | USA | Env | 90.4 | 100 | 0.4 | 98.2522 | 5111018 | 45.24 | 47 | 457752 |
| SAMN13283463-GCA_015779605.1 | CFSAN095256 | SAMN13283463 | GCA_015779605.1 | NonPC | Public | 2013 | USA | Env | 91.4 | 100 | 1.35 | 98.2112 | 5423819 | 45.2 | 44 | 459246 |
| SAMN13283464-GCA_015781335.1 | CFSAN095155 | SAMN13283464 | GCA_015781335.1 | NonPC | Public | 2013 | USA | Env | 91.7 | 100 | 2.07 | 98.3634 | 5344750 | 45.24 | 39 | 402843 |
| SAMN13283469-GCA_015781815.1 | CFSAN095226 | SAMN13283469 | GCA_015781815.1 | NonPC | Public | 2013 | USA | Env | 89.6 | 100 | 1.24 | 98.1181 | 5228421 | 45.15 | 62 | 232835 |
| SAMN13283471-GCA_015779305.1 | CFSAN095159 | SAMN13283471 | GCA_015779305.1 | NonPC | Public | 2013 | USA | Env | 91.6 | 100 | 0.3 | 98.3672 | 5262161 | 45.17 | 40 | 466269 |
| SAMN13283472-GCA_015779165.1 | CFSAN095220 | SAMN13283472 | GCA_015779165.1 | NonPC | Public | 2013 | USA | Env | 89.5 | 100 | 0.08 | 98.1552 | 5143825 | 45.22 | 168 | 69557 |
| SAMN13283473-GCA_015779045.1 | CFSAN095162 | SAMN13283473 | GCA_015779045.1 | NonPC | Public | 2013 | USA | Env | 91.6 | 100 | 1.52 | 98.3874 | 5298949 | 45.18 | 40 | 466533 |
| SAMN13283474-GCA_015779065.1 | CFSAN095265 | SAMN13283474 | GCA_015779065.1 | NonPC | Public | 2013 | USA | Env | 89.9 | 100 | 0.18 | 98.2715 | 5233403 | 45.18 | 36 | 464329 |
| SAMN13283475-GCA_015779025.1 | CFSAN095173 | SAMN13283475 | GCA_015779025.1 | NonPC | Public | 2013 | USA | Env | 91.1 | 100 | 0.75 | 98.2219 | 5115892 | 45.25 | 33 | 595564 |
| SAMN13283476-GCA_016223425.1 | CFSAN095263 | SAMN13283476 | GCA_016223425.1 | NonPC | Public | 2013 | USA | Env | 89.7 | 100 | 0.24 | 98.2509 | 5228496 | 45.18 | 50 | 349623 |
| SAMN13283477-GCA_015779145.1 | CFSAN095229 | SAMN13283477 | GCA_015779145.1 | NonPC | Public | 2013 | USA | Env | 88.9 | 100 | 0.68 | 98.1321 | 4988328 | 45.27 | 59 | 335819 |
| SAMN13283478-GCA_015781835.1 | CFSAN095231 | SAMN13283478 | GCA_015781835.1 | NonPC | Public | 2013 | USA | Env | 88.8 | 100 | 0.27 | 98.1734 | 4978080 | 45.27 | 58 | 335997 |
| SAMN13283480-GCA_015778925.1 | CFSAN095195 | SAMN13283480 | GCA_015778925.1 | NonPC | Public | 2013 | USA | Env | 90.4 | 100 | 0.73 | 98.2904 | 5179982 | 45.22 | 35 | 525326 |
| SAMN13283482-GCA_015779105.1 | CFSAN095164 | SAMN13283482 | GCA_015779105.1 | NonPC | Public | 2013 | USA | Env | 91.1 | 100 | 0.86 | 98.2827 | 5322358 | 45.26 | 38 | 476209 |
| SAMN13283483-GCA_015779085.1 | CFSAN095268 | SAMN13283483 | GCA_015779085.1 | NonPC | Public | 2013 | USA | Env | 89.8 | 100 | 0.11 | 98.3023 | 5246454 | 45.18 | 36 | 464232 |
| SAMN13283484-GCA_015778985.1 | CFSAN095259 | SAMN13283484 | GCA_015778985.1 | NonPC | Public | 2013 | USA | Env | 89.7 | 100 | 0.4 | 98.317 | 5174920 | 45.2 | 42 | 394945 |
| SAMN13283485-GCA_015779005.1 | CFSAN095163 | SAMN13283485 | GCA_015779005.1 | NonPC | Public | 2013 | USA | Env | 91.1 | 100 | 0.68 | 98.2577 | 5317673 | 45.26 | 41 | 399672 |
| SAMN13283486-GCA_015779115.1 | CFSAN095227 | SAMN13283486 | GCA_015779115.1 | NonPC | Public | 2013 | USA | Env | 89.6 | 100 | 2.13 | 98.1037 | 5211069 | 45.14 | 64 | 222356 |
| SAMN13283488-GCA_015781755.1 | CFSAN095230 | SAMN13283488 | GCA_015781755.1 | NonPC | Public | 2013 | USA | Env | 88.8 | 100 | 0.5 | 98.1342 | 4989412 | 45.28 | 59 | 286692 |
| SAMN13283490-GCA_015781775.1 | CFSAN095154 | SAMN13283490 | GCA_015781775.1 | NonPC | Public | 2013 | USA | Env | 91.6 | 100 | 4.04 | 98.3489 | 5389729 | 45.33 | 132 | 93454 |
| SAMN13283491-GCA_015781715.1 | CFSAN095217 | SAMN13283491 | GCA_015781715.1 | NonPC | Public | 2013 | USA | Env | 90.6 | 100 | 0.27 | 98.2971 | 5148860 | 45.3 | 130 | 88849 |
| SAMN13283492-GCA_015779185.1 | CFSAN095161 | SAMN13283492 | GCA_015779185.1 | NonPC | Public | 2013 | USA | Env | 91.6 | 100 | 0.99 | 98.3626 | 5262795 | 45.17 | 41 | 466269 |
| SAMN13283494-GCA_015781895.1 | CFSAN095196 | SAMN13283494 | GCA_015781895.1 | NonPC | Public | 2013 | USA | Env | 90.4 | 100 | 0.77 | 98.2781 | 5169984 | 45.22 | 42 | 459401 |
| SAMN13283497-GCA_015779225.1 | CFSAN095198 | SAMN13283497 | GCA_015779225.1 | NonPC | Public | 2013 | USA | Env | 90.5 | 100 | 0.61 | 98.2486 | 5292134 | 45.12 | 46 | 410594 |
| SAMN13283498-GCA_015779245.1 | CFSAN095160 | SAMN13283498 | GCA_015779245.1 | NonPC | Public | 2013 | USA | Env | 91.6 | 100 | 4.78 | 98.3883 | 5306540 | 45.27 | 37 | 466429 |
| SAMN13283499-GCA_015781675.1 | CFSAN095225 | SAMN13283499 | GCA_015781675.1 | NonPC | Public | 2013 | USA | Env | 89.6 | 100 | 2.53 | 98.1097 | 5221558 | 45.15 | 68 | 203682 |
| SAMN13283500-GCA_015779205.1 | CFSAN095255 | SAMN13283500 | GCA_015779205.1 | NonPC | Public | 2013 | USA | Env | 91.3 | 100 | 0.69 | 98.2507 | 5040918 | 45.37 | 43 | 456345 |
| SAMN13283501-GCA_015779265.1 | CFSAN095212 | SAMN13283501 | GCA_015779265.1 | NonPC | Public | 2013 | USA | Env | 90.7 | 100 | 2.11 | 98.3289 | 5194970 | 45.29 | 125 | 83672 |
| SAMN13283504-GCA_015778965.1 | CFSAN095260 | SAMN13283504 | GCA_015778965.1 | NonPC | Public | 2013 | USA | Env | 91.1 | 100 | 0.68 | 98.2386 | 5345405 | 45.26 | 31 | 595816 |
| SAMN13283505-GCA_015781735.1 | CFSAN095266 | SAMN13283505 | GCA_015781735.1 | NonPC | Public | 2013 | USA | Env | 89.8 | 100 | 3.04 | 98.2454 | 5353965 | 45.2 | 38 | 464329 |
| SAMN13283506-GCA_015781685.1 | CFSAN095223 | SAMN13283506 | GCA_015781685.1 | NonPC | Public | 2013 | USA | Env | 89.5 | 100 | 4.99 | 98.1579 | 5371406 | 45.23 | 131 | 86399 |
| SAMN13283507-GCA_015781915.1 | CFSAN095224 | SAMN13283507 | GCA_015781915.1 | NonPC | Public | 2013 | USA | Env | 89.5 | 100 | 3.53 | 98.1419 | 5284622 | 45.22 | 134 | 93997 |
| SAMN13283509-GCA_015779465.1 | CFSAN095264 | SAMN13283509 | GCA_015779465.1 | NonPC | Public | 2013 | USA | Env | 89.9 | 100 | 0.11 | 98.27 | 5227749 | 45.19 | 39 | 464319 |
| SAMN13283512-GCA_015781435.1 | CFSAN095213 | SAMN13283512 | GCA_015781435.1 | NonPC | Public | 2013 | USA | Env | 90.6 | 100 | 0.95 | 98.2847 | 5145367 | 45.31 | 158 | 77184 |
| SAMN13283514-GCA_015779645.1 | CFSAN095214 | SAMN13283514 | GCA_015779645.1 | NonPC | Public | 2013 | USA | Env | 90.6 | 100 | 0.72 | 98.2703 | 5162958 | 45.31 | 120 | 97523 |
| SAMN13283515-GCA_015781475.1 | CFSAN095156 | SAMN13283515 | GCA_015781475.1 | NonPC | Public | 2013 | USA | Env | 91.7 | 100 | 1.76 | 98.3645 | 5332179 | 45.24 | 41 | 385501 |
| SAMN13283516-GCA_015781495.1 | CFSAN095167 | SAMN13283516 | GCA_015781495.1 | NonPC | Public | 2013 | USA | Env | 91 | 100 | 0.89 | 98.2879 | 5331716 | 45.26 | 38 | 476221 |
| SAMN13283519-GCA_015781455.1 | CFSAN095218 | SAMN13283519 | GCA_015781455.1 | NonPC | Public | 2013 | USA | Env | 90.6 | 100 | 0.68 | 98.2946 | 5162300 | 45.3 | 136 | 83402 |
| SAMN13283520-GCA_015781555.1 | CFSAN095211 | SAMN13283520 | GCA_015781555.1 | NonPC | Public | 2013 | USA | Env | 90.9 | 100 | 0.54 | 98.3177 | 5139420 | 45.4 | 128 | 94505 |
| SAMN13283524-GCA_015779625.1 | CFSAN095192 | SAMN13283524 | GCA_015779625.1 | NonPC | Public | 2013 | USA | Env | 90.2 | 100 | 0.82 | 98.3087 | 5184239 | 45.19 | 32 | 487738 |
| SAMN13283526-GCA_015781975.1 | CFSAN095194 | SAMN13283526 | GCA_015781975.1 | NonPC | Public | 2013 | USA | Env | 90.4 | 100 | 0.77 | 98.284 | 5182037 | 45.22 | 41 | 525326 |

| | | | | | | | | | | | | | | | | |
|---|---|---|---|---|---|---|---|---|---|---|---|---|---|---|---|---|
| SAMN13283527-GCA_015779335.1 | CFSAN095215 | SAMN13283527 | GCA_015779335.1 | NonPC | Public | 2013 | USA | Env | 90.6 | 100 | 1.36 | 98.2921 | 5160732 | 45.3 | 148 | 83788 |
| SAMN13283540-GCA_015781395.1 | CFSAN095222 | SAMN13283540 | GCA_015781395.1 | NonPC | Public | 2013 | USA | Env | 89.5 | 100 | 3.4 | 98.1744 | 5181466 | 45.31 | 145 | 83160 |
| SAMN13283544-GCA_015781415.1 | CFSAN095216 | SAMN13283544 | GCA_015781415.1 | NonPC | Public | 2013 | USA | Env | 90.6 | 100 | 0.83 | 98.2631 | 5185880 | 45.3 | 128 | 87257 |
| SAMN13283549-GCA_015779365.1 | CFSAN095221 | SAMN13283549 | GCA_015779365.1 | NonPC | Public | 2013 | USA | Env | 89.4 | 100 | 0.15 | 98.1797 | 5153379 | 45.25 | 183 | 60470 |
| SAMN13388561-SRR10531460 | 2 | SAMN13388561 | SRR10531460 | PC | Public | 2019 | UK | Clinical | 96.5 | 100 | 0.27 | 99.9742 | 5014646 | 45.51 | 384 | 26201 |
| SAMN13426200-GCF_009734325.1 | 19-021-D1 | SAMN13426200 | GCF_009734325.1 | NonPC | Public | 2019 | South_Korea | Env | 92.8 | 100 | 0.27 | 98.2673 | 5579380 | 45.31 | 4 | 3577848 |
| SAMN13481054-GCA_015779405.1 | 19MO10SH04-V1 | SAMN13481054 | GCA_015779405.1 | NonPC | Public | 2019 | USA | Env | 89.9 | 100 | 0.19 | 98.3324 | 5321667 | 45.5 | 354 | 29126 |
| SAMN13481055-GCA_015779425.1 | 19MO10SH04-V2 | SAMN13481055 | GCA_015779425.1 | NonPC | Public | 2019 | USA | Env | 88.3 | 100 | 1.36 | 98.0066 | 5661613 | 45.15 | 251 | 45525 |
| SAMN13481057-GCA_016216265.1 | 19MO10SH05-V1 | SAMN13481057 | GCA_016216265.1 | NonPC | Public | 2019 | USA | Env | 89.3 | 100 | 0.83 | 98.3116 | 5181875 | 45.37 | 181 | 67171 |
| SAMN13481061-GCA_015779445.1 | 19MO10SH07-V1 | SAMN13481061 | GCA_015779445.1 | NonPC | Public | 2019 | USA | Env | 91.4 | 100 | 1.74 | 98.2553 | 5141396 | 45.39 | 171 | 67860 |
| SAMN13481063-GCA_015780315.1 | 19MO10SH02-V1 | SAMN13481063 | GCA_015780315.1 | NonPC | Public | 2019 | USA | Env | 87.7 | 100 | 1.62 | 98.1922 | 5466522 | 45.48 | 188 | 60617 |
| SAMN13500485-GCF_014525115.1 | 2013V-1125 | SAMN13500485 | GCF_014525115.1 | NonPC | Public | NA | USA | Clinical | 92 | 99.13 | 0 | 98.3032 | 5166012 | 45.33 | 5 | 1580243 |
| SAMN13500486-GCF_014525225.1 | 2013V-1220 | SAMN13500486 | GCF_014525225.1 | NonPC | Public | NA | USA | Clinical | 90.5 | 100 | 0.11 | 98.3273 | 5082201 | 45.44 | 3 | 3143777 |
| SAMN13560645-GCA_015781935.1 | PNUSAV001147 | SAMN13560645 | GCA_015781935.1 | NonPC | Public | NA | USA | Clinical | 90.5 | 100 | 0.75 | 98.3112 | 5159030 | 45.24 | 67 | 313657 |
| SAMN13564661-GCA_017312095.1 | 1.004-13 | SAMN13564661 | GCA_017312095.1 | NonPC | Public | 2012 | Peru | Clinical | 94.3 | 98.64 | 1.79 | 98.3121 | 5473199 | 45.14 | 298 | 286275 |
| SAMN13621685-GCA_015780635.1 | PNUSAV001145 | SAMN13621685 | GCA_015780635.1 | NonPC | Public | 2019 | USA | Clinical | 89.7 | 100 | 3.43 | 98.2766 | 5217542 | 45.34 | 118 | 89459 |
| SAMN13621686-GCA_015780655.1 | PNUSAV001142 | SAMN13621686 | GCA_015780655.1 | NonPC | Public | 2019 | USA | Clinical | 89.5 | 100 | 1.67 | 98.244 | 5044949 | 45.42 | 61 | 228524 |
| SAMN13621687-GCA_015780395.1 | PNUSAV001143 | SAMN13621687 | GCA_015780395.1 | NonPC | Public | 2019 | USA | Clinical | 88.8 | 100 | 2.18 | 98.1609 | 5135024 | 45.32 | 138 | 84059 |
| SAMN13621688-GCA_015780695.1 | PNUSAV001144 | SAMN13621688 | GCA_015780695.1 | NonPC | Public | 2019 | USA | Clinical | 89.8 | 100 | 1.7 | 98.3421 | 5014285 | 45.42 | 105 | 105351 |
| SAMN13621689-GCA_015780435.1 | PNUSAV001146 | SAMN13621689 | GCA_015780435.1 | NonPC | Public | 2019 | USA | Clinical | 89 | 100 | 0.82 | 98.3486 | 4979868 | 45.44 | 102 | 122446 |
| SAMN13633981-SRR10738143 | NA | SAMN13633981 | SRR10738143 | NonPC | Public | 2013 | USA | Clinical | 91.5 | 100 | 0.03 | 98.3079 | 5089012 | 45.26 | 76 | 260702 |
| SAMN13633983-GCA_015780375.1 | CTVP29C | SAMN13633983 | GCA_015780375.1 | NonPC | Public | 2013 | USA | Clinical | 88.7 | 100 | 0.64 | 98.1469 | 4915809 | 45.45 | 86 | 192139 |
| SAMN13633984-GCA_015780495.1 | G61 | SAMN13633984 | GCA_015780495.1 | NonPC | Public | 2007 | USA | Env | 89.6 | 100 | 0.08 | 98.2315 | 5094346 | 45.29 | 60 | 265701 |
| SAMN13633985-GCA_015780355.1 | G325 | SAMN13633985 | GCA_015780355.1 | NonPC | Public | 2008 | USA | NA | 89.6 | 100 | 1.16 | 98.3008 | 5001266 | 45.43 | 93 | 116812 |
| SAMN13633986-SRR10738125 | G363 | SAMN13633986 | SRR10738125 | NonPC | Public | 2008 | USA | Env | 90.5 | 100 | 0.27 | 98.2931 | 5142905 | 45.21 | 28 | 702748 |
| SAMN13633987-GCA_015780595.1 | G445 | SAMN13633987 | GCA_015780595.1 | NonPC | Public | 2008 | USA | Env | 87.2 | 100 | 3.24 | 98.2607 | 5130373 | 45.25 | 89 | 213696 |
| SAMN13633988-GCA_015780575.1 | G729 | SAMN13633988 | GCA_015780575.1 | NonPC | Public | 2008 | USA | Env | 89.8 | 100 | 0.85 | 98.3443 | 4983988 | 45.39 | 104 | 100333 |
| SAMN13633989-SRR10738122 | NA | SAMN13633989 | SRR10738122 | NonPC | Public | 2009 | USA | Env | 90.4 | 100 | 0.27 | 98.2753 | 5132725 | 45.2 | 36 | 538449 |
| SAMN13633990-GCA_015780555.1 | G6499 | SAMN13633990 | GCA_015780555.1 | NonPC | Public | 2015 | USA | Env | 89.4 | 100 | 0.34 | 98.3124 | 4992279 | 45.48 | 175 | 58014 |
| SAMN13633992-SRR10738140 | NA | SAMN13633992 | SRR10738140 | NonPC | Public | 2013 | USA | Clinical | 91.8 | 100 | 0 | 98.2603 | 5271487 | 45.13 | 78 | 436434 |
| SAMN13633993-GCA_015780295.1 | MAVP-15 | SAMN13633993 | GCA_015780295.1 | NonPC | Public | 2013 | NA | Clinical | 89.2 | 100 | 0.35 | 98.2402 | 5154743 | 45.2 | 298 | 38425 |
| SAMN13633997-GCA_015780615.1 | MAVP-H | SAMN13633997 | GCA_015780615.1 | NonPC | Public | 2011 | USA | Clinical | 91.6 | 100 | 0.08 | 98.272 | 5089503 | 45.33 | 103 | 100859 |
| SAMN13633998-SRR10738134 | NA | SAMN13633998 | SRR10738134 | NonPC | Public | 2011 | USA | Clinical | 89.5 | 100 | 0.08 | 98.2645 | 5124837 | 45.18 | 86 | 232803 |
| SAMN13634000-GCA_015780675.1 | MAVP-Y | SAMN13634000 | GCA_015780675.1 | NonPC | Public | 2011 | USA | Clinical | 89.2 | 100 | 2.45 | 98.0445 | 5130994 | 45.29 | 79 | 164193 |
| SAMN13634001-GCA_015780275.1 | MDOH-04-5M732 | SAMN13634001 | GCA_015780275.1 | PC | Public | 2004 | USA | Clinical | 97.2 | 100 | 3.81 | 99.935 | 5308279 | 45.14 | 70 | 227203 |
| SAMN13634002-GCA_015780455.1 | G747 | SAMN13634002 | GCA_015780455.1 | NonPC | Public | 2008 | USA | Env | 89 | 100 | 0.07 | 98.323 | 4954519 | 45.44 | 263 | 36989 |
| SAMN13634003-GCA_015780475.1 | G755 | SAMN13634003 | GCA_015780475.1 | NonPC | Public | 2008 | USA | Env | 91 | 100 | 0.89 | 98.3334 | 5090677 | 45.32 | 77 | 316176 |
| SAMN13634675-GCA_015780115.1 | CTVP16C | SAMN13634675 | GCA_015780115.1 | NonPC | Public | 2012 | USA | Clinical | 91.2 | 100 | 0 | 98.3394 | 5085921 | 45.27 | 74 | 204772 |
| SAMN13634676-SRR10740835 | CT4287 | SAMN13634676 | SRR10740835 | NonPC | Public | 2013 | USA | NA | 90.6 | 100 | 0 | 98.2639 | 5228435 | 45.19 | 53 | 462048 |
| SAMN13634677-SRR10740826 | CT4291 | SAMN13634677 | SRR10740826 | NonPC | Public | 2013 | USA | NA | 90.6 | 100 | 0 | 98.3103 | 5228989 | 45.18 | 38 | 457403 |
| SAMN13634679-GCA_015779855.1 | G145 | SAMN13634679 | GCA_015779855.1 | NonPC | Public | 2007 | USA | NA | 89.7 | 100 | 2.38 | 98.2948 | 5135179 | 45.32 | 76 | 161757 |
| SAMN13634680-GCA_015780175.1 | G151 | SAMN13634680 | GCA_015780175.1 | NonPC | Public | 2007 | USA | Env | 91.5 | 100 | 0.07 | 98.3929 | 5013628 | 45.35 | 53 | 314879 |
| SAMN13634681-GCA_015779975.1 | G320 | SAMN13634681 | GCA_015779975.1 | NonPC | Public | 2008 | USA | NA | 89.4 | 99.81 | 0.08 | 98.2566 | 5070889 | 45.24 | 151 | 84158 |
| SAMN13634682-GCA_015780235.1 | G441 | SAMN13634682 | GCA_015780235.1 | NonPC | Public | 2008 | USA | NA | 89.8 | 100 | 0.42 | 98.2868 | 5009812 | 45.35 | 78 | 190189 |
| SAMN13634683-GCA_015780095.1 | G524 | SAMN13634683 | GCA_015780095.1 | NonPC | Public | 2008 | USA | NA | 89.6 | 100 | 0.64 | 98.3323 | 4995738 | 45.35 | 49 | 345759 |
| SAMN13634684-GCA_015779995.1 | G640 | SAMN13634684 | GCA_015779995.1 | NonPC | Public | 2008 | USA | NA | 89.5 | 100 | 0.07 | 98.3506 | 4970535 | 45.39 | 102 | 116548 |
| SAMN13634685-GCA_015779795.1 | G650 | SAMN13634685 | GCA_015779795.1 | NonPC | Public | 2008 | USA | Env | 89.5 | 100 | 1.68 | 98.3558 | 5133660 | 45.27 | 49 | 438601 |
| SAMN13634686-GCA_015780035.1 | G653 | SAMN13634686 | GCA_015780035.1 | NonPC | Public | 2008 | USA | Env | 89.6 | 100 | 1.7 | 98.3433 | 5068720 | 45.35 | 80 | 165092 |

| | | | | | | | | | | | | | | | | |
|---|---|---|---|---|---|---|---|---|---|---|---|---|---|---|---|---|
| SAMN13634687-GCA_015779955.1 | G756 | SAMN13634687 | GCA_015779955.1 | NonPC | Public | 2008 | USA | Env | 89.6 | 100 | 0.9 | 98.336 | 5016363 | 45.35 | 47 | 369177 |
| SAMN13634688-GCA_015779935.1 | G760 | SAMN13634688 | GCA_015779935.1 | NonPC | Public | 2008 | USA | Env | 89.8 | 100 | 0.94 | 98.3107 | 5015190 | 45.33 | 61 | 286335 |
| SAMN13634690-GCA_015780015.1 | G1355 | SAMN13634690 | GCA_015780015.1 | NonPC | Public | 2009 | USA | Env | 91.6 | 100 | 4.13 | 98.402 | 5286658 | 45.31 | 57 | 250825 |
| SAMN13634691-SRR10740828 | G3673 | SAMN13634691 | SRR10740828 | NonPC | Public | 2013 | USA | Env | 90.5 | 100 | 0.14 | 98.1875 | 5042125 | 45.2 | 35 | 525258 |
| SAMN13634692-GCA_015779835.1 | G6928 | SAMN13634692 | GCA_015779835.1 | NonPC | Public | 2015 | USA | Env | 89.3 | 100 | 0.65 | 98.2559 | 5216429 | 45.11 | 66 | 266168 |
| SAMN13649479-GCA_015777925.1 | G1386 | SAMN13649479 | GCA_015777925.1 | NonPC | Public | 2009 | USA | Env | 89.5 | 100 | 0.72 | 98.2955 | 5012983 | 45.35 | 37 | 420186 |
| SAMN13649480-GCA_015778865.1 | G1393 | SAMN13649480 | GCA_015778865.1 | NonPC | Public | 2009 | USA | Env | 91 | 100 | 2.32 | 98.3496 | 5084951 | 45.31 | 75 | 367850 |
| SAMN13649481-GCA_015778085.1 | G1445 | SAMN13649481 | GCA_015778085.1 | NonPC | Public | 2009 | USA | Env | 88.1 | 100 | 1.27 | 98.1828 | 5234682 | 45.16 | 66 | 413920 |
| SAMN13649482-GCA_015778325.1 | G1487 | SAMN13649482 | GCA_015778325.1 | NonPC | Public | 2009 | USA | Env | 89.6 | 100 | 2.45 | 98.3242 | 5071106 | 45.35 | 53 | 397626 |
| SAMN13649483-SRR10744602 | G3654 | SAMN13649483 | SRR10744602 | NonPC | Public | 2013 | USA | Env | 88.4 | 100 | 0.03 | 98.0825 | 5108309 | 45.26 | 35 | 441430 |
| SAMN13649484-SRR10744601 | G3599 | SAMN13649484 | SRR10744601 | NonPC | Public | 2013 | USA | Env | 90.4 | 100 | 0 | 98.273 | 5219537 | 45.16 | 45 | 764309 |
| SAMN13649485-SRR10744600 | G4026 | SAMN13649485 | SRR10744600 | NonPC | Public | 2013 | USA | Env | 88.6 | 100 | 0.03 | 98.1097 | 5038562 | 45.29 | 52 | 556600 |
| SAMN13649489-GCA_015778845.1 | MA118 | SAMN13649489 | GCA_015778845.1 | NonPC | Public | 2014 | USA | NA | 88.5 | 100 | 3.34 | 98.0512 | 5048519 | 45.36 | 74 | 292700 |
| SAMN13649494-GCA_015778065.1 | MA157 | SAMN13649494 | GCA_015778065.1 | NonPC | Public | 2014 | USA | NA | 90.8 | 100 | 4.12 | 98.1992 | 5334621 | 45.31 | 66 | 330427 |
| SAMN13649497-GCA_015778185.1 | MA267 | SAMN13649497 | GCA_015778185.1 | NonPC | Public | 2015 | USA | NA | 90.6 | 99.5 | 1.39 | 98.3339 | 5383477 | 45.06 | 271 | 68024 |
| SAMN13649498-GCA_015778745.1 | MA271 | SAMN13649498 | GCA_015778745.1 | NonPC | Public | 2015 | USA | NA | 91.3 | 100 | 1.38 | 98.2545 | 5253572 | 45.17 | 71 | 218563 |
| SAMN13649501-GCA_015778165.1 | MA371 | SAMN13649501 | GCA_015778165.1 | NonPC | Public | 2015 | USA | NA | 91.3 | 100 | 2.21 | 98.276 | 5249449 | 45.17 | 68 | 303687 |
| SAMN13649502-GCA_015778885.1 | MA414 | SAMN13649502 | GCA_015778885.1 | NonPC | Public | 2015 | USA | NA | 90.5 | 100 | 3.71 | 98.2793 | 5294929 | 45.19 | 59 | 451726 |
| SAMN13649503-GCA_015778045.1 | MA432 | SAMN13649503 | GCA_015778045.1 | NonPC | Public | 2015 | USA | NA | 91.2 | 100 | 1.37 | 98.2447 | 5239254 | 45.17 | 74 | 194403 |
| SAMN13649504-GCA_015778305.1 | MA441 | SAMN13649504 | GCA_015778305.1 | NonPC | Public | 2015 | USA | NA | 90.5 | 100 | 4.51 | 98.2781 | 5281360 | 45.19 | 62 | 451850 |
| SAMN13649505-GCA_015778285.1 | MA448 | SAMN13649505 | GCA_015778285.1 | NonPC | Public | 2015 | USA | NA | 91.3 | 100 | 0.5 | 98.2438 | 5221016 | 45.17 | 67 | 218563 |
| SAMN13649506-GCA_015778005.1 | MA459 | SAMN13649506 | GCA_015778005.1 | NonPC | Public | 2015 | USA | NA | 90.9 | 100 | 4.03 | 98.3372 | 5244616 | 45.17 | 64 | 276989 |
| SAMN13649507-GCA_015777945.1 | MA505 | SAMN13649507 | GCA_015777945.1 | NonPC | Public | 2015 | USA | NA | 91 | 100 | 2.18 | 98.3066 | 5262638 | 45.14 | 59 | 334666 |
| SAMN13649508-GCA_015778385.1 | MAVP-65 | SAMN13649508 | GCA_015778385.1 | NonPC | Public | 2014 | USA | Clinical | 90.9 | 100 | 2.58 | 98.3403 | 5239904 | 45.19 | 60 | 386036 |
| SAMN13649509-GCA_015778265.1 | MAVP-69 | SAMN13649509 | GCA_015778265.1 | NonPC | Public | 2014 | USA | Clinical | 89.3 | 100 | 1.98 | 98.0987 | 5092251 | 45.27 | 83 | 206892 |
| SAMN13649510-SRR10744610 | MAVP-I | SAMN13649510 | SRR10744610 | NonPC | Public | 2011 | USA | Clinical | 91.5 | 100 | 0.03 | 98.3196 | 5227917 | 45.18 | 96 | 260978 |
| SAMN13649511-GCA_015778205.1 | MAVP-K | SAMN13649511 | GCA_015778205.1 | NonPC | Public | 2011 | USA | Clinical | 91.2 | 100 | 0.07 | 98.3868 | 5117335 | 45.15 | 70 | 185371 |
| SAMN13649512-GCA_015778405.1 | MAVP-N | SAMN13649512 | GCA_015778405.1 | NonPC | Public | 2011 | USA | Clinical | 91 | 100 | 1.08 | 98.4372 | 5141067 | 45.2 | 49 | 375529 |
| SAMN13649513-GCA_015778485.1 | MEVP-1 | SAMN13649513 | GCA_015778485.1 | NonPC | Public | 2013 | USA | Clinical | 88.6 | 100 | 0.88 | 98.1717 | 5091807 | 45.16 | 85 | 138329 |
| SAMN13649514-GCA_015778465.1 | MEVP-2 | SAMN13649514 | GCA_015778465.1 | NonPC | Public | 2013 | USA | Clinical | 88.6 | 100 | 0.79 | 98.1463 | 5090262 | 45.16 | 86 | 154745 |
| SAMN13649515-GCA_015778505.1 | MEVP-3 | SAMN13649515 | GCA_015778505.1 | NonPC | Public | 2013 | NA | Clinical | 91.4 | 100 | 1.19 | 98.2705 | 5278870 | 45.17 | 74 | 189031 |
| SAMN13649516-GCA_015778445.1 | MEVP-5 | SAMN13649516 | GCA_015778445.1 | NonPC | Public | 2013 | USA | Clinical | 90.8 | 100 | 1.1 | 98.2954 | 5213105 | 45.25 | 56 | 210700 |
| SAMN13649517-GCA_015778525.1 | MEVP-6 | SAMN13649517 | GCA_015778525.1 | NonPC | Public | 2013 | USA | Clinical | 87.3 | 100 | 1.33 | 98.0523 | 5078946 | 45.23 | 74 | 190972 |
| SAMN13706796-SRR11823772 | 20150718005-1 | SAMN13706796 | SRR11823772 | NonPC | Public | 2015 | China | Env | 89.9 | 100 | 0.46 | 98.2144 | 5454718 | 45.26 | 96 | 244882 |
| SAMN13706797-SRR11823773 | 20150421008 | SAMN13706797 | SRR11823773 | NonPC | Public | 2015 | China | Env | 87.1 | 100 | 0.65 | 98.2874 | 5495950 | 45.27 | 79 | 476908 |
| SAMN13706798-SRR11823740 | 20150421004 | SAMN13706798 | SRR11823740 | NonPC | Public | 2015 | China | Env | 89.9 | 100 | 0.46 | 98.2486 | 5448812 | 45.26 | 107 | 232170 |
| SAMN13706799-SRR11823770 | 20150421031-3 | SAMN13706799 | SRR11823770 | NonPC | Public | 2015 | China | Env | 89.9 | 100 | 0.46 | 98.2246 | 5446345 | 45.27 | 105 | 244882 |
| SAMN13706800-SRR11823771 | 20150421021-3 | SAMN13706800 | SRR11823771 | NonPC | Public | 2015 | China | Env | 89.9 | 100 | 0.46 | 98.3023 | 5449334 | 45.26 | 109 | 203765 |
| SAMN13706801-SRR11823743 | 20140722003-4 | SAMN13706801 | SRR11823743 | NonPC | Public | 2014 | China | Env | 91.2 | 100 | 0.61 | 98.313 | 5351260 | 45.28 | 75 | 299750 |
| SAMN13706802-SRR11823744 | 20140722001-2 | SAMN13706802 | SRR11823744 | NonPC | Public | 2014 | China | Env | 91.3 | 100 | 0.61 | 98.3076 | 5364774 | 45.27 | 87 | 344670 |
| SAMN13706803-SRR11823778 | CZ17 | SAMN13706803 | SRR11823778 | NonPC | Public | 2014 | China | Clinical | 89.2 | 100 | 0.61 | 98.2667 | 5365481 | 45.26 | 81 | 289333 |
| SAMN13706804-SRR11823769 | 20140723001-1 | SAMN13706804 | SRR11823769 | NonPC | Public | 2014 | China | Env | 91.2 | 100 | 0.33 | 98.3109 | 5516956 | 45.17 | 102 | 181890 |
| SAMN13706806-SRR11823757 | 20150731001-2 | SAMN13706806 | SRR11823757 | NonPC | Public | 2015 | China | Env | 92 | 100 | 0.49 | 98.3181 | 5477445 | 45.22 | 75 | 308992 |
| SAMN13706807-SRR11823768 | 20140723013-2 | SAMN13706807 | SRR11823768 | NonPC | Public | 2014 | China | Env | 91.3 | 100 | 0.61 | 98.3046 | 5293312 | 45.29 | 80 | 395239 |
| SAMN13706808-SRR11823755 | 20150731007 | SAMN13706808 | SRR11823755 | NonPC | Public | 2015 | China | Env | 91.2 | 100 | 0.61 | 98.2819 | 5402812 | 45.3 | 79 | 219025 |
| SAMN13706809-SRR11823754 | 20150731021-1 | SAMN13706809 | SRR11823754 | NonPC | Public | 2015 | China | Env | 91.9 | 100 | 0.49 | 98.3135 | 5451466 | 45.23 | 58 | 212027 |
| SAMN13706810-SRR11823753 | 20150731024 | SAMN13706810 | SRR11823753 | NonPC | Public | 2015 | China | Env | 91.2 | 100 | 0.46 | 98.3129 | 5475892 | 45.15 | 124 | 187221 |
| SAMN13706811-SRR11823741 | 20140829001-1 | SAMN13706811 | SRR11823741 | NonPC | Public | 2014 | China | Env | 89.5 | 100 | 0.54 | 98.2703 | 5329073 | 45.34 | 57 | 429474 |

| | | | | | | | | | | | | | | | | |
|---|---|---|---|---|---|---|---|---|---|---|---|---|---|---|---|---|
| SAMN13706812-SRR11823746 | 20140624009-1 | SAMN13706812 | SRR11823746 | NonPC | Public | 2014 | China | Env | 90.4 | 100 | 0.67 | 98.2212 | 5415764 | 45.32 | 55 | 604198 |
| SAMN13706813-SRR11823803 | 20140124215 | SAMN13706813 | SRR11823803 | NonPC | Public | 2014 | China | Env | 90.4 | 100 | 0.7 | 98.2029 | 5419689 | 45.32 | 56 | 451366 |
| SAMN13706814-SRR11823742 | 20140829005-2 | SAMN13706814 | SRR11823742 | NonPC | Public | 2014 | China | Env | 89.6 | 100 | 0.54 | 98.2563 | 5331429 | 45.35 | 57 | 462526 |
| SAMN13706815-SRR11823751 | 20151113002-3 | SAMN13706815 | SRR11823751 | NonPC | Public | 2015 | China | Env | 89.3 | 100 | 0.05 | 98.2185 | 5463454 | 45.28 | 74 | 260222 |
| SAMN13706816-SRR11823817 | ZJ-2 | SAMN13706816 | SRR11823817 | NonPC | Public | 2016 | China | Env | 91.9 | 100 | 0 | 98.3696 | 5115346 | 45.3 | 33 | 450529 |
| SAMN13706817-SRR11823739 | ZJ-17 | SAMN13706817 | SRR11823739 | NonPC | Public | 2016 | China | Env | 89.1 | 100 | 0 | 98.3155 | 5025894 | 45.33 | 36 | 464833 |
| SAMN13706818-SRR11823738 | ZJ-22 | SAMN13706818 | SRR11823738 | NonPC | Public | 2016 | China | Env | 91.3 | 100 | 0.54 | 98.3244 | 5280242 | 45.28 | 51 | 497530 |
| SAMN13706819-SRR11823737 | ZJ-23 | SAMN13706819 | SRR11823737 | NonPC | Public | 2016 | China | Env | 90.8 | 100 | 0.4 | 98.2115 | 5317902 | 45.16 | 90 | 293571 |
| SAMN13706820-SRR11823818 | ZJ-29 | SAMN13706820 | SRR11823818 | NonPC | Public | 2016 | China | Env | 90.8 | 100 | 0.4 | 98.2513 | 5315847 | 45.15 | 85 | 465767 |
| SAMN13706821-SRR11823815 | ZJ-30 | SAMN13706821 | SRR11823815 | NonPC | Public | 2016 | China | Env | 91.4 | 100 | 0.22 | 98.2939 | 5262470 | 45.27 | 51 | 336409 |
| SAMN13706823-SRR11823749 | 20150708004-1 | SAMN13706823 | SRR11823749 | NonPC | Public | 2016 | China | Env | 89.4 | 100 | 0.74 | 98.3357 | 5459903 | 45.21 | 93 | 314508 |
| SAMN13706824-SRR11823752 | 20100612002 | SAMN13706824 | SRR11823752 | NonPC | Public | 2010 | China | Env | 89.4 | 100 | 0.32 | 98.3193 | 5299715 | 45.18 | 57 | 332390 |
| SAMN13706825-SRR11823761 | 20150710007-1 | SAMN13706825 | SRR11823761 | NonPC | Public | 2015 | China | Env | 91.2 | 100 | 0.46 | 98.3284 | 5478976 | 45.16 | 92 | 214026 |
| SAMN13706826-SRR11823759 | 20150710010-1 | SAMN13706826 | SRR11823759 | NonPC | Public | 2015 | China | Env | 91.2 | 100 | 0.74 | 98.32 | 5519603 | 45.2 | 96 | 203545 |
| SAMN13706827-SRR11823760 | 20150710009-1 | SAMN13706827 | SRR11823760 | NonPC | Public | 2015 | China | Env | 91.9 | 100 | 0.49 | 98.252 | 5478455 | 45.21 | 74 | 231027 |
| SAMN13706829-SRR11823762 | 20150710003-1 | SAMN13706829 | SRR11823762 | NonPC | Public | 2015 | China | Env | 91.3 | 100 | 0.46 | 98.3514 | 5509790 | 45.14 | 104 | 276279 |
| SAMN13706830-SRR11823766 | VP131 | SAMN13706830 | SRR11823766 | NonPC | Public | 2014 | China | Env | 91.2 | 100 | 0.61 | 98.2961 | 5364914 | 45.27 | 79 | 469154 |
| SAMN13706831-SRR11823748 | 20150708007-1 | SAMN13706831 | SRR11823748 | NonPC | Public | 2015 | China | Env | 91.2 | 100 | 0.74 | 98.3264 | 5518829 | 45.2 | 99 | 214026 |
| SAMN13706832-SRR11823747 | 20150718005-1 | SAMN13706832 | SRR11823747 | NonPC | Public | 2014 | China | Env | 90.9 | 100 | 0.54 | 98.3222 | 5297757 | 45.32 | 122 | 168057 |
| SAMN13706833-SRR11823765 | VP135 | SAMN13706833 | SRR11823765 | NonPC | Public | 2014 | China | Env | 91.2 | 100 | 0.33 | 98.3578 | 5212564 | 45.22 | 54 | 276279 |
| SAMN13706834-SRR11823764 | 20150709032-2 | SAMN13706834 | SRR11823764 | NonPC | Public | 2015 | China | Env | 91.2 | 100 | 0.46 | 98.3543 | 5480972 | 45.16 | 92 | 216914 |
| SAMN13706835-SRR11823763 | 20150709034-2 | SAMN13706835 | SRR11823763 | NonPC | Public | 2015 | China | Env | 91.2 | 100 | 0.46 | 98.3529 | 5480457 | 45.16 | 92 | 239997 |
| SAMN13706836-SRR11823758 | 20150710011-1 | SAMN13706836 | SRR11823758 | NonPC | Public | 2015 | China | Env | 90.9 | 100 | 0.76 | 98.2364 | 5488703 | 45.27 | 99 | 356454 |
| SAMN13706838-SRR11823786 | SH-2 | SAMN13706838 | SRR11823786 | NonPC | Public | 2012 | China | Env | 91.2 | 100 | 0.2 | 98.3167 | 5380970 | 45.31 | 96 | 329413 |
| SAMN13706839-SRR11823783 | SH-3 | SAMN13706839 | SRR11823783 | NonPC | Public | 2012 | China | Env | 91.8 | 100 | 0.26 | 98.3215 | 5444372 | 45 | 47 | 535598 |
| SAMN13706840-SRR11823781 | SH-4 | SAMN13706840 | SRR11823781 | NonPC | Public | 2013 | China | Env | 91.7 | 100 | 0.26 | 98.351 | 5432202 | 44.95 | 37 | 535598 |
| SAMN13706841-SRR11823780 | SH-5 | SAMN13706841 | SRR11823780 | NonPC | Public | 2013 | China | Env | 91.8 | 100 | 0.01 | 98.3176 | 5175751 | 45.27 | 47 | 641531 |
| SAMN13706842-SRR11823779 | SH-7 | SAMN13706842 | SRR11823779 | NonPC | Public | 2014 | China | Env | 91.3 | 100 | 0.33 | 98.3602 | 5134099 | 45.25 | 47 | 409209 |
| SAMN13706843-SRR11823777 | SH-8 | SAMN13706843 | SRR11823777 | NonPC | Public | 2014 | China | Env | 91.3 | 100 | 0.33 | 98.3632 | 5148196 | 45.28 | 63 | 729746 |
| SAMN13706844-SRR11823799 | SH-10 | SAMN13706844 | SRR11823799 | NonPC | Public | 2014 | China | Env | 91.9 | 100 | 0.27 | 98.3392 | 5382094 | 45.12 | 75 | 460478 |
| SAMN13706845-SRR11823798 | SH-11 | SAMN13706845 | SRR11823798 | NonPC | Public | 2014 | China | Env | 91.9 | 100 | 0.27 | 98.2726 | 5399100 | 45.12 | 87 | 460478 |
| SAMN13706846-SRR11823797 | SH-15 | SAMN13706846 | SRR11823797 | NonPC | Public | 2014 | China | Env | 91.9 | 100 | 0.27 | 98.2958 | 5377650 | 45.11 | 72 | 460478 |
| SAMN13706847-SRR11823796 | SH-17 | SAMN13706847 | SRR11823796 | NonPC | Public | 2014 | China | Env | 91.9 | 100 | 0.27 | 98.2965 | 5376878 | 45.12 | 73 | 421894 |
| SAMN13706848-SRR11823795 | SH-18 | SAMN13706848 | SRR11823795 | NonPC | Public | 2014 | China | Env | 91.8 | 100 | 0.54 | 98.2805 | 5379868 | 45.13 | 74 | 460478 |
| SAMN13706849-SRR11823794 | SH-19 | SAMN13706849 | SRR11823794 | NonPC | Public | 2014 | China | Env | 91.8 | 100 | 0.27 | 98.3323 | 5378363 | 45.12 | 74 | 421901 |
| SAMN13706850-SRR11823793 | SH-20 | SAMN13706850 | SRR11823793 | NonPC | Public | 2014 | China | Env | 91.8 | 100 | 0.23 | 98.3371 | 5194616 | 45.38 | 70 | 641531 |
| SAMN13706852-SRR11823791 | SH-25 | SAMN13706852 | SRR11823791 | NonPC | Public | 2014 | China | Env | 91.4 | 100 | 0.27 | 98.2888 | 5361161 | 45.12 | 74 | 460478 |
| SAMN13706853-SRR11823788 | SH-27 | SAMN13706853 | SRR11823788 | PC | Public | 2015 | China | Env | 98.5 | 100 | 0 | 99.9864 | 5092728 | 45.29 | 52 | 425661 |
| SAMN13706854-SRR11823787 | SH-28 | SAMN13706854 | SRR11823787 | NonPC | Public | 2015 | China | Env | 91.8 | 100 | 0 | 98.3321 | 5155367 | 45.26 | 28 | 641510 |
| SAMN13706855-SRR11823785 | SH-30 | SAMN13706855 | SRR11823785 | PC | Public | 2015 | China | Env | 98.3 | 100 | 0 | 99.9849 | 5083049 | 45.3 | 51 | 447155 |
| SAMN13706856-SRR11823784 | SH-31 | SAMN13706856 | SRR11823784 | NonPC | Public | 2014 | China | Env | 91.4 | 100 | 0.33 | 98.2864 | 5321119 | 45.21 | 56 | 403107 |
| SAMN13706857-SRR11823782 | SH48 | SAMN13706857 | SRR11823782 | NonPC | Public | 2016 | China | Env | 91.7 | 100 | 0 | 98.3053 | 5247065 | 45.18 | 27 | 755481 |
| SAMN13706858-SRR11823811 | Avp1 | SAMN13706858 | SRR11823811 | NonPC | Public | 2013 | China | Env | 91.4 | 100 | 0.22 | 98.3453 | 5285290 | 45.25 | 71 | 528197 |
| SAMN13706859-SRR11823800 | Avp2 | SAMN13706859 | SRR11823800 | NonPC | Public | 2013 | China | Env | 91.4 | 100 | 0.22 | 98.3422 | 5278900 | 45.25 | 72 | 528198 |
| SAMN13706860-SRR11823789 | Avp3 | SAMN13706860 | SRR11823789 | NonPC | Public | 2013 | China | Env | 91.4 | 100 | 0.22 | 98.3396 | 5281528 | 45.25 | 77 | 528198 |
| SAMN13706861-SRR11823812 | AP7 | SAMN13706861 | SRR11823812 | NonPC | Public | 2014 | China | Env | 91.2 | 100 | 0.54 | 98.347 | 5144430 | 45.39 | 64 | 490958 |
| SAMN13706862-SRR11823804 | HZ-8 | SAMN13706862 | SRR11823804 | NonPC | Public | 2014 | China | Env | 91.3 | 100 | 0.37 | 98.3618 | 5442112 | 45.31 | 49 | 207995 |
| SAMN13706865-SRR11823745 | HZ-15 | SAMN13706865 | SRR11823745 | NonPC | Public | 2015 | China | Env | 91 | 100 | 0.68 | 98.357 | 5403265 | 45.17 | 60 | 329259 |

| | | | | | | | | | | | | | | | | |
|---|---|---|---|---|---|---|---|---|---|---|---|---|---|---|---|---|
| SAMN13706866-SRR11823816 | HZ-18 | SAMN13706866 | SRR11823816 | NonPC | Public | 2015 | China | Env | 90.9 | 100 | 0 | 98.3334 | 5297475 | 45.35 | 59 | 400853 |
| SAMN13706867-SRR11823813 | HZ-49 | SAMN13706867 | SRR11823813 | NonPC | Public | 2016 | China | Env | 90 | 99.46 | 0.51 | 98.3765 | 5409802 | 45.33 | 64 | 205109 |
| SAMN13706868-SRR11823810 | HZ-50 | SAMN13706868 | SRR11823810 | NonPC | Public | 2016 | China | Env | 90.9 | 100 | 0 | 98.3528 | 5295868 | 45.35 | 65 | 395353 |
| SAMN13706870-SRR11823809 | HZ-51 | SAMN13706870 | SRR11823809 | NonPC | Public | 2016 | China | Env | 90.9 | 100 | 0 | 98.3166 | 5297010 | 45.35 | 71 | 359620 |
| SAMN13706871-SRR11823808 | HZ-54 | SAMN13706871 | SRR11823808 | NonPC | Public | 2016 | China | Env | 91.3 | 100 | 0.22 | 98.3043 | 5518294 | 45.38 | 91 | 250956 |
| SAMN13706872-SRR11823807 | HZ-64 | SAMN13706872 | SRR11823807 | NonPC | Public | 2016 | China | Env | 91.9 | 100 | 0.68 | 98.3528 | 5402543 | 45.17 | 65 | 389646 |
| SAMN13706873-SRR11823806 | HZ-66 | SAMN13706873 | SRR11823806 | NonPC | Public | 2016 | China | Env | 91.9 | 100 | 0.68 | 98.3726 | 5401455 | 45.17 | 59 | 389646 |
| SAMN13706874-SRR11823805 | HZ-79 | SAMN13706874 | SRR11823805 | NonPC | Public | 2016 | China | Env | 91.3 | 100 | 0.37 | 98.366 | 5442744 | 45.31 | 51 | 207762 |
| SAMN13706879-SRR11823802 | JS-9 | SAMN13706879 | SRR11823802 | NonPC | Public | 2014 | China | Env | 91.4 | 100 | 0.22 | 98.3487 | 5284994 | 45.26 | 76 | 528198 |
| SAMN13706880-SRR11823801 | NT-120 | SAMN13706880 | SRR11823801 | NonPC | Public | 2016 | China | Env | 89.6 | 100 | 0.2 | 98.2452 | 5515561 | 45.05 | 75 | 251791 |
| SAMN13706881-SRR11823767 | FJ-47 | SAMN13706881 | SRR11823767 | NonPC | Public | 2016 | China | Env | 90.1 | 100 | 0.19 | 98.2555 | 5246603 | 45.22 | 31 | 892505 |
| SAMN13706883-SRR11823814 | ZP8 | SAMN13706883 | SRR11823814 | NonPC | Public | 2011 | China | Env | 91.3 | 100 | 0.49 | 98.3148 | 5400622 | 45.29 | 76 | 184335 |
| SAMN13706884-SRR11823775 | VE07 | SAMN13706884 | SRR11823775 | NonPC | Public | 2008 | Viet_Nam | Env | 91.3 | 100 | 0.56 | 98.3293 | 5346548 | 45.29 | 66 | 311816 |
| SAMN13706885-SRR11823774 | VE10-3 | SAMN13706885 | SRR11823774 | NonPC | Public | 2010 | Viet_Nam | Env | 91.4 | 100 | 0.32 | 98.3372 | 5300070 | 45.18 | 59 | 342487 |
| SAMN13706886-SRR11823750 | 20151114011-4 | SAMN13706886 | SRR11823750 | NonPC | Public | 2015 | China | Env | 89.9 | 100 | 0.22 | 98.3153 | 5404607 | 45.22 | 75 | 263466 |
| SAMN13746682-GCA_015775955.1 | PNUSAV001089 | SAMN13746682 | GCA_015775955.1 | PC | Public | NA | USA | Clinical | 96.8 | 100 | 0.07 | 99.9807 | 5121760 | 45.53 | 282 | 35392 |
| SAMN13746683-GCA_015776075.1 | PNUSAV001090 | SAMN13746683 | GCA_015776075.1 | PC | Public | NA | USA | Clinical | 97.8 | 100 | 1.71 | 99.9851 | 5224997 | 45.35 | 43 | 461511 |
| SAMN13746684-GCA_015776055.1 | PNUSAV001072 | SAMN13746684 | GCA_015776055.1 | NonPC | Public | NA | USA | Clinical | 91.1 | 100 | 0.24 | 98.3889 | 5150145 | 45.29 | 103 | 115746 |
| SAMN13746685-GCA_015775995.1 | PNUSAV001071 | SAMN13746685 | GCA_015775995.1 | NonPC | Public | NA | USA | Clinical | 90.2 | 100 | 1.29 | 98.2144 | 5237924 | 45.24 | 142 | 93555 |
| SAMN13746687-GCA_015775915.1 | PNUSAV001063 | SAMN13746687 | GCA_015775915.1 | NonPC | Public | NA | USA | Clinical | 91.2 | 100 | 0.54 | 98.3165 | 5105253 | 45.34 | 148 | 71832 |
| SAMN13746688-GCA_015776095.1 | PNUSAV001055 | SAMN13746688 | GCA_015776095.1 | NonPC | Public | NA | USA | Clinical | 91.4 | 100 | 3.18 | 98.3639 | 5289476 | 45.15 | 58 | 186681 |
| SAMN13746689-GCA_015776035.1 | PNUSAV001057 | SAMN13746689 | GCA_015776035.1 | NonPC | Public | NA | USA | Clinical | 91.5 | 100 | 2.53 | 98.3498 | 5144474 | 45.29 | 55 | 217520 |
| SAMN13746690-GCA_015775935.1 | PNUSAV000878 | SAMN13746690 | GCA_015775935.1 | NonPC | Public | NA | USA | Clinical | 91 | 100 | 3.81 | 98.3812 | 5168432 | 45.38 | 89 | 102183 |
| SAMN13746694-GCA_015776015.1 | PNUSAV001115 | SAMN13746694 | GCA_015776015.1 | NonPC | Public | NA | USA | Clinical | 91.4 | 100 | 0 | 98.3357 | 5133920 | 45.24 | 39 | 808514 |
| SAMN13746697-GCA_016231605.1 | PNUSAV001134 | SAMN13746697 | GCA_016231605.1 | NonPC | Public | NA | USA | Clinical | 90.6 | 100 | 4.23 | 98.2833 | 5137928 | 45.33 | 121 | 99949 |
| SAMN13746698-GCA_015775795.1 | PNUSAV001109 | SAMN13746698 | GCA_015775795.1 | NonPC | Public | NA | USA | Clinical | 91.4 | 100 | 3.36 | 98.3033 | 5229168 | 45.29 | 70 | 163566 |
| SAMN13746699-GCA_015775895.1 | PNUSAV001114 | SAMN13746699 | GCA_015775895.1 | NonPC | Public | NA | USA | Clinical | 91.4 | 100 | 3.13 | 98.3129 | 5204913 | 45.25 | 38 | 719058 |
| SAMN13747307-GCA_016231505.1 | PNUSAV001096 | SAMN13747307 | GCA_016231505.1 | NonPC | Public | NA | USA | Clinical | 90.1 | 100 | 0.22 | 98.3156 | 5095402 | 45.34 | 238 | 45784 |
| SAMN13763540-GCA_015775715.1 | PNUSAV001137 | SAMN13763540 | GCA_015775715.1 | NonPC | Public | NA | USA | Clinical | 91.1 | 100 | 1.19 | 98.2918 | 5040198 | 45.3 | 66 | 209062 |
| SAMN13763541-GCA_015775595.1 | PNUSAV000758 | SAMN13763541 | GCA_015775595.1 | NonPC | Public | NA | USA | Clinical | 91.2 | 100 | 1.36 | 98.3023 | 5149190 | 45.3 | 121 | 94038 |
| SAMN13763542-GCA_015775675.1 | PNUSAV000614 | SAMN13763542 | GCA_015775675.1 | NonPC | Public | NA | USA | Clinical | 90.8 | 100 | 1.96 | 98.2865 | 5263878 | 45.2 | 116 | 99861 |
| SAMN13763572-GCA_015775535.1 | PNUSAV000667 | SAMN13763572 | GCA_015775535.1 | NonPC | Public | NA | USA | Clinical | 91.4 | 100 | 0.42 | 98.3043 | 5147566 | 45.25 | 44 | 564948 |
| SAMN13763575-GCA_015775575.1 | PNUSAV000658 | SAMN13763575 | GCA_015775575.1 | NonPC | Public | NA | USA | Clinical | 88.9 | 100 | 1.63 | 98.0922 | 5085853 | 45.36 | 133 | 89992 |
| SAMN13763586-GCA_015775495.1 | PNUSAV001058 | SAMN13763586 | GCA_015775495.1 | NonPC | Public | NA | USA | Clinical | 91.4 | 100 | 3.54 | 98.3245 | 5322347 | 45.26 | 41 | 564960 |
| SAMN13793688-SRR11823776 | TJ150 | SAMN13793688 | SRR11823776 | NonPC | Public | 2016 | China | Env | 88.3 | 100 | 0.34 | 98.1869 | 5489180 | 45.18 | 57 | 533828 |
| SAMN13808681-GCA_015775475.1 | CFSAN095190 | SAMN13808681 | GCA_015775475.1 | NonPC | Public | 2013 | USA | Env | 90.1 | 100 | 4.49 | 98.3079 | 5377677 | 45.19 | 33 | 501862 |
| SAMN13808683-GCA_015775635.1 | CFSAN095280 | SAMN13808683 | GCA_015775635.1 | NonPC | Public | 2013 | USA | Env | 91.9 | 100 | 0.45 | 98.403 | 5090022 | 45.31 | 153 | 75502 |
| SAMN13808684-GCA_015775455.1 | CFSAN095296 | SAMN13808684 | GCA_015775455.1 | PrePC | Public | 2013 | USA | Env | 94.9 | 100 | 4.99 | 99.8213 | 5389080 | 45.32 | 74 | 161304 |
| SAMN13808686-GCA_015775395.1 | CFSAN095283 | SAMN13808686 | GCA_015775395.1 | NonPC | Public | 2013 | USA | Env | 89.9 | 100 | 0.35 | 98.2611 | 5048260 | 45.48 | 153 | 68718 |
| SAMN13808687-GCA_015775375.1 | CFSAN095168 | SAMN13808687 | GCA_015775375.1 | NonPC | Public | 2013 | USA | Env | 91 | 100 | 3.67 | 98.2776 | 5483672 | 45.26 | 44 | 368503 |
| SAMN13808689-GCA_015775415.1 | CFSAN095294 | SAMN13808689 | GCA_015775415.1 | NonPC | Public | 2013 | USA | Env | 90.8 | 100 | 0.28 | 98.2815 | 5158412 | 45.23 | 75 | 151111 |
| SAMN13808711-SRR10857358 | CFSAN095284 | SAMN13808711 | SRR10857358 | NonPC | Public | 2013 | USA | Env | 90.9 | 100 | 0.14 | 98.2527 | 5144158 | 45.25 | 91 | 131078 |
| SAMN13808808-GCA_015775335.1 | CFSAN095275 | SAMN13808808 | GCA_015775335.1 | NonPC | Public | 2013 | USA | Env | 91.3 | 100 | 1.41 | 98.3392 | 5180073 | 45.32 | 120 | 80817 |
| SAMN13808827-GCA_015775235.1 | CFSAN095289 | SAMN13808827 | GCA_015775235.1 | NonPC | Public | 2013 | USA | Env | 90.5 | 100 | 0.11 | 98.2717 | 5152318 | 45.29 | 138 | 83692 |
| SAMN13808830-GCA_015775305.1 | CFSAN095291 | SAMN13808830 | GCA_015775305.1 | NonPC | Public | 2013 | USA | Env | 90.8 | 100 | 3.26 | 98.2667 | 5254819 | 45.24 | 70 | 170766 |
| SAMN13808833-GCA_015775285.1 | CFSAN095276 | SAMN13808833 | GCA_015775285.1 | NonPC | Public | 2013 | USA | Env | 91.4 | 100 | 4.17 | 98.3485 | 5274589 | 45.31 | 103 | 104257 |
| SAMN13808835-GCA_015775255.1 | CFSAN095293 | SAMN13808835 | GCA_015775255.1 | NonPC | Public | 2013 | USA | Env | 90.9 | 100 | 1.02 | 98.28 | 5180640 | 45.22 | 49 | 192409 |
| SAMN13808836-SRR10857468 | CFSAN095278 | SAMN13808836 | SRR10857468 | NonPC | Public | 2013 | USA | Env | 91.4 | 100 | 0.27 | 98.278 | 5008765 | 45.41 | 70 | 167698 |

| | | | | | | | | | | | | | | | | |
|---|---|---|---|---|---|---|---|---|---|---|---|---|---|---|---|---|
| SAMN13808837-SRR10857470 | CFSAN095279 | SAMN13808837 | SRR10857470 | NonPC | Public | 2013 | USA | Env | 91.3 | 100 | 0.27 | 98.2793 | 5007786 | 45.42 | 80 | 133851 |
| SAMN13809857-GCA_015775165.1 | CFSAN095292 | SAMN13809857 | GCA_015775165.1 | NonPC | Public | 2013 | USA | Env | 90.8 | 100 | 2.79 | 98.276 | 5248561 | 45.25 | 73 | 128073 |
| SAMN13809860-GCA_015775215.1 | CFSAN095281 | SAMN13809860 | GCA_015775215.1 | NonPC | Public | 2013 | USA | Env | 90 | 100 | 0.55 | 98.2784 | 5143728 | 45.31 | 124 | 100269 |
| SAMN13809951-GCA_015775075.1 | CFSAN095287 | SAMN13809951 | GCA_015775075.1 | NonPC | Public | 2013 | USA | Env | 90.9 | 100 | 1.7 | 98.2458 | 5176006 | 45.31 | 88 | 98819 |
| SAMN13809952-GCA_015775015.1 | CFSAN095236 | SAMN13809952 | GCA_015775015.1 | NonPC | Public | 2013 | USA | Env | 89.2 | 100 | 0.67 | 98.2782 | 5117402 | 45.39 | 250 | 45749 |
| SAMN13809954-GCA_015775095.1 | CFSAN095286 | SAMN13809954 | GCA_015775095.1 | NonPC | Public | 2013 | USA | Env | 90.2 | 100 | 0.87 | 98.2821 | 5080158 | 45.43 | 91 | 130714 |
| SAMN13809955-GCA_015775115.1 | CFSAN095274 | SAMN13809955 | GCA_015775115.1 | NonPC | Public | 2013 | USA | Env | 88.7 | 100 | 0.58 | 98.1214 | 5034405 | 45.33 | 144 | 80342 |
| SAMN13825963-GCA_015774995.1 | PNUSAV000661 | SAMN13825963 | GCA_015774995.1 | NonPC | Public | NA | USA | Clinical | 91.4 | 100 | 1.27 | 98.2776 | 5180670 | 45.26 | 70 | 179194 |
| SAMN13825965-GCA_015774975.1 | PNUSAV000659 | SAMN13825965 | GCA_015774975.1 | NonPC | Public | NA | USA | Clinical | 89.2 | 100 | 1.73 | 98.1055 | 5136759 | 45.29 | 95 | 123287 |
| SAMN13825967-GCA_015960875.1 | PNUSAV000619 | SAMN13825967 | GCA_015960875.1 | NonPC | Public | NA | USA | Clinical | 90.5 | 100 | 2.59 | 98.2547 | 5202044 | 45.22 | 73 | 148789 |
| SAMN13869827-GCA_015774935.1 | PNUSAV001153 | SAMN13869827 | GCA_015774935.1 | NonPC | Public | NA | USA | Clinical | 91.3 | 100 | 0.27 | 98.3345 | 5184321 | 45.19 | 50 | 354345 |
| SAMN13878427-GCA_015960765.1 | PNUSAV001094 | SAMN13878427 | GCA_015960765.1 | NonPC | Public | NA | USA | Clinical | 89.8 | 100 | 3.58 | 98.258 | 5329468 | 45.21 | 55 | 206446 |
| SAMN13893117-GCF_010692765.1 | CAIM_1400 | SAMN13893117 | GCF_010692765.1 | PC | Public | 2004 | Mexico | Clinical | 98.5 | 100 | 0.76 | 99.9818 | 5290587 | 45.11 | 83 | 204016 |
| SAMN13893118-GCF_009936575.1 | CICESE-170 | SAMN13893118 | GCF_009936575.1 | PC | Public | 1998 | Mexico | Clinical | 98.2 | 100 | 0 | 99.9797 | 5105390 | 45.25 | 88 | 242398 |
| SAMN13893119-GCF_009936605.1 | CICESE-186 | SAMN13893119 | GCF_009936605.1 | PC | Public | 1999 | Mexico | Clinical | 97.8 | 100 | 0.12 | 99.9805 | 5073833 | 45.43 | 380 | 25898 |
| SAMN13893120-GCF_010692775.1 | CICESE-187 | SAMN13893120 | GCF_010692775.1 | PC | Public | 2000 | Mexico | Clinical | 96.6 | 99.95 | 0.06 | 99.9691 | 4988243 | 45.57 | 584 | 16672 |
| SAMN13893121-GCF_010671575.1 | CICESE-188 | SAMN13893121 | GCF_010671575.1 | PC | Public | 2009 | Mexico | Clinical | 91.8 | 99.06 | 0.03 | 99.985 | 4672642 | 46.06 | 1287 | 6064 |
| SAMN13893122-GCF_009936625.1 | CICESE-273 | SAMN13893122 | GCF_009936625.1 | PC | Public | 2012 | Mexico | Env | 96.1 | 100 | 0 | 99.9684 | 5013960 | 45.63 | 838 | 10353 |
| SAMN13901173-GCA_015960705.1 | PNUSAV001155 | SAMN13901173 | GCA_015960705.1 | NonPC | Public | NA | USA | Clinical | 91.3 | 100 | 2.53 | 98.3081 | 5258932 | 45.25 | 43 | 556731 |
| SAMN13909686-GCA_015960685.1 | 19MO12SH02-V | SAMN13909686 | GCA_015960685.1 | NonPC | Public | 2019 | USA | Env | 88.7 | 100 | 3.45 | 98.0901 | 5193060 | 45.3 | 59 | 261724 |
| SAMN13909687-GCA_015960725.1 | 19MO12SH01-V1 | SAMN13909687 | GCA_015960725.1 | NonPC | Public | 2019 | USA | Env | 90.1 | 100 | 3.64 | 98.2891 | 5412224 | 45.18 | 72 | 400399 |
| SAMN14064563-GCA_015960565.1 | CFSAN095300 | SAMN14064563 | GCA_015960565.1 | NonPC | Public | 2013 | USA | Env | 91.5 | 100 | 3.49 | 98.3643 | 5399376 | 45.4 | 97 | 122815 |
| SAMN14064736-GCA_015960785.1 | CFSAN095282 | SAMN14064736 | GCA_015960785.1 | NonPC | Public | 2013 | USA | Env | 90 | 100 | 0.77 | 98.2756 | 5156475 | 45.33 | 170 | 72398 |
| SAMN14066391-SRR11041488 | CFSAN095285 | SAMN14066391 | SRR11041488 | NonPC | Public | 2013 | USA | Env | 91.1 | 100 | 0.35 | 98.2442 | 5089713 | 45.3 | 92 | 108799 |
| SAMN14069582-GCA_015960515.1 | CFSAN095170 | SAMN14069582 | GCA_015960515.1 | NonPC | Public | 2013 | USA | Env | 91 | 100 | 4.16 | 98.2847 | 5584066 | 45.29 | 43 | 367566 |
| SAMN14080886-GCA_014836315.1 | M0802 | SAMN14080886 | GCA_014836315.1 | NonPC | Public | 2013 | Mexico | Env | 91.2 | 97.67 | 1.43 | 98.1406 | 5532117 | 45.25 | 110 | 108659 |
| SAMN14081008-GCA_014836295.1 | M0803 | SAMN14081008 | GCA_014836295.1 | NonPC | Public | 2013 | Mexico | Env | 91.2 | 98.89 | 1.34 | 98.1367 | 5561299 | 45.26 | 265 | 45226 |
| SAMN14081013-GCF_014836305.1 | M0904 | SAMN14081013 | GCF_014836305.1 | NonPC | Public | 2013 | Mexico | Env | 91.3 | 100 | 0.68 | 98.1669 | 5582418 | 45.31 | 90 | 224838 |
| SAMN14083307-GCA_015960405.1 | 19MO11SH01-V | SAMN14083307 | GCA_015960405.1 | NonPC | Public | 2019 | USA | Env | 88.8 | 100 | 3.57 | 98.0588 | 5410172 | 45.32 | 58 | 227592 |
| SAMN14121487-GCA_014833025.1 | M0905 | SAMN14121487 | GCA_014833025.1 | NonPC | Public | 2013 | Mexico | Env | 90.9 | 97.21 | 0.84 | 98.1438 | 5356302 | 45.26 | 202 | 46892 |
| SAMN14121488-GCF_014833015.1 | M2401 | SAMN14121488 | GCF_014833015.1 | NonPC | Public | 2019 | Mexico | Env | 90.7 | 100 | 0.07 | 98.2665 | 5338883 | 45.34 | 206 | 67938 |
| SAMN14121636-GCF_014833075.1 | M2411 | SAMN14121636 | GCF_014833075.1 | NonPC | Public | 2019 | Mexico | Env | 90.7 | 100 | 0.07 | 98.2705 | 5371069 | 45.34 | 189 | 78046 |
| SAMN14121637-GCF_014833085.1 | M2413 | SAMN14121637 | GCF_014833085.1 | NonPC | Public | 2019 | Mexico | Env | 91 | 100 | 0.34 | 98.382 | 5412491 | 45.3 | 293 | 45086 |
| SAMN14122045-GCF_014833125.1 | M2415 | SAMN14122045 | GCF_014833125.1 | NonPC | Public | 2019 | Mexico | Env | 91 | 100 | 0.43 | 98.3528 | 5402258 | 45.29 | 370 | 30846 |
| SAMN14147082-GCA_015960185.1 | PNUSAV001173 | SAMN14147082 | GCA_015960185.1 | NonPC | Public | 2020 | USA | Env | 91.1 | 100 | 0.82 | 98.2282 | 5189898 | 45.32 | 135 | 92392 |
| SAMN14147083-GCA_015960085.1 | PNUSAV001174 | SAMN14147083 | GCA_015960085.1 | NonPC | Public | 2020 | USA | Env | 91 | 100 | 3.27 | 98.2349 | 5340546 | 45.28 | 82 | 170123 |
| SAMN14147084-GCA_015960225.1 | PNUSAV001171 | SAMN14147084 | GCA_015960225.1 | NonPC | Public | 2020 | USA | Env | 91 | 100 | 1.99 | 98.2769 | 5259776 | 45.2 | 92 | 176493 |
| SAMN14147085-GCA_015960045.1 | PNUSAV001175 | SAMN14147085 | GCA_015960045.1 | NonPC | Public | 2020 | USA | Env | 90.7 | 100 | 3.22 | 98.333 | 5159957 | 45.38 | 54 | 445885 |
| SAMN14147086-GCA_015960245.1 | PNUSAV001160 | SAMN14147086 | GCA_015960245.1 | NonPC | Public | 2020 | USA | Env | 89.7 | 100 | 0.51 | 98.0802 | 5136773 | 45.35 | 169 | 64204 |
| SAMN14147088-GCA_015960065.1 | PNUSAV001168 | SAMN14147088 | GCA_015960065.1 | NonPC | Public | 2020 | USA | Env | 89.8 | 100 | 3.14 | 98.3011 | 5265594 | 45.31 | 37 | 449181 |
| SAMN14147090-GCA_015960205.1 | PNUSAV001166 | SAMN14147090 | GCA_015960205.1 | NonPC | Public | 2020 | USA | Env | 89.8 | 100 | 1.63 | 98.3178 | 5147245 | 45.26 | 98 | 116084 |
| SAMN14147700-GCA_015960265.1 | PNUSAV001161 | SAMN14147700 | GCA_015960265.1 | NonPC | Public | 2020 | USA | Env | 91 | 100 | 1.54 | 98.3719 | 5173038 | 45.36 | 122 | 93726 |
| SAMN14214392-GCF_014833115.1 | M0607 | SAMN14214392 | GCF_014833115.1 | NonPC | Public | 2013 | Mexico | Env | 90.6 | 100 | 0.07 | 98.1737 | 5291564 | 45.35 | 135 | 116511 |
| SAMN14266004-GCA_018086005.1 | PNUSAV001181 | SAMN14266004 | GCA_018086005.1 | NonPC | Public | NA | USA | Clinical | 91.5 | 100 | 1.1 | 98.3227 | 5098847 | 45.38 | 175 | 63654 |
| SAMN14266007-GCA_015774455.1 | PNUSAV001182 | SAMN14266007 | GCA_015774455.1 | PC | Public | NA | USA | Clinical | 97.1 | 100 | 3.54 | 99.9447 | 5302184 | 45.42 | 166 | 63818 |
| SAMN14306763-GCA_015843795.1 | 2014V-1063_FLEX_500_Vibrio | SAMN14306763 | GCA_015843795.1 | NonPC | Public | NA | USA | Clinical | 88.8 | 100 | 0.89 | 98.0629 | 5114276 | 45.35 | 157 | 71860 |
| SAMN14306766-SRR17442126 | 2013V-1178_FLEX_500_Vibrio | SAMN14306766 | SRR17442126 | NonPC | Public | NA | USA | NA | 89.8 | 100 | 0.27 | 98.314 | 5036341 | 45.36 | 38 | 642245 |

| | | | | | | | | | | | | | | | | | |
|---|---|---|---|---|---|---|---|---|---|---|---|---|---|---|---|---|---|
| SAMN14306773-GCA_015844055.1 | 2013V-1125_FLEX_500_Vibrio | SAMN14306773 | GCA_015844055.1 | NonPC | Public | | NA | USA | Clinical | 91.2 | 100 | 1.16 | 98.2943 | 5177720 | 45.37 | 151 | 68503 |
| SAMN14352316-GCA_015774435.1 | PNUSAV001165 | SAMN14352316 | GCA_015774435.1 | NonPC | Public | | 2020 | USA | Env | 89.3 | 100 | 1.29 | 98.3286 | 5072862 | 45.45 | 119 | 82691 |
| SAMN14352317-GCA_015774415.1 | PNUSAV001169 | SAMN14352317 | GCA_015774415.1 | NonPC | Public | | 2020 | USA | Env | 90.8 | 100 | 0.08 | 98.3192 | 5215600 | 45.26 | 146 | 79259 |
| SAMN14396416-GCA_018089465.1 | PNUSAV001185 | SAMN14396416 | GCA_018089465.1 | PC | Public | | NA | USA | Clinical | 97.9 | 100 | 1.59 | 99.9882 | 5182812 | 45.28 | 64 | 288811 |
| SAMN14404426-SRR11350400 | PNUSAV001162_1 | SAMN14404426 | SRR11350400 | NonPC | Public | | 2020 | USA | Env | 91.9 | 100 | 0 | 98.313 | 5143785 | 45.33 | 56 | 214480 |
| SAMN14411226-GCF_014922295.1 | L1 | SAMN14411226 | GCF_014922295.1 | PC | Public | | 2015 | China | Clinical | 98.6 | 100 | 0 | 99.9904 | 5103150 | 45.3 | 44 | 428235 |
| SAMN14411227-GCF_014922265.1 | L2 | SAMN14411227 | GCF_014922265.1 | PC | Public | | 2015 | China | Clinical | 98.5 | 100 | 0 | 99.9806 | 5102539 | 45.3 | 48 | 428121 |
| SAMN14411228-GCF_014922235.1 | L3 | SAMN14411228 | GCF_014922235.1 | PC | Public | | 2015 | China | Clinical | 98.6 | 100 | 0 | 99.989 | 5101026 | 45.29 | 44 | 428130 |
| SAMN14411230-GCF_014922245.1 | L6 | SAMN14411230 | GCF_014922245.1 | PC | Public | | 2015 | China | Clinical | 98.6 | 100 | 0 | 99.9895 | 5105020 | 45.3 | 45 | 445755 |
| SAMN14411231-GCF_014922205.1 | L7 | SAMN14411231 | GCF_014922205.1 | PC | Public | | 2015 | China | Clinical | 98.6 | 100 | 0 | 99.9827 | 5102313 | 45.29 | 44 | 428235 |
| SAMN14411232-GCF_014922185.1 | L8 | SAMN14411232 | GCF_014922185.1 | PC | Public | | 2015 | China | Clinical | 98.6 | 100 | 0 | 99.9896 | 5101613 | 45.29 | 46 | 441946 |
| SAMN14411233-GCF_014922155.1 | r16 | SAMN14411233 | GCF_014922155.1 | NonPC | Public | | 2018 | China | Clinical | 91.3 | 99.86 | 0 | 98.2155 | 5261654 | 45.16 | 35 | 546683 |
| SAMN14411234-GCF_014922125.1 | r46 | SAMN14411234 | GCF_014922125.1 | PC | Public | | 2018 | China | Clinical | 98.5 | 100 | 0.14 | 99.9833 | 5099286 | 45.3 | 51 | 507129 |
| SAMN14411235-GCF_014922075.1 | r75 | SAMN14411235 | GCF_014922075.1 | PC | Public | | 2017 | China | Clinical | 98.4 | 99.84 | 0.14 | 99.9844 | 5097582 | 45.29 | 44 | 826678 |
| SAMN14411236-GCF_014922115.1 | r79-2 | SAMN14411236 | GCF_014922115.1 | PC | Public | | 2017 | China | Clinical | 98.4 | 100 | 0.02 | 99.9859 | 5094493 | 45.29 | 40 | 826802 |
| SAMN14411237-GCF_014922095.1 | V10 | SAMN14411237 | GCF_014922095.1 | NonPC | Public | | 2015 | China | Env | 91.5 | 99.86 | 0.82 | 98.2747 | 5366656 | 45.34 | 77 | 534012 |
| SAMN14411238-GCA_014964205.1 | V101 | SAMN14411238 | GCA_014964205.1 | NonPC | Public | | 2015 | China | Env | 90 | 99.86 | 0.06 | 98.3094 | 5114750 | 45.42 | 53 | 530507 |
| SAMN14411239-GCF_014922055.1 | V11 | SAMN14411239 | GCF_014922055.1 | NonPC | Public | | 2015 | China | Env | 90.7 | 99.93 | 0.15 | 98.2354 | 5067345 | 45.24 | 61 | 650917 |
| SAMN14411240-GCF_014921975.1 | V110 | SAMN14411240 | GCF_014921975.1 | NonPC | Public | | 2016 | China | Env | 91.5 | 99.91 | 0.03 | 98.3719 | 5062156 | 45.34 | 46 | 805312 |
| SAMN14411241-GCF_014921965.1 | V111 | SAMN14411241 | GCF_014921965.1 | NonPC | Public | | 2016 | China | Env | 91.4 | 100 | 0.34 | 98.3257 | 5118642 | 45.31 | 46 | 534949 |
| SAMN14411242-GCF_014921995.1 | V112 | SAMN14411242 | GCF_014921995.1 | NonPC | Public | | 2016 | China | Env | 91.4 | 99.91 | 0.48 | 98.306 | 5195127 | 45.31 | 40 | 534396 |
| SAMN14411243-GCF_014921955.1 | V113 | SAMN14411243 | GCF_014921955.1 | NonPC | Public | | 2016 | China | Env | 91 | 100 | 0.05 | 98.3516 | 5007940 | 45.4 | 45 | 880483 |
| SAMN14411244-GCF_014922025.1 | V114 | SAMN14411244 | GCF_014922025.1 | NonPC | Public | | 2016 | China | Env | 91.4 | 99.93 | 0.38 | 98.3504 | 5118293 | 45.31 | 44 | 943134 |
| SAMN14411245-GCF_014921915.1 | V116 | SAMN14411245 | GCF_014921915.1 | NonPC | Public | | 2016 | China | Env | 91.4 | 99.66 | 0 | 98.3278 | 5156623 | 45.3 | 50 | 533823 |
| SAMN14411246-SRR11357184 | V118 | SAMN14411246 | SRR11357184 | NonPC | Public | | 2016 | China | Env | 91.4 | 99.8 | 0.3 | 98.3732 | 5182025 | 45.22 | 89 | 469992 |
| SAMN14411247-SRR11357183 | V12 | SAMN14411247 | SRR11357183 | NonPC | Public | | 2015 | China | Env | 90.7 | 99.51 | 0.33 | 98.2898 | 5094418 | 45.41 | 48 | 373100 |
| SAMN14411248-GCF_014921855.1 | V120 | SAMN14411248 | GCF_014921855.1 | NonPC | Public | | 2016 | China | Env | 88.4 | 99.93 | 0.43 | 98.1906 | 5269885 | 45.21 | 51 | 763089 |
| SAMN14411249-SRR11357181 | V122 | SAMN14411249 | SRR11357181 | NonPC | Public | | 2016 | China | Env | 91.5 | 99.32 | 0.03 | 98.3636 | 5123029 | 45.24 | 72 | 401888 |
| SAMN14411250-GCF_014921795.1 | V126 | SAMN14411250 | GCF_014921795.1 | NonPC | Public | | 2016 | China | Env | 91.7 | 99.93 | 0.09 | 98.3689 | 5050393 | 45.29 | 63 | 449687 |
| SAMN14411251-GCF_014921775.1 | V127 | SAMN14411251 | GCF_014921775.1 | NonPC | Public | | 2016 | China | Env | 91.1 | 99.93 | 0.32 | 98.3038 | 5149492 | 45.27 | 49 | 530815 |
| SAMN14411252-GCF_014921825.1 | V128 | SAMN14411252 | GCF_014921825.1 | NonPC | Public | | 2016 | China | Env | 88.4 | 99.93 | 0.48 | 98.2748 | 5113075 | 45.27 | 53 | 1024383 |
| SAMN14411253-GCF_014921785.1 | V13 | SAMN14411253 | GCF_014921785.1 | NonPC | Public | | 2015 | China | Env | 91.2 | 99.19 | 0.17 | 98.3698 | 5113645 | 45.31 | 37 | 453692 |
| SAMN14411254-SRR11357176 | V130 | SAMN14411254 | SRR11357176 | NonPC | Public | | 2016 | China | Env | 87.7 | 98.98 | 0.41 | 98.1907 | 5076924 | 45.22 | 67 | 379940 |
| SAMN14411255-GCF_014921725.1 | V131 | SAMN14411255 | GCF_014921725.1 | NonPC | Public | | 2016 | China | Env | 88.3 | 99.91 | 0.83 | 98.2365 | 4986855 | 45.34 | 43 | 510240 |
| SAMN14411256-GCF_014921705.1 | V132 | SAMN14411256 | GCF_014921705.1 | NonPC | Public | | 2016 | China | Env | 88.3 | 99.61 | 0.71 | 98.2233 | 5245124 | 45.11 | 98 | 459659 |
| SAMN14411257-GCF_014921695.1 | V135 | SAMN14411257 | GCF_014921695.1 | NonPC | Public | | 2016 | China | Env | 88.4 | 99.84 | 0.39 | 98.2667 | 5039132 | 45.34 | 74 | 779246 |
| SAMN14411258-GCF_014921665.1 | V136 | SAMN14411258 | GCF_014921665.1 | NonPC | Public | | 2016 | China | Env | 88.2 | 99.93 | 0.42 | 98.207 | 5108868 | 45.28 | 71 | 409302 |
| SAMN14411259-GCF_014921615.1 | V14 | SAMN14411259 | GCF_014921615.1 | NonPC | Public | | 2015 | China | Env | 87.9 | 100 | 0.3 | 98.2555 | 5157407 | 45.19 | 75 | 570548 |
| SAMN14411260-GCF_014921655.1 | V140 | SAMN14411260 | GCF_014921655.1 | NonPC | Public | | 2016 | China | Env | 91.7 | 99.93 | 0.33 | 98.3462 | 5164225 | 45.26 | 57 | 432225 |
| SAMN14411261-GCF_014921625.1 | V150 | SAMN14411261 | GCF_014921625.1 | NonPC | Public | | 2016 | China | Env | 87.6 | 99.95 | 0.4 | 98.2055 | 5093657 | 45.29 | 45 | 702089 |
| SAMN14411262-GCF_014921595.1 | V154 | SAMN14411262 | GCF_014921595.1 | NonPC | Public | | 2016 | China | Env | 91.3 | 99.93 | 0.1 | 98.3759 | 5011076 | 45.35 | 41 | 522503 |
| SAMN14411263-GCF_014921555.1 | V164 | SAMN14411263 | GCF_014921555.1 | NonPC | Public | | 2016 | China | Env | 88.3 | 99.88 | 0.5 | 98.2369 | 5001010 | 45.28 | 48 | 321061 |
| SAMN14411264-GCF_014921545.1 | V165 | SAMN14411264 | GCF_014921545.1 | NonPC | Public | | 2016 | China | Env | 88.1 | 99.77 | 0.66 | 98.2419 | 5067328 | 45.19 | 80 | 452360 |
| SAMN14411265-GCF_014921525.1 | V166 | SAMN14411265 | GCF_014921525.1 | NonPC | Public | | 2016 | China | Env | 89.9 | 99.82 | 0.27 | 98.2753 | 5125964 | 45.29 | 51 | 1159229 |
| SAMN14411266-SRR11357123 | V168 | SAMN14411266 | SRR11357123 | NonPC | Public | | 2016 | China | Env | 88.3 | 99.88 | 0.4 | 98.2839 | 5041000 | 45.3 | 47 | 387525 |
| SAMN14411267-GCF_014921475.1 | V169 | SAMN14411267 | GCF_014921475.1 | NonPC | Public | | 2016 | China | Env | 88.2 | 99.93 | 0.39 | 98.2326 | 5038574 | 45.28 | 51 | 551788 |
| SAMN14411268-GCF_014921435.1 | V170 | SAMN14411268 | GCF_014921435.1 | NonPC | Public | | 2016 | China | Env | 89.6 | 99.93 | 1.13 | 98.2979 | 5293317 | 45 | 28 | 1265052 |
| SAMN14411269-SRR11357120 | V173 | SAMN14411269 | SRR11357120 | NonPC | Public | | 2016 | China | Env | 88.5 | 99.77 | 0.63 | 98.2538 | 4962930 | 45.34 | 47 | 277700 |

| | | | | | | | | | | | | | | | | |
|---|---|---|---|---|---|---|---|---|---|---|---|---|---|---|---|---|
| SAMN14411270-GCF_014921455.1 | V174 | SAMN14411270 | GCF_014921455.1 | NonPC | Public | 2016 | China | Env | 88.2 | 100 | 0.46 | 98.177 | 5153708 | 45.15 | 66 | 802818 |
| SAMN14411271-GCF_014921415.1 | V176 | SAMN14411271 | GCF_014921415.1 | NonPC | Public | 2016 | China | Env | 88.5 | 99.97 | 0.56 | 98.2798 | 5002506 | 45.35 | 53 | 563945 |
| SAMN14411272-GCF_014921395.1 | V177 | SAMN14411272 | GCF_014921395.1 | NonPC | Public | 2016 | China | Env | 88.7 | 99.81 | 0.37 | 98.2401 | 5029222 | 45.27 | 45 | 714866 |
| SAMN14411273-GCF_014921355.1 | V180 | SAMN14411273 | GCF_014921355.1 | NonPC | Public | 2016 | China | Env | 91.8 | 99.7 | 0.41 | 98.4166 | 4961354 | 45.39 | 36 | 682216 |
| SAMN14411274-GCA_014964255.1 | V182 | SAMN14411274 | GCA_014964255.1 | NonPC | Public | 2016 | China | Env | 88.2 | 100 | 0.73 | 98.253 | 5062238 | 45.28 | 62 | 428153 |
| SAMN14411275-GCA_014964165.1 | V183 | SAMN14411275 | GCA_014964165.1 | NonPC | Public | 2016 | China | Env | 88.7 | 99.8 | 0.46 | 98.2155 | 5074608 | 45.22 | 64 | 572532 |
| SAMN14411276-GCA_014964105.1 | V188 | SAMN14411276 | GCA_014964105.1 | NonPC | Public | 2016 | China | Env | 87.6 | 99.97 | 0.3 | 98.1534 | 5065830 | 45.24 | 60 | 438783 |
| SAMN14411277-GCA_014964185.1 | V189 | SAMN14411277 | GCA_014964185.1 | NonPC | Public | 2016 | China | Env | 90 | 99.86 | 0.05 | 98.3351 | 5368892 | 45.08 | 60 | 377416 |
| SAMN14411278-GCA_014964125.1 | V191 | SAMN14411278 | GCA_014964125.1 | NonPC | Public | 2016 | China | Env | 88.2 | 99.8 | 0.48 | 98.255 | 5054173 | 45.22 | 63 | 346630 |
| SAMN14411279-GCA_014964145.1 | V192 | SAMN14411279 | GCA_014964145.1 | NonPC | Public | 2016 | China | Env | 88.3 | 100 | 0.49 | 98.2461 | 5077523 | 45.29 | 46 | 768426 |
| SAMN14411280-GCA_014964085.1 | V193 | SAMN14411280 | GCA_014964085.1 | NonPC | Public | 2016 | China | Env | 88.4 | 99.66 | 0.41 | 98.2444 | 5096269 | 45.27 | 62 | 619282 |
| SAMN14411281-GCA_014964045.1 | V194 | SAMN14411281 | GCA_014964045.1 | NonPC | Public | 2016 | China | Env | 92.1 | 100 | 0.13 | 98.2706 | 5218439 | 45.14 | 56 | 747013 |
| SAMN14411282-GCA_014964055.1 | V195 | SAMN14411282 | GCA_014964055.1 | NonPC | Public | 2016 | China | Env | 89.9 | 100 | 0 | 98.2271 | 5200432 | 45.23 | 66 | 695863 |
| SAMN14411283-GCA_014963985.1 | V197 | SAMN14411283 | GCA_014963985.1 | NonPC | Public | 2016 | China | Env | 87.6 | 99.93 | 0.37 | 98.1497 | 5079839 | 45.24 | 69 | 439477 |
| SAMN14411284-GCA_014964005.1 | V198 | SAMN14411284 | GCA_014964005.1 | NonPC | Public | 2016 | China | Env | 89.5 | 99.86 | 0.2 | 98.3219 | 5021350 | 45.35 | 63 | 768242 |
| SAMN14411285-GCA_014964025.1 | V199 | SAMN14411285 | GCA_014964025.1 | NonPC | Public | 2016 | China | Env | 88.3 | 99.8 | 0.34 | 98.312 | 5053543 | 45.24 | 56 | 736704 |
| SAMN14411286-GCA_014963945.1 | V200 | SAMN14411286 | GCA_014963945.1 | NonPC | Public | 2016 | China | Env | 88.1 | 99.77 | 0.46 | 98.1949 | 4955925 | 45.32 | 35 | 709357 |
| SAMN14411287-GCA_014963925.1 | V201 | SAMN14411287 | GCA_014963925.1 | NonPC | Public | 2016 | China | Env | 91.7 | 99.93 | 0.41 | 98.3522 | 5081625 | 45.26 | 33 | 654217 |
| SAMN14411288-GCA_014963965.1 | V202 | SAMN14411288 | GCA_014963965.1 | NonPC | Public | 2016 | China | Env | 92.2 | 100 | 0.1 | 98.3272 | 5214878 | 45.14 | 52 | 572848 |
| SAMN14411289-GCA_014963905.1 | V203 | SAMN14411289 | GCA_014963905.1 | NonPC | Public | 2016 | China | Env | 88.2 | 99.93 | 0.63 | 98.2485 | 5101478 | 45.06 | 40 | 876880 |
| SAMN14411290-GCA_014963885.1 | V204 | SAMN14411290 | GCA_014963885.1 | NonPC | Public | 2016 | China | Env | 88.5 | 99.66 | 0.39 | 98.2349 | 5152292 | 45.19 | 79 | 580928 |
| SAMN14411291-GCA_014963845.1 | V205 | SAMN14411291 | GCA_014963845.1 | NonPC | Public | 2016 | China | Env | 88.4 | 99.73 | 0.58 | 98.1877 | 5167123 | 45.23 | 56 | 881884 |
| SAMN14411292-SRR11430680 | V206 | SAMN14411292 | SRR11430680 | NonPC | Public | 2015 | China | Env | 88.4 | 99.56 | 0.3 | 98.1755 | 5067128 | 45.24 | 70 | 384926 |
| SAMN14411293-GCF_014921315.1 | V207 | SAMN14411293 | GCF_014921315.1 | NonPC | Public | 2015 | China | Env | 91.5 | 100 | 0.11 | 98.3486 | 5010336 | 45.29 | 53 | 417438 |
| SAMN14411294-GCF_014921335.1 | V208 | SAMN14411294 | GCF_014921335.1 | NonPC | Public | 2015 | China | Env | 88.3 | 99.77 | 0.34 | 98.1636 | 5065843 | 45.24 | 66 | 543153 |
| SAMN14411295-GCF_014921295.1 | V210 | SAMN14411295 | GCF_014921295.1 | NonPC | Public | 2016 | China | Env | 88.4 | 100 | 0.39 | 98.166 | 5174801 | 45.17 | 39 | 910695 |
| SAMN14411296-GCF_014921245.1 | V22 | SAMN14411296 | GCF_014921245.1 | NonPC | Public | 2015 | China | Env | 90.2 | 99.67 | 0.31 | 98.2542 | 5126499 | 45.24 | 77 | 428740 |
| SAMN14411297-GCF_014921275.1 | V23 | SAMN14411297 | GCF_014921275.1 | NonPC | Public | 2015 | China | Env | 88.2 | 99.93 | 0.39 | 98.2331 | 5068650 | 45.28 | 82 | 563192 |
| SAMN14411298-GCF_014921215.1 | V24 | SAMN14411298 | GCF_014921215.1 | NonPC | Public | 2015 | China | Env | 90.1 | 99.81 | 0.07 | 98.3172 | 5149271 | 45.22 | 80 | 400530 |
| SAMN14411299-GCF_014921205.1 | V26 | SAMN14411299 | GCF_014921205.1 | NonPC | Public | 2015 | China | Env | 89 | 99.68 | 0.04 | 98.3389 | 5057729 | 45.31 | 50 | 722416 |
| SAMN14411300-GCF_014921195.1 | V27 | SAMN14411300 | GCF_014921195.1 | NonPC | Public | 2015 | China | Env | 89 | 99.91 | 0.28 | 98.3508 | 5058492 | 45.31 | 51 | 722980 |
| SAMN14411301-GCF_014921125.1 | V3 | SAMN14411301 | GCF_014921125.1 | NonPC | Public | 2015 | China | Env | 91.9 | 99.93 | 0 | 98.3766 | 5106875 | 45.26 | 45 | 651574 |
| SAMN14411302-GCF_014921165.1 | V30 | SAMN14411302 | GCF_014921165.1 | NonPC | Public | 2015 | China | Env | 88.6 | 100 | 0.44 | 98.214 | 5080781 | 45.26 | 45 | 555200 |
| SAMN14411303-GCF_014921155.1 | V33 | SAMN14411303 | GCF_014921155.1 | NonPC | Public | 2015 | China | Env | 88.4 | 99.58 | 0.55 | 98.2261 | 5087379 | 45.17 | 84 | 543205 |
| SAMN14411304-GCF_014921095.1 | V34 | SAMN14411304 | GCF_014921095.1 | NonPC | Public | 2015 | China | Env | 91.3 | 100 | 0.05 | 98.3424 | 5108648 | 45.17 | 43 | 448593 |
| SAMN14411305-SRR11357228 | V35 | SAMN14411305 | SRR11357228 | NonPC | Public | 2015 | China | Env | 88.5 | 99.93 | 0.77 | 98.2037 | 5191001 | 45.18 | 54 | 541670 |
| SAMN14411306-GCF_014921045.1 | V36 | SAMN14411306 | GCF_014921045.1 | NonPC | Public | 2015 | China | Env | 91.3 | 100 | 0.33 | 98.3795 | 5102323 | 45.17 | 36 | 910905 |
| SAMN14411307-GCF_014921075.1 | V39 | SAMN14411307 | GCF_014921075.1 | NonPC | Public | 2015 | China | Env | 91.3 | 99.88 | 0.13 | 98.354 | 4987232 | 45.36 | 42 | 404868 |
| SAMN14411308-GCF_014920975.1 | V41 | SAMN14411308 | GCF_014920975.1 | NonPC | Public | 2015 | China | Env | 88.2 | 99.8 | 0.58 | 98.2351 | 5263420 | 45.25 | 69 | 503349 |
| SAMN14411310-GCF_014921005.1 | V48 | SAMN14411310 | GCF_014921005.1 | NonPC | Public | 2015 | China | Env | 91.4 | 100 | 0.54 | 98.2585 | 5327336 | 45.25 | 88 | 253130 |
| SAMN14411311-GCF_014920985.1 | V50 | SAMN14411311 | GCF_014920985.1 | NonPC | Public | 2015 | China | Env | 90 | 100 | 0.41 | 98.2773 | 5416880 | 45.26 | 94 | 543106 |
| SAMN14411312-GCF_014920935.1 | V52 | SAMN14411312 | GCF_014920935.1 | NonPC | Public | 2015 | China | Env | 89.9 | 100 | 0.14 | 98.2866 | 5374332 | 45.25 | 96 | 284447 |
| SAMN14411313-SRR11357220 | V53 | SAMN14411313 | SRR11357220 | NonPC | Public | 2015 | China | Env | 91.4 | 99.56 | 0.34 | 98.2691 | 5235712 | 45.24 | 82 | 300459 |
| SAMN14411314-SRR11357219 | V55 | SAMN14411314 | SRR11357219 | NonPC | Public | 2015 | China | Env | 91.5 | 99.69 | 0.27 | 98.2188 | 5115956 | 45.33 | 52 | 486269 |
| SAMN14411315-GCF_014920925.1 | V56 | SAMN14411315 | GCF_014920925.1 | NonPC | Public | 2015 | China | Env | 89.9 | 99.73 | 0.53 | 98.2207 | 5287258 | 45.11 | 35 | 535342 |
| SAMN14411316-GCF_014920885.1 | V58 | SAMN14411316 | GCF_014920885.1 | NonPC | Public | 2015 | China | Env | 91.4 | 99.93 | 0.18 | 98.2728 | 5006777 | 45.37 | 45 | 401783 |
| SAMN14411317-SRR11357216 | V64 | SAMN14411317 | SRR11357216 | NonPC | Public | 2015 | China | Env | 91.6 | 99.39 | 0.33 | 98.35 | 5412111 | 45.13 | 67 | 461365 |
| SAMN14411318-GCF_014920825.1 | V65 | SAMN14411318 | GCF_014920825.1 | NonPC | Public | 2015 | China | Env | 91.5 | 99.68 | 0.12 | 98.3342 | 5349950 | 45.2 | 58 | 831926 |

| | | | | | | | | | | | | | | | | |
|---|---|---|---|---|---|---|---|---|---|---|---|---|---|---|---|---|
| SAMN14411319-GCF_014920785.1 | V66 | SAMN14411319 | GCF_014920785.1 | NonPC | Public | 2015 | China | Env | 89.3 | 99.91 | 0.05 | 98.3577 | 5046351 | 45.26 | 55 | 250418 |
| SAMN14411320-SRR11357212 | V67 | SAMN14411320 | SRR11357212 | NonPC | Public | 2015 | China | Env | 90.8 | 99.64 | 0.43 | 98.3149 | 5391801 | 45.34 | 77 | 237750 |
| SAMN14411321-GCF_014920815.1 | V69 | SAMN14411321 | GCF_014920815.1 | NonPC | Public | 2015 | China | Env | 89.8 | 99.84 | 0.16 | 98.3221 | 5010182 | 45.39 | 25 | 1004077 |
| SAMN14411322-GCF_014920755.1 | V7 | SAMN14411322 | GCF_014920755.1 | NonPC | Public | 2015 | China | Env | 91.6 | 99.86 | 0.27 | 98.3562 | 5077355 | 45.32 | 54 | 517850 |
| SAMN14411323-SRR11357209 | V73 | SAMN14411323 | SRR11357209 | NonPC | Public | 2015 | China | Env | 91.4 | 99.16 | 0.2 | 98.3361 | 5119139 | 45.18 | 91 | 519015 |
| SAMN14411324-GCF_014920675.1 | V75 | SAMN14411324 | GCF_014920675.1 | NonPC | Public | 2015 | China | Env | 91.9 | 99.93 | 0 | 98.3121 | 5057340 | 45.34 | 38 | 549270 |
| SAMN14411325-GCF_014920685.1 | V76 | SAMN14411325 | GCF_014920685.1 | NonPC | Public | 2015 | China | Env | 91.4 | 99.8 | 0.03 | 98.3464 | 5005622 | 45.36 | 52 | 406934 |
| SAMN14411326-GCF_014920695.1 | V77 | SAMN14411326 | GCF_014920695.1 | NonPC | Public | 2015 | China | Env | 89.9 | 99.73 | 0.06 | 98.2982 | 5112214 | 45.42 | 47 | 1067175 |
| SAMN14411327-SRR11430668 | V78 | SAMN14411327 | SRR11430668 | NonPC | Public | 2015 | China | Env | 88.7 | 99.66 | 0.39 | 98.1667 | 5071019 | 45.22 | 71 | 553449 |
| SAMN14411329-GCF_014920595.1 | V81 | SAMN14411329 | GCF_014920595.1 | NonPC | Public | 2015 | China | Env | 88.6 | 99.93 | 0.36 | 98.258 | 5014878 | 45.3 | 37 | 639588 |
| SAMN14411330-GCF_014920565.1 | V82 | SAMN14411330 | GCF_014920565.1 | NonPC | Public | 2015 | China | Env | 89.7 | 100 | 0.09 | 98.2374 | 5110069 | 45.38 | 44 | 527619 |
| SAMN14411331-GCF_014920605.1 | V85 | SAMN14411331 | GCF_014920605.1 | NonPC | Public | 2015 | China | Env | 89.8 | 99.93 | 0.27 | 98.2971 | 4963027 | 45.38 | 59 | 807999 |
| SAMN14411332-GCF_014920545.1 | V86 | SAMN14411332 | GCF_014920545.1 | NonPC | Public | 2015 | China | Env | 91.1 | 100 | 0.05 | 98.2748 | 5146385 | 45.29 | 48 | 536635 |
| SAMN14411333-SRR11357163 | V9 | SAMN14411333 | SRR11357163 | NonPC | Public | 2015 | China | Env | 91.5 | 99.18 | 0.54 | 98.271 | 5365168 | 45.33 | 75 | 533912 |
| SAMN14411334-GCF_014920495.1 | V90 | SAMN14411334 | GCF_014920495.1 | NonPC | Public | 2015 | China | Env | 91.9 | 100 | 0.35 | 98.305 | 5022265 | 45.33 | 48 | 317097 |
| SAMN14411335-GCF_014920505.1 | V93 | SAMN14411335 | GCF_014920505.1 | NonPC | Public | 2015 | China | Env | 90.7 | 100 | 0.05 | 98.2414 | 5075398 | 45.28 | 41 | 782398 |
| SAMN14411336-SRR11357160 | Vp101 | SAMN14411336 | SRR11357160 | NonPC | Public | 2013 | China | Env | 88.2 | 99.77 | 0.36 | 98.2079 | 5004714 | 45.33 | 47 | 448794 |
| SAMN14411337-GCF_014920455.1 | Vp103 | SAMN14411337 | GCF_014920455.1 | NonPC | Public | 2013 | China | Env | 89.8 | 100 | 0.09 | 98.2949 | 5068965 | 45.38 | 45 | 400904 |
| SAMN14411338-GCF_014920415.1 | Vp125 | SAMN14411338 | GCF_014920415.1 | NonPC | Public | 2013 | China | Env | 90.2 | 100 | 0.42 | 98.2467 | 5124271 | 45.34 | 39 | 464075 |
| SAMN14411339-GCF_014920375.1 | Vp129 | SAMN14411339 | GCF_014920375.1 | NonPC | Public | 2013 | China | Env | 89.6 | 99.93 | 0.05 | 98.282 | 4968209 | 45.43 | 38 | 778110 |
| SAMN14411340-SRR11357155 | Vp132 | SAMN14411340 | SRR11357155 | NonPC | Public | 2013 | China | Env | 89 | 98.67 | 0.33 | 98.2478 | 5126759 | 45.35 | 50 | 1025514 |
| SAMN14411341-GCF_014920385.1 | Vp145 | SAMN14411341 | GCF_014920385.1 | NonPC | Public | 2013 | China | Env | 91.6 | 99.93 | 0.59 | 98.2896 | 5288142 | 45.19 | 69 | 300635 |
| SAMN14411342-GCF_014920315.1 | Vp146 | SAMN14411342 | GCF_014920315.1 | NonPC | Public | 2013 | China | Env | 91.1 | 100 | 0 | 98.3642 | 4988652 | 45.37 | 51 | 1029259 |
| SAMN14411343-GCF_014920355.1 | Vp154 | SAMN14411343 | GCF_014920355.1 | NonPC | Public | 2013 | China | Env | 88.6 | 100 | 0.61 | 98.1856 | 5049534 | 45.31 | 62 | 329043 |
| SAMN14411344-GCF_014920335.1 | Vp155 | SAMN14411344 | GCF_014920335.1 | NonPC | Public | 2013 | China | Env | 91.2 | 99.81 | 0.39 | 98.2884 | 5285155 | 45.2 | 79 | 300636 |
| SAMN14411345-GCF_014920275.1 | Vp156 | SAMN14411345 | GCF_014920275.1 | NonPC | Public | 2013 | China | Env | 91.8 | 100 | 0 | 98.3207 | 5327958 | 45.19 | 78 | 315593 |
| SAMN14411346-SRR11357149 | Vp158 | SAMN14411346 | SRR11357149 | NonPC | Public | 2013 | China | Env | 88.6 | 99.93 | 0.92 | 98.1841 | 5176283 | 45.29 | 96 | 398175 |
| SAMN14411347-SRR11357148 | Vp161 | SAMN14411347 | SRR11357148 | NonPC | Public | 2013 | China | Env | 89.8 | 99.12 | 0 | 98.3056 | 5053802 | 45.3 | 52 | 466942 |
| SAMN14411348-GCF_014920235.1 | Vp165 | SAMN14411348 | GCF_014920235.1 | NonPC | Public | 2013 | China | Env | 90.6 | 99.97 | 0.54 | 98.3379 | 5145733 | 45.25 | 63 | 333693 |
| SAMN14411349-SRR11357145 | Vp166 | SAMN14411349 | SRR11357145 | NonPC | Public | 2013 | China | Env | 91.7 | 99.64 | 0 | 98.3624 | 5030387 | 45.42 | 44 | 536543 |
| SAMN14411350-SRR11357144 | Vp18 | SAMN14411350 | SRR11357144 | NonPC | Public | 2012 | China | Env | 88 | 100 | 0.4 | 98.237 | 5008957 | 45.42 | 84 | 543594 |
| SAMN14411351-GCF_014920135.1 | Vp181 | SAMN14411351 | GCF_014920135.1 | NonPC | Public | 2013 | China | Env | 88.5 | 100 | 0.3 | 98.1927 | 5091061 | 45.25 | 32 | 1350939 |
| SAMN14411352-SRR11357142 | Vp184 | SAMN14411352 | SRR11357142 | NonPC | Public | 2013 | China | Env | 88.2 | 99.18 | 1.02 | 98.1878 | 5094329 | 45.2 | 84 | 390338 |
| SAMN14411353-SRR11357141 | Vp185 | SAMN14411353 | SRR11357141 | NonPC | Public | 2013 | China | Env | 88.3 | 99.86 | 0.77 | 98.1904 | 5209474 | 45.15 | 72 | 486339 |
| SAMN14411354-SRR11357140 | Vp194 | SAMN14411354 | SRR11357140 | NonPC | Public | 2013 | China | Env | 91.6 | 99.63 | 0.09 | 98.3842 | 5201001 | 45.27 | 52 | 522860 |
| SAMN14411355-SRR11357139 | Vp198 | SAMN14411355 | SRR11357139 | NonPC | Public | 2013 | China | Env | 88.3 | 99.13 | 0.3 | 98.1915 | 5114896 | 45.18 | 53 | 564648 |
| SAMN14411356-SRR11357138 | Vp203 | SAMN14411356 | SRR11357138 | NonPC | Public | 2013 | China | Env | 91.2 | 98.81 | 0.08 | 98.2182 | 5102195 | 45.23 | 52 | 652261 |
| SAMN14411357-GCF_014920045.1 | Vp224 | SAMN14411357 | GCF_014920045.1 | NonPC | Public | 2013 | China | Env | 89.9 | 100 | 0.14 | 98.2393 | 5340383 | 45.28 | 47 | 477839 |
| SAMN14411358-GCF_014920055.1 | Vp229 | SAMN14411358 | GCF_014920055.1 | NonPC | Public | 2013 | China | Env | 88.3 | 100 | 0.49 | 98.1959 | 5150433 | 45.2 | 31 | 1764997 |
| SAMN14411359-GCF_014919975.1 | Vp232 | SAMN14411359 | GCF_014919975.1 | NonPC | Public | 2013 | China | Env | 90.1 | 100 | 0 | 98.2654 | 5178188 | 45.28 | 65 | 405841 |
| SAMN14411360-GCF_014919935.1 | Vp234 | SAMN14411360 | GCF_014919935.1 | NonPC | Public | 2013 | China | Env | 91.9 | 100 | 0.54 | 98.3093 | 5273428 | 45.31 | 53 | 518447 |
| SAMN14411361-GCF_014920015.1 | Vp239 | SAMN14411361 | GCF_014920015.1 | NonPC | Public | 2013 | China | Env | 91.9 | 99.73 | 0.05 | 98.3892 | 5054176 | 45.4 | 54 | 429738 |
| SAMN14411362-GCF_014919965.1 | Vp242 | SAMN14411362 | GCF_014919965.1 | NonPC | Public | 2013 | China | Env | 89.8 | 99.86 | 0 | 98.3069 | 5044917 | 45.31 | 72 | 389860 |
| SAMN14411363-SRR11357134 | Vp248 | SAMN14411363 | SRR11357134 | NonPC | Public | 2013 | China | Env | 89 | 99.8 | 0.7 | 98.1859 | 5157564 | 45.31 | 85 | 358205 |
| SAMN14411364-SRR11357133 | Vp249 | SAMN14411364 | SRR11357133 | NonPC | Public | 2013 | China | Env | 88.2 | 99.8 | 0.39 | 98.191 | 4976272 | 45.36 | 78 | 452856 |
| SAMN14411365-SRR11357132 | Vp251 | SAMN14411365 | SRR11357132 | NonPC | Public | 2013 | China | Env | 89.5 | 99.39 | 0 | 98.2193 | 5333808 | 45.01 | 65 | 462285 |
| SAMN14411366-GCF_014919845.1 | Vp269 | SAMN14411366 | GCF_014919845.1 | NonPC | Public | 2013 | China | Env | 88.2 | 100 | 0.39 | 98.1855 | 5137374 | 45.22 | 67 | 492536 |
| SAMN14411367-GCF_014919865.1 | Vp271 | SAMN14411367 | GCF_014919865.1 | NonPC | Public | 2013 | China | Env | 88.3 | 99.73 | 0.49 | 98.2066 | 5120577 | 45.24 | 78 | 599014 |

| | | | | | | | | | | | | | | | | |
|---|---|---|---|---|---|---|---|---|---|---|---|---|---|---|---|---|
| SAMN14411368-SRR11357129 | Vp272 | SAMN14411368 | SRR11357129 | NonPC | Public | 2013 | China | Env | 88.9 | 99.49 | 0.58 | 98.3135 | 5227865 | 45.37 | 99 | 233183 |
| SAMN14411369-GCF_014919815.1 | Vp277 | SAMN14411369 | GCF_014919815.1 | NonPC | Public | 2013 | China | Env | 89.7 | 100 | 0.27 | 98.4096 | 4968153 | 45.43 | 48 | 470405 |
| SAMN14411370-GCF_014919795.1 | Vp278 | SAMN14411370 | GCF_014919795.1 | NonPC | Public | 2013 | China | Env | 88.3 | 100 | 0.39 | 98.1875 | 5076953 | 45.24 | 72 | 492523 |
| SAMN14411371-SRR11357127 | Vp279 | SAMN14411371 | SRR11357127 | NonPC | Public | 2013 | China | Env | 88.2 | 98.94 | 1.07 | 98.1684 | 5055975 | 45.23 | 55 | 366888 |
| SAMN14411372-GCF_014919745.1 | Vp284 | SAMN14411372 | GCF_014919745.1 | NonPC | Public | 2013 | China | Env | 91.3 | 100 | 0 | 98.3151 | 5123642 | 45.17 | 53 | 1069292 |
| SAMN14411373-GCF_014919755.1 | Vp286 | SAMN14411373 | GCF_014919755.1 | NonPC | Public | 2013 | China | Env | 91.7 | 99.93 | 0 | 98.3025 | 5163314 | 45.25 | 49 | 957867 |
| SAMN14411374-GCF_014919715.1 | Vp287 | SAMN14411374 | GCF_014919715.1 | NonPC | Public | 2013 | China | Env | 88.6 | 100 | 0.64 | 98.1844 | 5254209 | 45.16 | 67 | 983952 |
| SAMN14411375-SRR11357271 | Vp290 | SAMN14411375 | SRR11357271 | NonPC | Public | 2013 | China | Env | 88.3 | 99.37 | 0.61 | 98.2647 | 5053769 | 45.26 | 60 | 445899 |
| SAMN14411376-GCF_014919655.1 | Vp293 | SAMN14411376 | GCF_014919655.1 | NonPC | Public | 2013 | China | Env | 92.4 | 100 | 0.32 | 98.393 | 5111649 | 45.36 | 33 | 1259329 |
| SAMN14411377-GCF_014919635.1 | Vp297 | SAMN14411377 | GCF_014919635.1 | NonPC | Public | 2013 | China | Env | 92.1 | 100 | 0.27 | 98.3429 | 5027506 | 45.34 | 42 | 910867 |
| SAMN14411378-SRR11357269 | Vp299 | SAMN14411378 | SRR11357269 | NonPC | Public | 2013 | China | Env | 88.2 | 99.44 | 0.69 | 98.2343 | 5038138 | 45.25 | 57 | 316334 |
| SAMN14411379-GCF_014919555.1 | Vp301 | SAMN14411379 | GCF_014919555.1 | NonPC | Public | 2013 | China | Env | 88.3 | 100 | 0.39 | 98.1911 | 5098618 | 45.3 | 57 | 662485 |
| SAMN14411380-GCA_014919615.1 | Vp306 | SAMN14411380 | GCA_014919615.1 | NonPC | Public | 2013 | China | Env | 88.3 | 100 | 0.43 | 98.2083 | 5177700 | 45.17 | 67 | 576427 |
| SAMN14411381-GCF_014919575.1 | Vp313 | SAMN14411381 | GCF_014919575.1 | NonPC | Public | 2013 | China | Env | 92.1 | 99.93 | 0.27 | 98.3683 | 5026718 | 45.34 | 53 | 421633 |
| SAMN14411382-SRR11430674 | Vp315 | SAMN14411382 | SRR11430674 | NonPC | Public | 2013 | China | Env | 88.3 | 99.05 | 0.36 | 98.2367 | 5075057 | 45.27 | 78 | 677268 |
| SAMN14411383-SRR11357267 | Vp320 | SAMN14411383 | SRR11357267 | NonPC | Public | 2013 | China | Env | 91.1 | 99.36 | 0 | 98.2205 | 5141138 | 45.18 | 38 | 400428 |
| SAMN14411384-SRR11357266 | Vp322 | SAMN14411384 | SRR11357266 | NonPC | Public | 2013 | China | Env | 87.6 | 99.93 | 0.73 | 98.2743 | 5039233 | 45.36 | 79 | 542917 |
| SAMN14411385-SRR11430673 | Vp324 | SAMN14411385 | SRR11430673 | NonPC | Public | 2013 | China | Env | 89.4 | 99.88 | 0.05 | 98.2491 | 5284636 | 45.16 | 87 | 638686 |
| SAMN14411386-GCF_014919485.1 | Vp326 | SAMN14411386 | GCF_014919485.1 | NonPC | Public | 2013 | China | Env | 89 | 99.93 | 0.43 | 98.1914 | 5144110 | 45.23 | 85 | 504646 |
| SAMN14411387-SRR11357264 | Vp328 | SAMN14411387 | SRR11357264 | NonPC | Public | 2013 | China | Env | 90.6 | 99.5 | 0.27 | 98.3316 | 5053302 | 45.31 | 42 | 862859 |
| SAMN14411388-SRR11357263 | Vp331 | SAMN14411388 | SRR11357263 | NonPC | Public | 2013 | China | Env | 90 | 99.36 | 0.12 | 98.2383 | 5186313 | 45.23 | 53 | 526110 |
| SAMN14411389-GCF_014919415.1 | Vp332 | SAMN14411389 | GCF_014919415.1 | NonPC | Public | 2013 | China | Env | 88.4 | 99.9 | 0.36 | 98.2603 | 5022714 | 45.29 | 56 | 563217 |
| SAMN14411390-GCF_014919395.1 | Vp333 | SAMN14411390 | GCF_014919395.1 | NonPC | Public | 2013 | China | Env | 90.5 | 99.86 | 0.15 | 98.2639 | 5097448 | 45.37 | 62 | 1032208 |
| SAMN14411391-SRR11430672 | Vp347 | SAMN14411391 | SRR11430672 | NonPC | Public | 2013 | China | Env | 91.4 | 99.8 | 0.3 | 98.3732 | 5182025 | 45.22 | 89 | 469992 |
| SAMN14411392-GCF_014919355.1 | Vp350 | SAMN14411392 | GCF_014919355.1 | NonPC | Public | 2013 | China | Env | 89.7 | 99.93 | 0.03 | 98.2067 | 5148138 | 45.29 | 42 | 523937 |
| SAMN14411394-GCF_014919305.1 | Vp359 | SAMN14411394 | GCF_014919305.1 | NonPC | Public | 2013 | China | Env | 91 | 99.93 | 0.27 | 98.377 | 5146044 | 45.25 | 69 | 320945 |
| SAMN14411395-SRR11430670 | Vp360 | SAMN14411395 | SRR11430670 | NonPC | Public | 2013 | China | Env | 89.6 | 98.08 | 0.34 | 98.1596 | 5038929 | 45.3 | 52 | 532019 |
| SAMN14411396-GCF_014919255.1 | Vp368 | SAMN14411396 | GCF_014919255.1 | NonPC | Public | 2013 | China | Env | 89.9 | 99.73 | 0 | 98.1826 | 5449191 | 45.18 | 114 | 469252 |
| SAMN14411397-GCF_014919195.1 | Vp370 | SAMN14411397 | GCF_014919195.1 | NonPC | Public | 2013 | China | Env | 91 | 99.86 | 0.05 | 98.2514 | 5065845 | 45.3 | 53 | 443671 |
| SAMN14411399-SRR11357255 | Vp374 | SAMN14411399 | SRR11357255 | NonPC | Public | 2013 | China | Env | 90 | 99.56 | 0.93 | 98.2197 | 5365504 | 45.21 | 337 | 487283 |
| SAMN14411401-GCA_014930295.1 | Vp387 | SAMN14411401 | GCA_014930295.1 | NonPC | Public | 2013 | China | Env | 91.6 | 100 | 0 | 98.3035 | 5136617 | 45.3 | 77 | 523457 |
| SAMN14411402-GCA_014930285.1 | Vp396 | SAMN14411402 | GCA_014930285.1 | NonPC | Public | 2014 | China | Env | 91.6 | 100 | 0 | 98.2814 | 5143367 | 45.21 | 56 | 417728 |
| SAMN14411403-GCA_014930275.1 | Vp397 | SAMN14411403 | GCA_014930275.1 | NonPC | Public | 2014 | China | Env | 91.2 | 100 | 0.09 | 98.2684 | 5391402 | 45.19 | 52 | 473734 |
| SAMN14411404-GCA_014930255.1 | Vp398 | SAMN14411404 | GCA_014930255.1 | NonPC | Public | 2014 | China | Env | 91.6 | 100 | 0 | 98.2646 | 5138911 | 45.2 | 41 | 452256 |
| SAMN14411406-GCF_014919165.1 | Vp412 | SAMN14411406 | GCF_014919165.1 | NonPC | Public | 2013 | China | Env | 91.5 | 100 | 0.54 | 98.3159 | 5243761 | 45.35 | 53 | 888918 |
| SAMN14411407-GCF_014919155.1 | Vp415 | SAMN14411407 | GCF_014919155.1 | NonPC | Public | 2013 | China | Env | 89.5 | 99.93 | 0.14 | 98.2915 | 5613924 | 45.32 | 87 | 384997 |
| SAMN14411408-SRR11357247 | Vp420 | SAMN14411408 | SRR11357247 | NonPC | Public | 2013 | China | Env | 88.1 | 98.57 | 0.58 | 98.1987 | 5113103 | 45.16 | 70 | 280679 |
| SAMN14411409-GCF_014919095.1 | Vp421 | SAMN14411409 | GCF_014919095.1 | NonPC | Public | 2013 | China | Env | 88.7 | 100 | 0.66 | 98.2098 | 5104224 | 45.22 | 48 | 737129 |
| SAMN14411410-GCF_014919105.1 | Vp425 | SAMN14411410 | GCF_014919105.1 | NonPC | Public | 2013 | China | Env | 89.8 | 100 | 0.3 | 98.2979 | 5159487 | 45.28 | 55 | 401256 |
| SAMN14411411-GCF_014919075.1 | Vp427 | SAMN14411411 | GCF_014919075.1 | NonPC | Public | 2013 | China | Env | 91.2 | 100 | 0.37 | 98.2504 | 5297940 | 45.29 | 51 | 357392 |
| SAMN14411412-GCF_014919055.1 | Vp428 | SAMN14411412 | GCF_014919055.1 | NonPC | Public | 2013 | China | Env | 88.8 | 100 | 0.71 | 98.1696 | 5149676 | 45.22 | 51 | 1180099 |
| SAMN14411413-GCF_014919015.1 | Vp431 | SAMN14411413 | GCF_014919015.1 | NonPC | Public | 2013 | China | Env | 91.1 | 100 | 0.1 | 98.3075 | 5002059 | 45.31 | 51 | 470620 |
| SAMN14411414-GCF_014919025.1 | Vp432 | SAMN14411414 | GCF_014919025.1 | NonPC | Public | 2013 | China | Env | 91.1 | 100 | 0 | 98.3721 | 4972592 | 45.4 | 67 | 462650 |
| SAMN14411415-SRR11357244 | Vp435 | SAMN14411415 | SRR11357244 | NonPC | Public | 2013 | China | Env | 88.4 | 99.77 | 0.66 | 98.2192 | 5052928 | 45.27 | 65 | 711084 |
| SAMN14411416-SRR11357243 | Vp436 | SAMN14411416 | SRR11357243 | NonPC | Public | 2013 | China | Env | 87.9 | 99.84 | 0.66 | 98.2131 | 5155480 | 45.16 | 79 | 424665 |
| SAMN14411417-GCF_014918955.1 | Vp439 | SAMN14411417 | GCF_014918955.1 | NonPC | Public | 2013 | China | Env | 90 | 100 | 0.33 | 98.2623 | 5256024 | 45.35 | 23 | 1069233 |
| SAMN14411418-GCF_014918925.1 | Vp441 | SAMN14411418 | GCF_014918925.1 | NonPC | Public | 2013 | China | Env | 91.6 | 100 | 0.54 | 98.3159 | 5252676 | 45.33 | 46 | 532296 |
| SAMN14411419-SRR11357240 | Vp442 | SAMN14411419 | SRR11357240 | NonPC | Public | 2013 | China | Env | 91.4 | 99.93 | 0.64 | 98.2979 | 5069032 | 45.36 | 101 | 540673 |

| | | | | | | | | | | | | | | | | |
|---|---|---|---|---|---|---|---|---|---|---|---|---|---|---|---|---|
| SAMN14411420-GCF_014918895.1 | Vp443 | SAMN14411420 | GCF_014918895.1 | NonPC | Public | 2013 | China | Env | 90.7 | 100 | 0.49 | 98.3587 | 5319730 | 45.32 | 39 | 506348 |
| SAMN14411421-SRR11357239 | Vp446 | SAMN14411421 | SRR11357239 | NonPC | Public | 2013 | China | Env | 89.7 | 99.08 | 0.29 | 98.3079 | 5151456 | 45.26 | 43 | 549862 |
| SAMN14411422-GCF_014918855.1 | Vp448 | SAMN14411422 | GCF_014918855.1 | NonPC | Public | 2013 | China | Env | 90 | 100 | 0 | 98.251 | 5289409 | 45.33 | 42 | 538974 |
| SAMN14411423-GCF_014918815.1 | Vp450 | SAMN14411423 | GCF_014918815.1 | NonPC | Public | 2013 | China | Env | 88 | 100 | 0.63 | 98.2565 | 5160419 | 45.17 | 70 | 663507 |
| SAMN14411424-SRR11357235 | Vp453 | SAMN14411424 | SRR11357235 | NonPC | Public | 2013 | China | Env | 88.2 | 99.74 | 0.43 | 98.2602 | 5049447 | 45.27 | 70 | 344239 |
| SAMN14411425-GCF_014918805.1 | Vp456 | SAMN14411425 | GCF_014918805.1 | NonPC | Public | 2013 | China | Env | 91.5 | 100 | 0 | 98.3662 | 4981294 | 45.37 | 34 | 517023 |
| SAMN14411426-GCF_014918765.1 | Vp459 | SAMN14411426 | GCF_014918765.1 | NonPC | Public | 2013 | China | Env | 91.4 | 100 | 0.37 | 98.3447 | 5324628 | 45.32 | 40 | 424632 |
| SAMN14411427-GCF_014918755.1 | Vp460 | SAMN14411427 | GCF_014918755.1 | NonPC | Public | 2013 | China | Env | 89.9 | 100 | 0 | 98.3054 | 5152638 | 45.39 | 51 | 817214 |
| SAMN14411428-SRR11357204 | Vp466 | SAMN14411428 | SRR11357204 | NonPC | Public | 2013 | China | Env | 89.5 | 99.5 | 0.54 | 98.2781 | 5100701 | 45.27 | 79 | 531875 |
| SAMN14411429-GCF_014918685.1 | Vp469 | SAMN14411429 | GCF_014918685.1 | NonPC | Public | 2013 | China | Env | 88.4 | 100 | 0.47 | 98.2127 | 5129626 | 45.21 | 66 | 592602 |
| SAMN14411430-SRR11357202 | Vp483 | SAMN14411430 | SRR11357202 | NonPC | Public | 2013 | China | Env | 88.4 | 98.59 | 0.7 | 98.2272 | 5087622 | 45.25 | 52 | 426200 |
| SAMN14411431-GCF_014918645.1 | Vp486 | SAMN14411431 | GCF_014918645.1 | NonPC | Public | 2013 | China | Env | 91.4 | 100 | 0.56 | 98.2957 | 5411239 | 45.28 | 103 | 332024 |
| SAMN14411432-GCF_014918665.1 | Vp489 | SAMN14411432 | GCF_014918665.1 | NonPC | Public | 2013 | China | Env | 89.7 | 100 | 0.14 | 98.2728 | 5056606 | 45.33 | 71 | 533204 |
| SAMN14411433-GCF_014918635.1 | Vp492 | SAMN14411433 | GCF_014918635.1 | NonPC | Public | 2013 | China | Env | 91.8 | 100 | 0.12 | 98.34 | 5360034 | 45.26 | 97 | 338344 |
| SAMN14411434-GCF_014918585.1 | Vp497 | SAMN14411434 | GCF_014918585.1 | NonPC | Public | 2013 | China | Env | 90 | 100 | 0 | 98.2921 | 5065831 | 45.36 | 44 | 523439 |
| SAMN14411435-GCF_014918605.1 | Vp498 | SAMN14411435 | GCF_014918605.1 | NonPC | Public | 2013 | China | Env | 91.4 | 100 | 0.7 | 98.2406 | 5451725 | 45.26 | 67 | 406152 |
| SAMN14411436-GCF_014918575.1 | Vp50 | SAMN14411436 | GCF_014918575.1 | NonPC | Public | 2012 | China | Env | 92.2 | 99.93 | 0.12 | 98.3824 | 5135349 | 45.22 | 67 | 664762 |
| SAMN14411437-SRR11357198 | Vp54 | SAMN14411437 | SRR11357198 | NonPC | Public | 2012 | China | Env | 91.7 | 99.36 | 0 | 98.2202 | 5246828 | 45.21 | 82 | 542988 |
| SAMN14411438-GCF_014918535.1 | Vp55 | SAMN14411438 | GCF_014918535.1 | NonPC | Public | 2012 | China | Env | 88.4 | 99.8 | 0.66 | 98.2619 | 5034206 | 45.27 | 75 | 502938 |
| SAMN14411439-SRR11357196 | Vp58 | SAMN14411439 | SRR11357196 | NonPC | Public | 2012 | China | Env | 89.4 | 99.93 | 0.05 | 98.2249 | 5341518 | 45.14 | 78 | 323972 |
| SAMN14411440-SRR11357194 | Vp63 | SAMN14411440 | SRR11357194 | NonPC | Public | 2012 | China | Env | 88.6 | 99.8 | 0.7 | 98.2247 | 5331312 | 45.26 | 136 | 368313 |
| SAMN14411441-SRR11357193 | Vp65 | SAMN14411441 | SRR11357193 | NonPC | Public | 2012 | China | Env | 90.8 | 99.05 | 0.05 | 98.3207 | 4955774 | 45.37 | 55 | 471114 |
| SAMN14411442-SRR11357192 | Vp68 | SAMN14411442 | SRR11357192 | NonPC | Public | 2012 | China | Env | 90 | 99.06 | 0.09 | 98.2737 | 5212649 | 45.26 | 62 | 445650 |
| SAMN14411443-GCF_014918435.1 | Vp7 | SAMN14411443 | GCF_014918435.1 | NonPC | Public | 2012 | China | Env | 91.3 | 99.91 | 0.05 | 98.3022 | 5080665 | 45.27 | 78 | 627698 |
| SAMN14411444-SRR11357191 | Vp75 | SAMN14411444 | SRR11357191 | NonPC | Public | 2012 | China | Env | 90.8 | 99.62 | 0.54 | 98.3503 | 5090059 | 45.26 | 63 | 325287 |
| SAMN14411445-SRR11357190 | Vp78 | SAMN14411445 | SRR11357190 | NonPC | Public | 2012 | China | Env | 88.3 | 99.17 | 0.71 | 98.229 | 5141346 | 45.23 | 88 | 561907 |
| SAMN14411446-SRR11357189 | Vp79 | SAMN14411446 | SRR11357189 | NonPC | Public | 2012 | China | Env | 88.3 | 99.51 | 0.71 | 98.2716 | 5136133 | 45.24 | 88 | 561963 |
| SAMN14411447-GCF_014918335.1 | Vp94 | SAMN14411447 | GCF_014918335.1 | NonPC | Public | 2013 | China | Env | 92.2 | 100 | 0.09 | 98.3714 | 5256813 | 45.29 | 59 | 922901 |
| SAMN14411448-SRR11357187 | Vp95 | SAMN14411448 | SRR11357187 | NonPC | Public | 2013 | China | Env | 91.6 | 99.61 | 0.05 | 98.3558 | 5224061 | 45.21 | 63 | 458856 |
| SAMN14524700-GCF_012956445.1 | Vb0744 | SAMN14524700 | GCF_012956445.1 | NonPC | Public | 2015 | China | Env | 90.4 | 100 | 4.56 | 98.2423 | 5342601 | 45.29 | 198 | 382440 |
| SAMN14524706-GCF_012956355.1 | Vb0880 | SAMN14524706 | GCF_012956355.1 | NonPC | Public | 2016 | China | Env | 90.2 | 100 | 4.65 | 98.2245 | 5300880 | 45.26 | 198 | 382512 |
| SAMN14572085-GCF_012845845.1 | RZM-01 | SAMN14572085 | GCF_012845845.1 | NonPC | Public | NA | China | Env | 91.3 | 100 | 0.14 | 98.3494 | 5371156 | 45.13 | 80 | 298487 |
| SAMN14689471-GCA_018085005.1 | ARA_0040407-41 | SAMN14689471 | GCA_018085005.1 | NonPC | Public | 2007 | USA | Env | 91.5 | 100 | 4.78 | 98.3377 | 5331950 | 45.33 | 40 | 402773 |
| SAMN14852540-GCF_013085265.1 | Oyster | SAMN14852540 | GCF_013085265.1 | NonPC | Public | 2003 | India | Env | 89.8 | 100 | 0 | 98.3095 | 5199822 | 45.36 | 67 | 314045 |
| SAMN14861780-SRR11774527 | CGVP22 | SAMN14861780 | SRR11774527 | NonPC | Public | 2015 | South_Korea | Env | 91.6 | 100 | 0.59 | 98.3165 | 5402113 | 45.3 | 85 | 281409 |
| SAMN14884116-SRR11775239 | CGVP3 | SAMN14884116 | SRR11775239 | NonPC | Public | 2015 | South_Korea | Env | 90.9 | 100 | 0.19 | 98.2614 | 5329367 | 45.27 | 108 | 400515 |
| SAMN14884117-SRR11775238 | CGVP8 | SAMN14884117 | SRR11775238 | NonPC | Public | 2015 | South_Korea | Env | 90.9 | 100 | 0.19 | 98.3101 | 5325726 | 45.28 | 114 | 347067 |
| SAMN14884118-SRR11775237 | CGVP10 | SAMN14884118 | SRR11775237 | NonPC | Public | 2015 | South_Korea | Env | 90.9 | 100 | 0.19 | 98.323 | 5318159 | 45.28 | 106 | 432756 |
| SAMN14884119-SRR11775236 | CAVP11 | SAMN14884119 | SRR11775236 | NonPC | Public | 2015 | South_Korea | Env | 90.9 | 100 | 0.19 | 98.2784 | 5327144 | 45.27 | 112 | 347067 |
| SAMN14884120-SRR11775235 | CGVP18 | SAMN14884120 | SRR11775235 | NonPC | Public | 2015 | South_Korea | Env | 90.9 | 100 | 0.19 | 98.2869 | 5325853 | 45.27 | 117 | 347067 |
| SAMN14941690-SRR13023679 | XMO16 | SAMN14941690 | SRR13023679 | NonPC | Public | 2018 | China | Env | 89.7 | 100 | 0.33 | 98.314 | 5045111 | 45.32 | 43 | 670084 |
| SAMN14988329-GCF_014705665.1 | R-14810/1 | SAMN14988329 | GCF_014705665.1 | NonPC | Public | 2016 | Russia | Clinical | 92.7 | 100 | 0.18 | 98.3604 | 5140702 | 45.19 | 40 | 304128 |
| SAMN15054260-GCA_015774675.1 | PNUSAV001188 | SAMN15054260 | GCA_015774675.1 | NonPC | Public | NA | USA | Clinical | 89.1 | 100 | 0.48 | 98.2843 | 5136105 | 45.38 | 248 | 56848 |
| SAMN15294466-GCA_019686035.1 | ICDC-VP01784 | SAMN15294466 | GCA_019686035.1 | PC | Public | 2016 | China | Clinical | 97.6 | 100 | 0 | 99.9725 | 5151398 | 45.27 | 41 | 783436 |
| SAMN15294467-GCA_019686095.1 | ICDC-VP01786 | SAMN15294467 | GCA_019686095.1 | PC | Public | 2016 | China | Clinical | 97.7 | 100 | 0 | 99.9727 | 5154616 | 45.27 | 38 | 783652 |
| SAMN15294468-GCA_019686045.1 | ICDC-VP01787 | SAMN15294468 | GCA_019686045.1 | PC | Public | 2016 | China | Clinical | 97.7 | 100 | 0 | 99.9742 | 5131369 | 45.28 | 38 | 783282 |
| SAMN15294469-GCA_019685975.1 | ICDC-VP01791 | SAMN15294469 | GCA_019685975.1 | PC | Public | 2016 | China | Clinical | 94.6 | 100 | 0 | 99.9752 | 5154327 | 45.27 | 42 | 783301 |
| SAMN15294470-GCA_019686015.1 | ICDC-VP01793 | SAMN15294470 | GCA_019686015.1 | PC | Public | 2017 | China | Clinical | 97.7 | 100 | 0 | 99.9749 | 5154498 | 45.27 | 41 | 783220 |

| | | | | | | | | | | | | | | | | |
|---|---|---|---|---|---|---|---|---|---|---|---|---|---|---|---|---|
| SAMN15294471-GCA_019685945.1 | ICDC-VP01794 | SAMN15294471 | GCA_019685945.1 | PC | Public | 2017 | China | Clinical | 94.6 | 100 | 0 | 99.969 | 5154053 | 45.27 | 40 | 580818 |
| SAMN15294472-GCA_019685995.1 | ICDC-VP01797 | SAMN15294472 | GCA_019685995.1 | PC | Public | 2017 | China | Clinical | 97.7 | 100 | 0 | 99.9744 | 5136476 | 45.26 | 39 | 580549 |
| SAMN15294473-GCA_019685915.1 | ICDC-VP01799 | SAMN15294473 | GCA_019685915.1 | PC | Public | 2017 | China | Clinical | 94.5 | 100 | 0 | 99.9747 | 5151323 | 45.27 | 43 | 783430 |
| SAMN15294474-GCA_019685895.1 | ICDC-VP01800 | SAMN15294474 | GCA_019685895.1 | PC | Public | 2017 | China | Clinical | 94.6 | 100 | 0 | 99.9737 | 5154942 | 45.27 | 39 | 783709 |
| SAMN15294475-GCA_019685935.1 | ICDC-VP01801 | SAMN15294475 | GCA_019685935.1 | PC | Public | 2017 | China | Clinical | 98 | 100 | 0 | 99.9919 | 5078685 | 45.28 | 49 | 542649 |
| SAMN15294476-GCA_019685865.1 | ICDC-VP01802 | SAMN15294476 | GCA_019685865.1 | PC | Public | 2017 | China | Clinical | 97.6 | 100 | 0 | 99.9726 | 5150755 | 45.27 | 44 | 580818 |
| SAMN15294477-GCA_019685835.1 | ICDC-VP01803 | SAMN15294477 | GCA_019685835.1 | PC | Public | 2017 | China | Clinical | 97.3 | 100 | 0 | 99.9728 | 5126347 | 45.28 | 45 | 580444 |
| SAMN15294478-GCA_019685805.1 | ICDC-VP01805 | SAMN15294478 | GCA_019685805.1 | PC | Public | 2018 | China | Clinical | 98.1 | 100 | 0 | 99.986 | 5079126 | 45.28 | 44 | 821369 |
| SAMN15294479-GCA_019685755.1 | ICDC-VP01806 | SAMN15294479 | GCA_019685755.1 | PC | Public | 2018 | China | Clinical | 98.1 | 100 | 0 | 99.9886 | 5078536 | 45.28 | 46 | 542649 |
| SAMN15294480-GCA_019685775.1 | ICDC-VP01809 | SAMN15294480 | GCA_019685775.1 | PC | Public | 2018 | China | Clinical | 97.8 | 100 | 0.27 | 99.9838 | 5179216 | 45.14 | 38 | 812153 |
| SAMN15294481-GCA_019685735.1 | ICDC-VP01811 | SAMN15294481 | GCA_019685735.1 | PC | Public | 2018 | China | Clinical | 98 | 100 | 0 | 99.9919 | 5077290 | 45.28 | 50 | 542656 |
| SAMN15294482-GCA_019685715.1 | ICDC-VP01812 | SAMN15294482 | GCA_019685715.1 | PC | Public | 2018 | China | Clinical | 97.6 | 100 | 0 | 99.9682 | 5142165 | 45.27 | 43 | 580817 |
| SAMN15294483-GCA_019685695.1 | ICDC-VP01813 | SAMN15294483 | GCA_019685695.1 | PC | Public | 2018 | China | Clinical | 97.7 | 100 | 0 | 99.9901 | 5071131 | 45.27 | 50 | 505463 |
| SAMN15294484-GCA_019685675.1 | ICDC-VP01814 | SAMN15294484 | GCA_019685675.1 | PC | Public | 2018 | China | Clinical | 94.6 | 100 | 0 | 99.9708 | 5146087 | 45.27 | 41 | 580818 |
| SAMN15294485-GCA_019685635.1 | ICDC-VP01815 | SAMN15294485 | GCA_019685635.1 | PC | Public | 2018 | China | Clinical | 97.6 | 100 | 0 | 99.9708 | 5144984 | 45.27 | 40 | 580817 |
| SAMN15294486-GCA_019685615.1 | ICDC-VP01816 | SAMN15294486 | GCA_019685615.1 | PC | Public | 2018 | China | Clinical | 97.6 | 100 | 0 | 99.9691 | 5143700 | 45.27 | 44 | 580442 |
| SAMN15331473-SRR12065278 | HN1 | SAMN15331473 | SRR12065278 | NonPC | Public | 2017 | China | Env | 92.5 | 100 | 0.54 | 98.3169 | 5356034 | 45.28 | 4 | 3277891 |
| SAMN15408096-GCF_013391065.1 | 3.394-17 | SAMN15408096 | GCF_013391065.1 | NonPC | Public | 2016 | Peru | Env | 86.7 | 98.29 | 0.68 | 98.1923 | 5139857 | 45.09 | 619 | 12995 |
| SAMN15408103-GCF_013390705.1 | 3.399-17 | SAMN15408103 | GCF_013390705.1 | NonPC | Public | 2016 | Peru | Env | 88.2 | 97.96 | 0.45 | 98.2709 | 5230974 | 45.06 | 442 | 20186 |
| SAMN15428938-GCF_016880155.1 | 404-00 | SAMN15428938 | GCF_016880155.1 | PC | Public | 2000 | Peru | Clinical | 98.3 | 100 | 0 | 99.9648 | 5154819 | 45.27 | 71 | 250941 |
| SAMN15428940-GCF_016880045.1 | 325-00 | SAMN15428940 | GCF_016880045.1 | PC | Public | 2000 | Peru | Clinical | 98.4 | 100 | 0 | 99.991 | 5171428 | 45.12 | 59 | 288853 |
| SAMN15428941-GCF_016880075.1 | 403-00 | SAMN15428941 | GCF_016880075.1 | PC | Public | 2000 | Peru | Clinical | 98.4 | 100 | 0.07 | 99.9903 | 5097932 | 45.27 | 69 | 288853 |
| SAMN15428942-GCF_016880085.1 | 327-00 | SAMN15428942 | GCF_016880085.1 | PC | Public | 2000 | Peru | Clinical | 98.4 | 100 | 0 | 99.9874 | 5171777 | 45.11 | 53 | 312582 |
| SAMN15428943-GCF_016880035.1 | 1027-00 | SAMN15428943 | GCF_016880035.1 | PC | Public | 2000 | Peru | Clinical | 98.6 | 100 | 0 | 99.9894 | 5195821 | 45.17 | 123 | 124878 |
| SAMN15428944-GCF_016880115.1 | 2434-00 | SAMN15428944 | GCF_016880115.1 | PC | Public | 2000 | Peru | Clinical | 98.8 | 100 | 0.14 | 99.9733 | 5244838 | 45.1 | 80 | 428063 |
| SAMN15428945-GCF_016880015.1 | 2568-00 | SAMN15428945 | GCF_016880015.1 | PC | Public | 2000 | Peru | Clinical | 98.8 | 100 | 0 | 99.9849 | 5127271 | 45.31 | 74 | 347141 |
| SAMN15428946-GCF_016879975.1 | 242-02 | SAMN15428946 | GCF_016879975.1 | NonPC | Public | 2002 | Peru | Clinical | 92.9 | 99.93 | 0 | 98.4344 | 5137014 | 45.22 | 47 | 329705 |
| SAMN15428947-GCF_016879995.1 | 1286-06 | SAMN15428947 | GCF_016879995.1 | NonPC | Public | 2006 | Peru | Clinical | 92.9 | 100 | 0 | 98.3119 | 5118185 | 45.28 | 66 | 522788 |
| SAMN15428948-GCF_016878735.1 | 1259-07 | SAMN15428948 | GCF_016878735.1 | PC | Public | 2007 | Peru | Clinical | 98.2 | 100 | 0 | 99.9811 | 5082137 | 45.3 | 42 | 312466 |
| SAMN15428949-GCF_016879955.1 | 1260-07 | SAMN15428949 | GCF_016879955.1 | PC | Public | 2007 | Peru | Clinical | 98.2 | 100 | 0 | 99.9794 | 5159908 | 45.13 | 67 | 271828 |
| SAMN15428950-GCF_016878715.1 | 1254-07 | SAMN15428950 | GCF_016878715.1 | PC | Public | 2007 | Peru | Clinical | 98.5 | 100 | 0 | 99.9794 | 5110095 | 45.32 | 70 | 187927 |
| SAMN15428951-GCF_016878665.1 | 1261-07 | SAMN15428951 | GCF_016878665.1 | NonPC | Public | 2007 | Peru | Clinical | 92.2 | 100 | 0.46 | 98.3484 | 5024846 | 45.24 | 50 | 470870 |
| SAMN15428952-GCF_016878675.1 | 243-09 | SAMN15428952 | GCF_016878675.1 | PC | Public | 2009 | Peru | Clinical | 98.5 | 100 | 0 | 99.9811 | 5175993 | 45.13 | 72 | 288855 |
| SAMN15428953-GCF_016878655.1 | 225-09 | SAMN15428953 | GCF_016878655.1 | NonPC | Public | 2009 | Peru | Clinical | 92.9 | 100 | 0 | 98.3929 | 5132236 | 45.21 | 49 | 329767 |
| SAMN15428954-GCF_016878635.1 | 260-09 | SAMN15428954 | GCF_016878635.1 | NonPC | Public | 2009 | Peru | Clinical | 92.9 | 100 | 0 | 98.4085 | 5127365 | 45.22 | 41 | 400468 |
| SAMN15428955-GCF_016878615.1 | 223-09 | SAMN15428955 | GCF_016878615.1 | NonPC | Public | 2009 | Peru | Clinical | 92.9 | 100 | 0 | 98.3857 | 5127305 | 45.22 | 48 | 312776 |
| SAMN15428956-GCF_016878595.1 | 227-09 | SAMN15428956 | GCF_016878595.1 | NonPC | Public | 2009 | Peru | Clinical | 92.9 | 100 | 0.01 | 98.3659 | 5220433 | 45.24 | 53 | 257547 |
| SAMN15428957-GCF_016878555.1 | 238-09 | SAMN15428957 | GCF_016878555.1 | NonPC | Public | 2009 | Peru | Clinical | 92.9 | 100 | 0 | 98.4283 | 5130724 | 45.22 | 42 | 329707 |
| SAMN15428958-GCF_016878565.1 | 221-09 | SAMN15428958 | GCF_016878565.1 | NonPC | Public | 2009 | Peru | Clinical | 93 | 100 | 0 | 98.3792 | 5180661 | 45.22 | 46 | 284797 |
| SAMN15428959-GCF_016878535.1 | 222-09 | SAMN15428959 | GCF_016878535.1 | NonPC | Public | 2009 | Peru | Clinical | 92.9 | 100 | 0.01 | 98.3819 | 5221305 | 45.25 | 52 | 312615 |
| SAMN15428960-GCF_016878515.1 | 242-09 | SAMN15428960 | GCF_016878515.1 | NonPC | Public | 2009 | Peru | Clinical | 92.9 | 100 | 0 | 98.4027 | 5128566 | 45.22 | 47 | 366680 |
| SAMN15428961-GCF_016878495.1 | 576-09 | SAMN15428961 | GCF_016878495.1 | NonPC | Public | 2009 | Peru | Clinical | 93.1 | 100 | 0 | 98.3856 | 5158626 | 45.23 | 56 | 360859 |
| SAMN15428962-GCF_016878475.1 | 1214-09 | SAMN15428962 | GCF_016878475.1 | NonPC | Public | 2009 | Peru | Clinical | 92.4 | 100 | 0.46 | 98.3596 | 5047278 | 45.25 | 52 | 568848 |
| SAMN15428963-GCF_016878435.1 | 203-10 | SAMN15428963 | GCF_016878435.1 | PC | Public | 2010 | Peru | Clinical | 98.5 | 100 | 0 | 99.994 | 5099091 | 45.29 | 59 | 312374 |
| SAMN15428964-GCF_016878455.1 | 091-10 | SAMN15428964 | GCF_016878455.1 | PC | Public | 2010 | Peru | Clinical | 98.4 | 100 | 0 | 99.9848 | 5175162 | 45.11 | 60 | 312452 |
| SAMN15428965-GCF_016878395.1 | 304-10 | SAMN15428965 | GCF_016878395.1 | PC | Public | 2010 | Peru | Clinical | 98.4 | 100 | 0 | 99.9799 | 5165450 | 45.12 | 52 | 312374 |
| SAMN15428966-GCF_016878405.1 | 293-10 | SAMN15428966 | GCF_016878405.1 | PC | Public | 2010 | Peru | Clinical | 98.5 | 100 | 0 | 99.9839 | 5099560 | 45.29 | 60 | 289055 |
| SAMN15428967-GCF_016878375.1 | 202-10 | SAMN15428967 | GCF_016878375.1 | PC | Public | 2010 | Peru | Clinical | 98.4 | 100 | 0 | 99.992 | 5094730 | 45.29 | 60 | 314066 |

| | | | | | | | | | | | | | | | | |
|---|---|---|---|---|---|---|---|---|---|---|---|---|---|---|---|---|
| SAMN15428968-GCF_016878355.1 | 092-10 | SAMN15428968 | GCF_016878355.1 | PC | Public | 2010 | Peru | Clinical | 98.5 | 100 | 0 | 99.9856 | 5168389 | 45.13 | 65 | 288857 |
| SAMN15428969-GCF_016878325.1 | 361-10 | SAMN15428969 | GCF_016878325.1 | PC | Public | 2010 | Peru | Clinical | 98.4 | 100 | 0.12 | 99.9939 | 5096580 | 45.28 | 61 | 267685 |
| SAMN15428970-GCF_016878315.1 | 454-10 | SAMN15428970 | GCF_016878315.1 | PC | Public | 2010 | Peru | Clinical | 98.5 | 100 | 0 | 99.9869 | 5168836 | 45.13 | 56 | 294788 |
| SAMN15428971-GCF_016878295.1 | 119-10 | SAMN15428971 | GCF_016878295.1 | PC | Public | 2010 | Peru | Clinical | 98.5 | 100 | 0 | 99.9898 | 5169782 | 45.13 | 57 | 394720 |
| SAMN15428972-GCF_016878275.1 | 223-10 | SAMN15428972 | GCF_016878275.1 | PC | Public | 2010 | Peru | Clinical | 98.3 | 99.68 | 0.05 | 99.9884 | 5173591 | 45.11 | 121 | 340010 |
| SAMN15428973-GCF_016878255.1 | 107-10 | SAMN15428973 | GCF_016878255.1 | NonPC | Public | 2010 | Peru | Clinical | 91.4 | 100 | 0.26 | 98.2968 | 5287380 | 45.25 | 51 | 420329 |
| SAMN15428974-GCF_016878215.1 | 1218-11 | SAMN15428974 | GCF_016878215.1 | PC | Public | 2011 | Peru | Clinical | 98.7 | 100 | 0 | 99.9822 | 5206821 | 45.13 | 90 | 221913 |
| SAMN15428975-GCF_016878195.1 | 326-11 | SAMN15428975 | GCF_016878195.1 | NonPC | Public | 2011 | Peru | Clinical | 92.9 | 100 | 0.05 | 98.3985 | 5128881 | 45.22 | 50 | 329811 |
| SAMN15428976-GCF_016878235.1 | 1021-11 | SAMN15428976 | GCF_016878235.1 | NonPC | Public | 2011 | Peru | Clinical | 93.1 | 100 | 0 | 98.377 | 5152045 | 45.23 | 61 | 330910 |
| SAMN15428977-GCF_016878175.1 | 181-11 | SAMN15428977 | GCF_016878175.1 | NonPC | Public | 2011 | Peru | Clinical | 90.8 | 99.81 | 0.36 | 98.3777 | 4947131 | 45.4 | 165 | 96309 |
| SAMN15428978-GCF_016878095.1 | 005-13 | SAMN15428978 | GCF_016878095.1 | NonPC | Public | 2013 | Peru | Clinical | 89.3 | 100 | 0 | 98.1868 | 5065587 | 45.19 | 71 | 305119 |
| SAMN15428979-GCF_016878155.1 | 582-14 | SAMN15428979 | GCF_016878155.1 | NonPC | Public | 2014 | Peru | Clinical | 91 | 100 | 0.16 | 98.3821 | 5527246 | 45.15 | 60 | 243860 |
| SAMN15428980-GCF_016878135.1 | 1835-14 | SAMN15428980 | GCF_016878135.1 | NonPC | Public | 2014 | Peru | Clinical | 91.2 | 100 | 0.07 | 98.3656 | 5172706 | 45.22 | 61 | 202517 |
| SAMN15428981-GCF_016878105.1 | 024-14 | SAMN15428981 | GCF_016878105.1 | NonPC | Public | 2014 | Peru | Clinical | 92.8 | 100 | 0.23 | 98.3995 | 5132117 | 45.22 | 40 | 488541 |
| SAMN15428982-GCF_016878075.1 | 277-15 | SAMN15428982 | GCF_016878075.1 | PC | Public | 2015 | Peru | Clinical | 98.4 | 100 | 0.01 | 99.9784 | 5260434 | 45.15 | 56 | 394629 |
| SAMN15428983-GCF_016878055.1 | G6 | SAMN15428983 | GCF_016878055.1 | PC | Public | 2015 | Peru | Clinical | 98.8 | 100 | 0.01 | 99.9824 | 5333373 | 45.13 | 141 | 428513 |
| SAMN15428984-GCF_016877995.1 | H11 | SAMN15428984 | GCF_016877995.1 | PC | Public | 2015 | Peru | Clinical | 98.8 | 100 | 0.09 | 99.9828 | 5271308 | 45.29 | 69 | 427965 |
| SAMN15428985-GCF_016877975.1 | 369-15 | SAMN15428985 | GCF_016877975.1 | NonPC | Public | 2015 | Peru | Env | 91.8 | 100 | 0 | 98.2511 | 5228452 | 45.17 | 54 | 390473 |
| SAMN15428986-GCF_016878015.1 | G10 | SAMN15428986 | GCF_016878015.1 | NonPC | Public | 2015 | Peru | Clinical | 91.6 | 100 | 0 | 98.2908 | 5240560 | 45.22 | 51 | 549002 |
| SAMN15428987-GCF_016878035.1 | H6 | SAMN15428987 | GCF_016878035.1 | NonPC | Public | 2015 | Peru | Clinical | 91.6 | 100 | 0 | 98.2821 | 5243568 | 45.22 | 51 | 562531 |
| SAMN15428988-GCF_016877955.1 | 416-15 | SAMN15428988 | GCF_016877955.1 | NonPC | Public | 2015 | Peru | Clinical | 91.6 | 100 | 0 | 98.2711 | 5244373 | 45.22 | 69 | 232353 |
| SAMN15428989-GCF_016877895.1 | 166-15 | SAMN15428989 | GCF_016877895.1 | NonPC | Public | 2015 | Peru | Clinical | 91.4 | 99.86 | 0 | 98.307 | 5180380 | 45.22 | 63 | 461592 |
| SAMN15428990-GCF_016877915.1 | H5 | SAMN15428990 | GCF_016877915.1 | NonPC | Public | 2015 | Peru | Clinical | 90.9 | 100 | 0.88 | 98.2764 | 5352082 | 45.11 | 65 | 558313 |
| SAMN15428991-GCF_016877935.1 | 368-15 | SAMN15428991 | GCF_016877935.1 | NonPC | Public | 2015 | Peru | Env | 91.2 | 100 | 0.68 | 98.3883 | 4967968 | 45.39 | 77 | 453928 |
| SAMN15428992-GCF_016877875.1 | 373-15 | SAMN15428992 | GCF_016877875.1 | NonPC | Public | 2015 | Peru | Env | 91.3 | 100 | 0.27 | 98.403 | 4973507 | 45.39 | 65 | 403437 |
| SAMN15428993-GCF_016877855.1 | 371-15 | SAMN15428993 | GCF_016877855.1 | NonPC | Public | 2015 | Peru | Env | 90.2 | 99.93 | 0.41 | 98.3964 | 4969992 | 45.39 | 72 | 454217 |
| SAMN15428994-GCF_016877835.1 | 3370-15 | SAMN15428994 | GCF_016877835.1 | NonPC | Public | 2015 | Peru | Env | 91.4 | 100 | 0.27 | 98.3998 | 4994848 | 45.39 | 59 | 540500 |
| SAMN15428995-GCF_016877795.1 | 306-15 | SAMN15428995 | GCF_016877795.1 | NonPC | Public | 2015 | Peru | Env | 90.3 | 100 | 0.35 | 98.2418 | 5226088 | 45.18 | 64 | 318362 |
| SAMN15428996-GCF_016877805.1 | 3264-15 | SAMN15428996 | GCF_016877805.1 | NonPC | Public | 2015 | Peru | Env | 90 | 100 | 0 | 98.3663 | 5177911 | 45.26 | 89 | 246190 |
| SAMN15428997-GCF_016877775.1 | 302-15 | SAMN15428997 | GCF_016877775.1 | NonPC | Public | 2015 | Peru | Clinical | 88.6 | 100 | 0.38 | 98.1998 | 5267551 | 45.13 | 65 | 302895 |
| SAMN15428998-GCF_016877755.1 | H2 | SAMN15428998 | GCF_016877755.1 | NonPC | Public | 2015 | Peru | Clinical | 88.8 | 100 | 0.38 | 98.2211 | 5274755 | 45.13 | 61 | 576952 |
| SAMN15428999-GCF_016877735.1 | 305-15 | SAMN15428999 | GCF_016877735.1 | NonPC | Public | 2015 | Peru | Clinical | 89.5 | 100 | 0.05 | 98.3238 | 5047992 | 45.32 | 43 | 443007 |
| SAMN15429000-GCF_016877695.1 | H8 | SAMN15429000 | GCF_016877695.1 | NonPC | Public | 2015 | Peru | Clinical | 89.7 | 100 | 0.05 | 98.3393 | 5077159 | 45.32 | 57 | 864892 |
| SAMN15429001-GCF_016877715.1 | 167-16 | SAMN15429001 | GCF_016877715.1 | PC | Public | 2016 | Peru | Clinical | 98.4 | 100 | 0.09 | 99.9879 | 5238650 | 45.26 | 69 | 294787 |
| SAMN15429002-GCF_016877675.1 | 165-16 | SAMN15429002 | GCF_016877675.1 | PC | Public | 2016 | Peru | Clinical | 98.4 | 100 | 0 | 99.9857 | 5115284 | 45.27 | 64 | 312423 |
| SAMN15429003-GCF_016877655.1 | G8 | SAMN15429003 | GCF_016877655.1 | PC | Public | 2016 | Peru | Clinical | 98.8 | 100 | 0.09 | 99.9829 | 5393115 | 45.13 | 268 | 402146 |
| SAMN15429004-GCF_016877605.1 | H12 | SAMN15429004 | GCF_016877605.1 | PC | Public | 2016 | Peru | Clinical | 98.7 | 100 | 0 | 99.9812 | 5160969 | 45.27 | 103 | 402206 |
| SAMN15429005-GCF_016877595.1 | 252-16 | SAMN15429005 | GCF_016877595.1 | NonPC | Public | 2016 | Peru | Clinical | 91.2 | 100 | 0 | 98.3206 | 5188239 | 45.24 | 174 | 69879 |
| SAMN15429006-GCF_016877615.1 | 210-16 | SAMN15429006 | GCF_016877615.1 | NonPC | Public | 2016 | Peru | Clinical | 91.6 | 99.86 | 0 | 98.273 | 5222501 | 45.17 | 129 | 205212 |
| SAMN15429007-GCF_016877575.1 | 281-16 | SAMN15429007 | GCF_016877575.1 | NonPC | Public | 2016 | Peru | Clinical | 90.7 | 99.73 | 0.05 | 98.3489 | 5073824 | 45.33 | 160 | 106246 |
| SAMN15429008-GCF_016877545.1 | 275-16 | SAMN15429008 | GCF_016877545.1 | NonPC | Public | 2016 | Peru | Clinical | 91 | 100 | 0.05 | 98.3603 | 5090798 | 45.33 | 65 | 278821 |
| SAMN15429009-GCF_016877535.1 | 111-16 | SAMN15429009 | GCF_016877535.1 | NonPC | Public | 2016 | Peru | Clinical | 91.1 | 100 | 0.05 | 98.3192 | 5125093 | 45.31 | 74 | 287808 |
| SAMN15429010-GCF_016877515.1 | 3276-16 | SAMN15429010 | GCF_016877515.1 | NonPC | Public | 2016 | Peru | Env | 91.2 | 100 | 0.05 | 98.3161 | 5124596 | 45.32 | 70 | 287808 |
| SAMN15429011-GCF_016877495.1 | INS3_40517 | SAMN15429011 | GCF_016877495.1 | NonPC | Public | 2016 | Peru | Env | 91.6 | 100 | 0 | 98.3754 | 5028215 | 45.3 | 46 | 451106 |
| SAMN15429012-GCF_016877475.1 | INSP00112 | SAMN15429012 | GCF_016877475.1 | NonPC | Public | 2016 | Peru | Env | 91.1 | 100 | 0.38 | 98.3096 | 5076153 | 45.31 | 124 | 222944 |
| SAMN15429013-GCF_016877455.1 | 271-16 | SAMN15429013 | GCF_016877455.1 | NonPC | Public | 2016 | Peru | Clinical | 90.6 | 99.77 | 0.5 | 98.3002 | 5090819 | 45.24 | 125 | 110377 |
| SAMN15429014-GCF_016877415.1 | H1 | SAMN15429014 | GCF_016877415.1 | NonPC | Public | 2016 | Peru | Clinical | 92.9 | 100 | 0 | 98.3424 | 5119600 | 45.27 | 64 | 523105 |
| SAMN15429015-GCF_016877435.1 | 260-16 | SAMN15429015 | GCF_016877435.1 | NonPC | Public | 2016 | Peru | Clinical | 88.4 | 100 | 0.3 | 98.2497 | 5248981 | 45.12 | 84 | 430116 |

| | | | | | | | | | | | | | | | | |
|---|---|---|---|---|---|---|---|---|---|---|---|---|---|---|---|---|
| SAMN15429016-GCF_016877395.1 | INS3T395A17 | SAMN15429016 | GCF_016877395.1 | NonPC | Public | 2016 | Peru | Env | 88.4 | 100 | 0.3 | 98.2238 | 5168711 | 45.17 | 62 | 388661 |
| SAMN15429017-GCF_016877375.1 | 256-16 | SAMN15429017 | GCF_016877375.1 | NonPC | Public | 2016 | Peru | Clinical | 89.7 | 98.77 | 0 | 98.3329 | 5134893 | 45.26 | 105 | 248149 |
| SAMN15429018-GCF_016877355.1 | 254-16 | SAMN15429018 | GCF_016877355.1 | NonPC | Public | 2016 | Peru | Clinical | 91.5 | 100 | 0.09 | 98.2644 | 5136129 | 45.41 | 59 | 316312 |
| SAMN15429019-GCF_016877335.1 | INS3T403A17 | SAMN15429019 | GCF_016877335.1 | NonPC | Public | 2016 | Peru | Env | 88.9 | 99.36 | 0.53 | 98.3044 | 4944350 | 45.37 | 172 | 56034 |
| SAMN15429020-GCF_016877295.1 | INSP0011 | SAMN15429020 | GCF_016877295.1 | NonPC | Public | 2016 | Peru | Env | 90.9 | 100 | 0 | 98.3872 | 5179172 | 45.25 | 59 | 405483 |
| SAMN15429021-GCF_016877315.1 | INSP0021 | SAMN15429021 | GCF_016877315.1 | NonPC | Public | 2016 | Peru | Env | 88 | 100 | 0.3 | 98.2784 | 4993994 | 45.28 | 118 | 346264 |
| SAMN15429022-GCF_016877265.1 | INSP00222 | SAMN15429022 | GCF_016877265.1 | NonPC | Public | 2016 | Peru | Env | 89.6 | 100 | 0.05 | 98.3211 | 4975147 | 45.35 | 46 | 582355 |
| SAMN15429023-GCF_016877235.1 | INSP00223 | SAMN15429023 | GCF_016877235.1 | NonPC | Public | 2016 | Peru | Env | 91 | 100 | 0 | 98.3811 | 5204121 | 45.24 | 64 | 313863 |
| SAMN15429024-GCF_016877255.1 | INSP00224 | SAMN15429024 | GCF_016877255.1 | NonPC | Public | 2016 | Peru | Env | 90.9 | 100 | 0 | 98.3639 | 5177689 | 45.25 | 79 | 313863 |
| SAMN15429025-GCF_016877215.1 | INSP00556 | SAMN15429025 | GCF_016877215.1 | NonPC | Public | 2016 | Peru | Env | 91.2 | 100 | 0.27 | 98.3221 | 5001636 | 45.36 | 48 | 534516 |
| SAMN15429026-GCF_016877175.1 | INSP00559 | SAMN15429026 | GCF_016877175.1 | NonPC | Public | 2016 | Peru | Env | 88 | 100 | 0.39 | 98.2432 | 4968464 | 45.29 | 97 | 275061 |
| SAMN15429027-GCF_016877195.1 | INSP00561 | SAMN15429027 | GCF_016877195.1 | NonPC | Public | 2016 | Peru | Env | 87.9 | 100 | 0.39 | 98.2544 | 4962294 | 45.29 | 99 | 427490 |
| SAMN15429028-GCF_016877155.1 | INST00671 | SAMN15429028 | GCF_016877155.1 | NonPC | Public | 2016 | Peru | Env | 91.2 | 100 | 0 | 98.2933 | 5029361 | 45.36 | 72 | 328843 |
| SAMN15429029-GCF_016877135.1 | INST008109 | SAMN15429029 | GCF_016877135.1 | NonPC | Public | 2016 | Peru | Env | 91 | 100 | 0.14 | 98.3554 | 4937067 | 45.36 | 50 | 475509 |
| SAMN15429030-GCF_016877115.1 | INST008110 | SAMN15429030 | GCF_016877115.1 | NonPC | Public | 2016 | Peru | Env | 89.5 | 100 | 0 | 98.2807 | 5016568 | 45.3 | 59 | 346810 |
| SAMN15429031-GCF_016877055.1 | INST00896 | SAMN15429031 | GCF_016877055.1 | NonPC | Public | 2016 | Peru | Env | 90.9 | 100 | 0.32 | 98.2787 | 4979737 | 45.33 | 60 | 524819 |
| SAMN15429032-GCF_016877095.1 | INSP0026 | SAMN15429032 | GCF_016877095.1 | NonPC | Public | 2017 | Peru | Env | 91.6 | 100 | 0.27 | 98.4132 | 4998754 | 45.36 | 34 | 525885 |
| SAMN15429033-GCF_016877035.1 | INSCH0151 | SAMN15429033 | GCF_016877035.1 | NonPC | Public | 2017 | Peru | Env | 90.8 | 100 | 0.36 | 98.3393 | 4952060 | 45.39 | 53 | 345297 |
| SAMN15429034-GCF_016877075.1 | INST022117 | SAMN15429034 | GCF_016877075.1 | NonPC | Public | 2017 | Peru | Env | 89.4 | 100 | 0 | 98.3406 | 4933528 | 45.36 | 56 | 384751 |
| SAMN15429035-GCF_016877015.1 | INS3_42617 | SAMN15429035 | GCF_016877015.1 | NonPC | Public | 2017 | Peru | Env | 91 | 100 | 0.05 | 98.3529 | 4954179 | 45.35 | 51 | 470258 |
| SAMN15429036-GCF_016876975.1 | INS3_41617 | SAMN15429036 | GCF_016876975.1 | NonPC | Public | 2017 | Peru | Env | 91.5 | 100 | 0 | 98.3766 | 5051928 | 45.29 | 39 | 451101 |
| SAMN15429037-GCF_016876935.1 | INST0262 | SAMN15429037 | GCF_016876935.1 | NonPC | Public | 2017 | Peru | Env | 90.6 | 99.73 | 0.34 | 98.3475 | 4988414 | 45.32 | 56 | 250298 |
| SAMN15429038-GCF_016876945.1 | INSP032117 | SAMN15429038 | GCF_016876945.1 | NonPC | Public | 2017 | Peru | Env | 90.3 | 99.73 | 0.33 | 98.3134 | 5050908 | 45.3 | 77 | 179042 |
| SAMN15429039-GCF_016876985.1 | INSP032717 | SAMN15429039 | GCF_016876985.1 | NonPC | Public | 2017 | Peru | Env | 91.1 | 100 | 0.05 | 98.3014 | 5103416 | 45.31 | 54 | 371957 |
| SAMN15429040-GCF_016876915.1 | INSP0051 | SAMN15429040 | GCF_016876915.1 | NonPC | Public | 2017 | Peru | Env | 88 | 100 | 0.39 | 98.2409 | 5006125 | 45.24 | 86 | 444735 |
| SAMN15429041-GCF_016876885.1 | INSCH0111 | SAMN15429041 | GCF_016876885.1 | NonPC | Public | 2017 | Peru | Env | 91.3 | 100 | 0.05 | 98.3507 | 5089989 | 45.34 | 44 | 486568 |
| SAMN15429042-GCF_016876855.1 | INSCH0073 | SAMN15429042 | GCF_016876855.1 | NonPC | Public | 2017 | Peru | Env | 88.2 | 100 | 0.4 | 98.2531 | 5040171 | 45.25 | 95 | 361248 |
| SAMN15429043-GCF_016876815.1 | INSCH0074 | SAMN15429043 | GCF_016876815.1 | NonPC | Public | 2017 | Peru | Env | 88 | 100 | 0.47 | 98.1992 | 5045670 | 45.19 | 90 | 365014 |
| SAMN15429044-GCF_016876875.1 | INSCH013717 | SAMN15429044 | GCF_016876875.1 | NonPC | Public | 2017 | Peru | Env | 88.8 | 100 | 0.27 | 98.2485 | 4985014 | 45.32 | 54 | 381404 |
| SAMN15429045-GCF_016876835.1 | INSCH0143 | SAMN15429045 | GCF_016876835.1 | NonPC | Public | 2017 | Peru | Env | 88.9 | 100 | 0.05 | 98.2503 | 5002098 | 45.25 | 60 | 552757 |
| SAMN15429046-GCF_016876795.1 | INSCH0153 | SAMN15429046 | GCF_016876795.1 | NonPC | Public | 2017 | Peru | Env | 91.4 | 100 | 0.36 | 98.3561 | 5015552 | 45.35 | 95 | 475022 |
| SAMN15429047-GCF_016876775.1 | INSCH0154 | SAMN15429047 | GCF_016876775.1 | NonPC | Public | 2017 | Peru | Env | 90.7 | 100 | 0.05 | 98.3359 | 5014018 | 45.32 | 87 | 303765 |
| SAMN15429048-GCF_016876715.1 | INSCH0155 | SAMN15429048 | GCF_016876715.1 | NonPC | Public | 2017 | Peru | Env | 89.8 | 100 | 0.05 | 98.3481 | 5030762 | 45.35 | 56 | 392097 |
| SAMN15429049-GCF_016876735.1 | INSCH0156 | SAMN15429049 | GCF_016876735.1 | NonPC | Public | 2017 | Peru | Env | 90.4 | 100 | 0 | 98.2599 | 5084185 | 45.27 | 70 | 410531 |
| SAMN15429050-GCF_016876745.1 | INSP0022317 | SAMN15429050 | GCF_016876745.1 | NonPC | Public | 2017 | Peru | Env | 89.9 | 100 | 0.27 | 98.3174 | 4941730 | 45.46 | 62 | 225148 |
| SAMN15429051-GCF_016876695.1 | INSP0034 | SAMN15429051 | GCF_016876695.1 | NonPC | Public | 2017 | Peru | Env | 91 | 100 | 0.27 | 98.3762 | 5000062 | 45.33 | 66 | 402751 |
| SAMN15429052-GCF_016876675.1 | INSP0035 | SAMN15429052 | GCF_016876675.1 | NonPC | Public | 2017 | Peru | Env | 90.4 | 100 | 0.05 | 98.2582 | 5114974 | 45.22 | 65 | 374542 |
| SAMN15429053-GCF_016876645.1 | INSP0041017 | SAMN15429053 | GCF_016876645.1 | NonPC | Public | 2017 | Peru | Env | 91.1 | 100 | 0.09 | 98.3674 | 5119696 | 45.19 | 85 | 352737 |
| SAMN15429054-GCF_016876635.1 | INSP0047 | SAMN15429054 | GCF_016876635.1 | NonPC | Public | 2017 | Peru | Env | 91.1 | 99.91 | 0.54 | 98.3356 | 4994566 | 45.38 | 68 | 284891 |
| SAMN15429055-GCF_016876615.1 | INSP0062 | SAMN15429055 | GCF_016876615.1 | NonPC | Public | 2017 | Peru | Env | 88.5 | 100 | 0.3 | 98.2474 | 5118450 | 45.24 | 66 | 348182 |
| SAMN15429056-GCF_016876595.1 | INSP0066 | SAMN15429056 | GCF_016876595.1 | NonPC | Public | 2017 | Peru | Env | 91 | 100 | 0.09 | 98.252 | 4922688 | 45.41 | 74 | 316335 |
| SAMN15429057-GCF_016876575.1 | INSP032217 | SAMN15429057 | GCF_016876575.1 | NonPC | Public | 2017 | Peru | Env | 91.3 | 100 | 0.03 | 98.3237 | 5042672 | 45.26 | 68 | 374440 |
| SAMN15429058-GCF_016876535.1 | INSP033117 | SAMN15429058 | GCF_016876535.1 | NonPC | Public | 2017 | Peru | Env | 91.2 | 100 | 0 | 98.3482 | 5012282 | 45.36 | 82 | 318093 |
| SAMN15429059-GCF_016876515.1 | INST0181 | SAMN15429059 | GCF_016876515.1 | NonPC | Public | 2017 | Peru | Env | 91.5 | 100 | 0.05 | 98.3268 | 5000393 | 45.39 | 54 | 516361 |
| SAMN15429060-GCF_016876545.1 | INST0184 | SAMN15429060 | GCF_016876545.1 | NonPC | Public | 2017 | Peru | Env | 91.7 | 100 | 0.04 | 98.4127 | 5003376 | 45.37 | 62 | 372119 |
| SAMN15429061-GCF_016876495.1 | INST0192 | SAMN15429061 | GCF_016876495.1 | NonPC | Public | 2017 | Peru | Env | 91.7 | 100 | 0.05 | 98.2286 | 5022310 | 45.36 | 85 | 374160 |
| SAMN15429062-GCF_016876475.1 | INST0205 | SAMN15429062 | GCF_016876475.1 | NonPC | Public | 2017 | Peru | Env | 88.7 | 100 | 0.39 | 98.263 | 5122618 | 45.21 | 70 | 492753 |
| SAMN15429063-GCF_016876435.1 | INST0211 | SAMN15429063 | GCF_016876435.1 | NonPC | Public | 2017 | Peru | Env | 89.5 | 100 | 0.36 | 98.2811 | 5174864 | 45.33 | 56 | 324064 |

| | | | | | | | | | | | | | | | | |
|---|---|---|---|---|---|---|---|---|---|---|---|---|---|---|---|---|
| SAMN15429064-GCF_016876415.1 | INST0212 | SAMN15429064 | GCF_016876415.1 | NonPC | Public | 2017 | Peru | Env | 88 | 100 | 0.36 | 98.2871 | 5028241 | 45.32 | 65 | 380734 |
| SAMN15429065-GCF_016876455.1 | INST022317 | SAMN15429065 | GCF_016876455.1 | NonPC | Public | 2017 | Peru | Env | 89.4 | 100 | 0.23 | 98.2808 | 4928594 | 45.46 | 50 | 329260 |
| SAMN15429066-GCF_016876375.1 | INST0225 | SAMN15429066 | GCF_016876375.1 | NonPC | Public | 2017 | Peru | Env | 91.3 | 100 | 0 | 98.3718 | 5046964 | 45.34 | 63 | 473838 |
| SAMN15429067-GCF_016876355.1 | INST0228 | SAMN15429067 | GCF_016876355.1 | NonPC | Public | 2017 | Peru | Env | 89.5 | 100 | 0 | 98.2529 | 4979812 | 45.35 | 68 | 275082 |
| SAMN15429068-GCF_016876395.1 | INST0231 | SAMN15429068 | GCF_016876395.1 | NonPC | Public | 2017 | Peru | Env | 91.1 | 100 | 0 | 98.2588 | 5052801 | 45.34 | 65 | 498719 |
| SAMN15429069-GCF_016876335.1 | INST0232 | SAMN15429069 | GCF_016876335.1 | NonPC | Public | 2017 | Peru | Env | 91.2 | 99.93 | 0 | 98.3335 | 4980503 | 45.37 | 59 | 255991 |
| SAMN15429070-GCF_016876315.1 | INST0233 | SAMN15429070 | GCF_016876315.1 | NonPC | Public | 2017 | Peru | Env | 91.1 | 100 | 0 | 98.2807 | 4981769 | 45.36 | 68 | 313773 |
| SAMN15429071-GCF_016876285.1 | INST0241 | SAMN15429071 | GCF_016876285.1 | NonPC | Public | 2017 | Peru | Env | 88 | 100 | 0.33 | 98.3571 | 4932294 | 45.36 | 84 | 440462 |
| SAMN15429072-GCF_016876275.1 | INST0253 | SAMN15429072 | GCF_016876275.1 | NonPC | Public | 2017 | Peru | Env | 91.2 | 100 | 0 | 98.3215 | 5000131 | 45.33 | 60 | 490486 |
| SAMN15429073-GCF_016876255.1 | INST027417 | SAMN15429073 | GCF_016876255.1 | NonPC | Public | 2017 | Peru | Env | 89.5 | 100 | 0.11 | 98.2582 | 4984165 | 45.34 | 59 | 429150 |
| SAMN15429074-GCF_016876235.1 | INST028217 | SAMN15429074 | GCF_016876235.1 | NonPC | Public | 2017 | Peru | Env | 89.2 | 100 | 0.19 | 98.2646 | 4939082 | 45.36 | 47 | 517198 |
| SAMN15429075-GCF_016876215.1 | VA-0007-17 | SAMN15429075 | GCF_016876215.1 | NonPC | Public | 2017 | Peru | Env | 87.6 | 100 | 0.36 | 98.3015 | 5014676 | 45.32 | 58 | 660150 |
| SAMN15429076-GCF_016876195.1 | VA-0569-17 | SAMN15429076 | GCF_016876195.1 | NonPC | Public | 2017 | Peru | Env | 89.5 | 100 | 0.05 | 98.1968 | 4945250 | 45.41 | 54 | 453626 |
| SAMN15429077-GCF_016876175.1 | VA-0662-17 | SAMN15429077 | GCF_016876175.1 | NonPC | Public | 2017 | Peru | Env | 91.7 | 99.73 | 0.41 | 98.1483 | 5456320 | 45.08 | 56 | 415557 |
| SAMN15429078-GCF_016876145.1 | VA-0693-17 | SAMN15429078 | GCF_016876145.1 | NonPC | Public | 2017 | Peru | Env | 88.3 | 100 | 0.47 | 98.1993 | 5083657 | 45.2 | 78 | 314081 |
| SAMN15455880-GCF_014057915.1 | VP42 | SAMN15455880 | GCF_014057915.1 | NonPC | Public | 2013 | China | Clinical | 93.1 | 100 | 0.05 | 98.3654 | 5053447 | 45.32 | 29 | 1239422 |
| SAMN15455881-GCF_014057925.1 | VP103 | SAMN15455881 | GCF_014057925.1 | NonPC | Public | 2015 | China | Clinical | 93 | 100 | 0.05 | 98.3724 | 5060597 | 45.3 | 44 | 424367 |
| SAMN15455882-GCF_014057895.1 | VP168 | SAMN15455882 | GCF_014057895.1 | NonPC | Public | 2016 | China | Clinical | 93 | 100 | 0.05 | 98.4053 | 5056981 | 45.31 | 47 | 418070 |
| SAMN15455883-GCF_014057865.1 | VP365 | SAMN15455883 | GCF_014057865.1 | NonPC | Public | 2013 | China | Clinical | 93 | 100 | 0.14 | 98.3465 | 5265334 | 45.25 | 45 | 484447 |
| SAMN15455884-GCF_014057875.1 | VP542 | SAMN15455884 | GCF_014057875.1 | NonPC | Public | 2014 | China | Clinical | 93 | 100 | 0.05 | 98.392 | 5055490 | 45.3 | 40 | 484322 |
| SAMN15455885-GCF_014057815.1 | VP595 | SAMN15455885 | GCF_014057815.1 | NonPC | Public | 2015 | China | Clinical | 93 | 100 | 0.05 | 98.359 | 5091235 | 45.28 | 38 | 463542 |
| SAMN15455886-GCF_014057775.1 | VP761 | SAMN15455886 | GCF_014057775.1 | NonPC | Public | 2016 | China | Clinical | 93 | 100 | 0.05 | 98.3726 | 5050845 | 45.3 | 38 | 484471 |
| SAMN15455887-GCF_014057825.1 | VP856 | SAMN15455887 | GCF_014057825.1 | NonPC | Public | 2017 | China | Clinical | 89.8 | 94.55 | 0.19 | 98.3711 | 4841955 | 45.39 | 13 | 513872 |
| SAMN15455888-GCF_014057765.1 | VP2502 | SAMN15455888 | GCF_014057765.1 | NonPC | Public | 2017 | China | Clinical | 93.1 | 100 | 0.6 | 98.3499 | 5073883 | 45.31 | 38 | 1239417 |
| SAMN15455889-GCF_014057805.1 | VP2571 | SAMN15455889 | GCF_014057805.1 | NonPC | Public | 2017 | China | Clinical | 93.1 | 100 | 0.19 | 98.358 | 5088045 | 45.29 | 35 | 463699 |
| SAMN15510356-GCA_019793345.1 | ISF-232-3 | SAMN15510356 | GCA_019793345.1 | NonPC | Public | 2016 | NA | Env | 88.6 | 100 | 0 | 98.0783 | 5103320 | 45.3 | 159 | 80238 |
| SAMN15510357-GCA_019793285.1 | ISF-238-3 | SAMN15510357 | GCA_019793285.1 | NonPC | Public | 2016 | NA | Env | 91.4 | 100 | 0.41 | 98.3564 | 5424961 | 45.3 | 84 | 195494 |
| SAMN15516036-SRR12206318 | C4 | SAMN15516036 | SRR12206318 | NonPC | Public | 2014 | China | Env | 93 | 100 | 0.05 | 98.3499 | 5060345 | 45.31 | 59 | 364045 |
| SAMN15594841-GCF_014396905.1 | VHT1_MFl | SAMN15594841 | GCF_014396905.1 | NonPC | Public | 2020 | USA | Env | 91.1 | 100 | 0.75 | 98.2533 | 5115539 | 45.25 | 57 | 807591 |
| SAMN15594842-GCF_014396885.1 | VHT2_MFl | SAMN15594842 | GCF_014396885.1 | NonPC | Public | 2020 | USA | Env | 89.2 | 100 | 0 | 98.258 | 5261907 | 45.13 | 271 | 531181 |
| SAMN15716808-GCA_015746425.1 | PNUSAV001225 | SAMN15716808 | GCA_015746425.1 | NonPC | Public | NA | USA | Clinical | 89.8 | 100 | 1.6 | 98.2521 | 5282341 | 45.22 | 53 | 423593 |
| SAMN15716834-GCA_015746345.1 | PNUSAV001229 | SAMN15716834 | GCA_015746345.1 | NonPC | Public | NA | USA | Clinical | 91.2 | 100 | 1.73 | 98.2866 | 5227270 | 45.29 | 115 | 112618 |
| SAMN15755100-GCF_014217295.1 | TJA114 | SAMN15755100 | GCF_014217295.1 | NonPC | Public | 2017 | China | Env | 92.3 | 100 | 1.01 | 98.3416 | 5568877 | 45.25 | 5 | 3522430 |
| SAMN15848043-GCA_018073145.1 | PNUSAV001235 | SAMN15848043 | GCA_018073145.1 | NonPC | Public | NA | USA | Clinical | 91.2 | 100 | 1.91 | 98.317 | 5183143 | 45.37 | 183 | 68976 |
| SAMN15848057-GCA_015746305.1 | PNUSAV001255 | SAMN15848057 | GCA_015746305.1 | NonPC | Public | NA | USA | Clinical | 90.5 | 100 | 3.15 | 98.3174 | 5268415 | 45.25 | 97 | 131070 |
| SAMN15848060-GCA_015746265.1 | PNUSAV001256 | SAMN15848060 | GCA_015746265.1 | NonPC | Public | NA | USA | Clinical | 89.7 | 100 | 1.54 | 98.3244 | 5198894 | 45.22 | 65 | 199425 |
| SAMN15848063-GCA_015746275.1 | PNUSAV001234 | SAMN15848063 | GCA_015746275.1 | NonPC | Public | NA | USA | Clinical | 91.3 | 100 | 2.09 | 98.3401 | 5361946 | 45.24 | 143 | 74978 |
| SAMN15857075-GCA_015746185.1 | PNUSAV001259 | SAMN15857075 | GCA_015746185.1 | NonPC | Public | NA | USA | Clinical | 90.6 | 100 | 1.41 | 98.3515 | 5206093 | 45.27 | 131 | 84442 |
| SAMN15857082-GCA_015746205.1 | PNUSAV001250 | SAMN15857082 | GCA_015746205.1 | NonPC | Public | NA | USA | Clinical | 88.3 | 100 | 1.67 | 98.1338 | 5124336 | 45.29 | 185 | 56250 |
| SAMN15889502-SRR13023685 | XMM658 | SAMN15889502 | SRR13023685 | NonPC | Public | 2019 | China | Env | 92.3 | 100 | 0.36 | 98.3142 | 5445537 | 45.22 | 83 | 402483 |
| SAMN15889503-SRR13023668 | XMS415 | SAMN15889503 | SRR13023668 | NonPC | Public | 2018 | China | Env | 91.4 | 100 | 0.22 | 98.3383 | 5280510 | 45.25 | 74 | 528198 |
| SAMN15889504-SRR13023689 | XMM1750 | SAMN15889504 | SRR13023689 | NonPC | Public | 2019 | China | Env | 91.4 | 100 | 0.41 | 98.2866 | 5370242 | 45.19 | 37 | 527734 |
| SAMN15889506-SRR13023675 | XMO693 | SAMN15889506 | SRR13023675 | NonPC | Public | 2018 | China | Env | 91.6 | 100 | 0 | 98.3687 | 5015228 | 45.35 | 56 | 447522 |
| SAMN15889509-GCA_023205955.1 | XMO116 | SAMN15889509 | GCA_023205955.1 | NonPC | Public | 2018 | China | Env | 89.6 | 100 | 0 | 97.9839 | 5466102 | 45.14 | 3 | 3517946 |
| SAMN15889509-SRR13022991 | XMO116 | SAMN15889509 | SRR13022991 | NonPC | Public | 2018 | China | Env | 89.7 | 100 | 0.33 | 98.3105 | 5045383 | 45.32 | 43 | 670356 |
| SAMN15889510-GCA_023205935.1 | XMM117 | SAMN15889510 | GCA_023205935.1 | NonPC | Public | 2019 | China | Env | 89.6 | 99.57 | 0.27 | 97.9697 | 5672846 | 45.02 | 4 | 3518178 |
| SAMN15889510-SRR13022981 | XMM117 | SAMN15889510 | SRR13022981 | NonPC | Public | 2019 | China | Env | 90 | 100 | 0.05 | 98.3452 | 5018485 | 45.38 | 40 | 529097 |
| SAMN15889511-SRR13023673 | XMS1061 | SAMN15889511 | SRR13023673 | NonPC | Public | 2018 | China | Env | 90.2 | 100 | 0.05 | 98.2829 | 5151135 | 45.17 | 78 | 188925 |

| | | | | | | | | | | | | | | | | |
|---|---|---|---|---|---|---|---|---|---|---|---|---|---|---|---|---|
| SAMN15889519-SRR13023670 | XMS1823 | SAMN15889519 | SRR13023670 | NonPC | Public | 2018 | China | Env | 91.9 | 100 | 0.27 | 98.2942 | 5378753 | 45.12 | 75 | 421894 |
| SAMN15889520-SRR13023676 | XMO1823 | SAMN15889520 | SRR13023676 | NonPC | Public | 2018 | China | Env | 91.8 | 100 | 0.27 | 98.278 | 5377162 | 45.11 | 72 | 460478 |
| SAMN15889529-SRR13022983 | DLS1073 | SAMN15889529 | SRR13022983 | NonPC | Public | 2018 | China | Env | 91.1 | 100 | 0 | 98.2761 | 5328272 | 45.22 | 45 | 584936 |
| SAMN15889530-GCA_016429965.1 | DLO1073 | SAMN15889530 | GCA_016429965.1 | NonPC | Public | 2018 | China | Env | 91 | 100 | 2.79 | 98.2856 | 5465467 | 45.24 | 56 | 443208 |
| SAMN15889534-SRR13023707 | DLM658 | SAMN15889534 | SRR13023707 | NonPC | Public | 2018 | China | Env | 92.2 | 100 | 0.36 | 98.2911 | 5446244 | 45.22 | 77 | 534152 |
| SAMN15889536-SRR13023700 | DLO396 | SAMN15889536 | SRR13023700 | NonPC | Public | 2018 | China | Env | 91.2 | 100 | 0 | 98.2839 | 5280494 | 45.17 | 54 | 309846 |
| SAMN15889538-SRR13023695 | DLS415 | SAMN15889538 | SRR13023695 | NonPC | Public | 2018 | China | Env | 91.7 | 100 | 0.6 | 98.3314 | 5349991 | 45.24 | 81 | 255370 |
| SAMN15889539-SRR13023699 | DLS1750 | SAMN15889539 | SRR13023699 | NonPC | Public | 2018 | China | Env | 91.4 | 100 | 0.41 | 98.3104 | 5450602 | 45.15 | 44 | 509057 |
| SAMN15889540-SRR13023710 | DLM1750 | SAMN15889540 | SRR13023710 | NonPC | Public | 2018 | China | Env | 91.4 | 100 | 0.41 | 98.2625 | 5566411 | 45.19 | 40 | 532411 |
| SAMN15889541-SRR13023703 | DLO1750 | SAMN15889541 | SRR13023703 | NonPC | Public | 2019 | China | Env | 91.4 | 100 | 0.41 | 98.2775 | 5566511 | 45.19 | 38 | 648171 |
| SAMN15889553-SRR13023709 | DLM180 | SAMN15889553 | SRR13023709 | NonPC | Public | 2019 | China | Env | 89.2 | 100 | 0.41 | 98.3349 | 5217761 | 45.29 | 36 | 335652 |
| SAMN15889554-SRR13023698 | DLS1799 | SAMN15889554 | SRR13023698 | NonPC | Public | 2019 | China | Env | 89.5 | 100 | 0.27 | 98.2625 | 5039699 | 45.32 | 30 | 573340 |
| SAMN15889555-GCA_023205915.1 | DLM1799 | SAMN15889555 | GCA_023205915.1 | NonPC | Public | 2019 | China | Env | 90.3 | 100 | 0.27 | 98.3001 | 5119321 | 45.38 | 2 | 3352135 |
| SAMN15889559-GCA_023205895.1 | DLM1805 | SAMN15889559 | GCA_023205895.1 | NonPC | Public | 2019 | China | Env | 92.4 | 100 | 0.03 | 98.3996 | 5091623 | 45.39 | 2 | 3328895 |
| SAMN15889561-GCA_018030985.1 | DHO1986 | SAMN15889561 | GCA_018030985.1 | NonPC | Public | 2018 | China | Env | 91.2 | 100 | 3.99 | 98.372 | 5441930 | 45.24 | 39 | 381778 |
| SAMN15889569-SRR13023666 | DHO1 | SAMN15889569 | SRR13023666 | NonPC | Public | 2018 | China | Env | 90.9 | 100 | 0.05 | 98.3919 | 5076176 | 45.26 | 39 | 522262 |
| SAMN15889573-SRR13023702 | DHM1 | SAMN15889573 | SRR13023702 | NonPC | Public | 2019 | China | Env | 90.1 | 100 | 0.19 | 98.2945 | 5238827 | 45.24 | 45 | 448892 |
| SAMN15889575-GCA_016430065.1 | DHM1630 | SAMN15889575 | GCA_016430065.1 | NonPC | Public | 2019 | China | Env | 91.2 | 100 | 3.01 | 98.3243 | 5498811 | 45.25 | 61 | 373536 |
| SAMN15889576-GCA_016429705.1 | DHO1785 | SAMN15889576 | GCA_016429705.1 | NonPC | Public | 2019 | China | Env | 90.9 | 100 | 3.32 | 98.3839 | 5482449 | 45.26 | 51 | 364579 |
| SAMN15889577-SRR13023667 | DHM658 | SAMN15889577 | SRR13023667 | NonPC | Public | 2019 | China | Env | 92.3 | 100 | 0.36 | 98.2932 | 5445198 | 45.22 | 81 | 534152 |
| SAMN15900785-GCA_015746065.1 | PNUSAV001239 | SAMN15900785 | GCA_015746065.1 | NonPC | Public | NA | USA | Clinical | 89.4 | 100 | 3.44 | 98.1155 | 5191463 | 45.34 | 55 | 295926 |
| SAMN15900789-GCA_015746105.1 | PNUSAV001240 | SAMN15900789 | GCA_015746105.1 | NonPC | Public | NA | USA | Clinical | 89.7 | 100 | 0.07 | 98.3641 | 5030725 | 45.34 | 42 | 460979 |
| SAMN15900791-GCA_015746165.1 | PNUSAV001244 | SAMN15900791 | GCA_015746165.1 | NonPC | Public | NA | USA | Clinical | 90.3 | 100 | 4.97 | 98.2943 | 5400740 | 45.32 | 77 | 116996 |
| SAMN15948301-GCA_015745965.1 | PNUSAV001269 | SAMN15948301 | GCA_015745965.1 | NonPC | Public | NA | USA | Clinical | 90.5 | 100 | 3.97 | 98.1531 | 5287442 | 45.17 | 59 | 372277 |
| SAMN15967179-GCA_018071165.1 | PNUSAV001277 | SAMN15967179 | GCA_018071165.1 | NonPC | Public | NA | USA | Clinical | 91.3 | 100 | 4.42 | 98.2366 | 5439583 | 45.35 | 139 | 87950 |
| SAMN15967488-GCA_015745905.1 | PNUSAV001275 | SAMN15967488 | GCA_015745905.1 | NonPC | Public | NA | USA | Clinical | 91.2 | 100 | 4.98 | 98.3184 | 5486855 | 45.33 | 51 | 520823 |
| SAMN16057162-GCA_020740855.1 | 16-VB00179 | SAMN16057162 | GCA_020740855.1 | NonPC | Public | 2016 | Germany | Env | 87.6 | 100 | 0.39 | 98.1499 | 5077070 | 45.39 | 88 | 324648 |
| SAMN16057163-GCA_020740875.1 | 16-VB00184 | SAMN16057163 | GCA_020740875.1 | NonPC | Public | 2016 | Germany | Env | 89.9 | 100 | 0.1 | 98.2918 | 5032590 | 45.39 | 62 | 390261 |
| SAMN16057386-GCA_020740915.1 | 16-VB00204 | SAMN16057386 | GCA_020740915.1 | NonPC | Public | 2016 | Germany | Env | 89.4 | 100 | 0.19 | 98.0624 | 5338597 | 45.06 | 81 | 340303 |
| SAMN16057388-GCA_020740925.1 | 17-VB00127 | SAMN16057388 | GCA_020740925.1 | NonPC | Public | 2017 | Germany | Env | 89.7 | 100 | 0.41 | 98.2454 | 5526021 | 45.23 | 90 | 176020 |
| SAMN16057396-GCA_020740955.1 | 17-VB00129 | SAMN16057396 | GCA_020740955.1 | NonPC | Public | 2017 | Germany | Env | 89.5 | 100 | 0.48 | 98.2461 | 5493587 | 45.18 | 97 | 164028 |
| SAMN16057399-GCA_020740975.1 | 17-VB00161 | SAMN16057399 | GCA_020740975.1 | NonPC | Public | 2017 | Germany | Env | 90.1 | 100 | 0.08 | 98.2701 | 4932209 | 45.32 | 65 | 266329 |
| SAMN16057400-GCA_020741035.1 | 17-VB00208 | SAMN16057400 | GCA_020741035.1 | NonPC | Public | 2017 | Germany | Env | 88.9 | 100 | 0 | 98.0328 | 5143071 | 45.22 | 78 | 246145 |
| SAMN16057402-GCA_020740995.1 | 17-VB00250 | SAMN16057402 | GCA_020740995.1 | NonPC | Public | 2017 | Germany | Env | 87.7 | 100 | 0.58 | 98.0757 | 4996732 | 45.36 | 73 | 289291 |
| SAMN16057403-GCA_020741055.1 | 17-VB00303 | SAMN16057403 | GCA_020741055.1 | NonPC | Public | 2017 | Germany | Env | 90.1 | 100 | 0.08 | 98.2828 | 4940235 | 45.31 | 78 | 275221 |
| SAMN16057404-GCA_020741075.1 | 17-VB00319 | SAMN16057404 | GCA_020741075.1 | NonPC | Public | 2017 | Germany | Env | 87.8 | 100 | 0.3 | 98.161 | 5001900 | 45.36 | 76 | 302023 |
| SAMN16057424-GCA_020741095.1 | 17-VB00330 | SAMN16057424 | GCA_020741095.1 | NonPC | Public | 2017 | Germany | Env | 87.8 | 100 | 0.38 | 98.1371 | 5087584 | 45.42 | 78 | 277671 |
| SAMN16057438-GCA_020741115.1 | 17-VB00358 | SAMN16057438 | GCA_020741115.1 | NonPC | Public | 2017 | Germany | Env | 88 | 100 | 0 | 98.1141 | 5256054 | 45.12 | 121 | 98244 |
| SAMN16057439-GCA_020741135.1 | 17-VB00364 | SAMN16057439 | GCA_020741135.1 | NonPC | Public | 2017 | Germany | Env | 89.3 | 100 | 0 | 98.325 | 5161316 | 45.51 | 39 | 584223 |
| SAMN16057440-GCA_020741165.1 | 17-VB00366 | SAMN16057440 | GCA_020741165.1 | NonPC | Public | 2017 | Germany | Env | 88.8 | 100 | 0 | 98.0445 | 5123519 | 45.21 | 70 | 236333 |
| SAMN16081350-GCA_018051465.1 | PNUSAV001273 | SAMN16081350 | GCA_018051465.1 | NonPC | Public | NA | USA | Clinical | 91.3 | 100 | 4.72 | 98.3122 | 5421998 | 45.29 | 51 | 269900 |
| SAMN16111715-GCA_015172915.1 | BTXS2 | SAMN16111715 | GCA_015172915.1 | NonPC | Public | 2019 | China | Env | 91.7 | 96.35 | 0.33 | 98.2703 | 5068431 | 45.49 | 3 | 3298213 |
| SAMN16133288-GCA_015745875.1 | 20MO07SH01-V1 | SAMN16133288 | GCA_015745875.1 | NonPC | Public | 2020 | USA | Env | 91.2 | 100 | 1.29 | 98.3065 | 5669694 | 45.28 | 63 | 456508 |
| SAMN16134149-GCA_015745865.1 | 20MO08SH01-V1 | SAMN16134149 | GCA_015745865.1 | NonPC | Public | 2020 | USA | Env | 91 | 100 | 1.36 | 98.398 | 5372996 | 45.36 | 34 | 549862 |
| SAMN16205401-GCF_015680975.1 | SH112 | SAMN16205401 | GCF_015680975.1 | PC | Public | 2019 | China | Clinical | 98.2 | 100 | 0 | 99.9864 | 5087431 | 45.28 | 79 | 373535 |
| SAMN16252940-SRR12704060 | 17-VB00214 | SAMN16252940 | SRR12704060 | NonPC | Public | 2017 | Germany | Env | 91.5 | 100 | 0.22 | 98.3563 | 5178387 | 45.28 | 2 | 3382781 |
| SAMN16380649-GCA_015745285.1 | PNCV000085 | SAMN16380649 | GCA_015745285.1 | NonPC | Public | 2020 | Canada | Clinical | 89.8 | 100 | 0.38 | 98.2668 | 5208413 | 45.18 | 50 | 259323 |
| SAMN16380650-GCA_015745065.1 | PNCV000086 | SAMN16380650 | GCA_015745065.1 | NonPC | Public | 2020 | Canada | Clinical | 89.9 | 100 | 0.77 | 98.252 | 5248772 | 45.19 | 44 | 445041 |

| | | | | | | | | | | | | | | | | |
|---|---|---|---|---|---|---|---|---|---|---|---|---|---|---|---|---|
| SAMN16380651-GCA_015745305.1 | PNCV000087 | SAMN16380651 | GCA_015745305.1 | NonPC | Public | 2020 | Canada | Clinical | 89.6 | 100 | 0.52 | 98.2986 | 5208384 | 45.18 | 48 | 313547 |
| SAMN16380652-GCA_015744985.1 | PNCV000088 | SAMN16380652 | GCA_015744985.1 | NonPC | Public | 2020 | Canada | Clinical | 89.7 | 100 | 0.24 | 98.2587 | 5191423 | 45.19 | 56 | 218912 |
| SAMN16380653-GCA_015745365.1 | PNCV000089 | SAMN16380653 | GCA_015745365.1 | NonPC | Public | 2020 | Canada | Clinical | 89.6 | 100 | 0.92 | 98.2188 | 5195352 | 45.18 | 128 | 92717 |
| SAMN16380654-GCA_015745405.1 | PNCV000090 | SAMN16380654 | GCA_015745405.1 | NonPC | Public | 2020 | Canada | Clinical | 89.6 | 100 | 0.8 | 98.2724 | 5204622 | 45.18 | 55 | 245711 |
| SAMN16380655-GCA_015745725.1 | PNCV000091 | SAMN16380655 | GCA_015745725.1 | NonPC | Public | 2020 | Canada | Clinical | 89.8 | 100 | 0.79 | 98.2512 | 5221959 | 45.18 | 42 | 459345 |
| SAMN16392350-GCA_015745825.1 | PNUSAV001257 | SAMN16392350 | GCA_015745825.1 | NonPC | Public | NA | USA | Clinical | 90.8 | 100 | 3.32 | 98.3254 | 5338105 | 45.2 | 55 | 254673 |
| SAMN16392372-GCA_018043385.1 | PNUSAV001333 | SAMN16392372 | GCA_018043385.1 | NonPC | Public | NA | USA | Clinical | 91.2 | 100 | 0.07 | 98.3226 | 5105541 | 45.32 | 195 | 59754 |
| SAMN16392373-GCA_018043585.1 | PNUSAV001332 | SAMN16392373 | GCA_018043585.1 | NonPC | Public | NA | USA | Clinical | 91.4 | 100 | 3 | 98.296 | 5238232 | 45.31 | 97 | 100534 |
| SAMN16392374-GCA_018043445.1 | PNUSAV001343 | SAMN16392374 | GCA_018043445.1 | NonPC | Public | NA | USA | Clinical | 91.4 | 100 | 4.41 | 98.3214 | 5282322 | 45.36 | 90 | 140882 |
| SAMN16392376-GCA_017978285.1 | PNUSAV001324 | SAMN16392376 | GCA_017978285.1 | NonPC | Public | NA | USA | Clinical | 90.9 | 100 | 2.23 | 98.3319 | 5162845 | 45.39 | 146 | 68745 |
| SAMN16392377-GCA_018043605.1 | PNUSAV001327 | SAMN16392377 | GCA_018043605.1 | NonPC | Public | NA | USA | Clinical | 90.8 | 100 | 0.89 | 98.2989 | 5117808 | 45.46 | 262 | 35152 |
| SAMN16392378-GCA_018043425.1 | PNUSAV001326 | SAMN16392378 | GCA_018043425.1 | NonPC | Public | NA | USA | Clinical | 91.3 | 100 | 2.36 | 98.2982 | 5207148 | 45.34 | 126 | 100766 |
| SAMN16392379-GCA_018043405.1 | PNUSAV001325 | SAMN16392379 | GCA_018043405.1 | NonPC | Public | NA | USA | Clinical | 91.1 | 100 | 1.09 | 98.2977 | 5170376 | 45.36 | 194 | 54731 |
| SAMN16392380-GCA_015745745.1 | PNUSAV001323 | SAMN16392380 | GCA_015745745.1 | NonPC | Public | NA | USA | Clinical | 91.3 | 100 | 3.54 | 98.29 | 5396923 | 45.29 | 67 | 216313 |
| SAMN16392381-GCA_018043565.1 | PNUSAV001272 | SAMN16392381 | GCA_018043565.1 | NonPC | Public | NA | USA | Clinical | 91.3 | 100 | 2.72 | 98.3363 | 5209630 | 45.28 | 73 | 166829 |
| SAMN16392382-GCA_018043365.1 | PNUSAV001270 | SAMN16392382 | GCA_018043365.1 | NonPC | Public | NA | USA | Clinical | 91.3 | 100 | 3.83 | 98.3131 | 5310162 | 45.29 | 87 | 119850 |
| SAMN16482544-GCA_021022935.1 | TAK_145 | SAMN16482544 | GCA_021022935.1 | NonPC | Public | 2017 | Nigeria | Env | 90.9 | 100 | 0.07 | 98.361 | 5177814 | 45.36 | 15 | 3261379 |
| SAMN16489435-GCA_018041165.1 | PNUSAV001369 | SAMN16489435 | GCA_018041165.1 | NonPC | Public | NA | USA | Clinical | 91.3 | 100 | 2.25 | 98.2741 | 5311756 | 45.32 | 85 | 122508 |
| SAMN16489444-GCA_018041045.1 | PNUSAV001386 | SAMN16489444 | GCA_018041045.1 | NonPC | Public | NA | USA | Clinical | 91.4 | 100 | 1.43 | 98.3041 | 5224219 | 45.31 | 75 | 221938 |
| SAMN16489445-GCA_018041125.1 | PNUSAV001388 | SAMN16489445 | GCA_018041125.1 | NonPC | Public | NA | USA | Clinical | 91.4 | 100 | 0.34 | 98.2928 | 5149510 | 45.25 | 62 | 286791 |
| SAMN16489450-GCA_018041025.1 | PNUSAV001336 | SAMN16489450 | GCA_018041025.1 | NonPC | Public | NA | USA | Clinical | 91.4 | 100 | 4.29 | 98.2705 | 5381490 | 45.29 | 48 | 385653 |
| SAMN16489452-GCA_018041145.1 | PNUSAV001368 | SAMN16489452 | GCA_018041145.1 | NonPC | Public | NA | USA | Clinical | 91.2 | 100 | 3.27 | 98.2768 | 5310340 | 45.33 | 99 | 112806 |
| SAMN16489455-GCA_018041085.1 | PNUSAV001384 | SAMN16489455 | GCA_018041085.1 | NonPC | Public | NA | USA | Clinical | 91 | 100 | 2.45 | 98.303 | 5195801 | 45.37 | 178 | 73215 |
| SAMN16489456-GCA_015745665.1 | PNUSAV001383 | SAMN16489456 | GCA_015745665.1 | NonPC | Public | NA | USA | Clinical | 89.7 | 100 | 2.42 | 98.2099 | 5338761 | 45.24 | 109 | 95061 |
| SAMN16489457-GCA_018041105.1 | PNUSAV001385 | SAMN16489457 | GCA_018041105.1 | NonPC | Public | NA | USA | Clinical | 91.3 | 100 | 1.63 | 98.3021 | 5223559 | 45.35 | 129 | 99306 |
| SAMN16651043-GCA_017980645.1 | PNUSAV001416 | SAMN16651043 | GCA_017980645.1 | NonPC | Public | NA | USA | Clinical | 91.3 | 100 | 4.63 | 98.3347 | 5461496 | 45.34 | 77 | 168087 |
| SAMN16676833-GCA_015744865.1 | 20MO09SH02-V1 | SAMN16676833 | GCA_015744865.1 | NonPC | Public | 2020 | USA | Env | 88.1 | 100 | 1.5 | 98.2106 | 5227663 | 45.23 | 53 | 417994 |
| SAMN16782947-GCA_016818355.1 | VP2 | SAMN16782947 | GCA_016818355.1 | NonPC | Public | 2008 | China | Clinical | 92.7 | 100 | 0 | 98.4231 | 5114789 | 45.19 | 56 | 386894 |
| SAMN16782948-GCA_016818375.1 | VP5 | SAMN16782948 | GCA_016818375.1 | NonPC | Public | 2010 | China | Env | 91.4 | 100 | 0.62 | 98.3756 | 4982571 | 45.36 | 56 | 459063 |
| SAMN16782949-GCA_016818295.1 | VP7 | SAMN16782949 | GCA_016818295.1 | NonPC | Public | 2008 | China | Env | 91.2 | 100 | 0 | 98.3671 | 5084405 | 45.29 | 73 | 244591 |
| SAMN16782950-GCA_016818275.1 | VP9 | SAMN16782950 | GCA_016818275.1 | NonPC | Public | 2009 | China | Clinical | 90.2 | 100 | 0.26 | 98.369 | 5118135 | 45.29 | 69 | 313086 |
| SAMN16782951-GCA_016818255.1 | VP10 | SAMN16782951 | GCA_016818255.1 | NonPC | Public | 2009 | China | Clinical | 90.9 | 100 | 0.05 | 98.3047 | 5068618 | 45.27 | 60 | 369835 |
| SAMN16782952-GCA_016818285.1 | VP11 | SAMN16782952 | GCA_016818285.1 | NonPC | Public | 2009 | China | Env | 91.3 | 100 | 0 | 98.3712 | 5042056 | 45.38 | 61 | 314920 |
| SAMN16782953-GCA_016818235.1 | VP13 | SAMN16782953 | GCA_016818235.1 | NonPC | Public | 2009 | China | Env | 91 | 100 | 0.07 | 98.2678 | 5300962 | 45.22 | 77 | 372998 |
| SAMN16782954-GCA_016818175.1 | VP14 | SAMN16782954 | GCA_016818175.1 | NonPC | Public | 2010 | China | Env | 90.7 | 100 | 0.05 | 98.3096 | 5106965 | 45.24 | 47 | 314864 |
| SAMN16782955-GCA_016818195.1 | VP15 | SAMN16782955 | GCA_016818195.1 | NonPC | Public | 2011 | China | Env | 90.3 | 100 | 0 | 98.3552 | 5005032 | 45.36 | 40 | 333934 |
| SAMN16782956-GCA_016818135.1 | VP17 | SAMN16782956 | GCA_016818135.1 | NonPC | Public | 2007 | China | Clinical | 92.1 | 100 | 0.46 | 98.3735 | 5120998 | 45.23 | 61 | 445271 |
| SAMN16782957-GCA_016818185.1 | VP18 | SAMN16782957 | GCA_016818185.1 | NonPC | Public | 2007 | China | Env | 90.6 | 100 | 0 | 98.3267 | 5184817 | 45.16 | 73 | 314857 |
| SAMN16782958-GCA_016818155.1 | VP19 | SAMN16782958 | GCA_016818155.1 | NonPC | Public | 2012 | China | Env | 92.6 | 100 | 0 | 98.4295 | 5117765 | 45.2 | 56 | 400897 |
| SAMN16782959-GCA_016818085.1 | VP20 | SAMN16782959 | GCA_016818085.1 | NonPC | Public | 2012 | China | Env | 92.1 | 100 | 0.46 | 98.3447 | 5109534 | 45.24 | 68 | 425915 |
| SAMN16782960-GCA_016818055.1 | VP21 | SAMN16782960 | GCA_016818055.1 | NonPC | Public | 2012 | China | Env | 92.1 | 100 | 0.46 | 98.3732 | 5127182 | 45.22 | 72 | 339959 |
| SAMN16782961-GCA_016818035.1 | VP22 | SAMN16782961 | GCA_016818035.1 | PC | Public | 2008 | China | Clinical | 98.3 | 100 | 0 | 99.9892 | 5155728 | 45.11 | 80 | 267792 |
| SAMN16782962-GCA_016818115.1 | VP25 | SAMN16782962 | GCA_016818115.1 | NonPC | Public | 2015 | China | Clinical | 91.1 | 100 | 0.25 | 98.4165 | 5275331 | 45.05 | 42 | 437195 |
| SAMN16782963-GCA_016818065.1 | VP26 | SAMN16782963 | GCA_016818065.1 | NonPC | Public | 2009 | China | Env | 90.8 | 100 | 0 | 98.4108 | 5133535 | 45.36 | 95 | 179819 |
| SAMN16782964-GCA_016817965.1 | VP27 | SAMN16782964 | GCA_016817965.1 | NonPC | Public | 2016 | China | Env | 88.6 | 100 | 0.48 | 98.2514 | 5093630 | 45.17 | 87 | 335116 |
| SAMN16782965-GCA_016817945.1 | VP28 | SAMN16782965 | GCA_016817945.1 | PC | Public | 2009 | China | Env | 97.4 | 100 | 0 | 99.9095 | 5090611 | 45.27 | 83 | 259602 |
| SAMN16782966-GCA_016818015.1 | VP29 | SAMN16782966 | GCA_016818015.1 | NonPC | Public | 2015 | China | Env | 88.7 | 100 | 0.05 | 98.3245 | 4981283 | 45.22 | 65 | 313257 |
| SAMN16782967-GCA_016817955.1 | VP31 | SAMN16782967 | GCA_016817955.1 | NonPC | Public | 2010 | China | Env | 88.2 | 100 | 0.41 | 98.2204 | 5269477 | 45.21 | 80 | 338315 |

| | | | | | | | | | | | | | | | | |
|---|---|---|---|---|---|---|---|---|---|---|---|---|---|---|---|---|
| SAMN16782968-GCA_016817935.1 | VP34 | SAMN16782968 | GCA_016817935.1 | NonPC | Public | 2008 | China | Env | 91.4 | 100 | 0.44 | 98.3319 | 5016921 | 45.36 | 56 | 279958 |
| SAMN16782969-GCA_016817905.1 | VP36 | SAMN16782969 | GCA_016817905.1 | NonPC | Public | 2010 | China | Clinical | 92.2 | 100 | 0.05 | 98.4007 | 5092632 | 45.28 | 37 | 535554 |
| SAMN16782970-GCA_016817845.1 | VP37 | SAMN16782970 | GCA_016817845.1 | NonPC | Public | 2010 | China | Env | 88.5 | 100 | 0.58 | 98.2576 | 5115080 | 45.16 | 93 | 336099 |
| SAMN16782971-GCA_016817875.1 | VP38 | SAMN16782971 | GCA_016817875.1 | NonPC | Public | 2008 | China | Env | 89.4 | 100 | 0 | 98.3168 | 4975899 | 45.44 | 62 | 388279 |
| SAMN16782972-GCA_016817835.1 | VP39 | SAMN16782972 | GCA_016817835.1 | NonPC | Public | 2008 | China | Env | 91.8 | 100 | 0.14 | 98.3479 | 5049587 | 45.25 | 87 | 415887 |
| SAMN16782973-GCA_016817885.1 | VP40 | SAMN16782973 | GCA_016817885.1 | NonPC | Public | 2010 | China | Clinical | 89.3 | 100 | 0.36 | 98.2635 | 5081764 | 45.05 | 61 | 380124 |
| SAMN16782974-GCA_016817815.1 | VP41 | SAMN16782974 | GCA_016817815.1 | NonPC | Public | 2008 | China | Env | 88.2 | 100 | 0.36 | 98.2614 | 5123036 | 45.16 | 138 | 318625 |
| SAMN16782975-GCA_016817795.1 | VP43 | SAMN16782975 | GCA_016817795.1 | NonPC | Public | 2009 | China | Clinical | 89.2 | 100 | 0 | 98.2768 | 4984275 | 45.37 | 66 | 420913 |
| SAMN16782976-GCA_016817765.1 | VP44 | SAMN16782976 | GCA_016817765.1 | NonPC | Public | 2010 | China | Env | 90.2 | 100 | 0.38 | 98.3217 | 5237246 | 45.18 | 78 | 292896 |
| SAMN16782977-GCA_016817755.1 | VP45 | SAMN16782977 | GCA_016817755.1 | NonPC | Public | 2010 | China | Env | 91 | 100 | 0.33 | 98.3058 | 5436978 | 44.96 | 97 | 229437 |
| SAMN16782978-GCA_016817735.1 | VP46 | SAMN16782978 | GCA_016817735.1 | NonPC | Public | 2010 | China | Env | 88.1 | 100 | 0.38 | 98.1985 | 5122756 | 45.22 | 73 | 239655 |
| SAMN16782979-GCA_016817715.1 | VP47 | SAMN16782979 | GCA_016817715.1 | NonPC | Public | 2010 | China | Env | 88.1 | 100 | 0.34 | 98.2339 | 5059985 | 45.2 | 46 | 384387 |
| SAMN16782980-GCA_016817695.1 | VP48 | SAMN16782980 | GCA_016817695.1 | NonPC | Public | 2010 | China | Env | 89.6 | 100 | 0 | 98.2924 | 4920954 | 45.42 | 42 | 420938 |
| SAMN16782981-GCA_016817645.1 | VP49 | SAMN16782981 | GCA_016817645.1 | NonPC | Public | 2010 | China | Env | 88.3 | 100 | 0.36 | 98.2675 | 4976068 | 45.3 | 64 | 346036 |
| SAMN16782982-GCA_016817675.1 | VP50 | SAMN16782982 | GCA_016817675.1 | NonPC | Public | 2010 | China | Env | 89.5 | 100 | 0.03 | 98.2255 | 5008041 | 45.32 | 57 | 385240 |
| SAMN16782983-GCA_016817635.1 | VP51 | SAMN16782983 | GCA_016817635.1 | NonPC | Public | 2010 | China | Env | 90 | 100 | 0.05 | 98.2699 | 5135986 | 45.25 | 67 | 314890 |
| SAMN16782984-GCA_016817595.1 | VP52 | SAMN16782984 | GCA_016817595.1 | NonPC | Public | 2010 | China | Env | 90.8 | 100 | 0.27 | 98.43 | 4904161 | 45.41 | 36 | 410562 |
| SAMN16782985-GCA_016817615.1 | VP53 | SAMN16782985 | GCA_016817615.1 | NonPC | Public | 2010 | China | Env | 88.3 | 100 | 0.39 | 98.1595 | 5052538 | 45.23 | 62 | 449240 |
| SAMN16782986-GCA_016817575.1 | VP54 | SAMN16782986 | GCA_016817575.1 | NonPC | Public | 2010 | China | Env | 88.1 | 100 | 0.41 | 98.294 | 5121681 | 45.24 | 64 | 341098 |
| SAMN16782987-GCA_016817555.1 | VP55 | SAMN16782987 | GCA_016817555.1 | NonPC | Public | 2010 | China | Env | 88.1 | 100 | 0.41 | 98.2734 | 5038982 | 45.32 | 59 | 395495 |
| SAMN16782988-GCA_016817535.1 | VP56 | SAMN16782988 | GCA_016817535.1 | NonPC | Public | 2010 | China | Env | 88.3 | 100 | 0.37 | 98.267 | 5179958 | 45.24 | 101 | 298370 |
| SAMN16782989-GCA_016817495.1 | VP57 | SAMN16782989 | GCA_016817495.1 | NonPC | Public | 2010 | China | Env | 89.7 | 100 | 0.05 | 98.3327 | 5020158 | 45.36 | 33 | 416546 |
| SAMN16782990-GCA_016817505.1 | VP58 | SAMN16782990 | GCA_016817505.1 | NonPC | Public | 2010 | China | Env | 89.7 | 100 | 0.05 | 98.3408 | 5020870 | 45.36 | 32 | 448833 |
| SAMN16782991-GCA_016817455.1 | VP60 | SAMN16782991 | GCA_016817455.1 | NonPC | Public | 2011 | China | Env | 89.5 | 100 | 0 | 98.2642 | 5048818 | 45.35 | 61 | 453623 |
| SAMN16782992-GCA_016817465.1 | VP61 | SAMN16782992 | GCA_016817465.1 | NonPC | Public | 2011 | China | Env | 89.8 | 100 | 0.27 | 98.2819 | 4999924 | 45.38 | 55 | 415181 |
| SAMN16782993-GCA_016817435.1 | VP62 | SAMN16782993 | GCA_016817435.1 | NonPC | Public | 2006 | China | Clinical | 91.1 | 100 | 0 | 98.3069 | 5025992 | 45.25 | 56 | 385461 |
| SAMN16782994-GCA_016817415.1 | VP63 | SAMN16782994 | GCA_016817415.1 | PC | Public | 2011 | China | Clinical | 97.8 | 100 | 0 | 99.9899 | 5052907 | 45.28 | 71 | 244120 |
| SAMN16782995-GCA_016817375.1 | VP64 | SAMN16782995 | GCA_016817375.1 | NonPC | Public | 2012 | China | Clinical | 91.2 | 100 | 0 | 98.3256 | 5158075 | 45.31 | 76 | 312695 |
| SAMN16782996-GCA_016817385.1 | VP66 | SAMN16782996 | GCA_016817385.1 | NonPC | Public | 2013 | China | Clinical | 89.6 | 100 | 0.07 | 98.3387 | 5100619 | 45.2 | 72 | 314880 |
| SAMN16782997-GCA_016827575.1 | VP68 | SAMN16782997 | GCA_016827575.1 | NonPC | Public | 2014 | China | Env | 91.2 | 100 | 0.05 | 98.3531 | 5011550 | 45.35 | 49 | 404463 |
| SAMN16782998-GCA_016827555.1 | VP69 | SAMN16782998 | GCA_016827555.1 | NonPC | Public | 2014 | China | Env | 91.2 | 100 | 0 | 98.3834 | 5015332 | 45.31 | 51 | 312701 |
| SAMN16782999-GCA_016827535.1 | VP70 | SAMN16782999 | GCA_016827535.1 | NonPC | Public | 2007 | China | Clinical | 88.1 | 100 | 0.41 | 98.2547 | 5058460 | 45.28 | 46 | 316310 |
| SAMN16783000-GCA_016827475.1 | VP71 | SAMN16783000 | GCA_016827475.1 | NonPC | Public | 2009 | China | Clinical | 90.7 | 100 | 0.05 | 98.3942 | 5007637 | 45.33 | 56 | 391848 |
| SAMN16783001-GCA_016827495.1 | VP73 | SAMN16783001 | GCA_016827495.1 | PC | Public | 2007 | China | Clinical | 97.3 | 100 | 0 | 99.9255 | 5094070 | 45.27 | 78 | 259608 |
| SAMN16783002-GCA_016827505.1 | VP74 | SAMN16783002 | GCA_016827505.1 | NonPC | Public | 2008 | China | Clinical | 92.1 | 100 | 0.05 | 98.4481 | 5063811 | 45.23 | 48 | 367683 |
| SAMN16783003-GCA_016827455.1 | VP75 | SAMN16783003 | GCA_016827455.1 | NonPC | Public | 2008 | China | Clinical | 92.1 | 100 | 0.06 | 98.4292 | 5154974 | 45.26 | 49 | 367683 |
| SAMN16783004-GCA_016827415.1 | VP76 | SAMN16783004 | GCA_016827415.1 | NonPC | Public | 2008 | China | Clinical | 92.1 | 100 | 0.06 | 98.4303 | 5154498 | 45.26 | 50 | 329680 |
| SAMN16783005-GCA_016827435.1 | VP77 | SAMN16783005 | GCA_016827435.1 | NonPC | Public | 2008 | China | Clinical | 92.1 | 100 | 0.05 | 98.4193 | 5063554 | 45.23 | 46 | 451197 |
| SAMN16783006-GCA_016827395.1 | VP78 | SAMN16783006 | GCA_016827395.1 | NonPC | Public | 2008 | China | Clinical | 92.1 | 100 | 0.05 | 98.4472 | 5062748 | 45.23 | 44 | 367797 |
| SAMN16783007-GCA_016827355.1 | VP79 | SAMN16783007 | GCA_016827355.1 | PC | Public | 2008 | China | Clinical | 97.6 | 100 | 0 | 99.9289 | 5147931 | 45.12 | 77 | 259614 |
| SAMN16783008-GCA_016827335.1 | VP80 | SAMN16783008 | GCA_016827335.1 | NonPC | Public | 2008 | China | Clinical | 92.1 | 100 | 0.05 | 98.4252 | 5063551 | 45.23 | 46 | 367554 |
| SAMN16783009-GCA_016827315.1 | VP81 | SAMN16783009 | GCA_016827315.1 | NonPC | Public | 2009 | China | Clinical | 92.4 | 100 | 0.65 | 98.4172 | 5111691 | 45.23 | 61 | 206930 |
| SAMN16783010-GCA_016827365.1 | VP82 | SAMN16783010 | GCA_016827365.1 | NonPC | Public | 2009 | China | Clinical | 92.1 | 100 | 0.05 | 98.4298 | 5062893 | 45.23 | 47 | 367554 |
| SAMN16783011-GCA_016827295.1 | VP83 | SAMN16783011 | GCA_016827295.1 | NonPC | Public | 2009 | China | Clinical | 96.9 | 100 | 1.41 | 98.5209 | 5390160 | 45.04 | 109 | 367420 |
| SAMN16783012-GCA_016827275.1 | VP84 | SAMN16783012 | GCA_016827275.1 | PC | Public | 2009 | China | Clinical | 97.1 | 100 | 0 | 99.9246 | 5093696 | 45.27 | 75 | 288515 |
| SAMN16783013-GCA_016827235.1 | VP85 | SAMN16783013 | GCA_016827235.1 | PC | Public | 2009 | China | Clinical | 98.1 | 100 | 0 | 99.9895 | 5065381 | 45.27 | 74 | 259608 |
| SAMN16783014-GCA_016827215.1 | VP86 | SAMN16783014 | GCA_016827215.1 | NonPC | Public | 2009 | China | Env | 91.4 | 100 | 0.27 | 98.4777 | 5083312 | 45.08 | 61 | 314780 |
| SAMN16783015-GCA_016827245.1 | VP87 | SAMN16783015 | GCA_016827245.1 | PC | Public | 2009 | China | Clinical | 97.3 | 100 | 0 | 99.9353 | 5093920 | 45.27 | 82 | 215453 |

| | | | | | | | | | | | | | | | | |
|---|---|---|---|---|---|---|---|---|---|---|---|---|---|---|---|---|
| SAMN16783016-GCA_016827165.1 | VP88 | SAMN16783016 | GCA_016827165.1 | PC | Public | 2009 | China | Clinical | 97.3 | 100 | 0 | 99.9339 | 5093474 | 45.27 | 78 | 259608 |
| SAMN16783017-GCA_016827195.1 | VP89 | SAMN16783017 | GCA_016827195.1 | PC | Public | 2009 | China | Clinical | 97.6 | 100 | 0 | 99.9349 | 5174491 | 45.09 | 80 | 243063 |
| SAMN16783018-GCA_016827145.1 | VP90 | SAMN16783018 | GCA_016827145.1 | PC | Public | 2009 | China | Clinical | 97.6 | 100 | 0 | 99.9331 | 5170783 | 45.09 | 84 | 243063 |
| SAMN16783019-GCA_016827115.1 | VP91 | SAMN16783019 | GCA_016827115.1 | PC | Public | 2009 | China | Clinical | 97.3 | 100 | 0.01 | 99.9365 | 5185195 | 45.29 | 84 | 215453 |
| SAMN16783020-GCA_016827125.1 | VP92 | SAMN16783020 | GCA_016827125.1 | PC | Public | 2009 | China | Clinical | 97.3 | 100 | 0 | 99.932 | 5154068 | 45.11 | 66 | 258756 |
| SAMN16783021-GCA_016827095.1 | VP93 | SAMN16783021 | GCA_016827095.1 | PC | Public | 2009 | China | Clinical | 97.3 | 100 | 0 | 99.9412 | 5154785 | 45.11 | 67 | 258756 |
| SAMN16783022-GCA_016827075.1 | VP95 | SAMN16783022 | GCA_016827075.1 | PC | Public | 2009 | China | Clinical | 97.6 | 100 | 0 | 99.9415 | 5173634 | 45.09 | 82 | 243063 |
| SAMN16783023-GCA_016827035.1 | VP96 | SAMN16783023 | GCA_016827035.1 | PC | Public | 2009 | China | Clinical | 97.2 | 100 | 0.07 | 99.92 | 5178016 | 45.26 | 89 | 215453 |
| SAMN16783024-GCA_016827045.1 | VP98 | SAMN16783024 | GCA_016827045.1 | NonPC | Public | 2009 | China | Clinical | 90.9 | 100 | 0.05 | 98.4171 | 4998521 | 45.26 | 44 | 367906 |
| SAMN16783025-GCA_016827015.1 | VP100 | SAMN16783025 | GCA_016827015.1 | PC | Public | 2010 | China | Clinical | 97.6 | 100 | 0 | 99.9317 | 5171769 | 45.09 | 84 | 243063 |
| SAMN16783026-GCA_016826995.1 | VP101 | SAMN16783026 | GCA_016826995.1 | PC | Public | 2010 | China | Clinical | 97.3 | 100 | 0 | 99.9248 | 5094680 | 45.27 | 79 | 243063 |
| SAMN16783027-GCA_016826955.1 | VP102 | SAMN16783027 | GCA_016826955.1 | PC | Public | 2011 | China | Env | 97.3 | 100 | 0 | 99.9066 | 5173184 | 45.29 | 69 | 259753 |
| SAMN16783028-GCA_016826975.1 | VP104 | SAMN16783028 | GCA_016826975.1 | PC | Public | 2010 | China | Clinical | 97.3 | 100 | 0 | 99.8942 | 5141212 | 45.28 | 72 | 259749 |
| SAMN16783029-GCA_016826915.1 | VP105 | SAMN16783029 | GCA_016826915.1 | PC | Public | 2007 | China | Clinical | 97.3 | 100 | 0 | 99.9321 | 5095433 | 45.27 | 82 | 243063 |
| SAMN16783030-GCA_016826895.1 | VP106 | SAMN16783030 | GCA_016826895.1 | PC | Public | 2007 | China | Clinical | 97.3 | 100 | 0 | 99.9362 | 5092600 | 45.27 | 81 | 259608 |
| SAMN16783031-GCA_016826905.1 | VP107 | SAMN16783031 | GCA_016826905.1 | NonPC | Public | 2007 | China | Env | 92.3 | 100 | 0.65 | 98.4042 | 5105799 | 45.23 | 63 | 209669 |
| SAMN16783032-GCA_016826875.1 | VP108 | SAMN16783032 | GCA_016826875.1 | NonPC | Public | 2007 | China | Clinical | 92.4 | 100 | 0.65 | 98.4163 | 5106555 | 45.23 | 62 | 209669 |
| SAMN16783033-GCA_016826855.1 | VP109 | SAMN16783033 | GCA_016826855.1 | NonPC | Public | 2008 | China | Clinical | 92.1 | 100 | 0.05 | 98.4174 | 5063982 | 45.23 | 46 | 451200 |
| SAMN16783034-GCA_016826835.1 | VP110 | SAMN16783034 | GCA_016826835.1 | NonPC | Public | 2010 | China | Clinical | 92.3 | 100 | 0.65 | 98.4069 | 5153182 | 45.18 | 63 | 203642 |
| SAMN16783035-GCA_016826795.1 | VP111 | SAMN16783035 | GCA_016826795.1 | PC | Public | 2012 | China | Clinical | 97.3 | 100 | 0 | 99.9117 | 5092468 | 45.28 | 72 | 259753 |
| SAMN16783036-GCA_016826815.1 | VP112 | SAMN16783036 | GCA_016826815.1 | PC | Public | 2011 | China | Env | 97.2 | 100 | 0 | 99.9316 | 5128059 | 45.26 | 76 | 259608 |
| SAMN16783037-GCA_016826775.1 | VP114 | SAMN16783037 | GCA_016826775.1 | NonPC | Public | 2008 | China | Clinical | 92.2 | 100 | 0.05 | 98.0826 | 5117798 | 45.23 | 42 | 446610 |
| SAMN16783038-GCA_016826735.1 | VP115 | SAMN16783038 | GCA_016826735.1 | NonPC | Public | 2008 | China | Clinical | 92.2 | 100 | 0.05 | 98.0527 | 5117116 | 45.23 | 44 | 433148 |
| SAMN16783039-GCA_016826715.1 | VP116 | SAMN16783039 | GCA_016826715.1 | NonPC | Public | 2008 | China | Clinical | 92.2 | 100 | 0.05 | 98.0714 | 5117062 | 45.23 | 45 | 337600 |
| SAMN16783040-GCA_016826755.1 | VP117 | SAMN16783040 | GCA_016826755.1 | NonPC | Public | 2008 | China | Clinical | 92.2 | 100 | 0.05 | 98.0645 | 5117588 | 45.23 | 42 | 433146 |
| SAMN16783041-GCA_016826695.1 | VP118 | SAMN16783041 | GCA_016826695.1 | NonPC | Public | 2008 | China | Clinical | 92.2 | 100 | 0.05 | 98.0843 | 5119931 | 45.23 | 44 | 337435 |
| SAMN16783042-GCA_016826655.1 | VP119 | SAMN16783042 | GCA_016826655.1 | NonPC | Public | 2008 | China | Clinical | 92.2 | 100 | 0.05 | 98.0936 | 5116198 | 45.23 | 45 | 433146 |
| SAMN16783043-GCA_016826675.1 | VP120 | SAMN16783043 | GCA_016826675.1 | NonPC | Public | 2008 | China | Clinical | 92.2 | 100 | 0.05 | 98.0539 | 5116767 | 45.23 | 44 | 433146 |
| SAMN16783044-GCA_016826635.1 | VP121 | SAMN16783044 | GCA_016826635.1 | PC | Public | 2009 | China | Clinical | 97.3 | 100 | 0 | 99.9118 | 5127247 | 45.28 | 71 | 259753 |
| SAMN16783045-GCA_016826575.1 | VP122 | SAMN16783045 | GCA_016826575.1 | NonPC | Public | 2012 | China | Env | 89.6 | 100 | 0 | 98.2866 | 4965934 | 45.36 | 62 | 421674 |
| SAMN16783046-GCA_016826615.1 | VP123 | SAMN16783046 | GCA_016826615.1 | PC | Public | 2009 | China | Clinical | 97.3 | 100 | 0 | 99.9022 | 5175683 | 45.29 | 74 | 259753 |
| SAMN16783047-GCA_016826585.1 | VP124 | SAMN16783047 | GCA_016826585.1 | PC | Public | 2009 | China | Clinical | 97.3 | 100 | 0 | 99.9053 | 5108202 | 45.27 | 72 | 259753 |
| SAMN16783048-GCA_016826555.1 | VP125 | SAMN16783048 | GCA_016826555.1 | PC | Public | 2010 | China | Clinical | 97.3 | 100 | 0 | 99.9194 | 5127467 | 45.28 | 69 | 267671 |
| SAMN16783049-GCA_016826515.1 | VP126 | SAMN16783049 | GCA_016826515.1 | PC | Public | 2010 | China | Clinical | 97.3 | 100 | 0 | 99.8978 | 5125745 | 45.28 | 70 | 259753 |
| SAMN16783050-GCA_016826535.1 | VP127 | SAMN16783050 | GCA_016826535.1 | NonPC | Public | 2009 | China | Env | 89.2 | 100 | 0 | 98.4148 | 4973249 | 45.4 | 75 | 336965 |
| SAMN16783051-GCA_016826455.1 | VP128 | SAMN16783051 | GCA_016826455.1 | NonPC | Public | 2010 | China | Env | 88.6 | 100 | 0.39 | 98.2251 | 5080797 | 45.23 | 76 | 383718 |
| SAMN16783052-GCA_016826485.1 | VP129 | SAMN16783052 | GCA_016826485.1 | PC | Public | 2010 | China | Clinical | 97.3 | 100 | 0 | 99.9039 | 5126085 | 45.28 | 72 | 259753 |
| SAMN16783053-GCA_016826475.1 | VP130 | SAMN16783053 | GCA_016826475.1 | PC | Public | 2008 | China | Clinical | 97.5 | 100 | 0 | 99.9329 | 5138020 | 45.21 | 67 | 259608 |
| SAMN16783054-GCA_016826435.1 | VP131 | SAMN16783054 | GCA_016826435.1 | PC | Public | 2008 | China | Clinical | 97.6 | 100 | 0.07 | 99.9348 | 5127674 | 45.26 | 78 | 259608 |
| SAMN16783055-GCA_016826405.1 | VP132 | SAMN16783055 | GCA_016826405.1 | PC | Public | 2008 | China | Clinical | 97.7 | 100 | 0 | 99.9411 | 5156502 | 45.12 | 78 | 259124 |
| SAMN16783056-GCA_016826395.1 | VP133 | SAMN16783056 | GCA_016826395.1 | PC | Public | 2008 | China | Clinical | 97.7 | 100 | 0 | 99.9156 | 5155044 | 45.12 | 81 | 259124 |
| SAMN16783057-GCA_016826375.1 | VP134 | SAMN16783057 | GCA_016826375.1 | PC | Public | 2008 | China | Clinical | 97.5 | 100 | 0 | 99.9319 | 5140357 | 45.21 | 69 | 259608 |
| SAMN16783058-GCA_016826355.1 | VP135 | SAMN16783058 | GCA_016826355.1 | NonPC | Public | 2008 | China | Clinical | 92.9 | 100 | 0.05 | 98.3984 | 5026572 | 45.31 | 38 | 411525 |
| SAMN16783059-GCA_016826335.1 | VP136 | SAMN16783059 | GCA_016826335.1 | NonPC | Public | 2008 | China | Clinical | 92.9 | 100 | 0.06 | 98.3744 | 5124455 | 45.33 | 40 | 411529 |
| SAMN16783060-GCA_016826315.1 | VP137 | SAMN16783060 | GCA_016826315.1 | PC | Public | 2008 | China | Clinical | 97.6 | 100 | 0 | 99.9314 | 5148483 | 45.12 | 77 | 259614 |
| SAMN16783061-GCA_016826295.1 | VP138 | SAMN16783061 | GCA_016826295.1 | NonPC | Public | 2008 | China | Clinical | 92.9 | 100 | 0.05 | 98.3876 | 5026016 | 45.31 | 41 | 411529 |
| SAMN16783062-GCA_016826255.1 | VP139 | SAMN16783062 | GCA_016826255.1 | PC | Public | 2008 | China | Clinical | 97.3 | 100 | 0 | 99.9351 | 5092057 | 45.27 | 82 | 259608 |
| SAMN16783063-GCA_016826275.1 | VP140 | SAMN16783063 | GCA_016826275.1 | PC | Public | 2009 | China | Clinical | 97.7 | 100 | 0 | 99.9277 | 5167108 | 45.1 | 74 | 294880 |

| | | | | | | | | | | | | | | | | |
|---|---|---|---|---|---|---|---|---|---|---|---|---|---|---|---|---|
| SAMN16783064-GCA_016826235.1 | VP141 | SAMN16783064 | GCA_016826235.1 | PC | Public | 2009 | China | Clinical | 97.5 | 100 | 0 | 99.9183 | 5071895 | 45.27 | 76 | 259608 |
| SAMN16783065-GCA_016826215.1 | VP142 | SAMN16783065 | GCA_016826215.1 | PC | Public | 2012 | China | Env | 97.6 | 100 | 0 | 99.9339 | 5168018 | 45.1 | 81 | 215452 |
| SAMN16783066-GCA_016826195.1 | VP143 | SAMN16783066 | GCA_016826195.1 | PC | Public | 2009 | China | Clinical | 97.3 | 100 | 0 | 99.9232 | 5106929 | 45.27 | 78 | 259608 |
| SAMN16783067-GCA_016826165.1 | VP144 | SAMN16783067 | GCA_016826165.1 | NonPC | Public | 2009 | China | Clinical | 92.5 | 100 | 0.05 | 98.3627 | 5044974 | 45.3 | 45 | 463468 |
| SAMN16783068-GCA_016826155.1 | VP145 | SAMN16783068 | GCA_016826155.1 | NonPC | Public | 2009 | China | Clinical | 92.9 | 100 | 0.05 | 98.3467 | 5053469 | 45.29 | 51 | 370781 |
| SAMN16783069-GCA_016826135.1 | VP146 | SAMN16783069 | GCA_016826135.1 | PC | Public | 2012 | China | Env | 97.7 | 100 | 0 | 99.9402 | 5149131 | 45.12 | 78 | 259614 |
| SAMN16783070-GCA_016826095.1 | VP147 | SAMN16783070 | GCA_016826095.1 | PC | Public | 2009 | China | Clinical | 97.7 | 100 | 0 | 99.9323 | 5159687 | 45.1 | 75 | 265748 |
| SAMN16783071-GCA_016826105.1 | VP148 | SAMN16783071 | GCA_016826105.1 | NonPC | Public | 2009 | China | Clinical | 92.9 | 100 | 0.05 | 98.3687 | 5033149 | 45.31 | 39 | 463452 |
| SAMN16783072-GCA_016826055.1 | VP149 | SAMN16783072 | GCA_016826055.1 | NonPC | Public | 2009 | China | Clinical | 92.6 | 100 | 0.05 | 98.3702 | 5052164 | 45.3 | 45 | 463468 |
| SAMN16783073-GCA_016826065.1 | VP150 | SAMN16783073 | GCA_016826065.1 | NonPC | Public | 2009 | China | Clinical | 89.4 | 100 | 0.05 | 98.3297 | 4902343 | 45.37 | 41 | 454841 |
| SAMN16783074-GCA_016826015.1 | VP151 | SAMN16783074 | GCA_016826015.1 | NonPC | Public | 2009 | China | Clinical | 92.9 | 100 | 0.05 | 98.3755 | 5052628 | 45.29 | 53 | 370781 |
| SAMN16783075-GCA_016826035.1 | VP152 | SAMN16783075 | GCA_016826035.1 | NonPC | Public | 2009 | China | Env | 91.6 | 100 | 0.33 | 98.385 | 4973915 | 45.37 | 75 | 336422 |
| SAMN16783076-GCA_016825975.1 | VP153 | SAMN16783076 | GCA_016825975.1 | NonPC | Public | 2010 | China | Env | 89.7 | 100 | 0.05 | 98.3407 | 5036877 | 45.24 | 46 | 450248 |
| SAMN16783077-GCA_016825955.1 | VP154 | SAMN16783077 | GCA_016825955.1 | NonPC | Public | 2010 | China | Clinical | 92.9 | 100 | 0.05 | 98.3407 | 5094303 | 45.25 | 55 | 411554 |
| SAMN16783078-GCA_016825995.1 | VP155 | SAMN16783078 | GCA_016825995.1 | PC | Public | 2010 | China | Clinical | 97.5 | 100 | 0 | 99.9161 | 5139874 | 45.11 | 77 | 259608 |
| SAMN16783079-GCA_016825915.1 | VP158 | SAMN16783079 | GCA_016825915.1 | PC | Public | 2010 | China | Clinical | 97.2 | 100 | 0 | 99.9351 | 5096634 | 45.27 | 78 | 259608 |
| SAMN16783080-GCA_016825855.1 | VP159 | SAMN16783080 | GCA_016825855.1 | PC | Public | 2010 | China | Clinical | 97.3 | 100 | 0 | 99.9309 | 5092035 | 45.27 | 81 | 243063 |
| SAMN16783081-GCA_016825905.1 | VP160 | SAMN16783081 | GCA_016825905.1 | PC | Public | 2010 | China | Clinical | 97.3 | 100 | 0 | 99.941 | 5095027 | 45.27 | 82 | 215453 |
| SAMN16783082-GCA_016825875.1 | VP161 | SAMN16783082 | GCA_016825875.1 | PC | Public | 2010 | China | Clinical | 97.3 | 100 | 0 | 99.9328 | 5098678 | 45.27 | 80 | 259608 |
| SAMN16783083-GCA_016825895.1 | VP162 | SAMN16783083 | GCA_016825895.1 | NonPC | Public | 2012 | China | Clinical | 92.9 | 100 | 0.05 | 98.388 | 5032439 | 45.31 | 36 | 463453 |
| SAMN16783084-GCA_016825835.1 | VP163 | SAMN16783084 | GCA_016825835.1 | PC | Public | 2010 | China | Clinical | 97.3 | 100 | 0 | 99.9215 | 5165376 | 45.11 | 80 | 243063 |
| SAMN16783085-GCA_016825795.1 | VP164 | SAMN16783085 | GCA_016825795.1 | PC | Public | 2010 | China | Clinical | 97.3 | 100 | 0 | 99.9357 | 5094328 | 45.27 | 77 | 259608 |
| SAMN16783086-GCA_016825785.1 | VP165 | SAMN16783086 | GCA_016825785.1 | PC | Public | 2010 | China | Clinical | 97.3 | 100 | 0 | 99.9248 | 5139295 | 45.22 | 83 | 215453 |
| SAMN16783087-GCA_016825775.1 | VP166 | SAMN16783087 | GCA_016825775.1 | PC | Public | 2010 | China | Clinical | 97.3 | 100 | 0 | 99.9247 | 5142233 | 45.22 | 84 | 215453 |
| SAMN16783088-GCA_016825745.1 | VP167 | SAMN16783088 | GCA_016825745.1 | PC | Public | 2010 | China | Clinical | 97.6 | 100 | 0 | 99.9325 | 5167303 | 45.09 | 72 | 258749 |
| SAMN16783089-GCA_016825735.1 | VP168 | SAMN16783089 | GCA_016825735.1 | NonPC | Public | 2010 | China | Clinical | 91.3 | 100 | 0 | 98.3425 | 5092427 | 45.27 | 91 | 296584 |
| SAMN16783090-GCA_016825685.1 | VP170 | SAMN16783090 | GCA_016825685.1 | PC | Public | 2012 | China | Env | 97.4 | 99.73 | 0 | 99.9167 | 5188074 | 45.22 | 73 | 288515 |
| SAMN16783091-GCA_016825675.1 | VP171 | SAMN16783091 | GCA_016825675.1 | NonPC | Public | 2012 | China | Env | 92.9 | 100 | 0.19 | 98.3806 | 5059902 | 45.28 | 41 | 411525 |
| SAMN16783092-GCA_016825615.1 | VP172 | SAMN16783092 | GCA_016825615.1 | NonPC | Public | 2012 | China | Env | 92.9 | 100 | 0.05 | 98.401 | 5027267 | 45.31 | 37 | 420783 |
| SAMN16783093-GCA_016825635.1 | VP173 | SAMN16783093 | GCA_016825635.1 | NonPC | Public | 2012 | China | Env | 92.9 | 100 | 0.05 | 98.3958 | 5104744 | 45.14 | 40 | 410431 |
| SAMN16783094-GCA_016825645.1 | VP174 | SAMN16783094 | GCA_016825645.1 | NonPC | Public | 2012 | China | Env | 92.9 | 100 | 0.05 | 98.396 | 5111482 | 45.13 | 38 | 463452 |
| SAMN16783095-GCA_016825715.1 | VP175 | SAMN16783095 | GCA_016825715.1 | NonPC | Public | 2011 | China | Env | 92.9 | 100 | 0.05 | 98.3734 | 5032639 | 45.31 | 38 | 463346 |
| SAMN16783096-GCA_016825595.1 | VP176 | SAMN16783096 | GCA_016825595.1 | PC | Public | 2011 | China | Env | 97.3 | 100 | 0 | 99.9393 | 5118072 | 45.26 | 78 | 259608 |
| SAMN16783097-GCA_016825575.1 | VP178 | SAMN16783097 | GCA_016825575.1 | PC | Public | 2011 | China | Env | 97.3 | 100 | 0 | 99.924 | 5117753 | 45.26 | 80 | 259608 |
| SAMN16783098-GCA_016825535.1 | VP179 | SAMN16783098 | GCA_016825535.1 | NonPC | Public | 2012 | China | Clinical | 91.2 | 100 | 0 | 98.3316 | 5156943 | 45.31 | 68 | 458124 |
| SAMN16783099-GCA_016825555.1 | VP180 | SAMN16783099 | GCA_016825555.1 | NonPC | Public | 2016 | China | Env | 92.6 | 100 | 0 | 98.4149 | 5137670 | 45.16 | 59 | 401209 |
| SAMN16783100-GCA_016825515.1 | VP181 | SAMN16783100 | GCA_016825515.1 | PC | Public | 2007 | China | Clinical | 97.5 | 100 | 0 | 99.9188 | 5069597 | 45.27 | 76 | 259614 |
| SAMN16783101-GCA_016825495.1 | VP182 | SAMN16783101 | GCA_016825495.1 | PC | Public | 2007 | China | Clinical | 97.6 | 100 | 0 | 99.9172 | 5159614 | 45.11 | 70 | 267638 |
| SAMN16783102-GCA_016825455.1 | VP183 | SAMN16783102 | GCA_016825455.1 | PC | Public | 2007 | China | Clinical | 97.5 | 100 | 0 | 99.9181 | 5070119 | 45.27 | 77 | 259614 |
| SAMN16783103-GCA_016825435.1 | VP184 | SAMN16783103 | GCA_016825435.1 | PC | Public | 2007 | China | Clinical | 97.3 | 100 | 0 | 99.9335 | 5093705 | 45.27 | 81 | 259608 |
| SAMN16783104-GCA_016825475.1 | VP185 | SAMN16783104 | GCA_016825475.1 | PC | Public | 2011 | China | Env | 97.3 | 100 | 0 | 99.9342 | 5102678 | 45.27 | 73 | 288515 |
| SAMN16783105-GCA_016825415.1 | VP186 | SAMN16783105 | GCA_016825415.1 | PC | Public | 2011 | China | Env | 97.3 | 100 | 0 | 99.9156 | 5132092 | 45.13 | 70 | 259608 |
| SAMN16783106-GCA_016825375.1 | VP187 | SAMN16783106 | GCA_016825375.1 | NonPC | Public | 2011 | China | Env | 92.1 | 100 | 0.19 | 98.3577 | 5040819 | 45.27 | 63 | 361486 |
| SAMN16783107-GCA_016825395.1 | VP188 | SAMN16783107 | GCA_016825395.1 | PC | Public | 2012 | China | Env | 98.1 | 100 | 0 | 99.9914 | 5065299 | 45.27 | 75 | 259608 |
| SAMN16783109-GCA_016825355.1 | VP191 | SAMN16783109 | GCA_016825355.1 | NonPC | Public | 2008 | China | Env | 90.9 | 100 | 0.1 | 98.3539 | 4965110 | 45.36 | 54 | 467283 |
| SAMN16783110-GCA_016825315.1 | VP193 | SAMN16783110 | GCA_016825315.1 | NonPC | Public | 2014 | China | Env | 88.1 | 100 | 0.33 | 98.2633 | 4960125 | 45.33 | 63 | 369589 |
| SAMN16783111-GCA_016825285.1 | VP194 | SAMN16783111 | GCA_016825285.1 | PC | Public | 2015 | China | Env | 97.3 | 100 | 0.07 | 99.9345 | 5147073 | 45.25 | 79 | 259608 |
| SAMN16783112-GCA_016825275.1 | VP197 | SAMN16783112 | GCA_016825275.1 | NonPC | Public | 2015 | China | Env | 91.1 | 100 | 0.05 | 98.2154 | 5168511 | 45.16 | 73 | 333775 |

| | | | | | | | | | | | | | | | | |
|---|---|---|---|---|---|---|---|---|---|---|---|---|---|---|---|---|
| SAMN16783113-GCA_016825255.1 | VP198 | SAMN16783113 | GCA_016825255.1 | NonPC | Public | 2015 | China | Clinical | 90.4 | 100 | 0 | 98.4237 | 4961678 | 45.38 | 48 | 331690 |
| SAMN16783114-GCA_016825195.1 | VP199 | SAMN16783114 | GCA_016825195.1 | NonPC | Public | 2009 | China | Clinical | 90.6 | 100 | 0.05 | 98.3514 | 5066863 | 45.26 | 71 | 231041 |
| SAMN16783115-GCA_016825175.1 | VP201 | SAMN16783115 | GCA_016825175.1 | NonPC | Public | 2008 | China | Env | 91.6 | 100 | 0.03 | 98.367 | 5082139 | 45.28 | 51 | 407565 |
| SAMN16783116-GCA_016825235.1 | VP204 | SAMN16783116 | GCA_016825235.1 | PC | Public | 2017 | China | Env | 98.1 | 100 | 0 | 99.9902 | 5066110 | 45.27 | 76 | 259608 |
| SAMN16783117-GCA_016825215.1 | VP207 | SAMN16783117 | GCA_016825215.1 | NonPC | Public | 2010 | China | Env | 91 | 100 | 0.54 | 98.3102 | 5103829 | 45.32 | 89 | 309677 |
| SAMN16783118-GCA_016825155.1 | VP209 | SAMN16783118 | GCA_016825155.1 | PC | Public | 2016 | China | Env | 97.3 | 100 | 0.01 | 99.8978 | 5280515 | 45.27 | 72 | 267671 |
| SAMN16783119-GCA_016825135.1 | VP210 | SAMN16783119 | GCA_016825135.1 | PC | Public | 2008 | China | Clinical | 97.3 | 100 | 0 | 99.9151 | 5174759 | 45.27 | 76 | 267671 |
| SAMN16783120-GCA_016825085.1 | VP211 | SAMN16783120 | GCA_016825085.1 | PC | Public | 2007 | China | Clinical | 97.3 | 100 | 0 | 99.9116 | 5127259 | 45.28 | 73 | 259753 |
| SAMN16783121-GCA_016825075.1 | VP212 | SAMN16783121 | GCA_016825075.1 | PC | Public | 2009 | China | Clinical | 97.3 | 100 | 0 | 99.8948 | 5127062 | 45.28 | 71 | 267671 |
| SAMN16783122-GCA_016825095.1 | VP213 | SAMN16783122 | GCA_016825095.1 | PC | Public | 2007 | China | Clinical | 97.3 | 100 | 0 | 99.8945 | 5126644 | 45.28 | 73 | 259753 |
| SAMN16783123-GCA_016825055.1 | VP214 | SAMN16783123 | GCA_016825055.1 | PC | Public | 2009 | China | Clinical | 97.3 | 100 | 0 | 99.9174 | 5127474 | 45.28 | 72 | 259749 |
| SAMN16783124-GCA_016825035.1 | VP215 | SAMN16783124 | GCA_016825035.1 | PC | Public | 2010 | China | Clinical | 97.3 | 100 | 0 | 99.8967 | 5134477 | 45.28 | 72 | 275888 |
| SAMN16783125-GCA_016825015.1 | VP216 | SAMN16783125 | GCA_016825015.1 | PC | Public | 2010 | China | Clinical | 97.3 | 100 | 0 | 99.8957 | 5140570 | 45.28 | 72 | 259753 |
| SAMN16783126-GCA_016824965.1 | VP217 | SAMN16783126 | GCA_016824965.1 | PC | Public | 2010 | China | Env | 97.3 | 100 | 3.35 | 99.872 | 5475022 | 44.77 | 218 | 243054 |
| SAMN16783127-GCA_016824975.1 | VP218 | SAMN16783127 | GCA_016824975.1 | PC | Public | 2010 | China | Clinical | 97.3 | 100 | 0 | 99.9016 | 5140365 | 45.28 | 70 | 259753 |
| SAMN16783128-GCA_016824955.1 | VP219 | SAMN16783128 | GCA_016824955.1 | PC | Public | 2010 | China | Clinical | 97.3 | 100 | 0 | 99.9008 | 5140707 | 45.28 | 69 | 259753 |
| SAMN16783129-GCA_016824875.1 | VP220 | SAMN16783129 | GCA_016824875.1 | PC | Public | 2010 | China | Clinical | 97.3 | 100 | 0 | 99.9173 | 5140912 | 45.28 | 69 | 259753 |
| SAMN16783130-GCA_016824885.1 | VP221 | SAMN16783130 | GCA_016824885.1 | PC | Public | 2010 | China | Clinical | 97.3 | 100 | 0 | 99.9007 | 5140040 | 45.28 | 70 | 267671 |
| SAMN16783131-GCA_016824925.1 | VP222 | SAMN16783131 | GCA_016824925.1 | PC | Public | 2010 | China | Clinical | 97.3 | 100 | 0 | 99.8804 | 5177888 | 45.27 | 74 | 267671 |
| SAMN16783132-GCA_016824915.1 | VP223 | SAMN16783132 | GCA_016824915.1 | PC | Public | 2010 | China | Clinical | 97.3 | 100 | 0 | 99.9064 | 5139736 | 45.28 | 71 | 259753 |
| SAMN16783133-GCA_016824855.1 | VP224 | SAMN16783133 | GCA_016824855.1 | PC | Public | 2010 | China | Clinical | 97.3 | 100 | 0 | 99.9062 | 5140720 | 45.28 | 70 | 259753 |
| SAMN16783134-GCA_016824775.1 | VP225 | SAMN16783134 | GCA_016824775.1 | PC | Public | 2010 | China | Clinical | 97.3 | 100 | 0 | 99.8964 | 5139831 | 45.28 | 71 | 259753 |
| SAMN16783135-GCA_016824785.1 | VP226 | SAMN16783135 | GCA_016824785.1 | PC | Public | 2010 | China | Clinical | 97.3 | 100 | 0 | 99.9018 | 5140935 | 45.28 | 69 | 259900 |
| SAMN16783136-GCA_016824755.1 | VP227 | SAMN16783136 | GCA_016824755.1 | PC | Public | 2010 | China | Clinical | 97.3 | 100 | 0 | 99.9273 | 5094083 | 45.27 | 82 | 215453 |
| SAMN16783137-GCA_016824835.1 | VP228 | SAMN16783137 | GCA_016824835.1 | PC | Public | 2010 | China | Clinical | 97.3 | 100 | 0 | 99.9013 | 5139439 | 45.28 | 72 | 259753 |
| SAMN16783138-GCA_016824795.1 | VP231 | SAMN16783138 | GCA_016824795.1 | NonPC | Public | 2015 | China | Env | 89.9 | 100 | 0.05 | 98.319 | 5218538 | 45.14 | 57 | 326993 |
| SAMN16783140-GCA_016824665.1 | VP234 | SAMN16783140 | GCA_016824665.1 | NonPC | Public | 2010 | China | Clinical | 92.1 | 100 | 0.46 | 98.3809 | 5166670 | 45.23 | 65 | 459961 |
| SAMN16783141-GCA_016824685.1 | VP236 | SAMN16783141 | GCA_016824685.1 | NonPC | Public | 2009 | China | Env | 90.6 | 100 | 0.18 | 98.3162 | 5394418 | 45.11 | 103 | 268711 |
| SAMN16783142-GCA_016824675.1 | VP237 | SAMN16783142 | GCA_016824675.1 | PC | Public | 2009 | China | Clinical | 97.3 | 100 | 0 | 99.9349 | 5101511 | 45.26 | 77 | 259608 |
| SAMN16783143-GCA_016824655.1 | VP240 | SAMN16783143 | GCA_016824655.1 | NonPC | Public | 2009 | China | Env | 88.4 | 100 | 0.43 | 98.208 | 5086043 | 45.27 | 56 | 393072 |
| SAMN16783144-GCA_016824635.1 | VP241 | SAMN16783144 | GCA_016824635.1 | NonPC | Public | 2009 | China | Env | 89.3 | 100 | 0.07 | 98.3007 | 4950887 | 45.36 | 39 | 422505 |
| SAMN16783145-GCA_016824585.1 | VP242 | SAMN16783145 | GCA_016824585.1 | NonPC | Public | 2010 | China | Env | 91 | 100 | 0 | 98.3436 | 5112730 | 45.34 | 70 | 210025 |
| SAMN16783146-GCA_016824615.1 | VP244 | SAMN16783146 | GCA_016824615.1 | NonPC | Public | 2016 | China | Env | 91.5 | 100 | 0.27 | 98.3385 | 5299022 | 45.15 | 58 | 450524 |
| SAMN16783147-GCA_016824575.1 | VP246 | SAMN16783147 | GCA_016824575.1 | NonPC | Public | 2008 | China | Env | 88.2 | 100 | 0.63 | 98.3006 | 5129554 | 45.15 | 89 | 418880 |
| SAMN16783148-GCA_016824555.1 | VP247 | SAMN16783148 | GCA_016824555.1 | NonPC | Public | 2008 | China | Clinical | 92.7 | 100 | 0 | 98.3957 | 5162436 | 45.13 | 52 | 434343 |
| SAMN16783149-GCA_016824535.1 | VP248 | SAMN16783149 | GCA_016824535.1 | NonPC | Public | 2008 | China | Clinical | 89.3 | 100 | 0.36 | 98.3135 | 5086053 | 45.05 | 62 | 371852 |
| SAMN16783150-GCA_016824485.1 | VP249 | SAMN16783150 | GCA_016824485.1 | NonPC | Public | 2008 | China | Clinical | 90.3 | 100 | 0 | 98.4154 | 5031649 | 45.19 | 53 | 369925 |
| SAMN16783151-GCA_016824515.1 | VP250 | SAMN16783151 | GCA_016824515.1 | NonPC | Public | 2009 | China | Clinical | 91.8 | 100 | 0 | 98.3764 | 5059750 | 45.23 | 53 | 453374 |
| SAMN16783152-GCA_016824475.1 | VP251 | SAMN16783152 | GCA_016824475.1 | NonPC | Public | 2009 | China | Clinical | 92.9 | 100 | 0 | 98.4008 | 5170551 | 45.17 | 50 | 369919 |
| SAMN16783153-GCA_016824445.1 | VP252 | SAMN16783153 | GCA_016824445.1 | NonPC | Public | 2010 | China | Clinical | 89.3 | 100 | 0.37 | 98.2722 | 5272247 | 44.98 | 66 | 379135 |
| SAMN16783154-GCA_016824375.1 | VP253 | SAMN16783154 | GCA_016824375.1 | NonPC | Public | 2009 | China | Clinical | 89.3 | 100 | 0.49 | 98.266 | 5088123 | 45.06 | 69 | 368499 |
| SAMN16783155-GCA_016824435.1 | VP254 | SAMN16783155 | GCA_016824435.1 | NonPC | Public | 2009 | China | Env | 88 | 100 | 0.36 | 98.2084 | 5055346 | 45.19 | 83 | 259070 |
| SAMN16783156-GCA_016824395.1 | VP255 | SAMN16783156 | GCA_016824395.1 | NonPC | Public | 2010 | China | Clinical | 92.7 | 100 | 0 | 98.402 | 5117270 | 45.21 | 52 | 368758 |
| SAMN16783157-GCA_016824415.1 | VP256 | SAMN16783157 | GCA_016824415.1 | NonPC | Public | 2011 | China | Env | 89.3 | 100 | 0.36 | 98.2626 | 5107346 | 45.04 | 60 | 379882 |
| SAMN16783158-GCA_016824335.1 | VP257 | SAMN16783158 | GCA_016824335.1 | NonPC | Public | 2009 | China | Env | 91.3 | 100 | 0 | 98.3845 | 5035728 | 45.32 | 56 | 314355 |
| SAMN16783159-GCA_016824345.1 | VP258 | SAMN16783159 | GCA_016824345.1 | NonPC | Public | 2009 | China | Clinical | 89.3 | 100 | 0.36 | 98.2705 | 5088407 | 45.06 | 60 | 379890 |
| SAMN16783160-GCA_016824285.1 | VP259 | SAMN16783160 | GCA_016824285.1 | NonPC | Public | 2009 | China | Clinical | 91.8 | 100 | 0 | 98.3929 | 5059428 | 45.23 | 53 | 453374 |
| SAMN16783161-GCA_016824315.1 | VP260 | SAMN16783161 | GCA_016824315.1 | NonPC | Public | 2011 | China | Env | 88.2 | 100 | 0.39 | 98.2373 | 5075252 | 45.17 | 87 | 450892 |

| | | | | | | | | | | | | | | | | |
|---|---|---|---|---|---|---|---|---|---|---|---|---|---|---|---|---|
| SAMN16783162-GCA_016824275.1 | VP261 | SAMN16783162 | GCA_016824275.1 | NonPC | Public | 2010 | China | Env | 88.3 | 99.98 | 0.3 | 98.3062 | 4912083 | 45.39 | 50 | 323720 |
| SAMN16783163-GCA_016824255.1 | VP262 | SAMN16783163 | GCA_016824255.1 | NonPC | Public | 2016 | China | Env | 91.3 | 100 | 0.05 | 98.3559 | 4972647 | 45.35 | 42 | 267267 |
| SAMN16783164-GCA_016824215.1 | VP263 | SAMN16783164 | GCA_016824215.1 | NonPC | Public | 2010 | China | Env | 88.1 | 100 | 0.37 | 98.2484 | 5136325 | 45.24 | 70 | 385288 |
| SAMN16783165-GCA_016824195.1 | VP264 | SAMN16783165 | GCA_016824195.1 | PC | Public | 2010 | China | Clinical | 98 | 100 | 0 | 99.9881 | 5065659 | 45.27 | 75 | 259608 |
| SAMN16783166-GCA_016824225.1 | VP265 | SAMN16783166 | GCA_016824225.1 | NonPC | Public | 2013 | China | Env | 89.3 | 100 | 0.36 | 98.2664 | 5099200 | 45.04 | 61 | 375996 |
| SAMN16783167-GCA_016824175.1 | VP266 | SAMN16783167 | GCA_016824175.1 | NonPC | Public | 2007 | China | Clinical | 92.7 | 100 | 0 | 98.4149 | 5107715 | 45.21 | 53 | 369925 |
| SAMN16783168-GCA_016824135.1 | VP267 | SAMN16783168 | GCA_016824135.1 | NonPC | Public | 2007 | China | Clinical | 89.3 | 100 | 0.36 | 98.2588 | 5076142 | 45.07 | 53 | 368499 |
| SAMN16783169-GCA_016824155.1 | VP268 | SAMN16783169 | GCA_016824155.1 | NonPC | Public | 2012 | China | Env | 92.7 | 100 | 0.01 | 98.3997 | 5205491 | 45.23 | 56 | 373486 |
| SAMN16783170-GCA_016824105.1 | VP269 | SAMN16783170 | GCA_016824105.1 | NonPC | Public | 2012 | China | Env | 87.8 | 100 | 0.4 | 98.2711 | 5077421 | 45.18 | 116 | 287762 |
| SAMN16783171-GCA_016824095.1 | VP270 | SAMN16783171 | GCA_016824095.1 | NonPC | Public | 2010 | China | Env | 89.5 | 100 | 0.02 | 98.2867 | 5022470 | 45.42 | 82 | 141959 |
| SAMN16783172-GCA_016824075.1 | VP271 | SAMN16783172 | GCA_016824075.1 | NonPC | Public | 2010 | China | Env | 88.4 | 100 | 0.4 | 98.2943 | 5065326 | 45.27 | 69 | 436271 |
| SAMN16783173-GCA_016824045.1 | VP272 | SAMN16783173 | GCA_016824045.1 | PC | Public | 2017 | China | Env | 98 | 100 | 0 | 99.9886 | 5068819 | 45.27 | 78 | 259608 |
| SAMN16783174-GCA_016824035.1 | VP273 | SAMN16783174 | GCA_016824035.1 | NonPC | Public | 2007 | China | Clinical | 92.1 | 100 | 0.19 | 98.3647 | 5028886 | 45.25 | 62 | 372807 |
| SAMN16783175-GCA_016823995.1 | VP274 | SAMN16783175 | GCA_016823995.1 | NonPC | Public | 2007 | China | Clinical | 92.1 | 100 | 0.19 | 98.3625 | 5027331 | 45.25 | 60 | 406273 |
| SAMN16783176-GCA_016824005.1 | VP275 | SAMN16783176 | GCA_016824005.1 | NonPC | Public | 2008 | China | Clinical | 92.3 | 100 | 0.05 | 98.3309 | 5052744 | 45.27 | 56 | 437858 |
| SAMN16783177-GCA_016823975.1 | VP276 | SAMN16783177 | GCA_016823975.1 | NonPC | Public | 2008 | China | Clinical | 91.8 | 100 | 0 | 98.4146 | 5058149 | 45.24 | 48 | 417268 |
| SAMN16783178-GCA_016823935.1 | VP277 | SAMN16783178 | GCA_016823935.1 | NonPC | Public | 2008 | China | Clinical | 90.9 | 100 | 0 | 98.4035 | 4939593 | 45.35 | 50 | 340461 |
| SAMN16783179-GCA_016823915.1 | VP278 | SAMN16783179 | GCA_016823915.1 | NonPC | Public | 2008 | China | Env | 90.8 | 100 | 0.03 | 98.3474 | 5088976 | 45.32 | 56 | 301825 |
| SAMN16783180-GCA_016823945.1 | VP279 | SAMN16783180 | GCA_016823945.1 | NonPC | Public | 2008 | China | Env | 92.3 | 100 | 0.05 | 98.3344 | 5007990 | 45.26 | 53 | 450167 |
| SAMN16783181-GCA_016823895.1 | VP280 | SAMN16783181 | GCA_016823895.1 | NonPC | Public | 2008 | China | Clinical | 91.7 | 100 | 0 | 98.4258 | 5042547 | 45.25 | 45 | 417262 |
| SAMN16783182-GCA_016823835.1 | VP281 | SAMN16783182 | GCA_016823835.1 | NonPC | Public | 2008 | China | Clinical | 91.7 | 100 | 0 | 98.4272 | 5042982 | 45.25 | 47 | 417262 |
| SAMN16783183-GCA_016823825.1 | VP282 | SAMN16783183 | GCA_016823825.1 | NonPC | Public | 2008 | China | Clinical | 91.7 | 100 | 0 | 98.4142 | 5042417 | 45.25 | 46 | 417262 |
| SAMN16783184-GCA_016823875.1 | VP283 | SAMN16783184 | GCA_016823875.1 | NonPC | Public | 2009 | China | Clinical | 92.1 | 100 | 0.19 | 98.3426 | 5033597 | 45.25 | 58 | 435976 |
| SAMN16783185-GCA_016823815.1 | VP284 | SAMN16783185 | GCA_016823815.1 | NonPC | Public | 2009 | China | Clinical | 92.2 | 100 | 0.19 | 98.3268 | 5045298 | 45.29 | 62 | 289540 |
| SAMN16783186-GCA_016823785.1 | VP285 | SAMN16783186 | GCA_016823785.1 | PC | Public | 2015 | China | Env | 98 | 100 | 0 | 99.9918 | 5064345 | 45.27 | 79 | 259608 |
| SAMN16783187-GCA_016823725.1 | VP286 | SAMN16783187 | GCA_016823725.1 | NonPC | Public | 2009 | China | Clinical | 92.2 | 100 | 0.19 | 98.3547 | 5011778 | 45.26 | 62 | 313843 |
| SAMN16783188-GCA_016823775.1 | VP287 | SAMN16783188 | GCA_016823775.1 | NonPC | Public | 2010 | China | Env | 91.2 | 100 | 0.14 | 98.3639 | 5048434 | 45.29 | 75 | 256777 |
| SAMN16783189-GCA_016823735.1 | VP288 | SAMN16783189 | GCA_016823735.1 | NonPC | Public | 2010 | China | Clinical | 92.1 | 100 | 0.19 | 98.3668 | 4994973 | 45.25 | 57 | 450167 |
| SAMN16783190-GCA_016823715.1 | VP289 | SAMN16783190 | GCA_016823715.1 | NonPC | Public | 2010 | China | Clinical | 91.7 | 100 | 0 | 98.41 | 5081608 | 45.24 | 44 | 417684 |
| SAMN16783191-GCA_016823695.1 | VP290 | SAMN16783191 | GCA_016823695.1 | NonPC | Public | 2010 | China | Clinical | 91.7 | 100 | 0 | 98.4163 | 5081557 | 45.24 | 44 | 417684 |
| SAMN16783192-GCA_016823635.1 | VP291 | SAMN16783192 | GCA_016823635.1 | NonPC | Public | 2010 | China | Clinical | 91.7 | 100 | 0 | 98.3973 | 5082394 | 45.24 | 43 | 417684 |
| SAMN16783193-GCA_016823645.1 | VP292 | SAMN16783193 | GCA_016823645.1 | NonPC | Public | 2010 | China | Clinical | 91.7 | 100 | 0 | 98.4149 | 5082259 | 45.24 | 45 | 417672 |
| SAMN16783194-GCA_016823675.1 | VP293 | SAMN16783194 | GCA_016823675.1 | NonPC | Public | 2010 | China | Clinical | 92.3 | 100 | 0.05 | 98.3844 | 5037558 | 45.29 | 55 | 335253 |
| SAMN16783195-GCA_016823615.1 | VP294 | SAMN16783195 | GCA_016823615.1 | NonPC | Public | 2010 | China | Clinical | 91.7 | 100 | 0 | 98.4163 | 5081611 | 45.24 | 47 | 415091 |
| SAMN16783196-GCA_016823595.1 | VP295 | SAMN16783196 | GCA_016823595.1 | NonPC | Public | 2010 | China | Clinical | 91.7 | 100 | 0 | 98.4109 | 5082168 | 45.24 | 45 | 423140 |
| SAMN16783197-GCA_016823535.1 | VP296 | SAMN16783197 | GCA_016823535.1 | NonPC | Public | 2010 | China | Clinical | 92.3 | 100 | 0.05 | 98.3387 | 5073342 | 45.26 | 57 | 403831 |
| SAMN16783198-GCA_016823555.1 | VP297 | SAMN16783198 | GCA_016823555.1 | NonPC | Public | 2010 | China | Clinical | 91.7 | 100 | 0 | 98.416 | 5081794 | 45.24 | 43 | 417684 |
| SAMN16783199-GCA_016823575.1 | VP298 | SAMN16783199 | GCA_016823575.1 | NonPC | Public | 2016 | China | Env | 92.1 | 100 | 0.19 | 98.3777 | 5011948 | 45.24 | 60 | 367559 |
| SAMN16783200-GCA_016823495.1 | VP299 | SAMN16783200 | GCA_016823495.1 | NonPC | Public | 2012 | China | Clinical | 88.4 | 100 | 0.05 | 98.3423 | 4934181 | 45.31 | 74 | 295680 |
| SAMN16783201-GCA_016823515.1 | VP300 | SAMN16783201 | GCA_016823515.1 | NonPC | Public | 2008 | China | Env | 92 | 100 | 0 | 98.3503 | 5074288 | 45.24 | 77 | 400530 |
| SAMN16783202-GCA_016823475.1 | VP301 | SAMN16783202 | GCA_016823475.1 | NonPC | Public | 2008 | China | Env | 89.5 | 100 | 0.14 | 98.245 | 4953023 | 45.38 | 46 | 455744 |
| SAMN16783203-GCA_016823455.1 | VP302 | SAMN16783203 | GCA_016823455.1 | PC | Public | 2015 | China | Env | 98.3 | 100 | 0 | 99.9883 | 5125457 | 45.25 | 76 | 259614 |
| SAMN16783204-GCA_016823435.1 | VP303 | SAMN16783204 | GCA_016823435.1 | NonPC | Public | 2008 | China | Env | 90.9 | 100 | 0.14 | 98.34 | 5049507 | 45.33 | 71 | 403682 |
| SAMN16783205-GCA_016823395.1 | VP306 | SAMN16783205 | GCA_016823395.1 | NonPC | Public | 2009 | China | Clinical | 91.2 | 100 | 0 | 98.349 | 5147707 | 45.26 | 70 | 189010 |
| SAMN16783206-GCA_016823345.1 | VP307 | SAMN16783206 | GCA_016823345.1 | NonPC | Public | 2009 | China | Env | 89.4 | 99.95 | 0 | 98.3899 | 4904885 | 45.4 | 46 | 382195 |
| SAMN16783207-GCA_016823415.1 | VP308 | SAMN16783207 | GCA_016823415.1 | NonPC | Public | 2009 | China | Env | 89.7 | 100 | 0 | 98.3108 | 5027642 | 45.3 | 68 | 344779 |
| SAMN16783208-GCA_016823335.1 | VP309 | SAMN16783208 | GCA_016823335.1 | NonPC | Public | 2009 | China | Clinical | 91 | 100 | 0.23 | 98.3622 | 5116315 | 45.17 | 61 | 273780 |
| SAMN16783209-GCA_016823375.1 | VP310 | SAMN16783209 | GCA_016823375.1 | NonPC | Public | 2010 | China | Env | 91.1 | 100 | 0.05 | 98.287 | 5069901 | 45.26 | 99 | 280703 |

| | | | | | | | | | | | | | | | | |
|---|---|---|---|---|---|---|---|---|---|---|---|---|---|---|---|---|
| SAMN16783210-GCA_016823315.1 | VP311 | SAMN16783210 | GCA_016823315.1 | NonPC | Public | 2010 | China | Env | 90.8 | 100 | 0.03 | 98.2887 | 5176557 | 45.19 | 71 | 310461 |
| SAMN16783211-GCA_016823275.1 | VP312 | SAMN16783211 | GCA_016823275.1 | NonPC | Public | 2012 | China | Env | 90.3 | 100 | 0.27 | 98.2916 | 5116659 | 45.3 | 43 | 344065 |
| SAMN16783212-GCA_016823295.1 | VP313 | SAMN16783212 | GCA_016823295.1 | NonPC | Public | 2010 | China | Env | 90.8 | 100 | 0 | 98.3883 | 5017483 | 45.25 | 63 | 265152 |
| SAMN16783213-GCA_016823225.1 | VP314 | SAMN16783213 | GCA_016823225.1 | NonPC | Public | 2010 | China | Env | 89.6 | 100 | 0.07 | 98.2923 | 5078327 | 45.19 | 57 | 402069 |
| SAMN16783214-GCA_016823255.1 | VP315 | SAMN16783214 | GCA_016823255.1 | NonPC | Public | 2008 | China | Env | 91 | 100 | 0 | 98.3605 | 5021325 | 45.3 | 64 | 460431 |
| SAMN16783215-GCA_016823215.1 | VP316 | SAMN16783215 | GCA_016823215.1 | NonPC | Public | 2010 | China | Env | 90.3 | 100 | 0 | 98.3688 | 5113769 | 45.34 | 71 | 210025 |
| SAMN16783216-GCA_016823195.1 | VP317 | SAMN16783216 | GCA_016823195.1 | NonPC | Public | 2010 | China | Env | 90.9 | 100 | 0.08 | 98.4314 | 4939306 | 45.37 | 63 | 313131 |
| SAMN16783218-GCA_016823155.1 | VP320 | SAMN16783218 | GCA_016823155.1 | NonPC | Public | 2007 | China | Clinical | 92.1 | 100 | 0.6 | 98.3429 | 5173592 | 45.22 | 60 | 339962 |
| SAMN16783219-GCA_016823115.1 | VP321 | SAMN16783219 | GCA_016823115.1 | NonPC | Public | 2008 | China | Clinical | 92.7 | 100 | 0.05 | 98.4044 | 5082722 | 45.28 | 38 | 519551 |
| SAMN16783220-GCA_016823135.1 | VP322 | SAMN16783220 | GCA_016823135.1 | NonPC | Public | 2008 | China | Env | 90.7 | 100 | 0.05 | 98.3883 | 5052720 | 45.25 | 62 | 257984 |
| SAMN16783221-GCA_016823095.1 | VP323 | SAMN16783221 | GCA_016823095.1 | NonPC | Public | 2010 | China | Clinical | 89.9 | 100 | 0 | 98.2983 | 4987743 | 45.35 | 52 | 415902 |
| SAMN16783222-GCA_016823055.1 | VP324 | SAMN16783222 | GCA_016823055.1 | PC | Public | 2015 | China | Env | 98.1 | 100 | 0 | 99.9907 | 5067177 | 45.27 | 77 | 259608 |
| SAMN16783223-GCA_016823025.1 | VP325 | SAMN16783223 | GCA_016823025.1 | NonPC | Public | 2012 | China | Clinical | 92.3 | 100 | 0.05 | 98.3807 | 5008626 | 45.26 | 52 | 372799 |
| SAMN16783224-GCA_016823065.1 | VP326 | SAMN16783224 | GCA_016823065.1 | NonPC | Public | 2012 | China | Clinical | 92.1 | 100 | 0.19 | 98.357 | 5014195 | 45.23 | 61 | 367822 |
| SAMN16783225-GCA_016823015.1 | VP327 | SAMN16783225 | GCA_016823015.1 | NonPC | Public | 2012 | China | Clinical | 88.5 | 100 | 0.19 | 98.3371 | 4964239 | 45.26 | 56 | 313257 |
| SAMN16783226-GCA_016822995.1 | VP328 | SAMN16783226 | GCA_016822995.1 | NonPC | Public | 2007 | China | Clinical | 92.1 | 100 | 0.19 | 98.3341 | 5013882 | 45.24 | 59 | 368045 |
| SAMN16783227-GCA_016822975.1 | VP329 | SAMN16783227 | GCA_016822975.1 | NonPC | Public | 2013 | China | Env | 88.7 | 100 | 0.05 | 98.3325 | 4980386 | 45.23 | 65 | 313257 |
| SAMN16783228-GCA_016822945.1 | VP330 | SAMN16783228 | GCA_016822945.1 | NonPC | Public | 2010 | China | Env | 91.7 | 100 | 0 | 98.4366 | 5070933 | 45.3 | 46 | 425371 |
| SAMN16783229-GCA_016822935.1 | VP332 | SAMN16783229 | GCA_016822935.1 | NonPC | Public | 2015 | China | Env | 88.7 | 100 | 0.05 | 98.307 | 4985373 | 45.22 | 66 | 313257 |
| SAMN16783230-GCA_016822915.1 | VP333 | SAMN16783230 | GCA_016822915.1 | NonPC | Public | 2015 | China | Env | 92.1 | 100 | 0.19 | 98.3381 | 5031177 | 45.22 | 61 | 367822 |
| SAMN16783231-GCA_016822895.1 | VP334 | SAMN16783231 | GCA_016822895.1 | NonPC | Public | 2016 | China | Env | 88.7 | 100 | 0.05 | 98.3264 | 4986301 | 45.21 | 66 | 313257 |
| SAMN16783232-GCA_016822865.1 | VP335 | SAMN16783232 | GCA_016822865.1 | NonPC | Public | 2016 | China | Env | 88.7 | 100 | 0.05 | 98.3243 | 5126527 | 45.25 | 56 | 245434 |
| SAMN16783233-GCA_016822835.1 | VP336 | SAMN16783233 | GCA_016822835.1 | NonPC | Public | 2011 | China | Clinical | 92.2 | 100 | 0.19 | 98.3483 | 5042504 | 45.26 | 60 | 369099 |
| SAMN16783234-GCA_016822855.1 | VP337 | SAMN16783234 | GCA_016822855.1 | NonPC | Public | 2011 | China | Clinical | 92.1 | 100 | 0.19 | 98.3453 | 4995728 | 45.25 | 59 | 372812 |
| SAMN16783235-GCA_016822815.1 | VP338 | SAMN16783235 | GCA_016822815.1 | NonPC | Public | 2012 | China | Clinical | 90.9 | 100 | 0 | 98.3978 | 4981501 | 45.35 | 47 | 340525 |
| SAMN16783236-GCA_016822795.1 | VP339 | SAMN16783236 | GCA_016822795.1 | NonPC | Public | 2012 | China | Clinical | 92.3 | 100 | 0.33 | 98.3385 | 5039992 | 45.24 | 51 | 406578 |
| SAMN16783237-GCA_016822775.1 | VP340 | SAMN16783237 | GCA_016822775.1 | NonPC | Public | 2012 | China | Clinical | 92.3 | 100 | 0.05 | 98.3354 | 5109502 | 45.25 | 58 | 450167 |
| SAMN16783238-GCA_016822735.1 | VP341 | SAMN16783238 | GCA_016822735.1 | PC | Public | 2017 | China | Env | 98.1 | 100 | 0 | 99.9869 | 5065803 | 45.27 | 75 | 259608 |
| SAMN16783239-GCA_016822755.1 | VP342 | SAMN16783239 | GCA_016822755.1 | PC | Public | 2017 | China | Env | 98 | 100 | 0.27 | 99.9937 | 5117424 | 45.27 | 76 | 259608 |
| SAMN16783240-GCA_016822715.1 | VP343 | SAMN16783240 | GCA_016822715.1 | PC | Public | 2017 | China | Env | 98 | 100 | 0.27 | 99.9936 | 5109567 | 45.27 | 77 | 259608 |
| SAMN16783241-GCA_016822695.1 | VP344 | SAMN16783241 | GCA_016822695.1 | NonPC | Public | 2016 | China | Env | 91.8 | 100 | 0.46 | 98.2137 | 5060094 | 45.2 | 58 | 450171 |
| SAMN16783242-GCA_016822665.1 | VP345 | SAMN16783242 | GCA_016822665.1 | NonPC | Public | 2013 | China | Clinical | 92.3 | 100 | 0.05 | 98.3257 | 5007976 | 45.26 | 53 | 450167 |
| SAMN16783243-GCA_016822625.1 | VP346 | SAMN16783243 | GCA_016822625.1 | NonPC | Public | 2013 | China | Clinical | 92.1 | 100 | 0.19 | 98.3719 | 5015928 | 45.23 | 61 | 450169 |
| SAMN16783244-GCA_016822655.1 | VP347 | SAMN16783244 | GCA_016822655.1 | NonPC | Public | 2013 | China | Clinical | 88.5 | 100 | 0.19 | 98.3674 | 4965261 | 45.27 | 56 | 313257 |
| SAMN16783245-GCA_016822615.1 | VP348 | SAMN16783245 | GCA_016822615.1 | NonPC | Public | 2017 | China | Env | 92.1 | 100 | 0.19 | 98.3369 | 5013266 | 45.24 | 62 | 313961 |
| SAMN16783246-GCA_016822595.1 | VP349 | SAMN16783246 | GCA_016822595.1 | PC | Public | 2016 | China | Env | 97.5 | 100 | 0 | 99.9019 | 5140350 | 45.12 | 69 | 259608 |
| SAMN16783247-GCA_016822575.1 | VP351 | SAMN16783247 | GCA_016822575.1 | NonPC | Public | 2016 | China | Env | 91.8 | 100 | 0 | 98.3804 | 5062986 | 45.27 | 89 | 330050 |
| SAMN16783248-GCA_016822535.1 | VP352 | SAMN16783248 | GCA_016822535.1 | NonPC | Public | 2008 | China | Env | 91.4 | 100 | 0 | 98.3203 | 5104514 | 45.17 | 47 | 343780 |
| SAMN16783249-GCA_016822515.1 | VP353 | SAMN16783249 | GCA_016822515.1 | NonPC | Public | 2008 | China | Clinical | 89 | 100 | 0.05 | 98.3378 | 5029118 | 45.38 | 47 | 464930 |
| SAMN16783250-GCA_016822545.1 | VP354 | SAMN16783250 | GCA_016822545.1 | PC | Public | 2008 | China | Env | 98.2 | 100 | 0 | 99.9928 | 5085353 | 45.27 | 75 | 259603 |
| SAMN16783251-GCA_016822495.1 | VP355 | SAMN16783251 | GCA_016822495.1 | PC | Public | 2013 | China | Env | 97.5 | 100 | 0 | 99.9321 | 5113820 | 45.28 | 81 | 243063 |
| SAMN16783253-GCA_016822455.1 | VP357 | SAMN16783253 | GCA_016822455.1 | PC | Public | 2008 | China | Env | 98.3 | 100 | 0 | 99.9922 | 5124973 | 45.25 | 80 | 259608 |
| SAMN16783254-GCA_016822395.1 | VP358 | SAMN16783254 | GCA_016822395.1 | NonPC | Public | 2009 | China | Env | 91.5 | 100 | 0.12 | 98.3272 | 4986875 | 45.31 | 40 | 424450 |
| SAMN16783255-GCA_016822415.1 | VP359 | SAMN16783255 | GCA_016822415.1 | NonPC | Public | 2013 | China | Env | 92.5 | 100 | 0 | 98.3368 | 5071845 | 45.23 | 58 | 322729 |
| SAMN16783256-GCA_016822425.1 | VP360 | SAMN16783256 | GCA_016822425.1 | NonPC | Public | 2014 | China | Clinical | 91.4 | 100 | 0.2 | 98.3131 | 5068802 | 45.3 | 61 | 335238 |
| SAMN16783257-GCA_016822375.1 | VP361 | SAMN16783257 | GCA_016822375.1 | NonPC | Public | 2009 | China | Env | 90.7 | 100 | 0 | 98.3453 | 5105681 | 45.29 | 55 | 401722 |
| SAMN16783258-GCA_016822355.1 | VP362 | SAMN16783258 | GCA_016822355.1 | NonPC | Public | 2009 | China | Clinical | 90.7 | 100 | 0 | 98.3464 | 5109175 | 45.27 | 72 | 313947 |
| SAMN16783259-GCA_016822335.1 | VP363 | SAMN16783259 | GCA_016822335.1 | PC | Public | 2015 | China | Env | 97.9 | 100 | 0 | 99.9917 | 5160184 | 45.19 | 78 | 259608 |

| | | | | | | | | | | | | | | | | |
|---|---|---|---|---|---|---|---|---|---|---|---|---|---|---|---|---|
| SAMN16783260-GCA_016822295.1 | VP364 | SAMN16783260 | GCA_016822295.1 | NonPC | Public | 2009 | China | Env | 88.2 | 100 | 0.36 | 98.2373 | 5007329 | 45.32 | 58 | 362511 |
| SAMN16783261-GCA_016822305.1 | VP365 | SAMN16783261 | GCA_016822305.1 | PC | Public | 2015 | China | Clinical | 97.6 | 100 | 0 | 99.962 | 5122822 | 45.27 | 68 | 244119 |
| SAMN16783262-GCA_016822235.1 | VP366 | SAMN16783262 | GCA_016822235.1 | NonPC | Public | 2009 | China | Env | 90.9 | 100 | 0 | 98.2879 | 5124469 | 45.19 | 61 | 314931 |
| SAMN16783263-GCA_016822275.1 | VP367 | SAMN16783263 | GCA_016822275.1 | NonPC | Public | 2009 | China | Env | 87.6 | 100 | 0.14 | 98.3827 | 4967719 | 45.37 | 103 | 91584 |
| SAMN16783264-GCA_016822255.1 | VP368 | SAMN16783264 | GCA_016822255.1 | NonPC | Public | 2009 | China | Clinical | 92.3 | 100 | 0.69 | 98.3088 | 5614453 | 44.81 | 161 | 348187 |
| SAMN16783265-GCA_016822215.1 | VP369 | SAMN16783265 | GCA_016822215.1 | PC | Public | 2009 | China | Env | 98.3 | 100 | 0 | 99.9888 | 5125520 | 45.26 | 75 | 259608 |
| SAMN16783266-GCA_016822195.1 | VP370 | SAMN16783266 | GCA_016822195.1 | NonPC | Public | 2009 | China | Env | 90.4 | 100 | 0.27 | 98.3644 | 4999190 | 45.35 | 35 | 442424 |
| SAMN16783267-GCA_016822135.1 | VP371 | SAMN16783267 | GCA_016822135.1 | NonPC | Public | 2010 | China | Clinical | 90.9 | 100 | 0 | 98.4017 | 5122743 | 45.24 | 88 | 367757 |
| SAMN16783268-GCA_016822155.1 | VP372 | SAMN16783268 | GCA_016822155.1 | PC | Public | 2010 | China | Clinical | 98.1 | 100 | 0 | 99.9928 | 5066921 | 45.27 | 80 | 259608 |
| SAMN16783269-GCA_016822145.1 | VP373 | SAMN16783269 | GCA_016822145.1 | PC | Public | 2010 | China | Clinical | 98.3 | 100 | 0 | 99.9897 | 5130937 | 45.25 | 78 | 259608 |
| SAMN16783270-GCA_016822115.1 | VP374 | SAMN16783270 | GCA_016822115.1 | PC | Public | 2015 | China | Env | 97.6 | 100 | 0 | 99.9609 | 5141052 | 45.25 | 76 | 244119 |
| SAMN16783271-GCA_016822095.1 | VP375 | SAMN16783271 | GCA_016822095.1 | NonPC | Public | 2010 | China | Clinical | 90.9 | 100 | 0.3 | 98.322 | 5401298 | 45.15 | 105 | 266970 |
| SAMN16783273-GCA_016822075.1 | VP377 | SAMN16783273 | GCA_016822075.1 | NonPC | Public | 2015 | China | Env | 92.1 | 100 | 0.19 | 98.3495 | 5014683 | 45.24 | 59 | 367931 |
| SAMN16783274-GCA_016822035.1 | VP379 | SAMN16783274 | GCA_016822035.1 | NonPC | Public | 2015 | China | Clinical | 91.8 | 100 | 0.46 | 98.2274 | 5053230 | 45.21 | 59 | 439343 |
| SAMN16783275-GCA_016822015.1 | VP380 | SAMN16783275 | GCA_016822015.1 | NonPC | Public | 2012 | China | Env | 92.3 | 100 | 0.05 | 98.3195 | 5108957 | 45.25 | 56 | 450167 |
| SAMN16783276-GCA_016821955.1 | VP381 | SAMN16783276 | GCA_016821955.1 | NonPC | Public | 2009 | China | Clinical | 91.6 | 100 | 0 | 98.4199 | 5057085 | 45.25 | 47 | 417275 |
| SAMN16783277-GCA_016821995.1 | VP382 | SAMN16783277 | GCA_016821995.1 | NonPC | Public | 2008 | China | Env | 90 | 100 | 0.14 | 98.312 | 5032888 | 45.33 | 48 | 428858 |
| SAMN16783278-GCA_016821965.1 | VP383 | SAMN16783278 | GCA_016821965.1 | NonPC | Public | 2016 | China | Env | 91.8 | 100 | 0.46 | 98.2495 | 5058026 | 45.2 | 60 | 439342 |
| SAMN16783279-GCA_016821935.1 | VP384 | SAMN16783279 | GCA_016821935.1 | PC | Public | 2013 | China | Env | 98.1 | 100 | 0 | 99.9924 | 5067092 | 45.27 | 74 | 259608 |
| SAMN16783280-GCA_016821855.1 | VP385 | SAMN16783280 | GCA_016821855.1 | NonPC | Public | 2015 | China | Env | 92.9 | 100 | 0.19 | 98.3594 | 5066191 | 45.28 | 55 | 419292 |
| SAMN16783281-GCA_016821875.1 | VP386 | SAMN16783281 | GCA_016821875.1 | NonPC | Public | 2010 | China | Env | 91.3 | 100 | 0.27 | 98.3854 | 5048905 | 45.31 | 58 | 259488 |
| SAMN16783282-GCA_016821915.1 | VP387 | SAMN16783282 | GCA_016821915.1 | PC | Public | 2009 | China | Env | 97.4 | 100 | 0 | 99.9182 | 5141642 | 45.25 | 81 | 259602 |
| SAMN16783283-GCA_016821865.1 | VP388 | SAMN16783283 | GCA_016821865.1 | NonPC | Public | 2007 | China | Env | 91.9 | 100 | 0.03 | 98.3554 | 5254640 | 45.14 | 107 | 169361 |
| SAMN16783284-GCA_016821835.1 | VP389 | SAMN16783284 | GCA_016821835.1 | NonPC | Public | 2011 | China | Env | 91 | 100 | 0 | 98.3429 | 5149670 | 45.21 | 80 | 309994 |
| SAMN16783285-GCA_016821755.1 | VP390 | SAMN16783285 | GCA_016821755.1 | PC | Public | 2011 | China | Env | 98.3 | 100 | 0 | 99.9929 | 5125200 | 45.25 | 74 | 259608 |
| SAMN16783286-GCA_016821775.1 | VP391 | SAMN16783286 | GCA_016821775.1 | PC | Public | 2007 | China | Env | 97.5 | 100 | 0 | 99.9888 | 5046396 | 45.3 | 77 | 259608 |
| SAMN16783287-GCA_016821815.1 | VP392 | SAMN16783287 | GCA_016821815.1 | PC | Public | 2007 | China | Env | 98.1 | 100 | 0 | 99.9918 | 5080322 | 45.27 | 78 | 259608 |
| SAMN16783288-GCA_016821765.1 | VP393 | SAMN16783288 | GCA_016821765.1 | NonPC | Public | 2008 | China | Env | 89.2 | 100 | 0 | 98.2761 | 4911706 | 45.43 | 47 | 305437 |
| SAMN16783289-GCA_016821735.1 | VP394 | SAMN16783289 | GCA_016821735.1 | PC | Public | 2008 | China | Clinical | 98.3 | 100 | 0 | 99.9913 | 5123570 | 45.25 | 76 | 259608 |
| SAMN16783290-GCA_016821715.1 | VP395 | SAMN16783290 | GCA_016821715.1 | PC | Public | 2009 | China | Clinical | 98.2 | 100 | 0 | 99.9901 | 5124736 | 45.25 | 68 | 259608 |
| SAMN16783291-GCA_016821655.1 | VP396 | SAMN16783291 | GCA_016821655.1 | PC | Public | 2016 | China | Env | 97.8 | 100 | 0 | 99.9874 | 5082624 | 45.3 | 66 | 259608 |
| SAMN16783292-GCA_016821685.1 | VP397 | SAMN16783292 | GCA_016821685.1 | NonPC | Public | 2011 | China | Env | 92.7 | 100 | 0 | 98.3796 | 5117937 | 45.21 | 51 | 460750 |
| SAMN16783293-GCA_016821665.1 | VP398 | SAMN16783293 | GCA_016821665.1 | PC | Public | 2016 | China | Env | 98 | 100 | 0 | 99.9859 | 5065832 | 45.27 | 75 | 259602 |
| SAMN16783294-GCA_016821635.1 | VP399 | SAMN16783294 | GCA_016821635.1 | PC | Public | 2016 | China | Clinical | 97.3 | 100 | 0.01 | 99.9116 | 5222606 | 45.26 | 70 | 259753 |
| SAMN16783295-GCA_016821615.1 | VP400 | SAMN16783295 | GCA_016821615.1 | NonPC | Public | 2012 | China | Env | 90.7 | 100 | 0 | 98.2821 | 5114772 | 45.28 | 65 | 404811 |
| SAMN16783296-GCA_016821595.1 | VP401 | SAMN16783296 | GCA_016821595.1 | PC | Public | 2012 | China | Env | 98.1 | 100 | 0.14 | 99.9845 | 5104329 | 45.25 | 72 | 259608 |
| SAMN16783297-GCA_016821575.1 | VP402 | SAMN16783297 | GCA_016821575.1 | PC | Public | 2012 | China | Clinical | 98.1 | 100 | 0 | 99.9917 | 5065311 | 45.27 | 76 | 259608 |
| SAMN16783298-GCA_016821555.1 | VP403 | SAMN16783298 | GCA_016821555.1 | PC | Public | 2012 | China | Env | 98 | 100 | 0 | 99.9921 | 5066067 | 45.27 | 74 | 259608 |
| SAMN16783299-GCA_016821535.1 | VP404 | SAMN16783299 | GCA_016821535.1 | PC | Public | 2012 | China | Clinical | 98 | 100 | 0 | 99.9773 | 5118113 | 45.26 | 76 | 259608 |
| SAMN16783300-GCA_016821515.1 | VP405 | SAMN16783300 | GCA_016821515.1 | PC | Public | 2017 | China | Env | 98.3 | 100 | 0 | 99.9896 | 5124719 | 45.25 | 76 | 259608 |
| SAMN16783301-GCA_016821475.1 | VP406 | SAMN16783301 | GCA_016821475.1 | NonPC | Public | 2011 | China | Env | 90.4 | 100 | 0.14 | 98.292 | 5074323 | 45.24 | 70 | 314915 |
| SAMN16783302-GCA_016821495.1 | VP408 | SAMN16783302 | GCA_016821495.1 | NonPC | Public | 2015 | China | Env | 93.1 | 100 | 0.19 | 98.3792 | 5147250 | 45.11 | 58 | 422538 |
| SAMN16783303-GCA_016821455.1 | VP409 | SAMN16783303 | GCA_016821455.1 | NonPC | Public | 2015 | China | Env | 92.9 | 100 | 0.05 | 98.3341 | 5049037 | 45.3 | 46 | 463453 |
| SAMN16783304-GCA_016821435.1 | VP410 | SAMN16783304 | GCA_016821435.1 | NonPC | Public | 2015 | China | Env | 91.8 | 100 | 0.46 | 98.2601 | 5052098 | 45.21 | 58 | 439457 |
| SAMN16783305-GCA_016821395.1 | VP411 | SAMN16783305 | GCA_016821395.1 | NonPC | Public | 2015 | China | Env | 92.9 | 100 | 0.05 | 98.346 | 5043737 | 45.32 | 50 | 417597 |
| SAMN16783306-GCA_016821375.1 | VP412 | SAMN16783306 | GCA_016821375.1 | NonPC | Public | 2015 | China | Env | 92.9 | 100 | 0.19 | 98.3806 | 5070655 | 45.28 | 54 | 419294 |
| SAMN16783307-GCA_016821415.1 | VP413 | SAMN16783307 | GCA_016821415.1 | NonPC | Public | 2015 | China | Env | 92.9 | 100 | 0.05 | 98.3392 | 5126002 | 45.14 | 48 | 463452 |
| SAMN16783308-GCA_016821355.1 | VP414 | SAMN16783308 | GCA_016821355.1 | NonPC | Public | 2015 | China | Env | 92.9 | 100 | 0.19 | 98.3847 | 5086777 | 45.27 | 50 | 419294 |

| | | | | | | | | | | | | | | | | |
|---|---|---|---|---|---|---|---|---|---|---|---|---|---|---|---|---|
| SAMN16783309-GCA_016821335.1 | VP415 | SAMN16783309 | GCA_016821335.1 | NonPC | Public | 2015 | China | Env | 92.9 | 100 | 0.05 | 98.3916 | 5050890 | 45.3 | 47 | 463453 |
| SAMN16783310-GCA_016821275.1 | VP416 | SAMN16783310 | GCA_016821275.1 | NonPC | Public | 2017 | China | Clinical | 88.7 | 100 | 0.05 | 98.3216 | 5007558 | 45.18 | 68 | 313257 |
| SAMN16783311-GCA_016821285.1 | VP417 | SAMN16783311 | GCA_016821285.1 | NonPC | Public | 2016 | China | Env | 91.8 | 100 | 0.46 | 98.2346 | 5057956 | 45.2 | 58 | 439571 |
| SAMN16783312-GCA_016821315.1 | VP418 | SAMN16783312 | GCA_016821315.1 | NonPC | Public | 2016 | China | Env | 91.8 | 100 | 0.46 | 98.243 | 5058798 | 45.2 | 56 | 439457 |
| SAMN16783313-GCA_016821255.1 | VP419 | SAMN16783313 | GCA_016821255.1 | NonPC | Public | 2016 | China | Env | 92.9 | 100 | 0.05 | 98.3427 | 5081307 | 45.28 | 51 | 419294 |
| SAMN16783314-GCA_016821235.1 | VP420 | SAMN16783314 | GCA_016821235.1 | NonPC | Public | 2016 | China | Env | 91.8 | 100 | 0.46 | 98.2711 | 5059547 | 45.2 | 58 | 313824 |
| SAMN16783315-GCA_016821195.1 | VP421 | SAMN16783315 | GCA_016821195.1 | NonPC | Public | 2017 | China | Env | 91.4 | 100 | 0 | 98.3039 | 5033643 | 45.34 | 65 | 403731 |
| SAMN16783316-GCA_016821215.1 | VP422 | SAMN16783316 | GCA_016821215.1 | NonPC | Public | 2016 | China | Env | 91.8 | 100 | 0.46 | 98.24 | 5059001 | 45.2 | 59 | 439347 |
| SAMN16783317-GCA_016821155.1 | VP423 | SAMN16783317 | GCA_016821155.1 | NonPC | Public | 2016 | China | Env | 91.8 | 100 | 0.46 | 98.2615 | 5058274 | 45.2 | 57 | 439347 |
| SAMN16783318-GCA_016821175.1 | VP424 | SAMN16783318 | GCA_016821175.1 | NonPC | Public | 2016 | China | Env | 91.8 | 100 | 0.46 | 98.2417 | 5057727 | 45.2 | 58 | 439343 |
| SAMN16783319-GCA_016821095.1 | VP425 | SAMN16783319 | GCA_016821095.1 | NonPC | Public | 2016 | China | Env | 92.9 | 100 | 0.05 | 98.3764 | 5049836 | 45.3 | 50 | 417596 |
| SAMN16783320-GCA_016821115.1 | VP426 | SAMN16783320 | GCA_016821115.1 | NonPC | Public | 2016 | China | Env | 92.9 | 100 | 0.19 | 98.3542 | 5084010 | 45.27 | 52 | 417597 |
| SAMN16783321-GCA_016821135.1 | VP427 | SAMN16783321 | GCA_016821135.1 | NonPC | Public | 2016 | China | Env | 93 | 100 | 0.05 | 98.3663 | 5071679 | 45.28 | 55 | 417597 |
| SAMN16783322-GCA_016821065.1 | VP428 | SAMN16783322 | GCA_016821065.1 | NonPC | Public | 2017 | China | Clinical | 92.3 | 100 | 0.05 | 98.3746 | 5114495 | 45.08 | 53 | 372927 |
| SAMN16783323-GCA_016821055.1 | VP429 | SAMN16783323 | GCA_016821055.1 | NonPC | Public | 2017 | China | Env | 90.7 | 100 | 0.33 | 98.2895 | 5155098 | 45.12 | 77 | 238193 |
| SAMN16783324-GCA_016821025.1 | VP430 | SAMN16783324 | GCA_016821025.1 | NonPC | Public | 2017 | China | Env | 91.8 | 100 | 0.46 | 98.2482 | 5060054 | 45.2 | 59 | 439582 |
| SAMN16783325-GCA_016820995.1 | VP431 | SAMN16783325 | GCA_016820995.1 | NonPC | Public | 2017 | China | Env | 91.8 | 100 | 0.46 | 98.277 | 5059631 | 45.2 | 59 | 439582 |
| SAMN16783326-GCA_016821015.1 | VP432 | SAMN16783326 | GCA_016821015.1 | NonPC | Public | 2017 | China | Env | 91.8 | 100 | 0.19 | 98.253 | 5046286 | 45.22 | 57 | 438399 |
| SAMN16783327-GCA_016820965.1 | VP433 | SAMN16783327 | GCA_016820965.1 | NonPC | Public | 2008 | China | Env | 90.7 | 100 | 0.27 | 98.3761 | 5000075 | 45.32 | 62 | 312541 |
| SAMN16783328-GCA_016820955.1 | VP434 | SAMN16783328 | GCA_016820955.1 | NonPC | Public | 2007 | China | Clinical | 89.6 | 100 | 0 | 98.2395 | 4984904 | 45.34 | 64 | 380935 |
| SAMN16783329-GCA_016820895.1 | VP435 | SAMN16783329 | GCA_016820895.1 | NonPC | Public | 2017 | China | Env | 91.8 | 100 | 0.46 | 98.2422 | 5057655 | 45.2 | 59 | 450168 |
| SAMN16783330-GCA_016820935.1 | VP436 | SAMN16783330 | GCA_016820935.1 | NonPC | Public | 2017 | China | Env | 92.8 | 100 | 0.19 | 98.4094 | 5062784 | 45.28 | 58 | 410040 |
| SAMN16783331-GCA_016820905.1 | VP437 | SAMN16783331 | GCA_016820905.1 | NonPC | Public | 2017 | China | Env | 91.8 | 100 | 0.74 | 98.2565 | 5101983 | 45.2 | 60 | 439347 |
| SAMN16783332-GCA_016820865.1 | VP438 | SAMN16783332 | GCA_016820865.1 | NonPC | Public | 2017 | China | Env | 91.8 | 100 | 0.19 | 98.2407 | 5049912 | 45.21 | 56 | 439343 |
| SAMN16783333-GCA_016820855.1 | VP439 | SAMN16783333 | GCA_016820855.1 | NonPC | Public | 2009 | China | Clinical | 92.8 | 100 | 0 | 98.3317 | 5236240 | 45.18 | 43 | 531317 |
| SAMN16783334-GCA_016820835.1 | VP440 | SAMN16783334 | GCA_016820835.1 | PC | Public | 2017 | China | Env | 98 | 100 | 0 | 99.9906 | 5064197 | 45.27 | 74 | 259608 |
| SAMN16783335-GCA_016820815.1 | VP441 | SAMN16783335 | GCA_016820815.1 | PC | Public | 2017 | China | Env | 97.7 | 100 | 0 | 99.9916 | 5059443 | 45.27 | 79 | 259608 |
| SAMN16783336-GCA_016820765.1 | VP442 | SAMN16783336 | GCA_016820765.1 | NonPC | Public | 2017 | China | Clinical | 92.1 | 100 | 0.19 | 98.3433 | 5012783 | 45.24 | 59 | 367817 |
| SAMN16783337-GCA_016820795.1 | VP443 | SAMN16783337 | GCA_016820795.1 | PC | Public | 2017 | China | Env | 97.2 | 100 | 0 | 99.8791 | 5184770 | 45.24 | 71 | 267671 |
| SAMN16783338-GCA_016820755.1 | VP444 | SAMN16783338 | GCA_016820755.1 | PC | Public | 2017 | China | Clinical | 97.5 | 100 | 0 | 99.9247 | 5141384 | 45.12 | 70 | 259608 |
| SAMN16783339-GCA_016820735.1 | VP445 | SAMN16783339 | GCA_016820735.1 | NonPC | Public | 2017 | China | Env | 92.9 | 100 | 0.19 | 98.3403 | 5068417 | 45.28 | 56 | 417596 |
| SAMN16783340-GCA_016819295.1 | VP446 | SAMN16783340 | GCA_016819295.1 | NonPC | Public | 2017 | China | Env | 90.7 | 100 | 0.33 | 98.3128 | 5062883 | 45.23 | 59 | 246957 |
| SAMN16783341-GCA_016819275.1 | VP1 | SAMN16783341 | GCA_016819275.1 | NonPC | Public | 2008 | China | Clinical | 92.6 | 100 | 0 | 98.4338 | 5121533 | 45.18 | 52 | 417283 |
| SAMN16783342-GCA_016820715.1 | VP3 | SAMN16783342 | GCA_016820715.1 | NonPC | Public | 2008 | China | Env | 90.7 | 100 | 0.72 | 98.3191 | 5281912 | 45.19 | 59 | 202148 |
| SAMN16783343-GCA_016820695.1 | VP4 | SAMN16783343 | GCA_016820695.1 | NonPC | Public | 2008 | China | Env | 91.1 | 100 | 0 | 98.3538 | 5007167 | 45.41 | 62 | 232703 |
| SAMN16783344-GCA_016820675.1 | VP6 | SAMN16783344 | GCA_016820675.1 | NonPC | Public | 2008 | China | Env | 91 | 100 | 0.03 | 98.4235 | 5015531 | 45.31 | 38 | 400733 |
| SAMN16783345-GCA_016819245.1 | VP8 | SAMN16783345 | GCA_016819245.1 | PC | Public | 2016 | China | Env | 97.4 | 100 | 0 | 99.9098 | 5090345 | 45.26 | 74 | 266233 |
| SAMN16783346-GCA_016819215.1 | VP12 | SAMN16783346 | GCA_016819215.1 | NonPC | Public | 2009 | China | Env | 89.4 | 100 | 0 | 98.379 | 5063917 | 45.22 | 53 | 340451 |
| SAMN16783347-GCA_016819235.1 | VP16 | SAMN16783347 | GCA_016819235.1 | PC | Public | 2015 | China | Env | 97.4 | 100 | 0 | 99.8954 | 5089321 | 45.27 | 83 | 259602 |
| SAMN16783348-GCA_016819175.1 | VP23 | SAMN16783348 | GCA_016819175.1 | PC | Public | 2015 | China | Env | 97.2 | 100 | 0 | 99.9063 | 5086972 | 45.27 | 75 | 258341 |
| SAMN16783349-GCA_016819195.1 | VP24 | SAMN16783349 | GCA_016819195.1 | PC | Public | 2015 | China | Env | 97.4 | 100 | 0 | 99.8896 | 5095234 | 45.26 | 75 | 266229 |
| SAMN16783350-GCA_016819145.1 | VP30 | SAMN16783350 | GCA_016819145.1 | NonPC | Public | 2010 | China | Env | 88 | 100 | 0.53 | 98.2512 | 5190028 | 45.17 | 115 | 240180 |
| SAMN16783351-GCA_016819135.1 | VP32 | SAMN16783351 | GCA_016819135.1 | NonPC | Public | 2012 | China | Env | 89.9 | 100 | 0 | 98.4171 | 4974343 | 45.36 | 69 | 314834 |
| SAMN16783352-GCA_016819115.1 | VP33 | SAMN16783352 | GCA_016819115.1 | NonPC | Public | 2011 | China | Clinical | 91.1 | 100 | 0 | 98.3638 | 5080542 | 45.3 | 59 | 401622 |
| SAMN16783353-GCA_016819095.1 | VP35 | SAMN16783353 | GCA_016819095.1 | NonPC | Public | 2008 | China | Env | 91.5 | 100 | 0 | 98.2964 | 5122468 | 45.19 | 60 | 317357 |
| SAMN16783354-GCA_016819075.1 | VP42 | SAMN16783354 | GCA_016819075.1 | NonPC | Public | 2009 | China | Clinical | 89.5 | 100 | 0.05 | 98.3463 | 4959292 | 45.4 | 45 | 419782 |
| SAMN16783355-GCA_016819035.1 | VP59 | SAMN16783355 | GCA_016819035.1 | NonPC | Public | 2007 | China | Env | 91.4 | 100 | 0 | 98.3355 | 4977838 | 45.35 | 59 | 332920 |
| SAMN16783356-GCA_016819045.1 | VP65 | SAMN16783356 | GCA_016819045.1 | PC | Public | 2013 | China | Env | 97.4 | 100 | 0 | 99.903 | 5077298 | 45.28 | 73 | 259608 |

| | | | | | | | | | | | | | | | | |
|---|---|---|---|---|---|---|---|---|---|---|---|---|---|---|---|---|
| SAMN16783357-GCA_016819015.1 | VP67 | SAMN16783357 | GCA_016819015.1 | NonPC | Public | 2013 | China | Clinical | 91.9 | 100 | 1.05 | 98.2037 | 5403556 | 44.99 | 166 | 231792 |
| SAMN16783358-GCA_016818995.1 | VP72 | SAMN16783358 | GCA_016818995.1 | PC | Public | 2017 | China | Env | 97.2 | 100 | 0 | 99.8998 | 5087030 | 45.27 | 77 | 257863 |
| SAMN16783359-GCA_016818955.1 | VP94 | SAMN16783359 | GCA_016818955.1 | NonPC | Public | 2009 | China | Clinical | 92.1 | 100 | 0.06 | 98.4324 | 5153010 | 45.26 | 46 | 451197 |
| SAMN16783360-GCA_016818975.1 | VP97 | SAMN16783360 | GCA_016818975.1 | NonPC | Public | 2009 | China | Env | 91.5 | 100 | 0.27 | 98.4609 | 5084574 | 45.08 | 62 | 314780 |
| SAMN16783361-GCA_016818935.1 | VP99 | SAMN16783361 | GCA_016818935.1 | NonPC | Public | 2010 | China | Env | 89.6 | 100 | 0 | 98.2325 | 5006763 | 45.3 | 44 | 375768 |
| SAMN16783362-GCA_016818915.1 | VP103 | SAMN16783362 | GCA_016818915.1 | PC | Public | 2010 | China | Clinical | 97.3 | 100 | 0 | 99.9173 | 5146939 | 45.28 | 69 | 259753 |
| SAMN16783363-GCA_016818885.1 | VP113 | SAMN16783363 | GCA_016818885.1 | NonPC | Public | 2008 | China | Clinical | 92.2 | 100 | 0.05 | 98.07 | 5116920 | 45.23 | 44 | 433146 |
| SAMN16783364-GCA_016818875.1 | VP169 | SAMN16783364 | GCA_016818875.1 | NonPC | Public | 2012 | China | Clinical | 92.9 | 100 | 0.05 | 98.3788 | 5031812 | 45.31 | 38 | 463453 |
| SAMN16783365-GCA_016818835.1 | VP177 | SAMN16783365 | GCA_016818835.1 | NonPC | Public | 2007 | China | Clinical | 91.7 | 100 | 0 | 98.3693 | 5140041 | 45.12 | 87 | 347280 |
| SAMN16783367-GCA_016818775.1 | VP192 | SAMN16783367 | GCA_016818775.1 | NonPC | Public | 2007 | China | Clinical | 89.5 | 100 | 0 | 98.2755 | 4983547 | 45.34 | 66 | 347728 |
| SAMN16783368-GCA_016818795.1 | VP195 | SAMN16783368 | GCA_016818795.1 | NonPC | Public | 2008 | China | Env | 91.8 | 100 | 0.45 | 98.3825 | 5058693 | 45.27 | 62 | 318128 |
| SAMN16783369-GCA_016818785.1 | VP196 | SAMN16783369 | GCA_016818785.1 | NonPC | Public | 2009 | China | Env | 91.9 | 100 | 0 | 98.2722 | 5148006 | 45.26 | 91 | 314826 |
| SAMN16783370-GCA_016818755.1 | VP200 | SAMN16783370 | GCA_016818755.1 | NonPC | Public | 2008 | China | Env | 91.1 | 100 | 0.1 | 98.3259 | 5129027 | 45.19 | 63 | 377214 |
| SAMN16783371-GCA_016818735.1 | VP202 | SAMN16783371 | GCA_016818735.1 | NonPC | Public | 2008 | China | Env | 91.1 | 100 | 0 | 98.3573 | 5081830 | 45.25 | 44 | 459051 |
| SAMN16783372-GCA_016818715.1 | VP203 | SAMN16783372 | GCA_016818715.1 | NonPC | Public | 2008 | China | Env | 90.9 | 100 | 0.05 | 98.3726 | 5135710 | 45.21 | 62 | 391001 |
| SAMN16783373-GCA_016818675.1 | VP205 | SAMN16783373 | GCA_016818675.1 | NonPC | Public | 2008 | China | Env | 90.8 | 100 | 0.34 | 98.3176 | 5144898 | 45.26 | 60 | 309625 |
| SAMN16783374-GCA_016818685.1 | VP206 | SAMN16783374 | GCA_016818685.1 | NonPC | Public | 2008 | China | Env | 90.9 | 99.91 | 0.09 | 98.2919 | 5252512 | 45.18 | 75 | 259059 |
| SAMN16783375-GCA_016818655.1 | VP208 | SAMN16783375 | GCA_016818655.1 | PC | Public | 2015 | China | Env | 97.4 | 100 | 0 | 99.9111 | 5090419 | 45.27 | 80 | 259602 |
| SAMN16783376-GCA_016818635.1 | VP229 | SAMN16783376 | GCA_016818635.1 | NonPC | Public | 2008 | China | Env | 91.2 | 100 | 0 | 98.354 | 5095973 | 45.25 | 71 | 315212 |
| SAMN16783377-GCA_016818595.1 | VP230 | SAMN16783377 | GCA_016818595.1 | PC | Public | 2012 | China | Env | 98 | 100 | 0 | 99.9902 | 5064365 | 45.27 | 76 | 259608 |
| SAMN16783378-GCA_016818605.1 | VP233 | SAMN16783378 | GCA_016818605.1 | NonPC | Public | 2008 | China | Env | 91.7 | 100 | 0 | 98.4126 | 5056888 | 45.25 | 48 | 417275 |
| SAMN16783379-GCA_016818535.1 | VP235 | SAMN16783379 | GCA_016818535.1 | PC | Public | 2007 | China | Clinical | 97.3 | 100 | 0 | 99.9251 | 5093206 | 45.27 | 80 | 259608 |
| SAMN16783380-GCA_016818545.1 | VP238 | SAMN16783380 | GCA_016818545.1 | NonPC | Public | 2010 | China | Env | 91 | 100 | 0 | 98.3806 | 5225526 | 45.19 | 68 | 407511 |
| SAMN16783381-GCA_016818575.1 | VP239 | SAMN16783381 | GCA_016818575.1 | NonPC | Public | 2009 | China | Env | 90.7 | 99.66 | 0.17 | 98.3002 | 5291864 | 45.06 | 94 | 305116 |
| SAMN16783382-GCA_016818495.1 | VP243 | SAMN16783382 | GCA_016818495.1 | NonPC | Public | 2010 | China | Env | 90.2 | 100 | 0 | 98.3323 | 5002780 | 45.36 | 91 | 384380 |
| SAMN16783383-GCA_016818505.1 | VP245 | SAMN16783383 | GCA_016818505.1 | NonPC | Public | 2008 | China | Env | 88.2 | 100 | 0.47 | 98.3073 | 4988493 | 45.23 | 82 | 416114 |
| SAMN16783384-GCA_016818435.1 | VP304 | SAMN16783384 | GCA_016818435.1 | NonPC | Public | 2008 | China | Clinical | 92.1 | 100 | 0.61 | 98.3574 | 5265899 | 45.25 | 59 | 445271 |
| SAMN16783385-GCA_016818455.1 | VP305 | SAMN16783385 | GCA_016818455.1 | NonPC | Public | 2008 | China | Clinical | 90.6 | 100 | 0.05 | 98.318 | 5066750 | 45.26 | 69 | 231278 |
| SAMN16783386-GCA_016818465.1 | VP319 | SAMN16783386 | GCA_016818465.1 | NonPC | Public | 2007 | China | Env | 89.2 | 100 | 0.27 | 98.3131 | 4958845 | 45.41 | 67 | 335202 |
| SAMN16783387-GCA_016818415.1 | VP331 | SAMN16783387 | GCA_016818415.1 | NonPC | Public | 2015 | China | Env | 92.1 | 100 | 0.19 | 98.355 | 5011747 | 45.24 | 59 | 367936 |
| SAMN16783388-GCA_016818335.1 | VP378 | SAMN16783388 | GCA_016818335.1 | NonPC | Public | 2012 | China | Env | 92.7 | 100 | 0 | 98.4269 | 5116211 | 45.21 | 51 | 460704 |
| SAMN16783389-GCA_016818365.1 | VP407 | SAMN16783389 | GCA_016818365.1 | PC | Public | 2017 | China | Clinical | 97.2 | 100 | 0 | 99.9058 | 5086748 | 45.27 | 79 | 257863 |
| SAMN16790655-GCA_015744885.1 | PNUSAV001426 | SAMN16790655 | GCA_015744885.1 | NonPC | Public | NA | USA | Clinical | 89.3 | 100 | 0.62 | 98.0759 | 5279213 | 45.27 | 133 | 98099 |
| SAMN16836862-GCA_018036505.1 | PNUSAV001449 | SAMN16836862 | GCA_018036505.1 | NonPC | Public | NA | USA | Clinical | 91.4 | 100 | 2.23 | 98.3103 | 5294241 | 45.28 | 42 | 719393 |
| SAMN16839181-GCA_018036425.1 | PNUSAV001432 | SAMN16839181 | GCA_018036425.1 | NonPC | Public | NA | USA | Clinical | 91.5 | 100 | 1.71 | 98.3247 | 5210506 | 45.25 | 43 | 718946 |
| SAMN16839183-GCA_018036405.1 | PNUSAV001450 | SAMN16839183 | GCA_018036405.1 | NonPC | Public | NA | USA | Clinical | 89.6 | 100 | 0.64 | 98.2429 | 5175321 | 45.27 | 177 | 65057 |
| SAMN16844329-GCF_016403045.1 | 81TDH2 | SAMN16844329 | GCF_016403045.1 | NonPC | Public | 2003 | India | Env | 89.5 | 100 | 0 | 98.3641 | 4879979 | 45.49 | 2 | 3083194 |
| SAMN16844529-GCF_016834555.1 | Diseased_Shrimp | SAMN16844529 | GCF_016834555.1 | NonPC | Public | 2017 | India | Env | 89.4 | 99.73 | 0.6 | 98.2626 | 5079924 | 45.32 | 2 | 3296982 |
| SAMN16844541-GCF_016132465.1 | Diseased_Shrimp_sample | SAMN16844541 | GCF_016132465.1 | NonPC | Public | 2017 | India | Env | 91.3 | 100 | 3.32 | 98.3787 | 5249063 | 45.36 | 4 | 3247960 |
| SAMN16844542-GCF_016132485.1 | Moribund_shrimp | SAMN16844542 | GCF_016132485.1 | NonPC | Public | 2017 | India | Env | 90.9 | 100 | 4.63 | 98.4065 | 5421490 | 45.29 | 6 | 3199074 |
| SAMN16954303-GCF_015779285.1 | 20-082A3 | SAMN16954303 | GCF_015779285.1 | NonPC | Public | 2020 | South_Korea | Env | 92.8 | 100 | 0.27 | 98.2973 | 5601613 | 45.3 | 4 | 3610915 |
| SAMN17041478-GCA_015908385.1 | PNUSAV001477 | SAMN17041478 | GCA_015908385.1 | NonPC | Public | 2020 | USA | Clinical | 89.7 | 100 | 2.27 | 98.3489 | 5213029 | 45.32 | 36 | 631994 |
| SAMN17041479-GCA_015910125.1 | PNUSAV001480 | SAMN17041479 | GCA_015910125.1 | NonPC | Public | 2020 | USA | Clinical | 90 | 100 | 1.36 | 98.2812 | 5239609 | 45.31 | 46 | 357525 |
| SAMN17041481-GCA_018033385.1 | PNUSAV001478 | SAMN17041481 | GCA_018033385.1 | PC | Public | 2020 | USA | Clinical | 98.2 | 100 | 0.41 | 99.9857 | 5207625 | 45.24 | 51 | 373379 |
| SAMN17041483-GCA_018033365.1 | PNUSAV001479 | SAMN17041483 | GCA_018033365.1 | NonPC | Public | 2020 | USA | Clinical | 91.3 | 100 | 0.4 | 98.2973 | 5262894 | 45.19 | 42 | 461018 |
| SAMN17078542-GCA_016146875.1 | 20MO11SH01-V2 | SAMN17078542 | GCA_016146875.1 | NonPC | Public | 2020 | USA | Env | 88.6 | 100 | 4.12 | 98.0163 | 5607453 | 45.19 | 78 | 190821 |
| SAMN17188270-SRR14684349 | HB12 | SAMN17188270 | SRR14684349 | NonPC | Public | 2017 | China | Env | 89.6 | 100 | 0.27 | 98.3217 | 4986533 | 45.35 | 43 | 400688 |
| SAMN17188271-SRR14684350 | HB28 | SAMN17188271 | SRR14684350 | NonPC | Public | 2017 | China | Env | 89.6 | 100 | 0.27 | 98.3143 | 4987119 | 45.35 | 47 | 400535 |

| | | | | | | | | | | | | | | | | |
|---|---|---|---|---|---|---|---|---|---|---|---|---|---|---|---|---|
| SAMN17188274-SRR14373906 | HB18 | SAMN17188274 | SRR14373906 | NonPC | Public | 2017 | China | Env | 89.6 | 100 | 0.27 | 98.3143 | 4987119 | 45.35 | 47 | 400535 |
| SAMN17188276-GCA_021729985.1 | LH80 | SAMN17188276 | GCA_021729985.1 | NonPC | Public | 2017 | China | Env | 91.4 | 100 | 0.19 | 98.2505 | 5361935 | 45.17 | 3 | 3422475 |
| SAMN17188276-SRR14373900 | LH80 | SAMN17188276 | SRR14373900 | NonPC | Public | 2017 | China | Env | 90.5 | 100 | 0.46 | 98.2647 | 5241151 | 45.15 | 87 | 275867 |
| SAMN17188277-SRR14373903 | LH36 | SAMN17188277 | SRR14373903 | NonPC | Public | 2017 | China | Env | 89.8 | 100 | 0 | 98.2972 | 5181114 | 45.19 | 39 | 462911 |
| SAMN17188280-SRR14373897 | NB118 | SAMN17188280 | SRR14373897 | NonPC | Public | 2015 | China | Env | 88.8 | 100 | 0 | 98.0266 | 5525597 | 44.98 | 120 | 143650 |
| SAMN17188281-SRR14684351 | NB80 | SAMN17188281 | SRR14684351 | NonPC | Public | 2017 | China | Env | 90.9 | 100 | 0.05 | 98.3818 | 5075563 | 45.26 | 41 | 447094 |
| SAMN17188283-SRR14373893 | NB189 | SAMN17188283 | SRR14373893 | NonPC | Public | 2017 | China | Env | 92.3 | 100 | 0.19 | 98.3744 | 5049138 | 45.25 | 43 | 471218 |
| SAMN17188288-SRR14373891 | NB315 | SAMN17188288 | SRR14373891 | NonPC | Public | 2017 | China | Env | 91.6 | 100 | 0.08 | 98.3281 | 5199505 | 45.19 | 40 | 366188 |
| SAMN17188289-SRR14373889 | NB327 | SAMN17188289 | SRR14373889 | PC | Public | 2017 | China | Env | 97.8 | 100 | 0 | 99.9444 | 5123309 | 45.29 | 49 | 399827 |
| SAMN17188290-SRR14684352 | NB102 | SAMN17188290 | SRR14684352 | NonPC | Public | 2017 | China | Env | 92 | 100 | 0.68 | 98.3205 | 5503242 | 45.22 | 111 | 371906 |
| SAMN17188294-GCA_021730045.1 | VP17 | SAMN17188294 | GCA_021730045.1 | NonPC | Public | 2013 | China | Env | 92.1 | 97.68 | 0 | 98.269 | 5648889 | 45.35 | 5 | 3567562 |
| SAMN17188296-GCA_021730085.1 | VP157 | SAMN17188296 | GCA_021730085.1 | NonPC | Public | 2015 | China | Env | 89.3 | 100 | 0.34 | 98.2138 | 5549882 | 45.23 | 6 | 3530958 |
| SAMN17188298-GCA_021730065.1 | VP120 | SAMN17188298 | GCA_021730065.1 | NonPC | Public | 2015 | China | Env | 91.5 | 100 | 0.05 | 98.3652 | 5209822 | 45.34 | 3 | 3270162 |
| SAMN17188300-SRR14373885 | TJ-107 | SAMN17188300 | SRR14373885 | NonPC | Public | 2016 | China | Env | 92 | 100 | 0 | 98.2919 | 5567698 | 45.27 | 77 | 387759 |
| SAMN17188301-SRR14373886 | TJ-102 | SAMN17188301 | SRR14373886 | NonPC | Public | 2016 | China | Env | 91.3 | 100 | 0.61 | 98.2899 | 5374858 | 45.27 | 82 | 219025 |
| SAMN17188302-GCA_021730005.1 | TJ-20 | SAMN17188302 | GCA_021730005.1 | NonPC | Public | 2016 | China | Env | 91.5 | 100 | 0.05 | 98.3734 | 5213611 | 45.34 | 3 | 3270162 |
| SAMN17188303-SRR14373884 | TJ-108 | SAMN17188303 | SRR14373884 | NonPC | Public | 2016 | China | Env | 88.3 | 100 | 0.34 | 98.1869 | 5489180 | 45.18 | 57 | 533828 |
| SAMN17188304-SRR14373882 | TJ-124 | SAMN17188304 | SRR14373882 | NonPC | Public | 2016 | China | Env | 91.3 | 100 | 0.61 | 98.3049 | 5364908 | 45.27 | 83 | 300879 |
| SAMN17188305-SRR14373880 | TJ-139 | SAMN17188305 | SRR14373880 | NonPC | Public | 2016 | China | Env | 89.7 | 100 | 0.18 | 98.3216 | 5087796 | 45.29 | 45 | 550917 |
| SAMN17188306-SRR14373881 | TJ-125 | SAMN17188306 | SRR14373881 | NonPC | Public | 2016 | China | Env | 90.9 | 100 | 0.4 | 98.2537 | 5318696 | 45.16 | 84 | 465767 |
| SAMN17188307-SRR14373879 | TJ-167 | SAMN17188307 | SRR14373879 | NonPC | Public | 2017 | China | Env | 89.9 | 99.73 | 0.33 | 98.2859 | 5309316 | 45.33 | 24 | 1054606 |
| SAMN17188308-SRR14373878 | TJ-183 | SAMN17188308 | SRR14373878 | NonPC | Public | 2017 | China | Env | 90.1 | 100 | 0.19 | 98.2665 | 5239474 | 45.24 | 45 | 448892 |
| SAMN17188309-GCA_021730025.1 | TJ-187 | SAMN17188309 | GCA_021730025.1 | NonPC | Public | 2017 | China | Env | 92.3 | 99.91 | 0.61 | 98.2624 | 5466908 | 45.31 | 5 | 3354580 |
| SAMN17188310-SRR14373904 | LH3 | SAMN17188310 | SRR14373904 | NonPC | Public | 2014 | China | Env | 91.3 | 100 | 0.62 | 98.3045 | 5371706 | 45.26 | 91 | 491588 |
| SAMN17188311-SRR14373873 | LH11 | SAMN17188311 | SRR14373873 | NonPC | Public | 2015 | China | Env | 91.7 | 100 | 0.54 | 98.3265 | 5372064 | 45.14 | 57 | 329020 |
| SAMN17188313-SRR14373902 | LH4 | SAMN17188313 | SRR14373902 | NonPC | Public | 2015 | China | Env | 91.3 | 100 | 0.62 | 98.3263 | 5371196 | 45.25 | 92 | 491588 |
| SAMN17188316-GCA_021729965.1 | LH24 | SAMN17188316 | GCA_021729965.1 | NonPC | Public | 2017 | China | Env | 92.3 | 99.91 | 0.61 | 98.2623 | 5447540 | 45.35 | 5 | 3354580 |
| SAMN17188317-SRR14373874 | LH10 | SAMN17188317 | SRR14373874 | NonPC | Public | 2015 | China | Env | 91.1 | 100 | 0 | 98.2683 | 5324047 | 45.23 | 47 | 584936 |
| SAMN17188319-SRR14373872 | LH19-6 | SAMN17188319 | SRR14373872 | NonPC | Public | 2015 | China | Env | 91.4 | 100 | 0.13 | 98.3575 | 5283456 | 45.23 | 33 | 481789 |
| SAMN17188320-SRR14373901 | LH6 | SAMN17188320 | SRR14373901 | NonPC | Public | 2015 | China | Env | 91.3 | 100 | 0.62 | 98.3459 | 5371783 | 45.26 | 96 | 458368 |
| SAMN17188321-SRR14373871 | LH19-8 | SAMN17188321 | SRR14373871 | NonPC | Public | 2016 | China | Env | 91.4 | 100 | 0.13 | 98.363 | 5283084 | 45.22 | 31 | 457885 |
| SAMN17214563-GCA_019846395.1 | MCCC_1A02609 | SAMN17214563 | GCA_019846395.1 | NonPC | Public | 2020 | China | NA | 90.9 | 100 | 0.05 | 98.3603 | 5093307 | 45.2 | 43 | 697612 |
| SAMN17215410-GCA_016541065.1 | PNUSAV001485 | SAMN17215410 | GCA_016541065.1 | NonPC | Public | 2020 | USA | Clinical | 90 | 100 | 1.36 | 98.2686 | 5323184 | 45.35 | 40 | 461339 |
| SAMN17225431-GCA_019846795.1 | NTY-EMS | SAMN17225431 | GCA_019846795.1 | NonPC | Public | 2020 | China | NA | 91.1 | 100 | 0.36 | 98.2178 | 5371012 | 45.24 | 66 | 337445 |
| SAMN17256631-GCF_017562405.1 | Patient_feces1 | SAMN17256631 | GCF_017562405.1 | NonPC | Public | 2016 | NA | Clinical | 92.8 | 99.93 | 2.99 | 98.3443 | 5242929 | 45.31 | 123 | 402030 |
| SAMN17256632-GCF_017562385.1 | Patient_feces2 | SAMN17256632 | GCF_017562385.1 | NonPC | Public | 2018 | NA | Clinical | 91.6 | 100 | 0.46 | 98.2642 | 5017968 | 45.2 | 47 | 544753 |
| SAMN17256633-GCF_017562355.1 | Patient_feces3 | SAMN17256633 | GCF_017562355.1 | NonPC | Public | 2016 | NA | Clinical | 90.5 | 100 | 0 | 98.2895 | 5204320 | 45.19 | 64 | 314860 |
| SAMN17256634-GCF_017562305.1 | Patient_feces4 | SAMN17256634 | GCF_017562305.1 | NonPC | Public | 2016 | NA | Clinical | 92.7 | 100 | 0.05 | 98.3591 | 5039209 | 45.31 | 50 | 513499 |
| SAMN17256635-GCF_017562335.1 | Patient_feces5 | SAMN17256635 | GCF_017562335.1 | NonPC | Public | 2016 | NA | Clinical | 92.6 | 100 | 0.05 | 98.353 | 5031755 | 45.3 | 53 | 514024 |
| SAMN17256636-GCF_017562285.1 | Patient_feces6 | SAMN17256636 | GCF_017562285.1 | NonPC | Public | 2016 | NA | Clinical | 92.8 | 100 | 0.14 | 98.3597 | 5253787 | 45.26 | 59 | 522582 |
| SAMN17256637-GCF_017562315.1 | Patient_feces7 | SAMN17256637 | GCF_017562315.1 | NonPC | Public | 2016 | NA | Clinical | 92.8 | 100 | 0.05 | 98.3435 | 5048295 | 45.31 | 56 | 501872 |
| SAMN17256638-GCF_017562235.1 | Patient_feces8 | SAMN17256638 | GCF_017562235.1 | NonPC | Public | 2018 | NA | Clinical | 92.8 | 100 | 0.05 | 98.3353 | 5053813 | 45.31 | 50 | 513559 |
| SAMN17256639-GCF_017562225.1 | Patient_feces9 | SAMN17256639 | GCF_017562225.1 | NonPC | Public | 2017 | NA | Clinical | 92.7 | 100 | 0.05 | 98.358 | 5035563 | 45.31 | 51 | 513507 |
| SAMN17295486-GCA_017980065.1 | PNUSAV001488 | SAMN17295486 | GCA_017980065.1 | NonPC | Public | 2020 | USA | Clinical | 89.6 | 100 | 2.3 | 98.3492 | 5251842 | 45.4 | 40 | 528801 |
| SAMN17311104-SRR13716869 | YK15 | SAMN17311104 | SRR13716869 | NonPC | Public | 2018 | China | Env | 89.6 | 100 | 0.54 | 98.3381 | 5188369 | 45.26 | 39 | 454033 |
| SAMN17311105-SRR13716860 | YK2 | SAMN17311105 | SRR13716860 | NonPC | Public | 2018 | China | Env | 91.3 | 100 | 0.68 | 98.3188 | 5372202 | 45.26 | 93 | 491588 |
| SAMN17311106-SRR13716870 | YK14 | SAMN17311106 | SRR13716870 | NonPC | Public | 2018 | China | Env | 89.9 | 100 | 0 | 98.2732 | 5034269 | 45.34 | 34 | 512194 |
| SAMN17311107-SRR13716867 | YK17 | SAMN17311107 | SRR13716867 | NonPC | Public | 2018 | China | Env | 89.9 | 100 | 0 | 98.265 | 5032537 | 45.33 | 34 | 482106 |

| | | | | | | | | | | | | | | | | |
|---|---|---|---|---|---|---|---|---|---|---|---|---|---|---|---|---|
| SAMN17311108-SRR13716858 | YK4 | SAMN17311108 | SRR13716858 | NonPC | Public | 2018 | China | Env | 91.2 | 100 | 0.41 | 98.2444 | 5214220 | 45.23 | 66 | 332443 |
| SAMN17311110-SRR13716876 | YK10-9 | SAMN17311110 | SRR13716876 | NonPC | Public | 2018 | China | Env | 91.5 | 100 | 0.07 | 98.3832 | 5194559 | 45.32 | 43 | 525058 |
| SAMN17311111-SRR13716864 | YK18-5 | SAMN17311111 | SRR13716864 | NonPC | Public | 2018 | China | Env | 91.4 | 100 | 0 | 98.358 | 5002501 | 45.4 | 43 | 408561 |
| SAMN17311112-SRR13716861 | YK18-9 | SAMN17311112 | SRR13716861 | NonPC | Public | 2018 | China | Env | 91.9 | 100 | 0.27 | 98.2909 | 5378129 | 45.12 | 74 | 421894 |
| SAMN17311113-SRR13716872 | YK12 | SAMN17311113 | SRR13716872 | NonPC | Public | 2018 | China | Env | 89.8 | 100 | 0 | 98.2858 | 5183094 | 45.2 | 37 | 570363 |
| SAMN17311114-SRR13716859 | YK20 | SAMN17311114 | SRR13716859 | NonPC | Public | 2019 | China | Env | 91.3 | 100 | 0.68 | 98.2954 | 5371987 | 45.26 | 96 | 458758 |
| SAMN17311115-SRR13716857 | YK80 | SAMN17311115 | SRR13716857 | NonPC | Public | 2019 | China | Env | 91.9 | 100 | 1.26 | 98.3694 | 5401004 | 45.37 | 66 | 315002 |
| SAMN17311116-SRR13716874 | YK114 | SAMN17311116 | SRR13716874 | NonPC | Public | 2019 | China | Env | 91.2 | 100 | 0.41 | 98.2184 | 5206709 | 45.23 | 70 | 230202 |
| SAMN17311118-SRR13716871 | YK135-2 | SAMN17311118 | SRR13716871 | NonPC | Public | 2019 | China | Env | 90 | 100 | 0.05 | 98.3072 | 5018595 | 45.38 | 38 | 841955 |
| SAMN17311120-SRR13716863 | YK188 | SAMN17311120 | SRR13716863 | NonPC | Public | 2019 | China | Env | 91.2 | 100 | 0.6 | 98.3492 | 5223551 | 45.26 | 41 | 833244 |
| SAMN17311122-SRR13716865 | YK181 | SAMN17311122 | SRR13716865 | NonPC | Public | 2019 | China | Env | 91.4 | 100 | 0.41 | 98.2789 | 5450114 | 45.15 | 41 | 530331 |
| SAMN17311123-SRR13716868 | YK160 | SAMN17311123 | SRR13716868 | NonPC | Public | 2019 | China | Env | 91.6 | 100 | 0.27 | 98.3549 | 5062586 | 45.29 | 39 | 529247 |
| SAMN17311125-SRR13716866 | YK1785 | SAMN17311125 | SRR13716866 | NonPC | Public | 2019 | China | Env | 89.2 | 100 | 0.41 | 98.3188 | 5214320 | 45.29 | 37 | 335662 |
| SAMN17311127-SRR13716852 | LH112 | SAMN17311127 | SRR13716852 | NonPC | Public | 2020 | China | Env | 88.9 | 100 | 0 | 98.3106 | 4985585 | 45.3 | 68 | 216294 |
| SAMN17311128-SRR13716848 | LH138 | SAMN17311128 | SRR13716848 | NonPC | Public | 2020 | China | Env | 89.1 | 100 | 0.2 | 98.2528 | 5264772 | 45.32 | 78 | 428576 |
| SAMN17311129-SRR13716851 | LH134 | SAMN17311129 | SRR13716851 | NonPC | Public | 2020 | China | Env | 90.8 | 100 | 0.14 | 98.2452 | 5418794 | 45.12 | 81 | 251968 |
| SAMN17311131-SRR13716854 | LH1 | SAMN17311131 | SRR13716854 | NonPC | Public | 2020 | China | Env | 90 | 99.93 | 0 | 98.3086 | 5236219 | 45.33 | 55 | 324791 |
| SAMN17311135-SRR13716878 | SDW6 | SAMN17311135 | SRR13716878 | NonPC | Public | 2018 | China | Env | 92 | 100 | 0 | 98.3927 | 5119302 | 45.31 | 35 | 471779 |
| SAMN17311136-SRR13716879 | SDW23 | SAMN17311136 | SRR13716879 | NonPC | Public | 2018 | China | Env | 89.5 | 100 | 0.14 | 98.2165 | 5117983 | 45.31 | 36 | 458460 |
| SAMN17311138-SRR13716850 | JS123 | SAMN17311138 | SRR13716850 | NonPC | Public | 2018 | China | Env | 91.7 | 100 | 0 | 98.1731 | 5499455 | 45.22 | 51 | 468549 |
| SAMN17311140-SRR13716847 | SDW1 | SAMN17311140 | SRR13716847 | NonPC | Public | 2019 | China | Env | 91 | 100 | 0 | 98.2843 | 5409751 | 45.19 | 59 | 475027 |
| SAMN17311142-SRR13716873 | JS128 | SAMN17311142 | SRR13716873 | NonPC | Public | 2019 | China | Env | 91.7 | 100 | 0 | 98.2501 | 5517249 | 45.21 | 51 | 492534 |
| SAMN17311143-SRR13716862 | JS131 | SAMN17311143 | SRR13716862 | NonPC | Public | 2019 | China | Env | 91 | 100 | 0 | 98.3186 | 5484163 | 45.18 | 70 | 615560 |
| SAMN17311144-SRR13716856 | JS132 | SAMN17311144 | SRR13716856 | NonPC | Public | 2019 | China | Env | 91 | 100 | 0 | 98.3201 | 5541471 | 45.19 | 74 | 615560 |
| SAMN17311145-SRR13716855 | JS229 | SAMN17311145 | SRR13716855 | NonPC | Public | 2019 | China | Env | 91 | 100 | 0 | 98.3068 | 5539829 | 45.19 | 69 | 615560 |
| SAMN17311146-SRR13716849 | JS124 | SAMN17311146 | SRR13716849 | NonPC | Public | 2019 | China | Env | 91.2 | 100 | 0.08 | 98.2673 | 5251079 | 45.23 | 84 | 301183 |
| SAMN17311147-SRR13716881 | SDW120 | SAMN17311147 | SRR13716881 | NonPC | Public | 2020 | China | Env | 91.7 | 100 | 0 | 98.2022 | 5517303 | 45.21 | 52 | 461215 |
| SAMN17311149-SRR13716880 | SDW128 | SAMN17311149 | SRR13716880 | NonPC | Public | 2020 | China | Env | 89.8 | 100 | 0 | 98.3046 | 5182343 | 45.2 | 38 | 570363 |
| SAMN17379931-GCA_016728585.1 | 20MO12SH02-V1 | SAMN17379931 | GCA_016728585.1 | NonPC | Public | 2020 | USA | Env | 88.1 | 100 | 3.57 | 98.0748 | 5662822 | 44.97 | 135 | 147207 |
| SAMN17379932-GCA_016728545.1 | 20MO12SH02-V2 | SAMN17379932 | GCA_016728545.1 | NonPC | Public | 2020 | USA | Env | 88.1 | 100 | 2.1 | 98.0279 | 5616635 | 44.99 | 135 | 184574 |
| SAMN17723471-GCA_017180615.1 | PNUSAV001381 | SAMN17723471 | GCA_017180615.1 | NonPC | Public | 2020 | USA | Clinical | 89.8 | 100 | 0.11 | 98.2477 | 5252913 | 45.19 | 44 | 372130 |
| SAMN17723478-GCA_017180575.1 | PNUSAV001382 | SAMN17723478 | GCA_017180575.1 | NonPC | Public | 2020 | USA | Clinical | 89.2 | 100 | 4.36 | 98.0893 | 5284296 | 45.28 | 95 | 155532 |
| SAMN17723484-GCA_017180655.1 | PNUSAV001389 | SAMN17723484 | GCA_017180655.1 | NonPC | Public | 2020 | USA | Clinical | 89.7 | 100 | 0.38 | 98.2701 | 5263890 | 45.19 | 51 | 256412 |
| SAMN17723489-GCA_017180535.1 | PNUSAV001279 | SAMN17723489 | GCA_017180535.1 | NonPC | Public | 2020 | USA | Clinical | 89.6 | 100 | 2.9 | 98.2692 | 5293064 | 45.26 | 134 | 76994 |
| SAMN17723497-GCA_018025155.1 | PNUSAV001280 | SAMN17723497 | GCA_018025155.1 | NonPC | Public | 2020 | USA | Clinical | 91.1 | 100 | 0.42 | 98.312 | 5121773 | 45.36 | 197 | 56838 |
| SAMN17723501-GCA_018025135.1 | PNUSAV001268 | SAMN17723501 | GCA_018025135.1 | NonPC | Public | 2020 | USA | Clinical | 91.3 | 100 | 2.11 | 98.2892 | 5277078 | 45.35 | 116 | 100990 |
| SAMN17723507-GCA_017180635.1 | PNUSAV001245 | SAMN17723507 | GCA_017180635.1 | NonPC | Public | 2020 | USA | Clinical | 88.8 | 100 | 3.06 | 98.1712 | 5220192 | 45.19 | 48 | 248954 |
| SAMN17799677-GCA_017126575.1 | PNUSAV001263 | SAMN17799677 | GCA_017126575.1 | NonPC | Public | 2020 | USA | Clinical | 88.7 | 99.68 | 0.4 | 98.4554 | 4970270 | 45.55 | 142 | 160536 |
| SAMN17808220-GCA_018126665.1 | VB18PR-0007-1 | SAMN17808220 | GCA_018126665.1 | NonPC | Public | NA | NA | Env | 90.6 | 100 | 0.54 | 98.3329 | 5134677 | 45.28 | 106 | 133801 |
| SAMN17808223-GCA_018126645.1 | VB18PR-0011-2 | SAMN17808223 | GCA_018126645.1 | NonPC | Public | NA | NA | Env | 91.7 | 100 | 1.13 | 98.3847 | 5479576 | 45.4 | 88 | 349537 |
| SAMN17808224-SRR13684777 | VB18PR-0012-1 | SAMN17808224 | SRR13684777 | NonPC | Public | NA | NA | Env | 87.8 | 100 | 1.49 | 98.0161 | 5466747 | 45.07 | 109 | 138751 |
| SAMN17808225-SRR13684778 | VB18PR-0012-2 | SAMN17808225 | SRR13684778 | NonPC | Public | NA | NA | Env | 91 | 100 | 0.11 | 98.3124 | 5117772 | 45.28 | 49 | 400967 |
| SAMN17808226-SRR13684779 | VB18PR-0013-1 | SAMN17808226 | SRR13684779 | NonPC | Public | NA | NA | Env | 88.6 | 100 | 0 | 98.0475 | 5280454 | 45.09 | 137 | 122014 |
| SAMN17808227-SRR13684780 | VB18PR-0020-1 | SAMN17808227 | SRR13684780 | NonPC | Public | NA | NA | Env | 90.1 | 100 | 0.18 | 98.2247 | 5389890 | 45.14 | 56 | 569134 |
| SAMN17808228-GCA_018118885.1 | VB18PR-0021-1 | SAMN17808228 | GCA_018118885.1 | NonPC | Public | NA | NA | Env | 88.3 | 100 | 0.53 | 98.2038 | 5271795 | 45.13 | 76 | 237791 |
| SAMN17808231-SRR13684784 | VB18PR-0023-2 | SAMN17808231 | SRR13684784 | NonPC | Public | NA | NA | Env | 91.6 | 100 | 0 | 98.3728 | 5051366 | 45.38 | 60 | 557409 |
| SAMN17808235-GCA_018119105.1 | VB18PR-0025-2 | SAMN17808235 | GCA_018119105.1 | NonPC | Public | NA | NA | Env | 90 | 100 | 0.34 | 98.3152 | 5012805 | 45.39 | 63 | 250317 |
| SAMN17808239-GCA_018126605.1 | VB18PR-0034-1 | SAMN17808239 | GCA_018126605.1 | NonPC | Public | NA | NA | Env | 91.2 | 100 | 0.3 | 98.3466 | 5328433 | 45.42 | 94 | 182811 |

| | | | | | | | | | | | | | | | | |
|---|---|---|---|---|---|---|---|---|---|---|---|---|---|---|---|---|
| SAMN17808240-GCA_018126585.1 | VB18PR-0035-1 | SAMN17808240 | GCA_018126585.1 | NonPC | Public | NA | NA | Env | 89.7 | 100 | 2.15 | 98.3032 | 5166915 | 45.32 | 56 | 342382 |
| SAMN17808241-GCA_018119065.1 | VB18PR-0035-2 | SAMN17808241 | GCA_018119065.1 | NonPC | Public | NA | NA | Env | 88.5 | 100 | 0.34 | 98.0439 | 5079172 | 45.26 | 86 | 190088 |
| SAMN17808242-SRR13684796 | VB18PR-0036-1 | SAMN17808242 | SRR13684796 | NonPC | Public | NA | NA | Env | 91.1 | 100 | 0.05 | 98.3394 | 5240378 | 45.23 | 62 | 303981 |
| SAMN17808243-GCA_018121575.1 | VB18PR-0036-2 | SAMN17808243 | GCA_018121575.1 | NonPC | Public | NA | NA | Env | 88.7 | 100 | 2.9 | 98.0708 | 5355906 | 45.21 | 74 | 220527 |
| SAMN17808244-GCA_018126365.1 | VB18PR-0037-1 | SAMN17808244 | GCA_018126365.1 | NonPC | Public | NA | NA | Env | 91 | 100 | 0.1 | 98.3713 | 5070511 | 45.34 | 59 | 297157 |
| SAMN17808245-GCA_018118705.1 | VB18PR-0039-1 | SAMN17808245 | GCA_018118705.1 | NonPC | Public | NA | NA | Env | 89.6 | 100 | 2.29 | 98.3055 | 5583456 | 45.18 | 61 | 270641 |
| SAMN17808247-GCA_018119305.1 | VB18PR-0045-1 | SAMN17808247 | GCA_018119305.1 | NonPC | Public | NA | NA | Env | 87.4 | 100 | 3 | 98.076 | 5389499 | 45.14 | 111 | 93272 |
| SAMN17808248-GCA_018122395.1 | VB18PR-0045-2 | SAMN17808248 | GCA_018122395.1 | NonPC | Public | NA | NA | Env | 90.8 | 100 | 0.2 | 98.3168 | 5037865 | 45.33 | 55 | 375878 |
| SAMN17808249-GCA_018126325.1 | VB18PR-0048-1 | SAMN17808249 | GCA_018126325.1 | NonPC | Public | NA | NA | Env | 88.6 | 100 | 0.66 | 98.0854 | 5147880 | 45.19 | 85 | 191914 |
| SAMN17808250-GCA_018126545.1 | VB18PR-0050-1 | SAMN17808250 | GCA_018126545.1 | NonPC | Public | NA | NA | Env | 88 | 100 | 3.8 | 98.0406 | 5227520 | 45.25 | 69 | 180222 |
| SAMN17808252-GCA_018126525.1 | VB18PR-0053-1 | SAMN17808252 | GCA_018126525.1 | NonPC | Public | NA | NA | Env | 91.5 | 100 | 0.87 | 98.3132 | 5587052 | 44.76 | 84 | 129939 |
| SAMN17808253-GCA_018118845.1 | VB18PR-0055-1 | SAMN17808253 | GCA_018118845.1 | NonPC | Public | NA | NA | Env | 91.4 | 100 | 0.82 | 98.3478 | 5213853 | 45.24 | 57 | 399592 |
| SAMN17808254-GCA_018126305.1 | VB18PR-0059-1 | SAMN17808254 | GCA_018126305.1 | NonPC | Public | NA | NA | Env | 87.3 | 100 | 0.82 | 97.9856 | 5367838 | 45.1 | 84 | 185162 |
| SAMN17808256-GCA_018119145.1 | VB18PR-0061-1 | SAMN17808256 | GCA_018119145.1 | NonPC | Public | NA | NA | Env | 90.9 | 100 | 1.17 | 98.3052 | 5535102 | 45.32 | 100 | 175025 |
| SAMN17808257-GCA_018126485.1 | VB18PR-0062-1 | SAMN17808257 | GCA_018126485.1 | NonPC | Public | NA | NA | Env | 89.6 | 100 | 0.41 | 98.202 | 5305886 | 45.09 | 74 | 158615 |
| SAMN17808258-GCA_018119325.1 | VB18PR-0066-1 | SAMN17808258 | GCA_018119325.1 | NonPC | Public | NA | NA | Env | 88.8 | 100 | 0.26 | 98.0595 | 5335945 | 45.19 | 111 | 166185 |
| SAMN17808261-SRR13684817 | VB18PR-0073-1 | SAMN17808261 | SRR13684817 | NonPC | Public | NA | NA | Env | 89.6 | 100 | 0.34 | 98.2636 | 5314521 | 45 | 62 | 236799 |
| SAMN17808262-GCA_018122415.1 | VB18PR-0074-1 | SAMN17808262 | GCA_018122415.1 | NonPC | Public | NA | NA | Env | 88.4 | 100 | 0.98 | 98.1901 | 5139249 | 45.24 | 77 | 301760 |
| SAMN17808266-GCA_018119225.1 | VB18PR-0090-2 | SAMN17808266 | GCA_018119225.1 | NonPC | Public | NA | NA | Env | 91.5 | 100 | 0.37 | 98.3579 | 5264881 | 45.31 | 80 | 278628 |
| SAMN17808269-GCA_018126425.1 | VB18PR-0095-1 | SAMN17808269 | GCA_018126425.1 | NonPC | Public | NA | NA | Env | 90.9 | 100 | 0.96 | 98.3757 | 5540449 | 45.26 | 120 | 193594 |
| SAMN17808270-GCA_018119125.1 | VB18PR-0096-1 | SAMN17808270 | GCA_018119125.1 | NonPC | Public | NA | NA | Env | 89 | 100 | 1.54 | 98.077 | 5220168 | 45.35 | 76 | 187880 |
| SAMN17808271-GCA_018127005.1 | VB18PR-0096-2 | SAMN17808271 | GCA_018127005.1 | NonPC | Public | NA | NA | Env | 89.8 | 100 | 1.17 | 98.3258 | 5126293 | 45.35 | 49 | 412295 |
| SAMN17808272-SRR13684683 | VB18PR-0105-1 | SAMN17808272 | SRR13684683 | NonPC | Public | NA | NA | Env | 87.4 | 100 | 0 | 98.1238 | 5367927 | 45.06 | 126 | 99234 |
| SAMN17808273-GCA_018118805.1 | VB18PR-0106-1 | SAMN17808273 | GCA_018118805.1 | NonPC | Public | NA | NA | Env | 89.1 | 100 | 0.34 | 98.3083 | 5201971 | 45.18 | 61 | 302749 |
| SAMN17808276-GCA_018121595.1 | VB18PR-0117-1 | SAMN17808276 | GCA_018121595.1 | NonPC | Public | NA | NA | Env | 91.5 | 100 | 1.3 | 98.3669 | 5312205 | 45.15 | 81 | 186164 |
| SAMN17808278-GCA_018119595.1 | VB18PR-0118-2 | SAMN17808278 | GCA_018119595.1 | NonPC | Public | NA | NA | Env | 88.7 | 100 | 0.4 | 98.0577 | 5481316 | 45.18 | 115 | 134761 |
| SAMN17808279-GCA_018118825.1 | VB18PR-0121-1 | SAMN17808279 | GCA_018118825.1 | NonPC | Public | NA | NA | Env | 89.4 | 100 | 0.44 | 98.3207 | 5234676 | 45.07 | 69 | 214646 |
| SAMN17808282-GCA_018119465.1 | VB18PR-0131-1 | SAMN17808282 | GCA_018119465.1 | NonPC | Public | NA | NA | Env | 88.5 | 100 | 0.27 | 98.0998 | 5544398 | 45.27 | 103 | 175655 |
| SAMN17808283-GCA_018126465.1 | VB18PR-0131-2 | SAMN17808283 | GCA_018126465.1 | NonPC | Public | NA | NA | Env | 91.1 | 100 | 1.29 | 98.3626 | 5326500 | 45.04 | 88 | 150644 |
| SAMN17808285-GCA_018119005.1 | VB18PR-0133-1 | SAMN17808285 | GCA_018119005.1 | NonPC | Public | NA | NA | Env | 88.9 | 100 | 1.54 | 98.0082 | 5220414 | 45.2 | 82 | 190729 |
| SAMN17808286-GCA_018118965.1 | VB18PR-0133-2 | SAMN17808286 | GCA_018118965.1 | NonPC | Public | NA | NA | Env | 88.2 | 100 | 2.45 | 98.2162 | 5373841 | 45.15 | 80 | 311787 |
| SAMN17808289-GCA_018126945.1 | VB18PR-0137-1 | SAMN17808289 | GCA_018126945.1 | NonPC | Public | NA | NA | Env | 89.8 | 100 | 3.01 | 98.3315 | 5159953 | 45.39 | 63 | 412942 |
| SAMN17808290-SRR13684703 | VB18PR-0144-1 | SAMN17808290 | SRR13684703 | NonPC | Public | NA | NA | Env | 91.6 | 100 | 0 | 98.3413 | 5480485 | 45.14 | 85 | 341514 |
| SAMN17808291-GCA_018126885.1 | VB18PR-0145-1 | SAMN17808291 | GCA_018126885.1 | NonPC | Public | NA | NA | Env | 88.9 | 100 | 1.17 | 98.0213 | 5247909 | 45.19 | 80 | 172741 |
| SAMN17808292-GCA_018119045.1 | VB18PR-0146-1 | SAMN17808292 | GCA_018119045.1 | NonPC | Public | NA | NA | Env | 89.2 | 100 | 0.34 | 98.2789 | 5448611 | 45.36 | 61 | 381815 |
| SAMN17808293-GCA_018118865.1 | VB18PR-0147-1 | SAMN17808293 | GCA_018118865.1 | NonPC | Public | NA | NA | Env | 88.8 | 100 | 1.73 | 98.0574 | 5166367 | 45.27 | 76 | 224044 |
| SAMN17808294-GCA_018121555.1 | VB18PR-0147-2 | SAMN17808294 | GCA_018121555.1 | NonPC | Public | NA | NA | Env | 90.1 | 100 | 3 | 98.3033 | 5591757 | 45.25 | 94 | 269959 |
| SAMN17808295-SRR13684709 | VB18PR-0148-1 | SAMN17808295 | SRR13684709 | NonPC | Public | NA | NA | Env | 89 | 100 | 0.19 | 98.0927 | 5340056 | 45.19 | 91 | 167347 |
| SAMN17808297-GCA_018122315.1 | VB18PR-0149-2 | SAMN17808297 | GCA_018122315.1 | NonPC | Public | NA | NA | Env | 91.8 | 100 | 2.12 | 98.3722 | 5462845 | 45.09 | 80 | 153106 |
| SAMN17808299-GCA_018126985.1 | VB18PR-0150-2 | SAMN17808299 | GCA_018126985.1 | NonPC | Public | NA | NA | Env | 89.8 | 100 | 0 | 98.3182 | 5077105 | 45.31 | 86 | 194613 |
| SAMN17808300-SRR13684714 | VB18PR-0151-1 | SAMN17808300 | SRR13684714 | NonPC | Public | NA | NA | Env | 91.3 | 100 | 0 | 98.3285 | 4946825 | 45.38 | 39 | 372648 |
| SAMN17808301-GCA_018118645.1 | VB18PR-0152-1 | SAMN17808301 | GCA_018118645.1 | NonPC | Public | NA | NA | Env | 88.1 | 100 | 0.2 | 98.0834 | 5235303 | 45.23 | 97 | 179521 |
| SAMN17808302-GCA_018119345.1 | VB18PR-0152-2 | SAMN17808302 | GCA_018119345.1 | NonPC | Public | NA | NA | Env | 90.7 | 100 | 1.31 | 98.3025 | 5341705 | 45.21 | 113 | 103260 |
| SAMN17808306-GCA_018118785.1 | VB18PR-0165-1 | SAMN17808306 | GCA_018118785.1 | NonPC | Public | NA | NA | Env | 89.7 | 100 | 0.89 | 98.2803 | 5269074 | 45.1 | 62 | 163206 |
| SAMN17808307-SRR13684722 | VB18PR-0170-1 | SAMN17808307 | SRR13684722 | NonPC | Public | NA | NA | Env | 88.2 | 100 | 0.3 | 98.0969 | 5164919 | 45.2 | 45 | 351671 |
| SAMN17808308-GCA_018126925.1 | VB18PR-0173-1 | SAMN17808308 | GCA_018126925.1 | NonPC | Public | NA | NA | Env | 91.5 | 100 | 0.89 | 98.2958 | 5211136 | 45.21 | 64 | 462118 |
| SAMN17808309-GCA_018118745.1 | VB18PR-0176-1 | SAMN17808309 | GCA_018118745.1 | NonPC | Public | NA | NA | Env | 88.3 | 100 | 0.48 | 98.0936 | 5125152 | 45.28 | 83 | 181462 |
| SAMN17808310-GCA_018118605.1 | VB18PR-0177-1 | SAMN17808310 | GCA_018118605.1 | NonPC | Public | NA | NA | Env | 88.3 | 100 | 0.37 | 98.1561 | 4993027 | 45.34 | 76 | 191601 |

| | | | | | | | | | | | | | | | | |
|---|---|---|---|---|---|---|---|---|---|---|---|---|---|---|---|---|
| SAMN17808311-GCA_018122905.1 | VB18PR-0177-2 | SAMN17808311 | GCA_018122905.1 | NonPC | Public | NA | NA | Env | 91 | 100 | 2.75 | 98.3322 | 5506848 | 45.23 | 79 | 371847 |
| SAMN17808312-GCA_018119025.1 | VB18PR-0178-1 | SAMN17808312 | GCA_018119025.1 | NonPC | Public | NA | NA | Env | 89.7 | 100 | 2.27 | 98.2807 | 5396540 | 45.14 | 57 | 199991 |
| SAMN17808313-GCA_018127065.1 | VB18PR-0178-2 | SAMN17808313 | GCA_018127065.1 | NonPC | Public | NA | NA | Env | 91.3 | 100 | 4.23 | 98.3682 | 5271100 | 45.41 | 56 | 464867 |
| SAMN17808315-GCA_018126845.1 | VB18PR-0181-1 | SAMN17808315 | GCA_018126845.1 | NonPC | Public | NA | NA | Env | 88.5 | 100 | 0.97 | 98.1055 | 5269700 | 45.17 | 136 | 82022 |
| SAMN17808316-GCA_018126865.1 | VB18PR-0182-1 | SAMN17808316 | GCA_018126865.1 | NonPC | Public | NA | NA | Env | 88.6 | 100 | 0.41 | 97.9981 | 5404346 | 44.98 | 139 | 106641 |
| SAMN17808317-GCA_018119085.1 | VB18PR-0183-1 | SAMN17808317 | GCA_018119085.1 | NonPC | Public | NA | NA | Env | 89.2 | 100 | 0.33 | 98.2589 | 5441028 | 45.13 | 100 | 177496 |
| SAMN17808318-SRR13684734 | VB18PR-0187-1 | SAMN17808318 | SRR13684734 | NonPC | Public | NA | NA | Env | 89.6 | 100 | 0.37 | 98.3254 | 5240706 | 45.08 | 67 | 237031 |
| SAMN17808319-SRR13684735 | VB18PR-0187-2 | SAMN17808319 | SRR13684735 | NonPC | Public | NA | NA | Env | 89.7 | 100 | 0.54 | 98.3013 | 5306203 | 45.2 | 59 | 206119 |
| SAMN17808320-GCA_018126825.1 | VB18PR-0188-1 | SAMN17808320 | GCA_018126825.1 | NonPC | Public | NA | NA | Env | 90.3 | 100 | 1.7 | 98.2328 | 5588878 | 45.17 | 62 | 453743 |
| SAMN17808322-GCA_018126785.1 | VB18PR-0192-1 | SAMN17808322 | GCA_018126785.1 | NonPC | Public | NA | NA | Env | 88.7 | 100 | 1.61 | 98.0303 | 5248226 | 45.23 | 102 | 134761 |
| SAMN17808323-GCA_018122575.1 | VB18PR-0193-1 | SAMN17808323 | GCA_018122575.1 | NonPC | Public | NA | NA | Env | 89.6 | 100 | 4.04 | 98.2367 | 5273602 | 45.23 | 53 | 367321 |
| SAMN17808324-GCA_018126445.1 | VB18PR-0193-2 | SAMN17808324 | GCA_018126445.1 | NonPC | Public | NA | NA | Env | 91.7 | 100 | 1.68 | 98.2785 | 5372027 | 45.23 | 68 | 264227 |
| SAMN17808327-GCA_018118625.1 | VB18PR-0198-2 | SAMN17808327 | GCA_018118625.1 | NonPC | Public | NA | NA | Env | 90.5 | 100 | 2.57 | 98.2305 | 5345512 | 45.22 | 79 | 363136 |
| SAMN17808328-GCA_018118725.1 | VB18PR-0199-1 | SAMN17808328 | GCA_018118725.1 | NonPC | Public | NA | NA | Env | 90.6 | 100 | 1.3 | 98.2826 | 5433767 | 45.17 | 73 | 313592 |
| SAMN17808329-GCA_018126405.1 | VB18PR-0199-2 | SAMN17808329 | GCA_018126405.1 | NonPC | Public | NA | NA | Env | 87.5 | 100 | 3.19 | 98.2784 | 5477627 | 45.36 | 55 | 405921 |
| SAMN17808332-SRR13684749 | VB18PR-0201-1 | SAMN17808332 | SRR13684749 | NonPC | Public | NA | NA | Env | 88.9 | 100 | 0.41 | 98.0542 | 5204416 | 45.25 | 91 | 211511 |
| SAMN17808333-GCA_018126765.1 | VB18PR-0201-2 | SAMN17808333 | GCA_018126765.1 | NonPC | Public | NA | NA | Env | 89.3 | 100 | 3.41 | 98.2512 | 5560050 | 45.12 | 73 | 330120 |
| SAMN17808334-GCA_018122355.1 | VB18PR-0202-1 | SAMN17808334 | GCA_018122355.1 | NonPC | Public | NA | NA | Env | 88.8 | 100 | 0.94 | 98.0462 | 5586773 | 45.22 | 109 | 168254 |
| SAMN17808335-GCA_018118925.1 | VB18PR-0202-2 | SAMN17808335 | GCA_018118925.1 | NonPC | Public | NA | NA | Env | 90.1 | 100 | 1.32 | 98.263 | 5223784 | 45.37 | 85 | 370460 |
| SAMN17808337-GCA_018127025.1 | VB18PR-0204-1 | SAMN17808337 | GCA_018127025.1 | NonPC | Public | NA | NA | Env | 91.3 | 100 | 1.14 | 98.2399 | 5327355 | 45.3 | 74 | 254428 |
| SAMN17808338-GCA_018126725.1 | VB18PR-0205-1 | SAMN17808338 | GCA_018126725.1 | NonPC | Public | NA | NA | Env | 91.4 | 100 | 1.98 | 98.3462 | 5420905 | 45.36 | 77 | 329164 |
| SAMN17808342-SRR13684760 | VB18PR-0210-2 | SAMN17808342 | SRR13684760 | NonPC | Public | NA | NA | Env | 88.5 | 100 | 0.44 | 98.2218 | 5193660 | 45.21 | 69 | 429194 |
| SAMN17808343-GCA_018119205.1 | VB18PR-0211-1 | SAMN17808343 | GCA_018119205.1 | NonPC | Public | NA | NA | Env | 87.4 | 99.73 | 0.65 | 98.3413 | 5168979 | 45.21 | 120 | 132815 |
| SAMN17808346-GCA_018119185.1 | VB18PR-0275-3 | SAMN17808346 | GCA_018119185.1 | NonPC | Public | NA | NA | Env | 88.1 | 100 | 0.9 | 98.1168 | 5415825 | 45.5 | 103 | 234524 |
| SAMN17808347-GCA_018122555.1 | VB18PR-0275-4 | SAMN17808347 | GCA_018122555.1 | NonPC | Public | NA | NA | Env | 88.1 | 100 | 1.97 | 98.2824 | 5601163 | 44.94 | 95 | 216363 |
| SAMN17808348-GCA_018118765.1 | VB18PR-0275-5 | SAMN17808348 | GCA_018118765.1 | NonPC | Public | NA | NA | Env | 91.9 | 100 | 0.05 | 98.3096 | 5384433 | 45.24 | 112 | 256092 |
| SAMN17808349-GCA_018126345.1 | VB18PR-0275-6 | SAMN17808349 | GCA_018126345.1 | NonPC | Public | NA | NA | Env | 88.1 | 100 | 1.53 | 98.2486 | 5571350 | 44.95 | 96 | 237598 |
| SAMN17808351-GCA_018126565.1 | VB18PR-0277-2 | SAMN17808351 | GCA_018126565.1 | NonPC | Public | NA | NA | Env | 89 | 100 | 3.72 | 98.0878 | 5528498 | 45.23 | 86 | 177060 |
| SAMN17808352-GCA_018126505.1 | VB18PR-0277-3 | SAMN17808352 | GCA_018126505.1 | NonPC | Public | NA | NA | Env | 89 | 100 | 1.45 | 98.081 | 5392226 | 45.22 | 89 | 177057 |
| SAMN17808354-GCA_018122885.1 | VB18PR-0277-5 | SAMN17808354 | GCA_018122885.1 | NonPC | Public | NA | NA | Env | 91.3 | 100 | 3.3 | 98.3699 | 5360318 | 45.27 | 48 | 313374 |
| SAMN17808359-GCA_018126805.1 | VB18PR-0278-4 | SAMN17808359 | GCA_018126805.1 | NonPC | Public | NA | NA | Env | 91 | 99.73 | 0.75 | 98.2982 | 5372766 | 45.14 | 55 | 401978 |
| SAMN17831969-GCA_019272895.1 | Colony52 | SAMN17831969 | GCA_019272895.1 | NonPC | Public | NA | Thailand | Env | 91.5 | 98.54 | 0 | 98.3715 | 5392094 | 45.7 | 2 | 3577848 |
| SAMN17831970-GCA_019272875.1 | Colony83 | SAMN17831970 | GCA_019272875.1 | NonPC | Public | 2019 | Thailand | Clinical | 91.5 | 98.54 | 0 | 98.3715 | 5392094 | 45.7 | 2 | 3577848 |
| SAMN17831971-GCA_019272855.1 | Colony89 | SAMN17831971 | GCA_019272855.1 | NonPC | Public | NA | Thailand | Env | 91.5 | 98.54 | 0 | 98.3715 | 5392094 | 45.7 | 2 | 3577848 |
| SAMN17831972-GCA_019272835.1 | Colony91 | SAMN17831972 | GCA_019272835.1 | NonPC | Public | NA | Thailand | Env | 91.5 | 98.54 | 0 | 98.3715 | 5392094 | 45.7 | 2 | 3577848 |
| SAMN17831973-GCA_019272815.1 | Colony95 | SAMN17831973 | GCA_019272815.1 | NonPC | Public | NA | Thailand | Env | 91.5 | 98.54 | 0 | 98.3715 | 5392094 | 45.7 | 2 | 3577848 |
| SAMN17831974-GCA_019272795.1 | Colony96 | SAMN17831974 | GCA_019272795.1 | NonPC | Public | NA | Thailand | Env | 91.5 | 98.54 | 0 | 98.3715 | 5392094 | 45.7 | 2 | 3577848 |
| SAMN17831975-GCA_019272775.1 | Colony97 | SAMN17831975 | GCA_019272775.1 | NonPC | Public | NA | Thailand | Env | 91.5 | 98.54 | 0 | 98.3715 | 5392094 | 45.7 | 2 | 3577848 |
| SAMN17831976-GCA_019272755.1 | Colony12 | SAMN17831976 | GCA_019272755.1 | NonPC | Public | 2019 | Thailand | Clinical | 91.5 | 98.54 | 0 | 98.3715 | 5392094 | 45.7 | 2 | 3577848 |
| SAMN17831977-GCA_019272715.1 | Colony15 | SAMN17831977 | GCA_019272715.1 | NonPC | Public | 2019 | Thailand | Clinical | 91.5 | 98.54 | 0 | 98.3715 | 5392094 | 45.7 | 2 | 3577848 |
| SAMN17831978-GCA_019272675.1 | Colony18 | SAMN17831978 | GCA_019272675.1 | NonPC | Public | 2019 | Thailand | Clinical | 91.5 | 98.54 | 0 | 98.3715 | 5392094 | 45.7 | 2 | 3577848 |
| SAMN17831979-GCA_019272575.1 | Colony19 | SAMN17831979 | GCA_019272575.1 | NonPC | Public | 2019 | Thailand | Clinical | 91.5 | 98.54 | 0 | 98.3715 | 5392094 | 45.7 | 2 | 3577848 |
| SAMN17831980-GCA_019272495.1 | Colony28 | SAMN17831980 | GCA_019272495.1 | NonPC | Public | 2019 | Thailand | Clinical | 91.5 | 98.54 | 0 | 98.3715 | 5392094 | 45.7 | 2 | 3577848 |
| SAMN17831981-GCA_019272405.1 | Colony33 | SAMN17831981 | GCA_019272405.1 | NonPC | Public | 2019 | Thailand | Clinical | 91.5 | 98.54 | 0 | 98.3715 | 5392094 | 45.7 | 2 | 3577848 |
| SAMN17831982-GCA_019272355.1 | Colony34 | SAMN17831982 | GCA_019272355.1 | NonPC | Public | 2019 | Thailand | Clinical | 91.5 | 98.54 | 0 | 98.3715 | 5392094 | 45.7 | 2 | 3577848 |
| SAMN17831983-GCA_019272235.1 | Colony35 | SAMN17831983 | GCA_019272235.1 | NonPC | Public | 2019 | Thailand | Clinical | 91.5 | 98.54 | 0 | 98.3715 | 5392094 | 45.7 | 2 | 3577848 |
| SAMN17831984-GCA_019272155.1 | Colony38 | SAMN17831984 | GCA_019272155.1 | NonPC | Public | 2019 | Thailand | Clinical | 91.5 | 98.54 | 0 | 98.3715 | 5392094 | 45.7 | 2 | 3577848 |
| SAMN17831985-GCA_019272095.1 | Colony39 | SAMN17831985 | GCA_019272095.1 | NonPC | Public | 2019 | Thailand | Clinical | 91.5 | 98.54 | 0 | 98.3715 | 5392094 | 45.7 | 2 | 3577848 |

| | | | | | | | | | | | | | | | | |
|---|---|---|---|---|---|---|---|---|---|---|---|---|---|---|---|---|
| SAMN17831986-GCA_019272075.1 | Colony43 | SAMN17831986 | GCA_019272075.1 | NonPC | Public | 2019 | Thailand | Clinical | 91.5 | 98.54 | 0 | 98.3715 | 5392094 | 45.7 | 2 | 3577848 |
| SAMN17831987-GCA_019272055.1 | Colony45 | SAMN17831987 | GCA_019272055.1 | NonPC | Public | 2019 | Thailand | Clinical | 91.5 | 98.54 | 0 | 98.3715 | 5392094 | 45.7 | 2 | 3577848 |
| SAMN17831988-GCA_019272035.1 | Colony46 | SAMN17831988 | GCA_019272035.1 | NonPC | Public | 2019 | Thailand | Clinical | 91.5 | 98.54 | 0 | 98.3715 | 5392094 | 45.7 | 2 | 3577848 |
| SAMN17831989-GCA_019272015.1 | Colony99 | SAMN17831989 | GCA_019272015.1 | NonPC | Public | NA | Thailand | Env | 91.5 | 98.54 | 0 | 98.3715 | 5392094 | 45.7 | 2 | 3577848 |
| SAMN17831990-GCA_019271995.1 | Colony265 | SAMN17831990 | GCA_019271995.1 | NonPC | Public | NA | Thailand | Env | 91.5 | 98.54 | 0 | 98.3715 | 5392094 | 45.7 | 2 | 3577848 |
| SAMN17831991-GCA_019271975.1 | Colony266 | SAMN17831991 | GCA_019271975.1 | NonPC | Public | NA | Thailand | Env | 91.5 | 98.54 | 0 | 98.3715 | 5392094 | 45.7 | 2 | 3577848 |
| SAMN17831992-GCA_019271955.1 | Colony521 | SAMN17831992 | GCA_019271955.1 | NonPC | Public | NA | Thailand | Env | 91.5 | 98.54 | 0 | 98.3715 | 5392094 | 45.7 | 2 | 3577848 |
| SAMN17831993-GCA_019271935.1 | Colony523 | SAMN17831993 | GCA_019271935.1 | NonPC | Public | NA | Thailand | Env | 89.5 | 98.64 | 0 | 98.2759 | 5392094 | 45.74 | 2 | 3577848 |
| SAMN17831994-GCA_019271915.1 | Colony525 | SAMN17831994 | GCA_019271915.1 | NonPC | Public | NA | Thailand | Env | 91 | 99.42 | 0 | 98.2832 | 5392094 | 45.65 | 2 | 3577848 |
| SAMN17831995-GCA_019271895.1 | Colony551 | SAMN17831995 | GCA_019271895.1 | NonPC | Public | NA | Thailand | Env | 91.2 | 98.99 | 0 | 98.3447 | 5392094 | 45.7 | 2 | 3577848 |
| SAMN17831996-GCA_019271875.1 | Colony184 | SAMN17831996 | GCA_019271875.1 | NonPC | Public | NA | Thailand | Env | 91.5 | 98.54 | 0 | 98.3715 | 5392094 | 45.7 | 2 | 3577848 |
| SAMN17831997-GCA_019271675.1 | Colony92 | SAMN17831997 | GCA_019271675.1 | NonPC | Public | NA | Thailand | Env | 91.6 | 98.76 | 0 | 98.37 | 5392094 | 45.72 | 2 | 3577848 |
| SAMN17831998-GCA_019271575.1 | Colony93 | SAMN17831998 | GCA_019271575.1 | NonPC | Public | NA | Thailand | Env | 89.8 | 98.65 | 0 | 98.4075 | 5392094 | 45.74 | 2 | 3577848 |
| SAMN17831999-GCA_019271755.1 | Colony269 | SAMN17831999 | GCA_019271755.1 | NonPC | Public | NA | Thailand | Env | 89.8 | 98.23 | 0 | 98.3199 | 5392094 | 45.74 | 2 | 3577848 |
| SAMN17832002-GCA_019271295.1 | Colony526 | SAMN17832002 | GCA_019271295.1 | NonPC | Public | NA | Thailand | Env | 89.4 | 99.46 | 0 | 98.3079 | 5392094 | 45.77 | 2 | 3577848 |
| SAMN17832003-GCA_019271255.1 | Colony545 | SAMN17832003 | GCA_019271255.1 | NonPC | Public | NA | Thailand | Env | 91.6 | 98.88 | 0 | 98.3595 | 5392094 | 45.69 | 2 | 3577848 |
| SAMN17832004-GCA_019271235.1 | Colony546 | SAMN17832004 | GCA_019271235.1 | NonPC | Public | NA | Thailand | Env | 89.8 | 99.25 | 0 | 98.2531 | 5392094 | 45.76 | 2 | 3577848 |
| SAMN17832005-GCA_019271215.1 | Colony547 | SAMN17832005 | GCA_019271215.1 | NonPC | Public | NA | Thailand | Env | 91.2 | 98.54 | 0 | 98.3789 | 5392094 | 45.72 | 2 | 3577848 |
| SAMN17832006-GCA_019271195.1 | Colony548 | SAMN17832006 | GCA_019271195.1 | NonPC | Public | NA | Thailand | Env | 91.2 | 99.34 | 0 | 98.4026 | 5392094 | 45.67 | 2 | 3577848 |
| SAMN17832007-GCA_019271175.1 | Colony549 | SAMN17832007 | GCA_019271175.1 | NonPC | Public | NA | Thailand | Env | 89.9 | 98.05 | 0 | 98.2845 | 5392094 | 45.73 | 2 | 3577848 |
| SAMN17832008-GCA_019271155.1 | Colony550 | SAMN17832008 | GCA_019271155.1 | PrePC | Public | NA | Thailand | Env | 91.9 | 98.44 | 0 | 99.4237 | 5392094 | 45.72 | 2 | 3577848 |
| SAMN17832009-GCA_019271135.1 | Colony552 | SAMN17832009 | GCA_019271135.1 | NonPC | Public | NA | Thailand | Env | 91.3 | 98.72 | 0 | 98.3273 | 5392094 | 45.71 | 2 | 3577848 |
| SAMN17832010-GCA_019271115.1 | Colony553 | SAMN17832010 | GCA_019271115.1 | NonPC | Public | NA | Thailand | Env | 91.3 | 98.16 | 0.05 | 98.3274 | 5392094 | 45.69 | 2 | 3577848 |
| SAMN17832011-GCA_019271095.1 | Colony554 | SAMN17832011 | GCA_019271095.1 | NonPC | Public | NA | Thailand | Env | 89.9 | 99.15 | 0 | 98.3704 | 5392094 | 45.76 | 2 | 3577848 |
| SAMN17832012-GCA_019271075.1 | Colony555 | SAMN17832012 | GCA_019271075.1 | NonPC | Public | NA | Thailand | Env | 91.4 | 98.67 | 0 | 98.3657 | 5392094 | 45.71 | 2 | 3577848 |
| SAMN17832013-GCA_019271055.1 | Colony556 | SAMN17832013 | GCA_019271055.1 | NonPC | Public | NA | Thailand | Env | 91.3 | 98.61 | 0.05 | 98.3088 | 5392094 | 45.71 | 2 | 3577848 |
| SAMN17832014-GCA_019271035.1 | Colony557 | SAMN17832014 | GCA_019271035.1 | NonPC | Public | NA | Thailand | Env | 90 | 98.6 | 0 | 98.3146 | 5392094 | 45.76 | 2 | 3577848 |
| SAMN17832015-GCA_019271015.1 | Colony558 | SAMN17832015 | GCA_019271015.1 | NonPC | Public | NA | Thailand | Env | 89.7 | 99.51 | 0 | 98.2994 | 5392094 | 45.76 | 2 | 3577848 |
| SAMN17832016-GCA_019270995.1 | Colony559 | SAMN17832016 | GCA_019270995.1 | NonPC | Public | NA | Thailand | Env | 91.2 | 98.05 | 0 | 98.3564 | 5392094 | 45.69 | 2 | 3577848 |
| SAMN17832017-GCA_019270875.1 | Colony601 | SAMN17832017 | GCA_019270875.1 | NonPC | Public | NA | Thailand | Env | 91.6 | 98.45 | 0 | 98.3993 | 5392094 | 45.69 | 2 | 3577848 |
| SAMN17832018-GCA_019270795.1 | Colony602 | SAMN17832018 | GCA_019270795.1 | NonPC | Public | NA | Thailand | Env | 91.3 | 98.81 | 0 | 98.3242 | 5392094 | 45.7 | 2 | 3577848 |
| SAMN17832019-GCA_019270695.1 | Colony607 | SAMN17832019 | GCA_019270695.1 | NonPC | Public | NA | Thailand | Env | 91.3 | 98.32 | 0 | 98.382 | 5392094 | 45.69 | 2 | 3577848 |
| SAMN17832020-GCA_019270675.1 | Colony544 | SAMN17832020 | GCA_019270675.1 | NonPC | Public | NA | Thailand | Env | 91.3 | 99.42 | 0 | 98.259 | 5392094 | 45.65 | 2 | 3577848 |
| SAMN17833952-GCA_018023935.1 | PNUSAV001342 | SAMN17833952 | GCA_018023935.1 | NonPC | Public | 2020 | USA | Clinical | 91.4 | 100 | 0.21 | 98.2955 | 5186027 | 45.24 | 42 | 556743 |
| SAMN17839803-GCA_019273755.1 | Colony90 | SAMN17839803 | GCA_019273755.1 | NonPC | Public | NA | Thailand | Env | 89.4 | 98.57 | 0 | 98.2747 | 5392094 | 45.76 | 2 | 3577848 |
| SAMN17839804-GCA_019273735.1 | Colony524 | SAMN17839804 | GCA_019273735.1 | NonPC | Public | NA | Thailand | Env | 91.1 | 98.43 | 0 | 98.3212 | 5392094 | 45.69 | 2 | 3577848 |
| SAMN17839805-GCA_019273715.1 | Colony608 | SAMN17839805 | GCA_019273715.1 | NonPC | Public | NA | Thailand | Env | 91.3 | 98.88 | 0 | 98.4034 | 5392094 | 45.71 | 2 | 3577848 |
| SAMN17839806-GCA_019273695.1 | Colony127 | SAMN17839806 | GCA_019273695.1 | NonPC | Public | 2019 | Thailand | Env | 90 | 98.85 | 0 | 98.3561 | 5392094 | 45.74 | 2 | 3577848 |
| SAMN17846525-SRR14373877 | ZJ47 | SAMN17846525 | SRR14373877 | NonPC | Public | 2018 | China | Env | 91.3 | 100 | 0.37 | 98.362 | 5371699 | 45.32 | 45 | 277697 |
| SAMN17917431-GCF_018799025.1 | Colony41 | SAMN17917431 | GCF_018799025.1 | NonPC | Public | NA | Thailand | Clinical | 91.5 | 98.54 | 0 | 98.3715 | 5392094 | 45.7 | 2 | 3577848 |
| SAMN18065262-GCA_017108585.1 | PNUSAV001345 | SAMN18065262 | GCA_017108585.1 | NonPC | Public | 2020 | USA | Clinical | 90.6 | 100 | 3.19 | 98.2634 | 5206895 | 45.31 | 132 | 82069 |
| SAMN18065263-SRR13790765 | PNUSAV001319 | SAMN18065263 | SRR13790765 | NonPC | Public | 2020 | USA | Clinical | 91.5 | 100 | 0 | 98.3338 | 5154094 | 45.23 | 35 | 808117 |
| SAMN18065265-SRR13790764 | PNUSAV001251 | SAMN18065265 | SRR13790764 | NonPC | Public | 2020 | USA | Clinical | 89.9 | 100 | 0.31 | 98.215 | 5115283 | 45.26 | 54 | 203568 |
| SAMN18102746-GCF_017313735.1 | Vb0825 | SAMN18102746 | GCF_017313735.1 | NonPC | Public | 2016 | China | Env | 91.5 | 100 | 0.89 | 98.3513 | 5447706 | 45.18 | 84 | 186809 |
| SAMN18102747-GCF_017313825.1 | Vb0849 | SAMN18102747 | GCF_017313825.1 | NonPC | Public | 2016 | China | Env | 91.5 | 100 | 3.5 | 98.3461 | 5456075 | 45.17 | 96 | 168624 |
| SAMN18102749-GCF_017313685.1 | Vb0915 | SAMN18102749 | GCF_017313685.1 | NonPC | Public | 2016 | China | Env | 90.9 | 100 | 0.34 | 98.3143 | 5270561 | 45.02 | 95 | 218527 |
| SAMN18102751-GCF_017313655.1 | Vb1207 | SAMN18102751 | GCF_017313655.1 | NonPC | Public | 2016 | China | Env | 88.3 | 100 | 0.33 | 98.1361 | 5335753 | 45.22 | 67 | 265168 |
| SAMN18102752-GCF_017313595.1 | Vb1382 | SAMN18102752 | GCF_017313595.1 | NonPC | Public | 2016 | China | Env | 88.3 | 100 | 0.27 | 98.1201 | 5231087 | 45.1 | 76 | 189192 |

| | | | | | | | | | | | | | | | | |
|---|---|---|---|---|---|---|---|---|---|---|---|---|---|---|---|---|
| SAMN18102754-GCF_017313545.1 | Vb2153 | SAMN18102754 | GCF_017313545.1 | NonPC | Public | 2017 | China | Env | 89 | 100 | 0.89 | 98.2922 | 5203373 | 45.1 | 95 | 191142 |
| SAMN18102755-GCF_017313585.1 | Vb2188 | SAMN18102755 | GCF_017313585.1 | NonPC | Public | 2017 | China | Env | 90.2 | 100 | 3.45 | 98.2486 | 5092299 | 45.34 | 49 | 277303 |
| SAMN18102756-GCF_017313565.1 | Vb2225 | SAMN18102756 | GCF_017313565.1 | NonPC | Public | 2017 | China | Env | 90.9 | 100 | 1.74 | 98.3403 | 5289322 | 45.03 | 101 | 205682 |
| SAMN18102759-GCF_017313485.1 | Vb2511 | SAMN18102759 | GCF_017313485.1 | NonPC | Public | 2017 | China | Env | 88.4 | 100 | 1.07 | 98.0442 | 5396696 | 45 | 73 | 269747 |
| SAMN18314262-GCA_017372055.1 | 210030 | SAMN18314262 | GCA_017372055.1 | NonPC | Public | 2016 | USA | Env | 89.2 | 100 | 4.62 | 98.0205 | 5181945 | 45.42 | 117 | 100475 |
| SAMN18394829-GCA_019321785.1 | 64 | SAMN18394829 | GCA_019321785.1 | NonPC | Public | 2018 | China | Env | 90.9 | 99.86 | 0.23 | 98.2501 | 5427006 | 45.29 | 4 | 3446671 |
| SAMN18509352-GCA_017604525.1 | 210041 | SAMN18509352 | GCA_017604525.1 | NonPC | Public | 2016 | USA | Env | 91.3 | 100 | 0.27 | 98.355 | 5099019 | 45.41 | 166 | 71701 |
| SAMN18509353-GCA_017604485.1 | 210040 | SAMN18509353 | GCA_017604485.1 | NonPC | Public | 2016 | USA | Env | 89.9 | 100 | 0.27 | 98.188 | 5034685 | 45.37 | 269 | 34315 |
| SAMN18509354-GCA_017604205.1 | 210042 | SAMN18509354 | GCA_017604205.1 | NonPC | Public | 2016 | USA | Env | 91 | 100 | 0.69 | 98.2256 | 5339725 | 45.31 | 203 | 51757 |
| SAMN18509355-GCA_017604185.1 | 210038 | SAMN18509355 | GCA_017604185.1 | NonPC | Public | 2016 | USA | Env | 91 | 100 | 0 | 98.3957 | 5092842 | 45.42 | 280 | 31351 |
| SAMN18509359-GCA_017604385.1 | 210039 | SAMN18509359 | GCA_017604385.1 | NonPC | Public | 2016 | USA | Env | 90.8 | 100 | 1.91 | 98.2638 | 5260867 | 45.29 | 161 | 76949 |
| SAMN18509367-GCA_018017275.1 | 210034 | SAMN18509367 | GCA_018017275.1 | NonPC | Public | 2016 | USA | Env | 91.5 | 100 | 1.76 | 98.2453 | 5367121 | 45.3 | 152 | 77722 |
| SAMN18509371-GCA_017604405.1 | 210035 | SAMN18509371 | GCA_017604405.1 | NonPC | Public | 2016 | USA | Env | 90.8 | 100 | 0.44 | 98.3244 | 5128014 | 45.35 | 184 | 68183 |
| SAMN18509374-GCA_017604465.1 | 210037 | SAMN18509374 | GCA_017604465.1 | NonPC | Public | 2016 | USA | Env | 89.8 | 100 | 1.51 | 98.1617 | 5183924 | 45.34 | 176 | 60599 |
| SAMN18509378-GCA_017604425.1 | 210033 | SAMN18509378 | GCA_017604425.1 | NonPC | Public | 2016 | USA | Env | 90.6 | 100 | 3 | 98.3401 | 5171281 | 45.39 | 145 | 74274 |
| SAMN18509452-GCA_017604445.1 | 210032 | SAMN18509452 | GCA_017604445.1 | NonPC | Public | 2016 | USA | Env | 91.5 | 100 | 4.36 | 98.3734 | 5496102 | 45.22 | 87 | 139616 |
| SAMN18509455-GCA_017604505.1 | 210031 | SAMN18509455 | GCA_017604505.1 | NonPC | Public | 2016 | USA | Env | 91 | 100 | 1.17 | 98.275 | 5225426 | 45.18 | 82 | 214850 |
| SAMN18509469-GCA_018017295.1 | 210044 | SAMN18509469 | GCA_018017295.1 | NonPC | Public | 2016 | USA | Env | 90.6 | 100 | 0.2 | 98.2696 | 5321016 | 45.3 | 227 | 51404 |
| SAMN18509470-GCA_017604115.1 | 210043 | SAMN18509470 | GCA_017604115.1 | NonPC | Public | 2016 | USA | Env | 87.7 | 100 | 1.94 | 98.2892 | 5172841 | 45.39 | 155 | 67959 |
| SAMN18510895-GCA_018017315.1 | 210085 | SAMN18510895 | GCA_018017315.1 | NonPC | Public | 2017 | USA | Env | 90.4 | 100 | 4.27 | 98.1811 | 5257092 | 45.25 | 45 | 305756 |
| SAMN18510896-GCA_017603325.1 | 210084 | SAMN18510896 | GCA_017603325.1 | NonPC | Public | 2017 | USA | Env | 90.9 | 100 | 2.59 | 98.3002 | 5213962 | 45.39 | 116 | 100814 |
| SAMN18510948-GCA_017603345.1 | 210079 | SAMN18510948 | GCA_017603345.1 | NonPC | Public | 2017 | USA | Env | 90.2 | 100 | 2.07 | 98.3157 | 5331649 | 45.25 | 101 | 120909 |
| SAMN18510949-GCA_017610715.1 | 210074 | SAMN18510949 | GCA_017610715.1 | NonPC | Public | 2017 | USA | Env | 91.2 | 100 | 3 | 98.3239 | 5390721 | 45.23 | 107 | 115996 |
| SAMN18510950-GCA_017603175.1 | 210075 | SAMN18510950 | GCA_017603175.1 | NonPC | Public | 2017 | USA | Env | 91.2 | 100 | 4.2 | 98.3301 | 5428835 | 45.27 | 122 | 91120 |
| SAMN18510951-GCA_017603215.1 | 210073 | SAMN18510951 | GCA_017603215.1 | NonPC | Public | 2017 | USA | Env | 91.2 | 100 | 4.75 | 98.2507 | 5571669 | 45.24 | 83 | 226898 |
| SAMN18510952-GCA_018017235.1 | 210076 | SAMN18510952 | GCA_018017235.1 | NonPC | Public | 2017 | USA | Env | 91.2 | 100 | 2.18 | 98.1776 | 5342156 | 45.19 | 45 | 226363 |
| SAMN18510953-GCA_017603285.1 | 210077 | SAMN18510953 | GCA_017603285.1 | NonPC | Public | 2017 | USA | Env | 88.1 | 100 | 2.65 | 98.2681 | 5059692 | 45.38 | 73 | 110226 |
| SAMN18510954-GCA_017603245.1 | 210081 | SAMN18510954 | GCA_017603245.1 | NonPC | Public | 2017 | USA | Env | 88.2 | 100 | 2.6 | 98.249 | 5233196 | 45.32 | 118 | 100572 |
| SAMN18510955-GCA_017603165.1 | 210080 | SAMN18510955 | GCA_017603165.1 | NonPC | Public | 2017 | USA | Env | 90.9 | 100 | 3.81 | 98.4107 | 5169113 | 45.45 | 107 | 100423 |
| SAMN18510956-GCA_017603385.1 | 210082 | SAMN18510956 | GCA_017603385.1 | NonPC | Public | 2017 | USA | Env | 89.6 | 100 | 3.54 | 98.3127 | 5152538 | 45.43 | 46 | 292042 |
| SAMN18510957-GCA_017603185.1 | 210083 | SAMN18510957 | GCA_017603185.1 | NonPC | Public | 2017 | USA | Env | 88.6 | 100 | 3.09 | 98.1212 | 5534465 | 45.46 | 192 | 62561 |
| SAMN18510958-GCA_017603365.1 | 210070 | SAMN18510958 | GCA_017603365.1 | NonPC | Public | 2017 | USA | Env | 90.8 | 100 | 0.34 | 98.2452 | 5123093 | 45.33 | 198 | 48638 |
| SAMN18510959-GCA_017603305.1 | 210078 | SAMN18510959 | GCA_017603305.1 | NonPC | Public | 2017 | USA | Env | 91.9 | 100 | 4.22 | 98.2875 | 5133516 | 45.45 | 105 | 89640 |
| SAMN18528220-GCA_017637785.1 | 210276 | SAMN18528220 | GCA_017637785.1 | NonPC | Public | 2018 | USA | Env | 91.2 | 100 | 2.04 | 98.2798 | 5434463 | 45.18 | 65 | 447450 |
| SAMN18528229-GCA_017637725.1 | 210292 | SAMN18528229 | GCA_017637725.1 | NonPC | Public | 2019 | USA | Env | 89.1 | 100 | 4.98 | 98.3607 | 5216044 | 45.36 | 51 | 321808 |
| SAMN18528234-GCA_017637645.1 | 210286 | SAMN18528234 | GCA_017637645.1 | NonPC | Public | 2019 | USA | Env | 90.5 | 100 | 1.45 | 98.3097 | 5176165 | 45.18 | 82 | 157031 |
| SAMN18528238-GCA_018017095.1 | 210289 | SAMN18528238 | GCA_018017095.1 | NonPC | Public | 2019 | USA | Env | 90.2 | 100 | 0.69 | 98.2476 | 5141529 | 45.3 | 58 | 478881 |
| SAMN18528241-GCA_017637665.1 | 210280 | SAMN18528241 | GCA_017637665.1 | NonPC | Public | 2018 | USA | Env | 90.5 | 100 | 2.99 | 98.1796 | 5302081 | 45.18 | 65 | 395365 |
| SAMN18528243-GCA_017637745.1 | 210290 | SAMN18528243 | GCA_017637745.1 | NonPC | Public | 2019 | USA | Env | 91.8 | 100 | 1.18 | 98.3361 | 5115724 | 45.27 | 50 | 434019 |
| SAMN18528244-GCA_017637685.1 | 210291 | SAMN18528244 | GCA_017637685.1 | NonPC | Public | 2019 | USA | Env | 89.8 | 100 | 0.94 | 98.3004 | 5153018 | 45.28 | 38 | 529950 |
| SAMN18528252-GCA_018017055.1 | 210274 | SAMN18528252 | GCA_018017055.1 | NonPC | Public | 2018 | USA | Env | 91.2 | 100 | 4.08 | 98.3002 | 5288086 | 45.28 | 50 | 212330 |
| SAMN18528254-GCA_017637525.1 | 210266 | SAMN18528254 | GCA_017637525.1 | NonPC | Public | 2018 | USA | Env | 88.8 | 100 | 2.27 | 98.1668 | 5137639 | 45.28 | 36 | 522247 |
| SAMN18528255-GCA_017637565.1 | 210272 | SAMN18528255 | GCA_017637565.1 | NonPC | Public | 2018 | USA | Env | 89.6 | 100 | 4.77 | 98.3004 | 5353484 | 45.41 | 35 | 488162 |
| SAMN18528260-GCA_017637705.1 | 210268 | SAMN18528260 | GCA_017637705.1 | NonPC | Public | 2018 | USA | Env | 91 | 100 | 1.5 | 98.1784 | 5196980 | 45.25 | 57 | 413073 |
| SAMN18530006-GCA_017657415.1 | 210072 | SAMN18530006 | GCA_017657415.1 | NonPC | Public | 2017 | USA | Env | 90.7 | 100 | 1.99 | 98.3375 | 5261520 | 45.29 | 191 | 54860 |
| SAMN18721142-GCF_018135645.1 | FB-11 | SAMN18721142 | GCF_018135645.1 | NonPC | Public | 2017 | China | Env | 92.1 | 100 | 0.14 | 98.3362 | 5134111 | 45.36 | 2 | 3328286 |
| SAMN18742552-GCA_017968025.1 | 210346 | SAMN18742552 | GCA_017968025.1 | NonPC | Public | 2020 | USA | Env | 91.5 | 100 | 0.58 | 98.3572 | 5084834 | 45.32 | 37 | 456153 |
| SAMN18742554-GCA_017968145.1 | 210347 | SAMN18742554 | GCA_017968145.1 | NonPC | Public | 2020 | USA | Env | 89.1 | 100 | 0.54 | 98.1482 | 5013311 | 45.43 | 52 | 532145 |

| | | | | | | | | | | | | | | | | |
|---|---|---|---|---|---|---|---|---|---|---|---|---|---|---|---|---|
| SAMN18742557-GCA_017968045.1 | 210345 | SAMN18742557 | GCA_017968045.1 | NonPC | Public | 2020 | USA | Env | 88.9 | 100 | 4.12 | 98.1298 | 5594528 | 45.3 | 59 | 337634 |
| SAMN18742569-GCA_017968065.1 | 210349 | SAMN18742569 | GCA_017968065.1 | NonPC | Public | 2020 | USA | Env | 88.2 | 100 | 3.49 | 98.2181 | 5212944 | 45.33 | 36 | 385776 |
| SAMN18762037-GCA_018081105.1 | 210355 | SAMN18762037 | GCA_018081105.1 | NonPC | Public | 2020 | USA | Env | 90.4 | 100 | 4.9 | 98.2113 | 5285784 | 45.27 | 58 | 306317 |
| SAMN18762040-GCA_018081085.1 | 210354 | SAMN18762040 | GCA_018081085.1 | NonPC | Public | 2020 | USA | Env | 89.2 | 100 | 3.27 | 98.1054 | 5176912 | 45.32 | 59 | 233285 |
| SAMN18762051-GCA_018081165.1 | 210352 | SAMN18762051 | GCA_018081165.1 | NonPC | Public | 2020 | USA | Env | 89.6 | 100 | 0.42 | 98.2351 | 5128115 | 45.26 | 46 | 318774 |
| SAMN18796120-SRR14272962 | Vp._1513 | SAMN18796120 | SRR14272962 | NonPC | Public | 2016 | China | Env | 89.2 | 100 | 0.1 | 98.3125 | 5285099 | 45.27 | 104 | 207429 |
| SAMN19488933-GCA_018620435.1 | PNUSAV001574 | SAMN19488933 | GCA_018620435.1 | NonPC | Public | 2021 | USA | Clinical | 89.6 | 100 | 0.89 | 98.1224 | 5192394 | 45.26 | 76 | 136634 |
| SAMN19595902-GCA_018771785.1 | PNUSAV001576 | SAMN19595902 | GCA_018771785.1 | NonPC | Public | 2021 | USA | Clinical | 91.3 | 99.93 | 1.27 | 98.3548 | 5129663 | 45.3 | 147 | 71599 |
| SAMN19769865-GCA_021890375.1 | VpL7-7 | SAMN19769865 | GCA_021890375.1 | NonPC | Public | 2017 | China | Env | 90.7 | 100 | 0 | 98.4311 | 4935297 | 45.36 | 49 | 314969 |
| SAMN19769866-GCA_021890315.1 | VpN1-22 | SAMN19769866 | GCA_021890315.1 | NonPC | Public | 2017 | China | Env | 90.7 | 100 | 0.63 | 98.3842 | 5053467 | 45.22 | 37 | 651479 |
| SAMN19769867-GCA_021890335.1 | VpN3-33 | SAMN19769867 | GCA_021890335.1 | NonPC | Public | 2017 | China | Env | 90.7 | 100 | 0.1 | 98.3573 | 5016485 | 45.33 | 53 | 329323 |
| SAMN19769868-GCA_021890355.1 | VpN4-46 | SAMN19769868 | GCA_021890355.1 | NonPC | Public | 2017 | China | Env | 91.2 | 100 | 0 | 98.3161 | 5044352 | 45.3 | 63 | 402855 |
| SAMN19769869-GCA_021890385.1 | VpN8-42 | SAMN19769869 | GCA_021890385.1 | NonPC | Public | 2017 | China | Env | 90.8 | 100 | 0.27 | 98.358 | 4997249 | 45.39 | 37 | 508421 |
| SAMN19769870-GCA_021890295.1 | VpQ8-15 | SAMN19769870 | GCA_021890295.1 | NonPC | Public | 2017 | China | Env | 91.1 | 100 | 0 | 98.3233 | 4984771 | 45.34 | 21 | 1320740 |
| SAMN19796157-GCA_018991165.1 | PNUSAV001583 | SAMN19796157 | GCA_018991165.1 | NonPC | Public | 2021 | USA | Env | 91.5 | 100 | 4.71 | 98.3823 | 5254422 | 45.34 | 42 | 368102 |
| SAMN19796160-GCA_019000345.1 | PNUSAV001586 | SAMN19796160 | GCA_019000345.1 | NonPC | Public | 2021 | USA | Env | 90.7 | 100 | 3.73 | 98.285 | 5354915 | 45.13 | 44 | 418029 |
| SAMN19796170-GCA_018992345.1 | PNUSAV001588 | SAMN19796170 | GCA_018992345.1 | NonPC | Public | 2021 | USA | Env | 89.2 | 100 | 4.71 | 98.3664 | 5672216 | 45.13 | 79 | 202154 |
| SAMN19796191-GCA_019000665.1 | PNUSAV001582 | SAMN19796191 | GCA_019000665.1 | NonPC | Public | 2021 | USA | Clinical | 91 | 100 | 1.9 | 98.1664 | 5282672 | 45.21 | 93 | 127249 |
| SAMN19796196-GCA_018996245.1 | PNUSAV001587 | SAMN19796196 | GCA_018996245.1 | NonPC | Public | 2021 | USA | Env | 91.6 | 100 | 3.86 | 98.2728 | 5224887 | 45.35 | 64 | 373590 |
| SAMN19796200-GCA_019000625.1 | PNUSAV001602 | SAMN19796200 | GCA_019000625.1 | NonPC | Public | 2021 | USA | Env | 91.1 | 100 | 0.91 | 98.368 | 5095035 | 45.35 | 64 | 176958 |
| SAMN19918798-SRR14970089 | PNUSAV001647 | SAMN19918798 | SRR14970089 | NonPC | Public | 2021 | USA | Clinical | 90.3 | 99.91 | 0 | 98.2774 | 5216603 | 45.18 | 70 | 184959 |
| SAMN19918801-SRR14977867 | PNUSAV001641 | SAMN19918801 | SRR14977867 | NonPC | Public | 2021 | USA | Env | 90.5 | 100 | 0.27 | 98.297 | 5158207 | 45.24 | 50 | 564114 |
| SAMN19918804-SRR14949218 | PNUSAV001640 | SAMN19918804 | SRR14949218 | NonPC | Public | 2021 | USA | Env | 92.1 | 100 | 0.35 | 98.3857 | 5235836 | 45.21 | 44 | 381088 |
| SAMN19918808-SRR14962703 | PNUSAV001633 | SAMN19918808 | SRR14962703 | NonPC | Public | 2021 | USA | Env | 90.9 | 100 | 0.14 | 98.2057 | 5081475 | 45.21 | 40 | 530409 |
| SAMN19918811-SRR14958947 | PNUSAV001644 | SAMN19918811 | SRR14958947 | NonPC | Public | 2021 | USA | Env | 91.1 | 100 | 0.35 | 98.2432 | 5072021 | 45.25 | 61 | 399129 |
| SAMN19918814-SRR14977890 | PNUSAV001631 | SAMN19918814 | SRR14977890 | NonPC | Public | 2021 | USA | Env | 89.7 | 100 | 0.05 | 98.3276 | 5014116 | 45.37 | 29 | 1078980 |
| SAMN19918823-SRR14979165 | PNUSAV001629 | SAMN19918823 | SRR14979165 | NonPC | Public | 2021 | USA | Env | 90.4 | 100 | 0.3 | 98.2418 | 5309598 | 45.11 | 42 | 522938 |
| SAMN19918826-SRR14959381 | PNUSAV001646 | SAMN19918826 | SRR14959381 | NonPC | Public | 2021 | USA | Env | 88.3 | 100 | 0.37 | 98.2136 | 5061963 | 45.26 | 59 | 562938 |
| SAMN19918827-SRR14935289 | PNUSAV001636 | SAMN19918827 | SRR14935289 | NonPC | Public | 2021 | USA | Env | 90.9 | 100 | 0.08 | 98.2733 | 5093233 | 45.22 | 32 | 547253 |
| SAMN19918828-SRR14944463 | PNUSAV001643 | SAMN19918828 | SRR14944463 | NonPC | Public | 2021 | USA | Env | 89.6 | 100 | 0 | 98.3492 | 4970699 | 45.37 | 38 | 1053952 |
| SAMN19918829-SRR14944235 | PNUSAV001634 | SAMN19918829 | SRR14944235 | NonPC | Public | 2021 | USA | Env | 91 | 100 | 0.27 | 98.3322 | 5028102 | 45.32 | 44 | 529006 |
| SAMN19918836-SRR14970029 | PNUSAV001642 | SAMN19918836 | SRR14970029 | NonPC | Public | 2021 | USA | Env | 91.6 | 100 | 0.05 | 98.363 | 5029183 | 45.34 | 44 | 739570 |
| SAMN19918848-SRR14978822 | PNUSAV001627 | SAMN19918848 | SRR14978822 | NonPC | Public | 2021 | USA | Env | 89 | 100 | 0.05 | 98.116 | 4945932 | 45.36 | 65 | 288833 |
| SAMN19918850-SRR14977702 | PNUSAV001630 | SAMN19918850 | SRR14977702 | NonPC | Public | 2021 | USA | Env | 91.1 | 100 | 0.3 | 98.2736 | 5307252 | 45.1 | 50 | 521388 |
| SAMN19918854-SRR14959424 | PNUSAV001635 | SAMN19918854 | SRR14959424 | NonPC | Public | 2021 | USA | Env | 90.9 | 100 | 0.08 | 98.3063 | 5066618 | 45.3 | 26 | 769803 |
| SAMN19918858-SRR14979008 | PNUSAV001639 | SAMN19918858 | SRR14979008 | NonPC | Public | 2021 | USA | Env | 92 | 100 | 0 | 98.3047 | 5037859 | 45.33 | 36 | 910884 |
| SAMN19918862-SRR14985912 | PNUSAV001632 | SAMN19918862 | SRR14985912 | NonPC | Public | 2021 | USA | Env | 91.3 | 100 | 0.33 | 98.363 | 5059682 | 45.3 | 42 | 531649 |
| SAMN19918865-SRR14994468 | PNUSAV001638 | SAMN19918865 | SRR14994468 | NonPC | Public | 2021 | USA | Env | 88.7 | 100 | 0.39 | 98.1916 | 5102498 | 45.22 | 45 | 555109 |
| SAMN19918901-GCA_019058655.1 | 21MO05SH02-V2 | SAMN19918901 | GCA_019058655.1 | NonPC | Public | 2021 | USA | Env | 89 | 100 | 0.61 | 98.058 | 5553654 | 45.19 | 87 | 235844 |
| SAMN19918904-GCA_019059295.1 | 21MO05SH02-V1 | SAMN19918904 | GCA_019059295.1 | NonPC | Public | 2021 | USA | Env | 89.8 | 100 | 0.74 | 98.2923 | 5283734 | 45.27 | 48 | 394221 |
| SAMN20007432-GCA_019105305.1 | PNUSAV001657 | SAMN20007432 | GCA_019105305.1 | NonPC | Public | 2021 | USA | Clinical | 90 | 100 | 4.22 | 98.263 | 5207543 | 45.35 | 103 | 175158 |
| SAMN20081126-GCA_019147655.1 | PNUSAV001648 | SAMN20081126 | GCA_019147655.1 | NonPC | Public | 2021 | USA | Clinical | 91.4 | 100 | 4.09 | 98.3064 | 5295451 | 45.3 | 65 | 199554 |
| SAMN20294544-GCA_019351375.1 | BH-Huang-1-01 | SAMN20294544 | GCA_019351375.1 | PC | Public | 2020 | China | Clinical | 97.5 | 100 | 0 | 99.9131 | 5100543 | 45.26 | 57 | 425528 |
| SAMN20305592-GCA_019291805.1 | PNUSAV001718 | SAMN20305592 | GCA_019291805.1 | NonPC | Public | 2021 | USA | Env | 91.1 | 100 | 2.38 | 98.3287 | 5244062 | 45.19 | 41 | 400881 |
| SAMN20305596-GCA_019290435.1 | PNUSAV001705 | SAMN20305596 | GCA_019290435.1 | NonPC | Public | 2021 | USA | Env | 88.4 | 100 | 0.69 | 98.0842 | 4997369 | 45.37 | 57 | 227662 |
| SAMN20305597-GCA_019290755.1 | PNUSAV001704 | SAMN20305597 | GCA_019290755.1 | NonPC | Public | 2021 | USA | Env | 88.7 | 100 | 3.89 | 98.0794 | 5074923 | 45.44 | 47 | 282254 |
| SAMN20305604-GCA_019291885.1 | PNUSAV001715 | SAMN20305604 | GCA_019291885.1 | NonPC | Public | 2021 | USA | Env | 88.6 | 100 | 4.23 | 98.069 | 5237476 | 45.31 | 85 | 191862 |
| SAMN20305607-GCA_019291605.1 | PNUSAV001722 | SAMN20305607 | GCA_019291605.1 | NonPC | Public | 2021 | USA | Env | 88.7 | 100 | 2.32 | 98.0725 | 5163827 | 45.3 | 60 | 561830 |

| | | | | | | | | | | | | | | | | |
|---|---|---|---|---|---|---|---|---|---|---|---|---|---|---|---|---|
| SAMN20329550-GCA_019301935.1 | PNUSAV001727 | SAMN20329550 | GCA_019301935.1 | NonPC | Public | 2021 | USA | Clinical | 89.5 | 100 | 3.02 | 98.2263 | 5608791 | 44.99 | 139 | 80573 |
| SAMN20329554-GCA_019301775.1 | PNUSAV001728 | SAMN20329554 | GCA_019301775.1 | NonPC | Public | 2021 | USA | Clinical | 89.5 | 100 | 4.41 | 98.2337 | 5528830 | 45.33 | 158 | 71737 |
| SAMN20357679-GCA_019333385.1 | PNUSAV001771 | SAMN20357679 | GCA_019333385.1 | NonPC | Public | 2021 | USA | Clinical | 89.6 | 100 | 1.96 | 98.2422 | 5464689 | 45.22 | 55 | 216824 |
| SAMN20357680-GCA_019332775.1 | PNUSAV001772 | SAMN20357680 | GCA_019332775.1 | NonPC | Public | 2021 | USA | Clinical | 89.2 | 100 | 3.54 | 98.0792 | 5216894 | 45.3 | 78 | 220026 |
| SAMN20357683-GCA_019332715.1 | PNUSAV001724 | SAMN20357683 | GCA_019332715.1 | NonPC | Public | 2021 | USA | Clinical | 91.4 | 100 | 0.2 | 98.3003 | 5145237 | 45.24 | 40 | 556737 |
| SAMN20357685-GCA_019332995.1 | PNUSAV001725 | SAMN20357685 | GCA_019332995.1 | NonPC | Public | 2021 | USA | Clinical | 91.4 | 100 | 2.53 | 98.27 | 5279663 | 45.23 | 43 | 564960 |
| SAMN20395455-GCA_019429585.1 | R-14810 | SAMN20395455 | GCA_019429585.1 | NonPC | Public | 2016 | Russia | Env | 92.7 | 100 | 0.14 | 98.4016 | 5114536 | 45.21 | 49 | 373274 |
| SAMN20399185-GCA_019358715.1 | PNUSAV001671 | SAMN20399185 | GCA_019358715.1 | NonPC | Public | 2021 | USA | Clinical | 91.2 | 100 | 3.27 | 98.3008 | 5292587 | 45.31 | 110 | 100041 |
| SAMN20399187-GCA_019358575.1 | PNUSAV001665 | SAMN20399187 | GCA_019358575.1 | NonPC | Public | 2021 | USA | Clinical | 88.8 | 100 | 2.04 | 98.0967 | 5144044 | 45.41 | 185 | 54532 |
| SAMN20432052-GCA_019377495.1 | PNUSAV001812 | SAMN20432052 | GCA_019377495.1 | NonPC | Public | 2021 | USA | Clinical | 89.5 | 100 | 0.39 | 98.2351 | 5428150 | 45.33 | 208 | 54082 |
| SAMN20432055-GCA_019377575.1 | PNUSAV001813 | SAMN20432055 | GCA_019377575.1 | NonPC | Public | 2021 | USA | Clinical | 90.5 | 100 | 3.49 | 98.303 | 5346271 | 45.29 | 99 | 116214 |
| SAMN20458861-GCA_019442865.1 | PNUSAV001830 | SAMN20458861 | GCA_019442865.1 | NonPC | Public | 2021 | USA | Env | 91.1 | 100 | 1.77 | 98.3397 | 5200860 | 45.3 | 46 | 457869 |
| SAMN20458862-GCA_019442625.1 | PNUSAV001829 | SAMN20458862 | GCA_019442625.1 | NonPC | Public | 2021 | USA | Env | 91.2 | 100 | 3.48 | 98.3449 | 5334161 | 45.26 | 44 | 411818 |
| SAMN20458863-GCA_019437985.1 | PNUSAV001828 | SAMN20458863 | GCA_019437985.1 | NonPC | Public | 2021 | USA | Env | 90.2 | 100 | 4.65 | 98.2709 | 5356847 | 45.3 | 54 | 516149 |
| SAMN20458874-GCA_019393695.1 | PNUSAV001823 | SAMN20458874 | GCA_019393695.1 | NonPC | Public | 2021 | USA | Env | 90.8 | 100 | 3.43 | 98.2289 | 5620987 | 45.23 | 61 | 452613 |
| SAMN20458877-GCA_019438425.1 | PNUSAV001820 | SAMN20458877 | GCA_019438425.1 | NonPC | Public | 2021 | USA | Env | 90.6 | 100 | 3.22 | 98.2884 | 5400499 | 45.23 | 39 | 456535 |
| SAMN20522348-GCA_019569255.1 | PNUSAV001867 | SAMN20522348 | GCA_019569255.1 | NonPC | Public | 2021 | USA | Clinical | 91.4 | 100 | 1.54 | 98.3126 | 5181008 | 45.25 | 44 | 762300 |
| SAMN20522350-GCA_019569755.1 | PNUSAV001866 | SAMN20522350 | GCA_019569755.1 | NonPC | Public | 2021 | USA | Clinical | 91.4 | 100 | 0.08 | 98.3362 | 5143401 | 45.24 | 44 | 564960 |
| SAMN20527485-GCA_019563515.1 | PNUSAV001775 | SAMN20527485 | GCA_019563515.1 | NonPC | Public | 2021 | USA | Clinical | 90.6 | 100 | 0.07 | 98.2878 | 5089565 | 45.29 | 42 | 586654 |
| SAMN20527487-GCA_019563495.1 | PNUSAV001880 | SAMN20527487 | GCA_019563495.1 | NonPC | Public | 2021 | USA | Clinical | 91.3 | 100 | 2.09 | 98.3526 | 5214947 | 45.33 | 79 | 145985 |
| SAMN20527495-GCA_019563415.1 | PNUSAV001878 | SAMN20527495 | GCA_019563415.1 | NonPC | Public | 2021 | USA | Clinical | 90.5 | 100 | 3.6 | 98.3065 | 5308096 | 45.26 | 86 | 164477 |
| SAMN20527503-GCA_019563675.1 | PNUSAV001831 | SAMN20527503 | GCA_019563675.1 | NonPC | Public | 2021 | USA | Clinical | 91.4 | 100 | 0.48 | 98.2915 | 5168523 | 45.25 | 42 | 765769 |
| SAMN20591109-GCA_019563195.1 | PNUSAV001881 | SAMN20591109 | GCA_019563195.1 | NonPC | Public | 2021 | USA | Clinical | 89.7 | 100 | 0.12 | 98.2861 | 5067438 | 45.38 | 244 | 45471 |
| SAMN20673774-GCA_019496925.1 | PNUSAV001751 | SAMN20673774 | GCA_019496925.1 | NonPC | Public | 2021 | USA | Clinical | 91.3 | 100 | 3.34 | 98.326 | 5407305 | 45.3 | 62 | 179192 |
| SAMN20673775-GCA_019496045.1 | PNUSAV001805 | SAMN20673775 | GCA_019496045.1 | NonPC | Public | 2021 | USA | Clinical | 89.1 | 100 | 2.72 | 98.0568 | 5255998 | 45.33 | 88 | 174447 |
| SAMN20673777-GCA_019496325.1 | PNUSAV001803 | SAMN20673777 | GCA_019496325.1 | NonPC | Public | 2021 | USA | Clinical | 89.2 | 100 | 4.36 | 98.0689 | 5280056 | 45.32 | 72 | 217353 |
| SAMN20673779-GCA_019496785.1 | PNUSAV001890 | SAMN20673779 | GCA_019496785.1 | NonPC | Public | 2021 | USA | Clinical | 88.6 | 100 | 3.88 | 98.1342 | 5224471 | 45.22 | 111 | 111231 |
| SAMN20804966-GCA_022290765.1 | PV1 | SAMN20804966 | GCA_022290765.1 | NonPC | Public | 2011 | Colombia | Clinical | 92.7 | 100 | 1.4 | 98.3835 | 5128525 | 45.23 | 51 | 240289 |
| SAMN20804967-GCA_022290605.1 | PV278 | SAMN20804967 | GCA_022290605.1 | PC | Public | 2019 | Colombia | Clinical | 97.4 | 100 | 0.03 | 99.9407 | 5103868 | 45.25 | 46 | 564561 |
| SAMN20804968-GCA_022290575.1 | PV280 | SAMN20804968 | GCA_022290575.1 | PC | Public | 2019 | Colombia | Clinical | 97.5 | 100 | 0.14 | 99.9461 | 5144753 | 45.31 | 42 | 442827 |
| SAMN20804969-GCA_022290415.1 | PV53 | SAMN20804969 | GCA_022290415.1 | PC | Public | 2018 | Colombia | Clinical | 95.9 | 100 | 0.3 | 99.9549 | 5133150 | 45.21 | 54 | 427167 |
| SAMN20804970-GCA_022290445.1 | PV85 | SAMN20804970 | GCA_022290445.1 | PC | Public | 2017 | Colombia | Clinical | 97.5 | 100 | 0.27 | 99.9457 | 5146309 | 45.3 | 47 | 426983 |
| SAMN20805029-GCA_022290955.1 | PV112 | SAMN20805029 | GCA_022290955.1 | NonPC | Public | 2018 | Colombia | Env | 89.4 | 99.93 | 2.92 | 98.2766 | 5495560 | 45.47 | 272 | 171457 |
| SAMN20805030-GCA_022290975.1 | PV11 | SAMN20805030 | GCA_022290975.1 | NonPC | Public | 2012 | Colombia | Env | 90.9 | 100 | 0.89 | 98.4033 | 5036340 | 45.37 | 84 | 164783 |
| SAMN20805031-GCA_022290935.1 | PV156 | SAMN20805031 | GCA_022290935.1 | NonPC | Public | 2019 | Colombia | Env | 88 | 100 | 1.82 | 98.2062 | 4997172 | 45.33 | 91 | 172955 |
| SAMN20805032-GCA_022290755.1 | PV213 | SAMN20805032 | GCA_022290755.1 | NonPC | Public | 2019 | Colombia | Env | 90.3 | 99.97 | 0.79 | 98.1838 | 5247527 | 45.24 | 129 | 152816 |
| SAMN20805033-GCA_022290675.1 | PV221 | SAMN20805033 | GCA_022290675.1 | NonPC | Public | 2019 | Colombia | Env | 88.3 | 100 | 0.77 | 98.2407 | 5053872 | 45.3 | 97 | 162813 |
| SAMN20805034-GCA_022290785.1 | PV235 | SAMN20805034 | GCA_022290785.1 | NonPC | Public | 2019 | Colombia | Env | 88.3 | 100 | 4.82 | 98.2505 | 5059799 | 45.43 | 133 | 209984 |
| SAMN20805035-GCA_022290635.1 | PV161 | SAMN20805035 | GCA_022290635.1 | NonPC | Public | 2019 | Colombia | Env | 88.2 | 100 | 2.37 | 98.2866 | 5040855 | 45.29 | 84 | 272749 |
| SAMN20805036-GCA_022290595.1 | PV17 | SAMN20805036 | GCA_022290595.1 | NonPC | Public | 2018 | Colombia | Env | 89.4 | 99.59 | 0.77 | 98.2266 | 5041174 | 45.5 | 155 | 110458 |
| SAMN20805037-GCA_022290535.1 | PV32 | SAMN20805037 | GCA_022290535.1 | NonPC | Public | 2018 | NA | Env | 89.6 | 100 | 1.87 | 98.289 | 5051964 | 45.34 | 82 | 199318 |
| SAMN20813334-GCA_019644695.1 | PNUSAV001894 | SAMN20813334 | GCA_019644695.1 | NonPC | Public | 2021 | USA | Clinical | 88.6 | 100 | 1.27 | 98.1442 | 5137163 | 45.26 | 172 | 63817 |
| SAMN20849617-GCA_019682055.1 | PNUSAV001945 | SAMN20849617 | GCA_019682055.1 | NonPC | Public | 2021 | USA | Env | 91.2 | 100 | 1.43 | 98.3404 | 5091882 | 45.27 | 34 | 476596 |
| SAMN20849626-GCA_019681915.1 | PNUSAV001937 | SAMN20849626 | GCA_019681915.1 | NonPC | Public | 2021 | USA | Env | 89 | 100 | 4.09 | 98.1126 | 5230646 | 45.39 | 58 | 285834 |
| SAMN20849631-GCA_019682755.1 | PNUSAV001753 | SAMN20849631 | GCA_019682755.1 | NonPC | Public | 2021 | USA | Clinical | 89.1 | 100 | 3.81 | 98.1049 | 5298504 | 45.36 | 97 | 112463 |
| SAMN20849633-GCA_019682495.1 | PNUSAV001954 | SAMN20849633 | GCA_019682495.1 | NonPC | Public | 2021 | USA | Clinical | 89.5 | 100 | 4.14 | 98.3027 | 5276168 | 45.29 | 83 | 122133 |
| SAMN20931408-GCA_019727535.1 | PNUSAV001931 | SAMN20931408 | GCA_019727535.1 | NonPC | Public | 2021 | USA | Clinical | 91.5 | 100 | 0.9 | 98.3747 | 5137869 | 45.28 | 56 | 202163 |
| SAMN20931424-GCA_019727435.1 | PNUSAV002003 | SAMN20931424 | GCA_019727435.1 | NonPC | Public | 2021 | USA | Clinical | 89.8 | 100 | 2.82 | 98.2078 | 5268823 | 45.28 | 59 | 203304 |

| | | | | | | | | | | | | | | | | |
|---|---|---|---|---|---|---|---|---|---|---|---|---|---|---|---|---|
| SAMN20931426-GCA_019727395.1 | PNUSAV002002 | SAMN20931426 | GCA_019727395.1 | NonPC | Public | 2021 | USA | Clinical | 91.4 | 100 | 3.18 | 98.2246 | 5261531 | 45.28 | 51 | 286424 |
| SAMN20970995-GCA_019758735.1 | PNUSAV002031 | SAMN20970995 | GCA_019758735.1 | NonPC | Public | 2021 | USA | Clinical | 91.4 | 100 | 1.63 | 98.2899 | 5220911 | 45.33 | 98 | 101013 |
| SAMN20970997-GCA_019759015.1 | PNUSAV001895 | SAMN20970997 | GCA_019759015.1 | NonPC | Public | 2021 | USA | Clinical | 88 | 100 | 4.56 | 98.2143 | 5402276 | 45.19 | 56 | 336943 |
| SAMN20975111-GCA_019773555.1 | PNUSAV001972 | SAMN20975111 | GCA_019773555.1 | NonPC | Public | 2021 | USA | Env | 91.3 | 100 | 2.09 | 98.2739 | 5158429 | 45.28 | 31 | 439817 |
| SAMN20975113-GCA_019773575.1 | PNUSAV001970 | SAMN20975113 | GCA_019773575.1 | NonPC | Public | 2021 | USA | Env | 89.2 | 100 | 4.23 | 98.2197 | 5419144 | 45.28 | 71 | 256939 |
| SAMN20975115-GCA_019773395.1 | PNUSAV001968 | SAMN20975115 | GCA_019773395.1 | NonPC | Public | 2021 | USA | Env | 91.4 | 100 | 2.9 | 98.2035 | 5286161 | 45.25 | 47 | 432582 |
| SAMN20975121-GCA_019772935.1 | PNUSAV001963 | SAMN20975121 | GCA_019772935.1 | NonPC | Public | 2021 | USA | Env | 88.5 | 100 | 3.97 | 98.1926 | 5259863 | 45.25 | 48 | 380212 |
| SAMN21037992-GCA_019808395.1 | PNUSAV001875 | SAMN21037992 | GCA_019808395.1 | NonPC | Public | 2021 | USA | Clinical | 89 | 100 | 2.79 | 98.0857 | 5321562 | 45.4 | 149 | 82574 |
| SAMN21038002-GCA_019804745.1 | PNUSAV002066 | SAMN21038002 | GCA_019804745.1 | NonPC | Public | 2021 | USA | Clinical | 91.3 | 100 | 3.41 | 98.3409 | 5251849 | 45.3 | 50 | 520690 |
| SAMN21161076-GCA_019865405.1 | PNUSAV002044 | SAMN21161076 | GCA_019865405.1 | NonPC | Public | 2021 | USA | Clinical | 88.8 | 100 | 0.27 | 98.1204 | 5052590 | 45.41 | 216 | 50116 |
| SAMN21161077-GCA_019835005.1 | PNUSAV002043 | SAMN21161077 | GCA_019835005.1 | NonPC | Public | 2021 | USA | Clinical | 90.5 | 100 | 1.52 | 98.2623 | 5484002 | 45.11 | 152 | 72139 |
| SAMN21169431-GCA_019846875.1 | PNUSAV002087 | SAMN21169431 | GCA_019846875.1 | NonPC | Public | 2021 | USA | Clinical | 89.7 | 100 | 0.88 | 98.2717 | 5214176 | 45.19 | 60 | 269053 |
| SAMN21330656-GCA_019902005.1 | PNUSAV001877 | SAMN21330656 | GCA_019902005.1 | NonPC | Public | 2021 | USA | Clinical | 89 | 100 | 3.54 | 98.0967 | 5251729 | 45.36 | 132 | 77135 |
| SAMN21330658-GCA_019902965.1 | PNUSAV001892 | SAMN21330658 | GCA_019902965.1 | NonPC | Public | 2021 | USA | Clinical | 91.3 | 100 | 4.9 | 98.3126 | 5384349 | 45.3 | 61 | 180914 |
| SAMN21330660-GCA_019902825.1 | PNUSAV002095 | SAMN21330660 | GCA_019902825.1 | NonPC | Public | 2021 | USA | Clinical | 91.4 | 100 | 4.9 | 98.3204 | 5437406 | 45.3 | 43 | 564960 |
| SAMN21330661-GCA_019902045.1 | PNUSAV002093 | SAMN21330661 | GCA_019902045.1 | NonPC | Public | 2021 | USA | Clinical | 90.6 | 100 | 2.26 | 98.2796 | 5232026 | 45.27 | 60 | 318753 |
| SAMN21365011-GCA_020041945.1 | 20-082E4 | SAMN21365011 | GCA_020041945.1 | NonPC | Public | 2021 | South_Korea | Env | 92.8 | 100 | 0.34 | 98.2539 | 5621370 | 45.29 | 5 | 3610911 |
| SAMN21385800-GCA_019956675.1 | PNUSAV002141 | SAMN21385800 | GCA_019956675.1 | NonPC | Public | NA | USA | Clinical | 87.7 | 100 | 4.1 | 98.092 | 5493291 | 45.28 | 67 | 229287 |
| SAMN21391397-GCA_019945295.1 | PNUSAV002125 | SAMN21391397 | GCA_019945295.1 | NonPC | Public | 2021 | USA | Clinical | 89.8 | 100 | 3.03 | 98.2873 | 5259102 | 45.3 | 85 | 129071 |
| SAMN21391399-GCA_019944595.1 | PNUSAV002124 | SAMN21391399 | GCA_019944595.1 | NonPC | Public | 2021 | USA | Clinical | 91.2 | 100 | 1.7 | 98.3673 | 5117652 | 45.35 | 151 | 82013 |
| SAMN21391400-GCA_019945525.1 | PNUSAV002123 | SAMN21391400 | GCA_019945525.1 | NonPC | Public | 2021 | USA | Clinical | 91.4 | 100 | 0.96 | 98.2984 | 5223393 | 45.3 | 84 | 145275 |
| SAMN21391402-GCA_019946455.1 | PNUSAV002122 | SAMN21391402 | GCA_019946455.1 | NonPC | Public | 2021 | USA | Clinical | 91.4 | 100 | 2.79 | 98.3205 | 5215007 | 45.3 | 101 | 102317 |
| SAMN21439355-GCA_020006745.1 | PNUSAV001893 | SAMN21439355 | GCA_020006745.1 | NonPC | Public | 2021 | USA | Clinical | 89.1 | 100 | 4.63 | 98.0734 | 5265448 | 45.33 | 104 | 122185 |
| SAMN21439356-GCA_020010845.1 | PNUSAV001935 | SAMN21439356 | GCA_020010845.1 | NonPC | Public | 2021 | USA | Clinical | 89.1 | 100 | 3.27 | 98.1099 | 5262812 | 45.34 | 101 | 115258 |
| SAMN21439358-GCA_020006805.1 | PNUSAV001933 | SAMN21439358 | GCA_020006805.1 | NonPC | Public | 2021 | USA | Clinical | 89 | 100 | 3.61 | 98.0832 | 5196205 | 45.35 | 147 | 68400 |
| SAMN21439361-GCA_020006785.1 | PNUSAV002145 | SAMN21439361 | GCA_020006785.1 | NonPC | Public | 2021 | USA | Clinical | 91.2 | 100 | 2.72 | 98.2931 | 5274513 | 45.36 | 159 | 67688 |
| SAMN21442671-GCA_020010345.1 | PNUSAV001836 | SAMN21442671 | GCA_020010345.1 | NonPC | Public | 2021 | USA | Clinical | 89.7 | 100 | 2.29 | 98.3052 | 5280828 | 45.22 | 52 | 219571 |
| SAMN21442673-GCA_020010425.1 | PNUSAV001834 | SAMN21442673 | GCA_020010425.1 | NonPC | Public | 2021 | USA | Clinical | 92.2 | 100 | 4.18 | 98.2706 | 5567354 | 45.2 | 69 | 196169 |
| SAMN21442675-GCA_020010325.1 | PNUSAV001871 | SAMN21442675 | GCA_020010325.1 | NonPC | Public | 2021 | USA | Clinical | 91.3 | 100 | 2.45 | 98.3124 | 5217610 | 45.32 | 75 | 179788 |
| SAMN21531120-GCA_020086975.1 | PNUSAV002038 | SAMN21531120 | GCA_020086975.1 | NonPC | Public | 2021 | USA | Clinical | 90.2 | 100 | 3.18 | 98.285 | 5193661 | 45.38 | 121 | 93066 |
| SAMN21531121-GCA_020087165.1 | PNUSAV002088 | SAMN21531121 | GCA_020087165.1 | NonPC | Public | 2021 | USA | Clinical | 91.2 | 100 | 4.63 | 98.2862 | 5386688 | 45.31 | 95 | 112572 |
| SAMN21531122-GCA_020087535.1 | PNUSAV002099 | SAMN21531122 | GCA_020087535.1 | NonPC | Public | 2021 | USA | Clinical | 89 | 100 | 3.54 | 98.1231 | 5272838 | 45.39 | 150 | 76221 |
| SAMN21531123-GCA_020086655.1 | PNUSAV002100 | SAMN21531123 | GCA_020086655.1 | NonPC | Public | 2021 | USA | Clinical | 91.1 | 100 | 3 | 98.2733 | 5337614 | 45.33 | 136 | 77437 |
| SAMN21531127-GCA_020086875.1 | PNUSAV002155 | SAMN21531127 | GCA_020086875.1 | NonPC | Public | 2020 | USA | Clinical | 91.3 | 100 | 4.9 | 98.3168 | 5286726 | 45.31 | 62 | 201532 |
| SAMN21531128-GCA_020087505.1 | PNUSAV002151 | SAMN21531128 | GCA_020087505.1 | NonPC | Public | 2020 | USA | Clinical | 91.3 | 100 | 1.54 | 98.3281 | 5183929 | 45.29 | 66 | 177609 |
| SAMN21531130-GCA_020086915.1 | PNUSAV002167 | SAMN21531130 | GCA_020086915.1 | NonPC | Public | 2021 | USA | Clinical | 91.3 | 100 | 4.09 | 98.3266 | 5285475 | 45.31 | 65 | 225109 |
| SAMN21531131-GCA_020087095.1 | PNUSAV002166 | SAMN21531131 | GCA_020087095.1 | NonPC | Public | 2021 | USA | Clinical | 91.1 | 100 | 2.72 | 98.3175 | 5199609 | 45.38 | 173 | 65318 |
| SAMN21531132-GCA_020087565.1 | PNUSAV002189 | SAMN21531132 | GCA_020087565.1 | NonPC | Public | 2021 | USA | Clinical | 91.4 | 100 | 3.86 | 98.3203 | 5338329 | 45.27 | 43 | 564960 |
| SAMN21578974-GCA_020124555.1 | PNUSAV002129 | SAMN21578974 | GCA_020124555.1 | NonPC | Public | 2021 | USA | Env | 91.1 | 100 | 1.77 | 98.2255 | 5277217 | 45.26 | 53 | 390331 |
| SAMN21578981-GCA_020124955.1 | PNUSAV002130 | SAMN21578981 | GCA_020124955.1 | NonPC | Public | 2021 | USA | Env | 90.2 | 100 | 1.89 | 98.1678 | 5537245 | 45.09 | 57 | 241621 |
| SAMN21578983-GCA_020124935.1 | PNUSAV002134 | SAMN21578983 | GCA_020124935.1 | NonPC | Public | 2021 | USA | Env | 90.9 | 100 | 2.23 | 98.2881 | 5318025 | 45.23 | 48 | 517090 |
| SAMN21578987-GCA_020125155.1 | PNUSAV002133 | SAMN21578987 | GCA_020125155.1 | NonPC | Public | 2021 | USA | Env | 91 | 100 | 2.07 | 98.2564 | 5357789 | 45.22 | 56 | 271916 |
| SAMN21578991-GCA_020125295.1 | PNUSAV002132 | SAMN21578991 | GCA_020125295.1 | NonPC | Public | 2021 | USA | Env | 90.1 | 100 | 1.55 | 98.1819 | 5345312 | 45.13 | 57 | 254962 |
| SAMN21578994-GCA_020124895.1 | PNUSAV002137 | SAMN21578994 | GCA_020124895.1 | NonPC | Public | 2021 | USA | Env | 91.3 | 100 | 1.92 | 98.282 | 5520855 | 45.01 | 89 | 220025 |
| SAMN21855653-GCA_020190785.1 | PNUSAV002188 | SAMN21855653 | GCA_020190785.1 | NonPC | Public | 2021 | USA | Clinical | 88.7 | 100 | 0.34 | 98.0573 | 5120227 | 45.3 | 192 | 51890 |
| SAMN21855654-GCA_020209705.1 | PNUSAV002220 | SAMN21855654 | GCA_020209705.1 | NonPC | Public | 2021 | USA | Clinical | 90.9 | 100 | 2.17 | 98.2881 | 5281658 | 45.29 | 49 | 449037 |
| SAMN21855655-GCA_020190745.1 | PNUSAV002219 | SAMN21855655 | GCA_020190745.1 | PC | Public | 2021 | USA | Clinical | 98.2 | 100 | 2.26 | 99.9897 | 5348087 | 45.13 | 53 | 265864 |
| SAMN21897405-GCA_020223155.1 | PNUSAV002224 | SAMN21897405 | GCA_020223155.1 | NonPC | Public | 2021 | USA | Clinical | 90.8 | 100 | 1.03 | 98.2659 | 5022821 | 45.33 | 79 | 189317 |

| | | | | | | | | | | | | | | | | |
|---|---|---|---|---|---|---|---|---|---|---|---|---|---|---|---|---|
| SAMN21897422-GCA_020222875.1 | PNUSAV002248 | SAMN21897422 | GCA_020222875.1 | NonPC | Public | 2021 | USA | Clinical | 90.4 | 100 | 1.88 | 98.3179 | 5090658 | 45.28 | 34 | 457208 |
| SAMN21974751-GCA_020238465.1 | PNUSAV002246 | SAMN21974751 | GCA_020238465.1 | NonPC | Public | 2021 | USA | Clinical | 89.5 | 100 | 0.85 | 98.2945 | 5078805 | 45.36 | 149 | 65898 |
| SAMN21974752-GCA_020260225.1 | PNUSAV002257 | SAMN21974752 | GCA_020260225.1 | NonPC | Public | 2021 | USA | Clinical | 91.4 | 100 | 1.16 | 98.2972 | 5264397 | 45.33 | 69 | 267226 |
| SAMN21974753-GCA_020260445.1 | PNUSAV002256 | SAMN21974753 | GCA_020260445.1 | NonPC | Public | 2021 | USA | Clinical | 90.5 | 100 | 0.82 | 98.2753 | 5159737 | 45.38 | 156 | 107389 |
| SAMN21974754-GCA_020261025.1 | PNUSAV002258 | SAMN21974754 | GCA_020261025.1 | NonPC | Public | 2021 | USA | Clinical | 88.8 | 100 | 1.56 | 98.1498 | 5143195 | 45.25 | 105 | 111278 |
| SAMN21974755-GCA_020276785.1 | PNUSAV002265 | SAMN21974755 | GCA_020276785.1 | NonPC | Public | 2021 | USA | Clinical | 89.9 | 100 | 0.58 | 98.2241 | 5210544 | 45.18 | 83 | 406583 |
| SAMN21974756-GCA_020261425.1 | PNUSAV002252 | SAMN21974756 | GCA_020261425.1 | NonPC | Public | 2021 | USA | Clinical | 89.7 | 100 | 0.24 | 98.2417 | 5216802 | 45.22 | 89 | 109456 |
| SAMN22043428-GCA_020280225.1 | PNUSAV002217 | SAMN22043428 | GCA_020280225.1 | NonPC | Public | 2021 | USA | Clinical | 90.9 | 100 | 0.89 | 98.3436 | 5200780 | 45.18 | 51 | 406519 |
| SAMN22043429-GCA_020280185.1 | PNUSAV002216 | SAMN22043429 | GCA_020280185.1 | NonPC | Public | 2021 | USA | Clinical | 89.9 | 100 | 0.12 | 98.2381 | 5391119 | 45.21 | 46 | 464232 |
| SAMN22043430-GCA_020281585.1 | PNUSAV002244 | SAMN22043430 | GCA_020281585.1 | NonPC | Public | 2021 | USA | Clinical | 91.5 | 100 | 1.16 | 98.2849 | 5174541 | 45.28 | 51 | 236871 |
| SAMN22043431-GCA_020281605.1 | PNUSAV002254 | SAMN22043431 | GCA_020281605.1 | NonPC | Public | 2021 | USA | Clinical | 89.8 | 100 | 2.84 | 98.2242 | 5557712 | 45.24 | 49 | 227933 |
| SAMN22163831-GCA_020407345.1 | PNUSAV002271 | SAMN22163831 | GCA_020407345.1 | NonPC | Public | NA | USA | Clinical | 90.7 | 100 | 3.74 | 98.3234 | 5341130 | 45.2 | 56 | 349954 |
| SAMN22165054-GCA_020407235.1 | PNUSAV002148 | SAMN22165054 | GCA_020407235.1 | NonPC | Public | 2021 | USA | Clinical | 91 | 100 | 1.14 | 98.2936 | 5095263 | 45.33 | 72 | 294166 |
| SAMN22168687-GCA_020413625.1 | 2018V-1201 | SAMN22168687 | GCA_020413625.1 | NonPC | Public | 2018 | USA | Clinical | 91 | 100 | 0.27 | 98.2763 | 5119973 | 45.33 | 178 | 74319 |
| SAMN22169081-GCA_020413545.1 | 2017V-1208 | SAMN22169081 | GCA_020413545.1 | NonPC | Public | 2017 | USA | Clinical | 91.3 | 100 | 4.9 | 98.2912 | 5311585 | 45.28 | 69 | 181089 |
| SAMN22169082-GCA_020413465.1 | 2017V-1207 | SAMN22169082 | GCA_020413465.1 | NonPC | Public | 2017 | USA | Clinical | 91.3 | 100 | 4.22 | 98.2849 | 5279438 | 45.27 | 75 | 185286 |
| SAMN22169084-GCA_020412765.1 | PNUSAV001340 | SAMN22169084 | GCA_020412765.1 | NonPC | Public | 2020 | USA | Clinical | 89.5 | 100 | 1.7 | 98.2426 | 5286997 | 45.19 | 48 | 302443 |
| SAMN22169085-GCA_020413525.1 | PNUSAV001341 | SAMN22169085 | GCA_020413525.1 | NonPC | Public | 2020 | USA | Clinical | 89.7 | 100 | 0.72 | 98.2469 | 5315754 | 45.21 | 77 | 186678 |
| SAMN22169086-GCA_020413565.1 | PNUSAV001675 | SAMN22169086 | GCA_020413565.1 | NonPC | Public | 2021 | USA | Clinical | 88.9 | 100 | 2.18 | 98.073 | 5163423 | 45.37 | 138 | 76886 |
| SAMN22169087-GCA_020413445.1 | PNUSAV001702 | SAMN22169087 | GCA_020413445.1 | NonPC | Public | 2021 | USA | Clinical | 90.2 | 100 | 3 | 98.2843 | 5178433 | 45.34 | 108 | 105156 |
| SAMN22169088-GCA_020413425.1 | PNUSAV001703 | SAMN22169088 | GCA_020413425.1 | NonPC | Public | 2021 | USA | Clinical | 90.4 | 100 | 4.09 | 98.3334 | 5300778 | 45.3 | 42 | 417178 |
| SAMN22169442-GCA_020412105.1 | PNUSAV001484 | SAMN22169442 | GCA_020412105.1 | NonPC | Public | 2020 | USA | Clinical | 88.2 | 100 | 3.87 | 98.085 | 5175463 | 45.34 | 97 | 102699 |
| SAMN22183809-GCA_020411165.1 | PNUSAV002026 | SAMN22183809 | GCA_020411165.1 | NonPC | Public | 2021 | USA | Clinical | 92 | 100 | 4.62 | 98.3548 | 5446539 | 45.11 | 37 | 440788 |
| SAMN22183811-GCA_020410905.1 | PNUSAV001371 | SAMN22183811 | GCA_020410905.1 | NonPC | Public | 2020 | USA | Clinical | 88.4 | 100 | 1.09 | 98.1114 | 5011386 | 45.41 | 193 | 55594 |
| SAMN22183812-GCA_020410985.1 | PNUSAV001372 | SAMN22183812 | GCA_020410985.1 | NonPC | Public | 2020 | USA | Clinical | 91.4 | 100 | 4.9 | 98.312 | 5340141 | 45.31 | 90 | 123361 |
| SAMN22183817-GCA_020411065.1 | PNUSAV001338 | SAMN22183817 | GCA_020411065.1 | NonPC | Public | 2020 | USA | Clinical | 91.3 | 100 | 3.27 | 98.2929 | 5257978 | 45.38 | 154 | 73157 |
| SAMN22183818-GCA_020411005.1 | PNUSAV001337 | SAMN22183818 | GCA_020411005.1 | NonPC | Public | 2020 | USA | Clinical | 91.2 | 100 | 0.7 | 98.2629 | 5236366 | 45.37 | 187 | 57041 |
| SAMN22183826-GCA_020410685.1 | PNUSAV001661 | SAMN22183826 | GCA_020410685.1 | NonPC | Public | 2021 | USA | Clinical | 89 | 100 | 3.89 | 98.1031 | 5281112 | 45.31 | 64 | 233237 |
| SAMN22183828-GCA_020410605.1 | PNUSAV001732 | SAMN22183828 | GCA_020410605.1 | NonPC | Public | 2021 | USA | Clinical | 91.3 | 100 | 1.71 | 98.3489 | 5190203 | 45.25 | 42 | 562891 |
| SAMN22183830-GCA_020410645.1 | PNUSAV002039 | SAMN22183830 | GCA_020410645.1 | NonPC | Public | 2021 | USA | Clinical | 89.2 | 100 | 3.68 | 98.0615 | 5322587 | 45.13 | 64 | 234049 |
| SAMN22183833-GCA_020410625.1 | PNUSAV002221 | SAMN22183833 | GCA_020410625.1 | NonPC | Public | 2021 | USA | Clinical | 89.9 | 100 | 4.08 | 98.2784 | 5419512 | 45.23 | 37 | 546959 |
| SAMN22183960-GCA_020410425.1 | PNUSAV001903 | SAMN22183960 | GCA_020410425.1 | NonPC | Public | 2021 | USA | Clinical | 90.4 | 100 | 2.78 | 98.2369 | 5246842 | 45.26 | 104 | 116002 |
| SAMN22184125-GCA_020410405.1 | PNUSAV002215 | SAMN22184125 | GCA_020410405.1 | NonPC | Public | 2021 | USA | Clinical | 89.1 | 100 | 3.81 | 98.1031 | 5282861 | 45.32 | 76 | 232904 |
| SAMN22211694-GCA_020782035.1 | FXH250 | SAMN22211694 | GCA_020782035.1 | NonPC | Public | 2021 | China | Env | 89.7 | 100 | 0 | 98.2792 | 5082011 | 45.3 | 77 | 459598 |
| SAMN22211696-GCA_020782215.1 | FXH-72 | SAMN22211696 | GCA_020782215.1 | NonPC | Public | 2021 | China | Env | 91.5 | 100 | 0 | 98.4308 | 5020685 | 45.37 | 59 | 457002 |
| SAMN22231096-GCA_020452565.1 | PNUSAV002091 | SAMN22231096 | GCA_020452565.1 | NonPC | Public | 2021 | USA | Clinical | 91.4 | 100 | 3.09 | 98.3084 | 5226534 | 45.29 | 88 | 141620 |
| SAMN22231102-GCA_020452465.1 | PNUSAV002186 | SAMN22231102 | GCA_020452465.1 | NonPC | Public | 2021 | USA | Clinical | 89.2 | 100 | 3 | 98.0802 | 5250975 | 45.32 | 78 | 205543 |
| SAMN22231103-GCA_020452385.1 | PNUSAV002274 | SAMN22231103 | GCA_020452385.1 | NonPC | Public | 2021 | USA | Clinical | 91.3 | 100 | 3.81 | 98.2843 | 5221418 | 45.35 | 137 | 74287 |
| SAMN22231113-GCA_020452185.1 | PNUSAV002280 | SAMN22231113 | GCA_020452185.1 | NonPC | Public | 2021 | USA | Clinical | 91.3 | 100 | 0.07 | 98.306 | 5116930 | 45.3 | 139 | 78855 |
| SAMN22231273-GCA_020451645.1 | PNUSAV001276 | SAMN22231273 | GCA_020451645.1 | NonPC | Public | 2020 | USA | Clinical | 91.3 | 100 | 0.07 | 98.3056 | 5101756 | 45.29 | 91 | 123839 |
| SAMN22231277-GCA_020451545.1 | PNUSAV001652 | SAMN22231277 | GCA_020451545.1 | NonPC | Public | 2021 | USA | Clinical | 91.4 | 100 | 0.34 | 98.2967 | 5140058 | 45.27 | 42 | 475295 |
| SAMN22231280-GCA_020451665.1 | PNUSAV001778 | SAMN22231280 | GCA_020451665.1 | NonPC | Public | 2021 | USA | Clinical | 91.3 | 100 | 0.36 | 98.2962 | 5142812 | 45.26 | 84 | 147925 |
| SAMN22231282-GCA_020451525.1 | PNUSAV001929 | SAMN22231282 | GCA_020451525.1 | NonPC | Public | 2021 | USA | Clinical | 91.4 | 100 | 0.91 | 98.3111 | 5204251 | 45.25 | 56 | 315090 |
| SAMN22231284-GCA_020451505.1 | PNUSAV001901 | SAMN22231284 | GCA_020451505.1 | NonPC | Public | 2021 | USA | Clinical | 90.4 | 100 | 0.54 | 98.2981 | 5092651 | 45.31 | 174 | 80338 |
| SAMN22231286-GCA_020451485.1 | PNUSAV002025 | SAMN22231286 | GCA_020451485.1 | PC | Public | 2021 | USA | Clinical | 98.2 | 100 | 2.25 | 99.988 | 5259394 | 45.32 | 55 | 274090 |
| SAMN22231287-GCA_020451625.1 | PNUSAV002089 | SAMN22231287 | GCA_020451625.1 | NonPC | Public | 2021 | USA | Clinical | 89.1 | 100 | 3.81 | 98.088 | 5295812 | 45.24 | 153 | 89708 |
| SAMN22231991-GCA_020451585.1 | PNUSAV002140 | SAMN22231991 | GCA_020451585.1 | NonPC | Public | 2021 | USA | Clinical | 91.3 | 100 | 0.7 | 98.3363 | 5119902 | 45.33 | 108 | 109269 |
| SAMN22231992-GCA_020451605.1 | PNUSAV002245 | SAMN22231992 | GCA_020451605.1 | NonPC | Public | 2021 | USA | Clinical | 91.4 | 100 | 1.09 | 98.3007 | 5211732 | 45.33 | 107 | 131847 |

| | | | | | | | | | | | | | | | | |
|---|---|---|---|---|---|---|---|---|---|---|---|---|---|---|---|---|
| SAMN22231994-GCA_020451565.1 | PNUSAV001900 | SAMN22231994 | GCA_020451565.1 | NonPC | Public | 2021 | USA | Clinical | 88.9 | 100 | 0.34 | 98.0935 | 5032941 | 45.34 | 174 | 67738 |
| SAMN22231995-GCA_020451285.1 | PNUSAV001782 | SAMN22231995 | GCA_020451285.1 | NonPC | Public | 2021 | USA | Clinical | 91.3 | 100 | 3.81 | 98.2929 | 5318159 | 45.3 | 70 | 153356 |
| SAMN22231996-GCA_020451425.1 | PNUSAV001723 | SAMN22231996 | GCA_020451425.1 | NonPC | Public | 2021 | USA | Clinical | 90.1 | 100 | 1.3 | 98.2152 | 5184371 | 45.2 | 139 | 88619 |
| SAMN22253494-SRR16324072 | PNUSAV001504 | SAMN22253494 | SRR16324072 | NonPC | Public | 2020 | USA | Env | 91.7 | 100 | 0.13 | 98.2127 | 5220406 | 45.18 | 41 | 608784 |
| SAMN22253495-SRR16324052 | PNUSAV001499 | SAMN22253495 | SRR16324052 | NonPC | Public | 2020 | USA | Env | 88.3 | 100 | 0.36 | 98.2528 | 5018108 | 45.31 | 25 | 521086 |
| SAMN22253496-SRR16324073 | PNUSAV001498 | SAMN22253496 | SRR16324073 | NonPC | Public | 2020 | USA | Env | 91.3 | 100 | 0.14 | 98.1894 | 5161698 | 45.21 | 46 | 606352 |
| SAMN22253497-SRR16324069 | PNUSAV001501 | SAMN22253497 | SRR16324069 | NonPC | Public | 2020 | USA | Env | 91.6 | 100 | 0.27 | 98.342 | 5041775 | 45.31 | 28 | 816461 |
| SAMN22253498-SRR16324061 | PNUSAV001502 | SAMN22253498 | SRR16324061 | NonPC | Public | 2020 | USA | Env | 89 | 100 | 0.44 | 98.0574 | 5391975 | 45.3 | 64 | 337890 |
| SAMN22253499-SRR16324059 | PNUSAV001503 | SAMN22253499 | SRR16324059 | NonPC | Public | 2020 | USA | Env | 91.3 | 100 | 0.14 | 98.2098 | 5182974 | 45.26 | 55 | 535811 |
| SAMN22253500-SRR16324063 | PNUSAV001507 | SAMN22253500 | SRR16324063 | NonPC | Public | 2020 | USA | Env | 90.1 | 100 | 0.19 | 98.283 | 5101304 | 45.36 | 31 | 1143840 |
| SAMN22253501-SRR16324060 | PNUSAV001506 | SAMN22253501 | SRR16324060 | NonPC | Public | 2020 | USA | Env | 92 | 100 | 0.14 | 98.2966 | 5009201 | 45.38 | 27 | 749222 |
| SAMN22253502-SRR16324077 | PNUSAV001505 | SAMN22253502 | SRR16324077 | NonPC | Public | 2020 | USA | Env | 91.2 | 100 | 0 | 98.2739 | 5069126 | 45.32 | 32 | 402561 |
| SAMN22253503-GCA_020502885.1 | PNUSAV000636 | SAMN22253503 | GCA_020502885.1 | NonPC | Public | 2019 | USA | Clinical | 91.3 | 100 | 3.68 | 98.3026 | 5272029 | 45.3 | 85 | 140952 |
| SAMN22253504-GCA_020509945.1 | PNUSAV001419 | SAMN22253504 | GCA_020509945.1 | NonPC | Public | 2020 | USA | Clinical | 89.6 | 100 | 4.59 | 98.1403 | 5309095 | 45.33 | 73 | 168494 |
| SAMN22330461-GCA_020513195.1 | PNUSAV000136 | SAMN22330461 | GCA_020513195.1 | NonPC | Public | 2017 | USA | Clinical | 87.1 | 100 | 0.66 | 98.0534 | 5028589 | 45.26 | 95 | 105679 |
| SAMN22330470-GCA_020514125.1 | PNUSAV002187 | SAMN22330470 | GCA_020514125.1 | NonPC | Public | 2021 | USA | Clinical | 90.6 | 100 | 1.51 | 98.3209 | 5260261 | 45.28 | 159 | 82162 |
| SAMN22330483-GCA_020520085.1 | PNUSAV002266 | SAMN22330483 | GCA_020520085.1 | NonPC | Public | 2021 | USA | Clinical | 91.2 | 100 | 3.35 | 98.2997 | 5278186 | 45.34 | 112 | 109042 |
| SAMN22366332-GCA_020521175.1 | PNUSAV002284 | SAMN22366332 | GCA_020521175.1 | NonPC | Public | 2021 | USA | Clinical | 89.5 | 100 | 3.59 | 98.2464 | 5536836 | 45.32 | 150 | 74223 |
| SAMN22426622-GCA_020586515.1 | PNUSAV002309 | SAMN22426622 | GCA_020586515.1 | NonPC | Public | 2021 | USA | Clinical | 89.7 | 100 | 1.14 | 98.2299 | 5461693 | 45.22 | 53 | 349737 |
| SAMN22481326-GCA_020550985.1 | PNUSAV002314 | SAMN22481326 | GCA_020550985.1 | NonPC | Public | 2021 | USA | Clinical | 92.3 | 100 | 2.18 | 98.3322 | 5313274 | 45.25 | 48 | 343053 |
| SAMN22560587-GCA_020614455.1 | PNUSAV002283 | SAMN22560587 | GCA_020614455.1 | NonPC | Public | 2021 | USA | Clinical | 89.7 | 100 | 2.29 | 98.283 | 5393368 | 45.22 | 48 | 341500 |
| SAMN22561829-GCA_020679505.1 | PNUSAV002336 | SAMN22561829 | GCA_020679505.1 | NonPC | Public | 2021 | USA | Clinical | 90.9 | 100 | 0.61 | 98.2928 | 5113817 | 45.38 | 197 | 65278 |
| SAMN22561830-GCA_020610635.1 | PNUSAV002335 | SAMN22561830 | GCA_020610635.1 | NonPC | Public | 2021 | USA | Clinical | 91 | 100 | 0.08 | 98.3019 | 5170203 | 45.45 | 149 | 93741 |
| SAMN22561831-GCA_020610675.1 | PNUSAV002342 | SAMN22561831 | GCA_020610675.1 | NonPC | Public | 2021 | USA | Clinical | 91.9 | 100 | 1.57 | 98.3126 | 5190765 | 45.24 | 98 | 138443 |
| SAMN22561832-GCA_020610535.1 | PNUSAV002341 | SAMN22561832 | GCA_020610535.1 | NonPC | Public | 2021 | USA | Clinical | 90.9 | 100 | 3.27 | 98.4417 | 5298911 | 45.34 | 112 | 82387 |
| SAMN22561833-GCA_020610575.1 | PNUSAV002340 | SAMN22561833 | GCA_020610575.1 | NonPC | Public | 2021 | USA | Clinical | 89.7 | 100 | 3.85 | 98.2536 | 5383872 | 45.25 | 97 | 105656 |
| SAMN22672838-GCA_020701435.1 | PNUSAV002343 | SAMN22672838 | GCA_020701435.1 | NonPC | Public | 2021 | USA | Clinical | 90.4 | 100 | 1.85 | 98.2593 | 5253114 | 45.14 | 167 | 81983 |
| SAMN22866281-GCA_020764655.1 | PNUSAV002218 | SAMN22866281 | GCA_020764655.1 | NonPC | Public | 2021 | USA | Clinical | 90.6 | 100 | 0 | 98.319 | 5053011 | 45.27 | 65 | 450563 |
| SAMN23007929-GCA_020812655.1 | PNUSAV002351 | SAMN23007929 | GCA_020812655.1 | NonPC | Public | 2021 | USA | Clinical | 89.7 | 100 | 1.81 | 98.2877 | 5281242 | 45.23 | 39 | 438995 |
| SAMN23007930-GCA_020812115.1 | PNUSAV002350 | SAMN23007930 | GCA_020812115.1 | NonPC | Public | 2021 | USA | Clinical | 89.8 | 100 | 1.53 | 98.2129 | 5232594 | 45.28 | 40 | 460843 |
| SAMN23012431-GCA_020825235.1 | PNUSAV002372 | SAMN23012431 | GCA_020825235.1 | NonPC | Public | 2021 | USA | Clinical | 89.3 | 100 | 2.23 | 98.2366 | 5426844 | 45.33 | 190 | 57347 |
| SAMN23012432-GCA_020825295.1 | PNUSAV002371 | SAMN23012432 | GCA_020825295.1 | NonPC | Public | 2021 | USA | Clinical | 91.5 | 100 | 3.54 | 98.2439 | 5262548 | 45.38 | 143 | 71116 |
| SAMN23166992-GCA_020958835.1 | PNUSAV002375 | SAMN23166992 | GCA_020958835.1 | NonPC | Public | 2021 | USA | Clinical | 90 | 100 | 4.68 | 98.2911 | 5397800 | 45.39 | 40 | 320649 |
| SAMN23211423-GCA_020944785.1 | PNUSAV002288 | SAMN23211423 | GCA_020944785.1 | NonPC | Public | 2021 | USA | Clinical | 91.4 | 100 | 3.61 | 98.3067 | 5330319 | 45.3 | 71 | 195042 |
| SAMN23215745-GCA_020943985.1 | PNUSAV002287 | SAMN23215745 | GCA_020943985.1 | NonPC | Public | 2021 | USA | Clinical | 90.6 | 100 | 0.67 | 98.3299 | 5234654 | 45.23 | 141 | 83208 |
| SAMN23215746-GCA_020944005.1 | PNUSAV002346 | SAMN23215746 | GCA_020944005.1 | PC | Public | 2021 | USA | Clinical | 98.1 | 100 | 1.91 | 99.9833 | 5252326 | 45.32 | 103 | 105834 |
| SAMN23284737-GCA_020896115.1 | 21MO06SH01-V1 | SAMN23284737 | GCA_020896115.1 | NonPC | Public | 2021 | NA | Env | 91.7 | 100 | 2.38 | 98.3759 | 5383190 | 45.12 | 68 | 194487 |
| SAMN23391200-GCA_020961675.1 | 21MO07SH01-V1 | SAMN23391200 | GCA_020961675.1 | NonPC | Public | 2021 | USA | Env | 89.7 | 100 | 4.58 | 98.2947 | 5513360 | 45.35 | 47 | 399580 |
| SAMN23407501-GCA_020972685.1 | 21MO09SH01-V1 | SAMN23407501 | GCA_020972685.1 | NonPC | Public | 2021 | USA | Env | 90.2 | 100 | 1.83 | 98.3242 | 5416507 | 45.16 | 46 | 463502 |
| SAMN23407504-GCA_020962915.1 | 21MO07SH02-V1 | SAMN23407504 | GCA_020962915.1 | NonPC | Public | 2021 | USA | Env | 88.7 | 100 | 2.63 | 98.0616 | 5642121 | 45.17 | 115 | 153333 |
| SAMN23413467-GCA_020966945.1 | PNUSAV002308 | SAMN23413467 | GCA_020966945.1 | NonPC | Public | 2020 | USA | Clinical | 89.9 | 100 | 4.5 | 98.3431 | 5369875 | 45.31 | 53 | 360798 |
| SAMN23413468-GCA_020966935.1 | PNUSAV002307 | SAMN23413468 | GCA_020966935.1 | NonPC | Public | 2020 | USA | Clinical | 89.9 | 100 | 3.86 | 98.3356 | 5272990 | 45.28 | 52 | 402635 |
| SAMN23413470-GCA_020970115.1 | PNUSAV002306 | SAMN23413470 | GCA_020970115.1 | NonPC | Public | 2020 | USA | Clinical | 89.7 | 100 | 4.94 | 98.255 | 5420787 | 45.28 | 48 | 273053 |
| SAMN23413471-GCA_020970095.1 | PNUSAV002303 | SAMN23413471 | GCA_020970095.1 | NonPC | Public | 2020 | USA | Clinical | 89.8 | 100 | 4.47 | 98.3077 | 5262266 | 45.39 | 50 | 241668 |
| SAMN23413473-GCA_020967495.1 | PNUSAV002290 | SAMN23413473 | GCA_020967495.1 | NonPC | Public | 2020 | USA | Clinical | 89.6 | 100 | 4.58 | 98.3034 | 5285703 | 45.39 | 54 | 182599 |
| SAMN23413475-GCA_020967555.1 | PNUSAV002294 | SAMN23413475 | GCA_020967555.1 | NonPC | Public | 2020 | USA | Clinical | 91.1 | 100 | 1.54 | 98.3259 | 5242867 | 45.29 | 49 | 234322 |
| SAMN23413917-GCA_020969355.1 | PNUSAV002446 | SAMN23413917 | GCA_020969355.1 | NonPC | Public | 2021 | USA | Clinical | 91.4 | 100 | 0.95 | 98.3195 | 5154131 | 45.26 | 58 | 366021 |
| SAMN23425222-GCA_020977645.1 | PNUSAV002406 | SAMN23425222 | GCA_020977645.1 | NonPC | Public | 2021 | USA | Clinical | 91 | 100 | 3.32 | 98.2358 | 5597766 | 45.05 | 126 | 175444 |

| | | | | | | | | | | | | | | | | |
|---|---|---|---|---|---|---|---|---|---|---|---|---|---|---|---|---|
| SAMN23429703-GCA_021026225.1 | ump001v | SAMN23429703 | GCA_021026225.1 | NonPC | Public | 2020 | Malaysia | Env | 90.7 | 100 | 0.17 | 98.3522 | 4992817 | 45.31 | 52 | 471855 |
| SAMN23430655-SRR17035889 | ST417 | SAMN23430655 | SRR17035889 | NonPC | Public | NA | Australia | Env | 90.9 | 100 | 0.33 | 98.3251 | 5112843 | 45.22 | 51 | 255423 |
| SAMN23430656-SRR17035888 | ST50 | SAMN23430656 | SRR17035888 | NonPC | Public | NA | Australia | Clinical | 91.2 | 100 | 0.32 | 98.2668 | 5183178 | 45.16 | 91 | 300171 |
| SAMN23491980-GCA_021002415.1 | PNUSAV002405 | SAMN23491980 | GCA_021002415.1 | NonPC | Public | 2021 | USA | Clinical | 91.3 | 100 | 3.81 | 98.3324 | 5260925 | 45.32 | 79 | 191743 |
| SAMN23852877-GCA_021164445.1 | PNUSAV002452 | SAMN23852877 | GCA_021164445.1 | NonPC | Public | 2021 | USA | Clinical | 90.8 | 100 | 1.93 | 98.2546 | 5262338 | 45.14 | 87 | 305745 |
| SAMN24103581-GCA_021209095.1 | PNUSAV002453 | SAMN24103581 | GCA_021209095.1 | NonPC | Public | 2021 | USA | Clinical | 91 | 100 | 0.07 | 98.3307 | 5080793 | 45.37 | 193 | 59115 |
| SAMN24103582-GCA_021207355.1 | PNUSAV002454 | SAMN24103582 | GCA_021207355.1 | NonPC | Public | 2021 | USA | Clinical | 91.2 | 100 | 0.27 | 98.2888 | 5092260 | 45.24 | 71 | 161381 |
| SAMN24283276-GCA_021260845.1 | PNUSAV002462 | SAMN24283276 | GCA_021260845.1 | NonPC | Public | 2021 | USA | Clinical | 89.2 | 100 | 2.05 | 98.0482 | 5239395 | 45.28 | 74 | 233238 |
| SAMN24283278-GCA_021268105.1 | PNUSAV002461 | SAMN24283278 | GCA_021268105.1 | NonPC | Public | 2021 | USA | Clinical | 91.3 | 100 | 3.12 | 98.2624 | 5262647 | 45.28 | 38 | 556737 |
| SAMN24283279-GCA_021260825.1 | PNUSAV002460 | SAMN24283279 | GCA_021260825.1 | NonPC | Public | 2021 | USA | Clinical | 90.3 | 100 | 1.99 | 98.2312 | 5209339 | 45.21 | 37 | 539657 |
| SAMN24298400-GCA_021271265.1 | PNUSAV002464 | SAMN24298400 | GCA_021271265.1 | NonPC | Public | 2021 | USA | Clinical | 89.8 | 100 | 0.39 | 98.2521 | 5361500 | 45.18 | 45 | 498840 |
| SAMN24611822-SRR17429197 | PNUSAV002463 | SAMN24611822 | SRR17429197 | NonPC | Public | 2021 | USA | NA | 88.5 | 100 | 0.05 | 98.3016 | 4920006 | 45.34 | 75 | 168632 |
| SAMN24777699-GCA_022397635.1 | M13-18718 | SAMN24777699 | GCA_022397635.1 | NonPC | Public | 2013 | Australia | Env | 91.7 | 100 | 0 | 98.3507 | 5138243 | 45.36 | 136 | 83036 |
| SAMN24777700-GCA_022397695.1 | M14-00202 | SAMN24777700 | GCA_022397695.1 | NonPC | Public | 2013 | Australia | Env | 91.3 | 100 | 0 | 98.3971 | 4995260 | 45.38 | 23 | 1161470 |
| SAMN24777701-GCA_022397605.1 | M14-00202b | SAMN24777701 | GCA_022397605.1 | NonPC | Public | 2013 | Australia | Env | 91.1 | 100 | 0.27 | 98.3581 | 5141381 | 45.27 | 30 | 1116173 |
| SAMN25128001-SRR17673829 | ST417 | SAMN25128001 | SRR17673829 | NonPC | Public | 2021 | Australia | Clinical | 90.9 | 100 | 0.33 | 98.3119 | 5114575 | 45.22 | 34 | 747198 |
| SAMN25128002-SRR17673828 | ST417 | SAMN25128002 | SRR17673828 | NonPC | Public | 2021 | Australia | Clinical | 90.9 | 100 | 0.33 | 98.3109 | 5114397 | 45.21 | 36 | 729969 |
| SAMN25134884-SRR17786634 | PNUSAV002475 | SAMN25134884 | SRR17786634 | NonPC | Public | 2022 | USA | Env | 91.5 | 100 | 0.27 | 98.1612 | 5117301 | 45.29 | 71 | 444194 |
| SAMN25134885-SRR17786632 | PNUSAV002470 | SAMN25134885 | SRR17786632 | NonPC | Public | 2022 | USA | Env | 89.5 | 100 | 0 | 98.1336 | 4952671 | 45.46 | 55 | 350847 |
| SAMN25134886-SRR17786631 | PNUSAV002467 | SAMN25134886 | SRR17786631 | NonPC | Public | 2022 | USA | Env | 88.7 | 100 | 0.05 | 98.0552 | 4981996 | 45.34 | 56 | 443379 |
| SAMN25134887-SRR17677415 | PNUSAV002477 | SAMN25134887 | SRR17677415 | NonPC | Public | 2022 | USA | Env | 90 | 100 | 0.27 | 98.1671 | 5201203 | 45.27 | 57 | 482936 |
| SAMN25134888-SRR17677414 | PNUSAV002482 | SAMN25134888 | SRR17677414 | NonPC | Public | 2022 | USA | Env | 91.1 | 100 | 0.05 | 98.1885 | 5231043 | 45.23 | 51 | 550419 |
| SAMN25134889-SRR17677417 | PNUSAV002474 | SAMN25134889 | SRR17677417 | NonPC | Public | 2022 | USA | Env | 90.5 | 100 | 0 | 98.1749 | 5122021 | 45.19 | 55 | 401832 |
| SAMN25134890-SRR17786665 | PNUSAV002483 | SAMN25134890 | SRR17786665 | NonPC | Public | 2022 | USA | Env | 88.5 | 100 | 0.33 | 98.1602 | 4901707 | 45.42 | 38 | 403516 |
| SAMN25134891-SRR17677416 | PNUSAV002468 | SAMN25134891 | SRR17677416 | NonPC | Public | 2022 | USA | Env | 90 | 100 | 0.27 | 98.181 | 5201552 | 45.27 | 64 | 402874 |
| SAMN25174351-GCA_023338245.1 | BH0084 | SAMN25174351 | GCA_023338245.1 | PC | Public | 2020 | China | Clinical | 97.5 | 100 | 0 | 99.9148 | 5099944 | 45.26 | 54 | 425528 |
| SAMN25174513-GCA_023338275.1 | BH0085 | SAMN25174513 | GCA_023338275.1 | PC | Public | 2020 | China | Clinical | 97.5 | 100 | 0 | 99.9172 | 5101034 | 45.27 | 54 | 425401 |
| SAMN25209733-GCA_023338305.1 | BH0086 | SAMN25209733 | GCA_023338305.1 | PC | Public | 2020 | China | Clinical | 97.6 | 100 | 0 | 99.9122 | 5104612 | 45.26 | 63 | 373195 |
| SAMN25209740-GCA_023338315.1 | BH0087 | SAMN25209740 | GCA_023338315.1 | PC | Public | 2020 | China | Clinical | 97.6 | 100 | 0 | 99.9085 | 5102937 | 45.27 | 54 | 425515 |
| SAMN25209765-GCA_023338345.1 | BH0088 | SAMN25209765 | GCA_023338345.1 | PC | Public | 2020 | China | Clinical | 97.6 | 100 | 0 | 99.9162 | 5101321 | 45.27 | 53 | 425515 |
| SAMN25209771-GCA_023338355.1 | BH0089 | SAMN25209771 | GCA_023338355.1 | PC | Public | 2020 | China | Clinical | 97.5 | 100 | 0 | 99.9156 | 5100603 | 45.26 | 55 | 373358 |
| SAMN25209820-GCA_023338385.1 | BH0090 | SAMN25209820 | GCA_023338385.1 | PC | Public | 2020 | China | Env | 97.6 | 100 | 0 | 99.9113 | 5101553 | 45.27 | 56 | 425414 |
| SAMN25210965-GCA_023338395.1 | FCG0121 | SAMN25210965 | GCA_023338395.1 | PC | Public | 2020 | China | Clinical | 97.5 | 100 | 0 | 99.9172 | 5098838 | 45.26 | 55 | 425528 |
| SAMN25211063-GCA_023338425.1 | FCG0124 | SAMN25211063 | GCA_023338425.1 | PC | Public | 2020 | China | Clinical | 97.6 | 100 | 0 | 99.917 | 5101319 | 45.27 | 54 | 425515 |
| SAMN25211114-GCA_023338435.1 | FCG0125 | SAMN25211114 | GCA_023338435.1 | PC | Public | 2020 | China | Clinical | 97.6 | 100 | 0 | 99.9167 | 5101376 | 45.27 | 54 | 425515 |
| SAMN25212004-GCA_023338465.1 | FCG0126 | SAMN25212004 | GCA_023338465.1 | PC | Public | 2020 | China | Clinical | 97.6 | 100 | 0 | 99.9145 | 5100947 | 45.26 | 52 | 425373 |
| SAMN25214717-GCA_023338475.1 | FCG0127 | SAMN25214717 | GCA_023338475.1 | PC | Public | 2020 | China | Clinical | 97.6 | 100 | 0 | 99.9141 | 5102195 | 45.27 | 55 | 425487 |
| SAMN25220545-GCA_023338535.1 | FCG0128 | SAMN25220545 | GCA_023338535.1 | PC | Public | 2020 | China | Clinical | 97.6 | 100 | 0 | 99.9141 | 5102014 | 45.27 | 52 | 425515 |
| SAMN25220608-GCA_023338555.1 | FCG0130 | SAMN25220608 | GCA_023338555.1 | PC | Public | 2020 | China | Clinical | 97.6 | 100 | 0 | 99.9181 | 5100626 | 45.27 | 64 | 339053 |
| SAMN25221410-GCA_023338575.1 | FCG0131 | SAMN25221410 | GCA_023338575.1 | PC | Public | 2020 | China | Clinical | 97.6 | 100 | 0 | 99.9155 | 5102854 | 45.27 | 54 | 425373 |
| SAMN25232486-GCA_023338545.1 | FCG0132 | SAMN25232486 | GCA_023338545.1 | PC | Public | 2020 | China | Clinical | 97.5 | 100 | 0 | 99.9138 | 5099947 | 45.26 | 59 | 373188 |
| SAMN25232487-GCA_023338565.1 | NN0207 | SAMN25232487 | GCA_023338565.1 | PC | Public | 2020 | China | Clinical | 97.5 | 100 | 0 | 99.9178 | 5114629 | 45.25 | 55 | 425486 |
| SAMN25232498-GCA_023338635.1 | QZ0184 | SAMN25232498 | GCA_023338635.1 | PC | Public | 2020 | China | Clinical | 97.5 | 100 | 0 | 99.9143 | 5099863 | 45.26 | 58 | 425528 |
| SAMN25285885-SRR17772392 | V21010066 | SAMN25285885 | SRR17772392 | NonPC | Public | 2021 | USA | Env | 91.5 | 100 | 0.45 | 98.3566 | 5055425 | 45.31 | 45 | 413053 |
| SAMN25285886-SRR17772387 | V21010067 | SAMN25285886 | SRR17772387 | NonPC | Public | 2021 | USA | Env | 91.7 | 100 | 0.14 | 98.2698 | 5109146 | 45.23 | 38 | 846154 |
| SAMN25285887-SRR17772388 | V21010068 | SAMN25285887 | SRR17772388 | NonPC | Public | 2021 | USA | Env | 91.8 | 100 | 0.44 | 98.3341 | 5243822 | 45.21 | 55 | 664460 |
| SAMN25285888-SRR17772390 | V21010069 | SAMN25285888 | SRR17772390 | NonPC | Public | 2021 | USA | Env | 89.9 | 100 | 0 | 98.2815 | 5374820 | 45.15 | 49 | 538522 |
| SAMN25285889-SRR17772399 | V21010052 | SAMN25285889 | SRR17772399 | NonPC | Public | 2021 | USA | Env | 89.8 | 100 | 0.04 | 98.2818 | 5014160 | 45.33 | 56 | 825334 |

| | | | | | | | | | | | | | | | | |
|---|---|---|---|---|---|---|---|---|---|---|---|---|---|---|---|---|
| SAMN25285890-SRR17772393 | V21010054 | SAMN25285890 | SRR17772393 | NonPC | Public | 2021 | USA | Env | 88.8 | 100 | 0.09 | 98.0015 | 4945311 | 45.41 | 44 | 544326 |
| SAMN25285891-SRR17772396 | V21010057 | SAMN25285891 | SRR17772396 | NonPC | Public | 2021 | USA | Env | 89.4 | 100 | 0.05 | 98.1727 | 5024519 | 45.32 | 63 | 218391 |
| SAMN25285892-SRR17772394 | V21010056 | SAMN25285892 | SRR17772394 | NonPC | Public | 2021 | USA | Env | 89.8 | 100 | 0.04 | 98.2689 | 5014354 | 45.33 | 58 | 714513 |
| SAMN25285893-SRR17772398 | V21010059 | SAMN25285893 | SRR17772398 | NonPC | Public | 2021 | USA | Env | 91.5 | 100 | 0.08 | 98.2388 | 5162849 | 45.19 | 41 | 625533 |
| SAMN25285894-SRR17772397 | V21010060 | SAMN25285894 | SRR17772397 | NonPC | Public | 2021 | USA | Env | 91.1 | 100 | 0.08 | 98.186 | 5168417 | 45.2 | 50 | 579193 |
| SAMN25285895-SRR17772389 | V21010058 | SAMN25285895 | SRR17772389 | NonPC | Public | 2021 | USA | Env | 91.3 | 100 | 0 | 98.3272 | 5146405 | 45.31 | 34 | 458266 |
| SAMN25285896-SRR17772395 | V21010061 | SAMN25285896 | SRR17772395 | NonPC | Public | 2021 | USA | Env | 90 | 100 | 0.35 | 98.1866 | 5041603 | 45.18 | 31 | 511437 |
| SAMN25285897-SRR17772391 | V21010062 | SAMN25285897 | SRR17772391 | NonPC | Public | 2021 | USA | Env | 91.2 | 100 | 0.27 | 98.3127 | 4966835 | 45.4 | 51 | 281338 |
| SAMN25285898-SRR17772400 | V21010063 | SAMN25285898 | SRR17772400 | NonPC | Public | 2021 | USA | Env | 91.4 | 100 | 0.3 | 98.2519 | 5296227 | 45.14 | 63 | 543368 |
| SAMN25285899-SRR17772402 | V21010073 | SAMN25285899 | SRR17772402 | NonPC | Public | 2021 | USA | Env | 89.4 | 100 | 0.33 | 98.3019 | 5174916 | 45.33 | 58 | 589887 |
| SAMN25285900-SRR17772403 | V21010078 | SAMN25285900 | SRR17772403 | NonPC | Public | 2021 | USA | Env | 91 | 100 | 0.08 | 98.2457 | 5112440 | 45.24 | 48 | 545898 |
| SAMN25285901-SRR17772401 | V21010077 | SAMN25285901 | SRR17772401 | NonPC | Public | 2021 | USA | Env | 91.7 | 100 | 0.13 | 98.2744 | 5150199 | 45.28 | 44 | 420525 |
| SAMN25285902-SRR17772408 | V21010074 | SAMN25285902 | SRR17772408 | NonPC | Public | 2021 | USA | Env | 89.6 | 100 | 0.19 | 98.2304 | 4990340 | 45.38 | 59 | 568215 |
| SAMN25336236-SRR17786319 | PNUSAV002487 | SAMN25336236 | SRR17786319 | NonPC | Public | 2022 | USA | NA | 91.6 | 100 | 0 | 98.2846 | 5103073 | 45.27 | 49 | 390220 |
| SAMN25338425-SRR17787797 | PNUSAV002500 | SAMN25338425 | SRR17787797 | NonPC | Public | 2021 | USA | Env | 91.5 | 100 | 0.45 | 98.3566 | 5055425 | 45.31 | 45 | 413053 |
| SAMN25338427-SRR17787798 | PNUSAV002499 | SAMN25338427 | SRR17787798 | NonPC | Public | 2021 | USA | Env | 91.7 | 100 | 0.14 | 98.2592 | 5109146 | 45.23 | 38 | 846154 |
| SAMN25338429-SRR17787800 | PNUSAV002498 | SAMN25338429 | SRR17787800 | NonPC | Public | 2021 | USA | Env | 91.8 | 100 | 0.44 | 98.3356 | 5243822 | 45.21 | 55 | 664460 |
| SAMN25338432-SRR17787885 | PNUSAV002497 | SAMN25338432 | SRR17787885 | NonPC | Public | 2021 | USA | Env | 89.9 | 100 | 0 | 98.275 | 5374820 | 45.15 | 49 | 538522 |
| SAMN25524171-SRR17838914 | V21010086 | SAMN25524171 | SRR17838914 | NonPC | Public | 2021 | USA | Env | 90.5 | 100 | 0.03 | 98.3399 | 5104004 | 45.17 | 57 | 541599 |
| SAMN25524172-SRR17838938 | V21010085 | SAMN25524172 | SRR17838938 | NonPC | Public | 2021 | USA | Env | 89.8 | 100 | 0.17 | 98.2446 | 5456571 | 44.85 | 34 | 570274 |
| SAMN25524173-SRR17838827 | V21010079 | SAMN25524173 | SRR17838827 | NonPC | Public | 2021 | USA | Env | 91.3 | 100 | 0.22 | 98.2252 | 5228080 | 45.17 | 53 | 348514 |
| SAMN25524174-SRR17838887 | V21010083 | SAMN25524174 | SRR17838887 | NonPC | Public | 2021 | USA | Env | 91.5 | 100 | 0 | 98.4065 | 5133255 | 45.22 | 39 | 520021 |
| SAMN25524175-SRR17838891 | V21010080 | SAMN25524175 | SRR17838891 | NonPC | Public | 2021 | USA | Env | 92 | 100 | 0 | 98.432 | 4955954 | 45.4 | 33 | 459036 |
| SAMN25656892-SRR17888655 | V21010128 | SAMN25656892 | SRR17888655 | NonPC | Public | 2021 | USA | Env | 88.2 | 100 | 0.58 | 98.1929 | 5182613 | 45.21 | 67 | 387274 |
| SAMN25656893-SRR17888656 | V21010114 | SAMN25656893 | SRR17888656 | NonPC | Public | 2021 | USA | Env | 91.4 | 100 | 0.08 | 98.3341 | 5067672 | 45.3 | 36 | 560643 |
| SAMN25656894-SRR17888657 | V21010126 | SAMN25656894 | SRR17888657 | NonPC | Public | 2021 | USA | Env | 91.6 | 100 | 0.15 | 98.3514 | 5041873 | 45.27 | 51 | 535457 |
| SAMN25656895-SRR17888659 | V21010127 | SAMN25656895 | SRR17888659 | NonPC | Public | 2021 | USA | Env | 91.2 | 100 | 0.14 | 98.3261 | 5115471 | 45.26 | 57 | 437600 |
| SAMN25656896-SRR17888658 | V21010124 | SAMN25656896 | SRR17888658 | NonPC | Public | 2021 | USA | Env | 88.7 | 100 | 0.36 | 98.2656 | 5055238 | 45.3 | 63 | 544484 |
| SAMN25656897-SRR17888661 | V21010108 | SAMN25656897 | SRR17888661 | NonPC | Public | 2021 | USA | Env | 87.5 | 100 | 0.3 | 98.2032 | 5119692 | 45.19 | 63 | 363309 |
| SAMN25656898-SRR17888664 | V21010113 | SAMN25656898 | SRR17888664 | NonPC | Public | 2021 | USA | Env | 89.7 | 100 | 0.23 | 98.2234 | 5258948 | 45.01 | 58 | 346199 |
| SAMN25656899-SRR17888667 | V21010111 | SAMN25656899 | SRR17888667 | NonPC | Public | 2021 | USA | Env | 90.5 | 100 | 0 | 98.2306 | 5137718 | 45.19 | 56 | 381706 |
| SAMN25656903-SRR17888668 | V21010109 | SAMN25656903 | SRR17888668 | NonPC | Public | 2021 | USA | Env | 90.3 | 100 | 0.03 | 98.2532 | 5112025 | 45.26 | 74 | 815652 |
| SAMN25656904-SRR17888665 | V21010089 | SAMN25656904 | SRR17888665 | NonPC | Public | 2021 | USA | Env | 91.3 | 100 | 0.27 | 98.261 | 5180763 | 45.21 | 35 | 558213 |
| SAMN25656905-SRR17888663 | V21010088 | SAMN25656905 | SRR17888663 | NonPC | Public | 2021 | USA | Env | 89.4 | 100 | 0.01 | 98.2409 | 5275128 | 45.26 | 66 | 273603 |
| SAMN25656906-SRR17888662 | V21010094 | SAMN25656906 | SRR17888662 | NonPC | Public | 2021 | USA | Env | 91.1 | 100 | 0.08 | 98.3179 | 4984040 | 45.33 | 34 | 444196 |
| SAMN25656907-SRR17888660 | V21010093 | SAMN25656907 | SRR17888660 | NonPC | Public | 2021 | USA | Env | 91.2 | 100 | 0 | 98.3495 | 5029771 | 45.35 | 37 | 470306 |
| SAMN25656908-SRR17888666 | V21010105 | SAMN25656908 | SRR17888666 | NonPC | Public | 2021 | USA | Env | 89.7 | 100 | 0 | 98.2689 | 5271590 | 45.05 | 42 | 456792 |
| SAMN25656909-SRR17888671 | V21010087 | SAMN25656909 | SRR17888671 | NonPC | Public | 2021 | USA | Env | 91.1 | 100 | 0.35 | 98.2728 | 5043553 | 45.29 | 73 | 406123 |
| SAMN25815102-SRR17960991 | PNUSAV002566 | SAMN25815102 | SRR17960991 | NonPC | Public | 2022 | USA | NA | 89 | 100 | 0 | 98.0592 | 5192595 | 45.05 | 68 | 318929 |
| SAMN25936682-SRR18015351 | PNUSAV002541 | SAMN25936682 | SRR18015351 | NonPC | Public | 2022 | USA | NA | 88.6 | 100 | 0 | 98.2881 | 4878488 | 45.42 | 59 | 255141 |
| SAMN25962175-GCA_022352195.1 | 460 | SAMN25962175 | GCA_022352195.1 | PC | Public | 2017 | China | Clinical | 97.4 | 100 | 0 | 99.934 | 5149679 | 45.12 | 71 | 294833 |
| SAMN25962176-GCA_022352305.1 | 469 | SAMN25962176 | GCA_022352305.1 | NonPC | Public | 2017 | China | Clinical | 91 | 100 | 0 | 98.3622 | 5058349 | 45.24 | 84 | 285699 |
| SAMN25962177-GCA_022352495.1 | vp41 | SAMN25962177 | GCA_022352495.1 | PC | Public | 2015 | China | NA | 97.8 | 100 | 0 | 99.9852 | 5095112 | 45.26 | 68 | 288258 |
| SAMN25962178-GCA_022352165.1 | vp44 | SAMN25962178 | GCA_022352165.1 | NonPC | Public | 2015 | China | NA | 87.8 | 100 | 0.39 | 98.1966 | 5021015 | 45.23 | 117 | 371103 |
| SAMN25965007-SRR18034811 | PNUSAV002455 | SAMN25965007 | SRR18034811 | NonPC | Public | 2021 | USA | NA | 90.6 | 100 | 0.16 | 98.225 | 5331391 | 44.98 | 29 | 582315 |
| SAMN26198547-SRR18508061 | 19MO03SH02 | SAMN26198547 | SRR18508061 | NonPC | Public | 2019 | USA | Env | 90.9 | 100 | 0.14 | 98.3465 | 5480245 | 45.25 | 63 | 406191 |
| SAMN26198549-SRR18508059 | 19MO03SH03-2 | SAMN26198549 | SRR18508059 | NonPC | Public | 2019 | USA | Env | 89.8 | 100 | 0 | 98.2193 | 5415338 | 45.2 | 101 | 468183 |
| SAMN26198550-SRR18508058 | 19MO03SH05 | SAMN26198550 | SRR18508058 | NonPC | Public | 2019 | USA | Env | 91.9 | 100 | 0.27 | 98.2983 | 5376483 | 45.27 | 50 | 545261 |

| | | | | | | | | | | | | | | | | |
|---|---|---|---|---|---|---|---|---|---|---|---|---|---|---|---|---|
| SAMN26198741-SRR18508005 | 19MO01SH02 | SAMN26198741 | SRR18508005 | NonPC | Public | 2019 | USA | Env | 89.6 | 100 | 0 | 98.2222 | 5279432 | 45.26 | 51 | 460255 |
| SAMN26198742-SRR18508004 | 19MO01SH03 | SAMN26198742 | SRR18508004 | NonPC | Public | 2019 | USA | Env | 88.8 | 100 | 0 | 98.0214 | 5478049 | 45.06 | 124 | 123156 |
| SAMN26198744-SRR18508002 | 19MO01SH06 | SAMN26198744 | SRR18508002 | NonPC | Public | 2019 | USA | Env | 91.4 | 100 | 0 | 98.3028 | 5391545 | 45.23 | 61 | 296498 |
| SAMN26198754-SRR18507991 | 19CA03SH03 | SAMN26198754 | SRR18507991 | NonPC | Public | 2019 | USA | Env | 92 | 100 | 0.27 | 98.3164 | 5283560 | 45.2 | 64 | 269869 |
| SAMN26198758-SRR18507987 | 19SD02SH07 | SAMN26198758 | SRR18507987 | NonPC | Public | 2019 | USA | Env | 88.8 | 100 | 0.27 | 98.0305 | 5462769 | 45.01 | 124 | 145245 |
| SAMN26198765-SRR18507851 | 19NY01SH07 | SAMN26198765 | SRR18507851 | NonPC | Public | 2019 | USA | Env | 91.8 | 100 | 0.95 | 98.3227 | 5302555 | 45.16 | 81 | 211308 |
| SAMN26198766-SRR18507850 | 19NY03SH02 | SAMN26198766 | SRR18507850 | NonPC | Public | 2019 | USA | Env | 91.8 | 100 | 0 | 98.3543 | 5247260 | 45.18 | 35 | 513470 |
| SAMN26198773-SRR18507842 | 19GA03SH04 | SAMN26198773 | SRR18507842 | NonPC | Public | 2019 | USA | Env | 91.4 | 100 | 0 | 98.3509 | 5164929 | 45.27 | 66 | 187900 |
| SAMN26198777-SRR18507838 | 19SD04SH02 | SAMN26198777 | SRR18507838 | NonPC | Public | 2019 | USA | Env | 89.6 | 100 | 0.54 | 98.2474 | 5218016 | 45.16 | 40 | 442140 |
| SAMN26504778-SRR18254625 | PNUSAV002567 | SAMN26504778 | SRR18254625 | NonPC | Public | 2022 | USA | NA | 91.5 | 100 | 0 | 98.3052 | 5133087 | 45.24 | 44 | 561739 |
| SAMN26523570-SRR18270328 | V21010140 | SAMN26523570 | SRR18270328 | NonPC | Public | 2021 | USA | Env | 91.2 | 100 | 0 | 98.2444 | 5104217 | 45.25 | 66 | 538050 |
| SAMN26523571-SRR18270575 | V21010129 | SAMN26523571 | SRR18270575 | NonPC | Public | 2021 | USA | Env | 88.8 | 100 | 0.3 | 98.1552 | 4921590 | 45.43 | 41 | 432082 |
| SAMN26523572-SRR18270724 | V21010138 | SAMN26523572 | SRR18270724 | NonPC | Public | 2021 | USA | Env | 91.9 | 100 | 0 | 98.3979 | 5060422 | 45.34 | 31 | 832778 |
| SAMN26523573-SRR18270787 | V21010139 | SAMN26523573 | SRR18270787 | NonPC | Public | 2021 | USA | Env | 90.2 | 100 | 0 | 98.2465 | 5143756 | 45.25 | 44 | 494257 |
| SAMN26523574-SRR18270663 | V21010137 | SAMN26523574 | SRR18270663 | NonPC | Public | 2021 | USA | Env | 91.9 | 100 | 0 | 98.3939 | 5059605 | 45.34 | 32 | 444613 |
| SAMN26523575-SRR18270662 | V21010132 | SAMN26523575 | SRR18270662 | NonPC | Public | 2021 | USA | Env | 91.3 | 100 | 0.14 | 98.2163 | 5152901 | 45.2 | 56 | 275167 |
| SAMN26523576-SRR18270786 | V21010130 | SAMN26523576 | SRR18270786 | NonPC | Public | 2021 | USA | Env | 91.4 | 100 | 0 | 98.3587 | 5094616 | 45.26 | 58 | 828723 |
| SAMN26523585-SRR18270796 | V21010141 | SAMN26523585 | SRR18270796 | NonPC | Public | 2021 | USA | Env | 90.7 | 100 | 0.05 | 98.2758 | 5042515 | 45.28 | 46 | 734430 |
| SAMN26550102-GCA_022603355.1 | MUVP9 | SAMN26550102 | GCA_022603355.1 | NonPC | Public | 2016 | Thailand | Env | 91.2 | 100 | 0 | 98.3475 | 5109371 | 45.2 | 53 | 382698 |
| SAMN26550103-GCA_022603315.1 | MUVP10 | SAMN26550103 | GCA_022603315.1 | NonPC | Public | 2016 | Thailand | Env | 90.1 | 100 | 0 | 98.2591 | 5071201 | 45.3 | 31 | 542922 |
| SAMN26550104-GCA_022603295.1 | MUVP20 | SAMN26550104 | GCA_022603295.1 | NonPC | Public | 2017 | Thailand | Env | 91.6 | 100 | 0 | 98.3941 | 5090795 | 45.27 | 52 | 402508 |
| SAMN26550105-GCA_022603325.1 | MUVP22 | SAMN26550105 | GCA_022603325.1 | NonPC | Public | 2017 | Thailand | Env | 92.1 | 100 | 0.19 | 98.3576 | 5023226 | 45.23 | 55 | 406579 |
| SAMN26655273-SRR18395847 | NA | SAMN26655273 | SRR18395847 | NonPC | Public | 2020 | Malaysia | Env | 90.9 | 100 | 0.17 | 98.3454 | 5006029 | 45.31 | 50 | 1260342 |
| SAMN26659663-SRR18326279 | V21010161 | SAMN26659663 | SRR18326279 | NonPC | Public | 2017 | USA | Env | 89 | 100 | 0.03 | 98.1011 | 5029083 | 45.32 | 58 | 530045 |
| SAMN26659665-SRR18326289 | V21010170 | SAMN26659665 | SRR18326289 | NonPC | Public | 2017 | USA | Env | 88.9 | 100 | 0 | 98.1042 | 5018432 | 45.27 | 36 | 505348 |
| SAMN26659668-SRR18326277 | V21010143 | SAMN26659668 | SRR18326277 | NonPC | Public | 2021 | USA | Env | 89.4 | 100 | 0.01 | 98.2254 | 5256044 | 45.18 | 71 | 268722 |
| SAMN26659669-SRR18326278 | V21010157 | SAMN26659669 | SRR18326278 | NonPC | Public | 2017 | USA | Env | 91.3 | 100 | 0.17 | 98.3181 | 5222077 | 45.23 | 70 | 800919 |
| SAMN26659670-SRR18326291 | V21010145 | SAMN26659670 | SRR18326291 | NonPC | Public | 2021 | USA | Env | 91.9 | 100 | 0.15 | 98.2913 | 5038460 | 45.36 | 42 | 850497 |
| SAMN26659671-SRR18326288 | V21010149 | SAMN26659671 | SRR18326288 | NonPC | Public | 2021 | USA | Env | 91.5 | 100 | 0.08 | 98.2398 | 5296035 | 45.13 | 54 | 257074 |
| SAMN26659672-SRR18326287 | V21010179 | SAMN26659672 | SRR18326287 | NonPC | Public | 2018 | USA | Env | 91.1 | 100 | 0.08 | 98.1979 | 5206723 | 45.2 | 45 | 545820 |
| SAMN26659673-SRR18326280 | V21010182 | SAMN26659673 | SRR18326280 | NonPC | Public | 2018 | USA | Env | 90.6 | 100 | 0.44 | 98.2332 | 5293593 | 45.22 | 51 | 612858 |
| SAMN26659674-SRR18326293 | V21010181 | SAMN26659674 | SRR18326293 | NonPC | Public | 2018 | USA | Env | 89.5 | 100 | 0.22 | 98.3111 | 4974416 | 45.38 | 39 | 386969 |
| SAMN26659675-SRR18326276 | V21010175 | SAMN26659675 | SRR18326276 | NonPC | Public | 2017 | USA | Env | 89.9 | 100 | 0.22 | 98.2236 | 5395406 | 45.2 | 62 | 408486 |
| SAMN26876016-SRR18435715 | PNUSAV002692 | SAMN26876016 | SRR18435715 | NonPC | Public | 2022 | USA | NA | 88.1 | 100 | 0.66 | 98.2003 | 5085273 | 45.2 | 67 | 558742 |
| SAMN26980445-SRR18490902 | V21010174 | SAMN26980445 | SRR18490902 | NonPC | Public | 2017 | USA | Env | 88.9 | 100 | 0 | 98.1219 | 5020605 | 45.27 | 34 | 522503 |
| SAMN27023183-SRR18326294 | V21010156 | SAMN27023183 | SRR18326294 | NonPC | Public | 2017 | USA | Env | 89.4 | 100 | 0.01 | 98.2102 | 5280713 | 45.26 | 71 | 268721 |
| SAMN27114194-SRR18548563 | PNUSAV002696 | SAMN27114194 | SRR18548563 | NonPC | Public | 2022 | USA | Env | 89.1 | 100 | 0.06 | 98.0534 | 4984032 | 45.39 | 52 | 376612 |
| SAMN27114195-SRR18548566 | PNUSAV002699 | SAMN27114195 | SRR18548566 | NonPC | Public | 2022 | USA | Env | 91.2 | 100 | 0 | 98.2701 | 5074115 | 45.29 | 36 | 761771 |
| SAMN27114196-SRR18548561 | PNUSAV002698 | SAMN27114196 | SRR18548561 | NonPC | Public | 2022 | USA | Env | 89.9 | 100 | 0.54 | 98.3145 | 5012427 | 45.35 | 40 | 773690 |
| SAMN27114197-SRR18548564 | PNUSAV002697 | SAMN27114197 | SRR18548564 | NonPC | Public | 2022 | USA | Env | 91.7 | 100 | 0.08 | 98.2167 | 5194336 | 45.21 | 54 | 381996 |
| SAMN27114198-SRR18548565 | PNUSAV002700 | SAMN27114198 | SRR18548565 | NonPC | Public | 2022 | USA | Env | 91.3 | 100 | 0 | 98.2265 | 5156058 | 45.16 | 68 | 487703 |
| SAMN27150375-SRR18560351 | PNUSAV002428 | SAMN27150375 | SRR18560351 | NonPC | Public | 2020 | USA | NA | 91.2 | 100 | 0.05 | 98.3766 | 5086348 | 45.2 | 77 | 195633 |
| SAMN27150376-SRR18560352 | PNUSAV002427 | SAMN27150376 | SRR18560352 | NonPC | Public | 2020 | USA | NA | 90.7 | 100 | 0.26 | 98.2104 | 5196089 | 45.14 | 92 | 185291 |
| SAMN27150377-SRR18560355 | PNUSAV002691 | SAMN27150377 | SRR18560355 | NonPC | Public | 2021 | USA | NA | 90.7 | 100 | 0.41 | 98.1323 | 5101793 | 45.16 | 58 | 236685 |
| SAMN27150378-SRR18560350 | PNUSAV002690 | SAMN27150378 | SRR18560350 | NonPC | Public | 2021 | USA | NA | 89.2 | 100 | 0 | 98.0624 | 5116545 | 45.24 | 49 | 274870 |
| SAMN27150379-SRR18560353 | PNUSAV002689 | SAMN27150379 | SRR18560353 | NonPC | Public | 2021 | USA | NA | 89.5 | 100 | 0.36 | 98.3421 | 4994485 | 45.38 | 31 | 526608 |
| SAMN27150385-SRR18560364 | PNUSAV002426 | SAMN27150385 | SRR18560364 | NonPC | Public | 2020 | USA | NA | 89.8 | 100 | 0.04 | 98.2275 | 5169324 | 45.21 | 51 | 256616 |
| SAMN27155686-SRR18573346 | PNUSAV002694 | SAMN27155686 | SRR18573346 | NonPC | Public | 2022 | USA | NA | 89 | 100 | 0 | 98.1247 | 5005985 | 45.31 | 116 | 101700 |

| | | | | | | | | | | | | | | | | |
|---|---|---|---|---|---|---|---|---|---|---|---|---|---|---|---|---|
| SAMN27297343-SRR18609780 | 22MO01SH01-V1 | SAMN27297343 | SRR18609780 | NonPC | Public | 2022 | USA | Env | 90.9 | 100 | 0.4 | 98.2537 | 5237316 | 45.22 | 95 | 397225 |
| SAMN27297347-SRR18609788 | 21MO11SH01-V1 | SAMN27297347 | SRR18609788 | NonPC | Public | 2021 | USA | Env | 90 | 100 | 0.07 | 98.2858 | 5323238 | 45.28 | 99 | 299901 |
| SAMN27514928-SRR18709557 | V21010190 | SAMN27514928 | SRR18709557 | NonPC | Public | 2018 | USA | Env | 88.3 | 100 | 0.36 | 98.2891 | 5064313 | 45.24 | 40 | 490060 |
| SAMN27514929-SRR18709555 | V21010193 | SAMN27514929 | SRR18709555 | NonPC | Public | 2018 | USA | Env | 91.2 | 100 | 0.09 | 98.3001 | 5287122 | 45.19 | 47 | 723580 |
| SAMN27514930-SRR18709560 | V21010194 | SAMN27514930 | SRR18709560 | NonPC | Public | 2018 | USA | Env | 91.1 | 100 | 0.13 | 98.2704 | 5137362 | 45.22 | 60 | 589522 |
| SAMN27514931-SRR18709553 | V21010187 | SAMN27514931 | SRR18709553 | NonPC | Public | 2018 | USA | Env | 91.8 | 100 | 0.49 | 98.1913 | 5210121 | 45.17 | 34 | 850317 |
| SAMN27543701-SRR18730489 | PNUSAV002727 | SAMN27543701 | SRR18730489 | NonPC | Public | 2022 | USA | NA | 89 | 100 | 0 | 98.0794 | 5106413 | 45.24 | 58 | 375455 |
| SAMN27583593-SRR18752183 | PNUSAV002724 | SAMN27583593 | SRR18752183 | NonPC | Public | 2022 | USA | NA | 89.6 | 100 | 0 | 98.2274 | 5083107 | 45.32 | 46 | 548802 |
| SAMN27583595-SRR18751195 | PNUSAV002723 | SAMN27583595 | SRR18751195 | NonPC | Public | 2022 | USA | NA | 91.5 | 100 | 0 | 98.356 | 5057793 | 45.31 | 40 | 413005 |
| SAMN27583596-SRR18752186 | PNUSAV002730 | SAMN27583596 | SRR18752186 | NonPC | Public | 2022 | USA | NA | 92.2 | 100 | 0.22 | 98.2694 | 5103322 | 45.35 | 111 | 95629 |
| SAMN27590317-SRR18762478 | PHLUSAVBR00316 | SAMN27590317 | SRR18762478 | NonPC | Public | 2021 | USA | Env | 91.1 | 100 | 0.41 | 98.2661 | 5282948 | 45.21 | 39 | 549101 |
| SAMN27590318-SRR18762467 | PHLUSAVBR00317 | SAMN27590318 | SRR18762467 | NonPC | Public | 2021 | USA | Env | 90.5 | 100 | 0.11 | 98.2068 | 5289302 | 45.13 | 49 | 478108 |
| SAMN27590319-SRR18762461 | PHLUSAVBR00318 | SAMN27590319 | SRR18762461 | NonPC | Public | 2021 | USA | Env | 90.6 | 100 | 0.14 | 98.2162 | 5147013 | 45.1 | 48 | 471458 |
| SAMN27590320-SRR18762460 | PHLUSAVBR00319 | SAMN27590320 | SRR18762460 | NonPC | Public | 2021 | USA | Env | 91.2 | 100 | 0 | 98.2477 | 5072920 | 45.33 | 43 | 420954 |
| SAMN27590321-SRR18762459 | PHLUSAVBR00320 | SAMN27590321 | SRR18762459 | NonPC | Public | 2021 | USA | Env | 91.8 | 100 | 0 | 98.3659 | 5233953 | 45.14 | 53 | 519426 |
| SAMN27590322-SRR18762458 | PHLUSAVBR00321 | SAMN27590322 | SRR18762458 | NonPC | Public | 2021 | USA | Env | 88.4 | 100 | 0.03 | 98.0623 | 5047647 | 45.25 | 37 | 547451 |
| SAMN27590323-SRR18762457 | PHLUSAVBR00322 | SAMN27590323 | SRR18762457 | NonPC | Public | 2021 | USA | Env | 90.6 | 100 | 0 | 98.2153 | 5264553 | 45.18 | 51 | 738253 |
| SAMN27590324-SRR18762456 | PHLUSAVBR00323 | SAMN27590324 | SRR18762456 | NonPC | Public | 2021 | USA | Env | 91.6 | 100 | 0 | 98.2974 | 5055438 | 45.33 | 51 | 946048 |
| SAMN27590325-SRR18762455 | PHLUSAVBR00324 | SAMN27590325 | SRR18762455 | NonPC | Public | 2021 | USA | Env | 91.9 | 100 | 0.27 | 98.3324 | 5423923 | 44.9 | 36 | 800279 |
| SAMN27590326-SRR18762477 | PHLUSAVBR00326 | SAMN27590326 | SRR18762477 | NonPC | Public | 2021 | USA | Env | 90.5 | 100 | 0 | 98.2789 | 5255312 | 45.17 | 58 | 498308 |
| SAMN27590327-SRR18762476 | PHLUSAVBR00327 | SAMN27590327 | SRR18762476 | NonPC | Public | 2021 | USA | Env | 88.4 | 100 | 0.03 | 98.1593 | 5032876 | 45.26 | 37 | 547656 |
| SAMN27590328-SRR18762475 | PHLUSAVBR00328 | SAMN27590328 | SRR18762475 | NonPC | Public | 2021 | USA | Env | 91.2 | 100 | 0.41 | 98.2867 | 5269972 | 45.23 | 45 | 290454 |
| SAMN27590329-SRR18762474 | PHLUSAVBR00329 | SAMN27590329 | SRR18762474 | NonPC | Public | 2021 | USA | Env | 88.6 | 100 | 0 | 98.1452 | 5063956 | 45.24 | 73 | 272253 |
| SAMN27590330-SRR18762473 | PHLUSAVBR00330 | SAMN27590330 | SRR18762473 | NonPC | Public | 2021 | USA | Env | 91.2 | 100 | 0.41 | 98.3856 | 5098728 | 45.21 | 33 | 468688 |
| SAMN27590331-SRR18762472 | PHLUSAVBR00331 | SAMN27590331 | SRR18762472 | NonPC | Public | 2021 | USA | Env | 90.6 | 100 | 0.14 | 98.211 | 5048574 | 45.19 | 39 | 542833 |
| SAMN27590332-SRR18762471 | PHLUSAVBR00332 | SAMN27590332 | SRR18762471 | NonPC | Public | 2021 | USA | Env | 88.7 | 100 | 0.35 | 98.1516 | 5042578 | 45.19 | 84 | 110893 |
| SAMN27590333-SRR18762470 | PHLUSAVBR00333 | SAMN27590333 | SRR18762470 | NonPC | Public | 2021 | USA | Env | 91.1 | 100 | 0 | 98.3079 | 5084925 | 45.23 | 53 | 454905 |
| SAMN27590334-SRR18762469 | PHLUSAVBR00335 | SAMN27590334 | SRR18762469 | NonPC | Public | 2021 | USA | Env | 92.6 | 100 | 0 | 98.2707 | 5332595 | 45.04 | 36 | 866413 |
| SAMN27590335-SRR18762468 | PHLUSAVBR00336 | SAMN27590335 | SRR18762468 | NonPC | Public | 2021 | USA | Env | 91.1 | 100 | 0.16 | 98.2734 | 5382403 | 45.13 | 56 | 412369 |
| SAMN27590336-SRR18762466 | PHLUSAVBR00337 | SAMN27590336 | SRR18762466 | NonPC | Public | 2021 | USA | Env | 89.6 | 100 | 0 | 98.2503 | 5082236 | 45.19 | 52 | 737145 |
| SAMN27590337-SRR18762465 | PHLUSAVBR00338 | SAMN27590337 | SRR18762465 | NonPC | Public | 2021 | USA | Env | 88.7 | 100 | 0 | 98.1355 | 5064689 | 45.25 | 64 | 407220 |
| SAMN27590338-SRR18762464 | PHLUSAVBR00339 | SAMN27590338 | SRR18762464 | NonPC | Public | 2021 | USA | Env | 91.5 | 100 | 0.37 | 98.3568 | 5211226 | 45.14 | 51 | 453020 |
| SAMN27590339-SRR18762463 | PHLUSAVBR00340 | SAMN27590339 | SRR18762463 | NonPC | Public | 2021 | USA | Env | 91.1 | 100 | 0.41 | 98.3163 | 5272127 | 45.23 | 48 | 401550 |
| SAMN27590639-SRR18763201 | PHLUSAVBR00286 | SAMN27590639 | SRR18763201 | NonPC | Public | 2021 | USA | Env | 91.7 | 100 | 0.35 | 98.2753 | 5143570 | 45.27 | 33 | 638565 |
| SAMN27590640-SRR18763200 | PHLUSAVBR00287 | SAMN27590640 | SRR18763200 | NonPC | Public | 2021 | USA | Env | 90.5 | 100 | 0.14 | 98.2323 | 5129364 | 45.11 | 37 | 525465 |
| SAMN27590641-SRR18763189 | PHLUSAVBR00288 | SAMN27590641 | SRR18763189 | NonPC | Public | 2021 | USA | Env | 91.8 | 100 | 0.08 | 98.3155 | 5245313 | 45.18 | 42 | 472126 |
| SAMN27590642-SRR18763183 | PHLUSAVBR00289 | SAMN27590642 | SRR18763183 | NonPC | Public | 2021 | USA | Env | 91.3 | 100 | 0 | 98.2367 | 5102354 | 45.29 | 41 | 458288 |
| SAMN27590643-SRR18763182 | PHLUSAVBR00290 | SAMN27590643 | SRR18763182 | NonPC | Public | 2021 | USA | Env | 90.5 | 100 | 0.11 | 98.2327 | 5286652 | 45.13 | 46 | 377118 |
| SAMN27590644-SRR18763181 | PHLUSAVBR00292 | SAMN27590644 | SRR18763181 | NonPC | Public | 2021 | USA | Env | 91.3 | 100 | 0.08 | 98.2154 | 5271353 | 45.21 | 43 | 401590 |
| SAMN27590646-SRR18763179 | PHLUSAVBR00294 | SAMN27590646 | SRR18763179 | NonPC | Public | 2021 | USA | Env | 89.7 | 100 | 0.27 | 98.2005 | 5184569 | 45.22 | 36 | 562022 |
| SAMN27590648-SRR18763177 | PHLUSAVBR00296 | SAMN27590648 | SRR18763177 | NonPC | Public | 2021 | USA | Env | 92.5 | 100 | 0 | 98.3424 | 5315142 | 45.05 | 35 | 545935 |
| SAMN27590649-SRR18763199 | PHLUSAVBR00297 | SAMN27590649 | SRR18763199 | NonPC | Public | 2021 | USA | Env | 91.2 | 100 | 0.41 | 98.2646 | 5325651 | 45.18 | 28 | 549077 |
| SAMN27590650-SRR18763198 | PHLUSAVBR00298 | SAMN27590650 | SRR18763198 | NonPC | Public | 2021 | USA | Env | 91.2 | 100 | 0.41 | 98.2799 | 5263500 | 45.23 | 28 | 556423 |
| SAMN27590651-SRR18763197 | PHLUSAVBR00300 | SAMN27590651 | SRR18763197 | NonPC | Public | 2021 | USA | Env | 92.5 | 100 | 0 | 98.294 | 5200204 | 45.23 | 30 | 526351 |
| SAMN27590652-SRR18763196 | PHLUSAVBR00301 | SAMN27590652 | SRR18763196 | NonPC | Public | 2021 | USA | Env | 92.6 | 100 | 0 | 98.3169 | 5328619 | 45.04 | 31 | 733078 |
| SAMN27590653-SRR18763195 | PHLUSAVBR00302 | SAMN27590653 | SRR18763195 | NonPC | Public | 2021 | USA | Env | 89.8 | 100 | 0.12 | 98.3253 | 5153396 | 45.21 | 46 | 702675 |
| SAMN27590654-SRR18763194 | PHLUSAVBR00303 | SAMN27590654 | SRR18763194 | NonPC | Public | 2021 | USA | Env | 91.3 | 100 | 0.08 | 98.2747 | 5298101 | 45.2 | 48 | 561923 |
| SAMN27590655-SRR18763193 | PHLUSAVBR00304 | SAMN27590655 | SRR18763193 | NonPC | Public | 2021 | USA | Env | 88.7 | 100 | 0.35 | 98.1741 | 4952325 | 45.35 | 37 | 521341 |

| | | | | | | | | | | | | | | | | |
|---|---|---|---|---|---|---|---|---|---|---|---|---|---|---|---|---|
| SAMN27590656-SRR18763192 | PHLUSAVBR00305 | SAMN27590656 | SRR18763192 | NonPC | Public | 2021 | USA | Env | 91.2 | 100 | 0.41 | 98.2892 | 5268996 | 45.23 | 30 | 549101 |
| SAMN27590658-SRR18763190 | PHLUSAVBR00307 | SAMN27590658 | SRR18763190 | NonPC | Public | 2021 | USA | Env | 91.2 | 100 | 0 | 98.2358 | 5168365 | 45.24 | 40 | 751045 |
| SAMN27590659-SRR18763188 | PHLUSAVBR00308 | SAMN27590659 | SRR18763188 | NonPC | Public | 2021 | USA | Env | 91.3 | 100 | 0.12 | 98.335 | 5313995 | 45.13 | 44 | 398027 |
| SAMN27590660-SRR18763187 | PHLUSAVBR00309 | SAMN27590660 | SRR18763187 | NonPC | Public | 2021 | USA | Env | 92.6 | 100 | 0 | 98.2964 | 5424899 | 44.97 | 36 | 542862 |
| SAMN27590661-SRR18763186 | PHLUSAVBR00310 | SAMN27590661 | SRR18763186 | NonPC | Public | 2021 | USA | Env | 88.8 | 100 | 0.47 | 98.2358 | 5112236 | 45.27 | 54 | 774803 |
| SAMN27590662-SRR18763185 | PHLUSAVBR00313 | SAMN27590662 | SRR18763185 | NonPC | Public | 2021 | USA | Env | 91.1 | 100 | 0.41 | 98.295 | 5267068 | 45.23 | 42 | 401469 |
| SAMN27590663-SRR18763184 | PHLUSAVBR00314 | SAMN27590663 | SRR18763184 | NonPC | Public | 2021 | USA | Env | 90.5 | 100 | 0.42 | 98.3176 | 5081931 | 45.27 | 50 | 461937 |
| SAMN27591386-SRR18819752 | PHLUSAVBR00259 | SAMN27591386 | SRR18819752 | NonPC | Public | 2021 | USA | Env | 91.2 | 100 | 0.41 | 98.3093 | 5264770 | 45.23 | 35 | 476430 |
| SAMN27591387-SRR18819751 | PHLUSAVBR00260 | SAMN27591387 | SRR18819751 | NonPC | Public | 2021 | USA | Env | 92.6 | 100 | 0 | 98.3291 | 5405400 | 45 | 43 | 730375 |
| SAMN27591388-SRR18819740 | PHLUSAVBR00261 | SAMN27591388 | SRR18819740 | NonPC | Public | 2021 | USA | Env | 90.6 | 100 | 0.14 | 98.2191 | 5043006 | 45.19 | 42 | 650554 |
| SAMN27591389-SRR18819734 | PHLUSAVBR00262 | SAMN27591389 | SRR18819734 | NonPC | Public | 2021 | USA | Env | 91.6 | 100 | 0.37 | 98.381 | 5208327 | 45.14 | 49 | 939704 |
| SAMN27591390-SRR18819733 | PHLUSAVBR00263 | SAMN27591390 | SRR18819733 | NonPC | Public | 2021 | USA | Env | 88.7 | 100 | 0.35 | 98.1585 | 5041396 | 45.19 | 46 | 269266 |
| SAMN27591391-SRR18819732 | PHLUSAVBR00264 | SAMN27591391 | SRR18819732 | NonPC | Public | 2021 | USA | Env | 89.8 | 100 | 0.27 | 98.2196 | 5262227 | 45.18 | 37 | 562047 |
| SAMN27591392-SRR18819731 | PHLUSAVBR00265 | SAMN27591392 | SRR18819731 | NonPC | Public | 2021 | USA | Env | 91.3 | 100 | 0 | 98.272 | 5260811 | 45.18 | 42 | 625490 |
| SAMN27591393-SRR18819730 | PHLUSAVBR00266 | SAMN27591393 | SRR18819730 | NonPC | Public | 2021 | USA | Env | 92.6 | 100 | 0 | 98.3315 | 5413464 | 44.97 | 32 | 530096 |
| SAMN27591394-SRR18819729 | PHLUSAVBR00267 | SAMN27591394 | SRR18819729 | NonPC | Public | 2021 | USA | Env | 91.9 | 100 | 0 | 98.3609 | 5233958 | 45.14 | 42 | 388346 |
| SAMN27591395-SRR18819728 | PHLUSAVBR00268 | SAMN27591395 | SRR18819728 | NonPC | Public | 2021 | USA | Env | 88.4 | 100 | 0.03 | 98.0631 | 5040334 | 45.25 | 33 | 441029 |
| SAMN27591396-SRR18819750 | PHLUSAVBR00269 | SAMN27591396 | SRR18819750 | NonPC | Public | 2021 | USA | Env | 91.4 | 100 | 0.27 | 98.2992 | 5140281 | 45.25 | 59 | 541210 |
| SAMN27591397-SRR18819749 | PHLUSAVBR00270 | SAMN27591397 | SRR18819749 | NonPC | Public | 2021 | USA | Env | 88.7 | 100 | 0.35 | 98.1695 | 5130794 | 45.11 | 52 | 475518 |
| SAMN27591398-SRR18819748 | PHLUSAVBR00271 | SAMN27591398 | SRR18819748 | NonPC | Public | 2021 | USA | Env | 91.2 | 100 | 0.41 | 98.2957 | 5264823 | 45.23 | 29 | 638643 |
| SAMN27591399-SRR18819747 | PHLUSAVBR00273 | SAMN27591399 | SRR18819747 | NonPC | Public | 2021 | USA | Env | 91.2 | 100 | 0.08 | 98.3019 | 5502164 | 44.99 | 77 | 266268 |
| SAMN27591400-SRR18819746 | PHLUSAVBR00274 | SAMN27591400 | SRR18819746 | NonPC | Public | 2021 | USA | Env | 91.5 | 100 | 0.22 | 98.3108 | 5271962 | 45.09 | 41 | 607156 |
| SAMN27591401-SRR18819745 | PHLUSAVBR00276 | SAMN27591401 | SRR18819745 | NonPC | Public | 2021 | USA | Env | 89.2 | 100 | 0.11 | 98.1643 | 4974756 | 45.3 | 67 | 265876 |
| SAMN27591402-SRR18819744 | PHLUSAVBR00277 | SAMN27591402 | SRR18819744 | NonPC | Public | 2021 | USA | Env | 91.7 | 100 | 0.68 | 98.2959 | 5104246 | 45.28 | 37 | 590666 |
| SAMN27591403-SRR18819743 | PHLUSAVBR00278 | SAMN27591403 | SRR18819743 | NonPC | Public | 2021 | USA | Env | 92.6 | 100 | 0 | 98.2851 | 5328401 | 45.04 | 35 | 733048 |
| SAMN27591405-SRR18819741 | PHLUSAVBR00280 | SAMN27591405 | SRR18819741 | NonPC | Public | 2021 | USA | Env | 91.5 | 100 | 0.3 | 98.2221 | 5255116 | 45.16 | 37 | 559070 |
| SAMN27591406-SRR18819739 | PHLUSAVBR00281 | SAMN27591406 | SRR18819739 | NonPC | Public | 2021 | USA | Env | 91.2 | 100 | 0.41 | 98.2979 | 5307878 | 45.19 | 34 | 517228 |
| SAMN27591407-SRR18819738 | PHLUSAVBR00282 | SAMN27591407 | SRR18819738 | NonPC | Public | 2021 | USA | Env | 91.8 | 100 | 0 | 98.2542 | 5161265 | 45.17 | 38 | 826103 |
| SAMN27591408-SRR18819737 | PHLUSAVBR00283 | SAMN27591408 | SRR18819737 | NonPC | Public | 2021 | USA | Env | 91.5 | 100 | 0.3 | 98.2422 | 5197217 | 45.17 | 31 | 580239 |
| SAMN27591409-SRR18819736 | PHLUSAVBR00284 | SAMN27591409 | SRR18819736 | NonPC | Public | 2021 | USA | Env | 88.4 | 100 | 0.03 | 98.1171 | 5041128 | 45.25 | 34 | 574397 |
| SAMN27591410-SRR18819735 | PHLUSAVBR00285 | SAMN27591410 | SRR18819735 | NonPC | Public | 2021 | USA | Env | 91.7 | 100 | 0.27 | 98.3302 | 5240159 | 45.14 | 39 | 820871 |
| SAMN27646125-SRR18823221 | PHLUSAVBR00233 | SAMN27646125 | SRR18823221 | NonPC | Public | 2021 | USA | Env | 91.2 | 100 | 0.05 | 98.2697 | 5153163 | 45.23 | 45 | 953059 |
| SAMN27646126-SRR18823220 | PHLUSAVBR00234 | SAMN27646126 | SRR18823220 | NonPC | Public | 2021 | USA | Env | 91.4 | 100 | 0.1 | 98.2413 | 5202386 | 45.25 | 49 | 411365 |
| SAMN27646127-SRR18823209 | PHLUSAVBR00235 | SAMN27646127 | SRR18823209 | NonPC | Public | 2021 | USA | Env | 91.8 | 100 | 0 | 98.3779 | 5234139 | 45.14 | 47 | 580151 |
| SAMN27646128-SRR18823203 | PHLUSAVBR00236 | SAMN27646128 | SRR18823203 | NonPC | Public | 2021 | USA | Env | 92.6 | 100 | 0 | 98.3124 | 5417348 | 44.97 | 37 | 872309 |
| SAMN27646129-SRR18823202 | PHLUSAVBR00237 | SAMN27646129 | SRR18823202 | NonPC | Public | 2021 | USA | Env | 90.8 | 100 | 0.14 | 98.1829 | 5158413 | 45.11 | 44 | 509128 |
| SAMN27646130-SRR18823201 | PHLUSAVBR00238 | SAMN27646130 | SRR18823201 | NonPC | Public | 2021 | USA | Env | 91.3 | 100 | 0 | 98.2884 | 5263294 | 45.15 | 77 | 491087 |
| SAMN27646131-SRR18823200 | PHLUSAVBR00239 | SAMN27646131 | SRR18823200 | NonPC | Public | 2021 | USA | Env | 89.2 | 100 | 0.11 | 98.1534 | 4971475 | 45.36 | 61 | 269802 |
| SAMN27646132-SRR18823199 | PHLUSAVBR00240 | SAMN27646132 | SRR18823199 | NonPC | Public | 2021 | USA | Env | 91.8 | 100 | 0 | 98.3185 | 5236152 | 45.14 | 43 | 580121 |
| SAMN27646133-SRR18823198 | PHLUSAVBR00242 | SAMN27646133 | SRR18823198 | NonPC | Public | 2021 | USA | Env | 92.6 | 100 | 0 | 98.3316 | 5326897 | 45.04 | 34 | 603211 |
| SAMN27646134-SRR18823197 | PHLUSAVBR00243 | SAMN27646134 | SRR18823197 | NonPC | Public | 2021 | USA | Env | 91.5 | 100 | 0 | 98.3009 | 5133220 | 45.24 | 48 | 565315 |
| SAMN27646135-SRR18823219 | PHLUSAVBR00244 | SAMN27646135 | SRR18823219 | NonPC | Public | 2021 | USA | Env | 91.1 | 100 | 0.2 | 98.2345 | 5167152 | 45.19 | 26 | 609288 |
| SAMN27646136-SRR18823218 | PHLUSAVBR00245 | SAMN27646136 | SRR18823218 | NonPC | Public | 2021 | USA | Env | 89.1 | 100 | 0.01 | 98.1401 | 5021988 | 45.4 | 63 | 288216 |
| SAMN27646137-SRR18823217 | PHLUSAVBR00246 | SAMN27646137 | SRR18823217 | NonPC | Public | 2021 | USA | Env | 91.6 | 100 | 0.3 | 98.2246 | 5253901 | 45.16 | 32 | 761378 |
| SAMN27646138-SRR18823216 | PHLUSAVBR00247 | SAMN27646138 | SRR18823216 | NonPC | Public | 2021 | USA | Env | 89.7 | 100 | 0.08 | 98.127 | 5093103 | 45.23 | 62 | 198422 |
| SAMN27646139-SRR18823215 | PHLUSAVBR00248 | SAMN27646139 | SRR18823215 | NonPC | Public | 2021 | USA | Env | 88.1 | 100 | 0.06 | 98.0847 | 5132071 | 45.29 | 61 | 232097 |
| SAMN27646140-SRR18823214 | PHLUSAVBR00249 | SAMN27646140 | SRR18823214 | NonPC | Public | 2021 | USA | Env | 89 | 100 | 0.01 | 98.125 | 5020239 | 45.4 | 63 | 310842 |
| SAMN27646141-SRR18823213 | PHLUSAVBR00250 | SAMN27646141 | SRR18823213 | NonPC | Public | 2021 | USA | Env | 91.5 | 100 | 0.46 | 98.2398 | 5574276 | 44.88 | 35 | 559070 |

| | | | | | | | | | | | | | | | | |
|---|---|---|---|---|---|---|---|---|---|---|---|---|---|---|---|---|
| SAMN27646142-SRR18823212 | PHLUSAVBR00251 | SAMN27646142 | SRR18823212 | NonPC | Public | 2021 | USA | Env | 90.8 | 100 | 0.14 | 98.1821 | 5155948 | 45.12 | 42 | 578209 |
| SAMN27646143-SRR18823211 | PHLUSAVBR00252 | SAMN27646143 | SRR18823211 | NonPC | Public | 2021 | USA | Env | 90.6 | 100 | 0.14 | 98.1774 | 5045098 | 45.19 | 43 | 547599 |
| SAMN27646144-SRR18823210 | PHLUSAVBR00253 | SAMN27646144 | SRR18823210 | NonPC | Public | 2021 | USA | Env | 88.7 | 100 | 0.35 | 98.1632 | 5132206 | 45.11 | 49 | 495818 |
| SAMN27646145-SRR18823208 | PHLUSAVBR00254 | SAMN27646145 | SRR18823208 | NonPC | Public | 2021 | USA | Env | 91.8 | 100 | 0 | 98.3616 | 5233800 | 45.14 | 46 | 520988 |
| SAMN27646146-SRR18823207 | PHLUSAVBR00255 | SAMN27646146 | SRR18823207 | NonPC | Public | 2021 | USA | Env | 90.6 | 100 | 0.14 | 98.2106 | 5043048 | 45.19 | 47 | 425825 |
| SAMN27646147-SRR18823206 | PHLUSAVBR00257 | SAMN27646147 | SRR18823206 | NonPC | Public | 2021 | USA | Env | 91.8 | 100 | 0 | 98.3237 | 5162093 | 45.29 | 45 | 500062 |
| SAMN27646148-SRR18823205 | PHLUSAVBR00258 | SAMN27646148 | SRR18823205 | NonPC | Public | 2021 | USA | Env | 89 | 100 | 0 | 98.0997 | 5072043 | 45.25 | 58 | 370610 |
| SAMN27646149-SRR18823204 | PHLUSAVBR00299 | SAMN27646149 | SRR18823204 | NonPC | Public | 2021 | USA | Env | 88.7 | 100 | 0 | 98.0988 | 5064999 | 45.24 | 66 | 430742 |
| SAMN27738894-SRR18889095 | PNUSAV002743 | SAMN27738894 | SRR18889095 | NonPC | Public | 2022 | USA | NA | 90.9 | 99.86 | 0 | 98.3016 | 5265990 | 45.15 | 58 | 335177 |
| SAMN27779342-SRR18921564 | PNUSAV002746 | SAMN27779342 | SRR18921564 | NonPC | Public | 2022 | NA | NA | 89.9 | 100 | 0.4 | 98.3062 | 5247873 | 45.17 | 34 | 520867 |
| SAMN27779343-SRR18922530 | PNUSAV002745 | SAMN27779343 | SRR18922530 | NonPC | Public | 2022 | NA | NA | 89.5 | 100 | 0 | 98.0798 | 5047873 | 45.26 | 69 | 318870 |
| SAMN27917784-SRR18940317 | PHLUSAVBR00241 | SAMN27917784 | SRR18940317 | NonPC | Public | 2021 | USA | Env | 91.8 | 100 | 0 | 98.3562 | 5232580 | 45.14 | 38 | 479440 |
| SAMN27917785-SRR18940316 | PHLUSAVBR00325 | SAMN27917785 | SRR18940316 | NonPC | Public | 2021 | USA | Env | 89.8 | 100 | 0.05 | 98.3374 | 5183468 | 45.22 | 48 | 370596 |
| SAMN28093304-SRR19086137 | PNUSAV002129 | SAMN28093304 | SRR19086137 | NonPC | Public | 2021 | USA | Env | 91.4 | 100 | 0.1 | 98.2423 | 5200233 | 45.25 | 46 | 522872 |
| SAMN28093305-SRR19086136 | PNUSAV002136 | SAMN28093305 | SRR19086136 | NonPC | Public | 2021 | USA | Env | 90.5 | 100 | 0.11 | 98.2058 | 5288091 | 45.13 | 52 | 330376 |
| SAMN28093306-SRR19086135 | PNUSAV002135 | SAMN28093306 | SRR19086135 | NonPC | Public | 2021 | USA | Env | 89.1 | 100 | 0.11 | 98.1823 | 4960388 | 45.34 | 68 | 269770 |
| SAMN28093307-SRR19086134 | PNUSAV002130 | SAMN28093307 | SRR19086134 | NonPC | Public | 2021 | USA | Env | 90.4 | 100 | 0.11 | 98.1903 | 5373720 | 45.06 | 54 | 543368 |
| SAMN28093308-SRR19086133 | PNUSAV002134 | SAMN28093308 | SRR19086133 | NonPC | Public | 2021 | USA | Env | 91.2 | 100 | 0.41 | 98.2815 | 5263325 | 45.23 | 33 | 549070 |
| SAMN28093309-SRR19086132 | PNUSAV002131 | SAMN28093309 | SRR19086132 | NonPC | Public | 2021 | USA | Env | 91.6 | 100 | 0.22 | 98.3458 | 5319359 | 44.94 | 34 | 801960 |
| SAMN28093310-SRR19086131 | PNUSAV002133 | SAMN28093310 | SRR19086131 | NonPC | Public | 2021 | USA | Env | 91.2 | 100 | 0.08 | 98.2454 | 5270579 | 45.21 | 48 | 519874 |
| SAMN28093311-SRR19086130 | PNUSAV002132 | SAMN28093311 | SRR19086130 | NonPC | Public | 2021 | USA | Env | 90.4 | 100 | 0.11 | 98.2337 | 5285865 | 45.13 | 49 | 543258 |
| SAMN28093312-SRR19086129 | PNUSAV002137 | SAMN28093312 | SRR19086129 | NonPC | Public | 2021 | USA | Env | 91.6 | 100 | 0.01 | 98.2887 | 5463217 | 45.01 | 68 | 274259 |
| SAMN28100960-SRR19094423 | PNUSAV002754 | SAMN28100960 | SRR19094423 | NonPC | Public | 2019 | Viet_Nam | Env | 90.9 | 100 | 0.07 | 98.302 | 5379192 | 45.37 | 48 | 940483 |
| SAMN28100961-SRR19094414 | PNUSAV002781 | SAMN28100961 | SRR19094414 | NonPC | Public | 2019 | Mexico | Env | 87.4 | 100 | 1.11 | 98.0199 | 5115405 | 45.25 | 48 | 300390 |
| SAMN28100962-SRR19094411 | PNUSAV002776 | SAMN28100962 | SRR19094411 | NonPC | Public | 2019 | Saudi_Arabia | Env | 91.8 | 100 | 0.27 | 98.2539 | 5464213 | 45.15 | 64 | 238182 |
| SAMN28100963-SRR19094406 | PNUSAV002783 | SAMN28100963 | SRR19094406 | NonPC | Public | 2020 | Ecuador | Env | 88.9 | 100 | 0.61 | 98.1045 | 5157412 | 45.24 | 65 | 171684 |
| SAMN28100964-SRR19094410 | PNUSAV002761 | SAMN28100964 | SRR19094410 | NonPC | Public | 2020 | India | Env | 89.1 | 100 | 0.19 | 98.0405 | 5419831 | 45.16 | 86 | 219325 |
| SAMN28100965-SRR19094409 | PNUSAV002782 | SAMN28100965 | SRR19094409 | NonPC | Public | 2020 | Mexico | Env | 91.5 | 100 | 0 | 98.3307 | 5273421 | 45.23 | 54 | 761872 |
| SAMN28100966-SRR19094412 | PNUSAV002751 | SAMN28100966 | SRR19094412 | NonPC | Public | 2020 | Viet_Nam | Env | 89.6 | 100 | 0.14 | 98.2769 | 5310903 | 45.27 | 60 | 236474 |
| SAMN28100967-SRR19094408 | PNUSAV002772 | SAMN28100967 | SRR19094408 | NonPC | Public | 2020 | Ecuador | Env | 89.1 | 100 | 0 | 98.073 | 5319697 | 45.16 | 121 | 122906 |
| SAMN28100968-SRR19094413 | PNUSAV002755 | SAMN28100968 | SRR19094413 | NonPC | Public | 2020 | Ecuador | Env | 88.2 | 99.93 | 0.05 | 98.1056 | 5158881 | 45.17 | 70 | 418839 |
| SAMN28100969-SRR19094404 | PNUSAV002753 | SAMN28100969 | SRR19094404 | NonPC | Public | 2020 | Saudi_Arabia | Env | 89.9 | 100 | 0.68 | 98.3399 | 5453494 | 45.13 | 45 | 458046 |
| SAMN28100970-SRR19094401 | PNUSAV002785 | SAMN28100970 | SRR19094401 | NonPC | Public | 2020 | India | Env | 89.6 | 100 | 0.05 | 98.2997 | 5042196 | 45.39 | 27 | 550288 |
| SAMN28100971-SRR19094415 | PNUSAV002797 | SAMN28100971 | SRR19094415 | NonPC | Public | 2020 | Thailand | Env | 91.7 | 100 | 0 | 98.3358 | 5347494 | 45.06 | 70 | 313064 |
| SAMN28207897-SRR19178872 | PNUSAV002813 | SAMN28207897 | SRR19178872 | NonPC | Public | 2022 | USA | Env | 88.3 | 100 | 0.39 | 98.2479 | 5183369 | 45.22 | 31 | 570365 |
| SAMN28207898-SRR19178871 | PNUSAV002818 | SAMN28207898 | SRR19178871 | NonPC | Public | 2022 | USA | Env | 91 | 100 | 0.08 | 98.2342 | 5056610 | 45.28 | 54 | 545832 |
| SAMN28207899-SRR19178873 | PNUSAV002812 | SAMN28207899 | SRR19178873 | NonPC | Public | 2022 | USA | Env | 89.6 | 100 | 0 | 98.287 | 5110375 | 45.26 | 35 | 722126 |
| SAMN28207900-SRR19178818 | PNUSAV002816 | SAMN28207900 | SRR19178818 | NonPC | Public | 2022 | USA | Env | 89.5 | 100 | 0.31 | 98.168 | 5455435 | 44.87 | 59 | 502704 |
| SAMN28207901-SRR19178875 | PNUSAV002817 | SAMN28207901 | SRR19178875 | NonPC | Public | 2022 | USA | Env | 89.5 | 100 | 0.31 | 98.1768 | 5455380 | 44.87 | 57 | 658744 |
| SAMN28207902-SRR19178820 | PNUSAV002810 | SAMN28207902 | SRR19178820 | NonPC | Public | 2022 | USA | Env | 90.5 | 100 | 0 | 98.1815 | 5109092 | 45.17 | 58 | 401126 |
| SAMN28207903-SRR19178870 | PNUSAV002809 | SAMN28207903 | SRR19178870 | NonPC | Public | 2022 | USA | Env | 89 | 100 | 0 | 98.1057 | 5410407 | 45.3 | 133 | 182516 |
| SAMN28232267-SRR19187035 | 19-VB00999 | SAMN28232267 | SRR19187035 | NonPC | Public | 2019 | Germany | Env | 90.7 | 100 | 0.22 | 98.3684 | 5123773 | 45.19 | 46 | 424318 |
| SAMN28232268-SRR19187034 | 17-VB00250 | SAMN28232268 | SRR19187034 | NonPC | Public | 2017 | Germany | Env | 87.8 | 100 | 0.58 | 98.0797 | 5018385 | 45.36 | 68 | 295483 |
| SAMN28551625-SRR19315439 | PNUSAV002831 | SAMN28551625 | SRR19315439 | NonPC | Public | 2022 | USA | NA | 90 | 100 | 0.09 | 98.2954 | 5105955 | 45.27 | 42 | 534863 |
| SAMN28816693-SRR19512859 | PNUSAV002833 | SAMN28816693 | SRR19512859 | PC | Public | 2022 | USA | NA | 97.9 | 100 | 0 | 99.9192 | 5118739 | 45.28 | 62 | 223280 |
| SAMN28860738-SRR19537971 | PNUSAV002450 | SAMN28860738 | SRR19537971 | NonPC | Public | 2021 | USA | NA | 91.5 | 99.86 | 0.05 | 98.2961 | 5097104 | 45.29 | 128 | 82166 |
| SAMN28860739-SRR19538276 | PNUSAV002725 | SAMN28860739 | SRR19538276 | NonPC | Public | 2022 | USA | NA | 91.1 | 100 | 0 | 98.2858 | 5032332 | 45.31 | 46 | 385792 |
| SAMN28860741-SRR19537972 | PNUSAV002858 | SAMN28860741 | SRR19537972 | NonPC | Public | 2022 | USA | NA | 89.1 | 100 | 0 | 98.0247 | 5092800 | 45.24 | 54 | 562383 |

| | | | | | | | | | | | | | | | | |
|---|---|---|---|---|---|---|---|---|---|---|---|---|---|---|---|---|
| SAMN28860742-SRR19537976 | PNUSAV002838 | SAMN28860742 | SRR19537976 | NonPC | Public | 2021 | USA | NA | 90 | 100 | 0 | 98.1782 | 5056107 | 45.35 | 38 | 297706 |
| SAMN28860743-SRR19538274 | PNUSAV002836 | SAMN28860743 | SRR19538274 | NonPC | Public | 2022 | USA | NA | 91 | 100 | 0 | 98.3428 | 5068218 | 45.26 | 55 | 728412 |
| SAMN28861108-SRR19539304 | PNUSAV002721 | SAMN28861108 | SRR19539304 | NonPC | Public | 2022 | USA | NA | 89.3 | 100 | 0.27 | 98.1103 | 4929776 | 45.4 | 58 | 483449 |
| SAMN28870982-SRR19543007 | PNUSAV002861 | SAMN28870982 | SRR19543007 | NonPC | Public | 2022 | USA | NA | 89.5 | 100 | 0 | 97.9995 | 5094627 | 45.24 | 61 | 361702 |
| SAMN29017552-SRR19641559 | PNUSAV002868 | SAMN29017552 | SRR19641559 | NonPC | Public | 2022 | USA | NA | 90.1 | 100 | 0.22 | 98.3544 | 5218233 | 45.14 | 49 | 502722 |
| SAMN29017553-SRR19641512 | PNUSAV002867 | SAMN29017553 | SRR19641512 | NonPC | Public | 2022 | USA | NA | 89.9 | 100 | 0.4 | 98.3061 | 5250918 | 45.17 | 35 | 551831 |
| SAMN29017554-SRR19641560 | PNUSAV002866 | SAMN29017554 | SRR19641560 | NonPC | Public | 2022 | USA | NA | 88.6 | 100 | 0.11 | 98.0522 | 4987740 | 45.32 | 32 | 534615 |
| SAMN29444629-SRR19910440 | PNUSAV002542 | SAMN29444629 | SRR19910440 | NonPC | Public | 2022 | USA | NA | 91.4 | 100 | 0 | 98.3163 | 5056707 | 45.31 | 38 | 465225 |
| SAMN29444630-SRR19910443 | PNUSAV002568 | SAMN29444630 | SRR19910443 | PC | Public | 2022 | USA | NA | 97.9 | 100 | 0 | 99.9131 | 5117059 | 45.28 | 52 | 369713 |
| SAMN29444631-SRR19910450 | PNUSAV002824 | SAMN29444631 | SRR19910450 | PC | Public | 2022 | USA | NA | 97.9 | 100 | 0 | 99.9141 | 5118878 | 45.28 | 55 | 399314 |
| SAMN29444632-SRR19910447 | PNUSAV002825 | SAMN29444632 | SRR19910447 | PC | Public | 2022 | USA | NA | 97.9 | 100 | 0 | 99.9203 | 5117652 | 45.28 | 56 | 383960 |
| SAMN29444633-SRR19910442 | PNUSAV002893 | SAMN29444633 | SRR19910442 | NonPC | Public | 2022 | USA | NA | 89.5 | 100 | 0 | 98.0782 | 5046086 | 45.27 | 65 | 318868 |
| SAMN29444868-SRR19912348 | PNUSAV002900 | SAMN29444868 | SRR19912348 | NonPC | Public | 2022 | USA | NA | 91.5 | 100 | 0 | 98.3173 | 5154911 | 45.27 | 123 | 93691 |
| SAMN29444869-SRR19912347 | PNUSAV002899 | SAMN29444869 | SRR19912347 | NonPC | Public | 2022 | USA | NA | 89.5 | 100 | 0.41 | 98.2544 | 5230433 | 45.22 | 150 | 96494 |
| SAMN29444870-SRR19912351 | PNUSAV002896 | SAMN29444870 | SRR19912351 | NonPC | Public | 2022 | USA | NA | 91.5 | 100 | 0 | 98.2601 | 5118449 | 45.21 | 49 | 574915 |
| SAMN29444871-SRR19912350 | PNUSAV002895 | SAMN29444871 | SRR19912350 | NonPC | Public | 2022 | USA | NA | 91.2 | 100 | 0.21 | 98.2289 | 5196425 | 45.26 | 27 | 552632 |
| SAMN29622003-SRR20068178 | 22MO06SH02-V1 | SAMN29622003 | SRR20068178 | NonPC | Public | 2022 | USA | Env | 89.4 | 100 | 0.54 | 98.2728 | 5253224 | 45.29 | 61 | 237648 |
| SAMN29622004-SRR20068177 | 22MO04SH02-V1 | SAMN29622004 | SRR20068177 | NonPC | Public | 2022 | USA | Env | 91.1 | 100 | 0.03 | 98.2869 | 5397107 | 45.19 | 89 | 510431 |
| SAMN29637699-SRR20083412 | 2019V-1199 | SAMN29637699 | SRR20083412 | NonPC | Public | 2019 | USA | NA | 91.5 | 100 | 0 | 98.2227 | 5130583 | 45.24 | 52 | 344098 |
| SAMN29637700-SRR20083416 | 2020V-1032 | SAMN29637700 | SRR20083416 | NonPC | Public | 2020 | USA | NA | 88.3 | 100 | 0.03 | 98.0994 | 5000447 | 45.47 | 69 | 209538 |
| SAMN29637701-SRR20083420 | 2020V-1122 | SAMN29637701 | SRR20083420 | NonPC | Public | 2020 | USA | NA | 90.1 | 100 | 0.03 | 98.2718 | 5051182 | 45.37 | 39 | 601187 |
| SAMN29637702-SRR20083419 | PNUSAV002911 | SAMN29637702 | SRR20083419 | NonPC | Public | 2022 | USA | NA | 89.6 | 100 | 0 | 98.2168 | 5217672 | 45.23 | 38 | 776746 |
| SAMN29658966-SRR20095042 | PNUSAV002907 | SAMN29658966 | SRR20095042 | NonPC | Public | 2022 | USA | NA | 90.9 | 100 | 0.33 | 98.3535 | 5159644 | 45.27 | 24 | 520110 |
| SAMN29658970-SRR20096757 | PNUSAV002905 | SAMN29658970 | SRR20096757 | NonPC | Public | 2022 | USA | NA | 88.3 | 100 | 0.3 | 98.2131 | 5102574 | 45.27 | 54 | 473950 |
| SAMN29658975-SRR20096754 | PNUSAV002904 | SAMN29658975 | SRR20096754 | NonPC | Public | 2022 | USA | NA | 91.2 | 100 | 0.27 | 98.3428 | 5054229 | 45.3 | 43 | 848599 |
| SAMN29658980-SRR20096759 | PNUSAV002910 | SAMN29658980 | SRR20096759 | NonPC | Public | 2022 | USA | NA | 91.7 | 100 | 0.29 | 98.3039 | 5112068 | 45.27 | 29 | 890040 |
| SAMN29662269-SRR20106138 | PNUSAV002915 | SAMN29662269 | SRR20106138 | NonPC | Public | NA | USA | NA | 91.5 | 100 | 0 | 98.3526 | 5093566 | 45.26 | 49 | 397992 |
| SAMN29843622-SRR20325127 | PNUSAV002942 | SAMN29843622 | SRR20325127 | NonPC | Public | 2022 | USA | NA | 91.6 | 100 | 0.4 | 98.2642 | 5403116 | 44.98 | 141 | 86934 |
| SAMN29843623-SRR20325128 | PNUSAV002941 | SAMN29843623 | SRR20325128 | NonPC | Public | 2022 | USA | NA | 89.6 | 100 | 0.11 | 98.3031 | 5183135 | 45.24 | 47 | 381295 |
| SAMN29844259-SRR20325461 | PNUSAV002919 | SAMN29844259 | SRR20325461 | NonPC | Public | 2022 | USA | NA | 88.8 | 100 | 0 | 98.097 | 4931906 | 45.38 | 95 | 165596 |
| SAMN29844260-SRR20325462 | PNUSAV002940 | SAMN29844260 | SRR20325462 | NonPC | Public | 2022 | USA | NA | 91.6 | 100 | 0 | 98.4024 | 5018212 | 45.34 | 39 | 280566 |
| SAMN29853357-SRR20334240 | PNUSAV002951 | SAMN29853357 | SRR20334240 | NonPC | Public | NA | USA | NA | 91.5 | 99.86 | 0 | 98.3217 | 5094530 | 45.27 | 55 | 461950 |
| SAMN29853359-SRR20334242 | PNUSAV002950 | SAMN29853359 | SRR20334242 | NonPC | Public | NA | USA | NA | 91.5 | 100 | 0 | 98.3377 | 5095264 | 45.27 | 52 | 548679 |
| SPVP110001 | SPVP110001 | SAMN20298569 | SRR15185230 | NonPC | Public | 2011 | China | Env | 88.2 | 100 | 0.39 | 98.2388 | 5026729 | 45.26 | 59 | 474301 |
| SPVP110002 | SPVP110002 | SAMN20298567 | SRR15185234 | NonPC | Public | 2011 | China | Env | 91.5 | 100 | 0 | 98.3711 | 5304752 | 45.18 | 83 | 268759 |
| SPVP110003 | SPVP110003 | SAMN20298568 | SRR15185233 | NonPC | Public | 2011 | China | Env | 91.1 | 100 | 0.37 | 98.322 | 5033409 | 45.34 | 51 | 376341 |
| SPVP110006 | SPVP110006 | SAMN20298566 | SRR15185235 | NonPC | Public | 2011 | China | Env | 91.4 | 100 | 0.14 | 98.2996 | 5338528 | 45.12 | 48 | 365838 |
| SPVP110007 | SPVP110007 | SAMN20298570 | SRR15185229 | NonPC | Public | 2011 | China | Env | 88.3 | 100 | 0.37 | 98.2235 | 5074559 | 45.31 | 76 | 326939 |
| SPVP110009 | SPVP110009 | SAMN20298571 | SRR15185228 | NonPC | Public | 2011 | China | Env | 88.3 | 100 | 0.34 | 98.1912 | 5247849 | 45.19 | 60 | 453013 |
| SPVP110012 | SPVP110012 | SAMN20298574 | SRR15185225 | NonPC | Public | 2011 | China | Env | 91.5 | 100 | 0.08 | 98.3701 | 5031635 | 45.33 | 55 | 459002 |
| SPVP110013 | SPVP110013 | SAMN20298573 | SRR15185226 | NonPC | Public | 2011 | China | Env | 88.7 | 100 | 0.39 | 98.2162 | 5080035 | 45.29 | 61 | 401891 |
| SPVP110014 | SPVP110014 | SAMN20298572 | SRR15185227 | NonPC | Public | 2011 | China | Env | 92 | 100 | 0.12 | 98.3532 | 5250897 | 45.31 | 64 | 259771 |
| SPVP110015 | SPVP110015 | SAMN20298575 | SRR15185224 | NonPC | Public | 2011 | China | Env | 91.4 | 100 | 0.03 | 98.4156 | 5032696 | 45.33 | 57 | 383133 |
| SPVP110016 | SPVP110016 | SAMN20352791 | SRR15219062 | NonPC | Public | 2014 | China | Env | 89.9 | 100 | 0.09 | 98.3197 | 4975743 | 45.44 | 32 | 528332 |
| SPVP110018 | SPVP110018 | SAMN20352793 | SRR15219060 | NonPC | Public | 2014 | China | Env | 91.2 | 100 | 0.27 | 98.2401 | 5284616 | 45.22 | 92 | 245534 |
| SPVP110019 | SPVP110019 | SAMN20352792 | SRR15219061 | NonPC | Public | 2014 | China | Env | 90.6 | 100 | 0.14 | 98.2824 | 5095120 | 45.25 | 54 | 511654 |
| SPVP110021 | SPVP110021 | SAMN20298578 | SRR15185221 | NonPC | Public | 2011 | China | Env | 91.6 | 100 | 0 | 98.3169 | 5062053 | 45.29 | 60 | 457948 |
| SPVP110022 | SPVP110022 | SAMN20298579 | SRR15185219 | NonPC | Public | 2011 | China | Env | 88.1 | 100 | 0.3 | 98.2362 | 5138585 | 45.24 | 36 | 538104 |

| | | | | | | | | | | | | | | | | |
|---|---|---|---|---|---|---|---|---|---|---|---|---|---|---|---|---|
| SPVP110025 | SPVP110025 | SAMN20298580 | SRR15185218 | NonPC | Public | 2011 | China | Env | 90.1 | 100 | 0.6 | 98.3597 | 5151522 | 45.28 | 50 | 503711 |
| SPVP110027 | SPVP110027 | SAMN20298581 | SRR15185217 | NonPC | Public | 2011 | China | Env | 88.5 | 100 | 0.39 | 98.2601 | 5031374 | 45.25 | 38 | 1167231 |
| SPVP110029 | SPVP110029 | SAMN20298584 | SRR15185214 | NonPC | Public | 2011 | China | Env | 88.4 | 100 | 0.39 | 98.2362 | 5054406 | 45.3 | 61 | 471620 |
| SPVP110030 | SPVP110030 | SAMN20298585 | SRR15185213 | NonPC | Public | 2011 | China | Env | 91.4 | 100 | 0.01 | 98.3457 | 5254423 | 45.22 | 55 | 516131 |
| SPVP110031 | SPVP110031 | SAMN20298583 | SRR15185215 | NonPC | Public | 2011 | China | Env | 88.3 | 100 | 0.46 | 98.2573 | 5026330 | 45.34 | 70 | 462849 |
| SPVP110032 | SPVP110032 | SAMN20298582 | SRR15185216 | NonPC | Public | 2011 | China | Env | 89.9 | 100 | 0 | 98.353 | 5083754 | 45.33 | 46 | 460763 |
| SPVP110033 | SPVP110033 | SAMN20298586 | SRR15185212 | NonPC | Public | 2011 | China | Env | 88.2 | 100 | 0.58 | 98.1511 | 5191744 | 45.18 | 71 | 391144 |
| SPVP110035 | SPVP110035 | SAMN20298591 | SRR15185206 | NonPC | Public | 2011 | China | Env | 91.7 | 100 | 0 | 98.3397 | 5087440 | 45.3 | 50 | 458236 |
| SPVP110036 | SPVP110036 | SAMN20298599 | SRR15185197 | NonPC | Public | 2011 | China | Env | 89.4 | 100 | 0.14 | 98.3441 | 4978693 | 45.38 | 27 | 643927 |
| SPVP110038 | SPVP110038 | SAMN20298600 | SRR15185196 | NonPC | Public | 2011 | China | Env | 91.3 | 100 | 0 | 98.3587 | 4935652 | 45.46 | 41 | 374699 |
| SPVP110039 | SPVP110039 | SAMN20352888 | SRR15218780 | NonPC | Public | 2014 | China | Env | 88.6 | 100 | 0.41 | 98.2322 | 5121499 | 45.2 | 55 | 370797 |
| SPVP110040 | SPVP110040 | SAMN20352901 | SRR15219373 | NonPC | Public | 2014 | China | Env | 90.1 | 100 | 0.09 | 98.3371 | 5067617 | 45.29 | 47 | 457102 |
| SPVP110041 | SPVP110041 | SAMN20298592 | SRR15185205 | NonPC | Public | 2011 | China | Env | 91.3 | 100 | 0 | 98.3852 | 5037706 | 45.31 | 47 | 452490 |
| SPVP110044 | SPVP110044 | SAMN20298593 | SRR15185204 | NonPC | Public | 2011 | China | Env | 89.6 | 100 | 0.07 | 98.3085 | 5165372 | 45.35 | 60 | 227964 |
| SPVP110046 | SPVP110046 | SAMN20298594 | SRR15185203 | NonPC | Public | 2011 | China | Env | 89.7 | 100 | 0 | 98.2256 | 4998672 | 45.32 | 43 | 418123 |
| SPVP110047 | SPVP110047 | SAMN20298596 | SRR15185201 | NonPC | Public | 2011 | China | Env | 91 | 100 | 0.05 | 98.356 | 5120829 | 45.26 | 46 | 451152 |
| SPVP110048 | SPVP110048 | SAMN20298595 | SRR15185202 | NonPC | Public | 2011 | China | Env | 87.5 | 100 | 0.3 | 98.2358 | 5038107 | 45.24 | 63 | 710672 |
| SPVP110050 | SPVP110050 | SAMN20352941 | SRR15219260 | NonPC | Public | 2014 | China | Env | 91.7 | 100 | 0.09 | 98.3913 | 5079113 | 45.28 | 41 | 410844 |
| SPVP110051 | SPVP110051 | SAMN20352942 | SRR15219259 | NonPC | Public | 2014 | China | Env | 89.8 | 100 | 0 | 98.2686 | 4948655 | 45.41 | 37 | 555360 |
| SPVP110052 | SPVP110052 | SAMN20352943 | SRR15219258 | NonPC | Public | 2014 | China | Env | 89.7 | 100 | 0.02 | 98.2851 | 5113923 | 45.26 | 58 | 547182 |
| SPVP110053 | SPVP110053 | SAMN20298754 | SRR15185037 | NonPC | Public | 2011 | China | Env | 88.2 | 100 | 0.39 | 98.2283 | 5027038 | 45.26 | 53 | 542356 |
| SPVP110054 | SPVP110054 | SAMN20298759 | SRR15185031 | NonPC | Public | 2011 | China | Env | 91.3 | 100 | 0.24 | 98.3658 | 5193991 | 45.19 | 60 | 420504 |
| SPVP110055 | SPVP110055 | SAMN20298760 | SRR15185030 | NonPC | Public | 2011 | China | Env | 91 | 100 | 0.03 | 98.2867 | 5138997 | 45.23 | 88 | 334380 |
| SPVP110056 | SPVP110056 | SAMN20298795 | SRR15184919 | NonPC | Public | 2011 | China | Env | 90.9 | 100 | 0.19 | 98.3337 | 5221240 | 45.26 | 85 | 429468 |
| SPVP110057 | SPVP110057 | SAMN20298796 | SRR15184918 | NonPC | Public | 2011 | China | Env | 90.9 | 100 | 0.19 | 98.3538 | 5221315 | 45.26 | 83 | 429468 |
| SPVP110058 | SPVP110058 | SAMN20298793 | SRR15184921 | NonPC | Public | 2011 | China | Env | 89.7 | 100 | 0.05 | 98.3431 | 5067974 | 45.2 | 45 | 452142 |
| SPVP110059 | SPVP110059 | SAMN20298794 | SRR15184920 | NonPC | Public | 2011 | China | Env | 90.9 | 100 | 0.07 | 98.3122 | 5032713 | 45.36 | 55 | 458188 |
| SPVP110061 | SPVP110061 | SAMN20298799 | SRR15184914 | NonPC | Public | 2011 | China | Env | 88.3 | 100 | 0.63 | 98.2484 | 5071932 | 45.2 | 51 | 560284 |
| SPVP110062 | SPVP110062 | SAMN20298800 | SRR15184913 | NonPC | Public | 2011 | China | Env | 88.8 | 100 | 0.63 | 98.2181 | 5002442 | 45.3 | 42 | 277648 |
| SPVP110063 | SPVP110063 | SAMN20298797 | SRR15184917 | NonPC | Public | 2011 | China | Env | 88.5 | 100 | 0.41 | 98.2347 | 5111386 | 45.24 | 57 | 380459 |
| SPVP110064 | SPVP110064 | SAMN20298801 | SRR15184912 | NonPC | Public | 2011 | China | Env | 90.9 | 100 | 0 | 98.3505 | 5139146 | 45.26 | 36 | 353987 |
| SPVP110065 | SPVP110065 | SAMN20298798 | SRR15184916 | NonPC | Public | 2011 | China | Env | 90 | 100 | 0.05 | 98.3428 | 5007476 | 45.31 | 39 | 781368 |
| SPVP110066 | SPVP110066 | SAMN20298787 | SRR15184928 | NonPC | Public | 2011 | China | Env | 91.1 | 100 | 0.54 | 98.2858 | 5224750 | 45.4 | 73 | 471778 |
| SPVP110068 | SPVP110068 | SAMN20298772 | SRR15185016 | NonPC | Public | 2011 | China | Env | 88.6 | 100 | 0.34 | 98.186 | 5157293 | 45.21 | 56 | 460672 |
| SPVP110069 | SPVP110069 | SAMN20298775 | SRR15185013 | NonPC | Public | 2011 | China | Env | 91.4 | 100 | 0.05 | 98.3528 | 5180522 | 45.22 | 67 | 306487 |
| SPVP110070 | SPVP110070 | SAMN20298776 | SRR15185012 | NonPC | Public | 2011 | China | Env | 88.6 | 100 | 0.65 | 98.191 | 5424028 | 45.09 | 59 | 683779 |
| SPVP110073 | SPVP110073 | SAMN20298788 | SRR15184927 | NonPC | Public | 2011 | China | Env | 91.1 | 100 | 0.54 | 98.2952 | 5224674 | 45.4 | 66 | 471778 |
| SPVP110074 | SPVP110074 | SAMN20298771 | SRR15185017 | NonPC | Public | 2011 | China | Env | 88.2 | 100 | 0.39 | 98.2427 | 4990641 | 45.3 | 48 | 316494 |
| SPVP110076 | SPVP110076 | SAMN20298777 | SRR15185011 | NonPC | Public | 2011 | China | Env | 88.3 | 100 | 0.36 | 98.2274 | 5055211 | 45.2 | 92 | 541480 |
| SPVP110077 | SPVP110077 | SAMN20298785 | SRR15184930 | NonPC | Public | 2011 | China | Env | 88.5 | 100 | 0.39 | 98.2475 | 5060083 | 45.26 | 76 | 426902 |
| SPVP110078 | SPVP110078 | SAMN20298786 | SRR15184929 | NonPC | Public | 2011 | China | Env | 88.5 | 100 | 0.39 | 98.1945 | 5060008 | 45.26 | 77 | 338839 |
| SPVP110079 | SPVP110079 | SAMN20298778 | SRR15185010 | NonPC | Public | 2011 | China | Env | 92.3 | 100 | 0 | 98.4174 | 5166992 | 45.31 | 35 | 492786 |
| SPVP110080 | SPVP110080 | SAMN20298770 | SRR15185018 | NonPC | Public | 2011 | China | Env | 89.9 | 100 | 0.05 | 98.3188 | 5183008 | 45.31 | 42 | 375562 |
| SPVP110081 | SPVP110081 | SAMN20298779 | SRR15185007 | NonPC | Public | 2011 | China | Env | 90.4 | 100 | 0 | 98.395 | 5015493 | 45.33 | 34 | 449751 |
| SPVP110082 | SPVP110082 | SAMN20298773 | SRR15185015 | NonPC | Public | 2011 | China | Env | 88.4 | 100 | 0.58 | 98.2818 | 5189686 | 45.21 | 99 | 450079 |
| SPVP110083 | SPVP110083 | SAMN20298780 | SRR15184935 | NonPC | Public | 2011 | China | Env | 91.4 | 100 | 0.11 | 98.3617 | 5016749 | 45.32 | 41 | 525386 |
| SPVP110085 | SPVP110085 | SAMN20298781 | SRR15184934 | NonPC | Public | 2011 | China | Env | 88.6 | 100 | 0.66 | 98.2286 | 5013722 | 45.29 | 59 | 427941 |
| SPVP110086 | SPVP110086 | SAMN20298782 | SRR15184933 | NonPC | Public | 2011 | China | Env | 92.5 | 100 | 0.54 | 98.3043 | 5272730 | 45.24 | 77 | 425333 |

| | | | | | | | | | | | | | | | | |
|---|---|---|---|---|---|---|---|---|---|---|---|---|---|---|---|---|
| SPVP110087 | SPVP110087 | SAMN20298783 | SRR15184932 | NonPC | Public | 2011 | China | Env | 91.2 | 100 | 0.6 | 98.3441 | 5383367 | 45.26 | 107 | 400867 |
| SPVP110088 | SPVP110088 | SAMN20298769 | SRR15185019 | NonPC | Public | 2011 | China | Env | 91.2 | 100 | 0.27 | 98.2467 | 5384549 | 45.2 | 111 | 333585 |
| SPVP110089 | SPVP110089 | SAMN20298784 | SRR15184931 | NonPC | Public | 2011 | China | Env | 88.3 | 100 | 0.39 | 98.2247 | 5049515 | 45.24 | 43 | 589113 |
| SPVP110090 | SPVP110090 | SAMN20298774 | SRR15185014 | NonPC | Public | 2011 | China | Env | 88.3 | 100 | 0.66 | 98.2827 | 5060153 | 45.23 | 80 | 544015 |
| SPVP110091 | SPVP110091 | SAMN20298809 | SRR15184903 | NonPC | Public | 2011 | China | Env | 89.5 | 100 | 0.07 | 98.2795 | 5050431 | 45.31 | 37 | 617757 |
| SPVP110092 | SPVP110092 | SAMN20298807 | SRR15184906 | NonPC | Public | 2011 | China | Env | 91.7 | 100 | 0 | 98.2999 | 5063868 | 45.33 | 57 | 373764 |
| SPVP110093 | SPVP110093 | SAMN20298806 | SRR15184907 | NonPC | Public | 2011 | China | Env | 91.4 | 100 | 0.1 | 98.351 | 5308028 | 45.13 | 61 | 404054 |
| SPVP110094 | SPVP110094 | SAMN20298810 | SRR15184901 | NonPC | Public | 2011 | China | Env | 91.3 | 100 | 0.27 | 98.3432 | 5068282 | 45.36 | 42 | 395341 |
| SPVP110095 | SPVP110095 | SAMN20298805 | SRR15184908 | NonPC | Public | 2011 | China | Env | 92.2 | 100 | 0.12 | 98.334 | 5143119 | 45.22 | 62 | 465106 |
| SPVP110096 | SPVP110096 | SAMN20298811 | SRR15184829 | NonPC | Public | 2011 | China | Env | 88.3 | 100 | 0.3 | 98.1844 | 5027563 | 45.29 | 55 | 363287 |
| SPVP110098 | SPVP110098 | SAMN20298812 | SRR15184828 | NonPC | Public | 2011 | China | Env | 88.2 | 100 | 0.36 | 98.2335 | 5025266 | 45.31 | 54 | 339621 |
| SPVP110099 | SPVP110099 | SAMN20298808 | SRR15184905 | NonPC | Public | 2011 | China | Env | 89.8 | 100 | 0.27 | 98.2855 | 5062438 | 45.38 | 61 | 175072 |
| SPVP110100 | SPVP110100 | SAMN20298813 | SRR15184827 | NonPC | Public | 2011 | China | Env | 91.5 | 100 | 0.27 | 98.3237 | 5158145 | 45.21 | 62 | 358040 |
| SPVP110101 | SPVP110101 | SAMN20298814 | SRR15184826 | NonPC | Public | 2011 | China | Env | 91.5 | 100 | 0.27 | 98.3408 | 5159329 | 45.21 | 59 | 363278 |
| SPVP130001 | SPVP130001 | SAMN20352521 | SRR15218995 | NonPC | Public | 2013 | China | Env | 89.6 | 100 | 0 | 98.3737 | 5088667 | 45.26 | 56 | 236642 |
| SPVP130002 | SPVP130002 | SAMN20352530 | SRR15218985 | NonPC | Public | 2013 | China | Env | 88.1 | 100 | 0.39 | 98.2292 | 5169307 | 45.23 | 35 | 746404 |
| SPVP130003 | SPVP130003 | SAMN20352533 | SRR15218982 | NonPC | Public | 2013 | China | Env | 88.3 | 100 | 0.33 | 98.3033 | 5086864 | 45.22 | 52 | 445620 |
| SPVP130004 | SPVP130004 | SAMN20352524 | SRR15218992 | NonPC | Public | 2013 | China | Env | 91.5 | 100 | 0.27 | 98.3235 | 5091944 | 45.32 | 49 | 537307 |
| SPVP130005 | SPVP130005 | SAMN20352535 | SRR15218980 | NonPC | Public | 2013 | China | Env | 88.2 | 100 | 0.39 | 98.2026 | 5059883 | 45.21 | 53 | 542137 |
| SPVP130006 | SPVP130006 | SAMN20352527 | SRR15218989 | NonPC | Public | 2013 | China | Env | 90.9 | 100 | 0 | 98.3223 | 5001969 | 45.36 | 39 | 401256 |
| SPVP130007 | SPVP130007 | SAMN20352537 | SRR15218978 | NonPC | Public | 2013 | China | Env | 88.2 | 100 | 0.39 | 98.1786 | 5071742 | 45.24 | 55 | 485301 |
| SPVP130008 | SPVP130008 | SAMN20352534 | SRR15218981 | NonPC | Public | 2013 | China | Env | 88.3 | 100 | 0.33 | 98.3046 | 5086443 | 45.22 | 48 | 445620 |
| SPVP130009 | SPVP130009 | SAMN20352536 | SRR15218979 | NonPC | Public | 2013 | China | Env | 88.2 | 100 | 0.39 | 98.1835 | 5048123 | 45.25 | 54 | 542137 |
| SPVP130010 | SPVP130010 | SAMN20352526 | SRR15218990 | NonPC | Public | 2013 | China | Env | 88.8 | 100 | 0.36 | 98.2311 | 5158746 | 45.18 | 61 | 500962 |
| SPVP130011 | SPVP130011 | SAMN20352547 | SRR15218931 | NonPC | Public | 2013 | China | Env | 88.3 | 100 | 0.39 | 98.2206 | 5072580 | 45.24 | 54 | 576177 |
| SPVP130012 | SPVP130012 | SAMN20352541 | SRR15218937 | NonPC | Public | 2013 | China | Env | 89.8 | 100 | 0.42 | 98.3226 | 4989492 | 45.4 | 35 | 528695 |
| SPVP130013 | SPVP130013 | SAMN20352542 | SRR15218936 | NonPC | Public | 2013 | China | Env | 91.3 | 100 | 0 | 98.3798 | 5039093 | 45.35 | 51 | 375797 |
| SPVP130014 | SPVP130014 | SAMN20352543 | SRR15218935 | NonPC | Public | 2013 | China | Env | 91 | 100 | 0.27 | 98.3784 | 5023959 | 45.28 | 32 | 448978 |
| SPVP130015 | SPVP130015 | SAMN20352544 | SRR15218934 | NonPC | Public | 2013 | China | Env | 90.1 | 100 | 0.07 | 98.338 | 5038430 | 45.34 | 67 | 405623 |
| SPVP130016 | SPVP130016 | SAMN20352545 | SRR15218933 | NonPC | Public | 2013 | China | Env | 89.7 | 100 | 0.12 | 98.3365 | 5026853 | 45.34 | 45 | 529981 |
| SPVP130017 | SPVP130017 | SAMN20352546 | SRR15218932 | NonPC | Public | 2013 | China | Env | 91.9 | 100 | 0.27 | 98.3939 | 5159190 | 45.22 | 84 | 403606 |
| SPVP130018 | SPVP130018 | SAMN20352528 | SRR15218988 | NonPC | Public | 2013 | China | Env | 89.6 | 100 | 0.27 | 98.3058 | 5070062 | 45.25 | 59 | 454338 |
| SPVP130019 | SPVP130019 | SAMN20352529 | SRR15218987 | NonPC | Public | 2013 | China | Env | 89.8 | 100 | 0.14 | 98.338 | 5126054 | 45.26 | 43 | 448058 |
| SPVP130020 | SPVP130020 | SAMN20352531 | SRR15218984 | NonPC | Public | 2013 | China | Env | 91.8 | 100 | 0.33 | 98.4039 | 5115259 | 45.28 | 66 | 463065 |
| SPVP130021 | SPVP130021 | SAMN20352525 | SRR15218991 | NonPC | Public | 2013 | China | Env | 92.1 | 100 | 0 | 98.3329 | 5039021 | 45.31 | 37 | 403061 |
| SPVP130022 | SPVP130022 | SAMN20352532 | SRR15218983 | NonPC | Public | 2013 | China | Env | 91.7 | 100 | 0.33 | 98.3732 | 5106542 | 45.31 | 101 | 182923 |
| SPVP130024 | SPVP130024 | SAMN20352551 | SRR15218926 | NonPC | Public | 2013 | China | Env | 88.2 | 100 | 0.4 | 98.2538 | 5135568 | 45.13 | 97 | 724202 |
| SPVP130026 | SPVP130026 | SAMN20352550 | SRR15218927 | NonPC | Public | 2013 | China | Env | 91.3 | 100 | 0 | 98.4312 | 4964763 | 45.41 | 41 | 449783 |
| SPVP130029 | SPVP130029 | SAMN20352549 | SRR15218929 | NonPC | Public | 2013 | China | Env | 87.9 | 100 | 0.34 | 98.288 | 5073099 | 45.32 | 49 | 454430 |
| SPVP130039 | SPVP130039 | SAMN20352606 | SRR15219337 | NonPC | Public | 2013 | China | Env | 88.9 | 100 | 0.68 | 98.3953 | 5184753 | 45.08 | 79 | 175967 |
| SPVP130040 | SPVP130040 | SAMN20352607 | SRR15219336 | NonPC | Public | 2013 | China | Env | 88.9 | 100 | 0.68 | 98.3769 | 5185373 | 45.08 | 78 | 176833 |
| SPVP130041 | SPVP130041 | SAMN20352601 | SRR15219342 | NonPC | Public | 2013 | China | Env | 91.6 | 100 | 0.09 | 98.3536 | 5118426 | 45.3 | 77 | 460510 |
| SPVP130042 | SPVP130042 | SAMN20352597 | SRR15219416 | NonPC | Public | 2013 | China | Env | 90.7 | 100 | 0.2 | 98.3437 | 5097425 | 45.31 | 53 | 227571 |
| SPVP130043 | SPVP130043 | SAMN20352602 | SRR15219341 | NonPC | Public | 2013 | China | Env | 89.2 | 100 | 0.07 | 98.3535 | 5005194 | 45.36 | 56 | 741961 |
| SPVP130044 | SPVP130044 | SAMN20352596 | SRR15219417 | NonPC | Public | 2013 | China | Env | 91.4 | 100 | 0.03 | 98.4306 | 5081906 | 45.29 | 43 | 449114 |
| SPVP130045 | SPVP130045 | SAMN20352603 | SRR15219340 | NonPC | Public | 2013 | China | Env | 88.1 | 100 | 0.93 | 98.194 | 5024568 | 45.32 | 59 | 175101 |
| SPVP130046 | SPVP130046 | SAMN20352605 | SRR15219338 | NonPC | Public | 2013 | China | Env | 92.2 | 100 | 0.14 | 98.3076 | 5306153 | 45.1 | 57 | 424897 |
| SPVP130047 | SPVP130047 | SAMN20352595 | SRR15219418 | NonPC | Public | 2013 | China | Env | 89.2 | 100 | 0 | 98.2774 | 5057338 | 45.3 | 42 | 383856 |

| | | | | | | | | | | | | | | | | |
|---|---|---|---|---|---|---|---|---|---|---|---|---|---|---|---|---|
| SPVP130048 | SPVP130048 | SAMN20352600 | SRR15219343 | NonPC | Public | 2013 | China | Env | 91.6 | 100 | 0.09 | 98.3298 | 5232390 | 45.27 | 66 | 451554 |
| SPVP130049 | SPVP130049 | SAMN20352594 | SRR15219419 | NonPC | Public | 2013 | China | Env | 91.1 | 100 | 0.41 | 98.1998 | 5205013 | 45.23 | 80 | 313254 |
| SPVP130050 | SPVP130050 | SAMN20352598 | SRR15219415 | NonPC | Public | 2013 | China | Env | 92.1 | 100 | 0 | 98.3251 | 5125456 | 45.29 | 59 | 353703 |
| SPVP130051 | SPVP130051 | SAMN20352604 | SRR15219339 | NonPC | Public | 2013 | China | Env | 89.9 | 100 | 0.14 | 98.2455 | 5064791 | 45.25 | 42 | 399797 |
| SPVP130053 | SPVP130053 | SAMN20352599 | SRR15219414 | NonPC | Public | 2013 | China | Env | 90.1 | 100 | 0.05 | 98.3772 | 5047074 | 45.27 | 44 | 566442 |
| SPVP130054 | SPVP130054 | SAMN20352608 | SRR15219335 | NonPC | Public | 2013 | China | Env | 88.2 | 100 | 0.39 | 98.1663 | 5041391 | 45.26 | 54 | 661479 |
| SPVP130055 | SPVP130055 | SAMN20352576 | SRR15219439 | NonPC | Public | 2013 | China | Env | 90 | 100 | 0.12 | 98.2892 | 5173540 | 45.35 | 48 | 400902 |
| SPVP130056 | SPVP130056 | SAMN20352581 | SRR15219433 | NonPC | Public | 2013 | China | Env | 88.1 | 100 | 0.3 | 98.2137 | 5140368 | 45.25 | 36 | 746404 |
| SPVP130057 | SPVP130057 | SAMN20352571 | SRR15219444 | NonPC | Public | 2013 | China | Env | 88.5 | 100 | 0.93 | 98.2779 | 5105712 | 45.33 | 55 | 500022 |
| SPVP130058 | SPVP130058 | SAMN20352577 | SRR15219438 | NonPC | Public | 2013 | China | Env | 88.6 | 100 | 0.78 | 98.2434 | 5090021 | 45.22 | 47 | 1146418 |
| SPVP130059 | SPVP130059 | SAMN20352578 | SRR15219437 | NonPC | Public | 2013 | China | Env | 87.5 | 99.93 | 0 | 98.2645 | 5027196 | 45.39 | 114 | 115718 |
| SPVP130060 | SPVP130060 | SAMN20352610 | SRR15219332 | NonPC | Public | 2013 | China | Env | 89.6 | 100 | 0.6 | 98.2896 | 5019943 | 45.37 | 23 | 1174679 |
| SPVP130061 | SPVP130061 | SAMN20352612 | SRR15219330 | NonPC | Public | 2013 | China | Env | 88.3 | 100 | 0.39 | 98.2104 | 5072844 | 45.21 | 53 | 661387 |
| SPVP130062 | SPVP130062 | SAMN20352611 | SRR15219331 | NonPC | Public | 2013 | China | Env | 88.3 | 100 | 0.66 | 98.2122 | 5074964 | 45.19 | 51 | 752462 |
| SPVP130063 | SPVP130063 | SAMN20352609 | SRR15219334 | NonPC | Public | 2013 | China | Env | 88.4 | 100 | 0.36 | 98.2132 | 5084819 | 45.21 | 63 | 539686 |
| SPVP130064 | SPVP130064 | SAMN20352506 | SRR15218488 | NonPC | Public | 2013 | China | Env | 91.3 | 100 | 0.27 | 98.3112 | 5094176 | 45.29 | 67 | 242100 |
| SPVP130065 | SPVP130065 | SAMN20352572 | SRR15219443 | NonPC | Public | 2013 | China | Env | 88.3 | 100 | 0.36 | 98.2633 | 5045359 | 45.31 | 43 | 543732 |
| SPVP130066 | SPVP130066 | SAMN20352573 | SRR15219442 | NonPC | Public | 2013 | China | Env | 92.2 | 100 | 0.14 | 98.3091 | 5328175 | 45.1 | 59 | 424897 |
| SPVP130068 | SPVP130068 | SAMN20352579 | SRR15219436 | NonPC | Public | 2013 | China | Env | 91.8 | 100 | 0.05 | 98.2578 | 5312165 | 45.19 | 45 | 362235 |
| SPVP130069 | SPVP130069 | SAMN20352570 | SRR15219445 | NonPC | Public | 2013 | China | Env | 89.9 | 100 | 0.33 | 98.3779 | 5105596 | 45.37 | 42 | 775213 |
| SPVP130070 | SPVP130070 | SAMN20352625 | SRR15219316 | NonPC | Public | 2013 | China | Env | 90.9 | 100 | 0.07 | 98.3318 | 5165777 | 45.25 | 71 | 361436 |
| SPVP130071 | SPVP130071 | SAMN20352626 | SRR15219315 | NonPC | Public | 2013 | China | Env | 90.9 | 100 | 0.07 | 98.334 | 5165567 | 45.25 | 70 | 361435 |
| SPVP130072 | SPVP130072 | SAMN20352634 | SRR15219200 | NonPC | Public | 2013 | China | Env | 90.4 | 100 | 0 | 98.3199 | 5132737 | 45.11 | 43 | 317965 |
| SPVP130073 | SPVP130073 | SAMN20352635 | SRR15219199 | NonPC | Public | 2013 | China | Env | 91.9 | 100 | 0 | 98.424 | 5070085 | 45.36 | 52 | 269851 |
| SPVP130074 | SPVP130074 | SAMN20352636 | SRR15219198 | NonPC | Public | 2013 | China | Env | 91.9 | 100 | 0 | 98.3573 | 5062128 | 45.33 | 55 | 403388 |
| SPVP130075 | SPVP130075 | SAMN20352632 | SRR15219202 | NonPC | Public | 2013 | China | Env | 91.1 | 100 | 0 | 98.3716 | 5121408 | 45.28 | 61 | 260888 |
| SPVP130076 | SPVP130076 | SAMN20352637 | SRR15219197 | NonPC | Public | 2013 | China | Env | 92.4 | 100 | 0.27 | 98.365 | 5017436 | 45.38 | 40 | 403599 |
| SPVP130078 | SPVP130078 | SAMN20352638 | SRR15219196 | NonPC | Public | 2013 | China | Env | 90 | 100 | 0.09 | 98.2806 | 4974313 | 45.42 | 45 | 512167 |
| SPVP130079 | SPVP130079 | SAMN20352633 | SRR15219201 | NonPC | Public | 2013 | China | Env | 87.8 | 100 | 0.36 | 98.2742 | 5070722 | 45.21 | 49 | 554090 |
| SPVP130080 | SPVP130080 | SAMN20352639 | SRR15219195 | NonPC | Public | 2013 | China | Env | 88.5 | 100 | 0.3 | 98.2024 | 5105452 | 45.18 | 66 | 524441 |
| SPVP130081 | SPVP130081 | SAMN20352640 | SRR15219192 | NonPC | Public | 2013 | China | Env | 88.7 | 100 | 0.46 | 98.1784 | 5094287 | 45.24 | 46 | 563061 |
| SPVP130082 | SPVP130082 | SAMN20352582 | SRR15219432 | NonPC | Public | 2013 | China | Env | 91.4 | 100 | 0.05 | 98.402 | 5149496 | 45.2 | 34 | 448653 |
| SPVP130083 | SPVP130083 | SAMN20352574 | SRR15219441 | NonPC | Public | 2013 | China | Env | 90.8 | 100 | 0.03 | 98.3717 | 4970174 | 45.34 | 50 | 314776 |
| SPVP130084 | SPVP130084 | SAMN20352580 | SRR15219434 | NonPC | Public | 2013 | China | Env | 88.2 | 100 | 0.3 | 98.2675 | 5006841 | 45.28 | 64 | 332038 |
| SPVP130085 | SPVP130085 | SAMN20352575 | SRR15219440 | NonPC | Public | 2013 | China | Env | 91.6 | 100 | 0.09 | 98.3796 | 5127044 | 45.29 | 67 | 409511 |
| SPVP130086 | SPVP130086 | SAMN20352617 | SRR15219325 | NonPC | Public | 2013 | China | Env | 90.8 | 100 | 0 | 98.3914 | 5073735 | 45.34 | 35 | 438873 |
| SPVP130087 | SPVP130087 | SAMN20352618 | SRR15219324 | NonPC | Public | 2013 | China | Env | 91.5 | 100 | 0.03 | 98.3549 | 5042724 | 45.3 | 49 | 449102 |
| SPVP130088 | SPVP130088 | SAMN20352619 | SRR15219323 | NonPC | Public | 2013 | China | Env | 91.4 | 100 | 0 | 98.3364 | 5169577 | 45.21 | 67 | 370645 |
| SPVP130089 | SPVP130089 | SAMN20352616 | SRR15219326 | NonPC | Public | 2013 | China | Env | 88.4 | 100 | 0.36 | 98.2942 | 5010360 | 45.35 | 41 | 451557 |
| SPVP130090 | SPVP130090 | SAMN20352642 | SRR15219190 | NonPC | Public | 2013 | China | Env | 91.1 | 100 | 0.03 | 98.294 | 5398212 | 45.04 | 51 | 401410 |
| SPVP130091 | SPVP130091 | SAMN20352643 | SRR15219189 | NonPC | Public | 2013 | China | Env | 89.5 | 100 | 0.08 | 98.2686 | 5034311 | 45.36 | 40 | 452665 |
| SPVP130092 | SPVP130092 | SAMN20352644 | SRR15219188 | NonPC | Public | 2013 | China | Env | 89.6 | 100 | 0.4 | 98.3179 | 5158466 | 45.26 | 42 | 450572 |
| SPVP130093 | SPVP130093 | SAMN20352641 | SRR15219191 | NonPC | Public | 2013 | China | Env | 90.6 | 100 | 0 | 98.3736 | 4959501 | 45.42 | 46 | 265905 |
| SPVP130094 | SPVP130094 | SAMN20352658 | SRR15219173 | NonPC | Public | 2013 | China | Env | 88.9 | 100 | 0.39 | 98.2703 | 4947329 | 45.36 | 61 | 565723 |
| SPVP130095 | SPVP130095 | SAMN20352659 | SRR15219172 | NonPC | Public | 2013 | China | Env | 88.8 | 100 | 0.3 | 98.3333 | 5074207 | 45.36 | 37 | 447732 |
| SPVP130096 | SPVP130096 | SAMN20352667 | SRR15219116 | NonPC | Public | 2013 | China | Env | 88.3 | 100 | 0.39 | 98.2048 | 5185236 | 45.2 | 59 | 361593 |
| SPVP130097 | SPVP130097 | SAMN20352660 | SRR15219123 | NonPC | Public | 2013 | China | Env | 91.3 | 100 | 0 | 98.3615 | 4946600 | 45.4 | 30 | 535010 |
| SPVP130098 | SPVP130098 | SAMN20352661 | SRR15219122 | NonPC | Public | 2013 | China | Env | 89.4 | 100 | 0.34 | 98.2827 | 5156252 | 45.24 | 57 | 399664 |

| | | | | | | | | | | | | | | | | |
|---|---|---|---|---|---|---|---|---|---|---|---|---|---|---|---|---|
| SPVP130099 | SPVP130099 | SAMN20352662 | SRR15219121 | NonPC | Public | 2013 | China | Env | 91.5 | 100 | 0.54 | 98.2838 | 5280723 | 45.26 | 83 | 280056 |
| SPVP130102 | SPVP130102 | SAMN20352656 | SRR15219175 | NonPC | Public | 2013 | China | Env | 88.1 | 100 | 0.33 | 98.3055 | 4980054 | 45.31 | 51 | 550952 |
| SPVP130103 | SPVP130103 | SAMN20352663 | SRR15219120 | NonPC | Public | 2013 | China | Env | 88.6 | 100 | 0.36 | 98.1846 | 5201738 | 45.11 | 62 | 465341 |
| SPVP130104 | SPVP130104 | SAMN20352668 | SRR15219115 | NonPC | Public | 2013 | China | Env | 89.8 | 100 | 0.27 | 98.3826 | 4964710 | 45.33 | 33 | 511793 |
| SPVP130105 | SPVP130105 | SAMN20352669 | SRR15219114 | NonPC | Public | 2013 | China | Env | 89.8 | 100 | 0.27 | 98.3357 | 4963193 | 45.33 | 30 | 567015 |
| SPVP130106 | SPVP130106 | SAMN20352664 | SRR15219119 | NonPC | Public | 2013 | China | Env | 89.8 | 100 | 0 | 98.3294 | 4998917 | 45.28 | 43 | 445714 |
| SPVP130107 | SPVP130107 | SAMN20352657 | SRR15219174 | NonPC | Public | 2013 | China | Env | 90 | 100 | 0.03 | 98.2387 | 5151003 | 45.27 | 78 | 388578 |
| SPVP130108 | SPVP130108 | SAMN20352665 | SRR15219118 | NonPC | Public | 2013 | China | Env | 89.5 | 100 | 0 | 98.2243 | 4956748 | 45.42 | 50 | 448356 |
| SPVP130109 | SPVP130109 | SAMN20352666 | SRR15219117 | NonPC | Public | 2013 | China | Env | 88.6 | 100 | 0.63 | 98.207 | 5139919 | 45.17 | 80 | 544208 |
| SPVP130111 | SPVP130111 | SAMN20352670 | SRR15218775 | NonPC | Public | 2013 | China | Env | 88.3 | 100 | 0.36 | 98.3055 | 4986060 | 45.27 | 68 | 463649 |
| SPVP130112 | SPVP130112 | SAMN20352672 | SRR15218773 | NonPC | Public | 2013 | China | Env | 88.8 | 100 | 0.36 | 98.2826 | 5121409 | 45.19 | 35 | 459569 |
| SPVP130113 | SPVP130113 | SAMN20352673 | SRR15218772 | NonPC | Public | 2013 | China | Env | 88.2 | 100 | 0.39 | 98.1977 | 5085574 | 45.27 | 51 | 542134 |
| SPVP130114 | SPVP130114 | SAMN20352675 | SRR15218770 | NonPC | Public | 2013 | China | Env | 91.2 | 100 | 0.07 | 98.3519 | 5019142 | 45.35 | 50 | 461886 |
| SPVP130115 | SPVP130115 | SAMN20352676 | SRR15218769 | NonPC | Public | 2013 | China | Env | 91.2 | 100 | 0.07 | 98.3556 | 5016806 | 45.35 | 51 | 461886 |
| SPVP130116 | SPVP130116 | SAMN20352674 | SRR15218771 | NonPC | Public | 2013 | China | Env | 88.2 | 100 | 0.39 | 98.1874 | 5174319 | 45.19 | 57 | 542137 |
| SPVP130117 | SPVP130117 | SAMN20352677 | SRR15218768 | NonPC | Public | 2013 | China | Env | 91.2 | 100 | 0.07 | 98.3085 | 5017316 | 45.35 | 52 | 461886 |
| SPVP130118 | SPVP130118 | SAMN20352678 | SRR15218767 | NonPC | Public | 2013 | China | Env | 91.2 | 100 | 0.07 | 98.3311 | 5016865 | 45.35 | 51 | 461886 |
| SPVP130119 | SPVP130119 | SAMN20352671 | SRR15218774 | NonPC | Public | 2013 | China | Env | 88.1 | 100 | 0.37 | 98.2886 | 5043701 | 45.3 | 34 | 452044 |
| SPVP130120 | SPVP130120 | SAMN20352648 | SRR15219184 | NonPC | Public | 2013 | China | Env | 88.2 | 100 | 0.4 | 98.2525 | 5135019 | 45.13 | 100 | 451011 |
| SPVP130121 | SPVP130121 | SAMN20352649 | SRR15219183 | NonPC | Public | 2013 | China | Env | 88.2 | 100 | 0.4 | 98.2658 | 5135139 | 45.13 | 97 | 544763 |
| SPVP130122 | SPVP130122 | SAMN20352650 | SRR15219181 | NonPC | Public | 2013 | China | Env | 88.2 | 100 | 0.4 | 98.2632 | 5134947 | 45.13 | 96 | 724236 |
| SPVP130123 | SPVP130123 | SAMN20352651 | SRR15219180 | NonPC | Public | 2013 | China | Env | 88.2 | 100 | 0.4 | 98.2236 | 5134439 | 45.13 | 98 | 451011 |
| SPVP130124 | SPVP130124 | SAMN20352652 | SRR15219179 | NonPC | Public | 2013 | China | Env | 88.2 | 100 | 0.4 | 98.2489 | 5132754 | 45.13 | 97 | 544763 |
| SPVP130125 | SPVP130125 | SAMN20352653 | SRR15219178 | NonPC | Public | 2013 | China | Env | 88.3 | 100 | 0.4 | 98.2638 | 5135702 | 45.13 | 99 | 451011 |
| SPVP130126 | SPVP130126 | SAMN20352680 | SRR15218764 | NonPC | Public | 2013 | China | Env | 91.3 | 100 | 0 | 98.399 | 4964502 | 45.41 | 39 | 449783 |
| SPVP130127 | SPVP130127 | SAMN20352681 | SRR15218763 | NonPC | Public | 2013 | China | Env | 91.2 | 100 | 0 | 98.3844 | 4982796 | 45.38 | 47 | 456637 |
| SPVP130128 | SPVP130128 | SAMN20352682 | SRR15218762 | NonPC | Public | 2013 | China | Env | 91.2 | 100 | 0 | 98.3812 | 4982991 | 45.38 | 49 | 456637 |
| SPVP130129 | SPVP130129 | SAMN20352679 | SRR15218766 | NonPC | Public | 2013 | China | Env | 87.9 | 100 | 0.34 | 98.2477 | 5074366 | 45.32 | 49 | 561401 |
| SPVP140001 | SPVP140001 | SAMN20352718 | SRR15218691 | NonPC | Public | 2014 | China | Env | 89.9 | 100 | 0 | 98.3273 | 4964181 | 45.37 | 56 | 389284 |
| SPVP140002 | SPVP140002 | SAMN20352719 | SRR15218690 | NonPC | Public | 2014 | China | Env | 91.4 | 100 | 0.38 | 98.3634 | 5017012 | 45.34 | 51 | 460502 |
| SPVP140003 | SPVP140003 | SAMN20352720 | SRR15218587 | NonPC | Public | 2014 | China | Env | 91.1 | 100 | 0 | 98.3057 | 4982666 | 45.32 | 51 | 332061 |
| SPVP140004 | SPVP140004 | SAMN20352721 | SRR15218586 | NonPC | Public | 2014 | China | Env | 90 | 100 | 0.4 | 98.3296 | 4984774 | 45.35 | 41 | 480891 |
| SPVP140005 | SPVP140005 | SAMN20352724 | SRR15218583 | NonPC | Public | 2014 | China | Env | 89.7 | 100 | 0.09 | 98.3782 | 5075919 | 45.34 | 56 | 528589 |
| SPVP140006 | SPVP140006 | SAMN20352722 | SRR15218585 | NonPC | Public | 2014 | China | Env | 91.6 | 100 | 0.14 | 98.4029 | 5190816 | 45.16 | 71 | 182443 |
| SPVP140007 | SPVP140007 | SAMN20352725 | SRR15218582 | NonPC | Public | 2014 | China | Env | 90.7 | 100 | 0.05 | 98.2236 | 5237520 | 45.17 | 63 | 435554 |
| SPVP140008 | SPVP140008 | SAMN20352723 | SRR15218584 | NonPC | Public | 2014 | China | Env | 91.7 | 100 | 0.05 | 98.3042 | 5033421 | 45.3 | 80 | 243312 |
| SPVP140009 | SPVP140009 | SAMN20352726 | SRR15218581 | NonPC | Public | 2014 | China | Env | 88.5 | 100 | 0.36 | 98.2432 | 5008684 | 45.3 | 55 | 732289 |
| SPVP140010 | SPVP140010 | SAMN20352727 | SRR15218580 | NonPC | Public | 2014 | China | Env | 91.1 | 100 | 0.31 | 98.2907 | 5476977 | 44.95 | 58 | 403527 |
| SPVP140011 | SPVP140011 | SAMN20352728 | SRR15218579 | NonPC | Public | 2014 | China | Env | 90.1 | 100 | 0.05 | 98.3126 | 5086307 | 45.32 | 46 | 530018 |
| SPVP140013 | SPVP140013 | SAMN20352733 | SRR15218573 | NonPC | Public | 2014 | China | Env | 91.6 | 100 | 0.09 | 98.3292 | 5097378 | 45.22 | 54 | 460723 |
| SPVP140021 | SPVP140021 | SAMN20352783 | SRR15219071 | NonPC | Public | 2014 | China | Env | 88.2 | 100 | 0.39 | 98.2095 | 5066012 | 45.22 | 54 | 542138 |
| SPVP140022 | SPVP140022 | SAMN20352751 | SRR15218552 | NonPC | Public | 2014 | China | Env | 88.7 | 100 | 0.61 | 98.2049 | 5082814 | 45.21 | 42 | 452085 |
| SPVP140023 | SPVP140023 | SAMN20352752 | SRR15218551 | NonPC | Public | 2014 | China | Env | 90 | 100 | 0.12 | 98.3272 | 5084346 | 45.33 | 72 | 443132 |
| SPVP140024 | SPVP140024 | SAMN20352784 | SRR15219070 | NonPC | Public | 2014 | China | Env | 88.3 | 100 | 0.39 | 98.2101 | 5067026 | 45.22 | 53 | 542137 |
| SPVP140025 | SPVP140025 | SAMN20352753 | SRR15218550 | NonPC | Public | 2014 | China | Env | 90.4 | 100 | 0.19 | 98.2888 | 5044966 | 45.32 | 37 | 604689 |
| SPVP140026 | SPVP140026 | SAMN20352754 | SRR15218549 | NonPC | Public | 2014 | China | Env | 90.1 | 100 | 0.07 | 98.3589 | 5000858 | 45.36 | 85 | 521735 |
| SPVP140027 | SPVP140027 | SAMN20352755 | SRR15218548 | NonPC | Public | 2014 | China | Env | 91.1 | 100 | 0.1 | 98.3499 | 4968382 | 45.42 | 43 | 465465 |
| SPVP140028 | SPVP140028 | SAMN20352785 | SRR15219069 | NonPC | Public | 2014 | China | Env | 88.2 | 100 | 0.67 | 98.2149 | 5178590 | 45.26 | 53 | 542137 |

| | | | | | | | | | | | | | | | | |
|---|---|---|---|---|---|---|---|---|---|---|---|---|---|---|---|---|
| SPVP140029 | SPVP140029 | SAMN20352756 | SRR15218547 | NonPC | Public | 2014 | China | Env | 87.5 | 100 | 0.37 | 98.2704 | 5087567 | 45.3 | 84 | 462225 |
| SPVP140030 | SPVP140030 | SAMN20352757 | SRR15218546 | NonPC | Public | 2014 | China | Env | 91.8 | 100 | 0.2 | 98.3045 | 5070394 | 45.34 | 75 | 338391 |
| SPVP140031 | SPVP140031 | SAMN20352758 | SRR15218545 | NonPC | Public | 2014 | China | Env | 91.8 | 100 | 0.31 | 98.344 | 5191770 | 45.26 | 50 | 317906 |
| SPVP140032 | SPVP140032 | SAMN20352759 | SRR15218544 | NonPC | Public | 2014 | China | Env | 90 | 100 | 0.06 | 98.2945 | 5169016 | 45.33 | 41 | 451916 |
| SPVP140033 | SPVP140033 | SAMN20352745 | SRR15218559 | NonPC | Public | 2014 | China | Env | 89.4 | 100 | 0 | 98.3745 | 5052719 | 45.29 | 32 | 764404 |
| SPVP140034 | SPVP140034 | SAMN20352739 | SRR15218567 | NonPC | Public | 2014 | China | Env | 91.9 | 100 | 0.14 | 98.4048 | 5224640 | 45.14 | 44 | 457509 |
| SPVP140035 | SPVP140035 | SAMN20352749 | SRR15218555 | NonPC | Public | 2014 | China | Env | 88.6 | 100 | 0.42 | 98.279 | 5069832 | 45.32 | 42 | 452829 |
| SPVP140036 | SPVP140036 | SAMN20352744 | SRR15218560 | NonPC | Public | 2014 | China | Env | 91 | 100 | 0.03 | 98.2854 | 5149175 | 45.22 | 89 | 264078 |
| SPVP140037 | SPVP140037 | SAMN20352747 | SRR15218557 | NonPC | Public | 2014 | China | Env | 92.1 | 100 | 0 | 98.43 | 5107450 | 45.28 | 37 | 450530 |
| SPVP140038 | SPVP140038 | SAMN20352760 | SRR15218542 | NonPC | Public | 2014 | China | Env | 89.3 | 100 | 0 | 98.229 | 5019443 | 45.33 | 68 | 408641 |
| SPVP140040 | SPVP140040 | SAMN20352761 | SRR15218541 | NonPC | Public | 2014 | China | Env | 89.6 | 100 | 0.5 | 98.2791 | 5010513 | 45.38 | 71 | 522561 |
| SPVP140042 | SPVP140042 | SAMN20352746 | SRR15218558 | NonPC | Public | 2014 | China | Env | 89.5 | 100 | 0.54 | 98.2288 | 5291135 | 45.38 | 64 | 448798 |
| SPVP140043 | SPVP140043 | SAMN20352762 | SRR15218540 | NonPC | Public | 2014 | China | Env | 90.5 | 100 | 0.22 | 98.263 | 5024425 | 45.32 | 65 | 459777 |
| SPVP140044 | SPVP140044 | SAMN20352763 | SRR15218539 | NonPC | Public | 2014 | China | Env | 90.9 | 100 | 0 | 98.353 | 5070583 | 45.3 | 77 | 300497 |
| SPVP140045 | SPVP140045 | SAMN20352780 | SRR15219074 | NonPC | Public | 2014 | China | Env | 88.9 | 100 | 0.39 | 98.1902 | 5125305 | 45.23 | 45 | 782740 |
| SPVP140046 | SPVP140046 | SAMN20352764 | SRR15218538 | NonPC | Public | 2014 | China | Env | 88.5 | 100 | 0.49 | 98.2196 | 5051208 | 45.24 | 78 | 429841 |
| SPVP140047 | SPVP140047 | SAMN20352765 | SRR15218537 | NonPC | Public | 2014 | China | Env | 91.7 | 100 | 0.27 | 98.3993 | 5401490 | 45.1 | 57 | 357217 |
| SPVP140048 | SPVP140048 | SAMN20352743 | SRR15218561 | NonPC | Public | 2014 | China | Env | 88.9 | 100 | 0.39 | 98.2053 | 5234702 | 45.02 | 48 | 462825 |
| SPVP140049 | SPVP140049 | SAMN20352766 | SRR15218536 | NonPC | Public | 2014 | China | Env | 88.7 | 100 | 0.79 | 98.1614 | 5552803 | 45.04 | 76 | 771313 |
| SPVP140050 | SPVP140050 | SAMN20352750 | SRR15218553 | NonPC | Public | 2014 | China | Env | 88.4 | 100 | 0.36 | 98.2012 | 5115372 | 45.27 | 74 | 456709 |
| SPVP140051 | SPVP140051 | SAMN20352767 | SRR15218535 | NonPC | Public | 2014 | China | Env | 88.5 | 100 | 0.43 | 98.2403 | 5056841 | 45.27 | 83 | 550077 |
| SPVP140052 | SPVP140052 | SAMN20352786 | SRR15219068 | NonPC | Public | 2014 | China | Env | 88.2 | 100 | 0.39 | 98.2049 | 5058309 | 45.23 | 53 | 661387 |
| SPVP140053 | SPVP140053 | SAMN20352787 | SRR15219067 | NonPC | Public | 2014 | China | Env | 88.3 | 100 | 0.39 | 98.2425 | 5066573 | 45.22 | 55 | 542137 |
| SPVP140054 | SPVP140054 | SAMN20352788 | SRR15219066 | NonPC | Public | 2014 | China | Env | 88.3 | 100 | 0.39 | 98.2431 | 5066480 | 45.22 | 53 | 542137 |
| SPVP140056 | SPVP140056 | SAMN20352748 | SRR15218556 | NonPC | Public | 2014 | China | Env | 90 | 100 | 0 | 98.3517 | 5071046 | 45.31 | 32 | 446840 |
| SPVP140057 | SPVP140057 | SAMN20352768 | SRR15218534 | NonPC | Public | 2014 | China | Env | 91.1 | 100 | 0.43 | 98.3833 | 4984190 | 45.38 | 34 | 461969 |
| SPVP140058 | SPVP140058 | SAMN20352789 | SRR15219065 | NonPC | Public | 2014 | China | Env | 89.9 | 100 | 0.05 | 98.3289 | 5008076 | 45.34 | 51 | 306335 |
| SPVP140059 | SPVP140059 | SAMN20352769 | SRR15218533 | NonPC | Public | 2014 | China | Env | 90.2 | 100 | 0 | 98.351 | 5048818 | 45.36 | 71 | 453358 |
| SPVP140060 | SPVP140060 | SAMN20352781 | SRR15219073 | NonPC | Public | 2014 | China | Env | 88.5 | 100 | 0.36 | 98.2833 | 5095893 | 45.26 | 49 | 451557 |
| SPVP140061 | SPVP140061 | SAMN20352770 | SRR15218531 | NonPC | Public | 2014 | China | Env | 91.4 | 100 | 0 | 98.2588 | 5109249 | 45.27 | 50 | 465683 |
| SPVP140062 | SPVP140062 | SAMN20352741 | SRR15218563 | NonPC | Public | 2014 | China | Env | 88.1 | 100 | 0.66 | 98.273 | 4942806 | 45.33 | 55 | 444011 |
| SPVP140063 | SPVP140063 | SAMN20352771 | SRR15218530 | NonPC | Public | 2014 | China | Env | 91.8 | 100 | 0.08 | 98.284 | 5444334 | 45.1 | 58 | 416793 |
| SPVP140064 | SPVP140064 | SAMN20352742 | SRR15218562 | NonPC | Public | 2014 | China | Env | 88.3 | 100 | 0.33 | 98.2076 | 5176462 | 45.14 | 75 | 453133 |
| SPVP140065 | SPVP140065 | SAMN20352740 | SRR15218564 | NonPC | Public | 2014 | China | Env | 88.4 | 100 | 0.36 | 98.2441 | 4997318 | 45.32 | 80 | 384834 |
| SPVP140066 | SPVP140066 | SAMN20352772 | SRR15218529 | NonPC | Public | 2014 | China | Env | 91.3 | 100 | 0.27 | 98.3253 | 5135639 | 45.3 | 41 | 403454 |
| SPVP140067 | SPVP140067 | SAMN20352773 | SRR15218528 | NonPC | Public | 2014 | China | Env | 88.2 | 100 | 0.39 | 98.2402 | 5151798 | 45.19 | 61 | 328640 |
| SPVP140068 | SPVP140068 | SAMN20352774 | SRR15218527 | NonPC | Public | 2014 | China | Env | 92.4 | 100 | 0.59 | 98.3733 | 5242367 | 45.22 | 64 | 375140 |
| SPVP140069 | SPVP140069 | SAMN20352782 | SRR15219072 | NonPC | Public | 2014 | China | Env | 88.5 | 100 | 0.36 | 98.2358 | 5096628 | 45.26 | 47 | 451557 |
| SPVP140070 | SPVP140070 | SAMN20352790 | SRR15219063 | NonPC | Public | 2014 | China | Env | 89.1 | 100 | 0.05 | 98.3599 | 5007635 | 45.34 | 51 | 306335 |
| SPVP140071 | SPVP140071 | SAMN20352775 | SRR15218526 | NonPC | Public | 2014 | China | Env | 89.3 | 100 | 0 | 98.3112 | 5025437 | 45.27 | 45 | 324899 |
| SPVP140072 | SPVP140072 | SAMN20352738 | SRR15218568 | NonPC | Public | 2014 | China | Env | 90.3 | 100 | 0.19 | 98.3152 | 5090093 | 45.14 | 62 | 220744 |
| SPVP140073 | SPVP140073 | SAMN20352776 | SRR15218525 | NonPC | Public | 2014 | China | Env | 88.2 | 100 | 0.3 | 98.235 | 5153285 | 45.14 | 69 | 395614 |
| SPVP140074 | SPVP140074 | SAMN20352777 | SRR15218523 | NonPC | Public | 2014 | China | Env | 87.3 | 100 | 0.53 | 98.2131 | 5133927 | 45.11 | 33 | 454400 |
| SPVP140075 | SPVP140075 | SAMN20352778 | SRR15219077 | NonPC | Public | 2014 | China | Env | 88.4 | 100 | 0.41 | 98.2553 | 5038412 | 45.27 | 67 | 351192 |
| SPVP140076 | SPVP140076 | SAMN20352779 | SRR15219076 | NonPC | Public | 2014 | China | Env | 88.8 | 100 | 0.39 | 98.2587 | 5062706 | 45.26 | 67 | 544136 |
| SPVP140078 | SPVP140078 | SAMN20352794 | SRR15219059 | NonPC | Public | 2014 | China | Env | 90.6 | 100 | 0 | 98.3593 | 4999208 | 45.45 | 43 | 413549 |
| SPVP140080 | SPVP140080 | SAMN20352802 | SRR15219050 | NonPC | Public | 2014 | China | Env | 89.4 | 100 | 0.23 | 98.319 | 4958424 | 45.42 | 47 | 460729 |
| SPVP140081 | SPVP140081 | SAMN20352798 | SRR15219055 | NonPC | Public | 2014 | China | Env | 88.4 | 100 | 0.39 | 98.2096 | 5085620 | 45.25 | 74 | 612195 |

| | | | | | | | | | | | | | | | | |
|---|---|---|---|---|---|---|---|---|---|---|---|---|---|---|---|---|
| SPVP140082 | SPVP140082 | SAMN20352796 | SRR15219057 | NonPC | Public | 2014 | China | Env | 88.6 | 100 | 0.58 | 98.1479 | 5136063 | 45.22 | 65 | 398080 |
| SPVP140083 | SPVP140083 | SAMN20352797 | SRR15219056 | NonPC | Public | 2014 | China | Env | 91.9 | 100 | 0.34 | 98.3848 | 5047687 | 45.39 | 56 | 403236 |
| SPVP140084 | SPVP140084 | SAMN20352734 | SRR15218572 | NonPC | Public | 2014 | China | Env | 90.7 | 100 | 0.05 | 98.293 | 5177870 | 45.24 | 80 | 410818 |
| SPVP140085 | SPVP140085 | SAMN20352732 | SRR15218574 | NonPC | Public | 2014 | China | Env | 88.3 | 100 | 0.3 | 98.2266 | 5260968 | 45.11 | 58 | 421526 |
| SPVP140087 | SPVP140087 | SAMN20352731 | SRR15218575 | NonPC | Public | 2014 | China | Env | 89.8 | 100 | 0.12 | 98.3436 | 5016769 | 45.35 | 29 | 488830 |
| SPVP140089 | SPVP140089 | SAMN20352735 | SRR15218571 | NonPC | Public | 2014 | China | Env | 89.6 | 100 | 0.57 | 98.2716 | 4972163 | 45.43 | 43 | 548935 |
| SPVP140090 | SPVP140090 | SAMN20352736 | SRR15218570 | NonPC | Public | 2014 | China | Env | 90.2 | 100 | 0.05 | 98.3231 | 5011170 | 45.36 | 52 | 451101 |
| SPVP140091 | SPVP140091 | SAMN20352795 | SRR15219058 | NonPC | Public | 2014 | China | Env | 91.3 | 100 | 0.27 | 98.3741 | 5002864 | 45.44 | 60 | 314161 |
| SPVP140092 | SPVP140092 | SAMN20352800 | SRR15219052 | NonPC | Public | 2014 | China | Env | 91.8 | 100 | 0 | 98.2985 | 5205204 | 45.22 | 72 | 240684 |
| SPVP140093 | SPVP140093 | SAMN20352801 | SRR15219051 | NonPC | Public | 2014 | China | Env | 89.2 | 100 | 0.68 | 98.3731 | 5012506 | 45.3 | 78 | 145330 |
| SPVP140094 | SPVP140094 | SAMN20352737 | SRR15218569 | NonPC | Public | 2014 | China | Env | 89.2 | 100 | 0.68 | 98.3781 | 5013411 | 45.3 | 78 | 143005 |
| SPVP140095 | SPVP140095 | SAMN20352799 | SRR15219054 | NonPC | Public | 2014 | China | Env | 90.1 | 100 | 0.05 | 98.2962 | 5144389 | 45.23 | 57 | 462160 |
| SPVP140096 | SPVP140096 | SAMN20352829 | SRR15218952 | NonPC | Public | 2014 | China | Env | 87.6 | 100 | 0.45 | 98.2234 | 5154772 | 45.14 | 94 | 450998 |
| SPVP140097 | SPVP140097 | SAMN20352834 | SRR15218946 | NonPC | Public | 2014 | China | Env | 90.3 | 100 | 0 | 98.3397 | 5036533 | 45.29 | 67 | 278978 |
| SPVP140098 | SPVP140098 | SAMN20352835 | SRR15218945 | NonPC | Public | 2014 | China | Env | 91.2 | 100 | 0 | 98.3918 | 5279702 | 45.24 | 69 | 202472 |
| SPVP140099 | SPVP140099 | SAMN20352825 | SRR15218956 | NonPC | Public | 2014 | China | Env | 88.6 | 100 | 0.61 | 98.1463 | 5111334 | 45.23 | 67 | 487834 |
| SPVP140100 | SPVP140100 | SAMN20352832 | SRR15218948 | NonPC | Public | 2014 | China | Env | 91.7 | 100 | 0 | 98.3447 | 5118285 | 45.34 | 58 | 373752 |
| SPVP140104 | SPVP140104 | SAMN20352833 | SRR15218947 | NonPC | Public | 2014 | China | Env | 88.1 | 100 | 0.37 | 98.268 | 5065486 | 45.33 | 44 | 366836 |
| SPVP140105 | SPVP140105 | SAMN20352858 | SRR15218813 | NonPC | Public | 2014 | China | Env | 88.4 | 100 | 0.4 | 98.217 | 5148954 | 45.22 | 47 | 583843 |
| SPVP140106 | SPVP140106 | SAMN20352859 | SRR15218812 | NonPC | Public | 2014 | China | Env | 91.5 | 100 | 0.27 | 98.3664 | 5118878 | 45.43 | 56 | 313375 |
| SPVP140107 | SPVP140107 | SAMN20352860 | SRR15218810 | NonPC | Public | 2014 | China | Env | 90.1 | 100 | 0 | 98.2872 | 5153673 | 45.34 | 56 | 528680 |
| SPVP140108 | SPVP140108 | SAMN20352861 | SRR15218809 | NonPC | Public | 2014 | China | Env | 88.5 | 100 | 0.36 | 98.2122 | 5192336 | 45.16 | 73 | 497604 |
| SPVP140109 | SPVP140109 | SAMN20352879 | SRR15218790 | NonPC | Public | 2014 | China | Env | 89.8 | 100 | 0.53 | 98.2662 | 5440856 | 45.28 | 45 | 745606 |
| SPVP140110 | SPVP140110 | SAMN20352855 | SRR15218816 | NonPC | Public | 2014 | China | Env | 91.2 | 100 | 0.27 | 98.375 | 5007275 | 45.39 | 41 | 858598 |
| SPVP140112 | SPVP140112 | SAMN20352881 | SRR15218787 | PC | Public | 2014 | China | Env | 98.1 | 100 | 0 | 99.9816 | 5072205 | 45.3 | 47 | 404135 |
| SPVP140113 | SPVP140113 | SAMN20352880 | SRR15218788 | NonPC | Public | 2014 | China | Env | 89.8 | 100 | 0.53 | 98.3192 | 5441129 | 45.28 | 47 | 546357 |
| SPVP140114 | SPVP140114 | SAMN20352853 | SRR15218818 | NonPC | Public | 2014 | China | Env | 87.8 | 100 | 0.45 | 98.2167 | 5240640 | 45.29 | 47 | 383629 |
| SPVP140115 | SPVP140115 | SAMN20352846 | SRR15218826 | NonPC | Public | 2014 | China | Env | 90.7 | 100 | 0 | 98.3586 | 4970074 | 45.43 | 47 | 314351 |
| SPVP140116 | SPVP140116 | SAMN20352862 | SRR15218808 | NonPC | Public | 2014 | China | Env | 91.7 | 100 | 0.14 | 98.3493 | 5099741 | 45.33 | 78 | 402085 |
| SPVP140117 | SPVP140117 | SAMN20352848 | SRR15218824 | NonPC | Public | 2014 | China | Env | 88.5 | 100 | 0.33 | 98.2323 | 5043176 | 45.31 | 47 | 835142 |
| SPVP140118 | SPVP140118 | SAMN20352847 | SRR15218825 | NonPC | Public | 2014 | China | Env | 88.3 | 100 | 0.42 | 98.2404 | 5100497 | 45.24 | 51 | 427725 |
| SPVP140119 | SPVP140119 | SAMN20352845 | SRR15218827 | NonPC | Public | 2014 | China | Env | 90.9 | 100 | 0.64 | 98.361 | 5095324 | 45.29 | 37 | 452888 |
| SPVP140120 | SPVP140120 | SAMN20352856 | SRR15218815 | NonPC | Public | 2014 | China | Env | 90.1 | 100 | 0.09 | 98.3791 | 5047597 | 45.27 | 36 | 713549 |
| SPVP140121 | SPVP140121 | SAMN20352863 | SRR15218807 | NonPC | Public | 2014 | China | Env | 91.4 | 100 | 0.03 | 98.4152 | 5057022 | 45.26 | 68 | 461816 |
| SPVP140122 | SPVP140122 | SAMN20352851 | SRR15218820 | NonPC | Public | 2014 | China | Env | 88.9 | 100 | 0.39 | 98.2294 | 5239234 | 45.03 | 43 | 568938 |
| SPVP140123 | SPVP140123 | SAMN20352864 | SRR15218806 | NonPC | Public | 2014 | China | Env | 88.3 | 100 | 0.3 | 98.208 | 5000992 | 45.32 | 59 | 452021 |
| SPVP140125 | SPVP140125 | SAMN20352877 | SRR15218792 | NonPC | Public | 2014 | China | Env | 88.2 | 100 | 0.39 | 98.208 | 5077545 | 45.19 | 53 | 542137 |
| SPVP140126 | SPVP140126 | SAMN20352865 | SRR15218805 | NonPC | Public | 2014 | China | Env | 89.5 | 100 | 0.07 | 98.3379 | 5001800 | 45.41 | 34 | 527073 |
| SPVP140127 | SPVP140127 | SAMN20352866 | SRR15218804 | NonPC | Public | 2014 | China | Env | 88.1 | 100 | 0.39 | 98.2147 | 5176261 | 45.23 | 62 | 262307 |
| SPVP140128 | SPVP140128 | SAMN20352850 | SRR15218821 | NonPC | Public | 2014 | China | Env | 88.3 | 100 | 0.44 | 98.3168 | 5036621 | 45.21 | 49 | 452369 |
| SPVP140129 | SPVP140129 | SAMN20352854 | SRR15218817 | NonPC | Public | 2014 | China | Env | 91.6 | 100 | 0.34 | 98.37 | 5010630 | 45.41 | 41 | 405080 |
| SPVP140130 | SPVP140130 | SAMN20352878 | SRR15218791 | NonPC | Public | 2014 | China | Env | 91.4 | 100 | 0.05 | 98.378 | 5151069 | 45.01 | 30 | 448653 |
| SPVP140131 | SPVP140131 | SAMN20352867 | SRR15218803 | NonPC | Public | 2014 | China | Env | 92.1 | 100 | 0 | 98.3385 | 5085428 | 45.33 | 66 | 298445 |
| SPVP140133 | SPVP140133 | SAMN20352868 | SRR15218802 | NonPC | Public | 2014 | China | Env | 89.7 | 100 | 0.34 | 98.2727 | 5138062 | 45.23 | 34 | 480801 |
| SPVP140134 | SPVP140134 | SAMN20352843 | SRR15218829 | NonPC | Public | 2014 | China | Env | 90.6 | 100 | 0.1 | 98.3408 | 5331522 | 45.37 | 59 | 260312 |
| SPVP140135 | SPVP140135 | SAMN20352836 | SRR15218944 | NonPC | Public | 2014 | China | Env | 90.3 | 100 | 0.27 | 98.2975 | 5025236 | 45.45 | 36 | 517253 |
| SPVP140136 | SPVP140136 | SAMN20352844 | SRR15218828 | NonPC | Public | 2014 | China | Env | 90.6 | 100 | 0.1 | 98.3395 | 5328873 | 45.38 | 55 | 291156 |
| SPVP140137 | SPVP140137 | SAMN20352837 | SRR15218943 | NonPC | Public | 2014 | China | Env | 89.6 | 100 | 0.09 | 98.3125 | 5112942 | 45.22 | 42 | 363770 |

| | | | | | | | | | | | | | | | | |
|---|---|---|---|---|---|---|---|---|---|---|---|---|---|---|---|---|
| SPVP140138 | SPVP140138 | SAMN20352823 | SRR15218958 | NonPC | Public | 2014 | China | Env | 90.9 | 100 | 0.24 | 98.3536 | 5236750 | 45.09 | 47 | 426924 |
| SPVP140139 | SPVP140139 | SAMN20352838 | SRR15218942 | NonPC | Public | 2014 | China | Env | 92 | 100 | 0.27 | 98.4204 | 4986658 | 45.42 | 40 | 403381 |
| SPVP140140 | SPVP140140 | SAMN20352826 | SRR15218955 | NonPC | Public | 2014 | China | Env | 88.5 | 100 | 0.6 | 98.2087 | 5124262 | 45.19 | 67 | 452977 |
| SPVP140142 | SPVP140142 | SAMN20352830 | SRR15218950 | NonPC | Public | 2014 | China | Env | 91.4 | 100 | 0.36 | 98.3348 | 5033353 | 45.33 | 45 | 535723 |
| SPVP140143 | SPVP140143 | SAMN20352827 | SRR15218954 | NonPC | Public | 2014 | China | Env | 87.8 | 100 | 0.34 | 98.2028 | 5143852 | 45.14 | 62 | 655920 |
| SPVP140144 | SPVP140144 | SAMN20352824 | SRR15218957 | NonPC | Public | 2014 | China | Env | 91.5 | 100 | 0 | 98.4136 | 5004879 | 45.35 | 42 | 449170 |
| SPVP140146 | SPVP140146 | SAMN20352828 | SRR15218953 | NonPC | Public | 2014 | China | Env | 87.8 | 100 | 0.6 | 98.2041 | 5143931 | 45.16 | 68 | 570489 |
| SPVP140147 | SPVP140147 | SAMN20352831 | SRR15218949 | NonPC | Public | 2014 | China | Env | 90.6 | 100 | 0 | 98.3291 | 5057552 | 45.26 | 49 | 454317 |
| SPVP140148 | SPVP140148 | SAMN20352839 | SRR15218835 | NonPC | Public | 2014 | China | Env | 90.1 | 100 | 0.05 | 98.3251 | 5058817 | 45.27 | 70 | 455438 |
| SPVP140149 | SPVP140149 | SAMN20352840 | SRR15218832 | NonPC | Public | 2014 | China | Env | 88.5 | 100 | 0.64 | 98.2766 | 4984870 | 45.33 | 49 | 347907 |
| SPVP140150 | SPVP140150 | SAMN20352841 | SRR15218831 | NonPC | Public | 2014 | China | Env | 88.2 | 100 | 0.3 | 98.2421 | 5107794 | 45.17 | 52 | 560808 |
| SPVP140151 | SPVP140151 | SAMN20352869 | SRR15218801 | NonPC | Public | 2014 | China | Env | 88.1 | 100 | 0.43 | 98.2268 | 5025161 | 45.35 | 43 | 383055 |
| SPVP140152 | SPVP140152 | SAMN20352857 | SRR15218814 | NonPC | Public | 2014 | China | Env | 90.8 | 100 | 0.03 | 98.3538 | 4983507 | 45.32 | 46 | 349876 |
| SPVP140153 | SPVP140153 | SAMN20352870 | SRR15218799 | NonPC | Public | 2014 | China | Env | 89.7 | 100 | 0.26 | 98.2807 | 5194729 | 45.31 | 54 | 789794 |
| SPVP140154 | SPVP140154 | SAMN20352871 | SRR15218798 | NonPC | Public | 2014 | China | Env | 91.6 | 100 | 0 | 98.356 | 5079639 | 45.29 | 99 | 401349 |
| SPVP140155 | SPVP140155 | SAMN20352872 | SRR15218797 | NonPC | Public | 2014 | China | Env | 89.7 | 100 | 0.11 | 98.287 | 5248701 | 45.19 | 67 | 259678 |
| SPVP140156 | SPVP140156 | SAMN20352852 | SRR15218819 | NonPC | Public | 2014 | China | Env | 89.4 | 100 | 0 | 98.3677 | 4984217 | 45.33 | 33 | 324674 |
| SPVP140157 | SPVP140157 | SAMN20352873 | SRR15218796 | NonPC | Public | 2014 | China | Env | 90 | 100 | 0 | 98.3575 | 5027186 | 45.35 | 46 | 462412 |
| SPVP140159 | SPVP140159 | SAMN20352874 | SRR15218795 | NonPC | Public | 2014 | China | Env | 90.4 | 100 | 0 | 98.2688 | 4983557 | 45.41 | 30 | 478403 |
| SPVP140160 | SPVP140160 | SAMN20352875 | SRR15218794 | NonPC | Public | 2014 | China | Env | 88.2 | 100 | 0.46 | 98.2743 | 5033141 | 45.27 | 53 | 395579 |
| SPVP140162 | SPVP140162 | SAMN20352876 | SRR15218793 | NonPC | Public | 2014 | China | Env | 88.3 | 100 | 0.39 | 98.2926 | 5108904 | 45.16 | 75 | 520599 |
| SPVP140163 | SPVP140163 | SAMN20352849 | SRR15218823 | NonPC | Public | 2014 | China | Env | 88.4 | 100 | 0.36 | 98.2465 | 5196780 | 45.18 | 44 | 451551 |
| SPVP140164 | SPVP140164 | SAMN20352893 | SRR15219451 | NonPC | Public | 2014 | China | Env | 89.9 | 100 | 0.07 | 98.3345 | 5018214 | 45.29 | 51 | 457952 |
| SPVP140165 | SPVP140165 | SAMN20352894 | SRR15219450 | NonPC | Public | 2014 | China | Env | 90.8 | 100 | 0 | 98.3751 | 5030266 | 45.35 | 36 | 511670 |
| SPVP140166 | SPVP140166 | SAMN20352895 | SRR15219449 | NonPC | Public | 2014 | China | Env | 90.1 | 100 | 0.31 | 98.2997 | 5107832 | 45.33 | 49 | 459621 |
| SPVP140167 | SPVP140167 | SAMN20352892 | SRR15219452 | NonPC | Public | 2014 | China | Env | 88 | 100 | 0.3 | 98.2882 | 5015011 | 45.33 | 51 | 421486 |
| SPVP140170 | SPVP140170 | SAMN20352896 | SRR15219448 | NonPC | Public | 2014 | China | Env | 89.5 | 100 | 0.41 | 98.36 | 5088321 | 45.43 | 127 | 92159 |
| SPVP140172 | SPVP140172 | SAMN20352907 | SRR15219367 | NonPC | Public | 2014 | China | Env | 89.5 | 100 | 0 | 98.3212 | 4981333 | 45.32 | 25 | 519652 |
| SPVP140173 | SPVP140173 | SAMN20352908 | SRR15219366 | NonPC | Public | 2014 | China | Env | 89.8 | 100 | 0.3 | 98.354 | 4936765 | 45.47 | 33 | 464260 |
| SPVP140174 | SPVP140174 | SAMN20352842 | SRR15218830 | NonPC | Public | 2014 | China | Env | 89.2 | 100 | 0.05 | 98.2498 | 4943100 | 45.35 | 36 | 504680 |
| SPVP140176 | SPVP140176 | SAMN20352909 | SRR15219365 | NonPC | Public | 2014 | China | Env | 88.6 | 100 | 0.41 | 98.2158 | 5057663 | 45.29 | 64 | 452392 |
| SPVP140177 | SPVP140177 | SAMN20352910 | SRR15219363 | NonPC | Public | 2014 | China | Env | 89.9 | 100 | 0 | 98.3134 | 5058070 | 45.34 | 49 | 530527 |
| SPVP140178 | SPVP140178 | SAMN20352912 | SRR15219361 | NonPC | Public | 2014 | China | Env | 91.3 | 100 | 0.27 | 98.325 | 5143900 | 45.3 | 78 | 334691 |
| SPVP140180 | SPVP140180 | SAMN20352913 | SRR15219360 | NonPC | Public | 2014 | China | Env | 90 | 100 | 0.24 | 98.3188 | 5061415 | 45.29 | 29 | 518575 |
| SPVP140182 | SPVP140182 | SAMN20352922 | SRR15219350 | NonPC | Public | 2014 | China | Env | 88.1 | 100 | 0.39 | 98.2144 | 5150333 | 45.25 | 34 | 746570 |
| SPVP140184 | SPVP140184 | SAMN20352931 | SRR15219272 | NonPC | Public | 2014 | China | Env | 91.3 | 100 | 0.41 | 98.3069 | 5105656 | 45.22 | 47 | 383020 |
| SPVP140185 | SPVP140185 | SAMN20352932 | SRR15219271 | NonPC | Public | 2014 | China | Env | 92.1 | 100 | 0 | 98.3818 | 5047102 | 45.35 | 51 | 522176 |
| SPVP140186 | SPVP140186 | SAMN20352933 | SRR15219270 | NonPC | Public | 2014 | China | Env | 90.9 | 100 | 0.14 | 98.2883 | 5345980 | 45.18 | 66 | 261634 |
| SPVP140187 | SPVP140187 | SAMN20352930 | SRR15219273 | NonPC | Public | 2014 | China | Env | 88.2 | 100 | 1.02 | 98.3067 | 5276448 | 44.85 | 73 | 206275 |
| SPVP140188 | SPVP140188 | SAMN20352929 | SRR15219276 | NonPC | Public | 2014 | China | Env | 88.8 | 100 | 0.27 | 98.2076 | 5112589 | 45.48 | 67 | 201674 |
| SPVP140190 | SPVP140190 | SAMN20352934 | SRR15219269 | NonPC | Public | 2014 | China | Env | 88.4 | 100 | 0.39 | 98.2135 | 5036489 | 45.29 | 73 | 451105 |
| SPVP140191 | SPVP140191 | SAMN20352923 | SRR15219349 | NonPC | Public | 2014 | China | Env | 89.2 | 100 | 0.05 | 98.3159 | 5041090 | 45.25 | 45 | 607179 |
| SPVP140193 | SPVP140193 | SAMN20352692 | SRR15218719 | NonPC | Public | 2014 | China | Env | 90 | 100 | 0.14 | 98.423 | 5013376 | 45.37 | 33 | 405238 |
| SPVP140194 | SPVP140194 | SAMN20352693 | SRR15218718 | NonPC | Public | 2014 | China | Env | 91.5 | 100 | 0.27 | 98.278 | 5053360 | 45.39 | 75 | 259490 |
| SPVP140195 | SPVP140195 | SAMN20352708 | SRR15218702 | NonPC | Public | 2014 | China | Env | 90.8 | 100 | 0.05 | 98.375 | 5073060 | 45.26 | 44 | 521202 |
| SPVP140196 | SPVP140196 | SAMN20352694 | SRR15218717 | NonPC | Public | 2014 | China | Env | 88.7 | 100 | 0.39 | 98.2279 | 5120088 | 45.28 | 69 | 544155 |
| SPVP140197 | SPVP140197 | SAMN20352695 | SRR15218716 | NonPC | Public | 2014 | China | Env | 89.9 | 100 | 0.05 | 98.3892 | 5003945 | 45.31 | 43 | 348947 |
| SPVP140198 | SPVP140198 | SAMN20352696 | SRR15218715 | NonPC | Public | 2014 | China | Env | 90.3 | 100 | 0.05 | 98.1918 | 5058703 | 45.31 | 51 | 459388 |

| | | | | | | | | | | | | | | | | |
|---|---|---|---|---|---|---|---|---|---|---|---|---|---|---|---|---|
| SPVP140199 | SPVP140199 | SAMN20352684 | SRR15218760 | NonPC | Public | 2014 | China | Env | 88.7 | 100 | 0.39 | 98.2266 | 5094176 | 45.22 | 82 | 335704 |
| SPVP140202 | SPVP140202 | SAMN20352971 | SRR15219157 | NonPC | Public | 2014 | China | Env | 88.3 | 100 | 0.3 | 98.2631 | 5016334 | 45.31 | 71 | 326078 |
| SPVP140203 | SPVP140203 | SAMN20352964 | SRR15219165 | NonPC | Public | 2014 | China | Env | 88.3 | 100 | 0.3 | 98.1806 | 5177591 | 45.21 | 51 | 380323 |
| SPVP140204 | SPVP140204 | SAMN20352998 | SRR15218744 | NonPC | Public | 2014 | China | Env | 88.7 | 100 | 0.39 | 98.233 | 5282541 | 45.08 | 69 | 568254 |
| SPVP140206 | SPVP140206 | SAMN20353004 | SRR15218737 | NonPC | Public | 2014 | China | Env | 90 | 100 | 0 | 98.2697 | 5005433 | 45.35 | 58 | 462036 |
| SPVP140207 | SPVP140207 | SAMN20353005 | SRR15218736 | NonPC | Public | 2014 | China | Env | 90 | 100 | 0 | 98.3044 | 5005508 | 45.35 | 59 | 462036 |
| SPVP140210 | SPVP140210 | SAMN20352953 | SRR15219247 | NonPC | Public | 2014 | China | Env | 89.7 | 100 | 0 | 98.3084 | 4970046 | 45.33 | 32 | 510000 |
| SPVP140211 | SPVP140211 | SAMN20352972 | SRR15219156 | NonPC | Public | 2014 | China | Env | 88.4 | 100 | 0.3 | 98.2047 | 5094709 | 45.29 | 43 | 442828 |
| SPVP140212 | SPVP140212 | SAMN20352973 | SRR15219155 | NonPC | Public | 2014 | China | Env | 89.7 | 100 | 0.11 | 98.3078 | 5094724 | 45.25 | 73 | 458408 |
| SPVP140213 | SPVP140213 | SAMN20353002 | SRR15218739 | NonPC | Public | 2014 | China | Env | 88.6 | 100 | 0.61 | 98.2577 | 5016334 | 45.27 | 58 | 427648 |
| SPVP140214 | SPVP140214 | SAMN20352999 | SRR15218743 | NonPC | Public | 2014 | China | Env | 88.9 | 100 | 0.39 | 98.2345 | 5260401 | 45.16 | 56 | 451610 |
| SPVP140215 | SPVP140215 | SAMN20352974 | SRR15219154 | NonPC | Public | 2014 | China | Env | 88.9 | 100 | 0.39 | 98.2315 | 5173945 | 45.04 | 67 | 553496 |
| SPVP140216 | SPVP140216 | SAMN20352997 | SRR15218745 | NonPC | Public | 2014 | China | Env | 88.2 | 100 | 0.39 | 98.1625 | 5072466 | 45.21 | 54 | 542149 |
| SPVP140217 | SPVP140217 | SAMN20352958 | SRR15219242 | NonPC | Public | 2014 | China | Env | 90.1 | 100 | 0.06 | 98.3578 | 5212997 | 45.36 | 45 | 379238 |
| SPVP140218 | SPVP140218 | SAMN20352975 | SRR15219153 | NonPC | Public | 2014 | China | Env | 91 | 100 | 0.05 | 98.3014 | 5287440 | 45.19 | 79 | 237458 |
| SPVP140219 | SPVP140219 | SAMN20352976 | SRR15219152 | NonPC | Public | 2014 | China | Env | 89.7 | 100 | 0.27 | 98.2984 | 4976790 | 45.42 | 41 | 457334 |
| SPVP140220 | SPVP140220 | SAMN20352977 | SRR15219151 | NonPC | Public | 2014 | China | Env | 88.1 | 100 | 0.33 | 98.2635 | 4973747 | 45.37 | 42 | 448232 |
| SPVP140221 | SPVP140221 | SAMN20352965 | SRR15219164 | NonPC | Public | 2014 | China | Env | 91.7 | 100 | 0 | 98.2835 | 5016827 | 45.32 | 40 | 451945 |
| SPVP140222 | SPVP140222 | SAMN20352978 | SRR15219150 | NonPC | Public | 2014 | China | Env | 91 | 100 | 0.54 | 98.4042 | 5151292 | 45.24 | 47 | 603727 |
| SPVP140223 | SPVP140223 | SAMN20352955 | SRR15219245 | NonPC | Public | 2014 | China | Env | 92.1 | 100 | 0.26 | 98.3573 | 5256709 | 45.1 | 57 | 336949 |
| SPVP140224 | SPVP140224 | SAMN20353003 | SRR15218738 | NonPC | Public | 2014 | China | Env | 88.6 | 100 | 0.61 | 98.2377 | 5056068 | 45.27 | 57 | 641866 |
| SPVP140225 | SPVP140225 | SAMN20352979 | SRR15219149 | PrePC | Public | 2014 | China | Env | 93.6 | 100 | 0 | 99.6703 | 5046686 | 45.4 | 51 | 401830 |
| SPVP140226 | SPVP140226 | SAMN20352980 | SRR15219147 | NonPC | Public | 2014 | China | Env | 91.7 | 100 | 0.07 | 98.2978 | 5270074 | 45.12 | 73 | 268776 |
| SPVP140227 | SPVP140227 | SAMN20352981 | SRR15219146 | NonPC | Public | 2014 | China | Env | 89.9 | 100 | 0.05 | 98.3354 | 5147991 | 45.31 | 51 | 520698 |
| SPVP140228 | SPVP140228 | SAMN20352982 | SRR15219145 | NonPC | Public | 2014 | China | Env | 93.1 | 100 | 0.05 | 98.3313 | 5125945 | 45.26 | 55 | 313446 |
| SPVP140229 | SPVP140229 | SAMN20352957 | SRR15219243 | NonPC | Public | 2014 | China | Env | 91.1 | 99.73 | 0.46 | 98.3395 | 5245999 | 45.23 | 59 | 388290 |
| SPVP140230 | SPVP140230 | SAMN20352983 | SRR15219144 | NonPC | Public | 2014 | China | Env | 89.8 | 100 | 0.05 | 98.3438 | 4968079 | 45.38 | 61 | 400032 |
| SPVP140232 | SPVP140232 | SAMN20352956 | SRR15219244 | NonPC | Public | 2014 | China | Env | 91.4 | 100 | 0 | 98.3934 | 5122015 | 45.28 | 37 | 449124 |
| SPVP140233 | SPVP140233 | SAMN20352984 | SRR15219143 | NonPC | Public | 2014 | China | Env | 90 | 100 | 0.14 | 98.3117 | 5075759 | 45.3 | 39 | 388469 |
| SPVP140234 | SPVP140234 | SAMN20353000 | SRR15218741 | NonPC | Public | 2014 | China | Env | 88.5 | 100 | 0.66 | 98.2604 | 5028720 | 45.23 | 83 | 544236 |
| SPVP140235 | SPVP140235 | SAMN20353001 | SRR15218740 | NonPC | Public | 2014 | China | Env | 88.6 | 100 | 0.66 | 98.2534 | 5031198 | 45.23 | 78 | 727135 |
| SPVP140237 | SPVP140237 | SAMN20352985 | SRR15219142 | NonPC | Public | 2014 | China | Env | 87.5 | 100 | 0.6 | 98.2252 | 5012888 | 45.28 | 66 | 322813 |
| SPVP140238 | SPVP140238 | SAMN20352986 | SRR15219141 | NonPC | Public | 2014 | China | Env | 91.9 | 100 | 0 | 98.3657 | 5094632 | 45.33 | 38 | 373609 |
| SPVP140239 | SPVP140239 | SAMN20352987 | SRR15219140 | NonPC | Public | 2014 | China | Env | 87.4 | 100 | 0.43 | 98.2196 | 5137636 | 45.17 | 111 | 381501 |
| SPVP140240 | SPVP140240 | SAMN20352954 | SRR15219246 | NonPC | Public | 2014 | China | Env | 89.7 | 100 | 0.05 | 98.3564 | 4913062 | 45.46 | 33 | 529112 |
| SPVP140243 | SPVP140243 | SAMN20352960 | SRR15219169 | NonPC | Public | 2014 | China | Env | 89.3 | 100 | 0.33 | 98.279 | 5069035 | 45.23 | 43 | 391386 |
| SPVP140244 | SPVP140244 | SAMN20352959 | SRR15219241 | NonPC | Public | 2014 | China | Env | 90 | 100 | 0.13 | 98.3278 | 5235954 | 45.36 | 57 | 375543 |
| SPVP140247 | SPVP140247 | SAMN20352988 | SRR15219139 | NonPC | Public | 2014 | China | Env | 90.1 | 100 | 0 | 98.3282 | 5075682 | 45.26 | 66 | 530400 |
| SPVP140249 | SPVP140249 | SAMN20352989 | SRR15218754 | NonPC | Public | 2014 | China | Env | 92.2 | 100 | 0.05 | 98.3403 | 5166635 | 45.25 | 46 | 391956 |
| SPVP140250 | SPVP140250 | SAMN20352990 | SRR15218752 | NonPC | Public | 2014 | China | Env | 92.6 | 100 | 0.33 | 98.2695 | 5447919 | 45.06 | 56 | 386022 |
| SPVP140251 | SPVP140251 | SAMN20352966 | SRR15219163 | NonPC | Public | 2014 | China | Env | 90.1 | 100 | 0 | 98.3422 | 5056887 | 45.3 | 50 | 463780 |
| SPVP140252 | SPVP140252 | SAMN20352962 | SRR15219167 | NonPC | Public | 2014 | China | Env | 91 | 100 | 0 | 98.3861 | 4966958 | 45.36 | 48 | 507958 |
| SPVP140254 | SPVP140254 | SAMN20352968 | SRR15219161 | NonPC | Public | 2014 | China | Env | 90.8 | 100 | 0.19 | 98.2572 | 5258786 | 45.24 | 78 | 437865 |
| SPVP140255 | SPVP140255 | SAMN20352991 | SRR15218751 | NonPC | Public | 2014 | China | Env | 91.4 | 100 | 0.13 | 98.3386 | 5243043 | 45.38 | 83 | 298397 |
| SPVP140256 | SPVP140256 | SAMN20352992 | SRR15218750 | NonPC | Public | 2014 | China | Env | 91.2 | 100 | 0.27 | 98.3445 | 5093592 | 45.29 | 76 | 463904 |
| SPVP140257 | SPVP140257 | SAMN20352967 | SRR15219162 | NonPC | Public | 2014 | China | Env | 88.4 | 100 | 0.39 | 98.1839 | 5121140 | 45.15 | 80 | 481637 |
| SPVP140258 | SPVP140258 | SAMN20352961 | SRR15219168 | NonPC | Public | 2014 | China | Env | 90.3 | 100 | 0.53 | 98.3007 | 5121333 | 45.31 | 61 | 314240 |
| SPVP140259 | SPVP140259 | SAMN20352970 | SRR15219158 | NonPC | Public | 2014 | China | Env | 88.2 | 100 | 0.4 | 98.2821 | 5140611 | 45.18 | 97 | 763314 |

| | | | | | | | | | | | | | | | | |
|---|---|---|---|---|---|---|---|---|---|---|---|---|---|---|---|---|
| SPVP140260 | SPVP140260 | SAMN20352993 | SRR15218749 | NonPC | Public | 2014 | China | Env | 91.7 | 100 | 0.04 | 98.3322 | 5446088 | 45.13 | 81 | 268898 |
| SPVP140261 | SPVP140261 | SAMN20352969 | SRR15219160 | NonPC | Public | 2014 | China | Env | 90.8 | 100 | 0.19 | 98.2399 | 5305929 | 45.25 | 90 | 434930 |
| SPVP140262 | SPVP140262 | SAMN20352994 | SRR15218748 | NonPC | Public | 2014 | China | Env | 91.4 | 100 | 0.1 | 98.3765 | 5013281 | 45.4 | 45 | 463707 |
| SPVP140263 | SPVP140263 | SAMN20352995 | SRR15218747 | NonPC | Public | 2014 | China | Env | 91.7 | 100 | 0.27 | 98.357 | 5357928 | 45.19 | 39 | 404482 |
| SPVP140265 | SPVP140265 | SAMN20352963 | SRR15219166 | NonPC | Public | 2014 | China | Env | 89.6 | 100 | 0 | 98.2956 | 5094885 | 45.22 | 65 | 454317 |
| SPVP140266 | SPVP140266 | SAMN20352996 | SRR15218746 | NonPC | Public | 2014 | China | Env | 89.5 | 100 | 0 | 98.2829 | 5059585 | 45.28 | 34 | 545590 |
| SPVP140268 | SPVP140268 | SAMN20352697 | SRR15218714 | NonPC | Public | 2014 | China | Env | 91.5 | 100 | 0.32 | 98.3501 | 5155887 | 45.3 | 75 | 313781 |
| SPVP140269 | SPVP140269 | SAMN20352690 | SRR15218721 | NonPC | Public | 2014 | China | Env | 88.4 | 100 | 0.53 | 98.2161 | 5050940 | 45.19 | 79 | 365543 |
| SPVP140270 | SPVP140270 | SAMN20352716 | SRR15218693 | NonPC | Public | 2014 | China | Env | 91.8 | 100 | 0 | 98.3037 | 5067715 | 45.3 | 52 | 458160 |
| SPVP140271 | SPVP140271 | SAMN20352685 | SRR15218759 | NonPC | Public | 2014 | China | Env | 88.3 | 100 | 0.42 | 98.1928 | 5069678 | 45.25 | 52 | 427729 |
| SPVP140272 | SPVP140272 | SAMN20352717 | SRR15218692 | NonPC | Public | 2014 | China | Env | 91.4 | 100 | 0.05 | 98.3931 | 5173133 | 44.99 | 32 | 448653 |
| SPVP140273 | SPVP140273 | SAMN20352698 | SRR15218713 | NonPC | Public | 2014 | China | Env | 89.9 | 100 | 0.05 | 98.2633 | 5065931 | 45.31 | 74 | 525697 |
| SPVP140274 | SPVP140274 | SAMN20352686 | SRR15218758 | NonPC | Public | 2014 | China | Env | 92 | 100 | 0 | 98.3934 | 5057983 | 45.31 | 33 | 544373 |
| SPVP140276 | SPVP140276 | SAMN20352699 | SRR15218712 | NonPC | Public | 2014 | China | Env | 91 | 100 | 0.1 | 98.2466 | 4975535 | 45.41 | 35 | 508533 |
| SPVP140277 | SPVP140277 | SAMN20352689 | SRR15218755 | NonPC | Public | 2014 | China | Env | 88.1 | 100 | 0.3 | 98.2712 | 5140662 | 45.25 | 37 | 581571 |
| SPVP140278 | SPVP140278 | SAMN20352688 | SRR15218756 | NonPC | Public | 2014 | China | Env | 88.5 | 100 | 0.3 | 98.1968 | 5080260 | 45.22 | 45 | 451762 |
| SPVP140279 | SPVP140279 | SAMN20352691 | SRR15218720 | NonPC | Public | 2014 | China | Env | 88.4 | 100 | 0.3 | 98.2658 | 5097081 | 45.21 | 92 | 450081 |
| SPVP140280 | SPVP140280 | SAMN20352687 | SRR15218757 | NonPC | Public | 2014 | China | Env | 88.9 | 100 | 0.39 | 98.2224 | 5123104 | 45.22 | 44 | 782750 |
| SPVP140281 | SPVP140281 | SAMN20352683 | SRR15218761 | NonPC | Public | 2014 | China | Env | 91.4 | 100 | 0.1 | 98.327 | 5219214 | 45.21 | 56 | 404054 |
| SPVP140282 | SPVP140282 | SAMN20352700 | SRR15218710 | NonPC | Public | 2014 | China | Env | 88.3 | 100 | 0.38 | 98.2346 | 5131933 | 45.26 | 72 | 355240 |
| SPVP140283 | SPVP140283 | SAMN20352701 | SRR15218709 | NonPC | Public | 2014 | China | Env | 88.4 | 100 | 0.3 | 98.2239 | 5022045 | 45.26 | 65 | 357214 |
| SPVP140284 | SPVP140284 | SAMN20352702 | SRR15218708 | NonPC | Public | 2014 | China | Env | 91.2 | 100 | 0 | 98.3315 | 5140842 | 45.25 | 40 | 487171 |
| SPVP140285 | SPVP140285 | SAMN20352703 | SRR15218707 | NonPC | Public | 2014 | China | Env | 91.3 | 100 | 0.03 | 98.2638 | 5326949 | 45.08 | 54 | 450611 |
| SPVP140286 | SPVP140286 | SAMN20352704 | SRR15218706 | NonPC | Public | 2014 | China | Env | 89.9 | 100 | 0.05 | 98.3189 | 5081625 | 45.32 | 55 | 521354 |
| SPVP140292 | SPVP140292 | SAMN20352705 | SRR15218705 | NonPC | Public | 2014 | China | Env | 88.2 | 100 | 0.52 | 98.1674 | 5133807 | 45.14 | 93 | 398419 |
| SPVP140293 | SPVP140293 | SAMN20352706 | SRR15218704 | NonPC | Public | 2014 | China | Env | 91.3 | 100 | 0 | 98.3224 | 5047142 | 45.31 | 51 | 516633 |
| SPVP150010 | SPVP150010 | SAMN20353036 | SRR15218669 | NonPC | Public | 2015 | China | Env | 88.4 | 100 | 0.3 | 98.2568 | 5020131 | 45.32 | 33 | 539678 |
| SPVP150011 | SPVP150011 | SAMN20353037 | SRR15218668 | NonPC | Public | 2015 | China | Env | 88.4 | 100 | 0.3 | 98.2615 | 5019978 | 45.32 | 36 | 539678 |
| SPVP150012 | SPVP150012 | SAMN20353038 | SRR15218667 | NonPC | Public | 2015 | China | Env | 88.3 | 100 | 0.3 | 98.257 | 5019532 | 45.32 | 34 | 743246 |
| SPVP150013 | SPVP150013 | SAMN20353039 | SRR15218666 | NonPC | Public | 2015 | China | Env | 88.4 | 100 | 0.3 | 98.2641 | 5019233 | 45.32 | 33 | 539678 |
| SPVP150014 | SPVP150014 | SAMN20353040 | SRR15218663 | NonPC | Public | 2015 | China | Env | 88.4 | 100 | 0.3 | 98.2466 | 5020000 | 45.32 | 33 | 743246 |
| SPVP150015 | SPVP150015 | SAMN20353041 | SRR15218662 | NonPC | Public | 2015 | China | Env | 88.4 | 100 | 0.3 | 98.2379 | 5019877 | 45.32 | 35 | 897995 |
| SPVP150016 | SPVP150016 | SAMN20353042 | SRR15218661 | NonPC | Public | 2015 | China | Env | 88.3 | 100 | 0.3 | 98.256 | 5020238 | 45.32 | 35 | 539678 |
| SPVP1500163 | SPVP1500163 | SAMN20353280 | SRR15219289 | NonPC | Public | 2015 | China | Env | 88.2 | 100 | 0.36 | 98.2393 | 5033159 | 45.29 | 82 | 407252 |
| SPVP1500164 | SPVP1500164 | SAMN20353281 | SRR15219288 | NonPC | Public | 2015 | China | Env | 91.1 | 100 | 0 | 98.343 | 5011393 | 45.32 | 34 | 456286 |
| SPVP1500165 | SPVP1500165 | SAMN20353279 | SRR15219291 | NonPC | Public | 2015 | China | Env | 91.3 | 100 | 0.12 | 98.352 | 5241604 | 45.35 | 49 | 456607 |
| SPVP150017 | SPVP150017 | SAMN20353035 | SRR15218670 | NonPC | Public | 2015 | China | Env | 90 | 100 | 0 | 98.3111 | 5001053 | 45.39 | 63 | 463791 |
| SPVP150018 | SPVP150018 | SAMN20353075 | SRR15218589 | NonPC | Public | 2015 | China | Env | 90 | 100 | 0 | 98.3715 | 5000201 | 45.39 | 63 | 463791 |
| SPVP150019 | SPVP150019 | SAMN20353063 | SRR15218602 | NonPC | Public | 2015 | China | Env | 90.8 | 100 | 0.05 | 98.2969 | 4996021 | 45.37 | 30 | 466549 |
| SPVP150021 | SPVP150021 | SAMN20353047 | SRR15218620 | NonPC | Public | 2015 | China | Env | 89.7 | 100 | 0 | 98.2928 | 4938930 | 45.37 | 48 | 625590 |
| SPVP150022 | SPVP150022 | SAMN20353048 | SRR15218619 | NonPC | Public | 2015 | China | Env | 89.6 | 100 | 0.05 | 98.2605 | 4993032 | 45.32 | 64 | 605648 |
| SPVP150024 | SPVP150024 | SAMN20353049 | SRR15218618 | NonPC | Public | 2015 | China | Env | 90.9 | 100 | 0 | 98.3261 | 4945327 | 45.4 | 41 | 461960 |
| SPVP150025 | SPVP150025 | SAMN20353050 | SRR15218616 | NonPC | Public | 2015 | China | Env | 89.6 | 100 | 0 | 98.3167 | 5024469 | 45.36 | 39 | 386894 |
| SPVP150026 | SPVP150026 | SAMN20353044 | SRR15218659 | NonPC | Public | 2015 | China | Env | 91.6 | 100 | 0.54 | 98.3171 | 5257365 | 45.25 | 56 | 242920 |
| SPVP150027 | SPVP150027 | SAMN20353043 | SRR15218660 | NonPC | Public | 2015 | China | Env | 90.4 | 100 | 0.05 | 98.3125 | 5142770 | 45.22 | 60 | 269713 |
| SPVP150029 | SPVP150029 | SAMN20353064 | SRR15218601 | NonPC | Public | 2015 | China | Env | 91.4 | 100 | 0.05 | 98.3549 | 5135272 | 45.2 | 57 | 395794 |
| SPVP150030 | SPVP150030 | SAMN20353065 | SRR15218600 | NonPC | Public | 2015 | China | Env | 91.3 | 100 | 0.32 | 98.3272 | 5006897 | 45.38 | 47 | 464415 |
| SPVP150031 | SPVP150031 | SAMN20353060 | SRR15218605 | NonPC | Public | 2015 | China | Env | 89.4 | 100 | 0.09 | 98.3859 | 4990478 | 45.34 | 33 | 361071 |

| | | | | | | | | | | | | | | | | |
|---|---|---|---|---|---|---|---|---|---|---|---|---|---|---|---|---|
| SPVP150032 | SPVP150032 | SAMN20353051 | SRR15218615 | NonPC | Public | 2015 | China | Env | 87.6 | 100 | 0.36 | 98.2564 | 5081641 | 45.23 | 62 | 552988 |
| SPVP150033 | SPVP150033 | SAMN20353052 | SRR15218614 | NonPC | Public | 2015 | China | Env | 90.8 | 100 | 0 | 98.3414 | 5143703 | 45.18 | 62 | 236589 |
| SPVP150034 | SPVP150034 | SAMN20353053 | SRR15218613 | NonPC | Public | 2015 | China | Env | 89.6 | 100 | 0.09 | 98.3137 | 5064705 | 45.28 | 30 | 450671 |
| SPVP150036 | SPVP150036 | SAMN20353073 | SRR15218591 | NonPC | Public | 2015 | China | Env | 89.8 | 100 | 0.27 | 98.2996 | 5051930 | 45.27 | 71 | 505435 |
| SPVP150037 | SPVP150037 | SAMN20353074 | SRR15218590 | NonPC | Public | 2015 | China | Env | 89.8 | 100 | 0.27 | 98.3235 | 5052190 | 45.27 | 71 | 505435 |
| SPVP150038 | SPVP150038 | SAMN20353058 | SRR15218608 | NonPC | Public | 2015 | China | Env | 91.7 | 100 | 0.27 | 98.3339 | 5186694 | 45.25 | 79 | 460482 |
| SPVP150039 | SPVP150039 | SAMN20353054 | SRR15218612 | NonPC | Public | 2015 | China | Env | 90.6 | 100 | 0 | 98.3102 | 5072235 | 45.27 | 77 | 315328 |
| SPVP150041 | SPVP150041 | SAMN20353055 | SRR15218611 | NonPC | Public | 2015 | China | Env | 91.4 | 100 | 0.09 | 98.3029 | 5046513 | 45.31 | 52 | 414082 |
| SPVP150042 | SPVP150042 | SAMN20353046 | SRR15218656 | NonPC | Public | 2015 | China | Env | 90.8 | 100 | 0.03 | 98.3529 | 5002994 | 45.29 | 49 | 314776 |
| SPVP150043 | SPVP150043 | SAMN20353045 | SRR15218658 | NonPC | Public | 2015 | China | Env | 88.4 | 100 | 0.39 | 98.2875 | 4973556 | 45.37 | 54 | 453453 |
| SPVP150044 | SPVP150044 | SAMN20353056 | SRR15218610 | NonPC | Public | 2015 | China | Env | 89.3 | 100 | 0 | 98.2734 | 4989821 | 45.37 | 50 | 408338 |
| SPVP150045 | SPVP150045 | SAMN20353066 | SRR15218599 | NonPC | Public | 2015 | China | Env | 90.1 | 100 | 0.03 | 98.257 | 5017325 | 45.29 | 42 | 519857 |
| SPVP150047 | SPVP150047 | SAMN20353067 | SRR15218598 | NonPC | Public | 2015 | China | Env | 89.9 | 100 | 0 | 98.308 | 5015290 | 45.33 | 55 | 371057 |
| SPVP150048 | SPVP150048 | SAMN20353068 | SRR15218597 | NonPC | Public | 2015 | China | Env | 91.3 | 100 | 0.05 | 98.3448 | 5072390 | 45.32 | 48 | 457856 |
| SPVP150049 | SPVP150049 | SAMN20353069 | SRR15218596 | NonPC | Public | 2015 | China | Env | 91.8 | 100 | 0.09 | 98.3169 | 5081138 | 45.33 | 44 | 461879 |
| SPVP150050 | SPVP150050 | SAMN20353057 | SRR15218609 | NonPC | Public | 2015 | China | Env | 91.4 | 100 | 0 | 98.3484 | 5174869 | 45.22 | 41 | 521826 |
| SPVP150051 | SPVP150051 | SAMN20353061 | SRR15218604 | NonPC | Public | 2015 | China | Env | 90.2 | 100 | 0 | 98.2873 | 5052441 | 45.35 | 42 | 463780 |
| SPVP150052 | SPVP150052 | SAMN20353072 | SRR15218592 | NonPC | Public | 2015 | China | Env | 91.8 | 100 | 0.27 | 98.2965 | 5105421 | 45.34 | 51 | 458160 |
| SPVP150055 | SPVP150055 | SAMN20353070 | SRR15218594 | NonPC | Public | 2015 | China | Env | 89.5 | 100 | 0 | 98.2886 | 5091988 | 45.32 | 59 | 527951 |
| SPVP150064 | SPVP150064 | SAMN20353059 | SRR15218607 | NonPC | Public | 2015 | China | Env | 91.8 | 100 | 0.33 | 98.3998 | 5135885 | 45.25 | 63 | 532748 |
| SPVP150065 | SPVP150065 | SAMN20353071 | SRR15218593 | NonPC | Public | 2015 | China | Env | 89.7 | 100 | 0 | 98.3514 | 4960874 | 45.34 | 35 | 550454 |
| SPVP150067 | SPVP150067 | SAMN20353062 | SRR15218603 | NonPC | Public | 2015 | China | Env | 88.4 | 100 | 0.53 | 98.24 | 5073060 | 45.16 | 78 | 314863 |
| SPVP150070 | SPVP150070 | SAMN20353083 | SRR15218479 | NonPC | Public | 2015 | China | Env | 91.4 | 100 | 0.05 | 98.2778 | 5057460 | 45.32 | 46 | 461373 |
| SPVP150071 | SPVP150071 | SAMN20353084 | SRR15218478 | NonPC | Public | 2015 | China | Env | 91.5 | 100 | 0.22 | 98.3179 | 5298392 | 45.27 | 63 | 350427 |
| SPVP150074 | SPVP150074 | SAMN20353023 | SRR15218683 | NonPC | Public | 2015 | China | Env | 89.5 | 100 | 0 | 98.3824 | 4941578 | 45.35 | 34 | 550424 |
| SPVP150075 | SPVP150075 | SAMN20353021 | SRR15218685 | NonPC | Public | 2015 | China | Env | 89.2 | 100 | 0.05 | 98.2642 | 5091778 | 45.31 | 35 | 522852 |
| SPVP150076 | SPVP150076 | SAMN20353011 | SRR15218729 | NonPC | Public | 2015 | China | Env | 88.3 | 100 | 0.4 | 98.2557 | 5062833 | 45.2 | 106 | 529264 |
| SPVP150077 | SPVP150077 | SAMN20353009 | SRR15218732 | NonPC | Public | 2015 | China | Env | 91.5 | 100 | 0.08 | 98.4094 | 5060388 | 45.3 | 46 | 401225 |
| SPVP150078 | SPVP150078 | SAMN20353024 | SRR15218682 | NonPC | Public | 2015 | China | Env | 89.6 | 100 | 0 | 98.3483 | 4941384 | 45.35 | 33 | 747308 |
| SPVP150079 | SPVP150079 | SAMN20353022 | SRR15218684 | NonPC | Public | 2015 | China | Env | 89.2 | 100 | 0.05 | 98.2721 | 5091529 | 45.31 | 34 | 522852 |
| SPVP150081 | SPVP150081 | SAMN20353126 | SRR15219088 | NonPC | Public | 2015 | China | Env | 88.2 | 100 | 0.68 | 98.1818 | 5216223 | 45.22 | 74 | 160120 |
| SPVP150083 | SPVP150083 | SAMN20353127 | SRR15219087 | NonPC | Public | 2015 | China | Env | 91.2 | 100 | 0 | 98.3033 | 5128564 | 45.32 | 43 | 430511 |
| SPVP150084 | SPVP150084 | SAMN20353112 | SRR15219103 | NonPC | Public | 2015 | China | Env | 89.8 | 100 | 0.26 | 98.37 | 5156964 | 45.28 | 50 | 454532 |
| SPVP150085 | SPVP150085 | SAMN20353128 | SRR15219086 | NonPC | Public | 2015 | China | Env | 91.7 | 100 | 0 | 98.3215 | 5116745 | 45.25 | 71 | 268807 |
| SPVP150087 | SPVP150087 | SAMN20353129 | SRR15219085 | NonPC | Public | 2015 | China | Env | 89.5 | 100 | 0.36 | 98.3287 | 5002751 | 45.37 | 39 | 533025 |
| SPVP150088 | SPVP150088 | SAMN20353119 | SRR15219096 | NonPC | Public | 2015 | China | Env | 88.2 | 100 | 0.36 | 98.263 | 4998641 | 45.33 | 43 | 582095 |
| SPVP150089 | SPVP150089 | SAMN20353130 | SRR15219083 | NonPC | Public | 2015 | China | Env | 91.7 | 100 | 0.42 | 98.3595 | 5188638 | 45.37 | 54 | 344817 |
| SPVP150090 | SPVP150090 | SAMN20353131 | SRR15219082 | NonPC | Public | 2015 | China | Env | 89.6 | 100 | 0 | 98.2998 | 5210446 | 45.35 | 39 | 440017 |
| SPVP150091 | SPVP150091 | SAMN20353122 | SRR15219092 | NonPC | Public | 2015 | China | Env | 88.5 | 100 | 0.36 | 98.2558 | 5152785 | 45.16 | 80 | 416281 |
| SPVP150092 | SPVP150092 | SAMN20353111 | SRR15219104 | NonPC | Public | 2015 | China | Env | 88.7 | 100 | 0.58 | 98.1945 | 5120156 | 45.23 | 66 | 452142 |
| SPVP150093 | SPVP150093 | SAMN20353132 | SRR15219081 | NonPC | Public | 2015 | China | Env | 91.8 | 100 | 0.09 | 98.3838 | 5022698 | 45.4 | 68 | 316584 |
| SPVP150094 | SPVP150094 | SAMN20353144 | SRR15219035 | NonPC | Public | 2015 | China | Env | 88.3 | 100 | 0.39 | 98.1684 | 5224037 | 45.22 | 66 | 380766 |
| SPVP150095 | SPVP150095 | SAMN20353116 | SRR15219099 | NonPC | Public | 2015 | China | Env | 88.9 | 100 | 0.39 | 98.2158 | 5214946 | 45.14 | 49 | 466586 |
| SPVP150096 | SPVP150096 | SAMN20353133 | SRR15219080 | NonPC | Public | 2015 | China | Env | 91.4 | 100 | 0.27 | 98.2878 | 5052942 | 45.3 | 75 | 461787 |
| SPVP150097 | SPVP150097 | SAMN20353134 | SRR15219079 | NonPC | Public | 2015 | China | Env | 91.3 | 100 | 0.46 | 98.3252 | 5010932 | 45.38 | 60 | 458255 |
| SPVP150098 | SPVP150098 | SAMN20353120 | SRR15219094 | NonPC | Public | 2015 | China | Env | 88.1 | 100 | 0.3 | 98.2301 | 5117974 | 45.25 | 38 | 581755 |
| SPVP150099 | SPVP150099 | SAMN20353113 | SRR15219102 | NonPC | Public | 2015 | China | Env | 88.5 | 100 | 0.36 | 98.235 | 5169705 | 45.1 | 49 | 451551 |
| SPVP150100 | SPVP150100 | SAMN20353135 | SRR15219078 | NonPC | Public | 2015 | China | Env | 90.3 | 100 | 0 | 98.333 | 4976592 | 45.4 | 70 | 332763 |

| | | | | | | | | | | | | | | | | |
|---|---|---|---|---|---|---|---|---|---|---|---|---|---|---|---|---|
| SPVP150101 | SPVP150101 | SAMN20353136 | SRR15219045 | NonPC | Public | 2015 | China | Env | 88.2 | 100 | 0.39 | 98.2323 | 5061765 | 45.28 | 72 | 449520 |
| SPVP150102 | SPVP150102 | SAMN20353125 | SRR15219089 | NonPC | Public | 2015 | China | Env | 91.2 | 100 | 0.09 | 98.3541 | 5184968 | 45.3 | 71 | 405844 |
| SPVP150103 | SPVP150103 | SAMN20353117 | SRR15219098 | NonPC | Public | 2015 | China | Env | 89.2 | 100 | 0.08 | 98.392 | 4982880 | 45.34 | 73 | 151891 |
| SPVP150104 | SPVP150104 | SAMN20353121 | SRR15219093 | NonPC | Public | 2015 | China | Env | 91.2 | 100 | 0.27 | 98.3898 | 4971936 | 45.45 | 41 | 452023 |
| SPVP150105 | SPVP150105 | SAMN20353137 | SRR15219044 | NonPC | Public | 2015 | China | Env | 88.2 | 100 | 0.07 | 98.2965 | 5050836 | 45.31 | 96 | 152126 |
| SPVP150106 | SPVP150106 | SAMN20353118 | SRR15219097 | NonPC | Public | 2015 | China | Env | 88.2 | 100 | 0 | 98.443 | 5172804 | 45.41 | 126 | 123327 |
| SPVP150107 | SPVP150107 | SAMN20353094 | SRR15218467 | NonPC | Public | 2015 | China | Env | 89.2 | 100 | 0 | 98.3809 | 5078367 | 45.35 | 118 | 98497 |
| SPVP150108 | SPVP150108 | SAMN20353093 | SRR15218468 | NonPC | Public | 2015 | China | Env | 89.8 | 100 | 0.03 | 98.2706 | 5055010 | 45.27 | 33 | 453573 |
| SPVP150109 | SPVP150109 | SAMN20353092 | SRR15218469 | NonPC | Public | 2015 | China | Env | 89.3 | 100 | 0.99 | 98.3311 | 5089156 | 45.23 | 77 | 202629 |
| SPVP150110 | SPVP150110 | SAMN20353089 | SRR15218473 | NonPC | Public | 2015 | China | Env | 89 | 100 | 0 | 98.2477 | 5021619 | 45.49 | 54 | 331374 |
| SPVP150111 | SPVP150111 | SAMN20353091 | SRR15218470 | NonPC | Public | 2015 | China | Env | 88.4 | 100 | 0.39 | 98.2257 | 5048553 | 45.25 | 51 | 903887 |
| SPVP150112 | SPVP150112 | SAMN20353090 | SRR15218471 | NonPC | Public | 2015 | China | Env | 91.7 | 100 | 0.54 | 98.3352 | 5288402 | 45.22 | 70 | 239390 |
| SPVP150113 | SPVP150113 | SAMN20353088 | SRR15218474 | NonPC | Public | 2015 | China | Env | 88.8 | 100 | 0.39 | 98.251 | 5058688 | 45.24 | 78 | 544369 |
| SPVP150114 | SPVP150114 | SAMN20353123 | SRR15219091 | NonPC | Public | 2015 | China | Env | 91.2 | 100 | 0 | 98.3825 | 5091637 | 45.24 | 88 | 239134 |
| SPVP150115 | SPVP150115 | SAMN20353114 | SRR15219101 | NonPC | Public | 2015 | China | Env | 92.1 | 100 | 0 | 98.3213 | 5039227 | 45.31 | 37 | 401793 |
| SPVP150116 | SPVP150116 | SAMN20353124 | SRR15219090 | NonPC | Public | 2015 | China | Env | 91.3 | 100 | 0.14 | 98.294 | 5225158 | 45.2 | 43 | 429661 |
| SPVP150117 | SPVP150117 | SAMN20353138 | SRR15219043 | NonPC | Public | 2015 | China | Env | 90.2 | 100 | 0.14 | 98.2047 | 5069476 | 45.38 | 42 | 458045 |
| SPVP150118 | SPVP150118 | SAMN20353110 | SRR15219105 | NonPC | Public | 2015 | China | Env | 91.4 | 100 | 0.05 | 98.3659 | 5089924 | 45.17 | 33 | 448653 |
| SPVP150119 | SPVP150119 | SAMN20353139 | SRR15219042 | NonPC | Public | 2015 | China | Env | 89.9 | 100 | 0.05 | 98.3319 | 4951430 | 45.39 | 49 | 451251 |
| SPVP150120 | SPVP150120 | SAMN20353145 | SRR15219034 | NonPC | Public | 2015 | China | Env | 91 | 100 | 0.24 | 98.3508 | 5301553 | 45.04 | 88 | 229122 |
| SPVP150121 | SPVP150121 | SAMN20353140 | SRR15219039 | NonPC | Public | 2015 | China | Env | 89.3 | 100 | 0 | 98.3406 | 5066369 | 45.32 | 42 | 379985 |
| SPVP150122 | SPVP150122 | SAMN20353115 | SRR15219100 | NonPC | Public | 2015 | China | Env | 88.2 | 100 | 0.36 | 98.2585 | 4997489 | 45.24 | 57 | 452357 |
| SPVP150123 | SPVP150123 | SAMN20353141 | SRR15219038 | NonPC | Public | 2015 | China | Env | 88.6 | 100 | 0.36 | 98.244 | 5140897 | 45.27 | 76 | 369978 |
| SPVP150124 | SPVP150124 | SAMN20353142 | SRR15219037 | NonPC | Public | 2015 | China | Env | 90.5 | 100 | 0.08 | 98.2336 | 5043614 | 45.33 | 60 | 461244 |
| SPVP150125 | SPVP150125 | SAMN20353143 | SRR15219036 | NonPC | Public | 2015 | China | Env | 91.9 | 100 | 0.68 | 98.3349 | 5265261 | 45.19 | 47 | 314867 |
| SPVP150126 | SPVP150126 | SAMN20353099 | SRR15218462 | NonPC | Public | 2015 | China | Env | 88.2 | 100 | 0.3 | 98.2526 | 5048128 | 45.24 | 52 | 380566 |
| SPVP150128 | SPVP150128 | SAMN20353100 | SRR15218460 | NonPC | Public | 2015 | China | Env | 89.8 | 100 | 0.05 | 98.3467 | 4949320 | 45.41 | 46 | 537749 |
| SPVP150129 | SPVP150129 | SAMN20353101 | SRR15218459 | NonPC | Public | 2015 | China | Env | 91.7 | 100 | 0.03 | 98.3355 | 5166070 | 45.27 | 65 | 420047 |
| SPVP150130 | SPVP150130 | SAMN20353102 | SRR15218458 | NonPC | Public | 2015 | China | Env | 88 | 100 | 0.58 | 98.2492 | 5021731 | 45.29 | 54 | 553987 |
| SPVP150131 | SPVP150131 | SAMN20353098 | SRR15218463 | NonPC | Public | 2015 | China | Env | 88.4 | 100 | 0.34 | 98.2044 | 5114585 | 45.2 | 89 | 452966 |
| SPVP150132 | SPVP150132 | SAMN20353103 | SRR15218457 | NonPC | Public | 2015 | China | Env | 89.7 | 100 | 0.27 | 98.342 | 5000903 | 45.3 | 59 | 461572 |
| SPVP150133 | SPVP150133 | SAMN20353104 | SRR15218456 | NonPC | Public | 2015 | China | Env | 90.6 | 100 | 0.05 | 98.237 | 5132987 | 45.26 | 55 | 415644 |
| SPVP150134 | SPVP150134 | SAMN20353105 | SRR15218455 | NonPC | Public | 2015 | China | Env | 91.1 | 100 | 0.54 | 98.3021 | 5149203 | 45.3 | 46 | 459044 |
| SPVP150135 | SPVP150135 | SAMN20353106 | SRR15218454 | NonPC | Public | 2015 | China | Env | 88.1 | 100 | 0.66 | 98.2604 | 4942141 | 45.32 | 44 | 416105 |
| SPVP150136 | SPVP150136 | SAMN20353096 | SRR15218465 | NonPC | Public | 2015 | China | Env | 88.6 | 100 | 0.33 | 98.2537 | 5239761 | 45.08 | 84 | 554102 |
| SPVP150137 | SPVP150137 | SAMN20353107 | SRR15219109 | NonPC | Public | 2015 | China | Env | 89.5 | 100 | 0 | 98.2815 | 5059523 | 45.34 | 48 | 461627 |
| SPVP150138 | SPVP150138 | SAMN20353097 | SRR15218464 | NonPC | Public | 2015 | China | Env | 89.7 | 100 | 0 | 98.3171 | 5026347 | 45.37 | 38 | 573394 |
| SPVP150139 | SPVP150139 | SAMN20353108 | SRR15219108 | NonPC | Public | 2015 | China | Env | 88.6 | 100 | 0.4 | 98.2325 | 5205783 | 45.27 | 45 | 391020 |
| SPVP150140 | SPVP150140 | SAMN20353109 | SRR15219107 | NonPC | Public | 2015 | China | Env | 91.4 | 100 | 0.64 | 98.3547 | 5041313 | 45.33 | 49 | 293315 |
| SPVP150143 | SPVP150143 | SAMN20353163 | SRR15219014 | NonPC | Public | 2015 | China | Env | 91.2 | 100 | 0.27 | 98.3978 | 5040320 | 45.33 | 40 | 447621 |
| SPVP150148 | SPVP150148 | SAMN20353171 | SRR15218900 | NonPC | Public | 2015 | China | Env | 88.6 | 100 | 0.6 | 98.1602 | 5202593 | 45.15 | 68 | 452964 |
| SPVP150149 | SPVP150149 | SAMN20353166 | SRR15218906 | NonPC | Public | 2015 | China | Env | 91.7 | 100 | 0.14 | 98.3695 | 5226732 | 45.12 | 37 | 460959 |
| SPVP150150 | SPVP150150 | SAMN20353167 | SRR15218905 | NonPC | Public | 2015 | China | Env | 89.7 | 100 | 0 | 98.2934 | 4973354 | 45.46 | 28 | 531724 |
| SPVP150151 | SPVP150151 | SAMN20353168 | SRR15218904 | NonPC | Public | 2015 | China | Env | 88.6 | 100 | 0.39 | 98.2406 | 5065427 | 45.24 | 41 | 735547 |
| SPVP150152 | SPVP150152 | SAMN20353169 | SRR15218903 | NonPC | Public | 2015 | China | Env | 91.2 | 100 | 0.14 | 98.3073 | 5069022 | 45.27 | 51 | 313757 |
| SPVP150153 | SPVP150153 | SAMN20353170 | SRR15218901 | NonPC | Public | 2015 | China | Env | 88.4 | 100 | 0.36 | 98.2289 | 5004309 | 45.3 | 46 | 451385 |
| SPVP150154 | SPVP150154 | SAMN20353178 | SRR15218893 | NonPC | Public | 2015 | China | Env | 89.4 | 100 | 0.3 | 98.2971 | 4940623 | 45.37 | 35 | 401486 |
| SPVP150155 | SPVP150155 | SAMN20353177 | SRR15218894 | NonPC | Public | 2015 | China | Env | 91.4 | 100 | 0.05 | 98.3157 | 5066522 | 45.31 | 50 | 337912 |

| | | | | | | | | | | | | | | | | |
|---|---|---|---|---|---|---|---|---|---|---|---|---|---|---|---|---|
| SPVP150156 | SPVP150156 | SAMN20353179 | SRR15218892 | NonPC | Public | 2015 | China | Env | 89.2 | 100 | 0 | 98.3245 | 5078817 | 45.35 | 124 | 94173 |
| SPVP150157 | SPVP150157 | SAMN20353173 | SRR15218898 | NonPC | Public | 2015 | China | Env | 91.5 | 100 | 0.05 | 98.2796 | 5163644 | 45.19 | 60 | 439296 |
| SPVP150158 | SPVP150158 | SAMN20353174 | SRR15218897 | NonPC | Public | 2015 | China | Env | 88.4 | 100 | 0.37 | 98.3066 | 5032193 | 45.26 | 56 | 451579 |
| SPVP150159 | SPVP150159 | SAMN20353175 | SRR15218896 | NonPC | Public | 2015 | China | Env | 89.6 | 100 | 0.08 | 98.2976 | 5187347 | 45.25 | 34 | 536161 |
| SPVP150160 | SPVP150160 | SAMN20353172 | SRR15218899 | NonPC | Public | 2015 | China | Env | 91.6 | 100 | 0.41 | 98.4003 | 5083723 | 45.34 | 56 | 438367 |
| SPVP150161 | SPVP150161 | SAMN20353264 | SRR15219307 | NonPC | Public | 2015 | China | Env | 88.7 | 100 | 0.39 | 98.2165 | 5244899 | 45.12 | 67 | 425402 |
| SPVP150162 | SPVP150162 | SAMN20353013 | SRR15218727 | NonPC | Public | 2015 | China | Env | 88.4 | 100 | 0.34 | 98.1953 | 5143171 | 45.2 | 56 | 765440 |
| SPVP150166 | SPVP150166 | SAMN20353252 | SRR15219387 | NonPC | Public | 2015 | China | Env | 88 | 100 | 0.37 | 98.2923 | 5021035 | 45.34 | 45 | 354011 |
| SPVP150167 | SPVP150167 | SAMN20353247 | SRR15219393 | NonPC | Public | 2015 | China | Env | 91 | 100 | 0.05 | 98.2315 | 5098395 | 45.27 | 42 | 531765 |
| SPVP150168 | SPVP150168 | SAMN20353253 | SRR15219386 | NonPC | Public | 2015 | China | Env | 88 | 100 | 0.37 | 98.3311 | 5021313 | 45.34 | 47 | 354347 |
| SPVP150169 | SPVP150169 | SAMN20353243 | SRR15219397 | NonPC | Public | 2015 | China | Env | 89.5 | 100 | 0 | 98.3462 | 5083891 | 45.13 | 52 | 401127 |
| SPVP150170 | SPVP150170 | SAMN20353248 | SRR15219392 | NonPC | Public | 2015 | China | Env | 89.4 | 100 | 0.06 | 98.336 | 5041034 | 45.27 | 48 | 536092 |
| SPVP150171 | SPVP150171 | SAMN20353240 | SRR15219400 | NonPC | Public | 2015 | China | Env | 87.7 | 100 | 0.47 | 98.2235 | 5306998 | 45.09 | 70 | 345793 |
| SPVP150172 | SPVP150172 | SAMN20353249 | SRR15219391 | NonPC | Public | 2015 | China | Env | 89 | 100 | 0 | 98.3394 | 5061306 | 45.32 | 58 | 194982 |
| SPVP150173 | SPVP150173 | SAMN20353242 | SRR15219398 | NonPC | Public | 2015 | China | Env | 89.7 | 100 | 0.14 | 98.3532 | 5197112 | 45.36 | 49 | 299984 |
| SPVP150174 | SPVP150174 | SAMN20353241 | SRR15219399 | NonPC | Public | 2015 | China | Env | 88.3 | 100 | 0.44 | 98.2764 | 5004533 | 45.28 | 44 | 452368 |
| SPVP150175 | SPVP150175 | SAMN20353245 | SRR15219395 | NonPC | Public | 2015 | China | Env | 88.5 | 100 | 0.59 | 98.3218 | 5136103 | 45.32 | 55 | 443716 |
| SPVP150176 | SPVP150176 | SAMN20353246 | SRR15219394 | NonPC | Public | 2015 | China | Env | 88.3 | 100 | 0.42 | 98.2404 | 5181158 | 45.34 | 44 | 404260 |
| SPVP150178 | SPVP150178 | SAMN20353250 | SRR15219389 | NonPC | Public | 2015 | China | Env | 91.4 | 100 | 0.33 | 98.3301 | 5130533 | 45.36 | 51 | 456605 |
| SPVP150179 | SPVP150179 | SAMN20353251 | SRR15219388 | NonPC | Public | 2015 | China | Env | 91.5 | 100 | 0.33 | 98.3307 | 5130105 | 45.36 | 51 | 456605 |
| SPVP150180 | SPVP150180 | SAMN20353239 | SRR15219403 | NonPC | Public | 2015 | China | Env | 87.8 | 100 | 0.3 | 98.1881 | 5246529 | 45.12 | 95 | 211364 |
| SPVP150181 | SPVP150181 | SAMN20353244 | SRR15219396 | NonPC | Public | 2015 | China | Env | 87.6 | 100 | 0.3 | 98.2173 | 5092563 | 45.18 | 47 | 431610 |
| SPVP150183 | SPVP150183 | SAMN20353282 | SRR15219287 | NonPC | Public | 2015 | China | Env | 90.8 | 100 | 0.05 | 98.3954 | 5072687 | 45.26 | 42 | 446965 |
| SPVP160001 | SPVP160001 | SAMN20353330 | SRR15219133 | NonPC | Public | 2016 | China | Env | 90.7 | 100 | 0.05 | 98.2552 | 4987908 | 45.37 | 53 | 402329 |
| SPVP160003 | SPVP160003 | SAMN20353327 | SRR15219137 | NonPC | Public | 2016 | China | Env | 88.7 | 100 | 0.33 | 98.2406 | 5079482 | 45.21 | 84 | 342661 |
| SPVP160004 | SPVP160004 | SAMN20353331 | SRR15219132 | NonPC | Public | 2016 | China | Env | 90.7 | 100 | 0.05 | 98.2571 | 4988040 | 45.37 | 52 | 453389 |
| SPVP160005 | SPVP160005 | SAMN20353326 | SRR15219138 | NonPC | Public | 2016 | China | Env | 88.8 | 100 | 0.58 | 98.1829 | 5149921 | 45.26 | 62 | 462596 |
| SPVP160006 | SPVP160006 | SAMN20353328 | SRR15219136 | NonPC | Public | 2016 | China | Env | 88.8 | 100 | 0.63 | 98.1964 | 5203686 | 45.22 | 58 | 756226 |
| SPVP160007 | SPVP160007 | SAMN20353329 | SRR15219135 | NonPC | Public | 2016 | China | Env | 88 | 100 | 0.7 | 98.2066 | 5111036 | 45.17 | 91 | 447436 |
| SPVP160008 | SPVP160008 | SAMN20371534 | SRR15239774 | NonPC | Public | 2016 | China | Env | 91 | 100 | 0.1 | 98.3264 | 5084320 | 45.22 | 71 | 229122 |
| SPVP160009 | SPVP160009 | SAMN20353335 | SRR15219128 | NonPC | Public | 2016 | China | Env | 88.4 | 100 | 0.3 | 98.2823 | 5034293 | 45.27 | 75 | 559670 |
| SPVP160010 | SPVP160010 | SAMN20353336 | SRR15219127 | NonPC | Public | 2016 | China | Env | 88.4 | 100 | 0.3 | 98.2342 | 5033796 | 45.27 | 72 | 559670 |
| SPVP160011 | SPVP160011 | SAMN20353334 | SRR15219129 | NonPC | Public | 2016 | China | Env | 90.4 | 100 | 0 | 98.2871 | 5137380 | 45.24 | 35 | 573362 |
| SPVP160012 | SPVP160012 | SAMN20371535 | SRR15239773 | NonPC | Public | 2016 | China | Env | 90.9 | 100 | 0.1 | 98.3591 | 5088007 | 45.22 | 67 | 270690 |
| SPVP160013 | SPVP160013 | SAMN20371536 | SRR15240197 | NonPC | Public | 2016 | China | Env | 90.9 | 100 | 0.1 | 98.3322 | 5088486 | 45.22 | 68 | 270691 |
| SPVP160014 | SPVP160014 | SAMN20353332 | SRR15219131 | NonPC | Public | 2016 | China | Env | 88.9 | 100 | 0.66 | 98.2223 | 5069738 | 45.31 | 44 | 461169 |
| SPVP160015 | SPVP160015 | SAMN20353337 | SRR15219126 | NonPC | Public | 2016 | China | Env | 88.3 | 100 | 0.39 | 98.2321 | 5225382 | 45.22 | 62 | 371652 |
| SPVP160016 | SPVP160016 | SAMN20353338 | SRR15219125 | NonPC | Public | 2016 | China | Env | 88.3 | 100 | 0.39 | 98.2106 | 5224534 | 45.22 | 63 | 474415 |
| SPVP160017 | SPVP160017 | SAMN20353339 | SRR15219124 | NonPC | Public | 2016 | China | Env | 88.2 | 100 | 0.39 | 98.1853 | 5223325 | 45.21 | 65 | 403299 |
| SPVP160018 | SPVP160018 | SAMN20353333 | SRR15219130 | NonPC | Public | 2016 | China | Env | 91.4 | 100 | 0.41 | 98.3261 | 5010198 | 45.45 | 47 | 462184 |
| SPVP170001 | SPVP170001 | SAMN20371750 | SRR15240178 | NonPC | Public | 2017 | China | Env | 91.2 | 100 | 0.08 | 98.2614 | 5210710 | 45.21 | 106 | 216892 |
| SPVP170002 | SPVP170002 | SAMN20371726 | SRR15240241 | NonPC | Public | 2017 | China | Env | 91.7 | 100 | 0.33 | 98.392 | 5140966 | 45.24 | 67 | 371310 |
| SPVP170003 | SPVP170003 | SAMN20371751 | SRR15240177 | NonPC | Public | 2017 | China | Env | 91.2 | 100 | 0.08 | 98.2786 | 5211764 | 45.21 | 107 | 216892 |
| SPVP170004 | SPVP170004 | SAMN20371752 | SRR15240176 | NonPC | Public | 2017 | China | Env | 91.2 | 100 | 0.08 | 98.2653 | 5212084 | 45.21 | 107 | 177812 |
| SPVP170005 | SPVP170005 | SAMN20371746 | SRR15240182 | NonPC | Public | 2017 | China | Env | 88.2 | 100 | 0.39 | 98.209 | 5072586 | 45.28 | 52 | 542136 |
| SPVP170006 | SPVP170006 | SAMN20371729 | SRR15240238 | NonPC | Public | 2017 | China | Env | 87.7 | 100 | 0.39 | 98.2314 | 5098187 | 45.28 | 55 | 748451 |
| SPVP170007 | SPVP170007 | SAMN20371722 | SRR15240246 | NonPC | Public | 2017 | China | Env | 89.8 | 100 | 0.01 | 98.3022 | 5074303 | 45.4 | 33 | 462912 |
| SPVP170008 | SPVP170008 | SAMN20371744 | SRR15240184 | NonPC | Public | 2017 | China | Env | 89.5 | 99.91 | 0.09 | 98.3216 | 5009003 | 45.32 | 53 | 450604 |

| | | | | | | | | | | | | | | | | |
|---|---|---|---|---|---|---|---|---|---|---|---|---|---|---|---|---|
| SPVP170009 | SPVP170009 | SAMN20371730 | SRR15240237 | NonPC | Public | 2017 | China | Env | 89.8 | 100 | 0.08 | 98.3342 | 5093784 | 45.27 | 74 | 460203 |
| SPVP170010 | SPVP170010 | SAMN20371731 | SRR15240200 | NonPC | Public | 2017 | China | Env | 88.6 | 100 | 0.3 | 98.1864 | 5097451 | 45.25 | 104 | 398012 |
| SPVP170011 | SPVP170011 | SAMN20371732 | SRR15240199 | NonPC | Public | 2017 | China | Env | 90 | 100 | 0.33 | 98.2819 | 5018521 | 45.35 | 56 | 453425 |
| SPVP170012 | SPVP170012 | SAMN20371745 | SRR15240183 | NonPC | Public | 2017 | China | Env | 89.5 | 99.91 | 0.09 | 98.3187 | 5008887 | 45.32 | 55 | 450604 |
| SPVP170013 | SPVP170013 | SAMN20371725 | SRR15240242 | NonPC | Public | 2017 | China | Env | 88.4 | 100 | 0.71 | 98.2634 | 5042124 | 45.26 | 46 | 368125 |
| SPVP170014 | SPVP170014 | SAMN20371733 | SRR15240198 | NonPC | Public | 2017 | China | Env | 88.6 | 100 | 0.36 | 98.2463 | 5161090 | 45.16 | 44 | 395971 |
| SPVP170015 | SPVP170015 | SAMN20371734 | SRR15240195 | NonPC | Public | 2017 | China | Env | 91.3 | 100 | 0 | 98.3748 | 4978545 | 45.36 | 53 | 410235 |
| SPVP170016 | SPVP170016 | SAMN20371724 | SRR15240243 | NonPC | Public | 2017 | China | Env | 86.8 | 100 | 0.34 | 98.2187 | 5128584 | 45.24 | 122 | 127360 |
| SPVP170017 | SPVP170017 | SAMN20371735 | SRR15240194 | NonPC | Public | 2017 | China | Env | 88.7 | 100 | 0.39 | 98.1983 | 5134662 | 45.26 | 50 | 504792 |
| SPVP170018 | SPVP170018 | SAMN20371736 | SRR15240193 | NonPC | Public | 2017 | China | Env | 88.2 | 100 | 0.34 | 98.2597 | 5152122 | 45.25 | 56 | 399887 |
| SPVP170019 | SPVP170019 | SAMN20371728 | SRR15240239 | NonPC | Public | 2017 | China | Env | 88.2 | 100 | 0.4 | 98.2976 | 5187981 | 45.21 | 66 | 398002 |
| SPVP170020 | SPVP170020 | SAMN20371743 | SRR15240186 | NonPC | Public | 2017 | China | Env | 90.9 | 100 | 0.1 | 98.3595 | 5087557 | 45.22 | 67 | 317135 |
| SPVP170021 | SPVP170021 | SAMN20371737 | SRR15240192 | NonPC | Public | 2017 | China | Env | 91.6 | 100 | 0 | 98.3786 | 5163377 | 45.3 | 85 | 142565 |
| SPVP170022 | SPVP170022 | SAMN20371723 | SRR15240245 | NonPC | Public | 2017 | China | Env | 88.2 | 100 | 0.3 | 98.2405 | 5134980 | 45.32 | 57 | 241713 |
| SPVP170025 | SPVP170025 | SAMN20371747 | SRR15240181 | NonPC | Public | 2017 | China | Env | 91.7 | 100 | 0 | 98.311 | 5071130 | 45.27 | 52 | 516972 |
| SPVP170026 | SPVP170026 | SAMN20371748 | SRR15240180 | NonPC | Public | 2017 | China | Env | 91.7 | 100 | 0 | 98.3043 | 5070559 | 45.27 | 52 | 516972 |
| SPVP170027 | SPVP170027 | SAMN20371749 | SRR15240179 | NonPC | Public | 2017 | China | Env | 91.8 | 100 | 0 | 98.3059 | 5071387 | 45.27 | 54 | 458161 |
| SPVP170028 | SPVP170028 | SAMN20371727 | SRR15240240 | NonPC | Public | 2017 | China | Env | 87.7 | 100 | 0.36 | 98.2542 | 5059809 | 45.32 | 41 | 546516 |
| SPVP170029 | SPVP170029 | SAMN20371738 | SRR15240191 | NonPC | Public | 2017 | China | Env | 90.8 | 100 | 0.03 | 98.3661 | 5128180 | 45.33 | 74 | 177901 |
| SPVP170030 | SPVP170030 | SAMN20371705 | SRR15240264 | NonPC | Public | 2017 | China | Env | 91.9 | 100 | 0 | 98.3483 | 5241970 | 45.11 | 82 | 184008 |
| SPVP170031 | SPVP170031 | SAMN20371718 | SRR15240250 | NonPC | Public | 2017 | China | Env | 88.2 | 100 | 0.39 | 98.2327 | 5083644 | 45.27 | 53 | 542137 |
| SPVP170032 | SPVP170032 | SAMN20371719 | SRR15240249 | NonPC | Public | 2017 | China | Env | 88.2 | 100 | 0.39 | 98.2139 | 5065000 | 45.22 | 53 | 542356 |
| SPVP170033 | SPVP170033 | SAMN20371707 | SRR15240262 | NonPC | Public | 2017 | China | Env | 89.9 | 100 | 0.03 | 98.4141 | 4958238 | 45.45 | 33 | 517067 |
| SPVP170034 | SPVP170034 | SAMN20371708 | SRR15240261 | NonPC | Public | 2017 | China | Env | 91.2 | 100 | 0.27 | 98.3225 | 5035597 | 45.29 | 50 | 312635 |
| SPVP170035 | SPVP170035 | SAMN20371739 | SRR15240190 | NonPC | Public | 2017 | China | Env | 87.9 | 100 | 0.67 | 98.2248 | 5173780 | 45.19 | 50 | 348797 |
| SPVP170036 | SPVP170036 | SAMN20371740 | SRR15240189 | NonPC | Public | 2017 | China | Env | 89.8 | 100 | 0.27 | 98.3395 | 4960382 | 45.39 | 44 | 454155 |
| SPVP170037 | SPVP170037 | SAMN20371721 | SRR15240247 | NonPC | Public | 2017 | China | Env | 90.6 | 100 | 0.27 | 98.3893 | 4972683 | 45.36 | 37 | 449015 |
| SPVP170038 | SPVP170038 | SAMN20371741 | SRR15240188 | NonPC | Public | 2017 | China | Env | 91.1 | 100 | 0.54 | 98.3096 | 5202926 | 45.1 | 70 | 334336 |
| SPVP170039 | SPVP170039 | SAMN20371742 | SRR15240187 | NonPC | Public | 2017 | China | Env | 88.4 | 100 | 0.41 | 98.2368 | 5067902 | 45.19 | 51 | 451695 |
| SPVP170040 | SPVP170040 | SAMN20371706 | SRR15240263 | NonPC | Public | 2017 | China | Env | 88.4 | 100 | 0.49 | 98.2859 | 5008656 | 45.25 | 62 | 538692 |
| SPVP170041 | SPVP170041 | SAMN20371715 | SRR15240253 | NonPC | Public | 2017 | China | Env | 91 | 100 | 0 | 98.3561 | 5127437 | 45.21 | 44 | 340509 |
| SPVP170042 | SPVP170042 | SAMN20371716 | SRR15240252 | NonPC | Public | 2017 | China | Env | 90.9 | 100 | 0 | 98.356 | 5112140 | 45.2 | 37 | 528811 |
| SPVP170043 | SPVP170043 | SAMN20371717 | SRR15240251 | NonPC | Public | 2017 | China | Env | 91 | 100 | 0 | 98.3285 | 5128185 | 45.21 | 47 | 340509 |
| SPVP170044 | SPVP170044 | SAMN20371709 | SRR15240260 | NonPC | Public | 2017 | China | Env | 91 | 100 | 0.05 | 98.2881 | 4997947 | 45.35 | 53 | 409938 |
| SPVP170045 | SPVP170045 | SAMN20371710 | SRR15240259 | NonPC | Public | 2017 | China | Env | 91.6 | 100 | 0.27 | 98.3481 | 5113788 | 45.25 | 65 | 461739 |
| SPVP170046 | SPVP170046 | SAMN20371711 | SRR15240258 | NonPC | Public | 2017 | China | Env | 88.2 | 100 | 0.36 | 98.2934 | 5076120 | 45.25 | 56 | 382554 |
| SPVP170047 | SPVP170047 | SAMN20371712 | SRR15240257 | NonPC | Public | 2017 | China | Env | 88.2 | 100 | 0.36 | 98.1751 | 5202334 | 45.19 | 122 | 121276 |
| SPVP170048 | SPVP170048 | SAMN20371714 | SRR15240254 | NonPC | Public | 2017 | China | Env | 91.5 | 100 | 0 | 98.3638 | 5048459 | 45.35 | 62 | 448508 |
| SPVP170049 | SPVP170049 | SAMN20371713 | SRR15240256 | NonPC | Public | 2017 | China | Env | 91.3 | 100 | 0.09 | 98.2745 | 5425347 | 45.07 | 56 | 433192 |
| SPVP170050 | SPVP170050 | SAMN20371720 | SRR15240248 | NonPC | Public | 2017 | China | Env | 88.3 | 100 | 0.39 | 98.219 | 5084494 | 45.27 | 54 | 542356 |
| SPVP170051 | SPVP170051 | SAMN20371804 | SRR15240046 | NonPC | Public | 2017 | China | Env | 88.7 | 100 | 0.39 | 98.2305 | 5050705 | 45.24 | 64 | 428306 |
| SPVP170052 | SPVP170052 | SAMN20371805 | SRR15240045 | NonPC | Public | 2017 | China | Env | 91.4 | 100 | 0.14 | 98.3443 | 5263263 | 45.11 | 38 | 335645 |
| SPVP170053 | SPVP170053 | SAMN20371772 | SRR15240082 | NonPC | Public | 2017 | China | Env | 88.2 | 100 | 0.58 | 98.2371 | 5130454 | 45.2 | 82 | 195395 |
| SPVP170054 | SPVP170054 | SAMN20371773 | SRR15240081 | NonPC | Public | 2017 | China | Env | 90.1 | 100 | 0 | 98.3155 | 5088087 | 45.35 | 89 | 175583 |
| SPVP170056 | SPVP170056 | SAMN20371774 | SRR15240079 | NonPC | Public | 2017 | China | Env | 88.2 | 100 | 0.39 | 98.2306 | 4981453 | 45.32 | 36 | 710271 |
| SPVP170060 | SPVP170060 | SAMN20371785 | SRR15240067 | NonPC | Public | 2017 | China | Env | 91.2 | 100 | 0.1 | 98.3727 | 5100442 | 45.23 | 57 | 402138 |
| SPVP170064 | SPVP170064 | SAMN20371781 | SRR15240072 | NonPC | Public | 2017 | China | Env | 88.5 | 100 | 0.6 | 98.2288 | 5119158 | 45.21 | 67 | 560467 |
| SPVP170065 | SPVP170065 | SAMN20371786 | SRR15240066 | NonPC | Public | 2017 | China | Env | 91.4 | 100 | 0.33 | 98.3352 | 5074882 | 45.34 | 45 | 530018 |

| | | | | | | | | | | | | | | | | |
|---|---|---|---|---|---|---|---|---|---|---|---|---|---|---|---|---|
| SPVP170066 | SPVP170066 | SAMN20371787 | SRR15240065 | NonPC | Public | 2017 | China | Env | 91.1 | 100 | 0.15 | 98.322 | 5115961 | 45.22 | 40 | 431265 |
| SPVP170067 | SPVP170067 | SAMN20371788 | SRR15240064 | NonPC | Public | 2017 | China | Env | 88.4 | 100 | 0.39 | 98.1857 | 5086546 | 45.26 | 55 | 558028 |
| SPVP170069 | SPVP170069 | SAMN20371784 | SRR15240068 | NonPC | Public | 2017 | China | Env | 91 | 100 | 0 | 98.3573 | 5072187 | 45.24 | 37 | 413348 |
| SPVP170070 | SPVP170070 | SAMN20371789 | SRR15240063 | NonPC | Public | 2017 | China | Env | 88.8 | 100 | 0.33 | 98.1805 | 5177638 | 45.15 | 61 | 582092 |
| SPVP170071 | SPVP170071 | SAMN20371790 | SRR15240062 | NonPC | Public | 2017 | China | Env | 89.8 | 100 | 0 | 98.2319 | 5273174 | 45.08 | 29 | 779034 |
| SPVP170073 | SPVP170073 | SAMN20371779 | SRR15240074 | NonPC | Public | 2017 | China | Env | 91.7 | 100 | 0.05 | 98.3701 | 5222065 | 44.99 | 32 | 526918 |
| SPVP170074 | SPVP170074 | SAMN20371791 | SRR15240061 | NonPC | Public | 2017 | China | Env | 88.3 | 100 | 0.3 | 98.2229 | 5019152 | 45.35 | 51 | 396622 |
| SPVP170076 | SPVP170076 | SAMN20371783 | SRR15240070 | NonPC | Public | 2017 | China | Env | 91 | 100 | 0 | 98.3766 | 5018706 | 45.31 | 95 | 247858 |
| SPVP170078 | SPVP170078 | SAMN20371800 | SRR15240051 | NonPC | Public | 2017 | China | Env | 92.2 | 100 | 0.12 | 98.3566 | 5076722 | 45.28 | 49 | 418347 |
| SPVP170079 | SPVP170079 | SAMN20371796 | SRR15240055 | NonPC | Public | 2017 | China | Env | 88.6 | 100 | 0.42 | 98.212 | 5099129 | 45.22 | 91 | 546332 |
| SPVP170080 | SPVP170080 | SAMN20371792 | SRR15240060 | NonPC | Public | 2017 | China | Env | 91.2 | 100 | 0 | 98.3084 | 5339865 | 45.35 | 86 | 400930 |
| SPVP170081 | SPVP170081 | SAMN20371798 | SRR15240053 | NonPC | Public | 2017 | China | Env | 88.4 | 100 | 0.36 | 98.2489 | 5093347 | 45.28 | 61 | 561826 |
| SPVP170083 | SPVP170083 | SAMN20371793 | SRR15240059 | NonPC | Public | 2017 | China | Env | 88.4 | 100 | 0.34 | 98.2441 | 5104091 | 45.28 | 68 | 452139 |
| SPVP170084 | SPVP170084 | SAMN20371799 | SRR15240052 | NonPC | Public | 2017 | China | Env | 88.5 | 100 | 0.36 | 98.2273 | 5094500 | 45.28 | 65 | 561826 |
| SPVP170085 | SPVP170085 | SAMN20371782 | SRR15240071 | NonPC | Public | 2017 | China | Env | 88.4 | 100 | 0.39 | 98.2011 | 5097665 | 45.17 | 74 | 442668 |
| SPVP170087 | SPVP170087 | SAMN20371794 | SRR15240057 | NonPC | Public | 2017 | China | Env | 91.1 | 100 | 0.27 | 98.3801 | 4990992 | 45.34 | 68 | 239043 |
| SPVP170089 | SPVP170089 | SAMN20371775 | SRR15240078 | NonPC | Public | 2017 | China | Env | 88.3 | 100 | 0.39 | 98.2095 | 5084752 | 45.28 | 64 | 453037 |
| SPVP170090 | SPVP170090 | SAMN20371770 | SRR15240084 | NonPC | Public | 2017 | China | Env | 91.7 | 100 | 0 | 98.2823 | 5116438 | 45.29 | 53 | 516964 |
| SPVP170091 | SPVP170091 | SAMN20371778 | SRR15240075 | NonPC | Public | 2017 | China | Env | 92.2 | 100 | 0.26 | 98.36 | 5249724 | 45.1 | 50 | 418476 |
| SPVP170092 | SPVP170092 | SAMN20371771 | SRR15240083 | NonPC | Public | 2017 | China | Env | 87.4 | 99.73 | 0.37 | 98.3441 | 5006268 | 45.33 | 64 | 203415 |
| SPVP170093 | SPVP170093 | SAMN20371776 | SRR15240077 | NonPC | Public | 2017 | China | Env | 88.3 | 100 | 0.46 | 98.2483 | 5118356 | 45.2 | 62 | 316430 |
| SPVP170094 | SPVP170094 | SAMN20371777 | SRR15240076 | NonPC | Public | 2017 | China | Env | 89 | 100 | 0.33 | 98.2226 | 4995291 | 45.36 | 72 | 381107 |
| SPVP170095 | SPVP170095 | SAMN20371780 | SRR15240073 | NonPC | Public | 2017 | China | Env | 88.8 | 100 | 0.39 | 98.2198 | 5255407 | 45.11 | 66 | 425608 |
| SPVP170098 | SPVP170098 | SAMN20371795 | SRR15240056 | NonPC | Public | 2017 | China | Env | 88.3 | 100 | 0.33 | 98.1906 | 5109305 | 45.21 | 84 | 506900 |
| SPVP170102 | SPVP170102 | SAMN20371797 | SRR15240054 | NonPC | Public | 2017 | China | Env | 88.1 | 100 | 0.3 | 98.235 | 5143273 | 45.25 | 33 | 746570 |
| SPVP170106 | SPVP170106 | SAMN20371811 | SRR15240039 | NonPC | Public | 2017 | China | Env | 88.3 | 100 | 0.36 | 98.27 | 5032786 | 45.27 | 46 | 554090 |
| SPVP170141 | SPVP170141 | SAMN20371900 | SRR15239899 | NonPC | Public | 2017 | China | Env | 89.9 | 100 | 0.05 | 98.3163 | 5018570 | 45.31 | 72 | 389440 |
| SPVP170142 | SPVP170142 | SAMN20371899 | SRR15239900 | NonPC | Public | 2017 | China | Env | 88.6 | 100 | 0.39 | 98.2511 | 4973646 | 45.37 | 68 | 741114 |
| SPVP170143 | SPVP170143 | SAMN20371901 | SRR15239898 | NonPC | Public | 2017 | China | Env | 90 | 100 | 0.05 | 98.3124 | 5048464 | 45.36 | 47 | 459840 |
| SPVP170146 | SPVP170146 | SAMN20371902 | SRR15239897 | NonPC | Public | 2017 | China | Env | 89.4 | 100 | 0.12 | 98.2426 | 5250461 | 45.19 | 82 | 320620 |
| SPVP170147 | SPVP170147 | SAMN20371903 | SRR15239896 | NonPC | Public | 2017 | China | Env | 88.8 | 100 | 0 | 98.3471 | 5096603 | 45.42 | 90 | 151704 |
| SPVP170148 | SPVP170148 | SAMN20371898 | SRR15239901 | NonPC | Public | 2017 | China | Env | 88.3 | 100 | 0.47 | 98.2491 | 5037966 | 45.22 | 57 | 452368 |
| SPVP170149 | SPVP170149 | SAMN20371904 | SRR15239894 | NonPC | Public | 2017 | China | Env | 89.4 | 100 | 0.03 | 98.3262 | 4944448 | 45.35 | 45 | 380366 |
| SPVP170150 | SPVP170150 | SAMN20371897 | SRR15239902 | NonPC | Public | 2017 | China | Env | 90.1 | 100 | 0.05 | 98.2794 | 5145042 | 45.31 | 41 | 375562 |
| SPVP170151 | SPVP170151 | SAMN20371895 | SRR15239904 | NonPC | Public | 2017 | China | Env | 91.3 | 100 | 0 | 98.3869 | 5135798 | 45.31 | 33 | 334802 |
| SPVP170152 | SPVP170152 | SAMN20371887 | SRR15239913 | NonPC | Public | 2017 | China | Env | 90.9 | 100 | 0 | 98.3145 | 5071081 | 45.29 | 61 | 339583 |
| SPVP170153 | SPVP170153 | SAMN20371886 | SRR15239914 | NonPC | Public | 2017 | China | Env | 91.8 | 100 | 0.01 | 98.2805 | 5270806 | 45.24 | 62 | 462106 |
| SPVP170154 | SPVP170154 | SAMN20371888 | SRR15239912 | NonPC | Public | 2017 | China | Env | 90.3 | 100 | 0.05 | 98.2803 | 5002022 | 45.33 | 83 | 540356 |
| SPVP170155 | SPVP170155 | SAMN20371889 | SRR15239911 | NonPC | Public | 2017 | China | Env | 90 | 100 | 0.11 | 98.2657 | 5079551 | 45.3 | 54 | 460593 |
| SPVP170156 | SPVP170156 | SAMN20371892 | SRR15239908 | NonPC | Public | 2017 | China | Env | 90.5 | 100 | 0 | 98.3226 | 5186667 | 45.22 | 38 | 573368 |
| SPVP170157 | SPVP170157 | SAMN20371893 | SRR15239907 | NonPC | Public | 2017 | China | Env | 88.4 | 100 | 0.36 | 98.2038 | 5096888 | 45.23 | 62 | 464169 |
| SPVP170158 | SPVP170158 | SAMN20371894 | SRR15239905 | NonPC | Public | 2017 | China | Env | 88.4 | 100 | 0.36 | 98.2092 | 5096243 | 45.23 | 61 | 548679 |
| SPVP170159 | SPVP170159 | SAMN20371884 | SRR15239921 | NonPC | Public | 2017 | China | Env | 91.3 | 100 | 0.1 | 98.3473 | 5197015 | 45.21 | 64 | 401040 |
| SPVP170160 | SPVP170160 | SAMN20371890 | SRR15239910 | NonPC | Public | 2017 | China | Env | 90.9 | 100 | 0 | 98.3429 | 5079390 | 45.27 | 37 | 407867 |
| SPVP170161 | SPVP170161 | SAMN20371885 | SRR15239916 | NonPC | Public | 2017 | China | Env | 88.7 | 100 | 0.42 | 98.2279 | 5086253 | 45.25 | 53 | 722589 |
| SPVP170162 | SPVP170162 | SAMN20371891 | SRR15239909 | NonPC | Public | 2017 | China | Env | 88.3 | 100 | 0.41 | 98.2435 | 5137603 | 45.19 | 52 | 469954 |
| SPVP170163 | SPVP170163 | SAMN20371905 | SRR15239893 | NonPC | Public | 2017 | China | Env | 88.4 | 100 | 0 | 98.4078 | 4932562 | 45.31 | 62 | 175141 |
| SPVP170164 | SPVP170164 | SAMN20371896 | SRR15239903 | NonPC | Public | 2017 | China | Env | 88.6 | 100 | 0.33 | 98.2661 | 5105080 | 45.22 | 52 | 455152 |

| | | | | | | | | | | | | | | | | |
|---|---|---|---|---|---|---|---|---|---|---|---|---|---|---|---|---|
| SPVP180001 | SPVP180001 | SAMN20371946 | SRR15239847 | NonPC | Public | 2018 | China | Env | 88.9 | 100 | 0 | 98.3042 | 5175773 | 45.22 | 53 | 333124 |
| SPVP180002 | SPVP180002 | SAMN20371945 | SRR15239848 | NonPC | Public | 2018 | China | Env | 87.6 | 100 | 0.63 | 98.2397 | 5002409 | 45.34 | 50 | 449519 |
| SPVP180003 | SPVP180003 | SAMN20371966 | SRR15239825 | NonPC | Public | 2018 | China | Env | 88.1 | 99.73 | 0.54 | 98.2156 | 5158711 | 45.11 | 62 | 368073 |
| SPVP180004 | SPVP180004 | SAMN20371967 | SRR15239824 | NonPC | Public | 2018 | China | Env | 88.3 | 100 | 0.63 | 98.188 | 5205237 | 45.24 | 47 | 314967 |
| SPVP180007 | SPVP180007 | SAMN20371968 | SRR15239823 | NonPC | Public | 2018 | China | Env | 88.2 | 100 | 0.42 | 98.2779 | 5065676 | 45.27 | 38 | 427720 |
| SPVP180009 | SPVP180009 | SAMN20371964 | SRR15239827 | NonPC | Public | 2018 | China | Env | 91.1 | 100 | 0.27 | 98.2773 | 5198947 | 45.32 | 96 | 386361 |
| SPVP180011 | SPVP180011 | SAMN20371969 | SRR15239822 | NonPC | Public | 2018 | China | Env | 91.9 | 100 | 0 | 98.3518 | 5060904 | 45.29 | 34 | 460366 |
| SPVP180012 | SPVP180012 | SAMN20371970 | SRR15239821 | NonPC | Public | 2018 | China | Env | 91.4 | 100 | 0.27 | 98.3887 | 5027486 | 45.35 | 62 | 254246 |
| SPVP180015 | SPVP180015 | SAMN20371971 | SRR15239820 | NonPC | Public | 2018 | China | Env | 88.1 | 100 | 0.3 | 98.2287 | 5138584 | 45.25 | 37 | 538312 |
| SPVP180016 | SPVP180016 | SAMN20371963 | SRR15239829 | NonPC | Public | 2018 | China | Env | 91.5 | 100 | 0 | 98.2839 | 5071746 | 45.27 | 51 | 458160 |
| SPVP180017 | SPVP180017 | SAMN20371965 | SRR15239826 | NonPC | Public | 2018 | China | Env | 91 | 100 | 0 | 98.3382 | 5083237 | 45.25 | 36 | 528921 |
| SPVP180018 | SPVP180018 | SAMN20371986 | SRR15239803 | NonPC | Public | 2018 | China | Env | 91.1 | 100 | 0 | 98.3152 | 5054140 | 45.24 | 56 | 264003 |
| SPVP180019 | SPVP180019 | SAMN20371988 | SRR15239801 | NonPC | Public | 2018 | China | Env | 91.4 | 100 | 0 | 98.3282 | 5265279 | 45.3 | 48 | 404877 |
| SPVP180020 | SPVP180020 | SAMN20371987 | SRR15239802 | NonPC | Public | 2018 | China | Env | 91.3 | 100 | 0 | 98.3239 | 5052725 | 45.35 | 56 | 461891 |
| SPVP180021 | SPVP180021 | SAMN20372027 | SRR15239754 | NonPC | Public | 2018 | China | Env | 90.9 | 100 | 0.67 | 98.2906 | 5331838 | 45.16 | 88 | 437865 |
| SPVP180022 | SPVP180022 | SAMN20372026 | SRR15239755 | NonPC | Public | 2018 | China | Env | 87.6 | 100 | 0.45 | 98.2455 | 5247090 | 45.07 | 94 | 451005 |
| SPVP180024 | SPVP180024 | SAMN20372024 | SRR15239757 | NonPC | Public | 2018 | China | Env | 88.5 | 100 | 0.3 | 98.2503 | 5021760 | 45.28 | 70 | 735052 |
| SPVP180025 | SPVP180025 | SAMN20372025 | SRR15239756 | NonPC | Public | 2018 | China | Env | 88.2 | 100 | 0.39 | 98.1959 | 5260533 | 45.12 | 65 | 568264 |
| SPVP180026 | SPVP180026 | SAMN20372028 | SRR15239753 | NonPC | Public | 2018 | China | Env | 88.3 | 100 | 0.33 | 98.2336 | 5081435 | 45.24 | 46 | 455543 |
| SPVP180036 | SPVP180036 | SAMN20372156 | SRR15240115 | NonPC | Public | 2018 | China | Env | 91.6 | 100 | 0 | 98.3623 | 5017941 | 45.32 | 26 | 487127 |
| SPVP20150002 | SPVP20150002 | SAMN20353014 | SRR15218726 | NonPC | Public | 2015 | China | Env | 89.9 | 100 | 0 | 98.3419 | 5076309 | 45.16 | 51 | 367652 |
| SPVP20150003 | SPVP20150003 | SAMN20353015 | SRR15218725 | NonPC | Public | 2015 | China | Env | 89.8 | 100 | 0 | 98.3623 | 5157898 | 45.36 | 32 | 510103 |
| SPVP20150004 | SPVP20150004 | SAMN20353016 | SRR15218724 | NonPC | Public | 2015 | China | Env | 91.9 | 100 | 0.05 | 98.347 | 5040389 | 45.37 | 52 | 375716 |
| SPVP20150005 | SPVP20150005 | SAMN20353017 | SRR15218723 | NonPC | Public | 2015 | China | Env | 91.5 | 100 | 0.1 | 98.3584 | 5123740 | 45.23 | 47 | 546270 |
| SPVP20150006 | SPVP20150006 | SAMN20353008 | SRR15218733 | NonPC | Public | 2015 | China | Env | 88.3 | 100 | 0.3 | 98.2296 | 5010541 | 45.27 | 35 | 539783 |
| SPVP20150009 | SPVP20150009 | SAMN20353018 | SRR15218689 | NonPC | Public | 2015 | China | Env | 91.4 | 100 | 0.36 | 98.3528 | 5064092 | 45.34 | 55 | 463444 |
| Vp_Peru_CFSAN018757_1997_Peru_C | CFSAN018757 | SAMN03941065 | GCF_001696015.1 | PC | Public | 1997 | Peru | Clinical | 98.5 | 100 | 0 | 99.987 | 5099401 | 45.29 | 72 | 288855 |
| Vp_Peru_GCA_000182345.1_ASM18234v1_NA_Peru_Na | Peru-466 | SAMN02436220 | GCA_000182345.1 | PC | Public | NA | Peru | NA | 97.2 | 95.47 | 0.61 | 99.9886 | 5037488 | 45.36 | 149 | 81497 |
| VP02003 | VP02003 | SAMN20167697 | SRR15099227 | PC | Public | 2002 | China | Clinical | 98.4 | 100 | 0 | 99.9881 | 5121607 | 45.28 | 65 | 260144 |
| VP02004 | VP02004 | SAMN20167698 | SRR15099226 | PC | Public | 2002 | China | Clinical | 98.5 | 100 | 0 | 99.9873 | 5132862 | 45.29 | 61 | 401474 |
| VP03011 | VP03011 | SAMN20167699 | SRR15099115 | PC | Public | 2003 | China | Clinical | 98.1 | 100 | 0 | 99.9221 | 5104027 | 45.29 | 62 | 440646 |
| VP03013 | VP03013 | SAMN20167706 | SRR15099715 | PC | Public | 2003 | China | Clinical | 98.1 | 100 | 0 | 99.9129 | 5109942 | 45.29 | 60 | 401619 |
| VP03014 | VP03014 | SAMN20167707 | SRR15099225 | PC | Public | 2003 | China | Clinical | 98.1 | 100 | 0 | 99.9346 | 5109780 | 45.29 | 66 | 373635 |
| VP03015 | VP03015 | SAMN20167700 | SRR15099004 | PC | Public | 2003 | China | Clinical | 98 | 100 | 0 | 99.9267 | 5109220 | 45.28 | 67 | 312854 |
| VP03021 | VP03021 | SAMN20167701 | SRR15098893 | PC | Public | 2003 | China | Clinical | 98 | 100 | 0 | 99.9249 | 5109069 | 45.28 | 64 | 373648 |
| VP03022 | VP03022 | SAMN20167702 | SRR15099538 | PC | Public | 2003 | China | Clinical | 98.1 | 100 | 0 | 99.9495 | 5109163 | 45.28 | 67 | 373647 |
| VP03023 | VP03023 | SAMN20167703 | SRR15099427 | PC | Public | 2003 | China | Clinical | 97.4 | 99.73 | 0 | 99.9001 | 5072873 | 45.31 | 57 | 448992 |
| VP03028 | VP03028 | SAMN20167704 | SRR15099316 | PC | Public | 2003 | China | Clinical | 98.1 | 100 | 0 | 99.9263 | 5110163 | 45.29 | 66 | 288859 |
| VP03029 | VP03029 | SAMN20167705 | SRR15099826 | PC | Public | 2003 | China | Clinical | 97.6 | 100 | 0 | 99.9323 | 5079858 | 45.28 | 70 | 288853 |
| VP03049 | VP03049 | SAMN20167708 | SRR15099214 | PC | Public | 2003 | China | Clinical | 98.3 | 99.95 | 0 | 99.9849 | 5114486 | 45.29 | 66 | 288662 |
| VP03050 | VP03050 | SAMN20167709 | SRR15099203 | PC | Public | 2003 | China | Clinical | 98.5 | 99.73 | 0 | 99.9752 | 5128635 | 45.29 | 57 | 431658 |
| VP03256 | VP03256 | SAMN20167754 | SRR15099460 | PC | Public | 2005 | China | Clinical | 97.6 | 100 | 0 | 99.8954 | 5165411 | 45.28 | 49 | 401604 |
| VP03257 | VP03257 | SAMN20167755 | SRR15099449 | PC | Public | 2005 | China | Clinical | 97.6 | 100 | 0 | 99.8944 | 5164171 | 45.28 | 48 | 477837 |
| VP03259 | VP03259 | SAMN20167756 | SRR15099438 | PC | Public | 2005 | China | Clinical | 97.5 | 100 | 0 | 99.8977 | 5162704 | 45.28 | 51 | 461740 |
| VP03261 | VP03261 | SAMN20167757 | SRR15099426 | PC | Public | 2005 | China | Clinical | 97.6 | 100 | 0.07 | 99.9081 | 5192075 | 45.31 | 57 | 461740 |
| VP03285 | VP03285 | SAMN20167752 | SRR15099482 | NonPC | Public | 2005 | China | Env | 91.2 | 100 | 0.05 | 98.3689 | 5135549 | 45.23 | 44 | 361938 |
| VP03287 | VP03287 | SAMN20167753 | SRR15099471 | NonPC | Public | 2005 | China | Env | 91.2 | 100 | 0.27 | 98.3568 | 5000714 | 45.35 | 60 | 458805 |
| VP04076 | VP04076 | SAMN20167715 | SRR15099137 | PC | Public | 2004 | China | Clinical | 98.3 | 100 | 0 | 99.993 | 5162007 | 45.11 | 68 | 288661 |

| | | | | | | | | | | | | | | | | |
|---|---|---|---|---|---|---|---|---|---|---|---|---|---|---|---|---|
| VP04078 | VP04078 | SAMN20167716 | SRR15099126 | PC | Public | 2004 | China | Clinical | 98.4 | 100 | 0 | 99.9855 | 5171532 | 45.12 | 65 | 425676 |
| VP04079 | VP04079 | SAMN20167717 | SRR15099114 | PC | Public | 2004 | China | Clinical | 98.3 | 100 | 0 | 99.9931 | 5088598 | 45.28 | 66 | 425763 |
| VP04080 | VP04080 | SAMN20167718 | SRR15099103 | PC | Public | 2004 | China | Clinical | 98.5 | 100 | 0 | 99.9938 | 5172850 | 45.12 | 69 | 288859 |
| VP04081 | VP04081 | SAMN20167719 | SRR15099092 | NonPC | Public | 2004 | China | Clinical | 92.2 | 100 | 0.19 | 98.3643 | 5011814 | 45.25 | 51 | 450397 |
| VP04082 | VP04082 | SAMN20167720 | SRR15099081 | PC | Public | 2004 | China | Clinical | 98.3 | 100 | 0.01 | 99.9789 | 5249898 | 45.14 | 77 | 266045 |
| VP04083 | VP04083 | SAMN20167721 | SRR15099070 | PC | Public | 2004 | China | Clinical | 98.2 | 99.46 | 0 | 99.9931 | 5083830 | 45.29 | 60 | 288661 |
| VP04086 | VP04086 | SAMN20167722 | SRR15099059 | PC | Public | 2004 | China | Clinical | 98.3 | 100 | 0.01 | 99.9746 | 5185815 | 45.33 | 61 | 411185 |
| VP04094 | VP04094 | SAMN20167728 | SRR15098992 | NonPC | Public | 2004 | China | Clinical | 90.9 | 100 | 0 | 98.3043 | 5119277 | 45.31 | 54 | 374815 |
| VP04097 | VP04097 | SAMN20167723 | SRR15099048 | PC | Public | 2004 | China | Clinical | 98.3 | 100 | 0 | 99.9889 | 5163553 | 45.12 | 75 | 288661 |
| VP04098 | VP04098 | SAMN20167727 | SRR15099003 | NonPC | Public | 2004 | China | Clinical | 92.9 | 100 | 0.14 | 98.3565 | 5385496 | 45.07 | 56 | 281396 |
| VP04099 | VP04099 | SAMN20167724 | SRR15099037 | PC | Public | 2004 | China | Clinical | 98.2 | 100 | 0 | 99.9649 | 5212988 | 45.11 | 72 | 312799 |
| VP04101 | VP04101 | SAMN20167725 | SRR15099026 | PC | Public | 2004 | China | Clinical | 97.8 | 99.51 | 0 | 99.9202 | 5091968 | 45.29 | 63 | 370760 |
| VP04102 | VP04102 | SAMN20167726 | SRR15099015 | PC | Public | 2004 | China | Clinical | 97.9 | 100 | 0 | 99.9341 | 5094715 | 45.29 | 65 | 427693 |
| VP04106 | VP04106 | SAMN20167714 | SRR15099148 | NonPC | Public | 2004 | China | Clinical | 88.5 | 100 | 0.36 | 98.2271 | 5033216 | 45.28 | 38 | 548186 |
| VP04108 | VP04108 | SAMN20167729 | SRR15098981 | PC | Public | 2004 | China | Clinical | 98.3 | 100 | 0 | 99.9893 | 5162843 | 45.12 | 72 | 288682 |
| VP04109 | VP04109 | SAMN20167730 | SRR15098970 | NonPC | Public | 2004 | China | Clinical | 92.9 | 100 | 0 | 98.4249 | 5172695 | 45.19 | 35 | 520341 |
| VP04110 | VP04110 | SAMN20167731 | SRR15098959 | PC | Public | 2004 | China | Clinical | 97.8 | 100 | 0 | 99.9477 | 5125131 | 45.3 | 65 | 289369 |
| VP04112 | VP04112 | SAMN20167732 | SRR15098948 | PC | Public | 2004 | China | Clinical | 98.1 | 100 | 0 | 99.9201 | 5141768 | 45.28 | 69 | 288859 |
| VP04113 | VP04113 | SAMN20167733 | SRR15098937 | PC | Public | 2004 | China | Clinical | 98.1 | 100 | 0 | 99.9392 | 5150189 | 45.29 | 72 | 289370 |
| VP04114 | VP04114 | SAMN20167734 | SRR15098926 | NonPC | Public | 2004 | China | Clinical | 90.8 | 100 | 0 | 98.2972 | 5142578 | 45.27 | 77 | 395731 |
| VP04116 | VP04116 | SAMN20167735 | SRR15098915 | PC | Public | 2004 | China | Clinical | 98.1 | 100 | 0.14 | 99.9239 | 5352838 | 45.12 | 80 | 288871 |
| VP04117 | VP04117 | SAMN20167736 | SRR15098904 | PC | Public | 2004 | China | Clinical | 98 | 100 | 0 | 99.9399 | 5146873 | 45.28 | 76 | 288859 |
| VP04120 | VP04120 | SAMN20167737 | SRR15098892 | PC | Public | 2004 | China | Clinical | 98 | 100 | 0 | 99.9374 | 5148791 | 45.28 | 71 | 427739 |
| VP04121 | VP04121 | SAMN20167738 | SRR15098881 | PC | Public | 2004 | China | Clinical | 97.7 | 100 | 0 | 99.9158 | 5111507 | 45.29 | 72 | 427716 |
| VP04123 | VP04123 | SAMN20167740 | SRR15098859 | PC | Public | 2004 | China | Clinical | 98.2 | 100 | 0 | 99.9899 | 5087348 | 45.3 | 69 | 288661 |
| VP04124 | VP04124 | SAMN20167741 | SRR15099604 | PC | Public | 2004 | China | Clinical | 98.2 | 99.91 | 0 | 99.9867 | 5088450 | 45.29 | 72 | 268169 |
| VP04125 | VP04125 | SAMN20167742 | SRR15099593 | PC | Public | 2004 | China | Clinical | 98.3 | 100 | 0 | 99.9912 | 5096492 | 45.3 | 66 | 302694 |
| VP04126 | VP04126 | SAMN20167743 | SRR15099582 | PC | Public | 2004 | China | Clinical | 97.9 | 100 | 0 | 99.9232 | 5145181 | 45.25 | 68 | 373487 |
| VP04127 | VP04127 | SAMN20167744 | SRR15099571 | PC | Public | 2004 | China | Clinical | 98 | 100 | 0 | 99.9229 | 5136895 | 45.25 | 57 | 394961 |
| VP04128 | VP04128 | SAMN20167748 | SRR15099526 | NonPC | Public | 2004 | China | Clinical | 90.9 | 100 | 0 | 98.295 | 5142084 | 45.26 | 68 | 405040 |
| VP04129 | VP04129 | SAMN20167745 | SRR15099560 | PC | Public | 2004 | China | Clinical | 98.3 | 100 | 0 | 99.9896 | 5164332 | 45.12 | 72 | 288660 |
| VP04130 | VP04130 | SAMN20167746 | SRR15099549 | PC | Public | 2004 | China | Clinical | 98.1 | 100 | 0 | 99.9931 | 5091238 | 45.3 | 69 | 288661 |
| VP04131 | VP04131 | SAMN20167747 | SRR15099537 | PC | Public | 2004 | China | Clinical | 98.1 | 100 | 0 | 99.9922 | 5085541 | 45.3 | 66 | 312650 |
| VP04132 | VP04132 | SAMN20167749 | SRR15099515 | NonPC | Public | 2004 | China | Clinical | 90.9 | 100 | 0 | 98.2708 | 5136076 | 45.27 | 59 | 395865 |
| VP04137 | VP04137 | SAMN20167739 | SRR15098870 | NonPC | Public | 2004 | China | Clinical | 91.9 | 100 | 0.19 | 98.3226 | 5008850 | 45.26 | 46 | 539190 |
| VP04138 | VP04138 | SAMN20167710 | SRR15099192 | NonPC | Public | 2004 | China | Clinical | 92.3 | 100 | 0.19 | 98.355 | 5011777 | 45.25 | 53 | 450398 |
| VP04141 | VP04141 | SAMN20167711 | SRR15099181 | PC | Public | 2004 | China | Clinical | 98.4 | 100 | 0 | 99.9725 | 5120151 | 45.28 | 70 | 288661 |
| VP04158 | VP04158 | SAMN20167713 | SRR15099159 | NonPC | Public | 2004 | China | Clinical | 90.4 | 99.95 | 0.27 | 98.2875 | 5104509 | 45.27 | 65 | 313948 |
| VP04159 | VP04159 | SAMN20167712 | SRR15099170 | PC | Public | 2004 | China | Clinical | 98.3 | 100 | 0 | 99.974 | 5170419 | 45.27 | 68 | 312799 |
| VP05213 | VP05213 | SAMN20167758 | SRR15099415 | PC | Public | 2005 | China | Clinical | 98.2 | 100 | 0 | 99.9908 | 5156765 | 45.11 | 71 | 265933 |
| VP05215 | VP05215 | SAMN20167759 | SRR15099404 | PC | Public | 2005 | China | Clinical | 98.2 | 100 | 0 | 99.9728 | 5157040 | 45.11 | 74 | 312653 |
| VP05217 | VP05217 | SAMN20167760 | SRR15099393 | PC | Public | 2005 | China | Clinical | 98.2 | 99.93 | 0 | 99.9872 | 5158057 | 45.11 | 68 | 312660 |
| VP05220 | VP05220 | SAMN20167761 | SRR15099382 | PC | Public | 2005 | China | Clinical | 98.4 | 99.95 | 0 | 99.987 | 5119584 | 45.28 | 70 | 288661 |
| VP05223 | VP05223 | SAMN20167762 | SRR15099371 | PC | Public | 2005 | China | Clinical | 98.5 | 100 | 0 | 99.9925 | 5122541 | 45.28 | 68 | 288661 |
| VP05229 | VP05229 | SAMN20167764 | SRR15099349 | NonPC | Public | 2005 | China | Clinical | 92.4 | 100 | 0.05 | 98.3875 | 5030640 | 45.32 | 47 | 450227 |
| VP05234 | VP05234 | SAMN20167763 | SRR15099360 | PC | Public | 2005 | China | Clinical | 97.6 | 100 | 0 | 99.9902 | 5056977 | 45.31 | 66 | 312671 |
| VP05239 | VP05239 | SAMN20167765 | SRR15099338 | PC | Public | 2005 | China | Clinical | 98.5 | 99.93 | 0 | 99.9829 | 5100874 | 45.28 | 65 | 288661 |
| VP05242 | VP05242 | SAMN20167766 | SRR15099327 | PC | Public | 2005 | China | Clinical | 98.2 | 99.67 | 0 | 99.97 | 5085741 | 45.29 | 62 | 288576 |

| | | | | | | | | | | | | | | | | |
|---|---|---|---|---|---|---|---|---|---|---|---|---|---|---|---|---|
| VP05244 | VP05244 | SAMN20167767 | SRR15099315 | PC | Public | 2005 | China | Clinical | 98.5 | 100 | 0 | 99.9902 | 5101965 | 45.28 | 66 | 312808 |
| VP05245 | VP05245 | SAMN20167768 | SRR15099304 | PC | Public | 2005 | China | Clinical | 98.3 | 100 | 0 | 99.9897 | 5091229 | 45.27 | 67 | 312806 |
| VP05246 | VP05246 | SAMN20167769 | SRR15099293 | PC | Public | 2005 | China | Clinical | 98.4 | 100 | 0 | 99.988 | 5100963 | 45.28 | 67 | 288682 |
| VP05255 | VP05255 | SAMN20167770 | SRR15099282 | PC | Public | 2005 | China | Clinical | 97.5 | 100 | 0 | 99.902 | 5164202 | 45.28 | 60 | 288862 |
| VP05258 | VP05258 | SAMN20167771 | SRR15099271 | PC | Public | 2005 | China | Clinical | 97.6 | 100 | 0 | 99.8897 | 5164293 | 45.28 | 56 | 318037 |
| VP05260 | VP05260 | SAMN20167772 | SRR15099260 | PC | Public | 2005 | China | Clinical | 97.6 | 100 | 0 | 99.8997 | 5164421 | 45.28 | 55 | 318008 |
| VP05265 | VP05265 | SAMN20167773 | SRR15099249 | PC | Public | 2005 | China | Clinical | 97.6 | 100 | 0 | 99.9052 | 5166730 | 45.28 | 55 | 318122 |
| VP05272 | VP05272 | SAMN20167774 | SRR15099238 | NonPC | Public | 2005 | China | Clinical | 92.4 | 100 | 0.05 | 98.3271 | 5040056 | 45.26 | 45 | 539070 |
| VP05274 | VP05274 | SAMN20167775 | SRR15099848 | NonPC | Public | 2005 | China | Clinical | 92.4 | 100 | 0.05 | 98.3592 | 5047595 | 45.25 | 47 | 539144 |
| VP05276 | VP05276 | SAMN20167776 | SRR15099837 | NonPC | Public | 2005 | China | Clinical | 92.5 | 100 | 0.05 | 98.3463 | 5047175 | 45.25 | 51 | 478722 |
| VP05277 | VP05277 | SAMN20167777 | SRR15099825 | PC | Public | 2005 | China | Clinical | 98.5 | 99.93 | 0 | 99.9928 | 5122036 | 45.28 | 65 | 312792 |
| VP05284 | VP05284 | SAMN20167751 | SRR15099493 | NonPC | Public | 2005 | China | Env | 89.8 | 100 | 0.33 | 98.3342 | 5068099 | 45.29 | 28 | 565172 |
| VP05286 | VP05286 | SAMN20167750 | SRR15099504 | NonPC | Public | 2005 | China | Env | 91.3 | 100 | 0 | 98.3582 | 5000624 | 45.43 | 40 | 439281 |
| VP06001 | VP06001 | SAMN20167778 | SRR15099814 | NonPC | Public | 2006 | China | Env | 91.7 | 100 | 0.14 | 98.3169 | 5089222 | 45.27 | 57 | 458165 |
| VP06002 | VP06002 | SAMN20167784 | SRR15099748 | NonPC | Public | 2006 | China | Env | 90.7 | 100 | 0.05 | 98.2905 | 5022496 | 45.31 | 66 | 411920 |
| VP06003 | VP06003 | SAMN20167803 | SRR15099218 | PC | Public | 2006 | China | Clinical | 98 | 100 | 0 | 99.9411 | 5104709 | 45.28 | 57 | 427704 |
| VP06004 | VP06004 | SAMN20167804 | SRR15099217 | PC | Public | 2006 | China | Clinical | 98 | 100 | 0 | 99.9277 | 5105675 | 45.28 | 57 | 394930 |
| VP06005 | VP06005 | SAMN20167805 | SRR15099216 | PC | Public | 2006 | China | Clinical | 98 | 100 | 0 | 99.9505 | 5104498 | 45.28 | 58 | 288859 |
| VP06006 | VP06006 | SAMN20167806 | SRR15099215 | PC | Public | 2006 | China | Clinical | 98 | 100 | 0 | 99.9137 | 5111424 | 45.26 | 60 | 373649 |
| VP06007 | VP06007 | SAMN20167807 | SRR15099213 | PC | Public | 2006 | China | Clinical | 98 | 100 | 0 | 99.9338 | 5111975 | 45.26 | 56 | 401488 |
| VP06008 | VP06008 | SAMN20167808 | SRR15099212 | PC | Public | 2006 | China | Clinical | 98 | 100 | 0 | 99.9373 | 5113131 | 45.27 | 55 | 401632 |
| VP06009 | VP06009 | SAMN20167800 | SRR15099221 | NonPC | Public | 2006 | China | Clinical | 89.6 | 100 | 0.05 | 98.3567 | 5016325 | 45.33 | 56 | 463810 |
| VP06010 | VP06010 | SAMN20167779 | SRR15099803 | NonPC | Public | 2006 | China | Env | 91.2 | 100 | 0.08 | 98.3529 | 5023477 | 45.35 | 41 | 423489 |
| VP06011 | VP06011 | SAMN20167801 | SRR15099220 | PC | Public | 2006 | China | Env | 98.2 | 99.18 | 0 | 99.9913 | 5081648 | 45.3 | 59 | 395132 |
| VP06012 | VP06012 | SAMN20167802 | SRR15099219 | PC | Public | 2006 | China | Env | 98.5 | 100 | 0 | 99.9833 | 5092435 | 45.29 | 59 | 394972 |
| VP06013 | VP06013 | SAMN20167812 | SRR15099208 | PC | Public | 2006 | China | Env | 98.6 | 100 | 0 | 99.9926 | 5132381 | 45.29 | 59 | 427746 |
| VP06014 | VP06014 | SAMN20167809 | SRR15099211 | PC | Public | 2006 | China | Clinical | 98.7 | 100 | 0 | 99.9937 | 5132278 | 45.29 | 59 | 334357 |
| VP06015 | VP06015 | SAMN20167810 | SRR15099210 | PC | Public | 2006 | China | Clinical | 98.7 | 100 | 0 | 99.9854 | 5133454 | 45.29 | 63 | 334357 |
| VP06016 | VP06016 | SAMN20167811 | SRR15099209 | PC | Public | 2006 | China | Clinical | 98.6 | 100 | 0 | 99.9838 | 5132643 | 45.29 | 61 | 394877 |
| VP06017 | VP06017 | SAMN20167780 | SRR15099792 | NonPC | Public | 2006 | China | Env | 90.6 | 100 | 0 | 98.3358 | 5025050 | 45.33 | 55 | 236586 |
| VP06018 | VP06018 | SAMN20167813 | SRR15099207 | NonPC | Public | 2006 | China | Clinical | 91.2 | 100 | 0.19 | 98.3924 | 5064395 | 45.28 | 23 | 934455 |
| VP06019 | VP06019 | SAMN20167814 | SRR15099206 | PC | Public | 2006 | China | Clinical | 97.5 | 100 | 0 | 99.9066 | 5120058 | 45.3 | 55 | 421024 |
| VP06020 | VP06020 | SAMN20167815 | SRR15099205 | PC | Public | 2006 | China | Clinical | 97.5 | 100 | 0 | 99.9173 | 5119046 | 45.3 | 58 | 312825 |
| VP06021 | VP06021 | SAMN20167818 | SRR15099201 | PC | Public | 2006 | China | Clinical | 98 | 100 | 0 | 99.9316 | 5174795 | 45.13 | 60 | 373649 |
| VP06022 | VP06022 | SAMN20167816 | SRR15099204 | PC | Public | 2006 | China | Clinical | 97.5 | 100 | 0 | 99.9086 | 5147244 | 45.28 | 56 | 373621 |
| VP06023 | VP06023 | SAMN20167817 | SRR15099202 | PC | Public | 2006 | China | Clinical | 97.5 | 100 | 0 | 99.9376 | 5146502 | 45.28 | 55 | 373621 |
| VP06024 | VP06024 | SAMN20167819 | SRR15099200 | PC | Public | 2006 | China | Clinical | 98 | 100 | 0 | 99.9207 | 5174668 | 45.13 | 62 | 425710 |
| VP06025 | VP06025 | SAMN20167781 | SRR15099781 | NonPC | Public | 2006 | China | Env | 88 | 100 | 0.49 | 98.2066 | 5351625 | 45.03 | 130 | 131644 |
| VP06026 | VP06026 | SAMN20167785 | SRR15099737 | NonPC | Public | 2006 | China | Env | 92.1 | 100 | 0 | 98.3676 | 5058750 | 45.29 | 78 | 401157 |
| VP06027 | VP06027 | SAMN20167821 | SRR15099198 | PC | Public | 2006 | China | Clinical | 98.6 | 100 | 0 | 99.9843 | 5132257 | 45.29 | 61 | 394875 |
| VP06028 | VP06028 | SAMN20167822 | SRR15099197 | PC | Public | 2006 | China | Clinical | 98.6 | 100 | 0 | 99.9839 | 5131384 | 45.29 | 58 | 401330 |
| VP06029 | VP06029 | SAMN20167823 | SRR15099196 | PC | Public | 2006 | China | Clinical | 98.6 | 100 | 0 | 99.9923 | 5130864 | 45.29 | 56 | 427218 |
| VP06030 | VP06030 | SAMN20167786 | SRR15099726 | NonPC | Public | 2006 | China | Env | 88.5 | 100 | 0.56 | 98.2816 | 5053830 | 45.26 | 42 | 590824 |
| VP06031 | VP06031 | SAMN20167782 | SRR15099770 | NonPC | Public | 2006 | China | Env | 91.8 | 100 | 0.03 | 98.2834 | 5052210 | 45.31 | 44 | 458847 |
| VP06032 | VP06032 | SAMN20167783 | SRR15099759 | NonPC | Public | 2006 | China | Env | 91.7 | 100 | 0.03 | 98.3049 | 5049663 | 45.31 | 47 | 458853 |
| VP06033 | VP06033 | SAMN20167825 | SRR15099194 | NonPC | Public | 2006 | China | Clinical | 89.3 | 100 | 0.69 | 98.2794 | 5065765 | 45.29 | 30 | 451106 |
| VP06034 | VP06034 | SAMN20167827 | SRR15099191 | PC | Public | 2006 | China | Clinical | 98 | 99.46 | 0 | 99.9894 | 5073319 | 45.3 | 50 | 425751 |
| VP06035 | VP06035 | SAMN20167839 | SRR15099178 | PC | Public | 2006 | China | Clinical | 98 | 100 | 0 | 99.9801 | 5079496 | 45.3 | 54 | 410660 |

| | | | | | | | | | | | | | | | | |
|---|---|---|---|---|---|---|---|---|---|---|---|---|---|---|---|---|
| VP06036 | VP06036 | SAMN20167838 | SRR15099179 | PC | Public | 2006 | China | Clinical | 98.6 | 100 | 0.27 | 99.9854 | 5152858 | 45.29 | 58 | 427332 |
| VP06037 | VP06037 | SAMN20167840 | SRR15099177 | PC | Public | 2006 | China | Clinical | 97.8 | 100 | 0 | 99.9835 | 5070396 | 45.31 | 62 | 408750 |
| VP06038 | VP06038 | SAMN20167842 | SRR15099175 | NonPC | Public | 2006 | China | Clinical | 92.3 | 100 | 0.46 | 98.3595 | 5200957 | 45.19 | 67 | 470315 |
| VP06039 | VP06039 | SAMN20167843 | SRR15099174 | NonPC | Public | 2006 | China | Clinical | 92.3 | 100 | 0.46 | 98.3618 | 5202173 | 45.19 | 68 | 327714 |
| VP06041 | VP06041 | SAMN20167841 | SRR15099176 | PC | Public | 2006 | China | Clinical | 98.1 | 100 | 0 | 99.9199 | 5182023 | 45.14 | 63 | 427747 |
| VP06042 | VP06042 | SAMN20167853 | SRR15099163 | PC | Public | 2006 | China | Clinical | 98.7 | 100 | 0 | 99.985 | 5113004 | 45.29 | 53 | 427446 |
| VP06043 | VP06043 | SAMN20167854 | SRR15099162 | PC | Public | 2006 | China | Clinical | 98.6 | 100 | 0 | 99.9847 | 5112307 | 45.28 | 56 | 427320 |
| VP06044 | VP06044 | SAMN20167860 | SRR15099155 | PC | Public | 2006 | China | Clinical | 98 | 100 | 0 | 99.9842 | 5079939 | 45.3 | 58 | 394969 |
| VP06045 | VP06045 | SAMN20167852 | SRR15099164 | PC | Public | 2006 | China | Clinical | 98.5 | 100 | 0 | 99.9839 | 5093559 | 45.29 | 57 | 425348 |
| VP06046 | VP06046 | SAMN20167845 | SRR15099172 | PC | Public | 2006 | China | Clinical | 98.5 | 100 | 0 | 99.9838 | 5093748 | 45.29 | 56 | 425349 |
| VP06047 | VP06047 | SAMN20167861 | SRR15099154 | PC | Public | 2006 | China | Clinical | 98 | 100 | 0 | 99.9896 | 5080500 | 45.3 | 60 | 410888 |
| VP06048 | VP06048 | SAMN20167849 | SRR15099167 | NonPC | Public | 2006 | China | Clinical | 91.9 | 100 | 0 | 98.4194 | 5035453 | 45.27 | 40 | 522556 |
| VP06050 | VP06050 | SAMN20167848 | SRR15099168 | NonPC | Public | 2006 | China | Clinical | 90.7 | 100 | 0.18 | 98.3715 | 5426524 | 45.13 | 96 | 386137 |
| VP06051 | VP06051 | SAMN20167850 | SRR15099166 | NonPC | Public | 2006 | China | Clinical | 92.9 | 100 | 0 | 98.3723 | 5181622 | 45.2 | 45 | 498054 |
| VP06052 | VP06052 | SAMN20167847 | SRR15099169 | NonPC | Public | 2006 | China | Clinical | 89.3 | 100 | 0.4 | 98.294 | 5037767 | 45.26 | 52 | 448669 |
| VP06053 | VP06053 | SAMN20167855 | SRR15099161 | PC | Public | 2006 | China | Clinical | 98.6 | 100 | 0 | 99.994 | 5111616 | 45.29 | 62 | 334369 |
| VP06054 | VP06054 | SAMN20167856 | SRR15099160 | PC | Public | 2006 | China | Clinical | 98.6 | 100 | 0 | 99.9849 | 5111551 | 45.28 | 58 | 401625 |
| VP06056 | VP06056 | SAMN20167857 | SRR15099158 | PC | Public | 2006 | China | Clinical | 98.6 | 100 | 0 | 99.9841 | 5110819 | 45.29 | 57 | 427434 |
| VP06057 | VP06057 | SAMN20167858 | SRR15099157 | PC | Public | 2006 | China | Clinical | 98.6 | 100 | 0 | 99.9834 | 5112260 | 45.29 | 56 | 426906 |
| VP06058 | VP06058 | SAMN20167859 | SRR15099156 | PC | Public | 2006 | China | Clinical | 98.6 | 100 | 0 | 99.9924 | 5110804 | 45.28 | 58 | 427020 |
| VP06059 | VP06059 | SAMN20167851 | SRR15099165 | NonPC | Public | 2006 | China | Clinical | 90.9 | 100 | 0 | 98.292 | 5122149 | 45.29 | 67 | 374815 |
| VP06060 | VP06060 | SAMN20167846 | SRR15099171 | NonPC | Public | 2006 | China | Clinical | 90.9 | 100 | 0 | 98.293 | 5136946 | 45.27 | 61 | 450909 |
| VP06061 | VP06061 | SAMN20167863 | SRR15099152 | PC | Public | 2006 | China | Clinical | 98.5 | 100 | 0 | 99.9924 | 5092617 | 45.29 | 54 | 425235 |
| VP06062 | VP06062 | SAMN20167864 | SRR15099151 | PC | Public | 2006 | China | Clinical | 98.5 | 100 | 0 | 99.9895 | 5092859 | 45.29 | 56 | 425349 |
| VP06063 | VP06063 | SAMN20167865 | SRR15099150 | PC | Public | 2006 | China | Clinical | 98.5 | 100 | 0 | 99.9834 | 5094361 | 45.3 | 61 | 394955 |
| VP06064 | VP06064 | SAMN20167866 | SRR15099149 | PC | Public | 2006 | China | Clinical | 98.5 | 100 | 0 | 99.9827 | 5093920 | 45.3 | 59 | 401604 |
| VP06065 | VP06065 | SAMN20167867 | SRR15099147 | PC | Public | 2006 | China | Clinical | 98.4 | 100 | 0 | 99.9834 | 5089860 | 45.29 | 61 | 425769 |
| VP06066 | VP06066 | SAMN20167862 | SRR15099153 | NonPC | Public | 2006 | China | Clinical | 89.3 | 100 | 0 | 98.3468 | 4938963 | 45.44 | 53 | 349703 |
| VP06067 | VP06067 | SAMN20167868 | SRR15099146 | PC | Public | 2006 | China | Clinical | 98.4 | 100 | 0 | 99.9901 | 5133723 | 45.34 | 63 | 425769 |
| VP06068 | VP06068 | SAMN20167787 | SRR15099714 | PC | Public | 2006 | China | Env | 98.2 | 100 | 0 | 99.979 | 5113107 | 45.26 | 42 | 427344 |
| VP06069 | VP06069 | SAMN20167889 | SRR15099123 | PC | Public | 2006 | China | Clinical | 97.7 | 100 | 0 | 99.8947 | 5251677 | 45.27 | 59 | 316808 |
| VP06070 | VP06070 | SAMN20167892 | SRR15099120 | NonPC | Public | 2006 | China | Clinical | 90.9 | 100 | 0 | 98.3066 | 5121767 | 45.29 | 68 | 374815 |
| VP06071 | VP06071 | SAMN20167890 | SRR15099122 | NonPC | Public | 2006 | China | Env | 88.4 | 100 | 0.94 | 98.2597 | 4993813 | 45.3 | 61 | 741982 |
| VP06072 | VP06072 | SAMN20167891 | SRR15099121 | NonPC | Public | 2006 | China | Env | 91.4 | 100 | 0 | 98.3983 | 5010360 | 45.33 | 65 | 346286 |
| VP06073 | VP06073 | SAMN20167893 | SRR15099119 | PC | Public | 2006 | China | Env | 98.5 | 100 | 0 | 99.9846 | 5093657 | 45.3 | 58 | 425451 |
| VP06075 | VP06075 | SAMN20167894 | SRR15099118 | PC | Public | 2006 | China | Env | 98.5 | 100 | 0 | 99.9927 | 5093191 | 45.29 | 58 | 425337 |
| VP06076 | VP06076 | SAMN20167895 | SRR15099117 | PC | Public | 2006 | China | Env | 98.5 | 100 | 0 | 99.9927 | 5093774 | 45.29 | 56 | 425337 |
| VP06077 | VP06077 | SAMN20167896 | SRR15099116 | PC | Public | 2006 | China | Env | 98.5 | 100 | 0 | 99.9829 | 5094021 | 45.3 | 57 | 401470 |
| VP06078 | VP06078 | SAMN20167897 | SRR15099113 | PC | Public | 2006 | China | Env | 98.5 | 100 | 0 | 99.9917 | 5093084 | 45.29 | 56 | 425337 |
| VP06079 | VP06079 | SAMN20167872 | SRR15099142 | PC | Public | 2006 | China | Clinical | 98.5 | 100 | 0 | 99.9851 | 5123461 | 45.29 | 61 | 394875 |
| VP06080 | VP06080 | SAMN20167873 | SRR15099141 | PC | Public | 2006 | China | Clinical | 98.5 | 100 | 0 | 99.9937 | 5123308 | 45.29 | 60 | 394877 |
| VP06081 | VP06081 | SAMN20167874 | SRR15099140 | PC | Public | 2006 | China | Clinical | 98.5 | 100 | 0 | 99.9851 | 5122411 | 45.29 | 65 | 288858 |
| VP06082 | VP06082 | SAMN20167875 | SRR15099139 | PC | Public | 2006 | China | Clinical | 98.5 | 100 | 0 | 99.9844 | 5121915 | 45.29 | 60 | 394875 |
| VP06083 | VP06083 | SAMN20167871 | SRR15099143 | NonPC | Public | 2006 | China | Clinical | 93.1 | 100 | 0.05 | 98.3818 | 5067591 | 45.3 | 49 | 463660 |
| VP06084 | VP06084 | SAMN20167877 | SRR15099136 | PC | Public | 2006 | China | Clinical | 98.6 | 100 | 0 | 99.9918 | 5130401 | 45.29 | 58 | 401463 |
| VP06085 | VP06085 | SAMN20167884 | SRR15099129 | PC | Public | 2006 | China | Clinical | 98.6 | 100 | 0 | 99.9825 | 5132749 | 45.29 | 56 | 427224 |
| VP06086 | VP06086 | SAMN20167882 | SRR15099131 | PC | Public | 2006 | China | Clinical | 98.2 | 100 | 0 | 99.991 | 5115027 | 45.26 | 50 | 347616 |
| VP06087 | VP06087 | SAMN20167885 | SRR15099128 | PC | Public | 2006 | China | Clinical | 98.6 | 100 | 0 | 99.9935 | 5133096 | 45.29 | 57 | 401463 |

| | | | | | | | | | | | | | | | | |
|---|---|---|---|---|---|---|---|---|---|---|---|---|---|---|---|---|
| VP06088 | VP06088 | SAMN20167878 | SRR15099135 | PC | Public | 2006 | China | Clinical | 97.9 | 100 | 0 | 99.9296 | 5170642 | 45.22 | 65 | 373621 |
| VP06089 | VP06089 | SAMN20167886 | SRR15099127 | PC | Public | 2006 | China | Clinical | 98.5 | 100 | 0 | 99.9896 | 5122651 | 45.29 | 56 | 401597 |
| VP06090 | VP06090 | SAMN20167887 | SRR15099125 | PC | Public | 2006 | China | Clinical | 98.7 | 100 | 0 | 99.9863 | 5133409 | 45.29 | 60 | 394877 |
| VP06091 | VP06091 | SAMN20167881 | SRR15099132 | NonPC | Public | 2006 | China | Clinical | 93.1 | 100 | 0.05 | 98.3936 | 5072245 | 45.29 | 45 | 515956 |
| VP06092 | VP06092 | SAMN20167883 | SRR15099130 | PC | Public | 2006 | China | Clinical | 97.8 | 100 | 0.27 | 99.9835 | 5112681 | 45.32 | 65 | 368358 |
| VP06093 | VP06093 | SAMN20167876 | SRR15099138 | NonPC | Public | 2006 | China | Clinical | 92.5 | 100 | 0.05 | 98.3474 | 5069737 | 45.27 | 46 | 450396 |
| VP06094 | VP06094 | SAMN20167879 | SRR15099134 | PC | Public | 2006 | China | Clinical | 97.9 | 100 | 0 | 99.9243 | 5132583 | 45.25 | 49 | 401611 |
| VP06095 | VP06095 | SAMN20167880 | SRR15099133 | PC | Public | 2006 | China | Clinical | 98.1 | 100 | 0 | 99.9208 | 5170229 | 45.15 | 60 | 427861 |
| VP06096 | VP06096 | SAMN20167888 | SRR15099124 | PC | Public | 2006 | China | Clinical | 98.5 | 100 | 0 | 99.994 | 5123042 | 45.29 | 61 | 394711 |
| VP06097 | VP06097 | SAMN20167869 | SRR15099145 | PC | Public | 2006 | China | Clinical | 97.2 | 100 | 0 | 99.9378 | 5126877 | 45.29 | 61 | 256883 |
| VP06098 | VP06098 | SAMN20167788 | SRR15099703 | PC | Public | 2006 | China | Clinical | 97.6 | 100 | 0 | 99.8908 | 5163947 | 45.28 | 51 | 477807 |
| VP06099 | VP06099 | SAMN20167789 | SRR15099692 | PC | Public | 2006 | China | Clinical | 97.5 | 100 | 0 | 99.9178 | 5165253 | 45.28 | 54 | 373587 |
| VP06100 | VP06100 | SAMN20167790 | SRR15099681 | PC | Public | 2006 | China | Clinical | 97.6 | 100 | 0 | 99.8901 | 5164124 | 45.28 | 54 | 318037 |
| VP06101 | VP06101 | SAMN20167791 | SRR15099670 | PC | Public | 2006 | China | Clinical | 97.6 | 100 | 0 | 99.9003 | 5164256 | 45.28 | 53 | 477808 |
| VP06102 | VP06102 | SAMN20167792 | SRR15099659 | PC | Public | 2006 | China | Clinical | 97.6 | 100 | 0 | 99.9214 | 5165406 | 45.28 | 53 | 440631 |
| VP06103 | VP06103 | SAMN20167793 | SRR15099648 | PC | Public | 2006 | China | Clinical | 97.6 | 100 | 0 | 99.8934 | 5164990 | 45.28 | 52 | 394947 |
| VP06104 | VP06104 | SAMN20167794 | SRR15099637 | PC | Public | 2006 | China | Clinical | 97.5 | 99.73 | 0 | 99.8942 | 5159839 | 45.28 | 50 | 401604 |
| VP06105 | VP06105 | SAMN20167795 | SRR15099626 | PC | Public | 2006 | China | Clinical | 97.6 | 100 | 0 | 99.8958 | 5164796 | 45.28 | 53 | 373621 |
| VP06106 | VP06106 | SAMN20167796 | SRR15099615 | PC | Public | 2006 | China | Clinical | 97.6 | 100 | 0 | 99.8982 | 5165071 | 45.28 | 52 | 401468 |
| VP06107 | VP06107 | SAMN20167797 | SRR15099224 | PC | Public | 2006 | China | Clinical | 97.6 | 100 | 0 | 99.8899 | 5165291 | 45.28 | 51 | 373621 |
| VP06108 | VP06108 | SAMN20167798 | SRR15099223 | PC | Public | 2006 | China | Clinical | 97.6 | 100 | 0 | 99.8997 | 5164605 | 45.28 | 53 | 373621 |
| VP06109 | VP06109 | SAMN20167899 | SRR15099111 | PC | Public | 2006 | China | Clinical | 98.5 | 100 | 0 | 99.9814 | 5123164 | 45.29 | 56 | 401463 |
| VP06110 | VP06110 | SAMN20167898 | SRR15099112 | NonPC | Public | 2006 | China | Clinical | 92.9 | 100 | 0 | 98.3945 | 5124756 | 45.22 | 43 | 498054 |
| VP06111 | VP06111 | SAMN20167913 | SRR15099096 | PC | Public | 2006 | China | Clinical | 98.6 | 100 | 0 | 99.9867 | 5132299 | 45.29 | 57 | 401464 |
| VP06112 | VP06112 | SAMN20167914 | SRR15099095 | PC | Public | 2006 | China | Clinical | 98.7 | 100 | 0 | 99.985 | 5132797 | 45.29 | 58 | 394875 |
| VP06113 | VP06113 | SAMN20167915 | SRR15099094 | PC | Public | 2006 | China | Clinical | 98.6 | 100 | 0 | 99.9939 | 5132214 | 45.29 | 60 | 427218 |
| VP06114 | VP06114 | SAMN20167908 | SRR15099101 | PC | Public | 2006 | China | Clinical | 98.6 | 100 | 0 | 99.9932 | 5133273 | 45.29 | 59 | 401464 |
| VP06115 | VP06115 | SAMN20167911 | SRR15099098 | PC | Public | 2006 | China | Clinical | 98.6 | 100 | 0 | 99.994 | 5132444 | 45.29 | 59 | 394877 |
| VP06116 | VP06116 | SAMN20167902 | SRR15099108 | PC | Public | 2006 | China | Clinical | 98.6 | 100 | 0 | 99.9854 | 5131808 | 45.29 | 58 | 395055 |
| VP06117 | VP06117 | SAMN20167900 | SRR15099110 | PC | Public | 2006 | China | Clinical | 98.6 | 100 | 0 | 99.9917 | 5132448 | 45.29 | 57 | 427218 |
| VP06118 | VP06118 | SAMN20167901 | SRR15099109 | PC | Public | 2006 | China | Clinical | 98.6 | 100 | 0 | 99.9845 | 5132869 | 45.29 | 55 | 401464 |
| VP06119 | VP06119 | SAMN20167903 | SRR15099107 | PC | Public | 2006 | China | Clinical | 98.6 | 100 | 0 | 99.9943 | 5131712 | 45.29 | 57 | 427218 |
| VP06121 | VP06121 | SAMN20167904 | SRR15099106 | PC | Public | 2006 | China | Clinical | 98.7 | 100 | 0 | 99.9846 | 5133629 | 45.29 | 56 | 401330 |
| VP06122 | VP06122 | SAMN20167905 | SRR15099105 | PC | Public | 2006 | China | Clinical | 98.7 | 100 | 0 | 99.9846 | 5132875 | 45.29 | 56 | 401464 |
| VP06123 | VP06123 | SAMN20167906 | SRR15099104 | PC | Public | 2006 | China | Clinical | 98.7 | 100 | 0 | 99.9945 | 5132178 | 45.29 | 59 | 394877 |
| VP06124 | VP06124 | SAMN20167909 | SRR15099100 | PC | Public | 2006 | China | Clinical | 98.6 | 100 | 0 | 99.9941 | 5132688 | 45.29 | 59 | 395055 |
| VP06125 | VP06125 | SAMN20167912 | SRR15099097 | PC | Public | 2006 | China | Clinical | 98.6 | 100 | 0 | 99.9835 | 5131794 | 45.29 | 59 | 394875 |
| VP06127 | VP06127 | SAMN20167907 | SRR15099102 | PC | Public | 2006 | China | Clinical | 98.6 | 100 | 0 | 99.9952 | 5132809 | 45.29 | 63 | 395053 |
| VP06128 | VP06128 | SAMN20167910 | SRR15099099 | PC | Public | 2006 | China | Clinical | 98.6 | 100 | 0 | 99.9844 | 5131084 | 45.29 | 61 | 394794 |
| VP06129 | VP06129 | SAMN20167918 | SRR15099090 | PC | Public | 2006 | China | Clinical | 98.6 | 100 | 0 | 99.9943 | 5132047 | 45.29 | 59 | 401597 |
| VP06130 | VP06130 | SAMN20167919 | SRR15099089 | PC | Public | 2006 | China | Clinical | 98.6 | 100 | 0 | 99.9941 | 5132379 | 45.29 | 62 | 427218 |
| VP06131 | VP06131 | SAMN20167920 | SRR15099088 | PC | Public | 2006 | China | Clinical | 98.6 | 100 | 0 | 99.986 | 5131422 | 45.29 | 60 | 401463 |
| VP06132 | VP06132 | SAMN20167916 | SRR15099093 | PC | Public | 2006 | China | Clinical | 98.6 | 100 | 0 | 99.9799 | 5151048 | 45.27 | 62 | 420264 |
| VP06133 | VP06133 | SAMN20167917 | SRR15099091 | PC | Public | 2006 | China | Clinical | 98.6 | 100 | 0.27 | 99.9855 | 5152731 | 45.29 | 57 | 401625 |
| VP06134 | VP06134 | SAMN20167937 | SRR15099069 | PC | Public | 2006 | China | Clinical | 98.6 | 100 | 0 | 99.9942 | 5111929 | 45.28 | 58 | 427296 |
| VP06135 | VP06135 | SAMN20167938 | SRR15099068 | PC | Public | 2006 | China | Clinical | 98.6 | 100 | 0 | 99.9857 | 5110988 | 45.29 | 58 | 427332 |
| VP06136 | VP06136 | SAMN20167939 | SRR15099067 | PC | Public | 2006 | China | Clinical | 98.6 | 100 | 0 | 99.9853 | 5112309 | 45.29 | 56 | 427446 |
| VP06137 | VP06137 | SAMN20167940 | SRR15099066 | PC | Public | 2006 | China | Clinical | 98.6 | 100 | 0 | 99.9935 | 5111187 | 45.28 | 56 | 401625 |

| | | | | | | | | | | | | | | | | |
|---|---|---|---|---|---|---|---|---|---|---|---|---|---|---|---|---|
| VP06138 | VP06138 | SAMN20167941 | SRR15099065 | PC | Public | 2006 | China | Clinical | 98.6 | 100 | 0 | 99.9858 | 5110735 | 45.28 | 57 | 401491 |
| VP06140 | VP06140 | SAMN20167942 | SRR15099064 | PC | Public | 2006 | China | Clinical | 98.6 | 100 | 0.27 | 99.9844 | 5152929 | 45.29 | 61 | 427218 |
| VP06141 | VP06141 | SAMN20167934 | SRR15099073 | PC | Public | 2006 | China | Clinical | 98.6 | 100 | 0 | 99.9937 | 5110968 | 45.28 | 58 | 427325 |
| VP06142 | VP06142 | SAMN20167943 | SRR15099063 | PC | Public | 2006 | China | Clinical | 98.6 | 100 | 0 | 99.9931 | 5110640 | 45.28 | 56 | 401625 |
| VP06143 | VP06143 | SAMN20167935 | SRR15099072 | PC | Public | 2006 | China | Clinical | 98.6 | 100 | 0 | 99.985 | 5111033 | 45.28 | 56 | 427326 |
| VP06144 | VP06144 | SAMN20167936 | SRR15099071 | PC | Public | 2006 | China | Clinical | 98.6 | 100 | 0 | 99.9943 | 5110100 | 45.28 | 59 | 401491 |
| VP06145 | VP06145 | SAMN20167932 | SRR15099075 | PC | Public | 2006 | China | Clinical | 97.7 | 100 | 0 | 99.8997 | 5147850 | 45.29 | 56 | 401618 |
| VP06146 | VP06146 | SAMN20167933 | SRR15099074 | PC | Public | 2006 | China | Clinical | 97.8 | 100 | 0 | 99.9147 | 5153308 | 45.3 | 53 | 420015 |
| VP06147 | VP06147 | SAMN20167922 | SRR15099086 | NonPC | Public | 2006 | China | Clinical | 92.3 | 100 | 0.19 | 98.3464 | 5024915 | 45.23 | 51 | 471205 |
| VP06148 | VP06148 | SAMN20167923 | SRR15099085 | NonPC | Public | 2006 | China | Clinical | 92.3 | 100 | 0.19 | 98.344 | 5024876 | 45.24 | 52 | 450296 |
| VP06149 | VP06149 | SAMN20167924 | SRR15099084 | NonPC | Public | 2006 | China | Clinical | 92.3 | 100 | 0.19 | 98.3502 | 5024818 | 45.24 | 51 | 450296 |
| VP06150 | VP06150 | SAMN20167925 | SRR15099083 | NonPC | Public | 2006 | China | Clinical | 92.3 | 100 | 0.19 | 98.3662 | 5025612 | 45.24 | 53 | 450393 |
| VP06151 | VP06151 | SAMN20167926 | SRR15099082 | NonPC | Public | 2006 | China | Clinical | 92.3 | 100 | 0.19 | 98.3659 | 5024795 | 45.23 | 50 | 450393 |
| VP06152 | VP06152 | SAMN20167927 | SRR15099080 | NonPC | Public | 2006 | China | Clinical | 92.3 | 100 | 0.19 | 98.3695 | 5028119 | 45.25 | 54 | 450296 |
| VP06153 | VP06153 | SAMN20167928 | SRR15099079 | NonPC | Public | 2006 | China | Clinical | 92.3 | 100 | 0.19 | 98.3469 | 5025299 | 45.24 | 52 | 450393 |
| VP06154 | VP06154 | SAMN20167929 | SRR15099078 | NonPC | Public | 2006 | China | Clinical | 92.3 | 100 | 0.19 | 98.3845 | 5024465 | 45.23 | 50 | 450296 |
| VP06155 | VP06155 | SAMN20167930 | SRR15099077 | NonPC | Public | 2006 | China | Clinical | 92.3 | 100 | 0.19 | 98.3697 | 5024298 | 45.23 | 51 | 450296 |
| VP06156 | VP06156 | SAMN20167931 | SRR15099076 | NonPC | Public | 2006 | China | Clinical | 92 | 100 | 0.19 | 98.3584 | 5026170 | 45.24 | 51 | 450393 |
| VP06157 | VP06157 | SAMN20167921 | SRR15099087 | NonPC | Public | 2006 | China | Clinical | 89 | 100 | 0.12 | 98.2703 | 5056982 | 45.3 | 45 | 548443 |
| VP06158 | VP06158 | SAMN20167951 | SRR15099054 | PC | Public | 2006 | China | Clinical | 98.5 | 100 | 0 | 99.9852 | 5093996 | 45.29 | 58 | 425463 |
| VP06159 | VP06159 | SAMN20167952 | SRR15099053 | PC | Public | 2006 | China | Clinical | 98.5 | 100 | 0 | 99.985 | 5093562 | 45.29 | 58 | 401604 |
| VP06160 | VP06160 | SAMN20167953 | SRR15099052 | PC | Public | 2006 | China | Clinical | 98.5 | 100 | 0 | 99.9914 | 5093863 | 45.29 | 57 | 394963 |
| VP06161 | VP06161 | SAMN20167950 | SRR15099055 | NonPC | Public | 2006 | China | Clinical | 91 | 100 | 0 | 98.3237 | 5026417 | 45.25 | 54 | 404966 |
| VP06162 | VP06162 | SAMN20167944 | SRR15099062 | PC | Public | 2006 | China | Clinical | 98.6 | 100 | 0 | 99.9938 | 5112445 | 45.28 | 57 | 427440 |
| VP06163 | VP06163 | SAMN20167945 | SRR15099061 | PC | Public | 2006 | China | Clinical | 98.7 | 100 | 0 | 99.9928 | 5112319 | 45.28 | 56 | 401491 |
| VP06164 | VP06164 | SAMN20167946 | SRR15099060 | PC | Public | 2006 | China | Clinical | 98.6 | 100 | 0 | 99.9935 | 5112575 | 45.29 | 57 | 427440 |
| VP06165 | VP06165 | SAMN20167947 | SRR15099058 | PC | Public | 2006 | China | Clinical | 98.6 | 100 | 0 | 99.9857 | 5111134 | 45.28 | 58 | 401625 |
| VP06166 | VP06166 | SAMN20167948 | SRR15099057 | PC | Public | 2006 | China | Clinical | 98.6 | 100 | 0 | 99.9938 | 5110407 | 45.29 | 59 | 347634 |
| VP06167 | VP06167 | SAMN20167949 | SRR15099056 | PC | Public | 2006 | China | Clinical | 98.6 | 100 | 0 | 99.9851 | 5111132 | 45.28 | 56 | 401342 |
| VP06168 | VP06168 | SAMN20167954 | SRR15099051 | NonPC | Public | 2006 | China | Clinical | 92.3 | 100 | 0.19 | 98.3491 | 5043935 | 45.25 | 52 | 471204 |
| VP06169 | VP06169 | SAMN20167955 | SRR15099050 | NonPC | Public | 2006 | China | Clinical | 92.3 | 100 | 0.19 | 98.3485 | 5044032 | 45.25 | 51 | 539189 |
| VP06170 | VP06170 | SAMN20167956 | SRR15099049 | NonPC | Public | 2006 | China | Clinical | 92.3 | 100 | 0.19 | 98.3673 | 5046651 | 45.25 | 52 | 539189 |
| VP06171 | VP06171 | SAMN20167957 | SRR15099047 | NonPC | Public | 2006 | China | Clinical | 92.3 | 100 | 0.19 | 98.3588 | 5046712 | 45.25 | 52 | 539189 |
| VP06172 | VP06172 | SAMN20167959 | SRR15099045 | PC | Public | 2006 | China | Clinical | 98.6 | 100 | 0 | 99.9898 | 5131802 | 45.29 | 60 | 394877 |
| VP06173 | VP06173 | SAMN20167960 | SRR15099044 | PC | Public | 2006 | China | Clinical | 98.6 | 100 | 0 | 99.995 | 5131465 | 45.29 | 57 | 401477 |
| VP06174 | VP06174 | SAMN20167961 | SRR15099043 | PC | Public | 2006 | China | Clinical | 98.6 | 100 | 0 | 99.984 | 5131800 | 45.29 | 55 | 401478 |
| VP06175 | VP06175 | SAMN20167962 | SRR15099042 | PC | Public | 2006 | China | Clinical | 98.6 | 100 | 0 | 99.9938 | 5136605 | 45.29 | 60 | 401611 |
| VP06176 | VP06176 | SAMN20167963 | SRR15099041 | PC | Public | 2006 | China | Clinical | 98.6 | 100 | 0 | 99.9912 | 5131802 | 45.29 | 56 | 441863 |
| VP06177 | VP06177 | SAMN20167964 | SRR15099040 | PC | Public | 2006 | China | Clinical | 98.6 | 100 | 0 | 99.9847 | 5131211 | 45.29 | 59 | 401611 |
| VP06178 | VP06178 | SAMN20167965 | SRR15099039 | PC | Public | 2006 | China | Clinical | 98.7 | 100 | 0 | 99.9849 | 5131994 | 45.29 | 55 | 401477 |
| VP06179 | VP06179 | SAMN20167966 | SRR15099038 | PC | Public | 2006 | China | Clinical | 98.6 | 100 | 0 | 99.9849 | 5131973 | 45.29 | 58 | 401344 |
| VP06181 | VP06181 | SAMN20167829 | SRR15099189 | NonPC | Public | 2006 | China | Clinical | 92.3 | 100 | 0.46 | 98.3585 | 5171821 | 45.22 | 70 | 323764 |
| VP06183 | VP06183 | SAMN20167835 | SRR15099183 | NonPC | Public | 2006 | China | Clinical | 90.9 | 100 | 0 | 98.2881 | 5125421 | 45.29 | 68 | 405678 |
| VP06184 | VP06184 | SAMN20167828 | SRR15099190 | NonPC | Public | 2006 | China | Clinical | 93.1 | 100 | 0.05 | 98.3914 | 5056098 | 45.31 | 43 | 515938 |
| VP06185 | VP06185 | SAMN20167830 | SRR15099188 | NonPC | Public | 2006 | China | Clinical | 91.2 | 100 | 0 | 98.2953 | 5097879 | 45.3 | 52 | 313668 |
| VP06186 | VP06186 | SAMN20167833 | SRR15099185 | NonPC | Public | 2006 | China | Env | 90.9 | 99.46 | 0 | 98.3592 | 5081324 | 45.3 | 48 | 402183 |
| VP06187 | VP06187 | SAMN20167834 | SRR15099184 | NonPC | Public | 2006 | China | Env | 91.2 | 100 | 0 | 98.3634 | 5079041 | 45.32 | 48 | 407473 |
| VP06188 | VP06188 | SAMN20167837 | SRR15099180 | PC | Public | 2006 | China | Env | 97.2 | 100 | 0 | 99.9194 | 5125860 | 45.28 | 62 | 256881 |

| | | | | | | | | | | | | | | | | |
|---|---|---|---|---|---|---|---|---|---|---|---|---|---|---|---|---|
| VP06189 | VP06189 | SAMN20167836 | SRR15099182 | PC | Public | 2006 | China | Clinical | 97.2 | 100 | 0 | 99.9209 | 5124190 | 45.29 | 53 | 394941 |
| VP06190 | VP06190 | SAMN20167831 | SRR15099187 | NonPC | Public | 2006 | China | Clinical | 91.2 | 100 | 0 | 98.3004 | 5079535 | 45.32 | 47 | 407473 |
| VP06191 | VP06191 | SAMN20167832 | SRR15099186 | NonPC | Public | 2006 | China | Clinical | 91.1 | 100 | 0 | 98.3079 | 5085167 | 45.3 | 52 | 402184 |
| VP06192 | VP06192 | SAMN20167799 | SRR15099222 | PC | Public | 2006 | China | Clinical | 98.6 | 100 | 0 | 99.9932 | 5132785 | 45.29 | 61 | 394875 |
| VP06193 | VP06193 | SAMN20167826 | SRR15099193 | PC | Public | 2006 | China | Clinical | 98.4 | 100 | 0.02 | 99.9797 | 5174107 | 45.14 | 66 | 425450 |
| VP06194 | VP06194 | SAMN20167820 | SRR15099199 | PC | Public | 2006 | China | Clinical | 98.6 | 100 | 0 | 99.985 | 5193751 | 45.21 | 68 | 312691 |
| VP06195 | VP06195 | SAMN20167824 | SRR15099195 | PC | Public | 2006 | China | Clinical | 98.6 | 100 | 0 | 99.9835 | 5132869 | 45.29 | 59 | 401463 |
| VP06196 | VP06196 | SAMN20167844 | SRR15099173 | PC | Public | 2006 | China | Clinical | 98.6 | 100 | 0 | 99.9856 | 5111216 | 45.28 | 57 | 347621 |
| VP06197 | VP06197 | SAMN20167958 | SRR15099046 | PC | Public | 2006 | China | Clinical | 98.6 | 100 | 0 | 99.9932 | 5254169 | 45.18 | 64 | 394851 |
| VP06198 | VP06198 | SAMN20167870 | SRR15099144 | PC | Public | 2006 | China | Clinical | 98.2 | 100 | 0 | 99.9901 | 5113246 | 45.26 | 49 | 427344 |
| VP07001 | VP07001 | SAMN20167976 | SRR15099027 | NonPC | Public | 2007 | China | Env | 92.1 | 100 | 0.19 | 98.3773 | 5032071 | 45.25 | 56 | 313852 |
| VP07002 | VP07002 | SAMN20167975 | SRR15099028 | NonPC | Public | 2007 | China | Clinical | 92.3 | 100 | 0.19 | 98.3646 | 5045903 | 45.26 | 51 | 471204 |
| VP07003 | VP07003 | SAMN20167974 | SRR15099029 | NonPC | Public | 2007 | China | Clinical | 89.7 | 100 | 0.11 | 98.282 | 5178516 | 45.12 | 38 | 454722 |
| VP07004 | VP07004 | SAMN20167977 | SRR15099025 | PC | Public | 2007 | China | Clinical | 98.5 | 100 | 0 | 99.9861 | 5092976 | 45.29 | 61 | 425859 |
| VP07005 | VP07005 | SAMN20167978 | SRR15099024 | PC | Public | 2007 | China | Clinical | 98.5 | 100 | 0 | 99.9845 | 5092230 | 45.29 | 59 | 425859 |
| VP07006 | VP07006 | SAMN20167979 | SRR15099023 | PC | Public | 2007 | China | Clinical | 98.5 | 100 | 0 | 99.9937 | 5092598 | 45.29 | 61 | 394826 |
| VP07007 | VP07007 | SAMN20167980 | SRR15099022 | PC | Public | 2007 | China | Clinical | 98.5 | 100 | 0 | 99.984 | 5093599 | 45.29 | 59 | 425445 |
| VP07008 | VP07008 | SAMN20167981 | SRR15099021 | PC | Public | 2007 | China | Clinical | 98.5 | 100 | 0 | 99.9945 | 5092549 | 45.29 | 61 | 401604 |
| VP07009 | VP07009 | SAMN20167982 | SRR15099020 | PC | Public | 2007 | China | Clinical | 98.5 | 100 | 0 | 99.9852 | 5100839 | 45.3 | 59 | 425859 |
| VP07010 | VP07010 | SAMN20167983 | SRR15099019 | PC | Public | 2007 | China | Clinical | 98.5 | 100 | 0 | 99.9853 | 5092714 | 45.29 | 57 | 425445 |
| VP07011 | VP07011 | SAMN20167968 | SRR15099035 | NonPC | Public | 2007 | China | Env | 90.6 | 100 | 0 | 98.3295 | 5184430 | 45.16 | 75 | 314851 |
| VP07012 | VP07012 | SAMN20167989 | SRR15099012 | PC | Public | 2007 | China | Clinical | 98.6 | 100 | 0 | 99.9747 | 5132911 | 45.29 | 58 | 427740 |
| VP07013 | VP07013 | SAMN20167990 | SRR15099011 | PC | Public | 2007 | China | Clinical | 98.6 | 100 | 0 | 99.9731 | 5132013 | 45.29 | 58 | 401607 |
| VP07014 | VP07014 | SAMN20167991 | SRR15099010 | PC | Public | 2007 | China | Clinical | 98.6 | 100 | 0 | 99.9743 | 5131680 | 45.29 | 59 | 401607 |
| VP07015 | VP07015 | SAMN20167984 | SRR15099018 | NonPC | Public | 2007 | China | Clinical | 91.7 | 100 | 0 | 98.3778 | 5138917 | 45.13 | 91 | 314777 |
| VP07016 | VP07016 | SAMN20167985 | SRR15099017 | PC | Public | 2007 | China | Clinical | 98.1 | 100 | 0 | 99.9358 | 5179664 | 45.14 | 56 | 442366 |
| VP07017 | VP07017 | SAMN20167986 | SRR15099016 | PC | Public | 2007 | China | Clinical | 98 | 100 | 0 | 99.9175 | 5174039 | 45.14 | 76 | 250156 |
| VP07018 | VP07018 | SAMN20167992 | SRR15099009 | PC | Public | 2007 | China | Clinical | 98.6 | 100 | 0 | 99.9741 | 5132571 | 45.29 | 59 | 401607 |
| VP07019 | VP07019 | SAMN20167987 | SRR15099014 | PC | Public | 2007 | China | Clinical | 98 | 100 | 0 | 99.938 | 5175471 | 45.13 | 59 | 425728 |
| VP07020 | VP07020 | SAMN20167993 | SRR15099008 | PC | Public | 2007 | China | Clinical | 98.7 | 100 | 0 | 99.9815 | 5132743 | 45.29 | 63 | 334357 |
| VP07021 | VP07021 | SAMN20167994 | SRR15099007 | PC | Public | 2007 | China | Clinical | 98.6 | 100 | 0 | 99.9817 | 5131147 | 45.29 | 58 | 427212 |
| VP07022 | VP07022 | SAMN20167988 | SRR15099013 | PC | Public | 2007 | China | Clinical | 98 | 100 | 0 | 99.9488 | 5103220 | 45.29 | 61 | 425734 |
| VP07023 | VP07023 | SAMN20167995 | SRR15099006 | PC | Public | 2007 | China | Clinical | 98.1 | 100 | 0 | 99.9894 | 5087093 | 45.27 | 81 | 265894 |
| VP07024 | VP07024 | SAMN20167996 | SRR15099005 | PC | Public | 2007 | China | Clinical | 98.5 | 100 | 0 | 99.9939 | 5106152 | 45.28 | 61 | 365149 |
| VP07025 | VP07025 | SAMN20167997 | SRR15099002 | PC | Public | 2007 | China | Clinical | 98.5 | 100 | 0 | 99.9932 | 5092284 | 45.29 | 58 | 401604 |
| VP07026 | VP07026 | SAMN20167967 | SRR15099036 | NonPC | Public | 2007 | China | Clinical | 89.6 | 100 | 0 | 98.2729 | 4990807 | 45.34 | 69 | 317784 |
| VP07027 | VP07027 | SAMN20167999 | SRR15099000 | PC | Public | 2007 | China | Clinical | 98.5 | 100 | 0 | 99.9755 | 5163011 | 45.27 | 59 | 394965 |
| VP07028 | VP07028 | SAMN20167998 | SRR15099001 | NonPC | Public | 2007 | China | Env | 91.9 | 100 | 0.03 | 98.3436 | 5255067 | 45.13 | 106 | 162659 |
| VP07029 | VP07029 | SAMN20168014 | SRR15098984 | PC | Public | 2007 | China | Clinical | 98.6 | 100 | 0 | 99.9844 | 5133046 | 45.29 | 59 | 395359 |
| VP07030 | VP07030 | SAMN20168010 | SRR15098988 | PC | Public | 2007 | China | Clinical | 98.5 | 100 | 0 | 99.9849 | 5160477 | 45.27 | 65 | 312853 |
| VP07031 | VP07031 | SAMN20168011 | SRR15098987 | PC | Public | 2007 | China | Clinical | 98.5 | 100 | 0 | 99.9632 | 5161790 | 45.27 | 69 | 312720 |
| VP07032 | VP07032 | SAMN20168012 | SRR15098986 | PC | Public | 2007 | China | Clinical | 98.5 | 100 | 0 | 99.9631 | 5161308 | 45.27 | 56 | 401632 |
| VP07034 | VP07034 | SAMN20168013 | SRR15098985 | PC | Public | 2007 | China | Clinical | 97.8 | 100 | 0 | 99.9924 | 5071899 | 45.32 | 60 | 414320 |
| VP07036 | VP07036 | SAMN20168021 | SRR15098976 | PC | Public | 2007 | China | Clinical | 97.7 | 100 | 0 | 99.9102 | 5148523 | 45.29 | 55 | 373635 |
| VP07037 | VP07037 | SAMN20168020 | SRR15098977 | PC | Public | 2007 | China | Clinical | 97.8 | 100 | 0 | 99.9793 | 5070602 | 45.31 | 59 | 408762 |
| VP07039 | VP07039 | SAMN20168018 | SRR15098979 | PC | Public | 2007 | China | Clinical | 97.9 | 100 | 0 | 99.9843 | 5079781 | 45.32 | 58 | 401611 |
| VP07040 | VP07040 | SAMN20168015 | SRR15098983 | PC | Public | 2007 | China | Clinical | 98.5 | 100 | 0 | 99.992 | 5106914 | 45.28 | 61 | 401603 |
| VP07042 | VP07042 | SAMN20168022 | SRR15098975 | PC | Public | 2007 | China | Clinical | 98.5 | 100 | 0 | 99.9845 | 5092712 | 45.29 | 59 | 401470 |

| | | | | | | | | | | | | | | | | |
|---|---|---|---|---|---|---|---|---|---|---|---|---|---|---|---|---|
| VP07043 | VP07043 | SAMN20168023 | SRR15098974 | PC | Public | 2007 | China | Clinical | 98.5 | 100 | 0 | 99.9924 | 5092799 | 45.29 | 60 | 394745 |
| VP07044 | VP07044 | SAMN20168024 | SRR15098973 | PC | Public | 2007 | China | Clinical | 98.5 | 100 | 0 | 99.9937 | 5093441 | 45.29 | 54 | 425355 |
| VP07045 | VP07045 | SAMN20168028 | SRR15098968 | PC | Public | 2007 | China | Clinical | 98 | 100 | 0 | 99.9781 | 5079645 | 45.3 | 58 | 410775 |
| VP07046 | VP07046 | SAMN20168025 | SRR15098972 | PC | Public | 2007 | China | Clinical | 98.5 | 100 | 0 | 99.9847 | 5093909 | 45.29 | 57 | 425355 |
| VP07047 | VP07047 | SAMN20168029 | SRR15098967 | PC | Public | 2007 | China | Clinical | 98 | 100 | 0 | 99.9794 | 5080335 | 45.3 | 62 | 410661 |
| VP07048 | VP07048 | SAMN20168026 | SRR15098971 | PC | Public | 2007 | China | Clinical | 98.5 | 100 | 0 | 99.9916 | 5091838 | 45.29 | 57 | 401604 |
| VP07049 | VP07049 | SAMN20168027 | SRR15098969 | PC | Public | 2007 | China | Clinical | 98.5 | 100 | 0 | 99.9915 | 5092394 | 45.29 | 58 | 425355 |
| VP07050 | VP07050 | SAMN20168019 | SRR15098978 | NonPC | Public | 2007 | China | Clinical | 90 | 100 | 0.48 | 98.2438 | 5111314 | 45.3 | 59 | 613157 |
| VP07051 | VP07051 | SAMN20167969 | SRR15099034 | NonPC | Public | 2007 | China | Clinical | 92.1 | 100 | 0.05 | 98.3277 | 5066474 | 45.27 | 48 | 478570 |
| VP07053 | VP07053 | SAMN20168038 | SRR15098957 | PC | Public | 2007 | China | Clinical | 97.8 | 100 | 0 | 99.9915 | 5070355 | 45.32 | 60 | 408756 |
| VP07054 | VP07054 | SAMN20168037 | SRR15098958 | PC | Public | 2007 | China | Clinical | 97.8 | 100 | 0 | 99.9829 | 5070571 | 45.31 | 61 | 401611 |
| VP07057 | VP07057 | SAMN20168046 | SRR15098949 | PC | Public | 2007 | China | Clinical | 97.8 | 100 | 0 | 99.9905 | 5070589 | 45.32 | 62 | 401477 |
| VP07058 | VP07058 | SAMN20168051 | SRR15098943 | PC | Public | 2007 | China | Clinical | 97.8 | 100 | 0 | 99.9924 | 5070167 | 45.31 | 64 | 401611 |
| VP07060 | VP07060 | SAMN20168048 | SRR15098946 | PC | Public | 2007 | China | Clinical | 98.5 | 100 | 0 | 99.9918 | 5182869 | 45.21 | 61 | 440641 |
| VP07061 | VP07061 | SAMN20168060 | SRR15098933 | PC | Public | 2007 | China | Clinical | 97.8 | 100 | 0 | 99.9803 | 5070318 | 45.31 | 58 | 415576 |
| VP07062 | VP07062 | SAMN20168059 | SRR15098934 | PC | Public | 2007 | China | Clinical | 98.6 | 100 | 0.27 | 99.9933 | 5153330 | 45.29 | 54 | 440953 |
| VP07063 | VP07063 | SAMN20168055 | SRR15098939 | PC | Public | 2007 | China | Clinical | 98.6 | 100 | 0.27 | 99.9856 | 5153389 | 45.29 | 57 | 401498 |
| VP07064 | VP07064 | SAMN20168058 | SRR15098935 | PC | Public | 2007 | China | Clinical | 98.5 | 100 | 0 | 99.9928 | 5092725 | 45.29 | 57 | 401604 |
| VP07065 | VP07065 | SAMN20168052 | SRR15098942 | PC | Public | 2007 | China | Clinical | 98.6 | 100 | 0 | 99.9928 | 5131540 | 45.29 | 57 | 427770 |
| VP07066 | VP07066 | SAMN20168054 | SRR15098940 | PC | Public | 2007 | China | Clinical | 98.5 | 100 | 0 | 99.9835 | 5123640 | 45.29 | 59 | 401597 |
| VP07067 | VP07067 | SAMN20168049 | SRR15098945 | PC | Public | 2007 | China | Clinical | 98.6 | 100 | 0 | 99.9846 | 5110801 | 45.28 | 55 | 401632 |
| VP07068 | VP07068 | SAMN20168053 | SRR15098941 | PC | Public | 2007 | China | Clinical | 98.6 | 100 | 0 | 99.9859 | 5118869 | 45.28 | 65 | 427176 |
| VP07069 | VP07069 | SAMN20168050 | SRR15098944 | PC | Public | 2007 | China | Clinical | 98.6 | 100 | 0 | 99.9943 | 5156690 | 45.24 | 58 | 427331 |
| VP07070 | VP07070 | SAMN20168044 | SRR15098951 | PC | Public | 2007 | China | Clinical | 98.5 | 100 | 0 | 99.9833 | 5091990 | 45.29 | 56 | 425355 |
| VP07071 | VP07071 | SAMN20168039 | SRR15098956 | NonPC | Public | 2007 | China | Clinical | 92.9 | 100 | 0 | 98.3759 | 5125178 | 45.22 | 40 | 572280 |
| VP07073 | VP07073 | SAMN20167972 | SRR15099031 | PC | Public | 2007 | China | Clinical | 98.6 | 100 | 0 | 99.9844 | 5110656 | 45.28 | 57 | 427404 |
| VP07075 | VP07075 | SAMN20168066 | SRR15098927 | PC | Public | 2007 | China | Clinical | 98.4 | 100 | 0 | 99.9842 | 5105752 | 45.28 | 61 | 365149 |
| VP07076 | VP07076 | SAMN20168064 | SRR15098929 | PC | Public | 2007 | China | Clinical | 97.8 | 100 | 0 | 99.991 | 5070753 | 45.32 | 61 | 408870 |
| VP07077 | VP07077 | SAMN20168063 | SRR15098930 | PC | Public | 2007 | China | Clinical | 97.8 | 100 | 0 | 99.979 | 5071602 | 45.31 | 61 | 409284 |
| VP07078 | VP07078 | SAMN20168062 | SRR15098931 | PC | Public | 2007 | China | Clinical | 98 | 100 | 0 | 99.9837 | 5080953 | 45.3 | 68 | 313489 |
| VP07079 | VP07079 | SAMN20168061 | SRR15098932 | PC | Public | 2007 | China | Clinical | 97.8 | 100 | 0 | 99.9875 | 5070112 | 45.31 | 60 | 401477 |
| VP07080 | VP07080 | SAMN20168065 | SRR15098928 | PC | Public | 2007 | China | Clinical | 98.4 | 100 | 0 | 99.9849 | 5093476 | 45.29 | 59 | 425235 |
| VP07081 | VP07081 | SAMN20168047 | SRR15098947 | PC | Public | 2007 | China | Clinical | 97.7 | 100 | 0 | 99.9273 | 5208587 | 45.21 | 64 | 401611 |
| VP07082 | VP07082 | SAMN20168042 | SRR15098953 | PC | Public | 2007 | China | Clinical | 98.5 | 100 | 0 | 99.9921 | 5093107 | 45.29 | 54 | 425241 |
| VP07083 | VP07083 | SAMN20168036 | SRR15098960 | PC | Public | 2007 | China | Clinical | 97.8 | 100 | 0 | 99.9799 | 5070806 | 45.32 | 60 | 401611 |
| VP07084 | VP07084 | SAMN20168045 | SRR15098950 | PC | Public | 2007 | China | Clinical | 98 | 100 | 0 | 99.9913 | 5079702 | 45.3 | 58 | 401611 |
| VP07085 | VP07085 | SAMN20168041 | SRR15098954 | PC | Public | 2007 | China | Clinical | 98.5 | 100 | 0 | 99.9845 | 5092016 | 45.29 | 58 | 401604 |
| VP07086 | VP07086 | SAMN20168043 | SRR15098952 | PC | Public | 2007 | China | Clinical | 98.5 | 100 | 0 | 99.9931 | 5092140 | 45.29 | 57 | 425355 |
| VP07087 | VP07087 | SAMN20168040 | SRR15098955 | PC | Public | 2007 | China | Clinical | 98.5 | 100 | 0 | 99.9925 | 5092440 | 45.29 | 56 | 425883 |
| VP07089 | VP07089 | SAMN20168072 | SRR15098920 | PC | Public | 2007 | China | Clinical | 97.8 | 100 | 0 | 99.9793 | 5071649 | 45.32 | 62 | 401611 |
| VP07090 | VP07090 | SAMN20168067 | SRR15098925 | PC | Public | 2007 | China | Clinical | 97.7 | 100 | 0 | 99.9794 | 5062352 | 45.32 | 63 | 414014 |
| VP07091 | VP07091 | SAMN20168073 | SRR15098919 | PC | Public | 2007 | China | Clinical | 97.8 | 100 | 0 | 99.9796 | 5071475 | 45.32 | 61 | 409284 |
| VP07092 | VP07092 | SAMN20168070 | SRR15098922 | PC | Public | 2007 | China | Clinical | 98.4 | 100 | 0 | 99.9839 | 5092355 | 45.29 | 58 | 425241 |
| VP07093 | VP07093 | SAMN20168069 | SRR15098923 | PC | Public | 2007 | China | Clinical | 98.4 | 100 | 0 | 99.9933 | 5092505 | 45.29 | 59 | 401604 |
| VP07094 | VP07094 | SAMN20168074 | SRR15098918 | PC | Public | 2007 | China | Clinical | 98.6 | 100 | 0 | 99.9848 | 5218946 | 45.21 | 59 | 440605 |
| VP07096 | VP07096 | SAMN20168076 | SRR15098916 | PC | Public | 2007 | China | Clinical | 97.8 | 100 | 0 | 99.9885 | 5070978 | 45.32 | 61 | 408642 |
| VP07097 | VP07097 | SAMN20168078 | SRR15098913 | PC | Public | 2007 | China | Clinical | 98.5 | 100 | 0 | 99.9831 | 5106122 | 45.29 | 60 | 401604 |
| VP07098 | VP07098 | SAMN20168071 | SRR15098921 | PC | Public | 2007 | China | Clinical | 98.5 | 100 | 0 | 99.9925 | 5105815 | 45.28 | 63 | 394970 |

| | | | | | | | | | | | | | | | | |
|---|---|---|---|---|---|---|---|---|---|---|---|---|---|---|---|---|
| VP07099 | VP07099 | SAMN20168077 | SRR15098914 | PC | Public | 2007 | China | Clinical | 98.6 | 100 | 0 | 99.9936 | 5135040 | 45.29 | 57 | 427200 |
| VP07100 | VP07100 | SAMN20168081 | SRR15098910 | PC | Public | 2007 | China | Clinical | 98.5 | 100 | 0 | 99.9831 | 5093156 | 45.29 | 58 | 401604 |
| VP07102 | VP07102 | SAMN20168084 | SRR15098907 | PC | Public | 2007 | China | Clinical | 97.8 | 100 | 0 | 99.9828 | 5075210 | 45.34 | 64 | 408756 |
| VP07103 | VP07103 | SAMN20168083 | SRR15098908 | PC | Public | 2007 | China | Clinical | 98.6 | 100 | 0 | 99.9861 | 5111203 | 45.28 | 58 | 427331 |
| VP07104 | VP07104 | SAMN20168079 | SRR15098912 | PC | Public | 2007 | China | Clinical | 97.8 | 100 | 0 | 99.9787 | 5071888 | 45.32 | 61 | 401478 |
| VP07105 | VP07105 | SAMN20168075 | SRR15098917 | PC | Public | 2007 | China | Clinical | 98.4 | 100 | 0 | 99.9923 | 5092819 | 45.29 | 59 | 401470 |
| VP07106 | VP07106 | SAMN20168068 | SRR15098924 | NonPC | Public | 2007 | China | Clinical | 89.7 | 100 | 0.05 | 98.3084 | 4911651 | 45.39 | 39 | 403135 |
| VP07107 | VP07107 | SAMN20168080 | SRR15098911 | PC | Public | 2007 | China | Clinical | 97.8 | 100 | 0 | 99.9893 | 5070732 | 45.32 | 68 | 373628 |
| VP07108 | VP07108 | SAMN20168085 | SRR15098906 | PC | Public | 2007 | China | Clinical | 97.3 | 100 | 0 | 99.8994 | 5128872 | 45.28 | 79 | 245637 |
| VP07109 | VP07109 | SAMN20168082 | SRR15098909 | PC | Public | 2007 | China | Clinical | 98.6 | 100 | 0 | 99.985 | 5148520 | 45.32 | 61 | 427326 |
| VP07110 | VP07110 | SAMN20168034 | SRR15098962 | NonPC | Public | 2007 | China | Clinical | 92.8 | 100 | 0 | 98.3924 | 5125140 | 45.22 | 41 | 572280 |
| VP07111 | VP07111 | SAMN20168033 | SRR15098963 | NonPC | Public | 2007 | China | Clinical | 89.3 | 100 | 0.36 | 98.2759 | 5077382 | 45.07 | 60 | 368511 |
| VP07112 | VP07112 | SAMN20168035 | SRR15098961 | PC | Public | 2007 | China | Clinical | 98.6 | 100 | 0.3 | 99.9931 | 5175008 | 45.29 | 60 | 401597 |
| VP07114 | VP07114 | SAMN20168151 | SRR15099588 | NonPC | Public | 2007 | China | Clinical | 92.3 | 100 | 0.19 | 98.3498 | 5008559 | 45.25 | 51 | 471218 |
| VP07115 | VP07115 | SAMN20168145 | SRR15099595 | NonPC | Public | 2007 | China | Env | 91.4 | 100 | 0 | 98.359 | 4980530 | 45.35 | 61 | 262439 |
| VP07116 | VP07116 | SAMN20168146 | SRR15099594 | NonPC | Public | 2007 | China | Env | 89.9 | 100 | 0 | 98.3351 | 5100607 | 45.29 | 64 | 449230 |
| VP07117 | VP07117 | SAMN20168144 | SRR15099596 | NonPC | Public | 2007 | China | Env | 89.1 | 100 | 0.27 | 98.3412 | 4957728 | 45.41 | 70 | 335281 |
| VP07118 | VP07118 | SAMN20168143 | SRR15099597 | NonPC | Public | 2007 | China | Clinical | 89.5 | 100 | 0.36 | 98.3064 | 5106427 | 45.05 | 49 | 452324 |
| VP07119 | VP07119 | SAMN20168095 | SRR15098895 | PC | Public | 2007 | China | Clinical | 97.6 | 100 | 0 | 99.9867 | 5058048 | 45.33 | 52 | 408756 |
| VP07120 | VP07120 | SAMN20168091 | SRR15098899 | PC | Public | 2007 | China | Clinical | 98.4 | 100 | 0 | 99.9843 | 5092916 | 45.29 | 60 | 394957 |
| VP07121 | VP07121 | SAMN20168093 | SRR15098897 | PC | Public | 2007 | China | Clinical | 97.6 | 100 | 0 | 99.9841 | 5059144 | 45.33 | 52 | 408756 |
| VP07122 | VP07122 | SAMN20168094 | SRR15098896 | PC | Public | 2007 | China | Clinical | 98.6 | 100 | 0 | 99.9937 | 5131807 | 45.29 | 59 | 401597 |
| VP07123 | VP07123 | SAMN20168097 | SRR15098891 | PC | Public | 2007 | China | Clinical | 97.6 | 100 | 0 | 99.9785 | 5058427 | 45.33 | 53 | 408642 |
| VP07124 | VP07124 | SAMN20168100 | SRR15098888 | PC | Public | 2007 | China | Clinical | 97.6 | 100 | 0 | 99.9898 | 5058856 | 45.33 | 54 | 401477 |
| VP07125 | VP07125 | SAMN20168102 | SRR15098886 | PC | Public | 2007 | China | Clinical | 98.4 | 100 | 0 | 99.9827 | 5121022 | 45.26 | 44 | 401641 |
| VP07126 | VP07126 | SAMN20168110 | SRR15098877 | PC | Public | 2007 | China | Clinical | 97.6 | 100 | 0 | 99.9902 | 5058948 | 45.33 | 53 | 408756 |
| VP07127 | VP07127 | SAMN20168125 | SRR15098861 | PC | Public | 2007 | China | Clinical | 97.8 | 100 | 0 | 99.9891 | 5072011 | 45.32 | 64 | 409284 |
| VP07128 | VP07128 | SAMN20168111 | SRR15098876 | PC | Public | 2007 | China | Clinical | 97.8 | 100 | 0 | 99.9913 | 5070714 | 45.32 | 64 | 401611 |
| VP07129 | VP07129 | SAMN20168131 | SRR15098854 | PC | Public | 2007 | China | Clinical | 97.6 | 100 | 0 | 99.991 | 5059246 | 45.33 | 51 | 368343 |
| VP07130 | VP07130 | SAMN20168122 | SRR15098864 | PC | Public | 2007 | China | Clinical | 97.6 | 100 | 0 | 99.9861 | 5062716 | 45.33 | 50 | 401478 |
| VP07131 | VP07131 | SAMN20168126 | SRR15098860 | PC | Public | 2007 | China | Clinical | 97.8 | 100 | 0 | 99.9888 | 5071576 | 45.32 | 65 | 368651 |
| VP07132 | VP07132 | SAMN20168132 | SRR15098853 | PC | Public | 2007 | China | Clinical | 98.6 | 100 | 0 | 99.9853 | 5131389 | 45.29 | 60 | 440605 |
| VP07133 | VP07133 | SAMN20168129 | SRR15098856 | PC | Public | 2007 | China | Clinical | 97.6 | 100 | 0 | 99.9776 | 5058351 | 45.33 | 55 | 401477 |
| VP07134 | VP07134 | SAMN20168142 | SRR15099598 | PC | Public | 2007 | China | Clinical | 97.8 | 100 | 0 | 99.9799 | 5071931 | 45.32 | 59 | 409284 |
| VP07135 | VP07135 | SAMN20168105 | SRR15098883 | PC | Public | 2007 | China | Clinical | 98.6 | 100 | 0 | 99.9844 | 5111108 | 45.28 | 57 | 427335 |
| VP07136 | VP07136 | SAMN20168108 | SRR15098879 | PC | Public | 2007 | China | Clinical | 97.6 | 100 | 0 | 99.9881 | 5059536 | 45.33 | 51 | 409284 |
| VP07137 | VP07137 | SAMN20168112 | SRR15098875 | PC | Public | 2007 | China | Clinical | 97.8 | 100 | 0 | 99.9833 | 5071043 | 45.31 | 59 | 409284 |
| VP07138 | VP07138 | SAMN20168104 | SRR15098884 | PC | Public | 2007 | China | Clinical | 98.4 | 100 | 0 | 99.9826 | 5092787 | 45.29 | 60 | 401604 |
| VP07139 | VP07139 | SAMN20168128 | SRR15098857 | PC | Public | 2007 | China | Clinical | 97.6 | 100 | 0 | 99.9873 | 5059538 | 45.33 | 52 | 408870 |
| VP07141 | VP07141 | SAMN20168135 | SRR15098850 | PC | Public | 2007 | China | Clinical | 97.5 | 100 | 0 | 99.93 | 5074964 | 45.28 | 83 | 268004 |
| VP07142 | VP07142 | SAMN20168140 | SRR15099600 | PC | Public | 2007 | China | Clinical | 97.8 | 100 | 0 | 99.9913 | 5071846 | 45.32 | 62 | 368284 |
| VP07143 | VP07143 | SAMN20168157 | SRR15099581 | PC | Public | 2007 | China | Clinical | 97.6 | 100 | 0 | 99.9789 | 5062681 | 45.34 | 54 | 409284 |
| VP07144 | VP07144 | SAMN20168087 | SRR15098903 | NonPC | Public | 2007 | China | Clinical | 92.5 | 100 | 0.65 | 98.4292 | 5120316 | 45.23 | 59 | 203747 |
| VP07145 | VP07145 | SAMN20168096 | SRR15098894 | PC | Public | 2007 | China | Clinical | 97.8 | 100 | 0 | 99.9891 | 5071033 | 45.32 | 63 | 401611 |
| VP07146 | VP07146 | SAMN20168153 | SRR15099586 | PC | Public | 2007 | China | Clinical | 97.6 | 100 | 0 | 99.979 | 5061788 | 45.33 | 52 | 409284 |
| VP07149 | VP07149 | SAMN20168158 | SRR15099580 | PC | Public | 2007 | China | Clinical | 97.6 | 100 | 0 | 99.988 | 5060157 | 45.33 | 53 | 401611 |
| VP07150 | VP07150 | SAMN20168156 | SRR15099583 | PC | Public | 2007 | China | Clinical | 97.7 | 100 | 0 | 99.9783 | 5060200 | 45.33 | 51 | 408870 |
| VP07151 | VP07151 | SAMN20168155 | SRR15099584 | PC | Public | 2007 | China | Clinical | 97.6 | 100 | 0 | 99.9801 | 5059347 | 45.33 | 51 | 408870 |

| | | | | | | | | | | | | | | | | |
|---|---|---|---|---|---|---|---|---|---|---|---|---|---|---|---|---|
| VP07152 | VP07152 | SAMN20167973 | SRR15099030 | PC | Public | 2007 | China | Clinical | 97.6 | 100 | 0 | 99.9776 | 5060316 | 45.33 | 52 | 401611 |
| VP07153 | VP07153 | SAMN20168106 | SRR15098882 | PC | Public | 2007 | China | Clinical | 98 | 100 | 0 | 99.9921 | 5080723 | 45.3 | 61 | 410889 |
| VP07154 | VP07154 | SAMN20168101 | SRR15098887 | PC | Public | 2007 | China | Clinical | 97.6 | 100 | 0 | 99.9306 | 5161844 | 45.11 | 77 | 288527 |
| VP07156 | VP07156 | SAMN20168124 | SRR15098862 | PC | Public | 2007 | China | Clinical | 97.6 | 100 | 0 | 99.9901 | 5062088 | 45.32 | 52 | 408756 |
| VP07157 | VP07157 | SAMN20168123 | SRR15098863 | PC | Public | 2007 | China | Clinical | 97.6 | 100 | 0 | 99.992 | 5059289 | 45.33 | 50 | 408756 |
| VP07158 | VP07158 | SAMN20168141 | SRR15099599 | PC | Public | 2007 | China | Clinical | 97.6 | 100 | 0 | 99.9786 | 5059861 | 45.33 | 52 | 368330 |
| VP07159 | VP07159 | SAMN20168092 | SRR15098898 | PC | Public | 2007 | China | Clinical | 97.8 | 100 | 0 | 99.9881 | 5071762 | 45.32 | 57 | 415884 |
| VP07160 | VP07160 | SAMN20168098 | SRR15098890 | PC | Public | 2007 | China | Clinical | 97.8 | 100 | 0 | 99.9844 | 5071150 | 45.32 | 62 | 408756 |
| VP07161 | VP07161 | SAMN20168099 | SRR15098889 | PC | Public | 2007 | China | Clinical | 97.6 | 100 | 0 | 99.9887 | 5058583 | 45.33 | 51 | 401477 |
| VP07162 | VP07162 | SAMN20168109 | SRR15098878 | PC | Public | 2007 | China | Clinical | 97.6 | 100 | 0 | 99.9871 | 5058894 | 45.33 | 51 | 408756 |
| VP07163 | VP07163 | SAMN20168114 | SRR15098873 | PC | Public | 2007 | China | Clinical | 97.6 | 100 | 0 | 99.9886 | 5058929 | 45.33 | 50 | 401611 |
| VP07164 | VP07164 | SAMN20168107 | SRR15098880 | PC | Public | 2007 | China | Clinical | 97.8 | 100 | 0 | 99.9793 | 5070922 | 45.31 | 60 | 408756 |
| VP07165 | VP07165 | SAMN20168115 | SRR15098872 | PC | Public | 2007 | China | Clinical | 97.8 | 100 | 0 | 99.9882 | 5070571 | 45.32 | 61 | 401610 |
| VP07166 | VP07166 | SAMN20168103 | SRR15098885 | PC | Public | 2007 | China | Clinical | 98.5 | 100 | 0 | 99.9841 | 5105010 | 45.28 | 60 | 394619 |
| VP07167 | VP07167 | SAMN20168139 | SRR15099601 | PC | Public | 2007 | China | Clinical | 97.8 | 100 | 0 | 99.9828 | 5071088 | 45.32 | 59 | 401611 |
| VP07168 | VP07168 | SAMN20168138 | SRR15099602 | PC | Public | 2007 | China | Clinical | 97.6 | 100 | 0 | 99.9836 | 5062596 | 45.32 | 52 | 408642 |
| VP07169 | VP07169 | SAMN20168130 | SRR15098855 | PC | Public | 2007 | China | Clinical | 97.6 | 100 | 0 | 99.9878 | 5059462 | 45.33 | 53 | 408756 |
| VP07170 | VP07170 | SAMN20168134 | SRR15098851 | PC | Public | 2007 | China | Clinical | 98.5 | 100 | 0 | 99.9925 | 5092193 | 45.29 | 56 | 425469 |
| VP07171 | VP07171 | SAMN20168127 | SRR15098858 | PC | Public | 2007 | China | Clinical | 97.8 | 100 | 0 | 99.9829 | 5070938 | 45.31 | 58 | 408870 |
| VP07173 | VP07173 | SAMN20168137 | SRR15099603 | PC | Public | 2007 | China | Clinical | 98.6 | 100 | 0.07 | 99.9931 | 5121587 | 45.28 | 58 | 427326 |
| VP07175 | VP07175 | SAMN20168136 | SRR15099605 | PC | Public | 2007 | China | Clinical | 98.6 | 100 | 0 | 99.9843 | 5110796 | 45.28 | 58 | 347371 |
| VP07177 | VP07177 | SAMN20168149 | SRR15099590 | PC | Public | 2007 | China | Clinical | 98.6 | 100 | 0 | 99.9933 | 5110218 | 45.28 | 54 | 427326 |
| VP07178 | VP07178 | SAMN20168150 | SRR15099589 | PC | Public | 2007 | China | Clinical | 97.8 | 100 | 0 | 99.9907 | 5070787 | 45.32 | 60 | 408756 |
| VP07179 | VP07179 | SAMN20168090 | SRR15098900 | PC | Public | 2007 | China | Clinical | 97.3 | 100 | 0 | 99.9294 | 5097778 | 45.27 | 86 | 215465 |
| VP07180 | VP07180 | SAMN20168154 | SRR15099585 | NonPC | Public | 2007 | China | Clinical | 92.1 | 100 | 0.46 | 98.3761 | 5122290 | 45.23 | 61 | 323277 |
| VP07181 | VP07181 | SAMN20168086 | SRR15098905 | PC | Public | 2007 | China | Clinical | 97.3 | 100 | 0 | 99.8995 | 5128788 | 45.28 | 76 | 260119 |
| VP07182 | VP07182 | SAMN20167971 | SRR15099032 | PC | Public | 2007 | China | Clinical | 97.3 | 100 | 0 | 99.9317 | 5098314 | 45.27 | 83 | 267776 |
| VP07183 | VP07183 | SAMN20168148 | SRR15099591 | NonPC | Public | 2007 | China | Clinical | 92.7 | 100 | 0 | 98.3686 | 5111458 | 45.21 | 47 | 370300 |
| VP07184 | VP07184 | SAMN20168119 | SRR15098867 | PC | Public | 2007 | China | Clinical | 97.3 | 100 | 0 | 99.9356 | 5097682 | 45.28 | 82 | 267662 |
| VP07185 | VP07185 | SAMN20168147 | SRR15099592 | PC | Public | 2007 | China | Env | 97.7 | 100 | 0 | 99.9439 | 5119005 | 45.29 | 63 | 373628 |
| VP07186 | VP07186 | SAMN20168113 | SRR15098874 | PC | Public | 2007 | China | Clinical | 97.8 | 100 | 0 | 99.9828 | 5071454 | 45.31 | 60 | 408750 |
| VP07187 | VP07187 | SAMN20168162 | SRR15099576 | PC | Public | 2007 | China | Clinical | 98.5 | 100 | 0 | 99.9792 | 5182697 | 45.11 | 60 | 425763 |
| VP07188 | VP07188 | SAMN20168161 | SRR15099577 | PC | Public | 2007 | China | Clinical | 98 | 100 | 0.27 | 99.9775 | 5122872 | 45.31 | 61 | 347558 |
| VP07189 | VP07189 | SAMN20168088 | SRR15098902 | NonPC | Public | 2007 | China | Clinical | 92.2 | 100 | 0.6 | 98.3647 | 5175992 | 45.23 | 61 | 323098 |
| VP07190 | VP07190 | SAMN20168120 | SRR15098866 | PC | Public | 2007 | China | Clinical | 97.3 | 100 | 0 | 99.9379 | 5099771 | 45.27 | 78 | 312483 |
| VP07192 | VP07192 | SAMN20168133 | SRR15098852 | NonPC | Public | 2007 | China | Clinical | 88.1 | 100 | 0.41 | 98.2551 | 5057761 | 45.28 | 43 | 384201 |
| VP07194 | VP07194 | SAMN20168118 | SRR15098868 | NonPC | Public | 2007 | China | Clinical | 90.1 | 100 | 0.05 | 98.2866 | 5116536 | 45.24 | 49 | 393111 |
| VP07197 | VP07197 | SAMN20168121 | SRR15098865 | PC | Public | 2007 | China | Clinical | 98.6 | 100 | 0 | 99.9853 | 5110627 | 45.28 | 57 | 427863 |
| VP07198 | VP07198 | SAMN20168159 | SRR15099579 | PC | Public | 2007 | China | Clinical | 98.6 | 100 | 0 | 99.9851 | 5132681 | 45.29 | 56 | 401597 |
| VP07199 | VP07199 | SAMN20168032 | SRR15098964 | PC | Public | 2007 | China | Clinical | 98.5 | 100 | 0 | 99.9851 | 5092337 | 45.29 | 56 | 425865 |
| VP07200 | VP07200 | SAMN20168160 | SRR15099578 | NonPC | Public | 2007 | China | Clinical | 92.4 | 100 | 0.05 | 98.3528 | 5068666 | 45.27 | 48 | 539107 |
| VP07201 | VP07201 | SAMN20168152 | SRR15099587 | PC | Public | 2007 | China | Clinical | 97.5 | 100 | 0 | 99.9326 | 5077044 | 45.28 | 80 | 268004 |
| VP07202 | VP07202 | SAMN20168089 | SRR15098901 | PC | Public | 2007 | China | Clinical | 97.3 | 100 | 0 | 99.935 | 5096757 | 45.27 | 81 | 268004 |
| VP07203 | VP07203 | SAMN20168170 | SRR15099567 | PC | Public | 2007 | China | Clinical | 97.8 | 100 | 0 | 99.9868 | 5081297 | 45.32 | 59 | 401612 |
| VP07204 | VP07204 | SAMN20168165 | SRR15099573 | PC | Public | 2007 | China | Clinical | 98.6 | 100 | 0 | 99.9938 | 5132385 | 45.29 | 60 | 401597 |
| VP07205 | VP07205 | SAMN20168168 | SRR15099569 | PC | Public | 2007 | China | Clinical | 98.6 | 100 | 0 | 99.9845 | 5196979 | 45.19 | 61 | 401505 |
| VP07206 | VP07206 | SAMN20168171 | SRR15099566 | PC | Public | 2007 | China | Clinical | 97.6 | 100 | 0 | 99.9791 | 5059083 | 45.33 | 52 | 414012 |
| VP07207 | VP07207 | SAMN20168172 | SRR15099565 | PC | Public | 2007 | China | Clinical | 97.8 | 100 | 0 | 99.9827 | 5071132 | 45.32 | 60 | 409284 |

| | | | | | | | | | | | | | | | | |
|---|---|---|---|---|---|---|---|---|---|---|---|---|---|---|---|---|
| VP07208 | VP07208 | SAMN20168166 | SRR15099572 | NonPC | Public | 2007 | China | Clinical | 92.4 | 100 | 0.65 | 98.4166 | 5107500 | 45.23 | 67 | 203659 |
| VP07209 | VP07209 | SAMN20168175 | SRR15099562 | PC | Public | 2007 | China | Clinical | 97.6 | 100 | 0 | 99.9902 | 5058772 | 45.33 | 48 | 414012 |
| VP07210 | VP07210 | SAMN20168176 | SRR15099561 | PC | Public | 2007 | China | Clinical | 97.8 | 100 | 0 | 99.9823 | 5071587 | 45.32 | 64 | 401611 |
| VP07211 | VP07211 | SAMN20168173 | SRR15099564 | PC | Public | 2007 | China | Clinical | 97.6 | 100 | 0 | 99.9787 | 5061337 | 45.33 | 53 | 408756 |
| VP07212 | VP07212 | SAMN20168164 | SRR15099574 | PC | Public | 2007 | China | Clinical | 98.5 | 100 | 0 | 99.9925 | 5115601 | 45.27 | 61 | 401625 |
| VP07213 | VP07213 | SAMN20168174 | SRR15099563 | PC | Public | 2007 | China | Clinical | 97.8 | 100 | 0 | 99.9836 | 5070939 | 45.32 | 59 | 408756 |
| VP07214 | VP07214 | SAMN20168169 | SRR15099568 | PC | Public | 2007 | China | Clinical | 97.9 | 100 | 0 | 99.9883 | 5072627 | 45.31 | 58 | 402890 |
| VP07215 | VP07215 | SAMN20168167 | SRR15099570 | PC | Public | 2007 | China | Clinical | 98.5 | 100 | 0 | 99.9849 | 5092479 | 45.29 | 57 | 425355 |
| VP07216 | VP07216 | SAMN20168183 | SRR15099553 | NonPC | Public | 2007 | China | Clinical | 92.1 | 100 | 0.19 | 98.3414 | 5028326 | 45.25 | 60 | 440170 |
| VP07217 | VP07217 | SAMN20168184 | SRR15099552 | PC | Public | 2007 | China | Clinical | 98.5 | 100 | 0 | 99.9883 | 5182781 | 45.11 | 59 | 425349 |
| VP07218 | VP07218 | SAMN20168185 | SRR15099551 | PC | Public | 2007 | China | Clinical | 98.5 | 100 | 0 | 99.9884 | 5182137 | 45.11 | 57 | 425235 |
| VP07219 | VP07219 | SAMN20168186 | SRR15099550 | PC | Public | 2007 | China | Clinical | 98.5 | 100 | 0 | 99.9881 | 5181966 | 45.11 | 63 | 394986 |
| VP07220 | VP07220 | SAMN20168182 | SRR15099554 | NonPC | Public | 2007 | China | Clinical | 92.4 | 100 | 0.65 | 98.4219 | 5107672 | 45.23 | 67 | 203654 |
| VP07226 | VP07226 | SAMN20168163 | SRR15099575 | PC | Public | 2007 | China | Clinical | 97.6 | 100 | 0 | 99.9914 | 5065842 | 45.32 | 51 | 414014 |
| VP07229 | VP07229 | SAMN20168181 | SRR15099555 | PC | Public | 2007 | China | Clinical | 98 | 100 | 0 | 99.9117 | 5164058 | 45.16 | 57 | 425434 |
| VP07233 | VP07233 | SAMN20168187 | SRR15099548 | PC | Public | 2007 | China | Clinical | 97.8 | 100 | 0 | 99.8967 | 5148457 | 45.29 | 53 | 347487 |
| VP07235 | VP07235 | SAMN20168189 | SRR15099546 | PC | Public | 2007 | China | Clinical | 98.4 | 100 | 0 | 99.9794 | 5183262 | 45.11 | 60 | 401477 |
| VP07236 | VP07236 | SAMN20168188 | SRR15099547 | PC | Public | 2007 | China | Clinical | 98 | 100 | 0 | 99.9783 | 5080211 | 45.3 | 56 | 401611 |
| VP07241 | VP07241 | SAMN20168190 | SRR15099545 | PC | Public | 2007 | China | Clinical | 98.6 | 100 | 0 | 99.9847 | 5131396 | 45.29 | 59 | 401597 |
| VP07244 | VP07244 | SAMN20168192 | SRR15099543 | NonPC | Public | 2007 | China | Clinical | 92.3 | 100 | 0.19 | 98.3669 | 5009004 | 45.25 | 52 | 450399 |
| VP07246 | VP07246 | SAMN20168191 | SRR15099544 | PC | Public | 2007 | China | Clinical | 97.7 | 100 | 0 | 99.9422 | 5120461 | 45.29 | 58 | 427338 |
| VP07247 | VP07247 | SAMN20168193 | SRR15099542 | NonPC | Public | 2007 | China | Clinical | 92.3 | 100 | 0.19 | 98.3507 | 5008313 | 45.25 | 48 | 539180 |
| VP07249 | VP07249 | SAMN20168194 | SRR15099541 | PC | Public | 2007 | China | Clinical | 98.4 | 100 | 0 | 99.989 | 5181433 | 45.11 | 59 | 425235 |
| VP07250 | VP07250 | SAMN20168198 | SRR15099535 | NonPC | Public | 2007 | China | Clinical | 91.7 | 100 | 0.24 | 98.4061 | 5145381 | 45.28 | 37 | 402057 |
| VP07251 | VP07251 | SAMN20168200 | SRR15099533 | NonPC | Public | 2007 | China | Clinical | 93.1 | 100 | 0.06 | 98.3617 | 5188392 | 45.28 | 49 | 515949 |
| VP07253 | VP07253 | SAMN20168199 | SRR15099534 | PC | Public | 2007 | China | Clinical | 98.5 | 100 | 0 | 99.9777 | 5117111 | 45.28 | 59 | 401470 |
| VP07254 | VP07254 | SAMN20168201 | SRR15099532 | NonPC | Public | 2007 | China | Clinical | 92.4 | 100 | 0.05 | 98.3558 | 5067426 | 45.27 | 48 | 478553 |
| VP07255 | VP07255 | SAMN20168202 | SRR15099531 | NonPC | Public | 2007 | China | Clinical | 92.3 | 100 | 0.19 | 98.3343 | 5043904 | 45.25 | 49 | 471218 |
| VP07256 | VP07256 | SAMN20168177 | SRR15099559 | PC | Public | 2007 | China | Clinical | 98 | 100 | 0 | 99.9909 | 5080968 | 45.3 | 62 | 410775 |
| VP07257 | VP07257 | SAMN20168178 | SRR15099558 | PC | Public | 2007 | China | Clinical | 98 | 100 | 0 | 99.9818 | 5080220 | 45.3 | 56 | 410775 |
| VP07258 | VP07258 | SAMN20168179 | SRR15099557 | PC | Public | 2007 | China | Clinical | 98 | 100 | 0 | 99.9859 | 5079641 | 45.3 | 62 | 401611 |
| VP07259 | VP07259 | SAMN20168180 | SRR15099556 | PC | Public | 2007 | China | Clinical | 98 | 100 | 0 | 99.982 | 5080209 | 45.3 | 60 | 411303 |
| VP07260 | VP07260 | SAMN20168195 | SRR15099540 | PC | Public | 2007 | China | Clinical | 97.8 | 100 | 0 | 99.9916 | 5070487 | 45.32 | 61 | 409290 |
| VP07261 | VP07261 | SAMN20168196 | SRR15099539 | PC | Public | 2007 | China | Clinical | 97.8 | 100 | 0 | 99.9837 | 5071649 | 45.32 | 64 | 408648 |
| VP07262 | VP07262 | SAMN20168197 | SRR15099536 | PC | Public | 2007 | China | Clinical | 97.5 | 100 | 0 | 99.9892 | 5050126 | 45.3 | 82 | 266114 |
| VP07263 | VP07263 | SAMN20168204 | SRR15099529 | PC | Public | 2007 | China | Env | 98 | 100 | 0 | 99.9783 | 5079676 | 45.3 | 61 | 411417 |
| VP07264 | VP07264 | SAMN20168203 | SRR15099530 | PC | Public | 2007 | China | Clinical | 98 | 100 | 0 | 99.9814 | 5080308 | 45.3 | 59 | 401611 |
| VP07265 | VP07265 | SAMN20168017 | SRR15098980 | PC | Public | 2007 | China | Clinical | 97.7 | 100 | 0 | 99.9149 | 5147999 | 45.29 | 54 | 347485 |
| VP07266 | VP07266 | SAMN20168016 | SRR15098982 | PC | Public | 2007 | China | Clinical | 98.5 | 100 | 0 | 99.9833 | 5093567 | 45.29 | 57 | 425355 |
| VP07267 | VP07267 | SAMN20168031 | SRR15098965 | PC | Public | 2007 | China | Clinical | 98.5 | 100 | 0 | 99.9936 | 5093032 | 45.29 | 58 | 401604 |
| VP07268 | VP07268 | SAMN20168030 | SRR15098966 | NonPC | Public | 2007 | China | Clinical | 92.3 | 100 | 0.61 | 98.333 | 5285508 | 45.26 | 44 | 472180 |
| VP07269 | VP07269 | SAMN20168056 | SRR15098938 | NonPC | Public | 2007 | China | Clinical | 92.3 | 100 | 0.61 | 98.3485 | 5278318 | 45.26 | 44 | 472346 |
| VP07270 | VP07270 | SAMN20168057 | SRR15098936 | PC | Public | 2007 | China | Clinical | 98.5 | 100 | 0.14 | 99.984 | 5265845 | 45.11 | 61 | 401604 |
| VP07271 | VP07271 | SAMN20168116 | SRR15098871 | PC | Public | 2007 | China | Clinical | 98.4 | 100 | 0 | 99.9844 | 5092576 | 45.29 | 57 | 425469 |
| VP07272 | VP07272 | SAMN20168117 | SRR15098869 | PC | Public | 2007 | China | Clinical | 98.5 | 100 | 0 | 99.985 | 5106863 | 45.28 | 60 | 394972 |
| VP07274 | VP07274 | SAMN20168207 | SRR15099525 | PC | Public | 2007 | China | Clinical | 98.4 | 100 | 0 | 99.9911 | 5087179 | 45.3 | 49 | 425355 |
| VP07276 | VP07276 | SAMN20168205 | SRR15099528 | NonPC | Public | 2007 | China | Clinical | 92.3 | 100 | 0.19 | 98.3559 | 5008628 | 45.25 | 53 | 471204 |
| VP07277 | VP07277 | SAMN20167970 | SRR15099033 | NonPC | Public | 2007 | China | Clinical | 92.3 | 100 | 0.19 | 98.3551 | 5009983 | 45.25 | 52 | 539180 |

| | | | | | | | | | | | | | | | |
|---|---|---|---|---|---|---|---|---|---|---|---|---|---|---|---|
| VP07278 | VP07278 | SAMN20168206 | SRR15099527 | NonPC | Public | 2007 | China | Clinical | 92.2 | 100 | 0.19 | 98.3864 | 5009720 | 45.25 | 53 | 450399 |
| VP08001 | VP08001 | SAMN20168292 | SRR15099432 | NonPC | Public | 2007 | China | Clinical | 92.3 | 100 | 0.19 | 98.3452 | 5038776 | 45.26 | 50 | 539180 |
| VP08002 | VP08002 | SAMN20168293 | SRR15099431 | NonPC | Public | 2007 | China | Clinical | 92.3 | 100 | 0.19 | 98.3743 | 5009506 | 45.25 | 51 | 539180 |
| VP08003 | VP08003 | SAMN20168228 | SRR15099502 | NonPC | Public | 2008 | China | Env | 91.1 | 100 | 0 | 98.3586 | 4968609 | 45.4 | 51 | 453585 |
| VP08004 | VP08004 | SAMN20168275 | SRR15099451 | NonPC | Public | 2008 | China | Env | 89.2 | 100 | 0 | 98.2965 | 4915250 | 45.43 | 43 | 304813 |
| VP08007 | VP08007 | SAMN20168298 | SRR15099424 | PC | Public | 2008 | China | Clinical | 98.5 | 100 | 0 | 99.9907 | 5169270 | 45.29 | 67 | 288661 |
| VP08008 | VP08008 | SAMN20168299 | SRR15099423 | PC | Public | 2008 | China | Clinical | 98.5 | 100 | 0 | 99.9941 | 5121058 | 45.28 | 68 | 288661 |
| VP08009 | VP08009 | SAMN20168296 | SRR15099428 | NonPC | Public | 2008 | China | Clinical | 90.9 | 100 | 0 | 98.3154 | 5076948 | 45.26 | 64 | 374781 |
| VP08010 | VP08010 | SAMN20168297 | SRR15099425 | PC | Public | 2008 | China | Clinical | 98.3 | 100 | 0 | 99.9911 | 5083178 | 45.28 | 65 | 288661 |
| VP08011 | VP08011 | SAMN20168300 | SRR15099422 | PC | Public | 2008 | China | Clinical | 98.5 | 100 | 0.01 | 99.9937 | 5212605 | 45.31 | 69 | 288661 |
| VP08012 | VP08012 | SAMN20168301 | SRR15099421 | PC | Public | 2008 | China | Clinical | 98.4 | 100 | 0 | 99.9891 | 5121002 | 45.28 | 67 | 288661 |
| VP08014 | VP08014 | SAMN20168285 | SRR15099440 | NonPC | Public | 2008 | China | Clinical | 92.3 | 100 | 0.19 | 98.3636 | 5013959 | 45.24 | 60 | 450300 |
| VP08015 | VP08015 | SAMN20168295 | SRR15099429 | PC | Public | 2008 | China | Clinical | 98.2 | 100 | 0 | 99.9913 | 5128092 | 45.26 | 79 | 267761 |
| VP08016 | VP08016 | SAMN20168302 | SRR15099420 | NonPC | Public | 2008 | China | Clinical | 89.5 | 100 | 0.36 | 98.2919 | 5106701 | 45.05 | 41 | 539812 |
| VP08017 | VP08017 | SAMN20168229 | SRR15099501 | NonPC | Public | 2008 | China | Env | 91.1 | 100 | 0.1 | 98.3245 | 5130229 | 45.19 | 60 | 313447 |
| VP08018 | VP08018 | SAMN20168303 | SRR15099419 | NonPC | Public | 2008 | China | Clinical | 92.3 | 100 | 0.05 | 98.3406 | 5056344 | 45.27 | 54 | 450186 |
| VP08019 | VP08019 | SAMN20168306 | SRR15099416 | PC | Public | 2008 | China | Clinical | 98.4 | 100 | 0 | 99.9836 | 5086290 | 45.3 | 46 | 425883 |
| VP08020 | VP08020 | SAMN20168305 | SRR15099417 | PC | Public | 2008 | China | Clinical | 98.4 | 100 | 0 | 99.9929 | 5086902 | 45.3 | 48 | 401604 |
| VP08021 | VP08021 | SAMN20168309 | SRR15099412 | PC | Public | 2008 | China | Clinical | 98.5 | 100 | 0 | 99.9902 | 5122082 | 45.28 | 68 | 288661 |
| VP08022 | VP08022 | SAMN20168308 | SRR15099413 | PC | Public | 2008 | China | Clinical | 98.6 | 100 | 0 | 99.9839 | 5132257 | 45.29 | 60 | 401597 |
| VP08023 | VP08023 | SAMN20168310 | SRR15099411 | PC | Public | 2008 | China | Clinical | 98.6 | 100 | 0 | 99.9849 | 5132455 | 45.29 | 61 | 401597 |
| VP08024 | VP08024 | SAMN20168314 | SRR15099407 | PC | Public | 2008 | China | Clinical | 98.5 | 100 | 0 | 99.9783 | 5092146 | 45.29 | 55 | 425661 |
| VP08025 | VP08025 | SAMN20168304 | SRR15099418 | PC | Public | 2008 | China | Clinical | 97.7 | 100 | 0 | 99.9065 | 5148274 | 45.29 | 56 | 419982 |
| VP08026 | VP08026 | SAMN20168000 | SRR15098999 | NonPC | Public | 2007 | China | Clinical | 92.3 | 100 | 0.05 | 98.0574 | 5127887 | 45.23 | 38 | 470454 |
| VP08027 | VP08027 | SAMN20168001 | SRR15098998 | NonPC | Public | 2007 | China | Clinical | 92.3 | 100 | 0.05 | 98.0725 | 5125233 | 45.23 | 35 | 470454 |
| VP08028 | VP08028 | SAMN20168002 | SRR15098997 | NonPC | Public | 2007 | China | Clinical | 92.2 | 100 | 0.05 | 98.0911 | 5118357 | 45.23 | 45 | 337351 |
| VP08029 | VP08029 | SAMN20168003 | SRR15098996 | NonPC | Public | 2007 | China | Clinical | 92.2 | 100 | 0.05 | 98.0737 | 5117337 | 45.23 | 46 | 337351 |
| VP08030 | VP08030 | SAMN20168004 | SRR15098995 | NonPC | Public | 2007 | China | Clinical | 92.2 | 100 | 0.05 | 98.1123 | 5117950 | 45.23 | 47 | 337590 |
| VP08031 | VP08031 | SAMN20168005 | SRR15098994 | NonPC | Public | 2007 | China | Clinical | 92.2 | 100 | 0.05 | 98.1096 | 5117724 | 45.23 | 48 | 337447 |
| VP08032 | VP08032 | SAMN20168006 | SRR15098993 | NonPC | Public | 2007 | China | Clinical | 92.2 | 100 | 0.05 | 98.0984 | 5119023 | 45.23 | 45 | 337590 |
| VP08033 | VP08033 | SAMN20168007 | SRR15098991 | NonPC | Public | 2007 | China | Clinical | 92.2 | 100 | 0.05 | 98.1056 | 5121607 | 45.23 | 47 | 337447 |
| VP08034 | VP08034 | SAMN20168008 | SRR15098990 | NonPC | Public | 2007 | China | Clinical | 92.2 | 100 | 0.05 | 98.0829 | 5118879 | 45.23 | 44 | 433160 |
| VP08035 | VP08035 | SAMN20168009 | SRR15098989 | NonPC | Public | 2007 | China | Clinical | 92.2 | 100 | 0.05 | 98.055 | 5119106 | 45.23 | 44 | 337590 |
| VP08036 | VP08036 | SAMN20168307 | SRR15099414 | PC | Public | 2008 | China | Clinical | 98.4 | 100 | 0 | 99.9855 | 5093600 | 45.29 | 58 | 425985 |
| VP08037 | VP08037 | SAMN20168338 | SRR15099380 | NonPC | Public | 2008 | China | Clinical | 92.8 | 100 | 0.05 | 98.3827 | 5083928 | 45.28 | 38 | 343046 |
| VP08038 | VP08038 | SAMN20168230 | SRR15099500 | NonPC | Public | 2008 | China | Env | 90.3 | 100 | 0.1 | 98.3083 | 5248534 | 45.26 | 38 | 463654 |
| VP08040 | VP08040 | SAMN20168340 | SRR15099378 | NonPC | Public | 2008 | China | Clinical | 92.1 | 100 | 0.05 | 98.4519 | 5065512 | 45.23 | 48 | 294932 |
| VP08041 | VP08041 | SAMN20168341 | SRR15099377 | NonPC | Public | 2008 | China | Clinical | 92.1 | 100 | 0.05 | 98.4414 | 5064506 | 45.23 | 49 | 294932 |
| VP08042 | VP08042 | SAMN20168339 | SRR15099379 | NonPC | Public | 2008 | China | Clinical | 92.7 | 100 | 0 | 98.4144 | 5122031 | 45.19 | 56 | 258500 |
| VP08043 | VP08043 | SAMN20168342 | SRR15099376 | PC | Public | 2008 | China | Clinical | 98.5 | 100 | 0 | 99.993 | 5123060 | 45.28 | 67 | 288661 |
| VP08045 | VP08045 | SAMN20168319 | SRR15099401 | PC | Public | 2008 | China | Clinical | 97.5 | 100 | 0 | 99.9248 | 5142681 | 45.22 | 72 | 267890 |
| VP08046 | VP08046 | SAMN20168313 | SRR15099408 | NonPC | Public | 2008 | China | Clinical | 92.8 | 100 | 0 | 98.4054 | 5114313 | 45.19 | 55 | 359640 |
| VP08048 | VP08048 | SAMN20168315 | SRR15099406 | PC | Public | 2008 | China | Clinical | 98.5 | 100 | 0 | 99.9941 | 5121979 | 45.28 | 68 | 288661 |
| VP08050 | VP08050 | SAMN20168316 | SRR15099405 | PC | Public | 2008 | China | Clinical | 98.6 | 100 | 0 | 99.993 | 5132899 | 45.29 | 59 | 427728 |
| VP08051 | VP08051 | SAMN20168317 | SRR15099403 | PC | Public | 2008 | China | Clinical | 98.5 | 100 | 0 | 99.9888 | 5121331 | 45.28 | 67 | 288661 |
| VP08052 | VP08052 | SAMN20168318 | SRR15099402 | PC | Public | 2008 | China | Clinical | 98.5 | 100 | 0 | 99.9907 | 5121998 | 45.28 | 68 | 288661 |
| VP08053 | VP08053 | SAMN20168322 | SRR15099398 | PC | Public | 2008 | China | Clinical | 98.3 | 100 | 0 | 99.9929 | 5082353 | 45.28 | 63 | 312778 |
| VP08054 | VP08054 | SAMN20168320 | SRR15099400 | PC | Public | 2008 | China | Clinical | 98.5 | 100 | 0 | 99.9928 | 5121188 | 45.28 | 69 | 288661 |

| | | | | | | | | | | | | | | | | |
|---|---|---|---|---|---|---|---|---|---|---|---|---|---|---|---|---|
| VP08055 | VP08055 | SAMN20168321 | SRR15099399 | PC | Public | 2008 | China | Clinical | 98.6 | 100 | 0 | 99.9857 | 5131012 | 45.29 | 59 | 427728 |
| VP08056 | VP08056 | SAMN20168323 | SRR15099397 | PC | Public | 2008 | China | Clinical | 98.3 | 100 | 0 | 99.9821 | 5082304 | 45.28 | 66 | 288661 |
| VP08057 | VP08057 | SAMN20168353 | SRR15099364 | NonPC | Public | 2008 | China | Clinical | 92.1 | 100 | 0.08 | 98.3445 | 5147550 | 45.22 | 65 | 312715 |
| VP08058 | VP08058 | SAMN20168312 | SRR15099409 | PC | Public | 2008 | China | Clinical | 97.6 | 100 | 0.07 | 99.9257 | 5130770 | 45.26 | 84 | 268004 |
| VP08059 | VP08059 | SAMN20168311 | SRR15099410 | PC | Public | 2008 | China | Clinical | 98 | 100 | 0 | 99.9912 | 5115717 | 45.34 | 56 | 312806 |
| VP08060 | VP08060 | SAMN20168331 | SRR15099388 | PC | Public | 2008 | China | Clinical | 98.5 | 100 | 0 | 99.9935 | 5122092 | 45.28 | 68 | 288661 |
| VP08061 | VP08061 | SAMN20168335 | SRR15099384 | PC | Public | 2008 | China | Clinical | 98.5 | 100 | 0 | 99.9911 | 5140129 | 45.26 | 67 | 288674 |
| VP08062 | VP08062 | SAMN20168332 | SRR15099387 | PC | Public | 2008 | China | Clinical | 98.5 | 100 | 0 | 99.9935 | 5122437 | 45.28 | 69 | 288661 |
| VP08063 | VP08063 | SAMN20168336 | SRR15099383 | PC | Public | 2008 | China | Clinical | 98.5 | 100 | 0 | 99.987 | 5140789 | 45.26 | 66 | 288674 |
| VP08064 | VP08064 | SAMN20168333 | SRR15099386 | PC | Public | 2008 | China | Clinical | 98.4 | 100 | 0 | 99.9944 | 5123004 | 45.28 | 69 | 288661 |
| VP08065 | VP08065 | SAMN20168334 | SRR15099385 | PC | Public | 2008 | China | Clinical | 98.5 | 100 | 0 | 99.9899 | 5122630 | 45.28 | 67 | 288661 |
| VP08066 | VP08066 | SAMN20168337 | SRR15099381 | PC | Public | 2008 | China | Clinical | 98.5 | 100 | 0 | 99.9901 | 5141350 | 45.26 | 67 | 288674 |
| VP08067 | VP08067 | SAMN20168330 | SRR15099389 | PC | Public | 2008 | China | Clinical | 97.7 | 100 | 0 | 99.9379 | 5152002 | 45.22 | 63 | 288661 |
| VP08068 | VP08068 | SAMN20168325 | SRR15099395 | NonPC | Public | 2008 | China | Clinical | 92.4 | 100 | 0.05 | 98.3291 | 5061866 | 45.27 | 51 | 450226 |
| VP08069 | VP08069 | SAMN20168326 | SRR15099394 | NonPC | Public | 2008 | China | Clinical | 92.4 | 100 | 0.05 | 98.3626 | 5061873 | 45.27 | 50 | 450226 |
| VP08070 | VP08070 | SAMN20168327 | SRR15099392 | NonPC | Public | 2008 | China | Clinical | 92.4 | 100 | 0.05 | 98.3172 | 5062224 | 45.27 | 50 | 450226 |
| VP08071 | VP08071 | SAMN20168324 | SRR15099396 | NonPC | Public | 2008 | China | Clinical | 91.7 | 100 | 0 | 98.4177 | 5053227 | 45.24 | 48 | 417280 |
| VP08073 | VP08073 | SAMN20168328 | SRR15099391 | NonPC | Public | 2008 | China | Clinical | 92.4 | 100 | 0.05 | 98.3255 | 5061673 | 45.27 | 51 | 450226 |
| VP08075 | VP08075 | SAMN20168329 | SRR15099390 | NonPC | Public | 2008 | China | Clinical | 92.4 | 100 | 0.05 | 98.3266 | 5062523 | 45.27 | 48 | 450226 |
| VP08076 | VP08076 | SAMN20168344 | SRR15099374 | PC | Public | 2008 | China | Clinical | 97.6 | 100 | 0 | 99.9143 | 5160071 | 45.12 | 84 | 268004 |
| VP08077 | VP08077 | SAMN20168343 | SRR15099375 | NonPC | Public | 2008 | China | Clinical | 92.3 | 100 | 0.46 | 98.3678 | 5207396 | 45.19 | 66 | 470315 |
| VP08078 | VP08078 | SAMN20168345 | SRR15099373 | PC | Public | 2008 | China | Clinical | 97.7 | 100 | 0 | 99.9271 | 5159968 | 45.12 | 84 | 267776 |
| VP08079 | VP08079 | SAMN20168346 | SRR15099372 | PC | Public | 2008 | China | Clinical | 98.4 | 100 | 0 | 99.9886 | 5140766 | 45.26 | 68 | 288661 |
| VP08080 | VP08080 | SAMN20168350 | SRR15099367 | PC | Public | 2008 | China | Clinical | 98.5 | 100 | 0 | 99.9937 | 5209936 | 45.2 | 70 | 288661 |
| VP08081 | VP08081 | SAMN20168351 | SRR15099366 | PC | Public | 2008 | China | Clinical | 98.3 | 100 | 0 | 99.9929 | 5084578 | 45.28 | 62 | 312778 |
| VP08082 | VP08082 | SAMN20168348 | SRR15099369 | PC | Public | 2008 | China | Clinical | 97.5 | 100 | 0 | 99.9216 | 5143129 | 45.21 | 72 | 267890 |
| VP08083 | VP08083 | SAMN20168349 | SRR15099368 | PC | Public | 2008 | China | Clinical | 98.5 | 100 | 0 | 99.9863 | 5102209 | 45.28 | 68 | 312806 |
| VP08084 | VP08084 | SAMN20168347 | SRR15099370 | NonPC | Public | 2008 | China | Clinical | 88.3 | 99.98 | 0.32 | 98.2242 | 5093924 | 45.24 | 41 | 588690 |
| VP08085 | VP08085 | SAMN20168280 | SRR15099445 | NonPC | Public | 2008 | China | Env | 91.9 | 100 | 0 | 98.4309 | 5033375 | 45.26 | 32 | 524939 |
| VP08086 | VP08086 | SAMN20168282 | SRR15099443 | NonPC | Public | 2008 | China | Env | 92.9 | 100 | 0 | 98.3968 | 5136396 | 45.21 | 38 | 535923 |
| VP08087 | VP08087 | SAMN20168231 | SRR15099499 | NonPC | Public | 2008 | China | Env | 91.6 | 100 | 0.03 | 98.385 | 5083365 | 45.28 | 52 | 370527 |
| VP08088 | VP08088 | SAMN20168232 | SRR15099498 | NonPC | Public | 2008 | China | Env | 91.8 | 100 | 0.45 | 98.3826 | 5063237 | 45.27 | 66 | 314270 |
| VP08089 | VP08089 | SAMN20168278 | SRR15099447 | NonPC | Public | 2008 | China | Env | 91.9 | 100 | 0 | 98.405 | 5097803 | 45.25 | 31 | 527114 |
| VP08090 | VP08090 | SAMN20168233 | SRR15099497 | NonPC | Public | 2008 | China | Env | 89.5 | 100 | 0 | 98.3244 | 4978513 | 45.44 | 62 | 383320 |
| VP08091 | VP08091 | SAMN20168234 | SRR15099496 | NonPC | Public | 2008 | China | Env | 90.6 | 100 | 0.05 | 98.335 | 5042822 | 45.28 | 43 | 467399 |
| VP08092 | VP08092 | SAMN20168235 | SRR15099495 | NonPC | Public | 2008 | China | Env | 89.9 | 100 | 0 | 98.3007 | 5000155 | 45.36 | 38 | 387981 |
| VP08093 | VP08093 | SAMN20168236 | SRR15099494 | NonPC | Public | 2008 | China | Env | 92 | 100 | 0 | 98.3679 | 5074598 | 45.23 | 81 | 315732 |
| VP08095 | VP08095 | SAMN20168213 | SRR15099519 | NonPC | Public | 2008 | China | Env | 93 | 100 | 0 | 98.4073 | 5132036 | 45.22 | 48 | 400650 |
| VP08096 | VP08096 | SAMN20168237 | SRR15099492 | NonPC | Public | 2008 | China | Env | 91.4 | 100 | 0 | 98.2951 | 5104527 | 45.17 | 50 | 343786 |
| VP08097 | VP08097 | SAMN20168238 | SRR15099491 | NonPC | Public | 2008 | China | Env | 91.4 | 100 | 0.03 | 98.2502 | 5513453 | 45.02 | 83 | 252234 |
| VP08098 | VP08098 | SAMN20168239 | SRR15099490 | NonPC | Public | 2008 | China | Env | 91.8 | 100 | 0.14 | 98.3509 | 5051368 | 45.25 | 91 | 415899 |
| VP08099 | VP08099 | SAMN20168240 | SRR15099489 | NonPC | Public | 2008 | China | Env | 91.6 | 100 | 0.05 | 98.3886 | 4994637 | 45.34 | 66 | 413389 |
| VP08100 | VP08100 | SAMN20168241 | SRR15099488 | NonPC | Public | 2008 | China | Env | 89.9 | 100 | 0.14 | 98.2938 | 5032475 | 45.33 | 48 | 428870 |
| VP08101 | VP08101 | SAMN20168242 | SRR15099487 | NonPC | Public | 2008 | China | Env | 90.7 | 100 | 0.72 | 98.2881 | 5282362 | 45.19 | 60 | 198199 |
| VP08102 | VP08102 | SAMN20168243 | SRR15099486 | NonPC | Public | 2008 | China | Env | 91.1 | 100 | 0.03 | 98.2934 | 5346685 | 45.09 | 50 | 443949 |
| VP08103 | VP08103 | SAMN20168244 | SRR15099485 | NonPC | Public | 2008 | China | Env | 91.1 | 100 | 0 | 98.3751 | 5009791 | 45.41 | 60 | 232715 |
| VP08104 | VP08104 | SAMN20168245 | SRR15099484 | NonPC | Public | 2008 | China | Env | 91.4 | 100 | 0.44 | 98.3052 | 5019368 | 45.36 | 58 | 314953 |
| VP08105 | VP08105 | SAMN20168246 | SRR15099483 | NonPC | Public | 2008 | China | Env | 89.5 | 100 | 0.14 | 98.2671 | 4952907 | 45.38 | 47 | 388191 |

| | | | | | | | | | | | | | | | | |
|---|---|---|---|---|---|---|---|---|---|---|---|---|---|---|---|---|
| VP08106 | VP08106 | SAMN20168210 | SRR15099522 | PC | Public | 2008 | China | Env | 97.7 | 100 | 0 | 99.9226 | 5154452 | 45.14 | 54 | 360193 |
| VP08107 | VP08107 | SAMN20168247 | SRR15099481 | NonPC | Public | 2008 | China | Env | 91.2 | 100 | 0 | 98.2905 | 5098583 | 45.25 | 70 | 258381 |
| VP08108 | VP08108 | SAMN20168248 | SRR15099480 | NonPC | Public | 2008 | China | Env | 91.6 | 100 | 0 | 98.2793 | 5122101 | 45.19 | 59 | 317369 |
| VP08109 | VP08109 | SAMN20168249 | SRR15099479 | NonPC | Public | 2008 | China | Env | 90.9 | 100 | 0.14 | 98.3492 | 5050202 | 45.33 | 69 | 403694 |
| VP08110 | VP08110 | SAMN20168219 | SRR15099512 | NonPC | Public | 2008 | China | Env | 88.2 | 100 | 0.47 | 98.2967 | 4988781 | 45.22 | 84 | 313775 |
| VP08111 | VP08111 | SAMN20168273 | SRR15099453 | NonPC | Public | 2008 | China | Env | 89.7 | 100 | 0.05 | 98.31 | 5031949 | 45.38 | 48 | 423043 |
| VP08112 | VP08112 | SAMN20168274 | SRR15099452 | NonPC | Public | 2008 | China | Env | 89.1 | 100 | 0.05 | 98.3295 | 5041153 | 45.38 | 38 | 475046 |
| VP08113 | VP08113 | SAMN20168218 | SRR15099513 | NonPC | Public | 2008 | China | Env | 88.4 | 100 | 0.47 | 98.2554 | 5129524 | 45.22 | 63 | 452965 |
| VP08114 | VP08114 | SAMN20168216 | SRR15099516 | NonPC | Public | 2008 | China | Clinical | 92.9 | 100 | 0.05 | 98.3708 | 5027193 | 45.31 | 44 | 411537 |
| VP08115 | VP08115 | SAMN20168287 | SRR15099437 | PC | Public | 2008 | China | Clinical | 97.3 | 100 | 0 | 99.8991 | 5176916 | 45.27 | 74 | 288527 |
| VP08116 | VP08116 | SAMN20168226 | SRR15099505 | NonPC | Public | 2008 | China | Clinical | 90.9 | 100 | 0 | 98.3536 | 4941044 | 45.35 | 52 | 340380 |
| VP08117 | VP08117 | SAMN20168227 | SRR15099503 | NonPC | Public | 2008 | China | Clinical | 89.7 | 100 | 0.04 | 98.2712 | 5107070 | 45.24 | 40 | 462824 |
| VP08118 | VP08118 | SAMN20168286 | SRR15099439 | NonPC | Public | 2008 | China | Clinical | 92.4 | 100 | 0.05 | 98.338 | 5080127 | 45.26 | 53 | 450226 |
| VP08119 | VP08119 | SAMN20168277 | SRR15099448 | NonPC | Public | 2008 | China | Clinical | 91.9 | 100 | 0 | 98.3961 | 5094039 | 45.24 | 33 | 524940 |
| VP08120 | VP08120 | SAMN20168290 | SRR15099434 | PC | Public | 2008 | China | Clinical | 98.5 | 100 | 0 | 99.9915 | 5140763 | 45.26 | 68 | 288674 |
| VP08121 | VP08121 | SAMN20168352 | SRR15099365 | PC | Public | 2008 | China | Clinical | 97.8 | 100 | 0 | 99.9854 | 5070970 | 45.29 | 71 | 288661 |
| VP08122 | VP08122 | SAMN20168357 | SRR15099359 | NonPC | Public | 2008 | China | Clinical | 91 | 100 | 0.03 | 98.3409 | 5105501 | 45.33 | 55 | 261046 |
| VP08123 | VP08123 | SAMN20168354 | SRR15099363 | PC | Public | 2008 | China | Clinical | 98.5 | 100 | 0 | 99.9833 | 5092798 | 45.29 | 58 | 401463 |
| VP08125 | VP08125 | SAMN20168370 | SRR15099345 | PC | Public | 2008 | China | Clinical | 98.2 | 100 | 0 | 99.9874 | 5160177 | 45.11 | 86 | 267783 |
| VP08126 | VP08126 | SAMN20168371 | SRR15099344 | PC | Public | 2008 | China | Clinical | 98.6 | 100 | 0 | 99.9753 | 5181777 | 45.13 | 62 | 427357 |
| VP08127 | VP08127 | SAMN20168360 | SRR15099356 | PC | Public | 2008 | China | Clinical | 98.2 | 100 | 0 | 99.9909 | 5076847 | 45.29 | 58 | 288661 |
| VP08128 | VP08128 | SAMN20168361 | SRR15099355 | PC | Public | 2008 | China | Clinical | 98.2 | 100 | 0 | 99.9927 | 5076990 | 45.29 | 58 | 288661 |
| VP08129 | VP08129 | SAMN20168362 | SRR15099354 | PC | Public | 2008 | China | Clinical | 98.2 | 100 | 0 | 99.9922 | 5076888 | 45.29 | 55 | 312650 |
| VP08130 | VP08130 | SAMN20168363 | SRR15099353 | PC | Public | 2008 | China | Clinical | 98.2 | 100 | 0 | 99.9886 | 5077338 | 45.29 | 55 | 425183 |
| VP08131 | VP08131 | SAMN20168364 | SRR15099352 | PC | Public | 2008 | China | Clinical | 98.2 | 100 | 0 | 99.9896 | 5078324 | 45.29 | 60 | 288661 |
| VP08132 | VP08132 | SAMN20168365 | SRR15099351 | PC | Public | 2008 | China | Clinical | 98.2 | 100 | 0 | 99.9912 | 5077911 | 45.29 | 56 | 288661 |
| VP08133 | VP08133 | SAMN20168366 | SRR15099350 | PC | Public | 2008 | China | Clinical | 98.2 | 100 | 0 | 99.9885 | 5078712 | 45.29 | 56 | 288661 |
| VP08134 | VP08134 | SAMN20168367 | SRR15099348 | PC | Public | 2008 | China | Clinical | 98.2 | 100 | 0 | 99.9915 | 5077400 | 45.29 | 56 | 312785 |
| VP08135 | VP08135 | SAMN20168368 | SRR15099347 | PC | Public | 2008 | China | Clinical | 98.2 | 100 | 0 | 99.9882 | 5077462 | 45.29 | 56 | 288661 |
| VP08136 | VP08136 | SAMN20168369 | SRR15099346 | PC | Public | 2008 | China | Clinical | 98.2 | 100 | 0 | 99.9922 | 5076297 | 45.29 | 58 | 288661 |
| VP08137 | VP08137 | SAMN20168372 | SRR15099343 | NonPC | Public | 2008 | China | Clinical | 92.1 | 100 | 0.61 | 98.3558 | 5264405 | 45.25 | 61 | 323098 |
| VP08138 | VP08138 | SAMN20168220 | SRR15099511 | NonPC | Public | 2008 | China | Env | 87.8 | 100 | 0.36 | 98.2874 | 5127273 | 45.2 | 62 | 555558 |
| VP08139 | VP08139 | SAMN20168373 | SRR15099342 | NonPC | Public | 2008 | China | Env | 88.2 | 100 | 0.63 | 98.3223 | 5133456 | 45.15 | 87 | 369461 |
| VP08140 | VP08140 | SAMN20168375 | SRR15099340 | NonPC | Public | 2008 | China | Env | 91.4 | 100 | 0 | 98.3583 | 5001188 | 45.4 | 52 | 351314 |
| VP08144 | VP08144 | SAMN20168384 | SRR15099330 | PC | Public | 2008 | China | Clinical | 98.3 | 100 | 0 | 99.9924 | 5095975 | 45.27 | 69 | 288661 |
| VP08145 | VP08145 | SAMN20168385 | SRR15099329 | PC | Public | 2008 | China | Clinical | 98.3 | 100 | 0 | 99.9929 | 5097463 | 45.28 | 70 | 265934 |
| VP08148 | VP08148 | SAMN20168386 | SRR15099328 | NonPC | Public | 2008 | China | Clinical | 92.2 | 100 | 0.19 | 98.3883 | 5003165 | 45.25 | 58 | 450229 |
| VP08150 | VP08150 | SAMN20168387 | SRR15099326 | NonPC | Public | 2008 | China | Clinical | 92.2 | 100 | 0.19 | 98.3966 | 5003344 | 45.25 | 55 | 450229 |
| VP08151 | VP08151 | SAMN20168388 | SRR15099325 | NonPC | Public | 2008 | China | Clinical | 92.3 | 100 | 0.19 | 98.3499 | 5007710 | 45.25 | 50 | 539180 |
| VP08153 | VP08153 | SAMN20168389 | SRR15099324 | NonPC | Public | 2008 | China | Clinical | 92.3 | 100 | 0.19 | 98.3688 | 5008992 | 45.25 | 49 | 539180 |
| VP08154 | VP08154 | SAMN20168390 | SRR15099323 | NonPC | Public | 2008 | China | Clinical | 92.3 | 100 | 0.19 | 98.3719 | 5008655 | 45.25 | 49 | 539180 |
| VP08155 | VP08155 | SAMN20168391 | SRR15099322 | NonPC | Public | 2008 | China | Clinical | 92.3 | 100 | 0.19 | 98.3384 | 5008753 | 45.25 | 51 | 471370 |
| VP08156 | VP08156 | SAMN20168392 | SRR15099321 | NonPC | Public | 2008 | China | Clinical | 92.2 | 100 | 0.19 | 98.3964 | 5003786 | 45.26 | 56 | 450229 |
| VP08157 | VP08157 | SAMN20168393 | SRR15099320 | NonPC | Public | 2008 | China | Clinical | 92.3 | 100 | 0.19 | 98.3623 | 5009018 | 45.25 | 50 | 471204 |
| VP08158 | VP08158 | SAMN20168394 | SRR15099319 | NonPC | Public | 2008 | China | Clinical | 92.3 | 100 | 0.19 | 98.3418 | 5007623 | 45.25 | 49 | 539180 |
| VP08159 | VP08159 | SAMN20168395 | SRR15099318 | NonPC | Public | 2008 | China | Clinical | 92.2 | 100 | 0.19 | 98.3586 | 5007887 | 45.25 | 50 | 539180 |
| VP08160 | VP08160 | SAMN20168396 | SRR15099317 | NonPC | Public | 2008 | China | Clinical | 92.2 | 100 | 0.19 | 98.3603 | 5003908 | 45.26 | 56 | 450229 |
| VP08162 | VP08162 | SAMN20168405 | SRR15099306 | NonPC | Public | 2008 | China | Env | 91.1 | 100 | 0 | 98.3455 | 5082874 | 45.25 | 45 | 352481 |

| | | | | | | | | | | | | | | | | |
|---|---|---|---|---|---|---|---|---|---|---|---|---|---|---|---|---|
| VP08163 | VP08163 | SAMN20168407 | SRR15099303 | PC | Public | 2008 | China | Clinical | 98 | 100 | 0 | 99.9841 | 5080295 | 45.3 | 63 | 411189 |
| VP08164 | VP08164 | SAMN20168410 | SRR15099300 | PC | Public | 2008 | China | Clinical | 98.6 | 100 | 0.27 | 99.9831 | 5192728 | 45.28 | 60 | 427734 |
| VP08165 | VP08165 | SAMN20168408 | SRR15099302 | PC | Public | 2008 | China | Clinical | 98 | 100 | 0 | 99.9842 | 5080583 | 45.3 | 63 | 411303 |
| VP08166 | VP08166 | SAMN20168409 | SRR15099301 | PC | Public | 2008 | China | Clinical | 98 | 100 | 0 | 99.9818 | 5080509 | 45.3 | 64 | 401611 |
| VP08167 | VP08167 | SAMN20168411 | SRR15099299 | PC | Public | 2008 | China | Clinical | 98.6 | 100 | 0.27 | 99.9915 | 5191766 | 45.28 | 59 | 420265 |
| VP08168 | VP08168 | SAMN20168412 | SRR15099298 | PC | Public | 2008 | China | Clinical | 98.6 | 100 | 0 | 99.9803 | 5150141 | 45.27 | 60 | 427734 |
| VP08169 | VP08169 | SAMN20168406 | SRR15099305 | PC | Public | 2008 | China | Clinical | 98.5 | 100 | 0.27 | 99.9935 | 5133991 | 45.3 | 61 | 401604 |
| VP08170 | VP08170 | SAMN20168397 | SRR15099314 | PC | Public | 2008 | China | Clinical | 98 | 100 | 0 | 99.9786 | 5080464 | 45.3 | 62 | 411303 |
| VP08171 | VP08171 | SAMN20168401 | SRR15099310 | PC | Public | 2008 | China | Clinical | 98 | 100 | 0 | 99.9791 | 5079276 | 45.3 | 61 | 411135 |
| VP08172 | VP08172 | SAMN20168403 | SRR15099308 | PC | Public | 2008 | China | Clinical | 98.5 | 100 | 0 | 99.9843 | 5107429 | 45.28 | 59 | 425770 |
| VP08173 | VP08173 | SAMN20168402 | SRR15099309 | PC | Public | 2008 | China | Clinical | 98.6 | 100 | 0 | 99.9848 | 5131601 | 45.29 | 64 | 373607 |
| VP08174 | VP08174 | SAMN20168398 | SRR15099313 | NonPC | Public | 2008 | China | Clinical | 91.9 | 100 | 0 | 98.4056 | 5093875 | 45.24 | 33 | 524940 |
| VP08175 | VP08175 | SAMN20168382 | SRR15099332 | NonPC | Public | 2008 | China | Clinical | 92.9 | 100 | 0.06 | 98.3798 | 5125386 | 45.33 | 46 | 411537 |
| VP08176 | VP08176 | SAMN20168383 | SRR15099331 | NonPC | Public | 2008 | China | Clinical | 90.9 | 100 | 0 | 98.2797 | 5121181 | 45.29 | 66 | 374815 |
| VP08177 | VP08177 | SAMN20168404 | SRR15099307 | NonPC | Public | 2008 | China | Clinical | 92.5 | 100 | 0.05 | 98.349 | 5068394 | 45.27 | 45 | 450396 |
| VP08180 | VP08180 | SAMN20168413 | SRR15099297 | NonPC | Public | 2008 | China | Clinical | 92.5 | 100 | 0.05 | 98.3434 | 5067013 | 45.27 | 42 | 539107 |
| VP08181 | VP08181 | SAMN20168414 | SRR15099296 | NonPC | Public | 2008 | China | Clinical | 92.7 | 100 | 0 | 98.3797 | 5161485 | 45.13 | 56 | 434710 |
| VP08182 | VP08182 | SAMN20168417 | SRR15099292 | NonPC | Public | 2008 | China | Clinical | 89.8 | 99.86 | 0 | 98.2949 | 5028524 | 45.33 | 37 | 379511 |
| VP08183 | VP08183 | SAMN20168415 | SRR15099295 | NonPC | Public | 2008 | China | Clinical | 89.2 | 100 | 0.05 | 98.2988 | 5052712 | 45.24 | 47 | 658696 |
| VP08184 | VP08184 | SAMN20168420 | SRR15099289 | PC | Public | 2008 | China | Clinical | 98.5 | 100 | 0 | 99.9927 | 5092968 | 45.29 | 58 | 425763 |
| VP08185 | VP08185 | SAMN20168418 | SRR15099291 | NonPC | Public | 2008 | China | Clinical | 91.1 | 100 | 0.27 | 98.3234 | 5220959 | 45.29 | 44 | 383492 |
| VP08186 | VP08186 | SAMN20168416 | SRR15099294 | NonPC | Public | 2008 | China | Clinical | 89.9 | 100 | 0.2 | 98.2774 | 5029364 | 45.37 | 59 | 450410 |
| VP08187 | VP08187 | SAMN20168424 | SRR15099285 | PC | Public | 2008 | China | Clinical | 98.5 | 100 | 0 | 99.9926 | 5102209 | 45.28 | 59 | 425770 |
| VP08188 | VP08188 | SAMN20168426 | SRR15099283 | PC | Public | 2008 | China | Clinical | 98.6 | 100 | 0 | 99.9831 | 5239310 | 45.19 | 64 | 420398 |
| VP08189 | VP08189 | SAMN20168423 | SRR15099286 | NonPC | Public | 2008 | China | Clinical | 91.9 | 100 | 0 | 98.4225 | 5056224 | 45.25 | 35 | 522556 |
| VP08190 | VP08190 | SAMN20168425 | SRR15099284 | PC | Public | 2008 | China | Clinical | 98.1 | 100 | 0 | 99.9855 | 5074263 | 45.29 | 46 | 425763 |
| VP08191 | VP08191 | SAMN20168428 | SRR15099280 | NonPC | Public | 2008 | China | Clinical | 90.9 | 100 | 0 | 98.2865 | 5252527 | 45.26 | 72 | 374815 |
| VP08192 | VP08192 | SAMN20168435 | SRR15099273 | NonPC | Public | 2008 | China | Clinical | 90.8 | 100 | 0 | 98.337 | 5176912 | 45.27 | 58 | 395865 |
| VP08193 | VP08193 | SAMN20168436 | SRR15099272 | PC | Public | 2008 | China | Clinical | 98.4 | 100 | 0 | 99.9923 | 5086295 | 45.3 | 48 | 425769 |
| VP08194 | VP08194 | SAMN20168431 | SRR15099277 | NonPC | Public | 2008 | China | Clinical | 93 | 100 | 0 | 98.3855 | 5175359 | 45.21 | 43 | 488339 |
| VP08195 | VP08195 | SAMN20168433 | SRR15099275 | NonPC | Public | 2008 | China | Clinical | 90.9 | 100 | 0 | 98.2756 | 5121574 | 45.29 | 66 | 375351 |
| VP08196 | VP08196 | SAMN20168432 | SRR15099276 | NonPC | Public | 2008 | China | Clinical | 90.9 | 100 | 0 | 98.3167 | 5122101 | 45.29 | 67 | 375351 |
| VP08197 | VP08197 | SAMN20168472 | SRR15099232 | PC | Public | 2008 | China | Clinical | 98.6 | 100 | 0 | 99.9915 | 5150280 | 45.27 | 64 | 288872 |
| VP08198 | VP08198 | SAMN20168473 | SRR15099231 | PC | Public | 2008 | China | Clinical | 98.6 | 100 | 0 | 99.9909 | 5151125 | 45.27 | 59 | 420264 |
| VP08199 | VP08199 | SAMN20168471 | SRR15099233 | PC | Public | 2008 | China | Clinical | 98.4 | 100 | 0 | 99.9843 | 5089296 | 45.29 | 57 | 394933 |
| VP08200 | VP08200 | SAMN20168419 | SRR15099290 | NonPC | Public | 2008 | China | Clinical | 89.3 | 100 | 0.36 | 98.2739 | 5089100 | 45.05 | 64 | 371862 |
| VP08201 | VP08201 | SAMN20168476 | SRR15099228 | PC | Public | 2008 | China | Clinical | 98 | 100 | 0 | 99.9889 | 5080318 | 45.3 | 65 | 411189 |
| VP08202 | VP08202 | SAMN20168478 | SRR15099846 | PC | Public | 2008 | China | Clinical | 98.5 | 100 | 0 | 99.9786 | 5106904 | 45.28 | 56 | 425770 |
| VP08203 | VP08203 | SAMN20168477 | SRR15099847 | PC | Public | 2008 | China | Clinical | 98.6 | 100 | 0 | 99.9859 | 5131081 | 45.29 | 61 | 401597 |
| VP08204 | VP08204 | SAMN20168250 | SRR15099478 | NonPC | Public | 2008 | China | Env | 88.2 | 100 | 0.36 | 98.2584 | 5125414 | 45.16 | 139 | 318699 |
| VP08205 | VP08205 | SAMN20168400 | SRR15099311 | PC | Public | 2008 | China | Clinical | 98.6 | 100 | 0 | 99.9914 | 5130217 | 45.29 | 58 | 427842 |
| VP08206 | VP08206 | SAMN20168399 | SRR15099312 | PC | Public | 2008 | China | Clinical | 98.4 | 100 | 0 | 99.9924 | 5086478 | 45.29 | 51 | 425997 |
| VP08207 | VP08207 | SAMN20168359 | SRR15099357 | PC | Public | 2008 | China | Clinical | 98.6 | 100 | 0 | 99.9939 | 5111277 | 45.28 | 60 | 427746 |
| VP08208 | VP08208 | SAMN20168355 | SRR15099362 | PC | Public | 2008 | China | Clinical | 98 | 100 | 0 | 99.9832 | 5080629 | 45.3 | 62 | 411189 |
| VP08209 | VP08209 | SAMN20168356 | SRR15099361 | PC | Public | 2008 | China | Clinical | 98.6 | 100 | 0 | 99.984 | 5131445 | 45.29 | 59 | 401597 |
| VP08210 | VP08210 | SAMN20168358 | SRR15099358 | PC | Public | 2008 | China | Clinical | 98.4 | 100 | 0 | 99.9861 | 5104844 | 45.28 | 63 | 373488 |
| VP08212 | VP08212 | SAMN20168536 | SRR15099782 | PC | Public | 2008 | China | Clinical | 97.6 | 100 | 0 | 99.9377 | 5152132 | 45.12 | 80 | 312475 |
| VP08213 | VP08213 | SAMN20168467 | SRR15099237 | NonPC | Public | 2008 | China | Clinical | 92.1 | 100 | 0.05 | 98.4557 | 5065385 | 45.23 | 50 | 294932 |

| | | | | | | | | | | | | | | | | |
|---|---|---|---|---|---|---|---|---|---|---|---|---|---|---|---|---|
| VP08214 | VP08214 | SAMN20168468 | SRR15099236 | NonPC | Public | 2008 | China | Clinical | 92.1 | 100 | 0.06 | 98.4463 | 5158104 | 45.26 | 48 | 294932 |
| VP08215 | VP08215 | SAMN20168469 | SRR15099235 | NonPC | Public | 2008 | China | Clinical | 92.1 | 100 | 0.05 | 98.4402 | 5065160 | 45.23 | 46 | 312889 |
| VP08216 | VP08216 | SAMN20168470 | SRR15099234 | NonPC | Public | 2008 | China | Clinical | 92.1 | 100 | 0.05 | 98.4252 | 5064037 | 45.23 | 50 | 294932 |
| VP08217 | VP08217 | SAMN20168294 | SRR15099430 | PC | Public | 2008 | China | Clinical | 98.6 | 100 | 0 | 99.9901 | 5150354 | 45.27 | 60 | 420264 |
| VP08218 | VP08218 | SAMN20168434 | SRR15099274 | NonPC | Public | 2008 | China | Clinical | 90.9 | 100 | 0 | 98.2721 | 5121458 | 45.29 | 68 | 374815 |
| VP08220 | VP08220 | SAMN20168430 | SRR15099278 | NonPC | Public | 2008 | China | Clinical | 92.9 | 100 | 0.05 | 98.3871 | 5027037 | 45.31 | 45 | 411537 |
| VP08221 | VP08221 | SAMN20168441 | SRR15099266 | NonPC | Public | 2008 | China | Clinical | 90.6 | 100 | 0.05 | 98.3542 | 5068874 | 45.26 | 69 | 231056 |
| VP08222 | VP08222 | SAMN20168448 | SRR15099258 | NonPC | Public | 2008 | China | Clinical | 90.9 | 100 | 0 | 98.3116 | 5112576 | 45.3 | 64 | 374815 |
| VP08223 | VP08223 | SAMN20168446 | SRR15099261 | NonPC | Public | 2008 | China | Clinical | 90.9 | 100 | 0 | 98.2998 | 5210895 | 45.21 | 71 | 375351 |
| VP08224 | VP08224 | SAMN20168447 | SRR15099259 | NonPC | Public | 2008 | China | Clinical | 90.9 | 100 | 0 | 98.3215 | 5122047 | 45.29 | 67 | 375210 |
| VP08225 | VP08225 | SAMN20168452 | SRR15099254 | PC | Public | 2008 | China | Clinical | 97.7 | 100 | 0 | 99.9214 | 5118393 | 45.29 | 61 | 401611 |
| VP08227 | VP08227 | SAMN20168429 | SRR15099279 | NonPC | Public | 2008 | China | Clinical | 92.4 | 99.93 | 0.05 | 98.3683 | 5054440 | 45.29 | 42 | 450397 |
| VP08228 | VP08228 | SAMN20168455 | SRR15099251 | PC | Public | 2008 | China | Clinical | 98.6 | 100 | 0 | 99.9924 | 5151859 | 45.27 | 64 | 392415 |
| VP08229 | VP08229 | SAMN20168454 | SRR15099252 | PC | Public | 2008 | China | Clinical | 98.5 | 100 | 0 | 99.9916 | 5092682 | 45.29 | 56 | 440647 |
| VP08230 | VP08230 | SAMN20168535 | SRR15099783 | PC | Public | 2008 | China | Clinical | 98.6 | 100 | 0 | 99.9885 | 5150998 | 45.27 | 62 | 420389 |
| VP08231 | VP08231 | SAMN20168458 | SRR15099247 | PC | Public | 2008 | China | Clinical | 98.6 | 100 | 0 | 99.9818 | 5150636 | 45.27 | 58 | 427734 |
| VP08232 | VP08232 | SAMN20168438 | SRR15099269 | PC | Public | 2008 | China | Clinical | 98.5 | 100 | 0 | 99.9925 | 5092656 | 45.29 | 56 | 425733 |
| VP08233 | VP08233 | SAMN20168457 | SRR15099248 | PC | Public | 2008 | China | Clinical | 98.6 | 100 | 0.09 | 99.9912 | 5162879 | 45.26 | 58 | 420398 |
| VP08234 | VP08234 | SAMN20168443 | SRR15099264 | PC | Public | 2008 | China | Clinical | 98.5 | 100 | 0 | 99.9884 | 5270318 | 45.03 | 67 | 394986 |
| VP08235 | VP08235 | SAMN20168456 | SRR15099250 | PC | Public | 2008 | China | Clinical | 98.6 | 100 | 0 | 99.9845 | 5149781 | 45.27 | 59 | 427734 |
| VP08236 | VP08236 | SAMN20168451 | SRR15099255 | PC | Public | 2008 | China | Clinical | 98.5 | 100 | 0 | 99.9842 | 5094195 | 45.29 | 58 | 401604 |
| VP08237 | VP08237 | SAMN20168444 | SRR15099263 | NonPC | Public | 2008 | China | Clinical | 90.9 | 100 | 0 | 98.2909 | 5120956 | 45.29 | 64 | 405034 |
| VP08240 | VP08240 | SAMN20168437 | SRR15099270 | PC | Public | 2008 | China | Clinical | 98.6 | 100 | 0 | 99.9885 | 5150468 | 45.27 | 60 | 394851 |
| VP08241 | VP08241 | SAMN20168291 | SRR15099433 | PC | Public | 2008 | China | Clinical | 98.6 | 100 | 0.27 | 99.9806 | 5191839 | 45.28 | 60 | 420398 |
| VP08242 | VP08242 | SAMN20168442 | SRR15099265 | PC | Public | 2008 | China | Clinical | 97.8 | 100 | 0 | 99.894 | 5148211 | 45.29 | 54 | 394770 |
| VP08243 | VP08243 | SAMN20168439 | SRR15099268 | PC | Public | 2008 | China | Clinical | 98 | 100 | 0 | 99.9294 | 5201249 | 45.19 | 60 | 427315 |
| VP08244 | VP08244 | SAMN20168445 | SRR15099262 | NonPC | Public | 2008 | China | Clinical | 90.9 | 100 | 0 | 98.3144 | 5124462 | 45.3 | 71 | 374815 |
| VP08247 | VP08247 | SAMN20168376 | SRR15099339 | PC | Public | 2008 | China | Clinical | 98.6 | 100 | 0.7 | 99.9841 | 5254489 | 45.26 | 62 | 288872 |
| VP08248 | VP08248 | SAMN20168374 | SRR15099341 | PC | Public | 2008 | China | Clinical | 97.7 | 100 | 0 | 99.942 | 5166852 | 45.26 | 59 | 401611 |
| VP08250 | VP08250 | SAMN20168377 | SRR15099337 | PC | Public | 2008 | China | Clinical | 98.6 | 100 | 0 | 99.9814 | 5150062 | 45.27 | 59 | 420264 |
| VP08251 | VP08251 | SAMN20168378 | SRR15099336 | PC | Public | 2008 | China | Clinical | 98.6 | 100 | 0.7 | 99.9848 | 5254222 | 45.26 | 62 | 392281 |
| VP08252 | VP08252 | SAMN20168379 | SRR15099335 | PC | Public | 2008 | China | Clinical | 98.6 | 100 | 0.7 | 99.9829 | 5253842 | 45.26 | 60 | 394853 |
| VP08253 | VP08253 | SAMN20168380 | SRR15099334 | PC | Public | 2008 | China | Clinical | 98.6 | 100 | 0.7 | 99.9804 | 5254233 | 45.26 | 61 | 427734 |
| VP08254 | VP08254 | SAMN20168381 | SRR15099333 | PC | Public | 2008 | China | Clinical | 98.6 | 100 | 0 | 99.9885 | 5150495 | 45.27 | 61 | 420398 |
| VP08255 | VP08255 | SAMN20168491 | SRR15099832 | PC | Public | 2008 | China | Clinical | 98.6 | 100 | 0 | 99.9939 | 5110590 | 45.28 | 58 | 401611 |
| VP08257 | VP08257 | SAMN20168492 | SRR15099831 | PC | Public | 2008 | China | Clinical | 98.6 | 100 | 0 | 99.992 | 5111004 | 45.28 | 57 | 427206 |
| VP08258 | VP08258 | SAMN20168493 | SRR15099830 | PC | Public | 2008 | China | Clinical | 98.6 | 100 | 0 | 99.987 | 5110165 | 45.28 | 59 | 401477 |
| VP08259 | VP08259 | SAMN20168494 | SRR15099829 | PC | Public | 2008 | China | Clinical | 98.6 | 100 | 0 | 99.9854 | 5110513 | 45.28 | 60 | 401611 |
| VP08260 | VP08260 | SAMN20168495 | SRR15099828 | PC | Public | 2008 | China | Clinical | 98.6 | 100 | 0 | 99.9942 | 5110892 | 45.28 | 59 | 427734 |
| VP08262 | VP08262 | SAMN20168496 | SRR15099827 | PC | Public | 2008 | China | Clinical | 98.2 | 100 | 0 | 99.9928 | 5088383 | 45.27 | 80 | 268004 |
| VP08263 | VP08263 | SAMN20168497 | SRR15099824 | PC | Public | 2008 | China | Clinical | 98.6 | 100 | 0 | 99.9945 | 5110408 | 45.28 | 58 | 427734 |
| VP08265 | VP08265 | SAMN20168507 | SRR15099813 | PC | Public | 2008 | China | Clinical | 98.6 | 100 | 0 | 99.9803 | 5150845 | 45.27 | 62 | 420398 |
| VP08266 | VP08266 | SAMN20168498 | SRR15099823 | PC | Public | 2008 | China | Clinical | 98.6 | 100 | 0 | 99.9861 | 5110944 | 45.28 | 58 | 427734 |
| VP08267 | VP08267 | SAMN20168499 | SRR15099822 | PC | Public | 2008 | China | Clinical | 98.5 | 100 | 0 | 99.9765 | 5160758 | 45.27 | 58 | 401611 |
| VP08268 | VP08268 | SAMN20168500 | SRR15099821 | PC | Public | 2008 | China | Clinical | 98.6 | 100 | 0 | 99.9856 | 5110385 | 45.28 | 59 | 427734 |
| VP08270 | VP08270 | SAMN20168501 | SRR15099820 | PC | Public | 2008 | China | Clinical | 98.6 | 100 | 0 | 99.9859 | 5110291 | 45.28 | 58 | 401477 |
| VP08271 | VP08271 | SAMN20168502 | SRR15099819 | PC | Public | 2008 | China | Clinical | 98.6 | 100 | 0 | 99.9838 | 5111042 | 45.29 | 59 | 401611 |
| VP08273 | VP08273 | SAMN20168503 | SRR15099818 | PC | Public | 2008 | China | Clinical | 98.6 | 100 | 0 | 99.9847 | 5111606 | 45.29 | 60 | 427734 |

| | | | | | | | | | | | | | | | | |
|---|---|---|---|---|---|---|---|---|---|---|---|---|---|---|---|---|
| VP08274 | VP08274 | SAMN20168504 | SRR15099817 | PC | Public | 2008 | China | Clinical | 98.6 | 100 | 0 | 99.9938 | 5110534 | 45.29 | 60 | 427734 |
| VP08275 | VP08275 | SAMN20168505 | SRR15099816 | PC | Public | 2008 | China | Clinical | 98.6 | 100 | 0 | 99.983 | 5110003 | 45.28 | 57 | 427848 |
| VP08276 | VP08276 | SAMN20168506 | SRR15099815 | PC | Public | 2008 | China | Clinical | 98.6 | 100 | 0 | 99.987 | 5109903 | 45.28 | 61 | 401611 |
| VP08278 | VP08278 | SAMN20168251 | SRR15099477 | NonPC | Public | 2008 | China | Env | 90.9 | 100 | 0.05 | 98.3907 | 5135529 | 45.21 | 69 | 257737 |
| VP08279 | VP08279 | SAMN20168270 | SRR15099456 | NonPC | Public | 2008 | China | Env | 89.4 | 100 | 0.03 | 98.3499 | 5090671 | 45.32 | 61 | 188844 |
| VP08280 | VP08280 | SAMN20168209 | SRR15099523 | NonPC | Public | 2008 | China | Env | 92.3 | 100 | 0.05 | 98.3533 | 5009540 | 45.26 | 54 | 313971 |
| VP08281 | VP08281 | SAMN20168252 | SRR15099476 | NonPC | Public | 2008 | China | Env | 91.5 | 100 | 0.27 | 98.3919 | 5058527 | 45.34 | 52 | 414275 |
| VP08282 | VP08282 | SAMN20168283 | SRR15099442 | NonPC | Public | 2008 | China | Env | 90.9 | 100 | 0.27 | 98.4085 | 5015448 | 45.32 | 50 | 447619 |
| VP08283 | VP08283 | SAMN20168253 | SRR15099475 | NonPC | Public | 2008 | China | Env | 91.5 | 100 | 0.03 | 98.3938 | 5039827 | 45.37 | 40 | 349637 |
| VP08284 | VP08284 | SAMN20168211 | SRR15099521 | NonPC | Public | 2008 | China | Env | 89.7 | 100 | 0 | 98.3156 | 4954962 | 45.4 | 40 | 529786 |
| VP08285 | VP08285 | SAMN20168254 | SRR15099474 | NonPC | Public | 2008 | China | Env | 88.3 | 100 | 0.36 | 98.2386 | 5012387 | 45.27 | 59 | 392640 |
| VP08286 | VP08286 | SAMN20168222 | SRR15099509 | NonPC | Public | 2008 | China | Env | 91.5 | 100 | 0.09 | 98.384 | 5120826 | 45.3 | 31 | 496486 |
| VP08288 | VP08288 | SAMN20168217 | SRR15099514 | NonPC | Public | 2008 | China | Env | 91.1 | 100 | 0 | 98.3731 | 5025097 | 45.3 | 65 | 460327 |
| VP08289 | VP08289 | SAMN20168255 | SRR15099473 | NonPC | Public | 2008 | China | Env | 88.4 | 100 | 0.3 | 98.2837 | 4988839 | 45.3 | 54 | 451496 |
| VP08290 | VP08290 | SAMN20168215 | SRR15099517 | NonPC | Public | 2008 | China | Env | 91 | 100 | 0.03 | 98.4298 | 5015082 | 45.31 | 40 | 314079 |
| VP08291 | VP08291 | SAMN20168221 | SRR15099510 | NonPC | Public | 2008 | China | Env | 87.9 | 100 | 0.36 | 98.2353 | 5104719 | 45.15 | 67 | 560889 |
| VP08292 | VP08292 | SAMN20168256 | SRR15099472 | NonPC | Public | 2008 | China | Env | 91.9 | 100 | 0.31 | 98.3463 | 5117646 | 45.3 | 60 | 348410 |
| VP08293 | VP08293 | SAMN20168257 | SRR15099470 | NonPC | Public | 2008 | China | Env | 89.9 | 100 | 0 | 98.3339 | 5077120 | 45.31 | 65 | 467781 |
| VP08294 | VP08294 | SAMN20168258 | SRR15099469 | NonPC | Public | 2008 | China | Env | 90.7 | 100 | 0 | 98.2123 | 5277689 | 45.38 | 73 | 441619 |
| VP08295 | VP08295 | SAMN20168259 | SRR15099468 | NonPC | Public | 2008 | China | Env | 91.2 | 100 | 0.03 | 98.3247 | 5070836 | 45.32 | 75 | 413423 |
| VP08296 | VP08296 | SAMN20168483 | SRR15099841 | NonPC | Public | 2008 | China | Clinical | 92.3 | 100 | 0.19 | 98.3356 | 5010468 | 45.25 | 49 | 450300 |
| VP08297 | VP08297 | SAMN20168284 | SRR15099441 | NonPC | Public | 2008 | China | Env | 90.8 | 100 | 0.27 | 98.3823 | 5002155 | 45.32 | 58 | 312544 |
| VP08298 | VP08298 | SAMN20168260 | SRR15099467 | NonPC | Public | 2008 | China | Env | 91.5 | 99.97 | 0.58 | 98.3363 | 5067453 | 45.29 | 48 | 423066 |
| VP08299 | VP08299 | SAMN20168261 | SRR15099466 | NonPC | Public | 2008 | China | Env | 91.5 | 100 | 0 | 98.3519 | 5092129 | 45.3 | 50 | 452987 |
| VP08300 | VP08300 | SAMN20168281 | SRR15099444 | NonPC | Public | 2008 | China | Env | 91.7 | 100 | 0 | 98.4 | 5059257 | 45.25 | 46 | 417572 |
| VP08301 | VP08301 | SAMN20168224 | SRR15099507 | NonPC | Public | 2008 | China | Env | 91.6 | 100 | 0.03 | 98.2718 | 5046542 | 45.31 | 35 | 451946 |
| VP08302 | VP08302 | SAMN20168212 | SRR15099520 | NonPC | Public | 2008 | China | Env | 89.8 | 100 | 0 | 98.2827 | 4935670 | 45.39 | 39 | 421307 |
| VP08304 | VP08304 | SAMN20168214 | SRR15099518 | NonPC | Public | 2008 | China | Env | 89.5 | 100 | 0.33 | 98.3051 | 5137222 | 45.33 | 35 | 724393 |
| VP08306 | VP08306 | SAMN20168262 | SRR15099465 | NonPC | Public | 2008 | China | Env | 89.9 | 100 | 0 | 98.3378 | 4999703 | 45.29 | 35 | 448929 |
| VP08309 | VP08309 | SAMN20168225 | SRR15099506 | NonPC | Public | 2008 | China | Env | 88.6 | 99.93 | 0.34 | 98.2129 | 5086169 | 45.23 | 46 | 460671 |
| VP08310 | VP08310 | SAMN20168263 | SRR15099464 | NonPC | Public | 2008 | China | Env | 91.2 | 100 | 0 | 98.3513 | 5086494 | 45.28 | 73 | 244603 |
| VP08311 | VP08311 | SAMN20168279 | SRR15099446 | NonPC | Public | 2008 | China | Clinical | 91.7 | 100 | 0 | 98.3966 | 5042645 | 45.25 | 47 | 417274 |
| VP08312 | VP08312 | SAMN20168264 | SRR15099463 | NonPC | Public | 2008 | China | Env | 88.3 | 100 | 0.36 | 98.2365 | 5059902 | 45.31 | 67 | 400260 |
| VP08313 | VP08313 | SAMN20168449 | SRR15099257 | NonPC | Public | 2008 | China | Clinical | 90.9 | 100 | 0 | 98.2816 | 5121590 | 45.29 | 66 | 375351 |
| VP08314 | VP08314 | SAMN20168465 | SRR15099240 | PC | Public | 2008 | China | Clinical | 98.6 | 100 | 0 | 99.982 | 5150896 | 45.27 | 60 | 394861 |
| VP08315 | VP08315 | SAMN20168462 | SRR15099243 | PC | Public | 2008 | China | Clinical | 97.7 | 100 | 0 | 99.9281 | 5151978 | 45.12 | 80 | 265868 |
| VP08316 | VP08316 | SAMN20168463 | SRR15099242 | PC | Public | 2008 | China | Clinical | 98.5 | 100 | 0 | 99.9834 | 5092743 | 45.29 | 56 | 425871 |
| VP08317 | VP08317 | SAMN20168459 | SRR15099246 | NonPC | Public | 2008 | China | Clinical | 92.1 | 100 | 0.05 | 98.4353 | 5064407 | 45.23 | 49 | 294932 |
| VP08318 | VP08318 | SAMN20168464 | SRR15099241 | PC | Public | 2008 | China | Clinical | 98.5 | 100 | 0 | 99.9873 | 5109774 | 45.28 | 59 | 425770 |
| VP08319 | VP08319 | SAMN20168450 | SRR15099256 | NonPC | Public | 2008 | China | Clinical | 92.3 | 100 | 0.19 | 98.3432 | 5013374 | 45.25 | 55 | 450300 |
| VP08320 | VP08320 | SAMN20168440 | SRR15099267 | NonPC | Public | 2008 | China | Clinical | 90.9 | 100 | 0 | 98.3113 | 5130736 | 45.29 | 65 | 374836 |
| VP08321 | VP08321 | SAMN20168475 | SRR15099229 | NonPC | Public | 2008 | China | Clinical | 90.9 | 100 | 0 | 98.2711 | 5120797 | 45.29 | 67 | 374815 |
| VP08322 | VP08322 | SAMN20168427 | SRR15099281 | PC | Public | 2008 | China | Clinical | 98.5 | 100 | 0 | 99.9929 | 5112563 | 45.26 | 59 | 440325 |
| VP08323 | VP08323 | SAMN20168453 | SRR15099253 | PC | Public | 2008 | China | Clinical | 98.4 | 100 | 0 | 99.9933 | 5091647 | 45.29 | 60 | 401604 |
| VP08324 | VP08324 | SAMN20168460 | SRR15099245 | NonPC | Public | 2008 | China | Clinical | 93 | 100 | 0.05 | 98.353 | 5034243 | 45.33 | 29 | 515934 |
| VP08325 | VP08325 | SAMN20168461 | SRR15099244 | NonPC | Public | 2008 | China | Clinical | 90.8 | 100 | 0 | 98.3503 | 5176084 | 45.27 | 57 | 395865 |
| VP08326 | VP08326 | SAMN20168466 | SRR15099239 | PC | Public | 2008 | China | Clinical | 98.6 | 100 | 0 | 99.9917 | 5150573 | 45.27 | 64 | 392408 |
| VP08327 | VP08327 | SAMN20168474 | SRR15099230 | PC | Public | 2008 | China | Clinical | 98 | 100 | 0 | 99.9241 | 5185144 | 45.12 | 59 | 401484 |

| | | | | | | | | | | | | | | | | |
|---|---|---|---|---|---|---|---|---|---|---|---|---|---|---|---|---|
| VP08329 | VP08329 | SAMN20168480 | SRR15099844 | PC | Public | 2008 | China | Clinical | 98.4 | 100 | 0.02 | 99.9828 | 5089903 | 45.3 | 51 | 401471 |
| VP08330 | VP08330 | SAMN20168479 | SRR15099845 | NonPC | Public | 2008 | China | Clinical | 92.3 | 100 | 0.19 | 98.3205 | 5012463 | 45.25 | 55 | 450300 |
| VP08331 | VP08331 | SAMN20168481 | SRR15099843 | PC | Public | 2008 | China | Clinical | 98.6 | 100 | 0 | 99.9922 | 5150143 | 45.27 | 61 | 395241 |
| VP08332 | VP08332 | SAMN20168484 | SRR15099840 | PC | Public | 2008 | China | Clinical | 97.3 | 100 | 0 | 99.9252 | 5095568 | 45.27 | 83 | 267662 |
| VP08333 | VP08333 | SAMN20168508 | SRR15099812 | NonPC | Public | 2008 | China | Clinical | 92.3 | 100 | 0.19 | 98.3582 | 5008363 | 45.25 | 49 | 539180 |
| VP08334 | VP08334 | SAMN20168513 | SRR15099807 | PC | Public | 2008 | China | Clinical | 98.6 | 100 | 0 | 99.9917 | 5222147 | 45.12 | 62 | 394853 |
| VP08335 | VP08335 | SAMN20168515 | SRR15099805 | PC | Public | 2008 | China | Clinical | 98.6 | 100 | 0 | 99.9816 | 5150992 | 45.27 | 59 | 420398 |
| VP08336 | VP08336 | SAMN20168514 | SRR15099806 | PC | Public | 2008 | China | Clinical | 98.6 | 100 | 0 | 99.9809 | 5150883 | 45.27 | 62 | 420398 |
| VP08337 | VP08337 | SAMN20168511 | SRR15099809 | PC | Public | 2008 | China | Clinical | 98.6 | 100 | 0 | 99.9905 | 5151097 | 45.27 | 57 | 440583 |
| VP08338 | VP08338 | SAMN20168510 | SRR15099810 | PC | Public | 2008 | China | Clinical | 98 | 100 | 0 | 99.9793 | 5079460 | 45.3 | 62 | 401611 |
| VP08339 | VP08339 | SAMN20168509 | SRR15099811 | PC | Public | 2008 | China | Clinical | 98.5 | 100 | 0.27 | 99.9936 | 5134402 | 45.3 | 61 | 401604 |
| VP08340 | VP08340 | SAMN20168512 | SRR15099808 | PC | Public | 2008 | China | Clinical | 98.6 | 100 | 0 | 99.9908 | 5151189 | 45.27 | 63 | 420398 |
| VP08341 | VP08341 | SAMN20168516 | SRR15099804 | PC | Public | 2008 | China | Clinical | 98 | 100 | 0 | 99.9786 | 5080026 | 45.3 | 68 | 373628 |
| VP08342 | VP08342 | SAMN20168546 | SRR15099771 | PC | Public | 2008 | China | Clinical | 98.6 | 100 | 0 | 99.9902 | 5150742 | 45.27 | 60 | 420265 |
| VP08343 | VP08343 | SAMN20168517 | SRR15099802 | NonPC | Public | 2008 | China | Clinical | 92.4 | 100 | 0.05 | 98.3408 | 5069299 | 45.27 | 47 | 450396 |
| VP08344 | VP08344 | SAMN20168522 | SRR15099797 | PC | Public | 2008 | China | Clinical | 98.6 | 100 | 0 | 99.983 | 5149913 | 45.27 | 63 | 394861 |
| VP08345 | VP08345 | SAMN20168532 | SRR15099786 | PC | Public | 2008 | China | Clinical | 98.6 | 100 | 0 | 99.9842 | 5131470 | 45.29 | 67 | 288859 |
| VP08346 | VP08346 | SAMN20168485 | SRR15099839 | PC | Public | 2008 | China | Clinical | 98.6 | 99.73 | 0 | 99.9836 | 5150056 | 45.27 | 62 | 420264 |
| VP08347 | VP08347 | SAMN20168487 | SRR15099836 | PC | Public | 2008 | China | Clinical | 98.6 | 100 | 0 | 99.9945 | 5110109 | 45.28 | 63 | 318788 |
| VP08348 | VP08348 | SAMN20168490 | SRR15099833 | PC | Public | 2008 | China | Clinical | 98.6 | 100 | 0 | 99.9805 | 5165325 | 45.27 | 66 | 312832 |
| VP08349 | VP08349 | SAMN20168288 | SRR15099436 | PC | Public | 2008 | China | Clinical | 98.6 | 100 | 0 | 99.986 | 5110863 | 45.28 | 65 | 373628 |
| VP08350 | VP08350 | SAMN20168488 | SRR15099835 | PC | Public | 2008 | China | Clinical | 98.6 | 100 | 0 | 99.9941 | 5112536 | 45.29 | 59 | 427848 |
| VP08351 | VP08351 | SAMN20168489 | SRR15099834 | PC | Public | 2008 | China | Clinical | 98.6 | 100 | 0 | 99.9846 | 5111637 | 45.29 | 59 | 401611 |
| VP08352 | VP08352 | SAMN20168486 | SRR15099838 | PC | Public | 2008 | China | Clinical | 98.6 | 100 | 0 | 99.9851 | 5110148 | 45.28 | 57 | 427734 |
| VP08353 | VP08353 | SAMN20168523 | SRR15099796 | NonPC | Public | 2008 | China | Clinical | 91.9 | 100 | 0 | 98.4255 | 5056034 | 45.25 | 33 | 524939 |
| VP08354 | VP08354 | SAMN20168524 | SRR15099795 | NonPC | Public | 2008 | China | Clinical | 91.9 | 100 | 0 | 98.4072 | 5060280 | 45.25 | 46 | 410462 |
| VP08355 | VP08355 | SAMN20168525 | SRR15099794 | NonPC | Public | 2008 | China | Clinical | 91.9 | 100 | 0 | 98.4042 | 5055735 | 45.25 | 37 | 524939 |
| VP08356 | VP08356 | SAMN20168526 | SRR15099793 | NonPC | Public | 2008 | China | Clinical | 91.9 | 100 | 0 | 98.431 | 5055649 | 45.25 | 37 | 524939 |
| VP08357 | VP08357 | SAMN20168531 | SRR15099787 | PC | Public | 2008 | China | Clinical | 98.6 | 100 | 0 | 99.9936 | 5111502 | 45.28 | 60 | 347619 |
| VP08358 | VP08358 | SAMN20168527 | SRR15099791 | NonPC | Public | 2008 | China | Clinical | 91.9 | 100 | 0 | 98.3946 | 5055277 | 45.25 | 45 | 351110 |
| VP08359 | VP08359 | SAMN20168528 | SRR15099790 | NonPC | Public | 2008 | China | Clinical | 91.9 | 100 | 0 | 98.4072 | 5055341 | 45.25 | 36 | 524939 |
| VP08360 | VP08360 | SAMN20168529 | SRR15099789 | NonPC | Public | 2008 | China | Clinical | 91.9 | 100 | 0 | 98.4192 | 5056091 | 45.25 | 37 | 522556 |
| VP08361 | VP08361 | SAMN20168530 | SRR15099788 | NonPC | Public | 2008 | China | Clinical | 91.7 | 100 | 0 | 98.3858 | 5043213 | 45.25 | 46 | 417274 |
| VP08362 | VP08362 | SAMN20168518 | SRR15099801 | PC | Public | 2008 | China | Clinical | 98.6 | 100 | 0 | 99.9832 | 5150925 | 45.27 | 63 | 420398 |
| VP08363 | VP08363 | SAMN20168519 | SRR15099800 | PC | Public | 2008 | China | Clinical | 98.5 | 100 | 0 | 99.963 | 5201902 | 45.26 | 60 | 420398 |
| VP08364 | VP08364 | SAMN20168520 | SRR15099799 | PC | Public | 2008 | China | Clinical | 98.6 | 100 | 0 | 99.9921 | 5150932 | 45.27 | 61 | 395231 |
| VP08365 | VP08365 | SAMN20168521 | SRR15099798 | PC | Public | 2008 | China | Clinical | 98.6 | 100 | 0 | 99.9808 | 5150375 | 45.27 | 60 | 420398 |
| VP08366 | VP08366 | SAMN20168482 | SRR15099842 | PC | Public | 2008 | China | Clinical | 98.5 | 100 | 0 | 99.9854 | 5092207 | 45.29 | 55 | 425763 |
| VP08367 | VP08367 | SAMN20168422 | SRR15099287 | PC | Public | 2008 | China | Clinical | 98.6 | 100 | 0 | 99.9919 | 5111359 | 45.28 | 62 | 427734 |
| VP08368 | VP08368 | SAMN20168421 | SRR15099288 | PC | Public | 2008 | China | Clinical | 98.6 | 100 | 0 | 99.9843 | 5110479 | 45.28 | 57 | 427734 |
| VP08369 | VP08369 | SAMN20168289 | SRR15099435 | PC | Public | 2008 | China | Clinical | 98.6 | 100 | 0 | 99.9934 | 5175005 | 45.34 | 59 | 401597 |
| VP08370 | VP08370 | SAMN20168539 | SRR15099778 | PC | Public | 2008 | China | Clinical | 98.6 | 100 | 0 | 99.9898 | 5150610 | 45.27 | 62 | 420264 |
| VP08371 | VP08371 | SAMN20168540 | SRR15099777 | PC | Public | 2008 | China | Clinical | 98.2 | 100 | 0 | 99.9932 | 5126940 | 45.26 | 79 | 267610 |
| VP08372 | VP08372 | SAMN20168541 | SRR15099776 | PC | Public | 2008 | China | Clinical | 98.6 | 100 | 0 | 99.9934 | 5150639 | 45.27 | 63 | 395026 |
| VP08373 | VP08373 | SAMN20168542 | SRR15099775 | PC | Public | 2008 | China | Clinical | 98.6 | 100 | 0 | 99.9821 | 5150851 | 45.27 | 61 | 395228 |
| VP08374 | VP08374 | SAMN20168534 | SRR15099784 | PC | Public | 2008 | China | Clinical | 98 | 100 | 0 | 99.9877 | 5079405 | 45.3 | 64 | 411417 |
| VP08375 | VP08375 | SAMN20168538 | SRR15099779 | PC | Public | 2008 | China | Clinical | 98.6 | 100 | 0 | 99.9817 | 5151378 | 45.27 | 59 | 420265 |
| VP08376 | VP08376 | SAMN20168545 | SRR15099772 | PC | Public | 2008 | China | Clinical | 98.6 | 100 | 0 | 99.9843 | 5150726 | 45.27 | 60 | 395039 |

| | | | | | | | | | | | | | | | | |
|---|---|---|---|---|---|---|---|---|---|---|---|---|---|---|---|---|
| VP08377 | VP08377 | SAMN20168544 | SRR15099773 | NonPC | Public | 2008 | China | Clinical | 91.7 | 100 | 0 | 98.3993 | 5044383 | 45.25 | 45 | 417274 |
| VP08378 | VP08378 | SAMN20168550 | SRR15099766 | PC | Public | 2008 | China | Clinical | 98 | 100 | 0 | 99.9831 | 5079890 | 45.3 | 61 | 411189 |
| VP08379 | VP08379 | SAMN20168549 | SRR15099767 | PC | Public | 2008 | China | Clinical | 97.9 | 100 | 0 | 99.9376 | 5166810 | 45.13 | 63 | 373655 |
| VP08380 | VP08380 | SAMN20168537 | SRR15099780 | PC | Public | 2008 | China | Clinical | 98.6 | 100 | 0 | 99.9826 | 5151105 | 45.27 | 61 | 394853 |
| VP08381 | VP08381 | SAMN20168543 | SRR15099774 | PC | Public | 2008 | China | Clinical | 98.6 | 100 | 0 | 99.992 | 5150391 | 45.27 | 60 | 420398 |
| VP08382 | VP08382 | SAMN20168265 | SRR15099462 | NonPC | Public | 2008 | China | Env | 90 | 100 | 0.34 | 98.3323 | 5145046 | 45.25 | 61 | 257807 |
| VP08383 | VP08383 | SAMN20168547 | SRR15099769 | NonPC | Public | 2008 | China | Clinical | 89.5 | 100 | 0.36 | 98.2799 | 5110299 | 45.05 | 41 | 545809 |
| VP08384 | VP08384 | SAMN20168551 | SRR15099765 | PC | Public | 2008 | China | Clinical | 98.6 | 100 | 0 | 99.9887 | 5222363 | 45.12 | 63 | 420398 |
| VP08385 | VP08385 | SAMN20168554 | SRR15099762 | PC | Public | 2008 | China | Clinical | 98.6 | 100 | 0 | 99.9837 | 5150529 | 45.27 | 61 | 420264 |
| VP08386 | VP08386 | SAMN20168562 | SRR15099753 | PC | Public | 2008 | China | Clinical | 98.6 | 100 | 0 | 99.9902 | 5151140 | 45.27 | 61 | 420264 |
| VP08387 | VP08387 | SAMN20168563 | SRR15099752 | PC | Public | 2008 | China | Clinical | 98.6 | 100 | 0 | 99.9915 | 5149787 | 45.27 | 62 | 394851 |
| VP08388 | VP08388 | SAMN20168558 | SRR15099757 | NonPC | Public | 2008 | China | Clinical | 92.4 | 100 | 0.19 | 98.334 | 5050827 | 45.29 | 42 | 539189 |
| VP08390 | VP08390 | SAMN20168559 | SRR15099756 | NonPC | Public | 2008 | China | Clinical | 92.4 | 100 | 0.19 | 98.3595 | 5051972 | 45.3 | 42 | 539189 |
| VP08391 | VP08391 | SAMN20168561 | SRR15099754 | NonPC | Public | 2008 | China | Clinical | 92.4 | 100 | 0.19 | 98.3476 | 5051926 | 45.3 | 41 | 539189 |
| VP08392 | VP08392 | SAMN20168560 | SRR15099755 | NonPC | Public | 2008 | China | Clinical | 92.4 | 100 | 0.19 | 98.3283 | 5051685 | 45.3 | 41 | 539189 |
| VP08393 | VP08393 | SAMN20168208 | SRR15099524 | NonPC | Public | 2008 | China | Clinical | 92.5 | 100 | 0.05 | 98.3475 | 5066683 | 45.27 | 45 | 450396 |
| VP08394 | VP08394 | SAMN20168533 | SRR15099785 | PC | Public | 2008 | China | Clinical | 98.5 | 100 | 0 | 99.9821 | 5182591 | 45.11 | 64 | 401618 |
| VP08395 | VP08395 | SAMN20168555 | SRR15099761 | PC | Public | 2008 | China | Clinical | 98.6 | 100 | 0 | 99.9894 | 5150986 | 45.27 | 59 | 420398 |
| VP08396 | VP08396 | SAMN20168556 | SRR15099760 | PC | Public | 2008 | China | Clinical | 98.6 | 100 | 0 | 99.9895 | 5150448 | 45.27 | 60 | 420264 |
| VP08397 | VP08397 | SAMN20168557 | SRR15099758 | PC | Public | 2008 | China | Clinical | 98.6 | 100 | 0 | 99.9823 | 5150647 | 45.27 | 59 | 420264 |
| VP08398 | VP08398 | SAMN20168548 | SRR15099768 | NonPC | Public | 2008 | China | Clinical | 90.3 | 100 | 0 | 98.412 | 5032677 | 45.19 | 53 | 453381 |
| VP08401 | VP08401 | SAMN20168552 | SRR15099764 | PC | Public | 2008 | China | Clinical | 97.8 | 100 | 0 | 99.9051 | 5149141 | 45.29 | 48 | 419328 |
| VP08402 | VP08402 | SAMN20168553 | SRR15099763 | PC | Public | 2008 | China | Clinical | 98.4 | 100 | 0 | 99.9827 | 5101023 | 45.29 | 49 | 427754 |
| VP08403 | VP08403 | SAMN20168266 | SRR15099461 | NonPC | Public | 2008 | China | Env | 90.9 | 100 | 0.1 | 98.3365 | 4966918 | 45.36 | 51 | 467292 |
| VP08404 | VP08404 | SAMN20168272 | SRR15099454 | NonPC | Public | 2008 | China | Env | 91 | 99.91 | 0.09 | 98.258 | 5254231 | 45.18 | 74 | 238408 |
| VP08405 | VP08405 | SAMN20168276 | SRR15099450 | NonPC | Public | 2008 | China | Env | 89.3 | 100 | 0 | 98.2606 | 4921958 | 45.43 | 30 | 367963 |
| VP08407 | VP08407 | SAMN20168267 | SRR15099459 | NonPC | Public | 2008 | China | Env | 90.8 | 100 | 0.27 | 98.3463 | 4965898 | 45.38 | 47 | 506679 |
| VP08409 | VP08409 | SAMN20168269 | SRR15099457 | NonPC | Public | 2008 | China | Env | 88.4 | 100 | 0.34 | 98.2216 | 5106250 | 45.21 | 64 | 452978 |
| VP08410 | VP08410 | SAMN20168271 | SRR15099455 | NonPC | Public | 2008 | China | Env | 91.6 | 100 | 0.12 | 98.3913 | 5002202 | 45.32 | 27 | 520011 |
| VP08412 | VP08412 | SAMN20168223 | SRR15099508 | NonPC | Public | 2008 | China | Env | 91.7 | 100 | 0.1 | 98.2684 | 5260942 | 45.08 | 55 | 332597 |
| VP08413 | VP08413 | SAMN20168268 | SRR15099458 | NonPC | Public | 2008 | China | Env | 91.3 | 100 | 0 | 98.3742 | 4950459 | 45.36 | 31 | 464015 |
| VP09001 | VP09001 | SAMN20168622 | SRR15099686 | PC | Public | 2009 | China | Clinical | 98.6 | 100 | 0 | 99.9837 | 5135376 | 45.29 | 62 | 401478 |
| VP09003 | VP09003 | SAMN20168581 | SRR15099732 | NonPC | Public | 2009 | China | Env | 89.7 | 100 | 0 | 98.3672 | 4910110 | 45.43 | 34 | 453237 |
| VP09005 | VP09005 | SAMN20168582 | SRR15099731 | NonPC | Public | 2009 | China | Env | 91.4 | 100 | 0.27 | 98.3418 | 5231235 | 45.21 | 46 | 464869 |
| VP09006 | VP09006 | SAMN20168598 | SRR15099712 | NonPC | Public | 2009 | China | Env | 88.5 | 100 | 0.34 | 98.2045 | 5106833 | 45.21 | 66 | 383506 |
| VP09007 | VP09007 | SAMN20168601 | SRR15099709 | NonPC | Public | 2009 | China | Env | 91.1 | 100 | 0.09 | 98.2698 | 5208316 | 45.16 | 71 | 259371 |
| VP09008 | VP09008 | SAMN20168600 | SRR15099710 | NonPC | Public | 2009 | China | Env | 91.5 | 100 | 0.12 | 98.3959 | 4987066 | 45.31 | 48 | 297392 |
| VP09009 | VP09009 | SAMN20168576 | SRR15099738 | NonPC | Public | 2009 | China | Env | 91.3 | 100 | 0.6 | 98.3807 | 5039409 | 45.35 | 44 | 524123 |
| VP09012 | VP09012 | SAMN20168624 | SRR15099684 | PC | Public | 2009 | China | Clinical | 97.3 | 100 | 0 | 99.899 | 5129880 | 45.28 | 75 | 260119 |
| VP09014 | VP09014 | SAMN20168623 | SRR15099685 | PC | Public | 2009 | China | Clinical | 98.3 | 100 | 0 | 99.9837 | 5093140 | 45.31 | 52 | 427818 |
| VP09015 | VP09015 | SAMN20168627 | SRR15099680 | NonPC | Public | 2009 | China | Clinical | 91.8 | 100 | 0 | 98.3844 | 5059329 | 45.23 | 57 | 453381 |
| VP09016 | VP09016 | SAMN20168577 | SRR15099736 | NonPC | Public | 2009 | China | Clinical | 89.8 | 100 | 0.05 | 98.2998 | 5037222 | 45.38 | 45 | 605253 |
| VP09017 | VP09017 | SAMN20168625 | SRR15099683 | PC | Public | 2009 | China | Clinical | 97.7 | 100 | 0 | 99.9153 | 5171141 | 45.11 | 82 | 268520 |
| VP09018 | VP09018 | SAMN20168629 | SRR15099678 | NonPC | Public | 2009 | China | Clinical | 92.9 | 100 | 0 | 98.3919 | 5170529 | 45.17 | 52 | 370286 |
| VP09019 | VP09019 | SAMN20168626 | SRR15099682 | NonPC | Public | 2009 | China | Clinical | 90.6 | 100 | 0.05 | 98.3433 | 5067902 | 45.26 | 67 | 231056 |
| VP09020 | VP09020 | SAMN20168628 | SRR15099679 | PC | Public | 2009 | China | Clinical | 98.2 | 100 | 0 | 99.9902 | 5127533 | 45.26 | 73 | 267776 |
| VP09021 | VP09021 | SAMN20168630 | SRR15099677 | PC | Public | 2009 | China | Clinical | 97.3 | 100 | 0 | 99.8964 | 5128374 | 45.28 | 73 | 260119 |
| VP09022 | VP09022 | SAMN20168662 | SRR15099642 | PC | Public | 2009 | China | Clinical | 98.4 | 100 | 0.02 | 99.9916 | 5137400 | 45.29 | 68 | 312801 |

| | | | | | | | | | | | | | | | | |
|---|---|---|---|---|---|---|---|---|---|---|---|---|---|---|---|---|
| VP09024 | VP09024 | SAMN20168641 | SRR15099665 | NonPC | Public | 2009 | China | Clinical | 92.5 | 100 | 0.19 | 98.3438 | 5061574 | 45.29 | 54 | 471191 |
| VP09025 | VP09025 | SAMN20168642 | SRR15099664 | PC | Public | 2009 | China | Clinical | 98.5 | 100 | 0 | 99.9811 | 5142121 | 45.27 | 59 | 420398 |
| VP09026 | VP09026 | SAMN20168643 | SRR15099663 | PC | Public | 2009 | China | Clinical | 98.5 | 100 | 0 | 99.99 | 5141186 | 45.27 | 60 | 427734 |
| VP09027 | VP09027 | SAMN20168644 | SRR15099662 | PC | Public | 2009 | China | Clinical | 98.5 | 100 | 0 | 99.9794 | 5141584 | 45.27 | 60 | 420398 |
| VP09028 | VP09028 | SAMN20168645 | SRR15099661 | PC | Public | 2009 | China | Clinical | 98.5 | 100 | 0 | 99.99 | 5144301 | 45.27 | 61 | 420398 |
| VP09029 | VP09029 | SAMN20168637 | SRR15099669 | PC | Public | 2009 | China | Clinical | 98.4 | 100 | 0 | 99.9911 | 5099803 | 45.28 | 57 | 425463 |
| VP09030 | VP09030 | SAMN20168635 | SRR15099672 | NonPC | Public | 2009 | China | Clinical | 92.1 | 100 | 0.19 | 98.3775 | 5035663 | 45.25 | 60 | 314105 |
| VP09031 | VP09031 | SAMN20168648 | SRR15099657 | PC | Public | 2009 | China | Clinical | 98.4 | 100 | 0 | 99.9847 | 5089854 | 45.29 | 57 | 425769 |
| VP09032 | VP09032 | SAMN20168654 | SRR15099651 | PC | Public | 2009 | China | Clinical | 98.5 | 100 | 0 | 99.99 | 5141830 | 45.27 | 60 | 420398 |
| VP09033 | VP09033 | SAMN20168569 | SRR15099745 | NonPC | Public | 2009 | China | Env | 92.1 | 100 | 0.05 | 98.3551 | 5064513 | 45.29 | 47 | 460595 |
| VP09034 | VP09034 | SAMN20168602 | SRR15099708 | NonPC | Public | 2009 | China | Env | 90.9 | 100 | 0 | 98.3486 | 5034735 | 45.35 | 54 | 348867 |
| VP09035 | VP09035 | SAMN20168571 | SRR15099743 | NonPC | Public | 2009 | China | Env | 90.3 | 100 | 0.03 | 98.3266 | 5109467 | 45.26 | 90 | 334269 |
| VP09036 | VP09036 | SAMN20168632 | SRR15099675 | NonPC | Public | 2009 | China | Clinical | 92.3 | 100 | 0.19 | 98.3214 | 5046329 | 45.29 | 64 | 289552 |
| VP09037 | VP09037 | SAMN20168634 | SRR15099673 | PC | Public | 2009 | China | Clinical | 98.5 | 100 | 0 | 99.9936 | 5093957 | 45.29 | 59 | 401597 |
| VP09038 | VP09038 | SAMN20168636 | SRR15099671 | NonPC | Public | 2009 | China | Clinical | 92.1 | 100 | 0.19 | 98.3283 | 5060249 | 45.29 | 50 | 539189 |
| VP09039 | VP09039 | SAMN20168647 | SRR15099658 | PC | Public | 2009 | China | Clinical | 98.6 | 100 | 0 | 99.9861 | 5110680 | 45.28 | 64 | 373614 |
| VP09040 | VP09040 | SAMN20168646 | SRR15099660 | PC | Public | 2009 | China | Clinical | 98.4 | 100 | 0 | 99.9822 | 5188001 | 45.21 | 60 | 425355 |
| VP09041 | VP09041 | SAMN20168649 | SRR15099656 | PC | Public | 2009 | China | Clinical | 98.5 | 100 | 0 | 99.9817 | 5142440 | 45.27 | 59 | 394853 |
| VP09044 | VP09044 | SAMN20168640 | SRR15099666 | PC | Public | 2009 | China | Clinical | 98 | 100 | 0 | 99.9861 | 5080586 | 45.28 | 60 | 426361 |
| VP09045 | VP09045 | SAMN20168633 | SRR15099674 | NonPC | Public | 2009 | China | Clinical | 92.5 | 100 | 0.19 | 98.334 | 5060622 | 45.29 | 50 | 471324 |
| VP09047 | VP09047 | SAMN20168631 | SRR15099676 | PC | Public | 2009 | China | Clinical | 98.6 | 100 | 0 | 99.9775 | 5181733 | 45.12 | 61 | 427763 |
| VP09048 | VP09048 | SAMN20168653 | SRR15099652 | NonPC | Public | 2009 | China | Clinical | 92 | 100 | 0 | 98.3901 | 5076004 | 45.24 | 41 | 572202 |
| VP09049 | VP09049 | SAMN20168650 | SRR15099655 | PC | Public | 2009 | China | Clinical | 98 | 100 | 0 | 99.9822 | 5080070 | 45.3 | 61 | 411189 |
| VP09050 | VP09050 | SAMN20168651 | SRR15099654 | PC | Public | 2009 | China | Clinical | 97.4 | 100 | 0 | 99.914 | 5146973 | 45.26 | 81 | 312479 |
| VP09053 | VP09053 | SAMN20168652 | SRR15099653 | NonPC | Public | 2009 | China | Clinical | 89.3 | 100 | 0 | 98.3115 | 5105381 | 45.28 | 57 | 334271 |
| VP09054 | VP09054 | SAMN20168638 | SRR15099668 | PC | Public | 2009 | China | Clinical | 97.5 | 100 | 0 | 99.9303 | 5077173 | 45.28 | 82 | 267776 |
| VP09055 | VP09055 | SAMN20168659 | SRR15099645 | NonPC | Public | 2009 | China | Clinical | 91.2 | 100 | 0 | 98.3453 | 5147543 | 45.26 | 75 | 151666 |
| VP09056 | VP09056 | SAMN20168639 | SRR15099667 | PC | Public | 2009 | China | Clinical | 98.6 | 100 | 0 | 99.9939 | 5109797 | 45.28 | 59 | 427842 |
| VP09057 | VP09057 | SAMN20168657 | SRR15099647 | PC | Public | 2009 | China | Clinical | 98 | 100 | 0 | 99.992 | 5082076 | 45.31 | 67 | 411303 |
| VP09058 | VP09058 | SAMN20168656 | SRR15099649 | NonPC | Public | 2009 | China | Clinical | 92.5 | 100 | 0.05 | 98.3375 | 5021682 | 45.26 | 42 | 539124 |
| VP09060 | VP09060 | SAMN20168583 | SRR15099730 | NonPC | Public | 2009 | China | Env | 91.2 | 100 | 0 | 98.3268 | 4982573 | 45.38 | 49 | 453588 |
| VP09062 | VP09062 | SAMN20168584 | SRR15099729 | NonPC | Public | 2009 | China | Env | 91.9 | 100 | 0 | 98.3229 | 5151644 | 45.26 | 90 | 334800 |
| VP09063 | VP09063 | SAMN20168585 | SRR15099728 | NonPC | Public | 2009 | China | Env | 89.4 | 99.95 | 0 | 98.3655 | 4904559 | 45.4 | 48 | 382207 |
| VP09064 | VP09064 | SAMN20168661 | SRR15099643 | NonPC | Public | 2009 | China | Clinical | 91.1 | 100 | 0.1 | 98.2362 | 5008158 | 45.36 | 51 | 403376 |
| VP09071 | VP09071 | SAMN20168655 | SRR15099650 | PC | Public | 2009 | China | Clinical | 98.6 | 100 | 0 | 99.9831 | 5151178 | 45.27 | 60 | 420264 |
| VP09072 | VP09072 | SAMN20168658 | SRR15099646 | PC | Public | 2009 | China | Clinical | 98 | 100 | 0 | 99.9935 | 5081502 | 45.28 | 61 | 425947 |
| VP09075 | VP09075 | SAMN20168578 | SRR15099735 | NonPC | Public | 2009 | China | Clinical | 89.5 | 100 | 0.05 | 98.3675 | 4960760 | 45.4 | 43 | 435792 |
| VP09076 | VP09076 | SAMN20168672 | SRR15099631 | NonPC | Public | 2009 | China | Clinical | 91.8 | 100 | 0.19 | 98.385 | 5050841 | 45.28 | 61 | 474672 |
| VP09077 | VP09077 | SAMN20168674 | SRR15099629 | PC | Public | 2009 | China | Clinical | 98.6 | 100 | 0 | 99.9898 | 5150840 | 45.27 | 56 | 420257 |
| VP09078 | VP09078 | SAMN20168675 | SRR15099628 | PC | Public | 2009 | China | Clinical | 98.6 | 100 | 0 | 99.9917 | 5150543 | 45.27 | 60 | 442145 |
| VP09079 | VP09079 | SAMN20168673 | SRR15099630 | PC | Public | 2009 | China | Clinical | 98.5 | 100 | 0 | 99.9843 | 5105612 | 45.28 | 60 | 394826 |
| VP09087 | VP09087 | SAMN20168660 | SRR15099644 | PC | Public | 2009 | China | Clinical | 98 | 100 | 0 | 99.9918 | 5079307 | 45.3 | 59 | 401477 |
| VP09088 | VP09088 | SAMN20168663 | SRR15099641 | NonPC | Public | 2009 | China | Clinical | 89.9 | 100 | 0.2 | 98.2876 | 5235134 | 45.18 | 27 | 683698 |
| VP09089 | VP09089 | SAMN20168666 | SRR15099638 | PC | Public | 2009 | China | Clinical | 98.4 | 100 | 0 | 99.9928 | 5168440 | 45.22 | 64 | 401597 |
| VP09090 | VP09090 | SAMN20168665 | SRR15099639 | NonPC | Public | 2009 | China | Clinical | 92.4 | 100 | 0.19 | 98.3561 | 5020893 | 45.26 | 51 | 471218 |
| VP09091 | VP09091 | SAMN20168667 | SRR15099636 | PC | Public | 2009 | China | Clinical | 98.4 | 100 | 0 | 99.9851 | 5164116 | 45.23 | 62 | 401597 |
| VP09092 | VP09092 | SAMN20168671 | SRR15099632 | PC | Public | 2009 | China | Clinical | 98.6 | 100 | 0 | 99.9827 | 5149219 | 45.27 | 62 | 420264 |
| VP09093 | VP09093 | SAMN20168668 | SRR15099635 | PC | Public | 2009 | China | Clinical | 98 | 100 | 0 | 99.9839 | 5081788 | 45.28 | 60 | 425719 |

| | | | | | | | | | | | | | | | | |
|---|---|---|---|---|---|---|---|---|---|---|---|---|---|---|---|---|
| VP09094 | VP09094 | SAMN20168664 | SRR15099640 | NonPC | Public | 2009 | China | Clinical | 93 | 100 | 0 | 98.328 | 5253552 | 45.18 | 33 | 380943 |
| VP09096 | VP09096 | SAMN20168586 | SRR15099727 | NonPC | Public | 2009 | China | Env | 89.8 | 100 | 0 | 98.2902 | 5032597 | 45.3 | 67 | 350367 |
| VP09097 | VP09097 | SAMN20168587 | SRR15099725 | NonPC | Public | 2009 | China | Env | 90 | 100 | 0.4 | 98.3158 | 5189527 | 45.29 | 48 | 325727 |
| VP09098 | VP09098 | SAMN20168588 | SRR15099724 | NonPC | Public | 2009 | China | Env | 88.3 | 100 | 0.3 | 98.2676 | 5116845 | 45.32 | 59 | 255699 |
| VP09100 | VP09100 | SAMN20168669 | SRR15099634 | PC | Public | 2009 | China | Clinical | 98.6 | 100 | 0 | 99.9839 | 5151309 | 45.27 | 65 | 395029 |
| VP09101 | VP09101 | SAMN20168678 | SRR15099624 | PC | Public | 2009 | China | Clinical | 98 | 100 | 0 | 99.9931 | 5080953 | 45.28 | 63 | 401470 |
| VP09102 | VP09102 | SAMN20168680 | SRR15099622 | NonPC | Public | 2009 | China | Clinical | 90.7 | 100 | 0 | 98.315 | 5109705 | 45.27 | 74 | 314040 |
| VP09103 | VP09103 | SAMN20168682 | SRR15099620 | PC | Public | 2009 | China | Clinical | 98.6 | 100 | 0 | 99.9828 | 5150386 | 45.27 | 60 | 420264 |
| VP09104 | VP09104 | SAMN20168681 | SRR15099621 | PC | Public | 2009 | China | Clinical | 98.4 | 100 | 0 | 99.9862 | 5089675 | 45.29 | 57 | 425769 |
| VP09105 | VP09105 | SAMN20168677 | SRR15099625 | NonPC | Public | 2009 | China | Clinical | 92.3 | 100 | 0.19 | 98.3423 | 5048165 | 45.25 | 48 | 539189 |
| VP09106 | VP09106 | SAMN20168670 | SRR15099633 | PC | Public | 2009 | China | Clinical | 98.6 | 100 | 0 | 99.9884 | 5151309 | 45.27 | 63 | 420398 |
| VP09107 | VP09107 | SAMN20168676 | SRR15099627 | PC | Public | 2009 | China | Clinical | 97.7 | 100 | 0 | 99.9404 | 5118537 | 45.28 | 64 | 347631 |
| VP09108 | VP09108 | SAMN20168679 | SRR15099623 | PC | Public | 2009 | China | Clinical | 93.1 | 100 | 0 | 99.9925 | 5151480 | 45.27 | 63 | 394861 |
| VP09110 | VP09110 | SAMN20168618 | SRR15099690 | PC | Public | 2009 | China | Clinical | 98.6 | 100 | 0 | 99.9842 | 5149412 | 45.27 | 61 | 394770 |
| VP09112 | VP09112 | SAMN20168686 | SRR15099616 | PC | Public | 2009 | China | Clinical | 98 | 100 | 0 | 99.9848 | 5080869 | 45.28 | 61 | 425833 |
| VP09113 | VP09113 | SAMN20168684 | SRR15099618 | PC | Public | 2009 | China | Clinical | 97.7 | 100 | 0 | 99.8861 | 5214542 | 45.27 | 58 | 269679 |
| VP09115 | VP09115 | SAMN20168688 | SRR15099613 | NonPC | Public | 2009 | China | Clinical | 90.9 | 100 | 0.26 | 98.3643 | 5119464 | 45.29 | 68 | 313080 |
| VP09117 | VP09117 | SAMN20168687 | SRR15099614 | NonPC | Public | 2009 | China | Clinical | 90.9 | 100 | 0.05 | 98.3448 | 5070023 | 45.27 | 60 | 449813 |
| VP09118 | VP09118 | SAMN20168689 | SRR15099612 | NonPC | Public | 2009 | China | Clinical | 89.3 | 100 | 0.36 | 98.2905 | 5088250 | 45.06 | 67 | 368517 |
| VP09122 | VP09122 | SAMN20168683 | SRR15099619 | NonPC | Public | 2009 | China | Clinical | 92.8 | 100 | 0 | 98.3197 | 5239718 | 45.18 | 48 | 327584 |
| VP09123 | VP09123 | SAMN20168685 | SRR15099617 | PC | Public | 2009 | China | Clinical | 98.6 | 100 | 0 | 99.9882 | 5149216 | 45.27 | 61 | 420265 |
| VP09125 | VP09125 | SAMN20168695 | SRR15099606 | PC | Public | 2009 | China | Clinical | 98.4 | 100 | 0 | 99.9853 | 5090007 | 45.29 | 60 | 401597 |
| VP09126 | VP09126 | SAMN20168693 | SRR15099608 | PC | Public | 2009 | China | Clinical | 98.6 | 100 | 0 | 99.9892 | 5149329 | 45.27 | 61 | 427206 |
| VP09128 | VP09128 | SAMN20297871 | SRR15184474 | NonPC | Public | 2009 | China | Clinical | 92.3 | 100 | 0.19 | 98.3589 | 5047864 | 45.25 | 49 | 471218 |
| VP09129 | VP09129 | SAMN20168692 | SRR15099609 | PC | Public | 2009 | China | Clinical | 98.6 | 100 | 0 | 99.989 | 5149127 | 45.27 | 63 | 420398 |
| VP09132 | VP09132 | SAMN20168619 | SRR15099689 | PC | Public | 2009 | China | Clinical | 98.6 | 100 | 0 | 99.9904 | 5151018 | 45.27 | 59 | 420398 |
| VP09133 | VP09133 | SAMN20168694 | SRR15099607 | PC | Public | 2009 | China | Clinical | 98.6 | 100 | 0 | 99.9873 | 5151395 | 45.27 | 60 | 420398 |
| VP09138 | VP09138 | SAMN20297895 | SRR15184406 | PC | Public | 2009 | China | Clinical | 98.6 | 100 | 0 | 99.9823 | 5149526 | 45.27 | 60 | 427734 |
| VP09139 | VP09139 | SAMN20297896 | SRR15185001 | PC | Public | 2009 | China | Clinical | 98.4 | 100 | 0 | 99.9845 | 5099082 | 45.28 | 58 | 401701 |
| VP09140 | VP09140 | SAMN20297894 | SRR15184417 | PC | Public | 2009 | China | Clinical | 98.6 | 100 | 0 | 99.9805 | 5149547 | 45.27 | 58 | 427734 |
| VP09141 | VP09141 | SAMN20297897 | SRR15184990 | PC | Public | 2009 | China | Clinical | 98.4 | 100 | 0 | 99.9894 | 5099474 | 45.28 | 58 | 425769 |
| VP09142 | VP09142 | SAMN20297893 | SRR15184428 | PC | Public | 2009 | China | Clinical | 98.4 | 100 | 0 | 99.9845 | 5099285 | 45.28 | 58 | 425769 |
| VP09143 | VP09143 | SAMN20297891 | SRR15184450 | NonPC | Public | 2009 | China | Clinical | 92.5 | 100 | 0.19 | 98.3382 | 5027188 | 45.26 | 49 | 539189 |
| VP09144 | VP09144 | SAMN20297874 | SRR15184856 | NonPC | Public | 2009 | China | Clinical | 92.8 | 100 | 0.27 | 98.3711 | 5144773 | 45.2 | 45 | 488314 |
| VP09145 | VP09145 | SAMN20297870 | SRR15185102 | NonPC | Public | 2009 | China | Clinical | 92 | 100 | 0 | 98.3882 | 5076300 | 45.24 | 39 | 572386 |
| VP09146 | VP09146 | SAMN20168690 | SRR15099611 | NonPC | Public | 2009 | China | Clinical | 92.2 | 100 | 0.19 | 98.3472 | 5049782 | 45.25 | 49 | 539189 |
| VP09147 | VP09147 | SAMN20168691 | SRR15099610 | PC | Public | 2009 | China | Clinical | 97.3 | 100 | 0 | 99.9348 | 5106603 | 45.27 | 83 | 267890 |
| VP09148 | VP09148 | SAMN20297890 | SRR15184461 | PC | Public | 2009 | China | Clinical | 98.6 | 100 | 0 | 99.9818 | 5149059 | 45.27 | 60 | 420398 |
| VP09149 | VP09149 | SAMN20168614 | SRR15099695 | PC | Public | 2009 | China | Clinical | 98.4 | 100 | 0 | 99.9931 | 5090108 | 45.29 | 59 | 425355 |
| VP09150 | VP09150 | SAMN20297892 | SRR15184439 | PC | Public | 2009 | China | Clinical | 98.6 | 100 | 0 | 99.9805 | 5148954 | 45.27 | 56 | 427734 |
| VP09151 | VP09151 | SAMN20297888 | SRR15184592 | NonPC | Public | 2009 | China | Clinical | 92 | 100 | 0 | 98.4053 | 5075853 | 45.24 | 41 | 572386 |
| VP09152 | VP09152 | SAMN20168615 | SRR15099694 | PC | Public | 2009 | China | Clinical | 98.4 | 100 | 0 | 99.9937 | 5098116 | 45.28 | 60 | 394954 |
| VP09154 | VP09154 | SAMN20297898 | SRR15184979 | PC | Public | 2009 | China | Clinical | 98.4 | 100 | 0.27 | 99.9856 | 5132347 | 45.29 | 63 | 394895 |
| VP09155 | VP09155 | SAMN20168589 | SRR15099723 | NonPC | Public | 2009 | China | Env | 87.9 | 100 | 0.36 | 98.1918 | 5056654 | 45.19 | 79 | 195893 |
| VP09156 | VP09156 | SAMN20168574 | SRR15099740 | NonPC | Public | 2009 | China | Env | 87.4 | 100 | 0.36 | 98.2156 | 5010003 | 45.32 | 50 | 383264 |
| VP09157 | VP09157 | SAMN20297872 | SRR15184898 | NonPC | Public | 2009 | China | Clinical | 89.2 | 100 | 0 | 98.3116 | 4986448 | 45.37 | 68 | 455403 |
| VP09158 | VP09158 | SAMN20297873 | SRR15184539 | NonPC | Public | 2009 | China | Clinical | 91.4 | 100 | 0 | 98.3382 | 4981465 | 45.34 | 51 | 419994 |
| VP09159 | VP09159 | SAMN20297877 | SRR15185121 | PC | Public | 2009 | China | Clinical | 98.5 | 100 | 0 | 99.9847 | 5092457 | 45.29 | 58 | 401471 |

| | | | | | | | | | | | | | | | | |
|---|---|---|---|---|---|---|---|---|---|---|---|---|---|---|---|---|
| VP09160 | VP09160 | SAMN20297878 | SRR15185021 | PC | Public | 2009 | China | Clinical | 98 | 99.46 | 0 | 99.9837 | 5064835 | 45.29 | 56 | 425877 |
| VP09161 | VP09161 | SAMN20297876 | SRR15185232 | PC | Public | 2009 | China | Clinical | 98.5 | 100 | 0 | 99.9858 | 5091697 | 45.29 | 56 | 425763 |
| VP09162 | VP09162 | SAMN20297886 | SRR15184614 | PC | Public | 2009 | China | Clinical | 98.5 | 100 | 0 | 99.9858 | 5216305 | 45.11 | 61 | 420398 |
| VP09163 | VP09163 | SAMN20297880 | SRR15185090 | PC | Public | 2009 | China | Clinical | 98.5 | 99.86 | 0 | 99.9888 | 5092520 | 45.29 | 61 | 394825 |
| VP09164 | VP09164 | SAMN20297884 | SRR15184708 | PC | Public | 2009 | China | Clinical | 98.6 | 100 | 0 | 99.9857 | 5224183 | 45.11 | 66 | 363679 |
| VP09165 | VP09165 | SAMN20297885 | SRR15184697 | PC | Public | 2009 | China | Clinical | 98.6 | 100 | 0 | 99.976 | 5225267 | 45.11 | 59 | 420398 |
| VP09166 | VP09166 | SAMN20297881 | SRR15185079 | PC | Public | 2009 | China | Clinical | 98.5 | 100 | 0 | 99.984 | 5092193 | 45.29 | 55 | 425877 |
| VP09167 | VP09167 | SAMN20297887 | SRR15184603 | PC | Public | 2009 | China | Clinical | 98.6 | 100 | 0 | 99.9792 | 5225351 | 45.11 | 62 | 427206 |
| VP09168 | VP09168 | SAMN20297879 | SRR15185101 | PC | Public | 2009 | China | Clinical | 98.5 | 100 | 0 | 99.9849 | 5092661 | 45.29 | 58 | 425763 |
| VP09169 | VP09169 | SAMN20297882 | SRR15185068 | PC | Public | 2009 | China | Clinical | 98.4 | 100 | 0 | 99.9835 | 5092195 | 45.29 | 57 | 401604 |
| VP09170 | VP09170 | SAMN20297875 | SRR15185343 | NonPC | Public | 2009 | China | Clinical | 90.9 | 100 | 0 | 98.3061 | 5142029 | 45.21 | 67 | 395323 |
| VP09171 | VP09171 | SAMN20297883 | SRR15184719 | PC | Public | 2009 | China | Clinical | 98.3 | 100 | 0 | 99.984 | 5143024 | 45.28 | 59 | 401470 |
| VP09173 | VP09173 | SAMN20297903 | SRR15184677 | PC | Public | 2009 | China | Clinical | 98.4 | 100 | 0 | 99.985 | 5090044 | 45.29 | 59 | 425241 |
| VP09174 | VP09174 | SAMN20297869 | SRR15185103 | PC | Public | 2009 | China | Clinical | 98.4 | 100 | 0 | 99.9836 | 5089398 | 45.29 | 60 | 401463 |
| VP09175 | VP09175 | SAMN20168611 | SRR15099698 | PC | Public | 2009 | China | Clinical | 98 | 100 | 0 | 99.9851 | 5081352 | 45.28 | 59 | 426247 |
| VP09176 | VP09176 | SAMN20297899 | SRR15184897 | PC | Public | 2009 | China | Clinical | 98.6 | 100 | 0 | 99.9808 | 5150690 | 45.27 | 60 | 420398 |
| VP09177 | VP09177 | SAMN20168620 | SRR15099688 | PC | Public | 2009 | China | Clinical | 98.6 | 100 | 0 | 99.9887 | 5150133 | 45.27 | 61 | 420264 |
| VP09178 | VP09178 | SAMN20297902 | SRR15184688 | NonPC | Public | 2009 | China | Clinical | 92.4 | 100 | 0.19 | 98.3063 | 5060906 | 45.29 | 48 | 539189 |
| VP09179 | VP09179 | SAMN20297901 | SRR15184875 | NonPC | Public | 2009 | China | Clinical | 91.7 | 100 | 0 | 98.4082 | 5057597 | 45.25 | 49 | 417287 |
| VP09180 | VP09180 | SAMN20297906 | SRR15184644 | PC | Public | 2009 | China | Clinical | 98.6 | 100 | 0 | 99.9895 | 5148727 | 45.27 | 72 | 331602 |
| VP09181 | VP09181 | SAMN20297907 | SRR15184633 | PC | Public | 2009 | China | Clinical | 98.6 | 100 | 0 | 99.9832 | 5151361 | 45.27 | 61 | 420265 |
| VP09182 | VP09182 | SAMN20297909 | SRR15184538 | PC | Public | 2009 | China | Clinical | 98.6 | 100 | 0 | 99.9837 | 5150063 | 45.27 | 61 | 394859 |
| VP09183 | VP09183 | SAMN20297910 | SRR15184527 | NonPC | Public | 2009 | China | Clinical | 92.5 | 100 | 0.05 | 98.3623 | 5045494 | 45.3 | 48 | 370793 |
| VP09184 | VP09184 | SAMN20297911 | SRR15184516 | PC | Public | 2009 | China | Clinical | 98.4 | 100 | 0.27 | 99.9832 | 5131115 | 45.29 | 65 | 288859 |
| VP09185 | VP09185 | SAMN20297913 | SRR15185063 | PC | Public | 2009 | China | Clinical | 98.6 | 100 | 0.27 | 99.9894 | 5191641 | 45.28 | 62 | 427206 |
| VP09186 | VP09186 | SAMN20297904 | SRR15184666 | PC | Public | 2009 | China | Clinical | 98.5 | 100 | 0 | 99.9683 | 5197821 | 45.26 | 60 | 420264 |
| VP09187 | VP09187 | SAMN20297905 | SRR15184655 | PC | Public | 2009 | China | Clinical | 98.6 | 100 | 0 | 99.9909 | 5150036 | 45.27 | 59 | 420264 |
| VP09188 | VP09188 | SAMN20297912 | SRR15184396 | PC | Public | 2009 | China | Clinical | 98.4 | 100 | 0 | 99.9944 | 5089686 | 45.29 | 60 | 425769 |
| VP09189-01 | VP09189-01 | SAMN20297914 | SRR15185052 | NonPC | Public | 2009 | China | Clinical | 92.3 | 100 | 0.19 | 98.3381 | 5048530 | 45.26 | 47 | 539189 |
| VP09190-01 | VP09190-01 | SAMN20297916 | SRR15184959 | NonPC | Public | 2009 | China | Clinical | 92.4 | 100 | 0.05 | 98.3419 | 5069194 | 45.28 | 47 | 450396 |
| VP09191 | VP09191 | SAMN20297915 | SRR15184970 | NonPC | Public | 2009 | China | Clinical | 92.4 | 100 | 0.65 | 98.4168 | 5111774 | 45.23 | 62 | 203654 |
| VP09192 | VP09192 | SAMN20297917 | SRR15184948 | PC | Public | 2009 | China | Clinical | 98.4 | 100 | 0 | 99.9943 | 5135704 | 45.24 | 137 | 86861 |
| VP09193 | VP09193 | SAMN20297918 | SRR15184937 | PC | Public | 2009 | China | Clinical | 98.6 | 100 | 0 | 99.9898 | 5146890 | 45.27 | 140 | 74328 |
| VP09194 | VP09194 | SAMN20297919 | SRR15184855 | NonPC | Public | 2009 | China | Clinical | 92.1 | 100 | 0.05 | 98.4149 | 5065598 | 45.23 | 47 | 312516 |
| VP09195 | VP09195 | SAMN20297921 | SRR15184833 | NonPC | Public | 2009 | China | Clinical | 92.3 | 100 | 0.19 | 98.3719 | 5049656 | 45.25 | 79 | 178211 |
| VP09196 | VP09196 | SAMN20297927 | SRR15185365 | PC | Public | 2009 | China | Clinical | 98.6 | 100 | 0 | 99.9913 | 5150365 | 45.27 | 95 | 158562 |
| VP09197 | VP09197 | SAMN20297925 | SRR15184726 | PC | Public | 2009 | China | Clinical | 98.4 | 100 | 0 | 99.9842 | 5174038 | 45.12 | 101 | 151382 |
| VP09198 | VP09198 | SAMN20297926 | SRR15185376 | PC | Public | 2009 | China | Clinical | 98.6 | 100 | 0 | 99.9917 | 5150964 | 45.27 | 186 | 60307 |
| VP09199-01 | VP09199-01 | SAMN20297930 | SRR15185331 | PC | Public | 2009 | China | Clinical | 98.6 | 100 | 0 | 99.985 | 5150481 | 45.27 | 64 | 288859 |
| VP09200 | VP09200 | SAMN20297931 | SRR15185320 | NonPC | Public | 2009 | China | Clinical | 92 | 100 | 0 | 98.3785 | 5075288 | 45.24 | 38 | 572301 |
| VP09201 | VP09201 | SAMN20297933 | SRR15185298 | PC | Public | 2009 | China | Clinical | 97.8 | 100 | 0.07 | 99.8832 | 5300359 | 45.28 | 68 | 288863 |
| VP09202 | VP09202 | SAMN20297929 | SRR15185342 | PC | Public | 2009 | China | Clinical | 97.3 | 100 | 0 | 99.9115 | 5127829 | 45.28 | 77 | 259761 |
| VP09203 | VP09203 | SAMN20297923 | SRR15184776 | NonPC | Public | 2009 | China | Clinical | 92.9 | 100 | 0 | 98.3866 | 5123299 | 45.22 | 42 | 498054 |
| VP09204 | VP09204 | SAMN20297936 | SRR15185265 | NonPC | Public | 2009 | China | Clinical | 92.3 | 100 | 0.19 | 98.3291 | 5010318 | 45.25 | 54 | 450300 |
| VP09205 | VP09205 | SAMN20297928 | SRR15185354 | PC | Public | 2009 | China | Clinical | 98.5 | 100 | 0 | 99.992 | 5106425 | 45.28 | 67 | 312556 |
| VP09206 | VP09206 | SAMN20297934 | SRR15185287 | NonPC | Public | 2009 | China | Clinical | 90.7 | 100 | 0.18 | 98.3286 | 5429318 | 45.13 | 91 | 269731 |
| VP09208 | VP09208 | SAMN20297937 | SRR15185254 | PC | Public | 2009 | China | Clinical | 98.4 | 100 | 0 | 99.9838 | 5089154 | 45.29 | 55 | 425769 |
| VP09209 | VP09209 | SAMN20297939 | SRR15185231 | NonPC | Public | 2009 | China | Clinical | 92.5 | 100 | 0.19 | 98.3055 | 5060735 | 45.29 | 59 | 262395 |

| | | | | | | | | | | | | | | | | |
|---|---|---|---|---|---|---|---|---|---|---|---|---|---|---|---|---|
| VP09210 | VP09210 | SAMN20297941 | SRR15185209 | PC | Public | 2009 | China | Clinical | 98 | 100 | 0 | 99.9905 | 5079800 | 45.3 | 88 | 179142 |
| VP09211 | VP09211 | SAMN20297942 | SRR15185198 | PC | Public | 2009 | China | Clinical | 98 | 100 | 0 | 99.9926 | 5080924 | 45.28 | 88 | 164413 |
| VP09212 | VP09212 | SAMN20297938 | SRR15185243 | PC | Public | 2009 | China | Clinical | 97.3 | 100 | 0 | 99.9062 | 5177034 | 45.29 | 76 | 259762 |
| VP09213 | VP09213 | SAMN20297940 | SRR15185220 | PC | Public | 2009 | China | Clinical | 98 | 100 | 0 | 99.9388 | 5183005 | 45.13 | 112 | 112776 |
| VP09214 | VP09214 | SAMN20297943 | SRR15185187 | PC | Public | 2009 | China | Clinical | 98.4 | 100 | 0.27 | 99.9844 | 5141385 | 45.29 | 73 | 255565 |
| VP09215 | VP09215 | SAMN20297949 | SRR15185120 | PC | Public | 2009 | China | Clinical | 98 | 100 | 0.09 | 99.9301 | 5217873 | 45.11 | 98 | 186425 |
| VP09216 | VP09216 | SAMN20297948 | SRR15185132 | NonPC | Public | 2009 | China | Clinical | 93 | 100 | 0 | 98.4098 | 5134824 | 45.22 | 51 | 400644 |
| VP09217 | VP09217 | SAMN20297946 | SRR15185154 | PC | Public | 2009 | China | Clinical | 98.2 | 100 | 0 | 99.9881 | 5094197 | 45.31 | 266 | 35303 |
| VP09218 | VP09218 | SAMN20297947 | SRR15185143 | PC | Public | 2009 | China | Clinical | 97.3 | 100 | 0 | 99.8906 | 5109292 | 45.27 | 78 | 259761 |
| VP09219 | VP09219 | SAMN20297952 | SRR15184565 | PC | Public | 2009 | China | Clinical | 98.6 | 100 | 0 | 99.9828 | 5149977 | 45.27 | 71 | 199839 |
| VP09220 | VP09220 | SAMN20168612 | SRR15099697 | PC | Public | 2009 | China | Clinical | 98 | 100 | 0 | 99.9946 | 5082808 | 45.28 | 89 | 179150 |
| VP09221-01 | VP09221-01 | SAMN20168613 | SRR15099696 | PC | Public | 2009 | China | Clinical | 98 | 100 | 0 | 99.9838 | 5083188 | 45.28 | 67 | 288859 |
| VP09227 | VP09227 | SAMN20168603 | SRR15099707 | NonPC | Public | 2009 | China | Clinical | 93.1 | 100 | 0.05 | 98.3849 | 5065454 | 45.29 | 49 | 424363 |
| VP09228 | VP09228 | SAMN20297935 | SRR15185276 | NonPC | Public | 2009 | China | Clinical | 92.9 | 100 | 0.05 | 98.3646 | 5053739 | 45.29 | 54 | 370793 |
| VP09229 | VP09229 | SAMN20297944 | SRR15185176 | PC | Public | 2009 | China | Clinical | 98 | 100 | 0 | 99.9789 | 5080413 | 45.3 | 76 | 190524 |
| VP09231 | VP09231 | SAMN20297951 | SRR15184576 | NonPC | Public | 2009 | China | Clinical | 92.4 | 100 | 0.05 | 98.3278 | 5067947 | 45.27 | 46 | 438749 |
| VP09232 | VP09232 | SAMN20168606 | SRR15099704 | NonPC | Public | 2009 | China | Clinical | 92.3 | 100 | 0.19 | 98.348 | 5015540 | 45.26 | 60 | 289560 |
| VP09233 | VP09233 | SAMN20297945 | SRR15185165 | PC | Public | 2009 | China | Clinical | 98.4 | 100 | 0 | 99.9847 | 5099198 | 45.28 | 83 | 188797 |
| VP09237 | VP09237 | SAMN20297954 | SRR15184507 | NonPC | Public | 2009 | China | Clinical | 92 | 100 | 0 | 98.3649 | 5078200 | 45.24 | 44 | 572734 |
| VP09238 | VP09238 | SAMN20297953 | SRR15184554 | PC | Public | 2009 | China | Clinical | 98.5 | 100 | 0 | 99.9829 | 5141093 | 45.28 | 62 | 394901 |
| VP09240 | VP09240 | SAMN20297969 | SRR15185100 | PC | Public | 2009 | China | Clinical | 98.6 | 100 | 0 | 99.9896 | 5150985 | 45.27 | 97 | 137557 |
| VP09241 | VP09241 | SAMN20297968 | SRR15184741 | PC | Public | 2009 | China | Clinical | 97.7 | 100 | 0 | 99.9201 | 5127954 | 45.28 | 88 | 187017 |
| VP09242 | VP09242 | SAMN20297964 | SRR15184821 | PC | Public | 2009 | China | Clinical | 98.4 | 100 | 0 | 99.9835 | 5089703 | 45.29 | 58 | 425769 |
| VP09243 | VP09243 | SAMN20297962 | SRR15184915 | PC | Public | 2009 | China | Clinical | 97.6 | 100 | 0 | 99.9434 | 5128141 | 45.28 | 56 | 427194 |
| VP09244 | VP09244 | SAMN20297963 | SRR15184904 | PC | Public | 2009 | China | Clinical | 97.1 | 100 | 0 | 99.9233 | 5098599 | 45.27 | 79 | 268004 |
| VP09245 | VP09245 | SAMN20297965 | SRR15184810 | PC | Public | 2009 | China | Clinical | 98.6 | 100 | 0 | 99.9824 | 5193391 | 45.32 | 59 | 440285 |
| VP09246 | VP09246 | SAMN20297973 | SRR15185096 | PC | Public | 2009 | China | Clinical | 98.6 | 100 | 0 | 99.9815 | 5194220 | 45.33 | 75 | 257142 |
| VP09247 | VP09247 | SAMN20297970 | SRR15185099 | NonPC | Public | 2009 | China | Clinical | 92.9 | 100 | 0 | 98.399 | 5123873 | 45.22 | 45 | 498054 |
| VP09250 | VP09250 | SAMN20297972 | SRR15185097 | PC | Public | 2009 | China | Clinical | 98.6 | 100 | 0 | 99.9941 | 5151213 | 45.27 | 75 | 232999 |
| VP09251 | VP09251 | SAMN20297971 | SRR15185098 | PC | Public | 2009 | China | Clinical | 98 | 100 | 0 | 99.984 | 5081635 | 45.28 | 91 | 156607 |
| VP09252 | VP09252 | SAMN20297924 | SRR15184765 | NonPC | Public | 2009 | China | Clinical | 92.5 | 100 | 0.05 | 98.3677 | 5067325 | 45.27 | 46 | 438749 |
| VP09253 | VP09253 | SAMN20297950 | SRR15185109 | NonPC | Public | 2009 | China | Clinical | 92.5 | 100 | 0.05 | 98.335 | 5068438 | 45.27 | 56 | 231526 |
| VP09254 | VP09254 | SAMN20297956 | SRR15184485 | PC | Public | 2009 | China | Clinical | 97.3 | 100 | 0 | 99.9258 | 5106617 | 45.27 | 80 | 267890 |
| VP09255-2 | VP09255-2 | SAMN20297957 | SRR15184473 | PC | Public | 2009 | China | Clinical | 98.1 | 100 | 0 | 99.992 | 5069402 | 45.28 | 81 | 266114 |
| VP09259 | VP09259 | SAMN20297955 | SRR15184496 | NonPC | Public | 2009 | China | Clinical | 92 | 100 | 0 | 98.3798 | 5076415 | 45.24 | 41 | 572202 |
| VP09263 | VP09263 | SAMN20297984 | SRR15185084 | NonPC | Public | 2009 | China | Clinical | 92.5 | 100 | 0.05 | 98.3178 | 5066592 | 45.27 | 44 | 450396 |
| VP09264 | VP09264 | SAMN20297985 | SRR15185083 | NonPC | Public | 2009 | China | Clinical | 92.4 | 100 | 0.05 | 98.3551 | 5068325 | 45.27 | 99 | 97779 |
| VP09265 | VP09265 | SAMN20297986 | SRR15185082 | PC | Public | 2009 | China | Clinical | 98.4 | 100 | 0.02 | 99.991 | 5094368 | 45.29 | 59 | 440655 |
| VP09266 | VP09266 | SAMN20297987 | SRR15185081 | PC | Public | 2009 | China | Clinical | 98.4 | 100 | 0 | 99.983 | 5092338 | 45.29 | 60 | 401597 |
| VP09267 | VP09267 | SAMN20297988 | SRR15185080 | PC | Public | 2009 | China | Clinical | 98.4 | 100 | 0 | 99.9823 | 5091568 | 45.29 | 58 | 401597 |
| VP09270 | VP09270 | SAMN20297983 | SRR15185085 | NonPC | Public | 2009 | China | Clinical | 88.7 | 100 | 0.3 | 98.2388 | 5197919 | 45.17 | 79 | 365858 |
| VP09271 | VP09271 | SAMN20297998 | SRR15185069 | PC | Public | 2009 | China | Clinical | 98.3 | 100 | 0 | 99.9934 | 5150363 | 45.27 | 115 | 107609 |
| VP09272 | VP09272 | SAMN20297997 | SRR15185070 | PC | Public | 2009 | China | Clinical | 98.6 | 100 | 0 | 99.9833 | 5135991 | 45.28 | 57 | 431670 |
| VP09273 | VP09273 | SAMN20297996 | SRR15185071 | PC | Public | 2009 | China | Clinical | 98.5 | 100 | 0 | 99.9935 | 5099209 | 45.29 | 84 | 153202 |
| VP09274 | VP09274 | SAMN20297995 | SRR15185072 | PC | Public | 2009 | China | Clinical | 98.5 | 100 | 0 | 99.9919 | 5100395 | 45.29 | 64 | 425769 |
| VP09276 | VP09276 | SAMN20297994 | SRR15185073 | PC | Public | 2009 | China | Clinical | 98.4 | 100 | 0.27 | 99.9938 | 5141922 | 45.29 | 67 | 288859 |
| VP09277 | VP09277 | SAMN20297993 | SRR15185074 | NonPC | Public | 2009 | China | Clinical | 91 | 100 | 0.17 | 98.2983 | 5308509 | 45.06 | 84 | 334379 |
| VP09283 | VP09283 | SAMN20168568 | SRR15099746 | NonPC | Public | 2009 | China | Env | 91.2 | 100 | 0 | 98.3903 | 5042402 | 45.38 | 64 | 314929 |

| | | | | | | | | | | | | | | | | |
|---|---|---|---|---|---|---|---|---|---|---|---|---|---|---|---|---|
| VP09284 | VP09284 | SAMN20168590 | SRR15099722 | NonPC | Public | 2009 | China | Env | 90.8 | 100 | 0 | 98.3513 | 5136707 | 45.36 | 92 | 179831 |
| VP09285 | VP09285 | SAMN20168570 | SRR15099744 | NonPC | Public | 2009 | China | Env | 88.5 | 100 | 0.6 | 98.1987 | 5164647 | 45.19 | 66 | 452967 |
| VP09286 | VP09286 | SAMN20168591 | SRR15099721 | NonPC | Public | 2009 | China | Env | 89.8 | 100 | 0.04 | 98.3182 | 5137780 | 45.34 | 42 | 407644 |
| VP09287 | VP09287 | SAMN20168599 | SRR15099711 | NonPC | Public | 2009 | China | Env | 88.7 | 100 | 0.3 | 98.2248 | 5200291 | 45.17 | 82 | 313221 |
| VP09288 | VP09288 | SAMN20168567 | SRR15099747 | NonPC | Public | 2009 | China | Env | 91 | 100 | 0 | 98.2987 | 5127905 | 45.19 | 64 | 345896 |
| VP09289 | VP09289 | SAMN20168604 | SRR15099706 | NonPC | Public | 2009 | China | Env | 90.7 | 99.66 | 0.17 | 98.2491 | 5291700 | 45.05 | 91 | 232630 |
| VP09290 | VP09290 | SAMN20168592 | SRR15099720 | NonPC | Public | 2009 | China | Env | 88.4 | 100 | 0.43 | 98.2454 | 5088327 | 45.27 | 58 | 418640 |
| VP09291 | VP09291 | SAMN20168593 | SRR15099719 | NonPC | Public | 2009 | China | Env | 87.7 | 100 | 0.14 | 98.4094 | 4970217 | 45.37 | 103 | 91604 |
| VP09293 | VP09293 | SAMN20168579 | SRR15099734 | NonPC | Public | 2009 | China | Clinical | 91.7 | 100 | 0 | 98.3722 | 5078069 | 45.25 | 61 | 460356 |
| VP09294 | VP09294 | SAMN20168580 | SRR15099733 | NonPC | Public | 2009 | China | Clinical | 89.2 | 100 | 0.07 | 98.307 | 4953632 | 45.36 | 37 | 446690 |
| VP09295 | VP09295 | SAMN20297966 | SRR15184799 | NonPC | Public | 2009 | China | Clinical | 92.3 | 100 | 0.19 | 98.3526 | 5049856 | 45.25 | 56 | 436885 |
| VP09297 | VP09297 | SAMN20297980 | SRR15185088 | NonPC | Public | 2009 | China | Clinical | 92.3 | 100 | 0.6 | 98.3592 | 5187419 | 45.23 | 52 | 344014 |
| VP09298 | VP09298 | SAMN20297889 | SRR15184472 | PC | Public | 2009 | China | Clinical | 98.4 | 100 | 0 | 99.9835 | 5099255 | 45.28 | 124 | 87852 |
| VP09300 | VP09300 | SAMN20297961 | SRR15184926 | PC | Public | 2009 | China | Clinical | 98.6 | 100 | 0 | 99.9942 | 5135473 | 45.29 | 83 | 183099 |
| VP09301 | VP09301 | SAMN20297967 | SRR15184752 | NonPC | Public | 2009 | China | Clinical | 92.3 | 100 | 0.19 | 98.3682 | 5043730 | 45.25 | 78 | 162974 |
| VP09302 | VP09302 | SAMN20297900 | SRR15184886 | NonPC | Public | 2009 | China | Clinical | 89.3 | 100 | 0.36 | 98.2416 | 5090156 | 45.06 | 61 | 379902 |
| VP09303 | VP09303 | SAMN20297932 | SRR15185309 | NonPC | Public | 2009 | China | Clinical | 92.5 | 100 | 0.05 | 98.3298 | 5067895 | 45.27 | 76 | 176764 |
| VP09304 | VP09304 | SAMN20297920 | SRR15184844 | NonPC | Public | 2009 | China | Clinical | 92.2 | 100 | 0.19 | 98.3383 | 5013795 | 45.26 | 75 | 164916 |
| VP09305 | VP09305 | SAMN20297922 | SRR15184787 | PC | Public | 2009 | China | Clinical | 98.4 | 100 | 0 | 99.9849 | 5090342 | 45.29 | 77 | 194807 |
| VP09308 | VP09308 | SAMN20297979 | SRR15185089 | PC | Public | 2009 | China | Clinical | 98.6 | 100 | 0 | 99.9798 | 5150942 | 45.27 | 60 | 420398 |
| VP09309 | VP09309 | SAMN20297975 | SRR15185094 | PC | Public | 2009 | China | Clinical | 98.4 | 100 | 0 | 99.9914 | 5102817 | 45.28 | 62 | 425355 |
| VP09310 | VP09310 | SAMN20297908 | SRR15184622 | NonPC | Public | 2009 | China | Clinical | 92.3 | 100 | 0.05 | 98.3456 | 5040603 | 45.24 | 49 | 450396 |
| VP09311 | VP09311 | SAMN20297959 | SRR15185020 | NonPC | Public | 2009 | China | Clinical | 92.3 | 100 | 0.05 | 98.354 | 5040558 | 45.24 | 48 | 450396 |
| VP09312 | VP09312 | SAMN20298000 | SRR15185066 | PC | Public | 2009 | China | Clinical | 98.2 | 100 | 0 | 99.9919 | 5080651 | 45.29 | 62 | 312684 |
| VP09313 | VP09313 | SAMN20298001 | SRR15185065 | PC | Public | 2009 | China | Clinical | 98.4 | 100 | 0 | 99.9825 | 5090115 | 45.29 | 57 | 425769 |
| VP09314 | VP09314 | SAMN20298002 | SRR15185064 | PC | Public | 2009 | China | Clinical | 98.4 | 100 | 0 | 99.9955 | 5090582 | 45.29 | 69 | 256964 |
| VP09315 | VP09315 | SAMN20297978 | SRR15185091 | PC | Public | 2009 | China | Clinical | 98.4 | 100 | 0 | 99.9954 | 5087949 | 45.29 | 133 | 86295 |
| VP09316 | VP09316 | SAMN20297976 | SRR15185093 | PC | Public | 2009 | China | Clinical | 98.6 | 100 | 0 | 99.9917 | 5150768 | 45.27 | 152 | 62854 |
| VP09317 | VP09317 | SAMN20168616 | SRR15099693 | PC | Public | 2009 | China | Clinical | 98.4 | 100 | 0 | 99.9932 | 5090529 | 45.29 | 56 | 425997 |
| VP09318 | VP09318 | SAMN20168617 | SRR15099691 | PC | Public | 2009 | China | Clinical | 98.3 | 100 | 0 | 99.9858 | 5087220 | 45.29 | 159 | 67579 |
| VP09319 | VP09319 | SAMN20168609 | SRR15099700 | PC | Public | 2009 | China | Clinical | 98 | 100 | 0 | 99.9786 | 5080671 | 45.3 | 64 | 373628 |
| VP09320 | VP09320 | SAMN20168607 | SRR15099702 | PC | Public | 2009 | China | Clinical | 98.4 | 100 | 0 | 99.985 | 5100629 | 45.28 | 56 | 425997 |
| VP09321 | VP09321 | SAMN20297958 | SRR15185032 | NonPC | Public | 2009 | China | Clinical | 92.5 | 100 | 0.19 | 98.3529 | 5028951 | 45.26 | 51 | 471204 |
| VP09322 | VP09322 | SAMN20298003 | SRR15184402 | PC | Public | 2009 | China | Clinical | 97.3 | 100 | 0 | 99.9253 | 5099817 | 45.28 | 85 | 268004 |
| VP09323 | VP09323 | SAMN20298005 | SRR15184723 | PC | Public | 2009 | China | Clinical | 98.4 | 100 | 0.14 | 99.9842 | 5263293 | 45.11 | 64 | 334348 |
| VP09324 | VP09324 | SAMN20298004 | SRR15184724 | PC | Public | 2009 | China | Clinical | 98.4 | 100 | 0.14 | 99.987 | 5262620 | 45.1 | 56 | 425355 |
| VP09325 | VP09325 | SAMN20298008 | SRR15184720 | PC | Public | 2009 | China | Clinical | 98.4 | 100 | 0.14 | 99.9837 | 5263525 | 45.1 | 61 | 397343 |
| VP09327 | VP09327 | SAMN20298009 | SRR15184718 | PC | Public | 2009 | China | Clinical | 98.4 | 100 | 0 | 99.9829 | 5090768 | 45.29 | 60 | 440625 |
| VP09328 | VP09328 | SAMN20298007 | SRR15184721 | NonPC | Public | 2009 | China | Clinical | 92.3 | 100 | 0.19 | 98.3423 | 5049766 | 45.25 | 53 | 450300 |
| VP09329 | VP09329 | SAMN20297992 | SRR15185075 | NonPC | Public | 2009 | China | Clinical | 92.3 | 100 | 0.19 | 98.37 | 5048445 | 45.25 | 51 | 539190 |
| VP09330 | VP09330 | SAMN20298022 | SRR15184704 | NonPC | Public | 2009 | China | Clinical | 92.3 | 100 | 0.19 | 98.3695 | 5048553 | 45.25 | 50 | 450300 |
| VP09331 | VP09331 | SAMN20297990 | SRR15185077 | NonPC | Public | 2009 | China | Clinical | 88.3 | 100 | 0.31 | 98.2341 | 5095615 | 45.3 | 37 | 539783 |
| VP09332 | VP09332 | SAMN20297991 | SRR15185076 | NonPC | Public | 2009 | China | Clinical | 90.9 | 100 | 0.23 | 98.3427 | 5119885 | 45.17 | 53 | 227573 |
| VP09333 | VP09333 | SAMN20298010 | SRR15184717 | PC | Public | 2009 | China | Clinical | 97.6 | 100 | 0 | 99.9245 | 5177880 | 45.1 | 83 | 215465 |
| VP09334 | VP09334 | SAMN20298020 | SRR15184706 | PC | Public | 2009 | China | Clinical | 97.6 | 100 | 0 | 99.9258 | 5174625 | 45.09 | 88 | 215465 |
| VP09336 | VP09336 | SAMN20298013 | SRR15184714 | NonPC | Public | 2009 | China | Clinical | 92.3 | 100 | 0.05 | 98.3492 | 5041126 | 45.24 | 51 | 450396 |
| VP09337 | VP09337 | SAMN20298018 | SRR15184709 | NonPC | Public | 2009 | China | Clinical | 92.3 | 100 | 0.05 | 98.3608 | 5040806 | 45.24 | 47 | 450396 |
| VP09338 | VP09338 | SAMN20298019 | SRR15184707 | NonPC | Public | 2009 | China | Clinical | 92.3 | 100 | 0.05 | 98.3936 | 5041816 | 45.25 | 51 | 450396 |

| | | | | | | | | | | | | | | | | |
|---|---|---|---|---|---|---|---|---|---|---|---|---|---|---|---|---|
| VP09339 | VP09339 | SAMN20298025 | SRR15184701 | PC | Public | 2009 | China | Clinical | 98.6 | 100 | 0 | 99.9891 | 5189032 | 45.32 | 57 | 461740 |
| VP09341 | VP09341 | SAMN20298021 | SRR15184705 | PC | Public | 2009 | China | Clinical | 98.4 | 100 | 0 | 99.9842 | 5090634 | 45.29 | 59 | 401597 |
| VP09342 | VP09342 | SAMN20297977 | SRR15185092 | PC | Public | 2009 | China | Clinical | 98.4 | 100 | 0 | 99.9846 | 5098257 | 45.28 | 56 | 425355 |
| VP09343 | VP09343 | SAMN20298016 | SRR15184711 | PC | Public | 2009 | China | Clinical | 98.4 | 100 | 0 | 99.9844 | 5089307 | 45.29 | 57 | 401597 |
| VP09344 | VP09344 | SAMN20298015 | SRR15184712 | PC | Public | 2009 | China | Clinical | 98.4 | 100 | 0 | 99.9934 | 5090863 | 45.29 | 62 | 394954 |
| VP09345 | VP09345 | SAMN20298011 | SRR15184716 | PC | Public | 2009 | China | Clinical | 98.6 | 100 | 0 | 99.9828 | 5136684 | 45.29 | 62 | 401611 |
| VP09346 | VP09346 | SAMN20298017 | SRR15184710 | PC | Public | 2009 | China | Clinical | 98.4 | 100 | 0 | 99.9842 | 5091616 | 45.29 | 59 | 401330 |
| VP09347 | VP09347 | SAMN20297981 | SRR15185087 | PC | Public | 2009 | China | Clinical | 98.4 | 100 | 0 | 99.9906 | 5132138 | 45.35 | 58 | 401597 |
| VP09348 | VP09348 | SAMN20297974 | SRR15185095 | NonPC | Public | 2009 | China | Clinical | 92.2 | 100 | 0.19 | 98.3385 | 5048438 | 45.25 | 53 | 471204 |
| VP09349 | VP09349 | SAMN20298028 | SRR15184698 | PC | Public | 2009 | China | Clinical | 98.6 | 100 | 0 | 99.9888 | 5150262 | 45.27 | 59 | 420398 |
| VP09350 | VP09350 | SAMN20298029 | SRR15184696 | PC | Public | 2009 | China | Clinical | 98.6 | 100 | 0 | 99.9817 | 5200131 | 45.28 | 62 | 420398 |
| VP09351 | VP09351 | SAMN20298034 | SRR15184691 | PC | Public | 2009 | China | Clinical | 97.3 | 100 | 0.01 | 99.9264 | 5189259 | 45.3 | 88 | 196784 |
| VP09352 | VP09352 | SAMN20298039 | SRR15184613 | PC | Public | 2009 | China | Clinical | 98.6 | 100 | 0 | 99.9792 | 5150618 | 45.27 | 61 | 420264 |
| VP09354 | VP09354 | SAMN20298055 | SRR15184596 | PC | Public | 2009 | China | Clinical | 98.6 | 100 | 0 | 99.9893 | 5202945 | 45.23 | 60 | 420398 |
| VP09355 | VP09355 | SAMN20298053 | SRR15184598 | PC | Public | 2009 | China | Clinical | 98.4 | 100 | 0 | 99.9798 | 5189386 | 45.21 | 58 | 440617 |
| VP09356 | VP09356 | SAMN20298052 | SRR15184599 | NonPC | Public | 2009 | China | Clinical | 92.3 | 100 | 0.19 | 98.3397 | 5075458 | 45.23 | 51 | 450299 |
| VP09357 | VP09357 | SAMN20298054 | SRR15184597 | PC | Public | 2009 | China | Clinical | 98.4 | 100 | 0 | 99.9903 | 5099146 | 45.28 | 57 | 401597 |
| VP09358 | VP09358 | SAMN20298056 | SRR15184595 | PC | Public | 2009 | China | Clinical | 98.4 | 100 | 0 | 99.9837 | 5090108 | 45.29 | 55 | 425769 |
| VP09359 | VP09359 | SAMN20298057 | SRR15184594 | PC | Public | 2009 | China | Clinical | 98.5 | 100 | 0 | 99.9726 | 5199643 | 45.26 | 57 | 461740 |
| VP09360 | VP09360 | SAMN20298060 | SRR15184590 | PC | Public | 2009 | China | Clinical | 97.3 | 100 | 0 | 99.9256 | 5156397 | 45.11 | 68 | 288527 |
| VP09361 | VP09361 | SAMN20298065 | SRR15184585 | PC | Public | 2009 | China | Clinical | 97.3 | 100 | 0 | 99.9259 | 5158370 | 45.11 | 70 | 288527 |
| VP09362 | VP09362 | SAMN20298066 | SRR15184584 | PC | Public | 2009 | China | Clinical | 98.5 | 100 | 0 | 99.9925 | 5106891 | 45.29 | 63 | 394911 |
| VP09363 | VP09363 | SAMN20298064 | SRR15184586 | NonPC | Public | 2009 | China | Clinical | 92.1 | 100 | 0 | 98.3909 | 5075406 | 45.24 | 42 | 572301 |
| VP09364 | VP09364 | SAMN20298067 | SRR15184583 | PC | Public | 2009 | China | Clinical | 98.4 | 100 | 0 | 99.9919 | 5091914 | 45.29 | 59 | 425241 |
| VP09365 | VP09365 | SAMN20298068 | SRR15184582 | NonPC | Public | 2009 | China | Clinical | 92.1 | 100 | 0.06 | 98.4387 | 5155860 | 45.26 | 49 | 312882 |
| VP09366 | VP09366 | SAMN20298058 | SRR15184593 | NonPC | Public | 2009 | China | Clinical | 89.8 | 100 | 0.43 | 98.2767 | 4980908 | 45.4 | 34 | 747130 |
| VP09368 | VP09368 | SAMN20298063 | SRR15184587 | PC | Public | 2009 | China | Clinical | 98.5 | 100 | 0.27 | 99.9911 | 5172647 | 45.29 | 60 | 347570 |
| VP09369 | VP09369 | SAMN20298062 | SRR15184588 | PC | Public | 2009 | China | Clinical | 98.6 | 100 | 0.27 | 99.9902 | 5194891 | 45.27 | 62 | 420398 |
| VP09370 | VP09370 | SAMN20298061 | SRR15184589 | PC | Public | 2009 | China | Clinical | 98.6 | 100 | 0 | 99.9818 | 5149606 | 45.27 | 63 | 395029 |
| VP09371 | VP09371 | SAMN20298030 | SRR15184695 | PC | Public | 2009 | China | Clinical | 98.6 | 100 | 0 | 99.981 | 5148563 | 45.27 | 60 | 427206 |
| VP09372 | VP09372 | SAMN20298027 | SRR15184699 | PC | Public | 2009 | China | Clinical | 97.6 | 100 | 0 | 99.9294 | 5163864 | 45.11 | 82 | 245636 |
| VP09373 | VP09373 | SAMN20298037 | SRR15184616 | PC | Public | 2009 | China | Clinical | 98.6 | 100 | 0 | 99.9813 | 5202982 | 45.23 | 60 | 427740 |
| VP09374 | VP09374 | SAMN20298032 | SRR15184693 | PC | Public | 2009 | China | Clinical | 97.6 | 100 | 0 | 99.9239 | 5175955 | 45.09 | 87 | 215465 |
| VP09375 | VP09375 | SAMN20298033 | SRR15184692 | PC | Public | 2009 | China | Clinical | 97.2 | 100 | 0.07 | 99.9181 | 5180311 | 45.26 | 94 | 197933 |
| VP09376 | VP09376 | SAMN20298035 | SRR15184690 | PC | Public | 2009 | China | Clinical | 98.4 | 100 | 0 | 99.9852 | 5089836 | 45.29 | 56 | 425883 |
| VP09377 | VP09377 | SAMN20298038 | SRR15184615 | PC | Public | 2009 | China | Clinical | 98.6 | 100 | 0 | 99.98 | 5201742 | 45.22 | 63 | 420398 |
| VP09378 | VP09378 | SAMN20298036 | SRR15184617 | PC | Public | 2009 | China | Clinical | 98.4 | 100 | 0 | 99.9844 | 5090000 | 45.29 | 55 | 425883 |
| VP09379 | VP09379 | SAMN20298070 | SRR15184470 | PC | Public | 2009 | China | Clinical | 98.4 | 100 | 0 | 99.9808 | 5099096 | 45.28 | 56 | 425997 |
| VP09380 | VP09380 | SAMN20298069 | SRR15184471 | PC | Public | 2009 | China | Clinical | 98.4 | 100 | 0 | 99.9856 | 5101101 | 45.28 | 61 | 401463 |
| VP09381 | VP09381 | SAMN20298040 | SRR15184612 | NonPC | Public | 2009 | China | Clinical | 92.3 | 100 | 0.19 | 98.3668 | 5043840 | 45.25 | 50 | 539183 |
| VP09382 | VP09382 | SAMN20298041 | SRR15184611 | NonPC | Public | 2009 | China | Clinical | 92.3 | 100 | 0.19 | 98.3505 | 5043673 | 45.25 | 48 | 539183 |
| VP09383 | VP09383 | SAMN20298042 | SRR15184610 | NonPC | Public | 2009 | China | Clinical | 92.3 | 100 | 0.19 | 98.3405 | 5044115 | 45.25 | 49 | 539183 |
| VP09384 | VP09384 | SAMN20298043 | SRR15184609 | NonPC | Public | 2009 | China | Clinical | 92.3 | 100 | 0.19 | 98.3564 | 5045518 | 45.26 | 52 | 471370 |
| VP09385 | VP09385 | SAMN20298044 | SRR15184608 | NonPC | Public | 2009 | China | Clinical | 92.3 | 100 | 0.19 | 98.3307 | 5012928 | 45.25 | 52 | 471370 |
| VP09386 | VP09386 | SAMN20298045 | SRR15184607 | NonPC | Public | 2009 | China | Clinical | 92.3 | 100 | 0.19 | 98.3598 | 5044145 | 45.25 | 49 | 539183 |
| VP09387 | VP09387 | SAMN20298046 | SRR15184606 | NonPC | Public | 2009 | China | Clinical | 92.3 | 100 | 0.19 | 98.3381 | 5044649 | 45.25 | 51 | 471370 |
| VP09388 | VP09388 | SAMN20298047 | SRR15184605 | NonPC | Public | 2009 | China | Clinical | 92.3 | 100 | 0.19 | 98.3528 | 5090156 | 45.14 | 56 | 471204 |
| VP09389 | VP09389 | SAMN20298048 | SRR15184604 | NonPC | Public | 2009 | China | Clinical | 92.3 | 100 | 0.19 | 98.3472 | 5043265 | 45.25 | 48 | 471370 |

| | | | | | | | | | | | | | | | | |
|---|---|---|---|---|---|---|---|---|---|---|---|---|---|---|---|---|
| VP09390 | VP09390 | SAMN20298049 | SRR15184602 | NonPC | Public | 2009 | China | Clinical | 92.3 | 100 | 0.19 | 98.3418 | 5044045 | 45.25 | 50 | 471204 |
| VP09391 | VP09391 | SAMN20298050 | SRR15184601 | NonPC | Public | 2009 | China | Clinical | 92.3 | 100 | 0.19 | 98.3386 | 5045040 | 45.25 | 53 | 471204 |
| VP09392 | VP09392 | SAMN20298051 | SRR15184600 | NonPC | Public | 2009 | China | Clinical | 92.3 | 100 | 0.19 | 98.3415 | 5043941 | 45.25 | 49 | 471370 |
| VP09394 | VP09394 | SAMN20297960 | SRR15185009 | PC | Public | 2009 | China | Clinical | 98.4 | 100 | 0 | 99.9808 | 5099685 | 45.28 | 58 | 401597 |
| VP09395 | VP09395 | SAMN20297982 | SRR15185086 | NonPC | Public | 2009 | China | Clinical | 92.9 | 100 | 0.05 | 98.4012 | 5032855 | 45.31 | 43 | 411537 |
| VP09396 | VP09396 | SAMN20297999 | SRR15185067 | PC | Public | 2009 | China | Clinical | 98.6 | 100 | 0 | 99.9938 | 5151292 | 45.27 | 65 | 288859 |
| VP09397 | VP09397 | SAMN20298006 | SRR15184722 | NonPC | Public | 2009 | China | Clinical | 92 | 100 | 0 | 98.3928 | 5076499 | 45.24 | 41 | 572386 |
| VP09398 | VP09398 | SAMN20298023 | SRR15184703 | PC | Public | 2009 | China | Clinical | 98.5 | 100 | 0.27 | 99.9729 | 5149038 | 45.29 | 65 | 401583 |
| VP09399 | VP09399 | SAMN20298059 | SRR15184591 | NonPC | Public | 2009 | China | Clinical | 92.5 | 100 | 0.05 | 98.3783 | 5052531 | 45.3 | 48 | 370793 |
| VP09400 | VP09400 | SAMN20168608 | SRR15099701 | PC | Public | 2009 | China | Clinical | 98.4 | 100 | 0 | 99.989 | 5098536 | 45.28 | 57 | 425769 |
| VP09401 | VP09401 | SAMN20168610 | SRR15099699 | PC | Public | 2009 | China | Clinical | 98.6 | 100 | 0 | 99.9835 | 5136192 | 45.28 | 60 | 347570 |
| VP09402 | VP09402 | SAMN20168621 | SRR15099687 | PC | Public | 2009 | China | Clinical | 98.6 | 100 | 0 | 99.9816 | 5202091 | 45.22 | 64 | 392281 |
| VP09403 | VP09403 | SAMN20298071 | SRR15184469 | PC | Public | 2009 | China | Clinical | 98.4 | 100 | 0 | 99.9927 | 5098575 | 45.28 | 57 | 401463 |
| VP09404 | VP09404 | SAMN20298072 | SRR15184468 | PC | Public | 2009 | China | Clinical | 98.4 | 100 | 0 | 99.9912 | 5100821 | 45.29 | 60 | 425883 |
| VP09405 | VP09405 | SAMN20298073 | SRR15184467 | PC | Public | 2009 | China | Clinical | 98.4 | 100 | 0 | 99.9927 | 5099575 | 45.28 | 58 | 401597 |
| VP09406 | VP09406 | SAMN20298074 | SRR15184466 | PC | Public | 2009 | China | Clinical | 98.4 | 100 | 0 | 99.9839 | 5098847 | 45.28 | 57 | 401731 |
| VP09408 | VP09408 | SAMN20298075 | SRR15184465 | PC | Public | 2009 | China | Clinical | 98.4 | 100 | 0 | 99.9836 | 5099488 | 45.28 | 58 | 373480 |
| VP09411 | VP09411 | SAMN20298076 | SRR15184464 | PC | Public | 2009 | China | Clinical | 98.4 | 100 | 0.27 | 99.9821 | 5141610 | 45.29 | 58 | 425883 |
| VP09412 | VP09412 | SAMN20298079 | SRR15184460 | PC | Public | 2009 | China | Clinical | 98.4 | 100 | 0.27 | 99.9939 | 5142718 | 45.29 | 62 | 334348 |
| VP09413 | VP09413 | SAMN20298081 | SRR15184458 | PC | Public | 2009 | China | Clinical | 98.4 | 100 | 0 | 99.9839 | 5090316 | 45.29 | 57 | 425355 |
| VP09414 | VP09414 | SAMN20298082 | SRR15184457 | PC | Public | 2009 | China | Clinical | 98.4 | 100 | 0 | 99.9916 | 5090137 | 45.29 | 58 | 425883 |
| VP09415 | VP09415 | SAMN20297989 | SRR15185078 | PC | Public | 2009 | China | Clinical | 98.4 | 100 | 0 | 99.9833 | 5090445 | 45.29 | 59 | 425883 |
| VP09416 | VP09416 | SAMN20298024 | SRR15184702 | PC | Public | 2009 | China | Clinical | 98.5 | 100 | 0.27 | 99.9826 | 5147563 | 45.29 | 63 | 365149 |
| VP09417 | VP09417 | SAMN20298096 | SRR15184442 | PC | Public | 2009 | China | Clinical | 98.6 | 100 | 0 | 99.9831 | 5147504 | 45.27 | 59 | 420398 |
| VP09418 | VP09418 | SAMN20298097 | SRR15184441 | PC | Public | 2009 | China | Clinical | 98.6 | 100 | 0 | 99.9821 | 5148644 | 45.27 | 58 | 420264 |
| VP09419 | VP09419 | SAMN20298095 | SRR15184443 | PC | Public | 2009 | China | Clinical | 98.6 | 100 | 0 | 99.9811 | 5151144 | 45.27 | 59 | 420265 |
| VP09420 | VP09420 | SAMN20298078 | SRR15184462 | NonPC | Public | 2009 | China | Clinical | 89.4 | 100 | 0.05 | 98.3514 | 4904125 | 45.37 | 46 | 448648 |
| VP09421 | VP09421 | SAMN20298083 | SRR15184456 | NonPC | Public | 2009 | China | Clinical | 92.3 | 100 | 0.19 | 98.3386 | 5047839 | 45.25 | 51 | 450300 |
| VP09422 | VP09422 | SAMN20298085 | SRR15184454 | NonPC | Public | 2009 | China | Clinical | 90.9 | 100 | 0.05 | 98.4739 | 5000923 | 45.26 | 46 | 312784 |
| VP09423 | VP09423 | SAMN20298088 | SRR15184451 | PC | Public | 2009 | China | Clinical | 98.6 | 100 | 0 | 99.9815 | 5148937 | 45.27 | 57 | 427734 |
| VP09424 | VP09424 | SAMN20298087 | SRR15184452 | PC | Public | 2009 | China | Clinical | 98.6 | 100 | 0 | 99.9808 | 5151427 | 45.27 | 57 | 427728 |
| VP09425 | VP09425 | SAMN20298092 | SRR15184446 | NonPC | Public | 2009 | China | Clinical | 93 | 100 | 0.01 | 98.3789 | 5266649 | 45.23 | 45 | 488339 |
| VP09426 | VP09426 | SAMN20298089 | SRR15184449 | NonPC | Public | 2009 | China | Clinical | 92.3 | 100 | 0.19 | 98.3046 | 5009586 | 45.25 | 49 | 450300 |
| VP09427 | VP09427 | SAMN20298090 | SRR15184448 | PC | Public | 2009 | China | Clinical | 97.7 | 100 | 0 | 99.9034 | 5148296 | 45.29 | 55 | 419874 |
| VP09428 | VP09428 | SAMN20298091 | SRR15184447 | NonPC | Public | 2009 | China | Clinical | 92 | 100 | 0 | 98.3813 | 5075842 | 45.24 | 44 | 488314 |
| VP09429 | VP09429 | SAMN20298094 | SRR15184444 | PC | Public | 2009 | China | Clinical | 98.4 | 100 | 0 | 99.9915 | 5100419 | 45.28 | 56 | 425997 |
| VP09430 | VP09430 | SAMN20298093 | SRR15184445 | NonPC | Public | 2009 | China | Clinical | 93 | 100 | 0 | 98.3713 | 5174813 | 45.2 | 42 | 488339 |
| VP09431 | VP09431 | SAMN20298098 | SRR15184440 | NonPC | Public | 2009 | China | Clinical | 92.9 | 100 | 0.05 | 98.363 | 5053029 | 45.29 | 56 | 370793 |
| VP09432 | VP09432 | SAMN20298099 | SRR15184438 | NonPC | Public | 2009 | China | Clinical | 92.3 | 100 | 0.19 | 98.3123 | 5009977 | 45.25 | 50 | 450300 |
| VP09433 | VP09433 | SAMN20298103 | SRR15184434 | NonPC | Public | 2009 | China | Clinical | 90.7 | 100 | 0.05 | 98.4299 | 5006706 | 45.33 | 59 | 336312 |
| VP09434 | VP09434 | SAMN20298106 | SRR15184431 | NonPC | Public | 2009 | China | Clinical | 92.3 | 100 | 0.19 | 98.366 | 5048617 | 45.25 | 50 | 450300 |
| VP09435 | VP09435 | SAMN20298104 | SRR15184433 | NonPC | Public | 2009 | China | Clinical | 92 | 100 | 0 | 98.3955 | 5076171 | 45.24 | 39 | 572202 |
| VP09436 | VP09436 | SAMN20298101 | SRR15184436 | PC | Public | 2009 | China | Clinical | 98.2 | 100 | 0 | 99.9656 | 5131372 | 45.3 | 49 | 461740 |
| VP09437 | VP09437 | SAMN20298111 | SRR15184425 | PC | Public | 2009 | China | Clinical | 98.6 | 100 | 0 | 99.9822 | 5150197 | 45.27 | 61 | 394859 |
| VP09438 | VP09438 | SAMN20298108 | SRR15184429 | NonPC | Public | 2009 | China | Clinical | 92.5 | 99.97 | 0.33 | 98.2696 | 5104474 | 45.24 | 85 | 293269 |
| VP09439 | VP09439 | SAMN20298110 | SRR15184426 | NonPC | Public | 2009 | China | Clinical | 92 | 100 | 0.19 | 98.3541 | 5104502 | 45.17 | 54 | 407480 |
| VP09440 | VP09440 | SAMN20298109 | SRR15184427 | NonPC | Public | 2009 | China | Clinical | 92 | 100 | 0 | 98.3817 | 5074769 | 45.24 | 44 | 488314 |
| VP09441 | VP09441 | SAMN20298102 | SRR15184435 | NonPC | Public | 2009 | China | Clinical | 91.7 | 100 | 0 | 98.3725 | 5121128 | 45.26 | 89 | 401646 |

| | | | | | | | | | | | | | | | | |
|---|---|---|---|---|---|---|---|---|---|---|---|---|---|---|---|---|
| VP09442 | VP09442 | SAMN20298012 | SRR15184715 | NonPC | Public | 2009 | China | Clinical | 92.3 | 100 | 0.19 | 98.4008 | 5049517 | 45.25 | 53 | 450397 |
| VP09443 | VP09443 | SAMN20298077 | SRR15184463 | PC | Public | 2009 | China | Clinical | 98.4 | 100 | 0 | 99.9907 | 5099119 | 45.28 | 55 | 425769 |
| VP09444 | VP09444 | SAMN20298084 | SRR15184455 | PC | Public | 2009 | China | Clinical | 98.6 | 100 | 0 | 99.9923 | 5150730 | 45.27 | 68 | 288859 |
| VP09445 | VP09445 | SAMN20298080 | SRR15184459 | PC | Public | 2009 | China | Clinical | 97.9 | 100 | 0.27 | 99.9775 | 5232634 | 45.26 | 61 | 505531 |
| VP09446 | VP09446 | SAMN20168572 | SRR15099742 | NonPC | Public | 2009 | China | Env | 91.6 | 100 | 0.33 | 98.3841 | 4973827 | 45.37 | 86 | 259223 |
| VP09447 | VP09447 | SAMN20168573 | SRR15099741 | NonPC | Public | 2009 | China | Env | 89.4 | 100 | 0 | 98.3815 | 5064154 | 45.22 | 56 | 340430 |
| VP09448 | VP09448 | SAMN20168605 | SRR15099705 | NonPC | Public | 2009 | China | Env | 91.1 | 100 | 0.27 | 98.3538 | 5220818 | 45.29 | 42 | 383492 |
| VP09449 | VP09449 | SAMN20168594 | SRR15099718 | NonPC | Public | 2009 | China | Env | 89.2 | 100 | 0 | 98.3962 | 4974797 | 45.4 | 78 | 336977 |
| VP09451 | VP09451 | SAMN20168595 | SRR15099717 | NonPC | Public | 2009 | China | Env | 90.2 | 100 | 0.27 | 98.2854 | 5273793 | 45.23 | 36 | 578458 |
| VP09452 | VP09452 | SAMN20168596 | SRR15099716 | NonPC | Public | 2009 | China | Env | 91 | 100 | 0.07 | 98.3017 | 5301981 | 45.22 | 80 | 372993 |
| VP09453 | VP09453 | SAMN20168575 | SRR15099739 | NonPC | Public | 2009 | China | Env | 91.6 | 100 | 0.14 | 98.3478 | 5092776 | 45.32 | 60 | 448487 |
| VP09455 | VP09455 | SAMN20168597 | SRR15099713 | NonPC | Public | 2009 | China | Env | 91.2 | 100 | 0 | 98.3187 | 5103616 | 45.38 | 57 | 459290 |
| VP09456 | VP09456 | SAMN20298112 | SRR15184424 | NonPC | Public | 2009 | China | Clinical | 93 | 100 | 0 | 98.3779 | 5175814 | 45.21 | 44 | 572322 |
| VP09457 | VP09457 | SAMN20298113 | SRR15184423 | PC | Public | 2009 | China | Clinical | 98.4 | 100 | 0 | 99.9927 | 5099909 | 45.28 | 56 | 425769 |
| VP09458 | VP09458 | SAMN20298100 | SRR15184437 | PC | Public | 2009 | China | Clinical | 98.6 | 100 | 0 | 99.9842 | 5135888 | 45.29 | 58 | 431778 |
| VP09459 | VP09459 | SAMN20298086 | SRR15184453 | NonPC | Public | 2009 | China | Clinical | 92.3 | 100 | 0.19 | 98.3445 | 5049264 | 45.25 | 53 | 450397 |
| VP09460 | VP09460 | SAMN20298119 | SRR15184416 | NonPC | Public | 2009 | China | Clinical | 92 | 100 | 0 | 98.3812 | 5073818 | 45.24 | 39 | 572734 |
| VP09461 | VP09461 | SAMN20298115 | SRR15184421 | PC | Public | 2009 | China | Clinical | 98.4 | 100 | 0 | 99.9909 | 5179269 | 45.1 | 61 | 425883 |
| VP09462 | VP09462 | SAMN20298117 | SRR15184419 | PC | Public | 2009 | China | Clinical | 98.5 | 100 | 0 | 99.9902 | 5132352 | 45.29 | 55 | 432656 |
| VP09463 | VP09463 | SAMN20298118 | SRR15184418 | PC | Public | 2009 | China | Clinical | 98.6 | 100 | 0.27 | 99.9828 | 5190779 | 45.27 | 60 | 420398 |
| VP09464 | VP09464 | SAMN20298116 | SRR15184420 | NonPC | Public | 2009 | China | Clinical | 90.9 | 100 | 0 | 98.3017 | 5111503 | 45.29 | 70 | 333256 |
| VP09465 | VP09465 | SAMN20168565 | SRR15099750 | NonPC | Public | 2009 | China | Env | 92 | 100 | 0.12 | 98.4108 | 5081984 | 45.27 | 62 | 465106 |
| VP09466 | VP09466 | SAMN20298014 | SRR15184713 | PC | Public | 2009 | China | Clinical | 98.4 | 100 | 0 | 99.9813 | 5099931 | 45.28 | 57 | 425883 |
| VP09467 | VP09467 | SAMN20298026 | SRR15184700 | PC | Public | 2009 | China | Clinical | 98.3 | 100 | 0 | 99.99 | 5133472 | 45.26 | 77 | 288548 |
| VP09468 | VP09468 | SAMN20298031 | SRR15184694 | NonPC | Public | 2009 | China | Clinical | 92.3 | 100 | 0.19 | 98.3467 | 5043885 | 45.25 | 51 | 450300 |
| VP09469 | VP09469 | SAMN20298105 | SRR15184432 | NonPC | Public | 2009 | China | Clinical | 91.8 | 100 | 0 | 98.4289 | 5060040 | 45.23 | 55 | 453381 |
| VP09470 | VP09470 | SAMN20298107 | SRR15184430 | PC | Public | 2009 | China | Clinical | 98.6 | 100 | 0.27 | 99.991 | 5178207 | 45.29 | 60 | 431142 |
| VP09471 | VP09471 | SAMN20298122 | SRR15184413 | PC | Public | 2009 | China | Clinical | 97.3 | 100 | 0 | 99.9276 | 5099223 | 45.28 | 86 | 197933 |
| VP09472 | VP09472 | SAMN20298120 | SRR15184415 | NonPC | Public | 2009 | China | Clinical | 92.3 | 100 | 0.19 | 98.3781 | 5048367 | 45.25 | 52 | 471370 |
| VP09473 | VP09473 | SAMN20298121 | SRR15184414 | PC | Public | 2009 | China | Clinical | 98.4 | 100 | 0 | 99.9847 | 5099227 | 45.29 | 55 | 461733 |
| VP09474 | VP09474 | SAMN20298123 | SRR15184412 | PC | Public | 2009 | China | Clinical | 98.4 | 100 | 0 | 99.9843 | 5089894 | 45.29 | 55 | 425769 |
| VP09475 | VP09475 | SAMN20298127 | SRR15184408 | NonPC | Public | 2009 | China | Clinical | 90.9 | 100 | 0 | 98.2957 | 5257430 | 45.43 | 70 | 374822 |
| VP09476 | VP09476 | SAMN20298114 | SRR15184422 | PC | Public | 2009 | China | Clinical | 98.4 | 100 | 0.27 | 99.9943 | 5132921 | 45.29 | 62 | 312789 |
| VP09477 | VP09477 | SAMN20298126 | SRR15184409 | PC | Public | 2009 | China | Clinical | 98.5 | 100 | 0 | 99.9815 | 5173340 | 45.12 | 61 | 425847 |
| VP09478 | VP09478 | SAMN20298125 | SRR15184410 | PC | Public | 2009 | China | Clinical | 98.4 | 100 | 0 | 99.978 | 5107840 | 45.28 | 56 | 401477 |
| VP09479 | VP09479 | SAMN20298124 | SRR15184411 | NonPC | Public | 2009 | China | Clinical | 92.5 | 100 | 0.33 | 98.3658 | 5052252 | 45.26 | 45 | 450396 |
| VP09481 | VP09481 | SAMN20168564 | SRR15099751 | PC | Public | 2009 | China | Clinical | 97.8 | 100 | 0 | 99.9312 | 5178465 | 45.12 | 67 | 321093 |
| VP09482 | VP09482 | SAMN20168566 | SRR15099749 | NonPC | Public | 2009 | China | Env | 90.4 | 100 | 0.27 | 98.3618 | 5000478 | 45.35 | 34 | 442553 |
| VP10001 | VP10001 | SAMN20298204 | SRR15184682 | PC | Public | 2010 | China | Clinical | 98.5 | 100 | 0 | 99.9924 | 5092799 | 45.29 | 58 | 425883 |
| VP10002 | VP10002 | SAMN20298128 | SRR15184407 | PC | Public | 2009 | China | Clinical | 98.6 | 100 | 0 | 99.9908 | 5148329 | 45.27 | 57 | 420264 |
| VP10003 | VP10003 | SAMN20298205 | SRR15184681 | NonPC | Public | 2010 | China | Clinical | 93 | 100 | 0.05 | 98.4156 | 5092136 | 45.28 | 41 | 312444 |
| VP10004 | VP10004 | SAMN20298150 | SRR15184988 | NonPC | Public | 2010 | China | Env | 88.5 | 100 | 0.33 | 98.2409 | 5176804 | 45.13 | 93 | 321369 |
| VP10005 | VP10005 | SAMN20298151 | SRR15184987 | NonPC | Public | 2010 | China | Env | 91 | 100 | 0.27 | 98.3583 | 5095509 | 45.24 | 58 | 198857 |
| VP10006 | VP10006 | SAMN20298152 | SRR15184986 | NonPC | Public | 2010 | China | Env | 91.1 | 100 | 0.27 | 98.4279 | 4960379 | 45.41 | 47 | 374224 |
| VP10007 | VP10007 | SAMN20298153 | SRR15184985 | NonPC | Public | 2010 | China | Env | 91.2 | 100 | 0.03 | 98.2991 | 5328707 | 45.35 | 76 | 238924 |
| VP10008 | VP10008 | SAMN20298154 | SRR15184984 | NonPC | Public | 2010 | China | Env | 88.5 | 100 | 0.58 | 98.184 | 5121121 | 45.16 | 86 | 307627 |
| VP10009 | VP10009 | SAMN20298155 | SRR15184983 | NonPC | Public | 2010 | China | Env | 91.3 | 100 | 0.14 | 98.3541 | 5055376 | 45.3 | 79 | 257364 |
| VP10010 | VP10010 | SAMN20298156 | SRR15184982 | NonPC | Public | 2010 | China | Env | 91.1 | 100 | 0.05 | 98.3319 | 5073484 | 45.26 | 102 | 200539 |

| | | | | | | | | | | | | | | | | |
|---|---|---|---|---|---|---|---|---|---|---|---|---|---|---|---|---|
| VP10011 | VP10011 | SAMN20298157 | SRR15184981 | NonPC | Public | 2010 | China | Env | 90.2 | 100 | 0.38 | 98.2996 | 5242605 | 45.19 | 78 | 292908 |
| VP10012 | VP10012 | SAMN20298158 | SRR15184980 | NonPC | Public | 2010 | China | Env | 88.5 | 100 | 0.39 | 98.2054 | 5082187 | 45.23 | 83 | 384002 |
| VP10014 | VP10014 | SAMN20298159 | SRR15184978 | NonPC | Public | 2010 | China | Env | 90.8 | 100 | 0.03 | 98.3004 | 5178114 | 45.2 | 77 | 261367 |
| VP10015 | VP10015 | SAMN20298132 | SRR15184401 | NonPC | Public | 2010 | China | Env | 88.1 | 100 | 0.38 | 98.1812 | 5125994 | 45.21 | 71 | 229867 |
| VP10016 | VP10016 | SAMN20298147 | SRR15184992 | NonPC | Public | 2010 | China | Env | 88.7 | 100 | 0.3 | 98.2621 | 5131785 | 45.22 | 85 | 313235 |
| VP10017 | VP10017 | SAMN20298133 | SRR15185008 | NonPC | Public | 2010 | China | Env | 88.1 | 100 | 0.34 | 98.2307 | 5062291 | 45.2 | 44 | 384399 |
| VP10018 | VP10018 | SAMN20298184 | SRR15184880 | NonPC | Public | 2010 | China | Env | 93.1 | 100 | 0.05 | 98.3501 | 5046761 | 45.32 | 31 | 515928 |
| VP10019 | VP10019 | SAMN20298160 | SRR15184977 | NonPC | Public | 2010 | China | Env | 89.6 | 100 | 0 | 98.2951 | 4921122 | 45.42 | 43 | 420334 |
| VP10020 | VP10020 | SAMN20298191 | SRR15184872 | NonPC | Public | 2010 | China | Env | 91.5 | 100 | 0.05 | 98.3372 | 5169549 | 45.24 | 44 | 424435 |
| VP10021 | VP10021 | SAMN20298192 | SRR15184871 | NonPC | Public | 2010 | China | Env | 91.5 | 100 | 0.05 | 98.3512 | 5171285 | 45.25 | 45 | 424435 |
| VP10022 | VP10022 | SAMN20298161 | SRR15184976 | NonPC | Public | 2010 | China | Env | 90.5 | 100 | 0.08 | 98.1918 | 5334131 | 45.14 | 60 | 570545 |
| VP10023 | VP10023 | SAMN20298145 | SRR15184994 | NonPC | Public | 2010 | China | Env | 90 | 100 | 0.11 | 98.2981 | 5042452 | 45.36 | 53 | 728980 |
| VP10024 | VP10024 | SAMN20298136 | SRR15185004 | NonPC | Public | 2010 | China | Env | 89.6 | 100 | 0 | 98.1875 | 5009835 | 45.3 | 38 | 386705 |
| VP10025 | VP10025 | SAMN20298182 | SRR15184882 | NonPC | Public | 2010 | China | Env | 90.3 | 100 | 0 | 98.3737 | 5114705 | 45.34 | 73 | 210037 |
| VP10026 | VP10026 | SAMN20298137 | SRR15185003 | NonPC | Public | 2010 | China | Env | 90.8 | 100 | 0 | 98.3736 | 5021237 | 45.25 | 62 | 260199 |
| VP10027 | VP10027 | SAMN20298162 | SRR15184975 | NonPC | Public | 2010 | China | Env | 90.9 | 100 | 0 | 98.425 | 5122998 | 45.24 | 90 | 258553 |
| VP10028 | VP10028 | SAMN20298163 | SRR15184974 | NonPC | Public | 2010 | China | Env | 88.4 | 100 | 0.4 | 98.294 | 5067551 | 45.27 | 74 | 380530 |
| VP10029 | VP10029 | SAMN20298185 | SRR15184879 | NonPC | Public | 2010 | China | Env | 93.1 | 100 | 0.05 | 98.3593 | 5046513 | 45.31 | 32 | 515928 |
| VP10030 | VP10030 | SAMN20298141 | SRR15184998 | NonPC | Public | 2010 | China | Env | 90.8 | 100 | 0.05 | 98.3329 | 5063018 | 45.3 | 35 | 511600 |
| VP10031 | VP10031 | SAMN20298164 | SRR15184973 | NonPC | Public | 2010 | China | Env | 92 | 100 | 0.27 | 98.3827 | 5056980 | 45.27 | 52 | 401125 |
| VP10032 | VP10032 | SAMN20298144 | SRR15184995 | NonPC | Public | 2010 | China | Env | 88.2 | 100 | 0.41 | 98.1904 | 5267953 | 45.21 | 82 | 313307 |
| VP10033 | VP10033 | SAMN20298148 | SRR15184991 | NonPC | Public | 2010 | China | Clinical | 88.5 | 100 | 0.36 | 98.2109 | 5053806 | 45.21 | 64 | 452278 |
| VP10034 | VP10034 | SAMN20298149 | SRR15184989 | NonPC | Public | 2010 | China | Clinical | 89.5 | 100 | 0.02 | 98.2854 | 5023452 | 45.42 | 87 | 132493 |
| VP10035 | VP10035 | SAMN20298179 | SRR15184885 | NonPC | Public | 2010 | China | Clinical | 92.1 | 100 | 0.5 | 98.3292 | 5368848 | 45.18 | 54 | 462786 |
| VP10036 | VP10036 | SAMN20298186 | SRR15184878 | NonPC | Public | 2010 | China | Env | 93.1 | 100 | 0.05 | 98.3696 | 5103560 | 45.29 | 44 | 463569 |
| VP10037 | VP10037 | SAMN20298165 | SRR15184972 | NonPC | Public | 2010 | China | Env | 88 | 100 | 0.53 | 98.2404 | 5193861 | 45.16 | 110 | 240192 |
| VP10038 | VP10038 | SAMN20298134 | SRR15185006 | NonPC | Public | 2010 | China | Env | 88.6 | 100 | 0.33 | 98.2204 | 5112755 | 45.22 | 56 | 554102 |
| VP10039 | VP10039 | SAMN20298166 | SRR15184902 | NonPC | Public | 2010 | China | Env | 89.6 | 100 | 0.07 | 98.2809 | 5078370 | 45.19 | 57 | 383330 |
| VP10040 | VP10040 | SAMN20298130 | SRR15184404 | NonPC | Public | 2010 | China | Env | 89.9 | 100 | 0 | 98.3155 | 5019893 | 45.37 | 33 | 462918 |
| VP10041 | VP10041 | SAMN20298167 | SRR15184900 | NonPC | Public | 2010 | China | Env | 91.4 | 100 | 0.62 | 98.4074 | 4983199 | 45.36 | 58 | 389993 |
| VP10042 | VP10042 | SAMN20298168 | SRR15184899 | NonPC | Public | 2010 | China | Env | 88.3 | 100 | 0.36 | 98.2469 | 4975975 | 45.3 | 62 | 506728 |
| VP10043 | VP10043 | SAMN20298142 | SRR15184997 | NonPC | Public | 2010 | China | Env | 90.6 | 100 | 0.05 | 98.3551 | 5106065 | 45.24 | 51 | 314988 |
| VP10044 | VP10044 | SAMN20298169 | SRR15184896 | NonPC | Public | 2010 | China | Env | 90.1 | 100 | 0 | 98.32 | 5002679 | 45.36 | 95 | 384392 |
| VP10045 | VP10045 | SAMN20298139 | SRR15185000 | NonPC | Public | 2010 | China | Env | 91.8 | 100 | 0 | 98.4181 | 5071935 | 45.3 | 49 | 425668 |
| VP10046 | VP10046 | SAMN20298188 | SRR15184876 | NonPC | Public | 2010 | China | Env | 92 | 100 | 0 | 98.3812 | 5080053 | 45.24 | 37 | 572351 |
| VP10047 | VP10047 | SAMN20298183 | SRR15184881 | NonPC | Public | 2010 | China | Env | 90.3 | 100 | 0 | 98.3275 | 5115755 | 45.34 | 75 | 203724 |
| VP10048 | VP10048 | SAMN20298131 | SRR15184403 | NonPC | Public | 2010 | China | Env | 88.3 | 99.98 | 0.3 | 98.2964 | 4913925 | 45.39 | 53 | 314573 |
| VP10049 | VP10049 | SAMN20298170 | SRR15184895 | NonPC | Public | 2010 | China | Env | 89.5 | 100 | 0.03 | 98.3043 | 5009859 | 45.32 | 62 | 385252 |
| VP10050 | VP10050 | SAMN20298171 | SRR15184894 | NonPC | Public | 2010 | China | Env | 90 | 100 | 0.05 | 98.2735 | 5139830 | 45.25 | 68 | 314891 |
| VP10051 | VP10051 | SAMN20298138 | SRR15185002 | NonPC | Public | 2010 | China | Env | 90.2 | 100 | 0.08 | 98.4098 | 4942128 | 45.37 | 61 | 313149 |
| VP10052 | VP10052 | SAMN20298190 | SRR15184873 | NonPC | Public | 2010 | China | Env | 92.9 | 100 | 0 | 98.3672 | 5124944 | 45.22 | 42 | 498054 |
| VP10053 | VP10053 | SAMN20298172 | SRR15184893 | NonPC | Public | 2010 | China | Env | 90.7 | 100 | 0.27 | 98.4195 | 4905675 | 45.41 | 43 | 276233 |
| VP10054 | VP10054 | SAMN20298180 | SRR15184884 | NonPC | Public | 2010 | China | Env | 89.9 | 100 | 0.42 | 98.3161 | 4927714 | 45.41 | 51 | 461915 |
| VP10055 | VP10055 | SAMN20298146 | SRR15184993 | NonPC | Public | 2010 | China | Env | 89 | 100 | 0.05 | 98.3038 | 5036582 | 45.24 | 53 | 388201 |
| VP10056 | VP10056 | SAMN20298173 | SRR15184892 | NonPC | Public | 2010 | China | Env | 88.4 | 99.93 | 0.58 | 98.2302 | 5129285 | 45.22 | 87 | 349741 |
| VP10057 | VP10057 | SAMN20298181 | SRR15184883 | NonPC | Public | 2010 | China | Env | 89.9 | 100 | 0.42 | 98.2993 | 4941682 | 45.42 | 46 | 728511 |
| VP10058 | VP10058 | SAMN20298174 | SRR15184891 | NonPC | Public | 2010 | China | Env | 90.2 | 100 | 0 | 98.3715 | 5225463 | 45.19 | 71 | 315748 |
| VP10059 | VP10059 | SAMN20298203 | SRR15184683 | PC | Public | 2010 | China | Env | 98.6 | 100 | 0.27 | 99.9901 | 5192335 | 45.28 | 62 | 420398 |

| | | | | | | | | | | | | | | | | |
|---|---|---|---|---|---|---|---|---|---|---|---|---|---|---|---|---|
| VP10060 | VP10060 | SAMN20298201 | SRR15184685 | PC | Public | 2010 | China | Env | 98.6 | 100 | 0.27 | 99.9829 | 5192610 | 45.28 | 63 | 395233 |
| VP10061 | VP10061 | SAMN20298175 | SRR15184890 | NonPC | Public | 2010 | China | Env | 91.9 | 99.93 | 0 | 98.3146 | 5325657 | 45.15 | 84 | 241028 |
| VP10062 | VP10062 | SAMN20298140 | SRR15184999 | NonPC | Public | 2010 | China | Env | 91.6 | 99.95 | 0.58 | 98.3056 | 5319958 | 45.11 | 63 | 312552 |
| VP10063 | VP10063 | SAMN20298206 | SRR15184680 | PC | Public | 2010 | China | Clinical | 98.4 | 100 | 0 | 99.9842 | 5089888 | 45.29 | 56 | 425883 |
| VP10064 | VP10064 | SAMN20298209 | SRR15184676 | PC | Public | 2010 | China | Clinical | 98.7 | 100 | 0.27 | 99.9811 | 5193493 | 45.28 | 60 | 427734 |
| VP10065 | VP10065 | SAMN20298207 | SRR15184679 | PC | Public | 2010 | China | Clinical | 98.6 | 100 | 0.27 | 99.9829 | 5191935 | 45.28 | 61 | 395233 |
| VP10066 | VP10066 | SAMN20298208 | SRR15184678 | PC | Public | 2010 | China | Clinical | 98.6 | 100 | 0.27 | 99.9831 | 5177358 | 45.29 | 60 | 401580 |
| VP10067 | VP10067 | SAMN20298211 | SRR15184674 | NonPC | Public | 2010 | China | Clinical | 92.1 | 100 | 0.19 | 98.3582 | 4998362 | 45.25 | 57 | 372725 |
| VP10068 | VP10068 | SAMN20298212 | SRR15184673 | PC | Public | 2010 | China | Clinical | 98.6 | 100 | 0.27 | 99.9842 | 5178808 | 45.29 | 61 | 431670 |
| VP10069 | VP10069 | SAMN20298213 | SRR15184672 | NonPC | Public | 2010 | China | Env | 88.2 | 100 | 0.39 | 98.1515 | 5052527 | 45.23 | 63 | 379067 |
| VP10070 | VP10070 | SAMN20298214 | SRR15184671 | NonPC | Public | 2010 | China | Env | 87.2 | 100 | 0.41 | 98.271 | 5123711 | 45.24 | 64 | 341110 |
| VP10071 | VP10071 | SAMN20298215 | SRR15184670 | NonPC | Public | 2010 | China | Env | 88 | 100 | 0.41 | 98.2918 | 5039074 | 45.32 | 63 | 341110 |
| VP10072 | VP10072 | SAMN20298216 | SRR15184669 | PC | Public | 2010 | China | Clinical | 98.5 | 100 | 0 | 99.9928 | 5112505 | 45.28 | 63 | 312692 |
| VP10073 | VP10073 | SAMN20298217 | SRR15184668 | PC | Public | 2010 | China | Clinical | 98.5 | 100 | 0 | 99.9828 | 5113302 | 45.28 | 61 | 394962 |
| VP10074 | VP10074 | SAMN20298218 | SRR15184667 | PC | Public | 2010 | China | Clinical | 98.5 | 100 | 0 | 99.9831 | 5112300 | 45.28 | 61 | 394962 |
| VP10075 | VP10075 | SAMN20298219 | SRR15184665 | PC | Public | 2010 | China | Clinical | 98.4 | 100 | 0 | 99.9722 | 5162320 | 45.28 | 60 | 394901 |
| VP10076 | VP10076 | SAMN20298220 | SRR15184664 | PC | Public | 2010 | China | Clinical | 98.5 | 100 | 0 | 99.9926 | 5201401 | 45.2 | 65 | 394977 |
| VP10077 | VP10077 | SAMN20298221 | SRR15184663 | PC | Public | 2010 | China | Clinical | 98.5 | 100 | 0 | 99.9938 | 5112284 | 45.28 | 63 | 312825 |
| VP10078 | VP10078 | SAMN20298222 | SRR15184662 | PC | Public | 2010 | China | Clinical | 98.5 | 100 | 0 | 99.9837 | 5201644 | 45.2 | 63 | 394962 |
| VP10080 | VP10080 | SAMN20298193 | SRR15184870 | NonPC | Public | 2010 | China | Env | 88.3 | 100 | 0.39 | 98.1916 | 5032741 | 45.26 | 56 | 542137 |
| VP10081 | VP10081 | SAMN20298226 | SRR15184658 | PC | Public | 2010 | China | Clinical | 97.3 | 100 | 0 | 99.9007 | 5130438 | 45.28 | 77 | 260119 |
| VP10082 | VP10082 | SAMN20298223 | SRR15184661 | NonPC | Public | 2010 | China | Clinical | 88.1 | 100 | 0 | 98.3331 | 5121247 | 45.25 | 73 | 293035 |
| VP10084 | VP10084 | SAMN20298225 | SRR15184659 | PC | Public | 2010 | China | Clinical | 97.3 | 100 | 0 | 99.8938 | 5127800 | 45.28 | 72 | 288527 |
| VP10085 | VP10085 | SAMN20298228 | SRR15184656 | NonPC | Public | 2010 | China | Clinical | 92.9 | 100 | 0.05 | 98.3766 | 5093977 | 45.25 | 59 | 420831 |
| VP10086 | VP10086 | SAMN20298227 | SRR15184657 | PC | Public | 2010 | China | Clinical | 98.4 | 100 | 0 | 99.9843 | 5090508 | 45.29 | 54 | 425883 |
| VP10087 | VP10087 | SAMN20298135 | SRR15185005 | NonPC | Public | 2010 | China | Clinical | 90 | 100 | 0.09 | 98.3073 | 5084351 | 45.29 | 62 | 450936 |
| VP10088 | VP10088 | SAMN20298233 | SRR15184650 | PC | Public | 2010 | China | Clinical | 98.5 | 100 | 0 | 99.993 | 5092988 | 45.29 | 55 | 401597 |
| VP10089 | VP10089 | SAMN20298235 | SRR15184648 | PC | Public | 2010 | China | Clinical | 97.3 | 100 | 0 | 99.9089 | 5128835 | 45.28 | 73 | 260119 |
| VP10090 | VP10090 | SAMN20298237 | SRR15184646 | NonPC | Public | 2010 | China | Clinical | 89.3 | 100 | 0 | 98.3122 | 4973041 | 45.31 | 35 | 383367 |
| VP10091 | VP10091 | SAMN20298239 | SRR15184643 | NonPC | Public | 2010 | China | Clinical | 92.4 | 100 | 0.05 | 98.3196 | 5066749 | 45.27 | 46 | 450396 |
| VP10092 | VP10092 | SAMN20298241 | SRR15184641 | PC | Public | 2010 | China | Clinical | 98.5 | 100 | 0 | 99.9838 | 5092453 | 45.29 | 57 | 401604 |
| VP10093 | VP10093 | SAMN20298238 | SRR15184645 | NonPC | Public | 2010 | China | Clinical | 92.9 | 100 | 0 | 98.3901 | 5132228 | 45.21 | 43 | 488300 |
| VP10094 | VP10094 | SAMN20298197 | SRR15184866 | PC | Public | 2010 | China | Clinical | 98.6 | 100 | 0 | 99.9922 | 5150119 | 45.27 | 62 | 395031 |
| VP10095 | VP10095 | SAMN20298242 | SRR15184640 | PC | Public | 2010 | China | Clinical | 98.5 | 100 | 0 | 99.9923 | 5092524 | 45.29 | 55 | 425847 |
| VP10096 | VP10096 | SAMN20298231 | SRR15184652 | PC | Public | 2010 | China | Clinical | 98.6 | 100 | 0 | 99.9824 | 5151682 | 45.27 | 59 | 420398 |
| VP10097 | VP10097 | SAMN20298236 | SRR15184647 | NonPC | Public | 2010 | China | Clinical | 92.5 | 100 | 0.05 | 98.3377 | 5066262 | 45.27 | 45 | 450396 |
| VP10098 | VP10098 | SAMN20298229 | SRR15184654 | PC | Public | 2010 | China | Clinical | 98.6 | 100 | 0 | 99.981 | 5149898 | 45.27 | 60 | 420412 |
| VP10099 | VP10099 | SAMN20298232 | SRR15184651 | PC | Public | 2010 | China | Clinical | 98.6 | 100 | 0 | 99.9809 | 5151408 | 45.27 | 60 | 427734 |
| VP10100 | VP10100 | SAMN20298244 | SRR15184638 | PC | Public | 2010 | China | Clinical | 98.5 | 100 | 0 | 99.9815 | 5130388 | 45.29 | 56 | 401613 |
| VP10101 | VP10101 | SAMN20298198 | SRR15184689 | PC | Public | 2010 | China | Clinical | 98.6 | 100 | 0 | 99.9806 | 5149714 | 45.27 | 62 | 420412 |
| VP10102 | VP10102 | SAMN20298250 | SRR15184631 | PC | Public | 2010 | China | Clinical | 98.5 | 100 | 0 | 99.9833 | 5093276 | 45.29 | 57 | 440649 |
| VP10103 | VP10103 | SAMN20298246 | SRR15184636 | PC | Public | 2010 | China | Clinical | 97.5 | 100 | 0 | 99.9277 | 5143473 | 45.12 | 83 | 267776 |
| VP10104 | VP10104 | SAMN20298245 | SRR15184637 | PC | Public | 2010 | China | Clinical | 98.4 | 100 | 0 | 99.984 | 5091228 | 45.29 | 61 | 440624 |
| VP10105 | VP10105 | SAMN20298249 | SRR15184632 | NonPC | Public | 2010 | China | Clinical | 92.9 | 100 | 0 | 98.3893 | 5130380 | 45.21 | 40 | 488300 |
| VP10106 | VP10106 | SAMN20298252 | SRR15184629 | PC | Public | 2010 | China | Clinical | 98 | 100 | 0 | 99.9936 | 5070011 | 45.28 | 81 | 266000 |
| VP10107 | VP10107 | SAMN20298251 | SRR15184630 | PC | Public | 2010 | China | Clinical | 97.3 | 100 | 0 | 99.9078 | 5136248 | 45.28 | 74 | 260005 |
| VP10108 | VP10108 | SAMN20298253 | SRR15184628 | PC | Public | 2010 | China | Clinical | 97.4 | 100 | 0 | 99.9278 | 5151583 | 45.12 | 71 | 268960 |
| VP10109 | VP10109 | SAMN20298254 | SRR15184627 | NonPC | Public | 2010 | China | Clinical | 92.4 | 100 | 0.65 | 98.421 | 5155151 | 45.18 | 66 | 195956 |

| | | | | | | | | | | | | | | | | |
|---|---|---|---|---|---|---|---|---|---|---|---|---|---|---|---|---|
| VP10110 | VP10110 | SAMN20298255 | SRR15184626 | NonPC | Public | 2010 | China | Clinical | 90.6 | 100 | 0.27 | 98.2644 | 5268463 | 45.27 | 36 | 449697 |
| VP10111 | VP10111 | SAMN20298199 | SRR15184687 | PC | Public | 2010 | China | Clinical | 98.6 | 100 | 0 | 99.9907 | 5150029 | 45.27 | 62 | 420278 |
| VP10112 | VP10112 | SAMN20298200 | SRR15184686 | PC | Public | 2010 | China | Clinical | 98.6 | 100 | 0.27 | 99.9924 | 5192754 | 45.28 | 66 | 332167 |
| VP10113 | VP10113 | SAMN20298210 | SRR15184675 | NonPC | Public | 2010 | China | Clinical | 91.9 | 100 | 0 | 98.4044 | 5094874 | 45.24 | 36 | 524940 |
| VP10114 | VP10114 | SAMN20298248 | SRR15184634 | PC | Public | 2010 | China | Clinical | 97.9 | 100 | 0 | 99.9298 | 5191824 | 45.11 | 57 | 401231 |
| VP10115 | VP10115 | SAMN20298259 | SRR15184621 | PC | Public | 2010 | China | Clinical | 98.6 | 100 | 0.09 | 99.9819 | 5152958 | 45.27 | 68 | 347606 |
| VP10116 | VP10116 | SAMN20298258 | SRR15184623 | PC | Public | 2010 | China | Clinical | 98.6 | 100 | 0 | 99.9833 | 5150311 | 45.27 | 61 | 331485 |
| VP10117 | VP10117 | SAMN20298266 | SRR15184542 | PC | Public | 2010 | China | Clinical | 98.4 | 100 | 0 | 99.9847 | 5089980 | 45.29 | 58 | 425889 |
| VP10118 | VP10118 | SAMN20298270 | SRR15184536 | NonPC | Public | 2010 | China | Clinical | 91.7 | 100 | 0 | 98.4194 | 5082591 | 45.24 | 45 | 417696 |
| VP10119 | VP10119 | SAMN20298260 | SRR15184620 | PC | Public | 2010 | China | Clinical | 98 | 100 | 0 | 99.9844 | 5171938 | 45.2 | 64 | 426367 |
| VP10121 | VP10121 | SAMN20298285 | SRR15184520 | PC | Public | 2010 | China | Clinical | 98.6 | 100 | 0 | 99.981 | 5157295 | 45.27 | 60 | 427206 |
| VP10123 | VP10123 | SAMN20298286 | SRR15184519 | PC | Public | 2010 | China | Clinical | 98.6 | 100 | 0 | 99.9885 | 5155927 | 45.26 | 59 | 427206 |
| VP10124 | VP10124 | SAMN20298289 | SRR15184515 | PC | Public | 2010 | China | Clinical | 98.6 | 100 | 0 | 99.9904 | 5157185 | 45.27 | 61 | 420398 |
| VP10125 | VP10125 | SAMN20298288 | SRR15184517 | PC | Public | 2010 | China | Clinical | 98.3 | 100 | 0 | 99.9915 | 5134752 | 45.25 | 80 | 267646 |
| VP10126 | VP10126 | SAMN20298287 | SRR15184518 | PC | Public | 2010 | China | Clinical | 98.6 | 100 | 0 | 99.9828 | 5157578 | 45.27 | 62 | 420398 |
| VP10127 | VP10127 | SAMN20298271 | SRR15184535 | PC | Public | 2010 | China | Clinical | 98.4 | 100 | 0.01 | 99.9922 | 5184165 | 45.32 | 62 | 401597 |
| VP10128 | VP10128 | SAMN20298202 | SRR15184684 | PC | Public | 2010 | China | Clinical | 98.6 | 100 | 0 | 99.9885 | 5201082 | 45.22 | 58 | 440583 |
| VP10129 | VP10129 | SAMN20298262 | SRR15184618 | PC | Public | 2010 | China | Clinical | 98.4 | 100 | 0 | 99.9851 | 5092268 | 45.29 | 58 | 425883 |
| VP10130 | VP10130 | SAMN20298263 | SRR15184546 | PC | Public | 2010 | China | Clinical | 98.6 | 100 | 0 | 99.9933 | 5137643 | 45.29 | 60 | 431772 |
| VP10131 | VP10131 | SAMN20298257 | SRR15184624 | PC | Public | 2010 | China | Clinical | 98.4 | 100 | 0 | 99.9933 | 5091528 | 45.29 | 60 | 401597 |
| VP10133 | VP10133 | SAMN20298300 | SRR15184394 | NonPC | Public | 2010 | China | Clinical | 91.7 | 100 | 0 | 98.4101 | 5082461 | 45.24 | 43 | 417696 |
| VP10134 | VP10134 | SAMN20298302 | SRR15184392 | PC | Public | 2010 | China | Clinical | 97.3 | 100 | 0 | 99.9101 | 5142517 | 45.28 | 75 | 259891 |
| VP10135 | VP10135 | SAMN20298301 | SRR15184393 | NonPC | Public | 2010 | China | Clinical | 91.9 | 100 | 0 | 98.3747 | 5108967 | 45.23 | 43 | 522568 |
| VP10136 | VP10136 | SAMN20298309 | SRR15185062 | PC | Public | 2010 | China | Clinical | 98.4 | 100 | 0.01 | 99.9918 | 5184266 | 45.32 | 60 | 425769 |
| VP10137 | VP10137 | SAMN20298303 | SRR15184391 | PC | Public | 2010 | China | Clinical | 97.3 | 100 | 0 | 99.9122 | 5144394 | 45.28 | 77 | 260119 |
| VP10138 | VP10138 | SAMN20298299 | SRR15184395 | NonPC | Public | 2010 | China | Clinical | 91.9 | 100 | 0 | 98.4421 | 5064265 | 45.24 | 35 | 522563 |
| VP10139 | VP10139 | SAMN20298304 | SRR15184390 | PC | Public | 2010 | China | Clinical | 97.3 | 100 | 0 | 99.9084 | 5142328 | 45.28 | 75 | 259777 |
| VP10140 | VP10140 | SAMN20298305 | SRR15184389 | PC | Public | 2010 | China | Clinical | 97.3 | 100 | 0 | 99.9018 | 5141845 | 45.28 | 74 | 259891 |
| VP10141 | VP10141 | SAMN20298310 | SRR15185061 | PC | Public | 2010 | China | Clinical | 98.4 | 100 | 0.01 | 99.9936 | 5182544 | 45.31 | 61 | 394895 |
| VP10143 | VP10143 | SAMN20298306 | SRR15184388 | PC | Public | 2010 | China | Clinical | 97.3 | 100 | 0 | 99.9096 | 5141155 | 45.28 | 73 | 260119 |
| VP10144 | VP10144 | SAMN20298267 | SRR15184541 | PC | Public | 2010 | China | Clinical | 98.7 | 100 | 0 | 99.9807 | 5151397 | 45.27 | 58 | 420398 |
| VP10145 | VP10145 | SAMN20298268 | SRR15184540 | PC | Public | 2010 | China | Clinical | 98.6 | 100 | 0 | 99.9799 | 5151686 | 45.27 | 60 | 427170 |
| VP10146 | VP10146 | SAMN20298307 | SRR15184387 | PC | Public | 2010 | China | Clinical | 97.3 | 100 | 0 | 99.909 | 5140441 | 45.28 | 72 | 260119 |
| VP10147 | VP10147 | SAMN20298296 | SRR15184399 | NonPC | Public | 2010 | China | Env | 89.4 | 100 | 0.03 | 98.2941 | 4984878 | 45.35 | 40 | 449508 |
| VP10148 | VP10148 | SAMN20298297 | SRR15184398 | NonPC | Public | 2010 | China | Env | 89.5 | 100 | 0.03 | 98.3212 | 5017492 | 45.29 | 32 | 1330560 |
| VP10149 | VP10149 | SAMN20298298 | SRR15184397 | NonPC | Public | 2010 | China | Env | 89.5 | 100 | 0.05 | 98.3554 | 4958479 | 45.39 | 58 | 462134 |
| VP10150 | VP10150 | SAMN20298308 | SRR15184386 | PC | Public | 2010 | China | Clinical | 98.6 | 100 | 0 | 99.9809 | 5150682 | 45.27 | 61 | 420398 |
| VP10151 | VP10151 | SAMN20298275 | SRR15184531 | NonPC | Public | 2010 | China | Clinical | 91.9 | 100 | 0 | 98.4084 | 5094042 | 45.24 | 34 | 524940 |
| VP10152 | VP10152 | SAMN20298274 | SRR15184532 | PC | Public | 2010 | China | Clinical | 97.7 | 100 | 0 | 99.9298 | 5154947 | 45.29 | 59 | 459027 |
| VP10153 | VP10153 | SAMN20298276 | SRR15184530 | NonPC | Public | 2010 | China | Clinical | 91.7 | 100 | 0 | 98.398 | 5083022 | 45.24 | 42 | 417696 |
| VP10154 | VP10154 | SAMN20298277 | SRR15184529 | NonPC | Public | 2010 | China | Clinical | 91.9 | 100 | 0 | 98.4035 | 5093532 | 45.24 | 34 | 522568 |
| VP10155 | VP10155 | SAMN20298278 | SRR15184528 | NonPC | Public | 2010 | China | Clinical | 91.9 | 100 | 0 | 98.4337 | 5093828 | 45.24 | 36 | 524940 |
| VP10156 | VP10156 | SAMN20298291 | SRR15184513 | NonPC | Public | 2010 | China | Clinical | 92.1 | 100 | 0.05 | 98.3284 | 5066685 | 45.27 | 46 | 450396 |
| VP10157 | VP10157 | SAMN20298284 | SRR15184521 | PC | Public | 2010 | China | Clinical | 98.6 | 100 | 0 | 99.9911 | 5150956 | 45.27 | 59 | 420186 |
| VP10158 | VP10158 | SAMN20298283 | SRR15184522 | PC | Public | 2010 | China | Clinical | 98.6 | 100 | 0 | 99.9904 | 5151111 | 45.27 | 59 | 420186 |
| VP10159 | VP10159 | SAMN20298280 | SRR15184525 | NonPC | Public | 2010 | China | Clinical | 91.7 | 100 | 0 | 98.384 | 5084016 | 45.24 | 44 | 417692 |
| VP10160 | VP10160 | SAMN20298187 | SRR15184877 | NonPC | Public | 2010 | China | Clinical | 91.9 | 100 | 0 | 98.4 | 5094953 | 45.24 | 40 | 353152 |
| VP10161 | VP10161 | SAMN20298290 | SRR15184514 | NonPC | Public | 2010 | China | Clinical | 92.3 | 100 | 0.19 | 98.372 | 5044359 | 45.25 | 51 | 450300 |

| | | | | | | | | | | | | | | | | |
|---|---|---|---|---|---|---|---|---|---|---|---|---|---|---|---|---|
| VP10162 | VP10162 | SAMN20298281 | SRR15184524 | NonPC | Public | 2010 | China | Clinical | 91.9 | 100 | 0 | 98.4187 | 5094383 | 45.24 | 33 | 524940 |
| VP10163 | VP10163 | SAMN20298282 | SRR15184523 | PC | Public | 2010 | China | Clinical | 98.6 | 100 | 0 | 99.9856 | 5149002 | 45.27 | 59 | 420412 |
| VP10164 | VP10164 | SAMN20298261 | SRR15184619 | PC | Public | 2010 | China | Clinical | 97.2 | 100 | 0 | 99.9338 | 5098735 | 45.27 | 83 | 267646 |
| VP10165 | VP10165 | SAMN20298279 | SRR15184526 | NonPC | Public | 2010 | China | Clinical | 91.9 | 100 | 0 | 98.392 | 5094450 | 45.24 | 35 | 522568 |
| VP10167 | VP10167 | SAMN20298264 | SRR15184544 | NonPC | Public | 2010 | China | Clinical | 91.9 | 100 | 0 | 98.3979 | 5094250 | 45.24 | 33 | 522568 |
| VP10168 | VP10168 | SAMN20298269 | SRR15184537 | NonPC | Public | 2010 | China | Clinical | 90 | 100 | 0.45 | 98.3122 | 5029989 | 45.29 | 57 | 391294 |
| VP10170 | VP10170 | SAMN20298294 | SRR15184510 | PC | Public | 2010 | China | Clinical | 98.4 | 100 | 0 | 99.992 | 5090427 | 45.29 | 57 | 425313 |
| VP10171 | VP10171 | SAMN20298292 | SRR15184512 | PC | Public | 2010 | China | Clinical | 98.5 | 100 | 0.01 | 99.9911 | 5198635 | 45.31 | 62 | 394955 |
| VP10172 | VP10172 | SAMN20298293 | SRR15184511 | NonPC | Public | 2010 | China | Clinical | 92.4 | 100 | 0.05 | 98.343 | 5052656 | 45.3 | 46 | 539131 |
| VP10174 | VP10174 | SAMN20298314 | SRR15185057 | PC | Public | 2010 | China | Clinical | 98 | 100 | 0 | 99.9926 | 5082616 | 45.28 | 62 | 401604 |
| VP10175 | VP10175 | SAMN20298295 | SRR15184400 | NonPC | Public | 2010 | China | Clinical | 91.9 | 100 | 0 | 98.4023 | 5093446 | 45.24 | 34 | 524940 |
| VP10176 | VP10176 | SAMN20298311 | SRR15185060 | PC | Public | 2010 | China | Clinical | 98.4 | 100 | 0 | 99.9905 | 5090346 | 45.29 | 57 | 425355 |
| VP10177 | VP10177 | SAMN20298323 | SRR15185047 | PC | Public | 2010 | China | Clinical | 98.4 | 100 | 0 | 99.9843 | 5089634 | 45.29 | 57 | 425355 |
| VP10178 | VP10178 | SAMN20298322 | SRR15185048 | NonPC | Public | 2010 | China | Clinical | 92.5 | 100 | 0.05 | 98.3307 | 5051905 | 45.3 | 44 | 539131 |
| VP10179 | VP10179 | SAMN20298321 | SRR15185049 | NonPC | Public | 2010 | China | Clinical | 91.9 | 100 | 0 | 98.4046 | 5093851 | 45.24 | 32 | 524940 |
| VP10180 | VP10180 | SAMN20298312 | SRR15185059 | NonPC | Public | 2010 | China | Clinical | 92.3 | 100 | 0.05 | 98.3625 | 5042154 | 45.29 | 53 | 373056 |
| VP10181 | VP10181 | SAMN20298313 | SRR15185058 | NonPC | Public | 2010 | China | Clinical | 92.4 | 100 | 0.05 | 98.3483 | 5051340 | 45.3 | 43 | 478570 |
| VP10182 | VP10182 | SAMN20298315 | SRR15185056 | PC | Public | 2010 | China | Clinical | 98.4 | 100 | 0 | 99.9845 | 5090397 | 45.29 | 59 | 401587 |
| VP10183 | VP10183 | SAMN20298319 | SRR15185051 | PC | Public | 2010 | China | Clinical | 97.3 | 100 | 0 | 99.892 | 5180063 | 45.27 | 75 | 288527 |
| VP10185 | VP10185 | SAMN20298329 | SRR15184969 | NonPC | Public | 2010 | China | Clinical | 91.9 | 100 | 0 | 98.4017 | 5094402 | 45.24 | 34 | 524940 |
| VP10186 | VP10186 | SAMN20298332 | SRR15184966 | NonPC | Public | 2010 | China | Clinical | 91.9 | 100 | 0 | 98.4153 | 5094486 | 45.24 | 33 | 524940 |
| VP10187 | VP10187 | SAMN20298331 | SRR15184967 | NonPC | Public | 2010 | China | Clinical | 91.9 | 100 | 0 | 98.3951 | 5102095 | 45.25 | 33 | 522568 |
| VP10188 | VP10188 | SAMN20298333 | SRR15184965 | NonPC | Public | 2010 | China | Clinical | 91.9 | 100 | 0 | 98.4019 | 5094264 | 45.24 | 34 | 524940 |
| VP10189 | VP10189 | SAMN20298327 | SRR15185043 | NonPC | Public | 2010 | China | Clinical | 90.8 | 100 | 0.33 | 98.3093 | 5232459 | 45.12 | 67 | 332141 |
| VP10190 | VP10190 | SAMN20298189 | SRR15184874 | NonPC | Public | 2010 | China | Clinical | 92.9 | 100 | 0 | 98.3799 | 5132183 | 45.21 | 41 | 453475 |
| VP10191 | VP10191 | SAMN20298230 | SRR15184653 | PC | Public | 2010 | China | Clinical | 98.4 | 100 | 0 | 99.9936 | 5090691 | 45.29 | 54 | 425355 |
| VP10192 | VP10192 | SAMN20298234 | SRR15184649 | PC | Public | 2010 | China | Clinical | 97.3 | 100 | 0 | 99.9069 | 5142185 | 45.28 | 74 | 259891 |
| VP10193 | VP10193 | SAMN20298224 | SRR15184660 | PC | Public | 2010 | China | Clinical | 97.6 | 100 | 0 | 99.9238 | 5176230 | 45.09 | 85 | 215465 |
| VP10194 | VP10194 | SAMN20298240 | SRR15184642 | PC | Public | 2010 | China | Clinical | 98.6 | 100 | 0 | 99.9808 | 5151151 | 45.27 | 61 | 394851 |
| VP10195 | VP10195 | SAMN20298256 | SRR15184625 | PC | Public | 2010 | China | Clinical | 98.5 | 100 | 0 | 99.9919 | 5093405 | 45.29 | 59 | 401604 |
| VP10196 | VP10196 | SAMN20298243 | SRR15184639 | PC | Public | 2010 | China | Clinical | 98.4 | 100 | 0 | 99.9824 | 5088295 | 45.29 | 55 | 401463 |
| VP10197 | VP10197 | SAMN20298247 | SRR15184635 | PC | Public | 2010 | China | Clinical | 98.6 | 100 | 0 | 99.9838 | 5136377 | 45.29 | 61 | 401632 |
| VP10199 | VP10199 | SAMN20298265 | SRR15184543 | PC | Public | 2010 | China | Clinical | 98.4 | 100 | 0 | 99.9935 | 5090577 | 45.29 | 57 | 401463 |
| VP10200 | VP10200 | SAMN20298272 | SRR15184534 | PC | Public | 2010 | China | Clinical | 98.4 | 100 | 0 | 99.9844 | 5090964 | 45.29 | 61 | 394810 |
| VP10201 | VP10201 | SAMN20298273 | SRR15184533 | PC | Public | 2010 | China | Clinical | 98.4 | 100 | 0 | 99.9848 | 5090686 | 45.29 | 58 | 394939 |
| VP10202 | VP10202 | SAMN20298316 | SRR15185055 | PC | Public | 2010 | China | Clinical | 98.7 | 100 | 0 | 99.9829 | 5154166 | 45.27 | 65 | 394853 |
| VP10203 | VP10203 | SAMN20298317 | SRR15185054 | PC | Public | 2010 | China | Clinical | 98.6 | 100 | 0 | 99.9809 | 5150804 | 45.27 | 59 | 420264 |
| VP10204 | VP10204 | SAMN20298318 | SRR15185053 | PC | Public | 2010 | China | Clinical | 98.6 | 100 | 0 | 99.9813 | 5150216 | 45.27 | 56 | 427206 |
| VP10206 | VP10206 | SAMN20298338 | SRR15184960 | PC | Public | 2010 | China | Clinical | 98.6 | 100 | 0 | 99.9914 | 5198153 | 45.21 | 61 | 395233 |
| VP10207 | VP10207 | SAMN20298337 | SRR15184961 | PC | Public | 2010 | China | Clinical | 98.5 | 100 | 0 | 99.9941 | 5093520 | 45.29 | 59 | 401604 |
| VP10208 | VP10208 | SAMN20298325 | SRR15185045 | PC | Public | 2010 | China | Clinical | 98 | 100 | 0 | 99.9915 | 5082309 | 45.28 | 61 | 401604 |
| VP10209 | VP10209 | SAMN20298330 | SRR15184968 | PC | Public | 2010 | China | Clinical | 98.6 | 100 | 0 | 99.9877 | 5199126 | 45.21 | 59 | 420398 |
| VP10210 | VP10210 | SAMN20298350 | SRR15184946 | PC | Public | 2010 | China | Clinical | 98.7 | 100 | 0 | 99.9883 | 5150843 | 45.27 | 61 | 394851 |
| VP10211 | VP10211 | SAMN20298351 | SRR15184945 | PC | Public | 2010 | China | Clinical | 98.6 | 100 | 0 | 99.9904 | 5240823 | 45.2 | 60 | 420131 |
| VP10212 | VP10212 | SAMN20298352 | SRR15184944 | PC | Public | 2010 | China | Clinical | 98.6 | 100 | 0 | 99.9873 | 5150962 | 45.27 | 61 | 394851 |
| VP10213 | VP10213 | SAMN20298340 | SRR15184957 | NonPC | Public | 2010 | China | Clinical | 92.3 | 100 | 0.2 | 98.346 | 5135103 | 45.28 | 54 | 450300 |
| VP10214 | VP10214 | SAMN20298344 | SRR15184953 | PC | Public | 2010 | China | Clinical | 97.3 | 100 | 0 | 99.9277 | 5095980 | 45.27 | 86 | 196784 |
| VP10215 | VP10215 | SAMN20298343 | SRR15184954 | PC | Public | 2010 | China | Clinical | 97.3 | 100 | 0 | 99.9318 | 5097183 | 45.27 | 85 | 197933 |

| | | | | | | | | | | | | | | | | |
|---|---|---|---|---|---|---|---|---|---|---|---|---|---|---|---|---|
| VP10216 | VP10216 | SAMN20298341 | SRR15184956 | NonPC | Public | 2010 | China | Clinical | 91.9 | 100 | 0 | 98.4111 | 5094505 | 45.24 | 34 | 524940 |
| VP10217 | VP10217 | SAMN20298324 | SRR15185046 | NonPC | Public | 2010 | China | Clinical | 91.7 | 100 | 0 | 98.4006 | 5083294 | 45.24 | 44 | 417704 |
| VP10218 | VP10218 | SAMN20298176 | SRR15184889 | NonPC | Public | 2010 | China | Env | 88.1 | 100 | 0.37 | 98.2139 | 5138031 | 45.24 | 67 | 385300 |
| VP10219 | VP10219 | SAMN20298194 | SRR15184869 | NonPC | Public | 2010 | China | Env | 88.3 | 100 | 0.39 | 98.2304 | 5116874 | 45.18 | 56 | 542137 |
| VP10221 | VP10221 | SAMN20298143 | SRR15184996 | NonPC | Public | 2010 | China | Env | 88.3 | 100 | 0.37 | 98.2062 | 5186026 | 45.24 | 99 | 298381 |
| VP10222 | VP10222 | SAMN20298177 | SRR15184888 | NonPC | Public | 2010 | China | Env | 88.4 | 100 | 0.3 | 98.2305 | 5124835 | 45.23 | 61 | 418867 |
| VP10224 | VP10224 | SAMN20298129 | SRR15184405 | NonPC | Public | 2010 | China | Env | 90.7 | 100 | 0.27 | 98.3762 | 4982273 | 45.36 | 36 | 449007 |
| VP10225 | VP10225 | SAMN20298346 | SRR15184951 | PC | Public | 2010 | China | Clinical | 97.6 | 100 | 0 | 99.9033 | 5161908 | 45.29 | 52 | 440598 |
| VP10226 | VP10226 | SAMN20298336 | SRR15184962 | PC | Public | 2010 | China | Clinical | 97.3 | 100 | 0 | 99.9003 | 5141070 | 45.28 | 75 | 259761 |
| VP10227 | VP10227 | SAMN20298349 | SRR15184947 | NonPC | Public | 2010 | China | Clinical | 91.9 | 100 | 0 | 98.4185 | 5093734 | 45.24 | 34 | 522568 |
| VP10228 | VP10228 | SAMN20298345 | SRR15184952 | NonPC | Public | 2010 | China | Clinical | 92.5 | 100 | 0.05 | 98.3388 | 5087583 | 45.26 | 45 | 450396 |
| VP10229 | VP10229 | SAMN20298347 | SRR15184950 | PC | Public | 2010 | China | Clinical | 98.5 | 100 | 0 | 99.9915 | 5092424 | 45.29 | 56 | 425847 |
| VP10230 | VP10230 | SAMN20298196 | SRR15184867 | PC | Public | 2010 | China | Clinical | 98.5 | 100 | 0 | 99.9927 | 5126523 | 45.25 | 58 | 401477 |
| VP10231 | VP10231 | SAMN20298354 | SRR15184942 | NonPC | Public | 2010 | China | Clinical | 91.9 | 100 | 0 | 98.3996 | 5094599 | 45.24 | 36 | 524940 |
| VP10232 | VP10232 | SAMN20298355 | SRR15184941 | PC | Public | 2010 | China | Clinical | 97.2 | 100 | 0 | 99.9347 | 5101960 | 45.27 | 85 | 267776 |
| VP10233 | VP10233 | SAMN20298353 | SRR15184943 | PC | Public | 2010 | China | Clinical | 97.3 | 100 | 0 | 99.9123 | 5141936 | 45.28 | 73 | 259987 |
| VP10235 | VP10235 | SAMN20298328 | SRR15184971 | NonPC | Public | 2010 | China | Clinical | 91.3 | 100 | 0.27 | 98.3958 | 5048947 | 45.31 | 64 | 240713 |
| VP10236 | VP10236 | SAMN20298357 | SRR15184939 | PC | Public | 2010 | China | Clinical | 97.3 | 100 | 0 | 99.9002 | 5143310 | 45.28 | 73 | 260119 |
| VP10237 | VP10237 | SAMN20298356 | SRR15184940 | PC | Public | 2010 | China | Clinical | 97.3 | 100 | 0 | 99.9394 | 5099078 | 45.28 | 85 | 197933 |
| VP10238 | VP10238 | SAMN20298358 | SRR15184938 | PC | Public | 2010 | China | Clinical | 97.3 | 100 | 0 | 99.9116 | 5142162 | 45.28 | 73 | 259777 |
| VP10239 | VP10239 | SAMN20298359 | SRR15184936 | PC | Public | 2010 | China | Clinical | 97.7 | 100 | 0 | 99.913 | 5160363 | 45.29 | 56 | 373635 |
| VP10240 | VP10240 | SAMN20298339 | SRR15184958 | NonPC | Public | 2010 | China | Clinical | 91.9 | 100 | 0 | 98.4244 | 5094259 | 45.24 | 33 | 524940 |
| VP10241 | VP10241 | SAMN20298365 | SRR15184860 | PC | Public | 2010 | China | Clinical | 97.3 | 100 | 0 | 99.9227 | 5168228 | 45.12 | 86 | 197933 |
| VP10242 | VP10242 | SAMN20298360 | SRR15184865 | PC | Public | 2010 | China | Clinical | 98.6 | 100 | 0 | 99.9823 | 5152778 | 45.28 | 61 | 420398 |
| VP10244 | VP10244 | SAMN20298335 | SRR15184963 | PC | Public | 2010 | China | Clinical | 98.4 | 100 | 0 | 99.9846 | 5089551 | 45.29 | 57 | 440623 |
| VP10245 | VP10245 | SAMN20298178 | SRR15184887 | NonPC | Public | 2010 | China | Clinical | 92.3 | 100 | 0.46 | 98.3393 | 5200408 | 45.19 | 67 | 470429 |
| VP10246 | VP10246 | SAMN20298362 | SRR15184863 | PC | Public | 2010 | China | Clinical | 98.6 | 100 | 0 | 99.9891 | 5155933 | 45.26 | 59 | 420264 |
| VP10247 | VP10247 | SAMN20298368 | SRR15184857 | PC | Public | 2010 | China | Clinical | 98.6 | 100 | 0 | 99.981 | 5151318 | 45.27 | 58 | 420264 |
| VP10248 | VP10248 | SAMN20298334 | SRR15184964 | PC | Public | 2010 | China | Clinical | 98.6 | 100 | 0 | 99.9873 | 5150084 | 45.27 | 59 | 427206 |
| VP10249 | VP10249 | SAMN20298320 | SRR15185050 | PC | Public | 2010 | China | Clinical | 98.4 | 100 | 0 | 99.9839 | 5134538 | 45.24 | 60 | 425355 |
| VP10250 | VP10250 | SAMN20298371 | SRR15184852 | NonPC | Public | 2010 | China | Clinical | 92.7 | 100 | 0 | 98.437 | 5116962 | 45.21 | 55 | 369125 |
| VP10251 | VP10251 | SAMN20298376 | SRR15184847 | PC | Public | 2010 | China | Clinical | 97.3 | 100 | 0 | 99.9245 | 5097577 | 45.27 | 82 | 268877 |
| VP10252 | VP10252 | SAMN20298369 | SRR15184854 | NonPC | Public | 2010 | China | Clinical | 92.9 | 100 | 0 | 98.3671 | 5135324 | 45.22 | 41 | 498068 |
| VP10253 | VP10253 | SAMN20298367 | SRR15184858 | PC | Public | 2010 | China | Clinical | 98 | 100 | 0 | 99.989 | 5067592 | 45.27 | 79 | 265756 |
| VP10254 | VP10254 | SAMN20298364 | SRR15184861 | NonPC | Public | 2010 | China | Clinical | 92.5 | 100 | 0.05 | 98.3095 | 5087835 | 45.26 | 45 | 539125 |
| VP10256 | VP10256 | SAMN20298366 | SRR15184859 | NonPC | Public | 2010 | China | Clinical | 91.9 | 100 | 0 | 98.4276 | 5094786 | 45.24 | 36 | 483538 |
| VP10257 | VP10257 | SAMN20298370 | SRR15184853 | NonPC | Public | 2010 | China | Clinical | 92.9 | 100 | 0 | 98.3838 | 5132980 | 45.21 | 45 | 572297 |
| VP10258 | VP10258 | SAMN20298374 | SRR15184849 | PC | Public | 2010 | China | Clinical | 98.6 | 100 | 0 | 99.991 | 5156713 | 45.27 | 64 | 420264 |
| VP10259 | VP10259 | SAMN20298384 | SRR15184838 | PC | Public | 2010 | China | Clinical | 98.4 | 100 | 0 | 99.9844 | 5090194 | 45.29 | 61 | 394954 |
| VP10260 | VP10260 | SAMN20298382 | SRR15184840 | PC | Public | 2010 | China | Clinical | 97.2 | 100 | 0 | 99.9272 | 5095969 | 45.27 | 85 | 196861 |
| VP10261 | VP10261 | SAMN20298380 | SRR15184842 | PC | Public | 2010 | China | Clinical | 98 | 100 | 0 | 99.9226 | 5187144 | 45.13 | 62 | 427747 |
| VP10262 | VP10262 | SAMN20298390 | SRR15184831 | PC | Public | 2010 | China | Clinical | 98.4 | 100 | 0 | 99.993 | 5089986 | 45.29 | 58 | 425355 |
| VP10263 | VP10263 | SAMN20298386 | SRR15184836 | NonPC | Public | 2010 | China | Clinical | 92.3 | 100 | 0.05 | 98.377 | 5077539 | 45.26 | 55 | 313852 |
| VP10264 | VP10264 | SAMN20298387 | SRR15184835 | NonPC | Public | 2010 | China | Clinical | 92.4 | 100 | 0.05 | 98.3512 | 5088142 | 45.26 | 46 | 450396 |
| VP10267 | VP10267 | SAMN20298399 | SRR15184786 | NonPC | Public | 2010 | China | Clinical | 91.9 | 100 | 0.19 | 98.3471 | 5048519 | 45.25 | 52 | 450397 |
| VP10268 | VP10268 | SAMN20298379 | SRR15184843 | PC | Public | 2010 | China | Clinical | 98.5 | 100 | 0 | 99.9841 | 5093555 | 45.29 | 59 | 394955 |
| VP10269 | VP10269 | SAMN20298378 | SRR15184845 | PC | Public | 2010 | China | Clinical | 98.6 | 100 | 0 | 99.9887 | 5157329 | 45.27 | 57 | 420398 |
| VP10271 | VP10271 | SAMN20298342 | SRR15184955 | PC | Public | 2010 | China | Clinical | 98.5 | 100 | 0 | 99.9832 | 5092485 | 45.29 | 62 | 312690 |

| | | | | | | | | | | | | | | | | |
|---|---|---|---|---|---|---|---|---|---|---|---|---|---|---|---|---|
| VP10273 | VP10273 | SAMN20298363 | SRR15184862 | PC | Public | 2010 | China | Clinical | 98.4 | 100 | 0 | 99.9811 | 5135551 | 45.28 | 58 | 431250 |
| VP10274 | VP10274 | SAMN20298373 | SRR15184850 | PC | Public | 2010 | China | Clinical | 98.6 | 100 | 0 | 99.9917 | 5149333 | 45.27 | 58 | 420398 |
| VP10275 | VP10275 | SAMN20298372 | SRR15184851 | PC | Public | 2010 | China | Clinical | 98.3 | 100 | 0 | 99.9833 | 5094737 | 45.31 | 53 | 427818 |
| VP10276 | VP10276 | SAMN20298398 | SRR15184788 | NonPC | Public | 2010 | China | Clinical | 92.3 | 100 | 0.19 | 98.3604 | 5046143 | 45.26 | 53 | 450300 |
| VP10277 | VP10277 | SAMN20298401 | SRR15184784 | PC | Public | 2010 | China | Clinical | 98.6 | 100 | 0 | 99.9883 | 5151408 | 45.27 | 58 | 420398 |
| VP10278 | VP10278 | SAMN20298408 | SRR15184777 | PC | Public | 2010 | China | Clinical | 98.5 | 100 | 0 | 99.9833 | 5094375 | 45.3 | 58 | 401597 |
| VP10279 | VP10279 | SAMN20298409 | SRR15184775 | PC | Public | 2010 | China | Clinical | 98.6 | 100 | 0 | 99.9892 | 5152269 | 45.27 | 58 | 427734 |
| VP10280 | VP10280 | SAMN20298406 | SRR15184779 | NonPC | Public | 2010 | China | Clinical | 92.3 | 100 | 0.19 | 98.3695 | 5050485 | 45.25 | 53 | 471370 |
| VP10281 | VP10281 | SAMN20298404 | SRR15184781 | NonPC | Public | 2010 | China | Clinical | 91.9 | 100 | 0 | 98.4286 | 5097859 | 45.24 | 37 | 522513 |
| VP10283 | VP10283 | SAMN20298410 | SRR15184774 | PC | Public | 2010 | China | Clinical | 98.6 | 100 | 0.01 | 99.9819 | 5315961 | 45.21 | 70 | 420398 |
| VP10284 | VP10284 | SAMN20298413 | SRR15184771 | PC | Public | 2010 | China | Clinical | 97.8 | 100 | 0 | 99.9835 | 5076138 | 45.29 | 49 | 458188 |
| VP10285 | VP10285 | SAMN20298415 | SRR15184769 | NonPC | Public | 2010 | China | Clinical | 91.9 | 100 | 0 | 98.3897 | 5094118 | 45.24 | 35 | 524940 |
| VP10286 | VP10286 | SAMN20298416 | SRR15184768 | NonPC | Public | 2010 | China | Clinical | 92.3 | 100 | 0.19 | 98.3523 | 5044472 | 45.25 | 51 | 450397 |
| VP10288 | VP10288 | SAMN20298326 | SRR15185044 | NonPC | Public | 2010 | China | Clinical | 91.7 | 100 | 0 | 98.4196 | 5082255 | 45.24 | 44 | 417696 |
| VP10289 | VP10289 | SAMN20298407 | SRR15184778 | PC | Public | 2010 | China | Clinical | 98.4 | 100 | 0 | 99.9931 | 5102890 | 45.3 | 65 | 425463 |
| VP10291 | VP10291 | SAMN20298417 | SRR15184767 | NonPC | Public | 2010 | China | Clinical | 92.3 | 100 | 0.19 | 98.3438 | 5044617 | 45.25 | 51 | 450300 |
| VP10292 | VP10292 | SAMN20298422 | SRR15184761 | PC | Public | 2010 | China | Clinical | 98.6 | 100 | 0 | 99.991 | 5153374 | 45.28 | 62 | 420398 |
| VP10293 | VP10293 | SAMN20298421 | SRR15184762 | PC | Public | 2010 | China | Clinical | 98.2 | 100 | 0 | 99.9835 | 5092664 | 45.31 | 51 | 401611 |
| VP10294 | VP10294 | SAMN20298425 | SRR15184731 | PC | Public | 2010 | China | Clinical | 98.7 | 100 | 0 | 99.991 | 5151824 | 45.27 | 57 | 427734 |
| VP10296 | VP10296 | SAMN20298438 | SRR15185377 | NonPC | Public | 2010 | China | Clinical | 91.9 | 100 | 0.19 | 98.3151 | 5044680 | 45.25 | 51 | 450300 |
| VP10297 | VP10297 | SAMN20298439 | SRR15185375 | NonPC | Public | 2010 | China | Clinical | 92.3 | 100 | 0.19 | 98.3371 | 5045021 | 45.25 | 52 | 450294 |
| VP10298 | VP10298 | SAMN20298389 | SRR15184832 | PC | Public | 2010 | China | Clinical | 98.4 | 100 | 0 | 99.9843 | 5135218 | 45.3 | 62 | 440625 |
| VP10299 | VP10299 | SAMN20298388 | SRR15184834 | PC | Public | 2010 | China | Clinical | 98.4 | 100 | 0.11 | 99.9646 | 5268961 | 44.86 | 66 | 440623 |
| VP10300 | VP10300 | SAMN20298391 | SRR15184830 | PC | Public | 2010 | China | Clinical | 98.6 | 100 | 0 | 99.9803 | 5150669 | 45.27 | 56 | 442145 |
| VP10301 | VP10301 | SAMN20298392 | SRR15184794 | NonPC | Public | 2010 | China | Clinical | 92.3 | 100 | 0.19 | 98.3318 | 5030024 | 45.23 | 52 | 450299 |
| VP10302 | VP10302 | SAMN20298396 | SRR15184790 | NonPC | Public | 2010 | China | Clinical | 92.3 | 100 | 0.19 | 98.3436 | 5050048 | 45.25 | 55 | 450397 |
| VP10303 | VP10303 | SAMN20298393 | SRR15184793 | NonPC | Public | 2010 | China | Clinical | 92.3 | 100 | 0.19 | 98.33 | 5048333 | 45.25 | 51 | 450300 |
| VP10304 | VP10304 | SAMN20298397 | SRR15184789 | PC | Public | 2010 | China | Clinical | 97.3 | 100 | 0 | 99.9113 | 5142794 | 45.28 | 75 | 259891 |
| VP10305 | VP10305 | SAMN20298395 | SRR15184791 | NonPC | Public | 2010 | China | Clinical | 92.3 | 100 | 0.19 | 98.3873 | 5048078 | 45.25 | 51 | 450300 |
| VP10306 | VP10306 | SAMN20298394 | SRR15184792 | NonPC | Public | 2010 | China | Clinical | 92.3 | 100 | 0.28 | 98.3271 | 5062725 | 45.23 | 55 | 450294 |
| VP10307 | VP10307 | SAMN20298400 | SRR15184785 | NonPC | Public | 2010 | China | Clinical | 89.7 | 100 | 0.36 | 98.2679 | 5128171 | 45.03 | 46 | 404449 |
| VP10309 | VP10309 | SAMN20298402 | SRR15184783 | PC | Public | 2010 | China | Clinical | 97.6 | 100 | 0 | 99.9116 | 5155204 | 45.3 | 50 | 419488 |
| VP10310 | VP10310 | SAMN20298403 | SRR15184782 | PC | Public | 2010 | China | Clinical | 98 | 100 | 0.15 | 99.9683 | 5208804 | 44.95 | 71 | 411303 |
| VP10311 | VP10311 | SAMN20298412 | SRR15184772 | PC | Public | 2010 | China | Clinical | 98.4 | 100 | 0 | 99.9835 | 5091339 | 45.29 | 58 | 425883 |
| VP10312 | VP10312 | SAMN20298411 | SRR15184773 | PC | Public | 2010 | China | Clinical | 98.6 | 100 | 0 | 99.9787 | 5150311 | 45.27 | 59 | 420398 |
| VP10313 | VP10313 | SAMN20298420 | SRR15184763 | NonPC | Public | 2010 | China | Clinical | 92.5 | 100 | 0.05 | 98.3556 | 5034620 | 45.26 | 48 | 407493 |
| VP10314 | VP10314 | SAMN20298414 | SRR15184770 | PC | Public | 2010 | China | Clinical | 98.6 | 100 | 0 | 99.9821 | 5150053 | 45.27 | 60 | 427740 |
| VP10315 | VP10315 | SAMN20298423 | SRR15184760 | PC | Public | 2010 | China | Clinical | 98.6 | 100 | 0 | 99.9673 | 5187082 | 45.28 | 60 | 420264 |
| VP10316 | VP10316 | SAMN20298377 | SRR15184846 | PC | Public | 2010 | China | Clinical | 97.3 | 100 | 0 | 99.9307 | 5146224 | 45.23 | 90 | 196784 |
| VP10317 | VP10317 | SAMN20298419 | SRR15184764 | NonPC | Public | 2010 | China | Clinical | 92.3 | 100 | 0.19 | 98.3159 | 5044224 | 45.25 | 51 | 450300 |
| VP10319 | VP10319 | SAMN20298440 | SRR15185374 | PC | Public | 2010 | China | Clinical | 98.7 | 100 | 0 | 99.9896 | 5152175 | 45.27 | 59 | 420398 |
| VP10321 | VP10321 | SAMN20298431 | SRR15185384 | PC | Public | 2010 | China | Clinical | 98.5 | 100 | 0 | 99.9834 | 5092480 | 45.29 | 55 | 425877 |
| VP10322 | VP10322 | SAMN20298428 | SRR15184727 | PC | Public | 2010 | China | Clinical | 98.5 | 100 | 0 | 99.9735 | 5139441 | 45.25 | 55 | 425882 |
| VP10324 | VP10324 | SAMN20298434 | SRR15185381 | NonPC | Public | 2010 | China | Clinical | 91.9 | 100 | 0 | 98.3958 | 5094960 | 45.24 | 33 | 522568 |
| VP10325 | VP10325 | SAMN20298437 | SRR15185378 | NonPC | Public | 2010 | China | Clinical | 92.3 | 100 | 0.19 | 98.3687 | 5044260 | 45.25 | 51 | 450294 |
| VP10328 | VP10328 | SAMN20298426 | SRR15184729 | PC | Public | 2010 | China | Clinical | 98.5 | 100 | 0 | 99.9841 | 5093057 | 45.29 | 59 | 401597 |
| VP10330 | VP10330 | SAMN20298348 | SRR15184949 | PC | Public | 2010 | China | Clinical | 98.5 | 100 | 0 | 99.9843 | 5092786 | 45.29 | 57 | 425877 |
| VP10331 | VP10331 | SAMN20298427 | SRR15184728 | PC | Public | 2010 | China | Clinical | 98.4 | 100 | 0 | 99.9844 | 5094959 | 45.3 | 58 | 425349 |

| | | | | | | | | | | | | | | | | |
|---|---|---|---|---|---|---|---|---|---|---|---|---|---|---|---|---|
| VP10333 | VP10333 | SAMN20298375 | SRR15184848 | PC | Public | 2010 | China | Clinical | 98 | 100 | 0 | 99.9853 | 5081144 | 45.28 | 63 | 401604 |
| VP10334 | VP10334 | SAMN20298381 | SRR15184841 | NonPC | Public | 2010 | China | Clinical | 91.7 | 100 | 0 | 98.4285 | 5084892 | 45.24 | 43 | 353286 |
| VP10335 | VP10335 | SAMN20298385 | SRR15184837 | PC | Public | 2010 | China | Clinical | 98.4 | 100 | 0 | 99.9845 | 5091302 | 45.29 | 59 | 394812 |
| VP10336 | VP10336 | SAMN20298424 | SRR15184759 | NonPC | Public | 2010 | China | Clinical | 92.9 | 100 | 0 | 98.3783 | 5132818 | 45.22 | 42 | 573717 |
| VP10338 | VP10338 | SAMN20298460 | SRR15185352 | NonPC | Public | 2010 | China | Clinical | 92.9 | 100 | 0 | 98.3897 | 5133650 | 45.22 | 42 | 572297 |
| VP10339 | VP10339 | SAMN20298458 | SRR15185355 | NonPC | Public | 2010 | China | Clinical | 92.2 | 100 | 0.46 | 98.3766 | 5167517 | 45.23 | 64 | 339983 |
| VP10340 | VP10340 | SAMN20298461 | SRR15185351 | PC | Public | 2010 | China | Clinical | 98.4 | 100 | 0 | 99.9846 | 5092299 | 45.29 | 57 | 425337 |
| VP10341 | VP10341 | SAMN20298471 | SRR15185339 | PC | Public | 2010 | China | Clinical | 98.5 | 100 | 0 | 99.9926 | 5182386 | 45.21 | 61 | 425349 |
| VP10342 | VP10342 | SAMN20298468 | SRR15185344 | PC | Public | 2010 | China | Clinical | 98.6 | 100 | 0 | 99.9905 | 5148664 | 45.27 | 60 | 334357 |
| VP10343 | VP10343 | SAMN20298476 | SRR15185334 | PC | Public | 2010 | China | Clinical | 98.5 | 100 | 0 | 99.9845 | 5093338 | 45.29 | 58 | 394962 |
| VP10344 | VP10344 | SAMN20298497 | SRR15185311 | PC | Public | 2010 | China | Clinical | 98.4 | 100 | 0.5 | 99.9701 | 5142822 | 45.32 | 61 | 425235 |
| VP10345 | VP10345 | SAMN20298480 | SRR15185329 | PC | Public | 2010 | China | Clinical | 98.6 | 100 | 0 | 99.9824 | 5137137 | 45.29 | 61 | 431244 |
| VP10348 | VP10348 | SAMN20298490 | SRR15185318 | NonPC | Public | 2010 | China | Clinical | 92.3 | 100 | 0.19 | 98.3779 | 5045209 | 45.25 | 56 | 450300 |
| VP10350 | VP10350 | SAMN20298491 | SRR15185317 | PC | Public | 2010 | China | Clinical | 98.3 | 100 | 0 | 99.9924 | 5093510 | 45.31 | 55 | 334350 |
| VP10352 | VP10352 | SAMN20298492 | SRR15185316 | NonPC | Public | 2010 | China | Clinical | 92.1 | 100 | 0.05 | 98.3469 | 5034182 | 45.26 | 46 | 407493 |
| VP10353 | VP10353 | SAMN20298496 | SRR15185312 | PC | Public | 2010 | China | Clinical | 98.3 | 100 | 0.27 | 99.9921 | 5134876 | 45.31 | 52 | 394984 |
| VP10354 | VP10354 | SAMN20298498 | SRR15185310 | PC | Public | 2010 | China | Clinical | 98.5 | 100 | 0 | 99.9825 | 5093576 | 45.29 | 52 | 442205 |
| VP10355 | VP10355 | SAMN20298494 | SRR15185314 | NonPC | Public | 2010 | China | Clinical | 92.9 | 100 | 0 | 98.4074 | 5176122 | 45.26 | 39 | 488300 |
| VP10356 | VP10356 | SAMN20298495 | SRR15185313 | NonPC | Public | 2010 | China | Clinical | 92.9 | 100 | 0 | 98.395 | 5174715 | 45.26 | 42 | 488297 |
| VP10357 | VP10357 | SAMN20298507 | SRR15185300 | NonPC | Public | 2010 | China | Clinical | 92.3 | 100 | 0.19 | 98.3207 | 5043803 | 45.25 | 51 | 450300 |
| VP10358 | VP10358 | SAMN20298508 | SRR15185299 | PC | Public | 2010 | China | Clinical | 98.4 | 100 | 0 | 99.9832 | 5089911 | 45.29 | 57 | 425883 |
| VP10359 | VP10359 | SAMN20298509 | SRR15185297 | PC | Public | 2010 | China | Clinical | 98.4 | 100 | 0 | 99.985 | 5089578 | 45.29 | 55 | 401597 |
| VP10361 | VP10361 | SAMN20298515 | SRR15185291 | NonPC | Public | 2010 | China | Clinical | 92.3 | 100 | 0.19 | 98.3203 | 5010193 | 45.25 | 51 | 450300 |
| VP10362 | VP10362 | SAMN20298513 | SRR15185293 | NonPC | Public | 2010 | China | Clinical | 89.3 | 100 | 0.37 | 98.2964 | 5276741 | 44.98 | 74 | 379147 |
| VP10363 | VP10363 | SAMN20298517 | SRR15185289 | PC | Public | 2010 | China | Clinical | 98.6 | 100 | 0 | 99.9922 | 5146899 | 45.27 | 58 | 461740 |
| VP10364 | VP10364 | SAMN20298519 | SRR15185286 | PC | Public | 2010 | China | Clinical | 97.6 | 100 | 0 | 99.9255 | 5172157 | 45.1 | 77 | 288528 |
| VP10366 | VP10366 | SAMN20298520 | SRR15185285 | PC | Public | 2010 | China | Clinical | 98.4 | 100 | 0.27 | 99.9826 | 5138236 | 45.28 | 50 | 425883 |
| VP10367 | VP10367 | SAMN20298511 | SRR15185295 | PC | Public | 2010 | China | Clinical | 98.5 | 100 | 0 | 99.9931 | 5097250 | 45.29 | 59 | 401047 |
| VP10368 | VP10368 | SAMN20298512 | SRR15185294 | PC | Public | 2010 | China | Clinical | 97.3 | 100 | 0 | 99.9007 | 5150617 | 45.28 | 76 | 260119 |
| VP10369 | VP10369 | SAMN20298521 | SRR15185284 | PC | Public | 2010 | China | Clinical | 98.6 | 100 | 0 | 99.9804 | 5150476 | 45.27 | 55 | 420264 |
| VP10370 | VP10370 | SAMN20298484 | SRR15185325 | NonPC | Public | 2010 | China | Clinical | 92 | 100 | 0 | 98.3907 | 5115412 | 45.22 | 43 | 572421 |
| VP10371 | VP10371 | SAMN20298483 | SRR15185326 | NonPC | Public | 2010 | China | Clinical | 92 | 100 | 0 | 98.4025 | 5117155 | 45.22 | 43 | 488295 |
| VP10372 | VP10372 | SAMN20298485 | SRR15185324 | NonPC | Public | 2010 | China | Clinical | 92 | 100 | 0 | 98.3928 | 5117148 | 45.22 | 41 | 488295 |
| VP10373 | VP10373 | SAMN20298482 | SRR15185327 | NonPC | Public | 2010 | China | Clinical | 92 | 100 | 0 | 98.3952 | 5115678 | 45.22 | 39 | 488295 |
| VP10374 | VP10374 | SAMN20298405 | SRR15184780 | NonPC | Public | 2010 | China | Clinical | 92 | 100 | 0 | 98.3973 | 5116706 | 45.22 | 40 | 572322 |
| VP10376 | VP10376 | SAMN20298418 | SRR15184766 | PC | Public | 2010 | China | Clinical | 98.5 | 100 | 0 | 99.9856 | 5093836 | 45.29 | 61 | 312685 |
| VP10377 | VP10377 | SAMN20298453 | SRR15185360 | PC | Public | 2010 | China | Clinical | 98.5 | 100 | 0 | 99.9926 | 5092740 | 45.29 | 56 | 425877 |
| VP10379 | VP10379 | SAMN20298455 | SRR15185358 | PC | Public | 2010 | China | Clinical | 98.5 | 100 | 0 | 99.9894 | 5127190 | 45.25 | 57 | 401611 |
| VP10380 | VP10380 | SAMN20298489 | SRR15185319 | PC | Public | 2010 | China | Clinical | 98.6 | 100 | 0 | 99.9803 | 5151265 | 45.27 | 60 | 420398 |
| VP10382 | VP10382 | SAMN20298488 | SRR15185321 | PC | Public | 2010 | China | Clinical | 98.6 | 100 | 0 | 99.9819 | 5151557 | 45.27 | 60 | 425727 |
| VP10385 | VP10385 | SAMN20298530 | SRR15185274 | PC | Public | 2010 | China | Clinical | 98.5 | 100 | 0 | 99.9833 | 5092473 | 45.29 | 58 | 425877 |
| VP10388 | VP10388 | SAMN20298474 | SRR15185336 | PC | Public | 2010 | China | Clinical | 98.6 | 100 | 0 | 99.9945 | 5135618 | 45.29 | 60 | 431788 |
| VP10391 | VP10391 | SAMN20298454 | SRR15185359 | PC | Public | 2010 | China | Clinical | 98.4 | 100 | 0 | 99.9807 | 5090245 | 45.29 | 57 | 401603 |
| VP10392 | VP10392 | SAMN20298459 | SRR15185353 | NonPC | Public | 2010 | China | Clinical | 91.9 | 100 | 0 | 98.4315 | 5062351 | 45.24 | 35 | 524940 |
| VP10393 | VP10393 | SAMN20298450 | SRR15185363 | NonPC | Public | 2010 | China | Clinical | 92.4 | 100 | 0.05 | 98.3407 | 5067959 | 45.27 | 47 | 450396 |
| VP10394 | VP10394 | SAMN20298464 | SRR15185348 | NonPC | Public | 2010 | China | Clinical | 92.3 | 100 | 0.19 | 98.3182 | 5044900 | 45.25 | 52 | 450300 |
| VP10395 | VP10395 | SAMN20298441 | SRR15185373 | NonPC | Public | 2010 | China | Clinical | 90 | 100 | 0 | 98.2502 | 4989512 | 45.35 | 49 | 390563 |
| VP10396 | VP10396 | SAMN20298449 | SRR15185364 | PC | Public | 2010 | China | Clinical | 98.4 | 100 | 0 | 99.9898 | 5094237 | 45.29 | 58 | 425997 |

| | | | | | | | | | | | | | | | | |
|---|---|---|---|---|---|---|---|---|---|---|---|---|---|---|---|---|
| VP10397 | VP10397 | SAMN20298448 | SRR15185366 | PC | Public | 2010 | China | Clinical | 98.6 | 100 | 0 | 99.9802 | 5148749 | 45.27 | 60 | 420398 |
| VP10398 | VP10398 | SAMN20298451 | SRR15185362 | PC | Public | 2010 | China | Clinical | 98.3 | 100 | 0 | 99.9928 | 5093486 | 45.31 | 53 | 347631 |
| VP10399 | VP10399 | SAMN20298452 | SRR15185361 | PC | Public | 2010 | China | Clinical | 98.6 | 100 | 0 | 99.9811 | 5151114 | 45.27 | 61 | 420265 |
| VP10400 | VP10400 | SAMN20298456 | SRR15185357 | PC | Public | 2010 | China | Clinical | 97.7 | 100 | 0 | 99.9079 | 5161294 | 45.29 | 57 | 401618 |
| VP10401 | VP10401 | SAMN20298463 | SRR15185349 | PC | Public | 2010 | China | Clinical | 98.5 | 100 | 0 | 99.9635 | 5200914 | 45.27 | 62 | 394853 |
| VP10403 | VP10403 | SAMN20298466 | SRR15185346 | NonPC | Public | 2010 | China | Clinical | 92.3 | 100 | 0.25 | 98.3516 | 5045720 | 45.26 | 52 | 450294 |
| VP10405 | VP10405 | SAMN20298479 | SRR15185330 | PC | Public | 2010 | China | Clinical | 98.3 | 100 | 0 | 99.9559 | 5140698 | 45.29 | 56 | 461740 |
| VP10406 | VP10406 | SAMN20298444 | SRR15185370 | PC | Public | 2010 | China | Clinical | 98.4 | 100 | 0 | 99.983 | 5101063 | 45.29 | 58 | 440623 |
| VP10407 | VP10407 | SAMN20298446 | SRR15185368 | PC | Public | 2010 | China | Clinical | 98.4 | 100 | 0 | 99.9836 | 5091424 | 45.29 | 54 | 442187 |
| VP10408 | VP10408 | SAMN20298443 | SRR15185371 | PC | Public | 2010 | China | Clinical | 98.4 | 100 | 0 | 99.9928 | 5090677 | 45.29 | 57 | 442187 |
| VP10410 | VP10410 | SAMN20298447 | SRR15185367 | PC | Public | 2010 | China | Clinical | 98.5 | 100 | 0 | 99.9732 | 5248328 | 45.27 | 60 | 394928 |
| VP10411 | VP10411 | SAMN20298445 | SRR15185369 | PC | Public | 2010 | China | Clinical | 98.4 | 100 | 0 | 99.9828 | 5090113 | 45.29 | 55 | 440625 |
| VP10413 | VP10413 | SAMN20298465 | SRR15185347 | PC | Public | 2010 | China | Clinical | 98.6 | 100 | 0 | 99.9903 | 5110441 | 45.28 | 53 | 427332 |
| VP10414 | VP10414 | SAMN20298430 | SRR15185385 | PC | Public | 2010 | China | Clinical | 98.7 | 100 | 0 | 99.9818 | 5152271 | 45.27 | 58 | 394850 |
| VP10415 | VP10415 | SAMN20298429 | SRR15184725 | NonPC | Public | 2010 | China | Clinical | 92.3 | 100 | 0.19 | 98.3644 | 5045418 | 45.25 | 52 | 450300 |
| VP10416 | VP10416 | SAMN20298432 | SRR15185383 | PC | Public | 2010 | China | Clinical | 98.4 | 100 | 0 | 99.9843 | 5089993 | 45.29 | 57 | 401597 |
| VP10417 | VP10417 | SAMN20298433 | SRR15185382 | PC | Public | 2010 | China | Clinical | 98.7 | 100 | 0 | 99.9773 | 5175102 | 45.14 | 61 | 427452 |
| VP10418 | VP10418 | SAMN20298435 | SRR15185380 | NonPC | Public | 2010 | China | Clinical | 92.9 | 100 | 0 | 98.3991 | 5133979 | 45.22 | 42 | 498068 |
| VP10419 | VP10419 | SAMN20298436 | SRR15185379 | NonPC | Public | 2010 | China | Clinical | 92.3 | 100 | 0.19 | 98.3329 | 5043884 | 45.25 | 53 | 450300 |
| VP10420 | VP10420 | SAMN20298469 | SRR15185341 | PC | Public | 2010 | China | Clinical | 98.5 | 100 | 0 | 99.9842 | 5092912 | 45.29 | 56 | 401597 |
| VP10421 | VP10421 | SAMN20298470 | SRR15185340 | PC | Public | 2010 | China | Clinical | 98.5 | 100 | 0 | 99.9845 | 5101313 | 45.29 | 57 | 425877 |
| VP10422 | VP10422 | SAMN20298472 | SRR15185338 | PC | Public | 2010 | China | Clinical | 98.7 | 100 | 0 | 99.9893 | 5157674 | 45.28 | 64 | 420398 |
| VP10423 | VP10423 | SAMN20298477 | SRR15185333 | PC | Public | 2010 | China | Clinical | 98.4 | 100 | 0 | 99.9825 | 5089522 | 45.29 | 55 | 425292 |
| VP10424 | VP10424 | SAMN20298475 | SRR15185335 | PC | Public | 2010 | China | Clinical | 98.5 | 100 | 0 | 99.9846 | 5101288 | 45.29 | 58 | 394957 |
| VP10425 | VP10425 | SAMN20298478 | SRR15185332 | PC | Public | 2010 | China | Clinical | 98.6 | 100 | 0 | 99.9826 | 5199384 | 45.21 | 59 | 420398 |
| VP10426 | VP10426 | SAMN20298481 | SRR15185328 | NonPC | Public | 2010 | China | Clinical | 92.3 | 100 | 0.19 | 98.354 | 5044543 | 45.25 | 54 | 450397 |
| VP10427 | VP10427 | SAMN20298493 | SRR15185315 | PC | Public | 2010 | China | Clinical | 98.5 | 100 | 0 | 99.9764 | 5186775 | 45.27 | 57 | 461740 |
| VP10429 | VP10429 | SAMN20298510 | SRR15185296 | PC | Public | 2010 | China | Clinical | 98.7 | 100 | 0.09 | 99.9886 | 5140308 | 45.38 | 62 | 427344 |
| VP10431 | VP10431 | SAMN20298516 | SRR15185290 | PC | Public | 2010 | China | Clinical | 98.4 | 100 | 0 | 99.9932 | 5090599 | 45.29 | 56 | 425997 |
| VP10434 | VP10434 | SAMN20298361 | SRR15184864 | PC | Public | 2010 | China | Clinical | 97.7 | 100 | 0 | 99.9276 | 5119096 | 45.28 | 65 | 347629 |
| VP10435 | VP10435 | SAMN20298383 | SRR15184839 | PC | Public | 2010 | China | Clinical | 98.3 | 100 | 0 | 99.9834 | 5092766 | 45.31 | 52 | 401611 |
| VP10436 | VP10436 | SAMN20298532 | SRR15185272 | PC | Public | 2010 | China | Clinical | 98.4 | 100 | 0 | 99.9897 | 5099237 | 45.28 | 56 | 425882 |
| VP10437 | VP10437 | SAMN20298543 | SRR15185260 | PC | Public | 2010 | China | Clinical | 98.6 | 100 | 0 | 99.9916 | 5150477 | 45.27 | 61 | 420398 |
| VP10439 | VP10439 | SAMN20298531 | SRR15185273 | NonPC | Public | 2010 | China | Clinical | 92.3 | 100 | 0.19 | 98.3609 | 5044399 | 45.25 | 50 | 450300 |
| VP10440 | VP10440 | SAMN20298462 | SRR15185350 | PC | Public | 2010 | China | Clinical | 97.3 | 100 | 0 | 99.9157 | 5099004 | 45.28 | 84 | 267776 |
| VP10441 | VP10441 | SAMN20298529 | SRR15185275 | NonPC | Public | 2010 | China | Clinical | 90.9 | 100 | 0.34 | 98.3651 | 5035445 | 45.34 | 62 | 402003 |
| VP10442 | VP10442 | SAMN20298518 | SRR15185288 | NonPC | Public | 2010 | China | Clinical | 92.9 | 100 | 0 | 98.3846 | 5134672 | 45.22 | 41 | 498058 |
| VP10443 | VP10443 | SAMN20298553 | SRR15185249 | PC | Public | 2010 | China | Clinical | 98.4 | 100 | 0 | 99.99 | 5090745 | 45.29 | 56 | 425883 |
| VP10444 | VP10444 | SAMN20298535 | SRR15185269 | NonPC | Public | 2010 | China | Clinical | 89.3 | 100 | 0.36 | 98.2937 | 5085120 | 45.06 | 63 | 368486 |
| VP10445 | VP10445 | SAMN20298541 | SRR15185262 | PC | Public | 2010 | China | Clinical | 98.5 | 100 | 0 | 99.9798 | 5126593 | 45.25 | 57 | 401611 |
| VP10446 | VP10446 | SAMN20298536 | SRR15185268 | NonPC | Public | 2010 | China | Env | 91.3 | 100 | 0 | 98.3597 | 5095689 | 45.27 | 95 | 296596 |
| VP10447 | VP10447 | SAMN20298542 | SRR15185261 | PC | Public | 2010 | China | Clinical | 98.6 | 100 | 0 | 99.9804 | 5101476 | 45.29 | 72 | 318883 |
| VP10448 | VP10448 | SAMN20298540 | SRR15185263 | PC | Public | 2010 | China | Clinical | 98.3 | 100 | 0.27 | 99.9639 | 5182638 | 45.29 | 58 | 401463 |
| VP10449 | VP10449 | SAMN20298538 | SRR15185266 | NonPC | Public | 2010 | China | Clinical | 91.9 | 100 | 0 | 98.4111 | 5094115 | 45.24 | 36 | 522632 |
| VP10450 | VP10450 | SAMN20298523 | SRR15185282 | PC | Public | 2010 | China | Clinical | 98.6 | 100 | 0.01 | 99.9819 | 5242248 | 45.3 | 65 | 331486 |
| VP10451 | VP10451 | SAMN20298524 | SRR15185281 | PC | Public | 2010 | China | Clinical | 98.6 | 100 | 0 | 99.9896 | 5150283 | 45.27 | 57 | 427740 |
| VP10452 | VP10452 | SAMN20298527 | SRR15185278 | PC | Public | 2010 | China | Clinical | 98.4 | 100 | 0 | 99.9921 | 5089868 | 45.29 | 54 | 401597 |
| VP10453 | VP10453 | SAMN20298525 | SRR15185280 | PC | Public | 2010 | China | Clinical | 98.6 | 100 | 0 | 99.9797 | 5150298 | 45.27 | 59 | 420399 |

| | | | | | | | | | | | | | | | | |
|---|---|---|---|---|---|---|---|---|---|---|---|---|---|---|---|---|
| VP10454 | VP10454 | SAMN20298526 | SRR15185279 | PC | Public | 2010 | China | Clinical | 98.6 | 100 | 0 | 99.9898 | 5151750 | 45.27 | 63 | 420399 |
| VP10455 | VP10455 | SAMN20298522 | SRR15185283 | PC | Public | 2010 | China | Clinical | 98.5 | 100 | 0 | 99.9841 | 5130765 | 45.29 | 54 | 347487 |
| VP10456 | VP10456 | SAMN20298504 | SRR15185303 | PC | Public | 2010 | China | Clinical | 98.4 | 100 | 0 | 99.9855 | 5091310 | 45.29 | 59 | 440929 |
| VP10457 | VP10457 | SAMN20298505 | SRR15185302 | PC | Public | 2010 | China | Clinical | 98.4 | 100 | 0 | 99.9935 | 5090507 | 45.29 | 57 | 440623 |
| VP10458 | VP10458 | SAMN20298503 | SRR15185304 | PC | Public | 2010 | China | Clinical | 98.4 | 100 | 0 | 99.9836 | 5090576 | 45.29 | 58 | 394954 |
| VP10459 | VP10459 | SAMN20298506 | SRR15185301 | PC | Public | 2010 | China | Clinical | 98.4 | 100 | 0 | 99.9932 | 5089946 | 45.29 | 55 | 425871 |
| VP10460 | VP10460 | SAMN20298502 | SRR15185305 | PC | Public | 2010 | China | Clinical | 98.4 | 100 | 0 | 99.9854 | 5091232 | 45.29 | 65 | 312685 |
| VP10461 | VP10461 | SAMN20298486 | SRR15185323 | PC | Public | 2010 | China | Clinical | 98.5 | 100 | 0 | 99.9938 | 5181778 | 45.21 | 57 | 425763 |
| VP10462 | VP10462 | SAMN20298442 | SRR15185372 | PC | Public | 2010 | China | Clinical | 98.4 | 100 | 0 | 99.991 | 5145087 | 45.33 | 62 | 401597 |
| VP10463 | VP10463 | SAMN20298457 | SRR15185356 | PC | Public | 2010 | China | Clinical | 98.6 | 100 | 0 | 99.9808 | 5151813 | 45.27 | 59 | 420398 |
| VP10464 | VP10464 | SAMN20298467 | SRR15185345 | PC | Public | 2010 | China | Clinical | 94.8 | 100 | 0 | 99.9916 | 4902234 | 45.42 | 52 | 425883 |
| VP10465 | VP10465 | SAMN20298473 | SRR15185337 | PC | Public | 2010 | China | Clinical | 94.8 | 100 | 0 | 99.9807 | 4902920 | 45.42 | 52 | 425883 |
| VP10466 | VP10466 | SAMN20298533 | SRR15185271 | PC | Public | 2010 | China | Clinical | 98.4 | 100 | 0 | 99.9852 | 5090103 | 45.29 | 54 | 440625 |
| VP10467 | VP10467 | SAMN20298534 | SRR15185270 | PC | Public | 2010 | China | Clinical | 98.4 | 100 | 0 | 99.9828 | 5098506 | 45.28 | 56 | 425883 |
| VP10468 | VP10468 | SAMN20298487 | SRR15185322 | PC | Public | 2010 | China | Clinical | 98.6 | 100 | 0 | 99.9929 | 5125163 | 45.27 | 57 | 430751 |
| VP10469 | VP10469 | SAMN20298500 | SRR15185307 | PC | Public | 2010 | China | Clinical | 98.4 | 100 | 0 | 99.9927 | 5090166 | 45.29 | 59 | 425877 |
| VP10470 | VP10470 | SAMN20298499 | SRR15185308 | PC | Public | 2010 | China | Clinical | 98.5 | 100 | 0 | 99.9921 | 5092596 | 45.29 | 56 | 426188 |
| VP10471 | VP10471 | SAMN20298514 | SRR15185292 | PC | Public | 2010 | China | Clinical | 98.4 | 100 | 0 | 99.9927 | 5089931 | 45.29 | 55 | 425883 |
| VP10472 | VP10472 | SAMN20298528 | SRR15185277 | PC | Public | 2010 | China | Clinical | 98.4 | 100 | 0 | 99.9828 | 5090182 | 45.29 | 57 | 425355 |
| VP10473 | VP10473 | SAMN20298539 | SRR15185264 | NonPC | Public | 2010 | China | Clinical | 92.3 | 100 | 0.19 | 98.3489 | 5053713 | 45.25 | 53 | 450300 |
| VP10474 | VP10474 | SAMN20298545 | SRR15185258 | PC | Public | 2010 | China | Clinical | 98.6 | 100 | 0 | 99.9797 | 5150480 | 45.27 | 60 | 420399 |
| VP10475 | VP10475 | SAMN20298544 | SRR15185259 | PC | Public | 2010 | China | Clinical | 98.3 | 100 | 0 | 99.9777 | 5139652 | 45.28 | 58 | 475497 |
| VP10476 | VP10476 | SAMN20298550 | SRR15185252 | PC | Public | 2010 | China | Clinical | 98.5 | 100 | 0 | 99.9849 | 5092581 | 45.29 | 59 | 401597 |
| VP10478 | VP10478 | SAMN20298549 | SRR15185253 | PC | Public | 2010 | China | Clinical | 98.5 | 100 | 0 | 99.9935 | 5092351 | 45.29 | 59 | 425763 |
| VP10480 | VP10480 | SAMN20298546 | SRR15185257 | NonPC | Public | 2010 | China | Clinical | 92.3 | 100 | 0.19 | 98.3454 | 5039165 | 45.26 | 45 | 450300 |
| VP10481 | VP10481 | SAMN20298547 | SRR15185256 | PC | Public | 2010 | China | Clinical | 98.5 | 100 | 0 | 99.9842 | 5093538 | 45.29 | 59 | 394909 |
| VP10482 | VP10482 | SAMN20298548 | SRR15185255 | PC | Public | 2010 | China | Clinical | 98.5 | 100 | 0 | 99.9849 | 5093439 | 45.29 | 57 | 425319 |
| VP10483 | VP10483 | SAMN20298501 | SRR15185306 | NonPC | Public | 2010 | China | Env | 91.4 | 100 | 0 | 98.3141 | 5091750 | 45.31 | 64 | 462103 |
| VP10484 | VP10484 | SAMN20298537 | SRR15185267 | NonPC | Public | 2010 | China | Env | 89.7 | 100 | 0.05 | 98.3247 | 5020670 | 45.36 | 36 | 416558 |
| VP10485 | VP10485 | SAMN20298195 | SRR15184868 | NonPC | Public | 2010 | China | Env | 89.7 | 100 | 0.05 | 98.3339 | 5019981 | 45.36 | 34 | 416558 |
| VP10490 | VP10490 | SAMN20298552 | SRR15185250 | NonPC | Public | 2010 | China | Clinical | 90.9 | 100 | 0 | 98.3025 | 5131227 | 45.29 | 69 | 374822 |
| VP10491 | VP10491 | SAMN20298551 | SRR15185251 | PC | Public | 2010 | China | Clinical | 98.6 | 100 | 0.27 | 99.9847 | 5173218 | 45.29 | 60 | 440605 |
| VP11007 | VP11007 | SAMN20298576 | SRR15185223 | NonPC | Public | 2011 | China | Clinical | 92.3 | 100 | 0.19 | 98.3603 | 5049786 | 45.25 | 54 | 450397 |
| VP11015 | VP11015 | SAMN20298577 | SRR15185222 | PC | Public | 2011 | China | Clinical | 98.6 | 100 | 0 | 99.9838 | 5151235 | 45.27 | 63 | 420398 |
| VP11024 | VP11024 | SAMN20298565 | SRR15185236 | NonPC | Public | 2011 | China | Env | 91.2 | 100 | 0.27 | 98.2089 | 5284721 | 45.22 | 92 | 401496 |
| VP11025 | VP11025 | SAMN20298564 | SRR15185237 | NonPC | Public | 2011 | China | Env | 90.4 | 100 | 0.14 | 98.2615 | 5080314 | 45.25 | 67 | 314917 |
| VP11026 | VP11026 | SAMN20298556 | SRR15185246 | NonPC | Public | 2011 | China | Env | 89.5 | 100 | 0 | 98.2565 | 5051984 | 45.35 | 61 | 459334 |
| VP11042 | VP11042 | SAMN20298587 | SRR15185211 | PC | Public | 2011 | China | Clinical | 98.6 | 100 | 0 | 99.9905 | 5152832 | 45.26 | 61 | 420264 |
| VP11046 | VP11046 | SAMN20298588 | SRR15185210 | PC | Public | 2011 | China | Clinical | 98.6 | 100 | 0 | 99.9908 | 5151927 | 45.27 | 59 | 394859 |
| VP11048 | VP11048 | SAMN20298562 | SRR15185239 | PC | Public | 2011 | China | Clinical | 98.6 | 100 | 0 | 99.9804 | 5149192 | 45.27 | 61 | 420405 |
| VP11052 | VP11052 | SAMN20298590 | SRR15185207 | NonPC | Public | 2011 | China | Clinical | 89.3 | 100 | 0.36 | 98.2974 | 5111976 | 45.04 | 65 | 379902 |
| VP11053 | VP11053 | SAMN20298589 | SRR15185208 | NonPC | Public | 2011 | China | Clinical | 88 | 100 | 0 | 98.3645 | 5151193 | 45.21 | 86 | 233281 |
| VP11067 | VP11067 | SAMN20298557 | SRR15185245 | NonPC | Public | 2011 | China | Env | 88.3 | 100 | 0.37 | 98.2372 | 5162727 | 45.24 | 39 | 557217 |
| VP11073 | VP11073 | SAMN20298597 | SRR15185200 | NonPC | Public | 2011 | China | Clinical | 92.9 | 100 | 0.05 | 98.3843 | 5032851 | 45.31 | 44 | 411537 |
| VP11082 | VP11082 | SAMN20298602 | SRR15185194 | PC | Public | 2011 | China | Clinical | 98.5 | 100 | 0 | 99.9819 | 5129399 | 45.29 | 56 | 431136 |
| VP11119 | VP11119 | SAMN20298611 | SRR15185184 | NonPC | Public | 2011 | China | Clinical | 90.9 | 100 | 0 | 98.3078 | 5130512 | 45.29 | 70 | 401951 |
| VP11121 | VP11121 | SAMN20298617 | SRR15185178 | PC | Public | 2011 | China | Clinical | 97.2 | 100 | 0 | 99.9244 | 5120795 | 45.26 | 83 | 268004 |
| VP11122 | VP11122 | SAMN20298610 | SRR15185185 | PC | Public | 2011 | China | Clinical | 98.3 | 100 | 0 | 99.9767 | 5140664 | 45.28 | 56 | 461740 |

| | | | | | | | | | | | | | | | | |
|---|---|---|---|---|---|---|---|---|---|---|---|---|---|---|---|---|
| VP11123 | VP11123 | SAMN20298618 | SRR15185177 | PC | Public | 2011 | China | Clinical | 98.6 | 100 | 0 | 99.9911 | 5151595 | 45.27 | 65 | 394845 |
| VP11124 | VP11124 | SAMN20298616 | SRR15185179 | PC | Public | 2011 | China | Clinical | 98.4 | 100 | 0 | 99.9826 | 5088796 | 45.29 | 58 | 401597 |
| VP11125 | VP11125 | SAMN20298614 | SRR15185181 | PC | Public | 2011 | China | Clinical | 98 | 100 | 0 | 99.9938 | 5082352 | 45.28 | 63 | 401583 |
| VP11127 | VP11127 | SAMN20298613 | SRR15185182 | PC | Public | 2011 | China | Clinical | 98.3 | 100 | 0 | 99.9745 | 5190966 | 45.28 | 59 | 401604 |
| VP11128 | VP11128 | SAMN20298609 | SRR15185186 | PC | Public | 2011 | China | Clinical | 98.5 | 100 | 0 | 99.9842 | 5092194 | 45.29 | 55 | 425319 |
| VP11129 | VP11129 | SAMN20298604 | SRR15185192 | PC | Public | 2011 | China | Clinical | 98.4 | 100 | 0 | 99.9831 | 5087705 | 45.29 | 53 | 425876 |
| VP11130 | VP11130 | SAMN20298626 | SRR15185168 | NonPC | Public | 2011 | China | Clinical | 92.9 | 100 | 0 | 98.4169 | 5148700 | 45.2 | 38 | 472918 |
| VP11131 | VP11131 | SAMN20298623 | SRR15185171 | PC | Public | 2011 | China | Clinical | 98.6 | 100 | 0 | 99.9814 | 5151434 | 45.27 | 59 | 420398 |
| VP11132 | VP11132 | SAMN20298625 | SRR15185169 | PC | Public | 2011 | China | Clinical | 98.4 | 100 | 0 | 99.9814 | 5090036 | 45.29 | 56 | 401463 |
| VP11133 | VP11133 | SAMN20298615 | SRR15185180 | PC | Public | 2011 | China | Clinical | 98.4 | 100 | 0 | 99.9855 | 5089650 | 45.29 | 59 | 425355 |
| VP11134 | VP11134 | SAMN20298598 | SRR15185199 | NonPC | Public | 2011 | China | Clinical | 92.7 | 100 | 0 | 98.4006 | 5117496 | 45.21 | 57 | 369125 |
| VP11135 | VP11135 | SAMN20298620 | SRR15185174 | PC | Public | 2011 | China | Clinical | 98.6 | 100 | 0 | 99.989 | 5193400 | 45.16 | 61 | 420271 |
| VP11136 | VP11136 | SAMN20298601 | SRR15185195 | PC | Public | 2011 | China | Clinical | 98.3 | 100 | 0 | 99.9665 | 5143528 | 45.29 | 60 | 461740 |
| VP11137 | VP11137 | SAMN20298605 | SRR15185191 | NonPC | Public | 2011 | China | Clinical | 91.2 | 100 | 0 | 98.3537 | 5097493 | 45.3 | 46 | 407467 |
| VP11138 | VP11138 | SAMN20298607 | SRR15185189 | PC | Public | 2011 | China | Clinical | 97.3 | 100 | 0 | 99.9061 | 5178445 | 45.29 | 73 | 259987 |
| VP11139 | VP11139 | SAMN20298629 | SRR15185164 | NonPC | Public | 2011 | China | Clinical | 88.7 | 100 | 0.39 | 98.2306 | 5202464 | 45.15 | 107 | 347986 |
| VP11140 | VP11140 | SAMN20298619 | SRR15185175 | PC | Public | 2011 | China | Clinical | 98 | 100 | 0 | 99.9944 | 5081668 | 45.28 | 68 | 395266 |
| VP11142 | VP11142 | SAMN20298621 | SRR15185173 | PC | Public | 2011 | China | Clinical | 98.6 | 100 | 0 | 99.9839 | 5150210 | 45.27 | 58 | 427206 |
| VP11143 | VP11143 | SAMN20298622 | SRR15185172 | PC | Public | 2011 | China | Clinical | 98.4 | 100 | 0 | 99.9936 | 5090515 | 45.29 | 56 | 425355 |
| VP11144 | VP11144 | SAMN20298624 | SRR15185170 | PC | Public | 2011 | China | Clinical | 98.2 | 100 | 0 | 99.9902 | 5129805 | 45.26 | 81 | 267776 |
| VP11145 | VP11145 | SAMN20298628 | SRR15185166 | PC | Public | 2011 | China | Clinical | 98.6 | 100 | 0 | 99.9894 | 5153271 | 45.27 | 63 | 394851 |
| VP11146 | VP11146 | SAMN20298630 | SRR15185163 | PC | Public | 2011 | China | Clinical | 98.4 | 100 | 0 | 99.9828 | 5091038 | 45.29 | 59 | 425355 |
| VP11147 | VP11147 | SAMN20298635 | SRR15185158 | PC | Public | 2011 | China | Clinical | 98.4 | 100 | 0 | 99.9893 | 5091524 | 45.29 | 61 | 425241 |
| VP11149 | VP11149 | SAMN20298638 | SRR15185155 | NonPC | Public | 2011 | China | Clinical | 92.1 | 100 | 0.19 | 98.4107 | 5041782 | 45.28 | 66 | 313853 |
| VP11150 | VP11150 | SAMN20298639 | SRR15185153 | PC | Public | 2011 | China | Clinical | 98.5 | 100 | 0 | 99.992 | 5115956 | 45.29 | 53 | 425223 |
| VP11152 | VP11152 | SAMN20298652 | SRR15185139 | PC | Public | 2011 | China | Clinical | 98.6 | 100 | 0 | 99.9843 | 5131331 | 45.29 | 60 | 401597 |
| VP11153 | VP11153 | SAMN20298648 | SRR15185144 | PC | Public | 2011 | China | Clinical | 98.6 | 100 | 0 | 99.9856 | 5132263 | 45.29 | 60 | 401464 |
| VP11154 | VP11154 | SAMN20298651 | SRR15185140 | PC | Public | 2011 | China | Clinical | 98.7 | 100 | 0 | 99.9787 | 5151329 | 45.27 | 59 | 420398 |
| VP11155 | VP11155 | SAMN20298644 | SRR15185148 | PC | Public | 2011 | China | Clinical | 98.5 | 100 | 0 | 99.9824 | 5126649 | 45.29 | 59 | 401580 |
| VP11156 | VP11156 | SAMN20298641 | SRR15185151 | PC | Public | 2011 | China | Clinical | 98.6 | 100 | 0 | 99.9843 | 5131587 | 45.29 | 59 | 440605 |
| VP11159 | VP11159 | SAMN20298627 | SRR15185167 | PC | Public | 2011 | China | Clinical | 98.4 | 100 | 0 | 99.9828 | 5092969 | 45.29 | 59 | 401470 |
| VP11160 | VP11160 | SAMN20298636 | SRR15185157 | PC | Public | 2011 | China | Clinical | 98.6 | 100 | 0 | 99.9795 | 5150739 | 45.27 | 56 | 440277 |
| VP11161 | VP11161 | SAMN20298647 | SRR15185145 | PC | Public | 2011 | China | Clinical | 98.7 | 100 | 0 | 99.983 | 5151761 | 45.27 | 59 | 420398 |
| VP11162 | VP11162 | SAMN20298646 | SRR15185146 | NonPC | Public | 2011 | China | Clinical | 92.2 | 100 | 0.19 | 98.3593 | 5045135 | 45.25 | 52 | 450397 |
| VP11163 | VP11163 | SAMN20298632 | SRR15185161 | PC | Public | 2011 | China | Clinical | 98.5 | 100 | 0 | 99.9884 | 5127229 | 45.25 | 58 | 401611 |
| VP11165 | VP11165 | SAMN20298633 | SRR15185160 | PC | Public | 2011 | China | Clinical | 98.4 | 100 | 0 | 99.9922 | 5092289 | 45.29 | 58 | 401604 |
| VP11166 | VP11166 | SAMN20298634 | SRR15185159 | PC | Public | 2011 | China | Clinical | 98.5 | 100 | 0 | 99.9811 | 5117650 | 45.3 | 61 | 408005 |
| VP11167 | VP11167 | SAMN20298643 | SRR15185149 | PC | Public | 2011 | China | Clinical | 98.5 | 100 | 0 | 99.9894 | 5147083 | 45.27 | 56 | 542843 |
| VP11168 | VP11168 | SAMN20298650 | SRR15185141 | PC | Public | 2011 | China | Clinical | 98.6 | 100 | 0 | 99.9914 | 5151401 | 45.27 | 59 | 442453 |
| VP11169 | VP11169 | SAMN20298645 | SRR15185147 | PC | Public | 2011 | China | Clinical | 98.5 | 100 | 0 | 99.9902 | 5151257 | 45.27 | 59 | 463422 |
| VP11170 | VP11170 | SAMN20298649 | SRR15185142 | PC | Public | 2011 | China | Clinical | 98.6 | 100 | 0 | 99.9815 | 5150499 | 45.27 | 60 | 420398 |
| VP11171 | VP11171 | SAMN20298655 | SRR15185136 | PC | Public | 2011 | China | Clinical | 98.6 | 100 | 0 | 99.9824 | 5153474 | 45.27 | 60 | 420264 |
| VP11174 | VP11174 | SAMN20298656 | SRR15185135 | PC | Public | 2011 | China | Clinical | 98.6 | 100 | 0 | 99.9818 | 5150644 | 45.27 | 58 | 420398 |
| VP11175 | VP11175 | SAMN20298654 | SRR15185137 | PC | Public | 2011 | China | Clinical | 98.3 | 100 | 0 | 99.9804 | 5081971 | 45.3 | 48 | 401597 |
| VP11176 | VP11176 | SAMN20298672 | SRR15185116 | PC | Public | 2011 | China | Clinical | 97.3 | 100 | 0 | 99.9352 | 5122063 | 45.26 | 83 | 268004 |
| VP11177 | VP11177 | SAMN20298673 | SRR15185115 | PC | Public | 2011 | China | Clinical | 98.5 | 100 | 0 | 99.9926 | 5093752 | 45.29 | 57 | 401463 |
| VP11178 | VP11178 | SAMN20298670 | SRR15185118 | PC | Public | 2011 | China | Clinical | 98.6 | 100 | 0 | 99.9901 | 5151190 | 45.27 | 58 | 394853 |
| VP11179 | VP11179 | SAMN20298669 | SRR15185119 | PC | Public | 2011 | China | Clinical | 98.5 | 100 | 0 | 99.9919 | 5093388 | 45.29 | 57 | 401604 |

| | | | | | | | | | | | | | | | | |
|---|---|---|---|---|---|---|---|---|---|---|---|---|---|---|---|---|
| VP11180 | VP11180 | SAMN20298666 | SRR15185124 | PC | Public | 2011 | China | Clinical | 97.3 | 100 | 0 | 99.9269 | 5135478 | 45.13 | 71 | 269050 |
| VP11181 | VP11181 | SAMN20298665 | SRR15185125 | PC | Public | 2011 | China | Clinical | 98.5 | 100 | 0 | 99.9924 | 5093225 | 45.29 | 55 | 425313 |
| VP11182 | VP11182 | SAMN20298664 | SRR15185126 | PC | Public | 2011 | China | Clinical | 98.6 | 100 | 0 | 99.9823 | 5151532 | 45.27 | 61 | 394853 |
| VP11183 | VP11183 | SAMN20298663 | SRR15185127 | PC | Public | 2011 | China | Clinical | 98.6 | 100 | 0 | 99.9927 | 5148428 | 45.27 | 58 | 394853 |
| VP11184 | VP11184 | SAMN20298661 | SRR15185129 | PC | Public | 2011 | China | Clinical | 98.6 | 100 | 0 | 99.9862 | 5150447 | 45.27 | 61 | 394851 |
| VP11185 | VP11185 | SAMN20298676 | SRR15185112 | NonPC | Public | 2011 | China | Clinical | 92 | 100 | 0.19 | 98.322 | 5066894 | 45.27 | 55 | 372965 |
| VP11186 | VP11186 | SAMN20298667 | SRR15185123 | PC | Public | 2011 | China | Clinical | 98.4 | 100 | 0 | 99.9842 | 5090982 | 45.29 | 60 | 401597 |
| VP11187 | VP11187 | SAMN20298662 | SRR15185128 | PC | Public | 2011 | China | Clinical | 98.6 | 100 | 0 | 99.9829 | 5151318 | 45.27 | 59 | 420398 |
| VP11188 | VP11188 | SAMN20298657 | SRR15185134 | PC | Public | 2011 | China | Clinical | 98.4 | 100 | 0 | 99.9926 | 5090421 | 45.29 | 60 | 394954 |
| VP11190 | VP11190 | SAMN20298642 | SRR15185150 | PC | Public | 2011 | China | Clinical | 98.6 | 100 | 0 | 99.9893 | 5151311 | 45.27 | 56 | 440583 |
| VP11191 | VP11191 | SAMN20298695 | SRR15184569 | PC | Public | 2011 | China | Clinical | 98.5 | 100 | 0 | 99.9844 | 5093001 | 45.29 | 58 | 425337 |
| VP11192 | VP11192 | SAMN20298696 | SRR15184568 | PC | Public | 2011 | China | Clinical | 98.6 | 100 | 0 | 99.9891 | 5150192 | 45.27 | 61 | 420398 |
| VP11193 | VP11193 | SAMN20298690 | SRR15184574 | PC | Public | 2011 | China | Clinical | 97.2 | 100 | 0 | 99.9289 | 5128789 | 45.26 | 84 | 267776 |
| VP11194 | VP11194 | SAMN20298693 | SRR15184571 | PC | Public | 2011 | China | Clinical | 98.4 | 100 | 0.14 | 99.9928 | 5266066 | 45.11 | 61 | 394917 |
| VP11195 | VP11195 | SAMN20298688 | SRR15184577 | PC | Public | 2011 | China | Clinical | 98.4 | 100 | 0 | 99.9811 | 5087651 | 45.29 | 51 | 425360 |
| VP11196 | VP11196 | SAMN20298683 | SRR15185104 | PC | Public | 2011 | China | Clinical | 97.2 | 100 | 0 | 99.9354 | 5105977 | 45.27 | 79 | 268004 |
| VP11197 | VP11197 | SAMN20298677 | SRR15185111 | PC | Public | 2011 | China | Clinical | 98.6 | 100 | 0 | 99.9929 | 5132693 | 45.29 | 58 | 401597 |
| VP11198 | VP11198 | SAMN20298675 | SRR15185113 | NonPC | Public | 2011 | China | Clinical | 90.9 | 100 | 0 | 98.2758 | 5131270 | 45.29 | 65 | 374822 |
| VP11199 | VP11199 | SAMN20298679 | SRR15185108 | PC | Public | 2011 | China | Clinical | 98 | 100 | 0 | 99.9827 | 5124543 | 45.33 | 60 | 425719 |
| VP11200 | VP11200 | SAMN20298603 | SRR15185193 | PC | Public | 2011 | China | Clinical | 98.6 | 100 | 0 | 99.9798 | 5150448 | 45.27 | 60 | 420398 |
| VP11201 | VP11201 | SAMN20298606 | SRR15185190 | PC | Public | 2011 | China | Clinical | 98.4 | 100 | 0.27 | 99.9835 | 5129965 | 45.29 | 56 | 401597 |
| VP11202 | VP11202 | SAMN20298608 | SRR15185188 | PC | Public | 2011 | China | Clinical | 98.5 | 100 | 0 | 99.9913 | 5094316 | 45.29 | 57 | 401330 |
| VP11203 | VP11203 | SAMN20298612 | SRR15185183 | NonPC | Public | 2011 | China | Clinical | 92.2 | 100 | 0.19 | 98.3339 | 5112551 | 45.09 | 48 | 450299 |
| VP11204 | VP11204 | SAMN20298701 | SRR15184562 | NonPC | Public | 2011 | China | Clinical | 92.3 | 100 | 0.19 | 98.3389 | 5043976 | 45.25 | 51 | 450300 |
| VP11205 | VP11205 | SAMN20298715 | SRR15184547 | PC | Public | 2011 | China | Clinical | 98.5 | 100 | 0 | 99.9851 | 5092284 | 45.29 | 56 | 440641 |
| VP11206 | VP11206 | SAMN20298702 | SRR15184561 | NonPC | Public | 2011 | China | Clinical | 92.3 | 100 | 0.19 | 98.367 | 5044316 | 45.25 | 50 | 450397 |
| VP11207 | VP11207 | SAMN20298721 | SRR15184504 | PC | Public | 2011 | China | Clinical | 98.4 | 100 | 0 | 99.981 | 5090224 | 45.29 | 55 | 425883 |
| VP11208 | VP11208 | SAMN20298720 | SRR15184505 | PC | Public | 2011 | China | Clinical | 98.4 | 100 | 0 | 99.9909 | 5090552 | 45.29 | 61 | 401463 |
| VP11209 | VP11209 | SAMN20298705 | SRR15184558 | PC | Public | 2011 | China | Clinical | 98.4 | 100 | 0 | 99.9924 | 5090177 | 45.29 | 54 | 425235 |
| VP11210 | VP11210 | SAMN20298703 | SRR15184560 | NonPC | Public | 2011 | China | Clinical | 92.3 | 100 | 0.19 | 98.3746 | 5045894 | 45.25 | 52 | 450300 |
| VP11211 | VP11211 | SAMN20298706 | SRR15184557 | PC | Public | 2011 | China | Clinical | 98.4 | 100 | 0 | 99.9838 | 5091612 | 45.29 | 62 | 394812 |
| VP11212 | VP11212 | SAMN20298707 | SRR15184556 | PC | Public | 2011 | China | Clinical | 98.4 | 100 | 0 | 99.9826 | 5090225 | 45.29 | 55 | 401597 |
| VP11213 | VP11213 | SAMN20298704 | SRR15184559 | NonPC | Public | 2011 | China | Clinical | 92.3 | 100 | 0.19 | 98.3459 | 5043926 | 45.25 | 50 | 450300 |
| VP11214 | VP11214 | SAMN20298708 | SRR15184555 | PC | Public | 2011 | China | Clinical | 98.4 | 100 | 0 | 99.9832 | 5090196 | 45.29 | 57 | 425349 |
| VP11215 | VP11215 | SAMN20298716 | SRR15184545 | PC | Public | 2011 | China | Clinical | 98.5 | 100 | 0 | 99.9845 | 5092825 | 45.29 | 57 | 425949 |
| VP11216 | VP11216 | SAMN20298717 | SRR15184509 | PC | Public | 2011 | China | Clinical | 98.5 | 100 | 0 | 99.9835 | 5093766 | 45.29 | 59 | 394828 |
| VP11217 | VP11217 | SAMN20298709 | SRR15184553 | PC | Public | 2011 | China | Clinical | 98.4 | 100 | 0 | 99.9938 | 5091027 | 45.29 | 57 | 425349 |
| VP11218 | VP11218 | SAMN20298710 | SRR15184552 | PC | Public | 2011 | China | Clinical | 98.4 | 100 | 0 | 99.9928 | 5090659 | 45.29 | 55 | 440625 |
| VP11219 | VP11219 | SAMN20298718 | SRR15184508 | PC | Public | 2011 | China | Clinical | 98.5 | 100 | 0 | 99.9929 | 5092341 | 45.29 | 58 | 401604 |
| VP11220 | VP11220 | SAMN20298698 | SRR15184566 | NonPC | Public | 2011 | China | Clinical | 92.5 | 100 | 0.05 | 98.4214 | 5111090 | 45.24 | 31 | 451298 |
| VP11221 | VP11221 | SAMN20298719 | SRR15184506 | PC | Public | 2011 | China | Clinical | 98.5 | 100 | 0 | 99.9928 | 5093625 | 45.29 | 57 | 395699 |
| VP11222 | VP11222 | SAMN20298712 | SRR15184550 | PC | Public | 2011 | China | Clinical | 98 | 100 | 0 | 99.9837 | 5081726 | 45.28 | 59 | 401583 |
| VP11223 | VP11223 | SAMN20298713 | SRR15184549 | PC | Public | 2011 | China | Clinical | 98 | 100 | 0 | 99.9932 | 5081294 | 45.28 | 63 | 394901 |
| VP11224 | VP11224 | SAMN20298711 | SRR15184551 | PC | Public | 2011 | China | Clinical | 98.4 | 100 | 0 | 99.9822 | 5090799 | 45.29 | 56 | 425877 |
| VP11225 | VP11225 | SAMN20298714 | SRR15184548 | PC | Public | 2011 | China | Clinical | 98 | 100 | 0 | 99.9835 | 5081599 | 45.28 | 56 | 440629 |
| VP11226 | VP11226 | SAMN20298722 | SRR15184503 | PC | Public | 2011 | China | Clinical | 98.4 | 100 | 0 | 99.9842 | 5089429 | 45.29 | 57 | 394895 |
| VP11227 | VP11227 | SAMN20298700 | SRR15184563 | PC | Public | 2011 | China | Clinical | 98.5 | 100 | 0 | 99.9837 | 5130413 | 45.29 | 55 | 431844 |
| VP11228 | VP11228 | SAMN20298699 | SRR15184564 | NonPC | Public | 2011 | China | Env | 91.9 | 100 | 0 | 98.3399 | 5131227 | 45.24 | 64 | 375347 |

| | | | | | | | | | | | | | | | | |
|---|---|---|---|---|---|---|---|---|---|---|---|---|---|---|---|---|
| VP11229 | VP11229 | SAMN20298637 | SRR15185156 | PC | Public | 2011 | China | Clinical | 98 | 100 | 0 | 99.9845 | 5083569 | 45.28 | 61 | 425953 |
| VP11230 | VP11230 | SAMN20298640 | SRR15185152 | PC | Public | 2011 | China | Clinical | 98 | 100 | 0 | 99.9824 | 5099152 | 45.28 | 63 | 394963 |
| VP11231 | VP11231 | SAMN20298687 | SRR15184578 | PC | Public | 2011 | China | Clinical | 98.6 | 100 | 0 | 99.9934 | 5120128 | 45.28 | 59 | 427325 |
| VP11232 | VP11232 | SAMN20298678 | SRR15185110 | PC | Public | 2011 | China | Clinical | 98.4 | 100 | 0 | 99.9839 | 5092393 | 45.29 | 57 | 425427 |
| VP11233 | VP11233 | SAMN20298658 | SRR15185133 | PC | Public | 2011 | China | Clinical | 98.5 | 100 | 0 | 99.984 | 5093393 | 45.29 | 57 | 425427 |
| VP11235 | VP11235 | SAMN20298671 | SRR15185117 | PC | Public | 2011 | China | Clinical | 98.4 | 100 | 0 | 99.991 | 5090092 | 45.29 | 56 | 425355 |
| VP11236 | VP11236 | SAMN20298660 | SRR15185130 | PC | Public | 2011 | China | Clinical | 98.5 | 100 | 0 | 99.9802 | 5164121 | 45.25 | 61 | 408005 |
| VP11237 | VP11237 | SAMN20298682 | SRR15185105 | NonPC | Public | 2011 | China | Clinical | 92.3 | 100 | 0.19 | 98.3422 | 5022934 | 45.25 | 54 | 399425 |
| VP11238 | VP11238 | SAMN20298680 | SRR15185107 | PC | Public | 2011 | China | Clinical | 98.5 | 100 | 0 | 99.9818 | 5119199 | 45.3 | 63 | 394753 |
| VP11239 | VP11239 | SAMN20298685 | SRR15184580 | PC | Public | 2011 | China | Clinical | 98.5 | 100 | 0 | 99.9836 | 5092654 | 45.29 | 59 | 425745 |
| VP11240 | VP11240 | SAMN20298686 | SRR15184579 | PC | Public | 2011 | China | Clinical | 98.4 | 100 | 0 | 99.982 | 5090186 | 45.29 | 56 | 425883 |
| VP11241 | VP11241 | SAMN20298684 | SRR15184581 | PC | Public | 2011 | China | Clinical | 98.6 | 100 | 0 | 99.982 | 5151560 | 45.27 | 60 | 420264 |
| VP11242 | VP11242 | SAMN20298689 | SRR15184575 | NonPC | Public | 2011 | China | Clinical | 92.3 | 100 | 0.19 | 98.3552 | 5051746 | 45.25 | 52 | 539185 |
| VP11243 | VP11243 | SAMN20298653 | SRR15185138 | PC | Public | 2011 | China | Clinical | 98.6 | 100 | 0 | 99.9896 | 5146992 | 45.27 | 56 | 461740 |
| VP11244 | VP11244 | SAMN20298555 | SRR15185247 | NonPC | Public | 2011 | China | Clinical | 91.4 | 100 | 0.05 | 98.3246 | 5405734 | 45.04 | 79 | 316946 |
| VP11245 | VP11245 | SAMN20298697 | SRR15184567 | PC | Public | 2011 | China | Clinical | 98.5 | 100 | 0 | 99.9814 | 5100240 | 45.29 | 57 | 425451 |
| VP11246 | VP11246 | SAMN20298692 | SRR15184572 | PC | Public | 2011 | China | Clinical | 98.5 | 100 | 0 | 99.9826 | 5093581 | 45.29 | 55 | 425319 |
| VP11247 | VP11247 | SAMN20298742 | SRR15184481 | PC | Public | 2011 | China | Clinical | 98.6 | 100 | 0 | 99.9823 | 5151382 | 45.27 | 57 | 420398 |
| VP11248 | VP11248 | SAMN20298740 | SRR15184483 | NonPC | Public | 2011 | China | Clinical | 90.2 | 100 | 0.05 | 98.2515 | 5324720 | 45.22 | 72 | 489553 |
| VP11249 | VP11249 | SAMN20298732 | SRR15184492 | PC | Public | 2011 | China | Clinical | 98.6 | 100 | 0 | 99.9906 | 5150861 | 45.27 | 57 | 420398 |
| VP11250 | VP11250 | SAMN20298725 | SRR15184500 | PC | Public | 2011 | China | Clinical | 98.6 | 100 | 0 | 99.9925 | 5136375 | 45.29 | 60 | 431136 |
| VP11251 | VP11251 | SAMN20298737 | SRR15184487 | PC | Public | 2011 | China | Clinical | 98.6 | 100 | 0 | 99.9906 | 5151212 | 45.27 | 60 | 420405 |
| VP11252 | VP11252 | SAMN20298741 | SRR15184482 | PC | Public | 2011 | China | Clinical | 98.5 | 100 | 0 | 99.9849 | 5125879 | 45.24 | 56 | 425827 |
| VP11253 | VP11253 | SAMN20298558 | SRR15185244 | NonPC | Public | 2011 | China | Env | 89.8 | 100 | 0.27 | 98.2711 | 5000893 | 45.38 | 57 | 415192 |
| VP11254 | VP11254 | SAMN20298559 | SRR15185242 | NonPC | Public | 2011 | China | Env | 88.2 | 100 | 0.39 | 98.2205 | 5076905 | 45.17 | 84 | 433995 |
| VP11256 | VP11256 | SAMN20298560 | SRR15185241 | NonPC | Public | 2011 | China | Env | 91.1 | 100 | 0 | 98.3585 | 5007093 | 45.36 | 40 | 493842 |
| VP11257 | VP11257 | SAMN20298554 | SRR15185248 | NonPC | Public | 2011 | China | Clinical | 92.3 | 100 | 0.19 | 98.3516 | 5015267 | 45.25 | 52 | 450300 |
| VP11258 | VP11258 | SAMN20298681 | SRR15185106 | PC | Public | 2011 | China | Clinical | 98.6 | 100 | 0 | 99.9793 | 5151910 | 45.27 | 60 | 420405 |
| VP11259 | VP11259 | SAMN20298659 | SRR15185131 | PC | Public | 2011 | China | Clinical | 98.3 | 100 | 0 | 99.9759 | 5142529 | 45.29 | 57 | 401470 |
| VP11260 | VP11260 | SAMN20298668 | SRR15185122 | PC | Public | 2011 | China | Clinical | 98.6 | 100 | 0 | 99.99 | 5152427 | 45.27 | 59 | 420398 |
| VP11261 | VP11261 | SAMN20298674 | SRR15185114 | PC | Public | 2011 | China | Clinical | 98.6 | 100 | 0 | 99.982 | 5151435 | 45.27 | 57 | 427734 |
| VP11262 | VP11262 | SAMN20298561 | SRR15185240 | NonPC | Public | 2011 | China | Env | 87.4 | 100 | 0.36 | 98.2501 | 5092516 | 45.34 | 53 | 542498 |
| VP11263 | VP11263 | SAMN20298751 | SRR15185040 | PC | Public | 2011 | China | Clinical | 98.4 | 100 | 0 | 99.9816 | 5091335 | 45.29 | 59 | 401597 |
| VP11264 | VP11264 | SAMN20298746 | SRR15184477 | PC | Public | 2011 | China | Clinical | 98.4 | 100 | 0 | 99.9916 | 5090077 | 45.29 | 55 | 425871 |
| VP11265 | VP11265 | SAMN20298747 | SRR15184476 | PC | Public | 2011 | China | Clinical | 98.4 | 100 | 0 | 99.9852 | 5177370 | 45.21 | 62 | 394887 |
| VP11266 | VP11266 | SAMN20298744 | SRR15184479 | PC | Public | 2011 | China | Clinical | 98 | 100 | 0 | 99.9939 | 5082097 | 45.28 | 62 | 401583 |
| VP11267 | VP11267 | SAMN20298752 | SRR15185039 | PC | Public | 2011 | China | Clinical | 98.4 | 100 | 0 | 99.9831 | 5091158 | 45.29 | 62 | 394956 |
| VP11268 | VP11268 | SAMN20298750 | SRR15185041 | PC | Public | 2011 | China | Clinical | 98.5 | 100 | 0 | 99.9919 | 5094157 | 45.29 | 57 | 401604 |
| VP11269 | VP11269 | SAMN20298753 | SRR15185038 | PC | Public | 2011 | China | Clinical | 98.6 | 100 | 0 | 99.9818 | 5195501 | 45.31 | 61 | 440575 |
| VP11271 | VP11271 | SAMN20298755 | SRR15185036 | NonPC | Public | 2011 | China | Clinical | 90.9 | 100 | 0 | 98.3321 | 5259923 | 45.4 | 66 | 374829 |
| VP11272 | VP11272 | SAMN20298757 | SRR15185034 | PC | Public | 2011 | China | Clinical | 98.4 | 100 | 0 | 99.9918 | 5089962 | 45.29 | 58 | 401597 |
| VP11275 | VP11275 | SAMN20298743 | SRR15184480 | PC | Public | 2011 | China | Clinical | 98.6 | 100 | 0 | 99.9902 | 5150484 | 45.27 | 58 | 420405 |
| VP11276 | VP11276 | SAMN20298727 | SRR15184498 | PC | Public | 2011 | China | Clinical | 98.4 | 100 | 0 | 99.9918 | 5090964 | 45.29 | 55 | 425883 |
| VP11278 | VP11278 | SAMN20298723 | SRR15184502 | PC | Public | 2011 | China | Clinical | 98.4 | 100 | 0 | 99.9925 | 5099310 | 45.28 | 57 | 401597 |
| VP11279 | VP11279 | SAMN20298726 | SRR15184499 | NonPC | Public | 2011 | China | Clinical | 91.1 | 100 | 0 | 98.3767 | 5082088 | 45.3 | 58 | 314949 |
| VP11280 | VP11280 | SAMN20298691 | SRR15184573 | PC | Public | 2011 | China | Clinical | 98 | 100 | 0 | 99.9922 | 5081656 | 45.28 | 62 | 401583 |
| VP11281 | VP11281 | SAMN20298694 | SRR15184570 | PC | Public | 2011 | China | Clinical | 98.5 | 100 | 0 | 99.9839 | 5115845 | 45.29 | 58 | 425217 |
| VP11282 | VP11282 | SAMN20298724 | SRR15184501 | PC | Public | 2011 | China | Clinical | 98.6 | 100 | 0 | 99.9793 | 5151814 | 45.27 | 57 | 420398 |

| | | | | | | | | | | | | | | | | |
|---|---|---|---|---|---|---|---|---|---|---|---|---|---|---|---|---|
| VP11283 | VP11283 | SAMN20298730 | SRR15184494 | PC | Public | 2011 | China | Clinical | 98.4 | 100 | 0 | 99.9917 | 5087323 | 45.29 | 54 | 425348 |
| VP11284 | VP11284 | SAMN20298731 | SRR15184493 | NonPC | Public | 2011 | China | Clinical | 92.2 | 100 | 0.19 | 98.3438 | 5043265 | 45.26 | 61 | 313839 |
| VP11285 | VP11285 | SAMN20298738 | SRR15184486 | PC | Public | 2011 | China | Clinical | 98.4 | 100 | 0 | 99.9835 | 5177705 | 45.21 | 58 | 425348 |
| VP11286 | VP11286 | SAMN20298745 | SRR15184478 | PC | Public | 2011 | China | Clinical | 98.5 | 100 | 0 | 99.9843 | 5093387 | 45.29 | 58 | 401604 |
| VP11287 | VP11287 | SAMN20298748 | SRR15184475 | PC | Public | 2011 | China | Clinical | 98.4 | 100 | 0 | 99.9807 | 5091089 | 45.29 | 57 | 425469 |
| VP11288 | VP11288 | SAMN20298749 | SRR15185042 | PC | Public | 2011 | China | Clinical | 98.4 | 100 | 0 | 99.9825 | 5089975 | 45.29 | 56 | 425355 |
| VP11289 | VP11289 | SAMN20298735 | SRR15184489 | PC | Public | 2011 | China | Clinical | 98.5 | 100 | 0 | 99.9827 | 5093001 | 45.29 | 57 | 425331 |
| VP11290 | VP11290 | SAMN20298563 | SRR15185238 | PC | Public | 2011 | China | Clinical | 98.6 | 100 | 0 | 99.9818 | 5150452 | 45.27 | 60 | 427207 |
| VP11291 | VP11291 | SAMN20298739 | SRR15184484 | PC | Public | 2011 | China | Clinical | 98.4 | 100 | 0 | 99.9828 | 5089961 | 45.29 | 58 | 401597 |
| VP11292 | VP11292 | SAMN20298756 | SRR15185035 | PC | Public | 2011 | China | Clinical | 98.4 | 100 | 0 | 99.9832 | 5093023 | 45.29 | 57 | 425865 |
| VP11293 | VP11293 | SAMN20298729 | SRR15184495 | PC | Public | 2011 | China | Clinical | 98.5 | 100 | 0 | 99.9912 | 5092317 | 45.29 | 55 | 425337 |
| VP11294 | VP11294 | SAMN20298728 | SRR15184497 | PC | Public | 2011 | China | Clinical | 97.2 | 100 | 0 | 99.9834 | 5040545 | 45.28 | 62 | 425839 |
| VP11295 | VP11295 | SAMN20298733 | SRR15184491 | PC | Public | 2011 | China | Clinical | 98.6 | 100 | 0 | 99.9935 | 5151302 | 45.27 | 60 | 394851 |
| VP11296 | VP11296 | SAMN20298631 | SRR15185162 | PC | Public | 2011 | China | Clinical | 98.6 | 100 | 0 | 99.9885 | 5150125 | 45.27 | 57 | 427206 |
| VP11297 | VP11297 | SAMN20298768 | SRR15185022 | PC | Public | 2011 | China | Clinical | 98.5 | 100 | 0 | 99.9826 | 5093509 | 45.29 | 56 | 425337 |
| VP11298 | VP11298 | SAMN20298767 | SRR15185023 | NonPC | Public | 2011 | China | Clinical | 92.1 | 100 | 0.19 | 98.3828 | 4995807 | 45.25 | 61 | 313839 |
| VP11299 | VP11299 | SAMN20298765 | SRR15185025 | PC | Public | 2011 | China | Clinical | 98.5 | 100 | 0 | 99.9909 | 5092397 | 45.29 | 55 | 425223 |
| VP11301 | VP11301 | SAMN20298762 | SRR15185028 | PC | Public | 2011 | China | Clinical | 98.4 | 100 | 0 | 99.9906 | 5089906 | 45.29 | 57 | 442493 |
| VP11302 | VP11302 | SAMN20298758 | SRR15185033 | PC | Public | 2011 | China | Clinical | 98.6 | 100 | 0.7 | 99.9814 | 5252979 | 45.27 | 68 | 331620 |
| VP11303 | VP11303 | SAMN20298734 | SRR15184490 | PC | Public | 2011 | China | Clinical | 98.5 | 100 | 0 | 99.9907 | 5092642 | 45.29 | 55 | 425337 |
| VP11304 | VP11304 | SAMN20298736 | SRR15184488 | PC | Public | 2011 | China | Clinical | 98.4 | 100 | 0 | 99.9846 | 5092305 | 45.29 | 59 | 425223 |
| VP11305 | VP11305 | SAMN20298766 | SRR15185024 | PC | Public | 2011 | China | Clinical | 98.4 | 100 | 0 | 99.9929 | 5089965 | 45.29 | 58 | 440625 |
| VP11306 | VP11306 | SAMN20298761 | SRR15185029 | PC | Public | 2011 | China | Clinical | 98.3 | 100 | 0 | 99.9725 | 5146945 | 45.3 | 59 | 394684 |
| VP11307 | VP11307 | SAMN20298791 | SRR15184923 | PC | Public | 2011 | China | Clinical | 98.5 | 100 | 0 | 99.9839 | 5092921 | 45.29 | 56 | 440641 |
| VP11308 | VP11308 | SAMN20298792 | SRR15184922 | PC | Public | 2011 | China | Clinical | 98.4 | 100 | 0 | 99.9835 | 5092163 | 45.29 | 57 | 401604 |
| VP11309 | VP11309 | SAMN20298764 | SRR15185026 | NonPC | Public | 2011 | China | Clinical | 89.1 | 100 | 0.03 | 98.2837 | 4953573 | 45.32 | 44 | 382472 |
| VP11311 | VP11311 | SAMN20298803 | SRR15184910 | PC | Public | 2011 | China | Clinical | 98.4 | 100 | 0 | 99.9811 | 5089100 | 45.29 | 55 | 401597 |
| VP11313 | VP11313 | SAMN20298789 | SRR15184925 | PC | Public | 2011 | China | Clinical | 97.8 | 100 | 0 | 99.9894 | 5055921 | 45.29 | 71 | 244486 |
| VP11314 | VP11314 | SAMN20298804 | SRR15184909 | NonPC | Public | 2011 | China | Clinical | 90.9 | 100 | 0 | 98.3019 | 5130074 | 45.28 | 61 | 374815 |
| VP11315 | VP11315 | SAMN20298763 | SRR15185027 | NonPC | Public | 2011 | China | Clinical | 90.9 | 100 | 0 | 98.286 | 5134889 | 45.27 | 61 | 374822 |
| VP11316 | VP11316 | SAMN20298802 | SRR15184911 | NonPC | Public | 2011 | China | Clinical | 92.3 | 100 | 0.19 | 98.3272 | 5051023 | 45.25 | 52 | 539185 |
| VP12001 | VP12001 | SAMN20298826 | SRR15184813 | PC | Public | 2012 | China | Clinical | 98.3 | 100 | 0 | 99.9658 | 5151284 | 45.3 | 60 | 401604 |
| VP12002 | VP12002 | SAMN20298830 | SRR15184808 | NonPC | Public | 2012 | China | Clinical | 92.3 | 100 | 0.19 | 98.3715 | 5051056 | 45.25 | 56 | 313925 |
| VP12003 | VP12003 | SAMN20298829 | SRR15184809 | PC | Public | 2012 | China | Clinical | 98.4 | 100 | 0 | 99.9796 | 5093393 | 45.29 | 60 | 401464 |
| VP12004 | VP12004 | SAMN20298827 | SRR15184812 | NonPC | Public | 2012 | China | Clinical | 90.5 | 100 | 0 | 98.2982 | 5131199 | 45.29 | 65 | 333256 |
| VP12005 | VP12005 | SAMN20298828 | SRR15184811 | PC | Public | 2012 | China | Clinical | 98.5 | 100 | 0 | 99.9827 | 5092970 | 45.29 | 59 | 401597 |
| VP12006 | VP12006 | SAMN20298790 | SRR15184924 | PC | Public | 2011 | China | Clinical | 98.5 | 100 | 0 | 99.9802 | 5092486 | 45.29 | 58 | 394978 |
| VP12008 | VP12008 | SAMN20298832 | SRR15184806 | NonPC | Public | 2012 | China | Clinical | 90.7 | 100 | 0 | 98.3004 | 5116766 | 45.28 | 67 | 333176 |
| VP12009 | VP12009 | SAMN20298815 | SRR15184825 | NonPC | Public | 2012 | China | Clinical | 90.6 | 100 | 0 | 98.2653 | 5202022 | 45.26 | 48 | 415224 |
| VP12010 | VP12010 | SAMN20298831 | SRR15184807 | NonPC | Public | 2012 | China | Clinical | 88.4 | 100 | 0.05 | 98.3457 | 4934873 | 45.31 | 75 | 256676 |
| VP12011 | VP12011 | SAMN20298840 | SRR15184797 | PC | Public | 2012 | China | Clinical | 97.9 | 100 | 0 | 99.945 | 5116585 | 45.28 | 66 | 427736 |
| VP12012 | VP12012 | SAMN20298837 | SRR15184801 | NonPC | Public | 2012 | China | Clinical | 89.6 | 100 | 0 | 98.2892 | 4965651 | 45.37 | 68 | 421683 |
| VP12013 | VP12013 | SAMN20298838 | SRR15184800 | NonPC | Public | 2012 | China | Clinical | 87.8 | 100 | 0.4 | 98.2256 | 5084307 | 45.18 | 113 | 249927 |
| VP12014 | VP12014 | SAMN20298839 | SRR15184798 | NonPC | Public | 2012 | China | Clinical | 89.5 | 100 | 0.04 | 98.3542 | 4937295 | 45.39 | 36 | 730958 |
| VP12015 | VP12015 | SAMN20298835 | SRR15184803 | NonPC | Public | 2012 | China | Clinical | 90.9 | 100 | 0 | 98.2538 | 5152857 | 45.25 | 70 | 395329 |
| VP12016 | VP12016 | SAMN20298833 | SRR15184805 | PC | Public | 2012 | China | Clinical | 97.4 | 99.73 | 0 | 99.9132 | 5191984 | 45.22 | 79 | 294889 |
| VP12017 | VP12017 | SAMN20298834 | SRR15184804 | PC | Public | 2012 | China | Clinical | 97.8 | 100 | 0 | 99.8987 | 5148048 | 45.29 | 49 | 401505 |
| VP12018 | VP12018 | SAMN20298836 | SRR15184802 | NonPC | Public | 2012 | China | Env | 89.9 | 100 | 0 | 98.4301 | 4977872 | 45.36 | 72 | 315057 |

| | | | | | | | | | | | | | | | | |
|---|---|---|---|---|---|---|---|---|---|---|---|---|---|---|---|---|
| VP12019 | VP12019 | SAMN20298842 | SRR15184795 | PC | Public | 2012 | China | Clinical | 98.5 | 100 | 0 | 99.983 | 5092049 | 45.29 | 56 | 425751 |
| VP12024 | VP12024 | SAMN20298841 | SRR15184796 | NonPC | Public | 2012 | China | Clinical | 92.6 | 100 | 0 | 98.3938 | 5118441 | 45.2 | 55 | 326337 |
| VP12026 | VP12026 | SAMN20298844 | SRR15184757 | PC | Public | 2012 | China | Clinical | 98.5 | 100 | 0 | 99.9935 | 5092771 | 45.29 | 59 | 401604 |
| VP12030 | VP12030 | SAMN20298821 | SRR15184818 | PC | Public | 2012 | China | Clinical | 98.4 | 100 | 0 | 99.9851 | 5090024 | 45.29 | 59 | 401605 |
| VP12031 | VP12031 | SAMN20298843 | SRR15184758 | NonPC | Public | 2012 | China | Clinical | 92.3 | 100 | 0.05 | 98.3692 | 5009222 | 45.26 | 58 | 313971 |
| VP12032 | VP12032 | SAMN20298846 | SRR15184755 | NonPC | Public | 2012 | China | Clinical | 91.4 | 100 | 0 | 98.3463 | 5218469 | 45.27 | 55 | 504107 |
| VP12033 | VP12033 | SAMN20298848 | SRR15184753 | PC | Public | 2012 | China | Clinical | 98.4 | 100 | 0 | 99.9933 | 5084093 | 45.3 | 56 | 425355 |
| VP12034 | VP12034 | SAMN20298850 | SRR15184750 | NonPC | Public | 2012 | China | Clinical | 89.5 | 100 | 0.36 | 98.3003 | 5099151 | 45.06 | 49 | 404449 |
| VP12035 | VP12035 | SAMN20298849 | SRR15184751 | PC | Public | 2012 | China | Clinical | 98.6 | 100 | 0 | 99.9871 | 5219015 | 45.12 | 61 | 394843 |
| VP12036 | VP12036 | SAMN20298854 | SRR15184746 | PC | Public | 2012 | China | Clinical | 98 | 100 | 0 | 99.9916 | 5081077 | 45.28 | 60 | 394978 |
| VP12037 | VP12037 | SAMN20298858 | SRR15184742 | PC | Public | 2012 | China | Clinical | 98.5 | 100 | 0 | 99.9919 | 5092615 | 45.29 | 57 | 425319 |
| VP12039 | VP12039 | SAMN20298845 | SRR15184756 | PC | Public | 2012 | China | Clinical | 98.4 | 100 | 0 | 99.9819 | 5090574 | 45.29 | 58 | 425355 |
| VP12040 | VP12040 | SAMN20298852 | SRR15184748 | PC | Public | 2012 | China | Clinical | 98 | 100 | 0 | 99.9826 | 5081572 | 45.28 | 58 | 401604 |
| VP12041 | VP12041 | SAMN20298853 | SRR15184747 | PC | Public | 2012 | China | Clinical | 98.5 | 100 | 0 | 99.9845 | 5092999 | 45.29 | 57 | 401604 |
| VP12043 | VP12043 | SAMN20298819 | SRR15184820 | PC | Public | 2012 | China | Clinical | 98.1 | 100 | 0 | 99.9809 | 5070985 | 45.3 | 45 | 403715 |
| VP12044 | VP12044 | SAMN20298855 | SRR15184745 | NonPC | Public | 2012 | China | Clinical | 92.3 | 100 | 0.19 | 98.3479 | 5052390 | 45.24 | 54 | 450300 |
| VP12045 | VP12045 | SAMN20298847 | SRR15184754 | NonPC | Public | 2012 | China | Clinical | 92.9 | 100 | 0.05 | 98.374 | 5033092 | 45.31 | 41 | 420802 |
| VP12046 | VP12046 | SAMN20352344 | SRR15218566 | PC | Public | 2012 | China | Clinical | 98.4 | 100 | 0 | 99.9836 | 5090369 | 45.29 | 66 | 312692 |
| VP12047 | VP12047 | SAMN20352343 | SRR15219194 | PC | Public | 2012 | China | Clinical | 98.1 | 100 | 0 | 99.9819 | 5072394 | 45.3 | 48 | 403715 |
| VP12048 | VP12048 | SAMN20352340 | SRR15219113 | PC | Public | 2012 | China | Clinical | 98.4 | 100 | 0.14 | 99.9813 | 5128702 | 45.27 | 59 | 401604 |
| VP12049 | VP12049 | SAMN20298866 | SRR15184733 | PC | Public | 2012 | China | Clinical | 98.4 | 100 | 0 | 99.9922 | 5089293 | 45.29 | 57 | 425355 |
| VP12050 | VP12050 | SAMN20298863 | SRR15184736 | PC | Public | 2012 | China | Clinical | 98.5 | 100 | 0 | 99.9836 | 5092952 | 45.29 | 57 | 425319 |
| VP12051 | VP12051 | SAMN20298862 | SRR15184737 | PC | Public | 2012 | China | Clinical | 98.1 | 100 | 0.14 | 99.9876 | 5109553 | 45.26 | 76 | 266486 |
| VP12052 | VP12052 | SAMN20298864 | SRR15184735 | PC | Public | 2012 | China | Clinical | 98.4 | 100 | 0 | 99.9923 | 5089433 | 45.29 | 56 | 394603 |
| VP12053 | VP12053 | SAMN20298861 | SRR15184738 | PC | Public | 2012 | China | Clinical | 98.2 | 100 | 0 | 99.98 | 5077229 | 45.3 | 46 | 401463 |
| VP12054 | VP12054 | SAMN20298860 | SRR15184739 | PC | Public | 2012 | China | Clinical | 98 | 100 | 0 | 99.9916 | 5080829 | 45.28 | 59 | 401470 |
| VP12055 | VP12055 | SAMN20352342 | SRR15218940 | PC | Public | 2012 | China | Clinical | 98.4 | 100 | 0.14 | 99.9879 | 5168451 | 45.22 | 61 | 425713 |
| VP12056 | VP12056 | SAMN20352345 | SRR15218834 | PC | Public | 2012 | China | Clinical | 98.4 | 100 | 0 | 99.9818 | 5089271 | 45.29 | 56 | 425241 |
| VP12057 | VP12057 | SAMN20352348 | SRR15219041 | PC | Public | 2012 | China | Clinical | 98.4 | 100 | 0 | 99.9913 | 5090013 | 45.29 | 56 | 425241 |
| VP12059 | VP12059 | SAMN20352349 | SRR15219402 | PC | Public | 2012 | China | Clinical | 98.4 | 100 | 0 | 99.9824 | 5091903 | 45.29 | 57 | 425361 |
| VP12060 | VP12060 | SAMN20352347 | SRR15218665 | NonPC | Public | 2012 | China | Clinical | 92.5 | 100 | 0.05 | 98.3436 | 5069754 | 45.27 | 45 | 450396 |
| VP12061 | VP12061 | SAMN20298865 | SRR15184734 | NonPC | Public | 2012 | China | Clinical | 89.5 | 100 | 0.36 | 98.3153 | 5127273 | 45.05 | 48 | 404449 |
| VP12062 | VP12062 | SAMN20352341 | SRR15219112 | PC | Public | 2012 | China | Clinical | 98.4 | 100 | 0 | 99.9843 | 5091893 | 45.29 | 58 | 425319 |
| VP12063 | VP12063 | SAMN20352358 | SRR15218997 | PC | Public | 2012 | China | Clinical | 98.4 | 100 | 0 | 99.9817 | 5089450 | 45.29 | 55 | 401471 |
| VP12064 | VP12064 | SAMN20352359 | SRR15218986 | PC | Public | 2012 | China | Clinical | 98.4 | 100 | 0 | 99.991 | 5089459 | 45.29 | 54 | 425355 |
| VP12065 | VP12065 | SAMN20352356 | SRR15218495 | PC | Public | 2012 | China | Clinical | 98.4 | 100 | 0 | 99.9826 | 5090730 | 45.29 | 57 | 401472 |
| VP12066 | VP12066 | SAMN20298867 | SRR15184732 | PC | Public | 2012 | China | Clinical | 98 | 100 | 0 | 99.963 | 5120749 | 45.29 | 45 | 461740 |
| VP12067 | VP12067 | SAMN20352352 | SRR15218636 | PC | Public | 2012 | China | Clinical | 98.1 | 100 | 0 | 99.9814 | 5071879 | 45.3 | 50 | 403715 |
| VP12068 | VP12068 | SAMN20352367 | SRR15219333 | PC | Public | 2012 | China | Clinical | 98.1 | 100 | 0 | 99.9819 | 5072088 | 45.3 | 47 | 401463 |
| VP12069 | VP12069 | SAMN20352366 | SRR15219412 | PC | Public | 2012 | China | Clinical | 98.6 | 100 | 0 | 99.9896 | 5102309 | 45.29 | 61 | 401344 |
| VP12070 | VP12070 | SAMN20352362 | SRR15218917 | PC | Public | 2012 | China | Clinical | 98.4 | 100 | 0 | 99.9803 | 5091002 | 45.29 | 56 | 401463 |
| VP12071 | VP12071 | SAMN20298822 | SRR15184817 | PC | Public | 2012 | China | Clinical | 98.4 | 100 | 0 | 99.9817 | 5089498 | 45.29 | 56 | 425355 |
| VP12072 | VP12072 | SAMN20352361 | SRR15218928 | PC | Public | 2012 | China | Clinical | 98.4 | 100 | 0 | 99.9914 | 5089857 | 45.29 | 57 | 425355 |
| VP12073 | VP12073 | SAMN20298851 | SRR15184749 | NonPC | Public | 2012 | China | Clinical | 92.3 | 100 | 0.19 | 98.3692 | 5020221 | 45.24 | 50 | 450300 |
| VP12074 | VP12074 | SAMN20298857 | SRR15184743 | NonPC | Public | 2012 | China | Clinical | 91 | 100 | 0 | 98.3169 | 5131358 | 45.29 | 72 | 374836 |
| VP12075 | VP12075 | SAMN20298859 | SRR15184740 | NonPC | Public | 2012 | China | Clinical | 92.3 | 100 | 0.79 | 98.3929 | 5311213 | 45.19 | 87 | 163445 |
| VP12076 | VP12076 | SAMN20352350 | SRR15219111 | PC | Public | 2012 | China | Clinical | 98.2 | 100 | 0 | 99.9895 | 5077606 | 45.3 | 49 | 401597 |
| VP12077 | VP12077 | SAMN20352365 | SRR15219424 | PC | Public | 2012 | China | Clinical | 97.7 | 100 | 0 | 99.9022 | 5112521 | 45.29 | 56 | 419886 |

| | | | | | | | | | | | | | | | | |
|---|---|---|---|---|---|---|---|---|---|---|---|---|---|---|---|---|
| VP12078 | VP12078 | SAMN20352360 | SRR15218939 | PC | Public | 2012 | China | Clinical | 98 | 100 | 0 | 99.9933 | 5070844 | 45.28 | 78 | 266114 |
| VP12082 | VP12082 | SAMN20352353 | SRR15218625 | NonPC | Public | 2012 | China | Clinical | 92.9 | 100 | 0.01 | 98.4022 | 5224621 | 45.24 | 44 | 468830 |
| VP12083 | VP12083 | SAMN20352354 | SRR15218517 | NonPC | Public | 2012 | China | Clinical | 92.7 | 100 | 0 | 98.3935 | 5117706 | 45.21 | 53 | 468000 |
| VP12084 | VP12084 | SAMN20352374 | SRR15218765 | PC | Public | 2012 | China | Clinical | 98.5 | 100 | 0 | 99.9915 | 5092638 | 45.29 | 53 | 425241 |
| VP12085 | VP12085 | SAMN20352372 | SRR15219171 | PC | Public | 2012 | China | Clinical | 98.4 | 100 | 0 | 99.9823 | 5092425 | 45.29 | 56 | 425241 |
| VP12086 | VP12086 | SAMN20352351 | SRR15218647 | PC | Public | 2012 | China | Clinical | 98.4 | 100 | 0 | 99.9805 | 5107061 | 45.27 | 59 | 394711 |
| VP12087 | VP12087 | SAMN20352355 | SRR15218506 | NonPC | Public | 2012 | China | Clinical | 90.9 | 100 | 0 | 98.3844 | 4981343 | 45.35 | 51 | 268471 |
| VP12088 | VP12088 | SAMN20352400 | SRR15219262 | PC | Public | 2012 | China | Clinical | 98.4 | 100 | 0 | 99.9919 | 5090445 | 45.29 | 60 | 401471 |
| VP12089 | VP12089 | SAMN20352399 | SRR15219274 | PC | Public | 2012 | China | Clinical | 98.4 | 100 | 0 | 99.9851 | 5090715 | 45.29 | 57 | 425355 |
| VP12090 | VP12090 | SAMN20352402 | SRR15219170 | PC | Public | 2012 | China | Clinical | 98.4 | 100 | 0 | 99.9855 | 5091555 | 45.29 | 59 | 425241 |
| VP12091 | VP12091 | SAMN20352391 | SRR15218822 | PC | Public | 2012 | China | Clinical | 98.4 | 100 | 0 | 99.9815 | 5090882 | 45.29 | 56 | 425355 |
| VP12092 | VP12092 | SAMN20352390 | SRR15218833 | PC | Public | 2012 | China | Clinical | 98.4 | 100 | 0 | 99.9872 | 5089040 | 45.29 | 58 | 425355 |
| VP12093 | VP12093 | SAMN20352392 | SRR15218811 | PC | Public | 2012 | China | Clinical | 98.4 | 100 | 0 | 99.9846 | 5090706 | 45.29 | 57 | 425235 |
| VP12094 | VP12094 | SAMN20352388 | SRR15218962 | PC | Public | 2012 | China | Clinical | 98.4 | 100 | 0 | 99.9905 | 5090342 | 45.29 | 59 | 425769 |
| VP12095 | VP12095 | SAMN20352385 | SRR15219064 | PC | Public | 2012 | China | Clinical | 98.4 | 100 | 0 | 99.9844 | 5179851 | 45.21 | 62 | 425355 |
| VP12096 | VP12096 | SAMN20352387 | SRR15218973 | PC | Public | 2012 | China | Clinical | 98.5 | 100 | 0 | 99.9835 | 5092912 | 45.29 | 58 | 425355 |
| VP12097 | VP12097 | SAMN20352393 | SRR15218800 | PC | Public | 2012 | China | Clinical | 98.4 | 100 | 0 | 99.9915 | 5090227 | 45.29 | 57 | 442189 |
| VP12098 | VP12098 | SAMN20352386 | SRR15219053 | PC | Public | 2012 | China | Clinical | 98.4 | 100 | 0 | 99.9919 | 5179071 | 45.21 | 71 | 312662 |
| VP12099 | VP12099 | SAMN20352389 | SRR15218951 | PC | Public | 2012 | China | Clinical | 98.4 | 100 | 0 | 99.9929 | 5089041 | 45.29 | 63 | 394954 |
| VP12100 | VP12100 | SAMN20352379 | SRR15218577 | PC | Public | 2012 | China | Clinical | 98.5 | 100 | 0 | 99.9921 | 5092487 | 45.29 | 54 | 425355 |
| VP12101 | VP12101 | SAMN20352376 | SRR15218711 | PC | Public | 2012 | China | Clinical | 98.4 | 100 | 0 | 99.9828 | 5090039 | 45.29 | 60 | 401471 |
| VP12102 | VP12102 | SAMN20298868 | SRR15184730 | PC | Public | 2012 | China | Clinical | 97.6 | 100 | 0 | 99.9179 | 5152020 | 45.12 | 80 | 266114 |
| VP12103 | VP12103 | SAMN20352375 | SRR15218722 | NonPC | Public | 2012 | China | Clinical | 92.3 | 100 | 0.33 | 98.3431 | 5040871 | 45.24 | 55 | 313971 |
| VP12104 | VP12104 | SAMN20352380 | SRR15218565 | PC | Public | 2012 | China | Clinical | 98.4 | 100 | 0 | 99.9932 | 5090348 | 45.29 | 57 | 401604 |
| VP12106 | VP12106 | SAMN20352364 | SRR15219435 | NonPC | Public | 2012 | China | Clinical | 90.3 | 100 | 0.27 | 98.2716 | 5116731 | 45.3 | 46 | 269910 |
| VP12107 | VP12107 | SAMN20352377 | SRR15218700 | PC | Public | 2012 | China | Clinical | 98.4 | 100 | 0 | 99.9822 | 5090246 | 45.29 | 59 | 394812 |
| VP12108 | VP12108 | SAMN20352394 | SRR15218789 | PC | Public | 2012 | China | Clinical | 98 | 100 | 0 | 99.9902 | 5070930 | 45.27 | 78 | 265772 |
| VP12110 | VP12110 | SAMN20352401 | SRR15219251 | PC | Public | 2012 | China | Clinical | 98.4 | 100 | 0 | 99.982 | 5127946 | 45.24 | 61 | 394895 |
| VP12111 | VP12111 | SAMN20352403 | SRR15219159 | PC | Public | 2012 | China | Clinical | 98 | 100 | 0 | 99.9943 | 5068395 | 45.27 | 81 | 266114 |
| VP12112 | VP12112 | SAMN20352410 | SRR15218664 | PC | Public | 2012 | China | Clinical | 98.4 | 100 | 0 | 99.9813 | 5090599 | 45.29 | 57 | 425355 |
| VP12113 | VP12113 | SAMN20352398 | SRR15219353 | PC | Public | 2012 | China | Clinical | 98.1 | 100 | 0 | 99.9818 | 5071771 | 45.3 | 51 | 394834 |
| VP12114 | VP12114 | SAMN20352368 | SRR15219322 | NonPC | Public | 2012 | China | Clinical | 90.5 | 100 | 0 | 98.2731 | 5170610 | 45.29 | 40 | 654679 |
| VP12115 | VP12115 | SAMN20352373 | SRR15218776 | PC | Public | 2012 | China | Clinical | 98.4 | 100 | 0 | 99.9914 | 5092484 | 45.29 | 57 | 401470 |
| VP12116 | VP12116 | SAMN20352363 | SRR15219447 | PC | Public | 2012 | China | Clinical | 98.5 | 100 | 0.27 | 99.9926 | 5135330 | 45.3 | 60 | 425313 |
| VP12117 | VP12117 | SAMN20352370 | SRR15219193 | PC | Public | 2012 | China | Clinical | 98.5 | 100 | 0 | 99.9921 | 5094288 | 45.29 | 60 | 395762 |
| VP12118 | VP12118 | SAMN20352369 | SRR15219311 | NonPC | Public | 2012 | China | Clinical | 92.3 | 100 | 0.05 | 98.317 | 5111216 | 45.25 | 60 | 321585 |
| VP12119 | VP12119 | SAMN20352378 | SRR15218588 | NonPC | Public | 2012 | China | Clinical | 92.3 | 100 | 0.05 | 98.316 | 5111297 | 45.25 | 59 | 321584 |
| VP12120 | VP12120 | SAMN20352419 | SRR15219084 | PC | Public | 2012 | China | Clinical | 98.4 | 100 | 0 | 99.9841 | 5090408 | 45.29 | 59 | 425355 |
| VP12121 | VP12121 | SAMN20298856 | SRR15184744 | NonPC | Public | 2012 | China | Clinical | 91.3 | 100 | 0 | 98.3216 | 5159326 | 45.31 | 72 | 312707 |
| VP12122 | VP12122 | SAMN20352447 | SRR15218650 | PC | Public | 2012 | China | Clinical | 97.8 | 100 | 0.27 | 99.9151 | 5286944 | 45.25 | 63 | 461740 |
| VP12123 | VP12123 | SAMN20352448 | SRR15218649 | PC | Public | 2012 | China | Clinical | 97.6 | 100 | 0 | 99.9277 | 5172382 | 45.1 | 85 | 197931 |
| VP12124 | VP12124 | SAMN20352406 | SRR15218742 | PC | Public | 2012 | China | Clinical | 98.4 | 100 | 0 | 99.9922 | 5090555 | 45.29 | 62 | 394954 |
| VP12125 | VP12125 | SAMN20352407 | SRR15218731 | PC | Public | 2012 | China | Clinical | 98.4 | 100 | 0 | 99.9941 | 5177952 | 45.21 | 71 | 312677 |
| VP12126 | VP12126 | SAMN20352411 | SRR15218617 | PC | Public | 2012 | China | Clinical | 98.5 | 100 | 0 | 99.9838 | 5099309 | 45.29 | 64 | 312691 |
| VP12127 | VP12127 | SAMN20352458 | SRR15218638 | PC | Public | 2012 | China | Clinical | 98.4 | 100 | 0 | 99.9834 | 5090032 | 45.29 | 59 | 425241 |
| VP12128 | VP12128 | SAMN20352465 | SRR15218630 | PC | Public | 2012 | China | Clinical | 98.2 | 100 | 0 | 99.9892 | 5077068 | 45.3 | 50 | 401463 |
| VP12129 | VP12129 | SAMN20352442 | SRR15218655 | PC | Public | 2012 | China | Clinical | 98.4 | 100 | 0 | 99.9841 | 5092693 | 45.29 | 65 | 313348 |
| VP12130 | VP12130 | SAMN20352371 | SRR15219182 | NonPC | Public | 2012 | China | Clinical | 92.9 | 100 | 0.05 | 98.3709 | 5032798 | 45.31 | 43 | 420799 |

| | | | | | | | | | | | | | | | | |
|---|---|---|---|---|---|---|---|---|---|---|---|---|---|---|---|---|
| VP12131 | VP12131 | SAMN20352384 | SRR15219075 | PC | Public | 2012 | China | Clinical | 97.3 | 100 | 0 | 99.9034 | 5094382 | 45.28 | 74 | 259891 |
| VP12132 | VP12132 | SAMN20352395 | SRR15218778 | PC | Public | 2012 | China | Clinical | 98.5 | 100 | 0 | 99.984 | 5092419 | 45.29 | 59 | 394986 |
| VP12133 | VP12133 | SAMN20352437 | SRR15219222 | NonPC | Public | 2012 | China | Clinical | 92.3 | 100 | 0.19 | 98.3349 | 5019751 | 45.24 | 50 | 450300 |
| VP12134 | VP12134 | SAMN20352436 | SRR15219233 | NonPC | Public | 2012 | China | Clinical | 92.3 | 100 | 0.19 | 98.3724 | 5020257 | 45.24 | 49 | 450300 |
| VP12135 | VP12135 | SAMN20352438 | SRR15219211 | PC | Public | 2012 | China | Clinical | 98.4 | 100 | 0 | 99.9828 | 5089377 | 45.29 | 55 | 425355 |
| VP12136 | VP12136 | SAMN20352432 | SRR15219379 | NonPC | Public | 2012 | China | Clinical | 90.3 | 100 | 0.14 | 98.3189 | 5372178 | 44.97 | 44 | 476197 |
| VP12137 | VP12137 | SAMN20352434 | SRR15219290 | PC | Public | 2012 | China | Clinical | 98.4 | 100 | 0 | 99.9855 | 5088367 | 45.29 | 57 | 401471 |
| VP12138 | VP12138 | SAMN20352435 | SRR15219279 | PC | Public | 2012 | China | Clinical | 98.4 | 100 | 0 | 99.9928 | 5178911 | 45.21 | 62 | 425241 |
| VP12139 | VP12139 | SAMN20352426 | SRR15218869 | PC | Public | 2012 | China | Clinical | 98.4 | 100 | 0 | 99.9846 | 5089727 | 45.29 | 58 | 425355 |
| VP12140 | VP12140 | SAMN20298823 | SRR15184816 | PC | Public | 2012 | China | Clinical | 98.4 | 100 | 0 | 99.9826 | 5090135 | 45.29 | 58 | 425355 |
| VP12141 | VP12141 | SAMN20352427 | SRR15218858 | PC | Public | 2012 | China | Clinical | 98.4 | 100 | 0 | 99.9854 | 5090468 | 45.29 | 58 | 394954 |
| VP12142 | VP12142 | SAMN20352429 | SRR15218836 | PC | Public | 2012 | China | Clinical | 98.4 | 100 | 0 | 99.9922 | 5089345 | 45.29 | 57 | 425469 |
| VP12143 | VP12143 | SAMN20352431 | SRR15219390 | PC | Public | 2012 | China | Clinical | 98.4 | 100 | 0.27 | 99.9899 | 5130393 | 45.29 | 56 | 425241 |
| VP12144 | VP12144 | SAMN20352425 | SRR15218880 | NonPC | Public | 2012 | China | Clinical | 92.4 | 100 | 0.33 | 98.3238 | 5051775 | 45.24 | 46 | 478570 |
| VP12145 | VP12145 | SAMN20352424 | SRR15218891 | NonPC | Public | 2012 | China | Clinical | 90.9 | 100 | 0 | 98.2875 | 5131228 | 45.29 | 63 | 374822 |
| VP12146 | VP12146 | SAMN20352428 | SRR15218847 | PC | Public | 2012 | China | Clinical | 98.4 | 100 | 0 | 99.991 | 5088996 | 45.29 | 57 | 401605 |
| VP12147 | VP12147 | SAMN20352430 | SRR15219401 | PC | Public | 2012 | China | Clinical | 98.4 | 100 | 0 | 99.9822 | 5090286 | 45.29 | 60 | 394895 |
| VP12148 | VP12148 | SAMN20352421 | SRR15219029 | PC | Public | 2012 | China | Clinical | 98.4 | 100 | 0 | 99.9837 | 5090624 | 45.29 | 62 | 394954 |
| VP12149 | VP12149 | SAMN20352420 | SRR15219040 | PC | Public | 2012 | China | Clinical | 98.4 | 100 | 0 | 99.9841 | 5089997 | 45.29 | 56 | 425241 |
| VP12150 | VP12150 | SAMN20352422 | SRR15219018 | PC | Public | 2012 | China | Clinical | 98.4 | 100 | 0 | 99.9843 | 5089734 | 45.29 | 58 | 425319 |
| VP12151 | VP12151 | SAMN20352416 | SRR15218461 | PC | Public | 2012 | China | Clinical | 98.4 | 100 | 0 | 99.9821 | 5090414 | 45.29 | 60 | 425355 |
| VP12152 | VP12152 | SAMN20352415 | SRR15218472 | PC | Public | 2012 | China | Clinical | 98.4 | 100 | 0 | 99.9902 | 5089364 | 45.29 | 57 | 425355 |
| VP12153 | VP12153 | SAMN20352417 | SRR15219106 | PC | Public | 2012 | China | Clinical | 98.4 | 100 | 0 | 99.9907 | 5090213 | 45.29 | 62 | 313349 |
| VP12155 | VP12155 | SAMN20352440 | SRR15219110 | PC | Public | 2012 | China | Clinical | 98.4 | 100 | 0 | 99.9922 | 5090258 | 45.29 | 58 | 394810 |
| VP12156 | VP12156 | SAMN20352413 | SRR15218595 | PC | Public | 2012 | China | Clinical | 98.4 | 100 | 0 | 99.991 | 5090556 | 45.29 | 59 | 394895 |
| VP12157 | VP12157 | SAMN20352412 | SRR15218606 | PC | Public | 2012 | China | Clinical | 98.4 | 100 | 0 | 99.984 | 5090304 | 45.29 | 61 | 334347 |
| VP12158 | VP12158 | SAMN20352408 | SRR15218687 | PC | Public | 2012 | China | Clinical | 98.4 | 100 | 0 | 99.9937 | 5089404 | 45.29 | 59 | 394941 |
| VP12159 | VP12159 | SAMN20352409 | SRR15218676 | PC | Public | 2012 | China | Clinical | 98.4 | 100 | 0 | 99.9819 | 5089424 | 45.29 | 56 | 401605 |
| VP12160 | VP12160 | SAMN20352404 | SRR15219148 | PC | Public | 2012 | China | Clinical | 98.4 | 100 | 0 | 99.9891 | 5089726 | 45.29 | 56 | 425319 |
| VP12161 | VP12161 | SAMN20298824 | SRR15184815 | PC | Public | 2012 | China | Clinical | 98.4 | 100 | 0 | 99.9914 | 5096418 | 45.29 | 61 | 395118 |
| VP12162 | VP12162 | SAMN20352460 | SRR15218635 | NonPC | Public | 2012 | China | Clinical | 93.1 | 100 | 0.19 | 98.3564 | 5071145 | 45.29 | 30 | 625759 |
| VP12163 | VP12163 | SAMN20352459 | SRR15218637 | NonPC | Public | 2012 | China | Clinical | 92.9 | 100 | 0.19 | 98.3752 | 5061349 | 45.29 | 43 | 313436 |
| VP12164 | VP12164 | SAMN20352456 | SRR15218640 | PC | Public | 2012 | China | Clinical | 98.4 | 100 | 0 | 99.9814 | 5187817 | 45.2 | 63 | 401597 |
| VP12165 | VP12165 | SAMN20352455 | SRR15218641 | PC | Public | 2012 | China | Clinical | 98.4 | 100 | 0 | 99.9926 | 5090272 | 45.29 | 56 | 442189 |
| VP12166 | VP12166 | SAMN20352454 | SRR15218642 | NonPC | Public | 2012 | China | Clinical | 92.5 | 100 | 0.33 | 98.3193 | 5053214 | 45.24 | 45 | 478570 |
| VP12168 | VP12168 | SAMN20352451 | SRR15218645 | PC | Public | 2012 | China | Clinical | 98.4 | 100 | 0 | 99.9834 | 5099585 | 45.28 | 56 | 425349 |
| VP12169 | VP12169 | SAMN20352445 | SRR15218652 | PC | Public | 2012 | China | Clinical | 98.4 | 100 | 0 | 99.9847 | 5090348 | 45.29 | 60 | 401605 |
| VP12170 | VP12170 | SAMN20352443 | SRR15218654 | NonPC | Public | 2012 | China | Clinical | 92.4 | 100 | 0.33 | 98.3352 | 5053213 | 45.24 | 45 | 478404 |
| VP12171 | VP12171 | SAMN20352418 | SRR15219095 | NonPC | Public | 2012 | China | Clinical | 92.1 | 100 | 0.19 | 98.3616 | 5016073 | 45.23 | 63 | 313971 |
| VP12172 | VP12172 | SAMN20352414 | SRR15218483 | PC | Public | 2012 | China | Clinical | 98.2 | 99.73 | 0 | 99.9905 | 5080322 | 45.29 | 57 | 425391 |
| VP12173 | VP12173 | SAMN20352452 | SRR15218644 | PC | Public | 2012 | China | Clinical | 98.5 | 100 | 0 | 99.9928 | 5093448 | 45.29 | 60 | 394947 |
| VP12174 | VP12174 | SAMN20352449 | SRR15218648 | NonPC | Public | 2012 | China | Clinical | 92.7 | 100 | 0.01 | 98.3812 | 5208216 | 45.23 | 58 | 370300 |
| VP12175 | VP12175 | SAMN20352444 | SRR15218653 | PC | Public | 2012 | China | Clinical | 98.4 | 100 | 0 | 99.9826 | 5090980 | 45.29 | 55 | 425355 |
| VP12176 | VP12176 | SAMN20352439 | SRR15219134 | PC | Public | 2012 | China | Clinical | 98.4 | 100 | 0 | 99.9929 | 5089983 | 45.29 | 58 | 425241 |
| VP12177 | VP12177 | SAMN20352441 | SRR15218657 | PC | Public | 2012 | China | Clinical | 98.4 | 100 | 0 | 99.9824 | 5089786 | 45.29 | 58 | 401605 |
| VP12178 | VP12178 | SAMN20352433 | SRR15219301 | PC | Public | 2012 | China | Clinical | 98.5 | 100 | 0 | 99.9897 | 5172403 | 45.12 | 59 | 394925 |
| VP12179 | VP12179 | SAMN20352453 | SRR15218643 | PC | Public | 2012 | China | Clinical | 98.2 | 100 | 0 | 99.9892 | 5082261 | 45.29 | 55 | 425355 |
| VP12180 | VP12180 | SAMN20352423 | SRR15218902 | PC | Public | 2012 | China | Clinical | 98.4 | 100 | 0 | 99.9856 | 5090200 | 45.29 | 61 | 401605 |

| | | | | | | | | | | | | | | | | |
|---|---|---|---|---|---|---|---|---|---|---|---|---|---|---|---|---|
| VP12181 | VP12181 | SAMN20352405 | SRR15218753 | PC | Public | 2012 | China | Clinical | 98.4 | 100 | 0 | 99.9838 | 5090444 | 45.29 | 58 | 394727 |
| VP12182 | VP12182 | SAMN20352396 | SRR15219375 | PC | Public | 2012 | China | Clinical | 98.4 | 100 | 0 | 99.9923 | 5092674 | 45.29 | 60 | 425241 |
| VP12183 | VP12183 | SAMN20298825 | SRR15184814 | PC | Public | 2012 | China | Clinical | 98.4 | 100 | 0 | 99.9937 | 5089752 | 45.29 | 60 | 401605 |
| VP12184 | VP12184 | SAMN20298816 | SRR15184824 | NonPC | Public | 2012 | China | Clinical | 90.9 | 100 | 0 | 98.2848 | 5131148 | 45.29 | 63 | 375224 |
| VP12185 | VP12185 | SAMN20352383 | SRR15218532 | PC | Public | 2012 | China | Clinical | 98.5 | 100 | 0 | 99.9917 | 5092465 | 45.29 | 59 | 425355 |
| VP12186 | VP12186 | SAMN20352382 | SRR15218543 | PC | Public | 2012 | China | Clinical | 98.4 | 100 | 0 | 99.981 | 5090599 | 45.29 | 54 | 425355 |
| VP12187 | VP12187 | SAMN20352381 | SRR15218554 | NonPC | Public | 2012 | China | Clinical | 89.5 | 100 | 0.36 | 98.2924 | 5116041 | 45.05 | 45 | 745056 |
| VP12189 | VP12189 | SAMN20352357 | SRR15219008 | PC | Public | 2012 | China | Clinical | 98.4 | 100 | 0.09 | 99.9675 | 5193282 | 45.04 | 61 | 425355 |
| VP12190 | VP12190 | SAMN20352346 | SRR15219263 | PC | Public | 2012 | China | Clinical | 98.1 | 100 | 0 | 99.9879 | 5080125 | 45.3 | 48 | 403721 |
| VP12191 | VP12191 | SAMN20352470 | SRR15218624 | PC | Public | 2012 | China | Clinical | 98.4 | 100 | 0 | 99.9918 | 5096438 | 45.29 | 57 | 401605 |
| VP12192 | VP12192 | SAMN20352468 | SRR15218627 | PC | Public | 2012 | China | Clinical | 98.4 | 100 | 0.27 | 99.9827 | 5131170 | 45.29 | 57 | 425469 |
| VP12193 | VP12193 | SAMN20352462 | SRR15218633 | PC | Public | 2012 | China | Clinical | 98.4 | 100 | 0 | 99.9837 | 5090070 | 45.29 | 58 | 401597 |
| VP12194 | VP12194 | SAMN20352463 | SRR15218632 | PC | Public | 2012 | China | Clinical | 98.4 | 100 | 0 | 99.9844 | 5089558 | 45.29 | 59 | 312826 |
| VP12195 | VP12195 | SAMN20352461 | SRR15218634 | NonPC | Public | 2012 | China | Clinical | 92.1 | 100 | 0.46 | 98.3726 | 5109425 | 45.24 | 72 | 322854 |
| VP12196 | VP12196 | SAMN20352473 | SRR15218621 | NonPC | Public | 2012 | China | Clinical | 92.9 | 100 | 0.05 | 98.3917 | 5027539 | 45.31 | 43 | 420802 |
| VP12198 | VP12198 | SAMN20352471 | SRR15218623 | PC | Public | 2012 | China | Clinical | 98.4 | 100 | 0 | 99.9852 | 5088524 | 45.29 | 58 | 425355 |
| VP12199 | VP12199 | SAMN20352469 | SRR15218626 | NonPC | Public | 2012 | China | Clinical | 92.5 | 100 | 0.05 | 98.3328 | 5126406 | 45.25 | 49 | 450396 |
| VP12200 | VP12200 | SAMN20352467 | SRR15218628 | PC | Public | 2012 | China | Clinical | 98.4 | 100 | 0 | 99.9843 | 5090060 | 45.29 | 58 | 425469 |
| VP12201 | VP12201 | SAMN20352466 | SRR15218629 | NonPC | Public | 2012 | China | Clinical | 88.5 | 100 | 0.19 | 98.3275 | 4966763 | 45.27 | 56 | 313276 |
| VP12204 | VP12204 | SAMN20352446 | SRR15218651 | PC | Public | 2012 | China | Clinical | 98.4 | 100 | 0 | 99.9811 | 5089655 | 45.29 | 58 | 425242 |
| VP12205 | VP12205 | SAMN20352397 | SRR15219364 | PC | Public | 2012 | China | Clinical | 98.7 | 100 | 0 | 99.9813 | 5103068 | 45.29 | 59 | 427338 |
| VP12206 | VP12206 | SAMN20352481 | SRR15218515 | PC | Public | 2012 | China | Clinical | 98.6 | 100 | 0 | 99.9841 | 5136742 | 45.29 | 60 | 347568 |
| VP12207 | VP12207 | SAMN20352493 | SRR15218502 | PC | Public | 2012 | China | Clinical | 98.4 | 100 | 0 | 99.9899 | 5090331 | 45.29 | 59 | 334348 |
| VP12208 | VP12208 | SAMN20352482 | SRR15218514 | NonPC | Public | 2012 | China | Clinical | 90.9 | 100 | 0.5 | 98.2581 | 5182774 | 45.16 | 70 | 333220 |
| VP12209 | VP12209 | SAMN20352490 | SRR15218505 | NonPC | Public | 2012 | China | Clinical | 92.1 | 100 | 0.46 | 98.3608 | 5126149 | 45.22 | 74 | 323098 |
| VP12210 | VP12210 | SAMN20352489 | SRR15218507 | PC | Public | 2012 | China | Clinical | 98.4 | 100 | 0 | 99.9932 | 5090152 | 45.29 | 61 | 425997 |
| VP12211 | VP12211 | SAMN20352450 | SRR15218646 | NonPC | Public | 2012 | China | Clinical | 92.3 | 100 | 0.46 | 98.3393 | 5145024 | 45.23 | 61 | 465676 |
| VP12213 | VP12213 | SAMN20352457 | SRR15218639 | PC | Public | 2012 | China | Clinical | 98.5 | 100 | 0 | 99.9934 | 5094343 | 45.29 | 60 | 313482 |
| VP12215 | VP12215 | SAMN20352477 | SRR15218520 | NonPC | Public | 2012 | China | Clinical | 92.4 | 100 | 0.33 | 98.3492 | 5052345 | 45.24 | 45 | 478570 |
| VP12216 | VP12216 | SAMN20352494 | SRR15218501 | PC | Public | 2012 | China | Clinical | 98.3 | 100 | 0 | 99.9822 | 5081368 | 45.3 | 46 | 403733 |
| VP12217 | VP12217 | SAMN20352488 | SRR15218508 | NonPC | Public | 2012 | China | Clinical | 93.1 | 100 | 0.05 | 98.3745 | 5064341 | 45.31 | 42 | 513728 |
| VP12218 | VP12218 | SAMN20352486 | SRR15218510 | PC | Public | 2012 | China | Clinical | 98.4 | 100 | 0 | 99.9844 | 5091295 | 45.29 | 60 | 394812 |
| VP12219 | VP12219 | SAMN20352485 | SRR15218511 | NonPC | Public | 2012 | China | Clinical | 92.5 | 100 | 0.33 | 98.3428 | 5084387 | 45.24 | 48 | 478570 |
| VP12220 | VP12220 | SAMN20352479 | SRR15218518 | PC | Public | 2012 | China | Clinical | 98.4 | 100 | 0 | 99.9828 | 5090702 | 45.29 | 66 | 312944 |
| VP12221 | VP12221 | SAMN20352478 | SRR15218519 | PC | Public | 2012 | China | Clinical | 98 | 100 | 0 | 99.9835 | 5080968 | 45.28 | 60 | 401470 |
| VP12222 | VP12222 | SAMN20352474 | SRR15218524 | NonPC | Public | 2012 | China | Clinical | 92.4 | 100 | 0.05 | 98.3034 | 5067343 | 45.27 | 45 | 450396 |
| VP12223 | VP12223 | SAMN20352476 | SRR15218521 | PC | Public | 2012 | China | Clinical | 98.4 | 100 | 0 | 99.9905 | 5090012 | 45.29 | 56 | 425241 |
| VP12224 | VP12224 | SAMN20352475 | SRR15218522 | PC | Public | 2012 | China | Clinical | 98.4 | 100 | 0 | 99.9834 | 5090325 | 45.29 | 60 | 394727 |
| VP12225 | VP12225 | SAMN20352464 | SRR15218631 | PC | Public | 2012 | China | Clinical | 98 | 100 | 0 | 99.9839 | 5080448 | 45.28 | 60 | 394917 |
| VP12226 | VP12226 | SAMN20352472 | SRR15218622 | PC | Public | 2012 | China | Clinical | 98.4 | 100 | 0 | 99.9904 | 5091076 | 45.29 | 58 | 425241 |
| VP12227 | VP12227 | SAMN20352487 | SRR15218509 | PC | Public | 2012 | China | Clinical | 98 | 100 | 0 | 99.9912 | 5079893 | 45.28 | 56 | 401604 |
| VP12228 | VP12228 | SAMN20352484 | SRR15218512 | PC | Public | 2012 | China | Clinical | 98.4 | 100 | 0 | 99.9922 | 5092750 | 45.29 | 57 | 425320 |
| VP12229 | VP12229 | SAMN20352480 | SRR15218516 | PC | Public | 2012 | China | Clinical | 98.4 | 100 | 0 | 99.9916 | 5089221 | 45.29 | 57 | 425355 |
| VP12230 | VP12230 | SAMN20298818 | SRR15184822 | PC | Public | 2012 | China | Clinical | 98.4 | 100 | 0 | 99.9825 | 5091946 | 45.29 | 56 | 425845 |
| VP12232 | VP12232 | SAMN20352497 | SRR15218498 | NonPC | Public | 2012 | China | Clinical | 92.9 | 100 | 0.05 | 98.4151 | 5105926 | 45.14 | 44 | 410439 |
| VP12233 | VP12233 | SAMN20352498 | SRR15218497 | NonPC | Public | 2012 | China | Clinical | 92.9 | 100 | 0.05 | 98.403 | 5111772 | 45.14 | 43 | 410439 |
| VP12234 | VP12234 | SAMN20298820 | SRR15184819 | PC | Public | 2012 | China | Env | 98.4 | 100 | 0 | 99.9848 | 5089262 | 45.29 | 57 | 425355 |
| VP12235 | VP12235 | SAMN20298817 | SRR15184823 | PC | Public | 2012 | China | Env | 98.1 | 100 | 0 | 99.9932 | 5071160 | 45.28 | 76 | 288535 |

| | | | | | | | | | | | | | | | | |
|---|---|---|---|---|---|---|---|---|---|---|---|---|---|---|---|---|
| VP12236 | VP12236 | SAMN20352483 | SRR15218513 | PC | Public | 2012 | China | Clinical | 97.9 | 100 | 0 | 99.9736 | 5120953 | 45.26 | 76 | 288527 |
| VP12237 | VP12237 | SAMN20352501 | SRR15218493 | NonPC | Public | 2012 | China | Clinical | 93.1 | 100 | 0.05 | 98.3598 | 5045756 | 45.31 | 29 | 515916 |
| VP12238 | VP12238 | SAMN20352502 | SRR15218492 | PC | Public | 2012 | China | Clinical | 98.4 | 100 | 0 | 99.9802 | 5093569 | 45.28 | 55 | 425235 |
| VP13001 | VP13001 | SAMN20352495 | SRR15218500 | NonPC | Public | 2012 | China | Clinical | 93.1 | 100 | 0.05 | 98.3336 | 5041853 | 45.32 | 31 | 484515 |
| VP13002 | VP13002 | SAMN20352500 | SRR15218494 | NonPC | Public | 2012 | China | Clinical | 93.1 | 100 | 0.05 | 98.355 | 5040328 | 45.32 | 32 | 485699 |
| VP13003 | VP13003 | SAMN20352499 | SRR15218496 | NonPC | Public | 2012 | China | Clinical | 90.5 | 100 | 0 | 98.2682 | 5169931 | 45.29 | 37 | 654544 |
| VP13004 | VP13004 | SAMN20352513 | SRR15219004 | PC | Public | 2013 | China | Clinical | 98.4 | 100 | 0 | 99.9842 | 5090186 | 45.29 | 55 | 401605 |
| VP13005 | VP13005 | SAMN20352491 | SRR15218504 | NonPC | Public | 2012 | China | Clinical | 93.1 | 100 | 0.05 | 98.3853 | 5058236 | 45.32 | 44 | 463552 |
| VP13006 | VP13006 | SAMN20352492 | SRR15218503 | PC | Public | 2012 | China | Clinical | 98.4 | 100 | 0 | 99.982 | 5090044 | 45.29 | 57 | 401463 |
| VP13007 | VP13007 | SAMN20352496 | SRR15218499 | NonPC | Public | 2012 | China | Clinical | 93.1 | 100 | 0.05 | 98.374 | 5066086 | 45.31 | 44 | 463552 |
| VP13008 | VP13008 | SAMN20352505 | SRR15218489 | PC | Public | 2013 | China | Clinical | 97.4 | 100 | 0 | 99.9217 | 5081141 | 45.28 | 78 | 205045 |
| VP13009 | VP13009 | SAMN20352514 | SRR15219003 | NonPC | Public | 2013 | China | Clinical | 90.9 | 100 | 0.14 | 98.3074 | 5142222 | 45.27 | 60 | 407422 |
| VP13010 | VP13010 | SAMN20352518 | SRR15218999 | NonPC | Public | 2013 | China | Clinical | 88.9 | 100 | 0.05 | 98.3083 | 4998411 | 45.23 | 55 | 313573 |
| VP13011 | VP13011 | SAMN20352515 | SRR15219002 | NonPC | Public | 2013 | China | Clinical | 89.3 | 100 | 0.36 | 98.2879 | 5103833 | 45.04 | 63 | 376016 |
| VP13012 | VP13012 | SAMN20352519 | SRR15218998 | PC | Public | 2013 | China | Clinical | 98.4 | 100 | 0 | 99.9921 | 5134061 | 45.34 | 61 | 401604 |
| VP13013 | VP13013 | SAMN20352522 | SRR15218994 | NonPC | Public | 2013 | China | Clinical | 92.3 | 100 | 0.05 | 98.3213 | 5009478 | 45.26 | 55 | 406810 |
| VP13014 | VP13014 | SAMN20352523 | SRR15218993 | PC | Public | 2013 | China | Clinical | 98.5 | 100 | 0 | 99.9836 | 5134515 | 45.34 | 59 | 425199 |
| VP13015 | VP13015 | SAMN20352538 | SRR15218976 | NonPC | Public | 2013 | China | Clinical | 89.6 | 100 | 0.07 | 98.2898 | 5104529 | 45.2 | 73 | 314897 |
| VP13016 | VP13016 | SAMN20352540 | SRR15218938 | PC | Public | 2013 | China | Clinical | 98.1 | 100 | 0 | 99.9897 | 5073092 | 45.3 | 46 | 403721 |
| VP13018 | VP13018 | SAMN20352520 | SRR15218996 | NonPC | Public | 2013 | China | Clinical | 92.1 | 100 | 0.19 | 98.357 | 5016412 | 45.23 | 61 | 313971 |
| VP13019 | VP13019 | SAMN20352555 | SRR15218922 | NonPC | Public | 2013 | China | Clinical | 91 | 100 | 0.05 | 98.2402 | 5171805 | 45.16 | 89 | 328919 |
| VP13020 | VP13020 | SAMN20352548 | SRR15218930 | NonPC | Public | 2013 | China | Clinical | 92.2 | 100 | 0.19 | 98.3707 | 5046843 | 45.25 | 54 | 450300 |
| VP13022 | VP13022 | SAMN20352553 | SRR15218924 | PC | Public | 2013 | China | Clinical | 98.5 | 100 | 0 | 99.9911 | 5091994 | 45.29 | 55 | 425355 |
| VP13023 | VP13023 | SAMN20352556 | SRR15218921 | PC | Public | 2013 | China | Clinical | 98.4 | 100 | 0 | 99.9833 | 5090768 | 45.29 | 55 | 425199 |
| VP13024 | VP13024 | SAMN20352557 | SRR15218920 | PC | Public | 2013 | China | Clinical | 98.4 | 100 | 0 | 99.9842 | 5091564 | 45.29 | 58 | 401470 |
| VP13026 | VP13026 | SAMN20352510 | SRR15219007 | PC | Public | 2013 | China | Clinical | 98.4 | 100 | 0.09 | 99.9811 | 5130028 | 45.38 | 54 | 425199 |
| VP13027 | VP13027 | SAMN20352561 | SRR15218915 | PC | Public | 2013 | China | Clinical | 98.5 | 100 | 0 | 99.9849 | 5092260 | 45.29 | 58 | 425841 |
| VP13028 | VP13028 | SAMN20352562 | SRR15218914 | PC | Public | 2013 | China | Clinical | 98.4 | 100 | 0 | 99.9818 | 5090066 | 45.29 | 56 | 425355 |
| VP13029 | VP13029 | SAMN20352563 | SRR15218913 | PC | Public | 2013 | China | Clinical | 98.4 | 100 | 0 | 99.9831 | 5176626 | 45.22 | 64 | 425313 |
| VP13030 | VP13030 | SAMN20352564 | SRR15218912 | PC | Public | 2013 | China | Clinical | 98.1 | 100 | 0 | 99.9933 | 5071301 | 45.28 | 78 | 266114 |
| VP13031 | VP13031 | SAMN20352569 | SRR15218907 | PC | Public | 2013 | China | Clinical | 98.1 | 100 | 0 | 99.977 | 5104619 | 45.31 | 45 | 435887 |
| VP13032 | VP13032 | SAMN20352568 | SRR15218908 | PC | Public | 2013 | China | Clinical | 98.4 | 100 | 0 | 99.9919 | 5091438 | 45.29 | 58 | 401470 |
| VP13033 | VP13033 | SAMN20352559 | SRR15218918 | PC | Public | 2013 | China | Clinical | 98.6 | 100 | 0 | 99.9779 | 5223686 | 45.11 | 58 | 427211 |
| VP13034 | VP13034 | SAMN20352554 | SRR15218923 | PC | Public | 2013 | China | Clinical | 98.4 | 100 | 0.27 | 99.9892 | 5132472 | 45.3 | 59 | 425241 |
| VP13035 | VP13035 | SAMN20352589 | SRR15219425 | PC | Public | 2013 | China | Clinical | 98.2 | 100 | 0 | 99.9793 | 5076447 | 45.3 | 47 | 403949 |
| VP13036 | VP13036 | SAMN20352566 | SRR15218910 | PC | Public | 2013 | China | Clinical | 98.4 | 100 | 0 | 99.9916 | 5091704 | 45.29 | 57 | 425205 |
| VP13038 | VP13038 | SAMN20352560 | SRR15218916 | PC | Public | 2013 | China | Clinical | 94.6 | 100 | 0 | 99.984 | 4895803 | 45.42 | 54 | 425199 |
| VP13039 | VP13039 | SAMN20352558 | SRR15218919 | PC | Public | 2013 | China | Clinical | 98.5 | 100 | 0 | 99.9822 | 5093454 | 45.29 | 53 | 425308 |
| VP13040 | VP13040 | SAMN20352552 | SRR15218925 | NonPC | Public | 2013 | China | Clinical | 92.5 | 100 | 0.05 | 98.3438 | 5087138 | 45.26 | 44 | 450396 |
| VP13042 | VP13042 | SAMN20352592 | SRR15219421 | PC | Public | 2013 | China | Clinical | 98.2 | 100 | 0 | 99.9871 | 5076765 | 45.3 | 48 | 403949 |
| VP13044 | VP13044 | SAMN20352565 | SRR15218911 | PC | Public | 2013 | China | Clinical | 98.5 | 100 | 0 | 99.9916 | 5093015 | 45.29 | 56 | 442195 |
| VP13049 | VP13049 | SAMN20352583 | SRR15219431 | PC | Public | 2013 | China | Clinical | 98.5 | 100 | 0 | 99.9849 | 5092088 | 45.29 | 61 | 401471 |
| VP13052 | VP13052 | SAMN20352588 | SRR15219426 | PC | Public | 2013 | China | Clinical | 98.2 | 100 | 0 | 99.9865 | 5076540 | 45.3 | 49 | 401597 |
| VP13054 | VP13054 | SAMN20352584 | SRR15219430 | PC | Public | 2013 | China | Clinical | 98.2 | 100 | 0 | 99.987 | 5076820 | 45.3 | 47 | 403949 |
| VP13055 | VP13055 | SAMN20352585 | SRR15219429 | PC | Public | 2013 | China | Clinical | 98.2 | 100 | 0 | 99.9874 | 5077891 | 45.3 | 49 | 403949 |
| VP13056 | VP13056 | SAMN20352586 | SRR15219428 | PC | Public | 2013 | China | Clinical | 98.2 | 100 | 0 | 99.9866 | 5077792 | 45.3 | 48 | 403949 |
| VP13057 | VP13057 | SAMN20352587 | SRR15219427 | PC | Public | 2013 | China | Clinical | 98.2 | 100 | 0 | 99.9781 | 5076729 | 45.3 | 46 | 403950 |
| VP13058 | VP13058 | SAMN20352567 | SRR15218909 | NonPC | Public | 2013 | China | Clinical | 88.7 | 100 | 0.05 | 98.3324 | 4982598 | 45.23 | 68 | 246580 |

| | | | | | | | | | | | | | | | | |
|---|---|---|---|---|---|---|---|---|---|---|---|---|---|---|---|---|
| VP13059 | VP13059 | SAMN20352516 | SRR15219001 | PC | Public | 2013 | China | Clinical | 98.5 | 100 | 0 | 99.9905 | 5127409 | 45.29 | 53 | 431220 |
| VP13060 | VP13060 | SAMN20352517 | SRR15219000 | PC | Public | 2013 | China | Clinical | 98.5 | 100 | 0 | 99.9915 | 5128560 | 45.29 | 53 | 401651 |
| VP13061 | VP13061 | SAMN20352539 | SRR15218941 | NonPC | Public | 2013 | China | Clinical | 92.3 | 100 | 0.19 | 98.3551 | 5039837 | 45.24 | 54 | 407322 |
| VP13064 | VP13064 | SAMN20352590 | SRR15219423 | PC | Public | 2013 | China | Clinical | 98.2 | 100 | 0 | 99.9875 | 5077369 | 45.3 | 49 | 403949 |
| VP13065 | VP13065 | SAMN20352591 | SRR15219422 | PC | Public | 2013 | China | Clinical | 98.2 | 100 | 0 | 99.9888 | 5078086 | 45.3 | 50 | 394663 |
| VP13067 | VP13067 | SAMN20352509 | SRR15219009 | PC | Public | 2013 | China | Clinical | 97.8 | 100 | 0 | 99.9332 | 5136312 | 45.3 | 57 | 401569 |
| VP13071 | VP13071 | SAMN20352615 | SRR15219327 | PC | Public | 2013 | China | Clinical | 97.5 | 100 | 0 | 99.9256 | 5118543 | 45.29 | 82 | 267762 |
| VP13072 | VP13072 | SAMN20352613 | SRR15219329 | PC | Public | 2013 | China | Clinical | 98.4 | 100 | 0.14 | 99.9845 | 5295668 | 45.13 | 68 | 425199 |
| VP13073 | VP13073 | SAMN20352593 | SRR15219420 | PC | Public | 2013 | China | Clinical | 98.2 | 100 | 0 | 99.9782 | 5076767 | 45.3 | 45 | 403949 |
| VP13076 | VP13076 | SAMN20352620 | SRR15219321 | PC | Public | 2013 | China | Clinical | 98.4 | 100 | 0.84 | 99.9939 | 5406344 | 45.13 | 79 | 312811 |
| VP13083 | VP13083 | SAMN20352627 | SRR15219314 | NonPC | Public | 2013 | China | Clinical | 88.5 | 100 | 0.19 | 98.3443 | 4967582 | 45.27 | 56 | 313276 |
| VP13084 | VP13084 | SAMN20352622 | SRR15219319 | PC | Public | 2013 | China | Clinical | 98.3 | 100 | 0 | 99.9828 | 5083431 | 45.3 | 57 | 401577 |
| VP13085 | VP13085 | SAMN20352623 | SRR15219318 | NonPC | Public | 2013 | China | Clinical | 92.2 | 100 | 0.19 | 98.3478 | 5022753 | 45.25 | 55 | 399699 |
| VP13088 | VP13088 | SAMN20352614 | SRR15219328 | PC | Public | 2013 | China | Clinical | 98.1 | 100 | 0 | 99.9692 | 5127311 | 45.28 | 46 | 461740 |
| VP13089 | VP13089 | SAMN20352628 | SRR15219313 | NonPC | Public | 2013 | China | Clinical | 91.9 | 100 | 0.19 | 98.3614 | 5022718 | 45.25 | 53 | 407286 |
| VP13091 | VP13091 | SAMN20352645 | SRR15219187 | PC | Public | 2013 | China | Clinical | 98.5 | 100 | 0 | 99.9923 | 5091581 | 45.29 | 66 | 312690 |
| VP13092 | VP13092 | SAMN20352507 | SRR15219011 | NonPC | Public | 2013 | China | Clinical | 92.9 | 100 | 0.05 | 98.3951 | 5006247 | 45.36 | 27 | 515916 |
| VP13093 | VP13093 | SAMN20352631 | SRR15219203 | NonPC | Public | 2013 | China | Clinical | 92.3 | 100 | 0.19 | 98.3529 | 5024023 | 45.25 | 58 | 373349 |
| VP13097 | VP13097 | SAMN20352621 | SRR15219320 | PC | Public | 2013 | China | Clinical | 98.4 | 100 | 0.14 | 99.9824 | 5297057 | 45.14 | 69 | 425198 |
| VP13099 | VP13099 | SAMN20352512 | SRR15219005 | PC | Public | 2013 | China | Clinical | 98.1 | 100 | 0 | 99.9784 | 5076693 | 45.3 | 47 | 403949 |
| VP13101 | VP13101 | SAMN20352624 | SRR15219317 | PC | Public | 2013 | China | Clinical | 98.2 | 100 | 0 | 99.9873 | 5076799 | 45.3 | 45 | 403949 |
| VP13103 | VP13103 | SAMN20352629 | SRR15219312 | PC | Public | 2013 | China | Clinical | 98.2 | 100 | 0 | 99.9784 | 5076948 | 45.3 | 46 | 403835 |
| VP13104 | VP13104 | SAMN20352630 | SRR15219205 | NonPC | Public | 2013 | China | Clinical | 92.5 | 100 | 0 | 98.3083 | 5074396 | 45.23 | 56 | 322749 |
| VP13106 | VP13106 | SAMN20352646 | SRR15219186 | PC | Public | 2013 | China | Clinical | 98.6 | 100 | 0 | 99.9819 | 5150186 | 45.27 | 59 | 420264 |
| VP13108 | VP13108 | SAMN20352647 | SRR15219185 | NonPC | Public | 2013 | China | Clinical | 93.1 | 100 | 0.05 | 98.3597 | 5066012 | 45.31 | 43 | 513728 |
| VP13109 | VP13109 | SAMN20352654 | SRR15219177 | PC | Public | 2013 | China | Clinical | 98.2 | 100 | 0 | 99.9784 | 5077110 | 45.3 | 44 | 440631 |
| VP13110 | VP13110 | SAMN20352655 | SRR15219176 | NonPC | Public | 2013 | China | Clinical | 93 | 100 | 0.05 | 98.3783 | 5037938 | 45.31 | 31 | 484209 |
| VP13122 | VP13122 | SAMN20352508 | SRR15219010 | NonPC | Public | 2013 | China | Clinical | 92.5 | 100 | 0.05 | 98.2931 | 5125339 | 45.25 | 46 | 496074 |
| VP13125 | VP13125 | SAMN20352503 | SRR15218491 | NonPC | Public | 2013 | China | Clinical | 92.2 | 100 | 0.19 | 98.357 | 5045587 | 45.25 | 58 | 450300 |
| VP13126 | VP13126 | SAMN20352511 | SRR15219006 | PC | Public | 2013 | China | Clinical | 98.4 | 100 | 0 | 99.9832 | 5091483 | 45.29 | 56 | 425312 |
| VP13127 | VP13127 | SAMN20352504 | SRR15218490 | NonPC | Public | 2013 | China | Clinical | 92.4 | 100 | 0.19 | 98.3618 | 5067718 | 45.28 | 53 | 372965 |
| VP14001 | VP14001 | SAMN20352730 | SRR15218576 | PC | Public | 2014 | China | Clinical | 98.4 | 100 | 0 | 99.9904 | 5090218 | 45.29 | 58 | 442187 |
| VP14006 | VP14006 | SAMN20352804 | SRR15219048 | PC | Public | 2014 | China | Clinical | 98.1 | 100 | 0 | 99.9813 | 5072308 | 45.3 | 48 | 403721 |
| VP14008 | VP14008 | SAMN20352809 | SRR15218974 | PC | Public | 2014 | China | Clinical | 98.5 | 100 | 0.01 | 99.9829 | 5182607 | 45.32 | 58 | 425313 |
| VP14009 | VP14009 | SAMN20352810 | SRR15218972 | PC | Public | 2014 | China | Clinical | 98.5 | 100 | 0 | 99.9826 | 5105446 | 45.29 | 60 | 401470 |
| VP14011 | VP14011 | SAMN20352811 | SRR15218971 | PC | Public | 2014 | China | Clinical | 98.5 | 100 | 0 | 99.9818 | 5104036 | 45.28 | 62 | 365145 |
| VP14013 | VP14013 | SAMN20352812 | SRR15218970 | PC | Public | 2014 | China | Clinical | 98.5 | 100 | 0 | 99.9824 | 5105355 | 45.29 | 63 | 394911 |
| VP14014 | VP14014 | SAMN20352813 | SRR15218969 | PC | Public | 2014 | China | Clinical | 98.5 | 100 | 0 | 99.9919 | 5104921 | 45.29 | 61 | 394909 |
| VP14015 | VP14015 | SAMN20352814 | SRR15218968 | PC | Public | 2014 | China | Clinical | 98.4 | 100 | 0 | 99.9868 | 5103815 | 45.28 | 63 | 394745 |
| VP14016 | VP14016 | SAMN20352807 | SRR15218977 | NonPC | Public | 2014 | China | Clinical | 91.9 | 100 | 0 | 98.3627 | 5004473 | 45.37 | 42 | 368294 |
| VP14017 | VP14017 | SAMN20352808 | SRR15218975 | NonPC | Public | 2014 | China | Clinical | 90.2 | 100 | 0 | 98.4019 | 5003350 | 45.33 | 57 | 401133 |
| VP14018 | VP14018 | SAMN20352816 | SRR15218966 | PC | Public | 2014 | China | Clinical | 98.1 | 100 | 0 | 99.9814 | 5073115 | 45.3 | 47 | 403715 |
| VP14020 | VP14020 | SAMN20352815 | SRR15218967 | PC | Public | 2014 | China | Clinical | 98.4 | 100 | 0 | 99.9893 | 5104575 | 45.29 | 59 | 401604 |
| VP14021 | VP14021 | SAMN20352805 | SRR15219047 | NonPC | Public | 2014 | China | Clinical | 92.3 | 100 | 0.19 | 98.3691 | 5070323 | 45.26 | 54 | 407322 |
| VP14056 | VP14056 | SAMN20352819 | SRR15218963 | PC | Public | 2014 | China | Clinical | 97.9 | 100 | 0.01 | 99.9339 | 5258401 | 45.16 | 71 | 336973 |
| VP14061 | VP14061 | SAMN20352806 | SRR15219046 | PC | Public | 2014 | China | Clinical | 98.1 | 100 | 0 | 99.9909 | 5072185 | 45.3 | 47 | 403607 |
| VP14070 | VP14070 | SAMN20352713 | SRR15218696 | PC | Public | 2014 | China | Clinical | 98.1 | 100 | 0 | 99.9894 | 5073011 | 45.3 | 48 | 403721 |
| VP14071 | VP14071 | SAMN20352887 | SRR15218781 | PC | Public | 2014 | China | Clinical | 98.3 | 100 | 0 | 99.9763 | 5142622 | 45.29 | 59 | 401604 |

| | | | | | | | | | | | | | | | | |
|---|---|---|---|---|---|---|---|---|---|---|---|---|---|---|---|---|
| VP14072 | VP14072 | SAMN20352883 | SRR15218785 | PC | Public | 2014 | China | Clinical | 98.1 | 100 | 0 | 99.991 | 5067016 | 45.31 | 48 | 403607 |
| VP14075 | VP14075 | SAMN20352897 | SRR15219446 | PC | Public | 2014 | China | Clinical | 98.5 | 100 | 0 | 99.982 | 5104441 | 45.28 | 56 | 401470 |
| VP14076 | VP14076 | SAMN20352898 | SRR15219377 | PC | Public | 2014 | China | Clinical | 98.5 | 100 | 0 | 99.9824 | 5105879 | 45.29 | 60 | 394955 |
| VP14077 | VP14077 | SAMN20352891 | SRR15219453 | NonPC | Public | 2014 | China | Env | 91.1 | 100 | 0.05 | 98.3829 | 5010214 | 45.35 | 53 | 333064 |
| VP14078 | VP14078 | SAMN20352882 | SRR15218786 | PC | Public | 2014 | China | Clinical | 98.4 | 100 | 0 | 99.99 | 5103473 | 45.28 | 57 | 401604 |
| VP14079 | VP14079 | SAMN20352803 | SRR15219049 | PC | Public | 2014 | China | Clinical | 98.3 | 100 | 0 | 99.9678 | 5140496 | 45.28 | 56 | 401611 |
| VP14080 | VP14080 | SAMN20352729 | SRR15218578 | NonPC | Public | 2014 | China | Clinical | 91.2 | 100 | 0 | 98.3823 | 5016513 | 45.31 | 51 | 312713 |
| VP14081 | VP14081 | SAMN20352820 | SRR15218961 | PC | Public | 2014 | China | Clinical | 97.9 | 100 | 0 | 99.9544 | 5140320 | 45.28 | 47 | 401463 |
| VP14082 | VP14082 | SAMN20352886 | SRR15218782 | PC | Public | 2014 | China | Clinical | 98.1 | 100 | 0 | 99.9916 | 5073516 | 45.3 | 52 | 334356 |
| VP14083 | VP14083 | SAMN20352899 | SRR15219376 | PC | Public | 2014 | China | Clinical | 98.2 | 100 | 0 | 99.9773 | 5083747 | 45.3 | 49 | 403721 |
| VP14084 | VP14084 | SAMN20352889 | SRR15218779 | PC | Public | 2014 | China | Clinical | 97.7 | 100 | 0 | 99.9004 | 5146140 | 45.29 | 53 | 419838 |
| VP14085 | VP14085 | SAMN20352890 | SRR15218777 | PC | Public | 2014 | China | Clinical | 98.1 | 100 | 0 | 99.9907 | 5072396 | 45.3 | 47 | 401597 |
| VP14086 | VP14086 | SAMN20352821 | SRR15218960 | PC | Public | 2014 | China | Clinical | 98.1 | 100 | 0 | 99.9902 | 5072424 | 45.3 | 49 | 401463 |
| VP14087 | VP14087 | SAMN20352818 | SRR15218964 | PC | Public | 2014 | China | Clinical | 98.1 | 100 | 0 | 99.99 | 5072093 | 45.3 | 46 | 404249 |
| VP14089 | VP14089 | SAMN20352817 | SRR15218965 | PC | Public | 2014 | China | Clinical | 98.1 | 100 | 0 | 99.9919 | 5072116 | 45.3 | 48 | 404249 |
| VP14090 | VP14090 | SAMN20352711 | SRR15218698 | PC | Public | 2014 | China | Clinical | 98.5 | 100 | 0 | 99.9806 | 5092834 | 45.29 | 58 | 401470 |
| VP14091 | VP14091 | SAMN20352714 | SRR15218695 | PC | Public | 2014 | China | Clinical | 98 | 100 | 0 | 99.9819 | 5069788 | 45.31 | 47 | 401597 |
| VP14093 | VP14093 | SAMN20352707 | SRR15218703 | NonPC | Public | 2014 | China | Clinical | 93.1 | 100 | 0.05 | 98.3517 | 5059293 | 45.31 | 47 | 424357 |
| VP14094 | VP14094 | SAMN20352900 | SRR15219374 | NonPC | Public | 2014 | China | Clinical | 93.1 | 100 | 0.05 | 98.4204 | 5055993 | 45.3 | 43 | 424357 |
| VP14095 | VP14095 | SAMN20352885 | SRR15218783 | PC | Public | 2014 | China | Clinical | 98.1 | 100 | 0 | 99.9899 | 5104024 | 45.32 | 50 | 403721 |
| VP14096 | VP14096 | SAMN20352712 | SRR15218697 | PC | Public | 2014 | China | Clinical | 98.5 | 100 | 0 | 99.9918 | 5104177 | 45.29 | 59 | 394666 |
| VP14099 | VP14099 | SAMN20352903 | SRR15219371 | PC | Public | 2014 | China | Clinical | 98.1 | 100 | 0 | 99.9821 | 5071756 | 45.3 | 45 | 401463 |
| VP14102 | VP14102 | SAMN20352715 | SRR15218694 | PC | Public | 2014 | China | Clinical | 98.2 | 100 | 0 | 99.9767 | 5082706 | 45.3 | 45 | 403721 |
| VP14105 | VP14105 | SAMN20352884 | SRR15218784 | PC | Public | 2014 | China | Clinical | 98.1 | 100 | 0 | 99.9915 | 5066217 | 45.31 | 48 | 401597 |
| VP14107 | VP14107 | SAMN20352906 | SRR15219368 | NonPC | Public | 2014 | China | Clinical | 92.3 | 100 | 0.19 | 98.342 | 5045603 | 45.23 | 53 | 539189 |
| VP14108 | VP14108 | SAMN20352905 | SRR15219369 | PC | Public | 2014 | China | Clinical | 98.1 | 100 | 0 | 99.9816 | 5125026 | 45.25 | 48 | 403721 |
| VP14109 | VP14109 | SAMN20352916 | SRR15219357 | PC | Public | 2014 | China | Clinical | 98.1 | 100 | 0 | 99.9812 | 5123619 | 45.25 | 48 | 403607 |
| VP14110 | VP14110 | SAMN20352919 | SRR15219354 | PC | Public | 2014 | China | Clinical | 98.1 | 100 | 0 | 99.9909 | 5072310 | 45.3 | 45 | 403721 |
| VP14111 | VP14111 | SAMN20352915 | SRR15219358 | PC | Public | 2014 | China | Clinical | 98.1 | 100 | 0 | 99.9817 | 5072238 | 45.3 | 49 | 401463 |
| VP14112 | VP14112 | SAMN20352925 | SRR15219347 | PC | Public | 2014 | China | Clinical | 98.1 | 100 | 0 | 99.981 | 5071712 | 45.3 | 46 | 403721 |
| VP14113 | VP14113 | SAMN20352926 | SRR15219346 | PC | Public | 2014 | China | Clinical | 98 | 100 | 0.14 | 99.9743 | 5295140 | 45.12 | 50 | 461740 |
| VP14114 | VP14114 | SAMN20352927 | SRR15219345 | PC | Public | 2014 | China | Clinical | 98.1 | 100 | 0.14 | 99.9902 | 5246400 | 45.12 | 47 | 403721 |
| VP14115 | VP14115 | SAMN20352928 | SRR15219344 | PC | Public | 2014 | China | Clinical | 98.1 | 100 | 0 | 99.9826 | 5072006 | 45.3 | 48 | 401597 |
| VP14117 | VP14117 | SAMN20352822 | SRR15218959 | PC | Public | 2014 | China | Clinical | 98 | 100 | 0 | 99.9726 | 5122527 | 45.3 | 45 | 461740 |
| VP14118 | VP14118 | SAMN20352914 | SRR15219359 | PC | Public | 2014 | China | Clinical | 98.5 | 100 | 0 | 99.9908 | 5092848 | 45.29 | 59 | 425295 |
| VP14119 | VP14119 | SAMN20352904 | SRR15219370 | PC | Public | 2014 | China | Clinical | 98 | 100 | 0 | 99.9638 | 5120997 | 45.3 | 48 | 461731 |
| VP14120 | VP14120 | SAMN20352921 | SRR15219351 | PC | Public | 2014 | China | Clinical | 98.5 | 100 | 0 | 99.9935 | 5093613 | 45.29 | 58 | 425193 |
| VP14121 | VP14121 | SAMN20352920 | SRR15219352 | NonPC | Public | 2014 | China | Clinical | 92.5 | 100 | 0.05 | 98.3664 | 5059399 | 45.29 | 45 | 539124 |
| VP14122 | VP14122 | SAMN20352918 | SRR15219355 | PC | Public | 2014 | China | Clinical | 98.1 | 100 | 0 | 99.9818 | 5071541 | 45.3 | 53 | 313341 |
| VP14124 | VP14124 | SAMN20352924 | SRR15219348 | PC | Public | 2014 | China | Clinical | 98.1 | 100 | 0 | 99.9821 | 5073057 | 45.31 | 50 | 403721 |
| VP14131 | VP14131 | SAMN20352902 | SRR15219372 | PC | Public | 2014 | China | Clinical | 98.1 | 100 | 0 | 99.9791 | 5107984 | 45.32 | 51 | 401446 |
| VP14133 | VP14133 | SAMN20352917 | SRR15219356 | PC | Public | 2014 | China | Clinical | 98.1 | 100 | 0 | 99.9814 | 5071296 | 45.3 | 46 | 404135 |
| VP14134 | VP14134 | SAMN20352911 | SRR15219362 | NonPC | Public | 2014 | China | Clinical | 92.3 | 100 | 0.19 | 98.3654 | 5044509 | 45.22 | 52 | 471204 |
| VP14141 | VP14141 | SAMN20352709 | SRR15218701 | PC | Public | 2014 | China | Clinical | 97.7 | 100 | 0.02 | 99.8958 | 5120242 | 45.29 | 71 | 394743 |
| VP14145 | VP14145 | SAMN20352938 | SRR15219265 | PC | Public | 2014 | China | Clinical | 97.9 | 100 | 0 | 99.9758 | 5141555 | 45.28 | 47 | 395944 |
| VP14148 | VP14148 | SAMN20352936 | SRR15219267 | PC | Public | 2014 | China | Clinical | 98.1 | 100 | 0.02 | 99.9818 | 5094145 | 45.28 | 67 | 394919 |
| VP14149 | VP14149 | SAMN20352939 | SRR15219264 | PC | Public | 2014 | China | Clinical | 98.5 | 100 | 0 | 99.9839 | 5214482 | 45.18 | 74 | 425319 |
| VP14150 | VP14150 | SAMN20352945 | SRR15219256 | PC | Public | 2014 | China | Clinical | 98.5 | 100 | 0 | 99.9815 | 5195443 | 45.21 | 64 | 394980 |

| | | | | | | | | | | | | | | | | |
|---|---|---|---|---|---|---|---|---|---|---|---|---|---|---|---|---|
| VP14153 | VP14153 | SAMN20352710 | SRR15218699 | PC | Public | 2014 | China | Clinical | 98.5 | 100 | 0 | 99.9922 | 5193482 | 45.2 | 60 | 401470 |
| VP14156 | VP14156 | SAMN20352940 | SRR15219261 | PC | Public | 2014 | China | Clinical | 98.5 | 100 | 0 | 99.992 | 5193409 | 45.2 | 60 | 425205 |
| VP14161 | VP14161 | SAMN20352946 | SRR15219255 | PC | Public | 2014 | China | Clinical | 97.9 | 100 | 0 | 99.9762 | 5139628 | 45.28 | 48 | 395140 |
| VP14162 | VP14162 | SAMN20352947 | SRR15219254 | PC | Public | 2014 | China | Clinical | 97.9 | 100 | 0 | 99.9626 | 5139600 | 45.28 | 43 | 401463 |
| VP14163 | VP14163 | SAMN20352948 | SRR15219253 | PC | Public | 2014 | China | Clinical | 97.9 | 100 | 0 | 99.9546 | 5139127 | 45.28 | 44 | 401464 |
| VP14165 | VP14165 | SAMN20352949 | SRR15219252 | PC | Public | 2014 | China | Clinical | 97.9 | 100 | 0 | 99.9398 | 5139006 | 45.28 | 45 | 401463 |
| VP14166 | VP14166 | SAMN20352950 | SRR15219250 | PC | Public | 2014 | China | Clinical | 97.9 | 100 | 0 | 99.9519 | 5138562 | 45.28 | 47 | 401464 |
| VP14167 | VP14167 | SAMN20352951 | SRR15219249 | PC | Public | 2014 | China | Clinical | 97.9 | 100 | 0 | 99.9747 | 5138851 | 45.28 | 45 | 401463 |
| VP14168 | VP14168 | SAMN20352952 | SRR15219248 | PC | Public | 2014 | China | Clinical | 97.9 | 100 | 0 | 99.9625 | 5139282 | 45.28 | 45 | 401463 |
| VP14171 | VP14171 | SAMN20352944 | SRR15219257 | PC | Public | 2014 | China | Clinical | 98.4 | 100 | 0 | 99.9852 | 5089415 | 45.29 | 56 | 425355 |
| VP14172 | VP14172 | SAMN20352937 | SRR15219266 | PC | Public | 2014 | China | Clinical | 98.1 | 100 | 0 | 99.9808 | 5071670 | 45.3 | 45 | 403703 |
| VP14173 | VP14173 | SAMN20352935 | SRR15219268 | PC | Public | 2014 | China | Clinical | 98.1 | 100 | 0 | 99.9812 | 5071719 | 45.3 | 46 | 403703 |
| VP15001 | VP15001 | SAMN20353010 | SRR15218730 | NonPC | Public | 2015 | China | Clinical | 91.3 | 100 | 0 | 98.3452 | 5029353 | 45.32 | 42 | 406900 |
| VP15002 | VP15002 | SAMN20353027 | SRR15218679 | PC | Public | 2015 | China | Clinical | 97.4 | 100 | 0 | 99.9218 | 5094812 | 45.28 | 82 | 197933 |
| VP15003 | VP15003 | SAMN20353006 | SRR15218735 | PC | Public | 2015 | China | Clinical | 98.4 | 100 | 0 | 99.9836 | 5194946 | 45.21 | 63 | 425319 |
| VP15004 | VP15004 | SAMN20353029 | SRR15218677 | PC | Public | 2015 | China | Clinical | 97.9 | 100 | 0 | 99.9925 | 5163732 | 45.19 | 84 | 266114 |
| VP15005 | VP15005 | SAMN20353028 | SRR15218678 | PC | Public | 2015 | China | Clinical | 98.4 | 100 | 0 | 99.9827 | 5092361 | 45.29 | 59 | 425193 |
| VP15006 | VP15006 | SAMN20353034 | SRR15218671 | PC | Public | 2015 | China | Clinical | 97.9 | 100 | 0.07 | 99.9084 | 5181377 | 45.26 | 61 | 394983 |
| VP15007 | VP15007 | SAMN20353076 | SRR15218487 | PC | Public | 2015 | China | Clinical | 98.2 | 100 | 0 | 99.9748 | 5137756 | 45.28 | 55 | 475491 |
| VP15008 | VP15008 | SAMN20353079 | SRR15218484 | PC | Public | 2015 | China | Clinical | 97.9 | 100 | 0 | 99.9512 | 5202113 | 45.27 | 48 | 394836 |
| VP15009 | VP15009 | SAMN20353081 | SRR15218481 | NonPC | Public | 2015 | China | Clinical | 91.1 | 100 | 0.05 | 98.2385 | 5170601 | 45.17 | 80 | 328920 |
| VP15010 | VP15010 | SAMN20353080 | SRR15218482 | NonPC | Public | 2015 | China | Clinical | 92.9 | 100 | 0.19 | 98.3626 | 5139999 | 45.12 | 53 | 410581 |
| VP15011 | VP15011 | SAMN20353082 | SRR15218480 | NonPC | Public | 2015 | China | Clinical | 89.9 | 100 | 0.05 | 98.3269 | 5221616 | 45.14 | 59 | 327013 |
| VP15012 | VP15012 | SAMN20353085 | SRR15218477 | PC | Public | 2015 | China | Clinical | 98.4 | 100 | 0 | 99.9907 | 5092508 | 45.29 | 56 | 425199 |
| VP15013 | VP15013 | SAMN20353077 | SRR15218486 | PC | Public | 2015 | China | Clinical | 98.2 | 100 | 0 | 99.967 | 5193786 | 45.31 | 58 | 401241 |
| VP15014 | VP15014 | SAMN20353078 | SRR15218485 | PC | Public | 2015 | China | Clinical | 98.3 | 100 | 0 | 99.9738 | 5194466 | 45.31 | 62 | 394836 |
| VP15015 | VP15015 | SAMN20353086 | SRR15218476 | NonPC | Public | 2015 | China | Clinical | 92.9 | 100 | 0.05 | 98.3473 | 5050988 | 45.3 | 50 | 417409 |
| VP15016 | VP15016 | SAMN20353095 | SRR15218466 | PC | Public | 2015 | China | Clinical | 98.6 | 100 | 0 | 99.9767 | 5134149 | 45.26 | 59 | 420398 |
| VP15019 | VP15019 | SAMN20353151 | SRR15219027 | PC | Public | 2015 | China | Clinical | 98.5 | 100 | 0 | 99.9853 | 5092068 | 45.29 | 60 | 425307 |
| VP15020 | VP15020 | SAMN20353150 | SRR15219028 | PC | Public | 2015 | China | Clinical | 97.9 | 100 | 0 | 99.9534 | 5150531 | 45.29 | 46 | 401463 |
| VP15021 | VP15021 | SAMN20353149 | SRR15219030 | PC | Public | 2015 | China | Clinical | 97.9 | 100 | 0 | 99.9757 | 5139367 | 45.28 | 46 | 401464 |
| VP15022 | VP15022 | SAMN20353148 | SRR15219031 | PC | Public | 2015 | China | Clinical | 97.9 | 100 | 0 | 99.9762 | 5148435 | 45.29 | 47 | 401463 |
| VP15023 | VP15023 | SAMN20353157 | SRR15219021 | NonPC | Public | 2015 | China | Clinical | 92.1 | 100 | 0.19 | 98.3647 | 5030625 | 45.22 | 60 | 368057 |
| VP15024 | VP15024 | SAMN20353152 | SRR15219026 | NonPC | Public | 2015 | China | Clinical | 91.8 | 100 | 0.46 | 98.2406 | 5057166 | 45.21 | 58 | 313978 |
| VP15026 | VP15026 | SAMN20353155 | SRR15219023 | PC | Public | 2015 | China | Clinical | 97.9 | 100 | 0 | 99.9676 | 5238344 | 45.21 | 48 | 401463 |
| VP15027 | VP15027 | SAMN20353154 | SRR15219024 | PC | Public | 2015 | China | Clinical | 98.5 | 100 | 0 | 99.986 | 5093080 | 45.29 | 55 | 425307 |
| VP15030 | VP15030 | SAMN20353153 | SRR15219025 | PC | Public | 2015 | China | Clinical | 98.5 | 100 | 0 | 99.9808 | 5092600 | 45.29 | 54 | 425193 |
| VP15031 | VP15031 | SAMN20353087 | SRR15218475 | NonPC | Public | 2015 | China | Clinical | 91.3 | 100 | 0.05 | 98.2875 | 5188966 | 45.17 | 64 | 348120 |
| VP15032 | VP15032 | SAMN20353165 | SRR15219012 | PC | Public | 2015 | China | Clinical | 98.5 | 100 | 0 | 99.9814 | 5092071 | 45.29 | 56 | 425307 |
| VP15033 | VP15033 | SAMN20353164 | SRR15219013 | PC | Public | 2015 | China | Clinical | 97.9 | 100 | 0 | 99.978 | 5149861 | 45.29 | 44 | 401597 |
| VP15034 | VP15034 | SAMN20353162 | SRR15219015 | PC | Public | 2015 | China | Clinical | 97.9 | 100 | 0 | 99.9696 | 5139235 | 45.28 | 49 | 394911 |
| VP15035 | VP15035 | SAMN20353158 | SRR15219020 | PC | Public | 2015 | China | Clinical | 97.9 | 100 | 0 | 99.9613 | 5149815 | 45.29 | 48 | 401597 |
| VP15036 | VP15036 | SAMN20353159 | SRR15219019 | PC | Public | 2015 | China | Clinical | 97.9 | 100 | 0 | 99.9778 | 5138751 | 45.28 | 47 | 401597 |
| VP15037 | VP15037 | SAMN20353161 | SRR15219016 | PC | Public | 2015 | China | Clinical | 97.9 | 100 | 0 | 99.9763 | 5139345 | 45.28 | 50 | 394911 |
| VP15038 | VP15038 | SAMN20353176 | SRR15218895 | PC | Public | 2015 | China | Clinical | 97.9 | 100 | 0 | 99.9673 | 5150068 | 45.29 | 45 | 401597 |
| VP15040 | VP15040 | SAMN20353180 | SRR15218890 | PC | Public | 2015 | China | Clinical | 97.9 | 100 | 0 | 99.9634 | 5150732 | 45.29 | 45 | 401463 |
| VP15043 | VP15043 | SAMN20353181 | SRR15218889 | PC | Public | 2015 | China | Clinical | 97.9 | 100 | 0 | 99.9679 | 5151204 | 45.29 | 46 | 401463 |
| VP15044 | VP15044 | SAMN20353182 | SRR15218888 | PC | Public | 2015 | China | Clinical | 96.4 | 100 | 0 | 99.9269 | 5123668 | 45.24 | 64 | 288863 |

| | | | | | | | | | | | | | | | | |
|---|---|---|---|---|---|---|---|---|---|---|---|---|---|---|---|---|
| VP15048 | VP15048 | SAMN20353156 | SRR15219022 | PC | Public | 2015 | China | Clinical | 98.4 | 100 | 0 | 99.9812 | 5092699 | 45.29 | 59 | 394980 |
| VP15049 | VP15049 | SAMN20353146 | SRR15219033 | NonPC | Public | 2015 | China | Clinical | 91.3 | 100 | 0.05 | 98.2332 | 5188213 | 45.17 | 67 | 335747 |
| VP15050 | VP15050 | SAMN20353147 | SRR15219032 | NonPC | Public | 2015 | China | Clinical | 91.3 | 100 | 0.05 | 98.1967 | 5281185 | 45.09 | 64 | 348293 |
| VP15052 | VP15052 | SAMN20353160 | SRR15219017 | PC | Public | 2015 | China | Clinical | 98.2 | 100 | 0 | 99.9895 | 5080916 | 45.3 | 48 | 442197 |
| VP15053 | VP15053 | SAMN20353186 | SRR15218884 | PC | Public | 2015 | China | Clinical | 97.9 | 100 | 0 | 99.954 | 5181617 | 45.29 | 50 | 418310 |
| VP15054 | VP15054 | SAMN20353188 | SRR15218882 | PC | Public | 2015 | China | Clinical | 97.9 | 100 | 0 | 99.9545 | 5149084 | 45.29 | 46 | 401463 |
| VP15055 | VP15055 | SAMN20353183 | SRR15218887 | PC | Public | 2015 | China | Clinical | 98.5 | 100 | 0 | 99.9826 | 5093920 | 45.29 | 58 | 401604 |
| VP15056 | VP15056 | SAMN20353184 | SRR15218886 | PC | Public | 2015 | China | Clinical | 98.5 | 100 | 0 | 99.9907 | 5094024 | 45.29 | 57 | 425307 |
| VP15058 | VP15058 | SAMN20353191 | SRR15218878 | PC | Public | 2015 | China | Clinical | 98.5 | 100 | 0 | 99.9812 | 5092876 | 45.29 | 57 | 425721 |
| VP15059 | VP15059 | SAMN20353192 | SRR15218877 | PC | Public | 2015 | China | Clinical | 97.9 | 100 | 0 | 99.9552 | 5149884 | 45.29 | 46 | 401597 |
| VP15060 | VP15060 | SAMN20353193 | SRR15218876 | PC | Public | 2015 | China | Clinical | 98.5 | 100 | 0 | 99.9898 | 5092560 | 45.29 | 59 | 425193 |
| VP15061 | VP15061 | SAMN20353197 | SRR15218872 | PC | Public | 2015 | China | Clinical | 97.9 | 100 | 0 | 99.978 | 5150093 | 45.29 | 48 | 394909 |
| VP15063 | VP15063 | SAMN20353200 | SRR15218868 | PC | Public | 2015 | China | Clinical | 97.9 | 100 | 0 | 99.9545 | 5151541 | 45.29 | 50 | 394911 |
| VP15064 | VP15064 | SAMN20353203 | SRR15218865 | PC | Public | 2015 | China | Clinical | 97.9 | 100 | 0 | 99.9772 | 5150316 | 45.29 | 46 | 401597 |
| VP15065 | VP15065 | SAMN20353209 | SRR15218859 | PC | Public | 2015 | China | Clinical | 97.9 | 100 | 0 | 99.9698 | 5150736 | 45.29 | 53 | 394970 |
| VP15066 | VP15066 | SAMN20353210 | SRR15218857 | PC | Public | 2015 | China | Clinical | 97.9 | 100 | 0 | 99.9644 | 5149528 | 45.29 | 53 | 334356 |
| VP15067 | VP15067 | SAMN20353208 | SRR15218860 | PC | Public | 2015 | China | Clinical | 97.9 | 100 | 0 | 99.9546 | 5151448 | 45.3 | 51 | 387121 |
| VP15068 | VP15068 | SAMN20353204 | SRR15218864 | PC | Public | 2015 | China | Clinical | 97.9 | 100 | 0 | 99.9622 | 5151148 | 45.29 | 50 | 387121 |
| VP15069 | VP15069 | SAMN20353205 | SRR15218863 | PC | Public | 2015 | China | Clinical | 97.9 | 100 | 0 | 99.9686 | 5180819 | 45.29 | 50 | 401463 |
| VP15070 | VP15070 | SAMN20353207 | SRR15218861 | PC | Public | 2015 | China | Clinical | 97.9 | 100 | 0 | 99.9548 | 5151939 | 45.29 | 52 | 334342 |
| VP15071 | VP15071 | SAMN20353206 | SRR15218862 | PC | Public | 2015 | China | Clinical | 97.9 | 100 | 0 | 99.9484 | 5150323 | 45.29 | 50 | 334342 |
| VP15072 | VP15072 | SAMN20353199 | SRR15218870 | PC | Public | 2015 | China | Clinical | 98.3 | 100 | 0 | 99.9671 | 5142056 | 45.28 | 60 | 397298 |
| VP15073 | VP15073 | SAMN20353198 | SRR15218871 | PC | Public | 2015 | China | Clinical | 97.9 | 100 | 0 | 99.9769 | 5148663 | 45.29 | 47 | 401597 |
| VP15074 | VP15074 | SAMN20353196 | SRR15218873 | PC | Public | 2015 | China | Clinical | 98.2 | 100 | 0 | 99.9833 | 5080146 | 45.3 | 49 | 394962 |
| VP15075 | VP15075 | SAMN20353195 | SRR15218874 | NonPC | Public | 2015 | China | Clinical | 93.1 | 100 | 0.05 | 98.3828 | 5141988 | 45.12 | 48 | 424378 |
| VP15076 | VP15076 | SAMN20353187 | SRR15218883 | PC | Public | 2015 | China | Clinical | 97.9 | 100 | 0 | 99.9638 | 5149672 | 45.29 | 58 | 312789 |
| VP15078 | VP15078 | SAMN20353012 | SRR15218728 | NonPC | Public | 2015 | China | Clinical | 91.9 | 100 | 0.05 | 98.4081 | 5065189 | 45.28 | 70 | 464904 |
| VP15079 | VP15079 | SAMN20353224 | SRR15218842 | PC | Public | 2015 | China | Clinical | 97.9 | 100 | 0 | 99.9643 | 5150337 | 45.29 | 53 | 394909 |
| VP15080 | VP15080 | SAMN20353215 | SRR15218852 | PC | Public | 2015 | China | Clinical | 97.9 | 100 | 0 | 99.9675 | 5149907 | 45.29 | 47 | 401463 |
| VP15081 | VP15081 | SAMN20353211 | SRR15218856 | PC | Public | 2015 | China | Clinical | 97.9 | 100 | 0 | 99.9555 | 5181843 | 45.29 | 58 | 313085 |
| VP15082 | VP15082 | SAMN20353220 | SRR15218846 | PC | Public | 2015 | China | Clinical | 97.9 | 100 | 0 | 99.9783 | 5149934 | 45.29 | 48 | 401597 |
| VP15083 | VP15083 | SAMN20353222 | SRR15218844 | PC | Public | 2015 | China | Clinical | 97.9 | 100 | 0 | 99.965 | 5150721 | 45.29 | 45 | 401463 |
| VP15084 | VP15084 | SAMN20353218 | SRR15218849 | PC | Public | 2015 | China | Clinical | 98.5 | 100 | 0 | 99.989 | 5092836 | 45.29 | 56 | 425307 |
| VP15085 | VP15085 | SAMN20353217 | SRR15218850 | PC | Public | 2015 | China | Clinical | 97.2 | 100 | 0 | 99.9032 | 5091751 | 45.28 | 80 | 197824 |
| VP15086 | VP15086 | SAMN20353219 | SRR15218848 | PC | Public | 2015 | China | Clinical | 98.5 | 100 | 0 | 99.9906 | 5092656 | 45.29 | 56 | 425307 |
| VP15087 | VP15087 | SAMN20353221 | SRR15218845 | PC | Public | 2015 | China | Clinical | 97.9 | 100 | 0 | 99.9625 | 5151935 | 45.29 | 46 | 440639 |
| VP15088 | VP15088 | SAMN20353225 | SRR15218841 | PC | Public | 2015 | China | Clinical | 97.9 | 100 | 0 | 99.9761 | 5150565 | 45.29 | 47 | 401597 |
| VP15089 | VP15089 | SAMN20353223 | SRR15218843 | NonPC | Public | 2015 | China | Clinical | 92.3 | 100 | 0.19 | 98.3783 | 5030026 | 45.24 | 50 | 450301 |
| VP15090 | VP15090 | SAMN20353226 | SRR15218840 | PC | Public | 2015 | China | Clinical | 97.9 | 100 | 0 | 99.9677 | 5181659 | 45.29 | 43 | 418838 |
| VP15091 | VP15091 | SAMN20353212 | SRR15218855 | PC | Public | 2015 | China | Clinical | 97.9 | 100 | 0 | 99.9531 | 5149403 | 45.29 | 45 | 401597 |
| VP15092 | VP15092 | SAMN20353213 | SRR15218854 | PC | Public | 2015 | China | Clinical | 97.9 | 100 | 0 | 99.952 | 5150335 | 45.29 | 49 | 401463 |
| VP15093 | VP15093 | SAMN20353185 | SRR15218885 | PC | Public | 2015 | China | Clinical | 97.9 | 100 | 0 | 99.9547 | 5138119 | 45.28 | 47 | 394957 |
| VP15095 | VP15095 | SAMN20353214 | SRR15218853 | PC | Public | 2015 | China | Clinical | 97.9 | 100 | 0 | 99.9749 | 5180789 | 45.29 | 49 | 401583 |
| VP15107 | VP15107 | SAMN20353229 | SRR15218837 | PC | Public | 2015 | China | Clinical | 98.5 | 100 | 0 | 99.9817 | 5134321 | 45.34 | 57 | 401604 |
| VP15108 | VP15108 | SAMN20353189 | SRR15218881 | PC | Public | 2015 | China | Clinical | 98.5 | 100 | 0.27 | 99.9824 | 5134680 | 45.3 | 62 | 302696 |
| VP15109 | VP15109 | SAMN20353190 | SRR15218879 | PC | Public | 2015 | China | Clinical | 98.5 | 100 | 0 | 99.9899 | 5093588 | 45.29 | 62 | 312690 |
| VP15110 | VP15110 | SAMN20353194 | SRR15218875 | PC | Public | 2015 | China | Clinical | 97.9 | 100 | 0 | 99.9684 | 5139538 | 45.28 | 52 | 373480 |
| VP15111 | VP15111 | SAMN20353201 | SRR15218867 | PC | Public | 2015 | China | Clinical | 97.9 | 100 | 0 | 99.9745 | 5149261 | 45.29 | 54 | 373614 |

| | | | | | | | | | | | | | | | | |
|---|---|---|---|---|---|---|---|---|---|---|---|---|---|---|---|---|
| VP15112 | VP15112 | SAMN20353216 | SRR15218851 | PC | Public | 2015 | China | Clinical | 97.6 | 100 | 0 | 99.9582 | 5127965 | 45.28 | 72 | 244257 |
| VP15113 | VP15113 | SAMN20353230 | SRR15219413 | NonPC | Public | 2015 | China | Clinical | 88.7 | 100 | 0.05 | 98.3047 | 4982934 | 45.22 | 65 | 246580 |
| VP15114 | VP15114 | SAMN20353019 | SRR15218688 | NonPC | Public | 2015 | China | Clinical | 92.9 | 100 | 0.19 | 98.3625 | 5064905 | 45.28 | 58 | 419311 |
| VP15115 | VP15115 | SAMN20353255 | SRR15219384 | NonPC | Public | 2015 | China | Clinical | 88.7 | 100 | 0.05 | 98.318 | 4991307 | 45.22 | 64 | 257521 |
| VP15119 | VP15119 | SAMN20353235 | SRR15219407 | PC | Public | 2015 | China | Clinical | 97.9 | 100 | 0 | 99.9756 | 5149703 | 45.29 | 52 | 387121 |
| VP15121 | VP15121 | SAMN20353254 | SRR15219385 | PC | Public | 2015 | China | Clinical | 97.9 | 100 | 0 | 99.9552 | 5141782 | 45.28 | 48 | 401597 |
| VP15122 | VP15122 | SAMN20353257 | SRR15219382 | PC | Public | 2015 | China | Clinical | 97.9 | 100 | 0 | 99.9624 | 5149198 | 45.29 | 46 | 401597 |
| VP15123 | VP15123 | SAMN20353256 | SRR15219383 | PC | Public | 2015 | China | Clinical | 97.9 | 100 | 0 | 99.9536 | 5150646 | 45.29 | 46 | 401597 |
| VP15125 | VP15125 | SAMN20353259 | SRR15219380 | NonPC | Public | 2015 | China | Clinical | 90.9 | 100 | 0.33 | 98.2922 | 5158770 | 45.17 | 58 | 315380 |
| VP15126 | VP15126 | SAMN20353260 | SRR15219378 | PC | Public | 2015 | China | Clinical | 97.6 | 100 | 0 | 99.9777 | 5144431 | 45.26 | 72 | 288535 |
| VP15127 | VP15127 | SAMN20353007 | SRR15218734 | PC | Public | 2015 | China | Clinical | 98.3 | 100 | 0 | 99.9784 | 5181075 | 45.27 | 45 | 461740 |
| VP15128 | VP15128 | SAMN20353234 | SRR15219408 | PC | Public | 2015 | China | Clinical | 97.9 | 100 | 0 | 99.963 | 5150549 | 45.29 | 54 | 334356 |
| VP15129 | VP15129 | SAMN20353233 | SRR15219409 | PC | Public | 2015 | China | Clinical | 97.9 | 100 | 0 | 99.9551 | 5138780 | 45.28 | 45 | 395142 |
| VP15130 | VP15130 | SAMN20353227 | SRR15218839 | PC | Public | 2015 | China | Clinical | 98.5 | 100 | 0 | 99.9921 | 5097557 | 45.29 | 63 | 426373 |
| VP15131 | VP15131 | SAMN20353228 | SRR15218838 | PC | Public | 2015 | China | Clinical | 97.9 | 100 | 0 | 99.9633 | 5150759 | 45.29 | 48 | 401597 |
| VP15132 | VP15132 | SAMN20353258 | SRR15219381 | PC | Public | 2015 | China | Clinical | 97.8 | 100 | 0 | 99.9643 | 5201343 | 45.29 | 46 | 461740 |
| VP15133 | VP15133 | SAMN20353236 | SRR15219406 | PC | Public | 2015 | China | Clinical | 97.9 | 100 | 0 | 99.9662 | 5181771 | 45.3 | 47 | 401597 |
| VP15134 | VP15134 | SAMN20353231 | SRR15219411 | PC | Public | 2015 | China | Clinical | 98.4 | 100 | 0 | 99.9837 | 5090016 | 45.29 | 61 | 394735 |
| VP15135 | VP15135 | SAMN20353262 | SRR15219309 | PC | Public | 2015 | China | Clinical | 97.9 | 100 | 0 | 99.952 | 5181039 | 45.29 | 47 | 401463 |
| VP15136 | VP15136 | SAMN20353261 | SRR15219310 | PC | Public | 2015 | China | Clinical | 98.5 | 100 | 0 | 99.9893 | 5093584 | 45.29 | 56 | 425307 |
| VP15137 | VP15137 | SAMN20353263 | SRR15219308 | PC | Public | 2015 | China | Clinical | 97.9 | 100 | 0 | 99.9618 | 5171001 | 45.29 | 45 | 418196 |
| VP15138 | VP15138 | SAMN20353267 | SRR15219304 | PC | Public | 2015 | China | Clinical | 98.4 | 100 | 0 | 99.9839 | 5089643 | 45.29 | 61 | 394964 |
| VP15139 | VP15139 | SAMN20353265 | SRR15219306 | PC | Public | 2015 | China | Clinical | 98.4 | 100 | 0 | 99.9862 | 5123389 | 45.29 | 58 | 401337 |
| VP15140 | VP15140 | SAMN20353266 | SRR15219305 | PC | Public | 2015 | China | Clinical | 97.9 | 100 | 0 | 99.9396 | 5150442 | 45.29 | 49 | 401597 |
| VP15141 | VP15141 | SAMN20353271 | SRR15219299 | NonPC | Public | 2015 | China | Clinical | 92.1 | 100 | 0.19 | 98.3624 | 5015922 | 45.24 | 60 | 450180 |
| VP15142 | VP15142 | SAMN20353232 | SRR15219410 | PC | Public | 2015 | China | Clinical | 97.5 | 99.18 | 0 | 99.9818 | 5042166 | 45.3 | 56 | 401471 |
| VP15143 | VP15143 | SAMN20353238 | SRR15219404 | PC | Public | 2015 | China | Clinical | 98.2 | 100 | 0 | 99.9881 | 5098303 | 45.28 | 58 | 425469 |
| VP15144 | VP15144 | SAMN20353269 | SRR15219302 | PC | Public | 2015 | China | Clinical | 98.4 | 100 | 0.14 | 99.9926 | 5305712 | 45.12 | 72 | 425301 |
| VP15145 | VP15145 | SAMN20353270 | SRR15219300 | PC | Public | 2015 | China | Clinical | 97.4 | 100 | 0 | 99.8915 | 5093526 | 45.27 | 78 | 205045 |
| VP15146 | VP15146 | SAMN20353272 | SRR15219298 | PC | Public | 2015 | China | Clinical | 98.3 | 100 | 0.14 | 99.9725 | 5318547 | 45.12 | 76 | 401605 |
| VP15147 | VP15147 | SAMN20353273 | SRR15219297 | NonPC | Public | 2015 | China | Clinical | 92.3 | 100 | 0.19 | 98.341 | 5029871 | 45.24 | 53 | 450301 |
| VP15148 | VP15148 | SAMN20353030 | SRR15218675 | PC | Public | 2015 | China | Clinical | 98.5 | 100 | 0 | 99.989 | 5094854 | 45.29 | 58 | 425835 |
| VP15149 | VP15149 | SAMN20353033 | SRR15218672 | PC | Public | 2015 | China | Clinical | 97.9 | 100 | 0 | 99.9674 | 5150777 | 45.29 | 48 | 401463 |
| VP15150 | VP15150 | SAMN20353202 | SRR15218866 | PC | Public | 2015 | China | Clinical | 97.9 | 100 | 0 | 99.9602 | 5181991 | 45.3 | 49 | 401463 |
| VP15151 | VP15151 | SAMN20353237 | SRR15219405 | PC | Public | 2015 | China | Clinical | 97.9 | 100 | 0 | 99.968 | 5149145 | 45.29 | 47 | 401597 |
| VP15152 | VP15152 | SAMN20353278 | SRR15219292 | PC | Public | 2015 | China | Clinical | 98.5 | 100 | 0 | 99.9899 | 5092247 | 45.29 | 56 | 425307 |
| VP15153 | VP15153 | SAMN20353275 | SRR15219295 | PC | Public | 2015 | China | Clinical | 97.9 | 100 | 0 | 99.9766 | 5141014 | 45.28 | 47 | 401470 |
| VP15154 | VP15154 | SAMN20353274 | SRR15219296 | PC | Public | 2015 | China | Clinical | 98.1 | 100 | 0 | 99.991 | 5108535 | 45.32 | 55 | 394917 |
| VP15156 | VP15156 | SAMN20353277 | SRR15219293 | PC | Public | 2015 | China | Clinical | 98.4 | 100 | 0 | 99.9816 | 5092067 | 45.29 | 56 | 425307 |
| VP15157 | VP15157 | SAMN20353268 | SRR15219303 | NonPC | Public | 2015 | China | Clinical | 93 | 100 | 0.05 | 98.3696 | 5047478 | 45.32 | 55 | 417624 |
| VP15159 | VP15159 | SAMN20353283 | SRR15219286 | PC | Public | 2015 | China | Clinical | 98.1 | 100 | 0 | 99.9935 | 5067876 | 45.27 | 81 | 266000 |
| VP15160 | VP15160 | SAMN20353284 | SRR15219285 | PC | Public | 2015 | China | Clinical | 98.6 | 100 | 0 | 99.9795 | 5149965 | 45.27 | 62 | 420398 |
| VP15161 | VP15161 | SAMN20353285 | SRR15219284 | PC | Public | 2015 | China | Clinical | 98.3 | 100 | 0 | 99.9889 | 5128957 | 45.26 | 81 | 288527 |
| VP15162 | VP15162 | SAMN20353286 | SRR15219283 | PC | Public | 2015 | China | Clinical | 98.1 | 100 | 0 | 99.9917 | 5071305 | 45.28 | 79 | 265886 |
| VP15163 | VP15163 | SAMN20353289 | SRR15219280 | PC | Public | 2015 | China | Clinical | 98.4 | 100 | 0 | 99.9804 | 5093149 | 45.29 | 57 | 425307 |
| VP15164 | VP15164 | SAMN20353276 | SRR15219294 | NonPC | Public | 2015 | China | Clinical | 93 | 100 | 0.05 | 98.3561 | 5128287 | 45.13 | 49 | 419408 |
| VP15165 | VP15165 | SAMN20353020 | SRR15218686 | NonPC | Public | 2015 | China | Clinical | 93.1 | 100 | 0.05 | 98.3432 | 5084328 | 45.29 | 47 | 424373 |
| VP15166 | VP15166 | SAMN20353025 | SRR15218681 | NonPC | Public | 2015 | China | Clinical | 92.1 | 100 | 0.19 | 98.36 | 5012958 | 45.24 | 62 | 313839 |

| | | | | | | | | | | | | | | | | |
|---|---|---|---|---|---|---|---|---|---|---|---|---|---|---|---|---|
| VP15167 | VP15167 | SAMN20353287 | SRR15219282 | PC | Public | 2015 | China | Clinical | 98.2 | 100 | 0 | 99.9913 | 5098475 | 45.28 | 59 | 425355 |
| VP15168 | VP15168 | SAMN20353296 | SRR15219237 | PC | Public | 2015 | China | Clinical | 98.5 | 100 | 0 | 99.9811 | 5093378 | 45.29 | 56 | 425307 |
| VP15169 | VP15169 | SAMN20353290 | SRR15219278 | PC | Public | 2015 | China | Clinical | 98.5 | 100 | 0 | 99.9903 | 5100173 | 45.29 | 61 | 401604 |
| VP15170 | VP15170 | SAMN20353026 | SRR15218680 | PC | Public | 2015 | China | Clinical | 98.3 | 100 | 0 | 99.9748 | 5192042 | 45.22 | 58 | 401463 |
| VP15171 | VP15171 | SAMN20353292 | SRR15219275 | PC | Public | 2015 | China | Clinical | 98.5 | 100 | 0 | 99.9827 | 5092935 | 45.29 | 56 | 425319 |
| VP15172 | VP15172 | SAMN20353291 | SRR15219277 | PC | Public | 2015 | China | Clinical | 98.5 | 100 | 0 | 99.981 | 5093754 | 45.29 | 56 | 425307 |
| VP15173 | VP15173 | SAMN20353293 | SRR15219240 | PC | Public | 2015 | China | Clinical | 98.5 | 100 | 0 | 99.9908 | 5092444 | 45.29 | 55 | 425193 |
| VP15174 | VP15174 | SAMN20353294 | SRR15219239 | PC | Public | 2015 | China | Clinical | 98.5 | 100 | 0 | 99.9817 | 5093270 | 45.29 | 57 | 425313 |
| VP15175 | VP15175 | SAMN20353295 | SRR15219238 | PC | Public | 2015 | China | Clinical | 98.5 | 100 | 0 | 99.9812 | 5093580 | 45.29 | 62 | 394971 |
| VP15176 | VP15176 | SAMN20353297 | SRR15219236 | PC | Public | 2015 | China | Clinical | 98.3 | 100 | 0 | 99.975 | 5192477 | 45.22 | 57 | 401597 |
| VP15177 | VP15177 | SAMN20353298 | SRR15219235 | NonPC | Public | 2015 | China | Clinical | 93 | 100 | 0.19 | 98.3306 | 5073794 | 45.28 | 54 | 419408 |
| VP15178 | VP15178 | SAMN20353299 | SRR15219234 | PC | Public | 2015 | China | Clinical | 98.5 | 100 | 0 | 99.9928 | 5092247 | 45.29 | 56 | 401611 |
| VP15179 | VP15179 | SAMN20353302 | SRR15219230 | PC | Public | 2015 | China | Clinical | 98.4 | 100 | 0 | 99.9815 | 5092437 | 45.29 | 55 | 425193 |
| VP15180 | VP15180 | SAMN20353301 | SRR15219231 | PC | Public | 2015 | China | Clinical | 98.5 | 100 | 0 | 99.9812 | 5092770 | 45.29 | 57 | 425307 |
| VP15181 | VP15181 | SAMN20353300 | SRR15219232 | NonPC | Public | 2015 | China | Clinical | 92.9 | 100 | 0.19 | 98.3904 | 5090339 | 45.27 | 51 | 313445 |
| VP15182 | VP15182 | SAMN20353303 | SRR15219229 | NonPC | Public | 2015 | China | Clinical | 92.9 | 100 | 0.05 | 98.3299 | 5053676 | 45.3 | 50 | 419408 |
| VP15183 | VP15183 | SAMN20353305 | SRR15219227 | PC | Public | 2015 | China | Clinical | 98.5 | 100 | 0 | 99.9818 | 5092723 | 45.29 | 55 | 425421 |
| VP15184 | VP15184 | SAMN20353288 | SRR15219281 | NonPC | Public | 2015 | China | Clinical | 92.3 | 100 | 0.19 | 98.3609 | 5027663 | 45.24 | 53 | 539189 |
| VP15185 | VP15185 | SAMN20353031 | SRR15218674 | PC | Public | 2015 | China | Clinical | 98.5 | 100 | 0 | 99.9799 | 5093474 | 45.29 | 56 | 425193 |
| VP15186 | VP15186 | SAMN20353032 | SRR15218673 | PC | Public | 2015 | China | Clinical | 98.5 | 100 | 0 | 99.9901 | 5092866 | 45.29 | 57 | 425307 |
| VP15187 | VP15187 | SAMN20353306 | SRR15219226 | PC | Public | 2015 | China | Clinical | 98.5 | 100 | 0 | 99.9838 | 5093944 | 45.29 | 59 | 425307 |
| VP15189 | VP15189 | SAMN20353304 | SRR15219228 | NonPC | Public | 2015 | China | Clinical | 92.3 | 100 | 0.19 | 98.3463 | 5027941 | 45.24 | 52 | 471325 |
| VP15190 | VP15190 | SAMN20353316 | SRR15219215 | PC | Public | 2015 | China | Clinical | 98.4 | 100 | 0 | 99.9911 | 5089350 | 45.29 | 57 | 425223 |
| VP15191 | VP15191 | SAMN20353313 | SRR15219218 | PC | Public | 2015 | China | Clinical | 97.9 | 100 | 0 | 99.9767 | 5139862 | 45.28 | 44 | 401597 |
| VP15192 | VP15192 | SAMN20353314 | SRR15219217 | PC | Public | 2015 | China | Clinical | 98.4 | 100 | 0.27 | 99.9806 | 5233141 | 45.33 | 60 | 425187 |
| VP15193 | VP15193 | SAMN20353312 | SRR15219219 | PC | Public | 2015 | China | Clinical | 97.9 | 100 | 0 | 99.9516 | 5149679 | 45.29 | 49 | 401597 |
| VP15194 | VP15194 | SAMN20353309 | SRR15219223 | PrePC | Public | 2015 | China | Clinical | 94.5 | 100 | 0 | 99.5142 | 5102309 | 45.34 | 69 | 461849 |
| VP15195 | VP15195 | SAMN20353315 | SRR15219216 | PC | Public | 2015 | China | Clinical | 98.5 | 100 | 0 | 99.9897 | 5094109 | 45.29 | 59 | 401337 |
| VP15196 | VP15196 | SAMN20353308 | SRR15219224 | NonPC | Public | 2015 | China | Clinical | 90.3 | 100 | 0 | 98.4274 | 4960664 | 45.38 | 51 | 259511 |
| VP15197 | VP15197 | SAMN20353317 | SRR15219214 | PC | Public | 2015 | China | Clinical | 98.4 | 100 | 0 | 99.9904 | 5089845 | 45.29 | 55 | 425223 |
| VP15198 | VP15198 | SAMN20353311 | SRR15219220 | NonPC | Public | 2015 | China | Env | 89.6 | 100 | 0 | 98.296 | 5125134 | 45.19 | 60 | 505329 |
| VP15199 | VP15199 | SAMN20353310 | SRR15219221 | NonPC | Public | 2015 | China | Clinical | 91.1 | 100 | 0.25 | 98.4155 | 5273997 | 45.05 | 43 | 371140 |
| VP15200 | VP15200 | SAMN20353307 | SRR15219225 | NonPC | Public | 2015 | China | Env | 88.5 | 99.98 | 0.3 | 98.2893 | 5109588 | 45.21 | 69 | 325526 |
| VP15201 | VP15201 | SAMN20353318 | SRR15219213 | NonPC | Public | 2015 | China | Clinical | 93.1 | 100 | 0.05 | 98.3502 | 5083268 | 45.29 | 48 | 424373 |
| VP15202 | VP15202 | SAMN20353320 | SRR15219210 | NonPC | Public | 2015 | China | Clinical | 93.1 | 100 | 0.19 | 98.3929 | 5075818 | 45.28 | 48 | 424378 |
| VP15203 | VP15203 | SAMN20353319 | SRR15219212 | NonPC | Public | 2015 | China | Clinical | 91.8 | 100 | 0.46 | 98.2932 | 5055023 | 45.21 | 60 | 450179 |
| VP15205 | VP15205 | SAMN20353321 | SRR15219209 | PC | Public | 2015 | China | Clinical | 97.3 | 100 | 0.07 | 99.9263 | 5150596 | 45.26 | 85 | 288527 |
| VP16001 | VP16001 | SAMN20353323 | SRR15219207 | PC | Public | 2016 | China | Clinical | 98.1 | 100 | 0 | 99.989 | 5106919 | 45.32 | 49 | 416372 |
| VP16003 | VP16003 | SAMN20353325 | SRR15219204 | NonPC | Public | 2016 | China | Clinical | 93.1 | 100 | 0.05 | 98.4021 | 5077383 | 45.29 | 44 | 463540 |
| VP16004 | VP16004 | SAMN20353324 | SRR15219206 | NonPC | Public | 2016 | China | Clinical | 91.8 | 100 | 0 | 98.4026 | 5065439 | 45.27 | 91 | 262004 |
| VP16005 | VP16005 | SAMN20371537 | SRR15239978 | PC | Public | 2016 | China | Clinical | 98.5 | 100 | 0 | 99.983 | 5093018 | 45.29 | 57 | 425301 |
| VP16006 | VP16006 | SAMN20371538 | SRR15239862 | NonPC | Public | 2016 | China | Clinical | 91.3 | 100 | 0.05 | 98.3368 | 4971003 | 45.34 | 47 | 266710 |
| VP16007 | VP16007 | SAMN20371539 | SRR15239747 | PC | Public | 2016 | China | Clinical | 98.1 | 100 | 0 | 99.9821 | 5106422 | 45.31 | 47 | 401580 |
| VP16008 | VP16008 | SAMN20371540 | SRR15240141 | PC | Public | 2016 | China | Clinical | 98.4 | 100 | 0 | 99.9839 | 5130952 | 45.34 | 57 | 401471 |
| VP16009 | VP16009 | SAMN20371541 | SRR15240020 | NonPC | Public | 2016 | China | Clinical | 91.8 | 100 | 0.46 | 98.2301 | 5060341 | 45.2 | 62 | 314098 |
| VP16012 | VP16012 | SAMN20371542 | SRR15240009 | NonPC | Public | 2016 | China | Clinical | 92.9 | 100 | 0.05 | 98.3923 | 5029415 | 45.32 | 43 | 463552 |
| VP16013 | VP16013 | SAMN20371543 | SRR15239998 | PC | Public | 2016 | China | Clinical | 98.4 | 100 | 0 | 99.9822 | 5088392 | 45.29 | 58 | 425307 |
| VP16014 | VP16014 | SAMN20371550 | SRR15240313 | PC | Public | 2016 | China | Clinical | 97.9 | 100 | 0 | 99.9475 | 5164268 | 45.28 | 48 | 401597 |

| | | | | | | | | | | | | | | | | |
|---|---|---|---|---|---|---|---|---|---|---|---|---|---|---|---|---|
| VP16015 | VP16015 | SAMN20371549 | SRR15240324 | PC | Public | 2016 | China | Clinical | 98.3 | 100 | 0 | 99.9801 | 5114015 | 45.3 | 44 | 401597 |
| VP16016 | VP16016 | SAMN20371551 | SRR15240266 | PC | Public | 2016 | China | Clinical | 98.4 | 100 | 0 | 99.9823 | 5091021 | 45.29 | 60 | 394915 |
| VP16017 | VP16017 | SAMN20371552 | SRR15240255 | PC | Public | 2016 | China | Clinical | 98.5 | 100 | 0 | 99.9887 | 5092397 | 45.29 | 55 | 425301 |
| VP16018 | VP16018 | SAMN20371547 | SRR15240346 | NonPC | Public | 2016 | China | Clinical | 92 | 100 | 0.46 | 98.2572 | 5072600 | 45.2 | 52 | 450300 |
| VP16019 | VP16019 | SAMN20371544 | SRR15239772 | NonPC | Public | 2016 | China | Clinical | 92 | 100 | 0.46 | 98.2687 | 5073619 | 45.2 | 50 | 450300 |
| VP16020 | VP16020 | SAMN20371545 | SRR15239725 | NonPC | Public | 2016 | China | Clinical | 92 | 100 | 0.46 | 98.2586 | 5074508 | 45.2 | 54 | 450397 |
| VP16021 | VP16021 | SAMN20371546 | SRR15240357 | NonPC | Public | 2016 | China | Clinical | 92 | 100 | 0.46 | 98.2666 | 5073899 | 45.2 | 53 | 450397 |
| VP16022 | VP16022 | SAMN20371548 | SRR15240335 | NonPC | Public | 2016 | China | Clinical | 92 | 100 | 0.46 | 98.3206 | 5074757 | 45.2 | 53 | 450116 |
| VP16024 | VP16024 | SAMN20371553 | SRR15240244 | PC | Public | 2016 | China | Clinical | 98.3 | 100 | 0 | 99.9769 | 5112256 | 45.3 | 45 | 435097 |
| VP16025 | VP16025 | SAMN20371557 | SRR15240091 | PC | Public | 2016 | China | Clinical | 97.5 | 100 | 0 | 99.9227 | 5144836 | 45.13 | 74 | 269050 |
| VP16026 | VP16026 | SAMN20371556 | SRR15240174 | PC | Public | 2016 | China | Clinical | 97.9 | 100 | 0 | 99.9526 | 5163700 | 45.28 | 46 | 401597 |
| VP16027 | VP16027 | SAMN20371555 | SRR15240185 | PC | Public | 2016 | China | Clinical | 97.9 | 100 | 0 | 99.9524 | 5163756 | 45.28 | 49 | 394980 |
| VP16028 | VP16028 | SAMN20371563 | SRR15239989 | NonPC | Public | 2016 | China | Clinical | 91.8 | 100 | 0.46 | 98.2581 | 5061473 | 45.2 | 59 | 313978 |
| VP16030 | VP16030 | SAMN20371554 | SRR15240196 | PC | Public | 2016 | China | Clinical | 97.9 | 100 | 0 | 99.9549 | 5161946 | 45.27 | 47 | 401597 |
| VP16031 | VP16031 | SAMN20371562 | SRR15240036 | PC | Public | 2016 | China | Clinical | 97.9 | 100 | 0 | 99.9532 | 5162435 | 45.27 | 43 | 401597 |
| VP16033 | VP16033 | SAMN20371565 | SRR15239966 | PC | Public | 2016 | China | Clinical | 98.4 | 100 | 0 | 99.9817 | 5093338 | 45.29 | 60 | 401241 |
| VP16042 | VP16042 | SAMN20371559 | SRR15240069 | PC | Public | 2016 | China | Clinical | 98.4 | 100 | 0 | 99.9922 | 5092614 | 45.29 | 58 | 401470 |
| VP16043 | VP16043 | SAMN20371561 | SRR15240047 | PC | Public | 2016 | China | Clinical | 96.7 | 99.18 | 0 | 99.9913 | 5004453 | 45.29 | 51 | 425839 |
| VP16044 | VP16044 | SAMN20371560 | SRR15240058 | PC | Public | 2016 | China | Clinical | 98.5 | 100 | 0 | 99.9829 | 5093302 | 45.29 | 57 | 425725 |
| VP16046 | VP16046 | SAMN20371558 | SRR15240080 | PC | Public | 2016 | China | Clinical | 98.4 | 100 | 0 | 99.9835 | 5093123 | 45.29 | 56 | 425725 |
| VP16048 | VP16048 | SAMN20371564 | SRR15239977 | NonPC | Public | 2016 | China | Clinical | 91.8 | 100 | 0.46 | 98.2258 | 5059746 | 45.2 | 64 | 313890 |
| VP16049 | VP16049 | SAMN20371566 | SRR15239955 | NonPC | Public | 2016 | China | Clinical | 92.2 | 100 | 0.19 | 98.3463 | 5027759 | 45.24 | 54 | 471204 |
| VP16050 | VP16050 | SAMN20371571 | SRR15239895 | PC | Public | 2016 | China | Clinical | 98.4 | 100 | 0 | 99.9799 | 5089611 | 45.29 | 58 | 425457 |
| VP16051 | VP16051 | SAMN20371569 | SRR15239922 | PC | Public | 2016 | China | Clinical | 98.4 | 100 | 0 | 99.9909 | 5094943 | 45.28 | 57 | 425349 |
| VP16053 | VP16053 | SAMN20371567 | SRR15239944 | PC | Public | 2016 | China | Clinical | 97.4 | 100 | 0 | 99.9172 | 5093553 | 45.27 | 79 | 205045 |
| VP16054 | VP16054 | SAMN20371568 | SRR15239933 | PC | Public | 2016 | China | Clinical | 98.5 | 100 | 0 | 99.9798 | 5099908 | 45.29 | 57 | 425265 |
| VP16055 | VP16055 | SAMN20371575 | SRR15239850 | PC | Public | 2016 | China | Clinical | 97.9 | 100 | 0 | 99.9684 | 5139811 | 45.28 | 47 | 394629 |
| VP16056 | VP16056 | SAMN20371576 | SRR15239839 | PC | Public | 2016 | China | Clinical | 98.4 | 100 | 0 | 99.982 | 5091370 | 45.29 | 58 | 401604 |
| VP16057 | VP16057 | SAMN20371577 | SRR15239828 | PC | Public | 2016 | China | Clinical | 98.4 | 100 | 0 | 99.9814 | 5092030 | 45.29 | 60 | 425313 |
| VP16058 | VP16058 | SAMN20371573 | SRR15239873 | NonPC | Public | 2016 | China | Clinical | 88.3 | 100 | 0.36 | 98.2472 | 5130803 | 45.22 | 115 | 414498 |
| VP16059 | VP16059 | SAMN20371580 | SRR15239795 | PC | Public | 2016 | China | Clinical | 97.9 | 100 | 0 | 99.9669 | 5162247 | 45.27 | 47 | 442213 |
| VP16061 | VP16061 | SAMN20371572 | SRR15239884 | PC | Public | 2016 | China | Clinical | 97.9 | 100 | 0 | 99.9766 | 5162514 | 45.27 | 50 | 394980 |
| VP16062 | VP16062 | SAMN20371574 | SRR15239861 | PC | Public | 2016 | China | Clinical | 98.5 | 100 | 0 | 99.9811 | 5091427 | 45.29 | 56 | 425187 |
| VP16064 | VP16064 | SAMN20371579 | SRR15239806 | NonPC | Public | 2016 | China | Clinical | 92.3 | 100 | 0.19 | 98.3602 | 5026904 | 45.23 | 50 | 539189 |
| VP16065 | VP16065 | SAMN20371581 | SRR15239784 | NonPC | Public | 2016 | China | Clinical | 93.1 | 100 | 0.05 | 98.3551 | 5070514 | 45.31 | 45 | 419561 |
| VP16066 | VP16066 | SAMN20371582 | SRR15239769 | NonPC | Public | 2016 | China | Clinical | 92.3 | 100 | 0.19 | 98.3749 | 5027364 | 45.23 | 50 | 539198 |
| VP16067 | VP16067 | SAMN20371583 | SRR15239758 | PC | Public | 2016 | China | Clinical | 97.9 | 100 | 0 | 99.9609 | 5138839 | 45.28 | 44 | 401463 |
| VP16068 | VP16068 | SAMN20371578 | SRR15239817 | NonPC | Public | 2016 | China | Clinical | 93 | 100 | 0.05 | 98.3791 | 5083987 | 45.28 | 50 | 419392 |
| VP16069 | VP16069 | SAMN20371584 | SRR15239746 | PC | Public | 2016 | China | Clinical | 98.5 | 100 | 0 | 99.9816 | 5092558 | 45.29 | 56 | 425307 |
| VP16070 | VP16070 | SAMN20371585 | SRR15239734 | PC | Public | 2016 | China | Clinical | 98.4 | 100 | 0 | 99.9897 | 5091652 | 45.29 | 59 | 425301 |
| VP16071 | VP16071 | SAMN20371586 | SRR15240295 | PC | Public | 2016 | China | Clinical | 98 | 100 | 0 | 99.9916 | 5080009 | 45.28 | 56 | 401604 |
| VP16073 | VP16073 | SAMN20371587 | SRR15240284 | PC | Public | 2016 | China | Clinical | 98.4 | 100 | 0 | 99.9918 | 5089792 | 45.29 | 57 | 425733 |
| VP16074 | VP16074 | SAMN20371590 | SRR15240218 | PC | Public | 2016 | China | Clinical | 98.4 | 100 | 0 | 99.9879 | 5090385 | 45.29 | 57 | 425337 |
| VP16075 | VP16075 | SAMN20371588 | SRR15240273 | PC | Public | 2016 | China | Clinical | 98.4 | 100 | 0 | 99.9893 | 5092136 | 45.29 | 55 | 425193 |
| VP16076 | VP16076 | SAMN20371570 | SRR15239906 | NonPC | Public | 2016 | China | Clinical | 91.8 | 100 | 0.46 | 98.2675 | 5062253 | 45.2 | 57 | 314248 |
| VP16085 | VP16085 | SAMN20371591 | SRR15240207 | NonPC | Public | 2016 | China | Clinical | 89 | 100 | 0 | 98.3395 | 4915376 | 45.37 | 29 | 773550 |
| VP16090 | VP16090 | SAMN20371589 | SRR15240229 | NonPC | Public | 2016 | China | Clinical | 91 | 100 | 0.77 | 98.3246 | 5045551 | 45.3 | 58 | 414232 |
| VP16092 | VP16092 | SAMN20371597 | SRR15240107 | PC | Public | 2016 | China | Clinical | 98.4 | 100 | 0 | 99.9867 | 5090509 | 45.29 | 55 | 425337 |

| | | | | | | | | | | | | | | | | |
|---|---|---|---|---|---|---|---|---|---|---|---|---|---|---|---|---|
| VP16093 | VP16093 | SAMN20371598 | SRR15240096 | NonPC | Public | 2016 | China | Clinical | 90.9 | 100 | 3.49 | 98.258 | 5286382 | 44.96 | 65 | 405028 |
| VP16095 | VP16095 | SAMN20371599 | SRR15240025 | PC | Public | 2016 | China | Clinical | 98.4 | 100 | 0 | 99.979 | 5090108 | 45.29 | 56 | 425871 |
| VP16096 | VP16096 | SAMN20371600 | SRR15240024 | NonPC | Public | 2016 | China | Clinical | 91.9 | 100 | 0.46 | 98.2562 | 5074462 | 45.2 | 50 | 450397 |
| VP16097 | VP16097 | SAMN20371601 | SRR15240023 | NonPC | Public | 2016 | China | Clinical | 92 | 100 | 0.46 | 98.2667 | 5074388 | 45.2 | 49 | 450301 |
| VP16098 | VP16098 | SAMN20371602 | SRR15240022 | NonPC | Public | 2016 | China | Clinical | 91.8 | 100 | 0.46 | 98.2637 | 5060891 | 45.2 | 58 | 314248 |
| VP16099 | VP16099 | SAMN20371603 | SRR15240021 | PC | Public | 2016 | China | Clinical | 98 | 100 | 0 | 99.9831 | 5079867 | 45.28 | 57 | 401604 |
| VP16100 | VP16100 | SAMN20371604 | SRR15240019 | PC | Public | 2016 | China | Clinical | 98.4 | 100 | 0 | 99.9807 | 5091373 | 45.29 | 56 | 425337 |
| VP16101 | VP16101 | SAMN20371606 | SRR15240017 | PC | Public | 2016 | China | Clinical | 98.5 | 100 | 0 | 99.9896 | 5091712 | 45.29 | 59 | 401604 |
| VP16102 | VP16102 | SAMN20371596 | SRR15240118 | PC | Public | 2016 | China | Clinical | 98.5 | 100 | 0 | 99.9804 | 5092671 | 45.29 | 54 | 425307 |
| VP16103 | VP16103 | SAMN20371593 | SRR15240152 | PC | Public | 2016 | China | Clinical | 98.4 | 100 | 0 | 99.9905 | 5096381 | 45.29 | 55 | 425319 |
| VP16104 | VP16104 | SAMN20371594 | SRR15240140 | PC | Public | 2016 | China | Clinical | 98.5 | 100 | 0 | 99.9804 | 5091740 | 45.29 | 55 | 425301 |
| VP16106 | VP16106 | SAMN20371608 | SRR15240015 | NonPC | Public | 2016 | China | Clinical | 91.8 | 100 | 0.46 | 98.2435 | 5061510 | 45.2 | 60 | 313844 |
| VP16107 | VP16107 | SAMN20371607 | SRR15240016 | PC | Public | 2016 | China | Clinical | 97.3 | 100 | 0.01 | 99.9077 | 5284293 | 45.27 | 72 | 312475 |
| VP16111 | VP16111 | SAMN20371592 | SRR15240163 | NonPC | Public | 2016 | China | Clinical | 91.8 | 100 | 0.46 | 98.2533 | 5062291 | 45.2 | 62 | 313980 |
| VP16112 | VP16112 | SAMN20371605 | SRR15240018 | PC | Public | 2016 | China | Clinical | 98 | 100 | 0 | 99.9917 | 5081126 | 45.28 | 58 | 401738 |
| VP16113 | VP16113 | SAMN20371629 | SRR15239920 | NonPC | Public | 2016 | China | Clinical | 92.3 | 100 | 0.19 | 98.361 | 5029001 | 45.24 | 52 | 471370 |
| VP16114 | VP16114 | SAMN20371630 | SRR15239919 | NonPC | Public | 2016 | China | Clinical | 92.1 | 100 | 0.19 | 98.3642 | 5014078 | 45.24 | 59 | 367943 |
| VP16115 | VP16115 | SAMN20371627 | SRR15239994 | NonPC | Public | 2016 | China | Clinical | 92.3 | 100 | 0.19 | 98.336 | 5028822 | 45.24 | 52 | 539189 |
| VP16116 | VP16116 | SAMN20371612 | SRR15240011 | PC | Public | 2016 | China | Clinical | 98.3 | 100 | 0 | 99.9802 | 5084350 | 45.29 | 62 | 401604 |
| VP16117 | VP16117 | SAMN20371613 | SRR15240010 | PC | Public | 2016 | China | Clinical | 98.5 | 100 | 0 | 99.9812 | 5169890 | 45.13 | 60 | 401604 |
| VP16119 | VP16119 | SAMN20371639 | SRR15239730 | PC | Public | 2016 | China | Clinical | 98.5 | 100 | 0 | 99.9778 | 5170517 | 45.13 | 59 | 425187 |
| VP16122 | VP16122 | SAMN20371595 | SRR15240129 | PC | Public | 2016 | China | Clinical | 98.5 | 100 | 0 | 99.9796 | 5093724 | 45.29 | 57 | 425415 |
| VP16124 | VP16124 | SAMN20371611 | SRR15240012 | PC | Public | 2016 | China | Clinical | 98.5 | 100 | 0 | 99.9825 | 5092865 | 45.29 | 58 | 425354 |
| VP16125 | VP16125 | SAMN20371610 | SRR15240013 | NonPC | Public | 2016 | China | Clinical | 92.3 | 100 | 0.19 | 98.353 | 5066054 | 45.23 | 53 | 450300 |
| VP16128 | VP16128 | SAMN20371636 | SRR15239733 | NonPC | Public | 2016 | China | Clinical | 92.3 | 100 | 0.19 | 98.3623 | 5027870 | 45.24 | 51 | 539189 |
| VP16129 | VP16129 | SAMN20371637 | SRR15239732 | NonPC | Public | 2016 | China | Clinical | 92.3 | 100 | 0.19 | 98.3415 | 5029576 | 45.24 | 50 | 539189 |
| VP16130 | VP16130 | SAMN20371638 | SRR15239731 | NonPC | Public | 2016 | China | Clinical | 92.3 | 100 | 0.19 | 98.3475 | 5027972 | 45.24 | 50 | 539189 |
| VP16131 | VP16131 | SAMN20371628 | SRR15239992 | NonPC | Public | 2016 | China | Clinical | 92.3 | 100 | 0.19 | 98.3588 | 5027602 | 45.24 | 51 | 539189 |
| VP16132 | VP16132 | SAMN20371640 | SRR15239729 | PC | Public | 2016 | China | Clinical | 98 | 100 | 0 | 99.9919 | 5081019 | 45.28 | 61 | 401604 |
| VP16133 | VP16133 | SAMN20371642 | SRR15239727 | PC | Public | 2016 | China | Clinical | 98.5 | 100 | 0 | 99.9808 | 5092810 | 45.29 | 57 | 401471 |
| VP16134 | VP16134 | SAMN20371645 | SRR15240366 | NonPC | Public | 2016 | China | Clinical | 92.3 | 100 | 0.19 | 98.3544 | 5027086 | 45.24 | 51 | 471206 |
| VP16135 | VP16135 | SAMN20371644 | SRR15239724 | NonPC | Public | 2016 | China | Clinical | 88.7 | 100 | 0.05 | 98.2916 | 5129270 | 45.26 | 59 | 242001 |
| VP16137 | VP16137 | SAMN20371646 | SRR15240365 | PC | Public | 2016 | China | Clinical | 98.4 | 100 | 0 | 99.9869 | 5090388 | 45.29 | 58 | 401590 |
| VP16138 | VP16138 | SAMN20371647 | SRR15240364 | NonPC | Public | 2016 | China | Clinical | 92.3 | 100 | 0.19 | 98.3375 | 5083789 | 45.22 | 51 | 471325 |
| VP16139 | VP16139 | SAMN20371649 | SRR15240362 | NonPC | Public | 2016 | China | Clinical | 93 | 100 | 0.05 | 98.393 | 5053290 | 45.3 | 55 | 417623 |
| VP16140 | VP16140 | SAMN20371614 | SRR15240008 | NonPC | Public | 2016 | China | Clinical | 93 | 100 | 0.19 | 98.362 | 5086963 | 45.27 | 53 | 417624 |
| VP16141 | VP16141 | SAMN20371615 | SRR15240007 | PC | Public | 2016 | China | Clinical | 98.4 | 100 | 0 | 99.9817 | 5089233 | 45.29 | 59 | 401456 |
| VP16142 | VP16142 | SAMN20371616 | SRR15240006 | PC | Public | 2016 | China | Clinical | 98.4 | 100 | 0 | 99.979 | 5090789 | 45.29 | 56 | 425457 |
| VP16144 | VP16144 | SAMN20371618 | SRR15240004 | PC | Public | 2016 | China | Clinical | 98.4 | 100 | 0 | 99.9776 | 5090422 | 45.29 | 64 | 334348 |
| VP16145 | VP16145 | SAMN20371620 | SRR15240002 | PC | Public | 2016 | China | Clinical | 98.4 | 100 | 0 | 99.9812 | 5091186 | 45.29 | 59 | 425337 |
| VP16146 | VP16146 | SAMN20371617 | SRR15240005 | PC | Public | 2016 | China | Clinical | 98.4 | 100 | 0 | 99.9797 | 5090919 | 45.29 | 60 | 394956 |
| VP16147 | VP16147 | SAMN20371619 | SRR15240003 | PC | Public | 2016 | China | Clinical | 98.4 | 100 | 0 | 99.9901 | 5090580 | 45.29 | 59 | 394812 |
| VP16148 | VP16148 | SAMN20371624 | SRR15239997 | PC | Public | 2016 | China | Clinical | 98.4 | 100 | 0 | 99.9905 | 5090085 | 45.29 | 65 | 312812 |
| VP16149 | VP16149 | SAMN20371625 | SRR15239996 | PC | Public | 2016 | China | Clinical | 98.4 | 100 | 0 | 99.9772 | 5089405 | 45.29 | 57 | 425337 |
| VP16150 | VP16150 | SAMN20371621 | SRR15240001 | PC | Public | 2016 | China | Clinical | 98.4 | 100 | 0 | 99.9897 | 5090952 | 45.29 | 59 | 394637 |
| VP16151 | VP16151 | SAMN20371626 | SRR15239995 | PC | Public | 2016 | China | Clinical | 98.4 | 99.97 | 0 | 99.9809 | 5090305 | 45.29 | 59 | 394954 |
| VP16152 | VP16152 | SAMN20371623 | SRR15239999 | PC | Public | 2016 | China | Clinical | 98.4 | 100 | 0 | 99.9813 | 5090761 | 45.29 | 62 | 394895 |
| VP16153 | VP16153 | SAMN20371622 | SRR15240000 | PC | Public | 2016 | China | Clinical | 98.4 | 100 | 0 | 99.9785 | 5139196 | 45.3 | 59 | 401590 |

| | | | | | | | | | | | | | | | | |
|---|---|---|---|---|---|---|---|---|---|---|---|---|---|---|---|---|
| VP16154 | VP16154 | SAMN20371609 | SRR15240014 | PC | Public | 2016 | China | Clinical | 98.4 | 100 | 0 | 99.9894 | 5090500 | 45.29 | 57 | 401590 |
| VP16155 | VP16155 | SAMN20371633 | SRR15239915 | NonPC | Public | 2016 | China | Clinical | 92.3 | 100 | 0.19 | 98.3705 | 5027147 | 45.24 | 51 | 539189 |
| VP16156 | VP16156 | SAMN20371634 | SRR15239771 | NonPC | Public | 2016 | China | Clinical | 92.3 | 100 | 0.19 | 98.3563 | 5038299 | 45.23 | 54 | 372958 |
| VP16157 | VP16157 | SAMN20371635 | SRR15239735 | NonPC | Public | 2016 | China | Clinical | 92.2 | 100 | 0.19 | 98.3561 | 5037925 | 45.23 | 58 | 314055 |
| VP16158 | VP16158 | SAMN20371632 | SRR15239917 | PC | Public | 2016 | China | Clinical | 98.4 | 100 | 0 | 99.9801 | 5091721 | 45.29 | 60 | 395118 |
| VP16159 | VP16159 | SAMN20371657 | SRR15240353 | PC | Public | 2016 | China | Clinical | 98.1 | 100 | 0 | 99.9906 | 5107411 | 45.32 | 60 | 312784 |
| VP16160 | VP16160 | SAMN20371660 | SRR15240350 | PC | Public | 2016 | China | Clinical | 98.1 | 100 | 0 | 99.9828 | 5107456 | 45.32 | 52 | 401580 |
| VP16161 | VP16161 | SAMN20371658 | SRR15240352 | PC | Public | 2016 | China | Clinical | 98.1 | 100 | 0 | 99.9829 | 5108230 | 45.32 | 59 | 288863 |
| VP16162 | VP16162 | SAMN20371661 | SRR15240349 | PC | Public | 2016 | China | Clinical | 98.1 | 100 | 0 | 99.9804 | 5108285 | 45.32 | 52 | 394782 |
| VP16163 | VP16163 | SAMN20371659 | SRR15240351 | PC | Public | 2016 | China | Clinical | 97.8 | 100 | 0 | 99.9926 | 5087660 | 45.3 | 70 | 196516 |
| VP16164 | VP16164 | SAMN20371656 | SRR15240354 | PC | Public | 2016 | China | Clinical | 98.1 | 100 | 0 | 99.9889 | 5107079 | 45.32 | 52 | 401580 |
| VP16165 | VP16165 | SAMN20371650 | SRR15240361 | NonPC | Public | 2016 | China | Clinical | 91.9 | 100 | 0.46 | 98.2657 | 5073623 | 45.2 | 52 | 450397 |
| VP16166 | VP16166 | SAMN20371641 | SRR15239728 | NonPC | Public | 2016 | China | Clinical | 92 | 100 | 1.17 | 98.2399 | 5175370 | 45.2 | 46 | 450300 |
| VP16167 | VP16167 | SAMN20371652 | SRR15240359 | NonPC | Public | 2016 | China | Clinical | 91.9 | 100 | 0.46 | 98.243 | 5070962 | 45.2 | 42 | 544882 |
| VP16168 | VP16168 | SAMN20371662 | SRR15240348 | PC | Public | 2016 | China | Clinical | 98.5 | 100 | 0 | 99.9899 | 5093572 | 45.29 | 58 | 425307 |
| VP16169 | VP16169 | SAMN20371653 | SRR15240358 | NonPC | Public | 2016 | China | Clinical | 92.2 | 100 | 0.19 | 98.3222 | 5038558 | 45.23 | 57 | 361536 |
| VP16170 | VP16170 | SAMN20371655 | SRR15240355 | PC | Public | 2016 | China | Clinical | 98 | 100 | 0 | 99.98 | 5088232 | 45.27 | 58 | 401604 |
| VP16171 | VP16171 | SAMN20371651 | SRR15240360 | NonPC | Public | 2016 | China | Clinical | 92.2 | 100 | 0.19 | 98.353 | 5026815 | 45.24 | 51 | 539189 |
| VP16172 | VP16172 | SAMN20371654 | SRR15240356 | NonPC | Public | 2016 | China | Clinical | 92.3 | 100 | 0.19 | 98.3568 | 5039622 | 45.23 | 58 | 361536 |
| VP16174 | VP16174 | SAMN20371664 | SRR15240345 | NonPC | Public | 2016 | China | Clinical | 92.3 | 100 | 0.19 | 98.3665 | 5027909 | 45.24 | 50 | 471218 |
| VP16175 | VP16175 | SAMN20371663 | SRR15240347 | PC | Public | 2016 | China | Clinical | 97.8 | 100 | 0 | 99.9216 | 5167085 | 45.24 | 69 | 373623 |
| VP16177 | VP16177 | SAMN20371631 | SRR15239918 | PC | Public | 2016 | China | Clinical | 98.4 | 100 | 0.18 | 99.9647 | 5108144 | 45.31 | 66 | 425337 |
| VP16178 | VP16178 | SAMN20371669 | SRR15240340 | PC | Public | 2016 | China | Clinical | 98.4 | 100 | 0 | 99.9882 | 5090555 | 45.29 | 57 | 425337 |
| VP16179 | VP16179 | SAMN20371670 | SRR15240339 | NonPC | Public | 2016 | China | Clinical | 92.3 | 100 | 0.19 | 98.379 | 5028595 | 45.24 | 53 | 471370 |
| VP16180 | VP16180 | SAMN20371673 | SRR15240336 | PC | Public | 2016 | China | Clinical | 98.4 | 100 | 0 | 99.9805 | 5093582 | 45.28 | 55 | 425235 |
| VP16181 | VP16181 | SAMN20371674 | SRR15240334 | PC | Public | 2016 | China | Clinical | 98.4 | 100 | 0 | 99.9898 | 5090488 | 45.29 | 55 | 425337 |
| VP16182 | VP16182 | SAMN20371672 | SRR15240337 | NonPC | Public | 2016 | China | Clinical | 91.6 | 100 | 0.27 | 98.3461 | 5301196 | 45.15 | 59 | 450341 |
| VP16183 | VP16183 | SAMN20371643 | SRR15239726 | PC | Public | 2016 | China | Clinical | 98.4 | 100 | 0 | 99.9901 | 5090226 | 45.29 | 61 | 425217 |
| VP16184 | VP16184 | SAMN20371666 | SRR15240343 | NonPC | Public | 2016 | China | Clinical | 93 | 100 | 0.05 | 98.3689 | 5074519 | 45.28 | 55 | 417624 |
| VP16185 | VP16185 | SAMN20371665 | SRR15240344 | NonPC | Public | 2016 | China | Clinical | 88.7 | 100 | 0.05 | 98.3404 | 4991527 | 45.22 | 67 | 247059 |
| VP16186 | VP16186 | SAMN20371648 | SRR15240363 | NonPC | Public | 2016 | China | Clinical | 93.1 | 100 | 0.05 | 98.3641 | 5076609 | 45.29 | 46 | 436101 |
| VP16187 | VP16187 | SAMN20371668 | SRR15240341 | PC | Public | 2016 | China | Clinical | 98.4 | 100 | 0 | 99.9799 | 5090294 | 45.29 | 57 | 401590 |
| VP16188 | VP16188 | SAMN20371667 | SRR15240342 | PC | Public | 2016 | China | Clinical | 98.5 | 100 | 0 | 99.9811 | 5099939 | 45.3 | 59 | 425301 |
| VP16189 | VP16189 | SAMN20371675 | SRR15240333 | PC | Public | 2016 | China | Clinical | 98.5 | 100 | 0 | 99.9935 | 5095546 | 45.3 | 61 | 394965 |
| VP16191 | VP16191 | SAMN20371671 | SRR15240338 | PC | Public | 2016 | China | Clinical | 98 | 100 | 0 | 99.955 | 5179088 | 45.26 | 49 | 394917 |
| VP16192 | VP16192 | SAMN20371676 | SRR15240332 | PC | Public | 2016 | China | Clinical | 98.6 | 100 | 0 | 99.9868 | 5148073 | 45.28 | 61 | 420398 |
| VP16193 | VP16193 | SAMN20371677 | SRR15240331 | PC | Public | 2016 | China | Clinical | 98 | 100 | 0 | 99.9836 | 5079871 | 45.28 | 59 | 401604 |
| VP16194 | VP16194 | SAMN20371681 | SRR15240327 | PC | Public | 2016 | China | Clinical | 98 | 100 | 0 | 99.9865 | 5069757 | 45.28 | 80 | 265886 |
| VP16195 | VP16195 | SAMN20371700 | SRR15240306 | NonPC | Public | 2016 | China | Clinical | 92.3 | 100 | 0.19 | 98.3538 | 5027491 | 45.24 | 53 | 471325 |
| VP16196 | VP16196 | SAMN20371680 | SRR15240328 | PC | Public | 2016 | China | Clinical | 98.5 | 100 | 0 | 99.9928 | 5098291 | 45.28 | 60 | 401597 |
| VP16197 | VP16197 | SAMN20371678 | SRR15240330 | PC | Public | 2016 | China | Clinical | 98 | 100 | 0 | 99.9634 | 5164428 | 45.28 | 46 | 394963 |
| VP16199 | VP16199 | SAMN20371683 | SRR15240325 | NonPC | Public | 2016 | China | Clinical | 92.3 | 100 | 0.19 | 98.3327 | 5044539 | 45.22 | 51 | 471325 |
| VP16200 | VP16200 | SAMN20371679 | SRR15240329 | PC | Public | 2016 | China | Clinical | 97.3 | 100 | 0.01 | 99.9015 | 5222385 | 45.26 | 74 | 259762 |
| VP16201 | VP16201 | SAMN20371682 | SRR15240326 | PC | Public | 2016 | China | Clinical | 98 | 100 | 0 | 99.9938 | 5082492 | 45.28 | 60 | 442533 |
| VP16204 | VP16204 | SAMN20371685 | SRR15240322 | NonPC | Public | 2016 | China | Clinical | 91.9 | 100 | 0.19 | 98.3607 | 5028278 | 45.24 | 53 | 471204 |
| VP16205 | VP16205 | SAMN20371686 | SRR15240321 | PC | Public | 2016 | China | Clinical | 98.3 | 100 | 0 | 99.981 | 5083425 | 45.29 | 56 | 425307 |
| VP16206 | VP16206 | SAMN20371684 | SRR15240323 | NonPC | Public | 2016 | China | Clinical | 92.3 | 100 | 0.19 | 98.3601 | 5026798 | 45.24 | 53 | 471325 |
| VP16207 | VP16207 | SAMN20371687 | SRR15240320 | NonPC | Public | 2016 | China | Clinical | 91.8 | 100 | 0.46 | 98.2412 | 5060704 | 45.2 | 60 | 439588 |

| | | | | | | | | | | | | | | | | |
|---|---|---|---|---|---|---|---|---|---|---|---|---|---|---|---|---|
| VP16208 | VP16208 | SAMN20371695 | SRR15240311 | PC | Public | 2016 | China | Clinical | 97.9 | 100 | 0 | 99.9631 | 5172146 | 45.28 | 46 | 418635 |
| VP16209 | VP16209 | SAMN20371688 | SRR15240319 | NonPC | Public | 2016 | China | Clinical | 88.6 | 100 | 0.48 | 98.2254 | 5095913 | 45.16 | 90 | 321756 |
| VP16210 | VP16210 | SAMN20353322 | SRR15219208 | NonPC | Public | 2016 | China | Clinical | 92.7 | 100 | 0 | 98.4013 | 5138398 | 45.16 | 57 | 258836 |
| VP16211 | VP16211 | SAMN20371697 | SRR15240309 | PC | Public | 2016 | China | Clinical | 98.5 | 100 | 0 | 99.9898 | 5093187 | 45.29 | 59 | 401604 |
| VP16212 | VP16212 | SAMN20371698 | SRR15240308 | PC | Public | 2016 | China | Clinical | 98.5 | 100 | 0 | 99.9818 | 5094142 | 45.29 | 60 | 425307 |
| VP16213 | VP16213 | SAMN20371699 | SRR15240307 | PC | Public | 2016 | China | Clinical | 98.5 | 100 | 0 | 99.9799 | 5094777 | 45.29 | 59 | 394963 |
| VP16214 | VP16214 | SAMN20371696 | SRR15240310 | NonPC | Public | 2016 | China | Clinical | 88.8 | 100 | 0.05 | 98.3287 | 5001063 | 45.23 | 48 | 426163 |
| VP17001 | VP17001 | SAMN20371689 | SRR15240318 | NonPC | Public | 2017 | China | Clinical | 92 | 100 | 0.46 | 98.2512 | 5072921 | 45.2 | 55 | 450300 |
| VP17002 | VP17002 | SAMN20371702 | SRR15240268 | PC | Public | 2017 | China | Clinical | 98.3 | 100 | 0 | 99.9912 | 5130099 | 45.26 | 80 | 267776 |
| VP17003 | VP17003 | SAMN20371703 | SRR15240267 | NonPC | Public | 2017 | China | Clinical | 92 | 100 | 0.19 | 98.2832 | 5049250 | 45.23 | 50 | 544888 |
| VP17004 | VP17004 | SAMN20371701 | SRR15240270 | NonPC | Public | 2017 | China | Clinical | 90.7 | 100 | 0.33 | 98.2514 | 5159470 | 45.12 | 79 | 238308 |
| VP17007 | VP17007 | SAMN20371704 | SRR15240265 | PC | Public | 2017 | China | Clinical | 97.2 | 100 | 0 | 99.9052 | 5093326 | 45.28 | 82 | 258237 |
| VP17008 | VP17008 | SAMN20371754 | SRR15240173 | PC | Public | 2017 | China | Clinical | 98.1 | 100 | 0.27 | 99.9737 | 5107730 | 45.32 | 52 | 409709 |
| VP17009 | VP17009 | SAMN20371753 | SRR15240175 | NonPC | Public | 2017 | China | Clinical | 88.7 | 100 | 0.05 | 98.3138 | 5010571 | 45.18 | 67 | 271491 |
| VP17010 | VP17010 | SAMN20371755 | SRR15240172 | PC | Public | 2017 | China | Clinical | 98.4 | 100 | 0.27 | 99.9949 | 5166351 | 45.35 | 61 | 425985 |
| VP17011 | VP17011 | SAMN20371758 | SRR15240169 | PC | Public | 2017 | China | Clinical | 98 | 100 | 0 | 99.9842 | 5080698 | 45.28 | 61 | 394850 |
| VP17012 | VP17012 | SAMN20371757 | SRR15240170 | PC | Public | 2017 | China | Clinical | 98 | 100 | 0 | 99.9931 | 5081602 | 45.28 | 62 | 442227 |
| VP17013 | VP17013 | SAMN20371761 | SRR15240094 | NonPC | Public | 2017 | China | Clinical | 92.3 | 100 | 0.05 | 98.3681 | 5117290 | 45.07 | 54 | 450182 |
| VP17014 | VP17014 | SAMN20371756 | SRR15240171 | NonPC | Public | 2017 | China | Clinical | 91.8 | 100 | 0.46 | 98.2784 | 5062450 | 45.2 | 58 | 450188 |
| VP17015 | VP17015 | SAMN20371760 | SRR15240095 | NonPC | Public | 2017 | China | Clinical | 91.8 | 100 | 0.46 | 98.2896 | 5062553 | 45.2 | 58 | 439604 |
| VP17016 | VP17016 | SAMN20371759 | SRR15240168 | PC | Public | 2017 | China | Clinical | 98 | 100 | 0 | 99.9831 | 5079704 | 45.28 | 55 | 401604 |
| VP17017 | VP17017 | SAMN20371762 | SRR15240093 | NonPC | Public | 2017 | China | Clinical | 91.8 | 100 | 0.19 | 98.2116 | 5049859 | 45.22 | 59 | 450187 |
| VP17018 | VP17018 | SAMN20371763 | SRR15240092 | PC | Public | 2017 | China | Clinical | 98.4 | 100 | 0.09 | 99.9819 | 5128856 | 45.38 | 60 | 401590 |
| VP17020 | VP17020 | SAMN20371766 | SRR15240088 | PC | Public | 2017 | China | Clinical | 97.9 | 100 | 0 | 99.9771 | 5171833 | 45.28 | 48 | 418635 |
| VP17022 | VP17022 | SAMN20371764 | SRR15240090 | NonPC | Public | 2017 | China | Clinical | 92.2 | 100 | 0.19 | 98.3946 | 5028365 | 45.24 | 51 | 539189 |
| VP17023 | VP17023 | SAMN20371768 | SRR15240086 | PC | Public | 2017 | China | Clinical | 98.4 | 100 | 0 | 99.9836 | 5090163 | 45.29 | 57 | 425343 |
| VP17024 | VP17024 | SAMN20371767 | SRR15240087 | PC | Public | 2017 | China | Clinical | 98.4 | 100 | 0 | 99.9903 | 5090726 | 45.29 | 55 | 425313 |
| VP17028 | VP17028 | SAMN20371803 | SRR15240048 | PC | Public | 2017 | China | Clinical | 98.4 | 100 | 0 | 99.9899 | 5090771 | 45.29 | 59 | 440609 |
| VP17029 | VP17029 | SAMN20371802 | SRR15240049 | PC | Public | 2017 | China | Clinical | 98.4 | 100 | 0 | 99.9935 | 5090842 | 45.29 | 56 | 401456 |
| VP17030 | VP17030 | SAMN20371769 | SRR15240085 | NonPC | Public | 2017 | China | Clinical | 93.1 | 100 | 0.05 | 98.3665 | 5076958 | 45.29 | 49 | 513716 |
| VP17031 | VP17031 | SAMN20371807 | SRR15240043 | PC | Public | 2017 | China | Clinical | 98.4 | 100 | 0 | 99.9917 | 5091018 | 45.29 | 55 | 425343 |
| VP17032 | VP17032 | SAMN20371815 | SRR15240034 | PC | Public | 2017 | China | Clinical | 98.4 | 100 | 0 | 99.9828 | 5090251 | 45.29 | 59 | 425343 |
| VP17033 | VP17033 | SAMN20371814 | SRR15240035 | PC | Public | 2017 | China | Clinical | 98.4 | 100 | 0 | 99.985 | 5090481 | 45.29 | 58 | 401456 |
| VP17034 | VP17034 | SAMN20371816 | SRR15240033 | PC | Public | 2017 | China | Clinical | 98 | 100 | 0 | 99.992 | 5081275 | 45.28 | 59 | 442235 |
| VP17035 | VP17035 | SAMN20371817 | SRR15240032 | PC | Public | 2017 | China | Clinical | 98 | 100 | 0 | 99.9927 | 5080936 | 45.28 | 61 | 395004 |
| VP17036 | VP17036 | SAMN20371810 | SRR15240040 | PC | Public | 2017 | China | Clinical | 98.4 | 100 | 0 | 99.9936 | 5090713 | 45.29 | 58 | 425457 |
| VP17037 | VP17037 | SAMN20371808 | SRR15240042 | PC | Public | 2017 | China | Clinical | 98.4 | 100 | 0.27 | 99.9913 | 5134716 | 45.29 | 57 | 425343 |
| VP17038 | VP17038 | SAMN20371809 | SRR15240041 | PC | Public | 2017 | China | Clinical | 98.4 | 100 | 0.54 | 99.9921 | 5145303 | 45.28 | 59 | 394711 |
| VP17039 | VP17039 | SAMN20371812 | SRR15240038 | PC | Public | 2017 | China | Clinical | 98.4 | 100 | 0.27 | 99.9844 | 5134322 | 45.29 | 62 | 312677 |
| VP17040 | VP17040 | SAMN20371765 | SRR15240089 | PC | Public | 2017 | China | Clinical | 98.1 | 100 | 0 | 99.9932 | 5071281 | 45.27 | 75 | 288535 |
| VP17041 | VP17041 | SAMN20371691 | SRR15240316 | PC | Public | 2017 | China | Clinical | 97.7 | 100 | 0 | 99.9935 | 5062734 | 45.27 | 76 | 288535 |
| VP17043 | VP17043 | SAMN20371821 | SRR15239993 | NonPC | Public | 2017 | China | Clinical | 92.1 | 100 | 0.19 | 98.3729 | 5016667 | 45.23 | 60 | 313845 |
| VP17044 | VP17044 | SAMN20371822 | SRR15239991 | PC | Public | 2017 | China | Clinical | 98.4 | 100 | 0 | 99.9807 | 5090083 | 45.29 | 58 | 396071 |
| VP17045 | VP17045 | SAMN20371820 | SRR15240029 | NonPC | Public | 2017 | China | Clinical | 91.8 | 100 | 0.74 | 98.2001 | 5104975 | 45.2 | 64 | 314098 |
| VP17046 | VP17046 | SAMN20371818 | SRR15240031 | PC | Public | 2017 | China | Clinical | 98.4 | 100 | 0.27 | 99.9832 | 5135455 | 45.29 | 59 | 425343 |
| VP17047 | VP17047 | SAMN20371819 | SRR15240030 | PC | Public | 2017 | China | Clinical | 98.4 | 100 | 0 | 99.9944 | 5091231 | 45.29 | 57 | 401590 |
| VP17049 | VP17049 | SAMN20371801 | SRR15240050 | PC | Public | 2017 | China | Clinical | 98.4 | 100 | 0 | 99.9826 | 5091364 | 45.29 | 58 | 425343 |
| VP17050 | VP17050 | SAMN20371813 | SRR15240037 | PC | Public | 2017 | China | Clinical | 98 | 100 | 0 | 99.9845 | 5080839 | 45.28 | 59 | 394852 |

| | | | | | | | | | | | | | | | | |
|---|---|---|---|---|---|---|---|---|---|---|---|---|---|---|---|---|
| VP17051 | VP17051 | SAMN20371823 | SRR15239990 | PC | Public | 2017 | China | Clinical | 98.5 | 100 | 0 | 99.9811 | 5092307 | 45.29 | 57 | 401094 |
| VP17052 | VP17052 | SAMN20371840 | SRR15239970 | PC | Public | 2017 | China | Clinical | 98.4 | 100 | 0.01 | 99.9931 | 5182531 | 45.31 | 55 | 425871 |
| VP17053 | VP17053 | SAMN20371845 | SRR15239964 | PC | Public | 2017 | China | Clinical | 98 | 100 | 0 | 99.9844 | 5081196 | 45.28 | 59 | 401470 |
| VP17054 | VP17054 | SAMN20371831 | SRR15239981 | PC | Public | 2017 | China | Clinical | 98.1 | 100 | 0.27 | 99.9919 | 5107846 | 45.32 | 49 | 409709 |
| VP17055 | VP17055 | SAMN20371843 | SRR15239967 | PC | Public | 2017 | China | Clinical | 98.4 | 100 | 0 | 99.9929 | 5096866 | 45.29 | 58 | 425343 |
| VP17056 | VP17056 | SAMN20371837 | SRR15239973 | NonPC | Public | 2017 | China | Clinical | 92.4 | 100 | 0.19 | 98.3355 | 5046763 | 45.24 | 46 | 399566 |
| VP17057 | VP17057 | SAMN20371846 | SRR15239963 | PC | Public | 2017 | China | Clinical | 98 | 100 | 0 | 99.9928 | 5080060 | 45.28 | 57 | 401604 |
| VP17058 | VP17058 | SAMN20371835 | SRR15239975 | NonPC | Public | 2017 | China | Clinical | 92.4 | 100 | 0.19 | 98.3155 | 5045673 | 45.24 | 47 | 407315 |
| VP17059 | VP17059 | SAMN20371833 | SRR15239979 | NonPC | Public | 2017 | China | Clinical | 92.4 | 100 | 0.19 | 98.341 | 5046655 | 45.24 | 49 | 407315 |
| VP17060 | VP17060 | SAMN20371844 | SRR15239965 | PC | Public | 2017 | China | Clinical | 98.4 | 100 | 0 | 99.9843 | 5089673 | 45.29 | 57 | 401590 |
| VP17061 | VP17061 | SAMN20371829 | SRR15239983 | NonPC | Public | 2017 | China | Clinical | 92.9 | 99.46 | 0.05 | 98.3578 | 5061740 | 45.3 | 49 | 419573 |
| VP17062 | VP17062 | SAMN20371849 | SRR15239960 | PC | Public | 2017 | China | Clinical | 98 | 100 | 0.01 | 99.9948 | 5215794 | 45.35 | 61 | 394852 |
| VP17063 | VP17063 | SAMN20371850 | SRR15239959 | PC | Public | 2017 | China | Clinical | 98 | 100 | 0 | 99.985 | 5080831 | 45.28 | 59 | 401604 |
| VP17064 | VP17064 | SAMN20371842 | SRR15239968 | PC | Public | 2017 | China | Clinical | 98.4 | 100 | 0 | 99.985 | 5090916 | 45.29 | 59 | 425343 |
| VP17065 | VP17065 | SAMN20371827 | SRR15239985 | NonPC | Public | 2017 | China | Clinical | 91.9 | 100 | 0.19 | 98.3342 | 5042631 | 45.24 | 52 | 471204 |
| VP17067 | VP17067 | SAMN20371834 | SRR15239976 | NonPC | Public | 2017 | China | Clinical | 92.4 | 100 | 0.19 | 98.3351 | 5046673 | 45.24 | 47 | 407315 |
| VP17068 | VP17068 | SAMN20371828 | SRR15239984 | NonPC | Public | 2017 | China | Clinical | 90.9 | 100 | 0 | 98.2864 | 5080325 | 45.26 | 62 | 405022 |
| VP17069 | VP17069 | SAMN20371836 | SRR15239974 | NonPC | Public | 2017 | China | Clinical | 92.4 | 100 | 0.19 | 98.3422 | 5046428 | 45.24 | 49 | 399699 |
| VP17070 | VP17070 | SAMN20371839 | SRR15239971 | NonPC | Public | 2017 | China | Clinical | 92.4 | 100 | 0.19 | 98.3322 | 5046846 | 45.24 | 48 | 407315 |
| VP17071 | VP17071 | SAMN20371841 | SRR15239969 | PC | Public | 2017 | China | Clinical | 98.4 | 100 | 0 | 99.9939 | 5090282 | 45.29 | 57 | 401590 |
| VP17073 | VP17073 | SAMN20371838 | SRR15239972 | NonPC | Public | 2017 | China | Clinical | 92.4 | 100 | 0.19 | 98.3163 | 5045653 | 45.24 | 47 | 407315 |
| VP17074 | VP17074 | SAMN20371848 | SRR15239961 | PC | Public | 2017 | China | Clinical | 98 | 100 | 0 | 99.9841 | 5080680 | 45.28 | 61 | 401604 |
| VP17076 | VP17076 | SAMN20371852 | SRR15239957 | PC | Public | 2017 | China | Clinical | 98 | 100 | 0 | 99.9824 | 5080301 | 45.28 | 59 | 401604 |
| VP17077 | VP17077 | SAMN20371851 | SRR15239958 | PC | Public | 2017 | China | Clinical | 97.9 | 100 | 0 | 99.9677 | 5129334 | 45.28 | 61 | 394981 |
| VP17078 | VP17078 | SAMN20371847 | SRR15239962 | PC | Public | 2017 | China | Clinical | 98 | 100 | 0 | 99.9831 | 5079629 | 45.28 | 55 | 401470 |
| VP17079 | VP17079 | SAMN20371832 | SRR15239980 | NonPC | Public | 2017 | China | Clinical | 92 | 100 | 0.46 | 98.2555 | 5074806 | 45.2 | 51 | 450300 |
| VP17080 | VP17080 | SAMN20371830 | SRR15239982 | NonPC | Public | 2017 | China | Env | 91.4 | 100 | 0 | 98.3374 | 5036709 | 45.34 | 66 | 426625 |
| VP17081 | VP17081 | SAMN20371854 | SRR15239954 | PC | Public | 2017 | China | Clinical | 98.4 | 100 | 0 | 99.9909 | 5094036 | 45.29 | 55 | 425313 |
| VP17082 | VP17082 | SAMN20371853 | SRR15239956 | NonPC | Public | 2017 | China | Clinical | 91.9 | 100 | 0.46 | 98.2419 | 5073211 | 45.2 | 53 | 450397 |
| VP17083 | VP17083 | SAMN20371859 | SRR15239949 | PC | Public | 2017 | China | Clinical | 97.9 | 100 | 0 | 99.9677 | 5131723 | 45.27 | 57 | 401597 |
| VP17084 | VP17084 | SAMN20371855 | SRR15239953 | NonPC | Public | 2017 | China | Clinical | 92 | 100 | 0.46 | 98.2544 | 5074049 | 45.2 | 49 | 450118 |
| VP17086 | VP17086 | SAMN20371826 | SRR15239986 | PC | Public | 2017 | China | Clinical | 98 | 100 | 0 | 99.9738 | 5099802 | 45.27 | 64 | 401470 |
| VP17087 | VP17087 | SAMN20371864 | SRR15239943 | PC | Public | 2017 | China | Clinical | 98.4 | 100 | 0 | 99.9938 | 5091017 | 45.29 | 56 | 425343 |
| VP17088 | VP17088 | SAMN20371862 | SRR15239946 | PC | Public | 2017 | China | Clinical | 98 | 100 | 0 | 99.9928 | 5080927 | 45.28 | 58 | 401604 |
| VP17089 | VP17089 | SAMN20371861 | SRR15239947 | PC | Public | 2017 | China | Clinical | 98.4 | 100 | 0.27 | 99.9829 | 5134177 | 45.29 | 58 | 425343 |
| VP17090 | VP17090 | SAMN20371863 | SRR15239945 | NonPC | Public | 2017 | China | Clinical | 92.8 | 100 | 0.19 | 98.3984 | 5065105 | 45.28 | 56 | 410056 |
| VP17091 | VP17091 | SAMN20371866 | SRR15239941 | PC | Public | 2017 | China | Clinical | 97.3 | 100 | 0 | 99.8997 | 5190987 | 45.24 | 67 | 312385 |
| VP17092 | VP17092 | SAMN20371865 | SRR15239942 | PC | Public | 2017 | China | Clinical | 97.5 | 100 | 0 | 99.9311 | 5147289 | 45.13 | 67 | 269183 |
| VP17093 | VP17093 | SAMN20371869 | SRR15239938 | PC | Public | 2017 | China | Clinical | 97.6 | 100 | 0 | 99.8989 | 5217169 | 45.25 | 60 | 373417 |
| VP17094 | VP17094 | SAMN20371867 | SRR15239940 | PC | Public | 2017 | China | Clinical | 97.6 | 100 | 0.01 | 99.9045 | 5299975 | 45.28 | 55 | 461741 |
| VP17095 | VP17095 | SAMN20371868 | SRR15239939 | PC | Public | 2017 | China | Clinical | 97.6 | 100 | 0 | 99.9144 | 5208751 | 45.26 | 55 | 319939 |
| VP17096 | VP17096 | SAMN20371860 | SRR15239948 | PC | Public | 2017 | China | Clinical | 98.4 | 100 | 0 | 99.9799 | 5090956 | 45.29 | 61 | 401590 |
| VP17097 | VP17097 | SAMN20371856 | SRR15239952 | PC | Public | 2017 | China | Clinical | 98.4 | 100 | 0.27 | 99.9916 | 5134204 | 45.29 | 61 | 425229 |
| VP17098 | VP17098 | SAMN20371824 | SRR15239988 | PC | Public | 2017 | China | Clinical | 98.4 | 100 | 0 | 99.9834 | 5090075 | 45.29 | 56 | 401590 |
| VP17099 | VP17099 | SAMN20371806 | SRR15240044 | NonPC | Public | 2017 | China | Clinical | 92.9 | 100 | 0.19 | 98.3408 | 5072019 | 45.27 | 54 | 417623 |
| VP17100 | VP17100 | SAMN20371825 | SRR15239987 | PC | Public | 2017 | China | Clinical | 98 | 100 | 0 | 99.984 | 5079249 | 45.28 | 56 | 401470 |
| VP17101 | VP17101 | SAMN20371857 | SRR15239951 | PC | Public | 2017 | China | Clinical | 98 | 100 | 0 | 99.9924 | 5080706 | 45.28 | 56 | 440631 |
| VP17102 | VP17102 | SAMN20371871 | SRR15239936 | PC | Public | 2017 | China | Clinical | 98.4 | 100 | 0 | 99.9845 | 5090194 | 45.29 | 59 | 401457 |

| | | | | | | | | | | | | | | | | |
|---|---|---|---|---|---|---|---|---|---|---|---|---|---|---|---|---|
| VP17103 | VP17103 | SAMN20371872 | SRR15239935 | PC | Public | 2017 | China | Clinical | 98.4 | 100 | 0.27 | 99.9913 | 5133362 | 45.29 | 58 | 425229 |
| VP17104 | VP17104 | SAMN20371874 | SRR15239932 | NonPC | Public | 2017 | China | Clinical | 91.8 | 100 | 0.19 | 98.2573 | 5052350 | 45.21 | 60 | 313845 |
| VP17105 | VP17105 | SAMN20371876 | SRR15239930 | PC | Public | 2017 | China | Clinical | 98.4 | 100 | 0 | 99.9826 | 5089952 | 45.29 | 57 | 401590 |
| VP17106 | VP17106 | SAMN20371878 | SRR15239928 | NonPC | Public | 2017 | China | Clinical | 91.9 | 100 | 0 | 98.3569 | 5016635 | 45.38 | 41 | 368305 |
| VP17108 | VP17108 | SAMN20371877 | SRR15239929 | PC | Public | 2017 | China | Clinical | 98 | 100 | 0.27 | 99.9915 | 5120294 | 45.27 | 80 | 265886 |
| VP17109 | VP17109 | SAMN20371873 | SRR15239934 | NonPC | Public | 2017 | China | Clinical | 92 | 100 | 0.46 | 98.2411 | 5075711 | 45.2 | 52 | 450397 |
| VP17112 | VP17112 | SAMN20371870 | SRR15239937 | PC | Public | 2017 | China | Clinical | 98 | 100 | 0 | 99.9921 | 5080215 | 45.28 | 61 | 401470 |
| VP17113 | VP17113 | SAMN20371875 | SRR15239931 | NonPC | Public | 2017 | China | Clinical | 91.8 | 100 | 0.46 | 98.237 | 5061833 | 45.2 | 57 | 450187 |
| VP17115 | VP17115 | SAMN20371883 | SRR15239923 | NonPC | Public | 2017 | China | Clinical | 90.7 | 100 | 0.33 | 98.3044 | 5065101 | 45.23 | 57 | 247072 |
| VP17116 | VP17116 | SAMN20371858 | SRR15239950 | PC | Public | 2017 | China | Clinical | 98 | 100 | 0 | 99.9924 | 5079983 | 45.28 | 60 | 401604 |
| VP17117 | VP17117 | SAMN20371913 | SRR15239885 | PC | Public | 2017 | China | Clinical | 98 | 100 | 0.27 | 99.9932 | 5113998 | 45.28 | 80 | 266000 |
| VP17118 | VP17118 | SAMN20371910 | SRR15239888 | PC | Public | 2017 | China | Clinical | 98 | 100 | 0.27 | 99.9889 | 5124703 | 45.28 | 62 | 394690 |
| VP17120 | VP17120 | SAMN20371907 | SRR15239891 | PC | Public | 2017 | China | Clinical | 98.4 | 100 | 0.01 | 99.9843 | 5189475 | 45.31 | 62 | 312684 |
| VP17121 | VP17121 | SAMN20371879 | SRR15239927 | PC | Public | 2017 | China | Clinical | 98.4 | 100 | 0 | 99.979 | 5094045 | 45.29 | 56 | 425343 |
| VP17122 | VP17122 | SAMN20371881 | SRR15239925 | PC | Public | 2017 | China | Clinical | 98.3 | 100 | 0 | 99.9823 | 5096107 | 45.28 | 58 | 440623 |
| VP17123 | VP17123 | SAMN20371908 | SRR15239890 | PC | Public | 2017 | China | Clinical | 98.4 | 100 | 0 | 99.9938 | 5090088 | 45.29 | 56 | 425343 |
| VP17125 | VP17125 | SAMN20371915 | SRR15239882 | PC | Public | 2017 | China | Clinical | 98 | 100 | 0.27 | 99.9837 | 5123738 | 45.28 | 62 | 322237 |
| VP17126 | VP17126 | SAMN20371914 | SRR15239883 | PC | Public | 2017 | China | Clinical | 98.1 | 100 | 0 | 99.9903 | 5068801 | 45.28 | 82 | 288527 |
| VP17127 | VP17127 | SAMN20371906 | SRR15239892 | NonPC | Public | 2017 | China | Clinical | 90.7 | 100 | 0.67 | 98.2697 | 5109261 | 45.25 | 58 | 298071 |
| VP17129 | VP17129 | SAMN20371912 | SRR15239886 | NonPC | Public | 2017 | China | Clinical | 92.3 | 100 | 0.19 | 98.3519 | 5027466 | 45.24 | 52 | 539196 |
| VP17130 | VP17130 | SAMN20371882 | SRR15239924 | PC | Public | 2017 | China | Clinical | 98.4 | 100 | 0 | 99.9802 | 5096030 | 45.29 | 60 | 401597 |
| VP17131 | VP17131 | SAMN20371880 | SRR15239926 | PC | Public | 2017 | China | Clinical | 98.4 | 100 | 0 | 99.9942 | 5090023 | 45.29 | 56 | 425229 |
| VP17132 | VP17132 | SAMN20371917 | SRR15239880 | PC | Public | 2017 | China | Clinical | 98.4 | 100 | 0 | 99.9829 | 5089637 | 45.29 | 56 | 425343 |
| VP17133 | VP17133 | SAMN20371916 | SRR15239881 | PC | Public | 2017 | China | Clinical | 98.4 | 100 | 0 | 99.9833 | 5089324 | 45.29 | 55 | 425229 |
| VP17135 | VP17135 | SAMN20371918 | SRR15239879 | PC | Public | 2017 | China | Clinical | 98 | 100 | 0 | 99.9912 | 5071403 | 45.27 | 81 | 269810 |
| VP17136 | VP17136 | SAMN20371920 | SRR15239877 | PC | Public | 2017 | China | Clinical | 98.4 | 100 | 0 | 99.9923 | 5089690 | 45.29 | 56 | 425343 |
| VP17137 | VP17137 | SAMN20371929 | SRR15239867 | PC | Public | 2017 | China | Clinical | 97.9 | 100 | 0 | 99.9775 | 5139338 | 45.28 | 49 | 394836 |
| VP17138 | VP17138 | SAMN20371926 | SRR15239870 | NonPC | Public | 2017 | China | Clinical | 92.4 | 100 | 0.05 | 98.3201 | 5133402 | 45.07 | 47 | 450392 |
| VP17139 | VP17139 | SAMN20371921 | SRR15239876 | PC | Public | 2017 | China | Clinical | 98.4 | 100 | 0.27 | 99.9925 | 5135572 | 45.29 | 58 | 394796 |
| VP17140 | VP17140 | SAMN20371909 | SRR15239889 | PC | Public | 2017 | China | Clinical | 98.4 | 100 | 0 | 99.9917 | 5089650 | 45.29 | 62 | 425343 |
| VP17141 | VP17141 | SAMN20371911 | SRR15239887 | PC | Public | 2017 | China | Clinical | 98.4 | 100 | 0 | 99.977 | 5096678 | 45.29 | 58 | 425469 |
| VP17142 | VP17142 | SAMN20371923 | SRR15239874 | PC | Public | 2017 | China | Clinical | 98.4 | 100 | 0 | 99.9897 | 5096371 | 45.29 | 62 | 401597 |
| VP17143 | VP17143 | SAMN20371922 | SRR15239875 | PC | Public | 2017 | China | Clinical | 98 | 100 | 0 | 99.9921 | 5126680 | 45.23 | 60 | 394981 |
| VP17144 | VP17144 | SAMN20371925 | SRR15239871 | PC | Public | 2017 | China | Clinical | 98.1 | 100 | 0 | 99.9897 | 5069781 | 45.28 | 83 | 265886 |
| VP17145 | VP17145 | SAMN20371693 | SRR15240314 | PC | Public | 2017 | China | Clinical | 98.4 | 100 | 0 | 99.9888 | 5089995 | 45.29 | 55 | 425871 |
| VP17146 | VP17146 | SAMN20371690 | SRR15240317 | PC | Public | 2017 | China | Clinical | 98.4 | 100 | 0 | 99.9913 | 5090771 | 45.29 | 58 | 425343 |
| VP17147 | VP17147 | SAMN20371694 | SRR15240312 | PC | Public | 2017 | China | Clinical | 98.3 | 100 | 0 | 99.9782 | 5095959 | 45.28 | 56 | 401463 |
| VP17148 | VP17148 | SAMN20371932 | SRR15239864 | PC | Public | 2017 | China | Clinical | 98.4 | 100 | 0 | 99.9834 | 5089530 | 45.29 | 60 | 425193 |
| VP17149 | VP17149 | SAMN20371930 | SRR15239866 | PC | Public | 2017 | China | Clinical | 98.4 | 100 | 0 | 99.98 | 5094523 | 45.29 | 57 | 401590 |
| VP17150 | VP17150 | SAMN20371919 | SRR15239878 | NonPC | Public | 2017 | China | Clinical | 90.9 | 100 | 0.33 | 98.2858 | 5140766 | 45.18 | 61 | 376979 |
| VP17151 | VP17151 | SAMN20371924 | SRR15239872 | NonPC | Public | 2017 | China | Clinical | 91.3 | 100 | 0 | 98.3045 | 5150150 | 45.25 | 47 | 426751 |
| VP17152 | VP17152 | SAMN20371927 | SRR15239869 | PC | Public | 2017 | China | Clinical | 97.9 | 100 | 0 | 99.9709 | 5153909 | 45.28 | 46 | 401597 |
| VP17155 | VP17155 | SAMN20371935 | SRR15239859 | NonPC | Public | 2017 | China | Clinical | 91.6 | 100 | 0.46 | 98.2432 | 5110544 | 45.2 | 60 | 440234 |
| VP17158 | VP17158 | SAMN20371936 | SRR15239858 | NonPC | Public | 2017 | China | Clinical | 92.3 | 100 | 0.19 | 98.3313 | 5027695 | 45.24 | 52 | 471204 |
| VP17160-2 | VP17160-2 | SAMN20371933 | SRR15239863 | PC | Public | 2017 | China | Clinical | 98 | 100 | 0 | 99.9932 | 5080146 | 45.28 | 57 | 401604 |
| VP17161 | VP17161 | SAMN20371931 | SRR15239865 | NonPC | Public | 2017 | China | Clinical | 92 | 100 | 0.46 | 98.2606 | 5073535 | 45.2 | 47 | 544881 |
| VP17162 | VP17162 | SAMN20371934 | SRR15239860 | PC | Public | 2017 | China | Clinical | 98.4 | 100 | 0 | 99.9743 | 5149259 | 45.26 | 58 | 401456 |
| VP17163 | VP17163 | SAMN20371928 | SRR15239868 | PC | Public | 2017 | China | Clinical | 98 | 100 | 0 | 99.9828 | 5080854 | 45.28 | 60 | 401597 |

| | | | | | | | | | | | | | | | | |
|---|---|---|---|---|---|---|---|---|---|---|---|---|---|---|---|---|
| VP17165 | VP17165 | SAMN20371938 | SRR15239856 | NonPC | Public | 2017 | China | Clinical | 89.4 | 100 | 0.27 | 98.3046 | 5026126 | 45.31 | 54 | 546793 |
| VP17166 | VP17166 | SAMN20371937 | SRR15239857 | PC | Public | 2017 | China | Clinical | 98.3 | 100 | 0 | 99.9672 | 5138078 | 45.28 | 58 | 473797 |
| VP17168 | VP17168 | SAMN20371692 | SRR15240315 | PC | Public | 2017 | China | Clinical | 97.9 | 100 | 0 | 99.9836 | 5077274 | 45.27 | 56 | 401470 |
| VP17169 | VP17169 | SAMN20371939 | SRR15239855 | PC | Public | 2017 | China | Clinical | 98.4 | 100 | 0 | 99.9912 | 5091955 | 45.29 | 58 | 425307 |
| VP17170 | VP17170 | SAMN20371940 | SRR15239854 | PC | Public | 2017 | China | Clinical | 98.4 | 100 | 0 | 99.9877 | 5099487 | 45.28 | 55 | 425355 |
| VP18001 | VP18001 | SAMN20371948 | SRR15239845 | PC | Public | 2018 | China | Clinical | 98.4 | 100 | 0 | 99.9907 | 5097934 | 45.28 | 59 | 425961 |
| VP18002 | VP18002 | SAMN20371942 | SRR15239852 | PC | Public | 2018 | China | Clinical | 98.1 | 100 | 0 | 99.9891 | 5107380 | 45.32 | 49 | 416372 |
| VP18003 | VP18003 | SAMN20371951 | SRR15239842 | NonPC | Public | 2018 | China | Clinical | 92.2 | 100 | 0.52 | 98.4132 | 5101184 | 45.23 | 55 | 209769 |
| VP18004 | VP18004 | SAMN20371941 | SRR15239853 | NonPC | Public | 2018 | China | Clinical | 92 | 100 | 0.46 | 98.2261 | 5074012 | 45.2 | 50 | 450300 |
| VP18005 | VP18005 | SAMN20371952 | SRR15239841 | NonPC | Public | 2018 | China | Clinical | 91.1 | 100 | 0 | 98.3255 | 5203360 | 45.26 | 71 | 291595 |
| VP18006 | VP18006 | SAMN20371953 | SRR15239840 | NonPC | Public | 2018 | China | Clinical | 92.2 | 100 | 0.52 | 98.4464 | 5102299 | 45.23 | 53 | 238781 |
| VP18007 | VP18007 | SAMN20371954 | SRR15239838 | PC | Public | 2018 | China | Clinical | 98.1 | 100 | 0 | 99.979 | 5126756 | 45.31 | 49 | 415922 |
| VP18008 | VP18008 | SAMN20371955 | SRR15239837 | NonPC | Public | 2018 | China | Clinical | 92 | 100 | 0.46 | 98.2598 | 5081656 | 45.2 | 53 | 450300 |
| VP18009 | VP18009 | SAMN20371956 | SRR15239836 | PC | Public | 2018 | China | Clinical | 98.4 | 100 | 0 | 99.991 | 5089833 | 45.29 | 57 | 401472 |
| VP18010 | VP18010 | SAMN20371957 | SRR15239835 | PC | Public | 2018 | China | Clinical | 98.4 | 100 | 0 | 99.9927 | 5090598 | 45.29 | 60 | 394877 |
| VP18011 | VP18011 | SAMN20371958 | SRR15239834 | PC | Public | 2018 | China | Clinical | 98.1 | 100 | 0.27 | 99.9701 | 5111202 | 45.32 | 54 | 401327 |
| VP18012 | VP18012 | SAMN20371962 | SRR15239830 | PC | Public | 2018 | China | Clinical | 98.1 | 100 | 0.27 | 99.989 | 5111213 | 45.32 | 52 | 409410 |
| VP18013 | VP18013 | SAMN20371960 | SRR15239832 | PC | Public | 2018 | China | Clinical | 98.1 | 100 | 0.27 | 99.991 | 5110417 | 45.32 | 54 | 409296 |
| VP18014 | VP18014 | SAMN20371961 | SRR15239831 | PC | Public | 2018 | China | Clinical | 98.1 | 100 | 0.27 | 99.9821 | 5112132 | 45.32 | 52 | 408768 |
| VP18015 | VP18015 | SAMN20371949 | SRR15239844 | PC | Public | 2018 | China | Clinical | 98.1 | 100 | 0.27 | 99.9812 | 5110819 | 45.32 | 53 | 408654 |
| VP18017 | VP18017 | SAMN20371959 | SRR15239833 | PC | Public | 2018 | China | Clinical | 98.1 | 100 | 0.27 | 99.9706 | 5111703 | 45.32 | 50 | 408882 |
| VP18019 | VP18019 | SAMN20371973 | SRR15239818 | PC | Public | 2018 | China | Clinical | 98.1 | 100 | 0.27 | 99.9893 | 5107178 | 45.32 | 48 | 401446 |
| VP18020 | VP18020 | SAMN20371972 | SRR15239819 | PC | Public | 2018 | China | Clinical | 98.4 | 100 | 0 | 99.9846 | 5108735 | 45.28 | 57 | 401470 |
| VP18023 | VP18023 | SAMN20371976 | SRR15239814 | PC | Public | 2018 | China | Clinical | 98 | 100 | 0 | 99.9935 | 5079797 | 45.28 | 58 | 394994 |
| VP18024 | VP18024 | SAMN20371980 | SRR15239810 | PC | Public | 2018 | China | Clinical | 98.4 | 100 | 0 | 99.978 | 5097971 | 45.28 | 57 | 425241 |
| VP18025 | VP18025 | SAMN20371979 | SRR15239811 | PC | Public | 2018 | China | Clinical | 98.1 | 100 | 0.27 | 99.9818 | 5111277 | 45.32 | 52 | 408768 |
| VP18026 | VP18026 | SAMN20371978 | SRR15239812 | NonPC | Public | 2018 | China | Clinical | 93.1 | 100 | 0.05 | 98.4008 | 5093233 | 45.28 | 51 | 443425 |
| VP18027 | VP18027 | SAMN20371981 | SRR15239809 | PC | Public | 2018 | China | Clinical | 98.4 | 100 | 0 | 99.9832 | 5108285 | 45.28 | 55 | 425241 |
| VP18028 | VP18028 | SAMN20371975 | SRR15239815 | PC | Public | 2018 | China | Clinical | 98.4 | 100 | 0 | 99.9842 | 5109435 | 45.28 | 54 | 401604 |
| VP18029 | VP18029 | SAMN20371974 | SRR15239816 | PC | Public | 2018 | China | Clinical | 98.4 | 100 | 0 | 99.9758 | 5109042 | 45.28 | 54 | 425241 |
| VP18030 | VP18030 | SAMN20371985 | SRR15239804 | PC | Public | 2018 | China | Clinical | 98.4 | 100 | 0 | 99.9782 | 5090425 | 45.29 | 57 | 425343 |
| VP18031 | VP18031 | SAMN20371943 | SRR15239851 | PC | Public | 2018 | China | Clinical | 98 | 100 | 0 | 99.9836 | 5081697 | 45.28 | 60 | 394901 |
| VP18033 | VP18033 | SAMN20371950 | SRR15239843 | PC | Public | 2018 | China | Clinical | 98.4 | 100 | 0 | 99.9825 | 5091457 | 45.29 | 56 | 425193 |
| VP18034 | VP18034 | SAMN20371982 | SRR15239808 | NonPC | Public | 2018 | China | Clinical | 91.1 | 100 | 0.21 | 98.3334 | 5281735 | 45.07 | 56 | 412524 |
| VP18035 | VP18035 | SAMN20371990 | SRR15239799 | PC | Public | 2018 | China | Clinical | 98.5 | 100 | 0 | 99.9836 | 5092759 | 45.29 | 56 | 425721 |
| VP18036 | VP18036 | SAMN20371989 | SRR15239800 | PC | Public | 2018 | China | Clinical | 98.4 | 100 | 0 | 99.9851 | 5109491 | 45.28 | 55 | 394810 |
| VP18037 | VP18037 | SAMN20371983 | SRR15239807 | PC | Public | 2018 | China | Clinical | 98.4 | 100 | 0 | 99.9831 | 5115141 | 45.28 | 55 | 425883 |
| VP18038 | VP18038 | SAMN20371984 | SRR15239805 | PC | Public | 2018 | China | Clinical | 98.4 | 100 | 0 | 99.9744 | 5108616 | 45.28 | 53 | 425241 |
| VP18042 | VP18042 | SAMN20371977 | SRR15239813 | NonPC | Public | 2018 | China | Clinical | 90.9 | 100 | 0.03 | 98.2552 | 5100531 | 45.29 | 55 | 408663 |
| VP18043 | VP18043 | SAMN20372006 | SRR15239781 | PC | Public | 2018 | China | Clinical | 98.4 | 100 | 0 | 99.9827 | 5109093 | 45.28 | 59 | 313348 |
| VP18044 | VP18044 | SAMN20372012 | SRR15239775 | PC | Public | 2018 | China | Clinical | 98.4 | 100 | 0 | 99.9736 | 5109074 | 45.28 | 53 | 425355 |
| VP18045 | VP18045 | SAMN20372007 | SRR15239780 | NonPC | Public | 2018 | China | Clinical | 93.1 | 100 | 0.19 | 98.3905 | 5079913 | 45.29 | 47 | 424375 |
| VP18047 | VP18047 | SAMN20372008 | SRR15239779 | PC | Public | 2018 | China | Clinical | 98.5 | 100 | 0 | 99.9923 | 5092501 | 45.29 | 58 | 401470 |
| VP18048 | VP18048 | SAMN20372009 | SRR15239778 | PC | Public | 2018 | China | Clinical | 98.5 | 100 | 0 | 99.9916 | 5092907 | 45.29 | 58 | 401604 |
| VP18049 | VP18049 | SAMN20372010 | SRR15239777 | PC | Public | 2018 | China | Env | 98.5 | 100 | 0 | 99.9923 | 5092812 | 45.29 | 59 | 394963 |
| VP18050 | VP18050 | SAMN20372016 | SRR15239766 | NonPC | Public | 2018 | China | Clinical | 91.9 | 100 | 0.46 | 98.266 | 5074013 | 45.2 | 50 | 450397 |
| VP18051 | VP18051 | SAMN20372013 | SRR15239770 | NonPC | Public | 2018 | China | Clinical | 90.5 | 100 | 0 | 98.266 | 5170392 | 45.29 | 39 | 641540 |
| VP18052 | VP18052 | SAMN20372014 | SRR15239768 | PC | Public | 2018 | China | Clinical | 98 | 100 | 0 | 99.9783 | 5101791 | 45.26 | 61 | 395294 |

| | | | | | | | | | | | | | | | | |
|---|---|---|---|---|---|---|---|---|---|---|---|---|---|---|---|---|
| VP18053 | VP18053 | SAMN20372011 | SRR15239776 | PC | Public | 2018 | China | Clinical | 98.5 | 100 | 0 | 99.9829 | 5093800 | 45.29 | 58 | 401470 |
| VP18054 | VP18054 | SAMN20372002 | SRR15239786 | PC | Public | 2018 | China | Clinical | 98.5 | 100 | 0 | 99.9833 | 5092920 | 45.29 | 59 | 401604 |
| VP18055 | VP18055 | SAMN20372003 | SRR15239785 | PC | Public | 2018 | China | Clinical | 98.5 | 100 | 0 | 99.9834 | 5092788 | 45.29 | 57 | 425271 |
| VP18056 | VP18056 | SAMN20372004 | SRR15239783 | PC | Public | 2018 | China | Clinical | 98.5 | 100 | 0 | 99.9838 | 5094071 | 45.29 | 55 | 425949 |
| VP18057 | VP18057 | SAMN20372005 | SRR15239782 | PC | Public | 2018 | China | Clinical | 98.5 | 100 | 0 | 99.9838 | 5181928 | 45.21 | 63 | 401470 |
| VP18058 | VP18058 | SAMN20372020 | SRR15239762 | NonPC | Public | 2018 | China | Clinical | 91.9 | 100 | 0.19 | 98.3579 | 5027243 | 45.24 | 50 | 539189 |
| VP18059 | VP18059 | SAMN20372021 | SRR15239761 | PC | Public | 2018 | China | Clinical | 98.3 | 100 | 0 | 99.9841 | 5109275 | 45.28 | 54 | 425355 |
| VP18060 | VP18060 | SAMN20372022 | SRR15239760 | PC | Public | 2018 | China | Clinical | 98.4 | 100 | 0 | 99.9844 | 5109580 | 45.28 | 58 | 425355 |
| VP18061 | VP18061 | SAMN20372018 | SRR15239764 | PC | Public | 2018 | China | Clinical | 98.4 | 100 | 0 | 99.9838 | 5109331 | 45.28 | 54 | 425997 |
| VP18062 | VP18062 | SAMN20372019 | SRR15239763 | PC | Public | 2018 | China | Clinical | 98.4 | 100 | 0 | 99.9813 | 5090062 | 45.29 | 57 | 401456 |
| VP18063 | VP18063 | SAMN20372030 | SRR15239751 | PC | Public | 2018 | China | Clinical | 98.5 | 100 | 0 | 99.9828 | 5092915 | 45.29 | 57 | 401470 |
| VP18064 | VP18064 | SAMN20372029 | SRR15239752 | PC | Public | 2018 | China | Clinical | 98.5 | 100 | 0 | 99.9933 | 5093317 | 45.29 | 57 | 394935 |
| VP18065 | VP18065 | SAMN20372015 | SRR15239767 | NonPC | Public | 2018 | China | Clinical | 92.3 | 100 | 0.19 | 98.3801 | 5028503 | 45.24 | 52 | 539189 |
| VP18066 | VP18066 | SAMN20372023 | SRR15239759 | PC | Public | 2018 | China | Clinical | 98.4 | 100 | 0 | 99.9924 | 5092520 | 45.29 | 58 | 425307 |
| VP18067 | VP18067 | SAMN20372043 | SRR15239736 | PC | Public | 2018 | China | Clinical | 98.5 | 100 | 0 | 99.9835 | 5092177 | 45.29 | 58 | 425307 |
| VP18068 | VP18068 | SAMN20372037 | SRR15239742 | PC | Public | 2018 | China | Clinical | 98.5 | 100 | 0 | 99.9831 | 5092615 | 45.29 | 57 | 401597 |
| VP18069 | VP18069 | SAMN20372035 | SRR15239744 | PC | Public | 2018 | China | Clinical | 98.5 | 100 | 0 | 99.9828 | 5093409 | 45.29 | 56 | 425949 |
| VP18070 | VP18070 | SAMN20372017 | SRR15239765 | PC | Public | 2018 | China | Clinical | 98.5 | 100 | 0 | 99.9847 | 5092220 | 45.29 | 57 | 425307 |
| VP18071 | VP18071 | SAMN20371944 | SRR15239849 | PC | Public | 2018 | China | Clinical | 98.1 | 100 | 0 | 99.9904 | 5107617 | 45.32 | 53 | 394944 |
| VP18072 | VP18072 | SAMN20371947 | SRR15239846 | NonPC | Public | 2018 | China | Env | 90.2 | 100 | 0.27 | 98.2704 | 4992511 | 45.4 | 44 | 405447 |
| VP18073 | VP18073 | SAMN20372049 | SRR15240300 | PC | Public | 2018 | China | Clinical | 98.5 | 100 | 0 | 99.9826 | 5093275 | 45.29 | 58 | 425193 |
| VP18074 | VP18074 | SAMN20372050 | SRR15240299 | PC | Public | 2018 | China | Clinical | 98.5 | 100 | 0 | 99.9835 | 5092422 | 45.29 | 56 | 425835 |
| VP18075 | VP18075 | SAMN20372051 | SRR15240298 | PC | Public | 2018 | China | Clinical | 98.5 | 100 | 0 | 99.992 | 5093082 | 45.29 | 59 | 394769 |
| VP18076 | VP18076 | SAMN20372048 | SRR15240301 | PC | Public | 2018 | China | Clinical | 98.5 | 100 | 0 | 99.9845 | 5091859 | 45.29 | 65 | 312826 |
| VP18077 | VP18077 | SAMN20372047 | SRR15240302 | PC | Public | 2018 | China | Clinical | 98.5 | 100 | 0 | 99.9927 | 5092187 | 45.29 | 64 | 288859 |
| VP18078 | VP18078 | SAMN20372052 | SRR15240297 | PC | Public | 2018 | China | Clinical | 98.5 | 100 | 0 | 99.9926 | 5092871 | 45.29 | 58 | 401604 |
| VP18079 | VP18079 | SAMN20372045 | SRR15240304 | PC | Public | 2018 | China | Clinical | 98.5 | 100 | 0 | 99.9878 | 5182040 | 45.21 | 66 | 312692 |
| VP18080 | VP18080 | SAMN20372046 | SRR15240303 | PC | Public | 2018 | China | Clinical | 98.5 | 100 | 0 | 99.9914 | 5092607 | 45.29 | 57 | 401604 |
| VP18081 | VP18081 | SAMN20372053 | SRR15240296 | PC | Public | 2018 | China | Env | 98.5 | 100 | 0 | 99.991 | 5093471 | 45.29 | 59 | 425835 |
| VP18083 | VP18083 | SAMN20372091 | SRR15240221 | PC | Public | 2018 | China | Clinical | 97.8 | 100 | 0 | 99.9789 | 5153075 | 45.28 | 44 | 580312 |
| VP18084 | VP18084 | SAMN20372099 | SRR15240212 | NonPC | Public | 2018 | China | Env | 88 | 100 | 0.33 | 98.2011 | 5156706 | 45.12 | 80 | 322532 |
| VP18086 | VP18086 | SAMN20372100 | SRR15240211 | NonPC | Public | 2018 | China | Env | 88.3 | 100 | 0.63 | 98.2901 | 4966412 | 45.34 | 55 | 439949 |
| VP18087 | VP18087 | SAMN20372101 | SRR15240210 | NonPC | Public | 2018 | China | Env | 87.8 | 100 | 0.37 | 98.2505 | 5055640 | 45.27 | 54 | 368628 |
| VP18088 | VP18088 | SAMN20372103 | SRR15240208 | NonPC | Public | 2018 | China | Clinical | 92.3 | 100 | 0.19 | 98.3453 | 5039597 | 45.23 | 49 | 450299 |
| VP18089 | VP18089 | SAMN20372102 | SRR15240209 | NonPC | Public | 2018 | China | Clinical | 92.3 | 100 | 0.19 | 98.3364 | 5040201 | 45.23 | 51 | 450299 |
| VP18090 | VP18090 | SAMN20372104 | SRR15240206 | PC | Public | 2018 | China | Clinical | 98.4 | 100 | 0 | 99.9752 | 5143119 | 45.29 | 57 | 401604 |
| VP18091 | VP18091 | SAMN20372098 | SRR15240213 | NonPC | Public | 2018 | China | Clinical | 92.1 | 100 | 0.05 | 98.4343 | 5000154 | 45.35 | 48 | 463574 |
| VP18094 | VP18094 | SAMN20372034 | SRR15239745 | PC | Public | 2018 | China | Clinical | 98.5 | 100 | 0 | 99.9909 | 5093350 | 45.29 | 59 | 394933 |
| VP18095 | VP18095 | SAMN20372032 | SRR15239749 | PC | Public | 2018 | China | Clinical | 98.5 | 100 | 0 | 99.9915 | 5094245 | 45.29 | 61 | 425193 |
| VP18096 | VP18096 | SAMN20372036 | SRR15239743 | PC | Public | 2018 | China | Clinical | 98.5 | 100 | 0 | 99.9831 | 5092320 | 45.29 | 58 | 401604 |
| VP18098 | VP18098 | SAMN20372090 | SRR15240222 | PC | Public | 2018 | China | Clinical | 98.5 | 100 | 0 | 99.9824 | 5091972 | 45.29 | 56 | 401604 |
| VP18099 | VP18099 | SAMN20372089 | SRR15240223 | PC | Public | 2018 | China | Clinical | 98.5 | 100 | 0 | 99.9823 | 5093106 | 45.29 | 57 | 401604 |
| VP18100 | VP18100 | SAMN20372097 | SRR15240214 | PC | Public | 2018 | China | Clinical | 98.5 | 100 | 0 | 99.9827 | 5092452 | 45.29 | 59 | 394994 |
| VP18101 | VP18101 | SAMN20372096 | SRR15240215 | PC | Public | 2018 | China | Clinical | 98.5 | 100 | 0 | 99.9919 | 5092873 | 45.29 | 56 | 425835 |
| VP18102 | VP18102 | SAMN20372092 | SRR15240220 | PC | Public | 2018 | China | Clinical | 98.4 | 100 | 0 | 99.9907 | 5089223 | 45.29 | 56 | 425355 |
| VP18103 | VP18103 | SAMN20372093 | SRR15240219 | NonPC | Public | 2018 | China | Clinical | 93.1 | 100 | 0.05 | 98.3538 | 5066630 | 45.3 | 43 | 463552 |
| VP18104 | VP18104 | SAMN20372106 | SRR15240204 | PC | Public | 2018 | China | Clinical | 98.5 | 100 | 0 | 99.9821 | 5092602 | 45.29 | 53 | 425721 |
| VP18107 | VP18107 | SAMN20372056 | SRR15240292 | PC | Public | 2018 | China | Clinical | 98.4 | 100 | 0 | 99.9898 | 5097861 | 45.28 | 56 | 401463 |

| | | | | | | | | | | | | | | | | |
|---|---|---|---|---|---|---|---|---|---|---|---|---|---|---|---|---|
| VP18109 | VP18109 | SAMN20372054 | SRR15240294 | PC | Public | 2018 | China | Clinical | 98.4 | 100 | 0 | 99.98 | 5097283 | 45.28 | 57 | 425883 |
| VP18110 | VP18110 | SAMN20372042 | SRR15239737 | PC | Public | 2018 | China | Clinical | 98.5 | 100 | 0 | 99.9925 | 5093274 | 45.29 | 57 | 425193 |
| VP18111 | VP18111 | SAMN20372041 | SRR15239738 | PC | Public | 2018 | China | Clinical | 98.5 | 100 | 0 | 99.9921 | 5092898 | 45.29 | 57 | 440663 |
| VP18112 | VP18112 | SAMN20372044 | SRR15240305 | PC | Public | 2018 | China | Clinical | 98.5 | 100 | 0 | 99.9837 | 5093036 | 45.29 | 59 | 334357 |
| VP18113 | VP18113 | SAMN20372040 | SRR15239739 | PC | Public | 2018 | China | Clinical | 98.5 | 100 | 0 | 99.9825 | 5093482 | 45.29 | 61 | 312808 |
| VP18114 | VP18114 | SAMN20372039 | SRR15239740 | PC | Public | 2018 | China | Clinical | 97.8 | 100 | 0.01 | 99.9037 | 5190866 | 45.32 | 69 | 312690 |
| VP18115 | VP18115 | SAMN20372055 | SRR15240293 | NonPC | Public | 2018 | China | Clinical | 93.4 | 100 | 0.05 | 98.3711 | 5095285 | 45.28 | 46 | 424363 |
| VP18117 | VP18117 | SAMN20372058 | SRR15240290 | PC | Public | 2018 | China | Clinical | 98.4 | 100 | 0 | 99.9929 | 5089489 | 45.29 | 56 | 401477 |
| VP18118 | VP18118 | SAMN20372118 | SRR15240158 | NonPC | Public | 2018 | China | Clinical | 92 | 100 | 0.46 | 98.273 | 5074181 | 45.2 | 51 | 450300 |
| VP18120 | VP18120 | SAMN20372111 | SRR15240166 | PC | Public | 2018 | China | Clinical | 98.4 | 100 | 0 | 99.9766 | 5108130 | 45.28 | 54 | 401470 |
| VP18122 | VP18122 | SAMN20372109 | SRR15240201 | PC | Public | 2018 | China | Clinical | 98.4 | 100 | 0 | 99.9752 | 5110097 | 45.28 | 55 | 425355 |
| VP18123 | VP18123 | SAMN20372110 | SRR15240167 | PC | Public | 2018 | China | Clinical | 98.5 | 100 | 0 | 99.9928 | 5096218 | 45.29 | 60 | 425307 |
| VP18124 | VP18124 | SAMN20372107 | SRR15240203 | PC | Public | 2018 | China | Clinical | 97.9 | 100 | 0 | 99.9513 | 5154375 | 45.27 | 45 | 401463 |
| VP18125 | VP18125 | SAMN20372112 | SRR15240165 | PC | Public | 2018 | China | Clinical | 98.4 | 100 | 0 | 99.9917 | 5090546 | 45.29 | 58 | 440633 |
| VP18127 | VP18127 | SAMN20372095 | SRR15240216 | PC | Public | 2018 | China | Clinical | 98.4 | 100 | 0 | 99.9829 | 5087343 | 45.29 | 59 | 401477 |
| VP18128 | VP18128 | SAMN20372105 | SRR15240205 | NonPC | Public | 2018 | China | Clinical | 92.3 | 100 | 0.19 | 98.3279 | 5039629 | 45.23 | 49 | 450299 |
| VP18129 | VP18129 | SAMN20372094 | SRR15240217 | NonPC | Public | 2018 | China | Clinical | 91.9 | 100 | 0.46 | 98.2663 | 5074355 | 45.2 | 48 | 450300 |
| VP18130 | VP18130 | SAMN20372113 | SRR15240164 | NonPC | Public | 2018 | China | Clinical | 92 | 100 | 0.46 | 98.2419 | 5073552 | 45.2 | 50 | 450300 |
| VP18131 | VP18131 | SAMN20372114 | SRR15240162 | NonPC | Public | 2018 | China | Clinical | 91.6 | 100 | 0.46 | 98.2433 | 5084494 | 45.19 | 52 | 450300 |
| VP18134 | VP18134 | SAMN20372108 | SRR15240202 | NonPC | Public | 2018 | China | Clinical | 92 | 100 | 0.46 | 98.2568 | 5082204 | 45.2 | 53 | 450300 |
| VP18135 | VP18135 | SAMN20372038 | SRR15239741 | PC | Public | 2018 | China | Clinical | 98.5 | 100 | 0 | 99.9847 | 5099903 | 45.3 | 59 | 401604 |
| VP18137 | VP18137 | SAMN20372119 | SRR15240157 | PC | Public | 2018 | China | Clinical | 98.5 | 100 | 0 | 99.9916 | 5093001 | 45.29 | 57 | 425721 |
| VP18138 | VP18138 | SAMN20372120 | SRR15240156 | PC | Public | 2018 | China | Clinical | 98.4 | 100 | 0 | 99.9934 | 5093005 | 45.29 | 58 | 425307 |
| VP18139 | VP18139 | SAMN20372116 | SRR15240160 | PC | Public | 2018 | China | Clinical | 98.5 | 100 | 0 | 99.9829 | 5092419 | 45.29 | 56 | 425721 |
| VP18140 | VP18140 | SAMN20372117 | SRR15240159 | NonPC | Public | 2018 | China | Clinical | 92.3 | 100 | 0.19 | 98.3705 | 5040420 | 45.23 | 52 | 450299 |
| VP18141 | VP18141 | SAMN20372127 | SRR15240148 | PC | Public | 2018 | China | Clinical | 98.5 | 100 | 0 | 99.9923 | 5092441 | 45.29 | 60 | 401604 |
| VP18142 | VP18142 | SAMN20372121 | SRR15240155 | NonPC | Public | 2018 | China | Clinical | 92 | 100 | 0.46 | 98.2783 | 5082992 | 45.2 | 50 | 450300 |
| VP18143 | VP18143 | SAMN20372123 | SRR15240153 | PC | Public | 2018 | China | Clinical | 98.4 | 100 | 0 | 99.9807 | 5090435 | 45.29 | 55 | 401590 |
| VP18144 | VP18144 | SAMN20372126 | SRR15240149 | PC | Public | 2018 | China | Clinical | 98.4 | 100 | 0 | 99.9898 | 5090832 | 45.29 | 57 | 425343 |
| VP18145 | VP18145 | SAMN20372128 | SRR15240147 | PC | Public | 2018 | China | Clinical | 98.3 | 100 | 0 | 99.9779 | 5108863 | 45.28 | 52 | 425241 |
| VP18146 | VP18146 | SAMN20372130 | SRR15240145 | PC | Public | 2018 | China | Clinical | 98.3 | 100 | 0 | 99.9818 | 5109009 | 45.28 | 56 | 425355 |
| VP18147 | VP18147 | SAMN20372132 | SRR15240143 | PC | Public | 2018 | China | Clinical | 98.3 | 100 | 0 | 99.9765 | 5108292 | 45.28 | 53 | 425355 |
| VP18148 | VP18148 | SAMN20372125 | SRR15240150 | PC | Public | 2018 | China | Clinical | 98.4 | 100 | 0 | 99.9895 | 5097154 | 45.28 | 59 | 425241 |
| VP18149 | VP18149 | SAMN20372129 | SRR15240146 | PC | Public | 2018 | China | Clinical | 98.4 | 100 | 0 | 99.9809 | 5090757 | 45.29 | 59 | 401590 |
| VP18150 | VP18150 | SAMN20372142 | SRR15240131 | NonPC | Public | 2018 | China | Clinical | 90.9 | 100 | 0 | 98.3014 | 5114971 | 45.21 | 65 | 374836 |
| VP18151 | VP18151 | SAMN20372122 | SRR15240154 | PC | Public | 2018 | China | Clinical | 98.4 | 100 | 0 | 99.9734 | 5109661 | 45.28 | 55 | 401604 |
| VP18152 | VP18152 | SAMN20372133 | SRR15240142 | NonPC | Public | 2018 | China | Clinical | 90.9 | 100 | 0 | 98.2815 | 5070567 | 45.26 | 63 | 374822 |
| VP18153 | VP18153 | SAMN20371991 | SRR15239798 | PC | Public | 2018 | China | Clinical | 98.5 | 100 | 0 | 99.9916 | 5092954 | 45.29 | 61 | 425307 |
| VP18154 | VP18154 | SAMN20371992 | SRR15239797 | PC | Public | 2018 | China | Clinical | 98.5 | 100 | 0 | 99.9926 | 5093406 | 45.29 | 59 | 401604 |
| VP18155 | VP18155 | SAMN20371993 | SRR15239796 | PC | Public | 2018 | China | Clinical | 98.5 | 100 | 0 | 99.9836 | 5100022 | 45.29 | 59 | 394769 |
| VP18156 | VP18156 | SAMN20371994 | SRR15239794 | PC | Public | 2018 | China | Clinical | 98.5 | 100 | 0 | 99.9917 | 5092795 | 45.29 | 57 | 401604 |
| VP18157 | VP18157 | SAMN20371995 | SRR15239793 | PC | Public | 2018 | China | Clinical | 98.5 | 100 | 0 | 99.9835 | 5092694 | 45.29 | 56 | 401470 |
| VP18158 | VP18158 | SAMN20371996 | SRR15239792 | PC | Public | 2018 | China | Clinical | 98.5 | 100 | 0 | 99.9831 | 5092983 | 45.29 | 57 | 425307 |
| VP18159 | VP18159 | SAMN20371997 | SRR15239791 | PC | Public | 2018 | China | Clinical | 98.5 | 100 | 0 | 99.991 | 5093336 | 45.29 | 57 | 425835 |
| VP18160 | VP18160 | SAMN20371998 | SRR15239790 | PC | Public | 2018 | China | Clinical | 98.5 | 100 | 0 | 99.982 | 5092360 | 45.29 | 57 | 425307 |
| VP18161 | VP18161 | SAMN20371999 | SRR15239789 | PC | Public | 2018 | China | Clinical | 98.5 | 100 | 0 | 99.9917 | 5181703 | 45.21 | 57 | 440665 |
| VP18162 | VP18162 | SAMN20372000 | SRR15239788 | PC | Public | 2018 | China | Clinical | 98.5 | 100 | 0 | 99.9847 | 5091674 | 45.29 | 57 | 425799 |
| VP18163 | VP18163 | SAMN20372001 | SRR15239787 | PC | Public | 2018 | China | Clinical | 98.3 | 100 | 0 | 99.9808 | 5083726 | 45.29 | 57 | 440663 |

| | | | | | | | | | | | | | | | | |
|---|---|---|---|---|---|---|---|---|---|---|---|---|---|---|---|---|
| VP18164 | VP18164 | SAMN20372137 | SRR15240136 | NonPC | Public | 2018 | China | Clinical | 92 | 100 | 0.46 | 98.252 | 5081718 | 45.2 | 50 | 450300 |
| VP18165 | VP18165 | SAMN20372138 | SRR15240135 | NonPC | Public | 2018 | China | Clinical | 92 | 100 | 0.46 | 98.2425 | 5099302 | 45.23 | 57 | 440228 |
| VP18168 | VP18168 | SAMN20372139 | SRR15240134 | NonPC | Public | 2018 | China | Clinical | 92 | 100 | 0.46 | 98.2588 | 5081616 | 45.2 | 55 | 450300 |
| VP18169 | VP18169 | SAMN20372134 | SRR15240139 | NonPC | Public | 2018 | China | Env | 90.4 | 100 | 0 | 98.3001 | 5125305 | 45.27 | 27 | 465583 |
| VP18170 | VP18170 | SAMN20372136 | SRR15240137 | NonPC | Public | 2018 | China | Env | 88.5 | 100 | 0.42 | 98.2391 | 5087526 | 45.22 | 92 | 460419 |
| VP18172 | VP18172 | SAMN20372135 | SRR15240138 | NonPC | Public | 2018 | China | Env | 90 | 100 | 0 | 98.3411 | 4974812 | 45.44 | 26 | 614978 |
| VP18173 | VP18173 | SAMN20372144 | SRR15240128 | PC | Public | 2018 | China | Clinical | 98.4 | 100 | 0 | 99.9753 | 5108202 | 45.28 | 53 | 425355 |
| VP18174 | VP18174 | SAMN20372033 | SRR15239748 | PC | Public | 2018 | China | Clinical | 98.5 | 100 | 0 | 99.984 | 5095242 | 45.29 | 58 | 401604 |
| VP18175 | VP18175 | SAMN20372031 | SRR15239750 | PC | Public | 2018 | China | Clinical | 98.5 | 100 | 0 | 99.9927 | 5093016 | 45.29 | 60 | 394978 |
| VP18176 | VP18176 | SAMN20372141 | SRR15240132 | PC | Public | 2018 | China | Clinical | 98.5 | 100 | 0 | 99.9822 | 5092314 | 45.29 | 65 | 312824 |
| VP18177 | VP18177 | SAMN20372057 | SRR15240291 | NonPC | Public | 2018 | China | Clinical | 91.2 | 100 | 0 | 98.2944 | 5178873 | 45.22 | 63 | 404051 |
| VP18178 | VP18178 | SAMN20372063 | SRR15240285 | PC | Public | 2018 | China | Clinical | 98.4 | 100 | 0 | 99.9849 | 5090074 | 45.29 | 53 | 425355 |
| VP18179 | VP18179 | SAMN20372067 | SRR15240280 | PC | Public | 2018 | China | Clinical | 98.4 | 100 | 0 | 99.9921 | 5090288 | 45.29 | 55 | 394635 |
| VP18180 | VP18180 | SAMN20372074 | SRR15240272 | PC | Public | 2018 | China | Clinical | 98.4 | 100 | 0 | 99.9833 | 5089838 | 45.29 | 54 | 401477 |
| VP18181 | VP18181 | SAMN20372077 | SRR15240236 | PC | Public | 2018 | China | Clinical | 98.4 | 100 | 0 | 99.9833 | 5090274 | 45.29 | 56 | 401611 |
| VP18182 | VP18182 | SAMN20372068 | SRR15240279 | PC | Public | 2018 | China | Clinical | 98.4 | 100 | 0 | 99.9811 | 5089032 | 45.29 | 56 | 425241 |
| VP18183 | VP18183 | SAMN20372069 | SRR15240278 | PC | Public | 2018 | China | Clinical | 98.4 | 100 | 0 | 99.9925 | 5089772 | 45.29 | 57 | 401611 |
| VP18184 | VP18184 | SAMN20372078 | SRR15240235 | PC | Public | 2018 | China | Clinical | 98.3 | 100 | 0 | 99.9705 | 5139445 | 45.28 | 58 | 401477 |
| VP18185 | VP18185 | SAMN20372082 | SRR15240231 | PC | Public | 2018 | China | Clinical | 98.4 | 100 | 0 | 99.9816 | 5089786 | 45.29 | 59 | 394954 |
| VP18186 | VP18186 | SAMN20372070 | SRR15240277 | PC | Public | 2018 | China | Clinical | 98.4 | 100 | 0 | 99.9905 | 5089155 | 45.29 | 58 | 401611 |
| VP18187 | VP18187 | SAMN20372079 | SRR15240234 | PC | Public | 2018 | China | Clinical | 98.4 | 100 | 0 | 99.9916 | 5089033 | 45.29 | 56 | 440623 |
| VP18188 | VP18188 | SAMN20372084 | SRR15240228 | PC | Public | 2018 | China | Clinical | 98.4 | 100 | 0 | 99.9899 | 5090250 | 45.29 | 57 | 401344 |
| VP18189 | VP18189 | SAMN20372081 | SRR15240232 | PC | Public | 2018 | China | Clinical | 98.4 | 100 | 0 | 99.9823 | 5097690 | 45.28 | 59 | 401477 |
| VP18190 | VP18190 | SAMN20372075 | SRR15240271 | PC | Public | 2018 | China | Clinical | 98.4 | 100 | 0 | 99.992 | 5089284 | 45.29 | 55 | 401477 |
| VP18191 | VP18191 | SAMN20372071 | SRR15240276 | PC | Public | 2018 | China | Clinical | 98.4 | 100 | 0 | 99.9831 | 5089986 | 45.29 | 57 | 425361 |
| VP18192 | VP18192 | SAMN20372083 | SRR15240230 | PC | Public | 2018 | China | Clinical | 98.4 | 100 | 0 | 99.9924 | 5090233 | 45.29 | 60 | 425487 |
| VP18193 | VP18193 | SAMN20372060 | SRR15240288 | PC | Public | 2018 | China | Clinical | 98.4 | 100 | 0 | 99.992 | 5090023 | 45.29 | 57 | 401477 |
| VP18194 | VP18194 | SAMN20372059 | SRR15240289 | PC | Public | 2018 | China | Clinical | 98.3 | 100 | 0 | 99.9771 | 5109000 | 45.28 | 56 | 394956 |
| VP18195 | VP18195 | SAMN20372085 | SRR15240227 | PC | Public | 2018 | China | Clinical | 98.4 | 100 | 0 | 99.9838 | 5090133 | 45.29 | 54 | 394954 |
| VP18196 | VP18196 | SAMN20372086 | SRR15240226 | PC | Public | 2018 | China | Clinical | 98.4 | 100 | 0 | 99.9856 | 5088851 | 45.29 | 55 | 394954 |
| VP18197 | VP18197 | SAMN20372087 | SRR15240225 | PC | Public | 2018 | China | Clinical | 98.4 | 100 | 0 | 99.9906 | 5091583 | 45.29 | 55 | 394956 |
| VP18198 | VP18198 | SAMN20372088 | SRR15240224 | PC | Public | 2018 | China | Clinical | 98.4 | 100 | 0 | 99.9828 | 5090961 | 45.29 | 58 | 401476 |
| VP18199 | VP18199 | SAMN20372072 | SRR15240275 | PC | Public | 2018 | China | Clinical | 98.4 | 100 | 0 | 99.9828 | 5090891 | 45.29 | 52 | 401344 |
| VP18200 | VP18200 | SAMN20372065 | SRR15240282 | PC | Public | 2018 | China | Clinical | 98.4 | 100 | 0 | 99.9823 | 5090338 | 45.29 | 57 | 394954 |
| VP18201 | VP18201 | SAMN20372061 | SRR15240287 | PC | Public | 2018 | China | Clinical | 98.4 | 100 | 0 | 99.9827 | 5090838 | 45.29 | 62 | 394956 |
| VP18202 | VP18202 | SAMN20372080 | SRR15240233 | PC | Public | 2018 | China | Clinical | 98.3 | 100 | 0 | 99.9564 | 5140499 | 45.28 | 57 | 394941 |
| VP18203 | VP18203 | SAMN20372064 | SRR15240283 | PC | Public | 2018 | China | Clinical | 98.4 | 100 | 0 | 99.9816 | 5089604 | 45.29 | 58 | 425205 |
| VP18204 | VP18204 | SAMN20372066 | SRR15240281 | PC | Public | 2018 | China | Clinical | 97.8 | 100 | 0 | 99.9833 | 5089951 | 45.29 | 53 | 401477 |
| VP18205 | VP18205 | SAMN20372062 | SRR15240286 | PC | Public | 2018 | China | Clinical | 98.4 | 100 | 0 | 99.9892 | 5089008 | 45.29 | 56 | 425355 |
| VP18207 | VP18207 | SAMN20372073 | SRR15240274 | PC | Public | 2018 | China | Clinical | 98.4 | 100 | 0 | 99.9822 | 5138713 | 45.3 | 60 | 394947 |
| VP18208 | VP18208 | SAMN20372076 | SRR15240269 | PC | Public | 2018 | China | Clinical | 98.3 | 100 | 0 | 99.974 | 5188887 | 45.29 | 59 | 401611 |
| VP18210 | VP18210 | SAMN20372115 | SRR15240161 | PC | Public | 2018 | China | Clinical | 98.4 | 100 | 0 | 99.9793 | 5098528 | 45.28 | 59 | 425769 |
| VP18211 | VP18211 | SAMN20372150 | SRR15240122 | PC | Public | 2018 | China | Clinical | 98.3 | 100 | 0 | 99.9683 | 5144459 | 45.29 | 62 | 334357 |
| VP18212 | VP18212 | SAMN20372148 | SRR15240124 | PC | Public | 2018 | China | Clinical | 98.3 | 100 | 0 | 99.9767 | 5143506 | 45.29 | 56 | 401470 |
| VP18213 | VP18213 | SAMN20372146 | SRR15240126 | NonPC | Public | 2018 | China | Clinical | 91.9 | 100 | 0.46 | 98.253 | 5113577 | 45.19 | 51 | 450397 |
| VP18214 | VP18214 | SAMN20372147 | SRR15240125 | PC | Public | 2018 | China | Clinical | 98.3 | 100 | 0 | 99.9749 | 5142544 | 45.29 | 56 | 461740 |
| VP18215 | VP18215 | SAMN20372131 | SRR15240144 | PC | Public | 2018 | China | Clinical | 98.3 | 100 | 0 | 99.9627 | 5193394 | 45.32 | 63 | 410725 |
| VP18216 | VP18216 | SAMN20372149 | SRR15240123 | PC | Public | 2018 | China | Clinical | 98.5 | 100 | 0 | 99.9839 | 5092409 | 45.29 | 56 | 425307 |

| | | | | | | | | | | | | | | | | |
|---|---|---|---|---|---|---|---|---|---|---|---|---|---|---|---|---|
| VP18218 | VP18218 | SAMN20372140 | SRR15240133 | PC | Public | 2018 | China | Clinical | 98.4 | 100 | 0 | 99.973 | 5108465 | 45.28 | 52 | 425883 |
| VP18219 | VP18219 | SAMN20372154 | SRR15240117 | NonPC | Public | 2018 | China | Clinical | 93.1 | 100 | 0.05 | 98.3615 | 5091559 | 45.27 | 50 | 463551 |
| VP18220 | VP18220 | SAMN20372152 | SRR15240120 | NonPC | Public | 2018 | China | Clinical | 93.1 | 100 | 0.05 | 98.3997 | 5075991 | 45.3 | 47 | 463552 |
| VP18221 | VP18221 | SAMN20372153 | SRR15240119 | PC | Public | 2018 | China | Clinical | 98.3 | 100 | 0 | 99.9828 | 5108292 | 45.28 | 55 | 401470 |
| VP18222 | VP18222 | SAMN20372145 | SRR15240127 | NonPC | Public | 2018 | China | Clinical | 92 | 100 | 0.47 | 98.262 | 5163199 | 45.22 | 55 | 314178 |
| VP18223 | VP18223 | SAMN20372158 | SRR15240113 | PC | Public | 2018 | China | Clinical | 98.4 | 100 | 0 | 99.9731 | 5109492 | 45.28 | 51 | 425241 |
| VP18224 | VP18224 | SAMN20372157 | SRR15240114 | PC | Public | 2018 | China | Clinical | 98.4 | 100 | 0 | 99.9754 | 5109212 | 45.28 | 54 | 425355 |
| VP18225 | VP18225 | SAMN20372155 | SRR15240116 | NonPC | Public | 2018 | China | Clinical | 91.5 | 100 | 0.07 | 98.365 | 5155596 | 45.25 | 38 | 476629 |
| VP18226 | VP18226 | SAMN20372151 | SRR15240121 | NonPC | Public | 2018 | China | Clinical | 93.1 | 100 | 0.05 | 98.3969 | 5059508 | 45.3 | 45 | 463546 |
| VP18227 | VP18227 | SAMN20372143 | SRR15240130 | PC | Public | 2018 | China | Clinical | 98.4 | 100 | 0 | 99.9905 | 5091708 | 45.29 | 55 | 425307 |
| VP18228 | VP18228 | SAMN20372160 | SRR15240111 | NonPC | Public | 2018 | China | Clinical | 92 | 100 | 0.46 | 98.2481 | 5113872 | 45.19 | 54 | 450300 |
| VP18229 | VP18229 | SAMN20372159 | SRR15240112 | PC | Public | 2018 | China | Clinical | 97.9 | 100 | 0 | 99.9646 | 5136973 | 45.27 | 57 | 401470 |
| VP18231 | VP18231 | SAMN20372124 | SRR15240151 | PC | Public | 2018 | China | Clinical | 98.4 | 100 | 0 | 99.9756 | 5116727 | 45.29 | 58 | 394941 |
| VP18232 | VP18232 | SAMN20372161 | SRR15240110 | PC | Public | 2018 | China | Clinical | 98.4 | 100 | 0 | 99.991 | 5089618 | 45.29 | 57 | 425355 |
| VP18233 | VP18233 | SAMN20372162 | SRR15240109 | NonPC | Public | 2018 | China | Clinical | 92.3 | 100 | 0.19 | 98.3438 | 5027339 | 45.24 | 54 | 471370 |
| VP18234 | VP18234 | SAMN20372163 | SRR15240108 | NonPC | Public | 2018 | China | Clinical | 92 | 100 | 0.46 | 98.2495 | 5075726 | 45.2 | 53 | 450300 |
| VP18235 | VP18235 | SAMN20372164 | SRR15240106 | PC | Public | 2018 | China | Clinical | 98.4 | 100 | 0.7 | 99.9909 | 5192439 | 45.28 | 62 | 394786 |
| VP18236 | VP18236 | SAMN20372173 | SRR15240097 | PC | Public | 2018 | China | Clinical | 98.4 | 100 | 0 | 99.9828 | 5090359 | 45.29 | 57 | 425343 |
| VP18237 | VP18237 | SAMN20372172 | SRR15240098 | PC | Public | 2018 | China | Clinical | 98.4 | 100 | 0 | 99.9888 | 5089090 | 45.29 | 56 | 425343 |
| VP18238 | VP18238 | SAMN20372170 | SRR15240100 | PC | Public | 2018 | China | Clinical | 98.4 | 100 | 0 | 99.9875 | 5090185 | 45.29 | 56 | 395938 |
| VP18239 | VP18239 | SAMN20372171 | SRR15240099 | PC | Public | 2018 | China | Clinical | 98.4 | 100 | 0 | 99.9789 | 5091784 | 45.29 | 60 | 394895 |
| VP18240 | VP18240 | SAMN20372168 | SRR15240102 | NonPC | Public | 2018 | China | Env | 90.9 | 100 | 0.27 | 98.355 | 4953008 | 45.44 | 48 | 436967 |
| VP18241 | VP18241 | SAMN20372169 | SRR15240101 | NonPC | Public | 2018 | China | Env | 90.2 | 100 | 0.03 | 98.2664 | 5143667 | 45.23 | 81 | 334330 |
| VP18242 | VP18242 | SAMN20372166 | SRR15240104 | NonPC | Public | 2018 | China | Clinical | 92 | 100 | 0.03 | 98.4155 | 4985542 | 45.35 | 65 | 295107 |
| VP18243 | VP18243 | SAMN20372167 | SRR15240103 | PC | Public | 2018 | China | Clinical | 98.4 | 100 | 0 | 99.9918 | 5090865 | 45.29 | 61 | 394869 |
| VP18244 | VP18244 | SAMN20372174 | SRR15240028 | NonPC | Public | 2018 | China | Clinical | 92.3 | 100 | 0.19 | 98.3439 | 5040146 | 45.23 | 52 | 450299 |
| VP18245 | VP18245 | SAMN20372165 | SRR15240105 | PC | Public | 2018 | China | Clinical | 97.6 | 100 | 0 | 99.9376 | 5154856 | 45.25 | 63 | 313341 |
| VP18246 | VP18246 | SAMN20372176 | SRR15240026 | NonPC | Public | 2018 | China | Clinical | 90.9 | 100 | 0 | 98.2429 | 5156581 | 45.25 | 61 | 404870 |
| VP18248 | VP18248 | SAMN20372175 | SRR15240027 | NonPC | Public | 2018 | China | Clinical | 91.9 | 100 | 0.46 | 98.2558 | 5074140 | 45.2 | 57 | 450300 |
| Vp_UK_AB_S24 | AB_S24 | TBD | TBD | PC | Baker-Austin et al, Food Control, 2020 | 2016 | UK(Colombia) | Clinical | 98.3 | 100 | 0 | 99.9506 | 5156445 | 45.28 | 74 | 402331 |
| Vp_UK_AD_S26 | AD_S26 | TBD | TBD | PC | Baker-Austin et al, Food Control, 2020 | 2015 | UK | Clinical | 98.3 | 100 | 0 | 99.9536 | 5149586 | 45.28 | 69 | 402331 |
| Vp_UK_AE_S27 | AE_S27 | TBD | TBD | PC | Baker-Austin et al, Food Control, 2020 | NA | UK | Clinical | 98.5 | 100 | 0 | 99.9885 | 5135348 | 45.32 | 48 | 404897 |
| Vp_UK_AF_S28 | AF_S28 | TBD | TBD | PC | Baker-Austin et al, Food Control, 2020 | NA | UK | Clinical | 98.7 | 100 | 0 | 99.9834 | 5188523 | 45.29 | 72 | 302290 |
| Vp_UK_AJ_S32 | AJ_S32 | TBD | TBD | PC | Baker-Austin et al, Food Control, 2020 | 2016 | UK(Kenya) | Clinical | 98.2 | 100 | 0.07 | 99.9006 | 5203033 | 45.28 | 56 | 476668 |
| Vp_UK_AL_S34 | AL_S34 | TBD | TBD | PC | Baker-Austin et al, Food Control, 2020 | 2015 | UK | Clinical | 98.9 | 100 | 0 | 99.9908 | 5224097 | 45.24 | 56 | 428496 |
| Vp_UK_B_S2 | B_S2 | TBD | TBD | PC | Baker-Austin et al, Food Control, 2020 | 2009 | UK(Vietnam) | Clinical | 98.8 | 100 | 0 | 99.9897 | 5120139 | 45.31 | 53 | 426609 |
| Vp_UK_C_S3 | C_S3 | TBD | TBD | PC | Baker-Austin et al, Food Control, 2020 | 2009 | UK(Vietnam) | Clinical | 99 | 100 | 0 | 99.9917 | 5177871 | 45.29 | 58 | 428496 |
| Vp_UK_D_S4 | D_S4 | TBD | TBD | PC | Baker-Austin et al, Food Control, 2020 | 2013 | UK(Thailand) | Clinical | 98.9 | 100 | 0 | 99.9816 | 5204074 | 45.29 | 57 | 432394 |
| Vp_UK_F_S6 | F_S6 | TBD | TBD | PC | Baker-Austin et al, Food Control, 2020 | 2010 | UK(Thailand) | Clinical | 98.7 | 100 | 0 | 99.9894 | 5147208 | 45.29 | 97 | 402331 |
| Vp_UK_H_S8 | H_S8 | TBD | TBD | PC | Baker-Austin et al, Food Control, 2020 | 2014 | UK(Cuba) | Clinical | 98.3 | 99.97 | 0 | 99.9881 | 5095355 | 45.33 | 49 | 540460 |
| Vp_UK_Hist_Env_P860 | Hist_Env_P860 | TBD | TBD | PC | Baker-Austin et al, Food Control, 2020 | NA | UK | NA | 97.8 | 100 | 0.27 | 99.9914 | 5084286 | 45.29 | 147 | 95820 |
| Vp_UK_Hist_Env_V12-024 | Hist_Env_V12-024 | TBD | TBD | PC | Baker-Austin et al, Food Control, 2020 | 2012 | UK | Clinical | 98.3 | 100 | 0.2 | 99.9856 | 5103231 | 45.27 | 84 | 125273 |
| Vp_UK_K_S10 | K_S10 | TBD | TBD | PC | Baker-Austin et al, Food Control, 2020 | 2014 | UK(Thailand) | Clinical | 96.8 | 100 | 0.01 | 99.8968 | 5162904 | 45.36 | 58 | 402256 |
| Vp_UK_L_S11 | L_S11 | TBD | TBD | PC | Baker-Austin et al, Food Control, 2020 | 2014 | UK(Thailand) | Clinical | 98 | 100 | 0.01 | 99.9066 | 5224518 | 45.34 | 61 | 443424 |
| Vp_UK_R_S14 | R_S14 | TBD | TBD | PC | Baker-Austin et al, Food Control, 2020 | 2013 | UK | Clinical | 98.5 | 100 | 0 | 99.9901 | 5112001 | 45.31 | 80 | 187082 |
| Vp_UK_S_S15 | S_S15 | TBD | TBD | PC | Baker-Austin et al, Food Control, 2020 | 2011 | UK(Thailand) | Clinical | 98.1 | 99.46 | 0 | 99.99 | 5142335 | 45.3 | 58 | 342419 |
| Vp_UK_V_S18 | V_S18 | TBD | TBD | PC | Baker-Austin et al, Food Control, 2020 | 2016 | UK(Thailand) | Clinical | 98.7 | 100 | 0 | 99.9898 | 5124338 | 45.3 | 69 | 538271 |
| Vp_UK_W_S19 | W_S19 | TBD | TBD | PC | Baker-Austin et al, Food Control, 2020 | NA | UK | Clinical | 98.8 | 100 | 0.01 | 99.9891 | 5270241 | 45.34 | 66 | 428628 |

| | | | | | | | | | | | | | | | | |
|---|---|---|---|---|---|---|---|---|---|---|---|---|---|---|---|---|
| Vp_UK_Y_S21 | Y_S21 | TBD | TBD | PC | Baker-Austin et al, Food Control, 2020 | 2015 | UK | Clinical | 98.7 | 100 | 0.01 | 99.9895 | 5268327 | 45.33 | 68 | 428592 |
