## Supplementary Table 2 for "Wave succession in the pandemic clone of *Vibrio parahaemolyticus* driven by gene loss"

### Supplementary Table 2: Metadata and Classification Information of *Vibrio parahaemolyticus* Pre-Pandemic and Pandemic Clone Strains.

| ID | Isolate_identifiers | Year | Country | Isolation_type | Geographical region | Serotype | ST | Classification | Classification detail | PC-representative |
|---|---|---|---|---|---|---|---|---|---|---|
| SAMN06077000-GCA_015818905.1 | CFSAN023549 | 1980 | Bangladesh | Clinical | Asia | O3:K6 | ST87 | Pre-PC | Pre-PC1 | |
| SAMN02338872-GCF_000491955.1 | S011 | 1990 | Thailand | Clinical | Asia | O3:K6 | ST- | Pre-PC | Pre-PC2 | |
| SAMN02338873-GCF_000491935.1 | S012 | 1990 | Thailand | Clinical | Asia | O3:K6 | ST- | Pre-PC | Pre-PC2 | |
| AQ4235_NA_Japan_NA_NA | AQ4235 | 1987 | Japan(Thailand) | Clinical | Asia | O3:K6 | ST91 | Pre-PC | Pre-PC2 | |
| AQ4853_NA_Japan_NA_NA | AQ4853 | 1993 | Japan(HongKong) | Clinical | Asia | O3:K6 | ST91 | Pre-PC | Pre-PC2 | |
| AQ4901_NA_Japan_NA_NA | AQ4901 | 1993 | Japan(Thailand) | Clinical | Asia | O3:K6 | ST91 | Pre-PC | Pre-PC2 | |
| SAMN02338867-GCF_000492055.1 | S003 | NA | China | Clinical | Asia | O3:K6 | ST91 | Pre-PC | Pre-PC2 | |
| SAMN02338869-GCF_000492015.1 | S005 | NA | Thailand | Clinical | Asia | O3:K6 | ST91 | Pre-PC | Pre-PC2 | |
| SAMN02338870-GCF_000491995.1 | S008 | 1987 | Thailand | Clinical | Asia | O3:K6 | ST91 | Pre-PC | Pre-PC2 | |
| SAMN02338871-GCF_000491975.1 | S009 | 1987 | Thailand | Clinical | Asia | O3:K6 | ST91 | Pre-PC | Pre-PC2 | |
| SAMN02338874-GCF_000491915.1 | S013 | 1996 | China | Clinical | Asia | O3:K65 | ST91 | Pre-PC | Pre-PC2 | |
| SAMN02338948-GCF_000490445.1 | S095 | 1996 | China | Clinical | Asia | O3:K6 | ST91 | Pre-PC | Pre-PC2 | |
| VP08414 | VP08414 | 2008 | China | NA | Asia | O3:K6 | ST91 | Pre-PC | Pre-PC2 | |
| AQ4019_NA_Japan_NA_NA | AQ4019 | 1985 | Japan(Maldive) | Clinical | Asia | O3:K6 | ST96 | Pre-PC | Pre-PC2 | |
| AQ4037_NA_Japan_NA_NA | AQ4037 | 1985 | Japan(Maldive) | Clinical | Asia | O3:K6 | ST96 | Pre-PC | Pre-PC2 | |
| AQ4093_NA_Japan_NA_NA | AQ4093 | 1986 | Japan(Maldive) | Clinical | Asia | O3:K6 | ST96 | Pre-PC | Pre-PC2 | |
| SAMN02338868-GCF_000492035.1 | S004 | 1985 | Maldives | Clinical | Asia | O3:K6 | ST96 | Pre-PC | Pre-PC2 | |
| SAMN07338226-GCF_006368255.1 | S010 | NA | NA | NA | NA | O3:K6 | ST- | Pre-PC | Pre-PC2 | |
| SAMN07338233-GCF_006368175.1 | S177 | NA | NA | NA | NA | O3:K6 | ST96 | Pre-PC | Pre-PC2 | |
| VP22_4B-8 | VP22_4B-8 | 2022 | China | Env | Asia | O3:K6 | ST2251 | Pre-PC | Pre-PC3a | |
| SAMN17832008-GCA_019271155.1 | Colony550 | NA | Thailand | Env | Asia | OUK3:K26 | ST672 | Pre-PC | Pre-PC3a | |
| VP15194 | VP15194 | 2015 | China | Clinical | Asia | O3:K6 | ST672 | Pre-PC | Pre-PC3a | |
| SPVP140225 | SPVP140225 | 2014 | China | Env | Asia | OUK1:K43 | ST- | Pre-PC | Pre-PC3b | |
| SAMN02712354-GCF_000958595.1 | AF91 | 2006 | USA | Env | US_Canada | O3:K6 | ST- | Pre-PC | Pre-PC4a | |
| SAMN09839615-GCA_015801235.1 | AK1822500155 | 2018 | USA | Env | US_Canada | O3:K6 | ST- | Pre-PC | Pre-PC4a | |
| SAMN12842757-GCA_015783205.1 | PNUSAV001026 | NA | USA | Clinical | US_Canada | O3:K6 | ST- | Pre-PC | Pre-PC4a | |
| SAMEA8103019-GCA_905332005.1 | ERS5790097 | 1980 | Japan | Clinical | Asia | O3:K6 | ST3 | Pre-PC | Pre-PC4b | |
| SAMN02338946-GCF_000490485.1 | S093 | 1998 | Japan | Clinical | Asia | O3:K65 | ST3 | Pre-PC | Pre-PC4b | |
| SAMN07337971-GCF_006374305.1 | C1_10 | 2014 | China | Env | Asia | O3:K6 | ST3 | Pre-PC | Pre-PC4b | |

| | | | | | | | | | | |
|---|---|---|---|---|---|---|---|---|---|---|
| SAMN13808684-GCA_015775455.1 | CFSAN095296 | 2013 | USA | Env | US_Canada | O3:K6 | ST- | Pre-PC | Pre-PC4b | |
| SAMN08469472-GCA_015811985.1 | CFSAN075738 | 2012 | USA | Env | US_Canada | O3:K6 | ST3 | Pre-PC | Pre-PC4b | |
| SAMN08469562-GCA_015812005.1 | CFSAN075750 | 2012 | USA | Env | US_Canada | O3:K6 | ST3 | Pre-PC | Pre-PC4b | |
| SAMN08902707-GCA_015810665.1 | CFSAN079455 | 2012 | USA | Env | US_Canada | O3:K6 | ST3 | Pre-PC | Pre-PC4b | |
| SAMN08902738-GCA_015810765.1 | CFSAN079437 | 2012 | USA | Env | US_Canada | O3:K65 | ST3 | Pre-PC | Pre-PC4b | |
| SAMN08902775-GCA_015810545.1 | CFSAN079443 | 2012 | USA | Env | US_Canada | O3:K6 | ST3 | Pre-PC | Pre-PC4b | |
| SAMN08965140-GCA_015807555.1 | CFSAN080155 | 2013 | USA | Env | US_Canada | O3:K6 | ST3 | Pre-PC | Pre-PC4b | |
| SAMN09074611-GCA_015808055.1 | CFSAN079428 | 2012 | USA | Env | US_Canada | O3:K6 | ST3 | Pre-PC | Pre-PC4b | |
| SAMN09242511-GCA_015806975.1 | CFSAN081454 | 2013 | USA | Env | US_Canada | O3:K6 | ST3 | Pre-PC | Pre-PC4b | |
| SAMN09242527-GCA_015805395.1 | CFSAN081437 | 2013 | USA | Env | US_Canada | O3:K65 | ST3 | Pre-PC | Pre-PC4b | |
| 405-00_NA_Peru_Clin_Clin | 405-00 | 2000 | Peru | Clinical | Latin America | O3:K6 | ST3 | PC | Wave-1 | Putative mutator |
| 518-01_NA_Peru_Clin_Clin | 518-01 | 2001 | Peru | Clinical | Latin America | O3:K6 | ST3 | PC | Wave-1 | Putative mutator |
| VP10447 | VP10447 | 2010 | China | Clinical | Asia | O3:K6 | ST3 | PC | Wave-1 | Putative outbreak cluster |
| VPCZ66 | VPCZ66 | 2013 | China | Clinical | Asia | O3:K6 | ST3 | PC | Wave-1 | Putative outbreak cluster |
| 461-00_NA_Peru_Clin_Clin | 461-00 | 2000 | Peru | Clinical | Latin America | O3:K6 | ST3 | PC | Wave-1 | Putative outbreak cluster |
| 49_5-V2_NA_Chile_NA_NA | 49_5-V2 | 2005 | Chile | NA | Latin America | O3:K6 | ST3 | PC | Wave-1 | Putative outbreak cluster |
| 57_5_NA_Chile_NA_NA | 57_5 | 2005 | Chile | NA | Latin America | O3:K6 | ST3 | PC | Wave-1 | Putative outbreak cluster |
| 875-97_NA_Peru_Clin_Clin | 875-97 | 1997 | Peru | Clinical | Latin America | O3:K6 | ST3 | PC | Wave-1 | Putative outbreak cluster |
| PMC18_7_NA_Chile_NA_NA | PMC18 | 2007 | Chile | NA | Latin America | O3:K6 | ST3 | PC | Wave-1 | Putative outbreak cluster |
| PMC28_7_NA_Chile_Amb_Amb | PMC28 | 2007 | Chile | Env | Latin America | O3:K6 | ST3 | PC | Wave-1 | Putative outbreak cluster |
| PMC50_7_NA_Chile_NA_NA | PMC50 | 2007 | Chile | NA | Latin America | O3:K6 | ST3 | PC | Wave-1 | Putative outbreak cluster |
| PMC51_7_NA_Chile_NA_NA | PMC51 | 2007 | Chile | NA | Latin America | O3:K6 | ST3 | PC | Wave-1 | Putative outbreak cluster |
| PMC60_7_NA_Chile_Amb_Amb | PMC60 | 2007 | Chile | Env | Latin America | O3:K6 | ST3 | PC | Wave-1 | Putative outbreak cluster |
| PMC70_7_NA_Chile_NA_NA | PMC70 | 2007 | Chile | NA | Latin America | O3:K6 | ST3 | PC | Wave-1 | Putative outbreak cluster |
| PMC73_7_NA_Chile_Amb_Amb | PMC73 | 2007 | Chile | Env | Latin America | O3:K6 | ST3 | PC | Wave-1 | Putative outbreak cluster |
| SAMN02436220-GCF_000182345.1 | Peru-466 | 1996 | Peru | Clinical | Latin America | O3:K6 | ST3 | PC | Wave-1 | Putative outbreak cluster |
| SAMN02781341-GCA_001270835.1 | PMA37.5 | 2005 | Chile | Env | Latin America | O3:K6 | ST3 | PC | Wave-1 | Putative outbreak cluster |
| SAMN08225457-GCA_015814805.1 | CFSAN029655 | 2009 | Peru | Env | Latin America | O3:K6 | ST3 | PC | Wave-1 | Putative outbreak cluster |
| SAMN08225458-GCA_015814745.1 | CFSAN029652 | 2009 | Peru | Env | Latin America | O3:K6 | ST3 | PC | Wave-1 | Putative outbreak cluster |
| SAMN12364841-GCF_009991745.1 | 249-15 | 2015 | Peru | Clinical | Latin America | O3:K6 | ST3 | PC | Wave-1 | Putative outbreak cluster |
| SAMN12364842-GCF_009991805.1 | 276-15 | 2015 | Peru | Clinical | Latin America | O3:K6 | ST3 | PC | Wave-1 | Putative outbreak cluster |
| SAMN12364846-GCF_009991685.1 | 164-16 | 2016 | Peru | Clinical | Latin America | O3:K6 | ST3 | PC | Wave-1 | Putative outbreak cluster |

| | | | | | | | | | | |
|---|---|---|---|---|---|---|---|---|---|---|
| SAMN15428938-GCF_016880155.1 | 404-00 | 2000 | Peru | Clinical | Latin America | O3:K6 | ST3 | PC | Wave-1 | Putative outbreak cluster |
| SAMN15428941-GCF_016880075.1 | 403-00 | 2000 | Peru | Clinical | Latin America | O3:K6 | ST3 | PC | Wave-1 | Putative outbreak cluster |
| SAMN15428943-GCF_016880035.1 | 1027-00 | 2000 | Peru | Clinical | Latin America | O3:K6 | ST3 | PC | Wave-1 | Putative outbreak cluster |
| SAMN15428944-GCF_016880115.1 | 2434-00 | 2000 | Peru | Clinical | Latin America | O3:K6 | ST3 | PC | Wave-1 | Putative outbreak cluster |
| SAMN15428948-GCF_016878735.1 | 1259-07 | 2007 | Peru | Clinical | Latin America | O3:K6 | ST3 | PC | Wave-1 | Putative outbreak cluster |
| SAMN15428950-GCF_016878715.1 | 1254-07 | 2007 | Peru | Clinical | Latin America | O3:K6 | ST3 | PC | Wave-1 | Putative outbreak cluster |
| SAMN15428952-GCF_016878675.1 | 243-09 | 2009 | Peru | Clinical | Latin America | O3:K6 | ST3 | PC | Wave-1 | Putative outbreak cluster |
| SAMN15428963-GCF_016878435.1 | 203-10 | 2010 | Peru | Clinical | Latin America | O3:K6 | ST3 | PC | Wave-1 | Putative outbreak cluster |
| SAMN15428964-GCF_016878455.1 | 091-10 | 2010 | Peru | Clinical | Latin America | O3:K6 | ST3 | PC | Wave-1 | Putative outbreak cluster |
| SAMN15428965-GCF_016878395.1 | 304-10 | 2010 | Peru | Clinical | Latin America | O3:K6 | ST3 | PC | Wave-1 | Putative outbreak cluster |
| SAMN15428966-GCF_016878405.1 | 293-10 | 2010 | Peru | Clinical | Latin America | O3:K6 | ST3 | PC | Wave-1 | Putative outbreak cluster |
| SAMN15428967-GCF_016878375.1 | 202-10 | 2010 | Peru | Clinical | Latin America | O3:K6 | ST3 | PC | Wave-1 | Putative outbreak cluster |
| SAMN15428968-GCF_016878355.1 | 092-10 | 2010 | Peru | Clinical | Latin America | O3:K6 | ST3 | PC | Wave-1 | Putative outbreak cluster |
| SAMN15428969-GCF_016878325.1 | 361-10 | 2010 | Peru | Clinical | Latin America | O3:K6 | ST3 | PC | Wave-1 | Putative outbreak cluster |
| SAMN15428970-GCF_016878315.1 | 454-10 | 2010 | Peru | Clinical | Latin America | O3:K6 | ST3 | PC | Wave-1 | Putative outbreak cluster |
| SAMN15428971-GCF_016878295.1 | 119-10 | 2010 | Peru | Clinical | Latin America | O3:K6 | ST3 | PC | Wave-1 | Putative outbreak cluster |
| SAMN15428974-GCF_016878215.1 | 1218-11 | 2011 | Peru | Clinical | Latin America | O3:K6 | ST3 | PC | Wave-1 | Putative outbreak cluster |
| SAMN15428982-GCF_016878075.1 | 277-15 | 2015 | Peru | Clinical | Latin America | O3:K6 | ST3 | PC | Wave-1 | Putative outbreak cluster |
| SAMN15429001-GCF_016877715.1 | 167-16 | 2016 | Peru | Clinical | Latin America | O3:K6 | ST3 | PC | Wave-1 | Putative outbreak cluster |
| SAMN15429002-GCF_016877675.1 | 165-16 | 2016 | Peru | Clinical | Latin America | O3:K6 | ST3 | PC | Wave-1 | Putative outbreak cluster |
| SAMN15429003-GCF_016877655.1 | G8 | 2016 | Peru | Clinical | Latin America | O3:K6 | ST3 | PC | Wave-1 | Putative outbreak cluster |
| Vp_Peru_568-01_Peru_2001_C | 568-01 | 2001 | Peru | Clinical | Latin America | O3:K6 | ST3 | PC | Wave-1 | Putative outbreak cluster |
| Vp_Peru_CFSAN018757_1997_Peru_C | CFSAN018757 | 1997 | Peru | Clinical | Latin America | O3:K6 | ST3 | PC | Wave-1 | Putative outbreak cluster |
| Vp_Peru_CFSAN062243_2000_Peru_C | CFSAN062243 | 2000 | Peru | Clinical | Latin America | O3:K6 | ST3 | PC | Wave-1 | Putative outbreak cluster |
| Vp_Peru_CFSAN062252_2007_Peru_C | CFSAN062252 | 2007 | Peru | Clinical | Latin America | O3:K6 | ST3 | PC | Wave-1 | Putative outbreak cluster |
| Vp_Peru_CFSAN062257_2000_Peru_C | CFSAN062257 | 2000 | Peru | Clinical | Latin America | O3:K6 | ST3 | PC | Wave-1 | Putative outbreak cluster |
| Vp_Peru_CFSAN062258_2007_Peru_C | CFSAN062258 | 2007 | Peru | Clinical | Latin America | O3:K6 | ST3 | PC | Wave-1 | Putative outbreak cluster |
| Vp_Peru_CFSAN062268_2007_Peru_C | CFSAN062268 | 2007 | Peru | Clinical | Latin America | O3:K6 | ST3 | PC | Wave-1 | Putative outbreak cluster |
| Vp_Peru_CFSAN062271_2015_Peru_C | CFSAN062271 | 2015 | Peru | Clinical | Latin America | O3:K6 | ST3 | PC | Wave-1 | Putative outbreak cluster |
| Vp_Peru_CFSAN062278_2010_Peru_C | CFSAN062278 | 2010 | Peru | Clinical | Latin America | O3:K6 | ST3 | PC | Wave-1 | Putative outbreak cluster |
| Vp_Peru_CFSAN062279_2015_Peru_C | CFSAN062279 | 2015 | Peru | Clinical | Latin America | O3:K6 | ST3 | PC | Wave-1 | Putative outbreak cluster |
| Vp_Peru_CFSAN062298_2015_Peru_C | CFSAN062298 | 2015 | Peru | Clinical | Latin America | O3:K6 | ST3 | PC | Wave-1 | Putative outbreak cluster |

| | | | | | | | | | | |
|---|---|---|---|---|---|---|---|---|---|---|
| Vp_Peru_CFSAN062299_2015_Peru_C | CFSAN062299 | 2015 | Peru | Clinical | Latin America | O3:K6 | ST3 | PC | Wave-1 | Putative outbreak cluster |
| Vp_Peru_CFSAN062327_2015_Peru_C | CFSAN062327 | 2015 | Peru | Clinical | Latin America | O3:K6 | ST3 | PC | Wave-1 | Putative outbreak cluster |
| Vp_Peru_CFSAN062329_2010_Peru_C | CFSAN062329 | 2010 | Peru | Clinical | Latin America | O3:K6 | ST3 | PC | Wave-1 | Putative outbreak cluster |
| Vp_Peru_CFSAN062337_2010_Peru_C | CFSAN062337 | 2010 | Peru | Clinical | Latin America | O3:K6 | ST3 | PC | Wave-1 | Putative outbreak cluster |
| Vp_Peru_CFSAN062348_2016_Peru_C | CFSAN062348 | 2016 | Peru | Clinical | Latin America | O3:K6 | ST3 | PC | Wave-1 | Putative outbreak cluster |
| Vp_Peru_CFSAN062349_2016_Peru_C | CFSAN062349 | 2016 | Peru | Clinical | Latin America | O3:K6 | ST3 | PC | Wave-1 | Putative outbreak cluster |
| Vp_Peru_CFSAN062351_2015_Peru_C | CFSAN062351 | 2015 | Peru | Clinical | Latin America | O3:K6 | ST3 | PC | Wave-1 | Putative outbreak cluster |
| Vp_Peru_CFSAN062354_2010_Peru_C | CFSAN062354 | 2010 | Peru | Clinical | Latin America | O3:K6 | ST3 | PC | Wave-1 | Putative outbreak cluster |
| Vp_Peru_CFSAN062355_2010_Peru_C | CFSAN062355 | 2010 | Peru | Clinical | Latin America | O3:K6 | ST3 | PC | Wave-1 | Putative outbreak cluster |
| Vp_Peru_CFSAN062364_2016_Peru_C | CFSAN062364 | 2016 | Peru | Clinical | Latin America | O3:K6 | ST3 | PC | Wave-1 | Putative outbreak cluster |
| Vp_Peru_CFSAN062365_2016_Peru_C | CFSAN062365 | 2016 | Peru | Clinical | Latin America | O3:K6 | ST3 | PC | Wave-1 | Putative outbreak cluster |
| Vp_Peru_CFSAN062368_2010_Peru_C | CFSAN062368 | 2010 | Peru | Clinical | Latin America | O3:K6 | ST3 | PC | Wave-1 | Putative outbreak cluster |
| Vp_Peru_G7_2015_Lima_C | G7 | 2015 | Peru | Clinical | Latin America | O3:K6 | ST3 | PC | Wave-1 | Putative outbreak cluster |
| Vp_Peru_Peru-288_2001_Peru_C | Peru-288 | 2001 | Peru | Clinical | Latin America | O3:K6 | ST3 | PC | Wave-1 | Putative outbreak cluster |
| Vp_Peru_vp1027-00_2000_NA_C | vp1027-00 | 2000 | Peru | Clinical | Latin America | O3:K6 | ST3 | PC | Wave-1 | Putative outbreak cluster |
| Vp_Peru_vp1202-11_2011_Peru_Lima_C | vp1202-11 | 2011 | Peru | Clinical | Latin America | O3:K6 | ST3 | PC | Wave-1 | Putative outbreak cluster |
| Vp_Peru_vp1254-07_2007_NA_C | vp1254-07 | 2007 | Peru | Clinical | Latin America | O3:K6 | ST3 | PC | Wave-1 | Putative outbreak cluster |
| Vp_Peru_vp202-11_2010_Peru_Lima_C | vp202-11 | 2010 | Peru | Clinical | Latin America | O3:K6 | ST3 | PC | Wave-1 | Putative outbreak cluster |
| Vp_Peru_vp21260-07_2007_Peru_Lambayeque_C | vp21260-07 | 2007 | Peru | Clinical | Latin America | O3:K6 | ST3 | PC | Wave-1 | Putative outbreak cluster |
| Vp_Peru_vp2568-00_2000_Peru_Loreto_C | vp2568-00 | 2000 | Peru | Clinical | Latin America | O3:K6 | ST3 | PC | Wave-1 | Putative outbreak cluster |
| Vp_Peru_vp276-15_2015_Peru_Lima_C | vp276-15 | 2015 | Peru | Clinical | Latin America | O3:K6 | ST3 | PC | Wave-1 | Putative outbreak cluster |
| SAMN03358828-GCF_001727815.1 | BCW_3172 | 2006 | USA | Clinical | US_Canada | O3:K6 | ST3 | PC | Wave-1 | Putative outbreak cluster |
| LS326_NA_Senegal_Amb_Prawns | LS326 | NA | Senegal | Env | Africa | O3:K6 | ST3 | PC | Wave-1 | Representative |
| LS328_NA_Senegal_Amb_Prawns | LS328 | NA | Senegal | Env | Africa | O3:K6 | ST3 | PC | Wave-1 | Representative |
| LS329_NA_Senegal_Amb_Prawns | LS329 | NA | Senegal | Env | Africa | O3:K6 | ST3 | PC | Wave-1 | Representative |
| SAMN01933054-GCF_000593285.2 | PVCHO_VP-48 | 1996 | India | Clinical | Asia | O3:K6 | ST- | PC | Wave-1 | Representative |
| SAMN02338923-GCF_000490935.1 | S066 | 1997 | China | Clinical | Asia | O3:K6 | ST3 | PC | Wave-1 | Representative |
| SAMN02338924-GCF_000490915.1 | S067 | 1997 | China | Clinical | Asia | O3:K6 | ST3 | PC | Wave-1 | Representative |
| SAMN02338937-GCF_000490655.1 | S081 | NA | South_Korea | Clinical | Asia | O3:K6 | ST3 | PC | Wave-1 | Representative |
| VP10417 | VP10417 | 2010 | China | Clinical | Asia | O3:K6 | ST3 | PC | Wave-1 | Representative |
| VP10429 | VP10429 | 2010 | China | Clinical | Asia | O3:K6 | ST3 | PC | Wave-1 | Representative |

| | | | | | | | | | | |
|---|---|---|---|---|---|---|---|---|---|---|
| VP12069 | VP12069 | 2012 | China | Clinical | Asia | O3:K6 | ST3 | PC | Wave-1 | Representative |
| VP12205 | VP12205 | 2012 | China | Clinical | Asia | O3:K6 | ST3 | PC | Wave-1 | Representative |
| VPCZ27 | VPCZ27 | 2013 | China | Clinical | Asia | O3:K6 | ST3 | PC | Wave-1 | Representative |
| VPCZ28 | VPCZ28 | 2014 | China | Clinical | Asia | O3:K6 | ST3 | PC | Wave-1 | Representative |
| SAMEA5540086-ERR3255956 | ERS3342147 | NA | UK | NA | Europe | O3:K6 | ST3 | PC | Wave-1 | Representative |
| SAMN06214611-GCA_015817515.1 | AMC_317 | 2016 | Spain | Clinical | Europe | O3:K6 | ST3 | PC | Wave-1 | Representative |
| SAMN07185550-GCA_015817425.1 | CFSAN029651 | 2010 | UK | Env | Europe | O3:K6 | ST3 | PC | Wave-1 | Representative |
| Vp_UK_Hist_Env_P860 | Hist_Env_P860 | NA | UK | NA | Europe | O3:K6 | ST3 | PC | Wave-1 | Representative |
| Vp_UK_Hist_Env_V12-024 | Hist_Env_V12-024 | 2012 | UK | Clinical | Europe | O3:K6 | ST3 | PC | Wave-1 | Representative |
| Vp_UK_W_S19 | W_S19 | NA | UK | Clinical | Europe | O3:K6 | ST3 | PC | Wave-1 | Representative |
| Vp_UK_Y_S21 | Y_S21 | 2015 | UK | Clinical | Europe | O3:K6 | ST3 | PC | Wave-1 | Representative |
| 004-02_NA_Peru_Clin_Clin | 004-02 | 2002 | Peru | Clinical | Latin America | O3:K6 | ST3 | PC | Wave-1 | Representative |
| 020-02_NA_Peru_Clin_Clin | 020-02 | 2002 | Peru | Clinical | Latin America | O3:K6 | ST3 | PC | Wave-1 | Representative |
| 056-01_NA_Peru_Clin_Clin | 056-01 | 2001 | Peru | Clinical | Latin America | O3:K6 | ST3 | PC | Wave-1 | Representative |
| 085-02_NA_Peru_Clin_Clin | 085-02 | 2002 | Peru | Clinical | Latin America | O3:K6 | ST3 | PC | Wave-1 | Representative |
| 182-02_NA_Peru_Clin_Clin | 182-02 | 2002 | Peru | Clinical | Latin America | O3:K6 | ST3 | PC | Wave-1 | Representative |
| 210_NA_Chile_NA_NA | 210 | NA | Chile | NA | Latin America | O3:K6 | ST3 | PC | Wave-1 | Representative |
| 224_NA_Chile_NA_NA | 224 | NA | Chile | NA | Latin America | O3:K6 | ST3 | PC | Wave-1 | Representative |
| 240-02_NA_Peru_Clin_Clin | 240-02 | 2002 | Peru | Clinical | Latin America | O3:K6 | ST3 | PC | Wave-1 | Representative |
| 275-99_NA_Peru_Clin_Clin | 275-99 | 1999 | Peru | Clinical | Latin America | O3:K6 | ST3 | PC | Wave-1 | Representative |
| 276-99_NA_Peru_Clin_Clin | 276-99 | 1999 | Peru | Clinical | Latin America | O3:K6 | ST3 | PC | Wave-1 | Representative |
| 278-99_NA_Peru_Clin_Clin | 278-99 | 1999 | Peru | Clinical | Latin America | O3:K6 | ST3 | PC | Wave-1 | Representative |
| 357-99_NA_Peru_Clin_Clin | 357-99 | 1999 | Peru | Clinical | Latin America | O3:K6 | ST3 | PC | Wave-1 | Representative |
| 39_5_NA_Chile_NA_NA | 39_5 | 2005 | Chile | NA | Latin America | O3:K6 | ST3 | PC | Wave-1 | Representative |
| 429-00_NA_Peru_Clin_Clin | 429-00 | 2000 | Peru | Clinical | Latin America | O3:K6 | ST3 | PC | Wave-1 | Representative |
| 462-00_NA_Peru_Clin_Clin | 462-00 | 2000 | Peru | Clinical | Latin America | O3:K6 | ST3 | PC | Wave-1 | Representative |
| 463-01_NA_Peru_Clin_Clin | 463-01 | 2001 | Peru | Clinical | Latin America | O3:K6 | ST3 | PC | Wave-1 | Representative |
| 48_1_NA_Chile_NA_NA | 48_1 | 2001 | Chile | NA | Latin America | O3:K6 | ST3 | PC | Wave-1 | Representative |
| 511-00_NA_Peru_Clin_Clin | 511-00 | 2000 | Peru | Clinical | Latin America | O3:K6 | ST3 | PC | Wave-1 | Representative |
| 568-00_NA_Peru_Clin_Clin | 568-00 | 2000 | Peru | Clinical | Latin America | O3:K6 | ST3 | PC | Wave-1 | Representative |
| 572-01_NA_Peru_Clin_Clin | 572-01 | 2001 | Peru | Clinical | Latin America | O3:K6 | ST3 | PC | Wave-1 | Representative |
| 61_NA_Chile_Amb_Amb | 61 | NA | Chile | NA | Latin America | O3:K6 | ST3 | PC | Wave-1 | Representative |

| | | | | | | | | | | |
|---|---|---|---|---|---|---|---|---|---|---|
| 61-V2_NA_Chile_Amb_NA | 61-V2 | NA | Chile | NA | Latin America | O3:K6 | ST3 | PC | Wave-1 | Representative |
| 698-99_NA_Peru_Clin_Clin | 698-99 | 1999 | Peru | Clinical | Latin America | O3:K6 | ST- | PC | Wave-1 | Representative |
| 706-00_NA_Peru_Clin_Clin | 706-00 | 2000 | Peru | Clinical | Latin America | O3:K6 | ST3 | PC | Wave-1 | Representative |
| 763-97_NA_Peru_Clin_Clin | 763-97 | 1997 | Peru | Clinical | Latin America | O3:K6 | ST3 | PC | Wave-1 | Representative |
| 776-00_NA_Peru_Clin_Clin | 776-00 | 2000 | Peru | Clinical | Latin America | O3:K6 | ST3 | PC | Wave-1 | Representative |
| 780-98_NA_Peru_Clin_Clin | 780-98 | 1998 | Peru | Clinical | Latin America | O3:K6 | ST3 | PC | Wave-1 | Representative |
| 784-98_NA_Peru_Clin_Clin | 784-98 | 1998 | Peru | Clinical | Latin America | O3:K6 | ST3 | PC | Wave-1 | Representative |
| 790-97_NA_Peru_Clin_Clin | 790-97 | 1997 | Peru | Clinical | Latin America | O3:K6 | ST3 | PC | Wave-1 | Representative |
| 859-07_NA_Peru_Clin_Clin | 859-07 | 2007 | Peru | Clinical | Latin America | O3:K6 | ST3 | PC | Wave-1 | Representative |
| 948-97_NA_Peru_Clin_Clin | 948-97 | 1997 | Peru | Clinical | Latin America | O3:K6 | ST3 | PC | Wave-1 | Representative |
| 971-98_NA_Peru_Clin_Clin | 971-98 | 1998 | Peru | Clinical | Latin America | O3:K6 | ST3 | PC | Wave-1 | Representative |
| C15_124-07__NA_Peru_Clin_Clin | C15_124-07 | 2007 | Peru | Clinical | Latin America | O3:K6 | ST3 | PC | Wave-1 | Representative |
| C20_NA_Peru_Clin_Clin | C20 | NA | Peru | Clinical | Latin America | O1:K25 | ST3 | PC | Wave-1 | Representative |
| C21_1262-07__NA_Peru_Clin_Clin | C21_1262-07 | 2007 | Peru | Clinical | Latin America | O3:K6 | ST3 | PC | Wave-1 | Representative |
| PM57_7_NA_Chile_Amb_Amb | PM57 | 2007 | Chile | Env | Latin America | O3:K6 | ST3 | PC | Wave-1 | Representative |
| PMC1_7_NA_Chile_NA_NA | PMC1 | 2007 | Chile | NA | Latin America | O3:K6 | ST3 | PC | Wave-1 | Representative |
| PMC16_7_NA_Chile_NA_NA | PMC16 | 2007 | Chile | NA | Latin America | O3:K6 | ST3 | PC | Wave-1 | Representative |
| PMC19_7_NA_Chile_Amb_Amb | PMC19 | 2007 | Chile | Env | Latin America | O3:K6 | ST3 | PC | Wave-1 | Representative |
| PMC29_7_NA_Chile_NA_NA | PMC29 | 2007 | Chile | NA | Latin America | O3:K6 | ST3 | PC | Wave-1 | Representative |
| SAMN01923801-GCF_000522065.2 | PVCHO_Peru-288 | 2001 | Peru | Clinical | Latin America | O3:K6 | ST3 | PC | Wave-1 | Representative |
| SAMN01924113-GCF_000558885.2 | PVCHO_V14/01 | 2001 | Chile | Clinical | Latin America | O3:K6 | ST3 | PC | Wave-1 | Representative |
| SAMN02781333-GCF_001270885.1 | ATC210 | 1998 | Chile | Clinical | Latin America | O3:K6 | ST3 | PC | Wave-1 | Representative |
| SAMN02781334-GCF_001270975.1 | ATC220 | 1998 | Chile | Clinical | Latin America | O3:K6 | ST3 | PC | Wave-1 | Representative |
| SAMN02781335-GCA_001270805.1 | PMA109.5 | 2005 | Chile | Env | Latin America | O3:K6 | ST- | PC | Wave-1 | Representative |
| SAMN02781336-GCF_001270895.1 | PMC14.7 | 2007 | Chile | Clinical | Latin America | O3:K6 | ST3 | PC | Wave-1 | Representative |
| SAMN02781337-GCF_001270905.1 | PMC48 | 2004 | Chile | Clinical | Latin America | O3:K6 | ST3 | PC | Wave-1 | Representative |
| SAMN02781338-GCF_001270815.1 | PMC58.5 | 2005 | Chile | Clinical | Latin America | O3:K6 | ST3 | PC | Wave-1 | Representative |
| SAMN02781339-GCF_001270825.1 | PMC58.7 | 2007 | Chile | Clinical | Latin America | O3:K6 | ST3 | PC | Wave-1 | Representative |
| SAMN03941065-GCF_001696015.1 | 906-97 | 1997 | Peru | Clinical | Latin America | O3:K6 | ST3 | PC | Wave-1 | Representative |
| SAMN04338417-SRR3002506 | NA | 2013 | Chile | Clinical | Latin America | O3:K6 | ST3 | PC | Wave-1 | Representative |
| SAMN05858273-GCF_002018625.1 | PMA37.5 | 2005 | Chile | Clinical | Latin America | O3:K6 | ST3 | PC | Wave-1 | Representative |
| SAMN06077005-GCA_015818715.1 | CFSAN023554 | 2005 | Chile | Env | Latin America | O3:K6 | ST3 | PC | Wave-1 | Representative |

| | | | | | | | | | | |
|---|---|---|---|---|---|---|---|---|---|---|
| SAMN06077007-GCA_015817935.1 | CFSAN023556 | 2005 | Chile | Env | Latin America | O3:K6 | ST3 | PC | Wave-1 | Representative |
| SAMN08225459-GCA_015814685.1 | CFSAN029657 | 2009 | Peru | Env | Latin America | O3:K6 | ST3 | PC | Wave-1 | Representative |
| SAMN12364836-GCF_009991845.1 | 1202-11 | 2011 | Peru | Clinical | Latin America | O3:K6 | ST3 | PC | Wave-1 | Representative |
| SAMN12364839-GCF_009991445.1 | G1 | 2014 | Peru | Clinical | Latin America | O3:K6 | ST3 | PC | Wave-1 | Representative |
| SAMN12364847-GCF_009991625.1 | 686-17 | 2017 | Peru | Clinical | Latin America | O1:K25 | ST3 | PC | Wave-1 | Representative |
| SAMN12364848-GCF_009991675.1 | 2214-17 | 2017 | Peru | Clinical | Latin America | O1:K25 | ST3 | PC | Wave-1 | Representative |
| SAMN15428940-GCF_016880045.1 | 325-00 | 2000 | Peru | Clinical | Latin America | O3:K6 | ST3 | PC | Wave-1 | Representative |
| SAMN15428942-GCF_016880085.1 | 327-00 | 2000 | Peru | Clinical | Latin America | O3:K6 | ST3 | PC | Wave-1 | Representative |
| SAMN15428945-GCF_016880015.1 | 2568-00 | 2000 | Peru | Clinical | Latin America | O3:K6 | ST3 | PC | Wave-1 | Representative |
| SAMN15428949-GCF_016879955.1 | 1260-07 | 2007 | Peru | Clinical | Latin America | O3:K6 | ST3 | PC | Wave-1 | Representative |
| SAMN15428972-GCF_016878275.1 | 223-10 | 2010 | Peru | Clinical | Latin America | O3:K6 | ST3 | PC | Wave-1 | Representative |
| SAMN15428983-GCF_016878055.1 | G6 | 2015 | Peru | Clinical | Latin America | O3:K6 | ST3 | PC | Wave-1 | Representative |
| SAMN15428984-GCF_016877995.1 | H11 | 2015 | Peru | Clinical | Latin America | O3:K6 | ST3 | PC | Wave-1 | Representative |
| SAMN15429004-GCF_016877605.1 | H12 | 2016 | Peru | Clinical | Latin America | O3:K6 | ST3 | PC | Wave-1 | Representative |
| Vp_Peru_498-01_Peru_2001_C | 498-01 | 2001 | Peru | Clinical | Latin America | O3:K6 | ST3 | PC | Wave-1 | Representative |
| Vp_Peru_706_Peru_2000_C | 706 | 2000 | Peru | Clinical | Latin America | O3:K6 | ST3 | PC | Wave-1 | Representative |
| Vp_Peru_CFSAN062237_2000_Peru_C | CFSAN062237 | 2000 | Peru | Clinical | Latin America | O3:K6 | ST3 | PC | Wave-1 | Representative |
| Vp_Peru_CFSAN062238_2000_Peru_C | CFSAN062238 | 2000 | Peru | Clinical | Latin America | O3:K6 | ST3 | PC | Wave-1 | Representative |
| Vp_Peru_CFSAN062240_2007_Peru_C | CFSAN062240 | 2007 | Peru | Clinical | Latin America | O3:K6 | ST3 | PC | Wave-1 | Representative |
| Vp_Peru_CFSAN062241_2000_Peru_C | CFSAN062241 | 2000 | Peru | Clinical | Latin America | O3:K6 | ST3 | PC | Wave-1 | Representative |
| Vp_Peru_CFSAN062244_2000_Peru_C | CFSAN062244 | 2000 | Peru | Clinical | Latin America | O3:K6 | ST3 | PC | Wave-1 | Representative |
| Vp_Peru_CFSAN062245_2000_Peru_C | CFSAN062245 | 2000 | Peru | Clinical | Latin America | O3:K6 | ST3 | PC | Wave-1 | Representative |
| Vp_Peru_CFSAN062251_2007_Peru_C | CFSAN062251 | 2007 | Peru | Clinical | Latin America | O1:K25 | ST3 | PC | Wave-1 | Representative |
| Vp_Peru_CFSAN062256_2000_Peru_C | CFSAN062256 | 2000 | Peru | Clinical | Latin America | O3:K6 | ST3 | PC | Wave-1 | Representative |
| Vp_Peru_CFSAN062267_2009_Peru_C | CFSAN062267 | 2009 | Peru | Clinical | Latin America | O3:K6 | ST3 | PC | Wave-1 | Representative |
| Vp_Peru_CFSAN062269_2000_Peru_C | CFSAN062269 | 2000 | Peru | Clinical | Latin America | O3:K6 | ST3 | PC | Wave-1 | Representative |
| Vp_Peru_CFSAN062270_2000_Peru_C | CFSAN062270 | 2000 | Peru | Clinical | Latin America | O3:K6 | ST3 | PC | Wave-1 | Representative |
| Vp_Peru_CFSAN062272_2010_Peru_C | CFSAN062272 | 2010 | Peru | Clinical | Latin America | O3:K6 | ST3 | PC | Wave-1 | Representative |
| Vp_Peru_CFSAN062297_2010_Peru_C | CFSAN062297 | 2010 | Peru | Clinical | Latin America | O3:K6 | ST3 | PC | Wave-1 | Representative |
| Vp_Peru_CFSAN062301_2016_Peru_C | CFSAN062301 | 2016 | Peru | Clinical | Latin America | O3:K6 | ST3 | PC | Wave-1 | Representative |
| Vp_Peru_CFSAN062318_2000_Peru_C | CFSAN062318 | 2000 | Peru | Clinical | Latin America | O3:K6 | ST3 | PC | Wave-1 | Representative |
| Vp_Peru_CFSAN062324_2010_Peru_C | CFSAN062324 | 2010 | Peru | Clinical | Latin America | O3:K6 | ST3 | PC | Wave-1 | Representative |

| | | | | | | | | | | |
|---|---|---|---|---|---|---|---|---|---|---|
| Vp_Peru_CFSAN062331_2010_Peru_C | CFSAN062331 | 2010 | Peru | Clinical | Latin America | O3:K6 | ST3 | PC | Wave-1 | Representative |
| Vp_Peru_CFSAN062333_2013_Peru_C | CFSAN062333 | 2013 | Peru | Clinical | Latin America | O3:K6 | ST3 | PC | Wave-1 | Representative |
| Vp_Peru_CFSAN062367_2010_Peru_C | CFSAN062367 | 2010 | Peru | Clinical | Latin America | O3:K6 | ST3 | PC | Wave-1 | Representative |
| Vp_Peru_GCA_000182345.1_ASM18234v1_NA_Peru_Na | GCA_000182345.1 | NA | Peru | NA | Latin America | O3:K6 | ST3 | PC | Wave-1 | Representative |
| Vp_Peru_H7_NA_NA_C | H7 | NA | Peru | Clinical | Latin America | O3:K6 | ST3 | PC | Wave-1 | Representative |
| Vp_Peru_P682_2009_Peru_E | P682 | 2009 | Peru | Env | Latin America | O3:K6 | ST3 | PC | Wave-1 | Representative |
| Vp_Peru_P729_2009_Peru_E | P729 | 2009 | Peru | Env | Latin America | O3:K6 | ST3 | PC | Wave-1 | Representative |
| Vp_Peru_P860_2009_Peru_E | P860 | 2009 | Peru | Env | Latin America | O3:K6 | ST3 | PC | Wave-1 | Representative |
| Vp_Peru_Peru-466_Peru_1996_NA | Peru-466 | 1996 | Peru | NA | Latin America | O3:K6 | ST3 | PC | Wave-1 | Representative |
| Vp_Peru_vp1218-11_2011_Peru_Lima_C | vp1218-11 | 2011 | Peru | Clinical | Latin America | O3:K6 | ST3 | PC | Wave-1 | Representative |
| Vp_Peru_vp1247-09_2009_Peru_Lima_C | vp1247-09 | 2009 | Peru | Clinical | Latin America | O3:K6 | ST3 | PC | Wave-1 | Representative |
| Vp_Peru_vp1437-07_2007_Peru_Lima_C | vp1437-07 | 2007 | Peru | Clinical | Latin America | O3:K6 | ST3 | PC | Wave-1 | Representative |
| Vp_Peru_vp21254-07_2007_NA_C | vp21254-07 | 2007 | Peru | Clinical | Latin America | O3:K6 | ST3 | PC | Wave-1 | Representative |
| Vp_Peru_vp2434-00_2000_NA_C | vp2434-00 | 2000 | Peru | Clinical | Latin America | O3:K6 | ST3 | PC | Wave-1 | Representative |
| Vp_Peru_vp437-07_2007_Peru_Lima_C | vp437-07 | 2007 | Peru | Clinical | Latin America | O3:K6 | ST3 | PC | Wave-1 | Representative |
| SAMN09280055-GCA_015805555.1 | 399423 | 2015 | NA | Clinical | NA | O3:K6 | ST3 | PC | Wave-1 | Representative |
| SAMN02368289-GCF_001728005.1 | BCW_3182 | 2007 | USA | Clinical | US_Canada | O3:K6 | ST3 | PC | Wave-1 | Representative |
| SAMN02368300-GCF_001728205.1 | BCW_3193 | 2007 | USA | Clinical | US_Canada | O3:K6 | ST3 | PC | Wave-1 | Representative |
| SAMN03358829-GCF_001727825.1 | BCW_3173 | 2006 | USA | Clinical | US_Canada | O3:K6 | ST3 | PC | Wave-1 | Representative |
| SAMN06076993-GCA_015819075.1 | BAC-98-3372 | 1998 | USA | Clinical | US_Canada | O3:K6 | ST3 | PC | Wave-1 | Representative |
| SAMN06076994-GCA_015819025.1 | BAC-98-3374 | 1998 | USA | Clinical | US_Canada | O3:K6 | ST3 | PC | Wave-1 | Representative |
| SAMN06076995-GCA_015818935.1 | BAC-98-4092 | 1998 | USA | Clinical | US_Canada | O3:K6 | ST3 | PC | Wave-1 | Representative |
| VP02004 | VP02004 | 2002 | China | Clinical | Asia | O3:K6 | ST3 | PC | Wave-2 | Putative mutator |
| VP03049 | VP03049 | 2003 | China | Clinical | Asia | O3:K6 | ST3 | PC | Wave-2 | Putative mutator |
| VP03050 | VP03050 | 2003 | China | Clinical | Asia | O3:K6 | ST3 | PC | Wave-2 | Putative mutator |
| VP05220 | VP05220 | 2005 | China | Clinical | Asia | O3:K6 | ST- | PC | Wave-2 | Putative mutator |
| SAMN13388561-SRR10531460 | 2 | 2019 | UK | Clinical | Europe | O3:K6 | ST3 | PC | Wave-2 | Putative mutator |
| SAMN05727869-GCF_001901565.1 | CICESE-170 | 1998 | Mexico | Clinical | Latin America | O3:K6 | ST3 | PC | Wave-2 | Putative mutator |
| SAMN13893121-GCF_010671575.1 | CICESE-188 | 2009 | Mexico | Clinical | Latin America | O3:K65 | ST3 | PC | Wave-2 | Putative mutator |
| SAMN07338227-GCF_006368375.1 | S061 | NA | NA | Clinical | NA | O3:K6 | ST- | PC | Wave-2 | Putative mutator |

| | | | | | | | | | | |
|---|---|---|---|---|---|---|---|---|---|---|
| SAMEA8103032-GCA_905331735.1 | ERS5790110 | 2016 | China | Clinical | Asia | O3:KUT7 | ST3 | PC | Wave-2 | Putative outbreak cluster |
| SAMN02471132-GCF_000500365.1 | AXNK | 2008 | China | Clinical | Asia | O3:K6 | ST3 | PC | Wave-2 | Putative outbreak cluster |
| SAMN09742583-GCF_006382895.1 | QP16233 | 2016 | China | Clinical | Asia | O1:KUT6 | ST3 | PC | Wave-2 | Putative outbreak cluster |
| SAMN16783001-GCA_016827495.1 | VP73 | 2007 | China | Clinical | Asia | O4:K68 | ST3 | PC | Wave-2 | Putative outbreak cluster |
| SAMN16783012-GCA_016827275.1 | VP84 | 2009 | China | Clinical | Asia | O4:K68 | ST3 | PC | Wave-2 | Putative outbreak cluster |
| SAMN16783015-GCA_016827245.1 | VP87 | 2009 | China | Clinical | Asia | O4:K68 | ST3 | PC | Wave-2 | Putative outbreak cluster |
| SAMN16783016-GCA_016827165.1 | VP88 | 2009 | China | Clinical | Asia | O4:K68 | ST3 | PC | Wave-2 | Putative outbreak cluster |
| SAMN16783017-GCA_016827195.1 | VP89 | 2009 | China | Clinical | Asia | O4:K68 | ST3 | PC | Wave-2 | Putative outbreak cluster |
| SAMN16783019-GCA_016827115.1 | VP91 | 2009 | China | Clinical | Asia | O4:K68 | ST3 | PC | Wave-2 | Putative outbreak cluster |
| SAMN16783020-GCA_016827125.1 | VP92 | 2009 | China | Clinical | Asia | O4:K68 | ST3 | PC | Wave-2 | Putative outbreak cluster |
| SAMN16783021-GCA_016827095.1 | VP93 | 2009 | China | Clinical | Asia | O4:K68 | ST3 | PC | Wave-2 | Putative outbreak cluster |
| SAMN16783022-GCA_016827075.1 | VP95 | 2009 | China | Clinical | Asia | O4:K68 | ST3 | PC | Wave-2 | Putative outbreak cluster |
| SAMN16783023-GCA_016827035.1 | VP96 | 2009 | China | Clinical | Asia | O4:K68 | ST3 | PC | Wave-2 | Putative outbreak cluster |
| SAMN16783026-GCA_016826995.1 | VP101 | 2010 | China | Clinical | Asia | O4:K68 | ST3 | PC | Wave-2 | Putative outbreak cluster |
| SAMN16783029-GCA_016826915.1 | VP105 | 2007 | China | Clinical | Asia | O4:K68 | ST3 | PC | Wave-2 | Putative outbreak cluster |
| SAMN16783030-GCA_016826895.1 | VP106 | 2007 | China | Clinical | Asia | O4:K68 | ST3 | PC | Wave-2 | Putative outbreak cluster |
| SAMN16783036-GCA_016826815.1 | VP112 | 2011 | China | NA | Asia | O4:K68 | ST3 | PC | Wave-2 | Putative outbreak cluster |
| SAMN16783053-GCA_016826475.1 | VP130 | 2008 | China | Clinical | Asia | O4:K68 | ST3 | PC | Wave-2 | Putative outbreak cluster |
| SAMN16783054-GCA_016826435.1 | VP131 | 2008 | China | Clinical | Asia | O4:K68 | ST3 | PC | Wave-2 | Putative outbreak cluster |
| SAMN16783057-GCA_016826375.1 | VP134 | 2008 | China | Clinical | Asia | O4:K68 | ST3 | PC | Wave-2 | Putative outbreak cluster |
| SAMN16783062-GCA_016826255.1 | VP139 | 2008 | China | Clinical | Asia | O4:K68 | ST3 | PC | Wave-2 | Putative outbreak cluster |
| SAMN16783066-GCA_016826195.1 | VP143 | 2009 | China | Clinical | Asia | O4:K68 | ST3 | PC | Wave-2 | Putative outbreak cluster |
| SAMN16783079-GCA_016825915.1 | VP158 | 2010 | China | Clinical | Asia | O4:K68 | ST3 | PC | Wave-2 | Putative outbreak cluster |
| SAMN16783080-GCA_016825855.1 | VP159 | 2010 | China | Clinical | Asia | O4:K68 | ST3 | PC | Wave-2 | Putative outbreak cluster |
| SAMN16783081-GCA_016825905.1 | VP160 | 2010 | China | Clinical | Asia | O4:K68 | ST3 | PC | Wave-2 | Putative outbreak cluster |
| SAMN16783082-GCA_016825875.1 | VP161 | 2010 | China | Clinical | Asia | O4:K68 | ST3 | PC | Wave-2 | Putative outbreak cluster |
| SAMN16783084-GCA_016825835.1 | VP163 | 2010 | China | Clinical | Asia | O4:K68 | ST3 | PC | Wave-2 | Putative outbreak cluster |
| SAMN16783085-GCA_016825795.1 | VP164 | 2010 | China | Clinical | Asia | O4:K68 | ST3 | PC | Wave-2 | Putative outbreak cluster |
| SAMN16783086-GCA_016825785.1 | VP165 | 2010 | China | Clinical | Asia | O4:K68 | ST3 | PC | Wave-2 | Putative outbreak cluster |
| SAMN16783087-GCA_016825775.1 | VP166 | 2010 | China | Clinical | Asia | O4:K68 | ST3 | PC | Wave-2 | Putative outbreak cluster |
| SAMN16783088-GCA_016825745.1 | VP167 | 2010 | China | Clinical | Asia | O4:K68 | ST3 | PC | Wave-2 | Putative outbreak cluster |
| SAMN16783090-GCA_016825685.1 | VP170 | 2012 | China | NA | Asia | O4:K68 | ST3 | PC | Wave-2 | Putative outbreak cluster |

| | | | | | | | | | | |
|---|---|---|---|---|---|---|---|---|---|---|
| SAMN16783096-GCA_016825595.1 | VP176 | 2011 | China | NA | Asia | O4:K68 | ST3 | PC | Wave-2 | Putative outbreak cluster |
| SAMN16783097-GCA_016825575.1 | VP178 | 2011 | China | NA | Asia | O4:K68 | ST3 | PC | Wave-2 | Putative outbreak cluster |
| SAMN16783103-GCA_016825435.1 | VP184 | 2007 | China | Clinical | Asia | O4:K68 | ST3 | PC | Wave-2 | Putative outbreak cluster |
| SAMN16783111-GCA_016825285.1 | VP194 | 2015 | China | NA | Asia | O4:K68 | ST3 | PC | Wave-2 | Putative outbreak cluster |
| SAMN16783136-GCA_016824755.1 | VP227 | 2010 | China | Clinical | Asia | O4:K68 | ST3 | PC | Wave-2 | Putative outbreak cluster |
| SAMN16783142-GCA_016824675.1 | VP237 | 2009 | China | Clinical | Asia | O4:K68 | ST3 | PC | Wave-2 | Putative outbreak cluster |
| SAMN16783203-GCA_016823455.1 | VP302 | 2015 | China | NA | Asia | O3:K6 | ST3 | PC | Wave-2 | Putative outbreak cluster |
| SAMN16783251-GCA_016822495.1 | VP355 | 2013 | China | NA | Asia | O1:KUK3 | ST3 | PC | Wave-2 | Putative outbreak cluster |
| SAMN16783253-GCA_016822455.1 | VP357 | 2008 | China | NA | Asia | O3:K6 | ST3 | PC | Wave-2 | Putative outbreak cluster |
| SAMN16783265-GCA_016822215.1 | VP369 | 2009 | China | NA | Asia | O3:K6 | ST3 | PC | Wave-2 | Putative outbreak cluster |
| SAMN16783269-GCA_016822145.1 | VP373 | 2010 | China | Clinical | Asia | O3:K6 | ST3 | PC | Wave-2 | Putative outbreak cluster |
| SAMN16783285-GCA_016821755.1 | VP390 | 2011 | China | NA | Asia | O3:K6 | ST3 | PC | Wave-2 | Putative outbreak cluster |
| SAMN16783289-GCA_016821735.1 | VP394 | 2008 | China | Clinical | Asia | O3:K6 | ST3 | PC | Wave-2 | Putative outbreak cluster |
| SAMN16783290-GCA_016821715.1 | VP395 | 2009 | China | Clinical | Asia | O3:K6 | ST3 | PC | Wave-2 | Putative outbreak cluster |
| SAMN16783291-GCA_016821655.1 | VP396 | 2016 | China | NA | Asia | O3:K6 | ST3 | PC | Wave-2 | Putative outbreak cluster |
| SAMN16783300-GCA_016821515.1 | VP405 | 2017 | China | NA | Asia | O3:K6 | ST3 | PC | Wave-2 | Putative outbreak cluster |
| SAMN16783379-GCA_016818535.1 | VP235 | 2007 | China | Clinical | Asia | O4:K68 | ST3 | PC | Wave-2 | Putative outbreak cluster |
| VP06014 | VP06014 | 2006 | China | Clinical | Asia | O3:K6 | ST3 | PC | Wave-2 | Putative outbreak cluster |
| VP06015 | VP06015 | 2006 | China | Clinical | Asia | O3:K6 | ST3 | PC | Wave-2 | Putative outbreak cluster |
| VP06016 | VP06016 | 2006 | China | Clinical | Asia | O3:K6 | ST3 | PC | Wave-2 | Putative outbreak cluster |
| VP06027 | VP06027 | 2006 | China | Clinical | Asia | O3:K6 | ST3 | PC | Wave-2 | Putative outbreak cluster |
| VP06028 | VP06028 | 2006 | China | Clinical | Asia | O3:K6 | ST3 | PC | Wave-2 | Putative outbreak cluster |
| VP06029 | VP06029 | 2006 | China | Clinical | Asia | O3:K6 | ST3 | PC | Wave-2 | Putative outbreak cluster |
| VP06079 | VP06079 | 2006 | China | Clinical | Asia | O3:K6 | ST3 | PC | Wave-2 | Putative outbreak cluster |
| VP06080 | VP06080 | 2006 | China | Clinical | Asia | O3:K6 | ST3 | PC | Wave-2 | Putative outbreak cluster |
| VP06081 | VP06081 | 2006 | China | Clinical | Asia | O3:K6 | ST3 | PC | Wave-2 | Putative outbreak cluster |
| VP06082 | VP06082 | 2006 | China | Clinical | Asia | O3:K6 | ST3 | PC | Wave-2 | Putative outbreak cluster |
| VP06087 | VP06087 | 2006 | China | Clinical | Asia | O3:K6 | ST3 | PC | Wave-2 | Putative outbreak cluster |
| VP06089 | VP06089 | 2006 | China | Clinical | Asia | O3:K6 | ST3 | PC | Wave-2 | Putative outbreak cluster |
| VP06090 | VP06090 | 2006 | China | Clinical | Asia | O3:K6 | ST3 | PC | Wave-2 | Putative outbreak cluster |
| VP06096 | VP06096 | 2006 | China | Clinical | Asia | O3:K6 | ST3 | PC | Wave-2 | Putative outbreak cluster |
| VP06109 | VP06109 | 2006 | China | Clinical | Asia | O3:K6 | ST3 | PC | Wave-2 | Putative outbreak cluster |

| | | | | | | | | | | |
|---|---|---|---|---|---|---|---|---|---|---|
| VP06111 | VP06111 | 2006 | China | Clinical | Asia | O3:K6 | ST3 | PC | Wave-2 | Putative outbreak cluster |
| VP06112 | VP06112 | 2006 | China | Clinical | Asia | O3:K6 | ST3 | PC | Wave-2 | Putative outbreak cluster |
| VP06114 | VP06114 | 2006 | China | Clinical | Asia | O3:K6 | ST3 | PC | Wave-2 | Putative outbreak cluster |
| VP06115 | VP06115 | 2006 | China | Clinical | Asia | O3:K6 | ST3 | PC | Wave-2 | Putative outbreak cluster |
| VP06116 | VP06116 | 2006 | China | Clinical | Asia | O3:K6 | ST3 | PC | Wave-2 | Putative outbreak cluster |
| VP06117 | VP06117 | 2006 | China | Clinical | Asia | O3:K6 | ST3 | PC | Wave-2 | Putative outbreak cluster |
| VP06118 | VP06118 | 2006 | China | Clinical | Asia | O3:K6 | ST3 | PC | Wave-2 | Putative outbreak cluster |
| VP06119 | VP06119 | 2006 | China | Clinical | Asia | O3:K6 | ST3 | PC | Wave-2 | Putative outbreak cluster |
| VP06121 | VP06121 | 2006 | China | Clinical | Asia | O3:K6 | ST3 | PC | Wave-2 | Putative outbreak cluster |
| VP06122 | VP06122 | 2006 | China | Clinical | Asia | O3:K6 | ST3 | PC | Wave-2 | Putative outbreak cluster |
| VP06123 | VP06123 | 2006 | China | Clinical | Asia | O3:K6 | ST3 | PC | Wave-2 | Putative outbreak cluster |
| VP06124 | VP06124 | 2006 | China | Clinical | Asia | O3:K6 | ST3 | PC | Wave-2 | Putative outbreak cluster |
| VP06125 | VP06125 | 2006 | China | Clinical | Asia | O3:K6 | ST3 | PC | Wave-2 | Putative outbreak cluster |
| VP06127 | VP06127 | 2006 | China | Clinical | Asia | O3:K6 | ST3 | PC | Wave-2 | Putative outbreak cluster |
| VP06128 | VP06128 | 2006 | China | Clinical | Asia | O3:K6 | ST3 | PC | Wave-2 | Putative outbreak cluster |
| VP06129 | VP06129 | 2006 | China | Clinical | Asia | O3:K6 | ST3 | PC | Wave-2 | Putative outbreak cluster |
| VP06130 | VP06130 | 2006 | China | Clinical | Asia | O3:K6 | ST3 | PC | Wave-2 | Putative outbreak cluster |
| VP06131 | VP06131 | 2006 | China | Clinical | Asia | O3:K6 | ST3 | PC | Wave-2 | Putative outbreak cluster |
| VP06172 | VP06172 | 2006 | China | Clinical | Asia | O3:K6 | ST3 | PC | Wave-2 | Putative outbreak cluster |
| VP06173 | VP06173 | 2006 | China | Clinical | Asia | O3:K6 | ST3 | PC | Wave-2 | Putative outbreak cluster |
| VP06174 | VP06174 | 2006 | China | Clinical | Asia | O3:K6 | ST3 | PC | Wave-2 | Putative outbreak cluster |
| VP06175 | VP06175 | 2006 | China | Clinical | Asia | O3:K6 | ST3 | PC | Wave-2 | Putative outbreak cluster |
| VP06177 | VP06177 | 2006 | China | Clinical | Asia | O3:K6 | ST3 | PC | Wave-2 | Putative outbreak cluster |
| VP06178 | VP06178 | 2006 | China | Clinical | Asia | O3:K6 | ST3 | PC | Wave-2 | Putative outbreak cluster |
| VP06179 | VP06179 | 2006 | China | Clinical | Asia | O3:K6 | ST3 | PC | Wave-2 | Putative outbreak cluster |
| VP07013 | VP07013 | 2007 | China | Clinical | Asia | O3:K6 | ST3 | PC | Wave-2 | Putative outbreak cluster |
| VP07014 | VP07014 | 2007 | China | Clinical | Asia | O3:K6 | ST3 | PC | Wave-2 | Putative outbreak cluster |
| VP07018 | VP07018 | 2007 | China | Clinical | Asia | O3:K6 | ST3 | PC | Wave-2 | Putative outbreak cluster |
| VP07020 | VP07020 | 2007 | China | Clinical | Asia | O3:K6 | ST3 | PC | Wave-2 | Putative outbreak cluster |
| VP07021 | VP07021 | 2007 | China | Clinical | Asia | O3:K6 | ST3 | PC | Wave-2 | Putative outbreak cluster |
| VP07029 | VP07029 | 2007 | China | Clinical | Asia | O3:K6 | ST3 | PC | Wave-2 | Putative outbreak cluster |
| VP07066 | VP07066 | 2007 | China | Clinical | Asia | O3:K6 | ST3 | PC | Wave-2 | Putative outbreak cluster |

| | | | | | | | | | | |
|---|---|---|---|---|---|---|---|---|---|---|
| VP07081 | VP07081 | 2007 | China | Clinical | Asia | O4:K68 | ST3 | PC | Wave-2 | Putative outbreak cluster |
| VP07094 | VP07094 | 2007 | China | Clinical | Asia | O3:K6 | ST3 | PC | Wave-2 | Putative outbreak cluster |
| VP07122 | VP07122 | 2007 | China | Clinical | Asia | O3:K6 | ST3 | PC | Wave-2 | Putative outbreak cluster |
| VP07179 | VP07179 | 2007 | China | Clinical | Asia | O4:K68 | ST3 | PC | Wave-2 | Putative outbreak cluster |
| VP07182 | VP07182 | 2007 | China | Clinical | Asia | O4:K68 | ST3 | PC | Wave-2 | Putative outbreak cluster |
| VP07184 | VP07184 | 2007 | China | Clinical | Asia | O4:K68 | ST3 | PC | Wave-2 | Putative outbreak cluster |
| VP07185 | VP07185 | 2007 | China | Env | Asia | O4:K68 | ST3 | PC | Wave-2 | Putative outbreak cluster |
| VP07190 | VP07190 | 2007 | China | Clinical | Asia | O4:K68 | ST3 | PC | Wave-2 | Putative outbreak cluster |
| VP07198 | VP07198 | 2007 | China | Clinical | Asia | O3:K6 | ST3 | PC | Wave-2 | Putative outbreak cluster |
| VP07202 | VP07202 | 2007 | China | Clinical | Asia | O4:K68 | ST3 | PC | Wave-2 | Putative outbreak cluster |
| VP08007 | VP08007 | 2008 | China | Clinical | Asia | O3:K6 | ST3 | PC | Wave-2 | Putative outbreak cluster |
| VP08008 | VP08008 | 2008 | China | Clinical | Asia | O3:K6 | ST3 | PC | Wave-2 | Putative outbreak cluster |
| VP08011 | VP08011 | 2008 | China | Clinical | Asia | O3:K6 | ST3 | PC | Wave-2 | Putative outbreak cluster |
| VP08012 | VP08012 | 2008 | China | Clinical | Asia | O3:K6 | ST3 | PC | Wave-2 | Putative outbreak cluster |
| VP08021 | VP08021 | 2008 | China | Clinical | Asia | O3:K6 | ST3 | PC | Wave-2 | Putative outbreak cluster |
| VP08022 | VP08022 | 2008 | China | Clinical | Asia | O3:K6 | ST3 | PC | Wave-2 | Putative outbreak cluster |
| VP08023 | VP08023 | 2008 | China | Clinical | Asia | O3:K6 | ST3 | PC | Wave-2 | Putative outbreak cluster |
| VP08043 | VP08043 | 2008 | China | Clinical | Asia | O3:K6 | ST3 | PC | Wave-2 | Putative outbreak cluster |
| VP08045 | VP08045 | 2008 | China | Clinical | Asia | O4:K68 | ST3 | PC | Wave-2 | Putative outbreak cluster |
| VP08048 | VP08048 | 2008 | China | Clinical | Asia | O3:K6 | ST3 | PC | Wave-2 | Putative outbreak cluster |
| VP08050 | VP08050 | 2008 | China | Clinical | Asia | O3:K6 | ST3 | PC | Wave-2 | Putative outbreak cluster |
| VP08051 | VP08051 | 2008 | China | Clinical | Asia | O3:K6 | ST3 | PC | Wave-2 | Putative outbreak cluster |
| VP08052 | VP08052 | 2008 | China | Clinical | Asia | O3:K6 | ST3 | PC | Wave-2 | Putative outbreak cluster |
| VP08054 | VP08054 | 2008 | China | Clinical | Asia | O3:K6 | ST3 | PC | Wave-2 | Putative outbreak cluster |
| VP08055 | VP08055 | 2008 | China | Clinical | Asia | O3:K6 | ST3 | PC | Wave-2 | Putative outbreak cluster |
| VP08060 | VP08060 | 2008 | China | Clinical | Asia | O3:K6 | ST3 | PC | Wave-2 | Putative outbreak cluster |
| VP08061 | VP08061 | 2008 | China | Clinical | Asia | O3:K6 | ST3 | PC | Wave-2 | Putative outbreak cluster |
| VP08062 | VP08062 | 2008 | China | Clinical | Asia | O3:K6 | ST3 | PC | Wave-2 | Putative outbreak cluster |
| VP08063 | VP08063 | 2008 | China | Clinical | Asia | O3:K6 | ST3 | PC | Wave-2 | Putative outbreak cluster |
| VP08064 | VP08064 | 2008 | China | Clinical | Asia | O3:K6 | ST3 | PC | Wave-2 | Putative outbreak cluster |
| VP08065 | VP08065 | 2008 | China | Clinical | Asia | O3:K6 | ST3 | PC | Wave-2 | Putative outbreak cluster |
| VP08066 | VP08066 | 2008 | China | Clinical | Asia | O3:K6 | ST3 | PC | Wave-2 | Putative outbreak cluster |

| | | | | | | | | | | |
|---|---|---|---|---|---|---|---|---|---|---|
| VP08080 | VP08080 | 2008 | China | Clinical | Asia | O3:K6 | ST3 | PC | Wave-2 | Putative outbreak cluster |
| VP08082 | VP08082 | 2008 | China | Clinical | Asia | O4:K68 | ST3 | PC | Wave-2 | Putative outbreak cluster |
| VP08120 | VP08120 | 2008 | China | Clinical | Asia | O3:K6 | ST3 | PC | Wave-2 | Putative outbreak cluster |
| VP08164 | VP08164 | 2008 | China | Clinical | Asia | O3:K6 | ST3 | PC | Wave-2 | Putative outbreak cluster |
| VP08167 | VP08167 | 2008 | China | Clinical | Asia | O3:K6 | ST3 | PC | Wave-2 | Putative outbreak cluster |
| VP08188 | VP08188 | 2008 | China | Clinical | Asia | O3:K6 | ST3 | PC | Wave-2 | Putative outbreak cluster |
| VP08197 | VP08197 | 2008 | China | Clinical | Asia | O3:K6 | ST3 | PC | Wave-2 | Putative outbreak cluster |
| VP08198 | VP08198 | 2008 | China | Clinical | Asia | O3:K6 | ST3 | PC | Wave-2 | Putative outbreak cluster |
| VP08209 | VP08209 | 2008 | China | Clinical | Asia | O3:K6 | ST3 | PC | Wave-2 | Putative outbreak cluster |
| VP08217 | VP08217 | 2008 | China | Clinical | Asia | O3:K6 | ST3 | PC | Wave-2 | Putative outbreak cluster |
| VP08231 | VP08231 | 2008 | China | Clinical | Asia | O3:K6 | ST3 | PC | Wave-2 | Putative outbreak cluster |
| VP08233 | VP08233 | 2008 | China | Clinical | Asia | O3:K6 | ST3 | PC | Wave-2 | Putative outbreak cluster |
| VP08235 | VP08235 | 2008 | China | Clinical | Asia | O3:K6 | ST3 | PC | Wave-2 | Putative outbreak cluster |
| VP08240 | VP08240 | 2008 | China | Clinical | Asia | O3:K6 | ST3 | PC | Wave-2 | Putative outbreak cluster |
| VP08241 | VP08241 | 2008 | China | Clinical | Asia | O3:K6 | ST3 | PC | Wave-2 | Putative outbreak cluster |
| VP08247 | VP08247 | 2008 | China | Clinical | Asia | O3:K6 | ST3 | PC | Wave-2 | Putative outbreak cluster |
| VP08250 | VP08250 | 2008 | China | Clinical | Asia | O3:K6 | ST3 | PC | Wave-2 | Putative outbreak cluster |
| VP08251 | VP08251 | 2008 | China | Clinical | Asia | O3:K6 | ST3 | PC | Wave-2 | Putative outbreak cluster |
| VP08252 | VP08252 | 2008 | China | Clinical | Asia | O3:K6 | ST3 | PC | Wave-2 | Putative outbreak cluster |
| VP08253 | VP08253 | 2008 | China | Clinical | Asia | O3:K6 | ST3 | PC | Wave-2 | Putative outbreak cluster |
| VP08254 | VP08254 | 2008 | China | Clinical | Asia | O3:K6 | ST3 | PC | Wave-2 | Putative outbreak cluster |
| VP08265 | VP08265 | 2008 | China | Clinical | Asia | O3:K6 | ST3 | PC | Wave-2 | Putative outbreak cluster |
| VP08314 | VP08314 | 2008 | China | Clinical | Asia | O3:K6 | ST3 | PC | Wave-2 | Putative outbreak cluster |
| VP08326 | VP08326 | 2008 | China | Clinical | Asia | O3:K6 | ST3 | PC | Wave-2 | Putative outbreak cluster |
| VP08331 | VP08331 | 2008 | China | Clinical | Asia | O3:K6 | ST3 | PC | Wave-2 | Putative outbreak cluster |
| VP08332 | VP08332 | 2008 | China | Clinical | Asia | O4:K68 | ST3 | PC | Wave-2 | Putative outbreak cluster |
| VP08334 | VP08334 | 2008 | China | Clinical | Asia | O3:K6 | ST3 | PC | Wave-2 | Putative outbreak cluster |
| VP08335 | VP08335 | 2008 | China | Clinical | Asia | O3:K6 | ST3 | PC | Wave-2 | Putative outbreak cluster |
| VP08336 | VP08336 | 2008 | China | Clinical | Asia | O3:K6 | ST3 | PC | Wave-2 | Putative outbreak cluster |
| VP08340 | VP08340 | 2008 | China | Clinical | Asia | O3:K6 | ST3 | PC | Wave-2 | Putative outbreak cluster |
| VP08342 | VP08342 | 2008 | China | Clinical | Asia | O3:K6 | ST3 | PC | Wave-2 | Putative outbreak cluster |
| VP08344 | VP08344 | 2008 | China | Clinical | Asia | O3:K6 | ST3 | PC | Wave-2 | Putative outbreak cluster |

| | | | | | | | | | | |
|---|---|---|---|---|---|---|---|---|---|---|
| VP08346 | VP08346 | 2008 | China | Clinical | Asia | O3:K6 | ST3 | PC | Wave-2 | Putative outbreak cluster |
| VP08362 | VP08362 | 2008 | China | Clinical | Asia | O3:K6 | ST3 | PC | Wave-2 | Putative outbreak cluster |
| VP08364 | VP08364 | 2008 | China | Clinical | Asia | O3:K6 | ST3 | PC | Wave-2 | Putative outbreak cluster |
| VP08365 | VP08365 | 2008 | China | Clinical | Asia | O3:K6 | ST3 | PC | Wave-2 | Putative outbreak cluster |
| VP08369 | VP08369 | 2008 | China | Clinical | Asia | O3:K6 | ST3 | PC | Wave-2 | Putative outbreak cluster |
| VP08371 | VP08371 | 2008 | China | Clinical | Asia | O3:K6 | ST3 | PC | Wave-2 | Putative outbreak cluster |
| VP08372 | VP08372 | 2008 | China | Clinical | Asia | O3:K6 | ST3 | PC | Wave-2 | Putative outbreak cluster |
| VP08373 | VP08373 | 2008 | China | Clinical | Asia | O3:K6 | ST3 | PC | Wave-2 | Putative outbreak cluster |
| VP08376 | VP08376 | 2008 | China | Clinical | Asia | O3:K6 | ST3 | PC | Wave-2 | Putative outbreak cluster |
| VP08380 | VP08380 | 2008 | China | Clinical | Asia | O3:K6 | ST3 | PC | Wave-2 | Putative outbreak cluster |
| VP08381 | VP08381 | 2008 | China | Clinical | Asia | O3:K6 | ST3 | PC | Wave-2 | Putative outbreak cluster |
| VP08384 | VP08384 | 2008 | China | Clinical | Asia | O3:K6 | ST3 | PC | Wave-2 | Putative outbreak cluster |
| VP08385 | VP08385 | 2008 | China | Clinical | Asia | O3:K6 | ST3 | PC | Wave-2 | Putative outbreak cluster |
| VP08386 | VP08386 | 2008 | China | Clinical | Asia | O3:K6 | ST3 | PC | Wave-2 | Putative outbreak cluster |
| VP08387 | VP08387 | 2008 | China | Clinical | Asia | O3:K6 | ST3 | PC | Wave-2 | Putative outbreak cluster |
| VP08395 | VP08395 | 2008 | China | Clinical | Asia | O3:K6 | ST3 | PC | Wave-2 | Putative outbreak cluster |
| VP08396 | VP08396 | 2008 | China | Clinical | Asia | O3:K6 | ST3 | PC | Wave-2 | Putative outbreak cluster |
| VP08397 | VP08397 | 2008 | China | Clinical | Asia | O3:K6 | ST3 | PC | Wave-2 | Putative outbreak cluster |
| VP09001 | VP09001 | 2009 | China | Clinical | Asia | O3:K6 | ST3 | PC | Wave-2 | Putative outbreak cluster |
| VP09025 | VP09025 | 2009 | China | Clinical | Asia | O3:K6 | ST3 | PC | Wave-2 | Putative outbreak cluster |
| VP09027 | VP09027 | 2009 | China | Clinical | Asia | O3:K6 | ST3 | PC | Wave-2 | Putative outbreak cluster |
| VP09028 | VP09028 | 2009 | China | Clinical | Asia | O3:K6 | ST3 | PC | Wave-2 | Putative outbreak cluster |
| VP09032 | VP09032 | 2009 | China | Clinical | Asia | O3:K6 | ST3 | PC | Wave-2 | Putative outbreak cluster |
| VP09041 | VP09041 | 2009 | China | Clinical | Asia | O3:K6 | ST3 | PC | Wave-2 | Putative outbreak cluster |
| VP09071 | VP09071 | 2009 | China | Clinical | Asia | O3:K6 | ST3 | PC | Wave-2 | Putative outbreak cluster |
| VP09077 | VP09077 | 2009 | China | Clinical | Asia | O3:K6 | ST3 | PC | Wave-2 | Putative outbreak cluster |
| VP09092 | VP09092 | 2009 | China | Clinical | Asia | O3:K6 | ST3 | PC | Wave-2 | Putative outbreak cluster |
| VP09100 | VP09100 | 2009 | China | Clinical | Asia | O3:K6 | ST3 | PC | Wave-2 | Putative outbreak cluster |
| VP09103 | VP09103 | 2009 | China | Clinical | Asia | O3:K6 | ST3 | PC | Wave-2 | Putative outbreak cluster |
| VP09106 | VP09106 | 2009 | China | Clinical | Asia | O3:K6 | ST3 | PC | Wave-2 | Putative outbreak cluster |
| VP09108 | VP09108 | 2009 | China | Clinical | Asia | O3:K6 | ST3 | PC | Wave-2 | Putative outbreak cluster |
| VP09123 | VP09123 | 2009 | China | Clinical | Asia | O3:K6 | ST3 | PC | Wave-2 | Putative outbreak cluster |

| | | | | | | | | | | |
|---|---|---|---|---|---|---|---|---|---|---|
| VP09126 | VP09126 | 2009 | China | Clinical | Asia | O3:K6 | ST3 | PC | Wave-2 | Putative outbreak cluster |
| VP09129 | VP09129 | 2009 | China | Clinical | Asia | O3:K6 | ST3 | PC | Wave-2 | Putative outbreak cluster |
| VP09132 | VP09132 | 2009 | China | Clinical | Asia | O3:K6 | ST3 | PC | Wave-2 | Putative outbreak cluster |
| VP09133 | VP09133 | 2009 | China | Clinical | Asia | O3:K6 | ST3 | PC | Wave-2 | Putative outbreak cluster |
| VP09138 | VP09138 | 2009 | China | Clinical | Asia | O3:K6 | ST3 | PC | Wave-2 | Putative outbreak cluster |
| VP09140 | VP09140 | 2009 | China | Clinical | Asia | O3:K6 | ST3 | PC | Wave-2 | Putative outbreak cluster |
| VP09148 | VP09148 | 2009 | China | Clinical | Asia | O3:K6 | ST3 | PC | Wave-2 | Putative outbreak cluster |
| VP09150 | VP09150 | 2009 | China | Clinical | Asia | O3:K6 | ST3 | PC | Wave-2 | Putative outbreak cluster |
| VP09162 | VP09162 | 2009 | China | Clinical | Asia | O3:K6 | ST3 | PC | Wave-2 | Putative outbreak cluster |
| VP09164 | VP09164 | 2009 | China | Clinical | Asia | O3:K6 | ST3 | PC | Wave-2 | Putative outbreak cluster |
| VP09165 | VP09165 | 2009 | China | Clinical | Asia | O3:K6 | ST3 | PC | Wave-2 | Putative outbreak cluster |
| VP09176 | VP09176 | 2009 | China | Clinical | Asia | O3:K6 | ST3 | PC | Wave-2 | Putative outbreak cluster |
| VP09177 | VP09177 | 2009 | China | Clinical | Asia | O3:K6 | ST3 | PC | Wave-2 | Putative outbreak cluster |
| VP09180 | VP09180 | 2009 | China | Clinical | Asia | O3:K6 | ST3 | PC | Wave-2 | Putative outbreak cluster |
| VP09181 | VP09181 | 2009 | China | Clinical | Asia | O3:K6 | ST3 | PC | Wave-2 | Putative outbreak cluster |
| VP09182 | VP09182 | 2009 | China | Clinical | Asia | O3:K6 | ST3 | PC | Wave-2 | Putative outbreak cluster |
| VP09186 | VP09186 | 2009 | China | Clinical | Asia | O3:K6 | ST3 | PC | Wave-2 | Putative outbreak cluster |
| VP09187 | VP09187 | 2009 | China | Clinical | Asia | O3:K6 | ST3 | PC | Wave-2 | Putative outbreak cluster |
| VP09196 | VP09196 | 2009 | China | Clinical | Asia | O3:K6 | ST3 | PC | Wave-2 | Putative outbreak cluster |
| VP09198 | VP09198 | 2009 | China | Clinical | Asia | O3:K6 | ST3 | PC | Wave-2 | Putative outbreak cluster |
| VP09199-01 | VP09199-01 | 2009 | China | Clinical | Asia | O3:K6 | ST3 | PC | Wave-2 | Putative outbreak cluster |
| VP09219 | VP09219 | 2009 | China | Clinical | Asia | O3:K6 | ST3 | PC | Wave-2 | Putative outbreak cluster |
| VP09241 | VP09241 | 2009 | China | Clinical | Asia | O4:K68 | ST3 | PC | Wave-2 | Putative outbreak cluster |
| VP09244 | VP09244 | 2009 | China | Clinical | Asia | O4:K68 | ST3 | PC | Wave-2 | Putative outbreak cluster |
| VP09246 | VP09246 | 2009 | China | Clinical | Asia | O3:K6 | ST3 | PC | Wave-2 | Putative outbreak cluster |
| VP09250 | VP09250 | 2009 | China | Clinical | Asia | O3:K6 | ST3 | PC | Wave-2 | Putative outbreak cluster |
| VP09254 | VP09254 | 2009 | China | Clinical | Asia | O4:K68 | ST3 | PC | Wave-2 | Putative outbreak cluster |
| VP09271 | VP09271 | 2009 | China | Clinical | Asia | O3:K6 | ST3 | PC | Wave-2 | Putative outbreak cluster |
| VP09333 | VP09333 | 2009 | China | Clinical | Asia | O4:K68 | ST3 | PC | Wave-2 | Putative outbreak cluster |
| VP09334 | VP09334 | 2009 | China | Clinical | Asia | O4:K68 | ST3 | PC | Wave-2 | Putative outbreak cluster |
| VP09345 | VP09345 | 2009 | China | Clinical | Asia | O3:K6 | ST3 | PC | Wave-2 | Putative outbreak cluster |
| VP09349 | VP09349 | 2009 | China | Clinical | Asia | O3:K6 | ST3 | PC | Wave-2 | Putative outbreak cluster |

| | | | | | | | | | | |
|---|---|---|---|---|---|---|---|---|---|---|
| VP09350 | VP09350 | 2009 | China | Clinical | Asia | O3:K6 | ST3 | PC | Wave-2 | Putative outbreak cluster |
| VP09351 | VP09351 | 2009 | China | Clinical | Asia | O4:K68 | ST3 | PC | Wave-2 | Putative outbreak cluster |
| VP09352 | VP09352 | 2009 | China | Clinical | Asia | O3:K6 | ST3 | PC | Wave-2 | Putative outbreak cluster |
| VP09354 | VP09354 | 2009 | China | Clinical | Asia | O3:K6 | ST3 | PC | Wave-2 | Putative outbreak cluster |
| VP09360 | VP09360 | 2009 | China | Clinical | Asia | O4:K68 | ST3 | PC | Wave-2 | Putative outbreak cluster |
| VP09369 | VP09369 | 2009 | China | Clinical | Asia | O3:K6 | ST3 | PC | Wave-2 | Putative outbreak cluster |
| VP09370 | VP09370 | 2009 | China | Clinical | Asia | O3:K6 | ST3 | PC | Wave-2 | Putative outbreak cluster |
| VP09371 | VP09371 | 2009 | China | Clinical | Asia | O3:K6 | ST3 | PC | Wave-2 | Putative outbreak cluster |
| VP09373 | VP09373 | 2009 | China | Clinical | Asia | O3:K6 | ST3 | PC | Wave-2 | Putative outbreak cluster |
| VP09374 | VP09374 | 2009 | China | Clinical | Asia | O4:K68 | ST3 | PC | Wave-2 | Putative outbreak cluster |
| VP09375 | VP09375 | 2009 | China | Clinical | Asia | O4:K68 | ST3 | PC | Wave-2 | Putative outbreak cluster |
| VP09377 | VP09377 | 2009 | China | Clinical | Asia | O3:K6 | ST3 | PC | Wave-2 | Putative outbreak cluster |
| VP09396 | VP09396 | 2009 | China | Clinical | Asia | O3:K6 | ST3 | PC | Wave-2 | Putative outbreak cluster |
| VP09401 | VP09401 | 2009 | China | Clinical | Asia | O3:K6 | ST3 | PC | Wave-2 | Putative outbreak cluster |
| VP09402 | VP09402 | 2009 | China | Clinical | Asia | O3:K6 | ST3 | PC | Wave-2 | Putative outbreak cluster |
| VP09418 | VP09418 | 2009 | China | Clinical | Asia | O3:K6 | ST3 | PC | Wave-2 | Putative outbreak cluster |
| VP09423 | VP09423 | 2009 | China | Clinical | Asia | O3:K6 | ST3 | PC | Wave-2 | Putative outbreak cluster |
| VP09437 | VP09437 | 2009 | China | Clinical | Asia | O3:K6 | ST3 | PC | Wave-2 | Putative outbreak cluster |
| VP09463 | VP09463 | 2009 | China | Clinical | Asia | O3:K6 | ST3 | PC | Wave-2 | Putative outbreak cluster |
| VP09470 | VP09470 | 2009 | China | Clinical | Asia | O3:K6 | ST3 | PC | Wave-2 | Putative outbreak cluster |
| VP09471 | VP09471 | 2009 | China | Clinical | Asia | O4:K68 | ST3 | PC | Wave-2 | Putative outbreak cluster |
| VP10002 | VP10002 | 2009 | China | Clinical | Asia | O3:K6 | ST3 | PC | Wave-2 | Putative outbreak cluster |
| VP10059 | VP10059 | 2010 | China | Env | Asia | O3:K6 | ST3 | PC | Wave-2 | Putative outbreak cluster |
| VP10060 | VP10060 | 2010 | China | Env | Asia | O3:K6 | ST3 | PC | Wave-2 | Putative outbreak cluster |
| VP10064 | VP10064 | 2010 | China | Clinical | Asia | O3:K6 | ST3 | PC | Wave-2 | Putative outbreak cluster |
| VP10066 | VP10066 | 2010 | China | Clinical | Asia | O3:K6 | ST3 | PC | Wave-2 | Putative outbreak cluster |
| VP10094 | VP10094 | 2010 | China | Clinical | Asia | O3:K6 | ST3 | PC | Wave-2 | Putative outbreak cluster |
| VP10096 | VP10096 | 2010 | China | Clinical | Asia | O3:K6 | ST3 | PC | Wave-2 | Putative outbreak cluster |
| VP10098 | VP10098 | 2010 | China | Clinical | Asia | O3:K6 | ST3 | PC | Wave-2 | Putative outbreak cluster |
| VP10099 | VP10099 | 2010 | China | Clinical | Asia | O3:K6 | ST3 | PC | Wave-2 | Putative outbreak cluster |
| VP10101 | VP10101 | 2010 | China | Clinical | Asia | O3:K6 | ST3 | PC | Wave-2 | Putative outbreak cluster |
| VP10111 | VP10111 | 2010 | China | Clinical | Asia | O3:K6 | ST3 | PC | Wave-2 | Putative outbreak cluster |

| | | | | | | | | | | |
|---|---|---|---|---|---|---|---|---|---|---|
| VP10112 | VP10112 | 2010 | China | Clinical | Asia | O3:K6 | ST3 | PC | Wave-2 | Putative outbreak cluster |
| VP10121 | VP10121 | 2010 | China | Clinical | Asia | O3:K6 | ST3 | PC | Wave-2 | Putative outbreak cluster |
| VP10123 | VP10123 | 2010 | China | Clinical | Asia | O3:K6 | ST3 | PC | Wave-2 | Putative outbreak cluster |
| VP10124 | VP10124 | 2010 | China | Clinical | Asia | O3:K6 | ST3 | PC | Wave-2 | Putative outbreak cluster |
| VP10125 | VP10125 | 2010 | China | Clinical | Asia | O3:K6 | ST3 | PC | Wave-2 | Putative outbreak cluster |
| VP10126 | VP10126 | 2010 | China | Clinical | Asia | O3:K6 | ST3 | PC | Wave-2 | Putative outbreak cluster |
| VP10144 | VP10144 | 2010 | China | Clinical | Asia | O3:K6 | ST3 | PC | Wave-2 | Putative outbreak cluster |
| VP10150 | VP10150 | 2010 | China | Clinical | Asia | O3:K6 | ST3 | PC | Wave-2 | Putative outbreak cluster |
| VP10157 | VP10157 | 2010 | China | Clinical | Asia | O3:K6 | ST3 | PC | Wave-2 | Putative outbreak cluster |
| VP10158 | VP10158 | 2010 | China | Clinical | Asia | O3:K6 | ST3 | PC | Wave-2 | Putative outbreak cluster |
| VP10164 | VP10164 | 2010 | China | Clinical | Asia | O4:K68 | ST3 | PC | Wave-2 | Putative outbreak cluster |
| VP10193 | VP10193 | 2010 | China | Clinical | Asia | O4:K68 | ST3 | PC | Wave-2 | Putative outbreak cluster |
| VP10194 | VP10194 | 2010 | China | Clinical | Asia | O3:K6 | ST3 | PC | Wave-2 | Putative outbreak cluster |
| VP10202 | VP10202 | 2010 | China | Clinical | Asia | O3:K6 | ST3 | PC | Wave-2 | Putative outbreak cluster |
| VP10203 | VP10203 | 2010 | China | Clinical | Asia | O3:K6 | ST3 | PC | Wave-2 | Putative outbreak cluster |
| VP10204 | VP10204 | 2010 | China | Clinical | Asia | O3:K6 | ST3 | PC | Wave-2 | Putative outbreak cluster |
| VP10209 | VP10209 | 2010 | China | Clinical | Asia | O3:K6 | ST3 | PC | Wave-2 | Putative outbreak cluster |
| VP10210 | VP10210 | 2010 | China | Clinical | Asia | O3:K6 | ST3 | PC | Wave-2 | Putative outbreak cluster |
| VP10211 | VP10211 | 2010 | China | Clinical | Asia | O3:K6 | ST3 | PC | Wave-2 | Putative outbreak cluster |
| VP10212 | VP10212 | 2010 | China | Clinical | Asia | O3:K6 | ST3 | PC | Wave-2 | Putative outbreak cluster |
| VP10214 | VP10214 | 2010 | China | Clinical | Asia | O4:K68 | ST3 | PC | Wave-2 | Putative outbreak cluster |
| VP10215 | VP10215 | 2010 | China | Clinical | Asia | O4:K68 | ST3 | PC | Wave-2 | Putative outbreak cluster |
| VP10232 | VP10232 | 2010 | China | Clinical | Asia | O4:K68 | ST3 | PC | Wave-2 | Putative outbreak cluster |
| VP10237 | VP10237 | 2010 | China | Clinical | Asia | O4:K68 | ST3 | PC | Wave-2 | Putative outbreak cluster |
| VP10241 | VP10241 | 2010 | China | Clinical | Asia | O4:K68 | ST3 | PC | Wave-2 | Putative outbreak cluster |
| VP10246 | VP10246 | 2010 | China | Clinical | Asia | O3:K6 | ST3 | PC | Wave-2 | Putative outbreak cluster |
| VP10247 | VP10247 | 2010 | China | Clinical | Asia | O3:K6 | ST3 | PC | Wave-2 | Putative outbreak cluster |
| VP10258 | VP10258 | 2010 | China | Clinical | Asia | O3:K6 | ST3 | PC | Wave-2 | Putative outbreak cluster |
| VP10260 | VP10260 | 2010 | China | Clinical | Asia | O4:K68 | ST3 | PC | Wave-2 | Putative outbreak cluster |
| VP10269 | VP10269 | 2010 | China | Clinical | Asia | O3:K6 | ST3 | PC | Wave-2 | Putative outbreak cluster |
| VP10274 | VP10274 | 2010 | China | Clinical | Asia | O3:K6 | ST3 | PC | Wave-2 | Putative outbreak cluster |
| VP10277 | VP10277 | 2010 | China | Clinical | Asia | O3:K6 | ST3 | PC | Wave-2 | Putative outbreak cluster |

| | | | | | | | | | | |
|---|---|---|---|---|---|---|---|---|---|---|
| VP10279 | VP10279 | 2010 | China | Clinical | Asia | O3:K6 | ST3 | PC | Wave-2 | Putative outbreak cluster |
| VP10283 | VP10283 | 2010 | China | Clinical | Asia | O3:K6 | ST3 | PC | Wave-2 | Putative outbreak cluster |
| VP10292 | VP10292 | 2010 | China | Clinical | Asia | O3:K6 | ST3 | PC | Wave-2 | Putative outbreak cluster |
| VP10293 | VP10293 | 2010 | China | Clinical | Asia | O3:K6 | ST431 | PC | Wave-2 | Putative outbreak cluster |
| VP10312 | VP10312 | 2010 | China | Clinical | Asia | O3:K6 | ST3 | PC | Wave-2 | Putative outbreak cluster |
| VP10314 | VP10314 | 2010 | China | Clinical | Asia | O3:K6 | ST3 | PC | Wave-2 | Putative outbreak cluster |
| VP10315 | VP10315 | 2010 | China | Clinical | Asia | O3:K6 | ST3 | PC | Wave-2 | Putative outbreak cluster |
| VP10316 | VP10316 | 2010 | China | Clinical | Asia | O4:K68 | ST3 | PC | Wave-2 | Putative outbreak cluster |
| VP10342 | VP10342 | 2010 | China | Clinical | Asia | O3:K6 | ST3 | PC | Wave-2 | Putative outbreak cluster |
| VP10345 | VP10345 | 2010 | China | Clinical | Asia | O3:K6 | ST3 | PC | Wave-2 | Putative outbreak cluster |
| VP10353 | VP10353 | 2010 | China | Clinical | Asia | O3:K6 | ST431 | PC | Wave-2 | Putative outbreak cluster |
| VP10369 | VP10369 | 2010 | China | Clinical | Asia | O3:K6 | ST3 | PC | Wave-2 | Putative outbreak cluster |
| VP10380 | VP10380 | 2010 | China | Clinical | Asia | O3:K6 | ST3 | PC | Wave-2 | Putative outbreak cluster |
| VP10388 | VP10388 | 2010 | China | Clinical | Asia | O3:K6 | ST3 | PC | Wave-2 | Putative outbreak cluster |
| VP10397 | VP10397 | 2010 | China | Clinical | Asia | O3:K6 | ST3 | PC | Wave-2 | Putative outbreak cluster |
| VP10398 | VP10398 | 2010 | China | Clinical | Asia | O3:K6 | ST431 | PC | Wave-2 | Putative outbreak cluster |
| VP10399 | VP10399 | 2010 | China | Clinical | Asia | O3:K6 | ST3 | PC | Wave-2 | Putative outbreak cluster |
| VP10401 | VP10401 | 2010 | China | Clinical | Asia | O3:K6 | ST3 | PC | Wave-2 | Putative outbreak cluster |
| VP10410 | VP10410 | 2010 | China | Clinical | Asia | O3:K6 | ST3 | PC | Wave-2 | Putative outbreak cluster |
| VP10425 | VP10425 | 2010 | China | Clinical | Asia | O3:K6 | ST3 | PC | Wave-2 | Putative outbreak cluster |
| VP10435 | VP10435 | 2010 | China | Clinical | Asia | O3:K6 | ST431 | PC | Wave-2 | Putative outbreak cluster |
| VP10437 | VP10437 | 2010 | China | Clinical | Asia | O3:K6 | ST3 | PC | Wave-2 | Putative outbreak cluster |
| VP10450 | VP10450 | 2010 | China | Clinical | Asia | O3:K6 | ST3 | PC | Wave-2 | Putative outbreak cluster |
| VP10453 | VP10453 | 2010 | China | Clinical | Asia | O3:K6 | ST3 | PC | Wave-2 | Putative outbreak cluster |
| VP10454 | VP10454 | 2010 | China | Clinical | Asia | O3:K6 | ST3 | PC | Wave-2 | Putative outbreak cluster |
| VP10474 | VP10474 | 2010 | China | Clinical | Asia | O3:K6 | ST3 | PC | Wave-2 | Putative outbreak cluster |
| VP11042 | VP11042 | 2011 | China | Clinical | Asia | O3:K6 | ST3 | PC | Wave-2 | Putative outbreak cluster |
| VP11046 | VP11046 | 2011 | China | Clinical | Asia | O3:K6 | ST3 | PC | Wave-2 | Putative outbreak cluster |
| VP11123 | VP11123 | 2011 | China | Clinical | Asia | O3:K6 | ST3 | PC | Wave-2 | Putative outbreak cluster |
| VP11131 | VP11131 | 2011 | China | Clinical | Asia | O3:K6 | ST3 | PC | Wave-2 | Putative outbreak cluster |
| VP11142 | VP11142 | 2011 | China | Clinical | Asia | O3:K6 | ST3 | PC | Wave-2 | Putative outbreak cluster |
| VP11144 | VP11144 | 2011 | China | Clinical | Asia | O3:K6 | ST3 | PC | Wave-2 | Putative outbreak cluster |

| | | | | | | | | | | |
|---|---|---|---|---|---|---|---|---|---|---|
| VP11145 | VP11145 | 2011 | China | Clinical | Asia | O3:K6 | ST3 | PC | Wave-2 | Putative outbreak cluster |
| VP11152 | VP11152 | 2011 | China | Clinical | Asia | O3:K6 | ST3 | PC | Wave-2 | Putative outbreak cluster |
| VP11153 | VP11153 | 2011 | China | Clinical | Asia | O3:K6 | ST3 | PC | Wave-2 | Putative outbreak cluster |
| VP11154 | VP11154 | 2011 | China | Clinical | Asia | O3:K6 | ST3 | PC | Wave-2 | Putative outbreak cluster |
| VP11161 | VP11161 | 2011 | China | Clinical | Asia | O3:K6 | ST3 | PC | Wave-2 | Putative outbreak cluster |
| VP11170 | VP11170 | 2011 | China | Clinical | Asia | O3:K6 | ST3 | PC | Wave-2 | Putative outbreak cluster |
| VP11171 | VP11171 | 2011 | China | Clinical | Asia | O3:K6 | ST3 | PC | Wave-2 | Putative outbreak cluster |
| VP11174 | VP11174 | 2011 | China | Clinical | Asia | O3:K6 | ST3 | PC | Wave-2 | Putative outbreak cluster |
| VP11176 | VP11176 | 2011 | China | Clinical | Asia | O4:K68 | ST3 | PC | Wave-2 | Putative outbreak cluster |
| VP11178 | VP11178 | 2011 | China | Clinical | Asia | O3:K6 | ST3 | PC | Wave-2 | Putative outbreak cluster |
| VP11182 | VP11182 | 2011 | China | Clinical | Asia | O3:K6 | ST3 | PC | Wave-2 | Putative outbreak cluster |
| VP11183 | VP11183 | 2011 | China | Clinical | Asia | O3:K6 | ST3 | PC | Wave-2 | Putative outbreak cluster |
| VP11184 | VP11184 | 2011 | China | Clinical | Asia | O3:K6 | ST3 | PC | Wave-2 | Putative outbreak cluster |
| VP11187 | VP11187 | 2011 | China | Clinical | Asia | O3:K6 | ST3 | PC | Wave-2 | Putative outbreak cluster |
| VP11192 | VP11192 | 2011 | China | Clinical | Asia | O3:K6 | ST3 | PC | Wave-2 | Putative outbreak cluster |
| VP11196 | VP11196 | 2011 | China | Clinical | Asia | O4:K68 | ST3 | PC | Wave-2 | Putative outbreak cluster |
| VP11197 | VP11197 | 2011 | China | Clinical | Asia | O3:K6 | ST3 | PC | Wave-2 | Putative outbreak cluster |
| VP11200 | VP11200 | 2011 | China | Clinical | Asia | O3:K6 | ST3 | PC | Wave-2 | Putative outbreak cluster |
| VP11241 | VP11241 | 2011 | China | Clinical | Asia | O3:K6 | ST3 | PC | Wave-2 | Putative outbreak cluster |
| VP11247 | VP11247 | 2011 | China | Clinical | Asia | O3:K6 | ST3 | PC | Wave-2 | Putative outbreak cluster |
| VP11249 | VP11249 | 2011 | China | Clinical | Asia | O3:K6 | ST3 | PC | Wave-2 | Putative outbreak cluster |
| VP11251 | VP11251 | 2011 | China | Clinical | Asia | O3:K6 | ST3 | PC | Wave-2 | Putative outbreak cluster |
| VP11258 | VP11258 | 2011 | China | Clinical | Asia | O3:K6 | ST3 | PC | Wave-2 | Putative outbreak cluster |
| VP11261 | VP11261 | 2011 | China | Clinical | Asia | O3:K6 | ST3 | PC | Wave-2 | Putative outbreak cluster |
| VP11275 | VP11275 | 2011 | China | Clinical | Asia | O3:K6 | ST3 | PC | Wave-2 | Putative outbreak cluster |
| VP11282 | VP11282 | 2011 | China | Clinical | Asia | O3:K6 | ST3 | PC | Wave-2 | Putative outbreak cluster |
| VP11290 | VP11290 | 2011 | China | Clinical | Asia | O3:K6 | ST3 | PC | Wave-2 | Putative outbreak cluster |
| VP11295 | VP11295 | 2011 | China | Clinical | Asia | O3:K6 | ST3 | PC | Wave-2 | Putative outbreak cluster |
| VP11296 | VP11296 | 2011 | China | Clinical | Asia | O3:K6 | ST3 | PC | Wave-2 | Putative outbreak cluster |
| VP12123 | VP12123 | 2012 | China | Clinical | Asia | O4:K68 | ST3 | PC | Wave-2 | Putative outbreak cluster |
| VP13060 | VP13060 | 2013 | China | Clinical | Asia | O3:K6 | ST3 | PC | Wave-2 | Putative outbreak cluster |
| VP13071 | VP13071 | 2013 | China | Clinical | Asia | O1:KUK3 | ST3 | PC | Wave-2 | Putative outbreak cluster |

| | | | | | | | | | | |
|---|---|---|---|---|---|---|---|---|---|---|
| VP15161 | VP15161 | 2015 | China | Clinical | Asia | O3:K6 | ST3 | PC | Wave-2 | Putative outbreak cluster |
| VP16007 | VP16007 | 2016 | China | Clinical | Asia | O3:K6 | ST3 | PC | Wave-2 | Putative outbreak cluster |
| VP16159 | VP16159 | 2016 | China | Clinical | Asia | O3:K6 | ST3 | PC | Wave-2 | Putative outbreak cluster |
| VP16160 | VP16160 | 2016 | China | Clinical | Asia | O3:K6 | ST3 | PC | Wave-2 | Putative outbreak cluster |
| VP16161 | VP16161 | 2016 | China | Clinical | Asia | O3:K6 | ST3 | PC | Wave-2 | Putative outbreak cluster |
| VP16162 | VP16162 | 2016 | China | Clinical | Asia | O3:K6 | ST3 | PC | Wave-2 | Putative outbreak cluster |
| VP16163 | VP16163 | 2016 | China | Clinical | Asia | O3:K6 | ST3 | PC | Wave-2 | Putative outbreak cluster |
| VP16164 | VP16164 | 2016 | China | Clinical | Asia | O3:K6 | ST3 | PC | Wave-2 | Putative outbreak cluster |
| VP17008 | VP17008 | 2017 | China | Clinical | Asia | O3:K6 | ST3 | PC | Wave-2 | Putative outbreak cluster |
| VP18011 | VP18011 | 2018 | China | Clinical | Asia | O3:K6 | ST3 | PC | Wave-2 | Putative outbreak cluster |
| VP18013 | VP18013 | 2018 | China | Clinical | Asia | O3:K6 | ST3 | PC | Wave-2 | Putative outbreak cluster |
| VP18014 | VP18014 | 2018 | China | Clinical | Asia | O3:K6 | ST3 | PC | Wave-2 | Putative outbreak cluster |
| VP18015 | VP18015 | 2018 | China | Clinical | Asia | O3:K6 | ST3 | PC | Wave-2 | Putative outbreak cluster |
| VP18017 | VP18017 | 2018 | China | Clinical | Asia | O3:K6 | ST3 | PC | Wave-2 | Putative outbreak cluster |
| VP18025 | VP18025 | 2018 | China | Clinical | Asia | O3:K6 | ST3 | PC | Wave-2 | Putative outbreak cluster |
| SAMN12084703-GCA_015787915.1 | PNUSAV000640 | NA | USA | Clinical | US_Canada | O3:K6 | ST3 | PC | Wave-2 | Putative outbreak cluster |
| SAMN12084709-GCA_015789555.1 | PNUSAV000638 | NA | USA | Clinical | US_Canada | O3:K6 | ST3 | PC | Wave-2 | Putative outbreak cluster |
| SAMN12084717-GCA_015789615.1 | PNUSAV000639 | NA | USA | Clinical | US_Canada | O3:K6 | ST3 | PC | Wave-2 | Putative outbreak cluster |
| SAMN12084723-GCA_015787895.1 | PNUSAV000637 | NA | USA | Clinical | US_Canada | O3:K6 | ST3 | PC | Wave-2 | Putative outbreak cluster |
| SAMN12084766-GCA_015787955.1 | PNUSAV000641 | NA | USA | Clinical | US_Canada | O3:K6 | ST3 | PC | Wave-2 | Putative outbreak cluster |
| SAMN12211985-GCA_015789815.1 | PNUSAV000654 | NA | USA | Clinical | US_Canada | O3:K6 | ST3 | PC | Wave-2 | Putative outbreak cluster |
| SAMN12716022-GCA_015784765.1 | PNUSAV000997 | NA | USA | Clinical | US_Canada | O3:K6 | ST3 | PC | Wave-2 | Putative outbreak cluster |
| SAMN13746682-GCA_015775955.1 | PNUSAV001089 | NA | USA | Clinical | US_Canada | O3:K6 | ST3 | PC | Wave-2 | Putative outbreak cluster |
| SAMN02597369-GCF_000516875.2 | AZKN | 2004 | Mozambique | Clinical | Africa | O3:K6 | ST3 | PC | Wave-2 | Representative |
| SAMEA8103029-GCA_905331775.1 | ERS5790107 | 2016 | China | Clinical | Asia | O1:KUT6 | ST3 | PC | Wave-2 | Representative |
| SAMN01923802-GCF_000571915.2 | EKP-021 | 2008 | Bangladesh | NA | Asia | O4:K68 | ST3 | PC | Wave-2 | Representative |
| SAMN02189718-GCF_000454265.1 | NIHCB0603 | 2006 | Bangladesh | NA | Asia | O3:K6 | ST3 | PC | Wave-2 | Representative |
| SAMN02189721-GCF_000454225.1 | PVCHO_VP250 | 1998 | India | Clinical | Asia | O1:KUK3 | ST3 | PC | Wave-2 | Representative |
| SAMN02190056-GCF_000454185.1 | PVCHO_VP232 | 1998 | India | Clinical | Asia | O4:K68 | ST3 | PC | Wave-2 | Representative |
| SAMN02338920-GCF_000490995.1 | S063 | 1998 | China | Clinical | Asia | O4:K68 | ST3 | PC | Wave-2 | Representative |
| SAMN02338921-GCF_000490975.1 | S064 | 1998 | China | Clinical | Asia | O3:K6 | ST3 | PC | Wave-2 | Representative |
| SAMN02338925-GCF_000490895.1 | S068 | 1997 | China | Clinical | Asia | O3:K6 | ST3 | PC | Wave-2 | Representative |

| | | | | | | | | | | |
|---|---|---|---|---|---|---|---|---|---|---|
| SAMN02338926-GCF_000490875.1 | S069 | NA | Thailand | Clinical | Asia | O3:K6 | ST- | PC | Wave-2 | Representative |
| SAMN02338927-GCF_000490855.1 | S070 | NA | Thailand | Clinical | Asia | O3:K6 | ST- | PC | Wave-2 | Representative |
| SAMN02338928-GCF_000490835.1 | S071 | 1998 | Bangladesh | Clinical | Asia | O1:KUK3 | ST3 | PC | Wave-2 | Representative |
| SAMN02338929-GCF_000490815.1 | S072 | 1998 | Bangladesh | Clinical | Asia | O3:K6 | ST3 | PC | Wave-2 | Representative |
| SAMN02338931-GCF_000490775.1 | S074 | 1997 | China | Clinical | Asia | O3:K6 | ST3 | PC | Wave-2 | Representative |
| SAMN02338932-GCF_000490755.1 | S075 | 1999 | China | Clinical | Asia | O3:K6 | ST3 | PC | Wave-2 | Representative |
| SAMN02338933-GCF_000490735.1 | S076 | 1999 | China | Clinical | Asia | O3:K6 | ST3 | PC | Wave-2 | Representative |
| SAMN02338934-GCF_000490715.1 | S077 | 1999 | China | Clinical | Asia | O3:K6 | ST3 | PC | Wave-2 | Representative |
| SAMN02338935-GCF_000490695.1 | S078 | 1999 | China | Clinical | Asia | O3:K6 | ST3 | PC | Wave-2 | Representative |
| SAMN02338936-GCF_000490675.1 | S079 | NA | Indonesia | Clinical | Asia | O3:K6 | ST3 | PC | Wave-2 | Representative |
| SAMN02338938-GCF_000490635.1 | S082 | NA | Thailand | Clinical | Asia | O3:K6 | ST- | PC | Wave-2 | Representative |
| SAMN02338939-GCF_000490615.1 | S083 | 1998 | Japan | Clinical | Asia | O3:K6 | ST3 | PC | Wave-2 | Representative |
| SAMN02338941-GCF_000490575.1 | S087 | 1998 | Singapore | Clinical | Asia | O4:K68 | ST- | PC | Wave-2 | Representative |
| SAMN02338942-GCF_000490555.1 | S088 | 1998 | Singapore | Clinical | Asia | O4:K68 | ST3 | PC | Wave-2 | Representative |
| SAMN02338943-GCF_000490535.1 | S090 | 1999 | China | Clinical | Asia | O4:K68 | ST- | PC | Wave-2 | Representative |
| SAMN02338944-GCF_000490515.1 | S091 | 1999 | India | Clinical | Asia | O4:K68 | ST3 | PC | Wave-2 | Representative |
| SAMN02338945-GCF_000493715.1 | S092 | 1996 | China | Clinical | Asia | O3:K6 | ST3 | PC | Wave-2 | Representative |
| SAMN02338975-GCF_000489915.1 | S126 | NA | China | Clinical | Asia | O3:K6 | ST3 | PC | Wave-2 | Representative |
| SAMN02338986-GCF_000489695.1 | S138 | 2007 | China | Clinical | Asia | O3:K6 | ST3 | PC | Wave-2 | Representative |
| SAMN02436183-GCF_000182465.1 | ACKB | 2005 | India | Clinical | Asia | O3:K6 | ST3 | PC | Wave-2 | Representative |
| SAMN02436265-GCF_000182385.1 | ACFO | 1998 | Bangladesh | Clinical | Asia | O4:K68 | ST3 | PC | Wave-2 | Representative |
| SAMN02471129-GCA_000500385.1 | VIP4-0445 | 2008 | China | Clinical | Asia | O3:K6 | ST- | PC | Wave-2 | Representative |
| SAMN04875528-GCF_002073775.2 | FDA:FDAARGOS_191 | 1996 | India | Clinical | Asia | O3:K6 | ST3 | PC | Wave-2 | Representative |
| SAMN05935536-GCF_001895435.1 | 100152 | 2010 | China | Clinical | Asia | O3:K6 | ST3 | PC | Wave-2 | Representative |
| SAMN06076986-GCA_015819055.1 | CFSAN023533 | 1996 | India | Clinical | Asia | O3:K6 | ST3 | PC | Wave-2 | Representative |
| SAMN06076990-GCA_015818955.1 | AN-8373 | 1998 | Bangladesh | Clinical | Asia | O3:K6 | ST3 | PC | Wave-2 | Representative |
| SAMN06076996-GCA_015818855.1 | AN-5034 | 1998 | Bangladesh | Clinical | Asia | O4:K68 | ST3 | PC | Wave-2 | Representative |
| SAMN06076999-GCA_015819015.1 | AN-16000 | 1998 | Bangladesh | Clinical | Asia | O1:KUK3 | ST3 | PC | Wave-2 | Representative |
| SAMN06077014-GCA_015817735.1 | CFSAN023563 | 1999 | Thailand | Clinical | Asia | O4:K68 | ST3 | PC | Wave-2 | Representative |
| SAMN06077016-GCA_015817775.1 | AN-2189 | 1998 | Bangladesh | Clinical | Asia | O4:K68 | ST3 | PC | Wave-2 | Representative |
| SAMN06077017-GCA_015817595.1 | AP-11243 | 2000 | Bangladesh | Clinical | Asia | O1:KUK3 | ST3 | PC | Wave-2 | Representative |
| SAMN09742582-GCF_006383385.1 | QP16184 | 2016 | China | Clinical | Asia | O3:KUT7 | ST3 | PC | Wave-2 | Representative |

| | | | | | | | | | | |
|---|---|---|---|---|---|---|---|---|---|---|
| SAMN15294480-GCA_019685775.1 | ICDC-VP01809 | 2018 | China | Clinical | Asia | O3:K6 | ST3 | PC | Wave-2 | Representative |
| SAMN16783018-GCA_016827145.1 | VP90 | 2009 | China | Clinical | Asia | O4:K68 | ST3 | PC | Wave-2 | Representative |
| SAMN16783025-GCA_016827015.1 | VP100 | 2010 | China | Clinical | Asia | O4:K68 | ST3 | PC | Wave-2 | Representative |
| SAMN16783065-GCA_016826215.1 | VP142 | 2012 | China | NA | Asia | O4:K68 | ST3 | PC | Wave-2 | Representative |
| SAMN16783104-GCA_016825475.1 | VP185 | 2011 | China | NA | Asia | O4:K68 | ST3 | PC | Wave-2 | Representative |
| SAMN17188289-SRR14373889 | NB327 | 2017 | China | Env | Asia | O4:K68 | ST3 | PC | Wave-2 | Representative |
| VP02003 | VP02003 | 2002 | China | Clinical | Asia | O3:K6 | ST3 | PC | Wave-2 | Representative |
| VP04083 | VP04083 | 2004 | China | Clinical | Asia | O3:K6 | ST3 | PC | Wave-2 | Representative |
| VP04121 | VP04121 | 2004 | China | Clinical | Asia | O4:K68 | ST3 | PC | Wave-2 | Representative |
| VP04126 | VP04126 | 2004 | China | Clinical | Asia | O4:K68 | ST3 | PC | Wave-2 | Representative |
| VP04127 | VP04127 | 2004 | China | Clinical | Asia | O4:K68 | ST3 | PC | Wave-2 | Representative |
| VP04141 | VP04141 | 2004 | China | Clinical | Asia | O3:K6 | ST3 | PC | Wave-2 | Representative |
| VP04159 | VP04159 | 2004 | China | Clinical | Asia | O3:K6 | ST3 | PC | Wave-2 | Representative |
| VP05223 | VP05223 | 2005 | China | Clinical | Asia | O3:K6 | ST3 | PC | Wave-2 | Representative |
| VP05277 | VP05277 | 2005 | China | Clinical | Asia | O3:K6 | ST3 | PC | Wave-2 | Representative |
| VP06013 | VP06013 | 2006 | China | Env | Asia | O3:K6 | ST3 | PC | Wave-2 | Representative |
| VP06084 | VP06084 | 2006 | China | Clinical | Asia | O3:K6 | ST3 | PC | Wave-2 | Representative |
| VP06085 | VP06085 | 2006 | China | Clinical | Asia | O3:K6 | ST3 | PC | Wave-2 | Representative |
| VP06088 | VP06088 | 2006 | China | Clinical | Asia | O1:KUK3 | ST3 | PC | Wave-2 | Representative |
| VP06094 | VP06094 | 2006 | China | Clinical | Asia | O4:K68 | ST3 | PC | Wave-2 | Representative |
| VP06113 | VP06113 | 2006 | China | Clinical | Asia | O3:K6 | ST3 | PC | Wave-2 | Representative |
| VP06132 | VP06132 | 2006 | China | Clinical | Asia | O3:K6 | ST3 | PC | Wave-2 | Representative |
| VP06176 | VP06176 | 2006 | China | Clinical | Asia | O3:K6 | ST3 | PC | Wave-2 | Representative |
| VP06192 | VP06192 | 2006 | China | Clinical | Asia | O3:K6 | ST3 | PC | Wave-2 | Representative |
| VP06194 | VP06194 | 2006 | China | Clinical | Asia | O3:K6 | ST3 | PC | Wave-2 | Representative |
| VP06195 | VP06195 | 2006 | China | Clinical | Asia | O3:K6 | ST3 | PC | Wave-2 | Representative |
| VP06197 | VP06197 | 2006 | China | Clinical | Asia | O3:K6 | ST3 | PC | Wave-2 | Representative |
| VP07012 | VP07012 | 2007 | China | Clinical | Asia | O3:K6 | ST3 | PC | Wave-2 | Representative |
| VP07065 | VP07065 | 2007 | China | Clinical | Asia | O3:K6 | ST3 | PC | Wave-2 | Representative |
| VP07099 | VP07099 | 2007 | China | Clinical | Asia | O3:K6 | ST3 | PC | Wave-2 | Representative |
| VP07112 | VP07112 | 2007 | China | Clinical | Asia | O3:K6 | ST3 | PC | Wave-2 | Representative |
| VP07132 | VP07132 | 2007 | China | Clinical | Asia | O3:K6 | ST3 | PC | Wave-2 | Representative |

| | | | | | | | | | | |
|---|---|---|---|---|---|---|---|---|---|---|
| VP07204 | VP07204 | 2007 | China | Clinical | Asia | O3:K6 | ST3 | PC | Wave-2 | Representative |
| VP07241 | VP07241 | 2007 | China | Clinical | Asia | O3:K6 | ST3 | PC | Wave-2 | Representative |
| VP07246 | VP07246 | 2007 | China | Clinical | Asia | O4:K68 | ST3 | PC | Wave-2 | Representative |
| VP07253 | VP07253 | 2007 | China | Clinical | Asia | O3:K6 | ST3 | PC | Wave-2 | Representative |
| VP08015 | VP08015 | 2008 | China | Clinical | Asia | O3:K6 | ST3 | PC | Wave-2 | Representative |
| VP08058 | VP08058 | 2008 | China | Clinical | Asia | O4:K68 | ST3 | PC | Wave-2 | Representative |
| VP08067 | VP08067 | 2008 | China | Clinical | Asia | O4:K68 | ST3 | PC | Wave-2 | Representative |
| VP08168 | VP08168 | 2008 | China | Clinical | Asia | O3:K6 | ST3 | PC | Wave-2 | Representative |
| VP08173 | VP08173 | 2008 | China | Clinical | Asia | O3:K6 | ST3 | PC | Wave-2 | Representative |
| VP08203 | VP08203 | 2008 | China | Clinical | Asia | O3:K6 | ST3 | PC | Wave-2 | Representative |
| VP08205 | VP08205 | 2008 | China | Clinical | Asia | O3:K6 | ST3 | PC | Wave-2 | Representative |
| VP08225 | VP08225 | 2008 | China | Clinical | Asia | O4:K68 | ST3 | PC | Wave-2 | Representative |
| VP08228 | VP08228 | 2008 | China | Clinical | Asia | O3:K6 | ST3 | PC | Wave-2 | Representative |
| VP08230 | VP08230 | 2008 | China | Clinical | Asia | O3:K6 | ST3 | PC | Wave-2 | Representative |
| VP08248 | VP08248 | 2008 | China | Clinical | Asia | O4:K68 | ST3 | PC | Wave-2 | Representative |
| VP08337 | VP08337 | 2008 | China | Clinical | Asia | O3:K6 | ST3 | PC | Wave-2 | Representative |
| VP08345 | VP08345 | 2008 | China | Clinical | Asia | O3:K6 | ST3 | PC | Wave-2 | Representative |
| VP08363 | VP08363 | 2008 | China | Clinical | Asia | O3:K6 | ST3 | PC | Wave-2 | Representative |
| VP08370 | VP08370 | 2008 | China | Clinical | Asia | O3:K6 | ST3 | PC | Wave-2 | Representative |
| VP08375 | VP08375 | 2008 | China | Clinical | Asia | O3:K6 | ST3 | PC | Wave-2 | Representative |
| VP09014 | VP09014 | 2009 | China | Clinical | Asia | O3:K6 | ST431 | PC | Wave-2 | Representative |
| VP09020 | VP09020 | 2009 | China | Clinical | Asia | O3:K6 | ST3 | PC | Wave-2 | Representative |
| VP09026 | VP09026 | 2009 | China | Clinical | Asia | O3:K6 | ST3 | PC | Wave-2 | Representative |
| VP09078 | VP09078 | 2009 | China | Clinical | Asia | O3:K6 | ST3 | PC | Wave-2 | Representative |
| VP09107 | VP09107 | 2009 | China | Clinical | Asia | O4:K68 | ST3 | PC | Wave-2 | Representative |
| VP09110 | VP09110 | 2009 | China | Clinical | Asia | O3:K6 | ST3 | PC | Wave-2 | Representative |
| VP09147 | VP09147 | 2009 | China | Clinical | Asia | O4:K68 | ST3 | PC | Wave-2 | Representative |
| VP09167 | VP09167 | 2009 | China | Clinical | Asia | O3:K6 | ST3 | PC | Wave-2 | Representative |
| VP09185 | VP09185 | 2009 | China | Clinical | Asia | O3:K6 | ST3 | PC | Wave-2 | Representative |
| VP09193 | VP09193 | 2009 | China | Clinical | Asia | O3:K6 | ST3 | PC | Wave-2 | Representative |
| VP09238 | VP09238 | 2009 | China | Clinical | Asia | O3:K6 | ST3 | PC | Wave-2 | Representative |
| VP09240 | VP09240 | 2009 | China | Clinical | Asia | O3:K6 | ST3 | PC | Wave-2 | Representative |

| | | | | | | | | | | |
|---|---|---|---|---|---|---|---|---|---|---|
| VP09243 | VP09243 | 2009 | China | Clinical | Asia | O4:K68 | ST3 | PC | Wave-2 | Representative |
| VP09245 | VP09245 | 2009 | China | Clinical | Asia | O3:K6 | ST3 | PC | Wave-2 | Representative |
| VP09272 | VP09272 | 2009 | China | Clinical | Asia | O3:K6 | ST3 | PC | Wave-2 | Representative |
| VP09300 | VP09300 | 2009 | China | Clinical | Asia | O3:K6 | ST3 | PC | Wave-2 | Representative |
| VP09308 | VP09308 | 2009 | China | Clinical | Asia | O3:K6 | ST3 | PC | Wave-2 | Representative |
| VP09316 | VP09316 | 2009 | China | Clinical | Asia | O3:K6 | ST3 | PC | Wave-2 | Representative |
| VP09322 | VP09322 | 2009 | China | Clinical | Asia | O4:K68 | ST3 | PC | Wave-2 | Representative |
| VP09339 | VP09339 | 2009 | China | Clinical | Asia | O3:K6 | ST3 | PC | Wave-2 | Representative |
| VP09359 | VP09359 | 2009 | China | Clinical | Asia | O3:K6 | ST3 | PC | Wave-2 | Representative |
| VP09361 | VP09361 | 2009 | China | Clinical | Asia | O4:K68 | ST3 | PC | Wave-2 | Representative |
| VP09368 | VP09368 | 2009 | China | Clinical | Asia | O3:K6 | ST3 | PC | Wave-2 | Representative |
| VP09417 | VP09417 | 2009 | China | Clinical | Asia | O3:K6 | ST3 | PC | Wave-2 | Representative |
| VP09419 | VP09419 | 2009 | China | Clinical | Asia | O3:K6 | ST3 | PC | Wave-2 | Representative |
| VP09424 | VP09424 | 2009 | China | Clinical | Asia | O3:K6 | ST3 | PC | Wave-2 | Representative |
| VP09444 | VP09444 | 2009 | China | Clinical | Asia | O3:K6 | ST3 | PC | Wave-2 | Representative |
| VP09458 | VP09458 | 2009 | China | Clinical | Asia | O3:K6 | ST3 | PC | Wave-2 | Representative |
| VP09462 | VP09462 | 2009 | China | Clinical | Asia | O3:K6 | ST3 | PC | Wave-2 | Representative |
| VP09467 | VP09467 | 2009 | China | Clinical | Asia | O3:K6 | ST3 | PC | Wave-2 | Representative |
| VP09478 | VP09478 | 2009 | China | Clinical | Asia | O3:K6 | ST3 | PC | Wave-2 | Representative |
| VP09481 | VP09481 | 2009 | China | Clinical | Asia | O1:KUK3 | ST3 | PC | Wave-2 | Representative |
| VP10065 | VP10065 | 2010 | China | Clinical | Asia | O3:K6 | ST3 | PC | Wave-2 | Representative |
| VP10068 | VP10068 | 2010 | China | Clinical | Asia | O3:K6 | ST3 | PC | Wave-2 | Representative |
| VP10100 | VP10100 | 2010 | China | Clinical | Asia | O3:K6 | ST3 | PC | Wave-2 | Representative |
| VP10114 | VP10114 | 2010 | China | Clinical | Asia | O4:K68 | ST3 | PC | Wave-2 | Representative |
| VP10116 | VP10116 | 2010 | China | Clinical | Asia | O3:K6 | ST3 | PC | Wave-2 | Representative |
| VP10128 | VP10128 | 2010 | China | Clinical | Asia | O3:K6 | ST3 | PC | Wave-2 | Representative |
| VP10130 | VP10130 | 2010 | China | Clinical | Asia | O3:K6 | ST3 | PC | Wave-2 | Representative |
| VP10145 | VP10145 | 2010 | China | Clinical | Asia | O3:K6 | ST3 | PC | Wave-2 | Representative |
| VP10152 | VP10152 | 2010 | China | Clinical | Asia | O4:K68 | ST3 | PC | Wave-2 | Representative |
| VP10163 | VP10163 | 2010 | China | Clinical | Asia | O3:K6 | ST3 | PC | Wave-2 | Representative |
| VP10197 | VP10197 | 2010 | China | Clinical | Asia | O3:K6 | ST3 | PC | Wave-2 | Representative |
| VP10206 | VP10206 | 2010 | China | Clinical | Asia | O3:K6 | ST3 | PC | Wave-2 | Representative |

| | | | | | | | | | | |
|---|---|---|---|---|---|---|---|---|---|---|
| VP10242 | VP10242 | 2010 | China | Clinical | Asia | O3:K6 | ST3 | PC | Wave-2 | Representative |
| VP10248 | VP10248 | 2010 | China | Clinical | Asia | O3:K6 | ST3 | PC | Wave-2 | Representative |
| VP10251 | VP10251 | 2010 | China | Clinical | Asia | O4:K68 | ST3 | PC | Wave-2 | Representative |
| VP10273 | VP10273 | 2010 | China | Clinical | Asia | O3:K6 | ST3 | PC | Wave-2 | Representative |
| VP10275 | VP10275 | 2010 | China | Clinical | Asia | O3:K6 | ST431 | PC | Wave-2 | Representative |
| VP10294 | VP10294 | 2010 | China | Clinical | Asia | O3:K6 | ST3 | PC | Wave-2 | Representative |
| VP10300 | VP10300 | 2010 | China | Clinical | Asia | O3:K6 | ST3 | PC | Wave-2 | Representative |
| VP10319 | VP10319 | 2010 | China | Clinical | Asia | O3:K6 | ST3 | PC | Wave-2 | Representative |
| VP10350 | VP10350 | 2010 | China | Clinical | Asia | O3:K6 | ST431 | PC | Wave-2 | Representative |
| VP10363 | VP10363 | 2010 | China | Clinical | Asia | O3:K6 | ST3 | PC | Wave-2 | Representative |
| VP10364 | VP10364 | 2010 | China | Clinical | Asia | O4:K68 | ST3 | PC | Wave-2 | Representative |
| VP10382 | VP10382 | 2010 | China | Clinical | Asia | O3:K6 | ST3 | PC | Wave-2 | Representative |
| VP10414 | VP10414 | 2010 | China | Clinical | Asia | O3:K6 | ST3 | PC | Wave-2 | Representative |
| VP10422 | VP10422 | 2010 | China | Clinical | Asia | O3:K6 | ST3 | PC | Wave-2 | Representative |
| VP10427 | VP10427 | 2010 | China | Clinical | Asia | O3:K6 | ST3 | PC | Wave-2 | Representative |
| VP10434 | VP10434 | 2010 | China | Clinical | Asia | O4:K68 | ST3 | PC | Wave-2 | Representative |
| VP10440 | VP10440 | 2010 | China | Clinical | Asia | O4:K68 | ST3 | PC | Wave-2 | Representative |
| VP10451 | VP10451 | 2010 | China | Clinical | Asia | O3:K6 | ST3 | PC | Wave-2 | Representative |
| VP10455 | VP10455 | 2010 | China | Clinical | Asia | O3:K6 | ST3 | PC | Wave-2 | Representative |
| VP10463 | VP10463 | 2010 | China | Clinical | Asia | O3:K6 | ST3 | PC | Wave-2 | Representative |
| VP10468 | VP10468 | 2010 | China | Clinical | Asia | O3:K6 | ST3 | PC | Wave-2 | Representative |
| VP10491 | VP10491 | 2010 | China | Clinical | Asia | O3:K6 | ST3 | PC | Wave-2 | Representative |
| VP11015 | VP11015 | 2011 | China | Clinical | Asia | O3:K6 | ST3 | PC | Wave-2 | Representative |
| VP11048 | VP11048 | 2011 | China | Clinical | Asia | O3:K6 | ST3 | PC | Wave-2 | Representative |
| VP11082 | VP11082 | 2011 | China | Clinical | Asia | O3:K6 | ST3 | PC | Wave-2 | Representative |
| VP11121 | VP11121 | 2011 | China | Clinical | Asia | O4:K68 | ST3 | PC | Wave-2 | Representative |
| VP11135 | VP11135 | 2011 | China | Clinical | Asia | O3:K6 | ST3 | PC | Wave-2 | Representative |
| VP11155 | VP11155 | 2011 | China | Clinical | Asia | O3:K6 | ST- | PC | Wave-2 | Representative |
| VP11156 | VP11156 | 2011 | China | Clinical | Asia | O3:K6 | ST3 | PC | Wave-2 | Representative |
| VP11160 | VP11160 | 2011 | China | Clinical | Asia | O3:K6 | ST3 | PC | Wave-2 | Representative |
| VP11167 | VP11167 | 2011 | China | Clinical | Asia | O3:K6 | ST3 | PC | Wave-2 | Representative |
| VP11168 | VP11168 | 2011 | China | Clinical | Asia | O3:K6 | ST3 | PC | Wave-2 | Representative |

| | | | | | | | | | | |
|---|---|---|---|---|---|---|---|---|---|---|
| VP11190 | VP11190 | 2011 | China | Clinical | Asia | O3:K6 | ST3 | PC | Wave-2 | Representative |
| VP11193 | VP11193 | 2011 | China | Clinical | Asia | O4:K68 | ST3 | PC | Wave-2 | Representative |
| VP11227 | VP11227 | 2011 | China | Clinical | Asia | O3:K6 | ST3 | PC | Wave-2 | Representative |
| VP11231 | VP11231 | 2011 | China | Clinical | Asia | O3:K6 | ST3 | PC | Wave-2 | Representative |
| VP11243 | VP11243 | 2011 | China | Clinical | Asia | O3:K6 | ST3 | PC | Wave-2 | Representative |
| VP11250 | VP11250 | 2011 | China | Clinical | Asia | O3:K6 | ST3 | PC | Wave-2 | Representative |
| VP11260 | VP11260 | 2011 | China | Clinical | Asia | O3:K6 | ST3 | PC | Wave-2 | Representative |
| VP11269 | VP11269 | 2011 | China | Clinical | Asia | O3:K6 | ST3 | PC | Wave-2 | Representative |
| VP11302 | VP11302 | 2011 | China | Clinical | Asia | O3:K6 | ST3 | PC | Wave-2 | Representative |
| VP12011 | VP12011 | 2012 | China | Clinical | Asia | O1:KUK3 | ST3 | PC | Wave-2 | Representative |
| VP12016 | VP12016 | 2012 | China | Clinical | Asia | O4:K68 | ST3 | PC | Wave-2 | Representative |
| VP12035 | VP12035 | 2012 | China | Clinical | Asia | O3:K6 | ST3 | PC | Wave-2 | Representative |
| VP12122 | VP12122 | 2012 | China | Clinical | Asia | O4:K68 | ST3 | PC | Wave-2 | Representative |
| VP12206 | VP12206 | 2012 | China | Clinical | Asia | O3:K6 | ST3 | PC | Wave-2 | Representative |
| VP13033 | VP13033 | 2013 | China | Clinical | Asia | O3:K6 | ST3 | PC | Wave-2 | Representative |
| VP13059 | VP13059 | 2013 | China | Clinical | Asia | O3:K6 | ST3 | PC | Wave-2 | Representative |
| VP13067 | VP13067 | 2013 | China | Clinical | Asia | O1:KUK3 | ST3 | PC | Wave-2 | Representative |
| VP13106 | VP13106 | 2013 | China | Clinical | Asia | O3:K6 | ST3 | PC | Wave-2 | Representative |
| VP14131 | VP14131 | 2014 | China | Clinical | Asia | O3:K6 | ST3 | PC | Wave-2 | Representative |
| VP15006 | VP15006 | 2015 | China | Clinical | Asia | O3:KUT7 | ST3 | PC | Wave-2 | Representative |
| VP15016 | VP15016 | 2015 | China | Clinical | Asia | O3:K6 | ST3 | PC | Wave-2 | Representative |
| VP15044 | VP15044 | 2015 | China | Clinical | Asia | O1:KUT6 | ST3 | PC | Wave-2 | Representative |
| VP15127 | VP15127 | 2015 | China | Clinical | Asia | O3:K6 | ST3 | PC | Wave-2 | Representative |
| VP15154 | VP15154 | 2015 | China | Clinical | Asia | O3:K6 | ST3 | PC | Wave-2 | Representative |
| VP15160 | VP15160 | 2015 | China | Clinical | Asia | O3:K6 | ST3 | PC | Wave-2 | Representative |
| VP15205 | VP15205 | 2015 | China | Clinical | Asia | O4:K68 | ST3 | PC | Wave-2 | Representative |
| VP16001 | VP16001 | 2016 | China | Clinical | Asia | O3:K6 | ST3 | PC | Wave-2 | Representative |
| VP16175 | VP16175 | 2016 | China | Clinical | Asia | O1:KUT6 | ST3 | PC | Wave-2 | Representative |
| VP16192 | VP16192 | 2016 | China | Clinical | Asia | O3:K6 | ST3 | PC | Wave-2 | Representative |
| VP17002 | VP17002 | 2017 | China | Clinical | Asia | O3:K6 | ST3 | PC | Wave-2 | Representative |
| VP17054 | VP17054 | 2017 | China | Clinical | Asia | O3:K6 | ST3 | PC | Wave-2 | Representative |
| VP18002 | VP18002 | 2018 | China | Clinical | Asia | O3:K6 | ST3 | PC | Wave-2 | Representative |

| | | | | | | | | | | |
|---|---|---|---|---|---|---|---|---|---|---|
| VP18007 | VP18007 | 2018 | China | Clinical | Asia | O3:K6 | ST3 | PC | Wave-2 | Representative |
| VP18012 | VP18012 | 2018 | China | Clinical | Asia | O3:K6 | ST3 | PC | Wave-2 | Representative |
| VP18019 | VP18019 | 2018 | China | Clinical | Asia | O3:K6 | ST3 | PC | Wave-2 | Representative |
| VP18071 | VP18071 | 2018 | China | Clinical | Asia | O3:K6 | ST3 | PC | Wave-2 | Representative |
| VP18215 | VP18215 | 2018 | China | Clinical | Asia | O3:K6 | ST3 | PC | Wave-2 | Representative |
| VP18245 | VP18245 | 2018 | China | Clinical | Asia | O1:KUT6 | ST- | PC | Wave-2 | Representative |
| VP19005 | VP19005 | 2019 | China | Clinical | Asia | O3:K6 | ST3 | PC | Wave-2 | Representative |
| VP19241 | VP19241 | 2019 | China | Clinical | Asia | O3:K6 | ST3 | PC | Wave-2 | Representative |
| VPCZ10 | VPCZ10 | 2014 | China | Clinical | Asia | O3:K6 | ST3 | PC | Wave-2 | Representative |
| VPCZ38 | VPCZ38 | 2013 | China | Clinical | Asia | O4:K68 | ST3 | PC | Wave-2 | Representative |
| VPCZ9 | VPCZ9 | 2013 | China | Clinical | Asia | O3:K6 | ST3 | PC | Wave-2 | Representative |
| SAMN03941062-GCF_001695945.1 | 44569 | 2004 | Spain | Clinical | Europe | O3:K6 | ST3 | PC | Wave-2 | Representative |
| Vp_UK_AF_S28 | AF_S28 | NA | UK | Clinical | Europe | O3:K6 | ST3 | PC | Wave-2 | Representative |
| Vp_UK_AJ_S32 | AJ_S32 | 2016 | UK(Kenya) | Clinical | Europe | O3:KUT7 | ST3 | PC | Wave-2 | Representative |
| Vp_UK_AL_S34 | AL_S34 | 2015 | UK | Clinical | Europe | O3:K6 | ST3 | PC | Wave-2 | Representative |
| Vp_UK_C_S3 | C_S3 | 2009 | UK(Vietnam) | Clinical | Europe | O3:K6 | ST3 | PC | Wave-2 | Representative |
| Vp_UK_D_S4 | D_S4 | 2013 | UK(Thailand) | Clinical | Europe | O3:K6 | ST3 | PC | Wave-2 | Representative |
| Vp_UK_S_S15 | S_S15 | 2011 | UK(Thailand) | Clinical | Europe | O3:K6 | ST3 | PC | Wave-2 | Representative |
| SAMN13893118-GCF_009936575.1 | CICESE-170 | 1998 | Mexico | Clinical | Latin America | O3:K6 | ST3 | PC | Wave-2 | Representative |
| SAMN13893119-GCF_009936605.1 | CICESE-186 | 1999 | Mexico | Clinical | Latin America | O3:K6 | ST3 | PC | Wave-2 | Representative |
| SAMN13893120-GCF_010692775.1 | CICESE-187 | 2000 | Mexico | Clinical | Latin America | O3:K65 | ST3 | PC | Wave-2 | Representative |
| SAMN02338979-GCF_000489835.1 | S131 | 2003 | NA | Clinical | NA | O3:K6 | ST3 | PC | Wave-2 | Representative |
| SAMN07338228-GCF_006368415.1 | S089 | NA | NA | Clinical | NA | O4:K68 | ST3 | PC | Wave-2 | Representative |
| SAMN09280045-GCA_015805655.1 | 218342 | 2015 | NA | Clinical | NA | O3:K6 | ST3 | PC | Wave-2 | Representative |
| SAMN09280046-GCA_015805805.1 | 218346 | 2015 | NA | Clinical | NA | O3:K6 | ST3 | PC | Wave-2 | Representative |
| SAMN09280052-GCA_015805795.1 | 245518 | 2016 | NA | Clinical | NA | O3:KUT7 | ST3 | PC | Wave-2 | Representative |
| SAMN09280073-GCA_015805875.1 | 539846 | 2017 | NA | Clinical | NA | O3:K6 | ST3 | PC | Wave-2 | Representative |
| SAMN13282666-SRR10441577 | 786602 | 2019 | NA | Clinical | NA | O3:K6 | ST3 | PC | Wave-2 | Representative |
| SAMN02368334-GCF_001728625.1 | BCW_3231 | 2007 | USA | Clinical | US_Canada | O4:K68 | ST3 | PC | Wave-2 | Representative |
| SAMN03452289-GCF_000975195.1 | 04-2551 | 2004 | Canada | Clinical | US_Canada | O3:K6 | ST3 | PC | Wave-2 | Representative |
| SAMN04377384-GCF_001609595.1 | A5Z273 | 2005 | Canada | Clinical | US_Canada | O3:K6 | ST- | PC | Wave-2 | Representative |
| SAMN04377388-GCF_001609525.1 | A5Z853 | 2005 | Canada | Clinical | US_Canada | O3:K6 | ST3 | PC | Wave-2 | Representative |

| | | | | | | | | | | |
|---|---|---|---|---|---|---|---|---|---|---|
| SAMN04422055-GCF_001608835.1 | F63267 | 2006 | Canada | Clinical | US_Canada | O3:K6 | ST3 | PC | Wave-2 | Representative |
| SAMN05220836-SRR3655233 | 4610 | 2007 | USA | Clinical | US_Canada | O4:K68 | ST3 | PC | Wave-2 | Representative |
| SAMN06009035-GCA_015818455.1 | PNUSAV000041 | NA | USA | Clinical | US_Canada | O4:K68 | ST3 | PC | Wave-2 | Representative |
| SAMN06076992-GCA_015818835.1 | CFSAN023541 | 1998 | USA | Clinical | US_Canada | O3:K6 | ST3 | PC | Wave-2 | Representative |
| SAMN08370015-GCA_015814625.1 | CFSAN026732 | 2014 | USA | Clinical | US_Canada | O3:K6 | ST3 | PC | Wave-2 | Representative |
| SAMN13746683-GCA_015776075.1 | PNUSAV001090 | NA | USA | Clinical | US_Canada | O3:K6 | ST3 | PC | Wave-2 | Representative |
| VP03023 | VP03023 | 2003 | China | Clinical | Asia | O1:K25 | ST- | PC | Wave-3 | Putative mutator |
| VP04101 | VP04101 | 2004 | China | Clinical | Asia | O1:K25 | ST- | PC | Wave-3 | Putative mutator |
| VP05239 | VP05239 | 2005 | China | Clinical | Asia | O3:K6 | ST- | PC | Wave-3 | Putative mutator |
| VP05242 | VP05242 | 2005 | China | Clinical | Asia | O3:K6 | ST3 | PC | Wave-3 | Putative mutator |
| VP05255 | VP05255 | 2005 | China | Clinical | Asia | O1:KUK62 | ST192 | PC | Wave-3 | Putative mutator |
| SAMN05935535-GCF_001895595.1 | 100151 | 2010 | China | Clinical | Asia | O1:K36 | ST3 | PC | Wave-3 | Putative outbreak cluster |
| SAMN06076989-GCA_015818875.1 | CFSAN023537 | 1997 | India | Clinical | Asia | O3:K6 | ST3 | PC | Wave-3 | Putative outbreak cluster |
| SAMN16782961-GCA_016818035.1 | VP22 | 2008 | China | Clinical | Asia | O3:K6 | ST3 | PC | Wave-3 | Putative outbreak cluster |
| SAMN16783007-GCA_016827355.1 | VP79 | 2008 | China | Clinical | Asia | O1:K25 | ST3 | PC | Wave-3 | Putative outbreak cluster |
| SAMN16783027-GCA_016826955.1 | VP102 | 2011 | China | NA | Asia | O1:K36 | ST3 | PC | Wave-3 | Putative outbreak cluster |
| SAMN16783028-GCA_016826975.1 | VP104 | 2010 | China | Clinical | Asia | O1:K36 | ST3 | PC | Wave-3 | Putative outbreak cluster |
| SAMN16783035-GCA_016826795.1 | VP111 | 2012 | China | Clinical | Asia | O1:K36 | ST3 | PC | Wave-3 | Putative outbreak cluster |
| SAMN16783044-GCA_016826635.1 | VP121 | 2009 | China | Clinical | Asia | O1:K36 | ST3 | PC | Wave-3 | Putative outbreak cluster |
| SAMN16783046-GCA_016826615.1 | VP123 | 2009 | China | Clinical | Asia | O1:K36 | ST3 | PC | Wave-3 | Putative outbreak cluster |
| SAMN16783047-GCA_016826585.1 | VP124 | 2009 | China | Clinical | Asia | O1:K36 | ST3 | PC | Wave-3 | Putative outbreak cluster |
| SAMN16783048-GCA_016826555.1 | VP125 | 2010 | China | Clinical | Asia | O1:K36 | ST3 | PC | Wave-3 | Putative outbreak cluster |
| SAMN16783049-GCA_016826515.1 | VP126 | 2010 | China | Clinical | Asia | O1:K36 | ST3 | PC | Wave-3 | Putative outbreak cluster |
| SAMN16783052-GCA_016826485.1 | VP129 | 2010 | China | Clinical | Asia | O1:K36 | ST3 | PC | Wave-3 | Putative outbreak cluster |
| SAMN16783055-GCA_016826405.1 | VP132 | 2008 | China | Clinical | Asia | O1:K25 | ST- | PC | Wave-3 | Putative outbreak cluster |
| SAMN16783056-GCA_016826395.1 | VP133 | 2008 | China | Clinical | Asia | O1:K25 | ST- | PC | Wave-3 | Putative outbreak cluster |
| SAMN16783060-GCA_016826315.1 | VP137 | 2008 | China | Clinical | Asia | O1:K25 | ST3 | PC | Wave-3 | Putative outbreak cluster |
| SAMN16783064-GCA_016826235.1 | VP141 | 2009 | China | Clinical | Asia | O1:K25 | ST3 | PC | Wave-3 | Putative outbreak cluster |
| SAMN16783069-GCA_016826135.1 | VP146 | 2012 | China | NA | Asia | O1:K25 | ST3 | PC | Wave-3 | Putative outbreak cluster |
| SAMN16783078-GCA_016825995.1 | VP155 | 2010 | China | Clinical | Asia | O1:K25 | ST3 | PC | Wave-3 | Putative outbreak cluster |
| SAMN16783100-GCA_016825515.1 | VP181 | 2007 | China | Clinical | Asia | O1:K25 | ST3 | PC | Wave-3 | Putative outbreak cluster |
| SAMN16783101-GCA_016825495.1 | VP182 | 2007 | China | Clinical | Asia | O1:K25 | ST3 | PC | Wave-3 | Putative outbreak cluster |

| | | | | | | | | | | |
|---|---|---|---|---|---|---|---|---|---|---|
| SAMN16783102-GCA_016825455.1 | VP183 | 2007 | China | Clinical | Asia | O1:K25 | ST3 | PC | Wave-3 | Putative outbreak cluster |
| SAMN16783105-GCA_016825415.1 | VP186 | 2011 | China | NA | Asia | O1:K25 | ST3 | PC | Wave-3 | Putative outbreak cluster |
| SAMN16783118-GCA_016825155.1 | VP209 | 2016 | China | NA | Asia | O1:K36 | ST3 | PC | Wave-3 | Putative outbreak cluster |
| SAMN16783119-GCA_016825135.1 | VP210 | 2008 | China | Clinical | Asia | O1:K36 | ST3 | PC | Wave-3 | Putative outbreak cluster |
| SAMN16783120-GCA_016825085.1 | VP211 | 2007 | China | Clinical | Asia | O1:K36 | ST3 | PC | Wave-3 | Putative outbreak cluster |
| SAMN16783121-GCA_016825075.1 | VP212 | 2009 | China | Clinical | Asia | O1:K36 | ST3 | PC | Wave-3 | Putative outbreak cluster |
| SAMN16783122-GCA_016825095.1 | VP213 | 2007 | China | Clinical | Asia | O1:K36 | ST3 | PC | Wave-3 | Putative outbreak cluster |
| SAMN16783123-GCA_016825055.1 | VP214 | 2009 | China | Clinical | Asia | O1:K36 | ST3 | PC | Wave-3 | Putative outbreak cluster |
| SAMN16783124-GCA_016825035.1 | VP215 | 2010 | China | Clinical | Asia | O1:K36 | ST3 | PC | Wave-3 | Putative outbreak cluster |
| SAMN16783125-GCA_016825015.1 | VP216 | 2010 | China | Clinical | Asia | O1:K36 | ST3 | PC | Wave-3 | Putative outbreak cluster |
| SAMN16783126-GCA_016824965.1 | VP217 | 2010 | China | NA | Asia | O1:K36 | ST3 | PC | Wave-3 | Putative outbreak cluster |
| SAMN16783127-GCA_016824975.1 | VP218 | 2010 | China | Clinical | Asia | O1:K36 | ST3 | PC | Wave-3 | Putative outbreak cluster |
| SAMN16783128-GCA_016824955.1 | VP219 | 2010 | China | Clinical | Asia | O1:K36 | ST3 | PC | Wave-3 | Putative outbreak cluster |
| SAMN16783129-GCA_016824875.1 | VP220 | 2010 | China | Clinical | Asia | O1:K36 | ST3 | PC | Wave-3 | Putative outbreak cluster |
| SAMN16783130-GCA_016824885.1 | VP221 | 2010 | China | Clinical | Asia | O1:K36 | ST3 | PC | Wave-3 | Putative outbreak cluster |
| SAMN16783131-GCA_016824925.1 | VP222 | 2010 | China | Clinical | Asia | O1:K36 | ST3 | PC | Wave-3 | Putative outbreak cluster |
| SAMN16783132-GCA_016824915.1 | VP223 | 2010 | China | Clinical | Asia | O1:K36 | ST3 | PC | Wave-3 | Putative outbreak cluster |
| SAMN16783133-GCA_016824855.1 | VP224 | 2010 | China | Clinical | Asia | O1:K36 | ST3 | PC | Wave-3 | Putative outbreak cluster |
| SAMN16783134-GCA_016824775.1 | VP225 | 2010 | China | Clinical | Asia | O1:K36 | ST3 | PC | Wave-3 | Putative outbreak cluster |
| SAMN16783135-GCA_016824785.1 | VP226 | 2010 | China | Clinical | Asia | O1:K36 | ST3 | PC | Wave-3 | Putative outbreak cluster |
| SAMN16783137-GCA_016824835.1 | VP228 | 2010 | China | Clinical | Asia | O1:K36 | ST3 | PC | Wave-3 | Putative outbreak cluster |
| SAMN16783246-GCA_016822595.1 | VP349 | 2016 | China | NA | Asia | O1:K25 | ST3 | PC | Wave-3 | Putative outbreak cluster |
| SAMN16783250-GCA_016822545.1 | VP354 | 2008 | China | NA | Asia | O3:K6 | ST3 | PC | Wave-3 | Putative outbreak cluster |
| SAMN16783282-GCA_016821915.1 | VP387 | 2009 | China | NA | Asia | O1:KUK62 | ST3 | PC | Wave-3 | Putative outbreak cluster |
| SAMN16783286-GCA_016821775.1 | VP391 | 2007 | China | NA | Asia | O3:K6 | ST3 | PC | Wave-3 | Putative outbreak cluster |
| SAMN16783294-GCA_016821635.1 | VP399 | 2016 | China | Clinical | Asia | O1:K36 | ST3 | PC | Wave-3 | Putative outbreak cluster |
| SAMN16783337-GCA_016820795.1 | VP443 | 2017 | China | NA | Asia | O1:K36 | ST3 | PC | Wave-3 | Putative outbreak cluster |
| SAMN16783338-GCA_016820755.1 | VP444 | 2017 | China | Clinical | Asia | O1:K25 | ST3 | PC | Wave-3 | Putative outbreak cluster |
| SAMN16783362-GCA_016818915.1 | VP103 | 2010 | China | Clinical | Asia | O1:K36 | ST- | PC | Wave-3 | Putative outbreak cluster |
| VP03014 | VP03014 | 2003 | China | Clinical | Asia | O1:K25 | ST3 | PC | Wave-3 | Putative outbreak cluster |
| VP03256 | VP03256 | 2005 | China | Clinical | Asia | O1:KUK62 | ST192 | PC | Wave-3 | Putative outbreak cluster |
| VP03261 | VP03261 | 2005 | China | Clinical | Asia | O1:KUK62 | ST192 | PC | Wave-3 | Putative outbreak cluster |

| | | | | | | | | | | |
|---|---|---|---|---|---|---|---|---|---|---|
| VP05260 | VP05260 | 2005 | China | Clinical | Asia | O1:KUK62 | ST192 | PC | Wave-3 | Putative outbreak cluster |
| VP06004 | VP06004 | 2006 | China | Clinical | Asia | O1:K25 | ST3 | PC | Wave-3 | Putative outbreak cluster |
| VP06005 | VP06005 | 2006 | China | Clinical | Asia | O1:K25 | ST3 | PC | Wave-3 | Putative outbreak cluster |
| VP06006 | VP06006 | 2006 | China | Clinical | Asia | O1:K25 | ST3 | PC | Wave-3 | Putative outbreak cluster |
| VP06007 | VP06007 | 2006 | China | Clinical | Asia | O1:K25 | ST3 | PC | Wave-3 | Putative outbreak cluster |
| VP06008 | VP06008 | 2006 | China | Clinical | Asia | O1:K25 | ST3 | PC | Wave-3 | Putative outbreak cluster |
| VP06020 | VP06020 | 2006 | China | Clinical | Asia | O1:KUK62 | ST192 | PC | Wave-3 | Putative outbreak cluster |
| VP06021 | VP06021 | 2006 | China | Clinical | Asia | O1:K25 | ST3 | PC | Wave-3 | Putative outbreak cluster |
| VP06022 | VP06022 | 2006 | China | Clinical | Asia | O1:KUK62 | ST192 | PC | Wave-3 | Putative outbreak cluster |
| VP06035 | VP06035 | 2006 | China | Clinical | Asia | O3:K6 | ST3 | PC | Wave-3 | Putative outbreak cluster |
| VP06036 | VP06036 | 2006 | China | Clinical | Asia | O3:K6 | ST3 | PC | Wave-3 | Putative outbreak cluster |
| VP06042 | VP06042 | 2006 | China | Clinical | Asia | O3:K6 | ST3 | PC | Wave-3 | Putative outbreak cluster |
| VP06043 | VP06043 | 2006 | China | Clinical | Asia | O3:K6 | ST3 | PC | Wave-3 | Putative outbreak cluster |
| VP06044 | VP06044 | 2006 | China | Clinical | Asia | O3:K6 | ST3 | PC | Wave-3 | Putative outbreak cluster |
| VP06053 | VP06053 | 2006 | China | Clinical | Asia | O3:K6 | ST3 | PC | Wave-3 | Putative outbreak cluster |
| VP06054 | VP06054 | 2006 | China | Clinical | Asia | O3:K6 | ST3 | PC | Wave-3 | Putative outbreak cluster |
| VP06056 | VP06056 | 2006 | China | Clinical | Asia | O3:K6 | ST3 | PC | Wave-3 | Putative outbreak cluster |
| VP06057 | VP06057 | 2006 | China | Clinical | Asia | O3:K6 | ST3 | PC | Wave-3 | Putative outbreak cluster |
| VP06058 | VP06058 | 2006 | China | Clinical | Asia | O3:K6 | ST3 | PC | Wave-3 | Putative outbreak cluster |
| VP06086 | VP06086 | 2006 | China | Clinical | Asia | O3:K6 | ST3 | PC | Wave-3 | Putative outbreak cluster |
| VP06092 | VP06092 | 2006 | China | Clinical | Asia | O3:K6 | ST3 | PC | Wave-3 | Putative outbreak cluster |
| VP06097 | VP06097 | 2006 | China | Clinical | Asia | O1:KUK62 | ST192 | PC | Wave-3 | Putative outbreak cluster |
| VP06099 | VP06099 | 2006 | China | Clinical | Asia | O1:KUK62 | ST192 | PC | Wave-3 | Putative outbreak cluster |
| VP06100 | VP06100 | 2006 | China | Clinical | Asia | O1:KUK62 | ST192 | PC | Wave-3 | Putative outbreak cluster |
| VP06102 | VP06102 | 2006 | China | Clinical | Asia | O1:KUK62 | ST192 | PC | Wave-3 | Putative outbreak cluster |
| VP06103 | VP06103 | 2006 | China | Clinical | Asia | O1:KUK62 | ST192 | PC | Wave-3 | Putative outbreak cluster |
| VP06106 | VP06106 | 2006 | China | Clinical | Asia | O1:KUK62 | ST192 | PC | Wave-3 | Putative outbreak cluster |
| VP06107 | VP06107 | 2006 | China | Clinical | Asia | O1:KUK62 | ST192 | PC | Wave-3 | Putative outbreak cluster |
| VP06133 | VP06133 | 2006 | China | Clinical | Asia | O3:K6 | ST3 | PC | Wave-3 | Putative outbreak cluster |
| VP06134 | VP06134 | 2006 | China | Clinical | Asia | O3:K6 | ST3 | PC | Wave-3 | Putative outbreak cluster |
| VP06135 | VP06135 | 2006 | China | Clinical | Asia | O3:K6 | ST3 | PC | Wave-3 | Putative outbreak cluster |
| VP06137 | VP06137 | 2006 | China | Clinical | Asia | O3:K6 | ST3 | PC | Wave-3 | Putative outbreak cluster |

| | | | | | | | | | | |
|---|---|---|---|---|---|---|---|---|---|---|
| VP06138 | VP06138 | 2006 | China | Clinical | Asia | O3:K6 | ST3 | PC | Wave-3 | Putative outbreak cluster |
| VP06140 | VP06140 | 2006 | China | Clinical | Asia | O3:K6 | ST3 | PC | Wave-3 | Putative outbreak cluster |
| VP06141 | VP06141 | 2006 | China | Clinical | Asia | O3:K6 | ST3 | PC | Wave-3 | Putative outbreak cluster |
| VP06142 | VP06142 | 2006 | China | Clinical | Asia | O3:K6 | ST3 | PC | Wave-3 | Putative outbreak cluster |
| VP06144 | VP06144 | 2006 | China | Clinical | Asia | O3:K6 | ST3 | PC | Wave-3 | Putative outbreak cluster |
| VP06145 | VP06145 | 2006 | China | Clinical | Asia | O1:K36 | ST3 | PC | Wave-3 | Putative outbreak cluster |
| VP06162 | VP06162 | 2006 | China | Clinical | Asia | O3:K6 | ST3 | PC | Wave-3 | Putative outbreak cluster |
| VP06163 | VP06163 | 2006 | China | Clinical | Asia | O3:K6 | ST3 | PC | Wave-3 | Putative outbreak cluster |
| VP06164 | VP06164 | 2006 | China | Clinical | Asia | O3:K6 | ST3 | PC | Wave-3 | Putative outbreak cluster |
| VP06165 | VP06165 | 2006 | China | Clinical | Asia | O3:K6 | ST3 | PC | Wave-3 | Putative outbreak cluster |
| VP06166 | VP06166 | 2006 | China | Clinical | Asia | O3:K6 | ST3 | PC | Wave-3 | Putative outbreak cluster |
| VP06167 | VP06167 | 2006 | China | Clinical | Asia | O3:K6 | ST3 | PC | Wave-3 | Putative outbreak cluster |
| VP06188 | VP06188 | 2006 | China | Env | Asia | O1:KUK62 | ST192 | PC | Wave-3 | Putative outbreak cluster |
| VP06198 | VP06198 | 2006 | China | Clinical | Asia | O3:K6 | ST3 | PC | Wave-3 | Putative outbreak cluster |
| VP07019 | VP07019 | 2007 | China | Clinical | Asia | O1:K25 | ST3 | PC | Wave-3 | Putative outbreak cluster |
| VP07022 | VP07022 | 2007 | China | Clinical | Asia | O1:K25 | ST3 | PC | Wave-3 | Putative outbreak cluster |
| VP07027 | VP07027 | 2007 | China | Clinical | Asia | O3:K6 | ST3 | PC | Wave-3 | Putative outbreak cluster |
| VP07030 | VP07030 | 2007 | China | Clinical | Asia | O3:K6 | ST3 | PC | Wave-3 | Putative outbreak cluster |
| VP07031 | VP07031 | 2007 | China | Clinical | Asia | O3:K6 | ST3 | PC | Wave-3 | Putative outbreak cluster |
| VP07032 | VP07032 | 2007 | China | Clinical | Asia | O3:K6 | ST3 | PC | Wave-3 | Putative outbreak cluster |
| VP07034 | VP07034 | 2007 | China | Clinical | Asia | O3:K6 | ST3 | PC | Wave-3 | Putative outbreak cluster |
| VP07037 | VP07037 | 2007 | China | Clinical | Asia | O3:K6 | ST3 | PC | Wave-3 | Putative outbreak cluster |
| VP07039 | VP07039 | 2007 | China | Clinical | Asia | O3:K6 | ST3 | PC | Wave-3 | Putative outbreak cluster |
| VP07045 | VP07045 | 2007 | China | Clinical | Asia | O3:K6 | ST3 | PC | Wave-3 | Putative outbreak cluster |
| VP07047 | VP07047 | 2007 | China | Clinical | Asia | O3:K6 | ST3 | PC | Wave-3 | Putative outbreak cluster |
| VP07053 | VP07053 | 2007 | China | Clinical | Asia | O3:K6 | ST3 | PC | Wave-3 | Putative outbreak cluster |
| VP07054 | VP07054 | 2007 | China | Clinical | Asia | O3:K6 | ST3 | PC | Wave-3 | Putative outbreak cluster |
| VP07057 | VP07057 | 2007 | China | Clinical | Asia | O3:K6 | ST3 | PC | Wave-3 | Putative outbreak cluster |
| VP07058 | VP07058 | 2007 | China | Clinical | Asia | O3:K6 | ST3 | PC | Wave-3 | Putative outbreak cluster |
| VP07061 | VP07061 | 2007 | China | Clinical | Asia | O3:K6 | ST3 | PC | Wave-3 | Putative outbreak cluster |
| VP07063 | VP07063 | 2007 | China | Clinical | Asia | O3:K6 | ST3 | PC | Wave-3 | Putative outbreak cluster |
| VP07067 | VP07067 | 2007 | China | Clinical | Asia | O3:K6 | ST3 | PC | Wave-3 | Putative outbreak cluster |

| | | | | | | | | | | |
|---|---|---|---|---|---|---|---|---|---|---|
| VP07068 | VP07068 | 2007 | China | Clinical | Asia | O3:K6 | ST3 | PC | Wave-3 | Putative outbreak cluster |
| VP07076 | VP07076 | 2007 | China | Clinical | Asia | O3:K6 | ST3 | PC | Wave-3 | Putative outbreak cluster |
| VP07077 | VP07077 | 2007 | China | Clinical | Asia | O3:K6 | ST3 | PC | Wave-3 | Putative outbreak cluster |
| VP07078 | VP07078 | 2007 | China | Clinical | Asia | O3:K6 | ST3 | PC | Wave-3 | Putative outbreak cluster |
| VP07079 | VP07079 | 2007 | China | Clinical | Asia | O3:K6 | ST3 | PC | Wave-3 | Putative outbreak cluster |
| VP07083 | VP07083 | 2007 | China | Clinical | Asia | O3:K6 | ST3 | PC | Wave-3 | Putative outbreak cluster |
| VP07084 | VP07084 | 2007 | China | Clinical | Asia | O3:K6 | ST3 | PC | Wave-3 | Putative outbreak cluster |
| VP07089 | VP07089 | 2007 | China | Clinical | Asia | O3:K6 | ST3 | PC | Wave-3 | Putative outbreak cluster |
| VP07091 | VP07091 | 2007 | China | Clinical | Asia | O3:K6 | ST3 | PC | Wave-3 | Putative outbreak cluster |
| VP07096 | VP07096 | 2007 | China | Clinical | Asia | O3:K6 | ST3 | PC | Wave-3 | Putative outbreak cluster |
| VP07102 | VP07102 | 2007 | China | Clinical | Asia | O3:K6 | ST3 | PC | Wave-3 | Putative outbreak cluster |
| VP07104 | VP07104 | 2007 | China | Clinical | Asia | O3:K6 | ST3 | PC | Wave-3 | Putative outbreak cluster |
| VP07107 | VP07107 | 2007 | China | Clinical | Asia | O3:K6 | ST3 | PC | Wave-3 | Putative outbreak cluster |
| VP07108 | VP07108 | 2007 | China | Clinical | Asia | O1:K36 | ST3 | PC | Wave-3 | Putative outbreak cluster |
| VP07109 | VP07109 | 2007 | China | Clinical | Asia | O3:K6 | ST3 | PC | Wave-3 | Putative outbreak cluster |
| VP07119 | VP07119 | 2007 | China | Clinical | Asia | O3:K6 | ST3 | PC | Wave-3 | Putative outbreak cluster |
| VP07121 | VP07121 | 2007 | China | Clinical | Asia | O3:K6 | ST3 | PC | Wave-3 | Putative outbreak cluster |
| VP07123 | VP07123 | 2007 | China | Clinical | Asia | O3:K6 | ST3 | PC | Wave-3 | Putative outbreak cluster |
| VP07124 | VP07124 | 2007 | China | Clinical | Asia | O3:K6 | ST3 | PC | Wave-3 | Putative outbreak cluster |
| VP07126 | VP07126 | 2007 | China | Clinical | Asia | O3:K6 | ST3 | PC | Wave-3 | Putative outbreak cluster |
| VP07127 | VP07127 | 2007 | China | Clinical | Asia | O3:K6 | ST3 | PC | Wave-3 | Putative outbreak cluster |
| VP07128 | VP07128 | 2007 | China | Clinical | Asia | O3:K6 | ST3 | PC | Wave-3 | Putative outbreak cluster |
| VP07129 | VP07129 | 2007 | China | Clinical | Asia | O3:K6 | ST3 | PC | Wave-3 | Putative outbreak cluster |
| VP07130 | VP07130 | 2007 | China | Clinical | Asia | O3:K6 | ST3 | PC | Wave-3 | Putative outbreak cluster |
| VP07131 | VP07131 | 2007 | China | Clinical | Asia | O3:K6 | ST3 | PC | Wave-3 | Putative outbreak cluster |
| VP07133 | VP07133 | 2007 | China | Clinical | Asia | O3:K6 | ST3 | PC | Wave-3 | Putative outbreak cluster |
| VP07134 | VP07134 | 2007 | China | Clinical | Asia | O3:K6 | ST3 | PC | Wave-3 | Putative outbreak cluster |
| VP07135 | VP07135 | 2007 | China | Clinical | Asia | O3:K6 | ST3 | PC | Wave-3 | Putative outbreak cluster |
| VP07136 | VP07136 | 2007 | China | Clinical | Asia | O3:K6 | ST3 | PC | Wave-3 | Putative outbreak cluster |
| VP07137 | VP07137 | 2007 | China | Clinical | Asia | O3:K6 | ST3 | PC | Wave-3 | Putative outbreak cluster |
| VP07139 | VP07139 | 2007 | China | Clinical | Asia | O3:K6 | ST3 | PC | Wave-3 | Putative outbreak cluster |
| VP07142 | VP07142 | 2007 | China | Clinical | Asia | O3:K6 | ST3 | PC | Wave-3 | Putative outbreak cluster |

| | | | | | | | | | | |
|---|---|---|---|---|---|---|---|---|---|---|
| VP07143 | VP07143 | 2007 | China | Clinical | Asia | O3:K6 | ST3 | PC | Wave-3 | Putative outbreak cluster |
| VP07145 | VP07145 | 2007 | China | Clinical | Asia | O3:K6 | ST3 | PC | Wave-3 | Putative outbreak cluster |
| VP07146 | VP07146 | 2007 | China | Clinical | Asia | O3:K6 | ST3 | PC | Wave-3 | Putative outbreak cluster |
| VP07149 | VP07149 | 2007 | China | Clinical | Asia | O3:K6 | ST3 | PC | Wave-3 | Putative outbreak cluster |
| VP07150 | VP07150 | 2007 | China | Clinical | Asia | O3:K6 | ST3 | PC | Wave-3 | Putative outbreak cluster |
| VP07151 | VP07151 | 2007 | China | Clinical | Asia | O3:K6 | ST3 | PC | Wave-3 | Putative outbreak cluster |
| VP07152 | VP07152 | 2007 | China | Clinical | Asia | O3:K6 | ST3 | PC | Wave-3 | Putative outbreak cluster |
| VP07153 | VP07153 | 2007 | China | Clinical | Asia | O3:K6 | ST3 | PC | Wave-3 | Putative outbreak cluster |
| VP07156 | VP07156 | 2007 | China | Clinical | Asia | O3:K6 | ST3 | PC | Wave-3 | Putative outbreak cluster |
| VP07157 | VP07157 | 2007 | China | Clinical | Asia | O3:K6 | ST3 | PC | Wave-3 | Putative outbreak cluster |
| VP07158 | VP07158 | 2007 | China | Clinical | Asia | O3:K6 | ST3 | PC | Wave-3 | Putative outbreak cluster |
| VP07160 | VP07160 | 2007 | China | Clinical | Asia | O3:K6 | ST3 | PC | Wave-3 | Putative outbreak cluster |
| VP07161 | VP07161 | 2007 | China | Clinical | Asia | O3:K6 | ST3 | PC | Wave-3 | Putative outbreak cluster |
| VP07162 | VP07162 | 2007 | China | Clinical | Asia | O3:K6 | ST3 | PC | Wave-3 | Putative outbreak cluster |
| VP07163 | VP07163 | 2007 | China | Clinical | Asia | O3:K6 | ST3 | PC | Wave-3 | Putative outbreak cluster |
| VP07164 | VP07164 | 2007 | China | Clinical | Asia | O3:K6 | ST3 | PC | Wave-3 | Putative outbreak cluster |
| VP07165 | VP07165 | 2007 | China | Clinical | Asia | O3:K6 | ST3 | PC | Wave-3 | Putative outbreak cluster |
| VP07167 | VP07167 | 2007 | China | Clinical | Asia | O3:K6 | ST3 | PC | Wave-3 | Putative outbreak cluster |
| VP07168 | VP07168 | 2007 | China | Clinical | Asia | O3:K6 | ST3 | PC | Wave-3 | Putative outbreak cluster |
| VP07169 | VP07169 | 2007 | China | Clinical | Asia | O3:K6 | ST3 | PC | Wave-3 | Putative outbreak cluster |
| VP07171 | VP07171 | 2007 | China | Clinical | Asia | O3:K6 | ST3 | PC | Wave-3 | Putative outbreak cluster |
| VP07173 | VP07173 | 2007 | China | Clinical | Asia | O3:K6 | ST3 | PC | Wave-3 | Putative outbreak cluster |
| VP07175 | VP07175 | 2007 | China | Clinical | Asia | O3:K6 | ST3 | PC | Wave-3 | Putative outbreak cluster |
| VP07177 | VP07177 | 2007 | China | Clinical | Asia | O3:K6 | ST3 | PC | Wave-3 | Putative outbreak cluster |
| VP07178 | VP07178 | 2007 | China | Clinical | Asia | O3:K6 | ST3 | PC | Wave-3 | Putative outbreak cluster |
| VP07181 | VP07181 | 2007 | China | Clinical | Asia | O1:K36 | ST3 | PC | Wave-3 | Putative outbreak cluster |
| VP07186 | VP07186 | 2007 | China | Clinical | Asia | O3:K6 | ST3 | PC | Wave-3 | Putative outbreak cluster |
| VP07188 | VP07188 | 2007 | China | Clinical | Asia | O3:K6 | ST3 | PC | Wave-3 | Putative outbreak cluster |
| VP07201 | VP07201 | 2007 | China | Clinical | Asia | O1:K25 | ST3 | PC | Wave-3 | Putative outbreak cluster |
| VP07203 | VP07203 | 2007 | China | Clinical | Asia | O3:K6 | ST3 | PC | Wave-3 | Putative outbreak cluster |
| VP07206 | VP07206 | 2007 | China | Clinical | Asia | O3:K6 | ST3 | PC | Wave-3 | Putative outbreak cluster |
| VP07207 | VP07207 | 2007 | China | Clinical | Asia | O3:K6 | ST3 | PC | Wave-3 | Putative outbreak cluster |

| | | | | | | | | | | |
|---|---|---|---|---|---|---|---|---|---|---|
| VP07209 | VP07209 | 2007 | China | Clinical | Asia | O3:K6 | ST3 | PC | Wave-3 | Putative outbreak cluster |
| VP07210 | VP07210 | 2007 | China | Clinical | Asia | O3:K6 | ST3 | PC | Wave-3 | Putative outbreak cluster |
| VP07211 | VP07211 | 2007 | China | Clinical | Asia | O3:K6 | ST3 | PC | Wave-3 | Putative outbreak cluster |
| VP07213 | VP07213 | 2007 | China | Clinical | Asia | O3:K6 | ST3 | PC | Wave-3 | Putative outbreak cluster |
| VP07214 | VP07214 | 2007 | China | Clinical | Asia | O3:K6 | ST3 | PC | Wave-3 | Putative outbreak cluster |
| VP07226 | VP07226 | 2007 | China | Clinical | Asia | O3:K6 | ST3 | PC | Wave-3 | Putative outbreak cluster |
| VP07236 | VP07236 | 2007 | China | Clinical | Asia | O3:K6 | ST3 | PC | Wave-3 | Putative outbreak cluster |
| VP07256 | VP07256 | 2007 | China | Clinical | Asia | O3:K6 | ST3 | PC | Wave-3 | Putative outbreak cluster |
| VP07257 | VP07257 | 2007 | China | Clinical | Asia | O3:K6 | ST3 | PC | Wave-3 | Putative outbreak cluster |
| VP07258 | VP07258 | 2007 | China | Clinical | Asia | O3:K6 | ST3 | PC | Wave-3 | Putative outbreak cluster |
| VP07259 | VP07259 | 2007 | China | Clinical | Asia | O3:K6 | ST3 | PC | Wave-3 | Putative outbreak cluster |
| VP07260 | VP07260 | 2007 | China | Clinical | Asia | O3:K6 | ST3 | PC | Wave-3 | Putative outbreak cluster |
| VP07261 | VP07261 | 2007 | China | Clinical | Asia | O3:K6 | ST3 | PC | Wave-3 | Putative outbreak cluster |
| VP07262 | VP07262 | 2007 | China | Clinical | Asia | O3:K6 | ST3 | PC | Wave-3 | Putative outbreak cluster |
| VP07264 | VP07264 | 2007 | China | Clinical | Asia | O3:K6 | ST3 | PC | Wave-3 | Putative outbreak cluster |
| VP07265 | VP07265 | 2007 | China | Clinical | Asia | O1:K36 | ST3 | PC | Wave-3 | Putative outbreak cluster |
| VP08059 | VP08059 | 2008 | China | Clinical | Asia | O3:K6 | ST3 | PC | Wave-3 | Putative outbreak cluster |
| VP08078 | VP08078 | 2008 | China | Clinical | Asia | O1:K25 | ST- | PC | Wave-3 | Putative outbreak cluster |
| VP08121 | VP08121 | 2008 | China | Clinical | Asia | O3:K6 | ST3 | PC | Wave-3 | Putative outbreak cluster |
| VP08125 | VP08125 | 2008 | China | Clinical | Asia | O3:K6 | ST3 | PC | Wave-3 | Putative outbreak cluster |
| VP08163 | VP08163 | 2008 | China | Clinical | Asia | O3:K6 | ST3 | PC | Wave-3 | Putative outbreak cluster |
| VP08165 | VP08165 | 2008 | China | Clinical | Asia | O3:K6 | ST3 | PC | Wave-3 | Putative outbreak cluster |
| VP08166 | VP08166 | 2008 | China | Clinical | Asia | O3:K6 | ST3 | PC | Wave-3 | Putative outbreak cluster |
| VP08170 | VP08170 | 2008 | China | Clinical | Asia | O3:K6 | ST3 | PC | Wave-3 | Putative outbreak cluster |
| VP08171 | VP08171 | 2008 | China | Clinical | Asia | O3:K6 | ST3 | PC | Wave-3 | Putative outbreak cluster |
| VP08201 | VP08201 | 2008 | China | Clinical | Asia | O3:K6 | ST3 | PC | Wave-3 | Putative outbreak cluster |
| VP08207 | VP08207 | 2008 | China | Clinical | Asia | O3:K6 | ST3 | PC | Wave-3 | Putative outbreak cluster |
| VP08208 | VP08208 | 2008 | China | Clinical | Asia | O3:K6 | ST3 | PC | Wave-3 | Putative outbreak cluster |
| VP08255 | VP08255 | 2008 | China | Clinical | Asia | O3:K6 | ST3 | PC | Wave-3 | Putative outbreak cluster |
| VP08257 | VP08257 | 2008 | China | Clinical | Asia | O3:K6 | ST3 | PC | Wave-3 | Putative outbreak cluster |
| VP08258 | VP08258 | 2008 | China | Clinical | Asia | O3:K6 | ST3 | PC | Wave-3 | Putative outbreak cluster |
| VP08259 | VP08259 | 2008 | China | Clinical | Asia | O3:K6 | ST3 | PC | Wave-3 | Putative outbreak cluster |

| | | | | | | | | | | |
|---|---|---|---|---|---|---|---|---|---|---|
| VP08260 | VP08260 | 2008 | China | Clinical | Asia | O3:K6 | ST3 | PC | Wave-3 | Putative outbreak cluster |
| VP08262 | VP08262 | 2008 | China | Clinical | Asia | O3:K6 | ST3 | PC | Wave-3 | Putative outbreak cluster |
| VP08263 | VP08263 | 2008 | China | Clinical | Asia | O3:K6 | ST3 | PC | Wave-3 | Putative outbreak cluster |
| VP08266 | VP08266 | 2008 | China | Clinical | Asia | O3:K6 | ST3 | PC | Wave-3 | Putative outbreak cluster |
| VP08267 | VP08267 | 2008 | China | Clinical | Asia | O3:K6 | ST3 | PC | Wave-3 | Putative outbreak cluster |
| VP08268 | VP08268 | 2008 | China | Clinical | Asia | O3:K6 | ST3 | PC | Wave-3 | Putative outbreak cluster |
| VP08270 | VP08270 | 2008 | China | Clinical | Asia | O3:K6 | ST3 | PC | Wave-3 | Putative outbreak cluster |
| VP08271 | VP08271 | 2008 | China | Clinical | Asia | O3:K6 | ST3 | PC | Wave-3 | Putative outbreak cluster |
| VP08273 | VP08273 | 2008 | China | Clinical | Asia | O3:K6 | ST3 | PC | Wave-3 | Putative outbreak cluster |
| VP08274 | VP08274 | 2008 | China | Clinical | Asia | O3:K6 | ST3 | PC | Wave-3 | Putative outbreak cluster |
| VP08276 | VP08276 | 2008 | China | Clinical | Asia | O3:K6 | ST3 | PC | Wave-3 | Putative outbreak cluster |
| VP08315 | VP08315 | 2008 | China | Clinical | Asia | O1:K25 | ST3 | PC | Wave-3 | Putative outbreak cluster |
| VP08338 | VP08338 | 2008 | China | Clinical | Asia | O3:K6 | ST3 | PC | Wave-3 | Putative outbreak cluster |
| VP08341 | VP08341 | 2008 | China | Clinical | Asia | O3:K6 | ST3 | PC | Wave-3 | Putative outbreak cluster |
| VP08347 | VP08347 | 2008 | China | Clinical | Asia | O3:K6 | ST3 | PC | Wave-3 | Putative outbreak cluster |
| VP08348 | VP08348 | 2008 | China | Clinical | Asia | O3:K6 | ST3 | PC | Wave-3 | Putative outbreak cluster |
| VP08349 | VP08349 | 2008 | China | Clinical | Asia | O3:K6 | ST3 | PC | Wave-3 | Putative outbreak cluster |
| VP08350 | VP08350 | 2008 | China | Clinical | Asia | O3:K6 | ST3 | PC | Wave-3 | Putative outbreak cluster |
| VP08351 | VP08351 | 2008 | China | Clinical | Asia | O3:K6 | ST3 | PC | Wave-3 | Putative outbreak cluster |
| VP08352 | VP08352 | 2008 | China | Clinical | Asia | O3:K6 | ST3 | PC | Wave-3 | Putative outbreak cluster |
| VP08357 | VP08357 | 2008 | China | Clinical | Asia | O3:K6 | ST3 | PC | Wave-3 | Putative outbreak cluster |
| VP08367 | VP08367 | 2008 | China | Clinical | Asia | O3:K6 | ST3 | PC | Wave-3 | Putative outbreak cluster |
| VP08368 | VP08368 | 2008 | China | Clinical | Asia | O3:K6 | ST3 | PC | Wave-3 | Putative outbreak cluster |
| VP09012 | VP09012 | 2009 | China | Clinical | Asia | O1:K36 | ST3 | PC | Wave-3 | Putative outbreak cluster |
| VP09017 | VP09017 | 2009 | China | Clinical | Asia | O1:K25 | ST3 | PC | Wave-3 | Putative outbreak cluster |
| VP09021 | VP09021 | 2009 | China | Clinical | Asia | O1:K36 | ST3 | PC | Wave-3 | Putative outbreak cluster |
| VP09039 | VP09039 | 2009 | China | Clinical | Asia | O3:K6 | ST3 | PC | Wave-3 | Putative outbreak cluster |
| VP09049 | VP09049 | 2009 | China | Clinical | Asia | O3:K6 | ST3 | PC | Wave-3 | Putative outbreak cluster |
| VP09087 | VP09087 | 2009 | China | Clinical | Asia | O3:K6 | ST3 | PC | Wave-3 | Putative outbreak cluster |
| VP09202 | VP09202 | 2009 | China | Clinical | Asia | O1:K36 | ST3 | PC | Wave-3 | Putative outbreak cluster |
| VP09210 | VP09210 | 2009 | China | Clinical | Asia | O3:K6 | ST3 | PC | Wave-3 | Putative outbreak cluster |
| VP09213 | VP09213 | 2009 | China | Clinical | Asia | O1:K25 | ST3 | PC | Wave-3 | Putative outbreak cluster |

| | | | | | | | | | | |
|---|---|---|---|---|---|---|---|---|---|---|
| VP09229 | VP09229 | 2009 | China | Clinical | Asia | O3:K6 | ST3 | PC | Wave-3 | Putative outbreak cluster |
| VP09319 | VP09319 | 2009 | China | Clinical | Asia | O3:K6 | ST3 | PC | Wave-3 | Putative outbreak cluster |
| VP09372 | VP09372 | 2009 | China | Clinical | Asia | O1:K25 | ST3 | PC | Wave-3 | Putative outbreak cluster |
| VP10081 | VP10081 | 2010 | China | Clinical | Asia | O1:K36 | ST3 | PC | Wave-3 | Putative outbreak cluster |
| VP10089 | VP10089 | 2010 | China | Clinical | Asia | O1:K36 | ST3 | PC | Wave-3 | Putative outbreak cluster |
| VP10107 | VP10107 | 2010 | China | Clinical | Asia | O1:K36 | ST3 | PC | Wave-3 | Putative outbreak cluster |
| VP10134 | VP10134 | 2010 | China | Clinical | Asia | O1:K36 | ST3 | PC | Wave-3 | Putative outbreak cluster |
| VP10137 | VP10137 | 2010 | China | Clinical | Asia | O1:K36 | ST3 | PC | Wave-3 | Putative outbreak cluster |
| VP10139 | VP10139 | 2010 | China | Clinical | Asia | O1:K36 | ST3 | PC | Wave-3 | Putative outbreak cluster |
| VP10140 | VP10140 | 2010 | China | Clinical | Asia | O1:K36 | ST3 | PC | Wave-3 | Putative outbreak cluster |
| VP10143 | VP10143 | 2010 | China | Clinical | Asia | O1:K36 | ST3 | PC | Wave-3 | Putative outbreak cluster |
| VP10146 | VP10146 | 2010 | China | Clinical | Asia | O1:K36 | ST3 | PC | Wave-3 | Putative outbreak cluster |
| VP10226 | VP10226 | 2010 | China | Clinical | Asia | O1:K36 | ST3 | PC | Wave-3 | Putative outbreak cluster |
| VP10233 | VP10233 | 2010 | China | Clinical | Asia | O1:K36 | ST3 | PC | Wave-3 | Putative outbreak cluster |
| VP10236 | VP10236 | 2010 | China | Clinical | Asia | O1:K36 | ST3 | PC | Wave-3 | Putative outbreak cluster |
| VP10238 | VP10238 | 2010 | China | Clinical | Asia | O1:K36 | ST3 | PC | Wave-3 | Putative outbreak cluster |
| VP10239 | VP10239 | 2010 | China | Clinical | Asia | O1:K36 | ST3 | PC | Wave-3 | Putative outbreak cluster |
| VP10304 | VP10304 | 2010 | China | Clinical | Asia | O1:K36 | ST3 | PC | Wave-3 | Putative outbreak cluster |
| VP10400 | VP10400 | 2010 | China | Clinical | Asia | O1:K36 | ST3 | PC | Wave-3 | Putative outbreak cluster |
| VP12131 | VP12131 | 2012 | China | Clinical | Asia | O1:K36 | ST3 | PC | Wave-3 | Putative outbreak cluster |
| VP17091 | VP17091 | 2017 | China | Clinical | Asia | O1:K36 | ST3 | PC | Wave-3 | Putative outbreak cluster |
| VP17093 | VP17093 | 2017 | China | Clinical | Asia | O1:K36 | ST3 | PC | Wave-3 | Putative outbreak cluster |
| VP17095 | VP17095 | 2017 | China | Clinical | Asia | O1:K36 | ST3 | PC | Wave-3 | Putative outbreak cluster |
| VP19019 | VP19019 | 2019 | China | Clinical | Asia | O1:K25 | ST3 | PC | Wave-3 | Putative outbreak cluster |
| VP19021 | VP19021 | 2019 | China | Clinical | Asia | O1:K25 | ST3 | PC | Wave-3 | Putative outbreak cluster |
| VP19037 | VP19037 | 2019 | China | Clinical | Asia | O1:K25 | ST3 | PC | Wave-3 | Putative outbreak cluster |
| VP19040 | VP19040 | 2019 | China | Clinical | Asia | O1:K25 | ST3 | PC | Wave-3 | Putative outbreak cluster |
| VP19042 | VP19042 | 2019 | China | Clinical | Asia | O1:K25 | ST3 | PC | Wave-3 | Putative outbreak cluster |
| VP19071 | VP19071 | 2019 | China | Clinical | Asia | O1:K25 | ST3 | PC | Wave-3 | Putative outbreak cluster |
| VP19252 | VP19252 | 2019 | China | Clinical | Asia | O1:K25 | ST3 | PC | Wave-3 | Putative outbreak cluster |
| SAMN02204307-GCF_000877485.2 | 863 | 2007 | USA | Env | US_Canada | O3:K6 | ST3 | PC | Wave-3 | Putative outbreak cluster |
| SAMN02597368-GCF_000519365.2 | 605 | 2006 | USA | Env | US_Canada | O3:K6 | ST3 | PC | Wave-3 | Putative outbreak cluster |

| | | | | | | | | | | |
|---|---|---|---|---|---|---|---|---|---|---|
| RIMD_2210633_chr12 | RIMD_2210633 | 1996 | Japan | Clinical | Asia | O3:K6 | ST3 | PC | Wave-3 | Representative |
| SAMEA8103027-GCA_905331785.1 | ERS5790105 | 2012 | China | Clinical | Asia | O1:K36 | ST3 | PC | Wave-3 | Representative |
| SAMN01924111-GCF_000521825.2 | IDH02640 | 2009 | India | Clinical | Asia | O1:K25 | ST3 | PC | Wave-3 | Representative |
| SAMN02190055-GCF_000454145.1 | PVCHO_VP-NY4 | 1997 | India | Clinical | Asia | O3:K6 | ST3 | PC | Wave-3 | Representative |
| SAMN02338919-GCF_000491015.1 | S062 | 1998 | Singapore | Clinical | Asia | O1:K25 | ST3 | PC | Wave-3 | Representative |
| SAMN02338922-GCF_000490955.1 | S065 | 1998 | China | Clinical | Asia | O1:K25 | ST3 | PC | Wave-3 | Representative |
| SAMN02338940-GCF_000490595.1 | S086 | 1999 | Thailand | Clinical | Asia | O1:K25 | ST3 | PC | Wave-3 | Representative |
| SAMN02338947-GCF_000490465.1 | S094 | 1996 | Thailand | Clinical | Asia | O3:K6 | ST3 | PC | Wave-3 | Representative |
| SAMN02338981-GCF_000489795.1 | S133 | 2005 | China | Clinical | Asia | O3:K6 | ST3 | PC | Wave-3 | Representative |
| SAMN02338983-GCF_000489755.1 | S135 | 2003 | China | Clinical | Asia | O3:K6 | ST3 | PC | Wave-3 | Representative |
| SAMN02338984-GCF_000489735.1 | S136 | 2004 | China | Clinical | Asia | O3:K6 | ST3 | PC | Wave-3 | Representative |
| SAMN02338985-GCF_000489715.1 | S137 | 2005 | China | Clinical | Asia | O3:K65 | ST3 | PC | Wave-3 | Representative |
| SAMN02471128-GCF_000500505.1 | AXNJ | 2007 | China | Clinical | Asia | O3:K6 | ST3 | PC | Wave-3 | Representative |
| SAMN04349747-GCF_001541615.1 | Gxfcg07_003_7 | 2007 | China | Clinical | Asia | O1:K36 | ST3 | PC | Wave-3 | Representative |
| SAMN05935524-GCF_001895665.1 | 100138 | 2008 | China | Clinical | Asia | O1:K25 | ST3 | PC | Wave-3 | Representative |
| SAMN05935534-GCF_001895425.1 | 100150 | 2008 | China | Clinical | Asia | O1:K25 | ST3 | PC | Wave-3 | Representative |
| SAMN06076987-GCA_015818995.1 | CFSAN023535 | 1996 | India | Clinical | Asia | O3:K6 | ST3 | PC | Wave-3 | Representative |
| SAMN06076988-GCA_015818975.1 | CFSAN023536 | 1996 | India | Clinical | Asia | O3:K6 | ST3 | PC | Wave-3 | Representative |
| SAMN06076991-GCA_015818795.1 | CFSAN023540 | 1998 | South_Korea | Clinical | Asia | O3:K6 | ST3 | PC | Wave-3 | Representative |
| SAMN06076997-GCA_015818815.1 | AO-24491 | 1999 | Bangladesh | Clinical | Asia | O1:K25 | ST3 | PC | Wave-3 | Representative |
| SAMN06076998-GCA_015818775.1 | CFSAN023547 | 1999 | Thailand | Clinical | Asia | O1:K25 | ST3 | PC | Wave-3 | Representative |
| SAMN06077015-GCA_015817835.1 | CFSAN023564 | 1998 | Japan | Clinical | Asia | O1:K25 | ST3 | PC | Wave-3 | Representative |
| SAMN13181012-SRR10413229 | NA | 2004 | Japan | NA | Asia | O3:K6 | ST3 | PC | Wave-3 | Representative |
| SAMN16783063-GCA_016826275.1 | VP140 | 2009 | China | Clinical | Asia | O1:K25 | ST3 | PC | Wave-3 | Representative |
| SAMN16783070-GCA_016826095.1 | VP147 | 2009 | China | Clinical | Asia | O1:K25 | ST3 | PC | Wave-3 | Representative |
| SAMN25962175-GCA_022352195.1 | 460 | 2017 | China | Clinical | Asia | O1:K25 | ST- | PC | Wave-3 | Representative |
| SAMN25962177-GCA_022352495.1 | vp41 | 2015 | China | Clinical | Asia | O3:K6 | ST3 | PC | Wave-3 | Representative |
| VP03011 | VP03011 | 2003 | China | Clinical | Asia | O1:K25 | ST3 | PC | Wave-3 | Representative |
| VP03013 | VP03013 | 2003 | China | Clinical | Asia | O1:K25 | ST3 | PC | Wave-3 | Representative |
| VP03015 | VP03015 | 2003 | China | Clinical | Asia | O1:K25 | ST3 | PC | Wave-3 | Representative |
| VP03021 | VP03021 | 2003 | China | Clinical | Asia | O1:K25 | ST3 | PC | Wave-3 | Representative |
| VP03022 | VP03022 | 2003 | China | Clinical | Asia | O1:K25 | ST3 | PC | Wave-3 | Representative |

| | | | | | | | | | | |
|---|---|---|---|---|---|---|---|---|---|---|
| VP03028 | VP03028 | 2003 | China | Clinical | Asia | O1:K25 | ST3 | PC | Wave-3 | Representative |
| VP03029 | VP03029 | 2003 | China | Clinical | Asia | O1:K25 | ST3 | PC | Wave-3 | Representative |
| VP03257 | VP03257 | 2005 | China | Clinical | Asia | O1:KUK62 | ST192 | PC | Wave-3 | Representative |
| VP03259 | VP03259 | 2005 | China | Clinical | Asia | O1:KUK62 | ST192 | PC | Wave-3 | Representative |
| VP04086 | VP04086 | 2004 | China | Clinical | Asia | O3:K6 | ST3 | PC | Wave-3 | Representative |
| VP04102 | VP04102 | 2004 | China | Clinical | Asia | O1:K25 | ST3 | PC | Wave-3 | Representative |
| VP04110 | VP04110 | 2004 | China | Clinical | Asia | O1:K25 | ST3 | PC | Wave-3 | Representative |
| VP04112 | VP04112 | 2004 | China | Clinical | Asia | O1:K25 | ST3 | PC | Wave-3 | Representative |
| VP04113 | VP04113 | 2004 | China | Clinical | Asia | O1:K25 | ST3 | PC | Wave-3 | Representative |
| VP04116 | VP04116 | 2004 | China | Clinical | Asia | O1:K25 | ST3 | PC | Wave-3 | Representative |
| VP04117 | VP04117 | 2004 | China | Clinical | Asia | O1:K25 | ST3 | PC | Wave-3 | Representative |
| VP04120 | VP04120 | 2004 | China | Clinical | Asia | O1:K25 | ST3 | PC | Wave-3 | Representative |
| VP04123 | VP04123 | 2004 | China | Clinical | Asia | O3:K6 | ST3 | PC | Wave-3 | Representative |
| VP04124 | VP04124 | 2004 | China | Clinical | Asia | O3:K6 | ST3 | PC | Wave-3 | Representative |
| VP04125 | VP04125 | 2004 | China | Clinical | Asia | O3:K6 | ST3 | PC | Wave-3 | Representative |
| VP04130 | VP04130 | 2004 | China | Clinical | Asia | O3:K6 | ST3 | PC | Wave-3 | Representative |
| VP04131 | VP04131 | 2004 | China | Clinical | Asia | O3:K6 | ST3 | PC | Wave-3 | Representative |
| VP05234 | VP05234 | 2005 | China | Clinical | Asia | O3:K6 | ST3 | PC | Wave-3 | Representative |
| VP05244 | VP05244 | 2005 | China | Clinical | Asia | O3:K6 | ST3 | PC | Wave-3 | Representative |
| VP05245 | VP05245 | 2005 | China | Clinical | Asia | O3:K6 | ST3 | PC | Wave-3 | Representative |
| VP05246 | VP05246 | 2005 | China | Clinical | Asia | O3:K6 | ST3 | PC | Wave-3 | Representative |
| VP05258 | VP05258 | 2005 | China | Clinical | Asia | O1:KUK62 | ST192 | PC | Wave-3 | Representative |
| VP05265 | VP05265 | 2005 | China | Clinical | Asia | O1:KUK62 | ST192 | PC | Wave-3 | Representative |
| VP06003 | VP06003 | 2006 | China | Clinical | Asia | O1:K25 | ST3 | PC | Wave-3 | Representative |
| VP06019 | VP06019 | 2006 | China | Clinical | Asia | O1:KUK62 | ST192 | PC | Wave-3 | Representative |
| VP06023 | VP06023 | 2006 | China | Clinical | Asia | O1:KUK62 | ST192 | PC | Wave-3 | Representative |
| VP06024 | VP06024 | 2006 | China | Clinical | Asia | O1:K25 | ST3 | PC | Wave-3 | Representative |
| VP06037 | VP06037 | 2006 | China | Clinical | Asia | O3:K6 | ST3 | PC | Wave-3 | Representative |
| VP06041 | VP06041 | 2006 | China | Clinical | Asia | O1:K25 | ST- | PC | Wave-3 | Representative |
| VP06047 | VP06047 | 2006 | China | Clinical | Asia | O3:K6 | ST3 | PC | Wave-3 | Representative |
| VP06068 | VP06068 | 2006 | China | Env | Asia | O3:K6 | ST3 | PC | Wave-3 | Representative |
| VP06069 | VP06069 | 2006 | China | Clinical | Asia | O1:K36 | ST3 | PC | Wave-3 | Representative |

| | | | | | | | | | | |
|---|---|---|---|---|---|---|---|---|---|---|
| VP06095 | VP06095 | 2006 | China | Clinical | Asia | O1:K25 | ST- | PC | Wave-3 | Representative |
| VP06098 | VP06098 | 2006 | China | Clinical | Asia | O1:KUK62 | ST192 | PC | Wave-3 | Representative |
| VP06101 | VP06101 | 2006 | China | Clinical | Asia | O1:KUK62 | ST192 | PC | Wave-3 | Representative |
| VP06104 | VP06104 | 2006 | China | Clinical | Asia | O1:KUK62 | ST192 | PC | Wave-3 | Representative |
| VP06105 | VP06105 | 2006 | China | Clinical | Asia | O1:KUK62 | ST192 | PC | Wave-3 | Representative |
| VP06108 | VP06108 | 2006 | China | Clinical | Asia | O1:KUK62 | ST192 | PC | Wave-3 | Representative |
| VP06136 | VP06136 | 2006 | China | Clinical | Asia | O3:K6 | ST3 | PC | Wave-3 | Representative |
| VP06143 | VP06143 | 2006 | China | Clinical | Asia | O3:K6 | ST3 | PC | Wave-3 | Representative |
| VP06146 | VP06146 | 2006 | China | Clinical | Asia | O1:K36 | ST3 | PC | Wave-3 | Representative |
| VP06189 | VP06189 | 2006 | China | Clinical | Asia | O1:KUK62 | ST192 | PC | Wave-3 | Representative |
| VP06196 | VP06196 | 2006 | China | Clinical | Asia | O3:K6 | ST3 | PC | Wave-3 | Representative |
| VP07016 | VP07016 | 2007 | China | Clinical | Asia | O1:K25 | ST3 | PC | Wave-3 | Representative |
| VP07017 | VP07017 | 2007 | China | Clinical | Asia | O1:K25 | ST3 | PC | Wave-3 | Representative |
| VP07036 | VP07036 | 2007 | China | Clinical | Asia | O1:K36 | ST3 | PC | Wave-3 | Representative |
| VP07062 | VP07062 | 2007 | China | Clinical | Asia | O3:K6 | ST3 | PC | Wave-3 | Representative |
| VP07069 | VP07069 | 2007 | China | Clinical | Asia | O3:K6 | ST3 | PC | Wave-3 | Representative |
| VP07073 | VP07073 | 2007 | China | Clinical | Asia | O3:K6 | ST3 | PC | Wave-3 | Representative |
| VP07090 | VP07090 | 2007 | China | Clinical | Asia | O3:K6 | ST3 | PC | Wave-3 | Representative |
| VP07103 | VP07103 | 2007 | China | Clinical | Asia | O3:K6 | ST3 | PC | Wave-3 | Representative |
| VP07125 | VP07125 | 2007 | China | Clinical | Asia | O3:K6 | ST3 | PC | Wave-3 | Representative |
| VP07141 | VP07141 | 2007 | China | Clinical | Asia | O1:K25 | ST3 | PC | Wave-3 | Representative |
| VP07154 | VP07154 | 2007 | China | Clinical | Asia | O1:K25 | ST3 | PC | Wave-3 | Representative |
| VP07159 | VP07159 | 2007 | China | Clinical | Asia | O3:K6 | ST3 | PC | Wave-3 | Representative |
| VP07197 | VP07197 | 2007 | China | Clinical | Asia | O3:K6 | ST3 | PC | Wave-3 | Representative |
| VP07205 | VP07205 | 2007 | China | Clinical | Asia | O3:K6 | ST3 | PC | Wave-3 | Representative |
| VP07212 | VP07212 | 2007 | China | Clinical | Asia | O3:K6 | ST3 | PC | Wave-3 | Representative |
| VP07229 | VP07229 | 2007 | China | Clinical | Asia | O1:K25 | ST3 | PC | Wave-3 | Representative |
| VP07233 | VP07233 | 2007 | China | Clinical | Asia | O1:K36 | ST3 | PC | Wave-3 | Representative |
| VP07263 | VP07263 | 2007 | China | Env | Asia | O3:K6 | ST3 | PC | Wave-3 | Representative |
| VP08025 | VP08025 | 2008 | China | Clinical | Asia | O1:K36 | ST3 | PC | Wave-3 | Representative |
| VP08076 | VP08076 | 2008 | China | Clinical | Asia | O1:K25 | ST- | PC | Wave-3 | Representative |
| VP08079 | VP08079 | 2008 | China | Clinical | Asia | O3:K6 | ST3 | PC | Wave-3 | Representative |

| | | | | | | | | | | |
|---|---|---|---|---|---|---|---|---|---|---|
| VP08083 | VP08083 | 2008 | China | Clinical | Asia | O3:K6 | ST3 | PC | Wave-3 | Representative |
| VP08106 | VP08106 | 2008 | China | Env | Asia | O1:K25 | ST3 | PC | Wave-3 | Representative |
| VP08115 | VP08115 | 2008 | China | Clinical | Asia | O1:K36 | ST3 | PC | Wave-3 | Representative |
| VP08126 | VP08126 | 2008 | China | Clinical | Asia | O3:K6 | ST3 | PC | Wave-3 | Representative |
| VP08212 | VP08212 | 2008 | China | Clinical | Asia | O1:K25 | ST3 | PC | Wave-3 | Representative |
| VP08242 | VP08242 | 2008 | China | Clinical | Asia | O1:K36 | ST3 | PC | Wave-3 | Representative |
| VP08243 | VP08243 | 2008 | China | Clinical | Asia | O1:K25 | ST3 | PC | Wave-3 | Representative |
| VP08275 | VP08275 | 2008 | China | Clinical | Asia | O3:K6 | ST3 | PC | Wave-3 | Representative |
| VP08322 | VP08322 | 2008 | China | Clinical | Asia | O3:K6 | ST3 | PC | Wave-3 | Representative |
| VP08327 | VP08327 | 2008 | China | Clinical | Asia | O1:K25 | ST3 | PC | Wave-3 | Representative |
| VP08374 | VP08374 | 2008 | China | Clinical | Asia | O3:K6 | ST3 | PC | Wave-3 | Representative |
| VP08378 | VP08378 | 2008 | China | Clinical | Asia | O3:K6 | ST3 | PC | Wave-3 | Representative |
| VP08379 | VP08379 | 2008 | China | Clinical | Asia | O1:K25 | ST3 | PC | Wave-3 | Representative |
| VP08401 | VP08401 | 2008 | China | Clinical | Asia | O1:K36 | ST3 | PC | Wave-3 | Representative |
| VP08402 | VP08402 | 2008 | China | Clinical | Asia | O3:K6 | ST3 | PC | Wave-3 | Representative |
| VP09047 | VP09047 | 2009 | China | Clinical | Asia | O3:K6 | ST3 | PC | Wave-3 | Representative |
| VP09050 | VP09050 | 2009 | China | Clinical | Asia | O1:KUK62 | ST3 | PC | Wave-3 | Representative |
| VP09054 | VP09054 | 2009 | China | Clinical | Asia | O1:K25 | ST3 | PC | Wave-3 | Representative |
| VP09056 | VP09056 | 2009 | China | Clinical | Asia | O3:K6 | ST3 | PC | Wave-3 | Representative |
| VP09057 | VP09057 | 2009 | China | Clinical | Asia | O3:K6 | ST3 | PC | Wave-3 | Representative |
| VP09113 | VP09113 | 2009 | China | Clinical | Asia | O1:K36 | ST3 | PC | Wave-3 | Representative |
| VP09201 | VP09201 | 2009 | China | Clinical | Asia | O1:K36 | ST3 | PC | Wave-3 | Representative |
| VP09212 | VP09212 | 2009 | China | Clinical | Asia | O1:K36 | ST3 | PC | Wave-3 | Representative |
| VP09215 | VP09215 | 2009 | China | Clinical | Asia | O1:K25 | ST3 | PC | Wave-3 | Representative |
| VP09218 | VP09218 | 2009 | China | Clinical | Asia | O1:K36 | ST3 | PC | Wave-3 | Representative |
| VP09427 | VP09427 | 2009 | China | Clinical | Asia | O1:K36 | ST3 | PC | Wave-3 | Representative |
| VP10084 | VP10084 | 2010 | China | Clinical | Asia | O1:K36 | ST3 | PC | Wave-3 | Representative |
| VP10103 | VP10103 | 2010 | China | Clinical | Asia | O1:K25 | ST3 | PC | Wave-3 | Representative |
| VP10108 | VP10108 | 2010 | China | Clinical | Asia | O1:K25 | ST3 | PC | Wave-3 | Representative |
| VP10115 | VP10115 | 2010 | China | Clinical | Asia | O3:K6 | ST3 | PC | Wave-3 | Representative |
| VP10183 | VP10183 | 2010 | China | Clinical | Asia | O1:K36 | ST3 | PC | Wave-3 | Representative |
| VP10192 | VP10192 | 2010 | China | Clinical | Asia | O1:K36 | ST3 | PC | Wave-3 | Representative |

| | | | | | | | | | | |
|---|---|---|---|---|---|---|---|---|---|---|
| VP10225 | VP10225 | 2010 | China | Clinical | Asia | O1:K36 | ST3 | PC | Wave-3 | Representative |
| VP10261 | VP10261 | 2010 | China | Clinical | Asia | O1:K25 | ST3 | PC | Wave-3 | Representative |
| VP10309 | VP10309 | 2010 | China | Clinical | Asia | O1:K36 | ST3 | PC | Wave-3 | Representative |
| VP10310 | VP10310 | 2010 | China | Clinical | Asia | O3:K6 | ST3 | PC | Wave-3 | Representative |
| VP10368 | VP10368 | 2010 | China | Clinical | Asia | O1:K36 | ST- | PC | Wave-3 | Representative |
| VP10413 | VP10413 | 2010 | China | Clinical | Asia | O3:K6 | ST3 | PC | Wave-3 | Representative |
| VP11138 | VP11138 | 2011 | China | Clinical | Asia | O1:K36 | ST3 | PC | Wave-3 | Representative |
| VP11180 | VP11180 | 2011 | China | Clinical | Asia | O1:K25 | ST3 | PC | Wave-3 | Representative |
| VP12017 | VP12017 | 2012 | China | Clinical | Asia | O1:K36 | ST3 | PC | Wave-3 | Representative |
| VP12077 | VP12077 | 2012 | China | Clinical | Asia | O1:K36 | ST3 | PC | Wave-3 | Representative |
| VP12102 | VP12102 | 2012 | China | Clinical | Asia | O1:K25 | ST3 | PC | Wave-3 | Representative |
| VP14056 | VP14056 | 2014 | China | Clinical | Asia | O1:K25 | ST3 | PC | Wave-3 | Representative |
| VP14084 | VP14084 | 2014 | China | Clinical | Asia | O1:K36 | ST3 | PC | Wave-3 | Representative |
| VP16025 | VP16025 | 2016 | China | Clinical | Asia | O1:K25 | ST3 | PC | Wave-3 | Representative |
| VP16107 | VP16107 | 2016 | China | Clinical | Asia | O1:K36 | ST3 | PC | Wave-3 | Representative |
| VP16200 | VP16200 | 2016 | China | Clinical | Asia | O1:K36 | ST3 | PC | Wave-3 | Representative |
| VP17092 | VP17092 | 2017 | China | Clinical | Asia | O1:K25 | ST3 | PC | Wave-3 | Representative |
| VP17094 | VP17094 | 2017 | China | Clinical | Asia | O1:K36 | ST3 | PC | Wave-3 | Representative |
| VP19041 | VP19041 | 2019 | China | Clinical | Asia | O1:K25 | ST3 | PC | Wave-3 | Representative |
| VP19244 | VP19244 | 2019 | China | Clinical | Asia | O1:K25 | ST3 | PC | Wave-3 | Representative |
| VPCZ97 | VPCZ97 | 2014 | China | Clinical | Asia | O1:K36 | ST3 | PC | Wave-3 | Representative |
| SAMN05591502-GCF_001720365.1 | Klin | 2010 | Sweden | Clinical | Europe | O1:K36 | ST3 | PC | Wave-3 | Representative |
| Vp_Peru_CFSAN062308_2010_Peru_C | CFSAN062308 | 2010 | Peru | Clinical | Latin America | O1:K36 | ST3 | PC | Wave-3 | Representative |
| SAMN07338231-GCF_006368595.1 | S175 | NA | NA | Clinical | NA | O3:K6 | ST3 | PC | Wave-3 | Representative |
| SAMN01923894-GCF_000524535.2 | 861 | 2006 | USA | NA | US_Canada | O3:K6 | ST3 | PC | Wave-3 | Representative |
| SAMN01924102-SRR748523 | 605 | 2006 | USA | NA | US_Canada | O3:K6 | ST3 | PC | Wave-3 | Representative |
| SAMN02190054-GCF_000454455.1 | 949 | 2006 | USA | NA | US_Canada | O3:K6 | ST3 | PC | Wave-3 | Representative |
| SAMN02204304-GCF_000877415.2 | VP551 | 2007 | USA | Env | US_Canada | O3:K6 | ST3 | PC | Wave-3 | Representative |
| SAMN02204308-GCF_000877475.2 | 930 | 2007 | USA | Env | US_Canada | O3:K6 | ST3 | PC | Wave-3 | Representative |
| SAMN02368298-GCF_001728155.1 | BCW_3191 | 2006 | USA | Clinical | US_Canada | O1:K25 | ST3 | PC | Wave-3 | Representative |
| SAMN02368299-GCF_001728175.1 | BCW_3192 | 2006 | USA | Clinical | US_Canada | O1:K25 | ST3 | PC | Wave-3 | Representative |
| SAMN03349598-GCF_000951795.1 | 04-2549 | 2004 | Canada | Clinical | US_Canada | O3:K6 | ST3 | PC | Wave-3 | Representative |

| | | | | | | | | | | |
|---|---|---|---|---|---|---|---|---|---|---|
| SAMN04327443-GCF_001595565.1 | A4EZ927 | 2004 | Canada | Clinical | US_Canada | O1:K25 | ST3 | PC | Wave-3 | Representative |
| SAMN04422047-GCF_001608785.1 | A3EZ136 | 2003 | Canada | Clinical | US_Canada | O3:K6 | ST3 | PC | Wave-3 | Representative |
| SAMN05194682-SRR3624770 | 920 | 2007 | USA | Env | US_Canada | O3:K6 | ST3 | PC | Wave-3 | Representative |
| SAMN05194683-SRR3624774 | 752 | 2007 | USA | Env | US_Canada | O3:K6 | ST3 | PC | Wave-3 | Representative |
| SAMN05195061-SRR3624775 | 783 | 2007 | USA | Env | US_Canada | O3:K6 | ST3 | PC | Wave-3 | Representative |
| SAMN05195070-SRR3628261 | 658 | 2007 | USA | Env | US_Canada | O3:K6 | ST3 | PC | Wave-3 | Representative |
| SAMN05195120-SRR3628271 | 765 | 2007 | USA | Env | US_Canada | O3:K6 | ST3 | PC | Wave-3 | Representative |
| SAMN13181013-SRR10413228 | NA | 2011 | USA | NA | US_Canada | O3:K6 | ST3 | PC | Wave-3 | Representative |
| SAMN13181014-SRR10413227 | NA | 2018 | USA | NA | US_Canada | O3:K6 | ST3 | PC | Wave-3 | Representative |
| SAMN13634001-GCA_015780275.1 | MDOH-04-5M732 | 2004 | USA | Clinical | US_Canada | O1:K25 | ST3 | PC | Wave-3 | Representative |
| SAMN07338146-GCF_006370915.1 | F3_2 | 2014 | China | Env | Asia | O3:K6 | ST3 | PC | Wave-4 | Putative mutator |
| SAMN07338147-GCF_006370895.1 | F3_3 | 2014 | China | Env | Asia | O3:K6 | ST3 | PC | Wave-4 | Putative mutator |
| SAMN07338148-GCF_006370855.1 | F3_4 | 2014 | China | Env | Asia | O3:K6 | ST3 | PC | Wave-4 | Putative mutator |
| SAMN07338149-GCF_006370835.1 | F3_5 | 2014 | China | Env | Asia | O3:K6 | ST3 | PC | Wave-4 | Putative mutator |
| SAMN07338211-GCF_006368725.1 | G1_10 | 2014 | China | Env | Asia | O3:K6 | ST3 | PC | Wave-4 | Putative mutator |
| SAMN07338214-GCF_006368495.1 | G1_4 | 2014 | China | Env | Asia | O3:K6 | ST3 | PC | Wave-4 | Putative mutator |
| SAMN07338215-GCF_006368335.1 | G1_5 | 2014 | China | Env | Asia | O3:K6 | ST3 | PC | Wave-4 | Putative mutator |
| SAMN07338220-GCF_006368645.1 | G2_1 | 2014 | China | Env | Asia | O3:K6 | ST3 | PC | Wave-4 | Putative mutator |
| VP04082 | VP04082 | 2004 | China | Clinical | Asia | O3:K6 | ST- | PC | Wave-4 | Putative mutator |
| VP05215 | VP05215 | 2005 | China | Clinical | Asia | O3:K6 | ST3 | PC | Wave-4 | Putative mutator |
| SAMN05935537-GCF_001895445.1 | 100153 | 2009 | China | Clinical | Asia | O3:K6 | ST3 | PC | Wave-4 | Putative outbreak cluster |
| SAMN06163187-GCF_001913745.1 | GIMxtfL65-2011.05 | 2011 | China | Clinical | Asia | O3:K6 | ST3 | PC | Wave-4 | Putative outbreak cluster |
| SAMN07338145-GCF_006370685.1 | F3_10 | 2014 | China | Env | Asia | O3:K6 | ST3 | PC | Wave-4 | Putative outbreak cluster |
| SAMN07338153-GCF_006370625.1 | F3_9 | 2014 | China | Env | Asia | O3:K6 | ST3 | PC | Wave-4 | Putative outbreak cluster |
| SAMN07338218-GCF_006368305.1 | G1_8 | 2014 | China | Env | Asia | O3:K6 | ST3 | PC | Wave-4 | Putative outbreak cluster |
| SAMN08667565-GCF_003056675.1 | VPF-1 | 2017 | Lebanon | Clinical | Asia | O3:K6 | ST3 | PC | Wave-4 | Putative outbreak cluster |
| SAMN08667566-GCF_003057295.1 | VPF-2 | 2017 | Lebanon | Clinical | Asia | O3:K6 | ST3 | PC | Wave-4 | Putative outbreak cluster |
| SAMN08667567-GCF_003056625.1 | VPF-3 | 2017 | Lebanon | Clinical | Asia | O3:K6 | ST3 | PC | Wave-4 | Putative outbreak cluster |
| SAMN08667654-GCF_003057315.1 | VPF-5 | 2017 | Lebanon | Clinical | Asia | O3:K6 | ST3 | PC | Wave-4 | Putative outbreak cluster |
| SAMN08667669-GCF_003056745.1 | VPF-7 | 2017 | Lebanon | Clinical | Asia | O3:K6 | ST3 | PC | Wave-4 | Putative outbreak cluster |
| SAMN13706853-SRR11823788 | SH-27 | 2015 | China | Env | Asia | O3:K6 | ST3 | PC | Wave-4 | Putative outbreak cluster |
| SAMN14411226-GCF_014922295.1 | L1 | 2015 | China | Clinical | Asia | O3:K6 | ST3 | PC | Wave-4 | Putative outbreak cluster |

| | | | | | | | | | | |
|---|---|---|---|---|---|---|---|---|---|---|
| SAMN14411227-GCF_014922265.1 | L2 | 2015 | China | Clinical | Asia | O3:K6 | ST3 | PC | Wave-4 | Putative outbreak cluster |
| SAMN14411228-GCF_014922235.1 | L3 | 2015 | China | Clinical | Asia | O3:K6 | ST3 | PC | Wave-4 | Putative outbreak cluster |
| SAMN14411231-GCF_014922205.1 | L7 | 2015 | China | Clinical | Asia | O3:K6 | ST3 | PC | Wave-4 | Putative outbreak cluster |
| SAMN14411232-GCF_014922185.1 | L8 | 2015 | China | Clinical | Asia | O3:K6 | ST3 | PC | Wave-4 | Putative outbreak cluster |
| SAMN14411235-GCF_014922075.1 | r75 | 2017 | China | Clinical | Asia | O3:K6 | ST3 | PC | Wave-4 | Putative outbreak cluster |
| SAMN15294466-GCA_019686035.1 | ICDC-VP01784 | 2016 | China | Clinical | Asia | O3:K6 | ST3 | PC | Wave-4 | Putative outbreak cluster |
| SAMN15294469-GCA_019685975.1 | ICDC-VP01791 | 2016 | China | Clinical | Asia | O3:K6 | ST3 | PC | Wave-4 | Putative outbreak cluster |
| SAMN15294470-GCA_019686015.1 | ICDC-VP01793 | 2017 | China | Clinical | Asia | O3:K6 | ST3 | PC | Wave-4 | Putative outbreak cluster |
| SAMN15294471-GCA_019685945.1 | ICDC-VP01794 | 2017 | China | Clinical | Asia | O3:K6 | ST3 | PC | Wave-4 | Putative outbreak cluster |
| SAMN15294473-GCA_019685915.1 | ICDC-VP01799 | 2017 | China | Clinical | Asia | O3:K6 | ST3 | PC | Wave-4 | Putative outbreak cluster |
| SAMN15294476-GCA_019685865.1 | ICDC-VP01802 | 2017 | China | Clinical | Asia | O3:K6 | ST3 | PC | Wave-4 | Putative outbreak cluster |
| SAMN15294485-GCA_019685635.1 | ICDC-VP01815 | 2018 | China | Clinical | Asia | O3:K6 | ST3 | PC | Wave-4 | Putative outbreak cluster |
| SAMN16782994-GCA_016817415.1 | VP63 | 2011 | China | Clinical | Asia | O3:K6 | ST3 | PC | Wave-4 | Putative outbreak cluster |
| SAMN16783013-GCA_016827235.1 | VP85 | 2009 | China | Clinical | Asia | O3:K6 | ST3 | PC | Wave-4 | Putative outbreak cluster |
| SAMN16783107-GCA_016825395.1 | VP188 | 2012 | China | NA | Asia | O3:K6 | ST3 | PC | Wave-4 | Putative outbreak cluster |
| SAMN16783116-GCA_016825235.1 | VP204 | 2017 | China | NA | Asia | O3:K6 | ST3 | PC | Wave-4 | Putative outbreak cluster |
| SAMN16783165-GCA_016824195.1 | VP264 | 2010 | China | Clinical | Asia | O3:K6 | ST3 | PC | Wave-4 | Putative outbreak cluster |
| SAMN16783173-GCA_016824045.1 | VP272 | 2017 | China | NA | Asia | O3:K6 | ST3 | PC | Wave-4 | Putative outbreak cluster |
| SAMN16783186-GCA_016823785.1 | VP285 | 2015 | China | NA | Asia | O3:K6 | ST3 | PC | Wave-4 | Putative outbreak cluster |
| SAMN16783222-GCA_016823055.1 | VP324 | 2015 | China | NA | Asia | O3:K6 | ST3 | PC | Wave-4 | Putative outbreak cluster |
| SAMN16783238-GCA_016822735.1 | VP341 | 2017 | China | NA | Asia | O3:K6 | ST3 | PC | Wave-4 | Putative outbreak cluster |
| SAMN16783239-GCA_016822755.1 | VP342 | 2017 | China | NA | Asia | O3:K6 | ST3 | PC | Wave-4 | Putative outbreak cluster |
| SAMN16783240-GCA_016822715.1 | VP343 | 2017 | China | NA | Asia | O3:K6 | ST3 | PC | Wave-4 | Putative outbreak cluster |
| SAMN16783259-GCA_016822335.1 | VP363 | 2015 | China | NA | Asia | O3:K6 | ST3 | PC | Wave-4 | Putative outbreak cluster |
| SAMN16783261-GCA_016822305.1 | VP365 | 2015 | China | Clinical | Asia | O3:K6 | ST3 | PC | Wave-4 | Putative outbreak cluster |
| SAMN16783268-GCA_016822155.1 | VP372 | 2010 | China | Clinical | Asia | O3:K6 | ST3 | PC | Wave-4 | Putative outbreak cluster |
| SAMN16783270-GCA_016822115.1 | VP374 | 2015 | China | NA | Asia | O3:K6 | ST3 | PC | Wave-4 | Putative outbreak cluster |
| SAMN16783279-GCA_016821935.1 | VP384 | 2013 | China | NA | Asia | O3:K6 | ST3 | PC | Wave-4 | Putative outbreak cluster |
| SAMN16783287-GCA_016821815.1 | VP392 | 2007 | China | NA | Asia | O3:K6 | ST3 | PC | Wave-4 | Putative outbreak cluster |
| SAMN16783293-GCA_016821665.1 | VP398 | 2016 | China | NA | Asia | O3:K6 | ST3 | PC | Wave-4 | Putative outbreak cluster |
| SAMN16783296-GCA_016821595.1 | VP401 | 2012 | China | NA | Asia | O3:K6 | ST3 | PC | Wave-4 | Putative outbreak cluster |
| SAMN16783297-GCA_016821575.1 | VP402 | 2012 | China | Clinical | Asia | O3:K6 | ST3 | PC | Wave-4 | Putative outbreak cluster |

| | | | | | | | | | | |
|---|---|---|---|---|---|---|---|---|---|---|
| SAMN16783298-GCA_016821555.1 | VP403 | 2012 | China | NA | Asia | O3:K6 | ST3 | PC | Wave-4 | Putative outbreak cluster |
| SAMN16783299-GCA_016821535.1 | VP404 | 2012 | China | Clinical | Asia | O3:K6 | ST3 | PC | Wave-4 | Putative outbreak cluster |
| SAMN16783334-GCA_016820835.1 | VP440 | 2017 | China | NA | Asia | O3:K6 | ST3 | PC | Wave-4 | Putative outbreak cluster |
| SAMN16783335-GCA_016820815.1 | VP441 | 2017 | China | NA | Asia | O3:K6 | ST3 | PC | Wave-4 | Putative outbreak cluster |
| SAMN16783358-GCA_016818995.1 | VP72 | 2017 | China | NA | Asia | O10:K60 | ST3 | PC | Wave-4 | Putative outbreak cluster |
| SAMN16783375-GCA_016818655.1 | VP208 | 2015 | China | NA | Asia | O10:K60 | ST3 | PC | Wave-4 | Putative outbreak cluster |
| SAMN16783377-GCA_016818595.1 | VP230 | 2012 | China | NA | Asia | O3:K6 | ST3 | PC | Wave-4 | Putative outbreak cluster |
| SAMN16783389-GCA_016818365.1 | VP407 | 2017 | China | Clinical | Asia | O10:K60 | ST3 | PC | Wave-4 | Putative outbreak cluster |
| SAMN20294544-GCA_019351375.1 | BH-Huang-1-01 | 2020 | China | Clinical | Asia | O10:K4 | ST3 | PC | Wave-4 | Putative outbreak cluster |
| SAMN25174351-GCA_023338245.1 | BH0084 | 2020 | China | Clinical | Asia | O10:K4 | ST3 | PC | Wave-4 | Putative outbreak cluster |
| SAMN25174513-GCA_023338275.1 | BH0085 | 2020 | China | Clinical | Asia | O10:K4 | ST3 | PC | Wave-4 | Putative outbreak cluster |
| SAMN25209733-GCA_023338305.1 | BH0086 | 2020 | China | Clinical | Asia | O10:K4 | ST3 | PC | Wave-4 | Putative outbreak cluster |
| SAMN25209740-GCA_023338315.1 | BH0087 | 2020 | China | Clinical | Asia | O10:K4 | ST3 | PC | Wave-4 | Putative outbreak cluster |
| SAMN25209765-GCA_023338345.1 | BH0088 | 2020 | China | Clinical | Asia | O10:K4 | ST3 | PC | Wave-4 | Putative outbreak cluster |
| SAMN25209771-GCA_023338355.1 | BH0089 | 2020 | China | Clinical | Asia | O10:K4 | ST3 | PC | Wave-4 | Putative outbreak cluster |
| SAMN25209820-GCA_023338385.1 | BH0090 | 2020 | China | Env | Asia | O10:K4 | ST3 | PC | Wave-4 | Putative outbreak cluster |
| SAMN25210965-GCA_023338395.1 | FCG0121 | 2020 | China | Clinical | Asia | O10:K4 | ST3 | PC | Wave-4 | Putative outbreak cluster |
| SAMN25211063-GCA_023338425.1 | FCG0124 | 2020 | China | Clinical | Asia | O10:K4 | ST3 | PC | Wave-4 | Putative outbreak cluster |
| SAMN25211114-GCA_023338435.1 | FCG0125 | 2020 | China | Clinical | Asia | O10:K4 | ST3 | PC | Wave-4 | Putative outbreak cluster |
| SAMN25212004-GCA_023338465.1 | FCG0126 | 2020 | China | Clinical | Asia | O10:K4 | ST3 | PC | Wave-4 | Putative outbreak cluster |
| SAMN25214717-GCA_023338475.1 | FCG0127 | 2020 | China | Clinical | Asia | O10:K4 | ST3 | PC | Wave-4 | Putative outbreak cluster |
| SAMN25220545-GCA_023338535.1 | FCG0128 | 2020 | China | Clinical | Asia | O10:K4 | ST3 | PC | Wave-4 | Putative outbreak cluster |
| SAMN25220608-GCA_023338555.1 | FCG0130 | 2020 | China | Clinical | Asia | O10:K4 | ST3 | PC | Wave-4 | Putative outbreak cluster |
| SAMN25221410-GCA_023338575.1 | FCG0131 | 2020 | China | Clinical | Asia | O10:K4 | ST3 | PC | Wave-4 | Putative outbreak cluster |
| SAMN25232486-GCA_023338545.1 | FCG0132 | 2020 | China | Clinical | Asia | O10:K4 | ST3 | PC | Wave-4 | Putative outbreak cluster |
| SAMN25232487-GCA_023338565.1 | NN0207 | 2020 | China | Clinical | Asia | O10:K4 | ST3 | PC | Wave-4 | Putative outbreak cluster |
| SAMN25232498-GCA_023338635.1 | QZ0184 | 2020 | China | Clinical | Asia | O10:K4 | ST3 | PC | Wave-4 | Putative outbreak cluster |
| SPVP140112 | SPVP140112 | 2014 | China | Env | Asia | O3:K6 | ST3 | PC | Wave-4 | Putative outbreak cluster |
| VP06012 | VP06012 | 2006 | China | Env | Asia | O3:K6 | ST3 | PC | Wave-4 | Putative outbreak cluster |
| VP06061 | VP06061 | 2006 | China | Clinical | Asia | O3:K6 | ST3 | PC | Wave-4 | Putative outbreak cluster |
| VP06062 | VP06062 | 2006 | China | Clinical | Asia | O3:K6 | ST3 | PC | Wave-4 | Putative outbreak cluster |
| VP06063 | VP06063 | 2006 | China | Clinical | Asia | O3:K6 | ST3 | PC | Wave-4 | Putative outbreak cluster |

| | | | | | | | | | | |
|---|---|---|---|---|---|---|---|---|---|---|
| VP06064 | VP06064 | 2006 | China | Clinical | Asia | O3:K6 | ST3 | PC | Wave-4 | Putative outbreak cluster |
| VP06067 | VP06067 | 2006 | China | Clinical | Asia | O3:K6 | ST3 | PC | Wave-4 | Putative outbreak cluster |
| VP06075 | VP06075 | 2006 | China | Env | Asia | O3:K6 | ST3 | PC | Wave-4 | Putative outbreak cluster |
| VP06076 | VP06076 | 2006 | China | Env | Asia | O3:K6 | ST3 | PC | Wave-4 | Putative outbreak cluster |
| VP06077 | VP06077 | 2006 | China | Env | Asia | O3:K6 | ST3 | PC | Wave-4 | Putative outbreak cluster |
| VP06078 | VP06078 | 2006 | China | Env | Asia | O3:K6 | ST3 | PC | Wave-4 | Putative outbreak cluster |
| VP06159 | VP06159 | 2006 | China | Clinical | Asia | O3:K6 | ST3 | PC | Wave-4 | Putative outbreak cluster |
| VP06160 | VP06160 | 2006 | China | Clinical | Asia | O3:K6 | ST3 | PC | Wave-4 | Putative outbreak cluster |
| VP07004 | VP07004 | 2007 | China | Clinical | Asia | O3:K6 | ST3 | PC | Wave-4 | Putative outbreak cluster |
| VP07005 | VP07005 | 2007 | China | Clinical | Asia | O3:K6 | ST3 | PC | Wave-4 | Putative outbreak cluster |
| VP07006 | VP07006 | 2007 | China | Clinical | Asia | O3:K6 | ST3 | PC | Wave-4 | Putative outbreak cluster |
| VP07007 | VP07007 | 2007 | China | Clinical | Asia | O3:K6 | ST3 | PC | Wave-4 | Putative outbreak cluster |
| VP07008 | VP07008 | 2007 | China | Clinical | Asia | O3:K6 | ST3 | PC | Wave-4 | Putative outbreak cluster |
| VP07009 | VP07009 | 2007 | China | Clinical | Asia | O3:K6 | ST3 | PC | Wave-4 | Putative outbreak cluster |
| VP07010 | VP07010 | 2007 | China | Clinical | Asia | O3:K6 | ST3 | PC | Wave-4 | Putative outbreak cluster |
| VP07023 | VP07023 | 2007 | China | Clinical | Asia | O3:K6 | ST3 | PC | Wave-4 | Putative outbreak cluster |
| VP07024 | VP07024 | 2007 | China | Clinical | Asia | O3:K6 | ST3 | PC | Wave-4 | Putative outbreak cluster |
| VP07025 | VP07025 | 2007 | China | Clinical | Asia | O3:K6 | ST3 | PC | Wave-4 | Putative outbreak cluster |
| VP07040 | VP07040 | 2007 | China | Clinical | Asia | O3:K6 | ST3 | PC | Wave-4 | Putative outbreak cluster |
| VP07042 | VP07042 | 2007 | China | Clinical | Asia | O3:K6 | ST3 | PC | Wave-4 | Putative outbreak cluster |
| VP07043 | VP07043 | 2007 | China | Clinical | Asia | O3:K6 | ST3 | PC | Wave-4 | Putative outbreak cluster |
| VP07044 | VP07044 | 2007 | China | Clinical | Asia | O3:K6 | ST3 | PC | Wave-4 | Putative outbreak cluster |
| VP07046 | VP07046 | 2007 | China | Clinical | Asia | O3:K6 | ST3 | PC | Wave-4 | Putative outbreak cluster |
| VP07048 | VP07048 | 2007 | China | Clinical | Asia | O3:K6 | ST3 | PC | Wave-4 | Putative outbreak cluster |
| VP07049 | VP07049 | 2007 | China | Clinical | Asia | O3:K6 | ST3 | PC | Wave-4 | Putative outbreak cluster |
| VP07064 | VP07064 | 2007 | China | Clinical | Asia | O3:K6 | ST3 | PC | Wave-4 | Putative outbreak cluster |
| VP07070 | VP07070 | 2007 | China | Clinical | Asia | O3:K6 | ST3 | PC | Wave-4 | Putative outbreak cluster |
| VP07075 | VP07075 | 2007 | China | Clinical | Asia | O3:K6 | ST3 | PC | Wave-4 | Putative outbreak cluster |
| VP07082 | VP07082 | 2007 | China | Clinical | Asia | O3:K6 | ST3 | PC | Wave-4 | Putative outbreak cluster |
| VP07085 | VP07085 | 2007 | China | Clinical | Asia | O3:K6 | ST3 | PC | Wave-4 | Putative outbreak cluster |
| VP07086 | VP07086 | 2007 | China | Clinical | Asia | O3:K6 | ST3 | PC | Wave-4 | Putative outbreak cluster |
| VP07087 | VP07087 | 2007 | China | Clinical | Asia | O3:K6 | ST3 | PC | Wave-4 | Putative outbreak cluster |

| | | | | | | | | | | |
|---|---|---|---|---|---|---|---|---|---|---|
| VP07092 | VP07092 | 2007 | China | Clinical | Asia | O3:K6 | ST3 | PC | Wave-4 | Putative outbreak cluster |
| VP07093 | VP07093 | 2007 | China | Clinical | Asia | O3:K6 | ST3 | PC | Wave-4 | Putative outbreak cluster |
| VP07098 | VP07098 | 2007 | China | Clinical | Asia | O3:K6 | ST3 | PC | Wave-4 | Putative outbreak cluster |
| VP07100 | VP07100 | 2007 | China | Clinical | Asia | O3:K6 | ST3 | PC | Wave-4 | Putative outbreak cluster |
| VP07105 | VP07105 | 2007 | China | Clinical | Asia | O3:K6 | ST3 | PC | Wave-4 | Putative outbreak cluster |
| VP07120 | VP07120 | 2007 | China | Clinical | Asia | O3:K6 | ST3 | PC | Wave-4 | Putative outbreak cluster |
| VP07138 | VP07138 | 2007 | China | Clinical | Asia | O3:K6 | ST3 | PC | Wave-4 | Putative outbreak cluster |
| VP07166 | VP07166 | 2007 | China | Clinical | Asia | O3:K6 | ST3 | PC | Wave-4 | Putative outbreak cluster |
| VP07170 | VP07170 | 2007 | China | Clinical | Asia | O3:K6 | ST- | PC | Wave-4 | Putative outbreak cluster |
| VP07215 | VP07215 | 2007 | China | Clinical | Asia | O3:K6 | ST3 | PC | Wave-4 | Putative outbreak cluster |
| VP07217 | VP07217 | 2007 | China | Clinical | Asia | O3:K6 | ST3 | PC | Wave-4 | Putative outbreak cluster |
| VP07218 | VP07218 | 2007 | China | Clinical | Asia | O3:K6 | ST3 | PC | Wave-4 | Putative outbreak cluster |
| VP07219 | VP07219 | 2007 | China | Clinical | Asia | O3:K6 | ST3 | PC | Wave-4 | Putative outbreak cluster |
| VP07235 | VP07235 | 2007 | China | Clinical | Asia | O3:K6 | ST3 | PC | Wave-4 | Putative outbreak cluster |
| VP07249 | VP07249 | 2007 | China | Clinical | Asia | O3:K6 | ST3 | PC | Wave-4 | Putative outbreak cluster |
| VP07266 | VP07266 | 2007 | China | Clinical | Asia | O3:K6 | ST3 | PC | Wave-4 | Putative outbreak cluster |
| VP07267 | VP07267 | 2007 | China | Clinical | Asia | O3:K6 | ST3 | PC | Wave-4 | Putative outbreak cluster |
| VP07270 | VP07270 | 2007 | China | Clinical | Asia | O3:K6 | ST3 | PC | Wave-4 | Putative outbreak cluster |
| VP07271 | VP07271 | 2007 | China | Clinical | Asia | O3:K6 | ST3 | PC | Wave-4 | Putative outbreak cluster |
| VP07272 | VP07272 | 2007 | China | Clinical | Asia | O3:K6 | ST3 | PC | Wave-4 | Putative outbreak cluster |
| VP07274 | VP07274 | 2007 | China | Clinical | Asia | O3:K6 | ST3 | PC | Wave-4 | Putative outbreak cluster |
| VP08019 | VP08019 | 2008 | China | Clinical | Asia | O3:K6 | ST3 | PC | Wave-4 | Putative outbreak cluster |
| VP08020 | VP08020 | 2008 | China | Clinical | Asia | O3:K6 | ST3 | PC | Wave-4 | Putative outbreak cluster |
| VP08053 | VP08053 | 2008 | China | Clinical | Asia | O3:K6 | ST3 | PC | Wave-4 | Putative outbreak cluster |
| VP08056 | VP08056 | 2008 | China | Clinical | Asia | O3:K6 | ST3 | PC | Wave-4 | Putative outbreak cluster |
| VP08081 | VP08081 | 2008 | China | Clinical | Asia | O3:K6 | ST3 | PC | Wave-4 | Putative outbreak cluster |
| VP08123 | VP08123 | 2008 | China | Clinical | Asia | O3:K6 | ST3 | PC | Wave-4 | Putative outbreak cluster |
| VP08127 | VP08127 | 2008 | China | Clinical | Asia | O3:K6 | ST3 | PC | Wave-4 | Putative outbreak cluster |
| VP08128 | VP08128 | 2008 | China | Clinical | Asia | O3:K6 | ST3 | PC | Wave-4 | Putative outbreak cluster |
| VP08129 | VP08129 | 2008 | China | Clinical | Asia | O3:K6 | ST3 | PC | Wave-4 | Putative outbreak cluster |
| VP08130 | VP08130 | 2008 | China | Clinical | Asia | O3:K6 | ST3 | PC | Wave-4 | Putative outbreak cluster |
| VP08131 | VP08131 | 2008 | China | Clinical | Asia | O3:K6 | ST3 | PC | Wave-4 | Putative outbreak cluster |

| | | | | | | | | | | |
|---|---|---|---|---|---|---|---|---|---|---|
| VP08132 | VP08132 | 2008 | China | Clinical | Asia | O3:K6 | ST3 | PC | Wave-4 | Putative outbreak cluster |
| VP08133 | VP08133 | 2008 | China | Clinical | Asia | O3:K6 | ST3 | PC | Wave-4 | Putative outbreak cluster |
| VP08134 | VP08134 | 2008 | China | Clinical | Asia | O3:K6 | ST3 | PC | Wave-4 | Putative outbreak cluster |
| VP08135 | VP08135 | 2008 | China | Clinical | Asia | O3:K6 | ST3 | PC | Wave-4 | Putative outbreak cluster |
| VP08136 | VP08136 | 2008 | China | Clinical | Asia | O3:K6 | ST3 | PC | Wave-4 | Putative outbreak cluster |
| VP08145 | VP08145 | 2008 | China | Clinical | Asia | O3:K6 | ST3 | PC | Wave-4 | Putative outbreak cluster |
| VP08169 | VP08169 | 2008 | China | Clinical | Asia | O3:K6 | ST3 | PC | Wave-4 | Putative outbreak cluster |
| VP08172 | VP08172 | 2008 | China | Clinical | Asia | O3:K6 | ST3 | PC | Wave-4 | Putative outbreak cluster |
| VP08184 | VP08184 | 2008 | China | Clinical | Asia | O3:K6 | ST3 | PC | Wave-4 | Putative outbreak cluster |
| VP08187 | VP08187 | 2008 | China | Clinical | Asia | O3:K6 | ST3 | PC | Wave-4 | Putative outbreak cluster |
| VP08193 | VP08193 | 2008 | China | Clinical | Asia | O3:K6 | ST3 | PC | Wave-4 | Putative outbreak cluster |
| VP08199 | VP08199 | 2008 | China | Clinical | Asia | O3:K6 | ST3 | PC | Wave-4 | Putative outbreak cluster |
| VP08316 | VP08316 | 2008 | China | Clinical | Asia | O3:K6 | ST3 | PC | Wave-4 | Putative outbreak cluster |
| VP08318 | VP08318 | 2008 | China | Clinical | Asia | O3:K6 | ST3 | PC | Wave-4 | Putative outbreak cluster |
| VP08323 | VP08323 | 2008 | China | Clinical | Asia | O3:K6 | ST3 | PC | Wave-4 | Putative outbreak cluster |
| VP08329 | VP08329 | 2008 | China | Clinical | Asia | O3:K6 | ST3 | PC | Wave-4 | Putative outbreak cluster |
| VP08339 | VP08339 | 2008 | China | Clinical | Asia | O3:K6 | ST3 | PC | Wave-4 | Putative outbreak cluster |
| VP09022 | VP09022 | 2009 | China | Clinical | Asia | O3:K6 | ST3 | PC | Wave-4 | Putative outbreak cluster |
| VP09029 | VP09029 | 2009 | China | Clinical | Asia | O3:K6 | ST3 | PC | Wave-4 | Putative outbreak cluster |
| VP09031 | VP09031 | 2009 | China | Clinical | Asia | O3:K6 | ST3 | PC | Wave-4 | Putative outbreak cluster |
| VP09037 | VP09037 | 2009 | China | Clinical | Asia | O3:K6 | ST3 | PC | Wave-4 | Putative outbreak cluster |
| VP09040 | VP09040 | 2009 | China | Clinical | Asia | O3:K6 | ST3 | PC | Wave-4 | Putative outbreak cluster |
| VP09072 | VP09072 | 2009 | China | Clinical | Asia | O3:K6 | ST3 | PC | Wave-4 | Putative outbreak cluster |
| VP09079 | VP09079 | 2009 | China | Clinical | Asia | O3:K6 | ST3 | PC | Wave-4 | Putative outbreak cluster |
| VP09089 | VP09089 | 2009 | China | Clinical | Asia | O3:K6 | ST3 | PC | Wave-4 | Putative outbreak cluster |
| VP09091 | VP09091 | 2009 | China | Clinical | Asia | O3:K6 | ST3 | PC | Wave-4 | Putative outbreak cluster |
| VP09093 | VP09093 | 2009 | China | Clinical | Asia | O3:K6 | ST3 | PC | Wave-4 | Putative outbreak cluster |
| VP09101 | VP09101 | 2009 | China | Clinical | Asia | O3:K6 | ST3 | PC | Wave-4 | Putative outbreak cluster |
| VP09104 | VP09104 | 2009 | China | Clinical | Asia | O3:K6 | ST3 | PC | Wave-4 | Putative outbreak cluster |
| VP09112 | VP09112 | 2009 | China | Clinical | Asia | O3:K6 | ST3 | PC | Wave-4 | Putative outbreak cluster |
| VP09125 | VP09125 | 2009 | China | Clinical | Asia | O3:K6 | ST3 | PC | Wave-4 | Putative outbreak cluster |
| VP09139 | VP09139 | 2009 | China | Clinical | Asia | O3:K6 | ST3 | PC | Wave-4 | Putative outbreak cluster |

| | | | | | | | | | | |
|---|---|---|---|---|---|---|---|---|---|---|
| VP09141 | VP09141 | 2009 | China | Clinical | Asia | O3:K6 | ST3 | PC | Wave-4 | Putative outbreak cluster |
| VP09149 | VP09149 | 2009 | China | Clinical | Asia | O3:K6 | ST3 | PC | Wave-4 | Putative outbreak cluster |
| VP09152 | VP09152 | 2009 | China | Clinical | Asia | O3:K6 | ST3 | PC | Wave-4 | Putative outbreak cluster |
| VP09154 | VP09154 | 2009 | China | Clinical | Asia | O3:K6 | ST3 | PC | Wave-4 | Putative outbreak cluster |
| VP09159 | VP09159 | 2009 | China | Clinical | Asia | O3:K6 | ST3 | PC | Wave-4 | Putative outbreak cluster |
| VP09161 | VP09161 | 2009 | China | Clinical | Asia | O3:K6 | ST3 | PC | Wave-4 | Putative outbreak cluster |
| VP09163 | VP09163 | 2009 | China | Clinical | Asia | O3:K6 | ST3 | PC | Wave-4 | Putative outbreak cluster |
| VP09166 | VP09166 | 2009 | China | Clinical | Asia | O3:K6 | ST3 | PC | Wave-4 | Putative outbreak cluster |
| VP09168 | VP09168 | 2009 | China | Clinical | Asia | O3:K6 | ST3 | PC | Wave-4 | Putative outbreak cluster |
| VP09169 | VP09169 | 2009 | China | Clinical | Asia | O3:K6 | ST3 | PC | Wave-4 | Putative outbreak cluster |
| VP09171 | VP09171 | 2009 | China | Clinical | Asia | O3:K6 | ST3 | PC | Wave-4 | Putative outbreak cluster |
| VP09173 | VP09173 | 2009 | China | Clinical | Asia | O3:K6 | ST3 | PC | Wave-4 | Putative outbreak cluster |
| VP09174 | VP09174 | 2009 | China | Clinical | Asia | O3:K6 | ST3 | PC | Wave-4 | Putative outbreak cluster |
| VP09175 | VP09175 | 2009 | China | Clinical | Asia | O3:K6 | ST3 | PC | Wave-4 | Putative outbreak cluster |
| VP09184 | VP09184 | 2009 | China | Clinical | Asia | O3:K6 | ST3 | PC | Wave-4 | Putative outbreak cluster |
| VP09188 | VP09188 | 2009 | China | Clinical | Asia | O3:K6 | ST3 | PC | Wave-4 | Putative outbreak cluster |
| VP09192 | VP09192 | 2009 | China | Clinical | Asia | O3:K6 | ST3 | PC | Wave-4 | Putative outbreak cluster |
| VP09197 | VP09197 | 2009 | China | Clinical | Asia | O3:K6 | ST3 | PC | Wave-4 | Putative outbreak cluster |
| VP09208 | VP09208 | 2009 | China | Clinical | Asia | O3:K6 | ST3 | PC | Wave-4 | Putative outbreak cluster |
| VP09211 | VP09211 | 2009 | China | Clinical | Asia | O3:K6 | ST3 | PC | Wave-4 | Putative outbreak cluster |
| VP09214 | VP09214 | 2009 | China | Clinical | Asia | O3:K6 | ST3 | PC | Wave-4 | Putative outbreak cluster |
| VP09220 | VP09220 | 2009 | China | Clinical | Asia | O3:K6 | ST3 | PC | Wave-4 | Putative outbreak cluster |
| VP09221-01 | VP09221-01 | 2009 | China | Clinical | Asia | O3:K6 | ST3 | PC | Wave-4 | Putative outbreak cluster |
| VP09233 | VP09233 | 2009 | China | Clinical | Asia | O3:K6 | ST3 | PC | Wave-4 | Putative outbreak cluster |
| VP09242 | VP09242 | 2009 | China | Clinical | Asia | O3:K6 | ST3 | PC | Wave-4 | Putative outbreak cluster |
| VP09251 | VP09251 | 2009 | China | Clinical | Asia | O3:K6 | ST3 | PC | Wave-4 | Putative outbreak cluster |
| VP09255-2 | VP09255-2 | 2009 | China | Clinical | Asia | O3:K6 | ST3 | PC | Wave-4 | Putative outbreak cluster |
| VP09266 | VP09266 | 2009 | China | Clinical | Asia | O3:K6 | ST3 | PC | Wave-4 | Putative outbreak cluster |
| VP09267 | VP09267 | 2009 | China | Clinical | Asia | O3:K6 | ST3 | PC | Wave-4 | Putative outbreak cluster |
| VP09273 | VP09273 | 2009 | China | Clinical | Asia | O3:K6 | ST3 | PC | Wave-4 | Putative outbreak cluster |
| VP09276 | VP09276 | 2009 | China | Clinical | Asia | O3:K6 | ST3 | PC | Wave-4 | Putative outbreak cluster |
| VP09298 | VP09298 | 2009 | China | Clinical | Asia | O3:K6 | ST3 | PC | Wave-4 | Putative outbreak cluster |

| | | | | | | | | | | |
|---|---|---|---|---|---|---|---|---|---|---|
| VP09305 | VP09305 | 2009 | China | Clinical | Asia | O3:K6 | ST3 | PC | Wave-4 | Putative outbreak cluster |
| VP09309 | VP09309 | 2009 | China | Clinical | Asia | O3:K6 | ST3 | PC | Wave-4 | Putative outbreak cluster |
| VP09312 | VP09312 | 2009 | China | Clinical | Asia | O3:K6 | ST3 | PC | Wave-4 | Putative outbreak cluster |
| VP09314 | VP09314 | 2009 | China | Clinical | Asia | O3:K6 | ST3 | PC | Wave-4 | Putative outbreak cluster |
| VP09317 | VP09317 | 2009 | China | Clinical | Asia | O3:K6 | ST3 | PC | Wave-4 | Putative outbreak cluster |
| VP09320 | VP09320 | 2009 | China | Clinical | Asia | O3:K6 | ST3 | PC | Wave-4 | Putative outbreak cluster |
| VP09323 | VP09323 | 2009 | China | Clinical | Asia | O3:K6 | ST3 | PC | Wave-4 | Putative outbreak cluster |
| VP09324 | VP09324 | 2009 | China | Clinical | Asia | O3:K6 | ST3 | PC | Wave-4 | Putative outbreak cluster |
| VP09325 | VP09325 | 2009 | China | Clinical | Asia | O3:K6 | ST3 | PC | Wave-4 | Putative outbreak cluster |
| VP09341 | VP09341 | 2009 | China | Clinical | Asia | O3:K6 | ST3 | PC | Wave-4 | Putative outbreak cluster |
| VP09342 | VP09342 | 2009 | China | Clinical | Asia | O3:K6 | ST3 | PC | Wave-4 | Putative outbreak cluster |
| VP09343 | VP09343 | 2009 | China | Clinical | Asia | O3:K6 | ST3 | PC | Wave-4 | Putative outbreak cluster |
| VP09344 | VP09344 | 2009 | China | Clinical | Asia | O3:K6 | ST3 | PC | Wave-4 | Putative outbreak cluster |
| VP09346 | VP09346 | 2009 | China | Clinical | Asia | O3:K6 | ST3 | PC | Wave-4 | Putative outbreak cluster |
| VP09347 | VP09347 | 2009 | China | Clinical | Asia | O3:K6 | ST3 | PC | Wave-4 | Putative outbreak cluster |
| VP09355 | VP09355 | 2009 | China | Clinical | Asia | O3:K6 | ST3 | PC | Wave-4 | Putative outbreak cluster |
| VP09357 | VP09357 | 2009 | China | Clinical | Asia | O3:K6 | ST3 | PC | Wave-4 | Putative outbreak cluster |
| VP09358 | VP09358 | 2009 | China | Clinical | Asia | O3:K6 | ST3 | PC | Wave-4 | Putative outbreak cluster |
| VP09362 | VP09362 | 2009 | China | Clinical | Asia | O3:K6 | ST3 | PC | Wave-4 | Putative outbreak cluster |
| VP09364 | VP09364 | 2009 | China | Clinical | Asia | O3:K6 | ST3 | PC | Wave-4 | Putative outbreak cluster |
| VP09376 | VP09376 | 2009 | China | Clinical | Asia | O3:K6 | ST3 | PC | Wave-4 | Putative outbreak cluster |
| VP09378 | VP09378 | 2009 | China | Clinical | Asia | O3:K6 | ST3 | PC | Wave-4 | Putative outbreak cluster |
| VP09379 | VP09379 | 2009 | China | Clinical | Asia | O3:K6 | ST3 | PC | Wave-4 | Putative outbreak cluster |
| VP09380 | VP09380 | 2009 | China | Clinical | Asia | O3:K6 | ST3 | PC | Wave-4 | Putative outbreak cluster |
| VP09394 | VP09394 | 2009 | China | Clinical | Asia | O3:K6 | ST3 | PC | Wave-4 | Putative outbreak cluster |
| VP09400 | VP09400 | 2009 | China | Clinical | Asia | O3:K6 | ST3 | PC | Wave-4 | Putative outbreak cluster |
| VP09403 | VP09403 | 2009 | China | Clinical | Asia | O3:K6 | ST3 | PC | Wave-4 | Putative outbreak cluster |
| VP09405 | VP09405 | 2009 | China | Clinical | Asia | O3:K6 | ST3 | PC | Wave-4 | Putative outbreak cluster |
| VP09406 | VP09406 | 2009 | China | Clinical | Asia | O3:K6 | ST3 | PC | Wave-4 | Putative outbreak cluster |
| VP09408 | VP09408 | 2009 | China | Clinical | Asia | O3:K6 | ST3 | PC | Wave-4 | Putative outbreak cluster |
| VP09411 | VP09411 | 2009 | China | Clinical | Asia | O3:K6 | ST3 | PC | Wave-4 | Putative outbreak cluster |
| VP09412 | VP09412 | 2009 | China | Clinical | Asia | O3:K6 | ST3 | PC | Wave-4 | Putative outbreak cluster |

| | | | | | | | | | | |
|---|---|---|---|---|---|---|---|---|---|---|
| VP09413 | VP09413 | 2009 | China | Clinical | Asia | O3:K6 | ST3 | PC | Wave-4 | Putative outbreak cluster |
| VP09414 | VP09414 | 2009 | China | Clinical | Asia | O3:K6 | ST3 | PC | Wave-4 | Putative outbreak cluster |
| VP09415 | VP09415 | 2009 | China | Clinical | Asia | O3:K6 | ST3 | PC | Wave-4 | Putative outbreak cluster |
| VP09416 | VP09416 | 2009 | China | Clinical | Asia | O3:K6 | ST3 | PC | Wave-4 | Putative outbreak cluster |
| VP09429 | VP09429 | 2009 | China | Clinical | Asia | O3:K6 | ST3 | PC | Wave-4 | Putative outbreak cluster |
| VP09443 | VP09443 | 2009 | China | Clinical | Asia | O3:K6 | ST3 | PC | Wave-4 | Putative outbreak cluster |
| VP09457 | VP09457 | 2009 | China | Clinical | Asia | O3:K6 | ST3 | PC | Wave-4 | Putative outbreak cluster |
| VP09461 | VP09461 | 2009 | China | Clinical | Asia | O3:K6 | ST3 | PC | Wave-4 | Putative outbreak cluster |
| VP09466 | VP09466 | 2009 | China | Clinical | Asia | O3:K6 | ST3 | PC | Wave-4 | Putative outbreak cluster |
| VP09476 | VP09476 | 2009 | China | Clinical | Asia | O3:K6 | ST3 | PC | Wave-4 | Putative outbreak cluster |
| VP10072 | VP10072 | 2010 | China | Clinical | Asia | O3:K6 | ST3 | PC | Wave-4 | Putative outbreak cluster |
| VP10073 | VP10073 | 2010 | China | Clinical | Asia | O3:K6 | ST3 | PC | Wave-4 | Putative outbreak cluster |
| VP10074 | VP10074 | 2010 | China | Clinical | Asia | O3:K6 | ST3 | PC | Wave-4 | Putative outbreak cluster |
| VP10075 | VP10075 | 2010 | China | Clinical | Asia | O3:K6 | ST3 | PC | Wave-4 | Putative outbreak cluster |
| VP10077 | VP10077 | 2010 | China | Clinical | Asia | O3:K6 | ST3 | PC | Wave-4 | Putative outbreak cluster |
| VP10078 | VP10078 | 2010 | China | Clinical | Asia | O3:K6 | ST3 | PC | Wave-4 | Putative outbreak cluster |
| VP10086 | VP10086 | 2010 | China | Clinical | Asia | O3:K6 | ST3 | PC | Wave-4 | Putative outbreak cluster |
| VP10088 | VP10088 | 2010 | China | Clinical | Asia | O3:K6 | ST3 | PC | Wave-4 | Putative outbreak cluster |
| VP10092 | VP10092 | 2010 | China | Clinical | Asia | O3:K6 | ST3 | PC | Wave-4 | Putative outbreak cluster |
| VP10106 | VP10106 | 2010 | China | Clinical | Asia | O3:K6 | ST3 | PC | Wave-4 | Putative outbreak cluster |
| VP10117 | VP10117 | 2010 | China | Clinical | Asia | O3:K6 | ST3 | PC | Wave-4 | Putative outbreak cluster |
| VP10119 | VP10119 | 2010 | China | Clinical | Asia | O3:K6 | ST3 | PC | Wave-4 | Putative outbreak cluster |
| VP10127 | VP10127 | 2010 | China | Clinical | Asia | O3:K6 | ST3 | PC | Wave-4 | Putative outbreak cluster |
| VP10131 | VP10131 | 2010 | China | Clinical | Asia | O3:K6 | ST3 | PC | Wave-4 | Putative outbreak cluster |
| VP10141 | VP10141 | 2010 | China | Clinical | Asia | O3:K6 | ST3 | PC | Wave-4 | Putative outbreak cluster |
| VP10176 | VP10176 | 2010 | China | Clinical | Asia | O3:K6 | ST3 | PC | Wave-4 | Putative outbreak cluster |
| VP10177 | VP10177 | 2010 | China | Clinical | Asia | O3:K6 | ST3 | PC | Wave-4 | Putative outbreak cluster |
| VP10182 | VP10182 | 2010 | China | Clinical | Asia | O3:K6 | ST3 | PC | Wave-4 | Putative outbreak cluster |
| VP10191 | VP10191 | 2010 | China | Clinical | Asia | O3:K6 | ST3 | PC | Wave-4 | Putative outbreak cluster |
| VP10196 | VP10196 | 2010 | China | Clinical | Asia | O3:K6 | ST3 | PC | Wave-4 | Putative outbreak cluster |
| VP10199 | VP10199 | 2010 | China | Clinical | Asia | O3:K6 | ST3 | PC | Wave-4 | Putative outbreak cluster |
| VP10200 | VP10200 | 2010 | China | Clinical | Asia | O3:K6 | ST3 | PC | Wave-4 | Putative outbreak cluster |

| | | | | | | | | | | |
|---|---|---|---|---|---|---|---|---|---|---|
| VP10201 | VP10201 | 2010 | China | Clinical | Asia | O3:K6 | ST3 | PC | Wave-4 | Putative outbreak cluster |
| VP10208 | VP10208 | 2010 | China | Clinical | Asia | O3:K6 | ST3 | PC | Wave-4 | Putative outbreak cluster |
| VP10230 | VP10230 | 2010 | China | Clinical | Asia | O3:K6 | ST3 | PC | Wave-4 | Putative outbreak cluster |
| VP10244 | VP10244 | 2010 | China | Clinical | Asia | O3:K6 | ST3 | PC | Wave-4 | Putative outbreak cluster |
| VP10249 | VP10249 | 2010 | China | Clinical | Asia | O3:K6 | ST3 | PC | Wave-4 | Putative outbreak cluster |
| VP10253 | VP10253 | 2010 | China | Clinical | Asia | O3:K6 | ST3 | PC | Wave-4 | Putative outbreak cluster |
| VP10259 | VP10259 | 2010 | China | Clinical | Asia | O3:K6 | ST3 | PC | Wave-4 | Putative outbreak cluster |
| VP10262 | VP10262 | 2010 | China | Clinical | Asia | O3:K6 | ST3 | PC | Wave-4 | Putative outbreak cluster |
| VP10268 | VP10268 | 2010 | China | Clinical | Asia | O3:K6 | ST3 | PC | Wave-4 | Putative outbreak cluster |
| VP10271 | VP10271 | 2010 | China | Clinical | Asia | O3:K6 | ST3 | PC | Wave-4 | Putative outbreak cluster |
| VP10278 | VP10278 | 2010 | China | Clinical | Asia | O3:K6 | ST3 | PC | Wave-4 | Putative outbreak cluster |
| VP10289 | VP10289 | 2010 | China | Clinical | Asia | O3:K6 | ST3 | PC | Wave-4 | Putative outbreak cluster |
| VP10298 | VP10298 | 2010 | China | Clinical | Asia | O3:K6 | ST3 | PC | Wave-4 | Putative outbreak cluster |
| VP10299 | VP10299 | 2010 | China | Clinical | Asia | O3:K6 | ST3 | PC | Wave-4 | Putative outbreak cluster |
| VP10311 | VP10311 | 2010 | China | Clinical | Asia | O3:K6 | ST3 | PC | Wave-4 | Putative outbreak cluster |
| VP10321 | VP10321 | 2010 | China | Clinical | Asia | O3:K6 | ST3 | PC | Wave-4 | Putative outbreak cluster |
| VP10328 | VP10328 | 2010 | China | Clinical | Asia | O3:K6 | ST3 | PC | Wave-4 | Putative outbreak cluster |
| VP10330 | VP10330 | 2010 | China | Clinical | Asia | O3:K6 | ST3 | PC | Wave-4 | Putative outbreak cluster |
| VP10331 | VP10331 | 2010 | China | Clinical | Asia | O3:K6 | ST3 | PC | Wave-4 | Putative outbreak cluster |
| VP10333 | VP10333 | 2010 | China | Clinical | Asia | O3:K6 | ST3 | PC | Wave-4 | Putative outbreak cluster |
| VP10335 | VP10335 | 2010 | China | Clinical | Asia | O3:K6 | ST3 | PC | Wave-4 | Putative outbreak cluster |
| VP10340 | VP10340 | 2010 | China | Clinical | Asia | O3:K6 | ST3 | PC | Wave-4 | Putative outbreak cluster |
| VP10341 | VP10341 | 2010 | China | Clinical | Asia | O3:K6 | ST3 | PC | Wave-4 | Putative outbreak cluster |
| VP10343 | VP10343 | 2010 | China | Clinical | Asia | O3:K6 | ST3 | PC | Wave-4 | Putative outbreak cluster |
| VP10344 | VP10344 | 2010 | China | Clinical | Asia | O3:K6 | ST3 | PC | Wave-4 | Putative outbreak cluster |
| VP10358 | VP10358 | 2010 | China | Clinical | Asia | O3:K6 | ST3 | PC | Wave-4 | Putative outbreak cluster |
| VP10359 | VP10359 | 2010 | China | Clinical | Asia | O3:K6 | ST3 | PC | Wave-4 | Putative outbreak cluster |
| VP10367 | VP10367 | 2010 | China | Clinical | Asia | O3:K6 | ST3 | PC | Wave-4 | Putative outbreak cluster |
| VP10376 | VP10376 | 2010 | China | Clinical | Asia | O3:K6 | ST3 | PC | Wave-4 | Putative outbreak cluster |
| VP10377 | VP10377 | 2010 | China | Clinical | Asia | O3:K6 | ST3 | PC | Wave-4 | Putative outbreak cluster |
| VP10385 | VP10385 | 2010 | China | Clinical | Asia | O3:K6 | ST3 | PC | Wave-4 | Putative outbreak cluster |
| VP10406 | VP10406 | 2010 | China | Clinical | Asia | O3:K6 | ST3 | PC | Wave-4 | Putative outbreak cluster |

| | | | | | | | | | | |
|---|---|---|---|---|---|---|---|---|---|---|
| VP10407 | VP10407 | 2010 | China | Clinical | Asia | O3:K6 | ST3 | PC | Wave-4 | Putative outbreak cluster |
| VP10408 | VP10408 | 2010 | China | Clinical | Asia | O3:K6 | ST3 | PC | Wave-4 | Putative outbreak cluster |
| VP10411 | VP10411 | 2010 | China | Clinical | Asia | O3:K6 | ST3 | PC | Wave-4 | Putative outbreak cluster |
| VP10416 | VP10416 | 2010 | China | Clinical | Asia | O3:K6 | ST3 | PC | Wave-4 | Putative outbreak cluster |
| VP10420 | VP10420 | 2010 | China | Clinical | Asia | O3:K6 | ST3 | PC | Wave-4 | Putative outbreak cluster |
| VP10421 | VP10421 | 2010 | China | Clinical | Asia | O3:K6 | ST3 | PC | Wave-4 | Putative outbreak cluster |
| VP10424 | VP10424 | 2010 | China | Clinical | Asia | O3:K6 | ST3 | PC | Wave-4 | Putative outbreak cluster |
| VP10431 | VP10431 | 2010 | China | Clinical | Asia | O3:K6 | ST3 | PC | Wave-4 | Putative outbreak cluster |
| VP10443 | VP10443 | 2010 | China | Clinical | Asia | O3:K6 | ST3 | PC | Wave-4 | Putative outbreak cluster |
| VP10445 | VP10445 | 2010 | China | Clinical | Asia | O3:K6 | ST3 | PC | Wave-4 | Putative outbreak cluster |
| VP10448 | VP10448 | 2010 | China | Clinical | Asia | O3:K6 | ST3 | PC | Wave-4 | Putative outbreak cluster |
| VP10452 | VP10452 | 2010 | China | Clinical | Asia | O3:K6 | ST3 | PC | Wave-4 | Putative outbreak cluster |
| VP10456 | VP10456 | 2010 | China | Clinical | Asia | O3:K6 | ST3 | PC | Wave-4 | Putative outbreak cluster |
| VP10457 | VP10457 | 2010 | China | Clinical | Asia | O3:K6 | ST3 | PC | Wave-4 | Putative outbreak cluster |
| VP10458 | VP10458 | 2010 | China | Clinical | Asia | O3:K6 | ST3 | PC | Wave-4 | Putative outbreak cluster |
| VP10459 | VP10459 | 2010 | China | Clinical | Asia | O3:K6 | ST3 | PC | Wave-4 | Putative outbreak cluster |
| VP10460 | VP10460 | 2010 | China | Clinical | Asia | O3:K6 | ST3 | PC | Wave-4 | Putative outbreak cluster |
| VP10461 | VP10461 | 2010 | China | Clinical | Asia | O3:K6 | ST3 | PC | Wave-4 | Putative outbreak cluster |
| VP10462 | VP10462 | 2010 | China | Clinical | Asia | O3:K6 | ST3 | PC | Wave-4 | Putative outbreak cluster |
| VP10464 | VP10464 | 2010 | China | Clinical | Asia | O3:K6 | ST3 | PC | Wave-4 | Putative outbreak cluster |
| VP10466 | VP10466 | 2010 | China | Clinical | Asia | O3:K6 | ST3 | PC | Wave-4 | Putative outbreak cluster |
| VP10467 | VP10467 | 2010 | China | Clinical | Asia | O3:K6 | ST3 | PC | Wave-4 | Putative outbreak cluster |
| VP10469 | VP10469 | 2010 | China | Clinical | Asia | O3:K6 | ST3 | PC | Wave-4 | Putative outbreak cluster |
| VP10471 | VP10471 | 2010 | China | Clinical | Asia | O3:K6 | ST3 | PC | Wave-4 | Putative outbreak cluster |
| VP10476 | VP10476 | 2010 | China | Clinical | Asia | O3:K6 | ST3 | PC | Wave-4 | Putative outbreak cluster |
| VP10478 | VP10478 | 2010 | China | Clinical | Asia | O3:K6 | ST3 | PC | Wave-4 | Putative outbreak cluster |
| VP10481 | VP10481 | 2010 | China | Clinical | Asia | O3:K6 | ST3 | PC | Wave-4 | Putative outbreak cluster |
| VP10482 | VP10482 | 2010 | China | Clinical | Asia | O3:K6 | ST3 | PC | Wave-4 | Putative outbreak cluster |
| VP11124 | VP11124 | 2011 | China | Clinical | Asia | O3:K6 | ST3 | PC | Wave-4 | Putative outbreak cluster |
| VP11125 | VP11125 | 2011 | China | Clinical | Asia | O3:K6 | ST3 | PC | Wave-4 | Putative outbreak cluster |
| VP11128 | VP11128 | 2011 | China | Clinical | Asia | O3:K6 | ST3 | PC | Wave-4 | Putative outbreak cluster |
| VP11132 | VP11132 | 2011 | China | Clinical | Asia | O3:K6 | ST3 | PC | Wave-4 | Putative outbreak cluster |

| | | | | | | | | | | |
|---|---|---|---|---|---|---|---|---|---|---|
| VP11133 | VP11133 | 2011 | China | Clinical | Asia | O3:K6 | ST3 | PC | Wave-4 | Putative outbreak cluster |
| VP11140 | VP11140 | 2011 | China | Clinical | Asia | O3:K6 | ST3 | PC | Wave-4 | Putative outbreak cluster |
| VP11143 | VP11143 | 2011 | China | Clinical | Asia | O3:K6 | ST3 | PC | Wave-4 | Putative outbreak cluster |
| VP11146 | VP11146 | 2011 | China | Clinical | Asia | O3:K6 | ST3 | PC | Wave-4 | Putative outbreak cluster |
| VP11147 | VP11147 | 2011 | China | Clinical | Asia | O3:K6 | ST3 | PC | Wave-4 | Putative outbreak cluster |
| VP11159 | VP11159 | 2011 | China | Clinical | Asia | O3:K6 | ST3 | PC | Wave-4 | Putative outbreak cluster |
| VP11166 | VP11166 | 2011 | China | Clinical | Asia | O3:K6 | ST3 | PC | Wave-4 | Putative outbreak cluster |
| VP11177 | VP11177 | 2011 | China | Clinical | Asia | O3:K6 | ST3 | PC | Wave-4 | Putative outbreak cluster |
| VP11181 | VP11181 | 2011 | China | Clinical | Asia | O3:K6 | ST3 | PC | Wave-4 | Putative outbreak cluster |
| VP11188 | VP11188 | 2011 | China | Clinical | Asia | O3:K6 | ST3 | PC | Wave-4 | Putative outbreak cluster |
| VP11191 | VP11191 | 2011 | China | Clinical | Asia | O3:K6 | ST3 | PC | Wave-4 | Putative outbreak cluster |
| VP11194 | VP11194 | 2011 | China | Clinical | Asia | O3:K6 | ST3 | PC | Wave-4 | Putative outbreak cluster |
| VP11195 | VP11195 | 2011 | China | Clinical | Asia | O3:K6 | ST3 | PC | Wave-4 | Putative outbreak cluster |
| VP11201 | VP11201 | 2011 | China | Clinical | Asia | O3:K6 | ST3 | PC | Wave-4 | Putative outbreak cluster |
| VP11202 | VP11202 | 2011 | China | Clinical | Asia | O3:K6 | ST3 | PC | Wave-4 | Putative outbreak cluster |
| VP11207 | VP11207 | 2011 | China | Clinical | Asia | O3:K6 | ST3 | PC | Wave-4 | Putative outbreak cluster |
| VP11208 | VP11208 | 2011 | China | Clinical | Asia | O3:K6 | ST3 | PC | Wave-4 | Putative outbreak cluster |
| VP11209 | VP11209 | 2011 | China | Clinical | Asia | O3:K6 | ST3 | PC | Wave-4 | Putative outbreak cluster |
| VP11211 | VP11211 | 2011 | China | Clinical | Asia | O3:K6 | ST3 | PC | Wave-4 | Putative outbreak cluster |
| VP11212 | VP11212 | 2011 | China | Clinical | Asia | O3:K6 | ST3 | PC | Wave-4 | Putative outbreak cluster |
| VP11214 | VP11214 | 2011 | China | Clinical | Asia | O3:K6 | ST3 | PC | Wave-4 | Putative outbreak cluster |
| VP11215 | VP11215 | 2011 | China | Clinical | Asia | O3:K6 | ST3 | PC | Wave-4 | Putative outbreak cluster |
| VP11216 | VP11216 | 2011 | China | Clinical | Asia | O3:K6 | ST3 | PC | Wave-4 | Putative outbreak cluster |
| VP11217 | VP11217 | 2011 | China | Clinical | Asia | O3:K6 | ST3 | PC | Wave-4 | Putative outbreak cluster |
| VP11218 | VP11218 | 2011 | China | Clinical | Asia | O3:K6 | ST3 | PC | Wave-4 | Putative outbreak cluster |
| VP11219 | VP11219 | 2011 | China | Clinical | Asia | O3:K6 | ST3 | PC | Wave-4 | Putative outbreak cluster |
| VP11221 | VP11221 | 2011 | China | Clinical | Asia | O3:K6 | ST3 | PC | Wave-4 | Putative outbreak cluster |
| VP11222 | VP11222 | 2011 | China | Clinical | Asia | O3:K6 | ST3 | PC | Wave-4 | Putative outbreak cluster |
| VP11223 | VP11223 | 2011 | China | Clinical | Asia | O3:K6 | ST3 | PC | Wave-4 | Putative outbreak cluster |
| VP11224 | VP11224 | 2011 | China | Clinical | Asia | O3:K6 | ST3 | PC | Wave-4 | Putative outbreak cluster |
| VP11226 | VP11226 | 2011 | China | Clinical | Asia | O3:K6 | ST3 | PC | Wave-4 | Putative outbreak cluster |
| VP11229 | VP11229 | 2011 | China | Clinical | Asia | O3:K6 | ST3 | PC | Wave-4 | Putative outbreak cluster |

| | | | | | | | | | | |
|---|---|---|---|---|---|---|---|---|---|---|
| VP11230 | VP11230 | 2011 | China | Clinical | Asia | O3:K6 | ST3 | PC | Wave-4 | Putative outbreak cluster |
| VP11232 | VP11232 | 2011 | China | Clinical | Asia | O3:K6 | ST3 | PC | Wave-4 | Putative outbreak cluster |
| VP11233 | VP11233 | 2011 | China | Clinical | Asia | O3:K6 | ST3 | PC | Wave-4 | Putative outbreak cluster |
| VP11235 | VP11235 | 2011 | China | Clinical | Asia | O3:K6 | ST3 | PC | Wave-4 | Putative outbreak cluster |
| VP11236 | VP11236 | 2011 | China | Clinical | Asia | O3:K6 | ST3 | PC | Wave-4 | Putative outbreak cluster |
| VP11238 | VP11238 | 2011 | China | Clinical | Asia | O3:K6 | ST3 | PC | Wave-4 | Putative outbreak cluster |
| VP11245 | VP11245 | 2011 | China | Clinical | Asia | O3:K6 | ST3 | PC | Wave-4 | Putative outbreak cluster |
| VP11259 | VP11259 | 2011 | China | Clinical | Asia | O3:K6 | ST3 | PC | Wave-4 | Putative outbreak cluster |
| VP11263 | VP11263 | 2011 | China | Clinical | Asia | O3:K6 | ST3 | PC | Wave-4 | Putative outbreak cluster |
| VP11265 | VP11265 | 2011 | China | Clinical | Asia | O3:K6 | ST3 | PC | Wave-4 | Putative outbreak cluster |
| VP11266 | VP11266 | 2011 | China | Clinical | Asia | O3:K6 | ST3 | PC | Wave-4 | Putative outbreak cluster |
| VP11267 | VP11267 | 2011 | China | Clinical | Asia | O3:K6 | ST3 | PC | Wave-4 | Putative outbreak cluster |
| VP11268 | VP11268 | 2011 | China | Clinical | Asia | O3:K6 | ST3 | PC | Wave-4 | Putative outbreak cluster |
| VP11276 | VP11276 | 2011 | China | Clinical | Asia | O3:K6 | ST3 | PC | Wave-4 | Putative outbreak cluster |
| VP11280 | VP11280 | 2011 | China | Clinical | Asia | O3:K6 | ST3 | PC | Wave-4 | Putative outbreak cluster |
| VP11281 | VP11281 | 2011 | China | Clinical | Asia | O3:K6 | ST3 | PC | Wave-4 | Putative outbreak cluster |
| VP11283 | VP11283 | 2011 | China | Clinical | Asia | O3:K6 | ST3 | PC | Wave-4 | Putative outbreak cluster |
| VP11285 | VP11285 | 2011 | China | Clinical | Asia | O3:K6 | ST3 | PC | Wave-4 | Putative outbreak cluster |
| VP11286 | VP11286 | 2011 | China | Clinical | Asia | O3:K6 | ST3 | PC | Wave-4 | Putative outbreak cluster |
| VP11287 | VP11287 | 2011 | China | Clinical | Asia | O3:K6 | ST3 | PC | Wave-4 | Putative outbreak cluster |
| VP11289 | VP11289 | 2011 | China | Clinical | Asia | O3:K6 | ST3 | PC | Wave-4 | Putative outbreak cluster |
| VP11291 | VP11291 | 2011 | China | Clinical | Asia | O3:K6 | ST3 | PC | Wave-4 | Putative outbreak cluster |
| VP11293 | VP11293 | 2011 | China | Clinical | Asia | O3:K6 | ST3 | PC | Wave-4 | Putative outbreak cluster |
| VP11294 | VP11294 | 2011 | China | Clinical | Asia | O3:K6 | ST3 | PC | Wave-4 | Putative outbreak cluster |
| VP11297 | VP11297 | 2011 | China | Clinical | Asia | O3:K6 | ST3 | PC | Wave-4 | Putative outbreak cluster |
| VP11299 | VP11299 | 2011 | China | Clinical | Asia | O3:K6 | ST3 | PC | Wave-4 | Putative outbreak cluster |
| VP11303 | VP11303 | 2011 | China | Clinical | Asia | O3:K6 | ST3 | PC | Wave-4 | Putative outbreak cluster |
| VP11304 | VP11304 | 2011 | China | Clinical | Asia | O3:K6 | ST3 | PC | Wave-4 | Putative outbreak cluster |
| VP11306 | VP11306 | 2011 | China | Clinical | Asia | O3:K6 | ST3 | PC | Wave-4 | Putative outbreak cluster |
| VP11307 | VP11307 | 2011 | China | Clinical | Asia | O3:K6 | ST3 | PC | Wave-4 | Putative outbreak cluster |
| VP11308 | VP11308 | 2011 | China | Clinical | Asia | O3:K6 | ST3 | PC | Wave-4 | Putative outbreak cluster |
| VP11311 | VP11311 | 2011 | China | Clinical | Asia | O3:K6 | ST3 | PC | Wave-4 | Putative outbreak cluster |

| | | | | | | | | | | |
|---|---|---|---|---|---|---|---|---|---|---|
| VP12005 | VP12005 | 2012 | China | Clinical | Asia | O3:K6 | ST3 | PC | Wave-4 | Putative outbreak cluster |
| VP12026 | VP12026 | 2012 | China | Clinical | Asia | O3:K6 | ST3 | PC | Wave-4 | Putative outbreak cluster |
| VP12030 | VP12030 | 2012 | China | Clinical | Asia | O3:K6 | ST3 | PC | Wave-4 | Putative outbreak cluster |
| VP12033 | VP12033 | 2012 | China | Clinical | Asia | O3:K6 | ST3 | PC | Wave-4 | Putative outbreak cluster |
| VP12036 | VP12036 | 2012 | China | Clinical | Asia | O3:K6 | ST3 | PC | Wave-4 | Putative outbreak cluster |
| VP12037 | VP12037 | 2012 | China | Clinical | Asia | O3:K6 | ST3 | PC | Wave-4 | Putative outbreak cluster |
| VP12039 | VP12039 | 2012 | China | Clinical | Asia | O3:K6 | ST3 | PC | Wave-4 | Putative outbreak cluster |
| VP12040 | VP12040 | 2012 | China | Clinical | Asia | O3:K6 | ST3 | PC | Wave-4 | Putative outbreak cluster |
| VP12041 | VP12041 | 2012 | China | Clinical | Asia | O3:K6 | ST3 | PC | Wave-4 | Putative outbreak cluster |
| VP12043 | VP12043 | 2012 | China | Clinical | Asia | O3:K6 | ST3 | PC | Wave-4 | Putative outbreak cluster |
| VP12046 | VP12046 | 2012 | China | Clinical | Asia | O3:K6 | ST3 | PC | Wave-4 | Putative outbreak cluster |
| VP12048 | VP12048 | 2012 | China | Clinical | Asia | O3:K6 | ST3 | PC | Wave-4 | Putative outbreak cluster |
| VP12049 | VP12049 | 2012 | China | Clinical | Asia | O3:K6 | ST3 | PC | Wave-4 | Putative outbreak cluster |
| VP12051 | VP12051 | 2012 | China | Clinical | Asia | O3:K6 | ST3 | PC | Wave-4 | Putative outbreak cluster |
| VP12052 | VP12052 | 2012 | China | Clinical | Asia | O3:K6 | ST3 | PC | Wave-4 | Putative outbreak cluster |
| VP12054 | VP12054 | 2012 | China | Clinical | Asia | O3:K6 | ST3 | PC | Wave-4 | Putative outbreak cluster |
| VP12056 | VP12056 | 2012 | China | Clinical | Asia | O3:K6 | ST3 | PC | Wave-4 | Putative outbreak cluster |
| VP12057 | VP12057 | 2012 | China | Clinical | Asia | O3:K6 | ST3 | PC | Wave-4 | Putative outbreak cluster |
| VP12063 | VP12063 | 2012 | China | Clinical | Asia | O3:K6 | ST3 | PC | Wave-4 | Putative outbreak cluster |
| VP12064 | VP12064 | 2012 | China | Clinical | Asia | O3:K6 | ST3 | PC | Wave-4 | Putative outbreak cluster |
| VP12065 | VP12065 | 2012 | China | Clinical | Asia | O3:K6 | ST3 | PC | Wave-4 | Putative outbreak cluster |
| VP12067 | VP12067 | 2012 | China | Clinical | Asia | O3:K6 | ST3 | PC | Wave-4 | Putative outbreak cluster |
| VP12068 | VP12068 | 2012 | China | Clinical | Asia | O3:K6 | ST3 | PC | Wave-4 | Putative outbreak cluster |
| VP12070 | VP12070 | 2012 | China | Clinical | Asia | O3:K6 | ST3 | PC | Wave-4 | Putative outbreak cluster |
| VP12071 | VP12071 | 2012 | China | Clinical | Asia | O3:K6 | ST3 | PC | Wave-4 | Putative outbreak cluster |
| VP12072 | VP12072 | 2012 | China | Clinical | Asia | O3:K6 | ST3 | PC | Wave-4 | Putative outbreak cluster |
| VP12078 | VP12078 | 2012 | China | Clinical | Asia | O3:K6 | ST3 | PC | Wave-4 | Putative outbreak cluster |
| VP12084 | VP12084 | 2012 | China | Clinical | Asia | O3:K6 | ST3 | PC | Wave-4 | Putative outbreak cluster |
| VP12085 | VP12085 | 2012 | China | Clinical | Asia | O3:K6 | ST3 | PC | Wave-4 | Putative outbreak cluster |
| VP12088 | VP12088 | 2012 | China | Clinical | Asia | O3:K6 | ST3 | PC | Wave-4 | Putative outbreak cluster |
| VP12089 | VP12089 | 2012 | China | Clinical | Asia | O3:K6 | ST3 | PC | Wave-4 | Putative outbreak cluster |
| VP12090 | VP12090 | 2012 | China | Clinical | Asia | O3:K6 | ST3 | PC | Wave-4 | Putative outbreak cluster |

| | | | | | | | | | | |
|---|---|---|---|---|---|---|---|---|---|---|
| VP12091 | VP12091 | 2012 | China | Clinical | Asia | O3:K6 | ST3 | PC | Wave-4 | Putative outbreak cluster |
| VP12092 | VP12092 | 2012 | China | Clinical | Asia | O3:K6 | ST3 | PC | Wave-4 | Putative outbreak cluster |
| VP12093 | VP12093 | 2012 | China | Clinical | Asia | O3:K6 | ST3 | PC | Wave-4 | Putative outbreak cluster |
| VP12094 | VP12094 | 2012 | China | Clinical | Asia | O3:K6 | ST3 | PC | Wave-4 | Putative outbreak cluster |
| VP12095 | VP12095 | 2012 | China | Clinical | Asia | O3:K6 | ST3 | PC | Wave-4 | Putative outbreak cluster |
| VP12096 | VP12096 | 2012 | China | Clinical | Asia | O3:K6 | ST3 | PC | Wave-4 | Putative outbreak cluster |
| VP12097 | VP12097 | 2012 | China | Clinical | Asia | O3:K6 | ST3 | PC | Wave-4 | Putative outbreak cluster |
| VP12098 | VP12098 | 2012 | China | Clinical | Asia | O3:K6 | ST3 | PC | Wave-4 | Putative outbreak cluster |
| VP12099 | VP12099 | 2012 | China | Clinical | Asia | O3:K6 | ST3 | PC | Wave-4 | Putative outbreak cluster |
| VP12100 | VP12100 | 2012 | China | Clinical | Asia | O3:K6 | ST3 | PC | Wave-4 | Putative outbreak cluster |
| VP12101 | VP12101 | 2012 | China | Clinical | Asia | O3:K6 | ST3 | PC | Wave-4 | Putative outbreak cluster |
| VP12107 | VP12107 | 2012 | China | Clinical | Asia | O3:K6 | ST3 | PC | Wave-4 | Putative outbreak cluster |
| VP12108 | VP12108 | 2012 | China | Clinical | Asia | O3:K6 | ST3 | PC | Wave-4 | Putative outbreak cluster |
| VP12110 | VP12110 | 2012 | China | Clinical | Asia | O3:K6 | ST3 | PC | Wave-4 | Putative outbreak cluster |
| VP12111 | VP12111 | 2012 | China | Clinical | Asia | O3:K6 | ST3 | PC | Wave-4 | Putative outbreak cluster |
| VP12112 | VP12112 | 2012 | China | Clinical | Asia | O3:K6 | ST3 | PC | Wave-4 | Putative outbreak cluster |
| VP12113 | VP12113 | 2012 | China | Clinical | Asia | O3:K6 | ST3 | PC | Wave-4 | Putative outbreak cluster |
| VP12115 | VP12115 | 2012 | China | Clinical | Asia | O3:K6 | ST3 | PC | Wave-4 | Putative outbreak cluster |
| VP12117 | VP12117 | 2012 | China | Clinical | Asia | O3:K6 | ST3 | PC | Wave-4 | Putative outbreak cluster |
| VP12124 | VP12124 | 2012 | China | Clinical | Asia | O3:K6 | ST3 | PC | Wave-4 | Putative outbreak cluster |
| VP12125 | VP12125 | 2012 | China | Clinical | Asia | O3:K6 | ST3 | PC | Wave-4 | Putative outbreak cluster |
| VP12126 | VP12126 | 2012 | China | Clinical | Asia | O3:K6 | ST3 | PC | Wave-4 | Putative outbreak cluster |
| VP12127 | VP12127 | 2012 | China | Clinical | Asia | O3:K6 | ST3 | PC | Wave-4 | Putative outbreak cluster |
| VP12128 | VP12128 | 2012 | China | Clinical | Asia | O3:K6 | ST3 | PC | Wave-4 | Putative outbreak cluster |
| VP12129 | VP12129 | 2012 | China | Clinical | Asia | O3:K6 | ST3 | PC | Wave-4 | Putative outbreak cluster |
| VP12132 | VP12132 | 2012 | China | Clinical | Asia | O3:K6 | ST3 | PC | Wave-4 | Putative outbreak cluster |
| VP12135 | VP12135 | 2012 | China | Clinical | Asia | O3:K6 | ST3 | PC | Wave-4 | Putative outbreak cluster |
| VP12137 | VP12137 | 2012 | China | Clinical | Asia | O3:K6 | ST3 | PC | Wave-4 | Putative outbreak cluster |
| VP12138 | VP12138 | 2012 | China | Clinical | Asia | O3:K6 | ST3 | PC | Wave-4 | Putative outbreak cluster |
| VP12139 | VP12139 | 2012 | China | Clinical | Asia | O3:K6 | ST3 | PC | Wave-4 | Putative outbreak cluster |
| VP12140 | VP12140 | 2012 | China | Clinical | Asia | O3:K6 | ST3 | PC | Wave-4 | Putative outbreak cluster |
| VP12141 | VP12141 | 2012 | China | Clinical | Asia | O3:K6 | ST3 | PC | Wave-4 | Putative outbreak cluster |

| | | | | | | | | | | |
|---|---|---|---|---|---|---|---|---|---|---|
| VP12142 | VP12142 | 2012 | China | Clinical | Asia | O3:K6 | ST3 | PC | Wave-4 | Putative outbreak cluster |
| VP12143 | VP12143 | 2012 | China | Clinical | Asia | O3:K6 | ST3 | PC | Wave-4 | Putative outbreak cluster |
| VP12146 | VP12146 | 2012 | China | Clinical | Asia | O3:K6 | ST3 | PC | Wave-4 | Putative outbreak cluster |
| VP12147 | VP12147 | 2012 | China | Clinical | Asia | O3:K6 | ST3 | PC | Wave-4 | Putative outbreak cluster |
| VP12148 | VP12148 | 2012 | China | Clinical | Asia | O3:K6 | ST3 | PC | Wave-4 | Putative outbreak cluster |
| VP12149 | VP12149 | 2012 | China | Clinical | Asia | O3:K6 | ST3 | PC | Wave-4 | Putative outbreak cluster |
| VP12150 | VP12150 | 2012 | China | Clinical | Asia | O3:K6 | ST3 | PC | Wave-4 | Putative outbreak cluster |
| VP12151 | VP12151 | 2012 | China | Clinical | Asia | O3:K6 | ST3 | PC | Wave-4 | Putative outbreak cluster |
| VP12152 | VP12152 | 2012 | China | Clinical | Asia | O3:K6 | ST3 | PC | Wave-4 | Putative outbreak cluster |
| VP12153 | VP12153 | 2012 | China | Clinical | Asia | O3:K6 | ST3 | PC | Wave-4 | Putative outbreak cluster |
| VP12155 | VP12155 | 2012 | China | Clinical | Asia | O3:K6 | ST3 | PC | Wave-4 | Putative outbreak cluster |
| VP12156 | VP12156 | 2012 | China | Clinical | Asia | O3:K6 | ST3 | PC | Wave-4 | Putative outbreak cluster |
| VP12157 | VP12157 | 2012 | China | Clinical | Asia | O3:K6 | ST3 | PC | Wave-4 | Putative outbreak cluster |
| VP12158 | VP12158 | 2012 | China | Clinical | Asia | O3:K6 | ST3 | PC | Wave-4 | Putative outbreak cluster |
| VP12159 | VP12159 | 2012 | China | Clinical | Asia | O3:K6 | ST3 | PC | Wave-4 | Putative outbreak cluster |
| VP12160 | VP12160 | 2012 | China | Clinical | Asia | O3:K6 | ST3 | PC | Wave-4 | Putative outbreak cluster |
| VP12161 | VP12161 | 2012 | China | Clinical | Asia | O3:K6 | ST3 | PC | Wave-4 | Putative outbreak cluster |
| VP12164 | VP12164 | 2012 | China | Clinical | Asia | O3:K6 | ST3 | PC | Wave-4 | Putative outbreak cluster |
| VP12169 | VP12169 | 2012 | China | Clinical | Asia | O3:K6 | ST3 | PC | Wave-4 | Putative outbreak cluster |
| VP12172 | VP12172 | 2012 | China | Clinical | Asia | O3:K6 | ST3 | PC | Wave-4 | Putative outbreak cluster |
| VP12175 | VP12175 | 2012 | China | Clinical | Asia | O3:K6 | ST3 | PC | Wave-4 | Putative outbreak cluster |
| VP12176 | VP12176 | 2012 | China | Clinical | Asia | O3:K6 | ST3 | PC | Wave-4 | Putative outbreak cluster |
| VP12177 | VP12177 | 2012 | China | Clinical | Asia | O3:K6 | ST3 | PC | Wave-4 | Putative outbreak cluster |
| VP12178 | VP12178 | 2012 | China | Clinical | Asia | O3:K6 | ST3 | PC | Wave-4 | Putative outbreak cluster |
| VP12179 | VP12179 | 2012 | China | Clinical | Asia | O3:K6 | ST3 | PC | Wave-4 | Putative outbreak cluster |
| VP12180 | VP12180 | 2012 | China | Clinical | Asia | O3:K6 | ST3 | PC | Wave-4 | Putative outbreak cluster |
| VP12181 | VP12181 | 2012 | China | Clinical | Asia | O3:K6 | ST3 | PC | Wave-4 | Putative outbreak cluster |
| VP12182 | VP12182 | 2012 | China | Clinical | Asia | O3:K6 | ST3 | PC | Wave-4 | Putative outbreak cluster |
| VP12183 | VP12183 | 2012 | China | Clinical | Asia | O3:K6 | ST3 | PC | Wave-4 | Putative outbreak cluster |
| VP12186 | VP12186 | 2012 | China | Clinical | Asia | O3:K6 | ST3 | PC | Wave-4 | Putative outbreak cluster |
| VP12191 | VP12191 | 2012 | China | Clinical | Asia | O3:K6 | ST3 | PC | Wave-4 | Putative outbreak cluster |
| VP12192 | VP12192 | 2012 | China | Clinical | Asia | O3:K6 | ST3 | PC | Wave-4 | Putative outbreak cluster |

| | | | | | | | | | | |
|---|---|---|---|---|---|---|---|---|---|---|
| VP12193 | VP12193 | 2012 | China | Clinical | Asia | O3:K6 | ST3 | PC | Wave-4 | Putative outbreak cluster |
| VP12194 | VP12194 | 2012 | China | Clinical | Asia | O3:K6 | ST3 | PC | Wave-4 | Putative outbreak cluster |
| VP12200 | VP12200 | 2012 | China | Clinical | Asia | O3:K6 | ST3 | PC | Wave-4 | Putative outbreak cluster |
| VP12204 | VP12204 | 2012 | China | Clinical | Asia | O3:K6 | ST3 | PC | Wave-4 | Putative outbreak cluster |
| VP12207 | VP12207 | 2012 | China | Clinical | Asia | O3:K6 | ST3 | PC | Wave-4 | Putative outbreak cluster |
| VP12210 | VP12210 | 2012 | China | Clinical | Asia | O3:K6 | ST3 | PC | Wave-4 | Putative outbreak cluster |
| VP12213 | VP12213 | 2012 | China | Clinical | Asia | O3:K6 | ST3 | PC | Wave-4 | Putative outbreak cluster |
| VP12218 | VP12218 | 2012 | China | Clinical | Asia | O3:K6 | ST3 | PC | Wave-4 | Putative outbreak cluster |
| VP12220 | VP12220 | 2012 | China | Clinical | Asia | O3:K6 | ST3 | PC | Wave-4 | Putative outbreak cluster |
| VP12221 | VP12221 | 2012 | China | Clinical | Asia | O3:K6 | ST3 | PC | Wave-4 | Putative outbreak cluster |
| VP12223 | VP12223 | 2012 | China | Clinical | Asia | O3:K6 | ST3 | PC | Wave-4 | Putative outbreak cluster |
| VP12224 | VP12224 | 2012 | China | Clinical | Asia | O3:K6 | ST3 | PC | Wave-4 | Putative outbreak cluster |
| VP12225 | VP12225 | 2012 | China | Clinical | Asia | O3:K6 | ST3 | PC | Wave-4 | Putative outbreak cluster |
| VP12226 | VP12226 | 2012 | China | Clinical | Asia | O3:K6 | ST3 | PC | Wave-4 | Putative outbreak cluster |
| VP12229 | VP12229 | 2012 | China | Clinical | Asia | O3:K6 | ST3 | PC | Wave-4 | Putative outbreak cluster |
| VP12236 | VP12236 | 2012 | China | Clinical | Asia | O3:K6 | ST3 | PC | Wave-4 | Putative outbreak cluster |
| VP13008 | VP13008 | 2013 | China | Clinical | Asia | O10:K60 | ST3 | PC | Wave-4 | Putative outbreak cluster |
| VP13012 | VP13012 | 2013 | China | Clinical | Asia | O3:K6 | ST3 | PC | Wave-4 | Putative outbreak cluster |
| VP13014 | VP13014 | 2013 | China | Clinical | Asia | O3:K6 | ST3 | PC | Wave-4 | Putative outbreak cluster |
| VP13016 | VP13016 | 2013 | China | Clinical | Asia | O3:K6 | ST3 | PC | Wave-4 | Putative outbreak cluster |
| VP13023 | VP13023 | 2013 | China | Clinical | Asia | O3:K6 | ST3 | PC | Wave-4 | Putative outbreak cluster |
| VP13024 | VP13024 | 2013 | China | Clinical | Asia | O3:K6 | ST3 | PC | Wave-4 | Putative outbreak cluster |
| VP13026 | VP13026 | 2013 | China | Clinical | Asia | O3:K6 | ST3 | PC | Wave-4 | Putative outbreak cluster |
| VP13027 | VP13027 | 2013 | China | Clinical | Asia | O3:K6 | ST3 | PC | Wave-4 | Putative outbreak cluster |
| VP13029 | VP13029 | 2013 | China | Clinical | Asia | O3:K6 | ST3 | PC | Wave-4 | Putative outbreak cluster |
| VP13030 | VP13030 | 2013 | China | Clinical | Asia | O3:K6 | ST3 | PC | Wave-4 | Putative outbreak cluster |
| VP13032 | VP13032 | 2013 | China | Clinical | Asia | O3:K6 | ST3 | PC | Wave-4 | Putative outbreak cluster |
| VP13034 | VP13034 | 2013 | China | Clinical | Asia | O3:K6 | ST3 | PC | Wave-4 | Putative outbreak cluster |
| VP13035 | VP13035 | 2013 | China | Clinical | Asia | O3:K6 | ST3 | PC | Wave-4 | Putative outbreak cluster |
| VP13036 | VP13036 | 2013 | China | Clinical | Asia | O3:K6 | ST3 | PC | Wave-4 | Putative outbreak cluster |
| VP13038 | VP13038 | 2013 | China | Clinical | Asia | O3:K6 | ST3 | PC | Wave-4 | Putative outbreak cluster |
| VP13039 | VP13039 | 2013 | China | Clinical | Asia | O3:K6 | ST3 | PC | Wave-4 | Putative outbreak cluster |

| | | | | | | | | | | |
|---|---|---|---|---|---|---|---|---|---|---|
| VP13042 | VP13042 | 2013 | China | Clinical | Asia | O3:K6 | ST3 | PC | Wave-4 | Putative outbreak cluster |
| VP13049 | VP13049 | 2013 | China | Clinical | Asia | O3:K6 | ST3 | PC | Wave-4 | Putative outbreak cluster |
| VP13052 | VP13052 | 2013 | China | Clinical | Asia | O3:K6 | ST3 | PC | Wave-4 | Putative outbreak cluster |
| VP13054 | VP13054 | 2013 | China | Clinical | Asia | O3:K6 | ST3 | PC | Wave-4 | Putative outbreak cluster |
| VP13055 | VP13055 | 2013 | China | Clinical | Asia | O3:K6 | ST3 | PC | Wave-4 | Putative outbreak cluster |
| VP13056 | VP13056 | 2013 | China | Clinical | Asia | O3:K6 | ST3 | PC | Wave-4 | Putative outbreak cluster |
| VP13057 | VP13057 | 2013 | China | Clinical | Asia | O3:K6 | ST3 | PC | Wave-4 | Putative outbreak cluster |
| VP13064 | VP13064 | 2013 | China | Clinical | Asia | O3:K6 | ST3 | PC | Wave-4 | Putative outbreak cluster |
| VP13065 | VP13065 | 2013 | China | Clinical | Asia | O3:K6 | ST3 | PC | Wave-4 | Putative outbreak cluster |
| VP13073 | VP13073 | 2013 | China | Clinical | Asia | O3:K6 | ST3 | PC | Wave-4 | Putative outbreak cluster |
| VP13091 | VP13091 | 2013 | China | Clinical | Asia | O3:K6 | ST3 | PC | Wave-4 | Putative outbreak cluster |
| VP13099 | VP13099 | 2013 | China | Clinical | Asia | O3:K6 | ST3 | PC | Wave-4 | Putative outbreak cluster |
| VP13101 | VP13101 | 2013 | China | Clinical | Asia | O3:K6 | ST3 | PC | Wave-4 | Putative outbreak cluster |
| VP13103 | VP13103 | 2013 | China | Clinical | Asia | O3:K6 | ST3 | PC | Wave-4 | Putative outbreak cluster |
| VP13109 | VP13109 | 2013 | China | Clinical | Asia | O3:K6 | ST3 | PC | Wave-4 | Putative outbreak cluster |
| VP14006 | VP14006 | 2014 | China | Clinical | Asia | O3:K6 | ST3 | PC | Wave-4 | Putative outbreak cluster |
| VP14009 | VP14009 | 2014 | China | Clinical | Asia | O3:K6 | ST3 | PC | Wave-4 | Putative outbreak cluster |
| VP14011 | VP14011 | 2014 | China | Clinical | Asia | O3:K6 | ST3 | PC | Wave-4 | Putative outbreak cluster |
| VP14013 | VP14013 | 2014 | China | Clinical | Asia | O3:K6 | ST3 | PC | Wave-4 | Putative outbreak cluster |
| VP14014 | VP14014 | 2014 | China | Clinical | Asia | O3:K6 | ST3 | PC | Wave-4 | Putative outbreak cluster |
| VP14015 | VP14015 | 2014 | China | Clinical | Asia | O3:K6 | ST3 | PC | Wave-4 | Putative outbreak cluster |
| VP14018 | VP14018 | 2014 | China | Clinical | Asia | O3:K6 | ST3 | PC | Wave-4 | Putative outbreak cluster |
| VP14061 | VP14061 | 2014 | China | Clinical | Asia | O3:K6 | ST3 | PC | Wave-4 | Putative outbreak cluster |
| VP14070 | VP14070 | 2014 | China | Clinical | Asia | O3:K6 | ST3 | PC | Wave-4 | Putative outbreak cluster |
| VP14071 | VP14071 | 2014 | China | Clinical | Asia | O3:K6 | ST3 | PC | Wave-4 | Putative outbreak cluster |
| VP14072 | VP14072 | 2014 | China | Clinical | Asia | O3:K6 | ST3 | PC | Wave-4 | Putative outbreak cluster |
| VP14075 | VP14075 | 2014 | China | Clinical | Asia | O3:K6 | ST3 | PC | Wave-4 | Putative outbreak cluster |
| VP14076 | VP14076 | 2014 | China | Clinical | Asia | O3:K6 | ST3 | PC | Wave-4 | Putative outbreak cluster |
| VP14078 | VP14078 | 2014 | China | Clinical | Asia | O3:K6 | ST3 | PC | Wave-4 | Putative outbreak cluster |
| VP14081 | VP14081 | 2014 | China | Clinical | Asia | O3:K6 | ST3 | PC | Wave-4 | Putative outbreak cluster |
| VP14082 | VP14082 | 2014 | China | Clinical | Asia | O3:K6 | ST3 | PC | Wave-4 | Putative outbreak cluster |
| VP14083 | VP14083 | 2014 | China | Clinical | Asia | O3:K6 | ST3 | PC | Wave-4 | Putative outbreak cluster |

| | | | | | | | | | | |
|---|---|---|---|---|---|---|---|---|---|---|
| VP14085 | VP14085 | 2014 | China | Clinical | Asia | O3:K6 | ST3 | PC | Wave-4 | Putative outbreak cluster |
| VP14086 | VP14086 | 2014 | China | Clinical | Asia | O3:K6 | ST3 | PC | Wave-4 | Putative outbreak cluster |
| VP14087 | VP14087 | 2014 | China | Clinical | Asia | O3:K6 | ST3 | PC | Wave-4 | Putative outbreak cluster |
| VP14089 | VP14089 | 2014 | China | Clinical | Asia | O3:K6 | ST3 | PC | Wave-4 | Putative outbreak cluster |
| VP14090 | VP14090 | 2014 | China | Clinical | Asia | O3:K6 | ST3 | PC | Wave-4 | Putative outbreak cluster |
| VP14091 | VP14091 | 2014 | China | Clinical | Asia | O3:K6 | ST3 | PC | Wave-4 | Putative outbreak cluster |
| VP14095 | VP14095 | 2014 | China | Clinical | Asia | O3:K6 | ST3 | PC | Wave-4 | Putative outbreak cluster |
| VP14096 | VP14096 | 2014 | China | Clinical | Asia | O3:K6 | ST3 | PC | Wave-4 | Putative outbreak cluster |
| VP14099 | VP14099 | 2014 | China | Clinical | Asia | O3:K6 | ST3 | PC | Wave-4 | Putative outbreak cluster |
| VP14102 | VP14102 | 2014 | China | Clinical | Asia | O3:K6 | ST3 | PC | Wave-4 | Putative outbreak cluster |
| VP14105 | VP14105 | 2014 | China | Clinical | Asia | O3:K6 | ST3 | PC | Wave-4 | Putative outbreak cluster |
| VP14108 | VP14108 | 2014 | China | Clinical | Asia | O3:K6 | ST3 | PC | Wave-4 | Putative outbreak cluster |
| VP14109 | VP14109 | 2014 | China | Clinical | Asia | O3:K6 | ST3 | PC | Wave-4 | Putative outbreak cluster |
| VP14110 | VP14110 | 2014 | China | Clinical | Asia | O3:K6 | ST3 | PC | Wave-4 | Putative outbreak cluster |
| VP14111 | VP14111 | 2014 | China | Clinical | Asia | O3:K6 | ST3 | PC | Wave-4 | Putative outbreak cluster |
| VP14112 | VP14112 | 2014 | China | Clinical | Asia | O3:K6 | ST3 | PC | Wave-4 | Putative outbreak cluster |
| VP14114 | VP14114 | 2014 | China | Clinical | Asia | O3:K6 | ST3 | PC | Wave-4 | Putative outbreak cluster |
| VP14115 | VP14115 | 2014 | China | Clinical | Asia | O3:K6 | ST3 | PC | Wave-4 | Putative outbreak cluster |
| VP14117 | VP14117 | 2014 | China | Clinical | Asia | O3:K6 | ST3 | PC | Wave-4 | Putative outbreak cluster |
| VP14119 | VP14119 | 2014 | China | Clinical | Asia | O3:K6 | ST3 | PC | Wave-4 | Putative outbreak cluster |
| VP14122 | VP14122 | 2014 | China | Clinical | Asia | O3:K6 | ST3 | PC | Wave-4 | Putative outbreak cluster |
| VP14124 | VP14124 | 2014 | China | Clinical | Asia | O3:K6 | ST3 | PC | Wave-4 | Putative outbreak cluster |
| VP14133 | VP14133 | 2014 | China | Clinical | Asia | O3:K6 | ST3 | PC | Wave-4 | Putative outbreak cluster |
| VP14145 | VP14145 | 2014 | China | Clinical | Asia | O3:K6 | ST3 | PC | Wave-4 | Putative outbreak cluster |
| VP14148 | VP14148 | 2014 | China | Clinical | Asia | O3:K6 | ST3 | PC | Wave-4 | Putative outbreak cluster |
| VP14150 | VP14150 | 2014 | China | Clinical | Asia | O3:K6 | ST3 | PC | Wave-4 | Putative outbreak cluster |
| VP14153 | VP14153 | 2014 | China | Clinical | Asia | O3:K6 | ST3 | PC | Wave-4 | Putative outbreak cluster |
| VP14156 | VP14156 | 2014 | China | Clinical | Asia | O3:K6 | ST3 | PC | Wave-4 | Putative outbreak cluster |
| VP14161 | VP14161 | 2014 | China | Clinical | Asia | O3:K6 | ST3 | PC | Wave-4 | Putative outbreak cluster |
| VP14162 | VP14162 | 2014 | China | Clinical | Asia | O3:K6 | ST3 | PC | Wave-4 | Putative outbreak cluster |
| VP14163 | VP14163 | 2014 | China | Clinical | Asia | O3:K6 | ST3 | PC | Wave-4 | Putative outbreak cluster |
| VP14165 | VP14165 | 2014 | China | Clinical | Asia | O3:K6 | ST3 | PC | Wave-4 | Putative outbreak cluster |

| | | | | | | | | | | |
|---|---|---|---|---|---|---|---|---|---|---|
| VP14166 | VP14166 | 2014 | China | Clinical | Asia | O3:K6 | ST3 | PC | Wave-4 | Putative outbreak cluster |
| VP14167 | VP14167 | 2014 | China | Clinical | Asia | O3:K6 | ST3 | PC | Wave-4 | Putative outbreak cluster |
| VP14168 | VP14168 | 2014 | China | Clinical | Asia | O3:K6 | ST3 | PC | Wave-4 | Putative outbreak cluster |
| VP14172 | VP14172 | 2014 | China | Clinical | Asia | O3:K6 | ST3 | PC | Wave-4 | Putative outbreak cluster |
| VP14173 | VP14173 | 2014 | China | Clinical | Asia | O3:K6 | ST3 | PC | Wave-4 | Putative outbreak cluster |
| VP15002 | VP15002 | 2015 | China | Clinical | Asia | O10:K60 | ST3 | PC | Wave-4 | Putative outbreak cluster |
| VP15014 | VP15014 | 2015 | China | Clinical | Asia | O3:K6 | ST3 | PC | Wave-4 | Putative outbreak cluster |
| VP15019 | VP15019 | 2015 | China | Clinical | Asia | O3:K6 | ST3 | PC | Wave-4 | Putative outbreak cluster |
| VP15020 | VP15020 | 2015 | China | Clinical | Asia | O3:K6 | ST3 | PC | Wave-4 | Putative outbreak cluster |
| VP15021 | VP15021 | 2015 | China | Clinical | Asia | O3:K6 | ST3 | PC | Wave-4 | Putative outbreak cluster |
| VP15022 | VP15022 | 2015 | China | Clinical | Asia | O3:K6 | ST3 | PC | Wave-4 | Putative outbreak cluster |
| VP15026 | VP15026 | 2015 | China | Clinical | Asia | O3:K6 | ST3 | PC | Wave-4 | Putative outbreak cluster |
| VP15027 | VP15027 | 2015 | China | Clinical | Asia | O3:K6 | ST3 | PC | Wave-4 | Putative outbreak cluster |
| VP15030 | VP15030 | 2015 | China | Clinical | Asia | O3:K6 | ST3 | PC | Wave-4 | Putative outbreak cluster |
| VP15032 | VP15032 | 2015 | China | Clinical | Asia | O3:K6 | ST3 | PC | Wave-4 | Putative outbreak cluster |
| VP15033 | VP15033 | 2015 | China | Clinical | Asia | O3:K6 | ST3 | PC | Wave-4 | Putative outbreak cluster |
| VP15034 | VP15034 | 2015 | China | Clinical | Asia | O3:K6 | ST3 | PC | Wave-4 | Putative outbreak cluster |
| VP15035 | VP15035 | 2015 | China | Clinical | Asia | O3:K6 | ST3 | PC | Wave-4 | Putative outbreak cluster |
| VP15036 | VP15036 | 2015 | China | Clinical | Asia | O3:K6 | ST3 | PC | Wave-4 | Putative outbreak cluster |
| VP15037 | VP15037 | 2015 | China | Clinical | Asia | O3:K6 | ST3 | PC | Wave-4 | Putative outbreak cluster |
| VP15038 | VP15038 | 2015 | China | Clinical | Asia | O3:K6 | ST3 | PC | Wave-4 | Putative outbreak cluster |
| VP15040 | VP15040 | 2015 | China | Clinical | Asia | O3:K6 | ST3 | PC | Wave-4 | Putative outbreak cluster |
| VP15043 | VP15043 | 2015 | China | Clinical | Asia | O3:K6 | ST3 | PC | Wave-4 | Putative outbreak cluster |
| VP15048 | VP15048 | 2015 | China | Clinical | Asia | O3:K6 | ST3 | PC | Wave-4 | Putative outbreak cluster |
| VP15053 | VP15053 | 2015 | China | Clinical | Asia | O3:K6 | ST3 | PC | Wave-4 | Putative outbreak cluster |
| VP15054 | VP15054 | 2015 | China | Clinical | Asia | O3:K6 | ST3 | PC | Wave-4 | Putative outbreak cluster |
| VP15055 | VP15055 | 2015 | China | Clinical | Asia | O3:K6 | ST3 | PC | Wave-4 | Putative outbreak cluster |
| VP15056 | VP15056 | 2015 | China | Clinical | Asia | O3:K6 | ST3 | PC | Wave-4 | Putative outbreak cluster |
| VP15058 | VP15058 | 2015 | China | Clinical | Asia | O3:K6 | ST3 | PC | Wave-4 | Putative outbreak cluster |
| VP15059 | VP15059 | 2015 | China | Clinical | Asia | O3:K6 | ST3 | PC | Wave-4 | Putative outbreak cluster |
| VP15060 | VP15060 | 2015 | China | Clinical | Asia | O3:K6 | ST3 | PC | Wave-4 | Putative outbreak cluster |
| VP15061 | VP15061 | 2015 | China | Clinical | Asia | O3:K6 | ST3 | PC | Wave-4 | Putative outbreak cluster |

| | | | | | | | | | | |
|---|---|---|---|---|---|---|---|---|---|---|
| VP15063 | VP15063 | 2015 | China | Clinical | Asia | O3:K6 | ST3 | PC | Wave-4 | Putative outbreak cluster |
| VP15064 | VP15064 | 2015 | China | Clinical | Asia | O3:K6 | ST3 | PC | Wave-4 | Putative outbreak cluster |
| VP15065 | VP15065 | 2015 | China | Clinical | Asia | O3:K6 | ST3 | PC | Wave-4 | Putative outbreak cluster |
| VP15066 | VP15066 | 2015 | China | Clinical | Asia | O3:K6 | ST3 | PC | Wave-4 | Putative outbreak cluster |
| VP15067 | VP15067 | 2015 | China | Clinical | Asia | O3:K6 | ST3 | PC | Wave-4 | Putative outbreak cluster |
| VP15068 | VP15068 | 2015 | China | Clinical | Asia | O3:K6 | ST3 | PC | Wave-4 | Putative outbreak cluster |
| VP15069 | VP15069 | 2015 | China | Clinical | Asia | O3:K6 | ST3 | PC | Wave-4 | Putative outbreak cluster |
| VP15070 | VP15070 | 2015 | China | Clinical | Asia | O3:K6 | ST3 | PC | Wave-4 | Putative outbreak cluster |
| VP15071 | VP15071 | 2015 | China | Clinical | Asia | O3:K6 | ST3 | PC | Wave-4 | Putative outbreak cluster |
| VP15073 | VP15073 | 2015 | China | Clinical | Asia | O3:K6 | ST3 | PC | Wave-4 | Putative outbreak cluster |
| VP15076 | VP15076 | 2015 | China | Clinical | Asia | O3:K6 | ST3 | PC | Wave-4 | Putative outbreak cluster |
| VP15079 | VP15079 | 2015 | China | Clinical | Asia | O3:K6 | ST3 | PC | Wave-4 | Putative outbreak cluster |
| VP15080 | VP15080 | 2015 | China | Clinical | Asia | O3:K6 | ST3 | PC | Wave-4 | Putative outbreak cluster |
| VP15081 | VP15081 | 2015 | China | Clinical | Asia | O3:K6 | ST3 | PC | Wave-4 | Putative outbreak cluster |
| VP15082 | VP15082 | 2015 | China | Clinical | Asia | O3:K6 | ST3 | PC | Wave-4 | Putative outbreak cluster |
| VP15083 | VP15083 | 2015 | China | Clinical | Asia | O3:K6 | ST3 | PC | Wave-4 | Putative outbreak cluster |
| VP15084 | VP15084 | 2015 | China | Clinical | Asia | O3:K6 | ST3 | PC | Wave-4 | Putative outbreak cluster |
| VP15085 | VP15085 | 2015 | China | Clinical | Asia | O10:K60 | ST3 | PC | Wave-4 | Putative outbreak cluster |
| VP15086 | VP15086 | 2015 | China | Clinical | Asia | O3:K6 | ST3 | PC | Wave-4 | Putative outbreak cluster |
| VP15087 | VP15087 | 2015 | China | Clinical | Asia | O3:K6 | ST3 | PC | Wave-4 | Putative outbreak cluster |
| VP15088 | VP15088 | 2015 | China | Clinical | Asia | O3:K6 | ST3 | PC | Wave-4 | Putative outbreak cluster |
| VP15090 | VP15090 | 2015 | China | Clinical | Asia | O3:K6 | ST3 | PC | Wave-4 | Putative outbreak cluster |
| VP15091 | VP15091 | 2015 | China | Clinical | Asia | O3:K6 | ST3 | PC | Wave-4 | Putative outbreak cluster |
| VP15092 | VP15092 | 2015 | China | Clinical | Asia | O3:K6 | ST3 | PC | Wave-4 | Putative outbreak cluster |
| VP15093 | VP15093 | 2015 | China | Clinical | Asia | O3:K6 | ST3 | PC | Wave-4 | Putative outbreak cluster |
| VP15095 | VP15095 | 2015 | China | Clinical | Asia | O3:K6 | ST3 | PC | Wave-4 | Putative outbreak cluster |
| VP15107 | VP15107 | 2015 | China | Clinical | Asia | O3:K6 | ST3 | PC | Wave-4 | Putative outbreak cluster |
| VP15108 | VP15108 | 2015 | China | Clinical | Asia | O3:K6 | ST3 | PC | Wave-4 | Putative outbreak cluster |
| VP15109 | VP15109 | 2015 | China | Clinical | Asia | O3:K6 | ST3 | PC | Wave-4 | Putative outbreak cluster |
| VP15110 | VP15110 | 2015 | China | Clinical | Asia | O3:K6 | ST3 | PC | Wave-4 | Putative outbreak cluster |
| VP15111 | VP15111 | 2015 | China | Clinical | Asia | O3:K6 | ST3 | PC | Wave-4 | Putative outbreak cluster |
| VP15112 | VP15112 | 2015 | China | Clinical | Asia | O3:K6 | ST3 | PC | Wave-4 | Putative outbreak cluster |

| | | | | | | | | | | |
|---|---|---|---|---|---|---|---|---|---|---|
| VP15121 | VP15121 | 2015 | China | Clinical | Asia | O3:K6 | ST3 | PC | Wave-4 | Putative outbreak cluster |
| VP15122 | VP15122 | 2015 | China | Clinical | Asia | O3:K6 | ST3 | PC | Wave-4 | Putative outbreak cluster |
| VP15123 | VP15123 | 2015 | China | Clinical | Asia | O3:K6 | ST3 | PC | Wave-4 | Putative outbreak cluster |
| VP15126 | VP15126 | 2015 | China | Clinical | Asia | O3:K6 | ST3 | PC | Wave-4 | Putative outbreak cluster |
| VP15128 | VP15128 | 2015 | China | Clinical | Asia | O3:K6 | ST3 | PC | Wave-4 | Putative outbreak cluster |
| VP15129 | VP15129 | 2015 | China | Clinical | Asia | O3:K6 | ST3 | PC | Wave-4 | Putative outbreak cluster |
| VP15131 | VP15131 | 2015 | China | Clinical | Asia | O3:K6 | ST3 | PC | Wave-4 | Putative outbreak cluster |
| VP15133 | VP15133 | 2015 | China | Clinical | Asia | O3:K6 | ST3 | PC | Wave-4 | Putative outbreak cluster |
| VP15134 | VP15134 | 2015 | China | Clinical | Asia | O3:K6 | ST3 | PC | Wave-4 | Putative outbreak cluster |
| VP15135 | VP15135 | 2015 | China | Clinical | Asia | O3:K6 | ST3 | PC | Wave-4 | Putative outbreak cluster |
| VP15137 | VP15137 | 2015 | China | Clinical | Asia | O3:K6 | ST3 | PC | Wave-4 | Putative outbreak cluster |
| VP15138 | VP15138 | 2015 | China | Clinical | Asia | O3:K6 | ST3 | PC | Wave-4 | Putative outbreak cluster |
| VP15139 | VP15139 | 2015 | China | Clinical | Asia | O3:K6 | ST3 | PC | Wave-4 | Putative outbreak cluster |
| VP15140 | VP15140 | 2015 | China | Clinical | Asia | O3:K6 | ST3 | PC | Wave-4 | Putative outbreak cluster |
| VP15142 | VP15142 | 2015 | China | Clinical | Asia | O3:K6 | ST- | PC | Wave-4 | Putative outbreak cluster |
| VP15145 | VP15145 | 2015 | China | Clinical | Asia | O10:K60 | ST3 | PC | Wave-4 | Putative outbreak cluster |
| VP15146 | VP15146 | 2015 | China | Clinical | Asia | O3:K6 | ST3 | PC | Wave-4 | Putative outbreak cluster |
| VP15148 | VP15148 | 2015 | China | Clinical | Asia | O3:K6 | ST3 | PC | Wave-4 | Putative outbreak cluster |
| VP15149 | VP15149 | 2015 | China | Clinical | Asia | O3:K6 | ST3 | PC | Wave-4 | Putative outbreak cluster |
| VP15150 | VP15150 | 2015 | China | Clinical | Asia | O3:K6 | ST3 | PC | Wave-4 | Putative outbreak cluster |
| VP15151 | VP15151 | 2015 | China | Clinical | Asia | O3:K6 | ST3 | PC | Wave-4 | Putative outbreak cluster |
| VP15152 | VP15152 | 2015 | China | Clinical | Asia | O3:K6 | ST3 | PC | Wave-4 | Putative outbreak cluster |
| VP15153 | VP15153 | 2015 | China | Clinical | Asia | O3:K6 | ST3 | PC | Wave-4 | Putative outbreak cluster |
| VP15156 | VP15156 | 2015 | China | Clinical | Asia | O3:K6 | ST3 | PC | Wave-4 | Putative outbreak cluster |
| VP15162 | VP15162 | 2015 | China | Clinical | Asia | O3:K6 | ST3 | PC | Wave-4 | Putative outbreak cluster |
| VP15163 | VP15163 | 2015 | China | Clinical | Asia | O3:K6 | ST3 | PC | Wave-4 | Putative outbreak cluster |
| VP15167 | VP15167 | 2015 | China | Clinical | Asia | O3:K6 | ST3 | PC | Wave-4 | Putative outbreak cluster |
| VP15168 | VP15168 | 2015 | China | Clinical | Asia | O3:K6 | ST3 | PC | Wave-4 | Putative outbreak cluster |
| VP15169 | VP15169 | 2015 | China | Clinical | Asia | O3:K6 | ST3 | PC | Wave-4 | Putative outbreak cluster |
| VP15170 | VP15170 | 2015 | China | Clinical | Asia | O3:K6 | ST3 | PC | Wave-4 | Putative outbreak cluster |
| VP15172 | VP15172 | 2015 | China | Clinical | Asia | O3:K6 | ST3 | PC | Wave-4 | Putative outbreak cluster |
| VP15173 | VP15173 | 2015 | China | Clinical | Asia | O3:K6 | ST3 | PC | Wave-4 | Putative outbreak cluster |

| | | | | | | | | | | |
|---|---|---|---|---|---|---|---|---|---|---|
| VP15174 | VP15174 | 2015 | China | Clinical | Asia | O3:K6 | ST3 | PC | Wave-4 | Putative outbreak cluster |
| VP15175 | VP15175 | 2015 | China | Clinical | Asia | O3:K6 | ST3 | PC | Wave-4 | Putative outbreak cluster |
| VP15178 | VP15178 | 2015 | China | Clinical | Asia | O3:K6 | ST3 | PC | Wave-4 | Putative outbreak cluster |
| VP15179 | VP15179 | 2015 | China | Clinical | Asia | O3:K6 | ST3 | PC | Wave-4 | Putative outbreak cluster |
| VP15180 | VP15180 | 2015 | China | Clinical | Asia | O3:K6 | ST3 | PC | Wave-4 | Putative outbreak cluster |
| VP15183 | VP15183 | 2015 | China | Clinical | Asia | O3:K6 | ST3 | PC | Wave-4 | Putative outbreak cluster |
| VP15185 | VP15185 | 2015 | China | Clinical | Asia | O3:K6 | ST3 | PC | Wave-4 | Putative outbreak cluster |
| VP15186 | VP15186 | 2015 | China | Clinical | Asia | O3:K6 | ST3 | PC | Wave-4 | Putative outbreak cluster |
| VP15187 | VP15187 | 2015 | China | Clinical | Asia | O3:K6 | ST3 | PC | Wave-4 | Putative outbreak cluster |
| VP15190 | VP15190 | 2015 | China | Clinical | Asia | O3:K6 | ST3 | PC | Wave-4 | Putative outbreak cluster |
| VP15192 | VP15192 | 2015 | China | Clinical | Asia | O3:K6 | ST3 | PC | Wave-4 | Putative outbreak cluster |
| VP15193 | VP15193 | 2015 | China | Clinical | Asia | O3:K6 | ST3 | PC | Wave-4 | Putative outbreak cluster |
| VP15195 | VP15195 | 2015 | China | Clinical | Asia | O3:K6 | ST3 | PC | Wave-4 | Putative outbreak cluster |
| VP16 | VP16 | NA | China | Env | Asia | O3:K6 | ST3 | PC | Wave-4 | Putative outbreak cluster |
| VP16005 | VP16005 | 2016 | China | Clinical | Asia | O3:K6 | ST3 | PC | Wave-4 | Putative outbreak cluster |
| VP16013 | VP16013 | 2016 | China | Clinical | Asia | O3:K6 | ST3 | PC | Wave-4 | Putative outbreak cluster |
| VP16014 | VP16014 | 2016 | China | Clinical | Asia | O3:K6 | ST3 | PC | Wave-4 | Putative outbreak cluster |
| VP16015 | VP16015 | 2016 | China | Clinical | Asia | O3:K6 | ST3 | PC | Wave-4 | Putative outbreak cluster |
| VP16016 | VP16016 | 2016 | China | Clinical | Asia | O3:K6 | ST3 | PC | Wave-4 | Putative outbreak cluster |
| VP16017 | VP16017 | 2016 | China | Clinical | Asia | O3:K6 | ST3 | PC | Wave-4 | Putative outbreak cluster |
| VP16026 | VP16026 | 2016 | China | Clinical | Asia | O3:K6 | ST3 | PC | Wave-4 | Putative outbreak cluster |
| VP16027 | VP16027 | 2016 | China | Clinical | Asia | O3:K6 | ST3 | PC | Wave-4 | Putative outbreak cluster |
| VP16030 | VP16030 | 2016 | China | Clinical | Asia | O3:K6 | ST3 | PC | Wave-4 | Putative outbreak cluster |
| VP16031 | VP16031 | 2016 | China | Clinical | Asia | O3:K6 | ST3 | PC | Wave-4 | Putative outbreak cluster |
| VP16033 | VP16033 | 2016 | China | Clinical | Asia | O3:K6 | ST3 | PC | Wave-4 | Putative outbreak cluster |
| VP16042 | VP16042 | 2016 | China | Clinical | Asia | O3:K6 | ST3 | PC | Wave-4 | Putative outbreak cluster |
| VP16044 | VP16044 | 2016 | China | Clinical | Asia | O3:K6 | ST3 | PC | Wave-4 | Putative outbreak cluster |
| VP16046 | VP16046 | 2016 | China | Clinical | Asia | O3:K6 | ST3 | PC | Wave-4 | Putative outbreak cluster |
| VP16050 | VP16050 | 2016 | China | Clinical | Asia | O3:K6 | ST3 | PC | Wave-4 | Putative outbreak cluster |
| VP16053 | VP16053 | 2016 | China | Clinical | Asia | O10:K60 | ST3 | PC | Wave-4 | Putative outbreak cluster |
| VP16055 | VP16055 | 2016 | China | Clinical | Asia | O3:K6 | ST3 | PC | Wave-4 | Putative outbreak cluster |
| VP16056 | VP16056 | 2016 | China | Clinical | Asia | O3:K6 | ST3 | PC | Wave-4 | Putative outbreak cluster |

| | | | | | | | | | | |
|---|---|---|---|---|---|---|---|---|---|---|
| VP16057 | VP16057 | 2016 | China | Clinical | Asia | O3:K6 | ST3 | PC | Wave-4 | Putative outbreak cluster |
| VP16059 | VP16059 | 2016 | China | Clinical | Asia | O3:K6 | ST3 | PC | Wave-4 | Putative outbreak cluster |
| VP16061 | VP16061 | 2016 | China | Clinical | Asia | O3:K6 | ST3 | PC | Wave-4 | Putative outbreak cluster |
| VP16062 | VP16062 | 2016 | China | Clinical | Asia | O3:K6 | ST3 | PC | Wave-4 | Putative outbreak cluster |
| VP16067 | VP16067 | 2016 | China | Clinical | Asia | O3:K6 | ST3 | PC | Wave-4 | Putative outbreak cluster |
| VP16069 | VP16069 | 2016 | China | Clinical | Asia | O3:K6 | ST3 | PC | Wave-4 | Putative outbreak cluster |
| VP16070 | VP16070 | 2016 | China | Clinical | Asia | O3:K6 | ST3 | PC | Wave-4 | Putative outbreak cluster |
| VP16071 | VP16071 | 2016 | China | Clinical | Asia | O3:K6 | ST3 | PC | Wave-4 | Putative outbreak cluster |
| VP16074 | VP16074 | 2016 | China | Clinical | Asia | O3:K6 | ST3 | PC | Wave-4 | Putative outbreak cluster |
| VP16075 | VP16075 | 2016 | China | Clinical | Asia | O3:K6 | ST3 | PC | Wave-4 | Putative outbreak cluster |
| VP16092 | VP16092 | 2016 | China | Clinical | Asia | O3:K6 | ST3 | PC | Wave-4 | Putative outbreak cluster |
| VP16099 | VP16099 | 2016 | China | Clinical | Asia | O3:K6 | ST3 | PC | Wave-4 | Putative outbreak cluster |
| VP16100 | VP16100 | 2016 | China | Clinical | Asia | O3:K6 | ST3 | PC | Wave-4 | Putative outbreak cluster |
| VP16101 | VP16101 | 2016 | China | Clinical | Asia | O3:K6 | ST3 | PC | Wave-4 | Putative outbreak cluster |
| VP16102 | VP16102 | 2016 | China | Clinical | Asia | O3:K6 | ST3 | PC | Wave-4 | Putative outbreak cluster |
| VP16104 | VP16104 | 2016 | China | Clinical | Asia | O3:K6 | ST3 | PC | Wave-4 | Putative outbreak cluster |
| VP16116 | VP16116 | 2016 | China | Clinical | Asia | O3:K6 | ST3 | PC | Wave-4 | Putative outbreak cluster |
| VP16117 | VP16117 | 2016 | China | Clinical | Asia | O3:K6 | ST3 | PC | Wave-4 | Putative outbreak cluster |
| VP16119 | VP16119 | 2016 | China | Clinical | Asia | O3:K6 | ST3 | PC | Wave-4 | Putative outbreak cluster |
| VP16132 | VP16132 | 2016 | China | Clinical | Asia | O3:K6 | ST3 | PC | Wave-4 | Putative outbreak cluster |
| VP16133 | VP16133 | 2016 | China | Clinical | Asia | O3:K6 | ST3 | PC | Wave-4 | Putative outbreak cluster |
| VP16137 | VP16137 | 2016 | China | Clinical | Asia | O3:K6 | ST3 | PC | Wave-4 | Putative outbreak cluster |
| VP16141 | VP16141 | 2016 | China | Clinical | Asia | O3:K6 | ST3 | PC | Wave-4 | Putative outbreak cluster |
| VP16144 | VP16144 | 2016 | China | Clinical | Asia | O3:K6 | ST3 | PC | Wave-4 | Putative outbreak cluster |
| VP16145 | VP16145 | 2016 | China | Clinical | Asia | O3:K6 | ST3 | PC | Wave-4 | Putative outbreak cluster |
| VP16146 | VP16146 | 2016 | China | Clinical | Asia | O3:K6 | ST3 | PC | Wave-4 | Putative outbreak cluster |
| VP16147 | VP16147 | 2016 | China | Clinical | Asia | O3:K6 | ST3 | PC | Wave-4 | Putative outbreak cluster |
| VP16148 | VP16148 | 2016 | China | Clinical | Asia | O3:K6 | ST3 | PC | Wave-4 | Putative outbreak cluster |
| VP16149 | VP16149 | 2016 | China | Clinical | Asia | O3:K6 | ST3 | PC | Wave-4 | Putative outbreak cluster |
| VP16150 | VP16150 | 2016 | China | Clinical | Asia | O3:K6 | ST3 | PC | Wave-4 | Putative outbreak cluster |
| VP16151 | VP16151 | 2016 | China | Clinical | Asia | O3:K6 | ST3 | PC | Wave-4 | Putative outbreak cluster |
| VP16152 | VP16152 | 2016 | China | Clinical | Asia | O3:K6 | ST3 | PC | Wave-4 | Putative outbreak cluster |

| | | | | | | | | | | |
|---|---|---|---|---|---|---|---|---|---|---|
| VP16153 | VP16153 | 2016 | China | Clinical | Asia | O3:K6 | ST3 | PC | Wave-4 | Putative outbreak cluster |
| VP16154 | VP16154 | 2016 | China | Clinical | Asia | O3:K6 | ST3 | PC | Wave-4 | Putative outbreak cluster |
| VP16168 | VP16168 | 2016 | China | Clinical | Asia | O3:K6 | ST3 | PC | Wave-4 | Putative outbreak cluster |
| VP16170 | VP16170 | 2016 | China | Clinical | Asia | O3:K6 | ST3 | PC | Wave-4 | Putative outbreak cluster |
| VP16177 | VP16177 | 2016 | China | Clinical | Asia | O3:K6 | ST3 | PC | Wave-4 | Putative outbreak cluster |
| VP16178 | VP16178 | 2016 | China | Clinical | Asia | O3:K6 | ST3 | PC | Wave-4 | Putative outbreak cluster |
| VP16180 | VP16180 | 2016 | China | Clinical | Asia | O3:K6 | ST3 | PC | Wave-4 | Putative outbreak cluster |
| VP16181 | VP16181 | 2016 | China | Clinical | Asia | O3:K6 | ST3 | PC | Wave-4 | Putative outbreak cluster |
| VP16183 | VP16183 | 2016 | China | Clinical | Asia | O3:K6 | ST3 | PC | Wave-4 | Putative outbreak cluster |
| VP16187 | VP16187 | 2016 | China | Clinical | Asia | O3:K6 | ST3 | PC | Wave-4 | Putative outbreak cluster |
| VP16188 | VP16188 | 2016 | China | Clinical | Asia | O3:K6 | ST3 | PC | Wave-4 | Putative outbreak cluster |
| VP16189 | VP16189 | 2016 | China | Clinical | Asia | O3:K6 | ST3 | PC | Wave-4 | Putative outbreak cluster |
| VP16191 | VP16191 | 2016 | China | Clinical | Asia | O3:K6 | ST3 | PC | Wave-4 | Putative outbreak cluster |
| VP16193 | VP16193 | 2016 | China | Clinical | Asia | O3:K6 | ST3 | PC | Wave-4 | Putative outbreak cluster |
| VP16194 | VP16194 | 2016 | China | Clinical | Asia | O3:K6 | ST3 | PC | Wave-4 | Putative outbreak cluster |
| VP16197 | VP16197 | 2016 | China | Clinical | Asia | O3:K6 | ST3 | PC | Wave-4 | Putative outbreak cluster |
| VP16211 | VP16211 | 2016 | China | Clinical | Asia | O3:K6 | ST3 | PC | Wave-4 | Putative outbreak cluster |
| VP16212 | VP16212 | 2016 | China | Clinical | Asia | O3:K6 | ST3 | PC | Wave-4 | Putative outbreak cluster |
| VP16213 | VP16213 | 2016 | China | Clinical | Asia | O3:K6 | ST3 | PC | Wave-4 | Putative outbreak cluster |
| VP17010 | VP17010 | 2017 | China | Clinical | Asia | O3:K6 | ST3 | PC | Wave-4 | Putative outbreak cluster |
| VP17011 | VP17011 | 2017 | China | Clinical | Asia | O3:K6 | ST3 | PC | Wave-4 | Putative outbreak cluster |
| VP17016 | VP17016 | 2017 | China | Clinical | Asia | O3:K6 | ST3 | PC | Wave-4 | Putative outbreak cluster |
| VP17018 | VP17018 | 2017 | China | Clinical | Asia | O3:K6 | ST3 | PC | Wave-4 | Putative outbreak cluster |
| VP17023 | VP17023 | 2017 | China | Clinical | Asia | O3:K6 | ST3 | PC | Wave-4 | Putative outbreak cluster |
| VP17028 | VP17028 | 2017 | China | Clinical | Asia | O3:K6 | ST3 | PC | Wave-4 | Putative outbreak cluster |
| VP17029 | VP17029 | 2017 | China | Clinical | Asia | O3:K6 | ST3 | PC | Wave-4 | Putative outbreak cluster |
| VP17031 | VP17031 | 2017 | China | Clinical | Asia | O3:K6 | ST3 | PC | Wave-4 | Putative outbreak cluster |
| VP17032 | VP17032 | 2017 | China | Clinical | Asia | O3:K6 | ST3 | PC | Wave-4 | Putative outbreak cluster |
| VP17033 | VP17033 | 2017 | China | Clinical | Asia | O3:K6 | ST3 | PC | Wave-4 | Putative outbreak cluster |
| VP17035 | VP17035 | 2017 | China | Clinical | Asia | O3:K6 | ST3 | PC | Wave-4 | Putative outbreak cluster |
| VP17036 | VP17036 | 2017 | China | Clinical | Asia | O3:K6 | ST3 | PC | Wave-4 | Putative outbreak cluster |
| VP17037 | VP17037 | 2017 | China | Clinical | Asia | O3:K6 | ST3 | PC | Wave-4 | Putative outbreak cluster |

| | | | | | | | | | | |
|---|---|---|---|---|---|---|---|---|---|---|
| VP17038 | VP17038 | 2017 | China | Clinical | Asia | O3:K6 | ST3 | PC | Wave-4 | Putative outbreak cluster |
| VP17039 | VP17039 | 2017 | China | Clinical | Asia | O3:K6 | ST3 | PC | Wave-4 | Putative outbreak cluster |
| VP17040 | VP17040 | 2017 | China | Clinical | Asia | O3:K6 | ST3 | PC | Wave-4 | Putative outbreak cluster |
| VP17041 | VP17041 | 2017 | China | Clinical | Asia | O3:K6 | ST3 | PC | Wave-4 | Putative outbreak cluster |
| VP17044 | VP17044 | 2017 | China | Clinical | Asia | O3:K6 | ST3 | PC | Wave-4 | Putative outbreak cluster |
| VP17046 | VP17046 | 2017 | China | Clinical | Asia | O3:K6 | ST3 | PC | Wave-4 | Putative outbreak cluster |
| VP17047 | VP17047 | 2017 | China | Clinical | Asia | O3:K6 | ST3 | PC | Wave-4 | Putative outbreak cluster |
| VP17049 | VP17049 | 2017 | China | Clinical | Asia | O3:K6 | ST3 | PC | Wave-4 | Putative outbreak cluster |
| VP17050 | VP17050 | 2017 | China | Clinical | Asia | O3:K6 | ST3 | PC | Wave-4 | Putative outbreak cluster |
| VP17052 | VP17052 | 2017 | China | Clinical | Asia | O3:K6 | ST3 | PC | Wave-4 | Putative outbreak cluster |
| VP17053 | VP17053 | 2017 | China | Clinical | Asia | O3:K6 | ST3 | PC | Wave-4 | Putative outbreak cluster |
| VP17055 | VP17055 | 2017 | China | Clinical | Asia | O3:K6 | ST3 | PC | Wave-4 | Putative outbreak cluster |
| VP17057 | VP17057 | 2017 | China | Clinical | Asia | O3:K6 | ST3 | PC | Wave-4 | Putative outbreak cluster |
| VP17060 | VP17060 | 2017 | China | Clinical | Asia | O3:K6 | ST3 | PC | Wave-4 | Putative outbreak cluster |
| VP17062 | VP17062 | 2017 | China | Clinical | Asia | O3:K6 | ST3 | PC | Wave-4 | Putative outbreak cluster |
| VP17063 | VP17063 | 2017 | China | Clinical | Asia | O3:K6 | ST3 | PC | Wave-4 | Putative outbreak cluster |
| VP17064 | VP17064 | 2017 | China | Clinical | Asia | O3:K6 | ST3 | PC | Wave-4 | Putative outbreak cluster |
| VP17071 | VP17071 | 2017 | China | Clinical | Asia | O3:K6 | ST3 | PC | Wave-4 | Putative outbreak cluster |
| VP17074 | VP17074 | 2017 | China | Clinical | Asia | O3:K6 | ST3 | PC | Wave-4 | Putative outbreak cluster |
| VP17076 | VP17076 | 2017 | China | Clinical | Asia | O3:K6 | ST3 | PC | Wave-4 | Putative outbreak cluster |
| VP17077 | VP17077 | 2017 | China | Clinical | Asia | O3:K6 | ST3 | PC | Wave-4 | Putative outbreak cluster |
| VP17078 | VP17078 | 2017 | China | Clinical | Asia | O3:K6 | ST3 | PC | Wave-4 | Putative outbreak cluster |
| VP17083 | VP17083 | 2017 | China | Clinical | Asia | O3:K6 | ST3 | PC | Wave-4 | Putative outbreak cluster |
| VP17086 | VP17086 | 2017 | China | Clinical | Asia | O3:K6 | ST3 | PC | Wave-4 | Putative outbreak cluster |
| VP17087 | VP17087 | 2017 | China | Clinical | Asia | O3:K6 | ST3 | PC | Wave-4 | Putative outbreak cluster |
| VP17088 | VP17088 | 2017 | China | Clinical | Asia | O3:K6 | ST3 | PC | Wave-4 | Putative outbreak cluster |
| VP17089 | VP17089 | 2017 | China | Clinical | Asia | O3:K6 | ST3 | PC | Wave-4 | Putative outbreak cluster |
| VP17097 | VP17097 | 2017 | China | Clinical | Asia | O3:K6 | ST3 | PC | Wave-4 | Putative outbreak cluster |
| VP17098 | VP17098 | 2017 | China | Clinical | Asia | O3:K6 | ST3 | PC | Wave-4 | Putative outbreak cluster |
| VP17100 | VP17100 | 2017 | China | Clinical | Asia | O3:K6 | ST3 | PC | Wave-4 | Putative outbreak cluster |
| VP17102 | VP17102 | 2017 | China | Clinical | Asia | O3:K6 | ST3 | PC | Wave-4 | Putative outbreak cluster |
| VP17103 | VP17103 | 2017 | China | Clinical | Asia | O3:K6 | ST3 | PC | Wave-4 | Putative outbreak cluster |

| | | | | | | | | | | |
|---|---|---|---|---|---|---|---|---|---|---|
| VP17105 | VP17105 | 2017 | China | Clinical | Asia | O3:K6 | ST3 | PC | Wave-4 | Putative outbreak cluster |
| VP17108 | VP17108 | 2017 | China | Clinical | Asia | O3:K6 | ST3 | PC | Wave-4 | Putative outbreak cluster |
| VP17112 | VP17112 | 2017 | China | Clinical | Asia | O3:K6 | ST3 | PC | Wave-4 | Putative outbreak cluster |
| VP17116 | VP17116 | 2017 | China | Clinical | Asia | O3:K6 | ST3 | PC | Wave-4 | Putative outbreak cluster |
| VP17117 | VP17117 | 2017 | China | Clinical | Asia | O3:K6 | ST3 | PC | Wave-4 | Putative outbreak cluster |
| VP17118 | VP17118 | 2017 | China | Clinical | Asia | O3:K6 | ST3 | PC | Wave-4 | Putative outbreak cluster |
| VP17123 | VP17123 | 2017 | China | Clinical | Asia | O3:K6 | ST3 | PC | Wave-4 | Putative outbreak cluster |
| VP17125 | VP17125 | 2017 | China | Clinical | Asia | O3:K6 | ST3 | PC | Wave-4 | Putative outbreak cluster |
| VP17126 | VP17126 | 2017 | China | Clinical | Asia | O3:K6 | ST3 | PC | Wave-4 | Putative outbreak cluster |
| VP17130 | VP17130 | 2017 | China | Clinical | Asia | O3:K6 | ST3 | PC | Wave-4 | Putative outbreak cluster |
| VP17131 | VP17131 | 2017 | China | Clinical | Asia | O3:K6 | ST3 | PC | Wave-4 | Putative outbreak cluster |
| VP17132 | VP17132 | 2017 | China | Clinical | Asia | O3:K6 | ST3 | PC | Wave-4 | Putative outbreak cluster |
| VP17133 | VP17133 | 2017 | China | Clinical | Asia | O3:K6 | ST3 | PC | Wave-4 | Putative outbreak cluster |
| VP17135 | VP17135 | 2017 | China | Clinical | Asia | O3:K6 | ST3 | PC | Wave-4 | Putative outbreak cluster |
| VP17136 | VP17136 | 2017 | China | Clinical | Asia | O3:K6 | ST3 | PC | Wave-4 | Putative outbreak cluster |
| VP17139 | VP17139 | 2017 | China | Clinical | Asia | O3:K6 | ST3 | PC | Wave-4 | Putative outbreak cluster |
| VP17140 | VP17140 | 2017 | China | Clinical | Asia | O3:K6 | ST3 | PC | Wave-4 | Putative outbreak cluster |
| VP17141 | VP17141 | 2017 | China | Clinical | Asia | O3:K6 | ST3 | PC | Wave-4 | Putative outbreak cluster |
| VP17142 | VP17142 | 2017 | China | Clinical | Asia | O3:K6 | ST3 | PC | Wave-4 | Putative outbreak cluster |
| VP17143 | VP17143 | 2017 | China | Clinical | Asia | O3:K6 | ST3 | PC | Wave-4 | Putative outbreak cluster |
| VP17146 | VP17146 | 2017 | China | Clinical | Asia | O3:K6 | ST3 | PC | Wave-4 | Putative outbreak cluster |
| VP17147 | VP17147 | 2017 | China | Clinical | Asia | O3:K6 | ST3 | PC | Wave-4 | Putative outbreak cluster |
| VP17148 | VP17148 | 2017 | China | Clinical | Asia | O3:K6 | ST3 | PC | Wave-4 | Putative outbreak cluster |
| VP17152 | VP17152 | 2017 | China | Clinical | Asia | O3:K6 | ST3 | PC | Wave-4 | Putative outbreak cluster |
| VP17160-2 | VP17160-2 | 2017 | China | Clinical | Asia | O3:K6 | ST3 | PC | Wave-4 | Putative outbreak cluster |
| VP17163 | VP17163 | 2017 | China | Clinical | Asia | O3:K6 | ST3 | PC | Wave-4 | Putative outbreak cluster |
| VP17169 | VP17169 | 2017 | China | Clinical | Asia | O3:K6 | ST3 | PC | Wave-4 | Putative outbreak cluster |
| VP18010 | VP18010 | 2018 | China | Clinical | Asia | O3:K6 | ST3 | PC | Wave-4 | Putative outbreak cluster |
| VP18020 | VP18020 | 2018 | China | Clinical | Asia | O3:K6 | ST3 | PC | Wave-4 | Putative outbreak cluster |
| VP18024 | VP18024 | 2018 | China | Clinical | Asia | O3:K6 | ST3 | PC | Wave-4 | Putative outbreak cluster |
| VP18027 | VP18027 | 2018 | China | Clinical | Asia | O3:K6 | ST3 | PC | Wave-4 | Putative outbreak cluster |
| VP18028 | VP18028 | 2018 | China | Clinical | Asia | O3:K6 | ST3 | PC | Wave-4 | Putative outbreak cluster |

| | | | | | | | | | | |
|---|---|---|---|---|---|---|---|---|---|---|
| VP18029 | VP18029 | 2018 | China | Clinical | Asia | O3:K6 | ST3 | PC | Wave-4 | Putative outbreak cluster |
| VP18030 | VP18030 | 2018 | China | Clinical | Asia | O3:K6 | ST3 | PC | Wave-4 | Putative outbreak cluster |
| VP18031 | VP18031 | 2018 | China | Clinical | Asia | O3:K6 | ST3 | PC | Wave-4 | Putative outbreak cluster |
| VP18033 | VP18033 | 2018 | China | Clinical | Asia | O3:K6 | ST3 | PC | Wave-4 | Putative outbreak cluster |
| VP18035 | VP18035 | 2018 | China | Clinical | Asia | O3:K6 | ST3 | PC | Wave-4 | Putative outbreak cluster |
| VP18036 | VP18036 | 2018 | China | Clinical | Asia | O3:K6 | ST3 | PC | Wave-4 | Putative outbreak cluster |
| VP18037 | VP18037 | 2018 | China | Clinical | Asia | O3:K6 | ST3 | PC | Wave-4 | Putative outbreak cluster |
| VP18038 | VP18038 | 2018 | China | Clinical | Asia | O3:K6 | ST3 | PC | Wave-4 | Putative outbreak cluster |
| VP18043 | VP18043 | 2018 | China | Clinical | Asia | O3:K6 | ST3 | PC | Wave-4 | Putative outbreak cluster |
| VP18044 | VP18044 | 2018 | China | Clinical | Asia | O3:K6 | ST3 | PC | Wave-4 | Putative outbreak cluster |
| VP18047 | VP18047 | 2018 | China | Clinical | Asia | O3:K6 | ST3 | PC | Wave-4 | Putative outbreak cluster |
| VP18048 | VP18048 | 2018 | China | Clinical | Asia | O3:K6 | ST3 | PC | Wave-4 | Putative outbreak cluster |
| VP18049 | VP18049 | 2018 | China | Env | Asia | O3:K6 | ST3 | PC | Wave-4 | Putative outbreak cluster |
| VP18053 | VP18053 | 2018 | China | Clinical | Asia | O3:K6 | ST3 | PC | Wave-4 | Putative outbreak cluster |
| VP18054 | VP18054 | 2018 | China | Clinical | Asia | O3:K6 | ST3 | PC | Wave-4 | Putative outbreak cluster |
| VP18055 | VP18055 | 2018 | China | Clinical | Asia | O3:K6 | ST3 | PC | Wave-4 | Putative outbreak cluster |
| VP18056 | VP18056 | 2018 | China | Clinical | Asia | O3:K6 | ST3 | PC | Wave-4 | Putative outbreak cluster |
| VP18057 | VP18057 | 2018 | China | Clinical | Asia | O3:K6 | ST3 | PC | Wave-4 | Putative outbreak cluster |
| VP18059 | VP18059 | 2018 | China | Clinical | Asia | O3:K6 | ST3 | PC | Wave-4 | Putative outbreak cluster |
| VP18060 | VP18060 | 2018 | China | Clinical | Asia | O3:K6 | ST3 | PC | Wave-4 | Putative outbreak cluster |
| VP18062 | VP18062 | 2018 | China | Clinical | Asia | O3:K6 | ST3 | PC | Wave-4 | Putative outbreak cluster |
| VP18063 | VP18063 | 2018 | China | Clinical | Asia | O3:K6 | ST3 | PC | Wave-4 | Putative outbreak cluster |
| VP18064 | VP18064 | 2018 | China | Clinical | Asia | O3:K6 | ST3 | PC | Wave-4 | Putative outbreak cluster |
| VP18066 | VP18066 | 2018 | China | Clinical | Asia | O3:K6 | ST3 | PC | Wave-4 | Putative outbreak cluster |
| VP18067 | VP18067 | 2018 | China | Clinical | Asia | O3:K6 | ST3 | PC | Wave-4 | Putative outbreak cluster |
| VP18068 | VP18068 | 2018 | China | Clinical | Asia | O3:K6 | ST3 | PC | Wave-4 | Putative outbreak cluster |
| VP18069 | VP18069 | 2018 | China | Clinical | Asia | O3:K6 | ST3 | PC | Wave-4 | Putative outbreak cluster |
| VP18070 | VP18070 | 2018 | China | Clinical | Asia | O3:K6 | ST3 | PC | Wave-4 | Putative outbreak cluster |
| VP18073 | VP18073 | 2018 | China | Clinical | Asia | O3:K6 | ST3 | PC | Wave-4 | Putative outbreak cluster |
| VP18074 | VP18074 | 2018 | China | Clinical | Asia | O3:K6 | ST3 | PC | Wave-4 | Putative outbreak cluster |
| VP18075 | VP18075 | 2018 | China | Clinical | Asia | O3:K6 | ST3 | PC | Wave-4 | Putative outbreak cluster |
| VP18076 | VP18076 | 2018 | China | Clinical | Asia | O3:K6 | ST3 | PC | Wave-4 | Putative outbreak cluster |

| | | | | | | | | | | |
|---|---|---|---|---|---|---|---|---|---|---|
| VP18077 | VP18077 | 2018 | China | Clinical | Asia | O3:K6 | ST3 | PC | Wave-4 | Putative outbreak cluster |
| VP18078 | VP18078 | 2018 | China | Clinical | Asia | O3:K6 | ST3 | PC | Wave-4 | Putative outbreak cluster |
| VP18079 | VP18079 | 2018 | China | Clinical | Asia | O3:K6 | ST3 | PC | Wave-4 | Putative outbreak cluster |
| VP18080 | VP18080 | 2018 | China | Clinical | Asia | O3:K6 | ST3 | PC | Wave-4 | Putative outbreak cluster |
| VP18081 | VP18081 | 2018 | China | Env | Asia | O3:K6 | ST3 | PC | Wave-4 | Putative outbreak cluster |
| VP18090 | VP18090 | 2018 | China | Clinical | Asia | O3:K6 | ST3 | PC | Wave-4 | Putative outbreak cluster |
| VP18094 | VP18094 | 2018 | China | Clinical | Asia | O3:K6 | ST3 | PC | Wave-4 | Putative outbreak cluster |
| VP18095 | VP18095 | 2018 | China | Clinical | Asia | O3:K6 | ST3 | PC | Wave-4 | Putative outbreak cluster |
| VP18096 | VP18096 | 2018 | China | Clinical | Asia | O3:K6 | ST3 | PC | Wave-4 | Putative outbreak cluster |
| VP18098 | VP18098 | 2018 | China | Clinical | Asia | O3:K6 | ST3 | PC | Wave-4 | Putative outbreak cluster |
| VP18099 | VP18099 | 2018 | China | Clinical | Asia | O3:K6 | ST3 | PC | Wave-4 | Putative outbreak cluster |
| VP18100 | VP18100 | 2018 | China | Clinical | Asia | O3:K6 | ST3 | PC | Wave-4 | Putative outbreak cluster |
| VP18101 | VP18101 | 2018 | China | Clinical | Asia | O3:K6 | ST3 | PC | Wave-4 | Putative outbreak cluster |
| VP18104 | VP18104 | 2018 | China | Clinical | Asia | O3:K6 | ST3 | PC | Wave-4 | Putative outbreak cluster |
| VP18107 | VP18107 | 2018 | China | Clinical | Asia | O3:K6 | ST3 | PC | Wave-4 | Putative outbreak cluster |
| VP18109 | VP18109 | 2018 | China | Clinical | Asia | O3:K6 | ST3 | PC | Wave-4 | Putative outbreak cluster |
| VP18110 | VP18110 | 2018 | China | Clinical | Asia | O3:K6 | ST3 | PC | Wave-4 | Putative outbreak cluster |
| VP18111 | VP18111 | 2018 | China | Clinical | Asia | O3:K6 | ST3 | PC | Wave-4 | Putative outbreak cluster |
| VP18112 | VP18112 | 2018 | China | Clinical | Asia | O3:K6 | ST3 | PC | Wave-4 | Putative outbreak cluster |
| VP18113 | VP18113 | 2018 | China | Clinical | Asia | O3:K6 | ST3 | PC | Wave-4 | Putative outbreak cluster |
| VP18117 | VP18117 | 2018 | China | Clinical | Asia | O3:K6 | ST3 | PC | Wave-4 | Putative outbreak cluster |
| VP18120 | VP18120 | 2018 | China | Clinical | Asia | O3:K6 | ST3 | PC | Wave-4 | Putative outbreak cluster |
| VP18122 | VP18122 | 2018 | China | Clinical | Asia | O3:K6 | ST3 | PC | Wave-4 | Putative outbreak cluster |
| VP18123 | VP18123 | 2018 | China | Clinical | Asia | O3:K6 | ST3 | PC | Wave-4 | Putative outbreak cluster |
| VP18124 | VP18124 | 2018 | China | Clinical | Asia | O3:K6 | ST3 | PC | Wave-4 | Putative outbreak cluster |
| VP18127 | VP18127 | 2018 | China | Clinical | Asia | O3:K6 | ST3 | PC | Wave-4 | Putative outbreak cluster |
| VP18135 | VP18135 | 2018 | China | Clinical | Asia | O3:K6 | ST3 | PC | Wave-4 | Putative outbreak cluster |
| VP18137 | VP18137 | 2018 | China | Clinical | Asia | O3:K6 | ST3 | PC | Wave-4 | Putative outbreak cluster |
| VP18138 | VP18138 | 2018 | China | Clinical | Asia | O3:K6 | ST3 | PC | Wave-4 | Putative outbreak cluster |
| VP18139 | VP18139 | 2018 | China | Clinical | Asia | O3:K6 | ST3 | PC | Wave-4 | Putative outbreak cluster |
| VP18141 | VP18141 | 2018 | China | Clinical | Asia | O3:K6 | ST3 | PC | Wave-4 | Putative outbreak cluster |
| VP18143 | VP18143 | 2018 | China | Clinical | Asia | O3:K6 | ST3 | PC | Wave-4 | Putative outbreak cluster |

| | | | | | | | | | | |
|---|---|---|---|---|---|---|---|---|---|---|
| VP18144 | VP18144 | 2018 | China | Clinical | Asia | O3:K6 | ST3 | PC | Wave-4 | Putative outbreak cluster |
| VP18145 | VP18145 | 2018 | China | Clinical | Asia | O3:K6 | ST3 | PC | Wave-4 | Putative outbreak cluster |
| VP18146 | VP18146 | 2018 | China | Clinical | Asia | O3:K6 | ST3 | PC | Wave-4 | Putative outbreak cluster |
| VP18147 | VP18147 | 2018 | China | Clinical | Asia | O3:K6 | ST3 | PC | Wave-4 | Putative outbreak cluster |
| VP18148 | VP18148 | 2018 | China | Clinical | Asia | O3:K6 | ST3 | PC | Wave-4 | Putative outbreak cluster |
| VP18149 | VP18149 | 2018 | China | Clinical | Asia | O3:K6 | ST3 | PC | Wave-4 | Putative outbreak cluster |
| VP18151 | VP18151 | 2018 | China | Clinical | Asia | O3:K6 | ST3 | PC | Wave-4 | Putative outbreak cluster |
| VP18153 | VP18153 | 2018 | China | Clinical | Asia | O3:K6 | ST3 | PC | Wave-4 | Putative outbreak cluster |
| VP18154 | VP18154 | 2018 | China | Clinical | Asia | O3:K6 | ST3 | PC | Wave-4 | Putative outbreak cluster |
| VP18155 | VP18155 | 2018 | China | Clinical | Asia | O3:K6 | ST3 | PC | Wave-4 | Putative outbreak cluster |
| VP18156 | VP18156 | 2018 | China | Clinical | Asia | O3:K6 | ST3 | PC | Wave-4 | Putative outbreak cluster |
| VP18157 | VP18157 | 2018 | China | Clinical | Asia | O3:K6 | ST3 | PC | Wave-4 | Putative outbreak cluster |
| VP18158 | VP18158 | 2018 | China | Clinical | Asia | O3:K6 | ST3 | PC | Wave-4 | Putative outbreak cluster |
| VP18159 | VP18159 | 2018 | China | Clinical | Asia | O3:K6 | ST3 | PC | Wave-4 | Putative outbreak cluster |
| VP18160 | VP18160 | 2018 | China | Clinical | Asia | O3:K6 | ST3 | PC | Wave-4 | Putative outbreak cluster |
| VP18161 | VP18161 | 2018 | China | Clinical | Asia | O3:K6 | ST3 | PC | Wave-4 | Putative outbreak cluster |
| VP18162 | VP18162 | 2018 | China | Clinical | Asia | O3:K6 | ST3 | PC | Wave-4 | Putative outbreak cluster |
| VP18163 | VP18163 | 2018 | China | Clinical | Asia | O3:K6 | ST3 | PC | Wave-4 | Putative outbreak cluster |
| VP18173 | VP18173 | 2018 | China | Clinical | Asia | O3:K6 | ST3 | PC | Wave-4 | Putative outbreak cluster |
| VP18174 | VP18174 | 2018 | China | Clinical | Asia | O3:K6 | ST3 | PC | Wave-4 | Putative outbreak cluster |
| VP18175 | VP18175 | 2018 | China | Clinical | Asia | O3:K6 | ST3 | PC | Wave-4 | Putative outbreak cluster |
| VP18178 | VP18178 | 2018 | China | Clinical | Asia | O3:K6 | ST3 | PC | Wave-4 | Putative outbreak cluster |
| VP18179 | VP18179 | 2018 | China | Clinical | Asia | O3:K6 | ST3 | PC | Wave-4 | Putative outbreak cluster |
| VP18180 | VP18180 | 2018 | China | Clinical | Asia | O3:K6 | ST3 | PC | Wave-4 | Putative outbreak cluster |
| VP18181 | VP18181 | 2018 | China | Clinical | Asia | O3:K6 | ST3 | PC | Wave-4 | Putative outbreak cluster |
| VP18182 | VP18182 | 2018 | China | Clinical | Asia | O3:K6 | ST3 | PC | Wave-4 | Putative outbreak cluster |
| VP18183 | VP18183 | 2018 | China | Clinical | Asia | O3:K6 | ST3 | PC | Wave-4 | Putative outbreak cluster |
| VP18184 | VP18184 | 2018 | China | Clinical | Asia | O3:K6 | ST3 | PC | Wave-4 | Putative outbreak cluster |
| VP18185 | VP18185 | 2018 | China | Clinical | Asia | O3:K6 | ST3 | PC | Wave-4 | Putative outbreak cluster |
| VP18186 | VP18186 | 2018 | China | Clinical | Asia | O3:K6 | ST3 | PC | Wave-4 | Putative outbreak cluster |
| VP18188 | VP18188 | 2018 | China | Clinical | Asia | O3:K6 | ST3 | PC | Wave-4 | Putative outbreak cluster |
| VP18189 | VP18189 | 2018 | China | Clinical | Asia | O3:K6 | ST3 | PC | Wave-4 | Putative outbreak cluster |

| | | | | | | | | | | |
|---|---|---|---|---|---|---|---|---|---|---|
| VP18190 | VP18190 | 2018 | China | Clinical | Asia | O3:K6 | ST3 | PC | Wave-4 | Putative outbreak cluster |
| VP18191 | VP18191 | 2018 | China | Clinical | Asia | O3:K6 | ST3 | PC | Wave-4 | Putative outbreak cluster |
| VP18192 | VP18192 | 2018 | China | Clinical | Asia | O3:K6 | ST3 | PC | Wave-4 | Putative outbreak cluster |
| VP18193 | VP18193 | 2018 | China | Clinical | Asia | O3:K6 | ST3 | PC | Wave-4 | Putative outbreak cluster |
| VP18194 | VP18194 | 2018 | China | Clinical | Asia | O3:K6 | ST3 | PC | Wave-4 | Putative outbreak cluster |
| VP18195 | VP18195 | 2018 | China | Clinical | Asia | O3:K6 | ST3 | PC | Wave-4 | Putative outbreak cluster |
| VP18196 | VP18196 | 2018 | China | Clinical | Asia | O3:K6 | ST3 | PC | Wave-4 | Putative outbreak cluster |
| VP18199 | VP18199 | 2018 | China | Clinical | Asia | O3:K6 | ST3 | PC | Wave-4 | Putative outbreak cluster |
| VP18200 | VP18200 | 2018 | China | Clinical | Asia | O3:K6 | ST3 | PC | Wave-4 | Putative outbreak cluster |
| VP18201 | VP18201 | 2018 | China | Clinical | Asia | O3:K6 | ST3 | PC | Wave-4 | Putative outbreak cluster |
| VP18202 | VP18202 | 2018 | China | Clinical | Asia | O3:K6 | ST3 | PC | Wave-4 | Putative outbreak cluster |
| VP18203 | VP18203 | 2018 | China | Clinical | Asia | O3:K6 | ST3 | PC | Wave-4 | Putative outbreak cluster |
| VP18204 | VP18204 | 2018 | China | Clinical | Asia | O3:K6 | ST3 | PC | Wave-4 | Putative outbreak cluster |
| VP18205 | VP18205 | 2018 | China | Clinical | Asia | O3:K6 | ST3 | PC | Wave-4 | Putative outbreak cluster |
| VP18207 | VP18207 | 2018 | China | Clinical | Asia | O3:K6 | ST3 | PC | Wave-4 | Putative outbreak cluster |
| VP18208 | VP18208 | 2018 | China | Clinical | Asia | O3:K6 | ST3 | PC | Wave-4 | Putative outbreak cluster |
| VP18210 | VP18210 | 2018 | China | Clinical | Asia | O3:K6 | ST3 | PC | Wave-4 | Putative outbreak cluster |
| VP18211 | VP18211 | 2018 | China | Clinical | Asia | O3:K6 | ST3 | PC | Wave-4 | Putative outbreak cluster |
| VP18212 | VP18212 | 2018 | China | Clinical | Asia | O3:K6 | ST3 | PC | Wave-4 | Putative outbreak cluster |
| VP18214 | VP18214 | 2018 | China | Clinical | Asia | O3:K6 | ST3 | PC | Wave-4 | Putative outbreak cluster |
| VP18216 | VP18216 | 2018 | China | Clinical | Asia | O3:K6 | ST3 | PC | Wave-4 | Putative outbreak cluster |
| VP18218 | VP18218 | 2018 | China | Clinical | Asia | O3:K6 | ST3 | PC | Wave-4 | Putative outbreak cluster |
| VP18221 | VP18221 | 2018 | China | Clinical | Asia | O3:K6 | ST3 | PC | Wave-4 | Putative outbreak cluster |
| VP18223 | VP18223 | 2018 | China | Clinical | Asia | O3:K6 | ST3 | PC | Wave-4 | Putative outbreak cluster |
| VP18224 | VP18224 | 2018 | China | Clinical | Asia | O3:K6 | ST3 | PC | Wave-4 | Putative outbreak cluster |
| VP18227 | VP18227 | 2018 | China | Clinical | Asia | O3:K6 | ST3 | PC | Wave-4 | Putative outbreak cluster |
| VP18231 | VP18231 | 2018 | China | Clinical | Asia | O3:K6 | ST3 | PC | Wave-4 | Putative outbreak cluster |
| VP18235 | VP18235 | 2018 | China | Clinical | Asia | O3:K6 | ST3 | PC | Wave-4 | Putative outbreak cluster |
| VP18236 | VP18236 | 2018 | China | Clinical | Asia | O3:K6 | ST3 | PC | Wave-4 | Putative outbreak cluster |
| VP18237 | VP18237 | 2018 | China | Clinical | Asia | O3:K6 | ST3 | PC | Wave-4 | Putative outbreak cluster |
| VP18238 | VP18238 | 2018 | China | Clinical | Asia | O3:K6 | ST3 | PC | Wave-4 | Putative outbreak cluster |
| VP18239 | VP18239 | 2018 | China | Clinical | Asia | O3:K6 | ST3 | PC | Wave-4 | Putative outbreak cluster |

| | | | | | | | | | | |
|---|---|---|---|---|---|---|---|---|---|---|
| VP18243 | VP18243 | 2018 | China | Clinical | Asia | O3:K6 | ST3 | PC | Wave-4 | Putative outbreak cluster |
| VP19002 | VP19002 | 2019 | China | Clinical | Asia | O3:K6 | ST3 | PC | Wave-4 | Putative outbreak cluster |
| VP19004 | VP19004 | 2019 | China | Clinical | Asia | O3:K6 | ST3 | PC | Wave-4 | Putative outbreak cluster |
| VP19007 | VP19007 | 2019 | China | Clinical | Asia | O3:K6 | ST3 | PC | Wave-4 | Putative outbreak cluster |
| VP19024 | VP19024 | 2019 | China | Clinical | Asia | O3:K6 | ST3 | PC | Wave-4 | Putative outbreak cluster |
| VP19025 | VP19025 | 2019 | China | Clinical | Asia | O3:K6 | ST3 | PC | Wave-4 | Putative outbreak cluster |
| VP19027 | VP19027 | 2019 | China | Clinical | Asia | O3:K6 | ST3 | PC | Wave-4 | Putative outbreak cluster |
| VP19029 | VP19029 | 2019 | China | Clinical | Asia | O3:K6 | ST3 | PC | Wave-4 | Putative outbreak cluster |
| VP19030 | VP19030 | 2019 | China | Clinical | Asia | O3:K6 | ST3 | PC | Wave-4 | Putative outbreak cluster |
| VP19031 | VP19031 | 2019 | China | Clinical | Asia | O3:K6 | ST3 | PC | Wave-4 | Putative outbreak cluster |
| VP19034 | VP19034 | 2019 | China | Clinical | Asia | O3:K6 | ST3 | PC | Wave-4 | Putative outbreak cluster |
| VP19038 | VP19038 | 2019 | China | Clinical | Asia | O3:K6 | ST3 | PC | Wave-4 | Putative outbreak cluster |
| VP19039 | VP19039 | 2019 | China | Clinical | Asia | O3:K6 | ST3 | PC | Wave-4 | Putative outbreak cluster |
| VP19044 | VP19044 | 2019 | China | Clinical | Asia | O3:K6 | ST3 | PC | Wave-4 | Putative outbreak cluster |
| VP19048 | VP19048 | 2019 | China | Clinical | Asia | O3:K6 | ST3 | PC | Wave-4 | Putative outbreak cluster |
| VP19049 | VP19049 | 2019 | China | Clinical | Asia | O3:K6 | ST3 | PC | Wave-4 | Putative outbreak cluster |
| VP19050 | VP19050 | 2019 | China | Clinical | Asia | O3:K6 | ST3 | PC | Wave-4 | Putative outbreak cluster |
| VP19075 | VP19075 | 2019 | China | Clinical | Asia | O3:K6 | ST3 | PC | Wave-4 | Putative outbreak cluster |
| VP19076 | VP19076 | 2019 | China | Clinical | Asia | O3:K6 | ST3 | PC | Wave-4 | Putative outbreak cluster |
| VP19077 | VP19077 | 2019 | China | Clinical | Asia | O3:K6 | ST3 | PC | Wave-4 | Putative outbreak cluster |
| VP19090 | VP19090 | 2019 | China | Clinical | Asia | O3:K6 | ST3 | PC | Wave-4 | Putative outbreak cluster |
| VP19092 | VP19092 | 2019 | China | Clinical | Asia | O3:K6 | ST3 | PC | Wave-4 | Putative outbreak cluster |
| VP19111 | VP19111 | 2019 | China | Clinical | Asia | O3:K6 | ST3 | PC | Wave-4 | Putative outbreak cluster |
| VP19137 | VP19137 | 2019 | China | Clinical | Asia | O3:K6 | ST3 | PC | Wave-4 | Putative outbreak cluster |
| VP19139 | VP19139 | 2019 | China | Clinical | Asia | O3:K6 | ST3 | PC | Wave-4 | Putative outbreak cluster |
| VP19143 | VP19143 | 2019 | China | Clinical | Asia | O3:K6 | ST3 | PC | Wave-4 | Putative outbreak cluster |
| VP19146 | VP19146 | 2019 | China | Clinical | Asia | O3:K6 | ST3 | PC | Wave-4 | Putative outbreak cluster |
| VP19166 | VP19166 | 2019 | China | Clinical | Asia | O3:K6 | ST3 | PC | Wave-4 | Putative outbreak cluster |
| VP19169 | VP19169 | 2019 | China | Clinical | Asia | O3:K6 | ST3 | PC | Wave-4 | Putative outbreak cluster |
| VP19171 | VP19171 | 2019 | China | Clinical | Asia | O3:K6 | ST3 | PC | Wave-4 | Putative outbreak cluster |
| VP19205 | VP19205 | 2019 | China | Clinical | Asia | O3:K6 | ST3 | PC | Wave-4 | Putative outbreak cluster |
| VP19206 | VP19206 | 2019 | China | Clinical | Asia | O3:K6 | ST3 | PC | Wave-4 | Putative outbreak cluster |

| | | | | | | | | | | |
|---|---|---|---|---|---|---|---|---|---|---|
| VP19235 | VP19235 | 2019 | China | Clinical | Asia | O3:K6 | ST3 | PC | Wave-4 | Putative outbreak cluster |
| VP19238 | VP19238 | 2019 | China | Clinical | Asia | O3:K6 | ST3 | PC | Wave-4 | Putative outbreak cluster |
| VP19247 | VP19247 | 2019 | China | Clinical | Asia | O3:K6 | ST3 | PC | Wave-4 | Putative outbreak cluster |
| VP19248 | VP19248 | 2019 | China | Clinical | Asia | O3:K6 | ST3 | PC | Wave-4 | Putative outbreak cluster |
| VP19249 | VP19249 | 2019 | China | Clinical | Asia | O3:K6 | ST3 | PC | Wave-4 | Putative outbreak cluster |
| VP19251 | VP19251 | 2019 | China | Clinical | Asia | O3:K6 | ST3 | PC | Wave-4 | Putative outbreak cluster |
| VP19253 | VP19253 | 2019 | China | Clinical | Asia | O3:K6 | ST3 | PC | Wave-4 | Putative outbreak cluster |
| VP19254 | VP19254 | 2019 | China | Clinical | Asia | O3:K6 | ST3 | PC | Wave-4 | Putative outbreak cluster |
| VP19255 | VP19255 | 2019 | China | Clinical | Asia | O3:K6 | ST3 | PC | Wave-4 | Putative outbreak cluster |
| VP19256 | VP19256 | 2019 | China | Clinical | Asia | O3:K6 | ST3 | PC | Wave-4 | Putative outbreak cluster |
| VP19258 | VP19258 | 2019 | China | Clinical | Asia | O3:K6 | ST3 | PC | Wave-4 | Putative outbreak cluster |
| VP19260 | VP19260 | 2019 | China | Clinical | Asia | O3:K6 | ST3 | PC | Wave-4 | Putative outbreak cluster |
| VP19263 | VP19263 | 2019 | China | Clinical | Asia | O3:K6 | ST3 | PC | Wave-4 | Putative outbreak cluster |
| VP19267 | VP19267 | 2019 | China | Clinical | Asia | O3:K6 | ST3 | PC | Wave-4 | Putative outbreak cluster |
| VP19276 | VP19276 | 2019 | China | Clinical | Asia | O3:K6 | ST3 | PC | Wave-4 | Putative outbreak cluster |
| VP20023 | VP20023 | 2020 | China | Clinical | Asia | O3:K6 | ST3 | PC | Wave-4 | Putative outbreak cluster |
| VP20025 | VP20025 | 2020 | China | Clinical | Asia | O3:K6 | ST3 | PC | Wave-4 | Putative outbreak cluster |
| VP20026 | VP20026 | 2020 | China | Clinical | Asia | O3:K6 | ST3 | PC | Wave-4 | Putative outbreak cluster |
| VP20029 | VP20029 | 2020 | China | Clinical | Asia | O3:K6 | ST3 | PC | Wave-4 | Putative outbreak cluster |
| VP20030 | VP20030 | 2020 | China | Clinical | Asia | O3:K6 | ST3 | PC | Wave-4 | Putative outbreak cluster |
| VP20031 | VP20031 | 2020 | China | Clinical | Asia | O3:K6 | ST3 | PC | Wave-4 | Putative outbreak cluster |
| VP20035 | VP20035 | 2020 | China | Clinical | Asia | O3:K6 | ST3 | PC | Wave-4 | Putative outbreak cluster |
| VP20036 | VP20036 | 2020 | China | Clinical | Asia | O3:K6 | ST3 | PC | Wave-4 | Putative outbreak cluster |
| VP20037 | VP20037 | 2020 | China | Clinical | Asia | O10:K4 | ST3 | PC | Wave-4 | Putative outbreak cluster |
| VP20039 | VP20039 | 2020 | China | Clinical | Asia | O10:K4 | ST3 | PC | Wave-4 | Putative outbreak cluster |
| VP20077 | VP20077 | 2020 | China | Clinical | Asia | O10:K4 | ST3 | PC | Wave-4 | Putative outbreak cluster |
| VP20083 | VP20083 | 2020 | China | Clinical | Asia | O10:K4 | ST3 | PC | Wave-4 | Putative outbreak cluster |
| VP20085 | VP20085 | 2020 | China | Clinical | Asia | O10:K4 | ST3 | PC | Wave-4 | Putative outbreak cluster |
| VP20090 | VP20090 | 2020 | China | Clinical | Asia | O3:K6 | ST3 | PC | Wave-4 | Putative outbreak cluster |
| VP20093 | VP20093 | 2020 | China | Clinical | Asia | O3:K6 | ST3 | PC | Wave-4 | Putative outbreak cluster |
| VP20098 | VP20098 | 2020 | China | Clinical | Asia | O3:K6 | ST3 | PC | Wave-4 | Putative outbreak cluster |
| VP20105 | VP20105 | 2020 | China | Clinical | Asia | O3:K6 | ST3 | PC | Wave-4 | Putative outbreak cluster |

| | | | | | | | | | | |
|---|---|---|---|---|---|---|---|---|---|---|
| VP20106 | VP20106 | 2020 | China | Clinical | Asia | O3:K6 | ST3 | PC | Wave-4 | Putative outbreak cluster |
| VP20107 | VP20107 | 2020 | China | Clinical | Asia | O10:K4 | ST3 | PC | Wave-4 | Putative outbreak cluster |
| VP20108 | VP20108 | 2020 | China | Clinical | Asia | O10:K4 | ST3 | PC | Wave-4 | Putative outbreak cluster |
| VP20110 | VP20110 | 2020 | China | Clinical | Asia | O10:K4 | ST3 | PC | Wave-4 | Putative outbreak cluster |
| VP20162 | VP20162 | 2020 | China | Clinical | Asia | O10:K4 | ST3 | PC | Wave-4 | Putative outbreak cluster |
| VP20171 | VP20171 | 2020 | China | Clinical | Asia | O10:K4 | ST3 | PC | Wave-4 | Putative outbreak cluster |
| VP21019 | VP21019 | 2021 | China | Clinical | Asia | O10:K4 | ST3 | PC | Wave-4 | Putative outbreak cluster |
| VP21021 | VP21021 | 2021 | China | Clinical | Asia | O3:K6 | ST3 | PC | Wave-4 | Putative outbreak cluster |
| VP21023 | VP21023 | 2021 | China | Clinical | Asia | O10:K4 | ST3 | PC | Wave-4 | Putative outbreak cluster |
| VP21024 | VP21024 | 2021 | China | Clinical | Asia | O10:K4 | ST3 | PC | Wave-4 | Putative outbreak cluster |
| VP21034 | VP21034 | 2021 | China | Clinical | Asia | O10:K4 | ST3 | PC | Wave-4 | Putative outbreak cluster |
| VP21057 | VP21057 | 2021 | China | Clinical | Asia | O10:K4 | ST3 | PC | Wave-4 | Putative outbreak cluster |
| VP21067 | VP21067 | 2021 | China | Clinical | Asia | O10:K4 | ST3 | PC | Wave-4 | Putative outbreak cluster |
| VP21071 | VP21071 | 2021 | China | Clinical | Asia | O10:K4 | ST3 | PC | Wave-4 | Putative outbreak cluster |
| VP21072 | VP21072 | 2021 | China | Clinical | Asia | O10:K4 | ST3 | PC | Wave-4 | Putative outbreak cluster |
| VP21074 | VP21074 | 2021 | China | Clinical | Asia | O10:K4 | ST3 | PC | Wave-4 | Putative outbreak cluster |
| VP21077 | VP21077 | 2021 | China | Clinical | Asia | O10:K4 | ST3 | PC | Wave-4 | Putative outbreak cluster |
| VP21078 | VP21078 | 2021 | China | Clinical | Asia | O10:K4 | ST3 | PC | Wave-4 | Putative outbreak cluster |
| VP21079 | VP21079 | 2021 | China | Clinical | Asia | O10:K4 | ST3 | PC | Wave-4 | Putative outbreak cluster |
| VP21080 | VP21080 | 2021 | China | Clinical | Asia | O10:K4 | ST3 | PC | Wave-4 | Putative outbreak cluster |
| VP21081 | VP21081 | 2021 | China | Clinical | Asia | O10:K4 | ST3 | PC | Wave-4 | Putative outbreak cluster |
| VP21082 | VP21082 | 2021 | China | Clinical | Asia | O10:K4 | ST3 | PC | Wave-4 | Putative outbreak cluster |
| VP21083 | VP21083 | 2021 | China | Clinical | Asia | O10:K4 | ST3 | PC | Wave-4 | Putative outbreak cluster |
| VP21084 | VP21084 | 2021 | China | Clinical | Asia | O10:K4 | ST3 | PC | Wave-4 | Putative outbreak cluster |
| VP21086 | VP21086 | 2021 | China | Clinical | Asia | O10:K4 | ST3 | PC | Wave-4 | Putative outbreak cluster |
| VP21087 | VP21087 | 2021 | China | Clinical | Asia | O10:K4 | ST3 | PC | Wave-4 | Putative outbreak cluster |
| VP21088 | VP21088 | 2021 | China | Clinical | Asia | O10:K4 | ST3 | PC | Wave-4 | Putative outbreak cluster |
| VP21089 | VP21089 | 2021 | China | Clinical | Asia | O10:K4 | ST3 | PC | Wave-4 | Putative outbreak cluster |
| VP21090 | VP21090 | 2021 | China | Clinical | Asia | O10:K4 | ST3 | PC | Wave-4 | Putative outbreak cluster |
| VP21091 | VP21091 | 2021 | China | Clinical | Asia | O10:K4 | ST3 | PC | Wave-4 | Putative outbreak cluster |
| VP21092 | VP21092 | 2021 | China | Clinical | Asia | O10:K4 | ST3 | PC | Wave-4 | Putative outbreak cluster |
| VP21095 | VP21095 | 2021 | China | Clinical | Asia | O10:K4 | ST3 | PC | Wave-4 | Putative outbreak cluster |

| | | | | | | | | | | |
|---|---|---|---|---|---|---|---|---|---|---|
| VP21097 | VP21097 | 2021 | China | Clinical | Asia | O10:K4 | ST3 | PC | Wave-4 | Putative outbreak cluster |
| VP21098 | VP21098 | 2021 | China | Clinical | Asia | O10:K4 | ST3 | PC | Wave-4 | Putative outbreak cluster |
| VP21099 | VP21099 | 2021 | China | Clinical | Asia | O10:K4 | ST3 | PC | Wave-4 | Putative outbreak cluster |
| VP21100 | VP21100 | 2021 | China | Clinical | Asia | O10:K4 | ST3 | PC | Wave-4 | Putative outbreak cluster |
| VP21101 | VP21101 | 2021 | China | Clinical | Asia | O10:K4 | ST3 | PC | Wave-4 | Putative outbreak cluster |
| VP21102 | VP21102 | 2021 | China | Clinical | Asia | O10:K4 | ST3 | PC | Wave-4 | Putative outbreak cluster |
| VP21103 | VP21103 | 2021 | China | Clinical | Asia | O10:K4 | ST3 | PC | Wave-4 | Putative outbreak cluster |
| VPCZ1 | VPCZ1 | 2013 | China | Clinical | Asia | O3:K6 | ST3 | PC | Wave-4 | Putative outbreak cluster |
| VPCZ107 | VPCZ107 | 2014 | China | Clinical | Asia | O10:K60 | ST3 | PC | Wave-4 | Putative outbreak cluster |
| VPCZ13 | VPCZ13 | 2014 | China | Clinical | Asia | O3:K6 | ST3 | PC | Wave-4 | Putative outbreak cluster |
| VPCZ14 | VPCZ14 | 2014 | China | Clinical | Asia | O3:K6 | ST3 | PC | Wave-4 | Putative outbreak cluster |
| VPCZ17 | VPCZ17 | 2014 | China | Clinical | Asia | O3:K6 | ST3 | PC | Wave-4 | Putative outbreak cluster |
| VPCZ18 | VPCZ18 | 2014 | China | Clinical | Asia | O3:K6 | ST3 | PC | Wave-4 | Putative outbreak cluster |
| VPCZ21 | VPCZ21 | 2014 | China | Clinical | Asia | O3:K6 | ST3 | PC | Wave-4 | Putative outbreak cluster |
| VPCZ23 | VPCZ23 | 2014 | China | Clinical | Asia | O3:K6 | ST3 | PC | Wave-4 | Putative outbreak cluster |
| VPCZ25 | VPCZ25 | 2014 | China | Clinical | Asia | O3:K6 | ST3 | PC | Wave-4 | Putative outbreak cluster |
| VPCZ29 | VPCZ29 | 2014 | China | Clinical | Asia | O3:K6 | ST3 | PC | Wave-4 | Putative outbreak cluster |
| VPCZ3 | VPCZ3 | 2013 | China | Clinical | Asia | O3:K6 | ST3 | PC | Wave-4 | Putative outbreak cluster |
| VPCZ31 | VPCZ31 | 2014 | China | Clinical | Asia | O3:K6 | ST3 | PC | Wave-4 | Putative outbreak cluster |
| VPCZ33 | VPCZ33 | 2014 | China | Clinical | Asia | O3:K6 | ST3 | PC | Wave-4 | Putative outbreak cluster |
| VPCZ35 | VPCZ35 | 2014 | China | Clinical | Asia | O3:K6 | ST3 | PC | Wave-4 | Putative outbreak cluster |
| VPCZ36 | VPCZ36 | 2014 | China | Clinical | Asia | O3:K6 | ST3 | PC | Wave-4 | Putative outbreak cluster |
| VPCZ4 | VPCZ4 | 2013 | China | Clinical | Asia | O3:K6 | ST3 | PC | Wave-4 | Putative outbreak cluster |
| VPCZ58 | VPCZ58 | 2014 | China | Clinical | Asia | O3:K6 | ST3 | PC | Wave-4 | Putative outbreak cluster |
| VPCZ82 | VPCZ82 | 2014 | China | Clinical | Asia | O3:K6 | ST3 | PC | Wave-4 | Putative outbreak cluster |
| SAMN02741386-GCF_000707765.2 | CFSAN007450 | 2012 | USA | Clinical | US_Canada | O3:K6 | ST3 | PC | Wave-4 | Putative outbreak cluster |
| SAMN07327345-GCA_015816975.1 | PNUSAV000060 | NA | USA | Clinical | US_Canada | O3:K6 | ST3 | PC | Wave-4 | Putative outbreak cluster |
| SAMN09488304-GCA_015806535.1 | PNUSAV000192 | NA | USA | Clinical | US_Canada | O3:K6 | ST3 | PC | Wave-4 | Putative outbreak cluster |
| SAMN09488305-GCA_015806375.1 | PNUSAV000190 | NA | USA | Clinical | US_Canada | O3:K6 | ST3 | PC | Wave-4 | Putative outbreak cluster |
| SAMN09488306-GCA_015806455.1 | PNUSAV000189 | NA | USA | Clinical | US_Canada | O3:K65 | ST3 | PC | Wave-4 | Putative outbreak cluster |
| SAMN09664793-GCA_015806315.1 | PNUSAV000194 | NA | USA | Clinical | US_Canada | O3:K6 | ST3 | PC | Wave-4 | Putative outbreak cluster |
| SAMN09671573-GCA_015804035.1 | PNUSAV000206 | NA | USA | Clinical | US_Canada | O3:K6 | ST3 | PC | Wave-4 | Putative outbreak cluster |

| | | | | | | | | | | |
|---|---|---|---|---|---|---|---|---|---|---|
| SAMN09671576-GCA_015805145.1 | PNUSAV000203 | NA | USA | Clinical | US_Canada | O3:K6 | ST3 | PC | Wave-4 | Putative outbreak cluster |
| SAMN09671577-GCA_015804095.1 | PNUSAV000202 | NA | USA | Clinical | US_Canada | O3:K6 | ST3 | PC | Wave-4 | Putative outbreak cluster |
| SAMN09671587-GCA_015804115.1 | PNUSAV000207 | NA | USA | Clinical | US_Canada | O3:K6 | ST3 | PC | Wave-4 | Putative outbreak cluster |
| SAMN09703859-GCA_015803795.1 | PNUSAV000222 | NA | USA | Clinical | US_Canada | O3:K6 | ST3 | PC | Wave-4 | Putative outbreak cluster |
| SAMN09726939-GCA_015804895.1 | PNUSAV000211 | NA | USA | Clinical | US_Canada | O3:K6 | ST3 | PC | Wave-4 | Putative outbreak cluster |
| SAMN09760593-GCA_015803735.1 | PNUSAV000249 | NA | USA | Clinical | US_Canada | O3:K6 | ST3 | PC | Wave-4 | Putative outbreak cluster |
| SAMN09768254-GCA_015803475.1 | PNUSAV000283 | NA | USA | Clinical | US_Canada | O3:K6 | ST3 | PC | Wave-4 | Putative outbreak cluster |
| SAMN09864371-GCA_015801555.1 | PNUSAV000336 | NA | USA | Clinical | US_Canada | O3:K6 | ST3 | PC | Wave-4 | Putative outbreak cluster |
| SAMN09878925-GCA_015801455.1 | 2018AW-0103 | NA | USA | Clinical | US_Canada | O3:K6 | ST3 | PC | Wave-4 | Putative outbreak cluster |
| SAMN09878927-GCA_015801375.1 | 2018V-1064 | NA | USA | Clinical | US_Canada | O3:K6 | ST3 | PC | Wave-4 | Putative outbreak cluster |
| SAMN09917443-GCA_015800035.1 | PNUSAV000356 | NA | USA | Clinical | US_Canada | O3:K6 | ST3 | PC | Wave-4 | Putative outbreak cluster |
| SAMN10095495-GCA_015800555.1 | PNUSAV000440 | NA | USA | Clinical | US_Canada | O3:K6 | ST3 | PC | Wave-4 | Putative outbreak cluster |
| SAMN10095500-GCA_015800435.1 | PNUSAV000446 | NA | USA | Clinical | US_Canada | O3:K6 | ST3 | PC | Wave-4 | Putative outbreak cluster |
| SAMN10095503-GCA_015800775.1 | PNUSAV000443 | NA | USA | Clinical | US_Canada | O3:K6 | ST3 | PC | Wave-4 | Putative outbreak cluster |
| SAMN10095505-GCA_015800495.1 | PNUSAV000441 | NA | USA | Clinical | US_Canada | O3:K6 | ST3 | PC | Wave-4 | Putative outbreak cluster |
| SAMN10163150-GCA_015799235.1 | PNUSAV000480 | NA | USA | Clinical | US_Canada | O3:K6 | ST3 | PC | Wave-4 | Putative outbreak cluster |
| SAMN10163151-GCA_015799215.1 | PNUSAV000479 | NA | USA | Clinical | US_Canada | O3:K6 | ST3 | PC | Wave-4 | Putative outbreak cluster |
| SAMN10221805-GCA_015798995.1 | PNUSAV000481 | NA | USA | Clinical | US_Canada | O3:K6 | ST3 | PC | Wave-4 | Putative outbreak cluster |
| SAMN10419344-GCA_015797495.1 | PNUSAV000525 | NA | USA | Clinical | US_Canada | O3:K6 | ST3 | PC | Wave-4 | Putative outbreak cluster |
| SAMN11159227-GCA_015789105.1 | PNUSAV000580 | NA | USA | Clinical | US_Canada | O3:K6 | ST3 | PC | Wave-4 | Putative outbreak cluster |
| SAMN12212336-GCA_015789775.1 | PNUSAV000253 | NA | USA | Clinical | US_Canada | O3:K6 | ST3 | PC | Wave-4 | Putative outbreak cluster |
| SAMN29444630-SRR19910443 | PNUSAV002568 | 2022 | USA | Clinical | US_Canada | O10:K4 | ST3 | PC | Wave-4 | Putative outbreak cluster |
| SAMEA8103018-GCA_905331765.1 | ERS5790096 | NA | Japan | Clinical | Asia | O10:K60 | ST3 | PC | Wave-4 | Representative |
| SAMN01923897-GCF_000522025.2 | IDH02189 | 2009 | India | Clinical | Asia | O3:K6 | ST3 | PC | Wave-4 | Representative |
| SAMN01923903-GCF_000522005.2 | EKP-028 | 2008 | Bangladesh | Env | Asia | O3:K6 | ST3 | PC | Wave-4 | Representative |
| SAMN02471133-GCF_000500405.1 | AXNM | 2008 | China | Clinical | Asia | O3:K6 | ST3 | PC | Wave-4 | Representative |
| SAMN02641511-GCF_000525005.2 | EKP-026 | 2008 | Bangladesh | Env | Asia | O3:K6 | ST3 | PC | Wave-4 | Representative |
| SAMN05890675-GCF_003408895.1 | L70 | 2016 | China | NA | Asia | O3:K6 | ST3 | PC | Wave-4 | Representative |
| SAMN05935529-GCF_001895695.1 | 100145 | 2013 | China | Clinical | Asia | O3:K6 | ST3 | PC | Wave-4 | Representative |
| SAMN06162268-GCF_001913735.1 | GIMxtfL61-2011.05 | 2011 | China | Clinical | Asia | O3:K6 | ST3 | PC | Wave-4 | Representative |
| SAMN07338135-GCF_006371215.1 | F2_10 | 2014 | China | Env | Asia | O3:K6 | ST3 | PC | Wave-4 | Representative |
| SAMN07338142-GCF_006371225.1 | F2_8 | 2014 | China | Env | Asia | O3:K6 | ST- | PC | Wave-4 | Representative |

| | | | | | | | | | | |
|---|---|---|---|---|---|---|---|---|---|---|
| SAMN07338143-GCF_006370655.1 | F2_9 | 2014 | China | Env | Asia | O3:K6 | ST3 | PC | Wave-4 | Representative |
| SAMN07338151-GCF_006370635.1 | F3_7 | 2014 | China | Env | Asia | O3:K6 | ST3 | PC | Wave-4 | Representative |
| SAMN07338152-GCF_006371125.1 | F3_8 | 2014 | China | Env | Asia | O3:K6 | ST3 | PC | Wave-4 | Representative |
| SAMN07338213-GCF_006368505.1 | G1_3 | 2014 | China | Env | Asia | O3:K6 | ST3 | PC | Wave-4 | Representative |
| SAMN07338216-GCF_006368345.1 | G1_6 | 2014 | China | Env | Asia | O3:K6 | ST3 | PC | Wave-4 | Representative |
| SAMN07338217-GCF_006368485.1 | G1_7 | 2014 | China | Env | Asia | O3:K6 | ST3 | PC | Wave-4 | Representative |
| SAMN07338219-GCF_006368435.1 | G1_9 | 2014 | China | Env | Asia | O3:K6 | ST3 | PC | Wave-4 | Representative |
| SAMN07338221-GCF_006368425.1 | G2_2 | 2014 | China | Env | Asia | O3:K6 | ST3 | PC | Wave-4 | Representative |
| SAMN08667585-GCF_003056645.1 | VPF-4 | 2017 | Lebanon | Clinical | Asia | O3:K6 | ST3 | PC | Wave-4 | Representative |
| SAMN08667664-GCF_003056665.1 | VPF-6 | 2017 | Lebanon | Clinical | Asia | O3:K6 | ST3 | PC | Wave-4 | Representative |
| SAMN09742584-GCF_006382955.1 | MH16341 | 2016 | China | Clinical | Asia | O10:K60 | ST3 | PC | Wave-4 | Representative |
| SAMN09874427-GCF_004006515.1 | VPD14 | 2012 | China | NA | Asia | O3:K6 | ST3 | PC | Wave-4 | Representative |
| SAMN10240147-GCA_015778625.1 | PJ35 | 2017 | China | Env | Asia | O3:K6 | ST- | PC | Wave-4 | Representative |
| SAMN11547090-GCF_009389765.1 | HZ18-116 | 2018 | China | Clinical | Asia | O3:K6 | ST3 | PC | Wave-4 | Representative |
| SAMN13706855-SRR11823785 | SH-30 | 2015 | China | Env | Asia | O3:K6 | ST3 | PC | Wave-4 | Representative |
| SAMN14411230-GCF_014922245.1 | L6 | 2015 | China | Clinical | Asia | O3:K6 | ST3 | PC | Wave-4 | Representative |
| SAMN14411234-GCF_014922125.1 | r46 | 2018 | China | Clinical | Asia | O3:K6 | ST3 | PC | Wave-4 | Representative |
| SAMN14411236-GCF_014922115.1 | r79-2 | 2017 | China | Clinical | Asia | O3:K6 | ST3 | PC | Wave-4 | Representative |
| SAMN15294467-GCA_019686095.1 | ICDC-VP01786 | 2016 | China | Clinical | Asia | O3:K6 | ST3 | PC | Wave-4 | Representative |
| SAMN15294468-GCA_019686045.1 | ICDC-VP01787 | 2016 | China | Clinical | Asia | O3:K6 | ST3 | PC | Wave-4 | Representative |
| SAMN15294472-GCA_019685995.1 | ICDC-VP01797 | 2017 | China | Clinical | Asia | O3:K6 | ST3 | PC | Wave-4 | Representative |
| SAMN15294474-GCA_019685895.1 | ICDC-VP01800 | 2017 | China | Clinical | Asia | O3:K6 | ST3 | PC | Wave-4 | Representative |
| SAMN15294475-GCA_019685935.1 | ICDC-VP01801 | 2017 | China | Clinical | Asia | O3:K6 | ST3 | PC | Wave-4 | Representative |
| SAMN15294477-GCA_019685835.1 | ICDC-VP01803 | 2017 | China | Clinical | Asia | O3:K6 | ST3 | PC | Wave-4 | Representative |
| SAMN15294478-GCA_019685805.1 | ICDC-VP01805 | 2018 | China | Clinical | Asia | O3:K6 | ST3 | PC | Wave-4 | Representative |
| SAMN15294479-GCA_019685755.1 | ICDC-VP01806 | 2018 | China | Clinical | Asia | O3:K6 | ST3 | PC | Wave-4 | Representative |
| SAMN15294481-GCA_019685735.1 | ICDC-VP01811 | 2018 | China | Clinical | Asia | O3:K6 | ST3 | PC | Wave-4 | Representative |
| SAMN15294482-GCA_019685715.1 | ICDC-VP01812 | 2018 | China | Clinical | Asia | O3:K6 | ST3 | PC | Wave-4 | Representative |
| SAMN15294483-GCA_019685695.1 | ICDC-VP01813 | 2018 | China | Clinical | Asia | O3:K6 | ST3 | PC | Wave-4 | Representative |
| SAMN15294484-GCA_019685675.1 | ICDC-VP01814 | 2018 | China | Clinical | Asia | O3:K6 | ST3 | PC | Wave-4 | Representative |
| SAMN15294486-GCA_019685615.1 | ICDC-VP01816 | 2018 | China | Clinical | Asia | O3:K6 | ST3 | PC | Wave-4 | Representative |
| SAMN16205401-GCF_015680975.1 | SH112 | 2019 | China | Clinical | Asia | O3:K6 | ST3 | PC | Wave-4 | Representative |

| | | | | | | | | | | |
|---|---|---|---|---|---|---|---|---|---|---|
| SAMN16782965-GCA_016817945.1 | VP28 | 2009 | China | NA | Asia | O10:K60 | ST3 | PC | Wave-4 | Representative |
| SAMN16783345-GCA_016819245.1 | VP8 | 2016 | China | NA | Asia | O10:K60 | ST3 | PC | Wave-4 | Representative |
| SAMN16783347-GCA_016819235.1 | VP16 | 2015 | China | NA | Asia | O10:K60 | ST3 | PC | Wave-4 | Representative |
| SAMN16783348-GCA_016819175.1 | VP23 | 2015 | China | NA | Asia | O10:K60 | ST3 | PC | Wave-4 | Representative |
| SAMN16783349-GCA_016819195.1 | VP24 | 2015 | China | NA | Asia | O10:K60 | ST3 | PC | Wave-4 | Representative |
| SAMN16783356-GCA_016819045.1 | VP65 | 2013 | China | NA | Asia | O10:K60 | ST3 | PC | Wave-4 | Representative |
| VP04076 | VP04076 | 2004 | China | Clinical | Asia | O3:K6 | ST3 | PC | Wave-4 | Representative |
| VP04078 | VP04078 | 2004 | China | Clinical | Asia | O3:K6 | ST3 | PC | Wave-4 | Representative |
| VP04079 | VP04079 | 2004 | China | Clinical | Asia | O3:K6 | ST3 | PC | Wave-4 | Representative |
| VP04080 | VP04080 | 2004 | China | Clinical | Asia | O3:K6 | ST3 | PC | Wave-4 | Representative |
| VP04097 | VP04097 | 2004 | China | Clinical | Asia | O3:K6 | ST3 | PC | Wave-4 | Representative |
| VP04099 | VP04099 | 2004 | China | Clinical | Asia | O3:K6 | ST3 | PC | Wave-4 | Representative |
| VP04108 | VP04108 | 2004 | China | Clinical | Asia | O3:K6 | ST3 | PC | Wave-4 | Representative |
| VP04129 | VP04129 | 2004 | China | Clinical | Asia | O3:K6 | ST3 | PC | Wave-4 | Representative |
| VP05213 | VP05213 | 2005 | China | Clinical | Asia | O3:K6 | ST3 | PC | Wave-4 | Representative |
| VP05217 | VP05217 | 2005 | China | Clinical | Asia | O3:K6 | ST- | PC | Wave-4 | Representative |
| VP06011 | VP06011 | 2006 | China | Env | Asia | O3:K6 | ST- | PC | Wave-4 | Representative |
| VP06034 | VP06034 | 2006 | China | Clinical | Asia | O3:K6 | ST- | PC | Wave-4 | Representative |
| VP06045 | VP06045 | 2006 | China | Clinical | Asia | O3:K6 | ST3 | PC | Wave-4 | Representative |
| VP06046 | VP06046 | 2006 | China | Clinical | Asia | O3:K6 | ST3 | PC | Wave-4 | Representative |
| VP06065 | VP06065 | 2006 | China | Clinical | Asia | O3:K6 | ST3 | PC | Wave-4 | Representative |
| VP06073 | VP06073 | 2006 | China | Env | Asia | O3:K6 | ST3 | PC | Wave-4 | Representative |
| VP06158 | VP06158 | 2006 | China | Clinical | Asia | O3:K6 | ST3 | PC | Wave-4 | Representative |
| VP06193 | VP06193 | 2006 | China | Clinical | Asia | O3:K6 | ST3 | PC | Wave-4 | Representative |
| VP07060 | VP07060 | 2007 | China | Clinical | Asia | O3:K6 | ST3 | PC | Wave-4 | Representative |
| VP07080 | VP07080 | 2007 | China | Clinical | Asia | O3:K6 | ST3 | PC | Wave-4 | Representative |
| VP07097 | VP07097 | 2007 | China | Clinical | Asia | O3:K6 | ST3 | PC | Wave-4 | Representative |
| VP07187 | VP07187 | 2007 | China | Clinical | Asia | O3:K6 | ST3 | PC | Wave-4 | Representative |
| VP07199 | VP07199 | 2007 | China | Clinical | Asia | O3:K6 | ST3 | PC | Wave-4 | Representative |
| VP08010 | VP08010 | 2008 | China | Clinical | Asia | O3:K6 | ST3 | PC | Wave-4 | Representative |
| VP08024 | VP08024 | 2008 | China | Clinical | Asia | O3:K6 | ST3 | PC | Wave-4 | Representative |
| VP08036 | VP08036 | 2008 | China | Clinical | Asia | O3:K6 | ST3 | PC | Wave-4 | Representative |

| | | | | | | | | | | |
|---|---|---|---|---|---|---|---|---|---|---|
| VP08144 | VP08144 | 2008 | China | Clinical | Asia | O3:K6 | ST3 | PC | Wave-4 | Representative |
| VP08190 | VP08190 | 2008 | China | Clinical | Asia | O3:K6 | ST3 | PC | Wave-4 | Representative |
| VP08202 | VP08202 | 2008 | China | Clinical | Asia | O3:K6 | ST3 | PC | Wave-4 | Representative |
| VP08206 | VP08206 | 2008 | China | Clinical | Asia | O3:K6 | ST3 | PC | Wave-4 | Representative |
| VP08210 | VP08210 | 2008 | China | Clinical | Asia | O3:K6 | ST3 | PC | Wave-4 | Representative |
| VP08229 | VP08229 | 2008 | China | Clinical | Asia | O3:K6 | ST3 | PC | Wave-4 | Representative |
| VP08232 | VP08232 | 2008 | China | Clinical | Asia | O3:K6 | ST3 | PC | Wave-4 | Representative |
| VP08234 | VP08234 | 2008 | China | Clinical | Asia | O3:K6 | ST3 | PC | Wave-4 | Representative |
| VP08236 | VP08236 | 2008 | China | Clinical | Asia | O3:K6 | ST3 | PC | Wave-4 | Representative |
| VP08366 | VP08366 | 2008 | China | Clinical | Asia | O3:K6 | ST3 | PC | Wave-4 | Representative |
| VP08394 | VP08394 | 2008 | China | Clinical | Asia | O3:K6 | ST3 | PC | Wave-4 | Representative |
| VP09044 | VP09044 | 2009 | China | Clinical | Asia | O3:K6 | ST3 | PC | Wave-4 | Representative |
| VP09142 | VP09142 | 2009 | China | Clinical | Asia | O3:K6 | ST3 | PC | Wave-4 | Representative |
| VP09160 | VP09160 | 2009 | China | Clinical | Asia | O3:K6 | ST3 | PC | Wave-4 | Representative |
| VP09205 | VP09205 | 2009 | China | Clinical | Asia | O3:K6 | ST3 | PC | Wave-4 | Representative |
| VP09217 | VP09217 | 2009 | China | Clinical | Asia | O3:K6 | ST3 | PC | Wave-4 | Representative |
| VP09265 | VP09265 | 2009 | China | Clinical | Asia | O3:K6 | ST3 | PC | Wave-4 | Representative |
| VP09274 | VP09274 | 2009 | China | Clinical | Asia | O3:K6 | ST3 | PC | Wave-4 | Representative |
| VP09313 | VP09313 | 2009 | China | Clinical | Asia | O3:K6 | ST3 | PC | Wave-4 | Representative |
| VP09315 | VP09315 | 2009 | China | Clinical | Asia | O3:K6 | ST3 | PC | Wave-4 | Representative |
| VP09318 | VP09318 | 2009 | China | Clinical | Asia | O3:K6 | ST3 | PC | Wave-4 | Representative |
| VP09327 | VP09327 | 2009 | China | Clinical | Asia | O3:K6 | ST3 | PC | Wave-4 | Representative |
| VP09398 | VP09398 | 2009 | China | Clinical | Asia | O3:K6 | ST3 | PC | Wave-4 | Representative |
| VP09404 | VP09404 | 2009 | China | Clinical | Asia | O3:K6 | ST3 | PC | Wave-4 | Representative |
| VP09436 | VP09436 | 2009 | China | Clinical | Asia | O3:K6 | ST3 | PC | Wave-4 | Representative |
| VP09445 | VP09445 | 2009 | China | Clinical | Asia | O3:K6 | ST3 | PC | Wave-4 | Representative |
| VP09473 | VP09473 | 2009 | China | Clinical | Asia | O3:K6 | ST3 | PC | Wave-4 | Representative |
| VP09474 | VP09474 | 2009 | China | Clinical | Asia | O3:K6 | ST3 | PC | Wave-4 | Representative |
| VP09477 | VP09477 | 2009 | China | Clinical | Asia | O3:K6 | ST3 | PC | Wave-4 | Representative |
| VP10001 | VP10001 | 2010 | China | Clinical | Asia | O3:K6 | ST3 | PC | Wave-4 | Representative |
| VP10063 | VP10063 | 2010 | China | Clinical | Asia | O3:K6 | ST3 | PC | Wave-4 | Representative |
| VP10076 | VP10076 | 2010 | China | Clinical | Asia | O3:K6 | ST3 | PC | Wave-4 | Representative |

| | | | | | | | | | | |
|---|---|---|---|---|---|---|---|---|---|---|
| VP10095 | VP10095 | 2010 | China | Clinical | Asia | O3:K6 | ST3 | PC | Wave-4 | Representative |
| VP10102 | VP10102 | 2010 | China | Clinical | Asia | O3:K6 | ST3 | PC | Wave-4 | Representative |
| VP10104 | VP10104 | 2010 | China | Clinical | Asia | O3:K6 | ST3 | PC | Wave-4 | Representative |
| VP10129 | VP10129 | 2010 | China | Clinical | Asia | O3:K6 | ST3 | PC | Wave-4 | Representative |
| VP10136 | VP10136 | 2010 | China | Clinical | Asia | O3:K6 | ST3 | PC | Wave-4 | Representative |
| VP10170 | VP10170 | 2010 | China | Clinical | Asia | O3:K6 | ST3 | PC | Wave-4 | Representative |
| VP10171 | VP10171 | 2010 | China | Clinical | Asia | O3:K6 | ST3 | PC | Wave-4 | Representative |
| VP10174 | VP10174 | 2010 | China | Clinical | Asia | O3:K6 | ST3 | PC | Wave-4 | Representative |
| VP10195 | VP10195 | 2010 | China | Clinical | Asia | O3:K6 | ST3 | PC | Wave-4 | Representative |
| VP10207 | VP10207 | 2010 | China | Clinical | Asia | O3:K6 | ST3 | PC | Wave-4 | Representative |
| VP10229 | VP10229 | 2010 | China | Clinical | Asia | O3:K6 | ST3 | PC | Wave-4 | Representative |
| VP10284 | VP10284 | 2010 | China | Clinical | Asia | O3:K6 | ST3 | PC | Wave-4 | Representative |
| VP10322 | VP10322 | 2010 | China | Clinical | Asia | O3:K6 | ST3 | PC | Wave-4 | Representative |
| VP10354 | VP10354 | 2010 | China | Clinical | Asia | O3:K6 | ST3 | PC | Wave-4 | Representative |
| VP10366 | VP10366 | 2010 | China | Clinical | Asia | O3:K6 | ST3 | PC | Wave-4 | Representative |
| VP10379 | VP10379 | 2010 | China | Clinical | Asia | O3:K6 | ST3 | PC | Wave-4 | Representative |
| VP10391 | VP10391 | 2010 | China | Clinical | Asia | O3:K6 | ST3 | PC | Wave-4 | Representative |
| VP10396 | VP10396 | 2010 | China | Clinical | Asia | O3:K6 | ST3 | PC | Wave-4 | Representative |
| VP10405 | VP10405 | 2010 | China | Clinical | Asia | O3:K6 | ST3 | PC | Wave-4 | Representative |
| VP10423 | VP10423 | 2010 | China | Clinical | Asia | O3:K6 | ST3 | PC | Wave-4 | Representative |
| VP10436 | VP10436 | 2010 | China | Clinical | Asia | O3:K6 | ST3 | PC | Wave-4 | Representative |
| VP10465 | VP10465 | 2010 | China | Clinical | Asia | O3:K6 | ST3 | PC | Wave-4 | Representative |
| VP10470 | VP10470 | 2010 | China | Clinical | Asia | O3:K6 | ST3 | PC | Wave-4 | Representative |
| VP10472 | VP10472 | 2010 | China | Clinical | Asia | O3:K6 | ST3 | PC | Wave-4 | Representative |
| VP10475 | VP10475 | 2010 | China | Clinical | Asia | O3:K6 | ST3 | PC | Wave-4 | Representative |
| VP11122 | VP11122 | 2011 | China | Clinical | Asia | O3:K6 | ST3 | PC | Wave-4 | Representative |
| VP11127 | VP11127 | 2011 | China | Clinical | Asia | O3:K6 | ST3 | PC | Wave-4 | Representative |
| VP11129 | VP11129 | 2011 | China | Clinical | Asia | O3:K6 | ST3 | PC | Wave-4 | Representative |
| VP11136 | VP11136 | 2011 | China | Clinical | Asia | O3:K6 | ST3 | PC | Wave-4 | Representative |
| VP11150 | VP11150 | 2011 | China | Clinical | Asia | O3:K6 | ST3 | PC | Wave-4 | Representative |
| VP11163 | VP11163 | 2011 | China | Clinical | Asia | O3:K6 | ST3 | PC | Wave-4 | Representative |
| VP11165 | VP11165 | 2011 | China | Clinical | Asia | O3:K6 | ST3 | PC | Wave-4 | Representative |

| | | | | | | | | | | |
|---|---|---|---|---|---|---|---|---|---|---|
| VP11169 | VP11169 | 2011 | China | Clinical | Asia | O3:K6 | ST3 | PC | Wave-4 | Representative |
| VP11175 | VP11175 | 2011 | China | Clinical | Asia | O3:K6 | ST3 | PC | Wave-4 | Representative |
| VP11179 | VP11179 | 2011 | China | Clinical | Asia | O3:K6 | ST3 | PC | Wave-4 | Representative |
| VP11186 | VP11186 | 2011 | China | Clinical | Asia | O3:K6 | ST3 | PC | Wave-4 | Representative |
| VP11199 | VP11199 | 2011 | China | Clinical | Asia | O3:K6 | ST3 | PC | Wave-4 | Representative |
| VP11205 | VP11205 | 2011 | China | Clinical | Asia | O3:K6 | ST3 | PC | Wave-4 | Representative |
| VP11225 | VP11225 | 2011 | China | Clinical | Asia | O3:K6 | ST3 | PC | Wave-4 | Representative |
| VP11239 | VP11239 | 2011 | China | Clinical | Asia | O3:K6 | ST3 | PC | Wave-4 | Representative |
| VP11240 | VP11240 | 2011 | China | Clinical | Asia | O3:K6 | ST3 | PC | Wave-4 | Representative |
| VP11246 | VP11246 | 2011 | China | Clinical | Asia | O3:K6 | ST3 | PC | Wave-4 | Representative |
| VP11252 | VP11252 | 2011 | China | Clinical | Asia | O3:K6 | ST3 | PC | Wave-4 | Representative |
| VP11264 | VP11264 | 2011 | China | Clinical | Asia | O3:K6 | ST3 | PC | Wave-4 | Representative |
| VP11272 | VP11272 | 2011 | China | Clinical | Asia | O3:K6 | ST3 | PC | Wave-4 | Representative |
| VP11278 | VP11278 | 2011 | China | Clinical | Asia | O3:K6 | ST3 | PC | Wave-4 | Representative |
| VP11288 | VP11288 | 2011 | China | Clinical | Asia | O3:K6 | ST3 | PC | Wave-4 | Representative |
| VP11292 | VP11292 | 2011 | China | Clinical | Asia | O3:K6 | ST3 | PC | Wave-4 | Representative |
| VP11301 | VP11301 | 2011 | China | Clinical | Asia | O3:K6 | ST3 | PC | Wave-4 | Representative |
| VP11305 | VP11305 | 2011 | China | Clinical | Asia | O3:K6 | ST3 | PC | Wave-4 | Representative |
| VP11313 | VP11313 | 2011 | China | Clinical | Asia | O3:K6 | ST3 | PC | Wave-4 | Representative |
| VP12001 | VP12001 | 2012 | China | Clinical | Asia | O3:K6 | ST3 | PC | Wave-4 | Representative |
| VP12003 | VP12003 | 2012 | China | Clinical | Asia | O3:K6 | ST3 | PC | Wave-4 | Representative |
| VP12006 | VP12006 | 2011 | China | Clinical | Asia | O3:K6 | ST3 | PC | Wave-4 | Representative |
| VP12019 | VP12019 | 2012 | China | Clinical | Asia | O3:K6 | ST3 | PC | Wave-4 | Representative |
| VP12047 | VP12047 | 2012 | China | Clinical | Asia | O3:K6 | ST3 | PC | Wave-4 | Representative |
| VP12050 | VP12050 | 2012 | China | Clinical | Asia | O3:K6 | ST3 | PC | Wave-4 | Representative |
| VP12053 | VP12053 | 2012 | China | Clinical | Asia | O3:K6 | ST3 | PC | Wave-4 | Representative |
| VP12055 | VP12055 | 2012 | China | Clinical | Asia | O3:K6 | ST3 | PC | Wave-4 | Representative |
| VP12059 | VP12059 | 2012 | China | Clinical | Asia | O3:K6 | ST3 | PC | Wave-4 | Representative |
| VP12062 | VP12062 | 2012 | China | Clinical | Asia | O3:K6 | ST3 | PC | Wave-4 | Representative |
| VP12066 | VP12066 | 2012 | China | Clinical | Asia | O3:K6 | ST3 | PC | Wave-4 | Representative |
| VP12076 | VP12076 | 2012 | China | Clinical | Asia | O3:K6 | ST3 | PC | Wave-4 | Representative |
| VP12086 | VP12086 | 2012 | China | Clinical | Asia | O3:K6 | ST3 | PC | Wave-4 | Representative |

| | | | | | | | | | | |
|---|---|---|---|---|---|---|---|---|---|---|
| VP12104 | VP12104 | 2012 | China | Clinical | Asia | O3:K6 | ST3 | PC | Wave-4 | Representative |
| VP12116 | VP12116 | 2012 | China | Clinical | Asia | O3:K6 | ST3 | PC | Wave-4 | Representative |
| VP12120 | VP12120 | 2012 | China | Clinical | Asia | O3:K6 | ST3 | PC | Wave-4 | Representative |
| VP12165 | VP12165 | 2012 | China | Clinical | Asia | O3:K6 | ST3 | PC | Wave-4 | Representative |
| VP12168 | VP12168 | 2012 | China | Clinical | Asia | O3:K6 | ST3 | PC | Wave-4 | Representative |
| VP12173 | VP12173 | 2012 | China | Clinical | Asia | O3:K6 | ST3 | PC | Wave-4 | Representative |
| VP12185 | VP12185 | 2012 | China | Clinical | Asia | O3:K6 | ST3 | PC | Wave-4 | Representative |
| VP12189 | VP12189 | 2012 | China | Clinical | Asia | O3:K6 | ST3 | PC | Wave-4 | Representative |
| VP12190 | VP12190 | 2012 | China | Clinical | Asia | O3:K6 | ST3 | PC | Wave-4 | Representative |
| VP12198 | VP12198 | 2012 | China | Clinical | Asia | O3:K6 | ST3 | PC | Wave-4 | Representative |
| VP12216 | VP12216 | 2012 | China | Clinical | Asia | O3:K6 | ST3 | PC | Wave-4 | Representative |
| VP12227 | VP12227 | 2012 | China | Clinical | Asia | O3:K6 | ST3 | PC | Wave-4 | Representative |
| VP12228 | VP12228 | 2012 | China | Clinical | Asia | O3:K6 | ST3 | PC | Wave-4 | Representative |
| VP12230 | VP12230 | 2012 | China | Clinical | Asia | O3:K6 | ST3 | PC | Wave-4 | Representative |
| VP12234 | VP12234 | 2012 | China | Env | Asia | O3:K6 | ST3 | PC | Wave-4 | Representative |
| VP12235 | VP12235 | 2012 | China | Env | Asia | O3:K6 | ST3 | PC | Wave-4 | Representative |
| VP12238 | VP12238 | 2012 | China | Clinical | Asia | O3:K6 | ST3 | PC | Wave-4 | Representative |
| VP13004 | VP13004 | 2013 | China | Clinical | Asia | O3:K6 | ST3 | PC | Wave-4 | Representative |
| VP13006 | VP13006 | 2012 | China | Clinical | Asia | O3:K6 | ST3 | PC | Wave-4 | Representative |
| VP13022 | VP13022 | 2013 | China | Clinical | Asia | O3:K6 | ST3 | PC | Wave-4 | Representative |
| VP13028 | VP13028 | 2013 | China | Clinical | Asia | O3:K6 | ST3 | PC | Wave-4 | Representative |
| VP13031 | VP13031 | 2013 | China | Clinical | Asia | O3:K6 | ST3 | PC | Wave-4 | Representative |
| VP13044 | VP13044 | 2013 | China | Clinical | Asia | O3:K6 | ST3 | PC | Wave-4 | Representative |
| VP13072 | VP13072 | 2013 | China | Clinical | Asia | O3:K6 | ST3 | PC | Wave-4 | Representative |
| VP13076 | VP13076 | 2013 | China | Clinical | Asia | O3:K6 | ST3 | PC | Wave-4 | Representative |
| VP13084 | VP13084 | 2013 | China | Clinical | Asia | O3:K6 | ST3 | PC | Wave-4 | Representative |
| VP13088 | VP13088 | 2013 | China | Clinical | Asia | O3:K6 | ST3 | PC | Wave-4 | Representative |
| VP13097 | VP13097 | 2013 | China | Clinical | Asia | O3:K6 | ST3 | PC | Wave-4 | Representative |
| VP13126 | VP13126 | 2013 | China | Clinical | Asia | O3:K6 | ST3 | PC | Wave-4 | Representative |
| VP14001 | VP14001 | 2014 | China | Clinical | Asia | O3:K6 | ST3 | PC | Wave-4 | Representative |
| VP14008 | VP14008 | 2014 | China | Clinical | Asia | O3:K6 | ST3 | PC | Wave-4 | Representative |
| VP14020 | VP14020 | 2014 | China | Clinical | Asia | O3:K6 | ST3 | PC | Wave-4 | Representative |

| | | | | | | | | | | |
|---|---|---|---|---|---|---|---|---|---|---|
| VP14079 | VP14079 | 2014 | China | Clinical | Asia | O3:K6 | ST3 | PC | Wave-4 | Representative |
| VP14113 | VP14113 | 2014 | China | Clinical | Asia | O3:K6 | ST3 | PC | Wave-4 | Representative |
| VP14118 | VP14118 | 2014 | China | Clinical | Asia | O3:K6 | ST3 | PC | Wave-4 | Representative |
| VP14120 | VP14120 | 2014 | China | Clinical | Asia | O3:K6 | ST3 | PC | Wave-4 | Representative |
| VP14141 | VP14141 | 2014 | China | Clinical | Asia | O10:K60 | ST3 | PC | Wave-4 | Representative |
| VP14149 | VP14149 | 2014 | China | Clinical | Asia | O3:K6 | ST3 | PC | Wave-4 | Representative |
| VP14171 | VP14171 | 2014 | China | Clinical | Asia | O3:K6 | ST3 | PC | Wave-4 | Representative |
| VP15003 | VP15003 | 2015 | China | Clinical | Asia | O3:K6 | ST3 | PC | Wave-4 | Representative |
| VP15004 | VP15004 | 2015 | China | Clinical | Asia | O3:K6 | ST3 | PC | Wave-4 | Representative |
| VP15005 | VP15005 | 2015 | China | Clinical | Asia | O3:K6 | ST3 | PC | Wave-4 | Representative |
| VP15007 | VP15007 | 2015 | China | Clinical | Asia | O3:K6 | ST3 | PC | Wave-4 | Representative |
| VP15008 | VP15008 | 2015 | China | Clinical | Asia | O3:K6 | ST3 | PC | Wave-4 | Representative |
| VP15012 | VP15012 | 2015 | China | Clinical | Asia | O3:K6 | ST3 | PC | Wave-4 | Representative |
| VP15013 | VP15013 | 2015 | China | Clinical | Asia | O3:K6 | ST3 | PC | Wave-4 | Representative |
| VP15052 | VP15052 | 2015 | China | Clinical | Asia | O3:K6 | ST3 | PC | Wave-4 | Representative |
| VP15072 | VP15072 | 2015 | China | Clinical | Asia | O3:K6 | ST3 | PC | Wave-4 | Representative |
| VP15074 | VP15074 | 2015 | China | Clinical | Asia | O3:K6 | ST3 | PC | Wave-4 | Representative |
| VP15119 | VP15119 | 2015 | China | Clinical | Asia | O3:K6 | ST3 | PC | Wave-4 | Representative |
| VP15130 | VP15130 | 2015 | China | Clinical | Asia | O3:K6 | ST3 | PC | Wave-4 | Representative |
| VP15132 | VP15132 | 2015 | China | Clinical | Asia | O3:K6 | ST3 | PC | Wave-4 | Representative |
| VP15136 | VP15136 | 2015 | China | Clinical | Asia | O3:K6 | ST3 | PC | Wave-4 | Representative |
| VP15143 | VP15143 | 2015 | China | Clinical | Asia | O3:K6 | ST3 | PC | Wave-4 | Representative |
| VP15144 | VP15144 | 2015 | China | Clinical | Asia | O3:K6 | ST3 | PC | Wave-4 | Representative |
| VP15159 | VP15159 | 2015 | China | Clinical | Asia | O3:K6 | ST3 | PC | Wave-4 | Representative |
| VP15171 | VP15171 | 2015 | China | Clinical | Asia | O3:K6 | ST3 | PC | Wave-4 | Representative |
| VP15176 | VP15176 | 2015 | China | Clinical | Asia | O3:K6 | ST3 | PC | Wave-4 | Representative |
| VP15191 | VP15191 | 2015 | China | Clinical | Asia | O3:K6 | ST3 | PC | Wave-4 | Representative |
| VP15197 | VP15197 | 2015 | China | Clinical | Asia | O3:K6 | ST3 | PC | Wave-4 | Representative |
| VP16008 | VP16008 | 2016 | China | Clinical | Asia | O3:K6 | ST3 | PC | Wave-4 | Representative |
| VP16024 | VP16024 | 2016 | China | Clinical | Asia | O3:K6 | ST3 | PC | Wave-4 | Representative |
| VP16043 | VP16043 | 2016 | China | Clinical | Asia | O3:K6 | ST- | PC | Wave-4 | Representative |
| VP16051 | VP16051 | 2016 | China | Clinical | Asia | O3:K6 | ST3 | PC | Wave-4 | Representative |

| | | | | | | | | | | |
|---|---|---|---|---|---|---|---|---|---|---|
| VP16054 | VP16054 | 2016 | China | Clinical | Asia | O3:K6 | ST3 | PC | Wave-4 | Representative |
| VP16073 | VP16073 | 2016 | China | Clinical | Asia | O3:K6 | ST3 | PC | Wave-4 | Representative |
| VP16095 | VP16095 | 2016 | China | Clinical | Asia | O3:K6 | ST3 | PC | Wave-4 | Representative |
| VP16103 | VP16103 | 2016 | China | Clinical | Asia | O3:K6 | ST3 | PC | Wave-4 | Representative |
| VP16112 | VP16112 | 2016 | China | Clinical | Asia | O3:K6 | ST3 | PC | Wave-4 | Representative |
| VP16122 | VP16122 | 2016 | China | Clinical | Asia | O3:K6 | ST3 | PC | Wave-4 | Representative |
| VP16124 | VP16124 | 2016 | China | Clinical | Asia | O3:K6 | ST3 | PC | Wave-4 | Representative |
| VP16142 | VP16142 | 2016 | China | Clinical | Asia | O3:K6 | ST3 | PC | Wave-4 | Representative |
| VP16158 | VP16158 | 2016 | China | Clinical | Asia | O3:K6 | ST3 | PC | Wave-4 | Representative |
| VP16196 | VP16196 | 2016 | China | Clinical | Asia | O3:K6 | ST3 | PC | Wave-4 | Representative |
| VP16201 | VP16201 | 2016 | China | Clinical | Asia | O3:K6 | ST3 | PC | Wave-4 | Representative |
| VP16205 | VP16205 | 2016 | China | Clinical | Asia | O3:K6 | ST3 | PC | Wave-4 | Representative |
| VP16208 | VP16208 | 2016 | China | Clinical | Asia | O3:K6 | ST3 | PC | Wave-4 | Representative |
| VP17007 | VP17007 | 2017 | China | Clinical | Asia | O10:K60 | ST3 | PC | Wave-4 | Representative |
| VP17012 | VP17012 | 2017 | China | Clinical | Asia | O3:K6 | ST3 | PC | Wave-4 | Representative |
| VP17020 | VP17020 | 2017 | China | Clinical | Asia | O3:K6 | ST3 | PC | Wave-4 | Representative |
| VP17024 | VP17024 | 2017 | China | Clinical | Asia | O3:K6 | ST3 | PC | Wave-4 | Representative |
| VP17034 | VP17034 | 2017 | China | Clinical | Asia | O3:K6 | ST3 | PC | Wave-4 | Representative |
| VP17051 | VP17051 | 2017 | China | Clinical | Asia | O3:K6 | ST3 | PC | Wave-4 | Representative |
| VP17081 | VP17081 | 2017 | China | Clinical | Asia | O3:K6 | ST3 | PC | Wave-4 | Representative |
| VP17096 | VP17096 | 2017 | China | Clinical | Asia | O3:K6 | ST3 | PC | Wave-4 | Representative |
| VP17101 | VP17101 | 2017 | China | Clinical | Asia | O3:K6 | ST3 | PC | Wave-4 | Representative |
| VP17120 | VP17120 | 2017 | China | Clinical | Asia | O3:K6 | ST3 | PC | Wave-4 | Representative |
| VP17121 | VP17121 | 2017 | China | Clinical | Asia | O3:K6 | ST3 | PC | Wave-4 | Representative |
| VP17122 | VP17122 | 2017 | China | Clinical | Asia | O3:K6 | ST3 | PC | Wave-4 | Representative |
| VP17137 | VP17137 | 2017 | China | Clinical | Asia | O3:K6 | ST3 | PC | Wave-4 | Representative |
| VP17144 | VP17144 | 2017 | China | Clinical | Asia | O3:K6 | ST3 | PC | Wave-4 | Representative |
| VP17145 | VP17145 | 2017 | China | Clinical | Asia | O3:K6 | ST3 | PC | Wave-4 | Representative |
| VP17149 | VP17149 | 2017 | China | Clinical | Asia | O3:K6 | ST3 | PC | Wave-4 | Representative |
| VP17162 | VP17162 | 2017 | China | Clinical | Asia | O3:K6 | ST3 | PC | Wave-4 | Representative |
| VP17166 | VP17166 | 2017 | China | Clinical | Asia | O3:K6 | ST3 | PC | Wave-4 | Representative |
| VP17168 | VP17168 | 2017 | China | Clinical | Asia | O3:K6 | ST3 | PC | Wave-4 | Representative |

| | | | | | | | | | | |
|---|---|---|---|---|---|---|---|---|---|---|
| VP17170 | VP17170 | 2017 | China | Clinical | Asia | O3:K6 | ST3 | PC | Wave-4 | Representative |
| VP18001 | VP18001 | 2018 | China | Clinical | Asia | O3:K6 | ST3 | PC | Wave-4 | Representative |
| VP18009 | VP18009 | 2018 | China | Clinical | Asia | O3:K6 | ST3 | PC | Wave-4 | Representative |
| VP18023 | VP18023 | 2018 | China | Clinical | Asia | O3:K6 | ST3 | PC | Wave-4 | Representative |
| VP18052 | VP18052 | 2018 | China | Clinical | Asia | O3:K6 | ST3 | PC | Wave-4 | Representative |
| VP18061 | VP18061 | 2018 | China | Clinical | Asia | O3:K6 | ST3 | PC | Wave-4 | Representative |
| VP18083 | VP18083 | 2018 | China | Clinical | Asia | O3:K6 | ST3 | PC | Wave-4 | Representative |
| VP18102 | VP18102 | 2018 | China | Clinical | Asia | O3:K6 | ST3 | PC | Wave-4 | Representative |
| VP18114 | VP18114 | 2018 | China | Clinical | Asia | O10:K60 | ST3 | PC | Wave-4 | Representative |
| VP18125 | VP18125 | 2018 | China | Clinical | Asia | O3:K6 | ST3 | PC | Wave-4 | Representative |
| VP18176 | VP18176 | 2018 | China | Clinical | Asia | O3:K6 | ST3 | PC | Wave-4 | Representative |
| VP18187 | VP18187 | 2018 | China | Clinical | Asia | O3:K6 | ST3 | PC | Wave-4 | Representative |
| VP18197 | VP18197 | 2018 | China | Clinical | Asia | O3:K6 | ST3 | PC | Wave-4 | Representative |
| VP18198 | VP18198 | 2018 | China | Clinical | Asia | O3:K6 | ST3 | PC | Wave-4 | Representative |
| VP18229 | VP18229 | 2018 | China | Clinical | Asia | O3:K6 | ST3 | PC | Wave-4 | Representative |
| VP18232 | VP18232 | 2018 | China | Clinical | Asia | O3:K6 | ST3 | PC | Wave-4 | Representative |
| VP19003 | VP19003 | 2019 | China | Clinical | Asia | O3:K6 | ST3 | PC | Wave-4 | Representative |
| VP19008 | VP19008 | 2019 | China | Clinical | Asia | O3:K6 | ST3 | PC | Wave-4 | Representative |
| VP19009 | VP19009 | 2019 | China | Clinical | Asia | O3:K6 | ST3 | PC | Wave-4 | Representative |
| VP19010 | VP19010 | 2019 | China | Clinical | Asia | O3:K6 | ST3 | PC | Wave-4 | Representative |
| VP19020 | VP19020 | 2019 | China | Clinical | Asia | O3:K6 | ST3 | PC | Wave-4 | Representative |
| VP19026 | VP19026 | 2019 | China | Clinical | Asia | O3:K6 | ST3 | PC | Wave-4 | Representative |
| VP19032 | VP19032 | 2019 | China | Clinical | Asia | O3:K6 | ST3 | PC | Wave-4 | Representative |
| VP19043 | VP19043 | 2019 | China | Clinical | Asia | O3:K6 | ST3 | PC | Wave-4 | Representative |
| VP19045 | VP19045 | 2019 | China | Clinical | Asia | O3:K6 | ST3 | PC | Wave-4 | Representative |
| VP19074 | VP19074 | 2019 | China | Clinical | Asia | O3:K6 | ST3 | PC | Wave-4 | Representative |
| VP19079 | VP19079 | 2019 | China | Clinical | Asia | O3:K6 | ST3 | PC | Wave-4 | Representative |
| VP19091 | VP19091 | 2019 | China | Clinical | Asia | O3:K6 | ST3 | PC | Wave-4 | Representative |
| VP19093 | VP19093 | 2019 | China | Clinical | Asia | O3:K6 | ST3 | PC | Wave-4 | Representative |
| VP19095 | VP19095 | 2019 | China | Clinical | Asia | O3:K6 | ST3 | PC | Wave-4 | Representative |
| VP19097 | VP19097 | 2019 | China | Clinical | Asia | O3:K6 | ST3 | PC | Wave-4 | Representative |
| VP19110 | VP19110 | 2019 | China | Clinical | Asia | O3:K6 | ST3 | PC | Wave-4 | Representative |

| | | | | | | | | | | |
|---|---|---|---|---|---|---|---|---|---|---|
| VP19142 | VP19142 | 2019 | China | Clinical | Asia | O3:K6 | ST3 | PC | Wave-4 | Representative |
| VP19164 | VP19164 | 2019 | China | Clinical | Asia | O3:K6 | ST3 | PC | Wave-4 | Representative |
| VP19167 | VP19167 | 2019 | China | Clinical | Asia | O10:K60 | ST3 | PC | Wave-4 | Representative |
| VP19170 | VP19170 | 2019 | China | Clinical | Asia | O3:K6 | ST3 | PC | Wave-4 | Representative |
| VP19211 | VP19211 | 2019 | China | Clinical | Asia | O3:K6 | ST3 | PC | Wave-4 | Representative |
| VP19237 | VP19237 | 2019 | China | Clinical | Asia | O3:K6 | ST3 | PC | Wave-4 | Representative |
| VP19250 | VP19250 | 2019 | China | Clinical | Asia | O3:K6 | ST3 | PC | Wave-4 | Representative |
| VP19257 | VP19257 | 2019 | China | Clinical | Asia | O3:K6 | ST3 | PC | Wave-4 | Representative |
| VP19273 | VP19273 | 2019 | China | Clinical | Asia | O3:K6 | ST3 | PC | Wave-4 | Representative |
| VP19275 | VP19275 | 2019 | China | Clinical | Asia | O3:K6 | ST3 | PC | Wave-4 | Representative |
| VP20006 | VP20006 | 2020 | China | Clinical | Asia | O3:K6 | ST3 | PC | Wave-4 | Representative |
| VP20016 | VP20016 | 2020 | China | Clinical | Asia | O3:K6 | ST3 | PC | Wave-4 | Representative |
| VP20021 | VP20021 | 2020 | China | Clinical | Asia | O3:K6 | ST3 | PC | Wave-4 | Representative |
| VP20022 | VP20022 | 2020 | China | Clinical | Asia | O3:K6 | ST3 | PC | Wave-4 | Representative |
| VP20024 | VP20024 | 2020 | China | Clinical | Asia | O3:K6 | ST3 | PC | Wave-4 | Representative |
| VP20032 | VP20032 | 2020 | China | Clinical | Asia | O3:K6 | ST3 | PC | Wave-4 | Representative |
| VP20033 | VP20033 | 2020 | China | Clinical | Asia | O3:K6 | ST3 | PC | Wave-4 | Representative |
| VP20092 | VP20092 | 2020 | China | Clinical | Asia | O3:K6 | ST3 | PC | Wave-4 | Representative |
| VP20094 | VP20094 | 2020 | China | Clinical | Asia | O3:K6 | ST3 | PC | Wave-4 | Representative |
| VP20095 | VP20095 | 2020 | China | Clinical | Asia | O10:K4 | ST3 | PC | Wave-4 | Representative |
| VP20097 | VP20097 | 2020 | China | Clinical | Asia | O3:K6 | ST3 | PC | Wave-4 | Representative |
| VP20109 | VP20109 | 2020 | China | Clinical | Asia | O3:K6 | ST3 | PC | Wave-4 | Representative |
| VP20158 | VP20158 | 2020 | China | Clinical | Asia | O3:K6 | ST3 | PC | Wave-4 | Representative |
| VP20159 | VP20159 | 2020 | China | Clinical | Asia | O3:K6 | ST3 | PC | Wave-4 | Representative |
| VP20160 | VP20160 | 2020 | China | Clinical | Asia | O10:K4 | ST3 | PC | Wave-4 | Representative |
| VP20161 | VP20161 | 2020 | China | Clinical | Asia | O3:K6 | ST3 | PC | Wave-4 | Representative |
| VP20164 | VP20164 | 2020 | China | Clinical | Asia | O3:K6 | ST3 | PC | Wave-4 | Representative |
| VP20165 | VP20165 | 2020 | China | Clinical | Asia | O3:K6 | ST3 | PC | Wave-4 | Representative |
| VP21015 | VP21015 | 2021 | China | Clinical | Asia | O3:K6 | ST3 | PC | Wave-4 | Representative |
| VP21026 | VP21026 | 2021 | China | Clinical | Asia | O10:K4 | ST3 | PC | Wave-4 | Representative |
| VP21032 | VP21032 | 2021 | China | Clinical | Asia | O3:K6 | ST3 | PC | Wave-4 | Representative |
| VP21039 | VP21039 | 2021 | China | Clinical | Asia | O3:K6 | ST3 | PC | Wave-4 | Representative |

| | | | | | | | | | | |
|---|---|---|---|---|---|---|---|---|---|---|
| VP21040 | VP21040 | 2021 | China | Clinical | Asia | O3:K6 | ST3 | PC | Wave-4 | Representative |
| VP21041 | VP21041 | 2021 | China | Clinical | Asia | O3:K6 | ST3 | PC | Wave-4 | Representative |
| VP21059 | VP21059 | 2021 | China | Clinical | Asia | O10:K4 | ST3 | PC | Wave-4 | Representative |
| VP21061 | VP21061 | 2021 | China | Clinical | Asia | O10:K4 | ST3 | PC | Wave-4 | Representative |
| VP21065 | VP21065 | 2021 | China | Clinical | Asia | O10:K4 | ST3 | PC | Wave-4 | Representative |
| VP21068 | VP21068 | 2021 | China | Clinical | Asia | O3:K6 | ST3 | PC | Wave-4 | Representative |
| VP21070 | VP21070 | 2021 | China | Clinical | Asia | O3:K6 | ST3 | PC | Wave-4 | Representative |
| VP21076 | VP21076 | 2021 | China | Clinical | Asia | O10:K4 | ST3 | PC | Wave-4 | Representative |
| VP21096 | VP21096 | 2021 | China | Clinical | Asia | O10:K4 | ST3 | PC | Wave-4 | Representative |
| VP4 | VP4 | NA | China | NA | Asia | O3:K6 | ST3 | PC | Wave-4 | Representative |
| VP5 | VP5 | NA | China | NA | Asia | O3:K6 | ST3 | PC | Wave-4 | Representative |
| VPCZ106 | VPCZ106 | 2014 | China | Clinical | Asia | O10:K60 | ST3 | PC | Wave-4 | Representative |
| VPCZ11 | VPCZ11 | 2014 | China | Clinical | Asia | O3:K6 | ST3 | PC | Wave-4 | Representative |
| VPCZ19 | VPCZ19 | 2014 | China | Clinical | Asia | O3:K6 | ST3 | PC | Wave-4 | Representative |
| VPCZ2 | VPCZ2 | 2013 | China | Clinical | Asia | O3:K6 | ST3 | PC | Wave-4 | Representative |
| VPCZ32 | VPCZ32 | 2014 | China | Clinical | Asia | O3:K6 | ST3 | PC | Wave-4 | Representative |
| VPCZ93 | VPCZ93 | 2014 | China | Clinical | Asia | O10:K60 | ST3 | PC | Wave-4 | Representative |
| SAMN06214610-GCA_015816435.1 | CFSAN053626 | 2016 | Spain | Clinical | Europe | O3:K6 | ST3 | PC | Wave-4 | Representative |
| Vp_UK_AB_S24 | AB_S24 | 2016 | UK(Colombia) | Clinical | Europe | O3:K6 | ST3 | PC | Wave-4 | Representative |
| Vp_UK_AD_S26 | AD_S26 | 2015 | UK | Clinical | Europe | O3:K6 | ST3 | PC | Wave-4 | Representative |
| Vp_UK_AE_S27 | AE_S27 | NA | UK | Clinical | Europe | O3:K6 | ST3 | PC | Wave-4 | Representative |
| Vp_UK_B_S2 | B_S2 | 2009 | UK(Vietnam) | Clinical | Europe | O3:K6 | ST3 | PC | Wave-4 | Representative |
| Vp_UK_F_S6 | F_S6 | 2010 | UK(Thailand) | Clinical | Europe | O3:K6 | ST3 | PC | Wave-4 | Representative |
| Vp_UK_H_S8 | H_S8 | 2014 | UK(Cuba) | Clinical | Europe | O3:K6 | ST3 | PC | Wave-4 | Representative |
| Vp_UK_K_S10 | K_S10 | 2014 | UK(Thailand) | Clinical | Europe | O10:K60 | ST3 | PC | Wave-4 | Representative |
| Vp_UK_L_S11 | L_S11 | 2014 | UK(Thailand) | Clinical | Europe | O10:K60 | ST3 | PC | Wave-4 | Representative |
| Vp_UK_R_S14 | R_S14 | 2013 | UK | Clinical | Europe | O3:K6 | ST3 | PC | Wave-4 | Representative |
| Vp_UK_V_S18 | V_S18 | 2016 | UK(Thailand) | Clinical | Europe | O3:K6 | ST3 | PC | Wave-4 | Representative |
| SAMN13893117-GCF_010692765.1 | CAIM_1400 | 2004 | Mexico | Clinical | Latin America | O3:K6 | ST3 | PC | Wave-4 | Representative |
| SAMN13893122-GCF_009936625.1 | CICESE-273 | 2012 | Mexico | Env | Latin America | O3:K65 | ST3 | PC | Wave-4 | Representative |
| SAMN20804967-GCA_022290605.1 | PV278 | 2019 | Colombia | Clinical | Latin America | O3:K6 | ST3 | PC | Wave-4 | Representative |
| SAMN20804968-GCA_022290575.1 | PV280 | 2019 | Colombia | Clinical | Latin America | O3:K6 | ST3 | PC | Wave-4 | Representative |

| | | | | | | | | | | |
|---|---|---|---|---|---|---|---|---|---|---|
| SAMN20804969-GCA_022290415.1 | PV53 | 2018 | Colombia | Clinical | Latin America | O3:K6 | ST3 | PC | Wave-4 | Representative |
| SAMN20804970-GCA_022290445.1 | PV85 | 2017 | Colombia | Clinical | Latin America | O3:K6 | ST3 | PC | Wave-4 | Representative |
| SAMN09280047-GCA_015805855.1 | 218355 | 2015 | NA | Clinical | NA | O3:K6 | ST3 | PC | Wave-4 | Representative |
| SAMN09280051-GCA_015805695.1 | 242385 | 2016 | NA | Clinical | NA | O3:K6 | ST3 | PC | Wave-4 | Representative |
| SAMN09280054-GCA_015805755.1 | 399421 | 2015 | NA | Clinical | NA | O3:K6 | ST3 | PC | Wave-4 | Representative |
| SAMN09280057-GCA_015805595.1 | 404285 | 2016 | NA | Clinical | NA | O3:K6 | ST3 | PC | Wave-4 | Representative |
| SAMN10338902-GCA_015798015.1 | 617910 | 2018 | NA | Clinical | NA | O3:K6 | ST3 | PC | Wave-4 | Representative |
| SAMN11890816-GCA_015790165.1 | 652104 | 2018 | NA | Clinical | NA | O3:K6 | ST3 | PC | Wave-4 | Representative |
| SAMN02741385-GCF_000707745.2 | CFSAN007449 | 2012 | USA | Clinical | US_Canada | O3:K6 | ST3 | PC | Wave-4 | Representative |
| SAMN02741387-GCF_000707805.2 | CFSAN007451 | 2012 | USA | Clinical | US_Canada | O3:K6 | ST3 | PC | Wave-4 | Representative |
| SAMN03349606-GCF_000960685.1 | 09-4435 | 2009 | Canada | Clinical | US_Canada | O3:K6 | ST3 | PC | Wave-4 | Representative |
| SAMN03349609-GCF_001006105.1 | 10-4251 | 2006 | Canada | Clinical | US_Canada | O3:K6 | ST3 | PC | Wave-4 | Representative |
| SAMN03452290-GCF_000972045.1 | 07-1339 | 2007 | Canada | Clinical | US_Canada | O3:K6 | ST3 | PC | Wave-4 | Representative |
| SAMN05220835-SRR3655232 | 4635 | 2007 | USA | Clinical | US_Canada | O3:K6 | ST3 | PC | Wave-4 | Representative |
| SAMN05220837-SRR3655243 | 4703 | 2007 | USA | Clinical | US_Canada | O3:K6 | ST3 | PC | Wave-4 | Representative |
| SAMN07327346-GCA_015816995.1 | PNUSAV000061 | NA | USA | Clinical | US_Canada | O3:K6 | ST3 | PC | Wave-4 | Representative |
| SAMN07559594-GCA_015816825.1 | PNUSAV000055 | NA | USA | Clinical | US_Canada | O3:K6 | ST3 | PC | Wave-4 | Representative |
| SAMN07682410-GCA_015816265.1 | PNUSAV000097 | NA | USA | Clinical | US_Canada | O3:K6 | ST3 | PC | Wave-4 | Representative |
| SAMN07838949-GCA_015815765.1 | PNUSAV000101 | NA | USA | Clinical | US_Canada | O3:K6 | ST3 | PC | Wave-4 | Representative |
| SAMN08113868-GCA_015814725.1 | PNUSAV000140 | NA | USA | Clinical | US_Canada | O3:K6 | ST3 | PC | Wave-4 | Representative |
| SAMN08627207-GCA_015812285.1 | PNUSAV000174 | NA | USA | Clinical | US_Canada | O3:K6 | ST3 | PC | Wave-4 | Representative |
| SAMN08627210-GCA_015812265.1 | PNUSAV000173 | NA | USA | Clinical | US_Canada | O3:K6 | ST3 | PC | Wave-4 | Representative |
| SAMN09689475-GCA_015803755.1 | PNUSAV000186 | NA | USA | Clinical | US_Canada | O3:K6 | ST3 | PC | Wave-4 | Representative |
| SAMN09703857-GCA_015803815.1 | PNUSAV000223 | NA | USA | Clinical | US_Canada | O3:K6 | ST3 | PC | Wave-4 | Representative |
| SAMN09703860-GCA_015804015.1 | PNUSAV000221 | NA | USA | Clinical | US_Canada | O3:K6 | ST3 | PC | Wave-4 | Representative |
| SAMN09745133-GCA_015804585.1 | PNUSAV000255 | NA | USA | Clinical | US_Canada | O3:K6 | ST3 | PC | Wave-4 | Representative |
| SAMN10095496-GCA_016303805.1 | PNUSAV000439 | NA | USA | Clinical | US_Canada | O3:K6 | ST3 | PC | Wave-4 | Representative |
| SAMN10095504-GCA_015800755.1 | PNUSAV000442 | NA | USA | Clinical | US_Canada | O3:K6 | ST3 | PC | Wave-4 | Representative |
| SAMN10148613-GCA_015799255.1 | PNUSAV000455 | NA | USA | Clinical | US_Canada | O3:K6 | ST3 | PC | Wave-4 | Representative |
| SAMN10391026-GCA_015797575.1 | PNUSAV000523 | NA | USA | Clinical | US_Canada | O3:K6 | ST3 | PC | Wave-4 | Representative |
| SAMN10794262-GCA_015791115.1 | PNUSAV000566 | NA | USA | Clinical | US_Canada | O3:K6 | ST3 | PC | Wave-4 | Representative |
| SAMN10854974-GCA_015790415.1 | PNUSAV000567 | NA | USA | Clinical | US_Canada | O3:K6 | ST3 | PC | Wave-4 | Representative |

| | | | | | | | | | | |
|---|---|---|---|---|---|---|---|---|---|---|
| SAMN12211964-GCA_015789655.1 | PNUSAV000618 | NA | USA | Clinical | US_Canada | O3:K6 | ST3 | PC | Wave-4 | Representative |
| SAMN12661289-GCA_015784365.1 | PNUSAV000950 | NA | USA | Clinical | US_Canada | O3:K6 | ST3 | PC | Wave-4 | Representative |
| SAMN12699060-GCA_015785195.1 | PNUSAV000970 | NA | USA | Clinical | US_Canada | O3:K6 | ST3 | PC | Wave-4 | Representative |
| SAMN14266007-GCA_015774455.1 | PNUSAV001182 | NA | USA | Clinical | US_Canada | O3:K6 | ST3 | PC | Wave-4 | Representative |
| SAMN14396416-GCA_018089465.1 | PNUSAV001185 | NA | USA | Clinical | US_Canada | O3:K6 | ST3 | PC | Wave-4 | Representative |
| SAMN17041481-GCA_018033385.1 | PNUSAV001478 | 2020 | USA | Clinical | US_Canada | O3:K6 | ST3 | PC | Wave-4 | Representative |
| SAMN21855655-GCA_020190745.1 | PNUSAV002219 | 2021 | USA | Clinical | US_Canada | O3:K6 | ST3 | PC | Wave-4 | Representative |
| SAMN22231286-GCA_020451485.1 | PNUSAV002025 | 2021 | USA | Clinical | US_Canada | O3:K6 | ST3 | PC | Wave-4 | Representative |
| SAMN23215746-GCA_020944005.1 | PNUSAV002346 | 2021 | USA | Clinical | US_Canada | O3:K6 | ST3 | PC | Wave-4 | Representative |
| SAMN28816693-SRR19512859 | PNUSAV002833 | 2022 | USA | Clinical | US_Canada | O10:K4 | ST3 | PC | Wave-4 | Representative |
| SAMN29444631-SRR19910450 | PNUSAV002824 | 2022 | USA | Clinical | US_Canada | O10:K4 | ST3 | PC | Wave-4 | Representative |
| SAMN29444632-SRR19910447 | PNUSAV002825 | 2022 | USA | Clinical | US_Canada | O10:K4 | ST3 | PC | Wave-4 | Representative |
